# GHT-SELEX demonstrates unexpectedly high intrinsic sequence specificity and complex DNA binding of many human transcription factors

**DOI:** 10.1101/2024.11.11.618478

**Authors:** Arttu Jolma, Aldo Hernandez-Corchado, Ally W.H. Yang, Ali Fathi, Kaitlin U. Laverty, Alexander Brechalov, Rozita Razavi, Mihai Albu, Hong Zheng, The Codebook Consortium, Ivan V. Kulakovskiy, Hamed S. Najafabadi, Timothy R. Hughes

## Abstract

Precise identification of transcription factor (TF) binding sites is a long standing challenge in human regulatory genomics: TF binding motifs are short and degenerate, while the genome is large. Motif scans, therefore, often produce excessive binding site predictions. By surveying 179 TFs across 25 families using >1,500 cyclic *in vitro* selection experiments with fragmented, naked, and unmodified genomic DNA – a method we term GHT-SELEX (Genomic HT-SELEX) – we find that many human TFs possess much higher sequence specificity than anticipated. Moreover, genomic binding regions from GHT-SELEX are often surprisingly similar to those obtained *in vivo* (i.e., ChIP-seq peaks). Contrary to conventional wisdom, we find that high specificity can also be obtained from motif scans, but performance is highly dependent on the derivation and use of the motifs, including accounting for multiple local matches. We also observe alternative engagement of multiple DNA-binding domains within the same protein: long C2H2 zinc finger proteins often utilize modular DNA recognition, engaging different subsets of their DNA-binding domain (DBD) arrays to recognize multiple types of distinct target sites, frequently evolving via internal duplication and divergence of one or more DBDs. Thus, it is common for TFs to possess sufficient intrinsic specificity to delineate a large fraction of *in vivo* genomic targets, independently of other cellular factors.

## INTRODUCTION

The DNA-binding sequence preference of a Transcription Factor (TF) is typically referred to as a motif, and is most commonly modeled as a position weight matrix (PWM), which describes the relative preference of the TF for each base in the binding site^1^. In humans, TF binding motifs are generally short and flexible; PWMs are typically 8-14 bases long^2–4^, and multiple bases can be tolerated at many positions^5,6^. Thus, a typical TF PWM scan with default parameters yields over a million potential binding sites in the 3-billion-base human genome, often with multiple high-scoring matches per gene. Very few of the potential target sites are utilized in cells^7^, however, and the actual number of bound sites, as measured by ChIP-seq^8–10^ or other assays^11^ is typically much lower than the number of motif matches.

This deficit in specificity has been resolved conceptually by the widespread cooperative binding and synergy among TFs^5,6,12,13^, and evidence that the chromatin landscape generally dominates TF binding site selection, such that TF motif matches only determine binding within permissible regions^14^. In the latter model, only a special class of “pioneer” TFs can access target sequences to control the local chromatin. Indeed, some TFs have been shown to have high inherent specificity: for example, CTCF binds the majority of its strongest motif matches in the genome^15^, and repositions the surrounding nucleosomes^16^. PRDM9, which controls recombination hotspots, has been reported to independently specify roughly half of its binding sites in the genome^17^.

Another possible explanation for the generally low apparent specificity of TF motifs, however, is that PWMs are inaccurate, or are used inappropriately, or that the PWM model is fundamentally flawed^18^. PWMs are often derived from a non-comprehensive set of bound vs. unbound sequences, and there is ongoing controversy regarding the best methods for derivation, underlying representation, and scanning of TF motifs^1,19^, as well as the impact of DNA shape^20^, dependencies among base positions^18,21^, multimeric binding^22,23^, and lower-affinity binding sites^24^. Proteins in the CXXC zinc finger (CXXC-zf) class, for example, bind primarily to clusters of unmethylated CG dinucleotides within CpG islands^25^, achieving high specificity despite an extremely simple monomeric binding site.

Many human TFs still lack binding motifs, and prominent among them are hundreds of C2H2 zinc finger (C2H2-zf) proteins^26^. These proteins recognize DNA sequences that approximate a concatenation of the three or four base specificities of their sequential constituent C2H2-zf domains^27,28^. Different C2H2-zf proteins can bind very different motifs due to both the malleability of the individual C2H2-zf domains and rearrangement of the individual C2H2-zf domains^29^. An enigmatic feature of the C2H2-zf proteins is their theoretical capacity to recognize very long sequences: the median number of C2H2-zf domains in human TFs is 11, which could contact up to 33 DNA bases, much more than would be needed to specifically recognize even a single target site in the genome, on average. Indeed, C2H2-zf proteins often use only a subset of their DBDs to contact DNA, and whether and how frequently human C2H2-zf proteins utilize different segments of the C2H2-zf domain array to bind different sequences has also been a long-standing question. In a well-studied example, CTCF binding sites appear to reflect a constitutive “core”, bound by fingers 4-7 of the 11 C2H2-zf domain array, flanked by sequences that are bound by alternative usage of upstream and/or downstream C2H2-zf domains^30,31^. In another example, mouse Znf335 was found to bind to two distinct motifs utilizing two separated C2H2-zf arrays^32^. Analysis of the DNA-binding of C2H2-zf proteins to the genome is also complicated by the fact that they often bind repeat elements such as endogenous retroelements^33^, and thus the target site similarity is derived both from DNA recognition and the shared ancestry of the binding sites. The limited resolution of ChIP-seq (>100bp) presents a related hindrance. These confounding factors, however, can be ameliorated by incorporating information about the bases that are likely preferred at each position of the binding site, as predicted by a C2H2-zf “recognition code” that relates the C2H2-zf amino acid sequences to their binding preferences. These machine learning-based predictions can assist in identifying the most plausible protein-DNA interactions in such cases, as our earlier work demonstrated^34^.

Methods for the study of TF sequence specificity *in vitro* are important because they remove any confounding impact of cellular factors beyond the TF of interest. Some *in vitro* approaches survey binding to randomly-generated sequences, typified by SELEX (Systematic Evolution of Ligands by EXponential enrichment)^35^, which uses multiple cycles of affinity capture interleaved with PCR amplification of the selected DNA ligands. HT-SELEX (High-throughput SELEX), in which the protein-bound DNA is immobilized in microwell plates and multiplexed Illumina sequencing is performed on all selection cycles, has been used to characterize DNA binding of hundreds of TFs in human and other species^36,37^. SELEX has the advantage that it has a more uniform k-mer distribution and potentially higher diversity than genomic DNA, but has a disadvantage as increasingly long sequences are present in a decreasing proportion of pool sequences. This issue is of particular concern for C2H2-zfs, due to the potential for very large binding sites. Indeed, long C2H2-zf proteins have a lower success rate in HT-SELEX than those with few C2H2-zf domains^36,38^.

An alternative to randomized DNA is to use fragmented genomic DNA. SELEX has previously been performed with *E. coli* genomic DNA with either a sequencing^39^ or microarray^40^ readout. More recent innovations, including CAP-seq^41^, Affinity-seq^17^, and DAP-seq^42^, have employed only a single round of selection, allowing direct comparison (and similar peak calling) to ChIP-seq. These approaches have revealed remarkably high specificity for some TFs. They are not cost-effective when applied to the human genome, however, due to its large size. To our knowledge, only two human proteins (PRDM9^17^ and Cfp1^41^ have been studied by these methods.

Here, we describe GHT-SELEX (Genomic DNA HT-SELEX), a novel implementation of the HT-SELEX^37^ method that is performed with fragmented human genomic DNA, and uses an associated new statistical analysis method, MAGIX (Model-based Analysis of Genomic Intervals with eXponential enrichment). GHT-SELEX combines the advantages of HT-SELEX (the use of barcoding, magnetic affinity beads and laboratory automation makes it possible to run GHT-SELEX in parallel with hundreds of samples over numerous cycles) with the strengths of genomic DNA selection protocols^17,42^ (representation of long genomic sites such as repeat sequences, and determination of individual genomic binding sites). We developed GHT-SELEX in the context of the Codebook consortium project^43^, which was aimed primarily at analysis of uncharacterized putative TFs, and provides comparison data from several other platforms for the same set of TFs (HT-SELEX, ChIP-seq, Protein Binding Microarrays^44^, and SMiLE-seq^45^). We successfully applied GHT-SELEX to 139 uncharacterized TFs and 40 control TFs. For dozens of TFs, including some that are considered well-characterized, GHT-SELEX peaks correspond closely to *in vivo* binding (measured by ChIP-seq). We also derived motif models that predict genomic binding with similar accuracy in most cases. GHT-SELEX is particularly effective for C2H2-zf proteins and shows that they often use alternative subsets of their C2H2-zf domains to engage with different genomic target sites. We explore both explanations and ramifications of these observations.

## RESULTS

### Development and testing of GHT-SELEX

We developed GHT-SELEX to run in parallel with HT-SELEX (**Figure 1a**, see **Table S1** for oligomer designs and **Table S2** for experimental parameters) in the context of the Codebook consortium^43^, a systematic effort to obtain DNA-binding motifs for 331 uncharacterized putative TFs (defined as proteins containing DBDs or having prior literature evidence but lacking known motifs^46^), as well as 61 control TFs with established motifs (**Table S3**). The Codebook clone set included both full-length proteins and DBDs, and three types of expression vectors, to enable production by three different expression systems (N-terminal eGFP fusions in both wheat germ extract and HEK293 cells, and N-terminal GST fusions in *E. coli* extracts; see **Methods**). The GHT-SELEX DNA pool used in this study was produced by nonspecific enzymatic fragmentation (Fragmentase) of HEK293 DNA to fragments with a median length of ∼64 bp. We used HEK293 DNA for compatibility with ChIP-seq data generated simultaneously (see accompanying manuscript^47^), and the length of the DNA was chosen to mimic standard HT-SELEX procedures and provide relatively high resolution.

**Figure 1.**
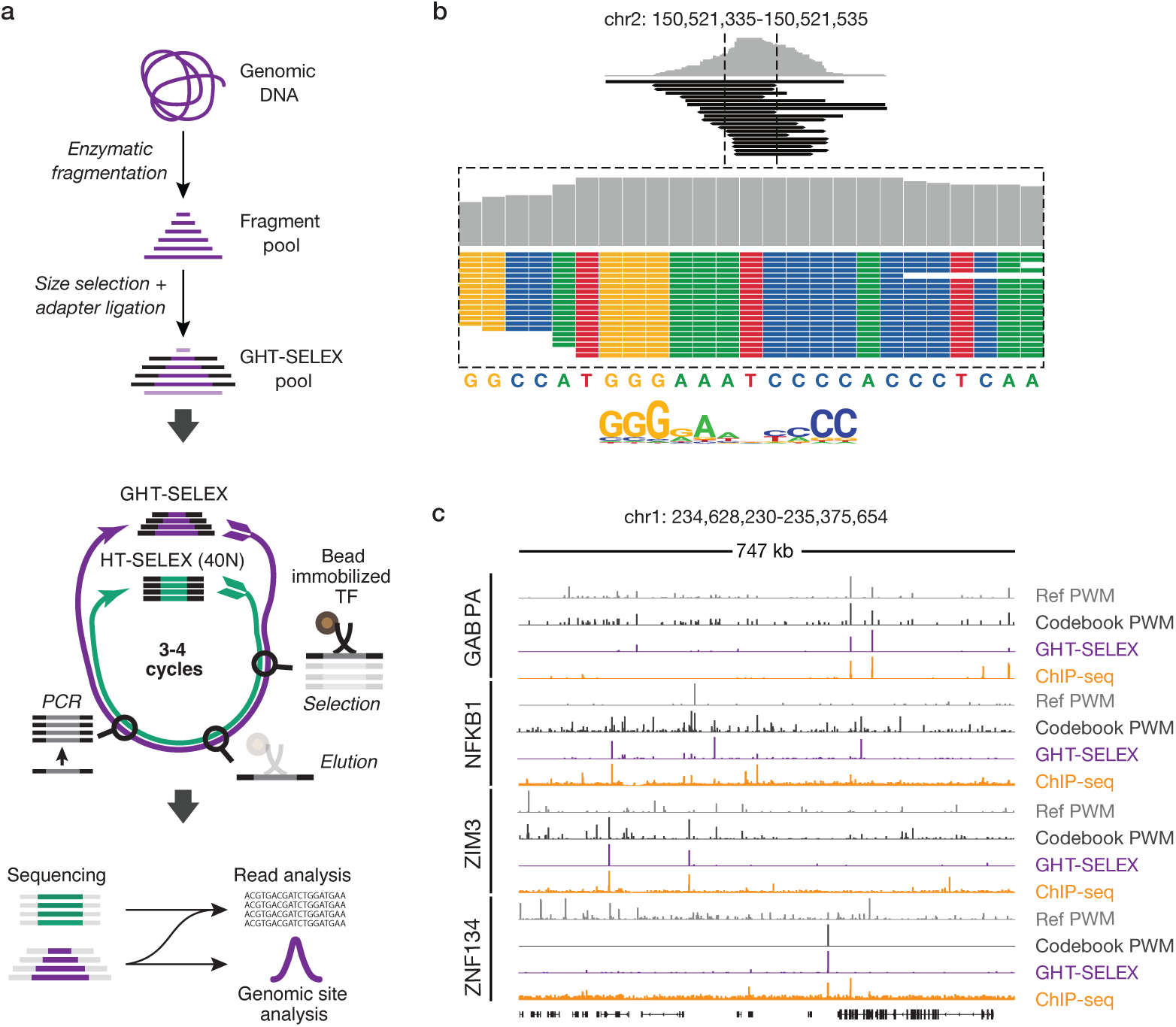
Overview of GHT-SELEX. **a.** Schematic of GHT-SELEX, showing parallels with HT-SELEX. **b.** Example of read accumulation over a TF motif match for NFKB1. **c.** Genomic binding for four positive control TFs on a genomic region showing (top to bottom) PWM scanning scores (moving average of affinity scores, from MOODS^69^ scan in linear domain, using a window of size 200bp) for literature PWMs (CIS-BP Identifiers: M0839, M03448, M08312 and M02996) and Codebook PWMs, followed by read coverage signal observed in GHT-SELEX and ChIP-seq.

We initially tested GHT-SELEX on the Codebook control proteins. Thirty of the control TFs represented a sampling of well-studied TFs with different classes of DBDs, most of which were previously analyzed using the independent *in vitro* SMiLE-seq platform^45^. An additional 31 controls were C2H2-zf proteins for which published ChIP-seq data yielded motifs^48^. Individual mapped reads typically accumulated at sites in which all reads overlap with what appears to be a motif match (**Figure 1b**). Moreover, the GHT-SELEX data typically had a strong resemblance to ChIP-seq data, forming strong peaks found sparsely across the genome, consistent with results from CAP-seq^41^, Affinity-seq^17^, and DAP-seq^42^. **Figure 1c** shows raw read density for four control TFs, comparing GHT-SELEX to ChIP-seq, and to target site predictions based on existing and newly derived (see below) PWM models for the TFs. GHT-SELEX experiments showed generally good reproducibility between replicates (examples in **Figure 2a**).

**Figure 2.**
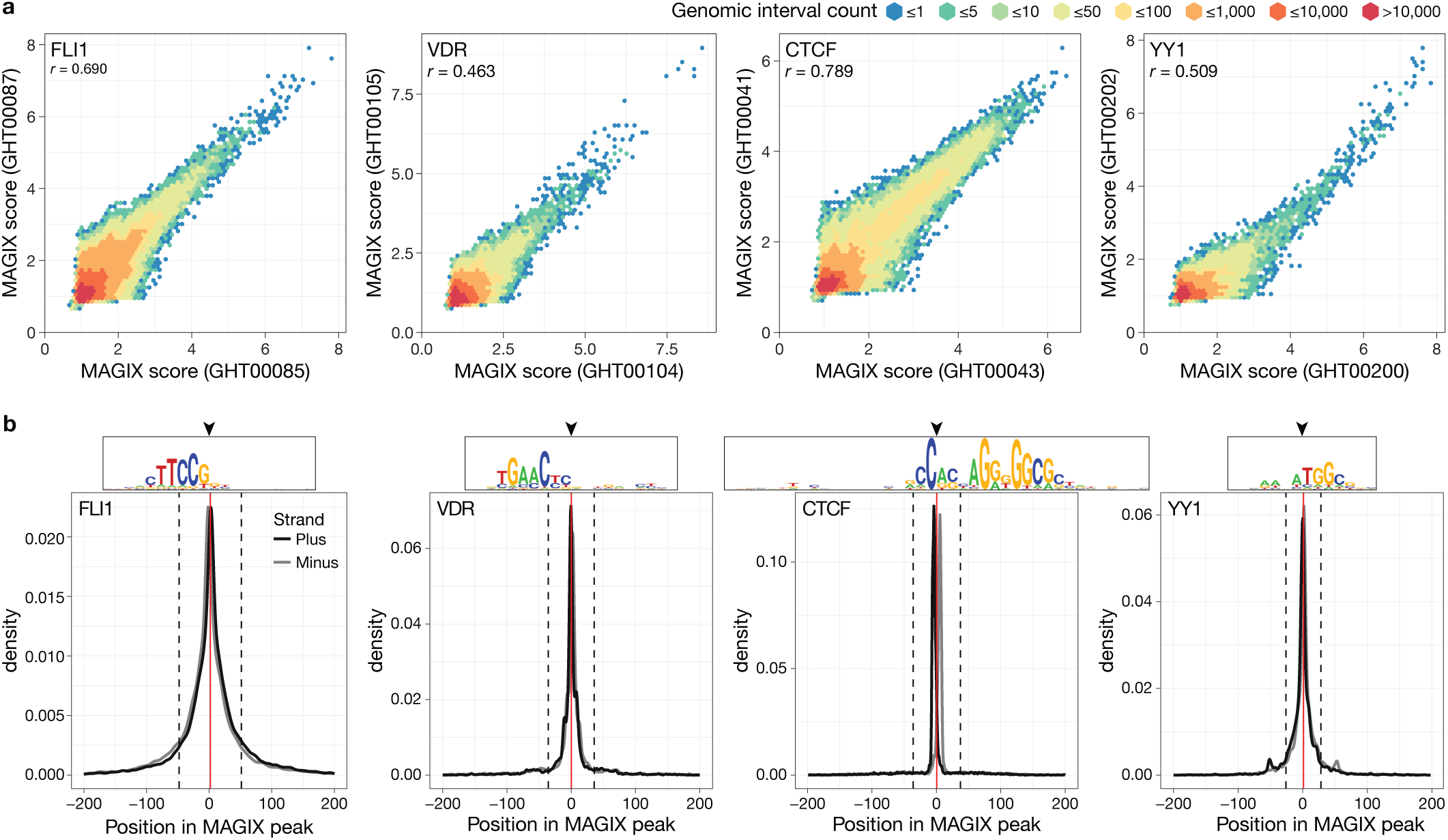
MAGIX method for interpretation of GHT-SELEX data. **a.** GHT-SELEX reproducibility. Scatterplots for four control TFs show MAGIX score values for all 200bp genomic bins between replicate GHT-SELEX experiments. **b.** Distribution of PWM hits for the top-ranked TF PWM (highest AUROC on GHT-peaks as determined in the accompanying study^49^ within the MAGIX peaks (up to the 10,000 highest scoring peaks). PWM hits were identified with MOODS^69^ (P < 0.0001) and the hit location is defined by the centre of the PWM, depicted above each plot. Solid red lines represent the mean PWM hit position within MAGIX peaks and dashed lines represent one standard deviation about the mean. Hits on the plus and minus strands are differentiated.

Peak calling from the GHT-SELEX data with conventional algorithms is confounded by the fact that different peaks have very different enrichment ratios across the cycles. This is presumably due to varying affinity of the TF for different sites, the overall increase in motif occurrences in the pool over the successive cycles, and simultaneous reduction in pool complexity, with the strongest binding sites dominating later cycles (**Extended Data Fig. 1a**). Consequently, enrichment information is distributed across the fragments from multiple selection cycles, with weaker peaks first appearing and disappearing, and the strongest peaks dominating in the later cycles **Extended Data Fig. 1b**). To adapt to these phenomena, we developed an analytical framework “MAGIX” (Model-based Analysis of Genomic Intervals with eXponential enrichment) that capitalizes on the added information gained from multiple SELEX cycles (**Extended Data Fig. 1b**; see **Methods** for details). The approach relies on a statistical method that explicitly models the exponential growth of TF-bound genomic regions over the SELEX cycles, which leads to a progressively higher proportion of TF-bound fragments relative to genomic background. The fragment abundances, in turn, are modeled as latent variables that determine the number of observed reads through a Poisson process. This hierarchical Bayesian model enables the integration of information across different selection cycles, experiments, and batches to calculate an estimated enrichment coefficient (MAGIX score) and associated FDR (**Extended Data Fig. 1b)**.

Among the 61 control proteins, we deemed 40 as successful in at least one GHT-SELEX experiment, based on enrichment of the expected motif at the peak centre (**Figure 2b**) (see **Methods** and accompanying manuscripts^43,49^ for details of how success was determined, and how PWMs were derived). The control proteins that were not successful in GHT-SELEX were mainly (15/21) from among the C2H2-zf proteins for which published ChIP-seq data yielded motifs^48^. These experiments were performed early in the study, prior to technical innovations that dramatically improved success rates with this class of proteins. The success rate for proteins for which previously-reported SMiLE-seq data yielded motifs may be a more realistic metric to gauge the success of GHT-SELEX in its current form (24/30, or 80%).

Among the 40 control TFs that were successful, 23 were represented by more than one successful experiment **(Table S4),** allowing us to assess the reproducibility of the combined GHT-SELEX/MAGIX pipeline. **Figure 2a** shows the enrichment values over genomic bins for replicates of FLI1, VDR, YY1, and CTCF. Similar plots for all successful replicates are given in **Supplementary Document S1**, and Pearson correlations for replicate pairs are in **Table S5.** We conclude that enrichment values across the genome are reproducible. MAGIX readily accommodates multiple experiments into a single analysis, and all subsequent analyses in this study utilize merged experimental data for each distinct protein.

Analysis of the data for the 40 successful controls by MAGIX resulted in between 926 and 82,045 peaks (median 9,927) with enrichment coefficient FDR less than 5% (see **Methods**). The number of strong PWM hits, on average, declines rapidly at ∼50 bp from peak centers, consistent with the utilized DNA fragment size (**Figure 2b)**; similar plots for all TFs analyzed are shown in **Supplementary Document S2**. In addition, higher PWM scores (which would, in theory, predict higher relative affinity) are clearly associated with a higher GHT-SELEX enrichment coefficient (see below).

### Application of GHT-SELEX to the full Codebook TF set

We next performed GHT-SELEX and, in parallel, HT-SELEX using random 40N ligands (**Table S1**) to assess DNA sequence specificity of 331 poorly characterized putative human TFs, as part of the Codebook project. We analyzed individual TFs with up to three types of constructs, and up to three protein expression strategies (two types of *in vitro* transcription–translation reactions, and expression in HEK293 cells, see **Methods**). We modulated several experimental variables over the course of the study (the protein production method, the number of washes, and digestion of single-stranded DNA with mung bean nuclease between cycles)in an effort to improve success rates. The protein production method appears to be most critical, particularly for C2H2-zf domain arrays (56%, 17% and 10% success rates using wheat germ extract, mammalian cell and *E. coli* extract-based expression systems, respectively; see **Methods, Table S2, and Extended data Fig. S2**).

For each TF, the constructs contained the full sequence of a representative isoform, or either all or a subset of its predicted DBDs. In total, we analyzed 1,315 constructs encompassing the 61 control TFs and 331 of the 332 putative TFs in the Codebook set of poorly characterized proteins. With these constructs, we performed 1,534 GHT-SELEX and 1,578 HT-SELEX experiments (see **Methods** and **Table S3**). The number of experiments per putative TF varied, due to the heterogeneity of the TFs and consequently heterogeneity in the clone collection; e.g., small proteins were represented with only a single insert, whereas larger, multi-DBD proteins were represented with both full-length and partial constructs. The vast majority (371/392, or 95%) of the proteins were tested two or more times in GHT-SELEX experiments, with a median of 4 experiments for each.

Importantly, 27% (420/1534) of the GHT-SELEX experiments were strict replicates (i.e., same construct, same expression system), covering a total of 198 potential TFs in 204 experiments (see **Methods**, **Table S3, and Table S4**), while the remainder were not strict replicates, allowing us to compare results from different expression systems.

Different protein production methods could potentially yield different DNA-binding activities for the same TF, e.g., via post-translational modifications, or if the intended protein associates with interaction partners that also bind DNA. It is also possible that the isolated DBD could bind different DNA sequences from the full-length protein. To ask whether we observe such differences, we evaluated 217 pairs of experiments where the same TF was expressed by two different expression systems (e.g., wheat germ vs. mammalian cells), and a further 72 pairs that used the same expression system, by plotting enrichment of 3, 5, 7, and 9 bp k-mers in the HT-SELEX data (a total of 289 instances encompassing 62 TFs; **Supplementary Document S3, Table S6**). We identified minor technical differences, including randomly-occurring low-level enrichment of homopolymers and concentration-dependent multimeric binding (see first page of **Supplementary Document S3** for full description), but we observed no evidence for cofactor contamination or unexpected differences in sequence specificity.

In separate parts of the Codebook project, this same set of poorly-characterized putative TFs was analyzed using ChIP-seq, Protein Binding Microarrays^44^, and SMiLE-seq^45^, as summarized in **Table S7** and described in the accompanying manuscripts ^43,47,50^. Because the binding motif was not known in advance, we gauged the success of each protein in each experiment, including the GHT-SELEX experiments, based on whether similar DNA-binding motifs (i.e., PWMs) were obtained from different types of experiments, with all data types considered in aggregate by a team of expert curators^49^. Selection of a single PWM for each TF for subsequent analyses is also described in accompanying study^43^; PWMs logos, and information about their derivation are available in accompanying study^43^ and online at https://codebook.ccbr.utoronto.ca, https://mex.autosome.org, and https://cisbp.ccbr.utoronto.ca^51^.

In total, 139 of the 331 Codebook putative TFs had at least one successful GHT-SELEX experiment, of which 131 were also successful in HT-SELEX, 108 in ChIP-seq, and 102 in all three (**Figure 3a** and **Table S3**). The 139 were comprised mainly of C2H2-zf proteins, which are prevalent in the Codebook set (**Figure 3b)**. A total of 24 types of DBDs were present among the successful experiments (**Figure 3b**), however, illustrating that the method can capture motif-containing genomic target site locations of diverse TF families. An additional 163 of the putative TFs did not yield motifs in any of these three assays, however, despite the use of multiple expression systems and multiple constructs. Of these 163, only an additional nine were successful when including data from other assays used in the Codebook project (SMiLE-seq and Protein Binding Microarrays)^43^ (**Table S7**). Successful and unsuccessful proteins produced as eGFP fusions yielded largely overlapping range of fluorescence signals (**Extended Data Fig 3, Table S3**), and on western blots, ∼90% of sampled proteins exhibited a single prominent band at the correct molecular weight (**Table S8 and Supplementary Document S4)** for both successful and unsuccessful samples, suggesting that the majority are expressed and properly folded. It is possible that some of the putative TFs require post-translational modifications or cofactors, that our data analysis, motif derivation, and testing methods did not capture their specificity, or that they are false-negatives for some other reason. However, it is also possible that many of the tested putative TF simply do not bind DNA with sequence specificity, as in the majority of cases, they were also unsuccessful with other methods (**Table S7)**; if we assume so, then the success rate of GHT-SELEX on the Codebook proteins is 78% (139/177). Nonetheless, a central conclusion of this study is the identification of binding motifs for over 100 poorly characterized human TFs. For most of them, we produced three types of data (HT-SELEX, GHT-SELEX, and ChIP-seq) that provide differing perspectives on their sequence specificity and DNA binding.

**Figure 3.**
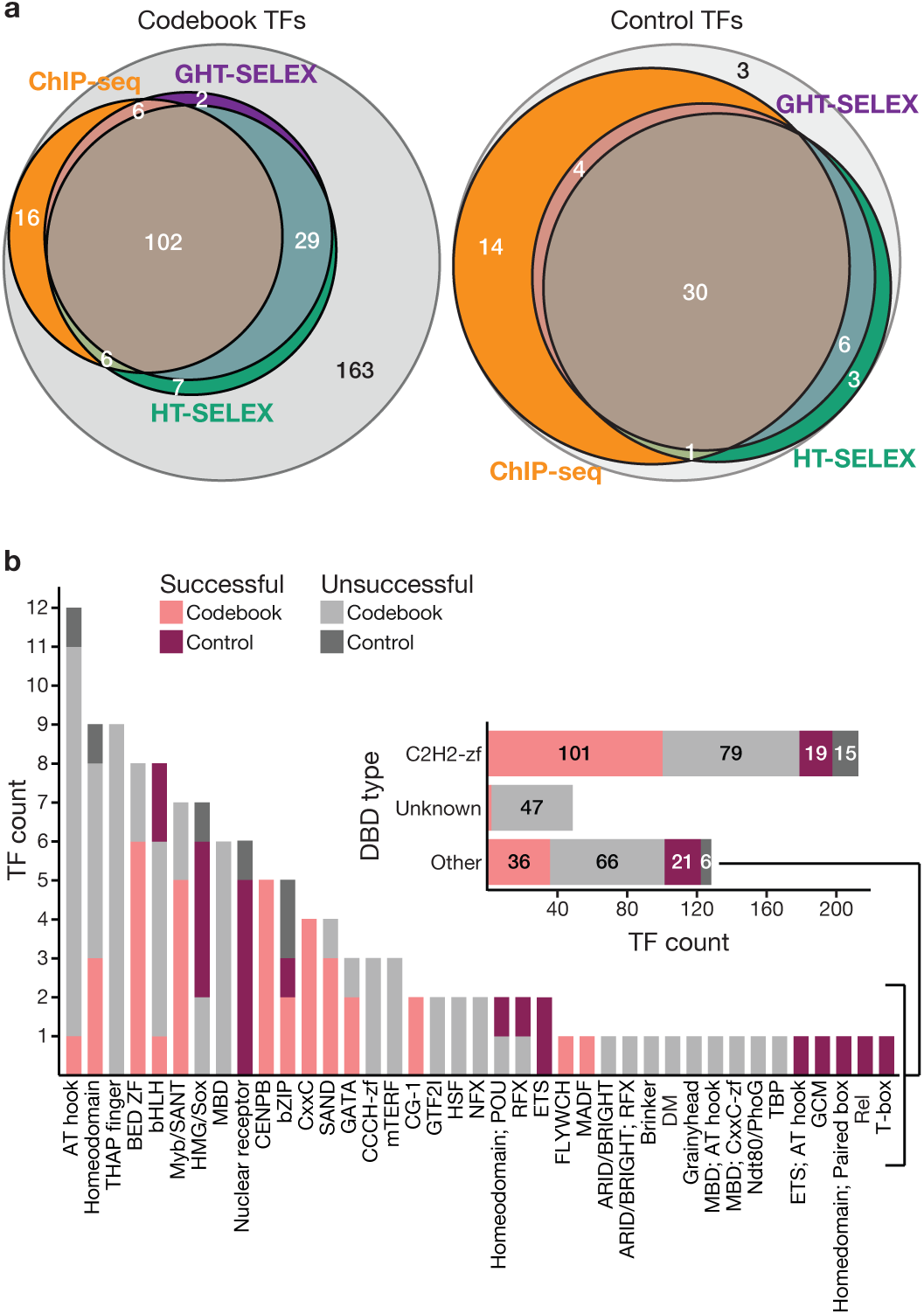
Analysis of 331 Codebook proteins and 61 control TFs using GHT-SELEX. **a.** Venn diagram displays the number of TFs with successful experiments in GHT-SELEX, HT-SELEX, and ChIP-seq for all Codebook TFs (left) and control TFs (right) assayed with GHT- and HT-SELEX. **b.** Bar chart shows the number of TFs with at least one successful GHT-SELEX experiment, categorized based by DBD type. C2H2-zf proteins and those with an unknown DBD (at the beginning of the project) are inset due to large numbers.

### Unexpectedly high overlap between TF binding to the genome *in vitro* and *in vivo*

We next sought to quantify the overlap between GHT-SELEX/MAGIX peaks and ChIP-seq peaks. For 137 TFs with GHT-SELEX data, encompassing 101 Codebook and 36 control TFs, ChIP-seq data in HEK293 cells were also available from a parallel Codebook study^52^, which employed heterologous expression in HEK293 cells. Scoring peak overlaps requires first defining peak sets, which is typically accomplished using a threshold on enrichment values or statistical significance. GHT-SELEX, like ChIP-seq, produces peaks with a continuum of enrichment coefficient values and other associated statistics, rather than a bimodal distribution that would more readily discriminate bound from unbound loci. The overlap with ChIP-seq peaks and the enrichment of PWM scores within peaks were also generally continuous (examples in **Figure 4a**; distributions for all TFs in **Supplementary Document S2**). Moreover, the MAGIX score and associated FDR values at which the overlap of GHT-SELEX peaks with either ChIP-seq peaks or PWM matches dropped to random values varied dramatically between TFs, such that no unifying thresholds emerged that could be generally applied convincingly to all TFs.

**Figure 4.**
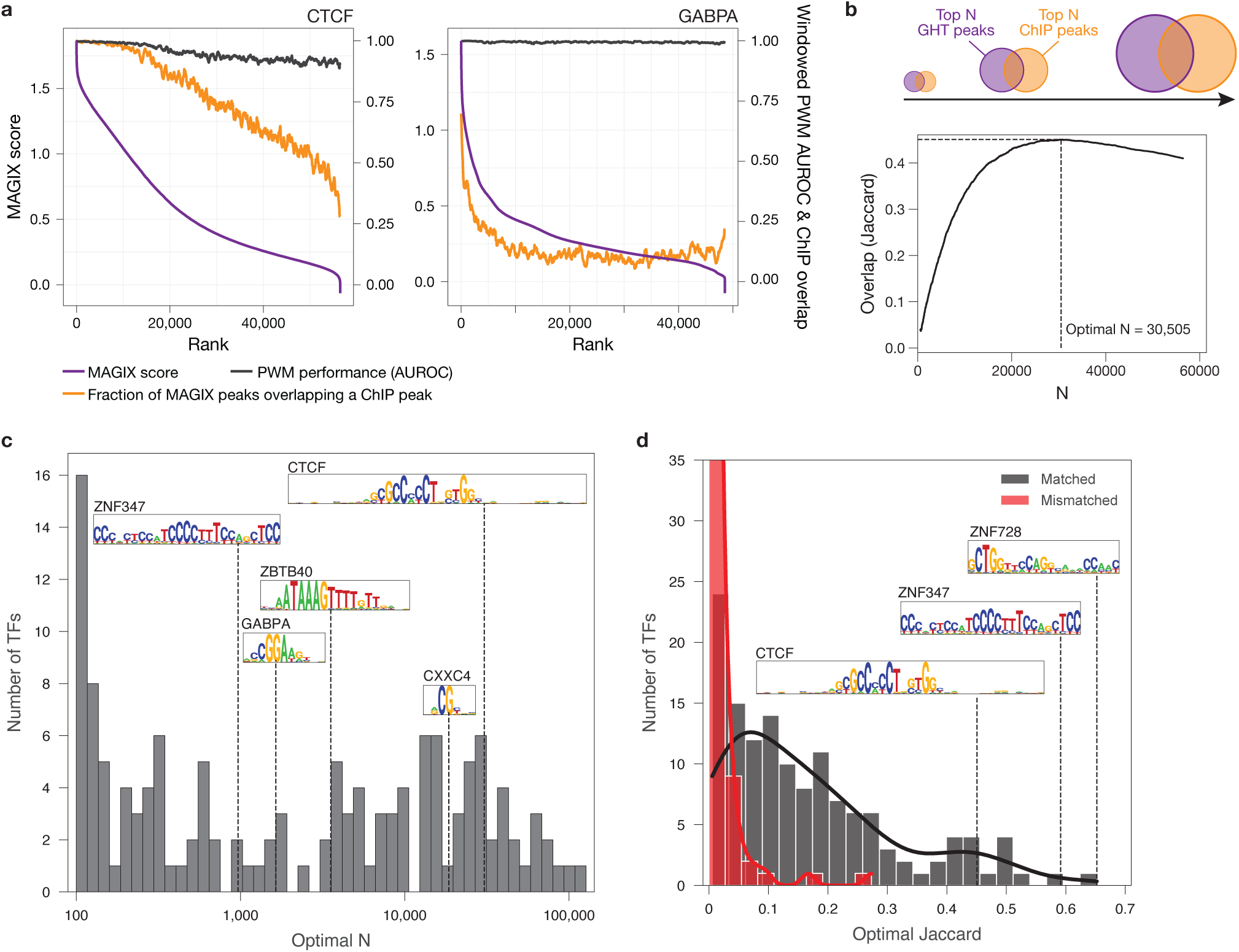
Correspondence between GHT-SELEX and ChIP-seq peaks. **a.** Enrichment of ChIP-seq peaks and PWM hits within MAGIX peaks, for two example control TFs. MAGIX peaks are sorted by their MAGIX score (purple, *left y-axis*). Orange line shows the proportion of peaks (in a sliding window of 500 peaks over the ranked peaks, with a step size of 50) that overlap with a ChIP-seq peak (at MACS threshold P < 0.001). Black line shows the AUROC for PWM affinity scores (calculated by AffiMx^52^) of MAGIX peaks in the same window vs. 500 random genomic sites. **b.** Illustration of peak number optimization (for CTCF as an example). **c.** Histogram of the optimal values of N (peak count) for the 137 TFs that have both GHT-SELEX and ChIP-seq peaks. **d.** Histogram of optimal Jaccard values, compared to the maximum Jaccard for mismatched TFs (i.e. between GHT-SELEX for one TF and ChIP-seq for a randomly selected TF).

As a workaround, we implemented a simple system to draw thresholds on the GHT-SELEX and ChIP-seq peak sets simultaneously and dynamically for each TF. This system identifies thresholds on a TF-by-TF basis by scanning through all numbers (N) of “Top N” peaks on a TF-specific basis, calculating the Jaccard statistic of overlap between the GHT-SELEX/MAGIX peaks and ChIP-seq peaks for each N, and choosing the peak number (N) with the maximum Jaccard value (**Figure 4b**). We posit that this approach accounts for unobserved TF-specific parameters in both GHT-SELEX and ChIP-seq assays, including different binding kinetics for both sequence-specific and nonspecific DNA binding, the effective concentration of the TF, and the ability of the TF to compete or cooperate with nucleosomes and other cofactors *in vivo*.

This approach yielded a very striking result, which is that for many TFs, a peak number can be identified with a surprisingly high Jaccard value (median 0.127 over all 137 TFs) (**Figure 4c,d**, and **Table S9**), meaning that almost 30% of GHT-SELEX peaks overlap with a ChIP-seq peak, and vice-versa. Peak overlap (i.e., Jaccard) is a demanding statistic, because random expectation (i.e., from choosing genomic regions at random) is near zero, as only a miniscule fraction of the genome is covered by the peaks in either data type, and both experimental variation and measurement noise will lead to fluctuation of the rank order of peaks. In permuted experiments (i.e., mismatched TFs), the Jaccard median is 0.00254 (Wilcoxon p=1.3×10^−39^) (**Figure 4c**). Using the same optimization process, the median Jaccard distance was 0.56 for ChIP-seq replicates^52^; this number presumably represents a typical upper bound. Overall, this outcome indicates that many TFs intrinsically (i.e. independently) specify many of their *in vivo* binding sites above a given TF-specific threshold. We do not expect the overlap to be perfect, due to the impact of chromatin and cofactors, but this result nonetheless contrasts with the traditional expectation that individual TF would not be able to independently specify the vast majority of their DNA targets in a genome^7^. The peak numbers yielding these high Jaccard values are often relatively low (**Figure 4d**), suggesting that the number of *in vivo* binding sites that are specified intrinsically by TFs is relatively small, and they correspond to a wide range of ChIP-seq p-value thresholds and MAGIX score values (**Table S9**), such that both GHT-SELEX/MAGIX and ChIP-seq data are needed to make these observations.

One potential explanation for this high overlap between GHT-SELEX and ChIP-seq peaks is that the Codebook dataset is dominated by C2H2-zfs, which often have long binding sites and thus intuitively high specificity. Indeed, among TFs with Jaccard > 0.1, 80% are C2H2-zf proteins (63 out of 79), vs 34% (20 out of 58) for those with Jaccard < 0.1, and overall, the median Jaccard value for C2H2-zf proteins is 0.1836, vs. 0.0559 for non-C2H2-zf proteins (Wilcoxon p=6.08×10^−9^). CTCF, a control TF that is known to possess large number of genomic target sites, unusually high intrinsic sequence specificity, and ability to control nucleosome positions^15,16^, is among those with high Jaccard values (0.45), although it is not the highest scoring in this dataset, and overall is clearly not unique in its capacity to independently specify where it binds in living cells. Counterintuitively, however, high Jaccard maxima were also obtained for a subset of TFs with relatively short motifs, including TERF1, ZNF48, ZBTB8A, NACC2, and several CXXC proteins, such as CXXC4 and KDM2A, that mainly bind CG dinucleotides, as expected^53^ (**Figure 5a**).

**Figure 5.**
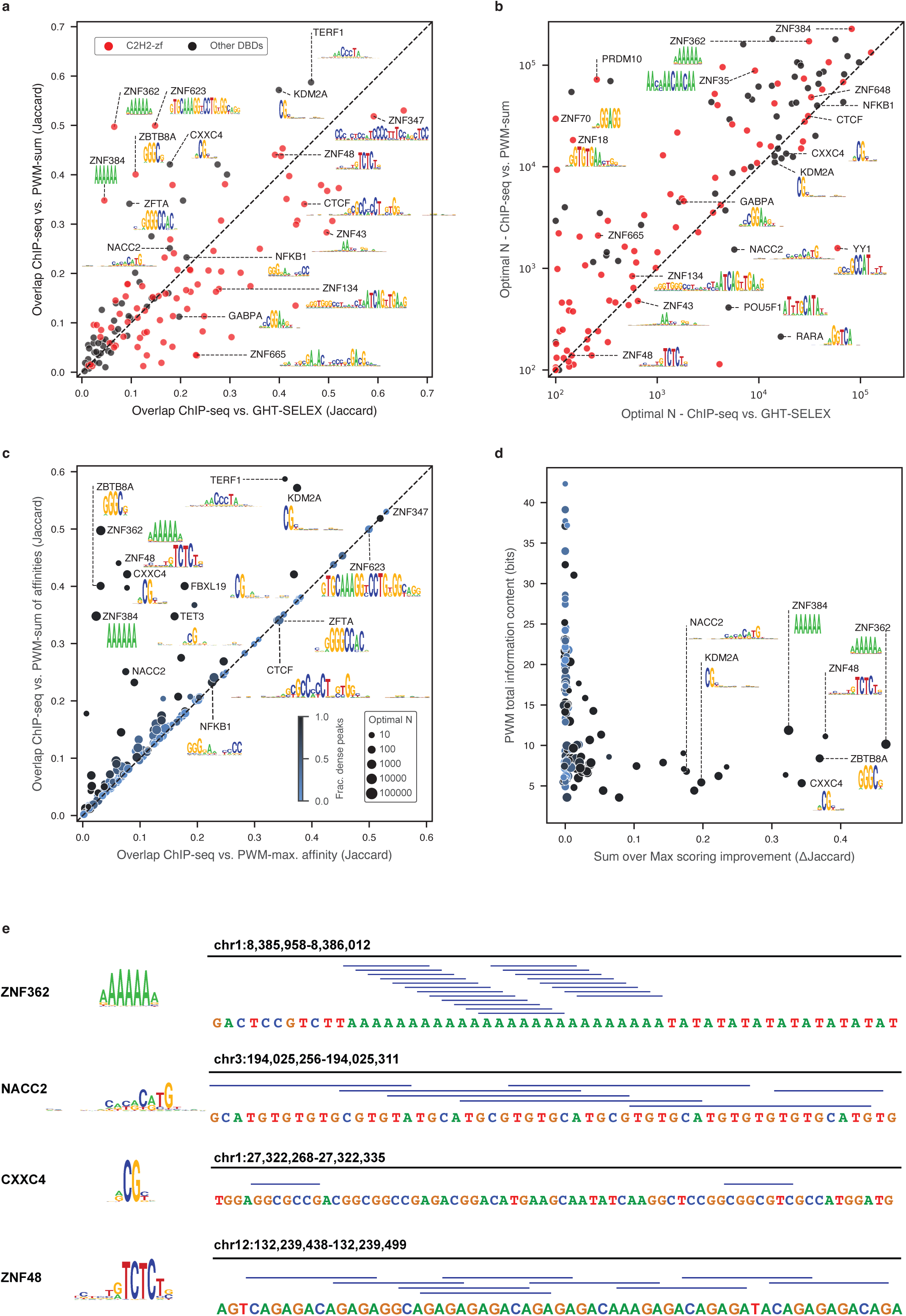
High quality PWMs often predict *in vivo* binding sites as effectively as GHT-SELEX peaks. **a.** Scatter plot of optimal Jaccard value between GHT-SELEX peaks and ChIP-seq peaks (x-axis) vs. optimal Jaccard value between PWM-predicted sites and ChIP-seq peaks (y-axis), for all 137 TFs (dots). **b.** Scatter plot of optimal N (peak number) for the same peak set comparisons shown in (**a**). **c.** Scatter plot showing optimal Jaccard value between PWM-predicted sites and ChIP-seq peaks, for maximum-affinity PWM scoring and sum-of-affinities PWM scoring. Points (TFs) are scaled based on the optimal number of peaks (in the sum scoring), and the color reflects the fraction of binding sites comprised of multiple PWM hits. **d.** Scatter plot of the improvement in the optimal Jaccard value associated with sum-of-affinities PWM scoring vs. information content of the PWM. Points’ size and color are the same as panel (**c**). **e.** Examples of four TFs with multiple motif matches within a single ChIP-seq peak.

### Multiple explanations for the high sequence specificity observed in GHT-SELEX

The traditional assumption that individual TFs have relatively low ability to specify their genomic targets is based largely on the results of motif scans^7^, which are dependent on the accuracy of the PWM. To revisit this assumption, we performed a maximization of the overlap between the Codebook PWM matches across the genome and the ChIP-seq peaks, using the same Jaccard-based method described above (see **Methods** for details). We found that the overlap between PWM predictions and ChIP-seq peaks is, in fact, similar to the overlap between GHT-SELEX/MAGIX and ChIP-seq peaks (**Figure 5a, Table S9**). The number of peaks at which the maximum Jaccard was obtained is also typically similar (**Figure 5b**). Thus, PWMs are often more accurate than generally believed, and comparable to the GHT-SELEX data. It is important to note that we selected the Codebook PWMs for their accuracy across datasets, i.e., other PWMs for the same TF would display lower overlap. Therefore, the PWM derivation method, including the experimental data employed, is critical for representing TF binding *in vivo*.

We also found that the motif scanning method is important to achieve <u>the</u> highest overlap between PWM matches and ChIP-seq peaks. Consistent with a recent report^54^, for some TFs, the sum of predicted affinity scores over a sequence window (i.e. the sum of the PWM probability scores at individual positions, rather than the log-odds that is output by most PWM scanning tools) results in a considerably higher maximum Jaccard value than taking the maximum or sum of log-odds PWM scores (which are generally thought to represent binding energy^55,56^) (**Figure 5c**). This Sum-of-affinity scoring presumably reflects the cooperation of multiple adjacent binding sites, traditionally referred to as “avidity”^57^. The effect is most striking for a subset of TFs that bind short or repetitive sequences, including CG dinucleotides and poly-A stretches (**Figure 5d**), but it also appears to underpin the specificity of NACC2 and ZNF48, which have unique, non-repetitive motifs (**Figure 5e**). Points above the diagonal in **Figure 5a**, in which PWM prediction shows higher overlap with ChIP than the GHT-SELEX, may therefore represent the impact of TF binding sites over a larger window, which would be detected in ChIP-seq but not GHT-SELEX (ChIP-seq fragments are 100-300 bp, and the PWM scans are performed with a 200 bp window, while the GHT-SELEX fragments are only ∼65 bp). For example, scanning 200-base windows with the short CG motif for CXXC4 may be better suited for the detection of CpG islands (which dominate the CXXC4 binding sites^43^, in which the CG dinucleotides will be distributed over a large region (by definition ≥200 bp).

In contrast – and despite evaluation of hundreds to thousands of PWMs for each TF - there were still many TFs for which we could not derive a PWM that rivals GHT-SELEX data in correspondence to ChIP-seq peaks (those below the diagonal in **Figure 5a**). These TFs are almost entirely proteins with a long array of C2H2-zf domains, which we examine more closely in the next section.

### Alternate usage of C2H2-zf domains within large arrays

The expansive collection of GHT-SELEX, HT-SELEX, and ChIP-seq data for C2H2-zf proteins provided an opportunity to examine the long-standing issue of the usage of individual C2H2-zf domains within large arrays. Anecdotally, we observed many instances where the motifs detected for C2H2-zf proteins were much shorter than expected based on the number of C2H2-zf domains, as well as examples in which multiple distinct motifs emerged, suggesting that the TFs might use partial subsets of their DBD array to engage DNA at different locations. Proving differential engagement of the specific C2H2-zf domains is challenging, however, due to low statistical power (there are many possible C2H2-zf domain sub-arrays, and a limited number of highly enriched peaks) and the fact that the genome is highly non-random and repeat-rich. To minimize the impact of these issues, we developed a new method that utilizes the C2H2-zf recognition code to assess which sets of C2H2-zf domains are likely to be engaged at any individual binding sites. We call this method RCADEEM (Recognition Code-Assisted Discovery of regulatory Elements by Expectation-Maximization) (see **Methods** for details). **Figure 6a** shows a schematic and the results of applying RCADEEM to CTCF, illustrating that it produces a “core” motif recognized by fingers 4-7 at all sites, and alternative usage of flanking C2H2-zf domains in a subset of sites, very similar to the differential usage of CTCF C2H2-zf domains that has been previously described^30^.

**Figure 6.**
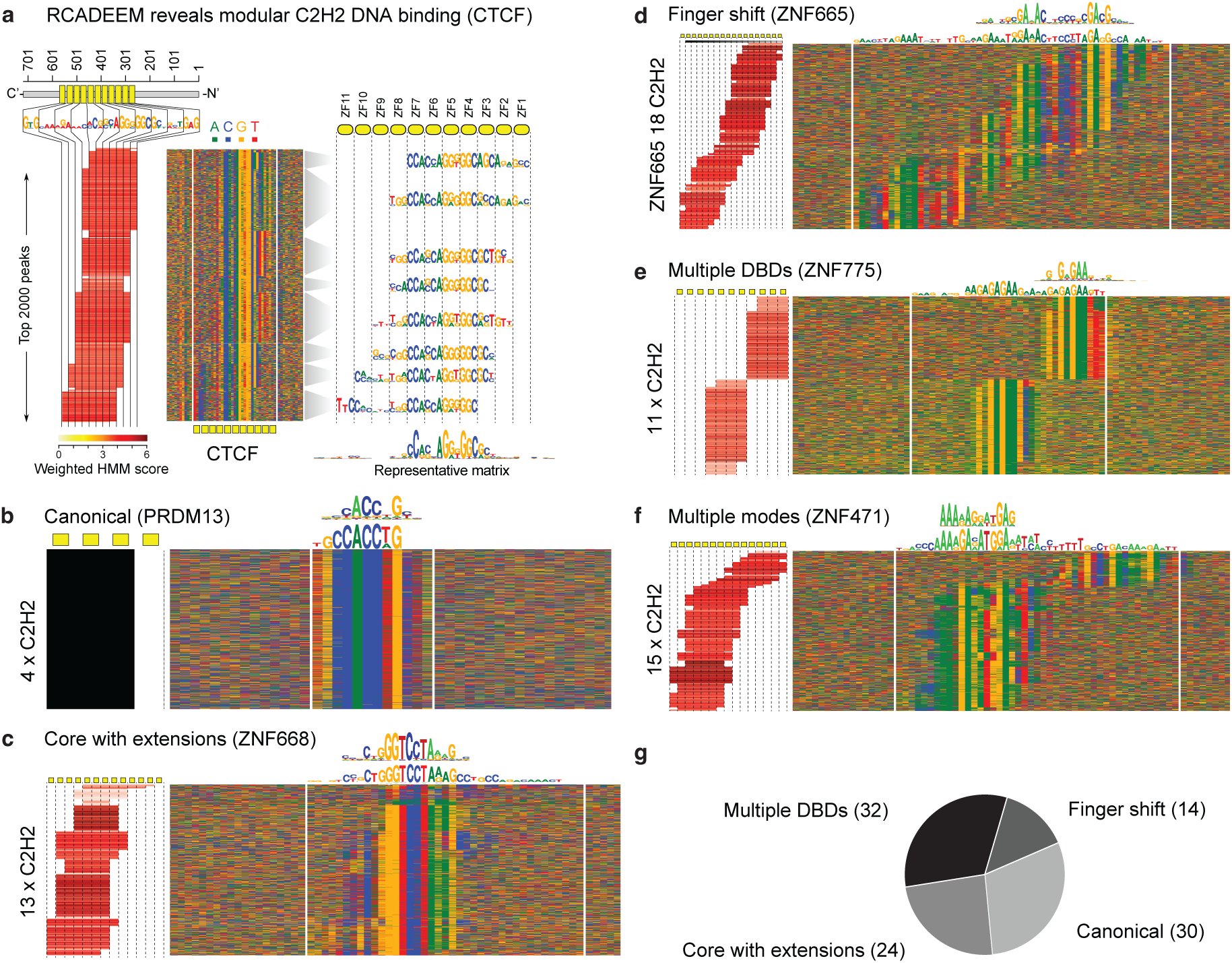
Alternative engagement of individual C2H2-zf domains at genomic binding sites inferred from the recognition code. **a.** RCADEEM applied to CTCF. *Middle* panel displays the top 2,000 nonrepetitive GHT-SELEX peaks. White vertical bars indicate the region that is expected to contact the DNA based on the assumption that each of the C2H2-zf domains define three contiguous bases. *Left* panel indicates which C2H2-zf domains are inferred to engage each DNA sequence, which is used to determine the row order in the figure. *Right* panel shows motifs for the major sub-sites, derived from base frequencies in the sequence alignment. **b-f,** Top 2,000 non-repeat peak sequences, as in (**a**), for representative TFs with different binding modes, as described in the main text. Above each is shown the sequence logo for the single representative Codebook PWM (*top*) and a motif generated by RCADEEM that represents all of the observed sequences (*bottom*). **g.** Number of occurrences of each category among all 86 C2H2-zf proteins for which RCADEEM yielded a significant outcome; note that a TF might appear in multiple or no categories.

We applied RCADEEM to all 120 C2H2-zf proteins for which we had successful data from GHT-SELEX (**Table S10).** We applied RCADEEM on GHT-SELEX data and separately, if available, on HT-SELEX and ChIP-seq; for GHT-SELEX and ChIP-seq, we applied it both with and without repeat sequences (i.e., removing any peaks that overlap with the UCSC RepeatMasker track). In total, we obtained RCADEEM predictions for 86 of them (**Table S10**), all of which are available via the web resources accompanying this paper (https://codebook.ccbr.utoronto.ca/). (For the remaining 34, the algorithm did not converge, suggesting that the sequence preferences of the protein do not closely follow the recognition code, and thus cannot be analyzed in this way). Most of the 86 displayed what appears to represent alternative usage of segments of the C2H2-zf domain array on different DNA molecules (e.g., different genomic loci) within the same experiment (**Supplementary Data S2**). These patterns could not be accounted for by truncations or other potential artifacts (**Document S4**).

We manually classified the apparent C2H2-zf domain usage into the following categories, examples of which are shown in **Figure 6b-f**. **Figure 6g** provides an overview of the descriptors and other properties of each of the C2H2-zf proteins. 1) *Canonical (30 instances)* follows the baseline assumption that a TF always uses the same set of C2H2-zf domains to recognize sites that can be described with a single PWM. 2) *Core with extensions (24 instances)*, where all sites share a sequence motif bound by a subset of the C2H2-zf domains, which is supplemented by recognition of flanking sequences by adjacent C2H2-zf domains at some binding sites. 3) *Finger shift (14 instances),* where the TF recognizes a range of tiled target sites by binding with variable subsets of adjacent C2H2-zf domains. 4) *Multiple DBDs (32 instances)*, in which subsets of the C2H2-zf domain array appear to function as independent DBDs. The last three binding modes are not mutually exclusive. For example, ZNF471 displays both multiple DBDs and core with extensions with one of the DBDs (**Figure 6f**), while the long finger shift in ZNF665 (**Figure 6d**) leads effectively to multiple DBDs, as the target sites of most N-terminal and C-terminal ends do not overlap with each other. **Table S10** lists the annotations for all 86 proteins.

### Evolution of C2H2-zf protein DNA-binding specificities via internal duplication

In the RCADEEM outputs, different segments of a C2H2-zf domain array (i.e., different DNA binding regions of the protein) are often predicted to bind similar yet distinct sets of sequences. For example, ZNF775 (**Figure 6e**) binds two types of sites that contain a shared GNWGAA consensus, followed by either TTT or GCA trinucleotides. RCADEEM predicts that these two sites are recognized by C2H2-zf domain arrays 1-4 and 5-8, respectively. Indeed, arrays 1-3 and 5-7, as well as 9-11, are homologous, on the basis of sequence identity (visualized at https://codebook.ccbr.utoronto.ca/details.php?TF=ZNF775), suggesting that they arose from duplications. All three arrays are present in mammals as distant as the Tasmanian devil, indicating that the duplications predate divergence from marsupials and have since been conserved. The cellular and physiological functions of this protein are unknown, to our knowledge, but this degree of sequence conservation suggests a conserved role across mammals.

Another example is ZNF721: RCADEEM indicates that it has three DNA-binding modes, with related but distinct motifs (**Figure 7a**), corresponding to homologous C2H2-zf domain arrays containing fingers 6-13, 12-16, and 18-22 (**Figure 7b**). The distinct sequence preferences of the duplicated ZNF721 arrays are supported by experimental data for partial “DBD1” and “DBD2” constructs, corresponding roughly to the first and second half of the full array, which recognize largely distinct subsets of the genomic sites bound by full length TF in GHT-SELEX (**Figure 7a**) and prefer almost entirely distinct 10-mers in HT-SELEX (**Figure 7c**). The function of ZNF721 has not been determined, but sequences recognized by the first (6-13) and third (18-22) duplicated C2H2-zf domain arrays of ZNF721 are found in the highly numerous Alpha repeats, which are fast-evolving elements found at primate centromeres^58^. ZNF721 itself is present only in primates. ZNF721 also binds thousands of unique loci outside known repeat elements, and associates physically with TRIM28/KAP1^59^, suggesting a role in gene silencing or heterochromatin formation.

**Figure 7.**
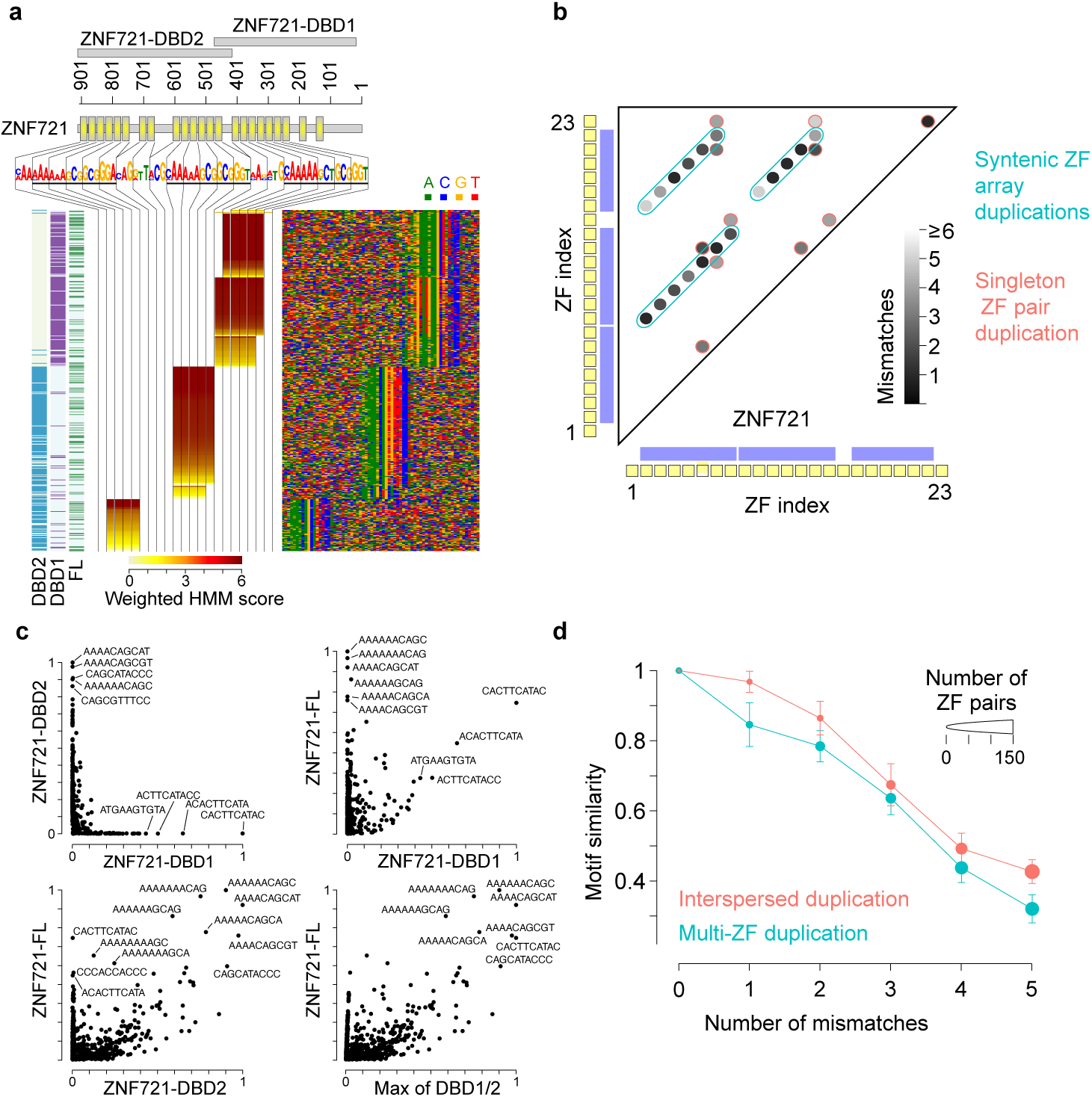
Evolution of C2H2-zf protein DNA-binding specificities through internal duplication of DBDs and DBD arrays. **a.** RCADEEM results for the pool of top 500 peaks from full-length ZNF721 and two DBD constructs, after removing peaks that overlap repeats. The construct in which the peak was observed is indicated on the left. **b.** Similarity of C2H2-zf domains of ZNF721 (based on the number of mismatches in the global alignment). Apparent duplicated arrays are encircled by blue border (i.e. syntenic duplications), while single pairs that may also be duplicates are circled in red. **c.** Scatterplots of the HT-SELEX *k*-mer scores^65^ (relative counts) across the three ZNF721 constructs **d.** Comparison of average per-base similarity (correlation of nucleotide frequency) in PWMs predicted by the recognition code, for those present in duplicated arrays vs. those duplicated as individual C2H2-zf domains, with duplicated C2H2-zf domains taken as pairs that are separated from each other by 5 or less edits. DBD pairs have been filtered to contain only combinations where both DBDs are likely to have retained their ability to bind DNA (have DNA binding functionality score^68^ >0.5). Error bars show standard error of the mean.

To survey the prevalence of internal duplication of C2H2-zf domains, we compared all pairs of individual human C2H2-zf domains occurring in the same protein and found that 185 human C2H2-zf proteins (∼25%) contain at least one pair of C2H2-zf domains that differ by 3 or fewer edits (substitutions, deletions or insertions; **Table S10**), suggesting that they are derived from recent duplications. Furthermore, as in ZNF775 and ZNF721, there are 140 proteins with apparent internal C2H2-zf domain array duplications, defined as two (or more) adjacent C2H2-zf domains (i.e., an array) related to a second such array with two (or more) C2H2-zf domains, with 5 or fewer edits per C2H2-zf domain. Based on recognition code predictions, C2H2-zf domain arrays within internal array duplications have more diverged sequence specificities from each other than individually duplicated C2H2-zf domains (**Figure 7d, Table S11**The prevalence and diversification of internal C2H2-zf domain array duplications suggest that they are a common modality for evolution of novel functional roles for this large class of proteins.

## DISCUSSION

GHT-SELEX assays direct and unassisted binding of single TFs to the unmodified and unchromatinized genome *in vitro*, revealing surprisingly specific intrinsic sequence preferences for many human TFs. The assay, and the associated MAGIX analysis pipeline, offers several technical advantages over alternatives, including smaller fragment size and compatibility with the same instrumentation used for HT-SELEX. GHT-SELEX data are often more similar to *in vivo* TF binding data than previous assessments would suggest they should be^7,26^, indicating that, for an apparently large subset of TFs, chromatin and cofactors have less critical influence on where binding occurs. This same observation implies that this subset of individual TFs may have a greater ability to overcome the chromatin state than is commonly believed.

The unexpectedly high overlap between ChIP-seq, GHT-SELEX, and PWM scans could be partly explained by technical shortcomings in previous PWM-based genome scans. HT-SELEX and other *in vitro* approaches utilizing random sequences are powerful in that they are unbiased in terms of sequence composition^60^, but they are inherently limited in sequence length and context that can be surveyed. ChIP-seq is invaluable because it can assay binding within cells, but it does not inherently discern direct, indirect, and non-specific binding. Thus, PWMs derived from ChIP-seq and other *in vivo* approaches are influenced by factors other than the TF, in addition to the biased sequence content of the genome. GHT-SELEX provides a powerful intermediate that can resolve ambiguities of both motif discovery and PWM scanning, and thus provides data that complements both ChIP-seq and *in vitro* assays that utilize random sequences. The PWM scanning method is of particular importance: taking the sum of predicted affinity over longer sequence windows is critical to capturing positions that are bound *in vitro* and *in vivo* by CXXC-zfs and other proteins that recognize clusters of simple sequence motifs.

GHT-SELEX is particularly effective with C2H2-zf proteins, and, together with RCADEEM, has an unprecedented ability to both obtain and dissect *in vitro* the multiple binding modes that are characteristic of this family, and inherently more difficult to represent as a single PWM. The existence of multiple binding modes also provides a potential explanation for the large number of C2H2-zf domains in each protein. In at least some cases, these large arrays derive from internal duplications of segments of the C2H2-zf domain arrays, possibly facilitating generation of evolutionary novelty via duplication and divergence.

In contrast to the C2H2-zf family, the most well-studied TFs tend to be in the TF classes such as homeodomain, bHLH, bZIP, nuclear receptor, and Sox TFs, because they are the most strongly conserved and often dictate specific biological processes (e.g., morphogenesis, body plan, lineage specification, etc.)^26^. Our study included some of these TFs (e.g., LEUTX, BATF2, RARA and SRY), and they displayed only limited overlap between GHT-SELEX and ChIP-seq peaks, indicating that many of them cannot independently specify *in vivo* binding locations and hence target genes. It has long been known that TFs controlling chromatin in yeast are largely distinct from those that regulate specific pathways; we speculate that a similar division may exist in human and other animals.

GHT-SELEX data, together with the larger Codebook dataset, provide an extensive new dataset of TF motifs (i.e., PWMs), encompassing most putative TFs currently lacking them. Accompanying papers provide a thorough analysis of the results of this project, which underscore many challenges and benefits of accurate motif representations. Representation of TF sequence specificity remains an open challenge, more than four decades after the introduction of the standard PWM model^61^. More accurate representations of multimeric binding and large and complex binding sites, in particular for C2H2-zf proteins, will be useful for a variety of purposes. We propose that obtaining data from GHT-SELEX for additional TFs with “known” motifs and genomic binding sites from ChIP-seq will produce a more detailed view of their intrinsic DNA binding abilities and how this intrinsic ability dictates TF-genome interactions in living cells.

## METHODS

### TFs and constructs

Selection of TFs, design of constructs for gene synthesis, and expression vectors are described in the accompanying study^43^. Sequences and other information are available as described below in Data Availability.

### Protein production and quality control

We used three protein expression systems, which we refer to in **Table S2** and below as *Lysate*, *IVT*, and *eGFP-IVT*, respectively. The *Lysate* system employed recombinant HEK293 cells, created in the accompanying study^47^, and in a previous study^48^, which express eGFP-tagged full-length proteins from a Tet-inducible promoter (plasmid backbones pTH13195^48^ and pTH12027). We induced expression by Doxycycline treatment for 24 hours prior to harvest, and confirmed via fluorescent microscopy. Whole cell lysates were then harvested from a 10cm plate (∼10 million cells) for each line using 1 ml of lysis buffer (50 mM Tris-Cl at pH 7.4 containing 150 mM NaCl and 1% Triton X-100), supplemented with protease-inhibitor cocktail (Roche cOmplete mini, 04693159001), as described previously^37^. Each of the SELEX cycles used 50 <u>µ</u>l of lysate. *IVT* used an *in vitro* transcription-translation reaction (PURExpress In Vitro Protein Synthesis Kit, NEB, Cat# E6800L) to express T7-driven, GST-tagged proteins (either full-length or DBDs) (plasmid backbone pTH6838^51^). *eGFP-IVT* employs the TNT SP6 High-Yield Wheat Germ Protein Expression System (Promega, Cat# L3260) to express SP6-driven, eGFP-tagged proteins (either full-length or DBDs) (plasmid backbone pTH16505, an SP6-promoter driven, N-terminal eGFP-tagged bacterial expression vector, modified from pF3A–eGFP ^45^ to contain AscI and SbfI restriction sites after the eGFP. For *IVT* and *eGFP-IVT* production systems, we performed reactions according to kit instructions, but using a smaller volume: 7.5 μl of *IVT* or 5 μl of *eGFP-IVT* reaction sample was used in each binding reaction of each SELEX cycle. Fluorescence levels shown in **Table S3** and **Extended Data Fig. S3** were measured using a BIOTEK Synergy Neo microplate reader from 10 μl volume of lysate or wheat germ extracts on a conical transparent 96-well plate using bottom optics and with a gain setting of 80. A subset of these proteins was also analyzed on Western blots (**Table S7**, **Supplementary Document S4)** with the same anti-GFP antibody (ab290, Abcam) as used in SELEX, and a secondary horseradish peroxidase fusion antibody (Cat Nr. Anti-Rabbit IgG HRP Conjugate, Cat W401B, Promega) and Immobilon Western Chemiluminescent HRP Substrate (WBKLS0050, Millipore Sigma). Blots were imaged with Odyssey Fc Imager (LICORbio).

### GHT-SELEX and HT-SELEX library preparation

We fragmented HEK293 genomic DNA (Genscript, USA; Cat. No. M00094) for 45 minutes using NEBNext dsDNA Fragmentase enzyme mix (NEB, M0348S), and then performed a size selection step to reduce the amounts of fragments larger than 200 bp. In the size selection we added 0.9X volume of bead suspension (magnetic SPRI beads, supplied with the kit, NEB, E7103S) to the fragmented DNA, mixed the reaction for a minute, and then removed the large DNA fragment bound beads with a magnet, after which we diluted the supernatant 5X with water, followed by purification with a PCR purification kit (NEB, T1030S), to recover fragments as small as 25 bp. Next, the fragments were converted to an Illumina sequencing compatible library using NEBNext® Ultra™ II DNA Library Prep kit (NEB: E7103S) and NEB E7350 adapters. After adapter ligation, we purified the library with a PCR purification kit (NEB, T1030S) and then amplified it for five PCR cycles to convert the partially single-stranded adapter flanks to fully double-stranded DNA, to increase the amount of the product and reduce the amount of methylated cytosine residues in the initial library. The ninety-six (96) HT-SELEX ligands were prepared as described^62^, with the exception that the reverse primer was replaced with a primer (5’-CTGGAGTTCAGACGTGTGCTCTTCCGATCT-3’) that does not contain a T7 promoter sequence. HT-SELEX ligands differ from each other by containing a well-specific variable region that flanks the randomized 40 bases indicated in the name of the experiments (e.g., AA40NCCAGTG contains 40 bases flanked by AA and CCAGTG sequences and partial Illumina adapter sequences). All primers and library preparation schemes are given in **Table S1**.

### HT-SELEX and GHT-SELEX

We modified protocols from a previously-described HT-SELEX procedure^37^. HT-SELEX and the GHT-SELEX ligands contain the same flanking constant regions and thus there were no differences in the selections or sequencing library preparations. We conducted the magnetic bead washing operations below using a Biotek 405TS plate washer fitted with a magnetic carrier. We performed 21 different batches of SELEX, which varied in some technical respects in order to accommodate the three protein production systems and to implement improvements developed during the study (See **Table S2** for description of conditions used in each experimental batch). Protein immobilization was carried out in buffers based either on Lysis buffer (150 mM NaCl and 1% Triton X100 in Tris-Cl, pH 8) or Low stringency binding buffer (LSBB)(140 mM KCl, 5 mM NaCl, 1 mM K2HPO4, 2 mM MgSO4, 100 µM EGTA, 1 mM ZnSO4, and 0.1% Tween20 in 20 mM HEPES-HCl (pH 7). All DNA-protein reactions used LSBB. For GST-tagged proteins, we used glutathione magnetic beads (Sigma-Aldrich G0924-1ML), and for GFP-protein immobilization, we used GFP-Trap Magnetic Agarose” (Chromotek, gtma-100) for initial batches, and Anti-GFP antibody (ab290, Abcam) immobilized to Protein G Mag Sepharose® Xtra (Cytiva, 28-9670-70) for later batches, as the latter showed <u>a</u> higher success rate. All selections used 1μl of the magnetic bead slurry, a volume that in majority of the cases, according to manufacturers’ information, contains excess protein binding capacity but is still visible in microwell plates, allowing quality control of the washing steps.

*SELEX process*: All of the protocols (described in **Table S2)** followed these general steps: 1) Affinity beads and 96-well plates were blocked with BSA for 15 minutes; 2) Beads and plates were washed to remove unbound BSA; 3) Protein was immobilized into beads for 1h on a shaker; 4) Beads were washed to remove nonspecific proteins and carryover DNA; 5) Protein coated beads were incubated with DNA ligand for 1h to allow the proteins to bind their target sites; 6) Unbound and weakly bound DNA ligands were removed with extensive washing; 7) DNA ligands were eluted by suspending the beads into heat elution buffer (0.4 μM forward and reverse primers, 1 mM EDTA and 1% Tween 20 in 10 mM Tris-Cl, pH 8) transferring the suspension into a conical PCR plate and heat treating it in a PCR machine using a program that cycled between temperatures of 98 and 60°C, in order to denature the proteins and DNA, use convection to drive the DNA into the solution, and to hybridize DNA to the amplification primers; 8) Bead suspension obtained from heat elution was used as template in PCR and qPCR reactions; 9) An additional DNA amplification cycle was performed with 2X more primers and dNTPs to ensure that majority of the ligands are in fully double-stranded state and 10) In batch YWO and an associated batch YWN, we tested a set of same proteins with (YWO) or without (YWN) mung bean nuclease treatment to digest partially single-stranded ligands from the experiments, in order to reduce enrichment of DNA with artifactual sites, such as potential aptamers. This strategy proved effective in HT-SELEX experiments and was used in the following batches (P, Q, R, and S). For these last four batches, the TFs were mainly in alphabetical order. In each mung bean nuclease reaction, the pH of the solution (PCR reaction) was first lowered by the addition of 1:10 volume of 100 mM acetic acid, followed by addition of 1μl (0.75 units) of the enzyme and incubation for one hour at 37°C.

### Sequencing

Samples were prepared for sequencing by performing a PCR reaction that indexes each sample and its selection cycle with a unique combination of i7 and i5 barcodes, followed by a double-stranding reaction with primers that target regions of DNA outside indices (**Table S1**). Following this step, DNA libraries were pooled, purified with a PCR purification kit (NEB, T1030S), and then subjected to Illumina sequencing with 60bp reads at ∼3M reads per sample (Donnelly Centre sequencing core facility).

### HT- and GHT-SELEX read processing and mapping

HT-SELEX reads were filtered by Phred quality score (Q >= 30 in at least 90% of bases). GHT-SELEX reads were parsed with Trimmomatic^63^ to remove the constant regions from genomic fragments that were shorter than the sequencing read length (options: ILLUMINACLIP:CustomAdapters.fa:2:5:5, LEADING:3, TRAILING:3 MINLEN:25). The custom adapters in the fasta file were AGATCGGAAGAGCACACGTCTGAACTCCAG and AGATCGGAAGAGCGTCGTGTAGGGAAAGAGTGTTA. For GHT-SELEX, we mapped trimmed reads to the human genome build hg38 with bowtie2 (options: --very-sensitive, --no-unal). The mapped reads were further filtered using Samtools (options: - F 1548, version 1.20)^64^. Data were processed further for MAGIX-based analyses: each cycle of individual experiments was filtered through, sorting mapped fragments by read name, deduplicating them by position (samtools command: markdup, option: -r), and keeping only properly paired reads (option: –f 2).

### HT-SELEX k-mer enrichment-based comparison of protein production methods

To identify binding specificity differences conferred by either protein production method or using a different clone type that contains all predicted DBDs (DBD or FL), we identified 62 TFs for which we had at least one pair of successful experiments performed with a lysate HT-SELEX experiment and another with either of the IVT systems. For each successful experiment performed for these 62 TFs, we counted occurrences of 3-, 5-, 7-, and 9-mers in the unique reads contained in the final two cycles, as well as in the corresponding DNA library (cycle 0) using jellyfish (v2.3.1)^65^. For all possible experiment comparisons, log2 fold change was compared for each k-mer size.

### MAGIX statistical framework

At the core of MAGIX is a generative model that explicitly connects the enrichment of TF-bound genomic intervals to the fragment counts observed across GHT-SELEX cycles. MAGIX models how TF-bound intervals progressively occupy a higher proportion of the selected fragments pool in each cycle relative to the genomic background. These fragment proportions, in turn, are treated as latent variables in the model that, together with a sample-specific library size factor, determine the number of observed reads through a Poisson process. Consider the genomic interval *j*∈[1,*G*], where *G* is the total number of unique genomic intervals that we are modeling. Assume that fragments originating from interval *j* have a starting abundance of *aj* in the library. We also assume an exponential enrichment for the fragments, that in each cycle of SELEX, the abundance of these fragments changes by a factor of *e^bj^*, where *bj* is the log fold-change in abundance per cycle (referred to as *enrichment coefficient*), conceptually associated with biophysical parameters such as binding energies. Therefore, at cycle *t*, the abundance of the fragment originating from interval *j* is given by:

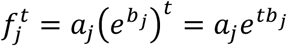

For convenience, we work with the logarithm of abundance, *y_j_*=log *f_j_*, transforming the exponential equation above to a linear equation as follows:

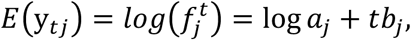

which can be seen as the linear multiplication of a feature vector **x***_i_* = [1 *t_i_*] for all the samples *i* (and across different cycles) corresponding to the TF of interest, and the interval-specific parameters **β**_*j*_ = [log *a_j_ b_j_*].

We note that to accurately model the enrichment of each fragment per cycle, other factors also need to be taken into consideration, such as background or batch effects, and therefore, the linear equation above needs to be fitted not only to the samples that correspond to the TF of interest, but also <u>to</u> samples from other experiments. We embed these dependencies in a design matrix ***X***∈ℝ*^N^*^×*K*^, where *N* is the total number of samples and *K* is the number of variables to consider, including an intercept term (whose coefficient will correspond to log *aj* above), a term for the SELEX cycle *t* (variable for the samples corresponding to a same TF), and other terms for batch and background effects. In addition to the variables included in ***X***, the abundance of each fragment in each sample depends on a sample-specific scaling factor that is often referred to as the library size. Assume that this library effect, for each sample *i*∈[1,*N*], is the scaling factor *si* (in logarithmic scale). Therefore:

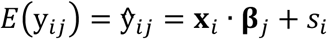

Here, *ŷ_ij_* corresponds to the expected logarithm of the abundance of interval *j* in sample *i*, **x***i*∈ℝ*^K^* is a vector representing the *i*’th row of the design matrix **X** (i.e., the sample-level variables for sample *i*), and **β***_j_*∈ℝ*^K^*is an interval-specific vector of coefficients for the *K* variables included in the model.

We note that the equation above does not have a unique solution. For example, any Δ**β** can be added to **β***_j_*, followed by subtraction of **X**Δ**β** from **s**, without any change in **ŷ***_j_*:

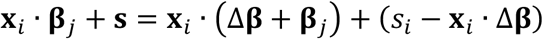

Therefore, to make the model identifiable, we limit **β***j* so that Σ_*j*∈[1,*G*]_**β***_j_*=**0**, where **0** is the zero vector of length *K*. This constraint is also useful since it means that, across all *G* intervals, the mean of each coefficient in **β**, including the coefficient for the SELEX cycle, is zero; in other words, the enrichment per cycle for each interval is calculated relative to the mean of all *G* intervals.

To incorporate the experimental noise in the logarithm of the abundance of interval *j* in sample *i* (i.e., building the real distribution of y*_ij_*), we modeled it as a Gaussian random variable whose mean is given by *ŷ_ij_* (the linear model above) with a sample-specific variance *σ^2^_i_*:

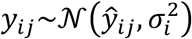

To complete the Bayesian framework, we also assume a multivariate Gaussian prior for **β***_j_*:

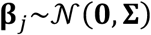

Here, **Σ**, the covariance matrix of the prior distribution, is shared across all intervals.

Altogether, the equations above form the following Bayesian model:

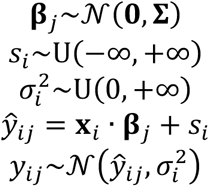

Assuming that the values for *y_ij_* are directly observed, we can obtain the maximum a posteriori (MAP) estimates of the parameters **β***_j_* (for *j*∈[1,*G*]), **s**, and *σ_i_*^2^ (for *i*∈[1,*N*]) through a block coordinate descent algorithm, as previously described^48^. The prior covariance matrix **Σ** is a hyperparameter that is obtained using an empirical Bayes approach. More specifically, we first obtain the maximum likelihood estimate (MLE) of the parameters **β***_j_*, *s_i_*, and *σi*^2^ without assuming any prior on **β***j*, and then estimate the values of *σ* ^2^, the variance of each element *k* in **β** , using the MLE solutions of all **β***_j_* coefficients. The covariance matrix **Σ** is then constructed as **Σ**=diag(*σ*_*β1*_^2^,…, *σ*_*βK*_^2^).

We note, however, that the log-abundance values *y_ij_*‘s are not directly observable in GHT-SELEX data. Instead, we observe *m_ij_*, the count of reads mapping to interval *j* in sample *i* (see **HT- and GHT-SELEX read processing and mapping**). This parameter adds another step to the framework, leading to the following hierarchical Bayesian model:

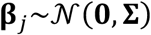

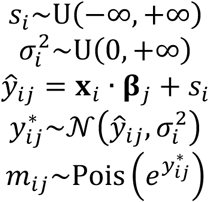

Here, *y*_ij_* is the logarithm of the true abundance of fragment *j* in sample *i*, which is latent. We obtain the MAP estimates of the parameters **β***_j_* (for *j*∈[1,*G*]), *si*, and *σ_i_*^2^ (for *i*∈[1,*N*]) using an expectation maximization (EM) algorithm, in which at each E-step we obtain the expected value of each *y*_ij_* given the observed read count *m_ij_* and the current model parameters, followed by re-estimation of the model parameters in the M-step, similar to a previous method established for EM optimization of Poisson-lognormal models^48^.

### Identifying GHT-SELEX peaks with MAGIX

The statistical framework described above calculates the rate of enrichment across GHT-SELEX cycles for a set of given genomic intervals, e.g., a set of candidate peak regions. Below, we describe how we identify such candidate peaks.

For each TF, an aggregate BAM file is created across replicates and cycles. For each genomic region with continuous non-zero coverage in this aggregate BAM file, the maximum coverage is calculated. Regions with a maximum coverage smaller than a threshold are discarded, with the threshold being set to the total fragment count in the aggregate BAM file divided by 2×10^6^. Then, per region, the coordinate with the highest coverage is defined as the summit. The candidate peaks are defined as the 200 bp regions centered on the summits.

Once the candidate peaks are identified, their count profiles are calculated from the original (unaggregated) BAM files (per cycle and replicate) and used as input to MAGIX to calculate MAGIX scores (enrichment coefficients), using prefixed library sizes that are estimated by fitting a similar MAGIX model to count profiles of 200-bp non-overlapping genomic intervals (a total of ∼13M bins). For each candidate peak, we also calculate a P-value, representing the statistical significance of the enrichment coefficient (<u>the</u> null hypothesis is that the enrichment coefficient is zero). To do so, we obtain<u>a</u> maximum likelihood estimate of the model coefficients and perform a likelihood ratio test (LRT) against a reduced model in which the enrichment coefficient is restricted to zero. The source code for MAGIX is available at https://github.com/csglab/MAGIX.

MAGIX peak sets analyzed throughout these papers and provided at codebook.ccbr.utoronto.ca are derived from a merge of all successful experiments that contain full set of TF’s DBDs (marked with “DBD” or “FL”) prior to running MAGIX. Experiments performed with specific subsets of DBDs (Marked with “DBD1”, “DBD2”, or “DBD3”) are analyzed separately.

### Selection of thresholds for peak sets

We sorted the GHT-SELEX peaks by their MAGIX score, or as named in the peaks BED files, *coefficient.br*, which estimates cycle enrichment. Similarly, we sorted the merged ChIP-seq peaks by P-value. Then, for different values of N (between 100 and the total number of peaks), we took the top N peaks for both peaks sets and calculated the Jaccard index (= O/(2N-O), in which O is the intersection of peaks). To eliminate the error in the cases when one peak in a set overlaps multiple peaks in another set, we used the average of the overlaps for the intersection (i.e., O=(O1+O2)/2, in which O1 is the number of peaks in set1 overlapping with any peaks in set2 and vice versa). The value of N that yielded the maximum Jaccard value was identified, and the threshold for each peak set was taken as that which yielded this maximum N. The same process was applied to compare PWM-predicted binding sites and ChIP-seq peaks.

### GHT-SELEX experiment overlap comparison

To investigate the reproducibility of GHT-SELEX profiles, we focused on experiment pairs with the same TF construct and protein production approach. Per replicate, we fitted genome-wide MAGIX models to 200-bp non-overlapping genomic intervals (∼13M genomic intervals), resulting in a model coefficient per genomic bin. The exponential of this coefficient represents the fold-enrichment per cycle, which we visualize as scatterplots and use to calculate the Pearson correlation between replicates.

### Motif derivation and assessment of experiment success

We evaluated the success of GHT- and HT-SELEX experiments systematically alongside other Codebook datasets generated with ChIP-seq, protein binding microarray, and SMiLE-seq experiments (See **Table S7** for a summary of all experiments), by performing motif discovery with nine different programs and then testing their performance in all experiments for the same TF. Experiments were classified by expert curation as successful if they produced motifs that scored well in data generated by other methods (preferably, and in most cases) and, in rare cases, in replicate data from the same experimental method, when other methods failed. All motifs can be browsed at https://mex.autosome.org. See accompanying article for detailed description of the process^66^.

### Representative motif selection

For each TF, a list of up to 60 motifs was automatically assembled based on auROC, auPRC, and motif centrality metrics performance across all successful experiments^66^. These motifs were then assessed manually by an assembly of authors to select a representative motif. Besides scoring across all data sets, we considered whether the motif appears likely to describe the inherent specificity of the TF, rather than, for example, a larger sequence that is partially derived from a genomic repeat element (a scenario common with KRAB zinc fingers). In the cases where several motifs displayed highly similar performance, we prioritized motifs with higher information content. See^66^ for further details and **Supplementary Data S1** for the representative motifs.

### Comparing PWM scoring methods

To create *in silico* predicted binding sites for a TF, we first scanned the genome using the generated PWM (see the Codebook overview manuscript for the details on PWM selection), using MOODS^67^ with a p-value threshold of 0.0001. We then merged the clusters of PWM hits with a distance less than 200bp between neighboring hits, since this is the median length of ChIP-seq fragments, and the task is predicting in vivo binding sites. Singleton PWM hits and boundary hits were also expanded to have a width of at least 200bp. The clusters of PWM hits were re-scored using sum-of-affinity (i.e., with PWM log-odds scores at each base converted to linear/probability space, prior to calculation of the sum) and maximum-affinity methods, by either applying a sum or maximum, respectively, over the PWM scores of the cluster members. The resulting sites were sorted by their new score and processed through the same optimization procedure described above for peaks, to maximize their overlap with ChIP-seq peaks.

### Modeling alternative C2H2-zf binding modes with RCADEEM

RCADEEM uses a hidden Markov model (HMM) to represent multiple, alternative DNA-binding motifs, each corresponding to the binding preference of a C2H2-zf array. Briefly, the DNA sequences (e.g., GHT-SELEX peaks) are modeled as sequences generated from a discrete Markov process with hidden states that include a background state (*S*0) and *M* motif states *Sm* (*m*∈[1,*M*]). The background state, with marginal probability π0, emits each nucleotide *n* with probability *b*0(*n*) (Σn*b*0(*n*)=1). The background state can transition to itself (i.e., consecutive DNA nucleotides can be generated from the background state) with probability *a*0,0, or to each motif state *Sm* (*m*∈[1,*M*]) with probability *a*0,*m* (*a*0,0+Σm*a*0,m=1). Each motif *m*, with marginal probability π*m*, generates a sequence of length *lm*, with each nucleotide *n* at position *i* emitted with probability *bm*,*i*(*n*) (Σn*bm*,*i*(*n*)=1 ∀*m*∈[1,*M*], *i*∈[1,*lm*]). Note that, for each *m*∈[1,*M*], the values *bm*,*i*(*n*) form a position-specific frequency matrix (PFM, i.e. the exponential of the classical log-odds PWM) with width *lm*, which is fixed to be 3 times the number of zinc finger domains in the array represented by motif *m*, as each zinc finger domain binds to three nucleotides. Finally, each motif state *Sm* transitions to the background state with probability *a*m,0=1.

We start the model by including the motifs representing all possible consecutive zinc finger domain arrays^50^. We initialize the emission probabilities *bm*,*i*(*n*) for each motif *m* using the PFM predicted for the associated zinc finger array by a previously created C2H2-zf recognition code^51^—this recognition code is a machine learning model that, given the sequence of a zinc finger array, predicts the expected binding preference. The HMM parameters, including all marginal state probabilities, state transition probabilities, and emission probabilities, are then optimized via expectation maximization using Baum–Welch algorithm. Then, each of the optimized PFMs is tested for (*i*) enrichment of the motif in actual sequences compared to dinucleotide-shuffled sequences, and (*ii*) similarity to the original recognition code-predicted PFM. To achieve (*i*), for each position *x* in each DNA sequence *k*, we calculate γ*k,x*(*Sm*), the probability that it was generated from motif state *Sm*, using the forward-backward algorithm. The motif score for DNA sequence *k* is then calculated as Σ*x*γ*k,x*(*Sm*)/*lm*, representing the expected number of times the state *Sm* is seen in sequence *k*. For each motif *m*, these scores are calculated both for actual GHT-SELEX peak sequences and their dinucleotide-shuffled version. Then, the top 100 sequences with the largest scores for each motif are tested to see whether they are enriched in the motif compared to shuffled sequences (Fisher’s exact test, FDR≤0.01). Motifs that do not pass this cutoff are removed from the model. To achieve (*ii*), each HMM-optimized PFM is first converted to log-scale (representing a PWM), followed by the calculation of the Pearson correlation of the PWM entries with those predicted by the recognition code. Pearson correlations are then converted using Fisher transformation in order to calculate a P-value, followed by removal of motifs that do not pass the FDR cut-off ≤0.01. The remaining motifs are then used to reconstruct a smaller HMM, similar to the procedure described above, followed by another round of EM optimization. This procedure is repeated until all motifs pass the cut-offs for enrichment in GHT-SELEX sequences while maintaining significant similarity to the original recognition code-predicted sequences.

To visualize the binding modes predicted by RCADEEM, the resulting PWMs are used to identify their best match in each of the input sequences using AffiMx^52^. Then, for each sequence, the PWM with the highest weighted HMM score on the best match is kept as the predicted binding mode. To align the sequences, offsets are calculated based on the corresponding C2H2-zf domains (**Figures 6a-f**). C2H2-zf proteins were categorized based on their alternative usage of C2H2-zf domains (i.e., Multiple DBDs, Finger shift, Canonical, and Core with extensions; **Figure 6**) through an expert-curated evaluation (**Table S10**). To make a motif model for each binding mode, we manually selected representative peaks corresponding to each bi<u>n</u>ding mode over the 2000 GHT-SELEX peaks with the highest enrichment coefficient. The sequence (already aligned by RCADEEM) and C2H2-zf domain array coordinates of these peaks were used to create PFMs. The resulting PFMs for those C2H2-zf TFs are available in **Supplementary Data S2** and online at https://cisbp.ccbr.utoronto.ca^51^. The logos, coordinates, selected sequences, annotated sequence heatmaps, and associated metadata are available online at https://codebook.ccbr.utoronto.ca. The source code for RCADEEM is available at https://github.com/csglab/RCADEEM.

### Comparison of C2H2 DBDs

C2H2 DBD similarities were compared by pairwise alignment with Needleman-Wunsch algorithm, as implemented in the R-package Biostrings and counting substitutions, insertions, and unmatched flanking bases as edits. DNA-binding functionality scores and predicted motif similarity for the DBDs were analyzed as described previously^68^.

## DATA AVAILABILITY

The sequencing raw data for the HT-SELEX and GHT-SELEX experiments have been deposited into the SRA database under identifiers PRJEB61115 (HT-SELEX) and PRJEB76622 (GHT-SELEX). Additionally, genomic interval information generated for the GHT-SELEX has been deposited into GEO under accession GSE278858. The entire Codebook data structure, with many accessory files and browsable results at is available at https://codebook.ccbr.utoronto.ca. Larger collection of motifs generated for these experiments in an accompanying study^49^ can be browsed at https://mex.autosome.org.

## CODE AVAILABILITY

Source code for MAGIX and RCADEEM is available in GitHub (https://github.com/csglab/MAGIX and https://github.com/csglab/RCADEEM).

## ACKNOWLEDGEMENTS

We thank the IT Group of the Institute of Computer Science at Halle University for computational resources, Maximilian Biermann for valuable technical support, Gherman Novakovsky for providing feedback, Berat Dogan for testing earlier versions of RCADEEM, and Debashish Ray for assistance with database depositions.

This work was supported by the following:

- Canadian Institutes of Health Research (CIHR) grants FDN-148403, PJT-186136, PJT-191768, and PJT-191802, and NIH grant R21HG012258 to T.R.H.
- CIHR grant PJT-191802 to T.R.H. and H.S.N.
- Natural Sciences and Engineering Research Council of Canada (NSERC) grant RGPIN-2018-05962 to H.S.N.
- Russian Science Foundation grant 24-14-20031 to F.A.K.
- MSHERF grant 075-15-2025-014 (prev. 075-15-2024-666)
- State assignment 125091010189-3 (FFRW-2025-010)
- Swiss National Science Foundation grant (no. 310030_197082) to B.D.
- Marie Skłodowska-Curie (no. 895426) and EMBO long-term (1139-2019) fellowships to J.F.K.
- NIH grants R01HG013328 and U24HG013078 to M.T.W., T.R.H., and Q.M.
- NIH grants R01AR073228, P30AR070549, and R01AI173314 to M.T.W.
- NIH grant P30CA008748 partially supported Q.M.
- Canada Research Chairs funded by CIHR to T.R.H. and H.S.N.
- Ontario Graduate Scholarships to K.U.L and I.Y.
- A.J. was supported by Vetenskapsrådet (Swedish Research Council) Postdoctoral Fellowship (2016-00158)
- The Billes Chair of Medical Research at the University of Toronto to T.R.H.
- EPFL Center for Imaging
- Institutional funding from EPFL
- Resource allocations from the Digital Research Alliance of Canada

## Author contributions

A.J. and T.R.H. designed the study. A.J. and A.W.H.Y. performed the SELEX experiments with assistance from H.Z., R.R., and A.B. A.H.C. and H.S.N. developed and performed MAGIX and RCADEEM. A.H.C., A.F., K.U.L. and A.J. performed sequence and data analyses with assistance from A.B.; A.J., K.U.L and A.H.C. prepared the illustrations. M.A. and I.V.K. orchestrated the data and motif repositories. A.J, A.H.C, A.F, K.U.L, H.S.N and T.R.H. wrote the paper. All authors contributed to data analysis and reviewed the manuscript.

---

**Extended Data Fig. 1:**
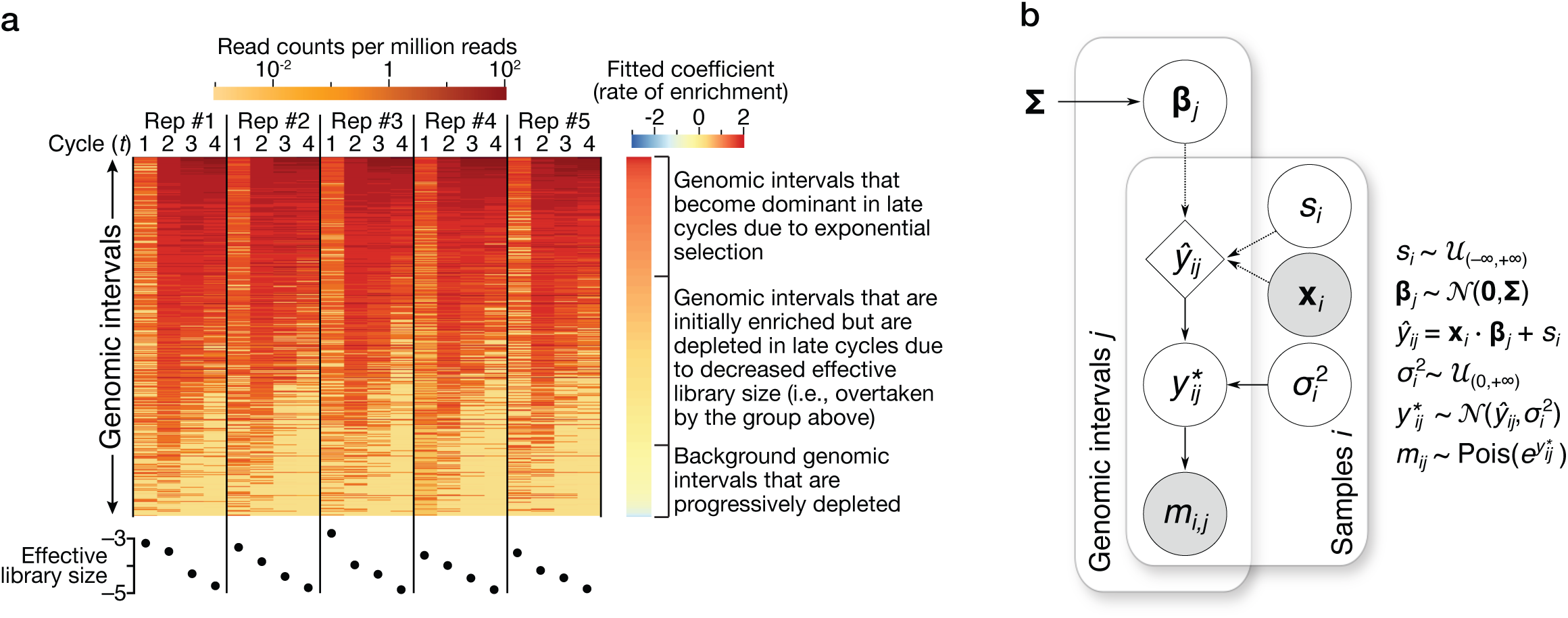
**a.** Example of actual read count data for CTCF over five replicates of four cycles, illustrating enrichment patterns, fitted coefficients (right), and estimated library sizes (bottom). **b.** A brief overview of the statistical framework of the generative model of MAGIX. Open circles, closed circles, and the diamonds represent latent variables, observed variables, and deterministic computations, respectively. *si*: library size for sample *i*; **x***i*: vector of sample-level variables for sample *i*, including an intercept term and a term for the SELEX cycle, in addition to other terms for batch and background effects; **β***j*: vector of model coefficients for interval *j*; *mij*: number of observed reads mapping to interval *j* in sample *i*. See **Methods** for description of other variables..

**Extended Data Fig. 2:**
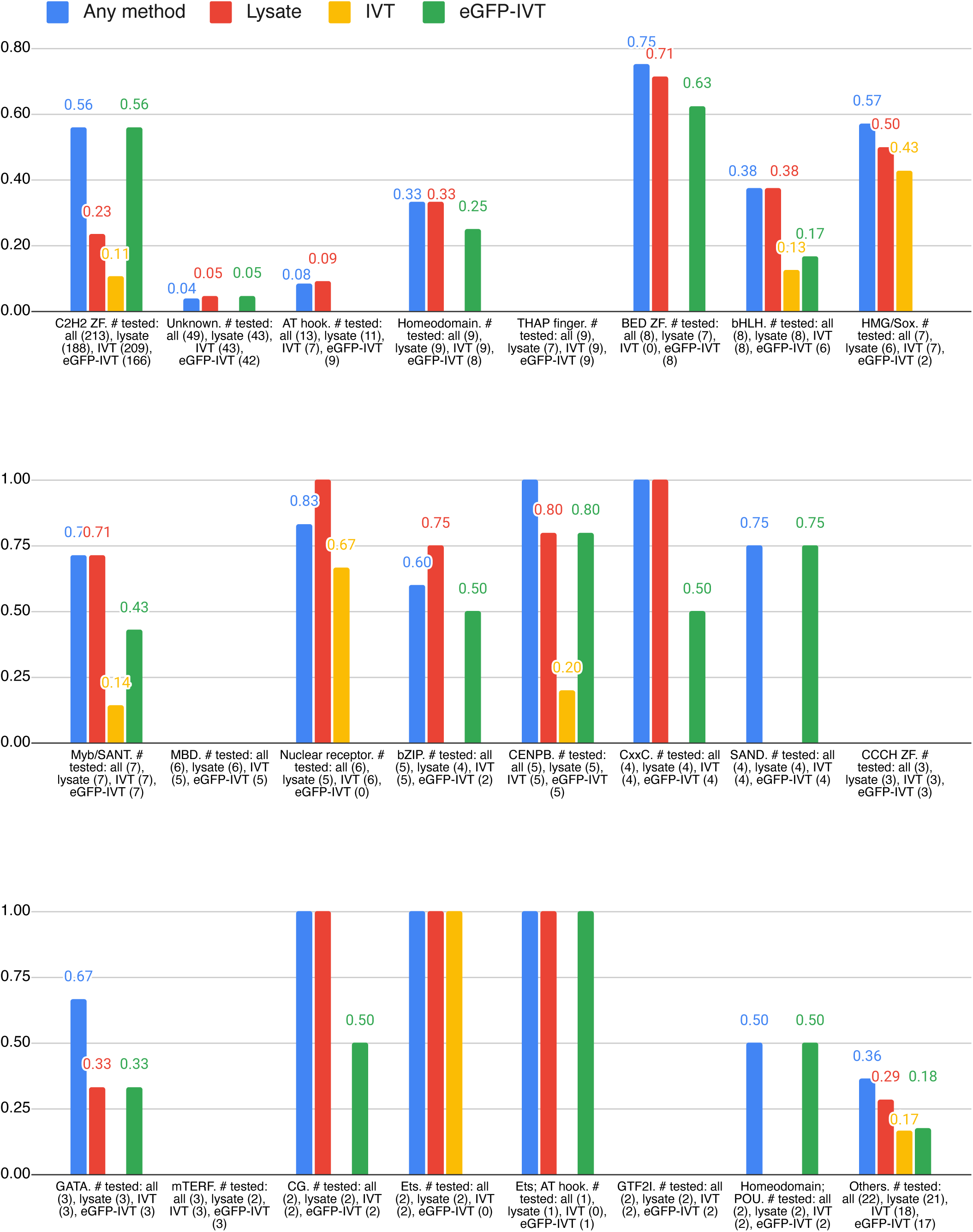
TF success rate by protein production method and structural class. Figure displays bar graphs of success rates for putative TFs based on their structural class showing a bar for TFs based on success on if the protein was successful in any of the production methods and then individually based on protein production methods. Number of TFs tested in each category is listed with the name of the structural class.

**Extended Data Fig. 3:**
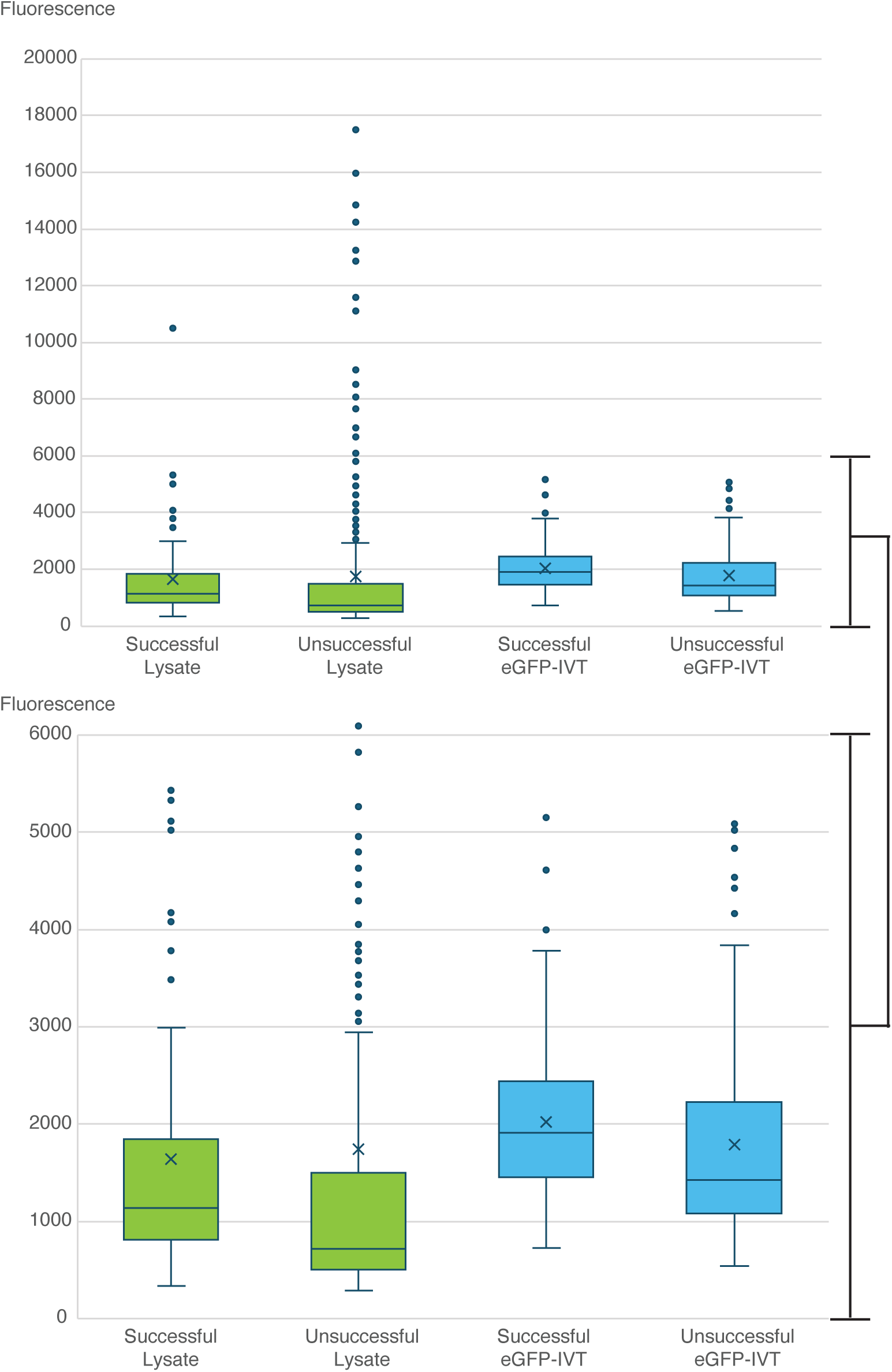
TF expression levels are plotted for four categories based on experiment success and protein production. Data is shown for the full range of fluorescence (*top)* and for the region with most of the signal (*bottom).* Note that even though the successful samples have generally higher expression levels in both protein production systems it is not an effective predictor of experiment success.

## Notes

https://cisbp.ccbr.utoronto.ca

https://codebook.ccbr.utoronto.ca

https://mex.autosome.org

### Competing Interest Statement

The authors have declared no competing interest.

### Summary of Updates

New version of the manuscript has been updated substantially to respond reviewers' feedback through: additional experiments, re-analysis of most datasets using updated computational approaches, addition of supplementary figures and tables, and through editing both text and main display items

## REFERENCES

1. Stormo, G.D. & Zhao, Y. Determining the specificity of protein-DNA interactions. Nat Rev Genet 11, 751–60 (2010).

2. Khan, A. et al. JASPAR 2018: update of the open-access database of transcription factor binding profiles and its web framework. Nucleic Acids Res 46, D1284 (2018).

3. Bernard, B., Thorsson, V., Rovira, H. & Shmulevich, I. Increasing coverage of transcription factor position weight matrices through domain-level homology. PLoS One 7, e42779 (2012).

4. Lambert, S.A. et al. Similarity regression predicts evolution of transcription factor sequence specificity. Nat Genet 51, 981–989 (2019).

5. Wunderlich, Z. & Mirny, L.A. Different gene regulation strategies revealed by analysis of binding motifs. Trends Genet 25, 434–40 (2009).

6. Patel, Z.M. & Hughes, T.R. Global properties of regulatory sequences are predicted by transcription factor recognition mechanisms. Genome Biol 22, 285 (2021).

7. Wasserman, W.W. & Sandelin, A. Applied bioinformatics for the identification of regulatory elements. Nat Rev Genet 5, 276–87 (2004).

8. Valouev, A. et al. Genome-wide analysis of transcription factor binding sites based on ChIP-Seq data. Nat Methods 5, 829–34 (2008).

9. Marinov, G.K., Kundaje, A., Park, P.J. & Wold, B.J. Large-Scale Quality Analysis of Published ChIP-seq Data. G3 (Bethesda) (2013).

10. Consortium, E.P. et al. Expanded encyclopaedias of DNA elements in the human and mouse genomes. Nature 583, 699–710 (2020).

11. Rhee, H.S. & Pugh, B.F. Comprehensive Genome-wide Protein-DNA Interactions Detected at Single-Nucleotide Resolution. Cell 147, 1408–19 (2011).

12. Long, H.K., Prescott, S.L. & Wysocka, J. Ever-Changing Landscapes: Transcriptional Enhancers in Development and Evolution. Cell 167, 1170–1187 (2016).

13. Zinzen, R.P., Girardot, C., Gagneur, J., Braun, M. & Furlong, E.E. Combinatorial binding predicts spatio-temporal cis-regulatory activity. Nature 462, 65–70 (2009).

14. Liu, X., Lee, C.K., Granek, J.A., Clarke, N.D. & Lieb, J.D. Whole-genome comparison of Leu3 binding in vitro and in vivo reveals the importance of nucleosome occupancy in target site selection. Genome Res 16, 1517–28 (2006).

15. Kim, T.H. et al. Analysis of the vertebrate insulator protein CTCF-binding sites in the human genome. Cell 128, 1231–45 (2007).

16. Fu, Y., Sinha, M., Peterson, C.L. & Weng, Z. The insulator binding protein CTCF positions 20 nucleosomes around its binding sites across the human genome. PLoS Genet 4, e1000138 (2008).

17. Walker, M. et al. Affinity-seq detects genome-wide PRDM9 binding sites and reveals the impact of prior chromatin modifications on mammalian recombination hotspot usage. Epigenetics Chromatin 8, 31 (2015).

18. Morgunova, E. et al. Two distinct DNA sequences recognized by transcription factors represent enthalpy and entropy optima. Elife 7(2018).

19. Stormo, G.D. DNA binding sites: representation and discovery. Bioinformatics 16, 16–23 (2000).

20. Rohs, R. et al. The role of DNA shape in protein-DNA recognition. Nature 461, 1248–53 (2009).

21. Zhao, Y., Ruan, S., Pandey, M. & Stormo, G.D. Improved models for transcription factor binding site identification using nonindependent interactions. Genetics 191, 781–90 (2012).

22. Jolma, A. et al. DNA-dependent formation of transcription factor pairs alters their binding specificity. Nature 527, 384–8 (2015).

23. Horton, C.A. et al. Short tandem repeats bind transcription factors to tune eukaryotic gene expression. Science 381, eadd1250 (2023).

24. Kribelbauer, J.F., Rastogi, C., Bussemaker, H.J. & Mann, R.S. Low-Affinity Binding Sites and the Transcription Factor Specificity Paradox in Eukaryotes. Annu Rev Cell Dev Biol 35, 357–379 (2019).

25. Thomson, J.P. et al. CpG islands influence chromatin structure via the CpG-binding protein Cfp1. Nature 464, 1082–6 (2010).

26. Lambert, S.A. et al. The Human Transcription Factors. Cell 175, 598–599 (2018).

27. Wolfe, S.A., Nekludova, L. & Pabo, C.O. DNA recognition by Cys2His2 zinc finger proteins. Annu Rev Biophys Biomol Struct 29, 183–212 (2000).

28. Klug, A. The discovery of zinc fingers and their applications in gene regulation and genome manipulation. Annu Rev Biochem 79, 213–31 (2010).

29. Stubbs, L., Sun, Y. & Caetano-Anolles, D. Function and Evolution of C2H2 Zinc Finger Arrays. Subcell Biochem 52, 75–94 (2011).

30. Nakahashi, H. et al. A genome-wide map of CTCF multivalency redefines the CTCF code. Cell Rep 3, 1678–1689 (2013).

31. Kieffer-Kwon, K.R. et al. Interactome maps of mouse gene regulatory domains reveal basic principles of transcriptional regulation. Cell 155, 1507–20 (2013).

32. Han, B.Y., Foo, C.S., Wu, S. & Cyster, J.G. The C2H2-ZF transcription factor Zfp335 recognizes two consensus motifs using separate zinc finger arrays. Genes Dev 30, 1509–14 (2016).

33. Najafabadi, H.S. et al. C2H2 zinc finger proteins greatly expand the human regulatory lexicon. Nat Biotechnol (2015).

34. Najafabadi, H.S., Albu, M. & Hughes, T.R. Identification of C2H2-ZF binding preferences from ChIP-seq data using RCADE. Bioinformatics 31, 2879–81 (2015).

35. Tuerk, C. & Gold, L. Systematic evolution of ligands by exponential enrichment: RNA ligands to bacteriophage T4 DNA polymerase. Science 249, 505–10 (1990).

36. Jolma, A. et al. DNA-Binding Specificities of Human Transcription Factors. Cell 152, 327–39 (2013).

37. Jolma, A. et al. Multiplexed massively parallel SELEX for characterization of human transcription factor binding specificities. Genome Res 20, 861–73 (2010).

38. Yin, Y. et al. Impact of cytosine methylation on DNA binding specificities of human transcription factors. Science 356(2017).

39. Reiss, D.J. & Mobley, H.L. Determination of target sequence bound by PapX, repressor of bacterial motility, in flhD promoter using systematic evolution of ligands by exponential enrichment (SELEX) and high throughput sequencing. J Biol Chem 286, 44726–38 (2011).

40. Ishihama, A., Shimada, T. & Yamazaki, Y. Transcription profile of Escherichia coli: genomic SELEX search for regulatory targets of transcription factors. Nucleic Acids Res 44, 2058–74 (2016).

41. Illingworth, R.S. et al. Orphan CpG islands identify numerous conserved promoters in the mammalian genome. PLoS Genet 6, e1001134 (2010).

42. O’Malley, R.C. et al. Cistrome and Epicistrome Features Shape the Regulatory DNA Landscape. Cell 165, 1280–1292 (2016).

43. Jolma, A. et al. Perspectives on Codebook: sequence specificity of uncharacterized human transcription factors. bioRxiv, 2024.11.11.622097 (2024).

44. Berger, M.F. et al. Compact, universal DNA microarrays to comprehensively determine transcription-factor binding site specificities. Nat Biotechnol 24, 1429–35 (2006).

45. Isakova, A. et al. SMiLE-seq identifies binding motifs of single and dimeric transcription factors. Nat Methods 14, 316–322 (2017).

46. Lambert, S.A. et al. The Human Transcription Factors. Cell 172, 650–665 (2018).

47. Razavi, R. et al. Extensive binding of uncharacterized human transcription factors to genomic dark matter. bioRxiv, 2024.11.11.622123 (2024).

48. Schmitges, F.W. et al. Multiparameter functional diversity of human C2H2 zinc finger proteins. Genome Res 26, 1742–1752 (2016).

49. Vorontsov, I.E. et al. Cross-platform DNA motif discovery and benchmarking to explore binding specificities of poorly studied human transcription factors. bioRxiv, 2024.11.11.619379 (2024).

50. Gralak, A.J. et al. Identification of methylation-sensitive human transcription factors using meSMiLE-seq. bioRxiv, 2024.11.11.619598 (2024).

51. Weirauch, M.T. et al. Determination and inference of eukaryotic transcription factor sequence specificity. Cell 158, 1431–43 (2014).

52. Razavi, R. et al. Extensive binding of uncharacterized human transcription factors to genomic dark matter. bioRxiv, 2024.11.11.622123 (2024).

53. Birke, M. et al. The MT domain of the proto-oncoprotein MLL binds to CpG-containing DNA and discriminates against methylation. Nucleic Acids Res 30, 958–65 (2002).

54. Khetan, S., Carroll, B.S. & Bulyk, M.L. Multiple overlapping binding sites determine transcription factor occupancy. Nature (2025).

55. Stormo, G.D. & Fields, D.S. Specificity, free energy and information content in protein-DNA interactions. Trends Biochem Sci 23, 109–13 (1998).

56. Weirauch, M.T. et al. Evaluation of methods for modeling transcription factor sequence specificity. Nat Biotechnol 31, 126–34 (2013).

57. Kuznetsov, V.A. Mathematical Modeling of Avidity Distribution and Estimating General Binding Properties of Transcription Factors from Genome-Wide Binding Profiles. Methods Mol Biol 1613, 193–276 (2017).

58. Alexandrov, I., Kazakov, A., Tumeneva, I., Shepelev, V. & Yurov, Y. Alpha-satellite DNA of primates: old and new families. Chromosoma 110, 253–66 (2001).

59. Huttlin, E.L. et al. Architecture of the human interactome defines protein communities and disease networks. Nature 545, 505–509 (2017).

60. de Boer, C.G. & Taipale, J. Hold out the genome: a roadmap to solving the cis-regulatory code. Nature 625, 41–50 (2024).

61. Stormo, G.D., Schneider, T.D., Gold, L. & Ehrenfeucht, A. Use of the ‘Perceptron’ algorithm to distinguish translational initiation sites in E. coli. Nucleic Acids Res 10, 2997–3011 (1982).

62. Laverty, K.U. et al. PRIESSTESS: interpretable, high-performing models of the sequence and structure preferences of RNA-binding proteins. Nucleic Acids Res 50, e111 (2022).

63. Bolger, A.M., Lohse, M. & Usadel, B. Trimmomatic: a flexible trimmer for Illumina sequence data. Bioinformatics 30, 2114–20 (2014).

64. Li, H. et al. The Sequence Alignment/Map format and SAMtools. Bioinformatics 25, 2078–9 (2009).

65. Marcais, G. & Kingsford, C. A fast, lock-free approach for efficient parallel counting of occurrences of k-mers. Bioinformatics 27, 764–70 (2011).

66. Vorontsov, I.E. et al. Cross-platform DNA motif discovery and benchmarking to explore binding specificities of poorly studied human transcription factors. bioRxiv, 2024.11.11.619379 (2024).

67. Korhonen, J., Martinmaki, P., Pizzi, C., Rastas, P. & Ukkonen, E. MOODS: fast search for position weight matrix matches in DNA sequences. Bioinformatics 25, 3181–2 (2009).

68. Najafabadi, H.S. et al. Non-base-contacting residues enable kaleidoscopic evolution of metazoan C2H2 zinc finger DNA binding. Genome Biol 18, 167 (2017).

69. Korhonen, J.H., Palin, K., Taipale, J. & Ukkonen, E. Fast motif matching revisited: high-order PWMs, SNPs and indels. Bioinformatics 33, 514–521 (2017).

