## Supplementary figures and images for "GHT-SELEX demonstrates unexpectedly high intrinsic sequence specificity and complex DNA binding of many human transcription factors"

### AC092835_PWM006586_logo.png

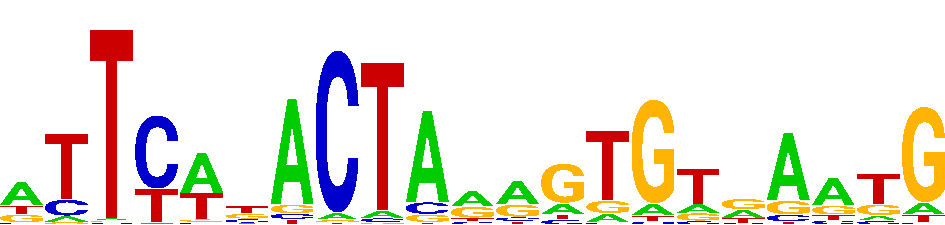

### BATF2_PWM000075_logo.png

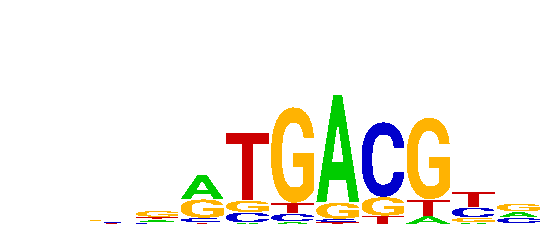

### CAMTA1_PWM000118_logo.png

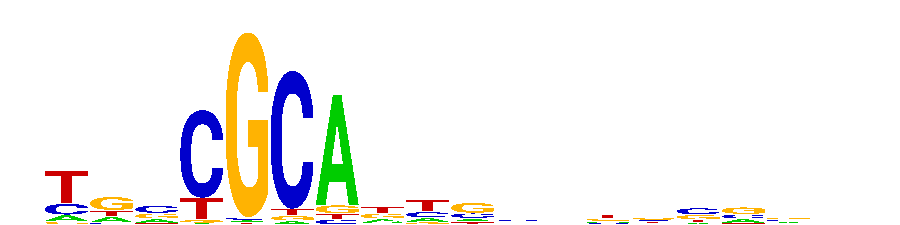

### CTCF_PWM000310_logo_REV.png

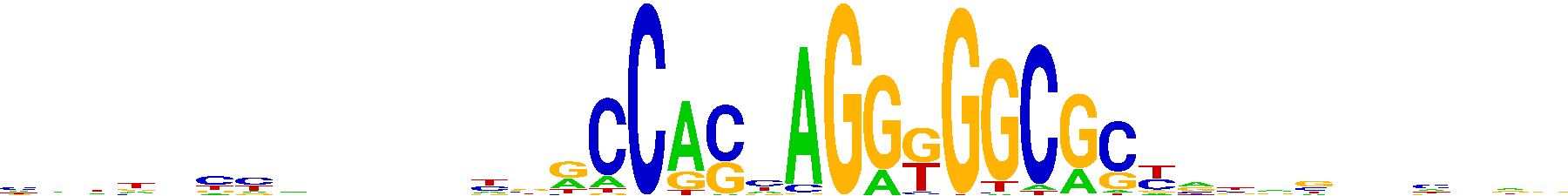

### DACH1_PWM020780_logo.png

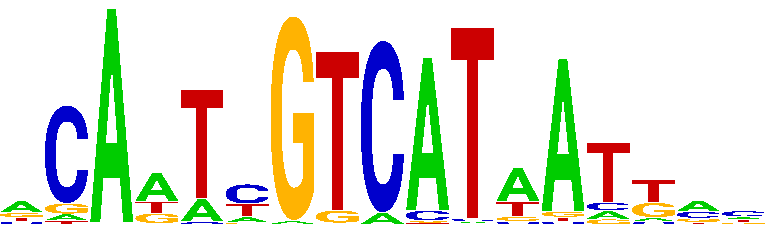

### DMTF1_PWM000380_logo_REV.png

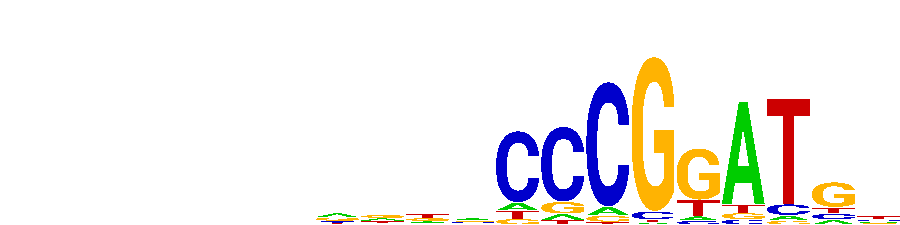

### DNTTIP1_PWM000419_logo.png

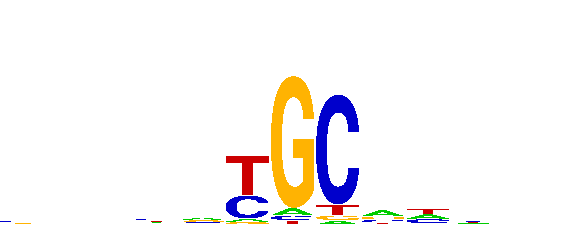

### DNTTIP1_PWM000419_logo_REV.png

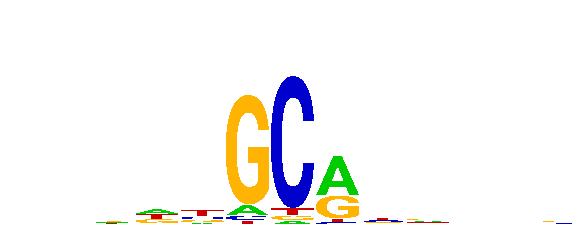

### FBXL19_PWM000531_logo.png

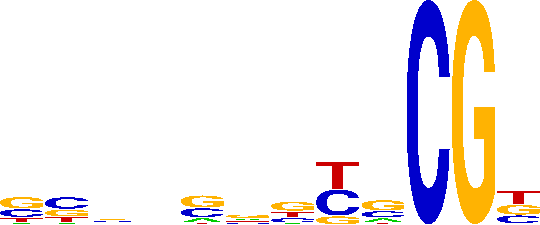

### FBXL19_PWM000531_logo_REV.png

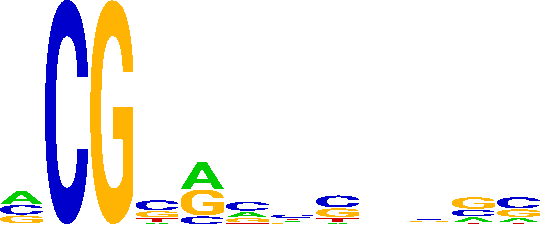

### FIZ1_PWM000556_logo_REV.png

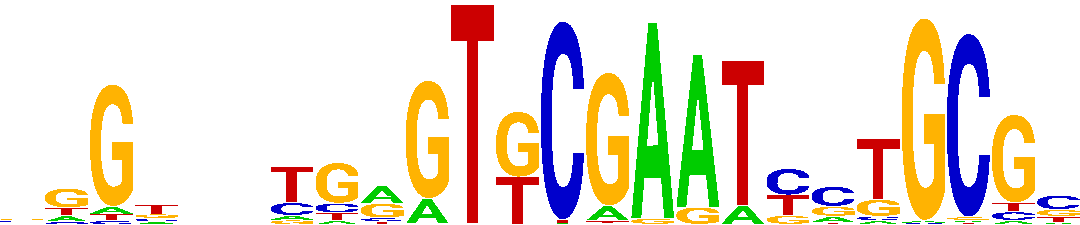

### FLYWCH1_PWM000599_logo_REV.png

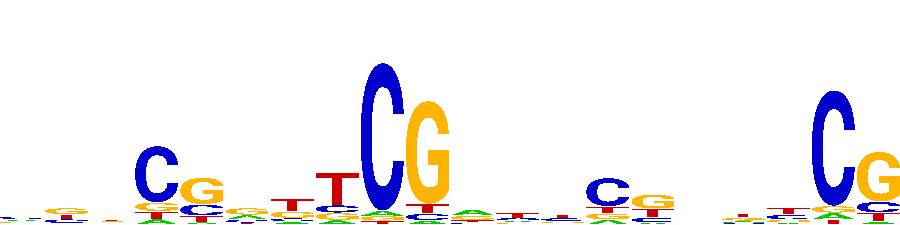

### FOSL2_PWM000627_logo.png

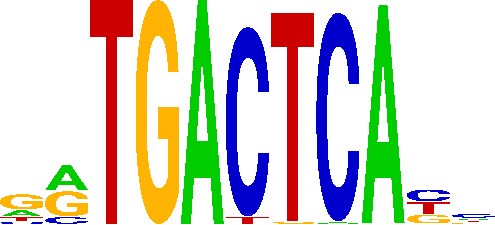

### GABPA_PWM000688_logo_REV.png

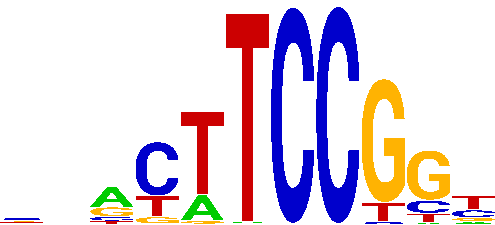

### GCM1_PWM000730_logo.png

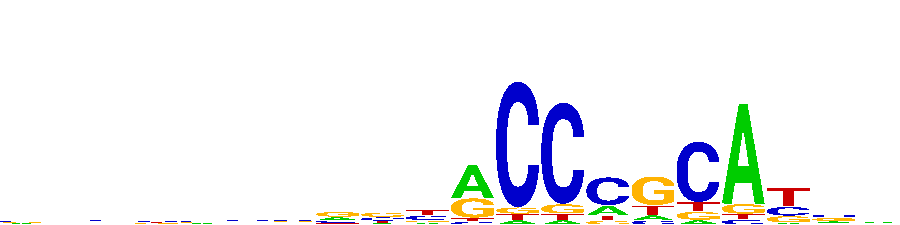

### GLYR1_PWM034466_logo.png

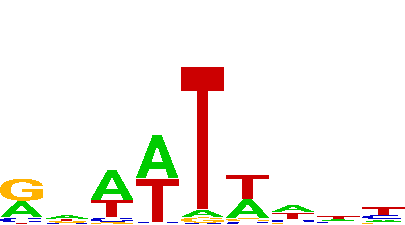

### GRHL3_PWM034802_logo.png

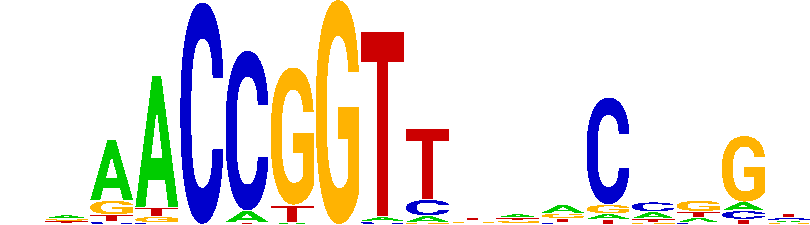

### GTF2IRD2_PWM000795_logo.png

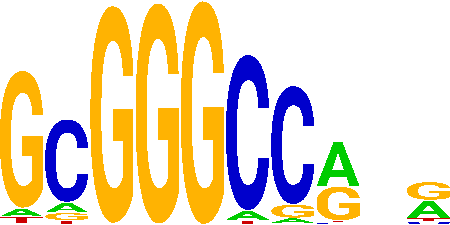

### HMGA1_PWM036826_logo.png

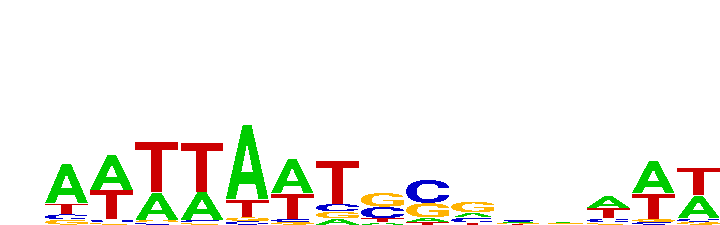

### JRK_PWM038292_logo_REV.png

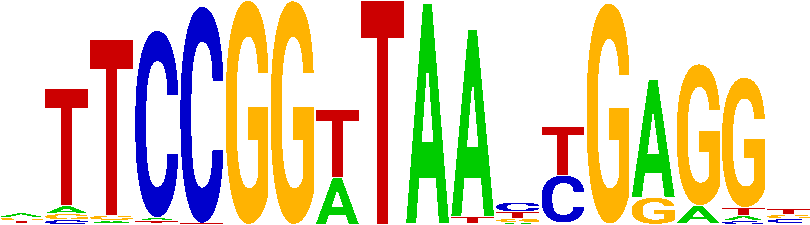

### JUN_PWM000851_logo.png

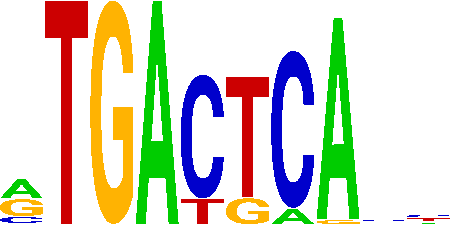

### KDM2A_PWM000899_logo_REV.png

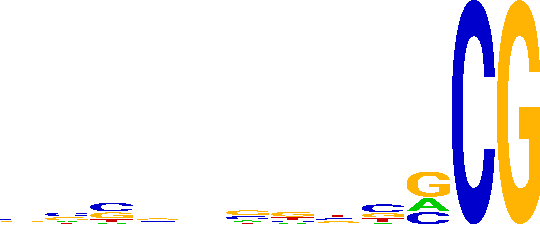

### LEUTX_PWM000962_logo.png

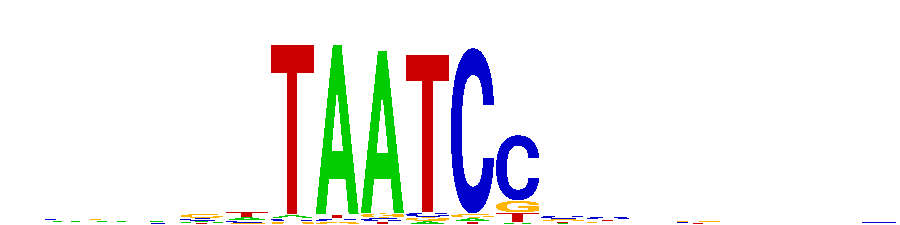

### LHX6_PWM000985_logo_REV.png

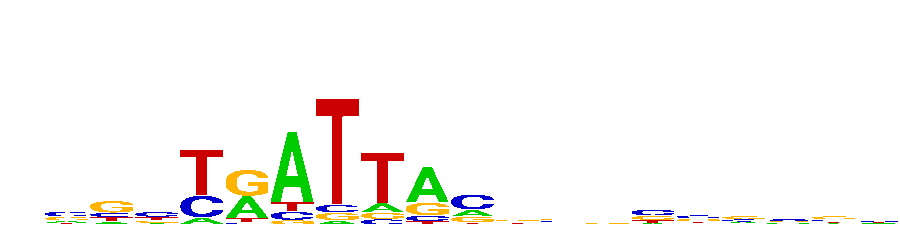

### MBD1_PWM001044_logo_REV.png

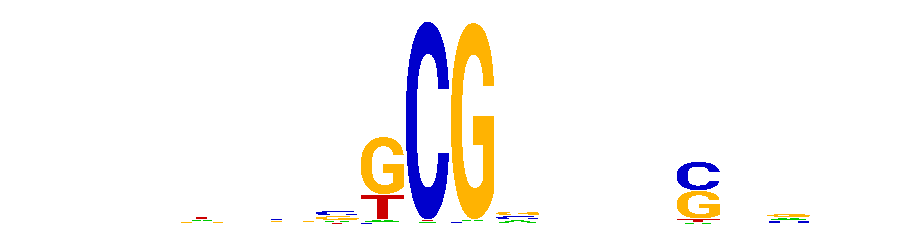

### MKX_PWM001152_logo_REV.png

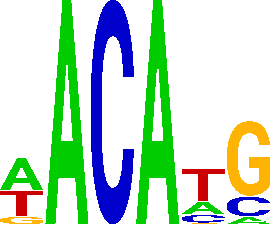

### MSANTD1_PWM001197_logo.png

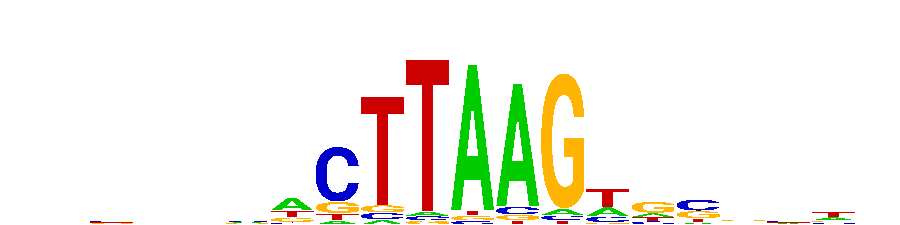

### MYF6_PWM001260_logo_REV.png

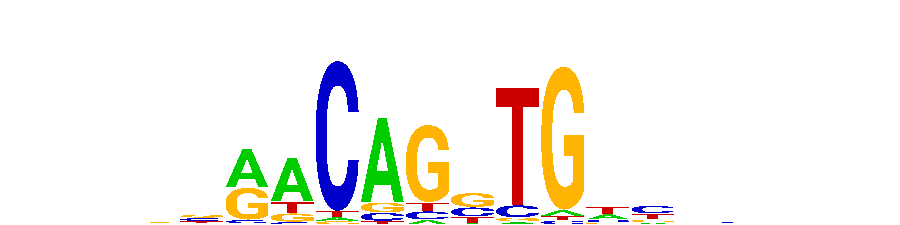

### MYRFL_PWM002425_logo.png

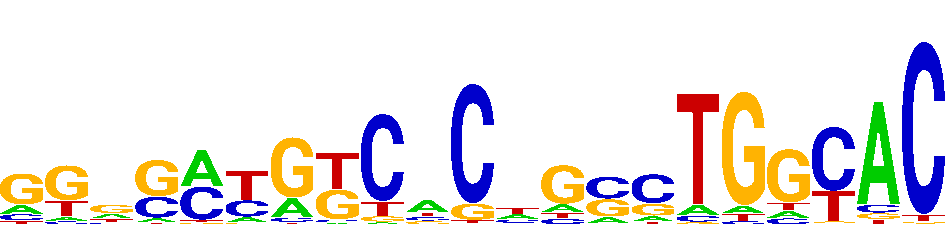

### MYRFL_PWM002425_logo_REV.png

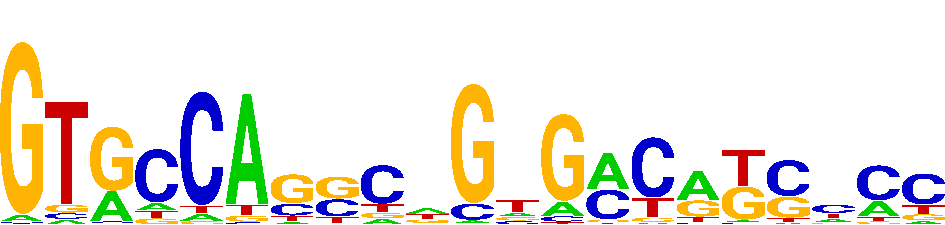

### NFKB1_PWM001438_logo.png

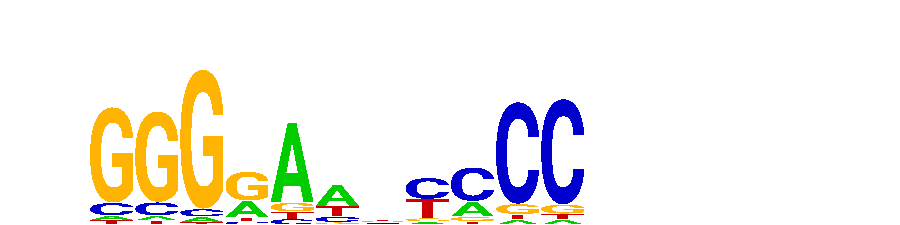

### NFKB1_PWM001438_logo_REV.png

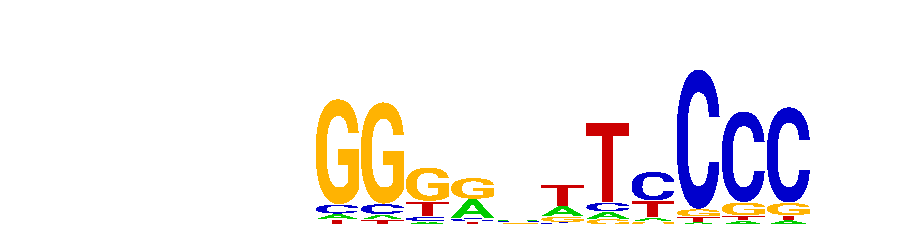

### NR1H4_PWM001456_logo.png

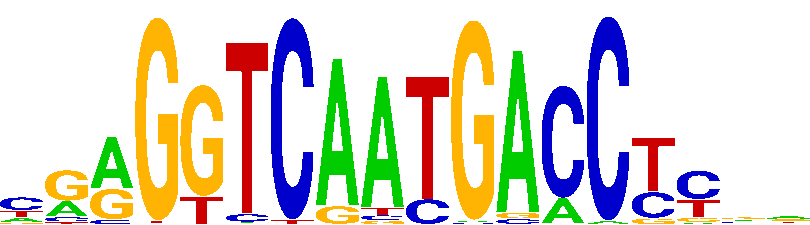

### NR4A2_PWM057625_logo.png

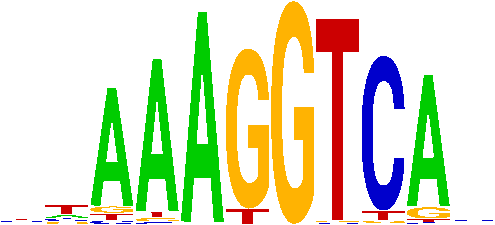

### NR4A2_PWM057625_logo_REV.png

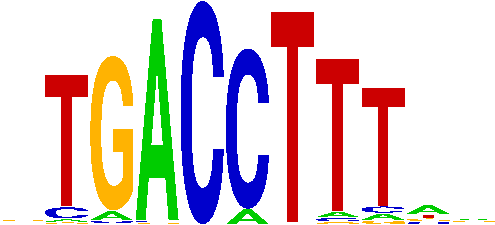

### PAX7_PWM001537_logo.png

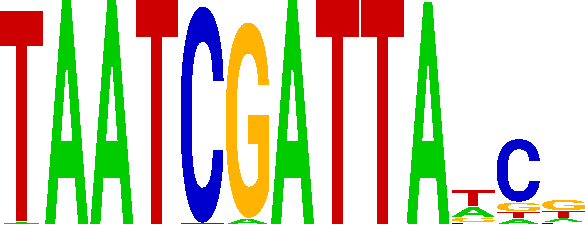

### POGK_PWM058987_logo.png

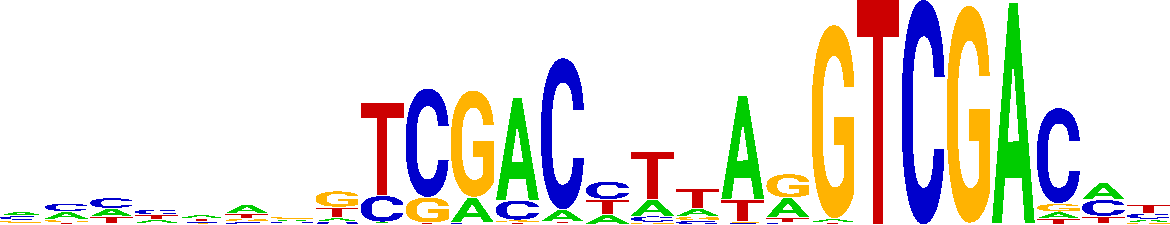

### PRDM13_PWM001636_logo_REV.png

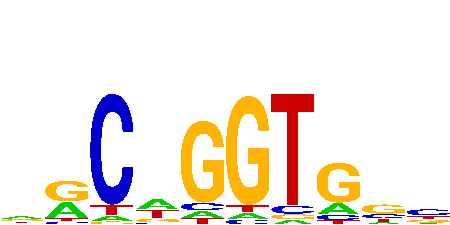

### SALL3_PWM001887_logo.png

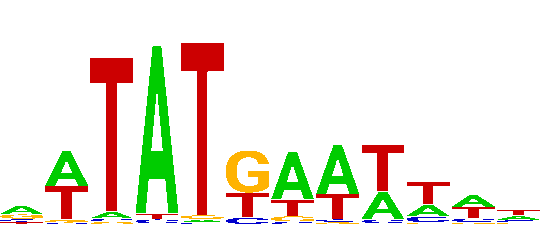

### SLC2A4RG_PWM001936_logo_REV.png

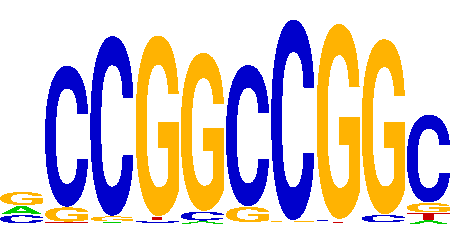

### SP140L_PWM002104_logo.png

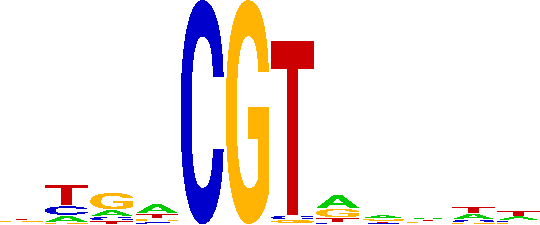

### Supplementary Document S1

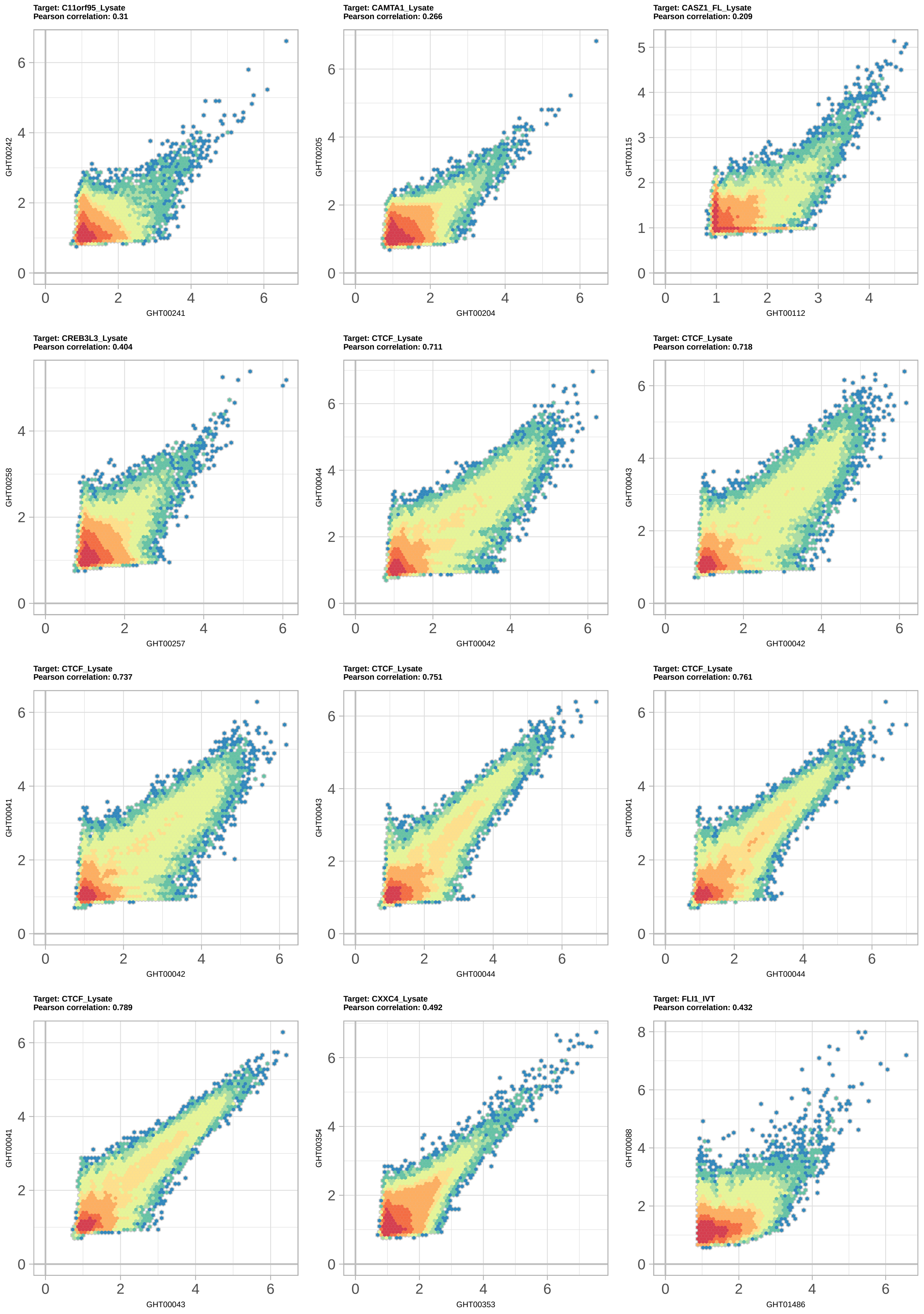

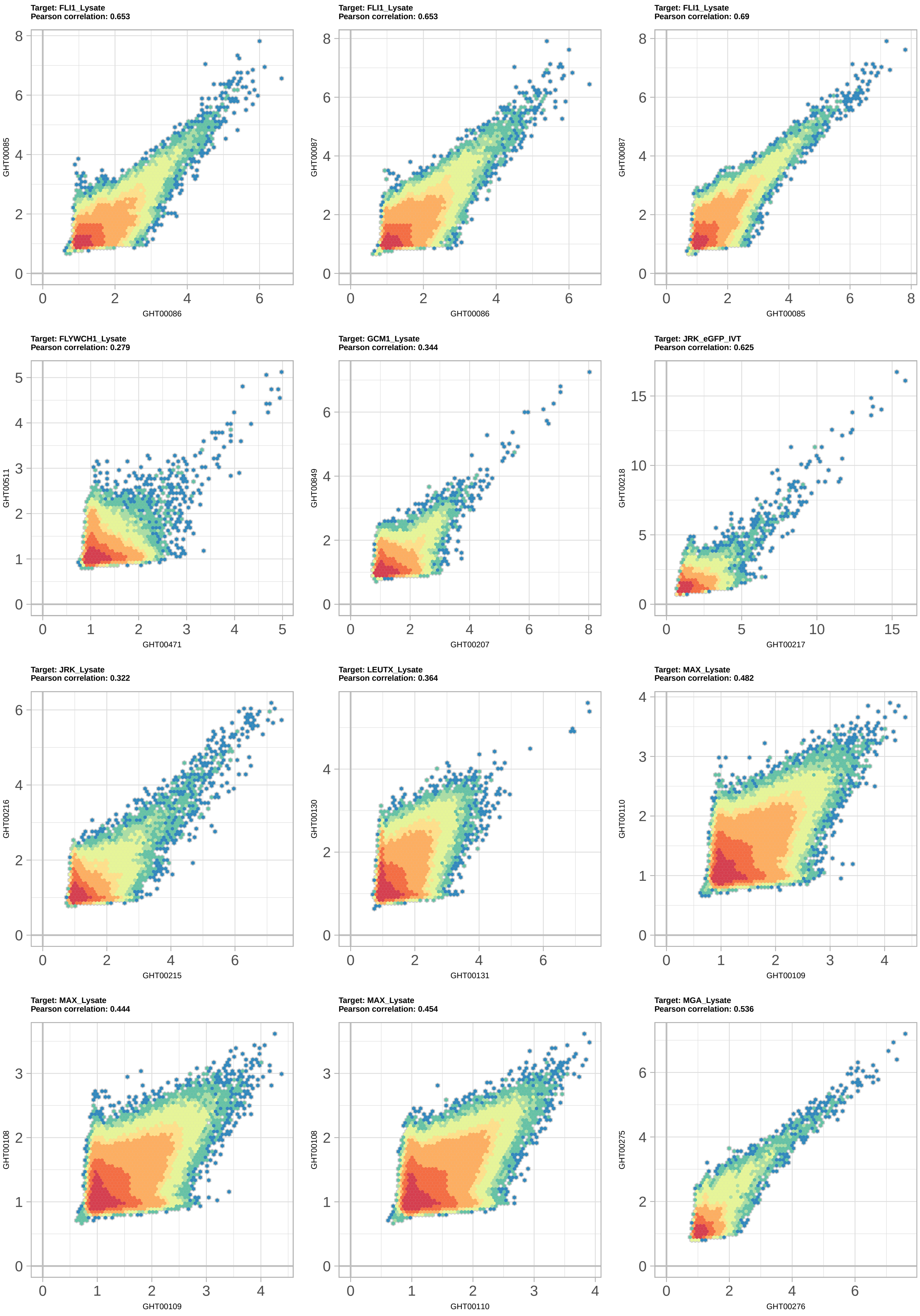

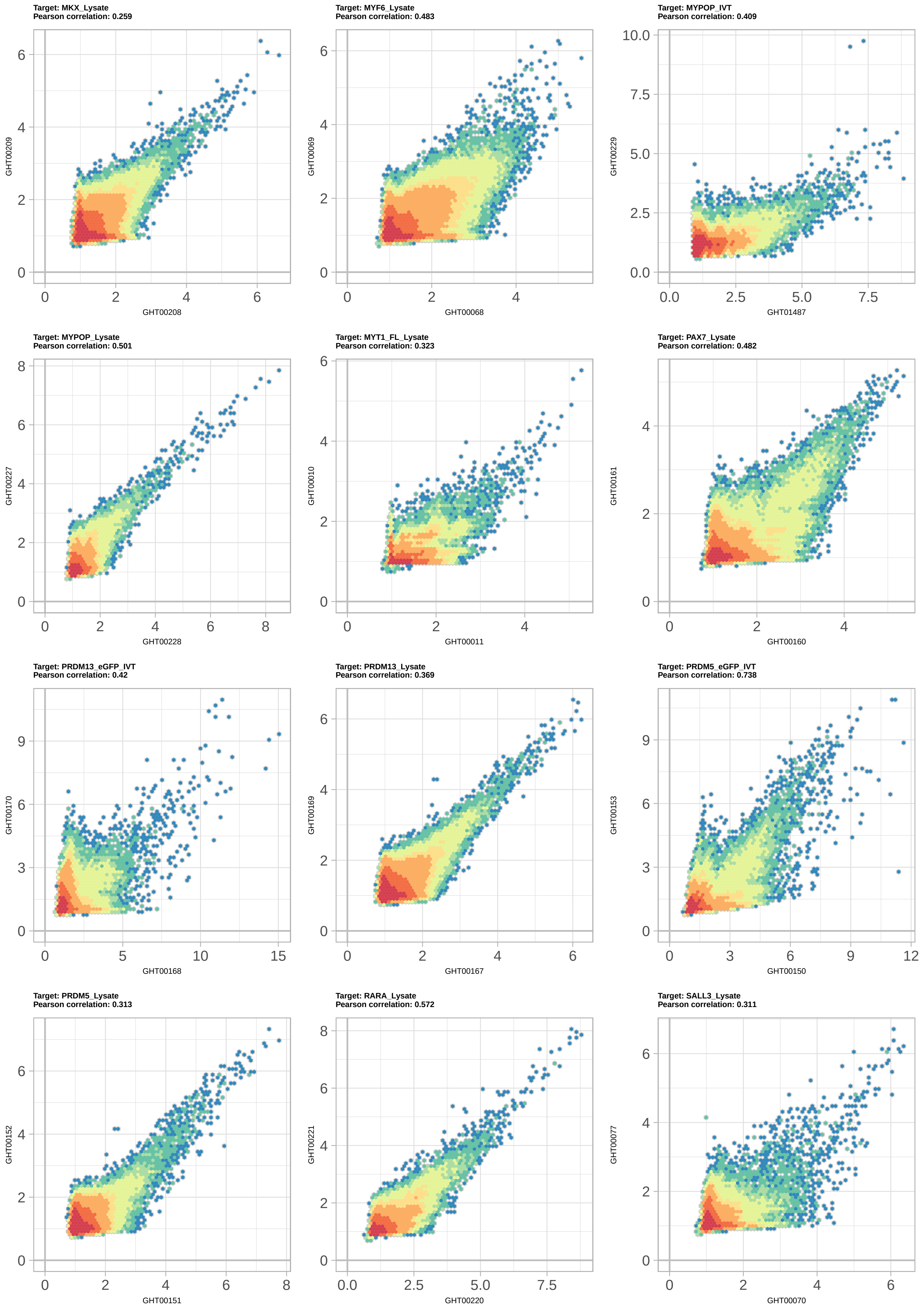

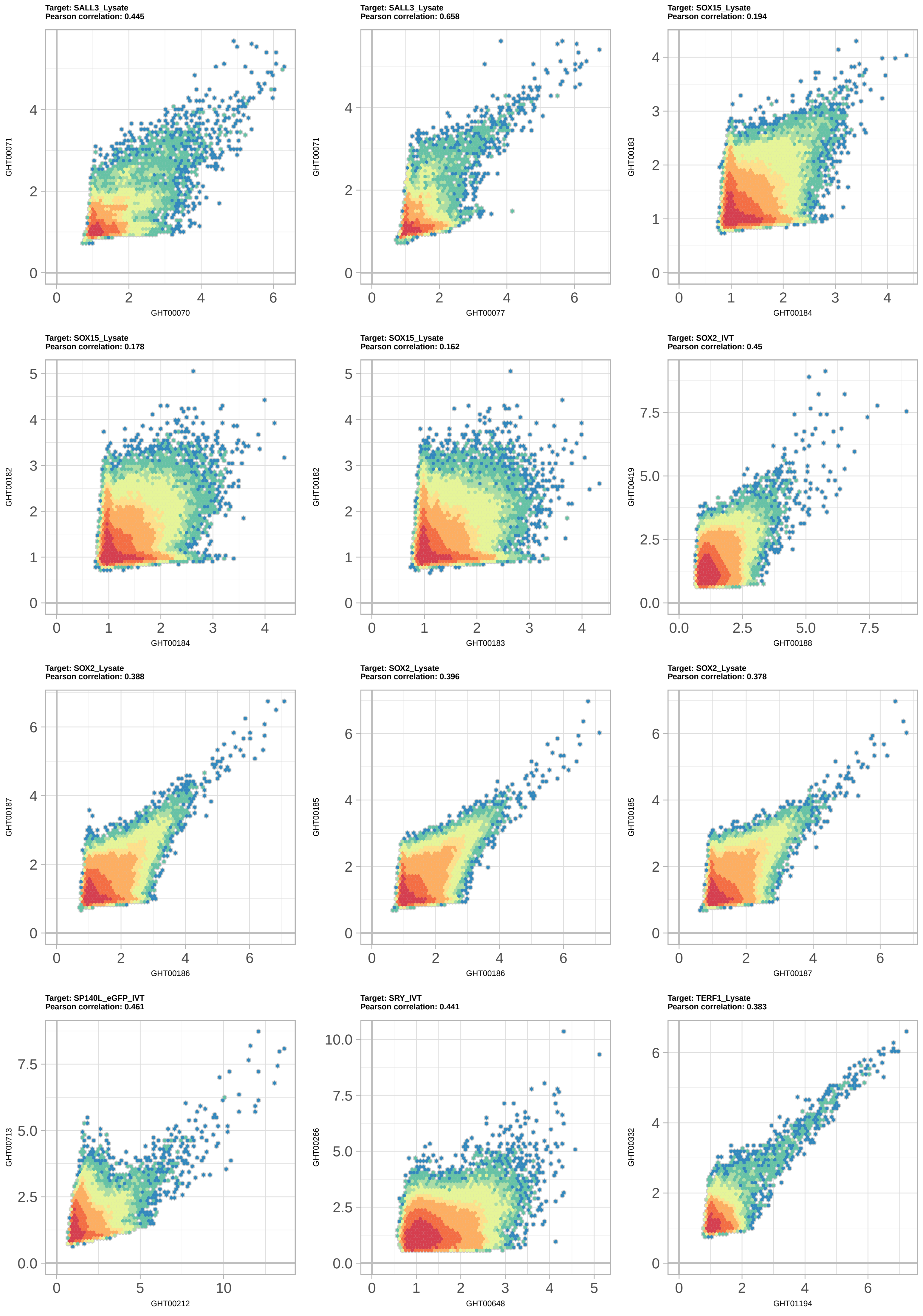

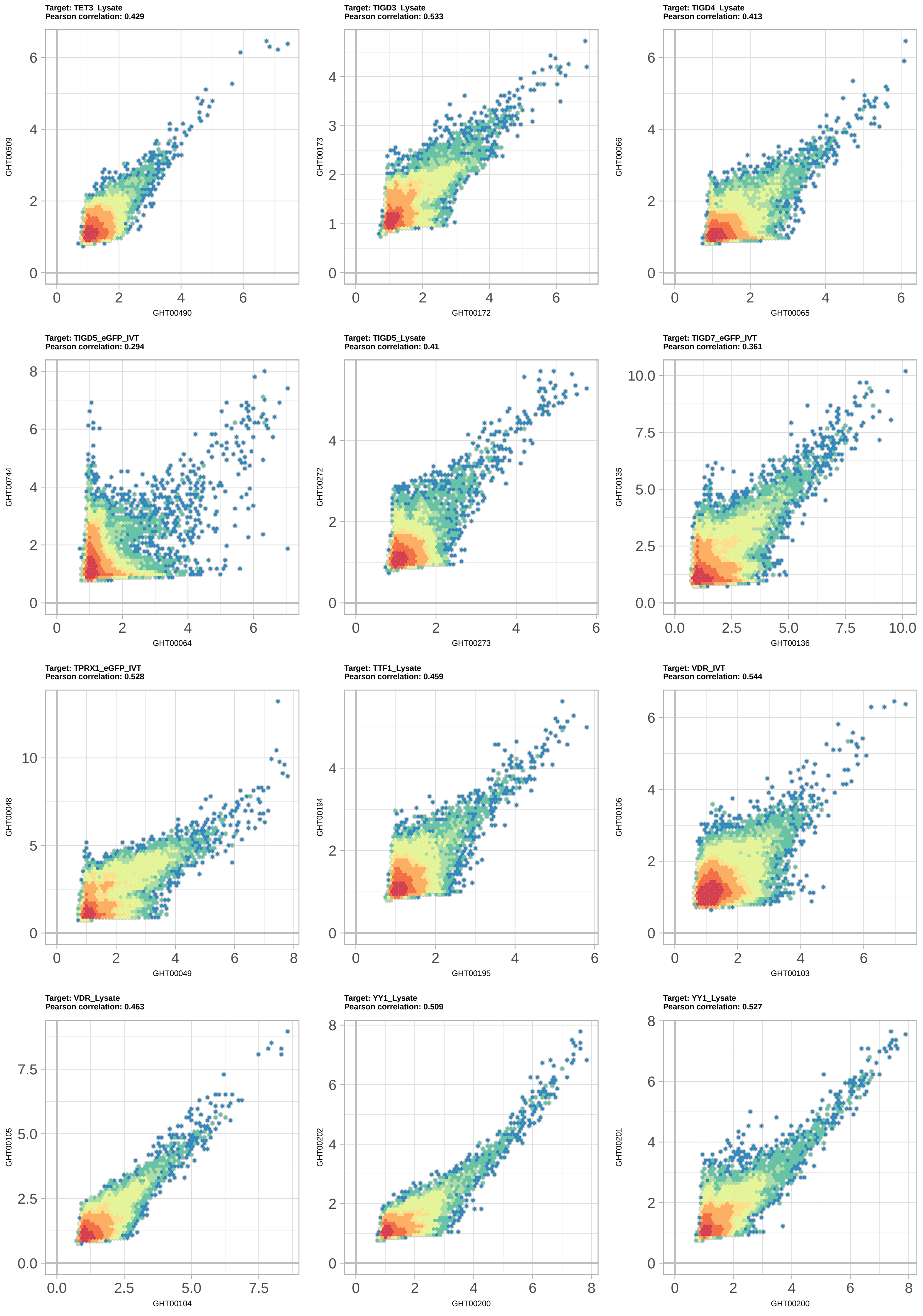

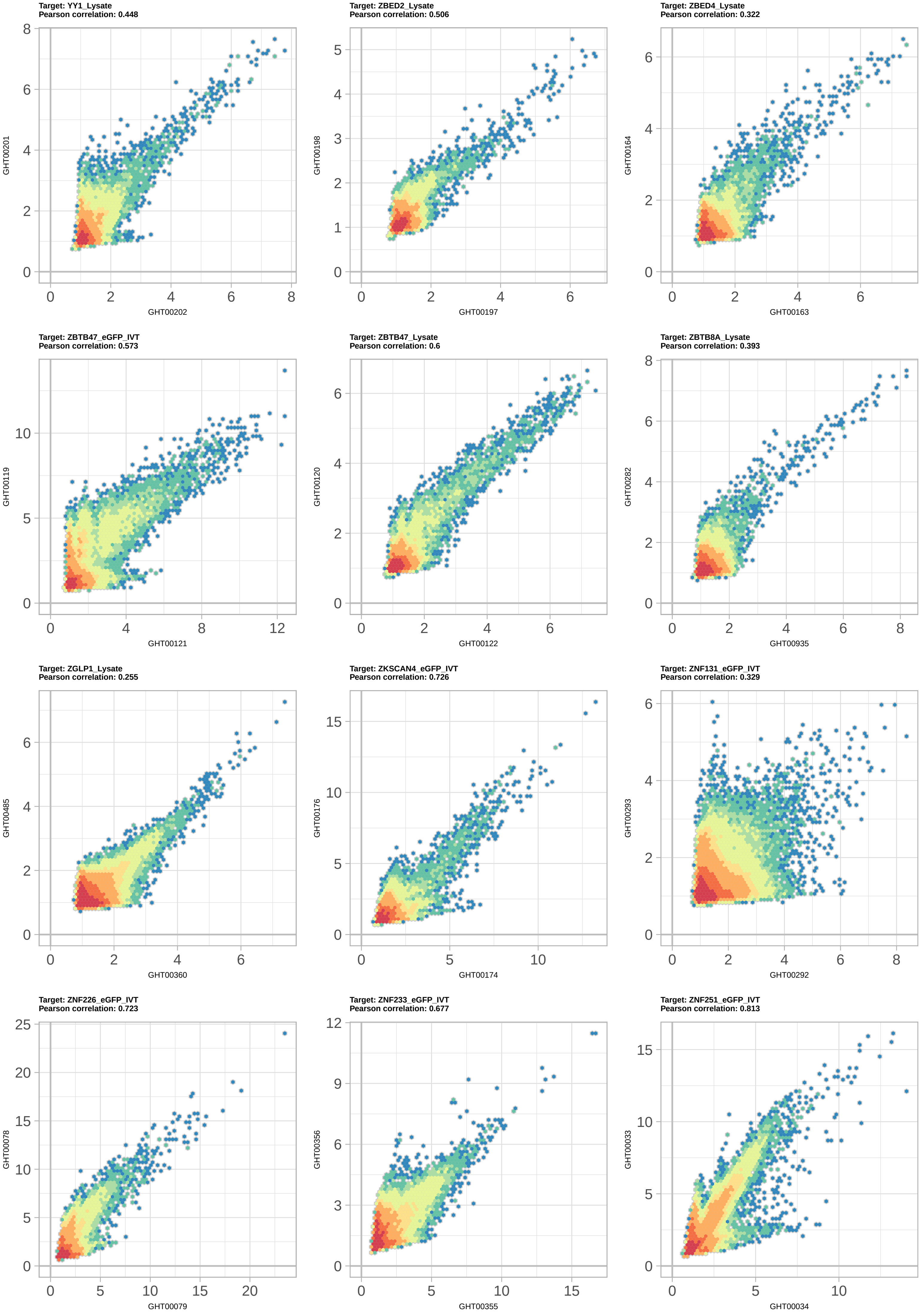

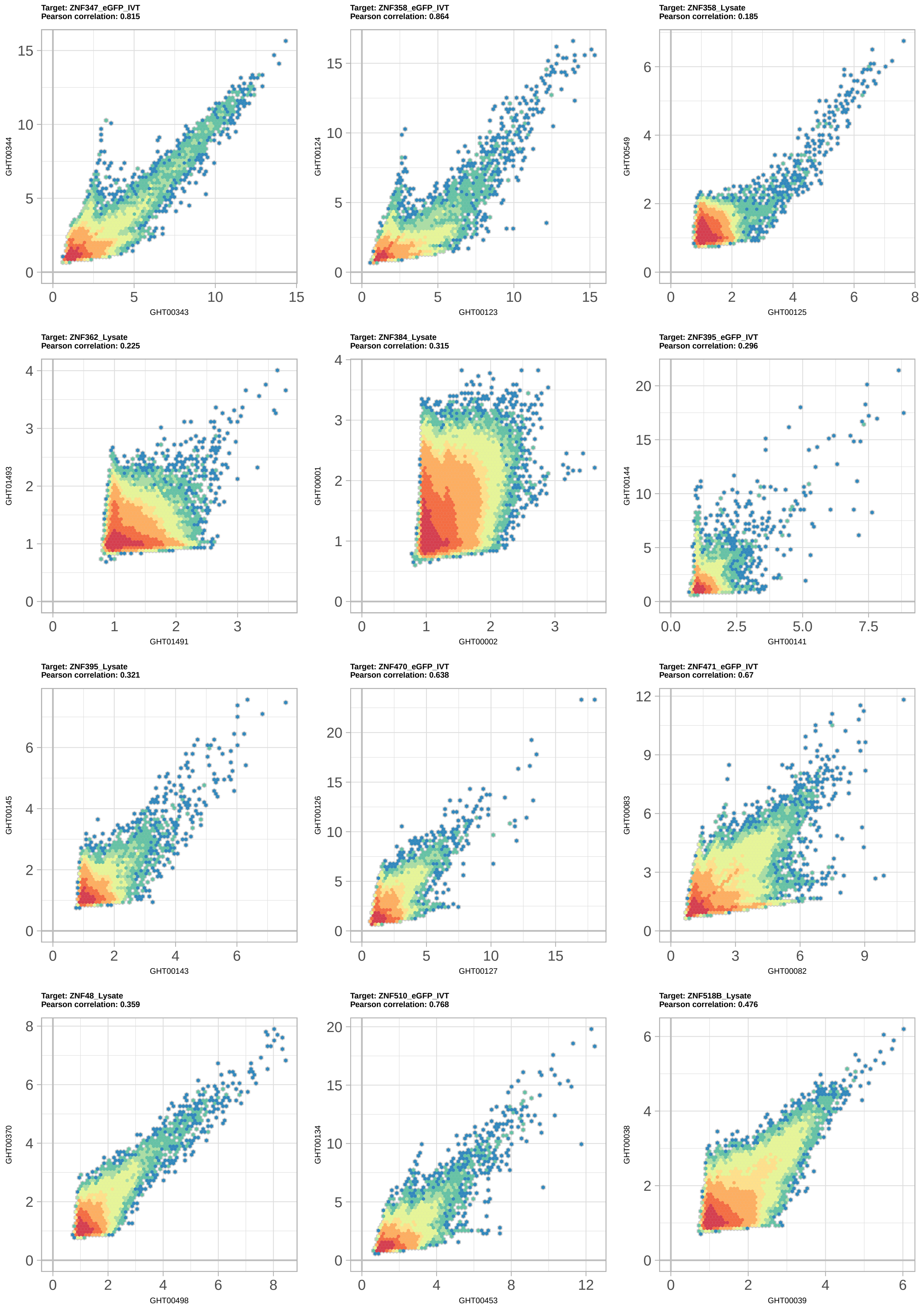

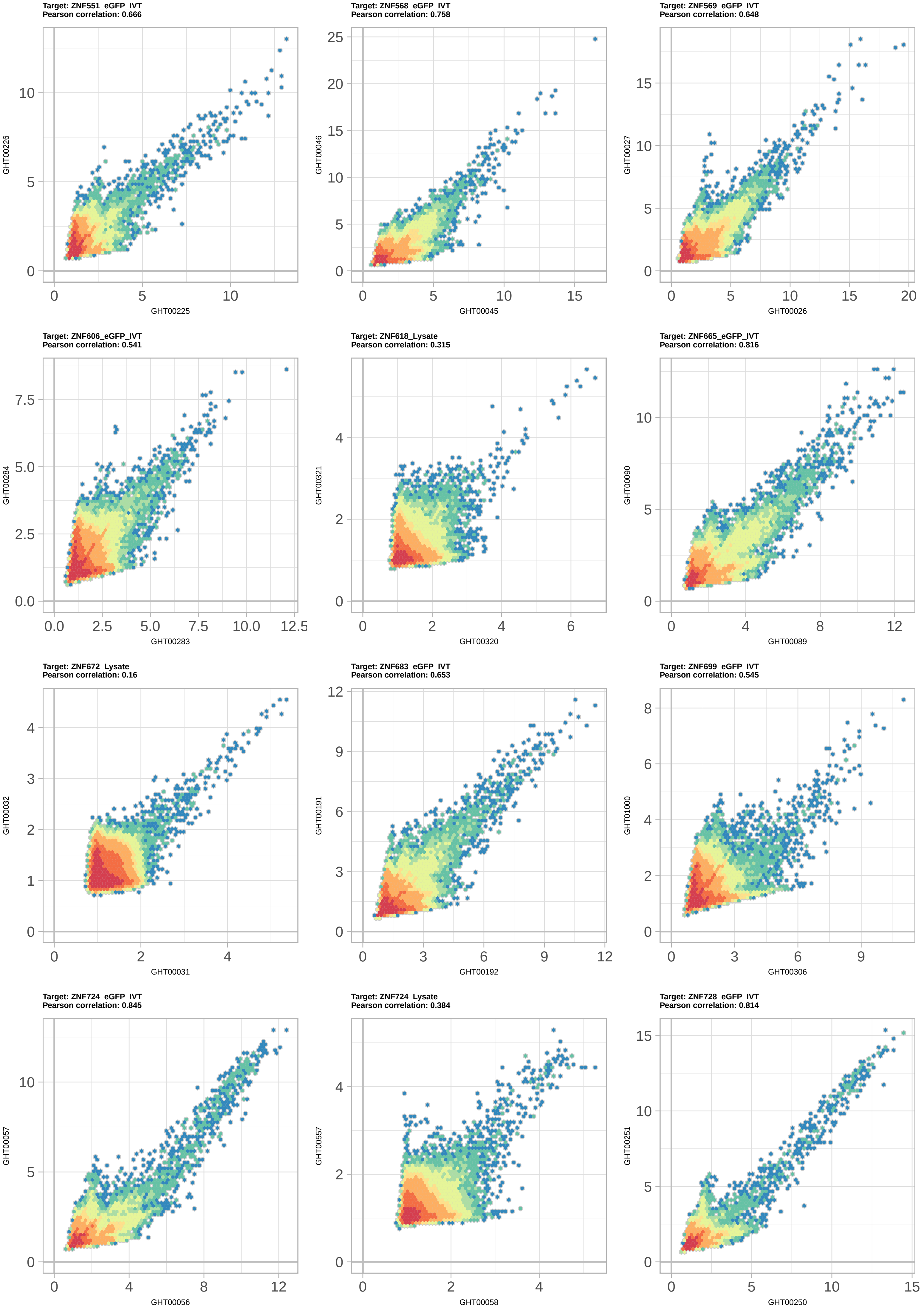

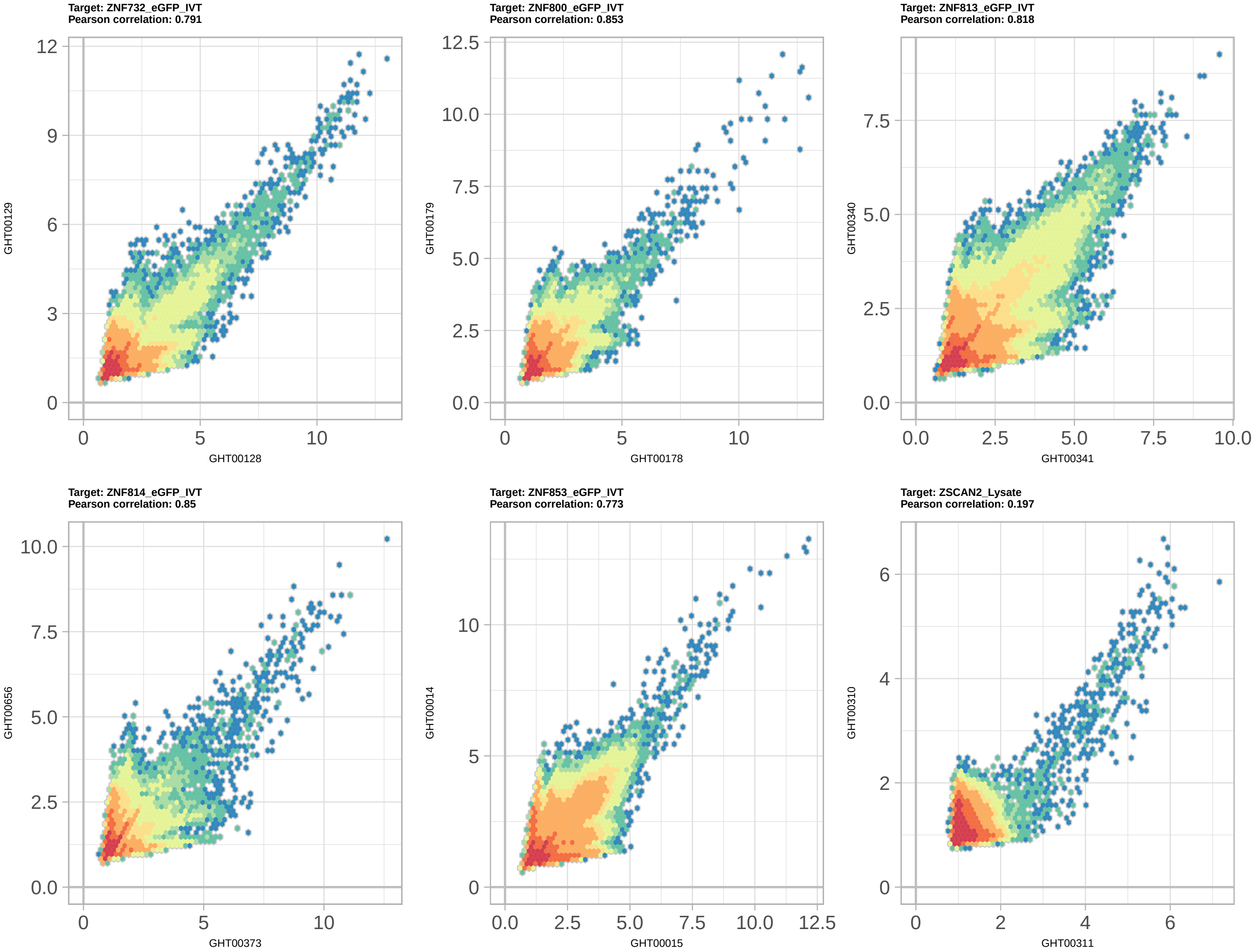

### TIGD3_PWM002267_logo_REV.png

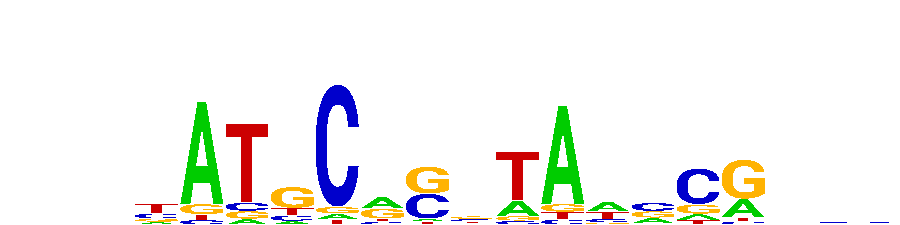

### TIGD4_PWM089799_logo_REV.png

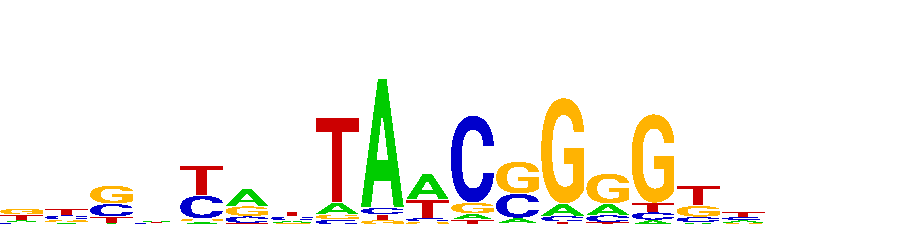

### TSHZ2_PWM002399_logo.png

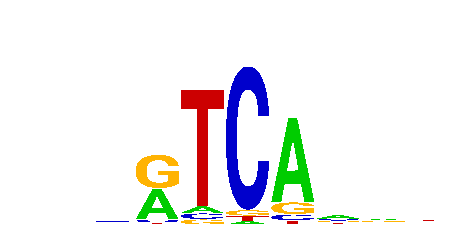

### TSHZ2_PWM002399_logo_REV.png

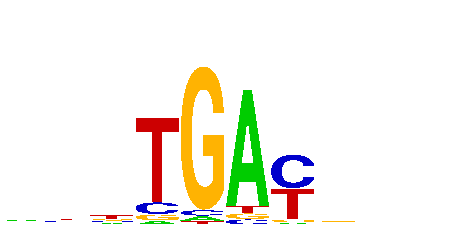

### TTF1_PWM001324_logo.png

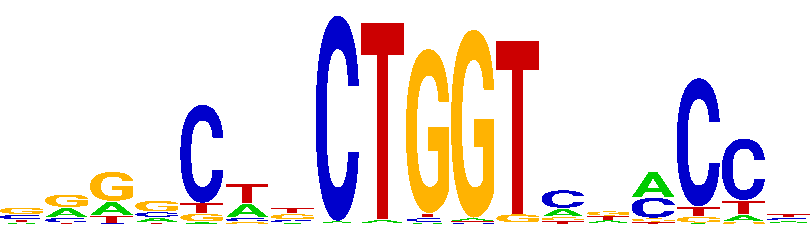

### ZBED2_PWM100759_logo.png

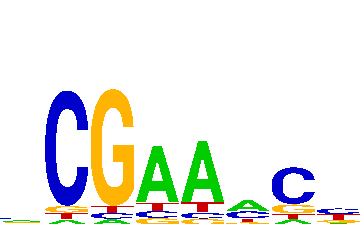

### ZBTB24_PWM002721_logo_REV.png

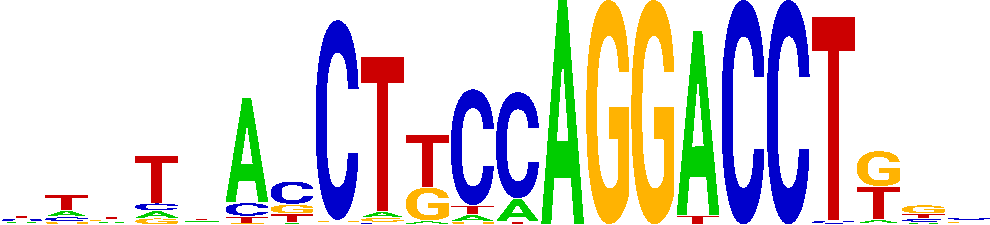

### ZBTB40_PWM002736_logo_REV.png

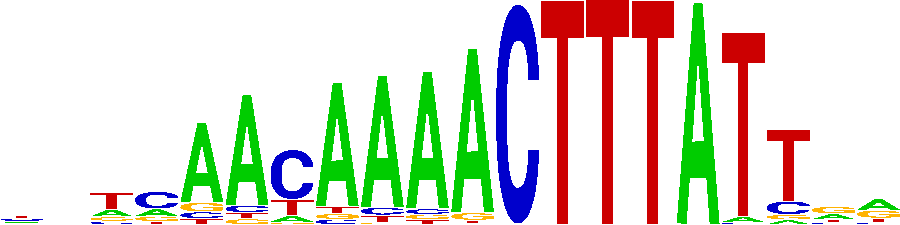

### ZFP3_PWM002952_logo_REV.png

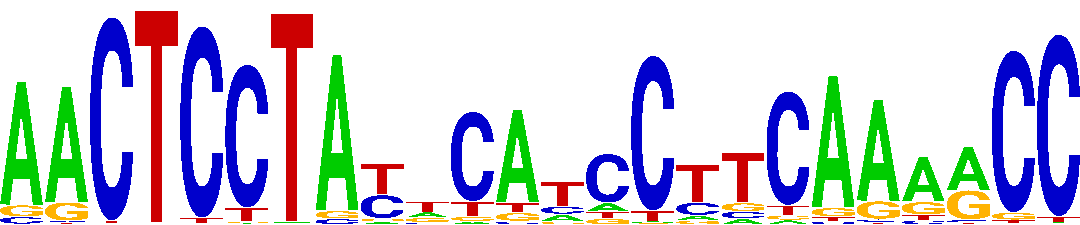

### ZFPM1_PWM114579_logo.png

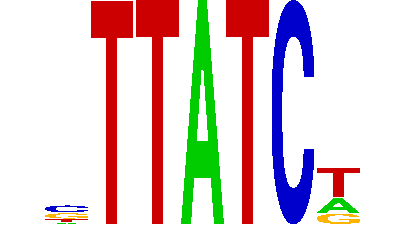

### ZFTA_PWM003007_logo.png

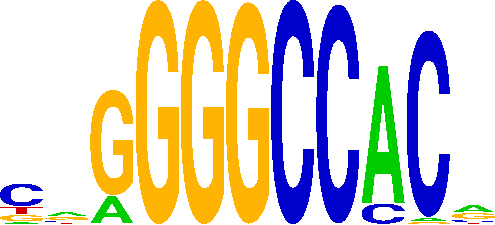

### ZIM3_PWM003081_logo_REV.png

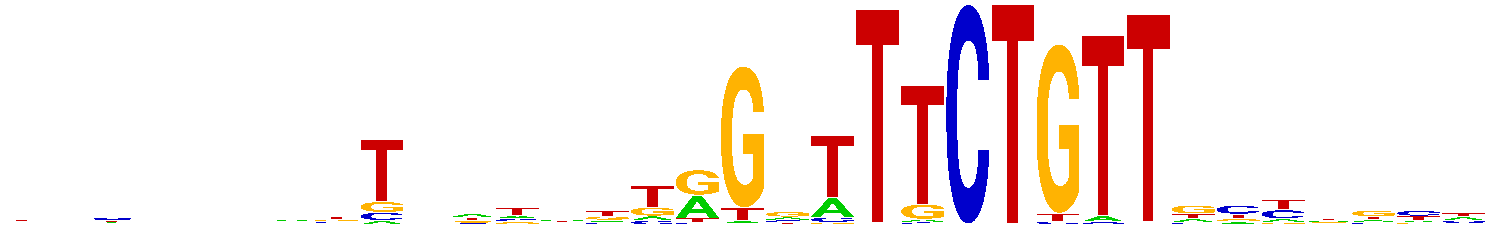

### ZKSCAN4_PWM003117_logo.png

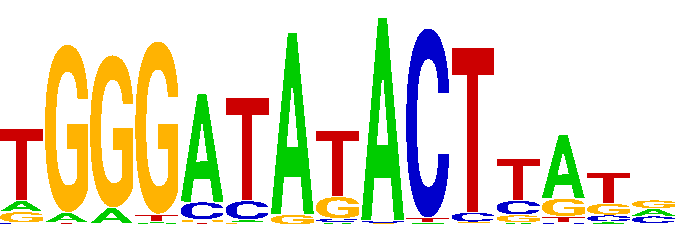

### ZNF8_PWM005820_logo.png

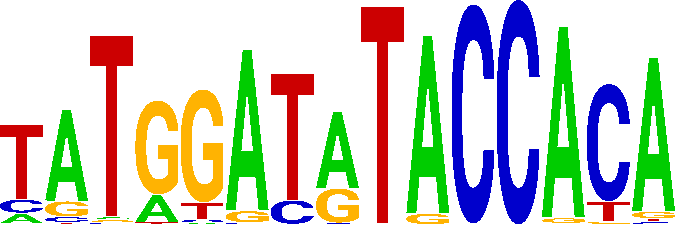

### ZNF18_PWM003405_logo.png

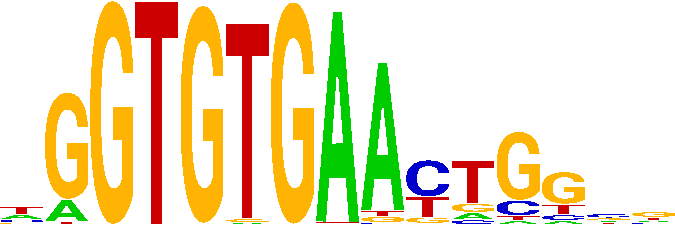

### ZNF43_PWM004214_logo.png

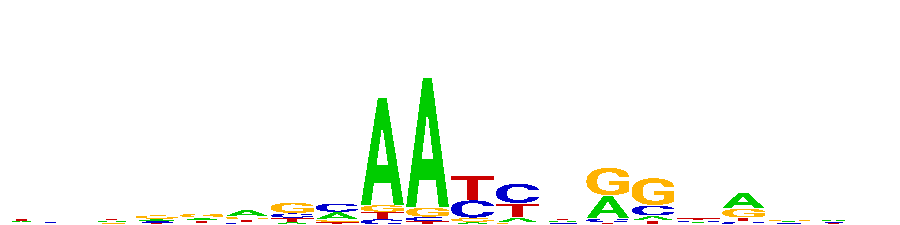

### ZNF134_PWM003246_logo_REV.png

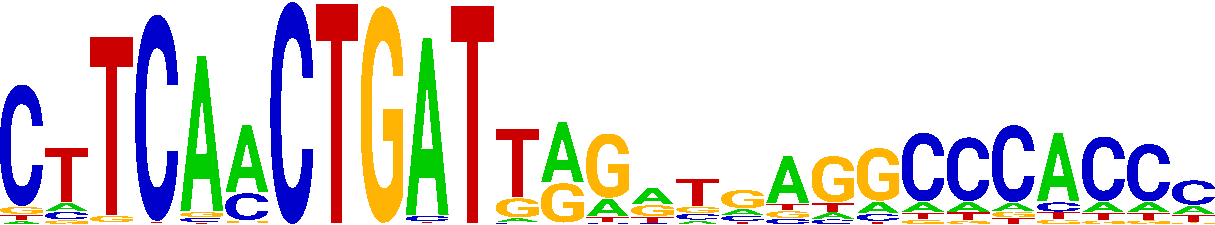

### ZNF140_PWM003275_logo.png

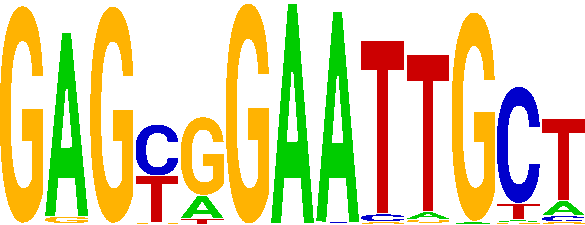

### ZNF142_PWM003289_logo.png

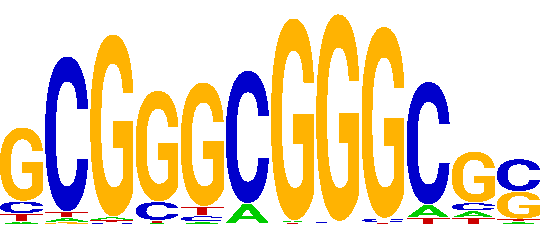

### ZNF142_PWM003289_logo_REV.png

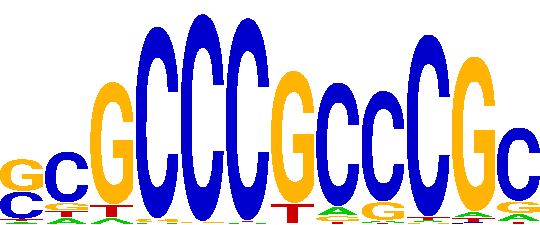

### ZNF146_PWM003340_logo_REV.png

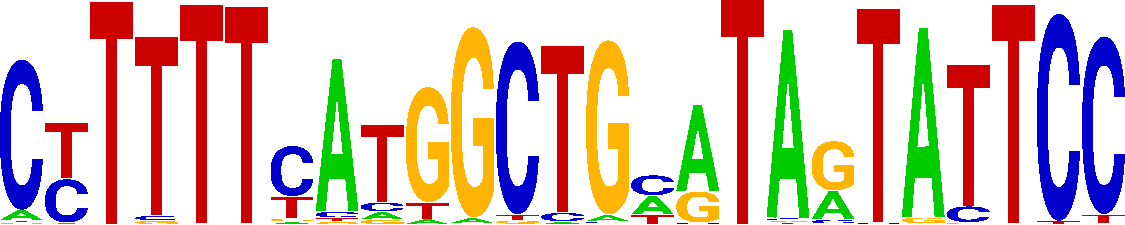

### ZNF208_PWM124632_logo.png

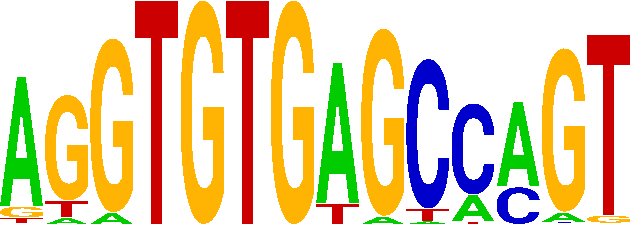

### ZNF215_PWM003472_logo.png

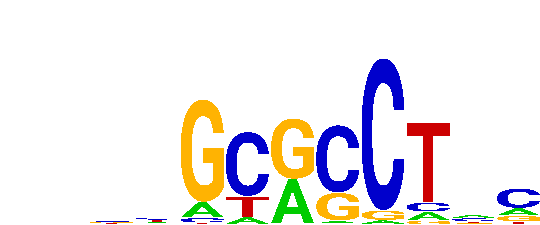

### ZNF226_PWM003524_logo.png

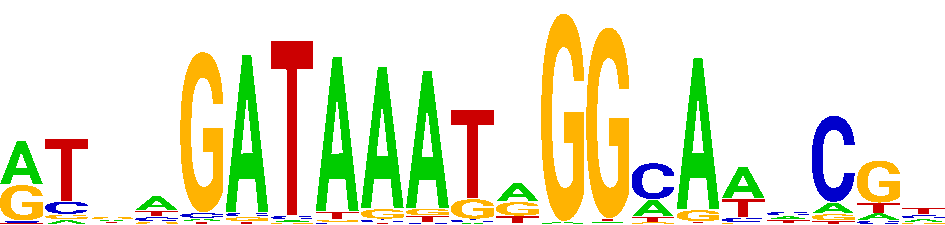

### ZNF229_PWM128094_logo.png

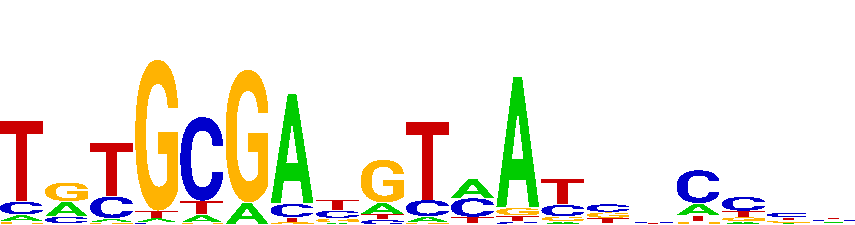

### ZNF250_PWM003653_logo_REV.png

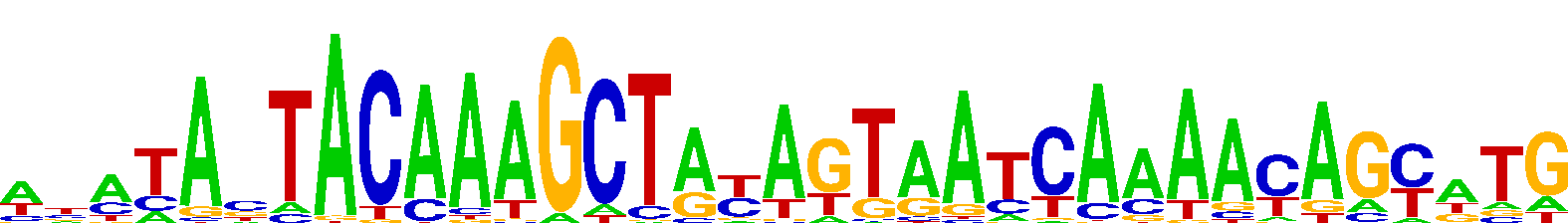

### ZNF260_PWM003684_logo_REV.png

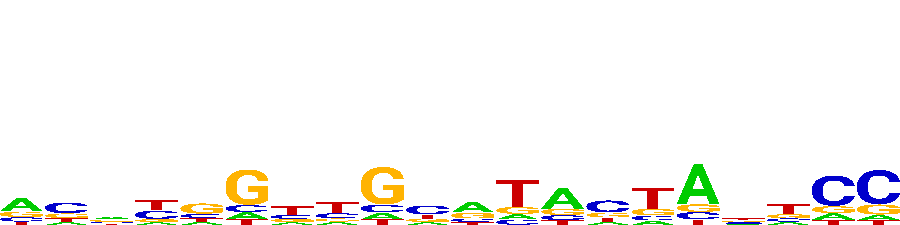

### ZNF275_PWM003748_logo.png

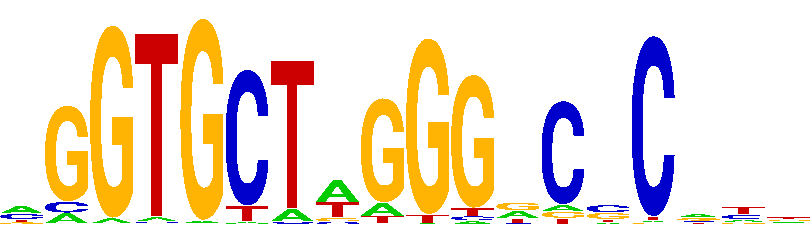

### ZNF275_PWM003748_logo_REV.png

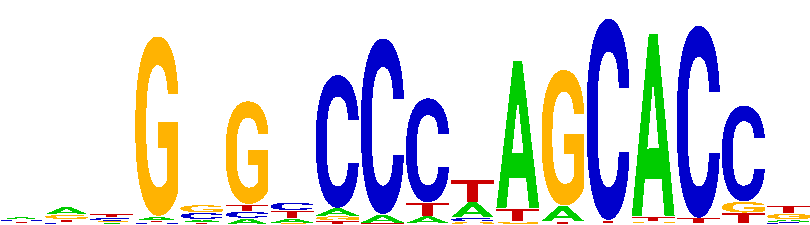

### ZNF286B_PWM003766_logo.png

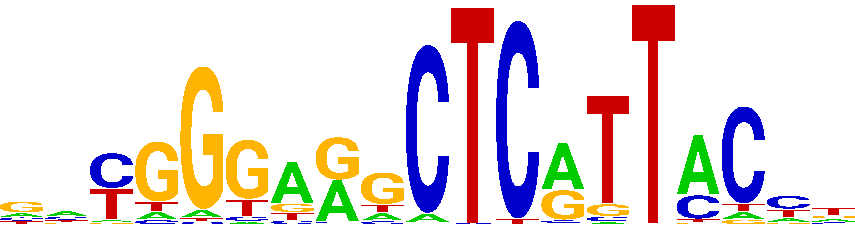

### ZNF347_PWM003932_logo.png

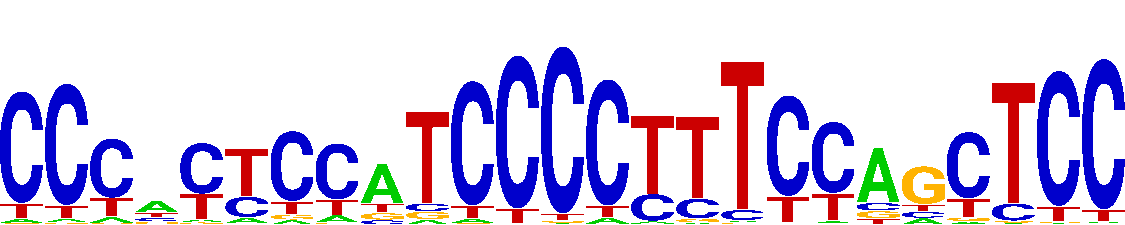

### ZNF347_PWM003932_logo_REV.png

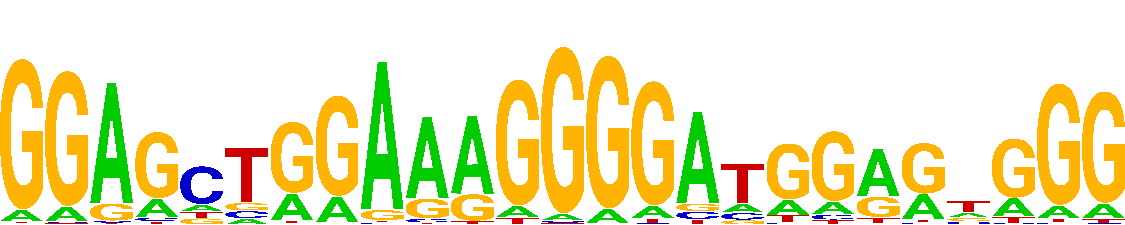

### ZNF358_PWM004007_logo_REV.png

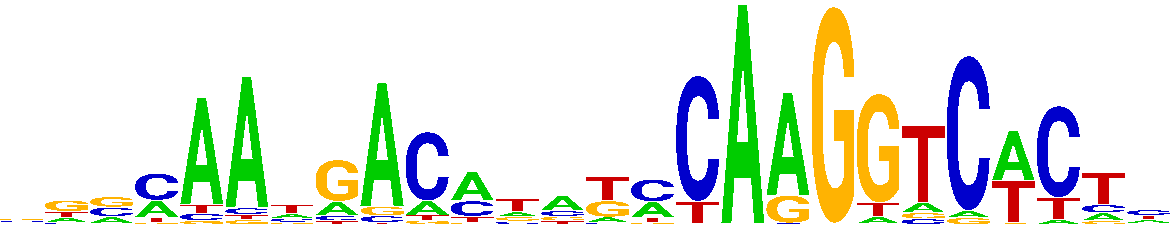

### ZNF362_PWM004025_logo.png

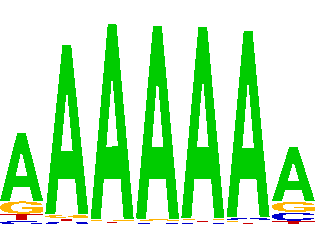

### ZNF384_PWM004138_logo_REV.png

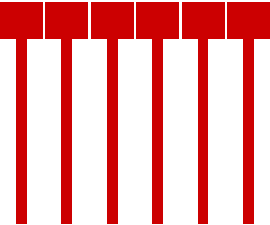

### ZNF395_PWM004142_logo.png

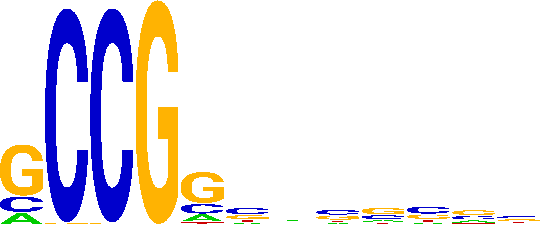

### ZNF436_PWM004241_logo_REV.png

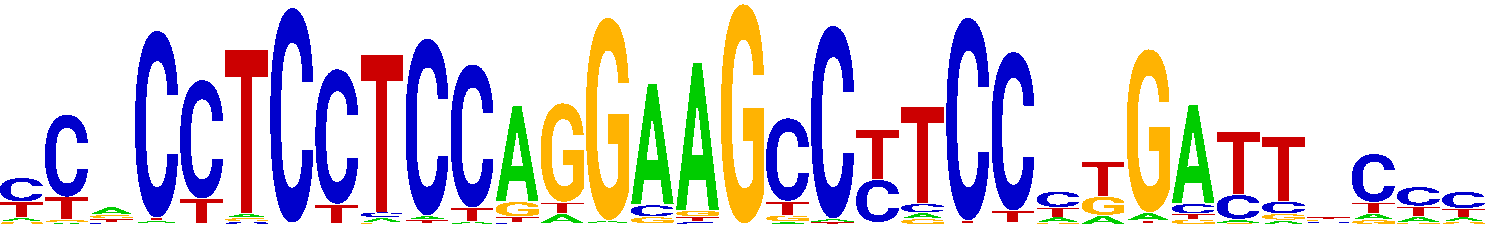

### ZNF470_PWM004290_logo_REV.png

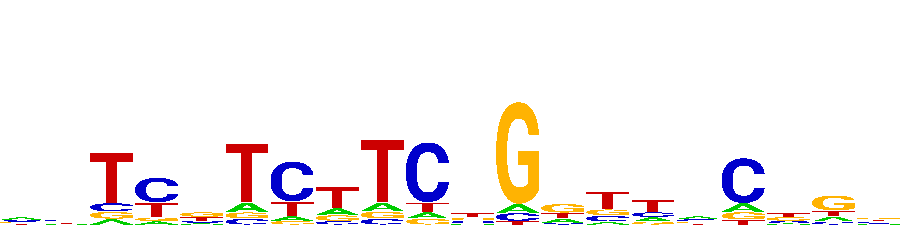

### ZNF471_PWM004322_logo.png

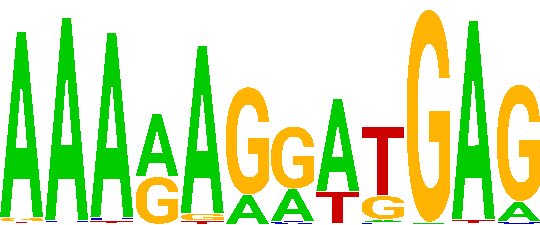

### ZNF493_PWM004421_logo_REV.png

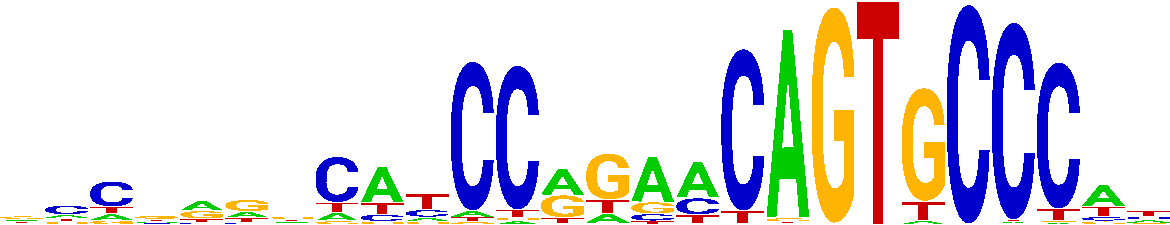

### ZNF500_PWM004464_logo.png

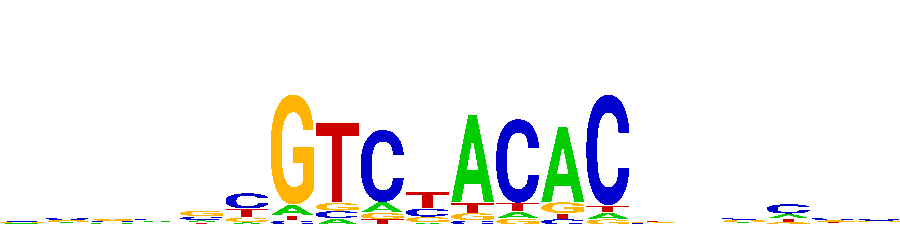

### ZNF503_PWM004483_logo_REV.png

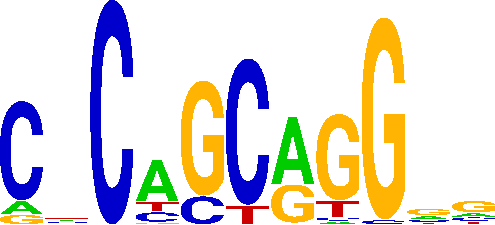

### ZNF526_PWM153677_logo.png

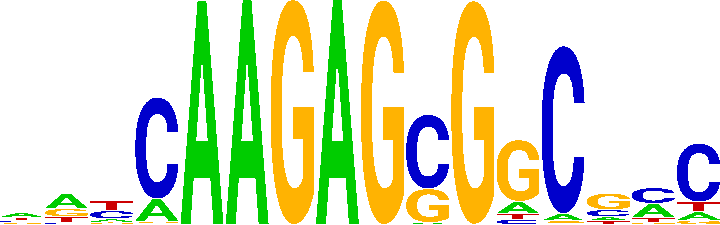

### ZNF569_PWM004692_logo.png

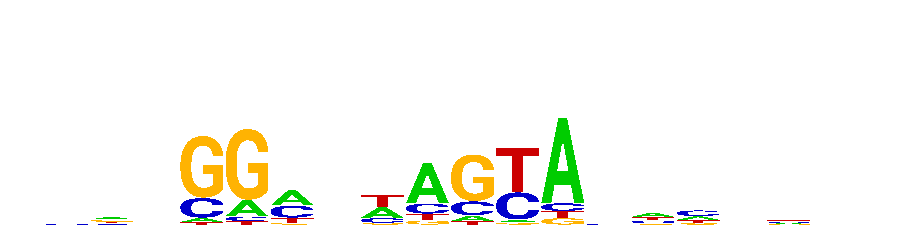

### ZNF575_PWM004745_logo.png

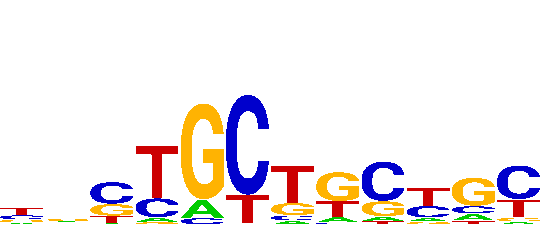

### ZNF618_PWM004873_logo.png

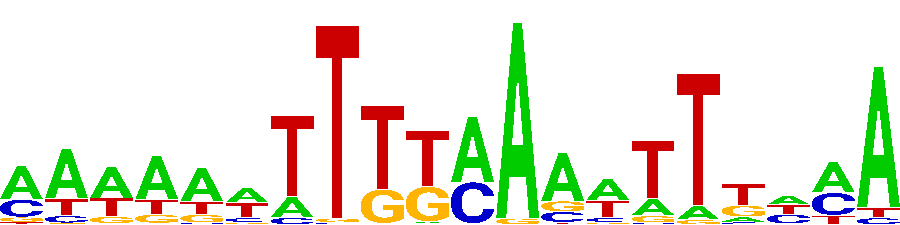

### ZNF668_PWM005084_logo_REV.png

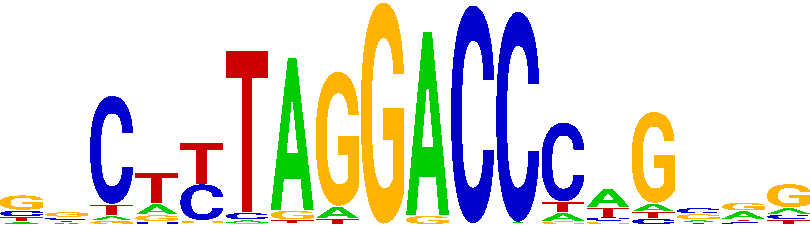

### ZNF672_PWM005120_logo_REV.png

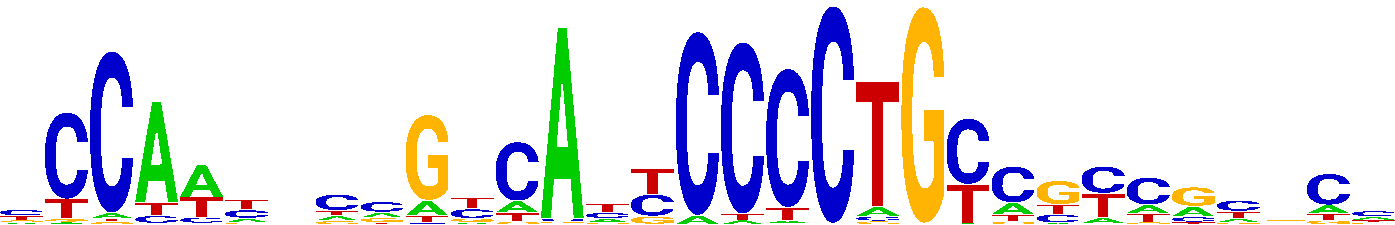

### ZNF689_PWM005261_logo_REV.png

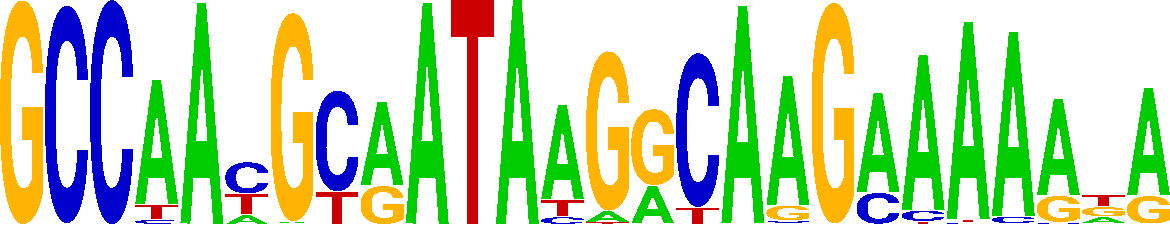

### ZNF770_PWM005686_logo.png

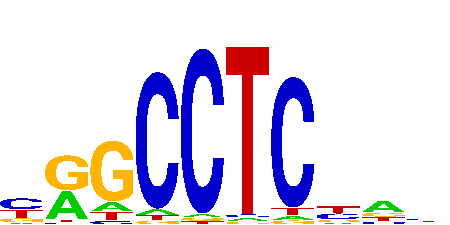

### ZNF775_PWM005778_logo_REV.png

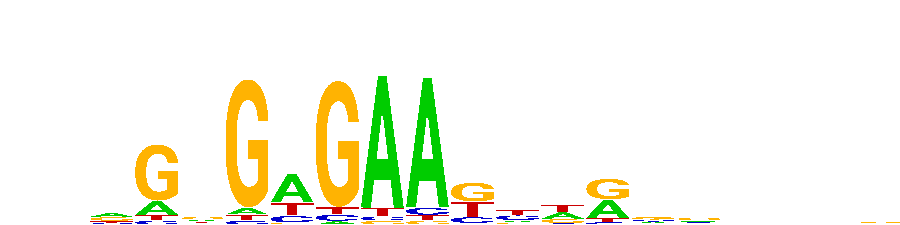

### ZNF780B_PWM005796_logo.png

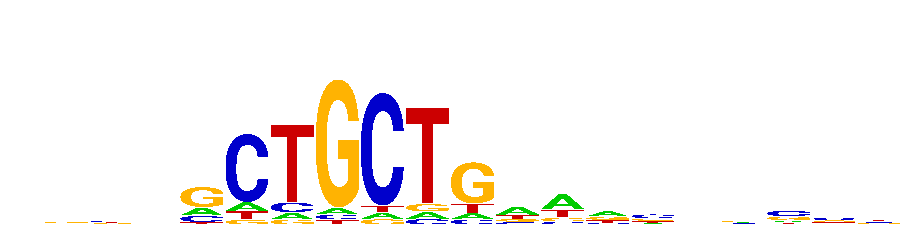

### ZNF788P_PWM185323_logo.png

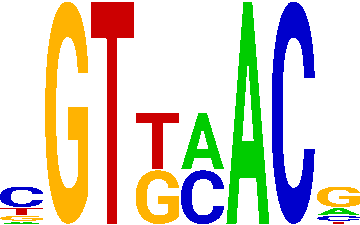

### ZNF836_PWM005990_logo.png

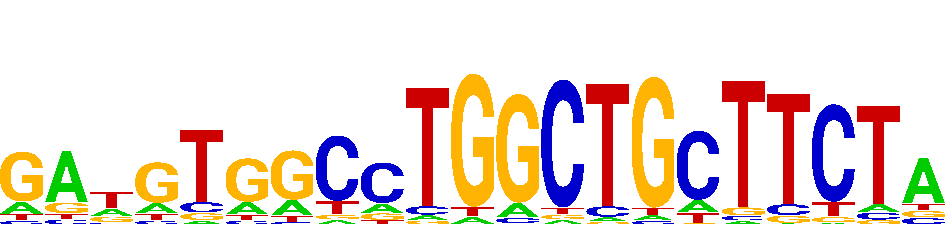

### ZNF845_PWM193356_logo_REV.png

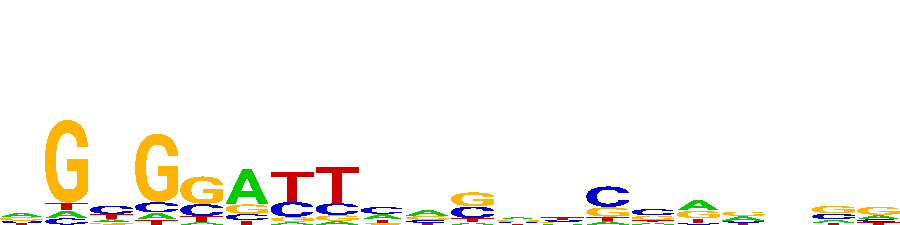

### ZNF888_PWM006179_logo_REV.png

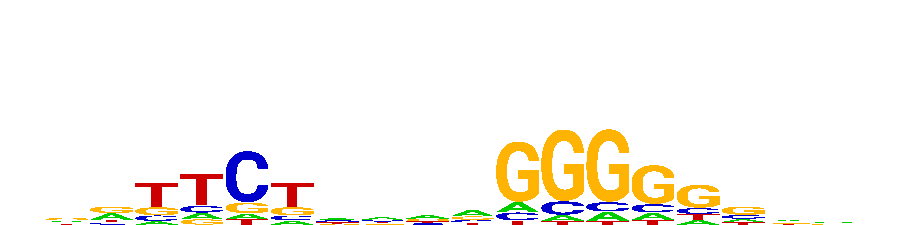

### ZSCAN25_PWM006286_logo_REV.png

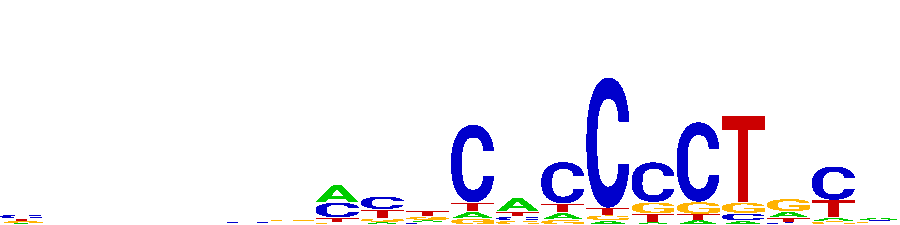

### ZXDC_PWM006370_logo.png

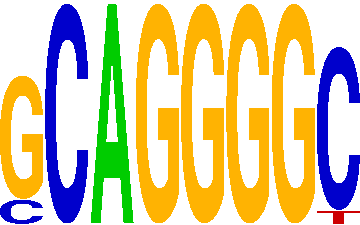
