## Supplementary Document S2 for "GHT-SELEX demonstrates unexpectedly high intrinsic sequence specificity and complex DNA binding of many human transcription factors"

### BATF2

Top GRECO-BIT/MEX PWM: PWM000070

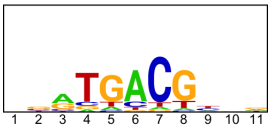


--------------------------------------------------------------------------------------------

Location of top GRECO-BIT/MEX PWM in MAGIX peaks (up to the top 10,000)

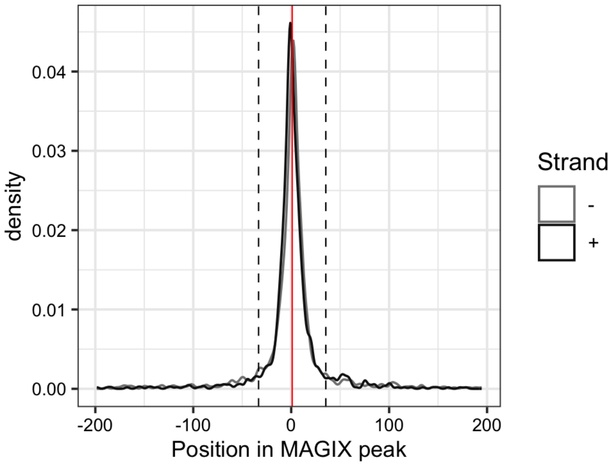


--------------------------------------------------------------------------------------------

AUROC of top GRECO-BIT/MEX PWM in sliding windows (500 peaks, 50 peak step) along the MAGIX peaks (up to the top 75,000) ranked by the MAGIX enrichment coefficient and fraction of overlapping ChIP peaks in the same window

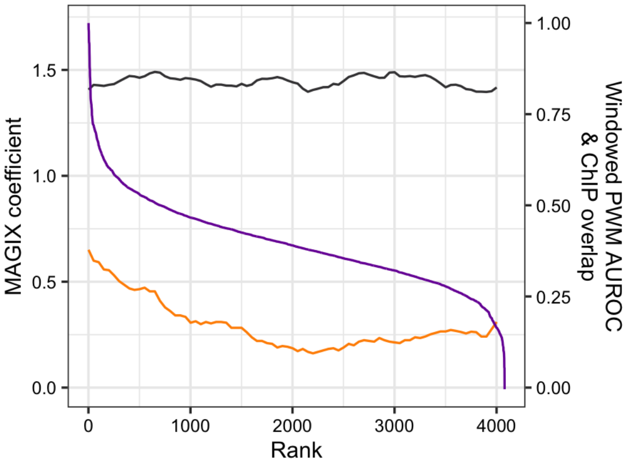

### CAMTA1

Top GRECO-BIT/MEX PWM: PWM000117

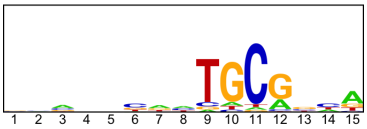


---

Location of top GRECO-BIT/MEX PWM in MAGIX peaks (up to the top 10,000)

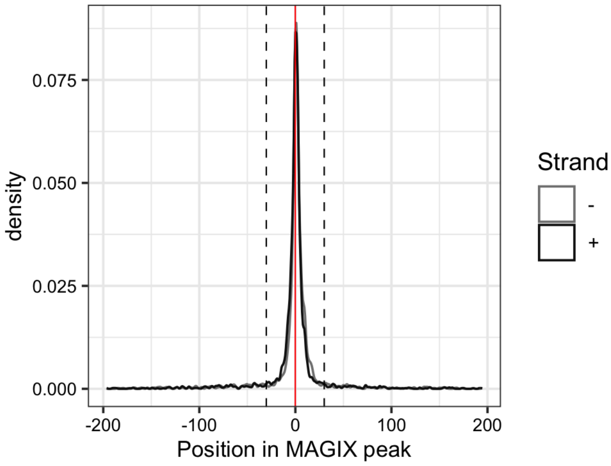


---

AUROC of top GRECO-BIT/MEX PWM in sliding windows (500 peaks, 50 peak step) along the MAGIX peaks (up to the top 75,000) ranked by the MAGIX enrichment coefficient and fraction of overlapping ChIP peaks in the same window

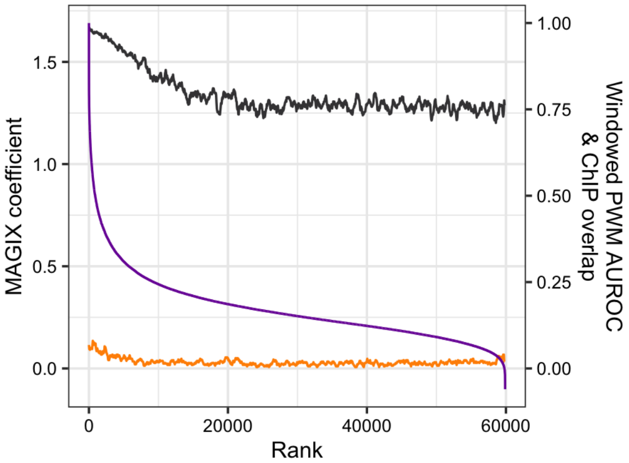

### CAMTA2

Top GRECO-BIT/MEX PWM: PWM000141

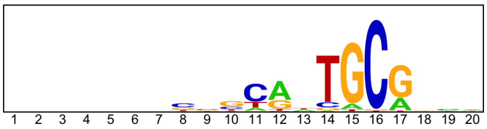


--------------------------------------------------------------------------------------------

Location of top GRECO-BIT/MEX PWM in MAGIX peaks (up to the top 10,000)

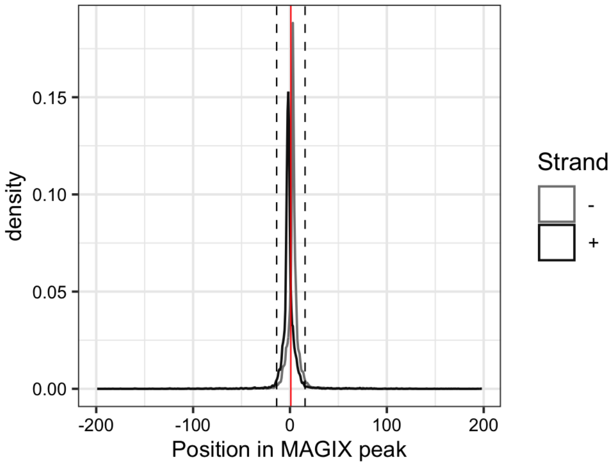


--------------------------------------------------------------------------------------------

AUROC of top GRECO-BIT/MEX PWM in sliding windows (500 peaks, 50 peak step) along the MAGIX peaks (up to the top 75,000) ranked by the MAGIX enrichment coefficient and fraction of overlapping ChIP peaks in the same window

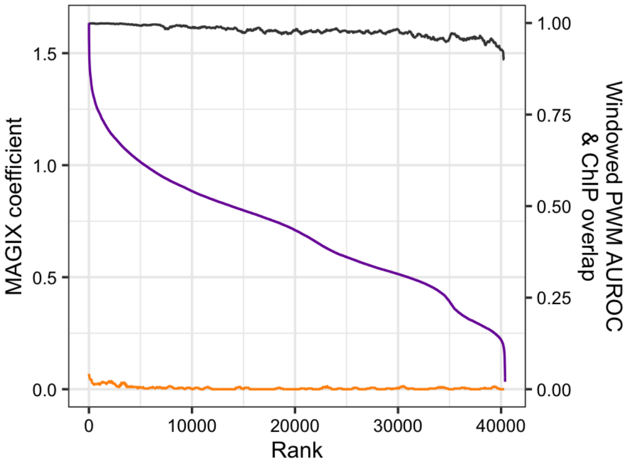

### CASZ1.DBD1

Top GRECO-BIT/MEX PWM: PWM000184

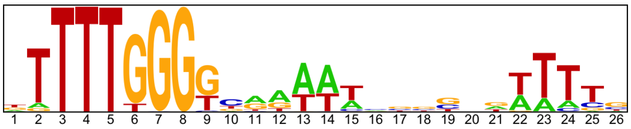


------------------------------------------------------------------------------------------------

Location of top GRECO-BIT/MEX PWM in MAGIX peaks (up to the top 10,000)

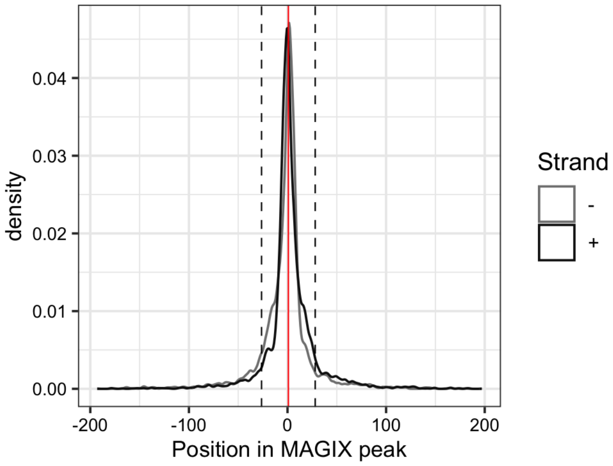

### CASZ1.FL

Top GRECO-BIT/MEX PWM: PWM000181

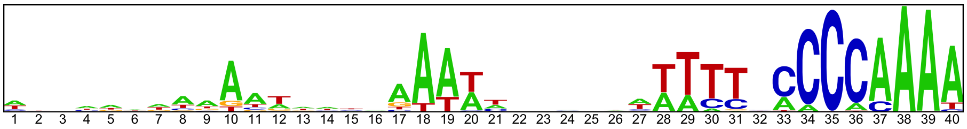


------------------------------------------------------------------------------------------------

Location of top GRECO-BIT/MEX PWM in MAGIX peaks (up to the top 10,000)

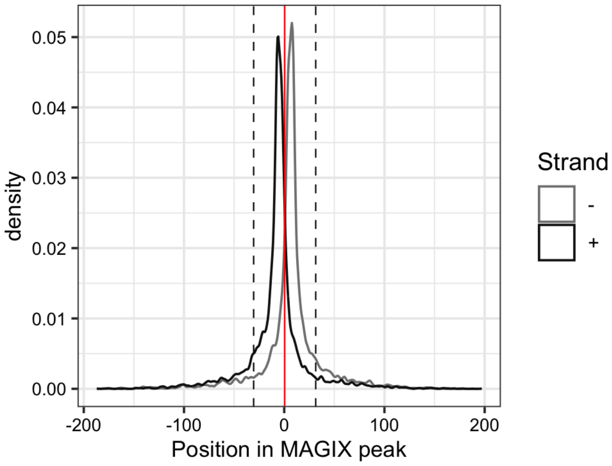

### CGGBP1

Top GRECO-BIT/MEX PWM: PWM000216

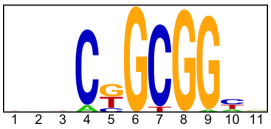


------------------------------------------------------------------------------------------------

Location of top GRECO-BIT/MEX PWM in MAGIX peaks (up to the top 10,000)

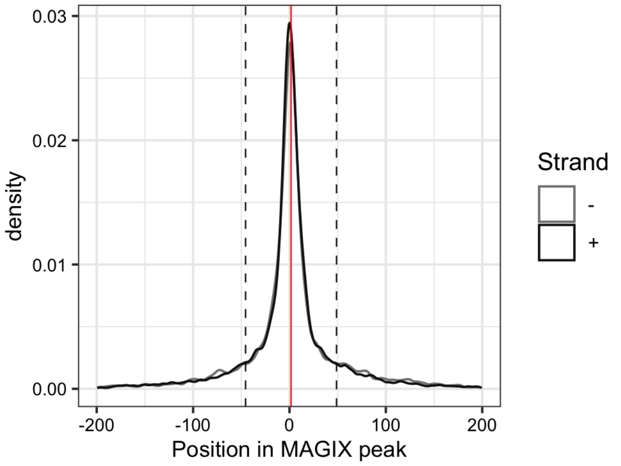


------------------------------------------------------------------------------------------------

AUROC of top GRECO-BIT/MEX PWM in sliding windows (500 peaks, 50 peak step) along the MAGIX peaks (up to the top 75,000) ranked by the MAGIX enrichment coefficient and fraction of overlapping ChIP peaks in the same window

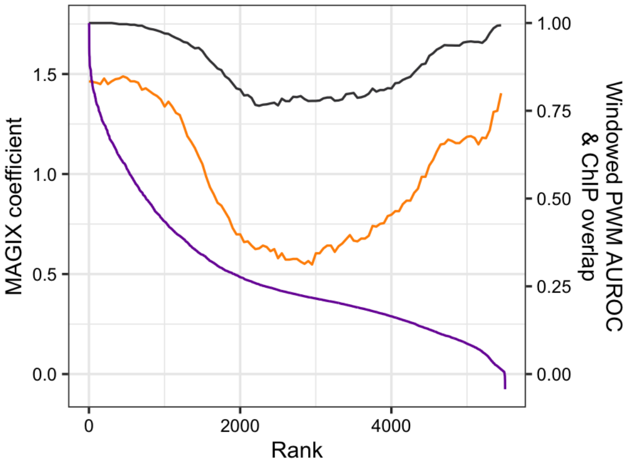

### CREB3L3

Top GRECO-BIT/MEX PWM: PWM000262

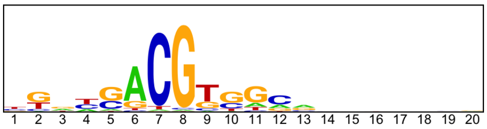


----------------------------------------------------------------------------------------------------

Location of top GRECO-BIT/MEX PWM in MAGIX peaks (up to the top 10,000)

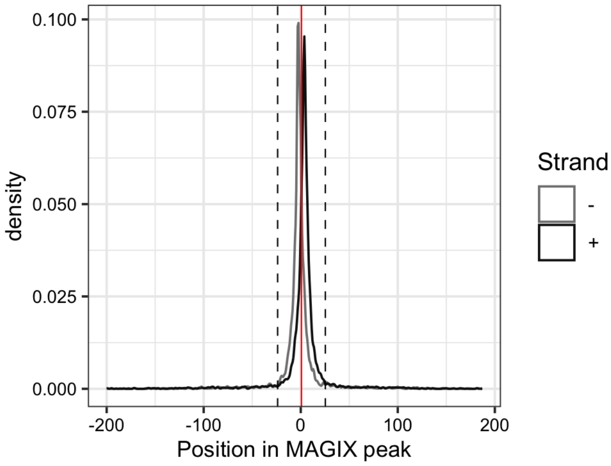


----------------------------------------------------------------------------------------------------

AUROC of top GRECO-BIT/MEX PWM in sliding windows (500 peaks, 50 peak step) along the MAGIX peaks (up to the top 75,000) ranked by the MAGIX enrichment coefficient and fraction of overlapping ChIP peaks in the same window

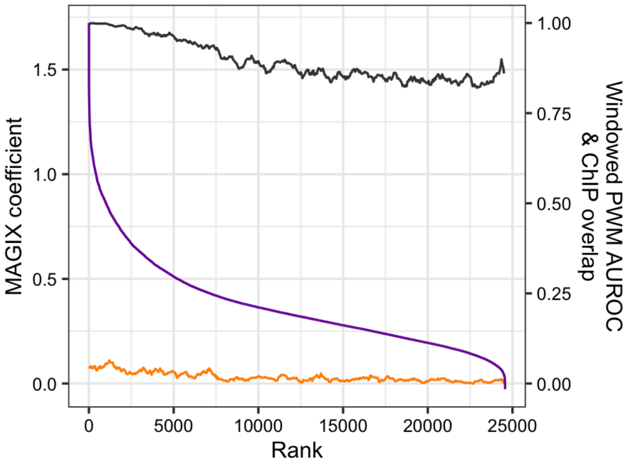

### CTCF

Top GRECO-BIT/MEX PWM: PWM000307

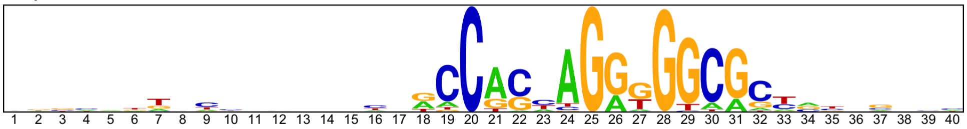

---

Location of top GRECO-BIT/MEX PWM in MAGIX peaks (up to the top 10,000)

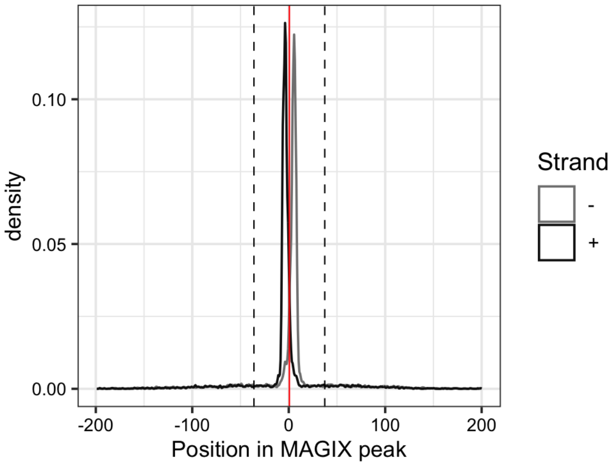


---

AUROC of top GRECO-BIT/MEX PWM in sliding windows (500 peaks, 50 peak step) along the MAGIX peaks (up to the top 75,000) ranked by the MAGIX enrichment coefficient and fraction of overlapping ChIP peaks in the same window

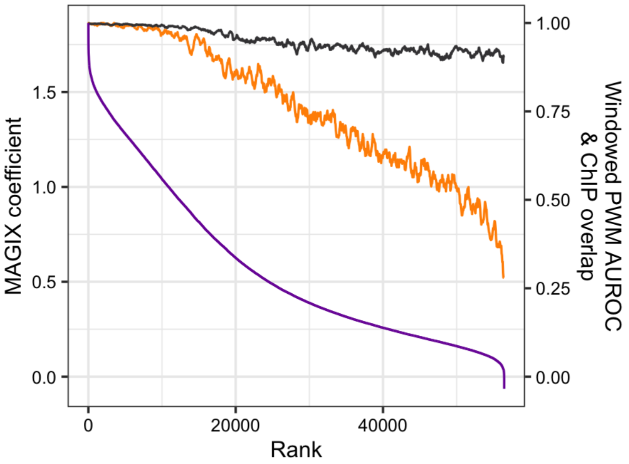

### CXXC4

Top GRECO-BIT/MEX PWM: PWM000326

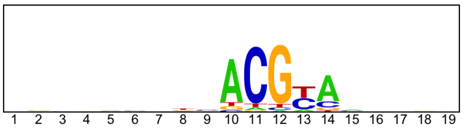


---

Location of top GRECO-BIT/MEX PWM in MAGIX peaks (up to the top 10,000)

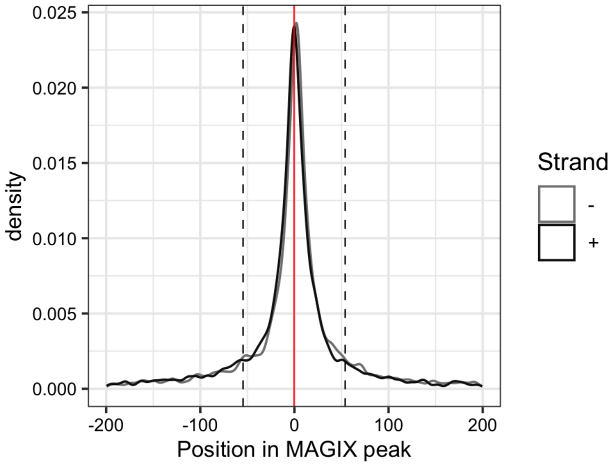


---

AUROC of top GRECO-BIT/MEX PWM in sliding windows (500 peaks, 50 peak step) along the MAGIX peaks (up to the top 75,000) ranked by the MAGIX enrichment coefficient and fraction of overlapping ChIP peaks in the same window

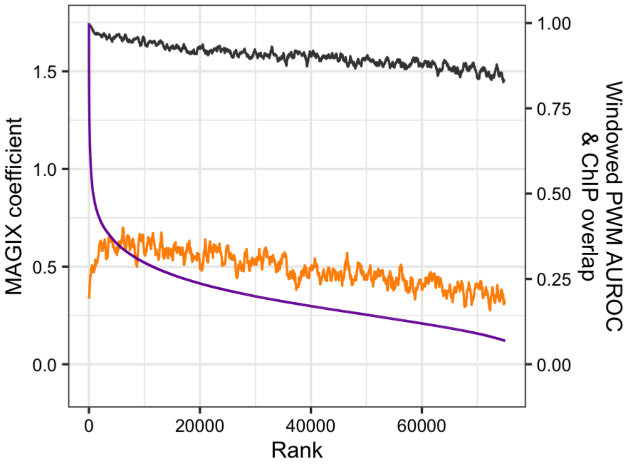

### DMTF1

Top GRECO-BIT/MEX PWM: PWM000383

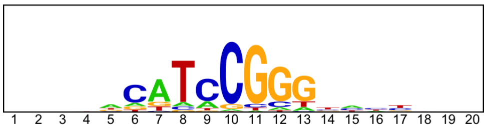


---

Location of top GRECO-BIT/MEX PWM in MAGIX peaks (up to the top 10,000)

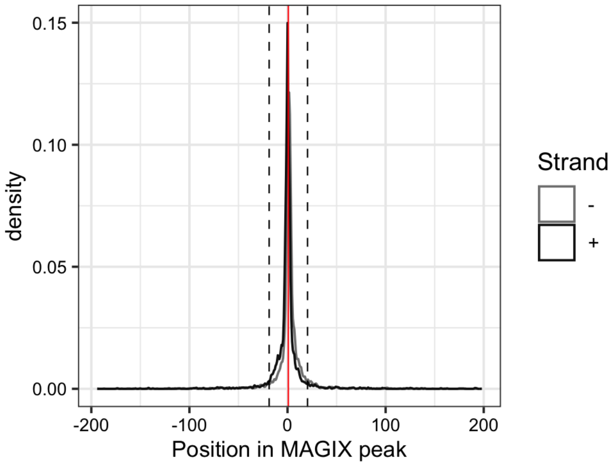


---

AUROC of top GRECO-BIT/MEX PWM in sliding windows (500 peaks, 50 peak step) along the MAGIX peaks (up to the top 75,000) ranked by the MAGIX enrichment coefficient and fraction of overlapping ChIP peaks in the same window

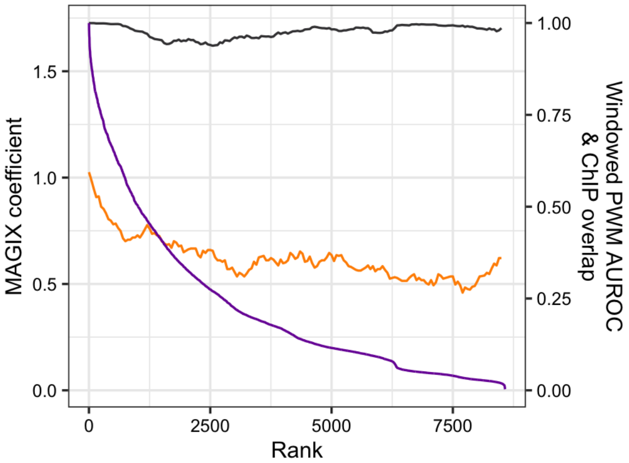

### DNTTIP1

Top GRECO-BIT/MEX PWM: PWM000422

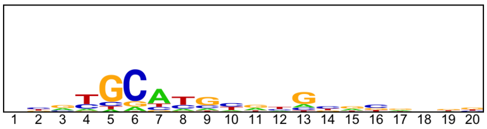


---

Location of top GRECO-BIT/MEX PWM in MAGIX peaks (up to the top 10,000)

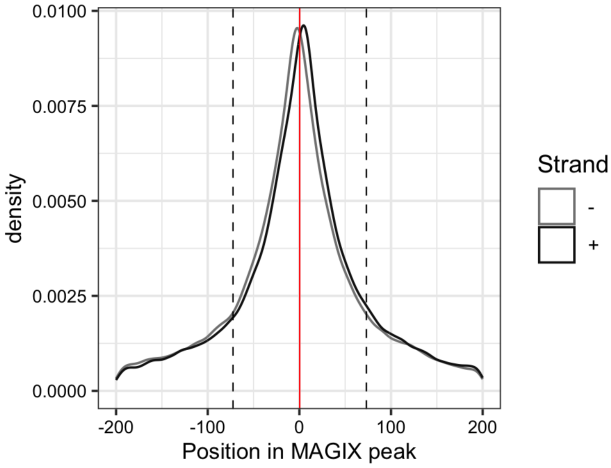


---

AUROC of top GRECO-BIT/MEX PWM in sliding windows (500 peaks, 50 peak step) along the MAGIX peaks (up to the top 75,000) ranked by the MAGIX enrichment coefficient and fraction of overlapping ChIP peaks in the same window

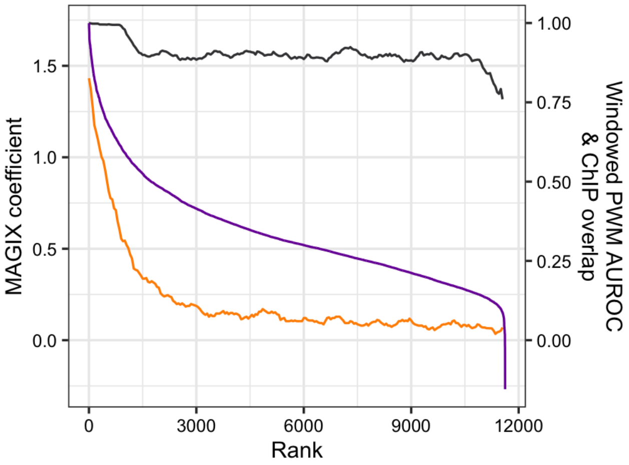

### ELF3

Top GRECO-BIT/MEX PWM: PWM000447

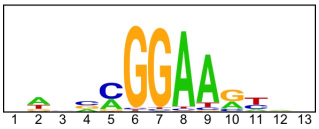


---

Location of top GRECO-BIT/MEX PWM in MAGIX peaks (up to the top 10,000)

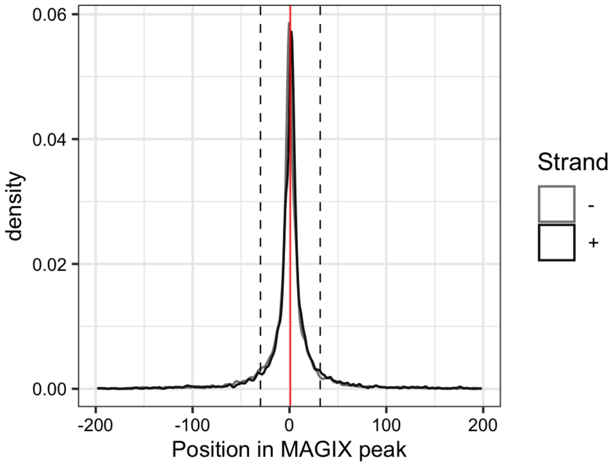


---

AUROC of top GRECO-BIT/MEX PWM in sliding windows (500 peaks, 50 peak step) along the MAGIX peaks (up to the top 75,000) ranked by the MAGIX enrichment coefficient and fraction of overlapping ChIP peaks in the same window

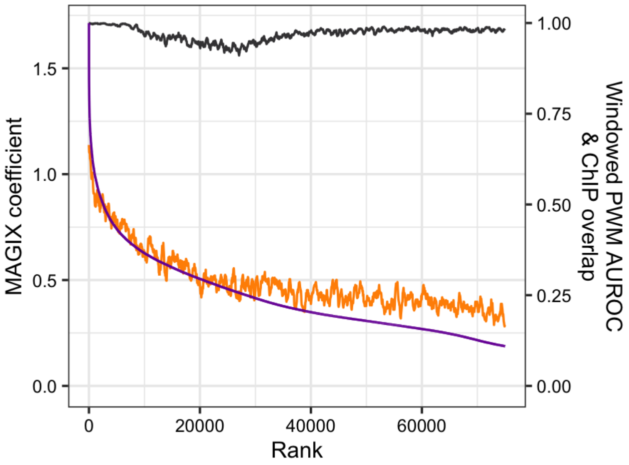

### FAM200B

Top GRECO-BIT/MEX PWM: PWM000481

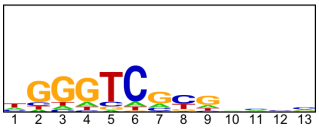


--------------------------------------------------------------------------------------------

Location of top GRECO-BIT/MEX PWM in MAGIX peaks (up to the top 10,000)

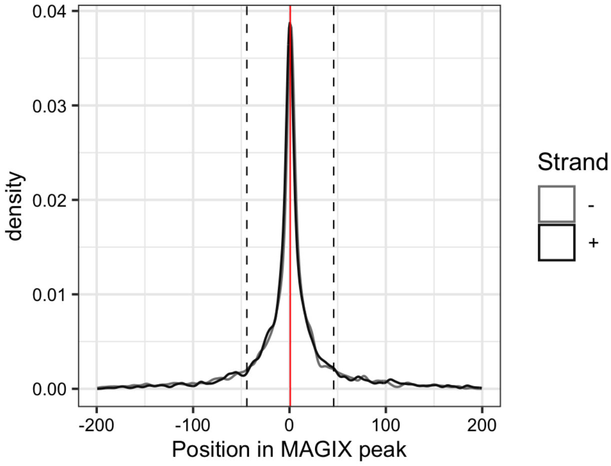


--------------------------------------------------------------------------------------------

AUROC of top GRECO-BIT/MEX PWM in sliding windows (500 peaks, 50 peak step) along the MAGIX peaks (up to the top 75,000) ranked by the MAGIX enrichment coefficient and fraction of overlapping ChIP peaks in the same window

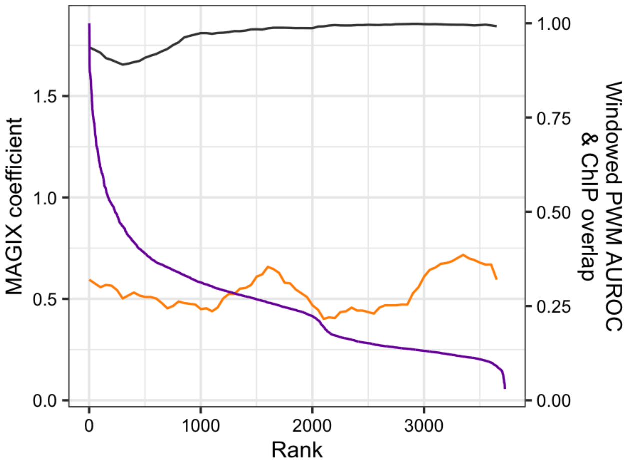

### FBXL19

Top GRECO-BIT/MEX PWM: PWM000507

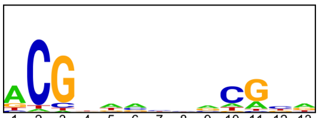


-----------------------------------------------------------------------------------------------

Location of top GRECO-BIT/MEX PWM in MAGIX peaks (up to the top 10,000)

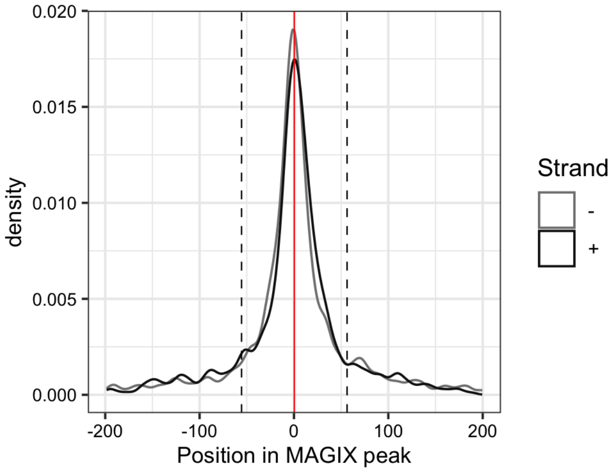


-----------------------------------------------------------------------------------------------

AUROC of top GRECO-BIT/MEX PWM in sliding windows (500 peaks, 50 peak step) along the MAGIX peaks (up to the top 75,000) ranked by the MAGIX enrichment coefficient and fraction of overlapping ChIP peaks in the same window

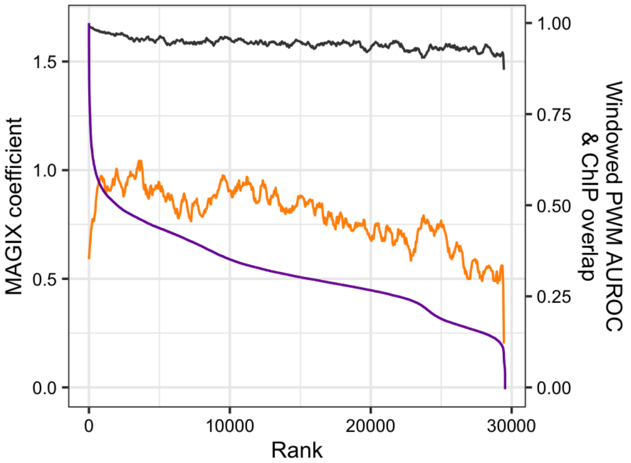

### FIZ1

Top GRECO-BIT/MEX PWM: PWM000541

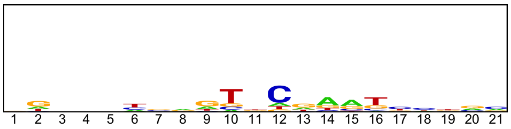


------------------------------------------------------------------------------------------

Location of top GRECO-BIT/MEX PWM in MAGIX peaks (up to the top 10,000)

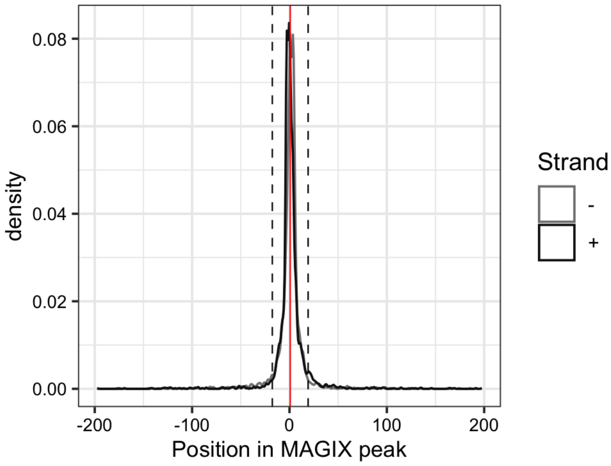


------------------------------------------------------------------------------------------

AUROC of top GRECO-BIT/MEX PWM in sliding windows (500 peaks, 50 peak step) along the MAGIX peaks (up to the top 75,000) ranked by the MAGIX enrichment coefficient and fraction of overlapping ChIP peaks in the same window

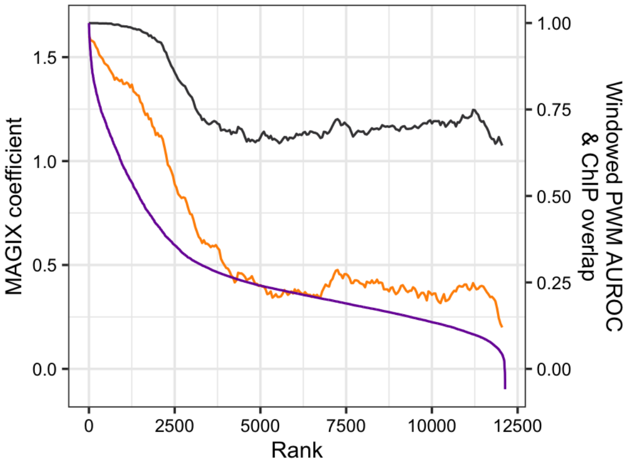

### FLI1

Top GRECO-BIT/MEX PWM: PWM000587

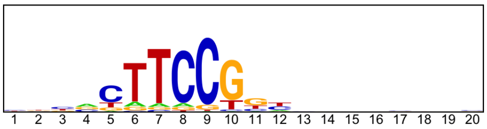


---

Location of top GRECO-BIT/MEX PWM in MAGIX peaks (up to the top 10,000)

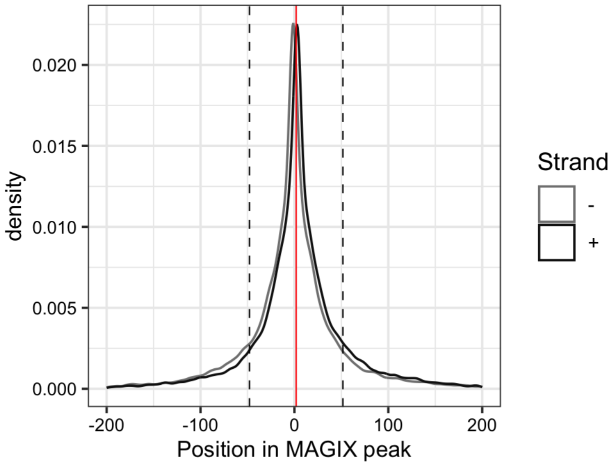


---

AUROC of top GRECO-BIT/MEX PWM in sliding windows (500 peaks, 50 peak step) along the MAGIX peaks (up to the top 75,000) ranked by the MAGIX enrichment coefficient and fraction of overlapping ChIP peaks in the same window

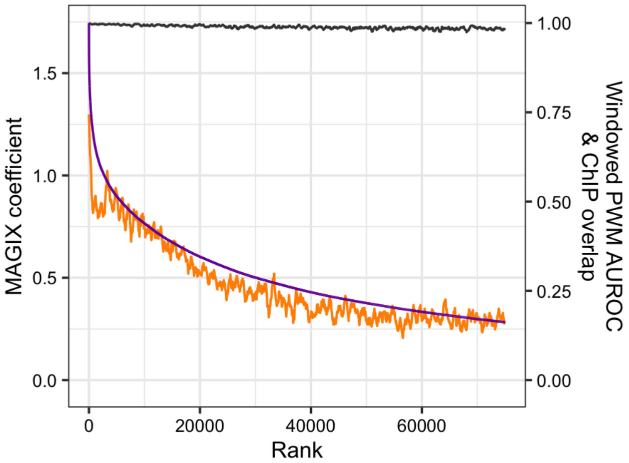

### FLYWCH1

Top GRECO-BIT/MEX PWM: PWM000607

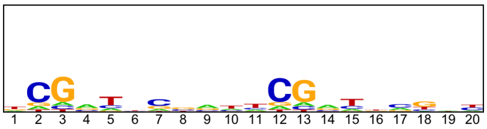


--------------------------------------------------------------------------------------------

Location of top GRECO-BIT/MEX PWM in MAGIX peaks (up to the top 10,000)

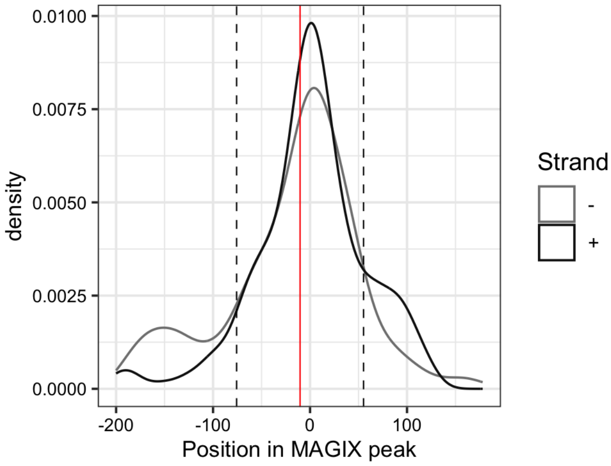


--------------------------------------------------------------------------------------------

AUROC of top GRECO-BIT/MEX PWM in sliding windows (500 peaks, 50 peak step) along the MAGIX peaks (up to the top 75,000) ranked by the MAGIX enrichment coefficient and fraction of overlapping ChIP peaks in the same window

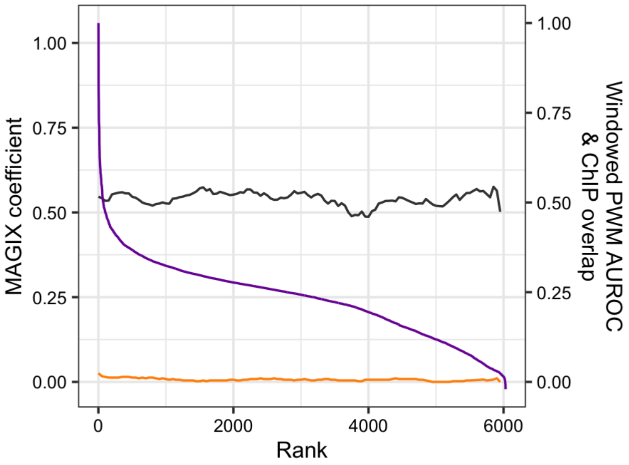

### FOSL2

Top GRECO-BIT/MEX PWM: PWM000649

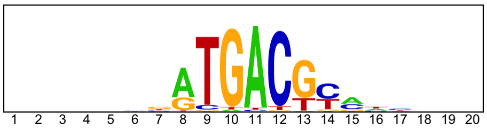


---

Location of top GRECO-BIT/MEX PWM in MAGIX peaks (up to the top 10,000)

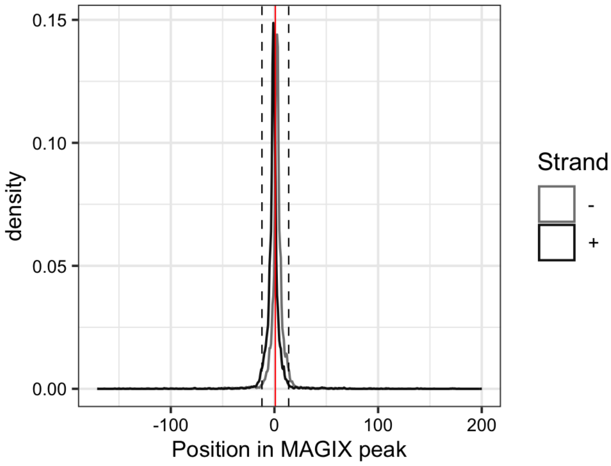


---

AUROC of top GRECO-BIT/MEX PWM in sliding windows (500 peaks, 50 peak step) along the MAGIX peaks (up to the top 75,000) ranked by the MAGIX enrichment coefficient and fraction of overlapping ChIP peaks in the same window

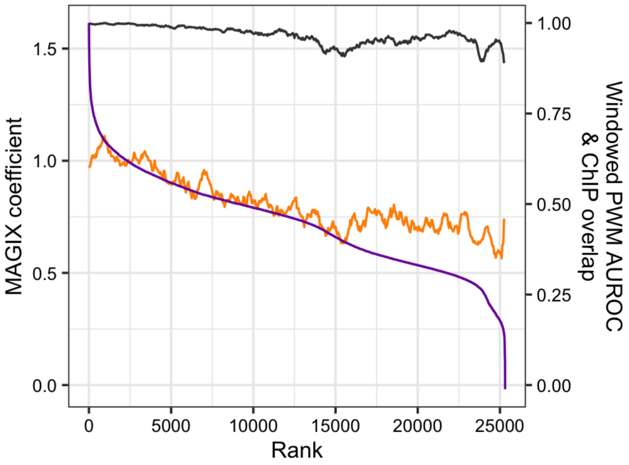

### GABPA

Top GRECO-BIT/MEX PWM: PWM000687

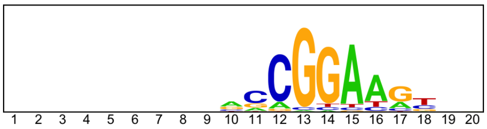


---

Location of top GRECO-BIT/MEX PWM in MAGIX peaks (up to the top 10,000)

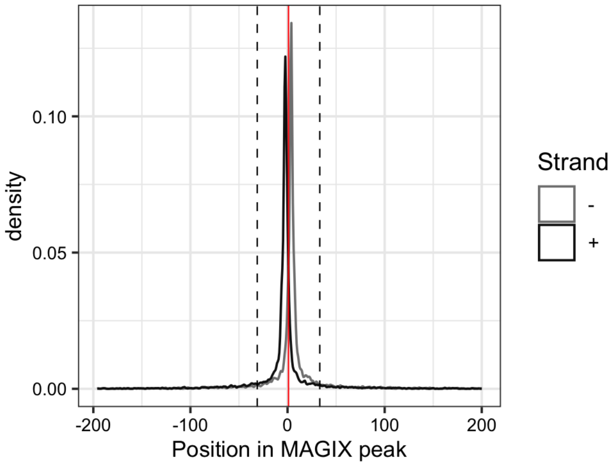


---

AUROC of top GRECO-BIT/MEX PWM in sliding windows (500 peaks, 50 peak step) along the MAGIX peaks (up to the top 75,000) ranked by the MAGIX enrichment coefficient and fraction of overlapping ChIP peaks in the same window

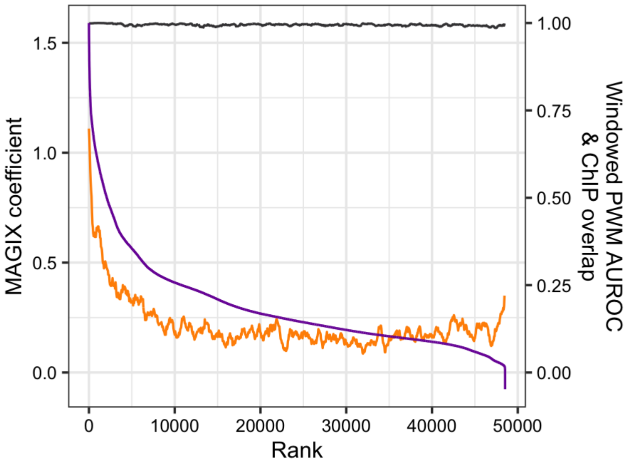

### GATAD2A

Top GRECO-BIT/MEX PWM: PWM000707

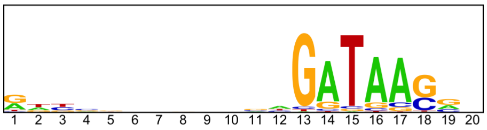


---

Location of top GRECO-BIT/MEX PWM in MAGIX peaks (up to the top 10,000)

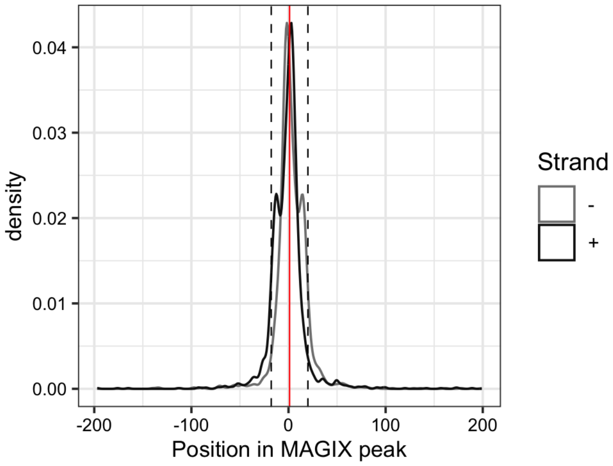

### GCM1

Top GRECO-BIT/MEX PWM: PWM000755

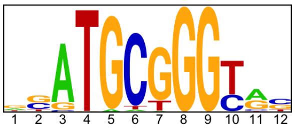


-----------------------------------------------------------------------------------------------

Location of top GRECO-BIT/MEX PWM in MAGIX peaks (up to the top 10,000)

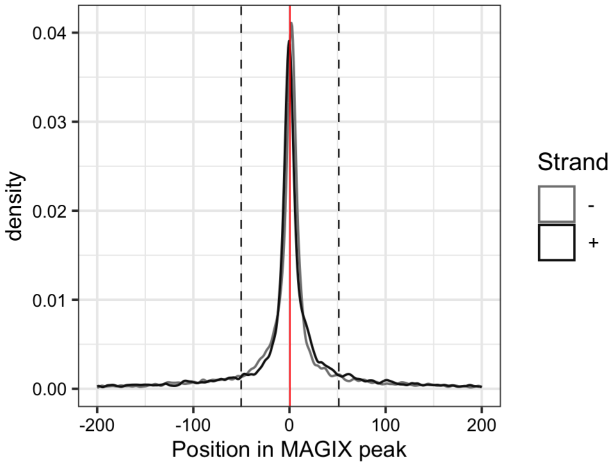


-----------------------------------------------------------------------------------------------

AUROC of top GRECO-BIT/MEX PWM in sliding windows (500 peaks, 50 peak step) along the MAGIX peaks (up to the top 75,000) ranked by the MAGIX enrichment coefficient and fraction of overlapping ChIP peaks in the same window

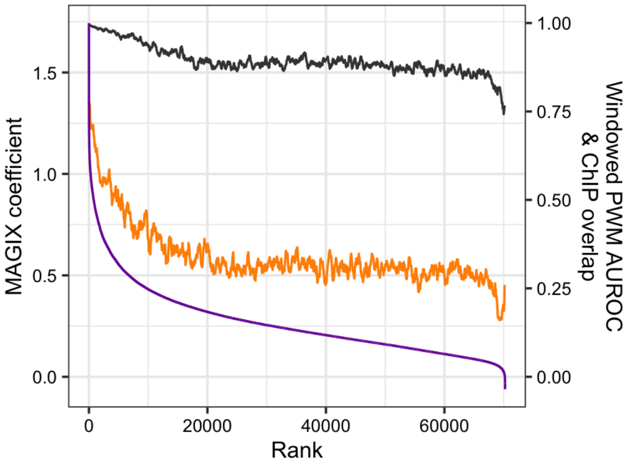

### GLI4

Top GRECO-BIT/MEX PWM: PWM000773

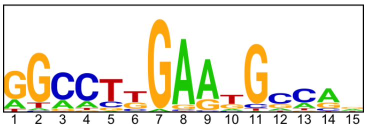


---

Location of top GRECO-BIT/MEX PWM in MAGIX peaks (up to the top 10,000)

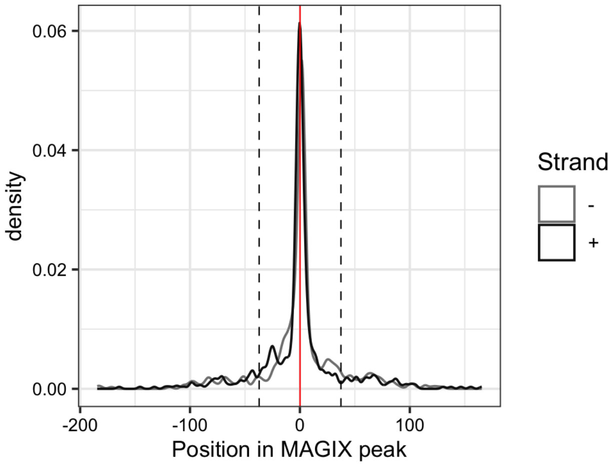


---

AUROC of top GRECO-BIT/MEX PWM in sliding windows (500 peaks, 50 peak step) along the MAGIX peaks (up to the top 75,000) ranked by the MAGIX enrichment coefficient and fraction of overlapping ChIP peaks in the same window

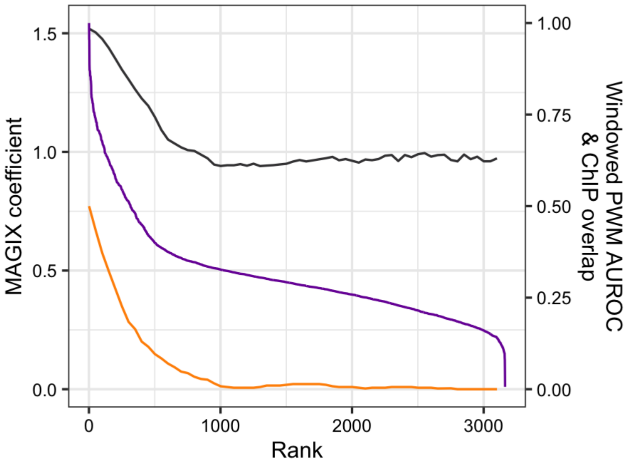

### JRK

Top GRECO-BIT/MEX PWM: PWM000822

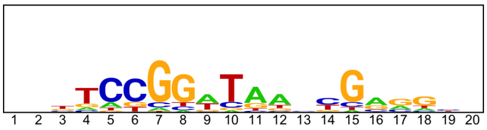


----------------------------------------------------------------------------------------------------

Location of top GRECO-BIT/MEX PWM in MAGIX peaks (up to the top 10,000)

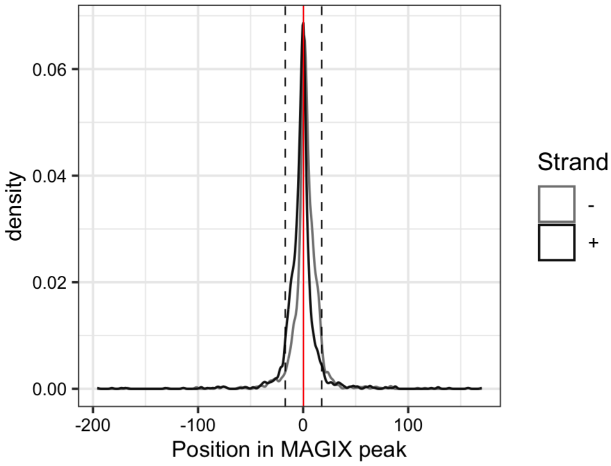


----------------------------------------------------------------------------------------------------

AUROC of top GRECO-BIT/MEX PWM in sliding windows (500 peaks, 50 peak step) along the MAGIX peaks (up to the top 75,000) ranked by the MAGIX enrichment coefficient and fraction of overlapping ChIP peaks in the same window

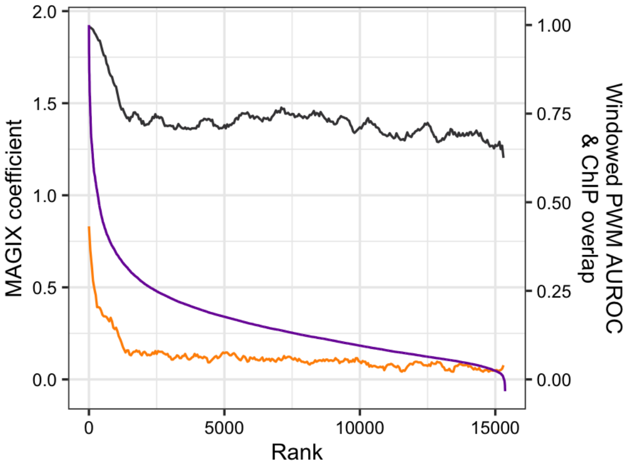

### KDM2A

Top GRECO-BIT/MEX PWM: PWM000893

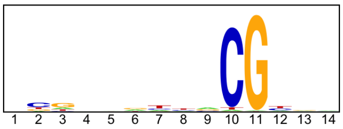


---

Location of top GRECO-BIT/MEX PWM in MAGIX peaks (up to the top 10,000)

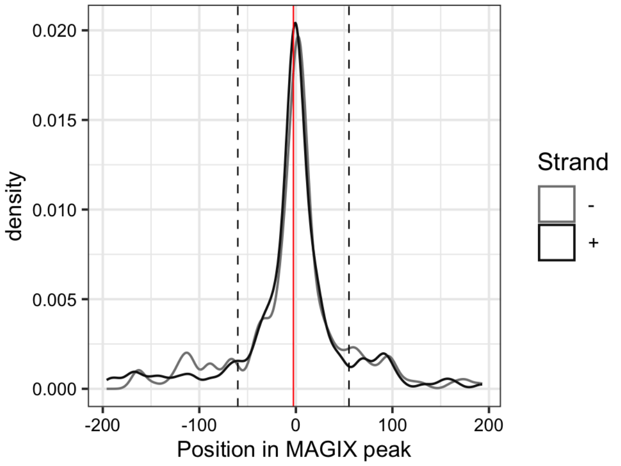


---

AUROC of top GRECO-BIT/MEX PWM in sliding windows (500 peaks, 50 peak step) along the MAGIX peaks (up to the top 75,000) ranked by the MAGIX enrichment coefficient and fraction of overlapping ChIP peaks in the same window

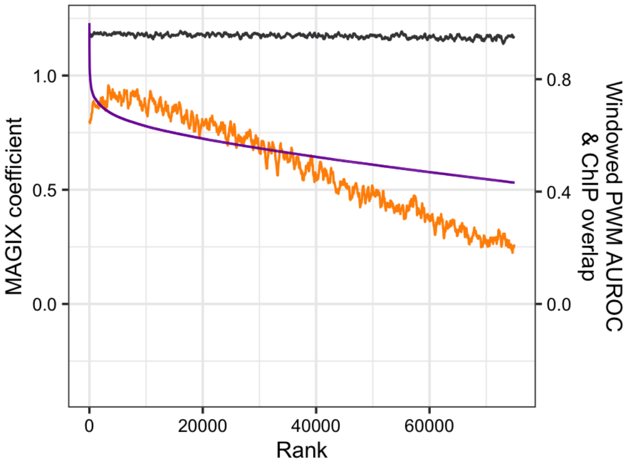

### LEF1

Top GRECO-BIT/MEX PWM: PWM000926

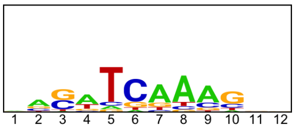


---

Location of top GRECO-BIT/MEX PWM in MAGIX peaks (up to the top 10,000)

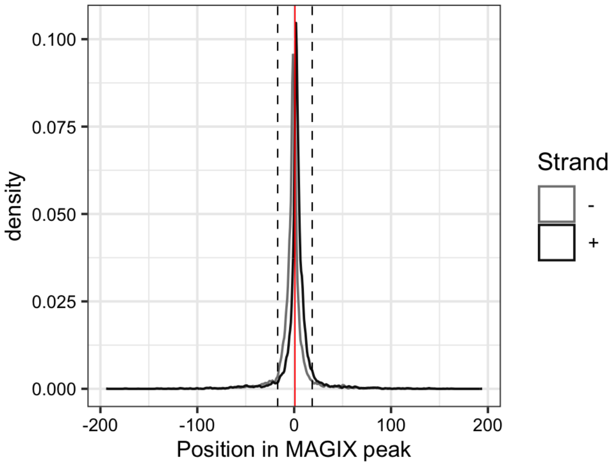


---

AUROC of top GRECO-BIT/MEX PWM in sliding windows (500 peaks, 50 peak step) along the MAGIX peaks (up to the top 75,000) ranked by the MAGIX enrichment coefficient and fraction of overlapping ChIP peaks in the same window

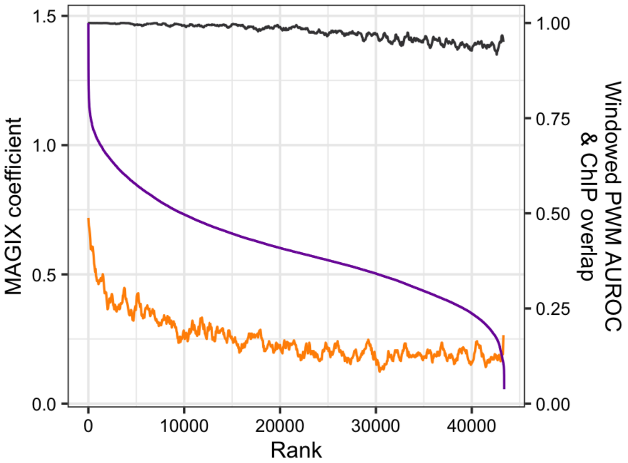

### LEUTX

Top GRECO-BIT/MEX PWM: PWM000952

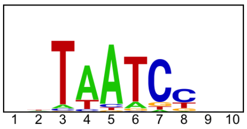


---

Location of top GRECO-BIT/MEX PWM in MAGIX peaks (up to the top 10,000)

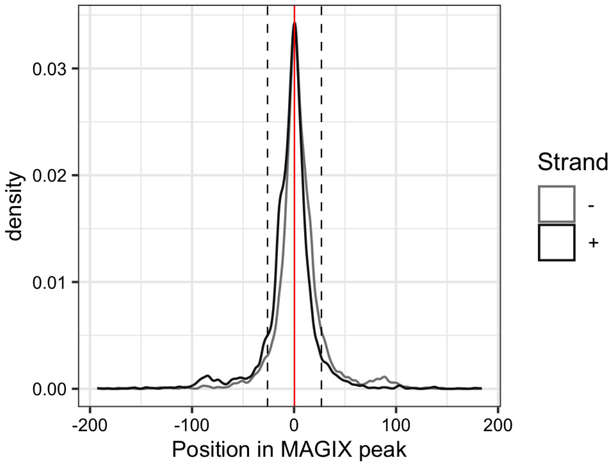


---

AUROC of top GRECO-BIT/MEX PWM in sliding windows (500 peaks, 50 peak step) along the MAGIX peaks (up to the top 75,000) ranked by the MAGIX enrichment coefficient and fraction of overlapping ChIP peaks in the same window

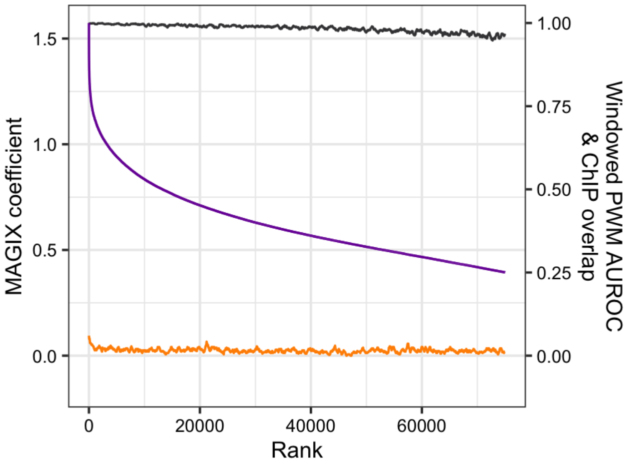

### MAX

Top GRECO-BIT/MEX PWM: PWM001030

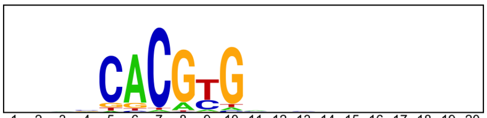


----------------------------------------------------------------------------------------------------

Location of top GRECO-BIT/MEX PWM in MAGIX peaks (up to the top 10,000)

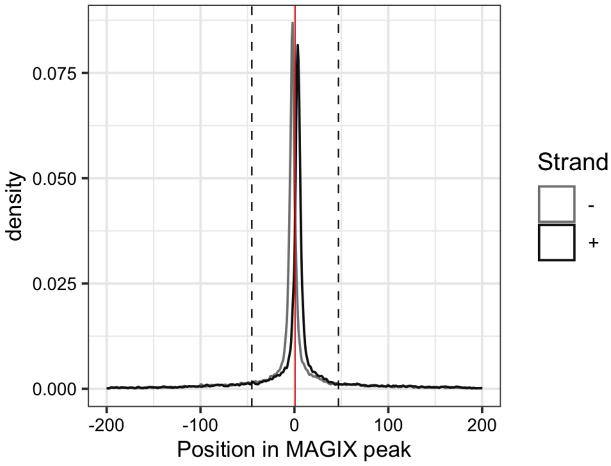


----------------------------------------------------------------------------------------------------

AUROC of top GRECO-BIT/MEX PWM in sliding windows (500 peaks, 50 peak step) along the MAGIX peaks (up to the top 75,000) ranked by the MAGIX enrichment coefficient and fraction of overlapping ChIP peaks in the same window

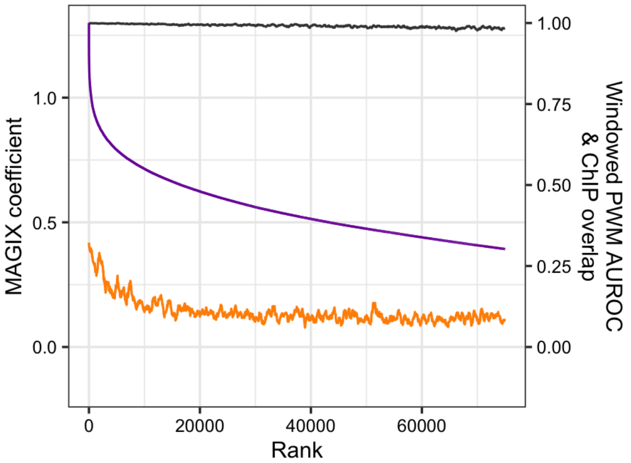

### MGA

Top GRECO-BIT/MEX PWM: PWM001108

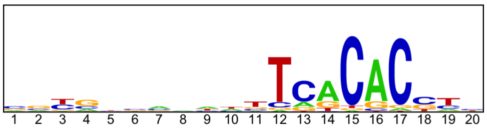


---

Location of top GRECO-BIT/MEX PWM in MAGIX peaks (up to the top 10,000)

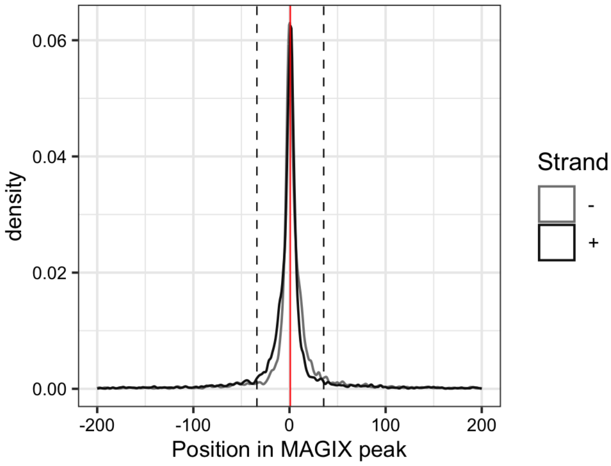


---

AUROC of top GRECO-BIT/MEX PWM in sliding windows (500 peaks, 50 peak step) along the MAGIX peaks (up to the top 75,000) ranked by the MAGIX enrichment coefficient and fraction of overlapping ChIP peaks in the same window

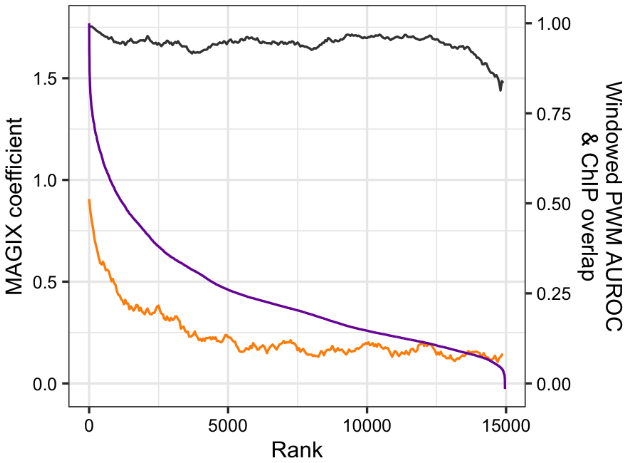

### MKX

Top GRECO-BIT/MEX PWM: PWM001148

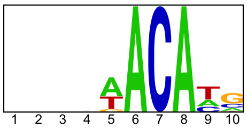


------------------------------------------------------------------------------------------

Location of top GRECO-BIT/MEX PWM in MAGIX peaks (up to the top 10,000)

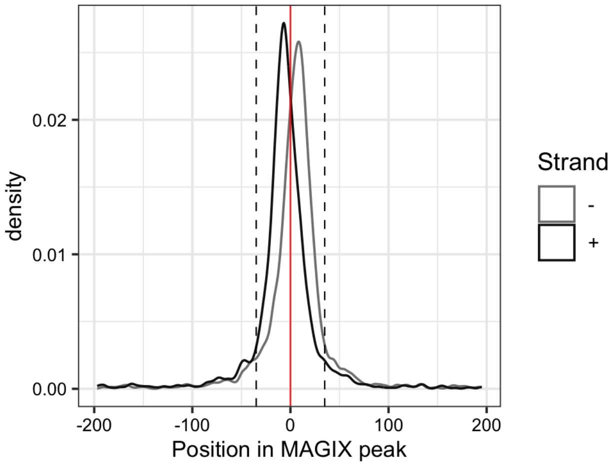


------------------------------------------------------------------------------------------

AUROC of top GRECO-BIT/MEX PWM in sliding windows (500 peaks, 50 peak step) along the MAGIX peaks (up to the top 75,000) ranked by the MAGIX enrichment coefficient and fraction of overlapping ChIP peaks in the same window

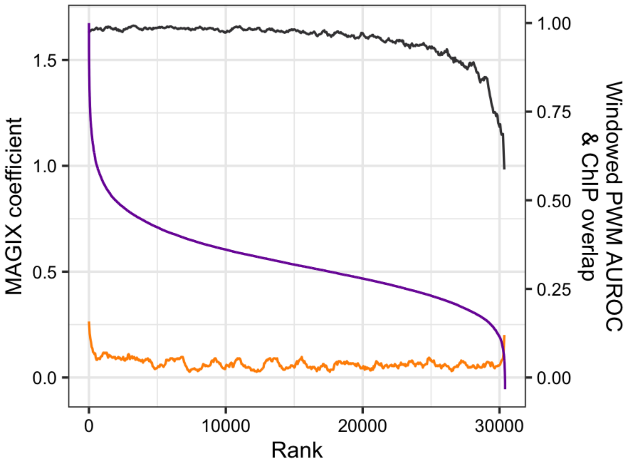

### MSANTD1

Top GRECO-BIT/MEX PWM: PWM001171

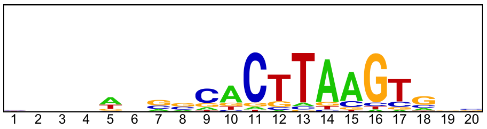

---

Location of top GRECO-BIT/MEX PWM in MAGIX peaks (up to the top 10,000)

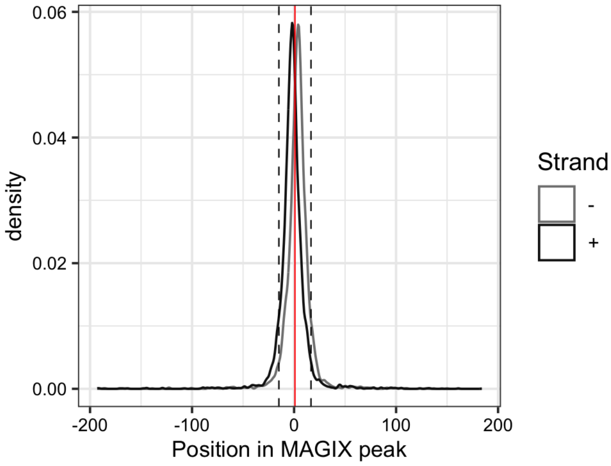


---

AUROC of top GRECO-BIT/MEX PWM in sliding windows (500 peaks, 50 peak step) along the MAGIX peaks (up to the top 75,000) ranked by the MAGIX enrichment coefficient and fraction of overlapping ChIP peaks in the same window

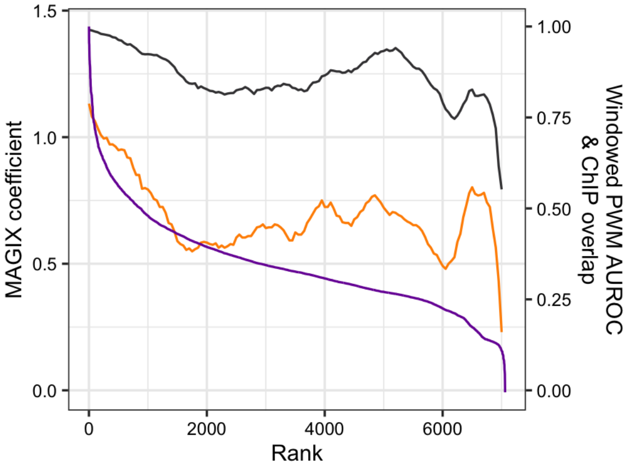

### MSANTD4

Top GRECO-BIT/MEX PWM: PWM001209

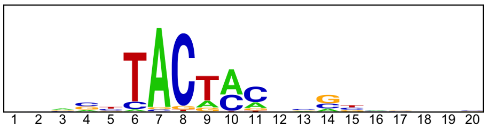


------------------------------------------------------------------------------------------------

Location of top GRECO-BIT/MEX PWM in MAGIX peaks (up to the top 10,000)

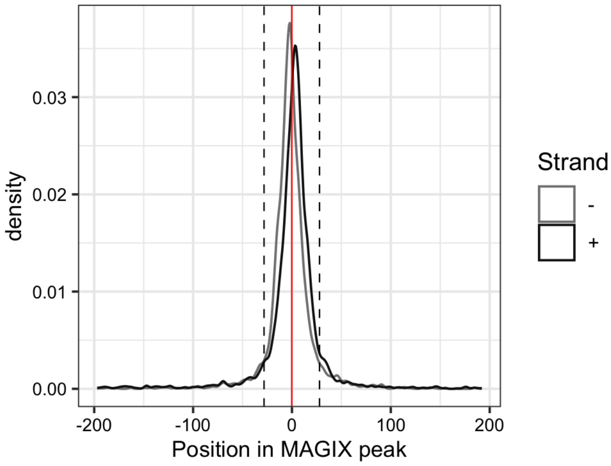

### MYF6

Top GRECO-BIT/MEX PWM: PWM001265

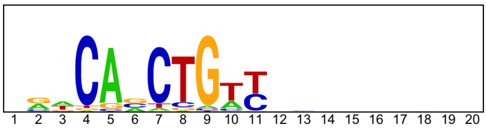


---

Location of top GRECO-BIT/MEX PWM in MAGIX peaks (up to the top 10,000)

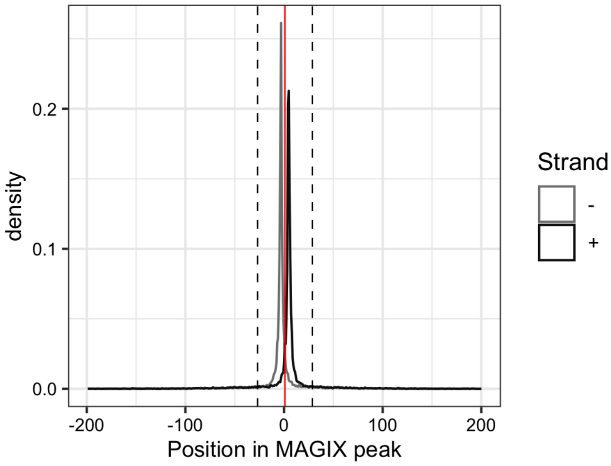


---

AUROC of top GRECO-BIT/MEX PWM in sliding windows (500 peaks, 50 peak step) along the MAGIX peaks (up to the top 75,000) ranked by the MAGIX enrichment coefficient and fraction of overlapping ChIP peaks in the same window

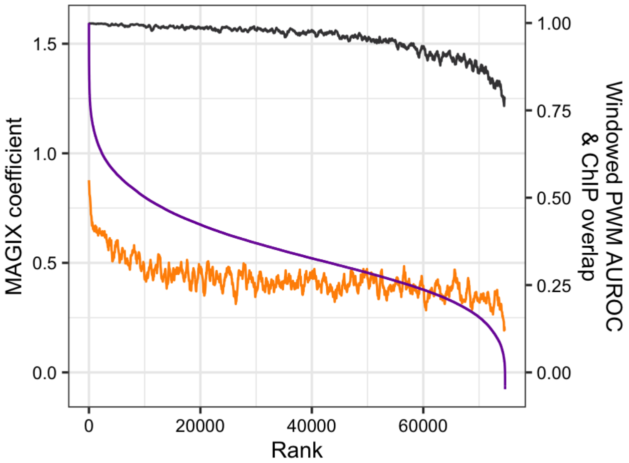

### MYPOP

Top GRECO-BIT/MEX PWM: PWM001292

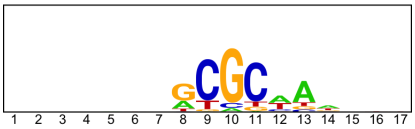


------------------------------------------------------------------------------------------

Location of top GRECO-BIT/MEX PWM in MAGIX peaks (up to the top 10,000)

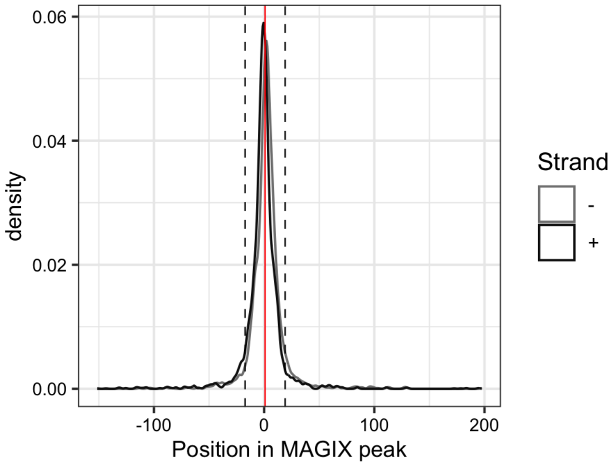


------------------------------------------------------------------------------------------

AUROC of top GRECO-BIT/MEX PWM in sliding windows (500 peaks, 50 peak step) along the MAGIX peaks (up to the top 75,000) ranked by the MAGIX enrichment coefficient and fraction of overlapping ChIP peaks in the same window

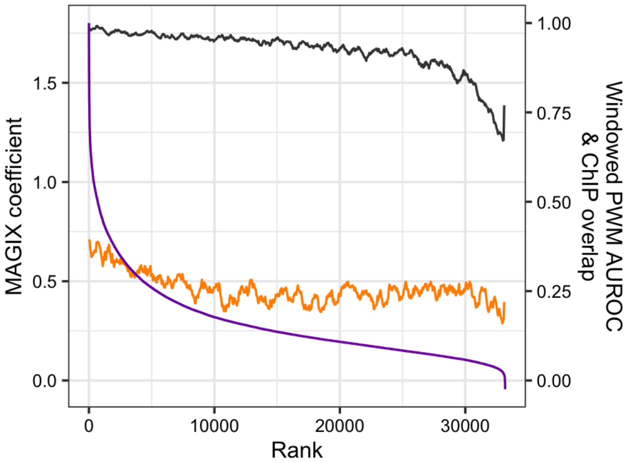

### MYRFL

Top GRECO-BIT/MEX PWM: PWM002431

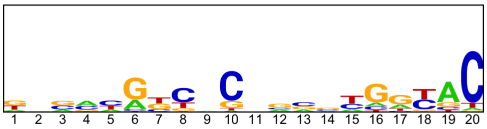


---

Location of top GRECO-BIT/MEX PWM in MAGIX peaks (up to the top 10,000)

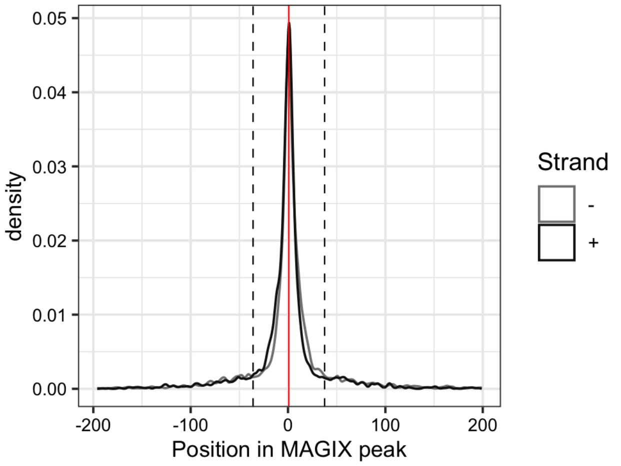


---

AUROC of top GRECO-BIT/MEX PWM in sliding windows (500 peaks, 50 peak step) along the MAGIX peaks (up to the top 75,000) ranked by the MAGIX enrichment coefficient and fraction of overlapping ChIP peaks in the same window

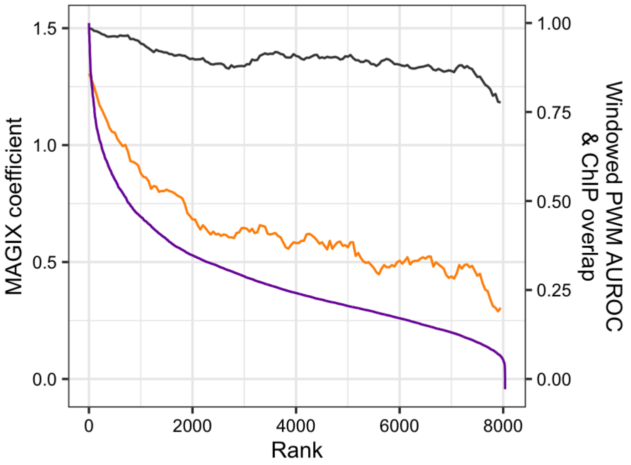

### MYT1.DBD1

Top GRECO-BIT/MEX PWM: PWM053556

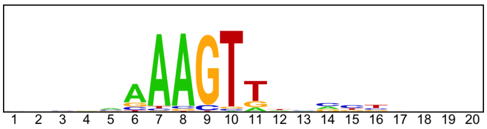


------------------------------------------------------------------------------------------------

Location of top GRECO-BIT/MEX PWM in MAGIX peaks (up to the top 10,000)

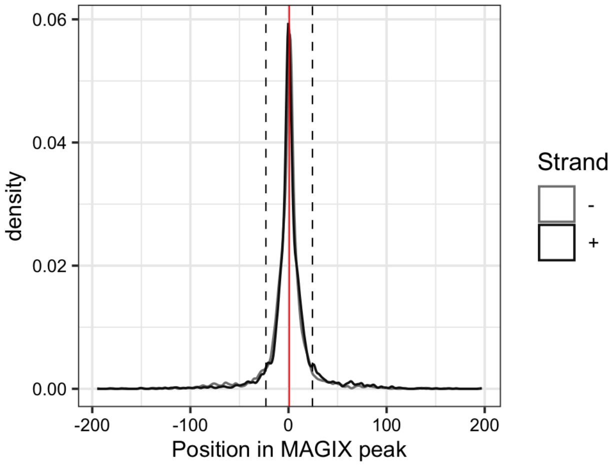

### MYT1.DBD2

Top GRECO-BIT/MEX PWM: PWM001364

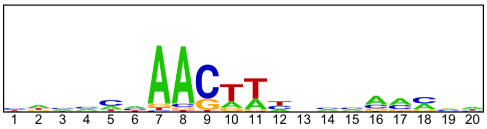


-----------------------------------------------------------------------------------------------

Location of top GRECO-BIT/MEX PWM in MAGIX peaks (up to the top 10,000)

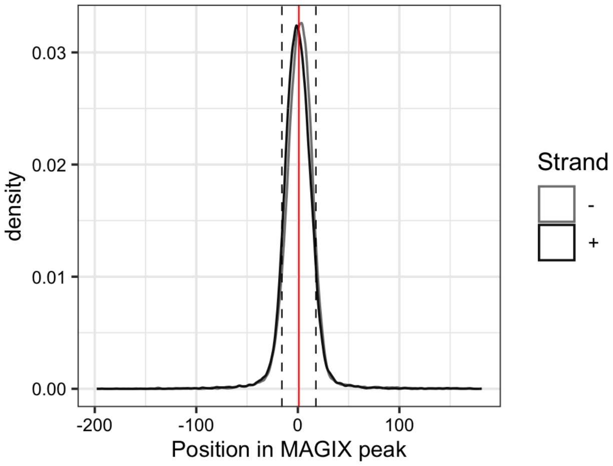

### MYT1.FL

Top GRECO-BIT/MEX PWM: PWM001366

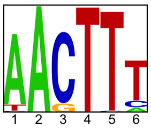


----------------------------------------------------------------------------------------------------

Location of top GRECO-BIT/MEX PWM in MAGIX peaks (up to the top 10,000)

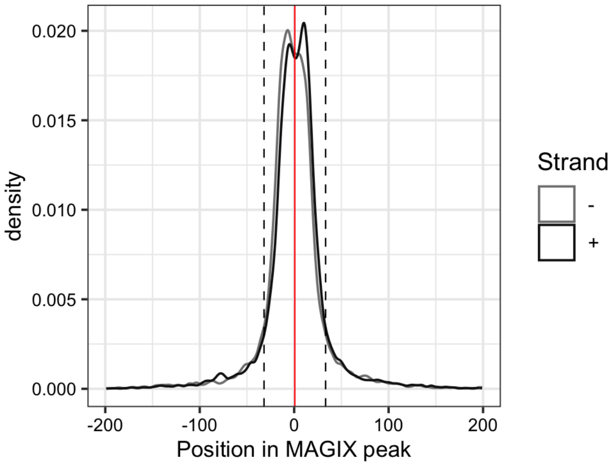

### NACC2

Top GRECO-BIT/MEX PWM: PWM001390

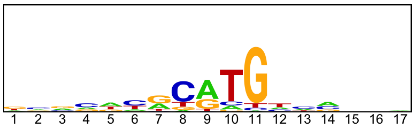


-----------------------------------------------------------------------------------------------

Location of top GRECO-BIT/MEX PWM in MAGIX peaks (up to the top 10,000)

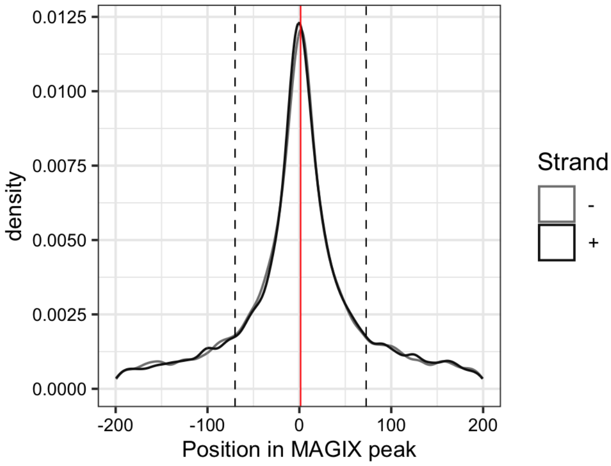


-----------------------------------------------------------------------------------------------

AUROC of top GRECO-BIT/MEX PWM in sliding windows (500 peaks, 50 peak step) along the MAGIX peaks (up to the top 75,000) ranked by the MAGIX enrichment coefficient and fraction of overlapping ChIP peaks in the same window

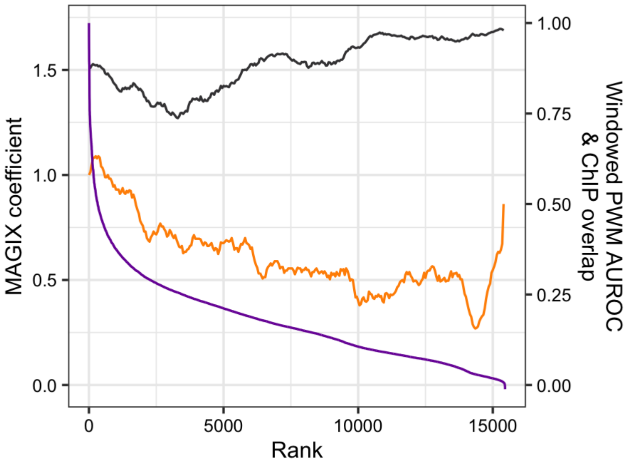

### NFKB1

Top GRECO-BIT/MEX PWM: PWM001436

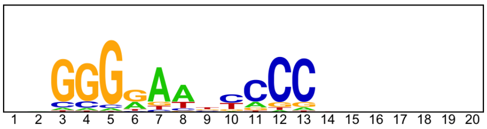


---

Location of top GRECO-BIT/MEX PWM in MAGIX peaks (up to the top 10,000)

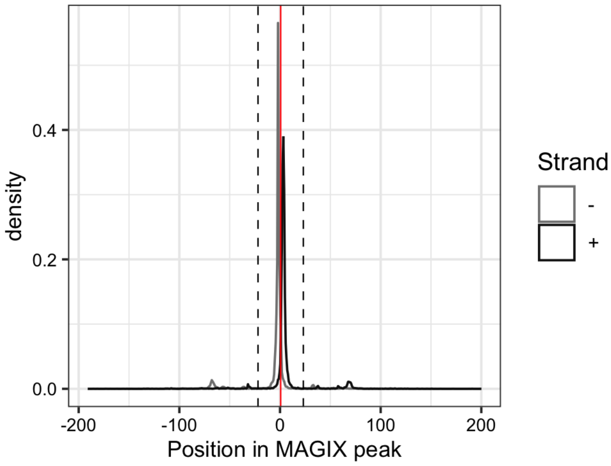


---

AUROC of top GRECO-BIT/MEX PWM in sliding windows (500 peaks, 50 peak step) along the MAGIX peaks (up to the top 75,000) ranked by the MAGIX enrichment coefficient and fraction of overlapping ChIP peaks in the same window

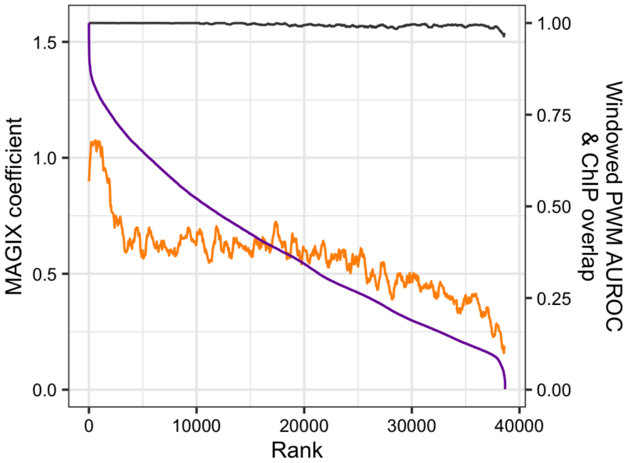

# NR1H4

Top GRECO-BIT/MEX PWM: PWM001473

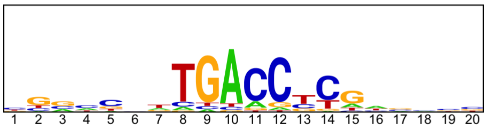


--------------------------------------------------------------------------------------------

Location of top GRECO-BIT/MEX PWM in MAGIX peaks (up to the top 10,000)

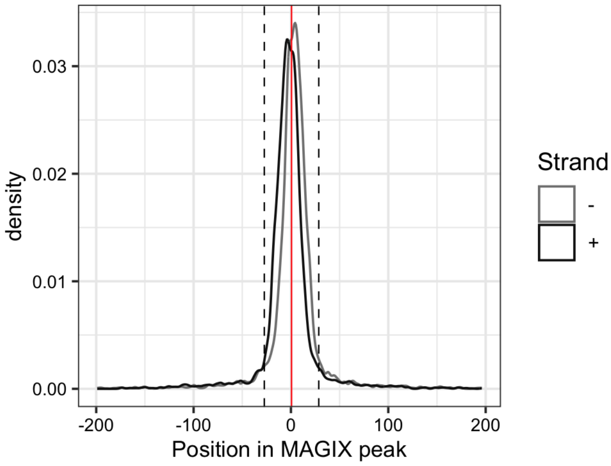


--------------------------------------------------------------------------------------------

AUROC of top GRECO-BIT/MEX PWM in sliding windows (500 peaks, 50 peak step) along the MAGIX peaks (up to the top 75,000) ranked by the MAGIX enrichment coefficient and fraction of overlapping ChIP peaks in the same window

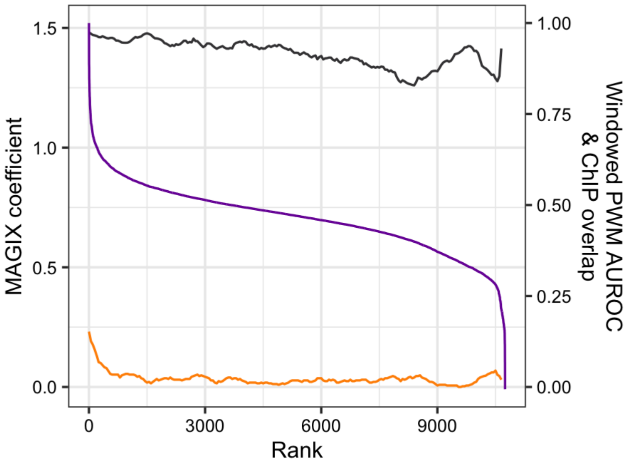

### PAX7

Top GRECO-BIT/MEX PWM: PWM001530

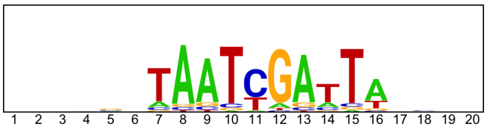


--------------------------------------------------------------------------------------------

Location of top GRECO-BIT/MEX PWM in MAGIX peaks (up to the top 10,000)

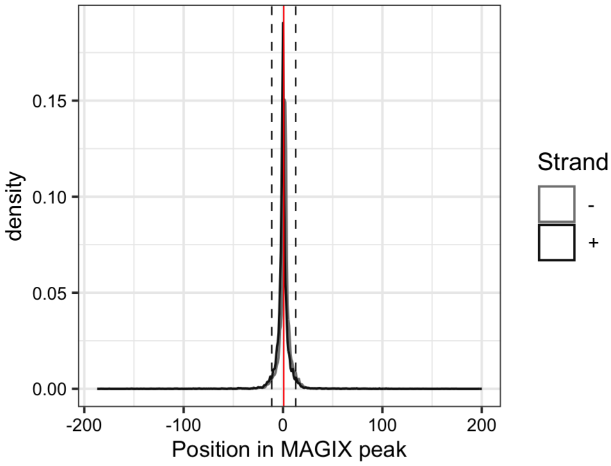


--------------------------------------------------------------------------------------------

AUROC of top GRECO-BIT/MEX PWM in sliding windows (500 peaks, 50 peak step) along the MAGIX peaks (up to the top 75,000) ranked by the MAGIX enrichment coefficient and fraction of overlapping ChIP peaks in the same window

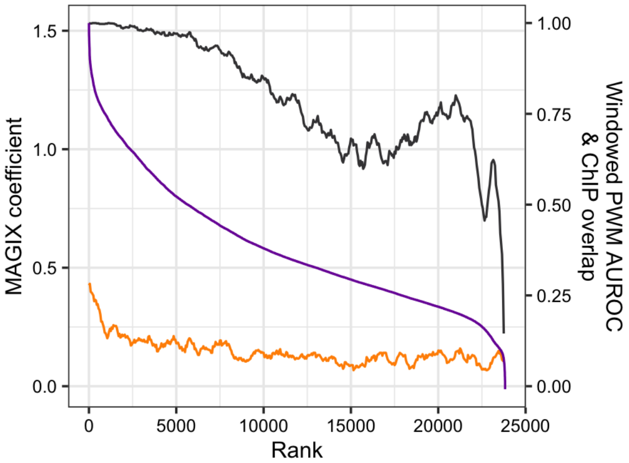

### POU5F1

Top GRECO-BIT/MEX PWM: PWM001566

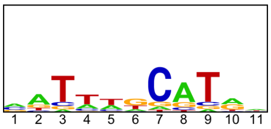

---

Location of top GRECO-BIT/MEX PWM in MAGIX peaks (up to the top 10,000)

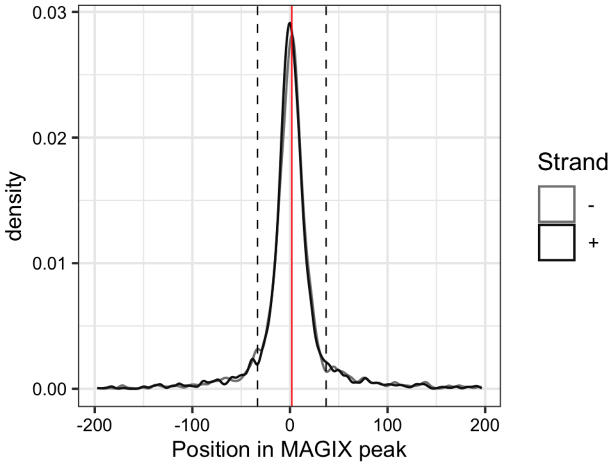

### PRDM10

Top GRECO-BIT/MEX PWM: PWM001598

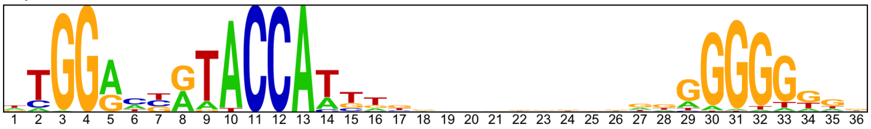


---

Location of top GRECO-BIT/MEX PWM in MAGIX peaks (up to the top 10,000)

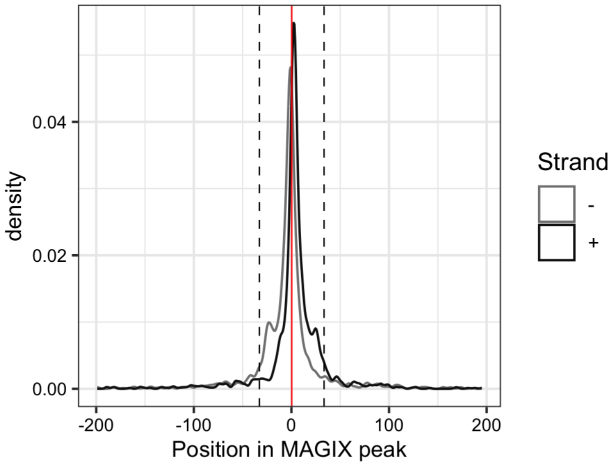


---

AUROC of top GRECO-BIT/MEX PWM in sliding windows (500 peaks, 50 peak step) along the MAGIX peaks (up to the top 75,000) ranked by the MAGIX enrichment coefficient and fraction of overlapping ChIP peaks in the same window

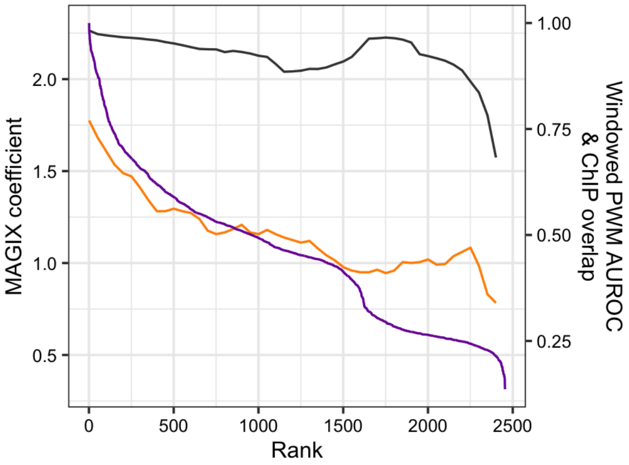

### PRDM13

Top GRECO-BIT/MEX PWM: PWM001641

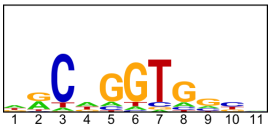


-----------------------------------------------------------------------------------------------

Location of top GRECO-BIT/MEX PWM in MAGIX peaks (up to the top 10,000)

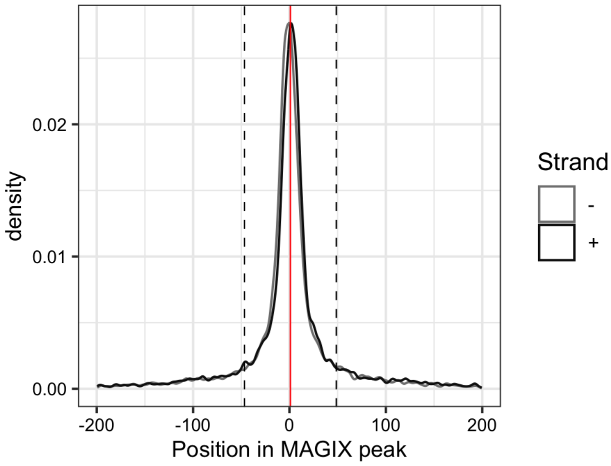


-----------------------------------------------------------------------------------------------

AUROC of top GRECO-BIT/MEX PWM in sliding windows (500 peaks, 50 peak step) along the MAGIX peaks (up to the top 75,000) ranked by the MAGIX enrichment coefficient and fraction of overlapping ChIP peaks in the same window

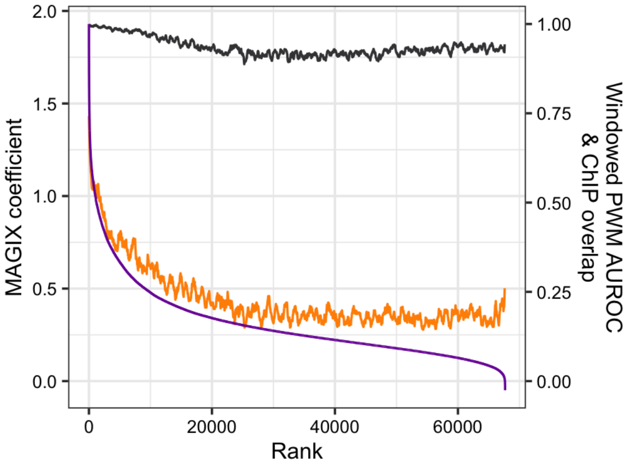

### PRDM5

Top GRECO-BIT/MEX PWM: PWM001672

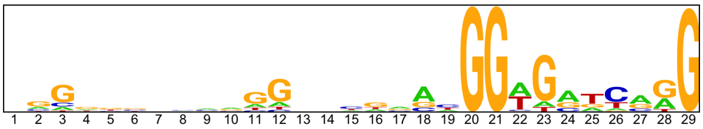


---

Location of top GRECO-BIT/MEX PWM in MAGIX peaks (up to the top 10,000)

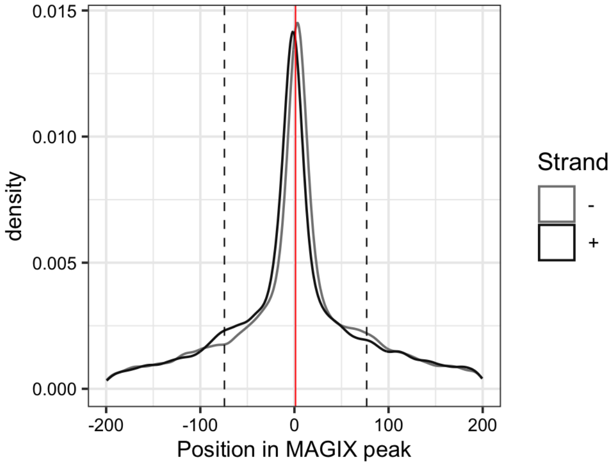


---

AUROC of top GRECO-BIT/MEX PWM in sliding windows (500 peaks, 50 peak step) along the MAGIX peaks (up to the top 75,000) ranked by the MAGIX enrichment coefficient and fraction of overlapping ChIP peaks in the same window

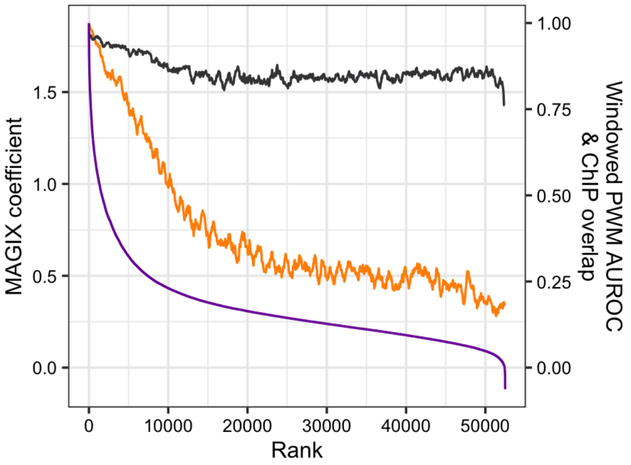

### RARA

Top GRECO-BIT/MEX PWM: PWM001684

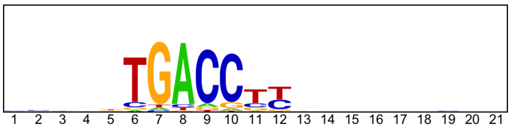


---

Location of top GRECO-BIT/MEX PWM in MAGIX peaks (up to the top 10,000)

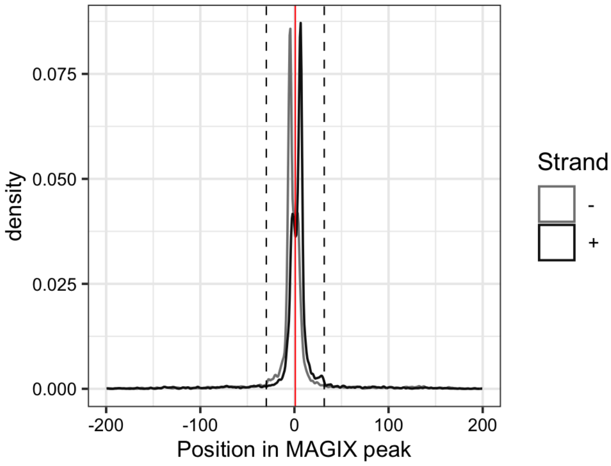

### RFX5

Top GRECO-BIT/MEX PWM: PWM001724

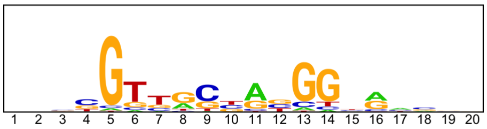


----------------------------------------------------------------------------------------------------

Location of top GRECO-BIT/MEX PWM in MAGIX peaks (up to the top 10,000)

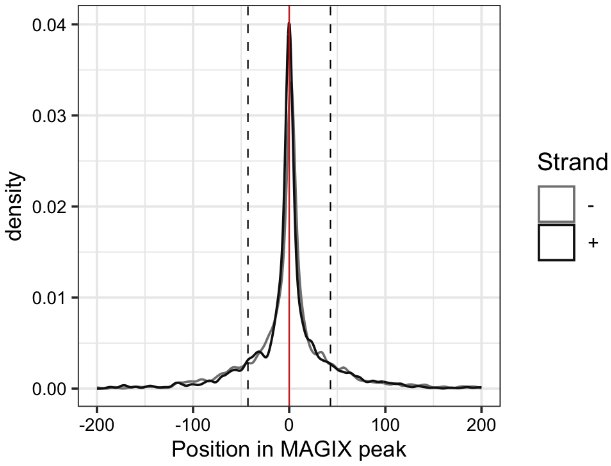


----------------------------------------------------------------------------------------------------

AUROC of top GRECO-BIT/MEX PWM in sliding windows (500 peaks, 50 peak step) along the MAGIX peaks (up to the top 75,000) ranked by the MAGIX enrichment coefficient and fraction of overlapping ChIP peaks in the same window

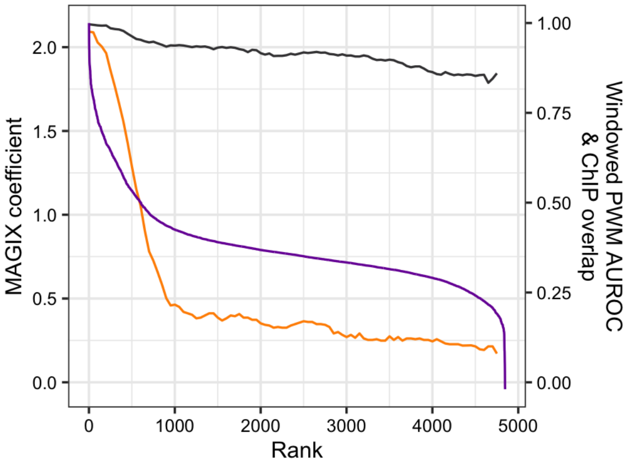

### RLF.DBD2

Top GRECO-BIT/MEX PWM: PWM001771

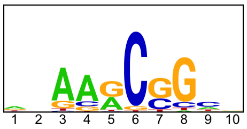


----------------------------------------------------------------------------------------------------

Location of top GRECO-BIT/MEX PWM in MAGIX peaks (up to the top 10,000)

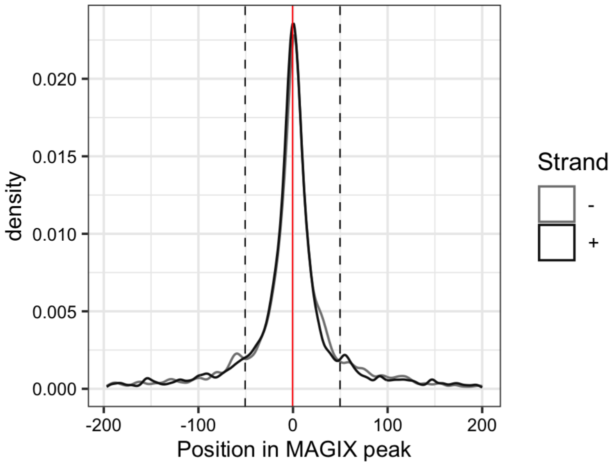

### RORB

Top GRECO-BIT/MEX PWM: PWM001817

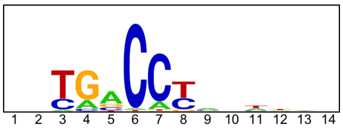


---

Location of top GRECO-BIT/MEX PWM in MAGIX peaks (up to the top 10,000)

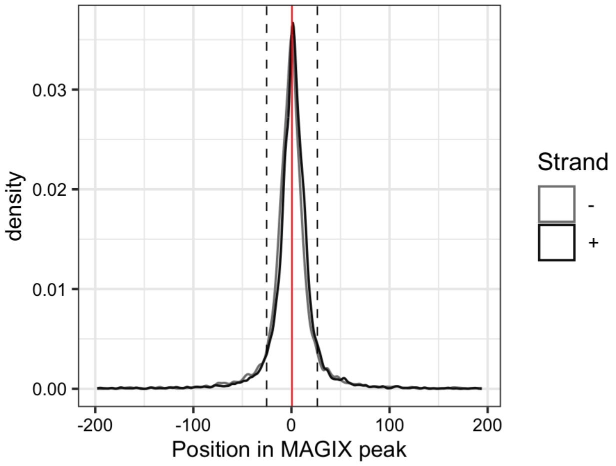


---

AUROC of top GRECO-BIT/MEX PWM in sliding windows (500 peaks, 50 peak step) along the MAGIX peaks (up to the top 75,000) ranked by the MAGIX enrichment coefficient and fraction of overlapping ChIP peaks in the same window

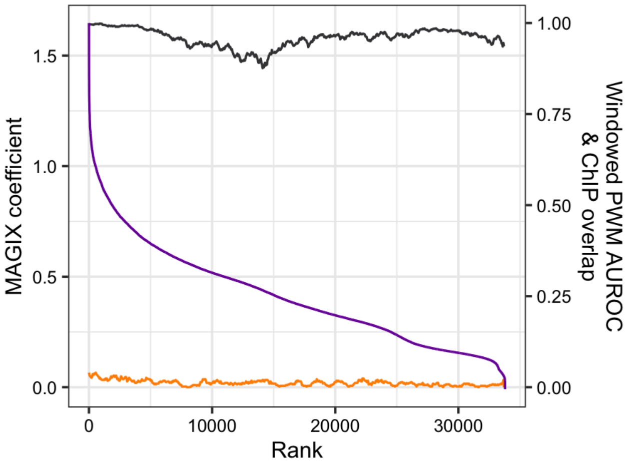

### RXRA

Top GRECO-BIT/MEX PWM: PWM001833

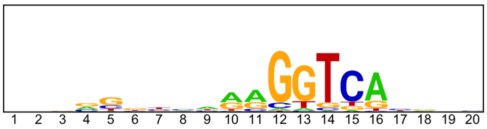


---

Location of top GRECO-BIT/MEX PWM in MAGIX peaks (up to the top 10,000)

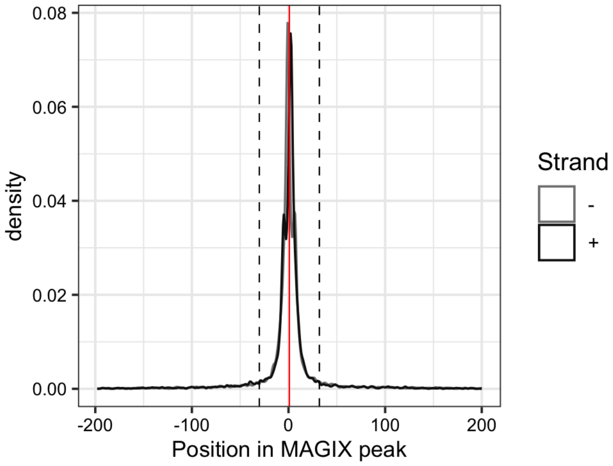

### SALL3.DBD1

Top GRECO-BIT/MEX PWM: PWM070151

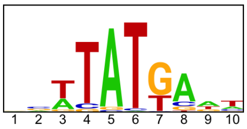


-----------------------------------------------------------------------------------------------

Location of top GRECO-BIT/MEX PWM in MAGIX peaks (up to the top 10,000)

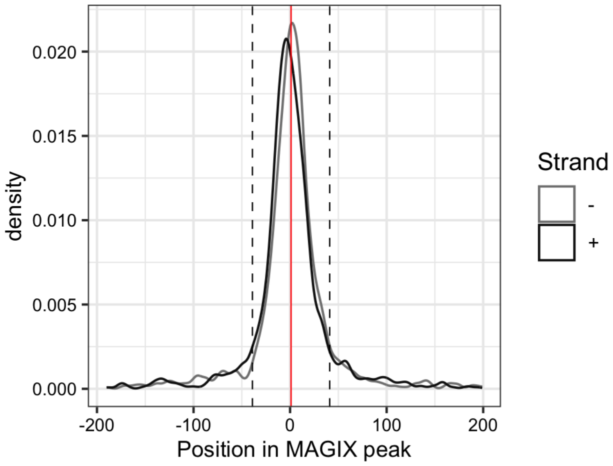

### SALL3.DBD2

Top GRECO-BIT/MEX PWM: PWM001862

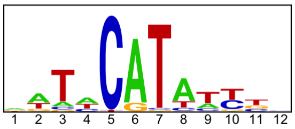


--------------------------------------------------------------------------------------------

Location of top GRECO-BIT/MEX PWM in MAGIX peaks (up to the top 10,000)

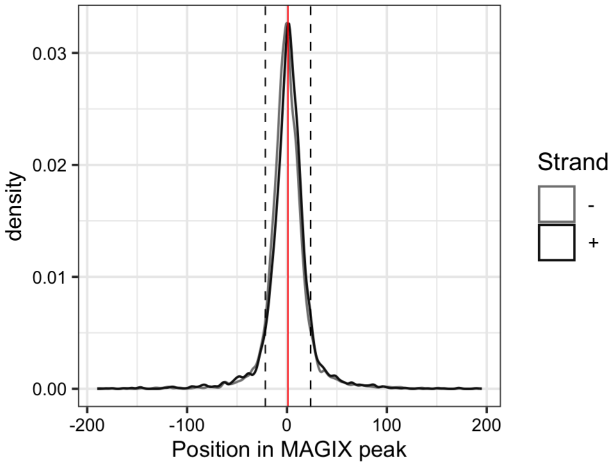

### SALL3.FL

Top GRECO-BIT/MEX PWM: PWM001879

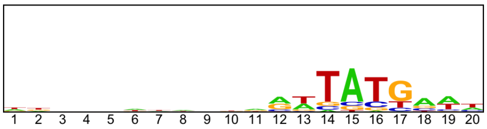


---------------------------------------------------------------------------------------------

Location of top GRECO-BIT/MEX PWM in MAGIX peaks (up to the top 10,000)

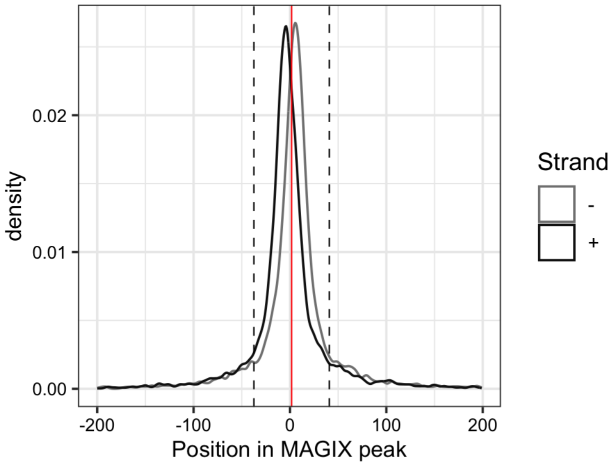


---------------------------------------------------------------------------------------------

AUROC of top GRECO-BIT/MEX PWM in sliding windows (500 peaks, 50 peak step) along the MAGIX peaks (up to the top 75,000) ranked by the MAGIX enrichment coefficient and fraction of overlapping ChIP peaks in the same window

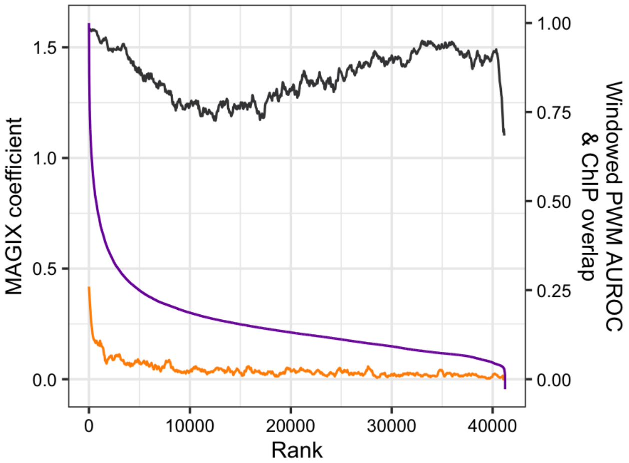

### SLC2A4RG

Top GRECO-BIT/MEX PWM: PWM001919

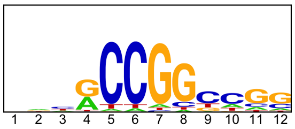


----------------------------------------------------------------------------------------------------

Location of top GRECO-BIT/MEX PWM in MAGIX peaks (up to the top 10,000)

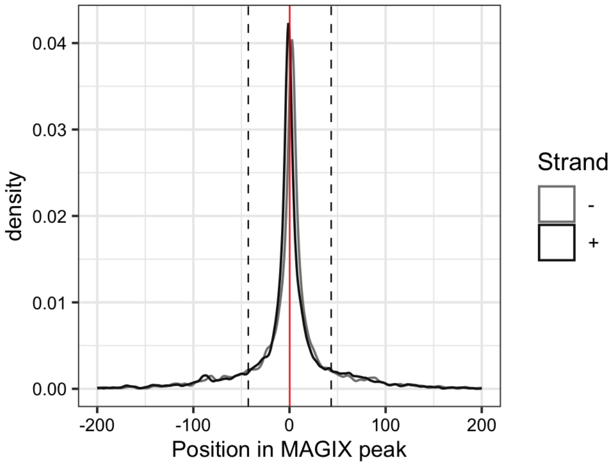


----------------------------------------------------------------------------------------------------

AUROC of top GRECO-BIT/MEX PWM in sliding windows (500 peaks, 50 peak step) along the MAGIX peaks (up to the top 75,000) ranked by the MAGIX enrichment coefficient and fraction of overlapping ChIP peaks in the same window

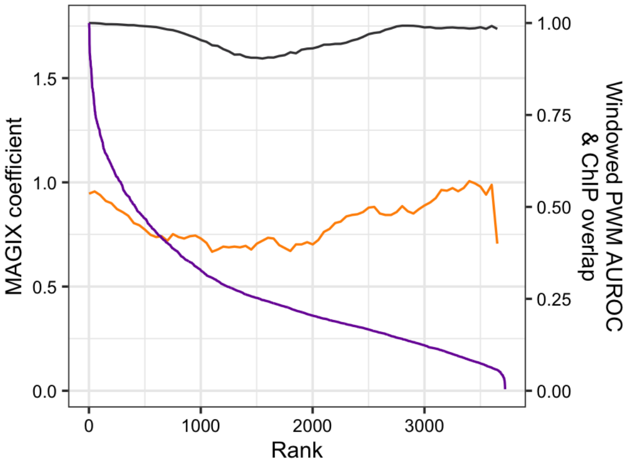

### SNAI1

Top GRECO-BIT/MEX PWM: PWM001937

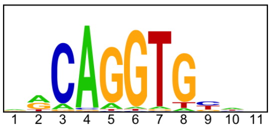


----------------------------------------------------------------------------------------------------

Location of top GRECO-BIT/MEX PWM in MAGIX peaks (up to the top 10,000)

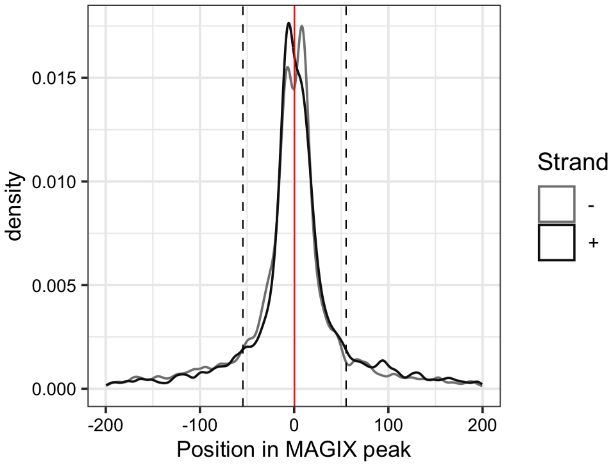

### SOX15

Top GRECO-BIT/MEX PWM: PWM001963

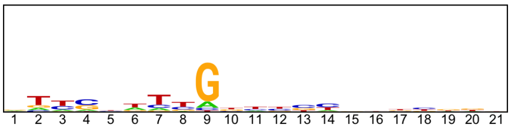


--------------------------------------------------------------------------------------------

Location of top GRECO-BIT/MEX PWM in MAGIX peaks (up to the top 10,000)

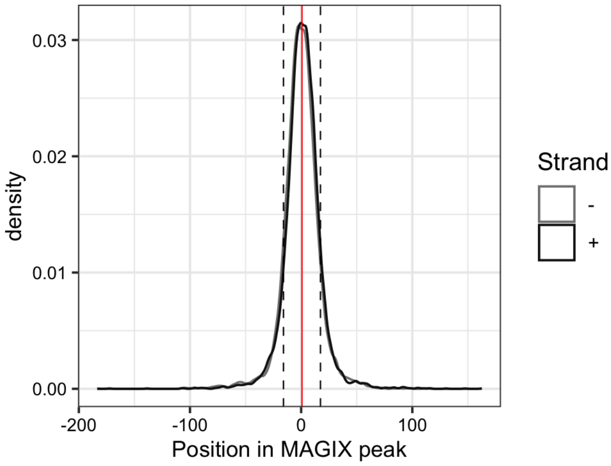


--------------------------------------------------------------------------------------------

AUROC of top GRECO-BIT/MEX PWM in sliding windows (500 peaks, 50 peak step) along the MAGIX peaks (up to the top 75,000) ranked by the MAGIX enrichment coefficient and fraction of overlapping ChIP peaks in the same window

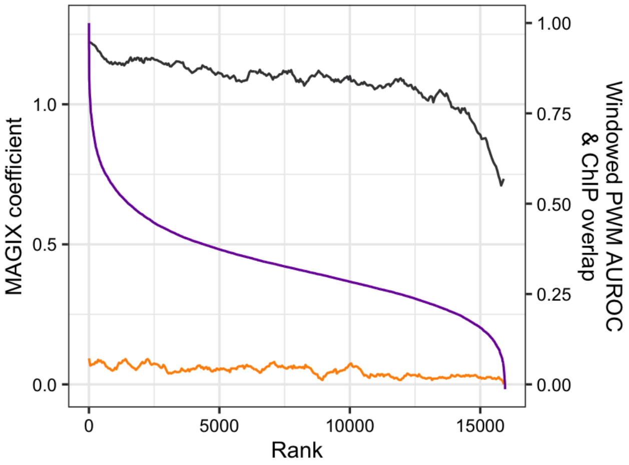

### SOX2

Top GRECO-BIT/MEX PWM: PWM002014

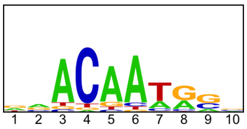


--------------------------------------------------------------------------------------------

Location of top GRECO-BIT/MEX PWM in MAGIX peaks (up to the top 10,000)

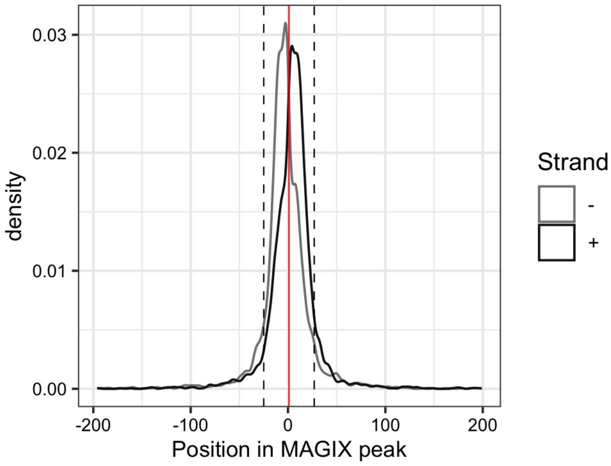


--------------------------------------------------------------------------------------------

AUROC of top GRECO-BIT/MEX PWM in sliding windows (500 peaks, 50 peak step) along the MAGIX peaks (up to the top 75,000) ranked by the MAGIX enrichment coefficient and fraction of overlapping ChIP peaks in the same window

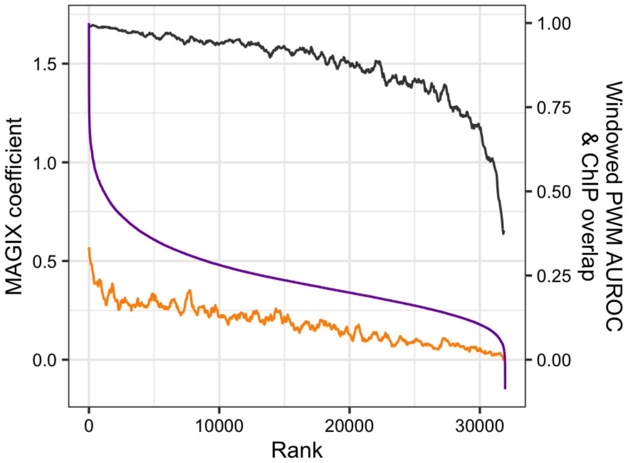

# SP100

Top GRECO-BIT/MEX PWM: PWM002035

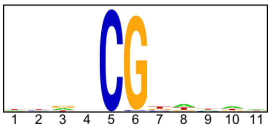


-----------------------------------------------------------------------------------------------

Location of top GRECO-BIT/MEX PWM in MAGIX peaks (up to the top 10,000)

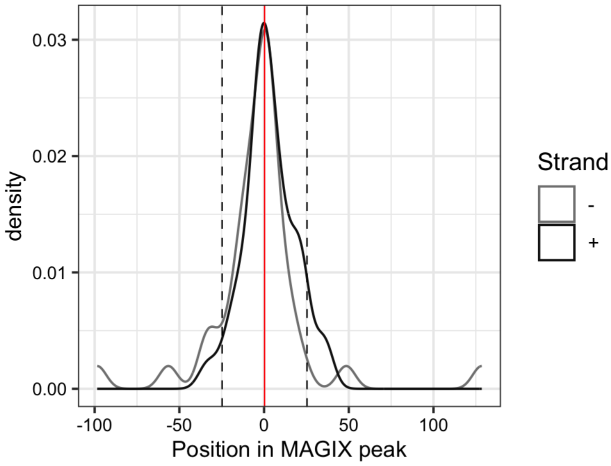


-----------------------------------------------------------------------------------------------

AUROC of top GRECO-BIT/MEX PWM in sliding windows (500 peaks, 50 peak step) along the MAGIX peaks (up to the top 75,000) ranked by the MAGIX enrichment coefficient and fraction of overlapping ChIP peaks in the same window

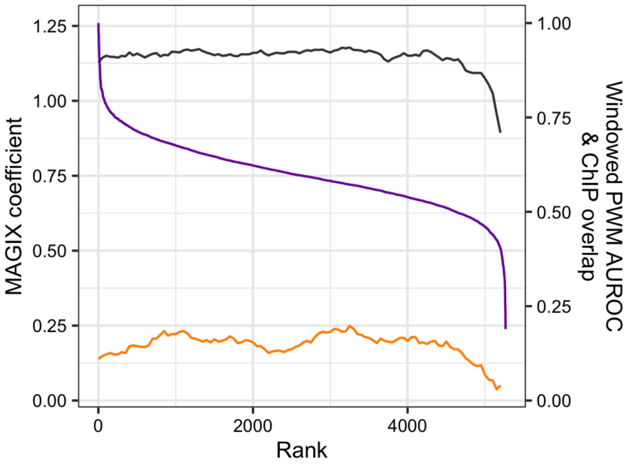

# SP140

Top GRECO-BIT/MEX PWM: PWM002066

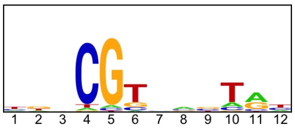


--------------------------------------------------------------------------------------------

Location of top GRECO-BIT/MEX PWM in MAGIX peaks (up to the top 10,000)

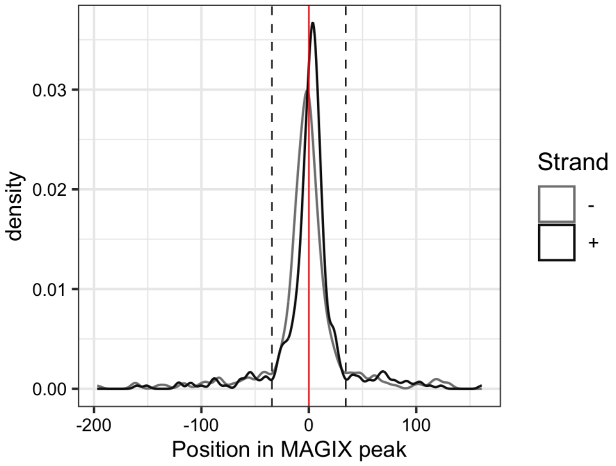


--------------------------------------------------------------------------------------------

AUROC of top GRECO-BIT/MEX PWM in sliding windows (500 peaks, 50 peak step) along the MAGIX peaks (up to the top 75,000) ranked by the MAGIX enrichment coefficient and fraction of overlapping ChIP peaks in the same window

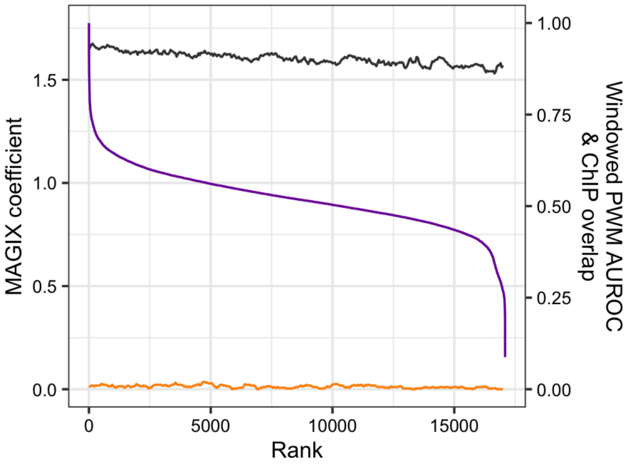

# SP140L

Top GRECO-BIT/MEX PWM: PWM002102

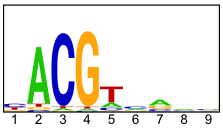


---

Location of top GRECO-BIT/MEX PWM in MAGIX peaks (up to the top 10,000)

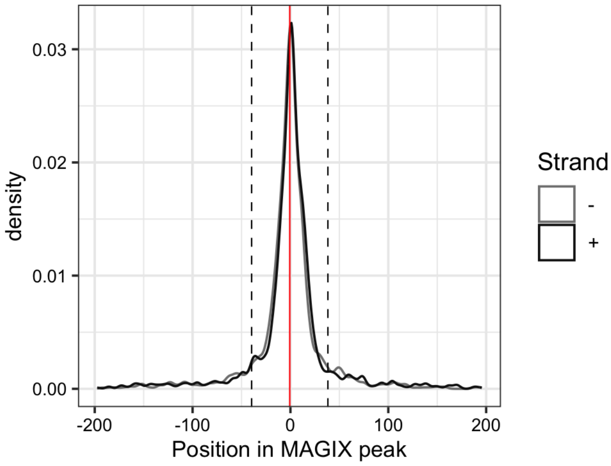


---

AUROC of top GRECO-BIT/MEX PWM in sliding windows (500 peaks, 50 peak step) along the MAGIX peaks (up to the top 75,000) ranked by the MAGIX enrichment coefficient and fraction of overlapping ChIP peaks in the same window

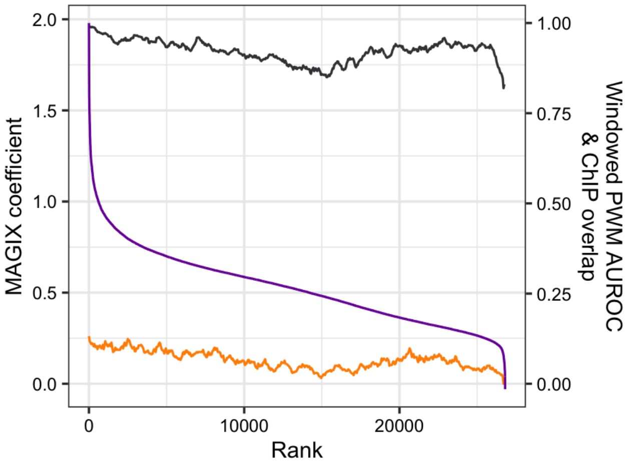

### SRY

Top GRECO-BIT/MEX PWM: PWM002143

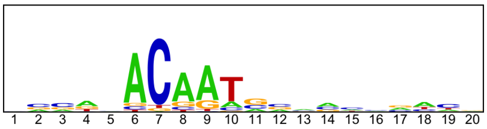

-----------------------------------------------------------------------------------------------

Location of top GRECO-BIT/MEX PWM in MAGIX peaks (up to the top 10,000)

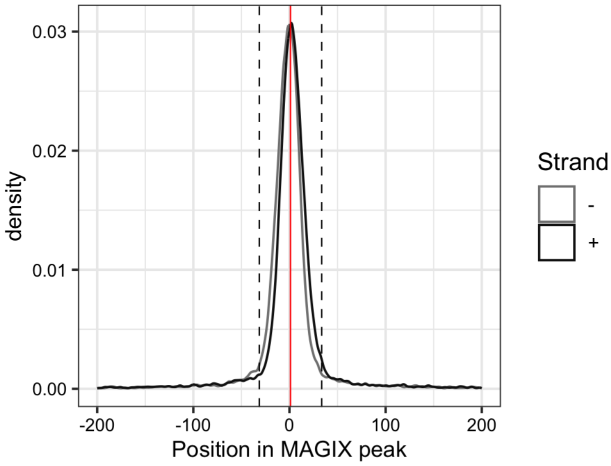


-----------------------------------------------------------------------------------------------

AUROC of top GRECO-BIT/MEX PWM in sliding windows (500 peaks, 50 peak step) along the MAGIX peaks (up to the top 75,000) ranked by the MAGIX enrichment coefficient and fraction of overlapping ChIP peaks in the same window

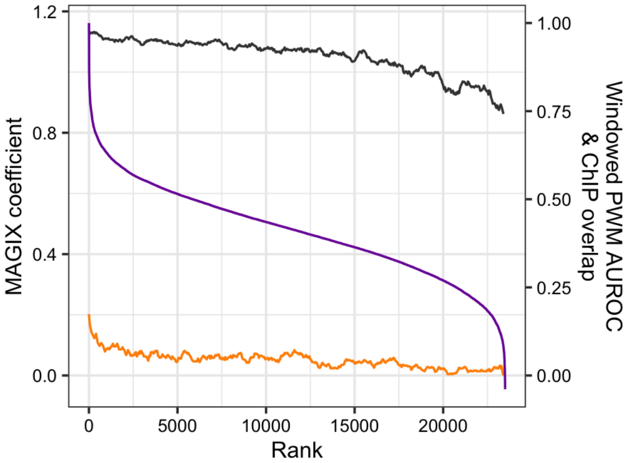

### TERF1

Top GRECO-BIT/MEX PWM: PWM002154

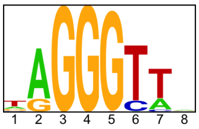


---

Location of top GRECO-BIT/MEX PWM in MAGIX peaks (up to the top 10,000)

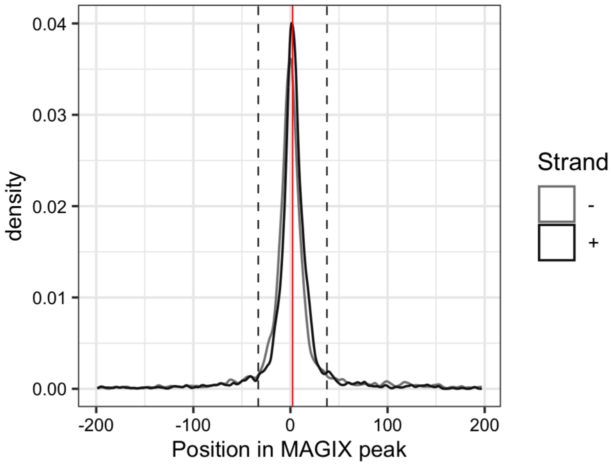


---

AUROC of top GRECO-BIT/MEX PWM in sliding windows (500 peaks, 50 peak step) along the MAGIX peaks (up to the top 75,000) ranked by the MAGIX enrichment coefficient and fraction of overlapping ChIP peaks in the same window

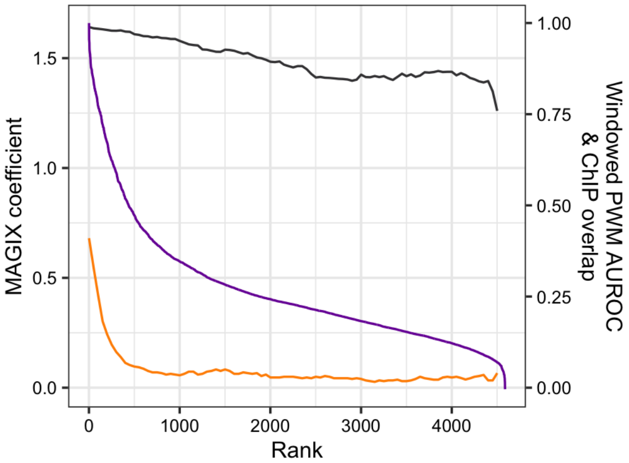

### TET3

Top GRECO-BIT/MEX PWM: PWM002217

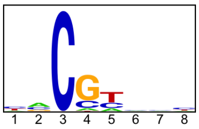


---

Location of top GRECO-BIT/MEX PWM in MAGIX peaks (up to the top 10,000)

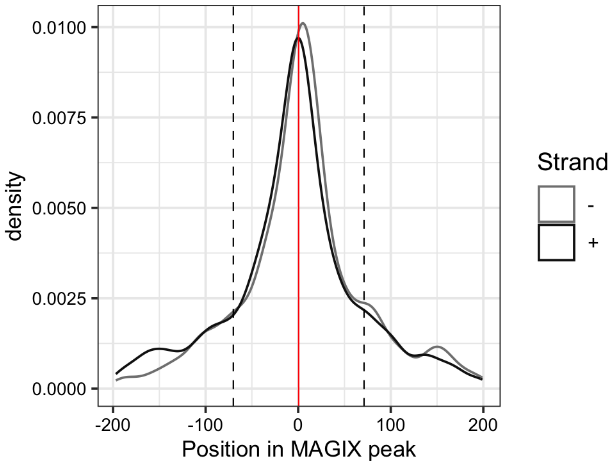


---

AUROC of top GRECO-BIT/MEX PWM in sliding windows (500 peaks, 50 peak step) along the MAGIX peaks (up to the top 75,000) ranked by the MAGIX enrichment coefficient and fraction of overlapping ChIP peaks in the same window

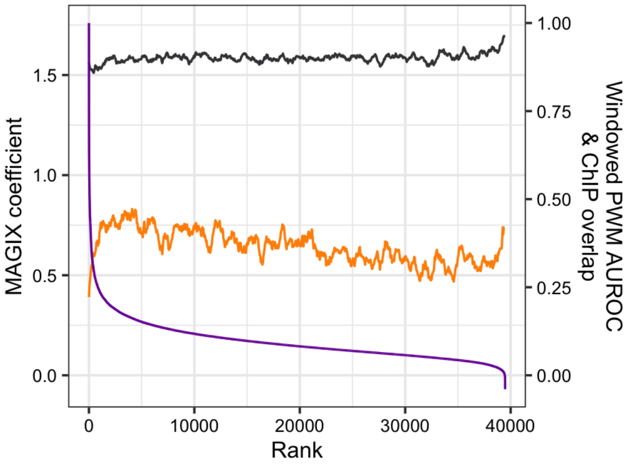

### TIGD3

Top GRECO-BIT/MEX PWM: PWM002247

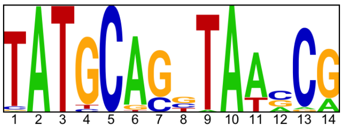


---

Location of top GRECO-BIT/MEX PWM in MAGIX peaks (up to the top 10,000)

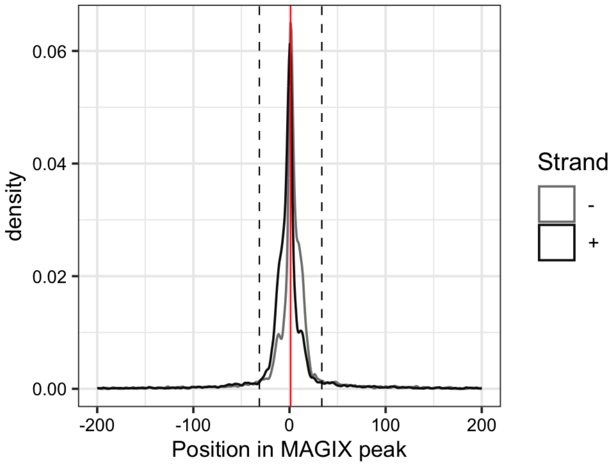


---

AUROC of top GRECO-BIT/MEX PWM in sliding windows (500 peaks, 50 peak step) along the MAGIX peaks (up to the top 75,000) ranked by the MAGIX enrichment coefficient and fraction of overlapping ChIP peaks in the same window

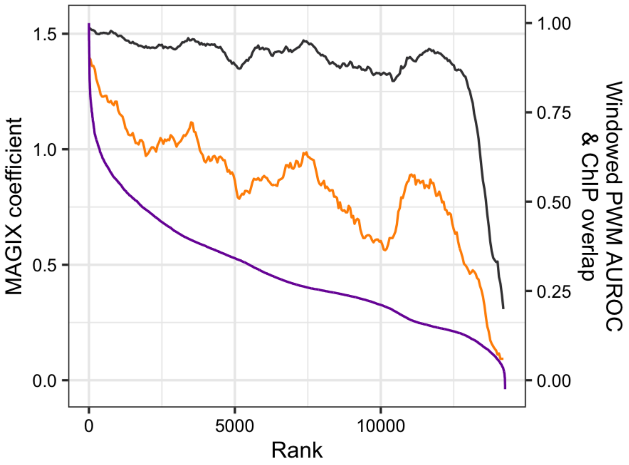

### TIGD4

Top GRECO-BIT/MEX PWM: PWM089770

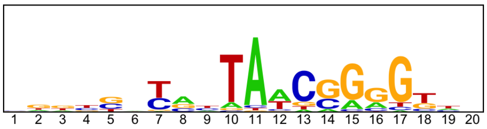


---

Location of top GRECO-BIT/MEX PWM in MAGIX peaks (up to the top 10,000)

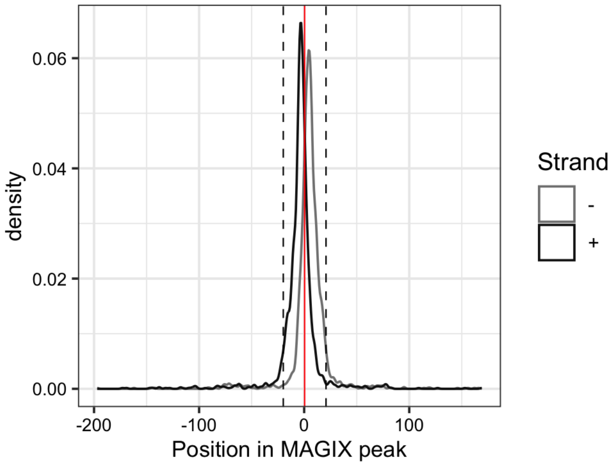

### TIGD5

Top GRECO-BIT/MEX PWM: PWM002303

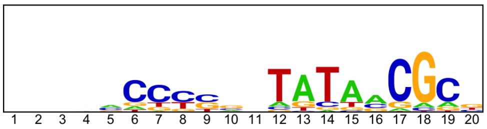


---

Location of top GRECO-BIT/MEX PWM in MAGIX peaks (up to the top 10,000)

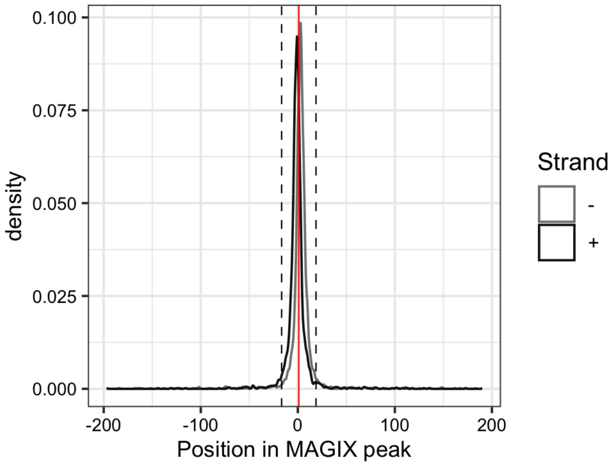


---

AUROC of top GRECO-BIT/MEX PWM in sliding windows (500 peaks, 50 peak step) along the MAGIX peaks (up to the top 75,000) ranked by the MAGIX enrichment coefficient and fraction of overlapping ChIP peaks in the same window

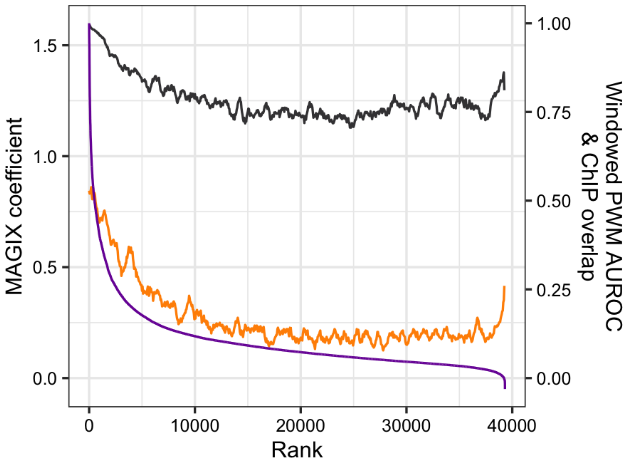

### TIGD7

Top GRECO-BIT/MEX PWM: PWM002334

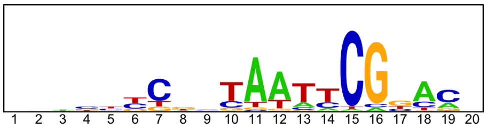


--------------------------------------------------------------------------------------------

Location of top GRECO-BIT/MEX PWM in MAGIX peaks (up to the top 10,000)

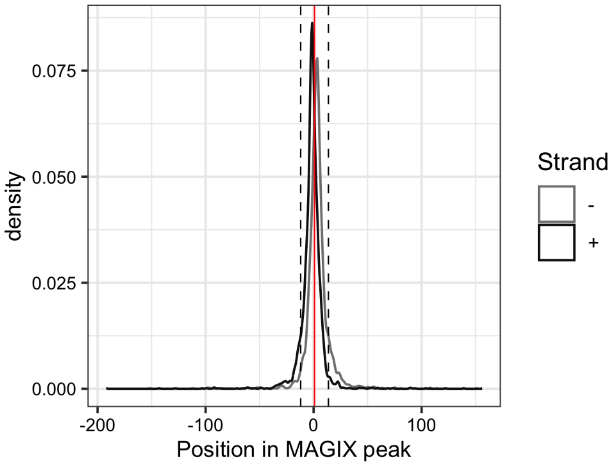

### TPRX1

Top GRECO-BIT/MEX PWM: PWM002363

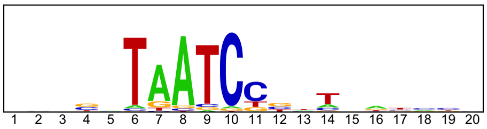


---

Location of top GRECO-BIT/MEX PWM in MAGIX peaks (up to the top 10,000)

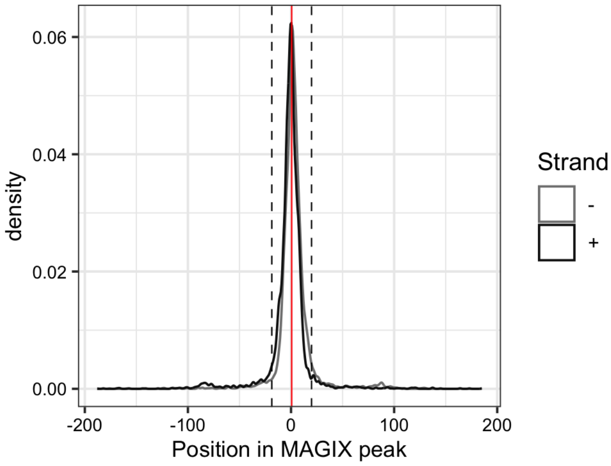


---

AUROC of top GRECO-BIT/MEX PWM in sliding windows (500 peaks, 50 peak step) along the MAGIX peaks (up to the top 75,000) ranked by the MAGIX enrichment coefficient and fraction of overlapping ChIP peaks in the same window

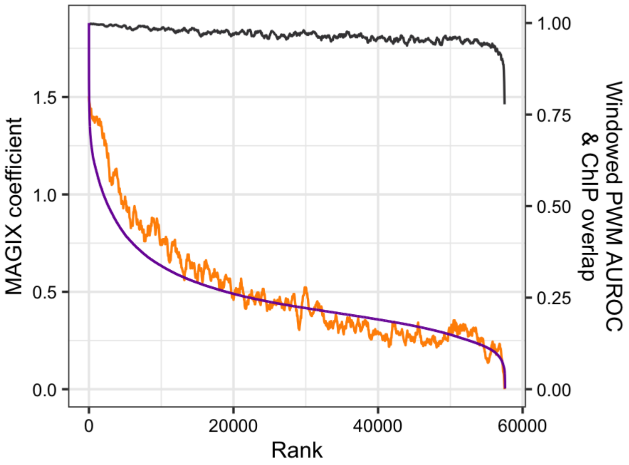

### TSHZ2.DBD2

Top GRECO-BIT/MEX PWM: PWM002395

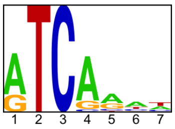


----------------------------------------------------------------------------------------------------

Location of top GRECO-BIT/MEX PWM in MAGIX peaks (up to the top 10,000)

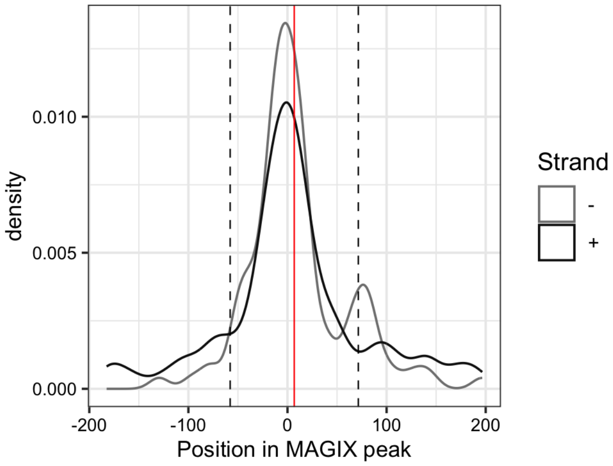

### USF3

Top GRECO-BIT/MEX PWM: PWM002447

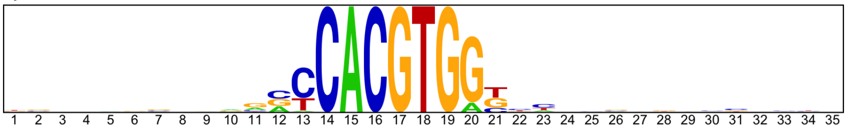

---

Location of top GRECO-BIT/MEX PWM in MAGIX peaks (up to the top 10,000)

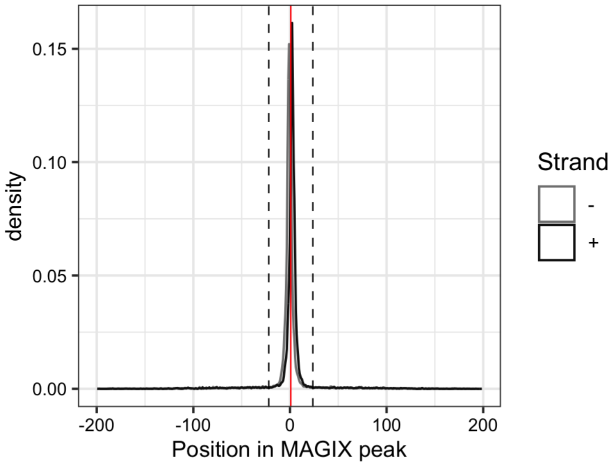


---

AUROC of top GRECO-BIT/MEX PWM in sliding windows (500 peaks, 50 peak step) along the MAGIX peaks (up to the top 75,000) ranked by the MAGIX enrichment coefficient and fraction of overlapping ChIP peaks in the same window

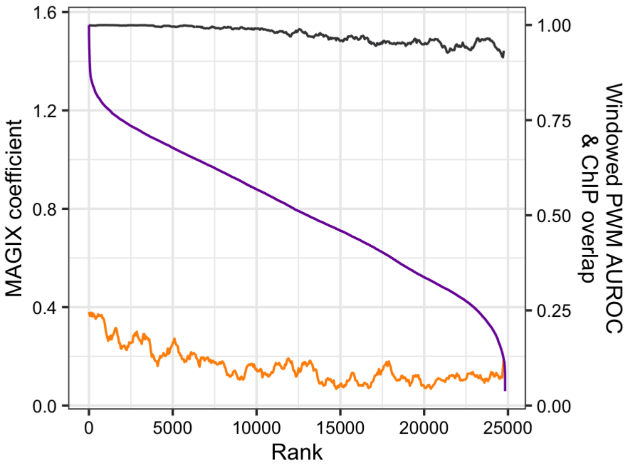

### VDR

Top GRECO-BIT/MEX PWM: PWM002503

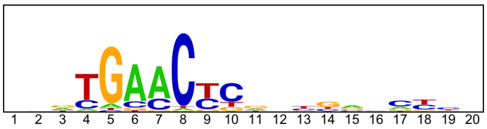


-------------------------------------------------------------------------------------------

Location of top GRECO-BIT/MEX PWM in MAGIX peaks (up to the top 10,000)

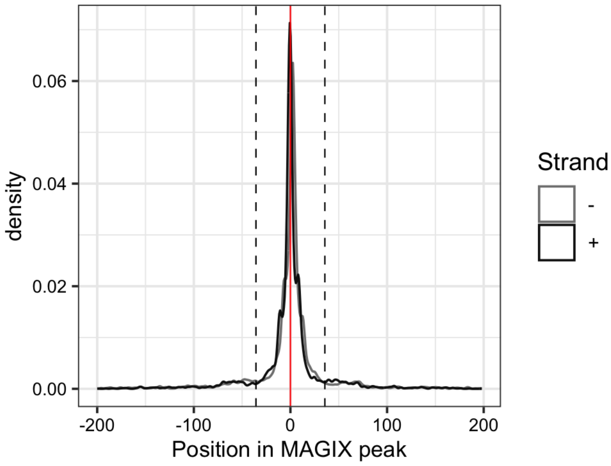


-------------------------------------------------------------------------------------------

AUROC of top GRECO-BIT/MEX PWM in sliding windows (500 peaks, 50 peak step) along the MAGIX peaks (up to the top 75,000) ranked by the MAGIX enrichment coefficient and fraction of overlapping ChIP peaks in the same window

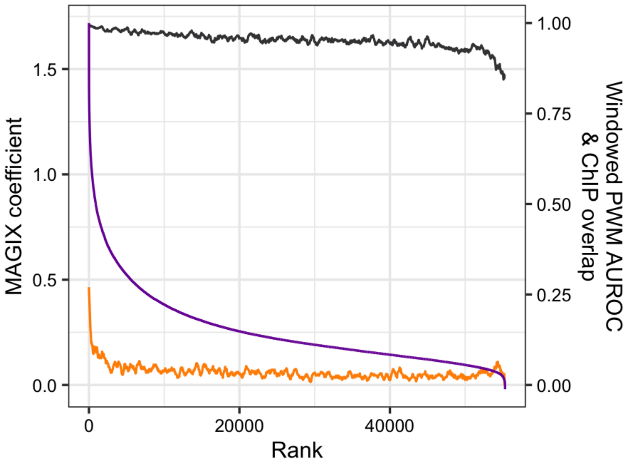

# YY1

Top GRECO-BIT/MEX PWM: PWM002528

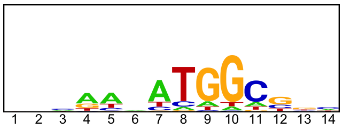


---

Location of top GRECO-BIT/MEX PWM in MAGIX peaks (up to the top 10,000)

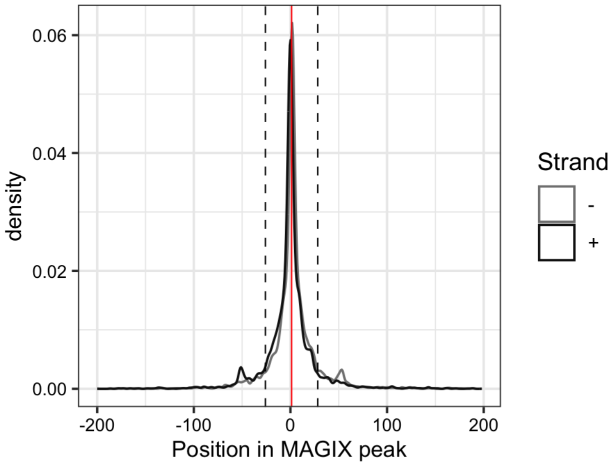


---

AUROC of top GRECO-BIT/MEX PWM in sliding windows (500 peaks, 50 peak step) along the MAGIX peaks (up to the top 75,000) ranked by the MAGIX enrichment coefficient and fraction of overlapping ChIP peaks in the same window

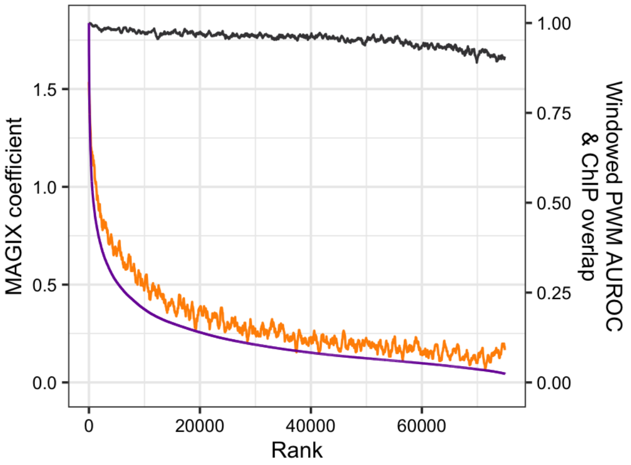

### ZBED2

Top GRECO-BIT/MEX PWM: PWM002573

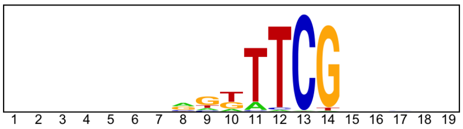


---

Location of top GRECO-BIT/MEX PWM in MAGIX peaks (up to the top 10,000)

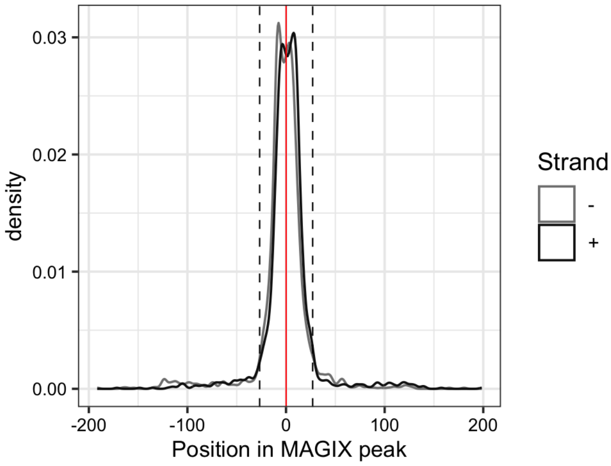


---

AUROC of top GRECO-BIT/MEX PWM in sliding windows (500 peaks, 50 peak step) along the MAGIX peaks (up to the top 75,000) ranked by the MAGIX enrichment coefficient and fraction of overlapping ChIP peaks in the same window

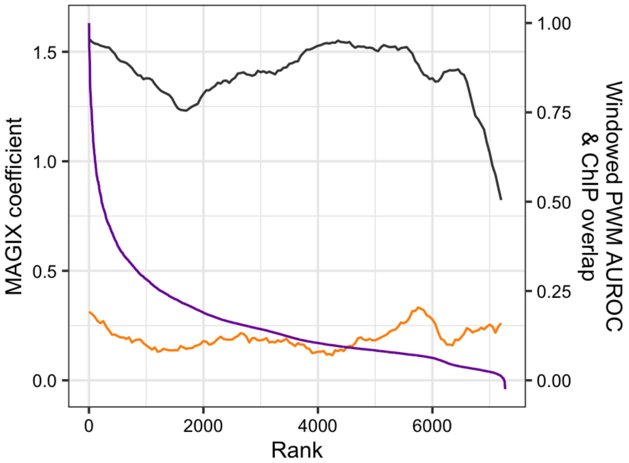

### ZBED4

Top GRECO-BIT/MEX PWM: PWM002593

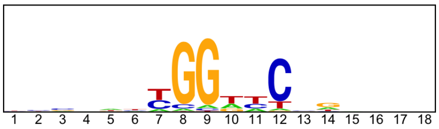


------------------------------------------------------------------------------------------------

Location of top GRECO-BIT/MEX PWM in MAGIX peaks (up to the top 10,000)

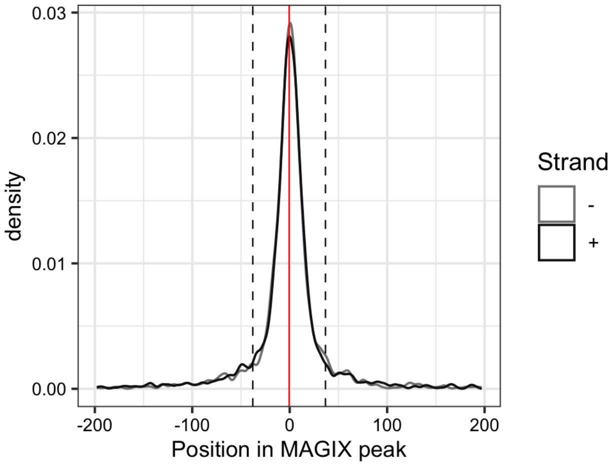

### ZBED5

Top GRECO-BIT/MEX PWM: PWM002641

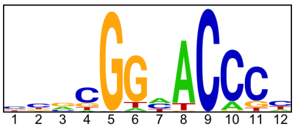


--------------------------------------------------------------------------------------------

Location of top GRECO-BIT/MEX PWM in MAGIX peaks (up to the top 10,000)

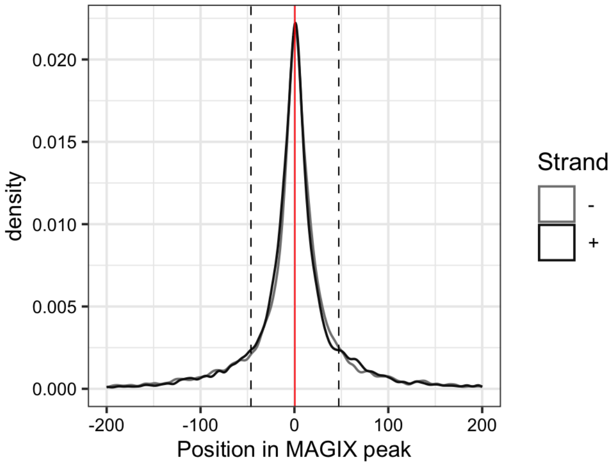


--------------------------------------------------------------------------------------------

AUROC of top GRECO-BIT/MEX PWM in sliding windows (500 peaks, 50 peak step) along the MAGIX peaks (up to the top 75,000) ranked by the MAGIX enrichment coefficient and fraction of overlapping ChIP peaks in the same window

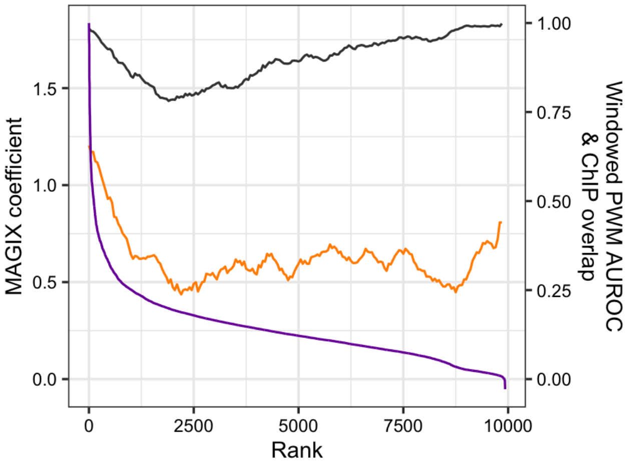

### ZBED9

Top GRECO-BIT/MEX PWM: PWM002687

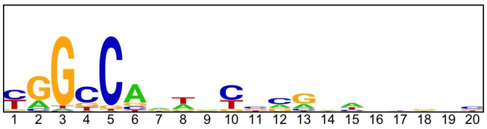


----------------------------------------------------------------------------------------------------

Location of top GRECO-BIT/MEX PWM in MAGIX peaks (up to the top 10,000)

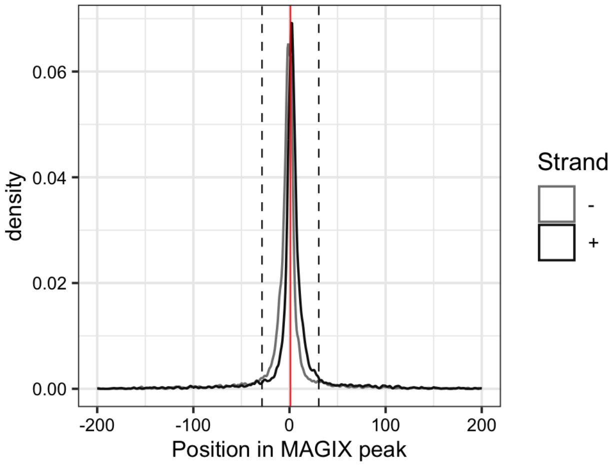


----------------------------------------------------------------------------------------------------

AUROC of top GRECO-BIT/MEX PWM in sliding windows (500 peaks, 50 peak step) along the MAGIX peaks (up to the top 75,000) ranked by the MAGIX enrichment coefficient and fraction of overlapping ChIP peaks in the same window

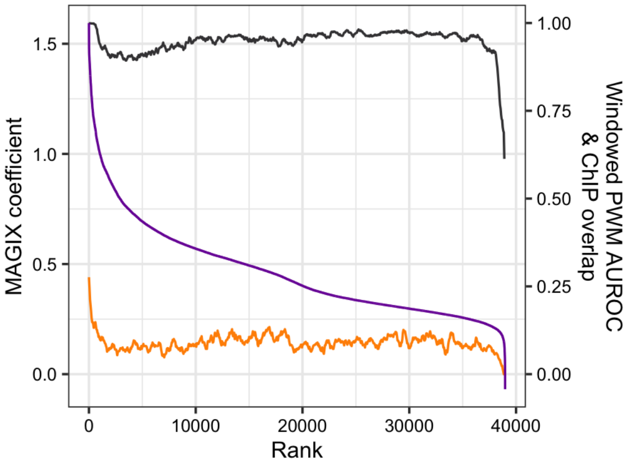

### ZBTB24

Top GRECO-BIT/MEX PWM: PWM002706

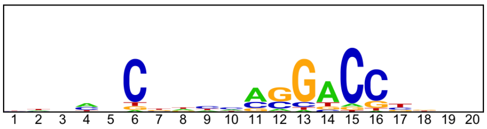


-----------------------------------------------------------------------------------------------

Location of top GRECO-BIT/MEX PWM in MAGIX peaks (up to the top 10,000)

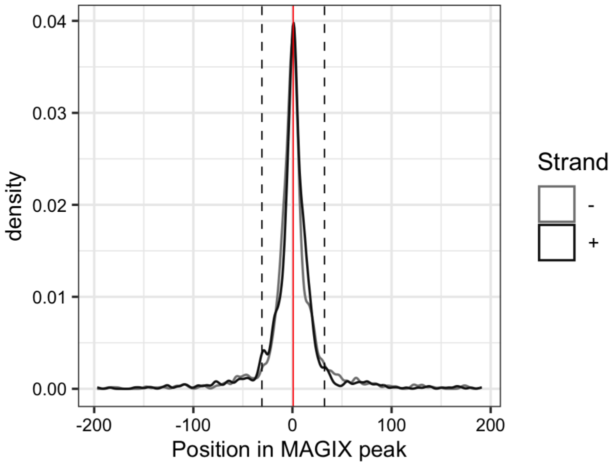

### ZBTB40

Top GRECO-BIT/MEX PWM: PWM002730

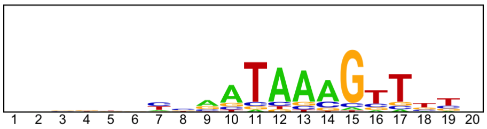


---

Location of top GRECO-BIT/MEX PWM in MAGIX peaks (up to the top 10,000)

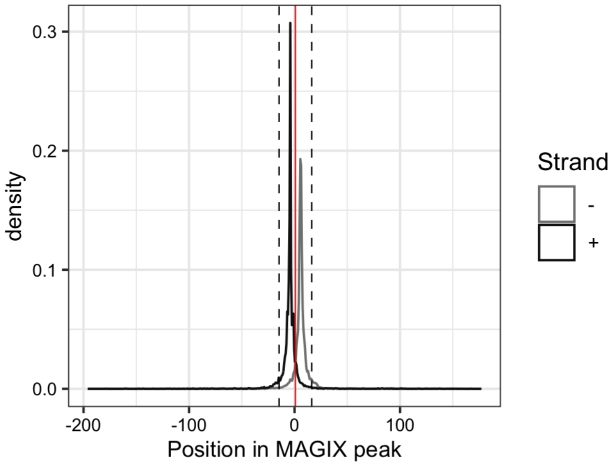


---

AUROC of top GRECO-BIT/MEX PWM in sliding windows (500 peaks, 50 peak step) along the MAGIX peaks (up to the top 75,000) ranked by the MAGIX enrichment coefficient and fraction of overlapping ChIP peaks in the same window

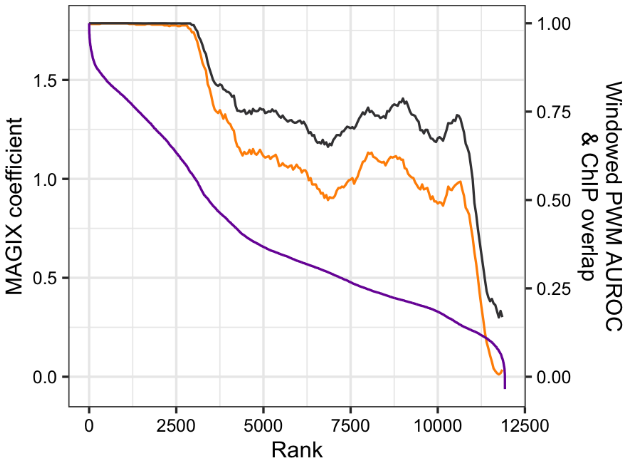

### ZBTB41

Top GRECO-BIT/MEX PWM: PWM002773

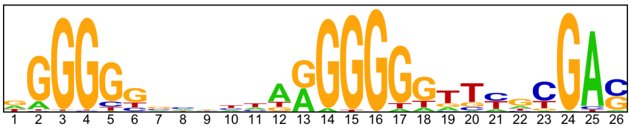


-----------------------------------------------------------------------------------------------

Location of top GRECO-BIT/MEX PWM in MAGIX peaks (up to the top 10,000)

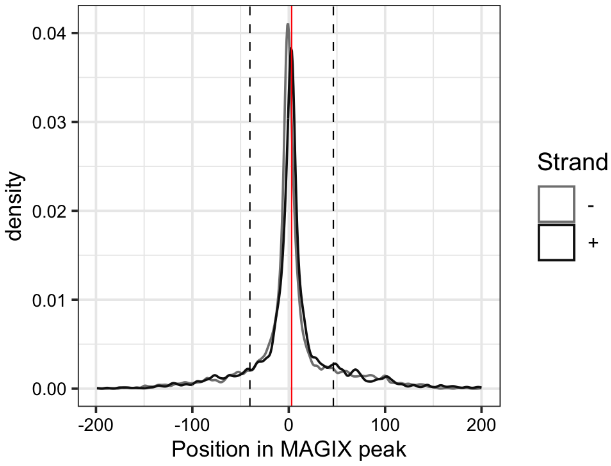


-----------------------------------------------------------------------------------------------

AUROC of top GRECO-BIT/MEX PWM in sliding windows (500 peaks, 50 peak step) along the MAGIX peaks (up to the top 75,000) ranked by the MAGIX enrichment coefficient and fraction of overlapping ChIP peaks in the same window

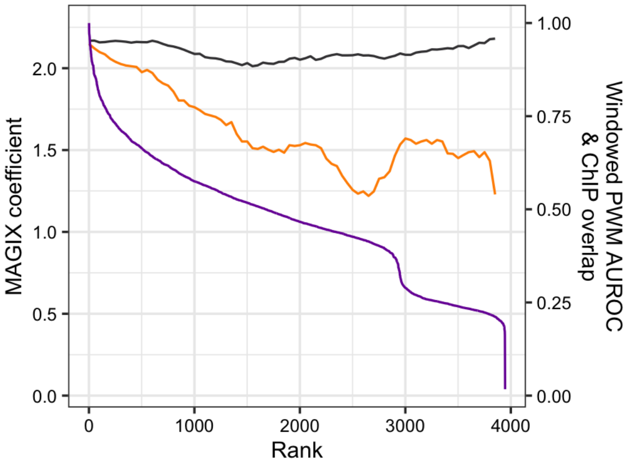

#### ZBTB47

Top GRECO-BIT/MEX PWM: PWM002799

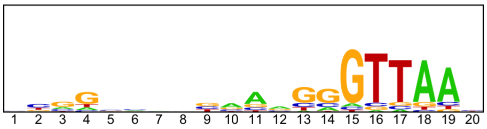


---

Location of top GRECO-BIT/MEX PWM in MAGIX peaks (up to the top 10,000)

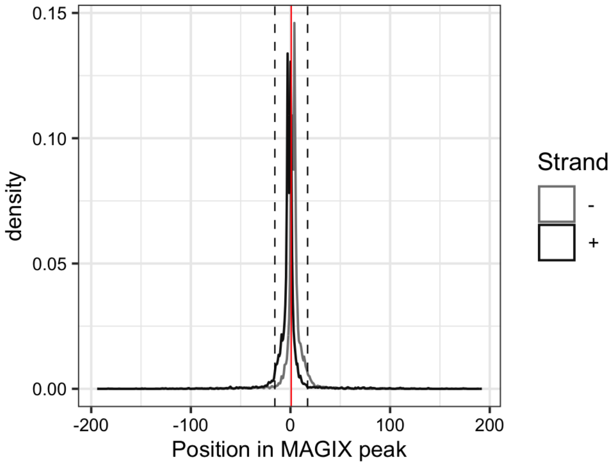


---

AUROC of top GRECO-BIT/MEX PWM in sliding windows (500 peaks, 50 peak step) along the MAGIX peaks (up to the top 75,000) ranked by the MAGIX enrichment coefficient and fraction of overlapping ChIP peaks in the same window

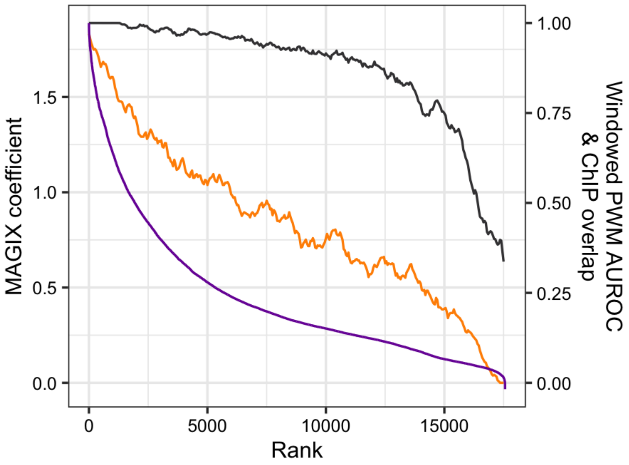

### ZBTB5

Top GRECO-BIT/MEX PWM: PWM002811

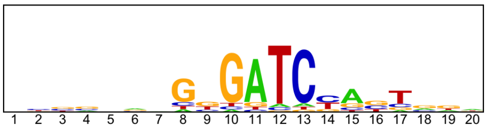


----------------------------------------------------------------------------------------------------

Location of top GRECO-BIT/MEX PWM in MAGIX peaks (up to the top 10,000)

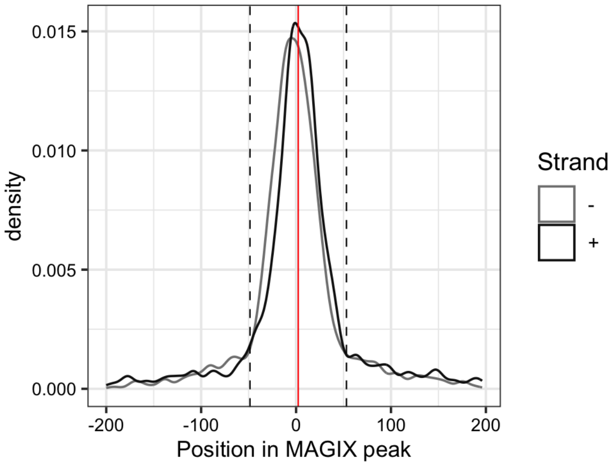

### ZBTB8A

Top GRECO-BIT/MEX PWM: PWM002842

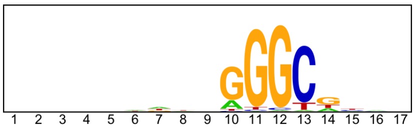


-----------------------------------------------------------------------------------------------

Location of top GRECO-BIT/MEX PWM in MAGIX peaks (up to the top 10,000)

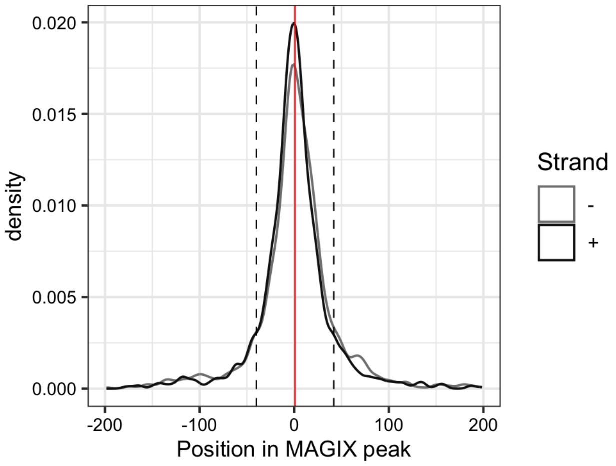


-----------------------------------------------------------------------------------------------

AUROC of top GRECO-BIT/MEX PWM in sliding windows (500 peaks, 50 peak step) along the MAGIX peaks (up to the top 75,000) ranked by the MAGIX enrichment coefficient and fraction of overlapping ChIP peaks in the same window

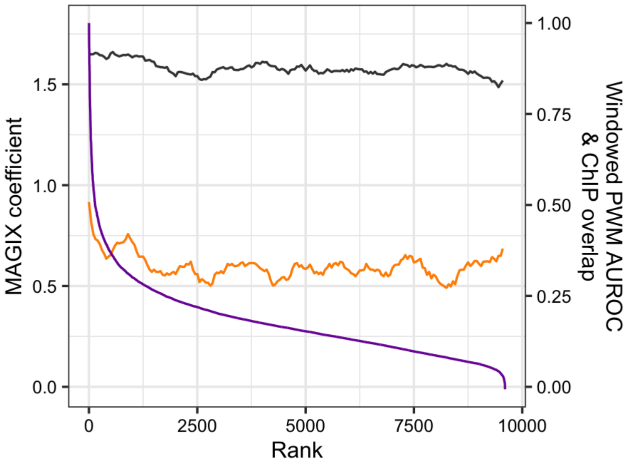

### ZBTB8B

Top GRECO-BIT/MEX PWM: PWM002878

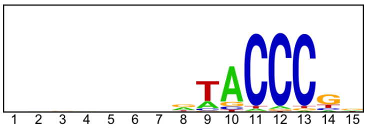


-----------------------------------------------------------------------------------------------

Location of top GRECO-BIT/MEX PWM in MAGIX peaks (up to the top 10,000)

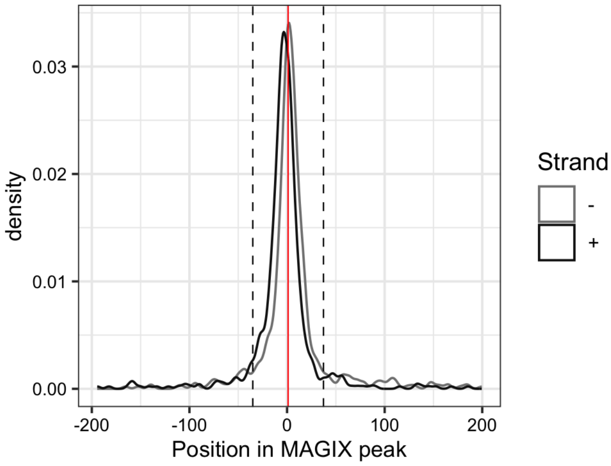


-----------------------------------------------------------------------------------------------

AUROC of top GRECO-BIT/MEX PWM in sliding windows (500 peaks, 50 peak step) along the MAGIX peaks (up to the top 75,000) ranked by the MAGIX enrichment coefficient and fraction of overlapping ChIP peaks in the same window

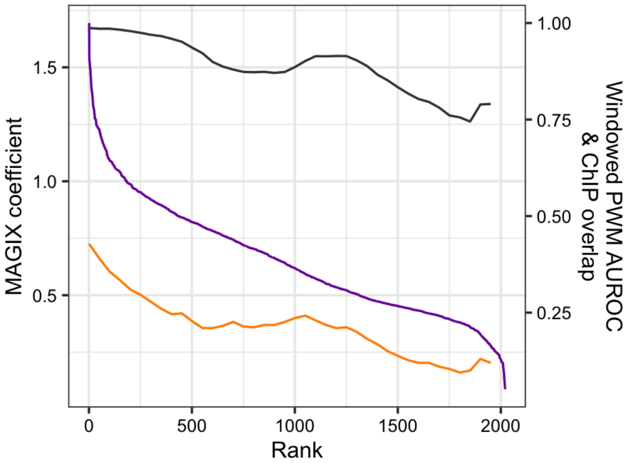

### ZFAT.DBD1

Top GRECO-BIT/MEX PWM: PWM002917

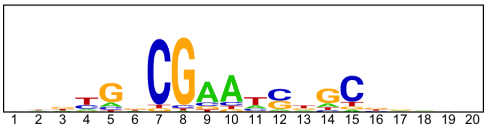


---------------------------------------------------------------------------------------------

Location of top GRECO-BIT/MEX PWM in MAGIX peaks (up to the top 10,000)

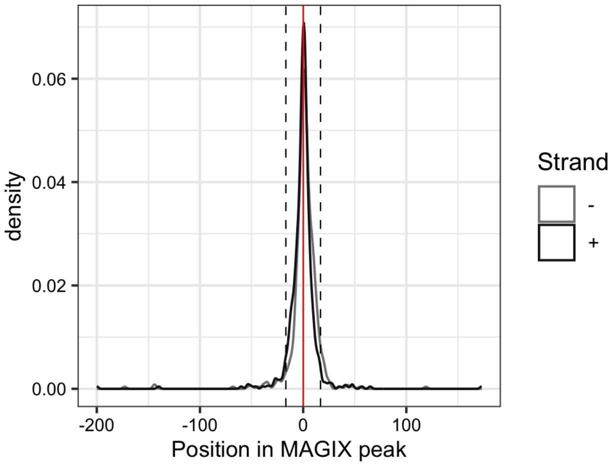

### ZFP3

Top GRECO-BIT/MEX PWM: PWM002973

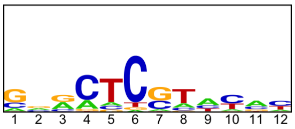


---

Location of top GRECO-BIT/MEX PWM in MAGIX peaks (up to the top 10,000)

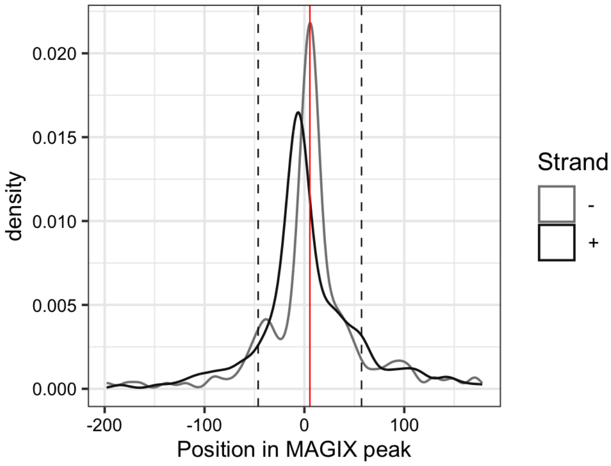


---

AUROC of top GRECO-BIT/MEX PWM in sliding windows (500 peaks, 50 peak step) along the MAGIX peaks (up to the top 75,000) ranked by the MAGIX enrichment coefficient and fraction of overlapping ChIP peaks in the same window

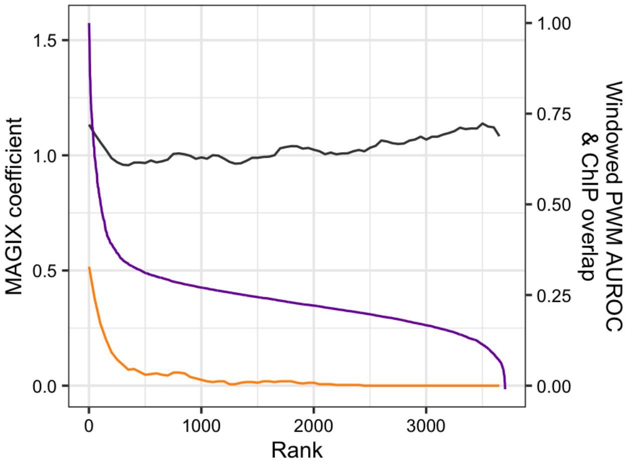

### ZFP91

Top GRECO-BIT/MEX PWM: PWM002984

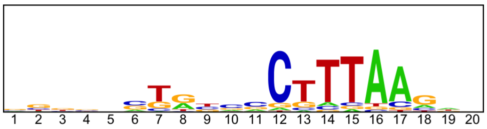


----------------------------------------------------------------------------------------------------

Location of top GRECO-BIT/MEX PWM in MAGIX peaks (up to the top 10,000)

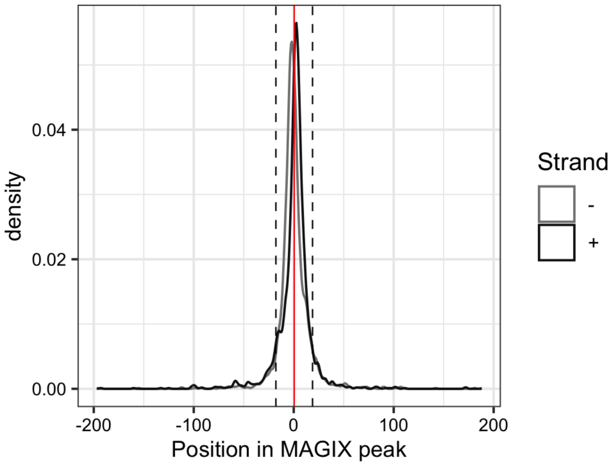

### ZFTA

Top GRECO-BIT/MEX PWM: PWM003020

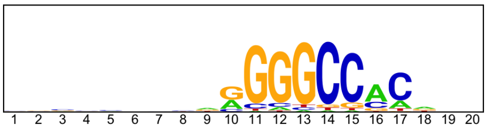

--------------------------------------------------------------------------------------------

Location of top GRECO-BIT/MEX PWM in MAGIX peaks (up to the top 10,000)

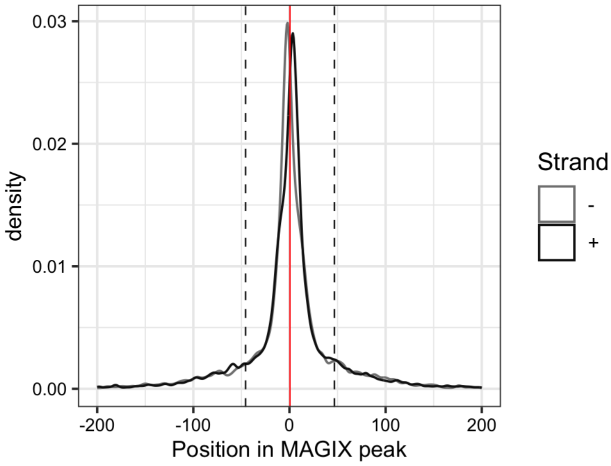


--------------------------------------------------------------------------------------------

AUROC of top GRECO-BIT/MEX PWM in sliding windows (500 peaks, 50 peak step) along the MAGIX peaks (up to the top 75,000) ranked by the MAGIX enrichment coefficient and fraction of overlapping ChIP peaks in the same window

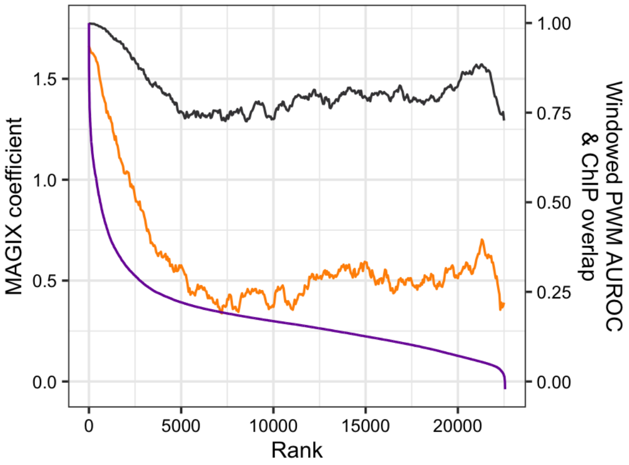

### ZGLP1

Top GRECO-BIT/MEX PWM: PWM003053

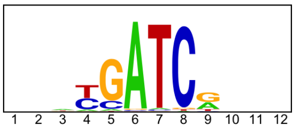


-----------------------------------------------------------------------------------------------

Location of top GRECO-BIT/MEX PWM in MAGIX peaks (up to the top 10,000)

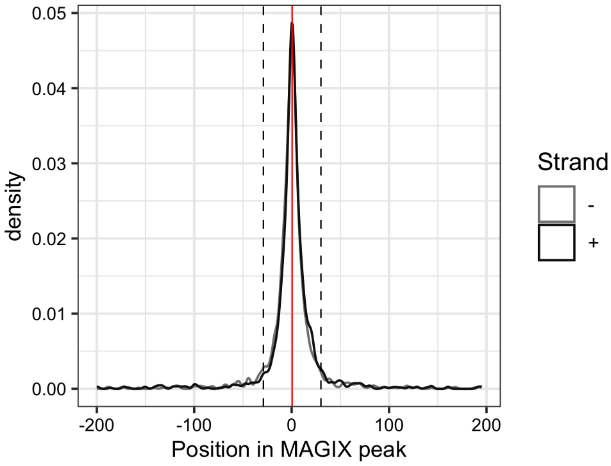

### ZIM3

Top GRECO-BIT/MEX PWM: PWM003079

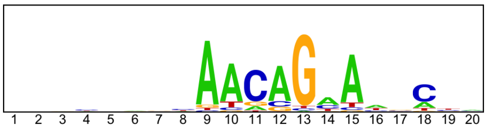


-----------------------------------------------------------------------------------------------

Location of top GRECO-BIT/MEX PWM in MAGIX peaks (up to the top 10,000)

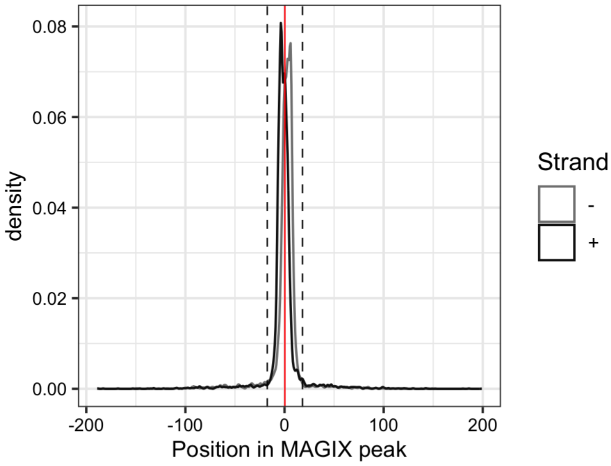


-----------------------------------------------------------------------------------------------

AUROC of top GRECO-BIT/MEX PWM in sliding windows (500 peaks, 50 peak step) along the MAGIX peaks (up to the top 75,000) ranked by the MAGIX enrichment coefficient and fraction of overlapping ChIP peaks in the same window

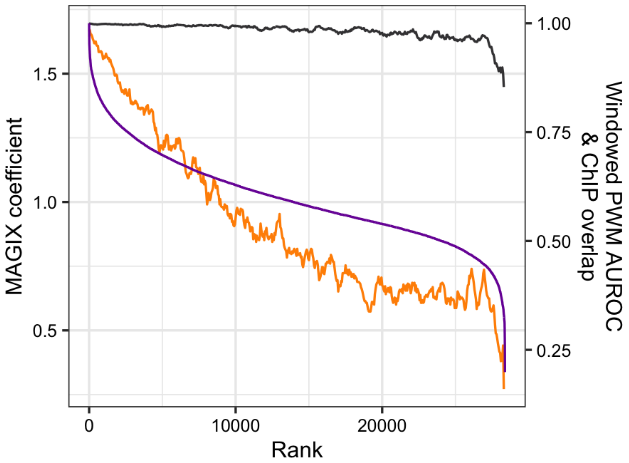

### ZKSCAN4

Top GRECO-BIT/MEX PWM: PWM003099

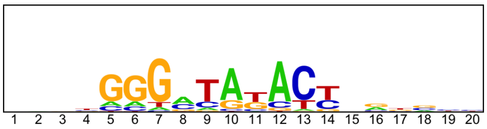

---

Location of top GRECO-BIT/MEX PWM in MAGIX peaks (up to the top 10,000)

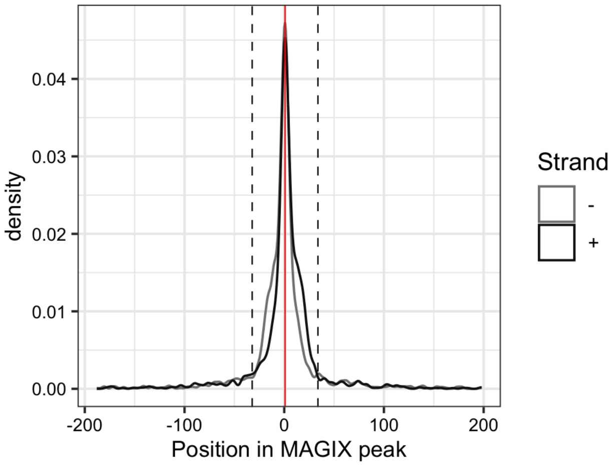

### ZNF107.DBD1

Top GRECO-BIT/MEX PWM: PWM003143

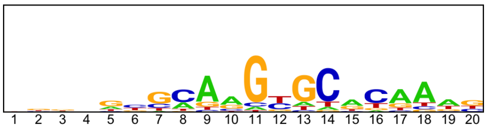


------------------------------------------------------------------------------------------------

Location of top GRECO-BIT/MEX PWM in MAGIX peaks (up to the top 10,000)

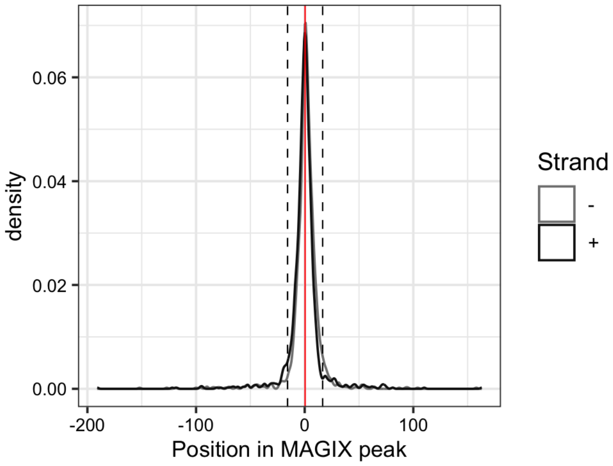

### ZNF107.DBD2

Top GRECO-BIT/MEX PWM: PWM003126

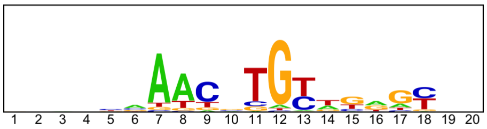


------------------------------------------------------------------------------------------------

Location of top GRECO-BIT/MEX PWM in MAGIX peaks (up to the top 10,000)

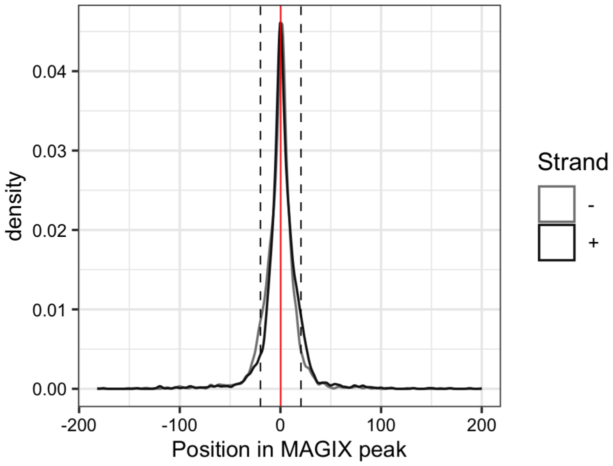

### ZNF131

Top GRECO-BIT/MEX PWM: PWM003208

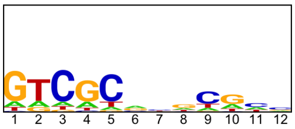


-----------------------------------------------------------------------------------------

Location of top GRECO-BIT/MEX PWM in MAGIX peaks (up to the top 10,000)

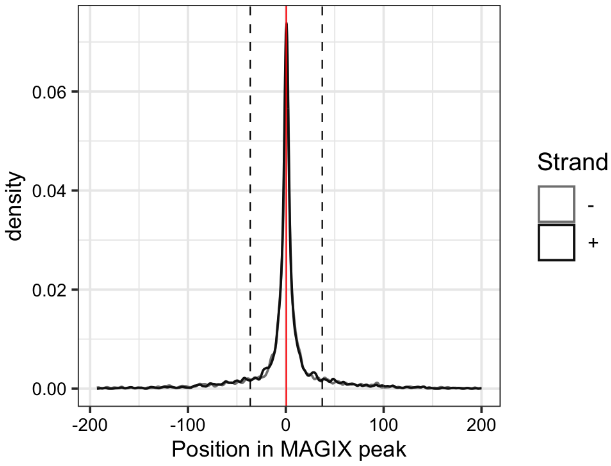


-----------------------------------------------------------------------------------------

AUROC of top GRECO-BIT/MEX PWM in sliding windows (500 peaks, 50 peak step) along the MAGIX peaks (up to the top 75,000) ranked by the MAGIX enrichment coefficient and fraction of overlapping ChIP peaks in the same window

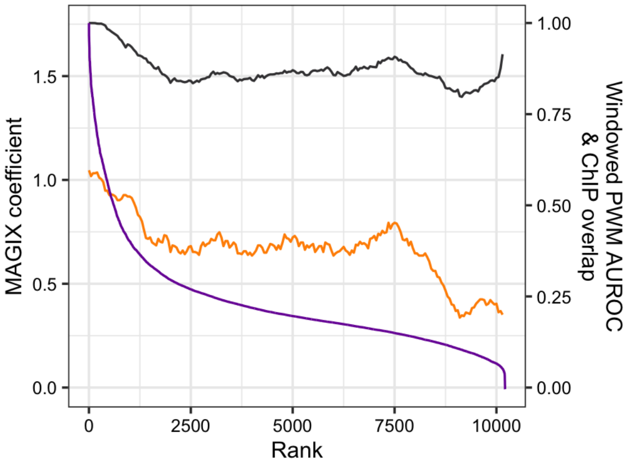

### ZNF134

Top GRECO-BIT/MEX PWM: PWM003228

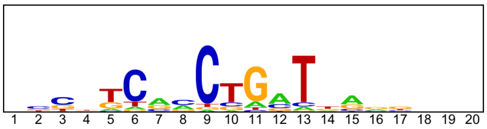


---

Location of top GRECO-BIT/MEX PWM in MAGIX peaks (up to the top 10,000)

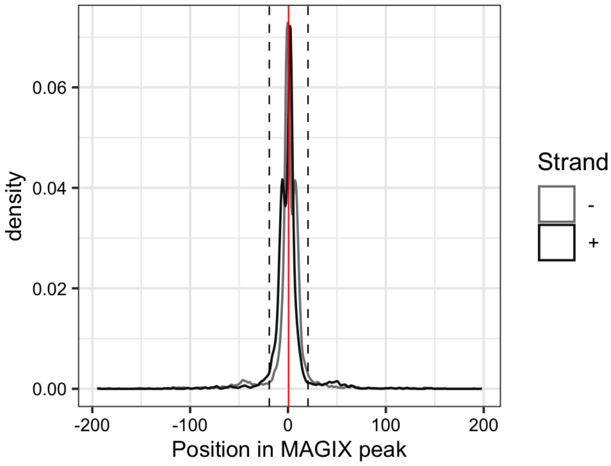


---

AUROC of top GRECO-BIT/MEX PWM in sliding windows (500 peaks, 50 peak step) along the MAGIX peaks (up to the top 75,000) ranked by the MAGIX enrichment coefficient and fraction of overlapping ChIP peaks in the same window

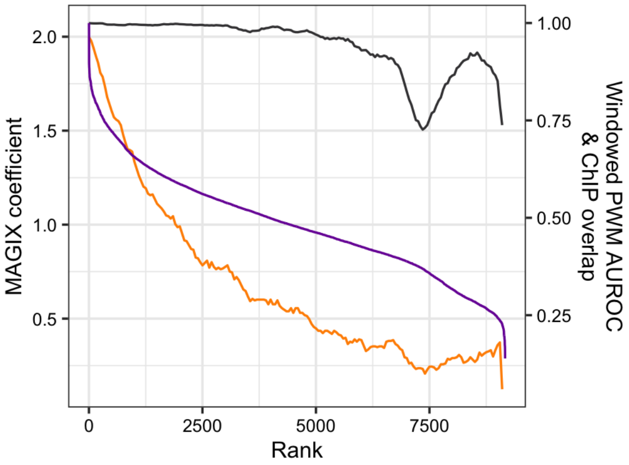

### ZNF142.DBD1

Top GRECO-BIT/MEX PWM: PWM003313

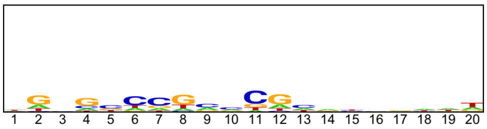


------------------------------------------------------------------------------------------------

Location of top GRECO-BIT/MEX PWM in MAGIX peaks (up to the top 10,000)

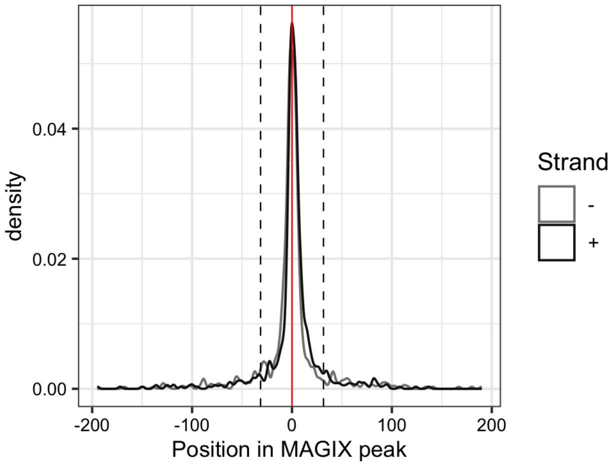

### ZNF16

Top GRECO-BIT/MEX PWM: PWM003349

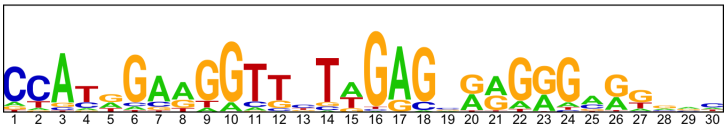


---

Location of top GRECO-BIT/MEX PWM in MAGIX peaks (up to the top 10,000)

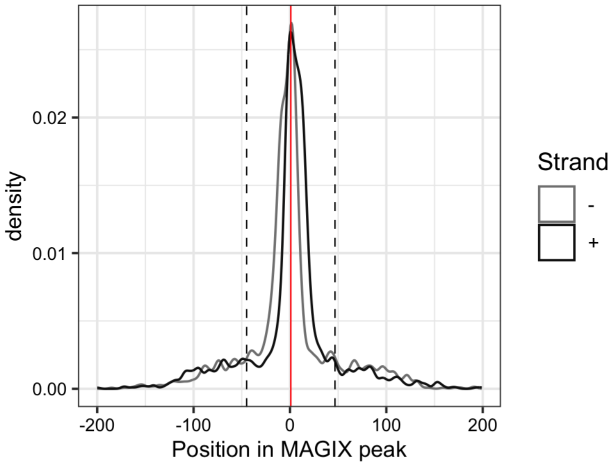

### ZNF160

Top GRECO-BIT/MEX PWM: PWM003369

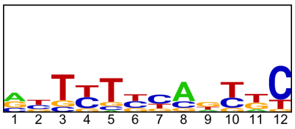


----------------------------------------------------------------------------------------------

Location of top GRECO-BIT/MEX PWM in MAGIX peaks (up to the top 10,000)

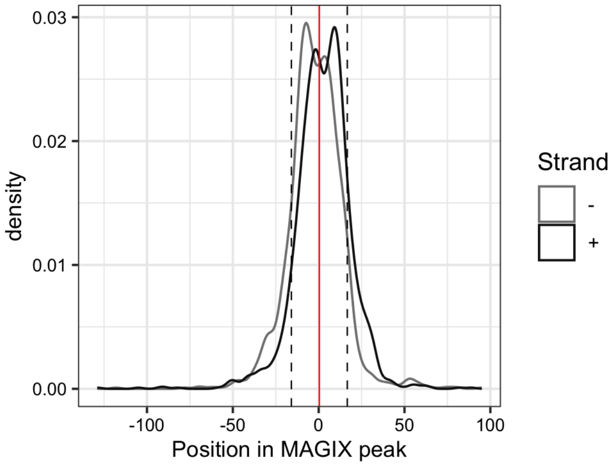

### ZNF18

Top GRECO-BIT/MEX PWM: PWM003393

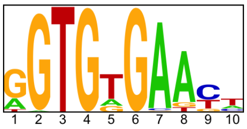


--------------------------------------------------------------------------------------------

Location of top GRECO-BIT/MEX PWM in MAGIX peaks (up to the top 10,000)

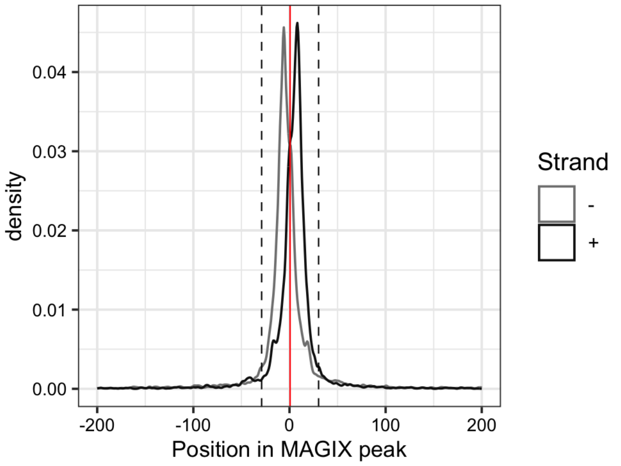


--------------------------------------------------------------------------------------------

AUROC of top GRECO-BIT/MEX PWM in sliding windows (500 peaks, 50 peak step) along the MAGIX peaks (up to the top 75,000) ranked by the MAGIX enrichment coefficient and fraction of overlapping ChIP peaks in the same window

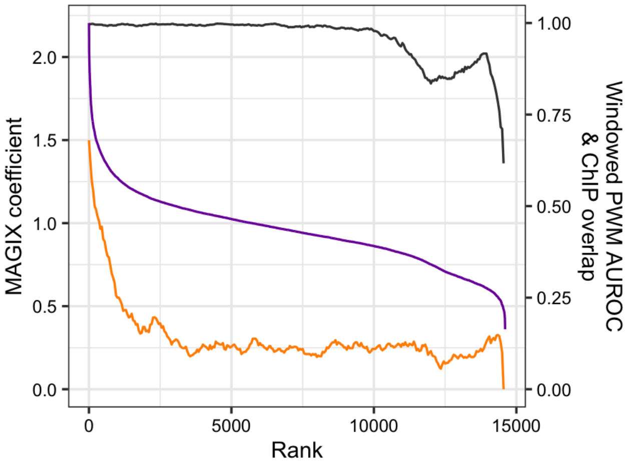

### ZNF20

Top GRECO-BIT/MEX PWM: PWM003422

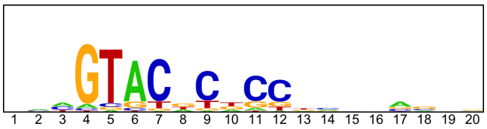


-----------------------------------------------------------------------------------------------

Location of top GRECO-BIT/MEX PWM in MAGIX peaks (up to the top 10,000)

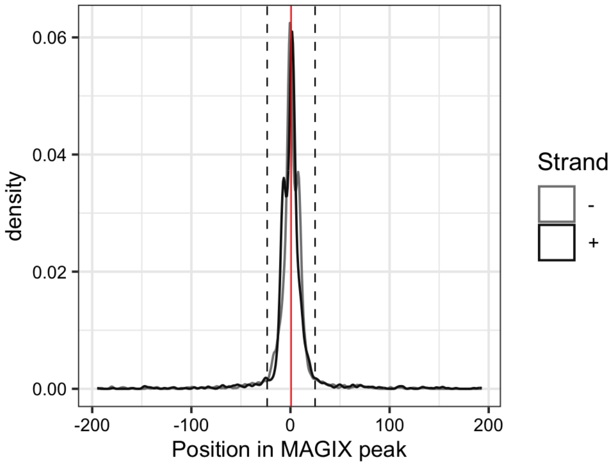


-----------------------------------------------------------------------------------------------

AUROC of top GRECO-BIT/MEX PWM in sliding windows (500 peaks, 50 peak step) along the MAGIX peaks (up to the top 75,000) ranked by the MAGIX enrichment coefficient and fraction of overlapping ChIP peaks in the same window

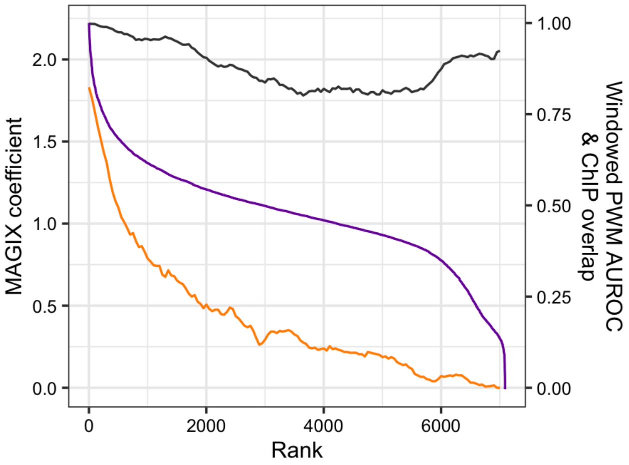

### ZNF208.DBD1

Top GRECO-BIT/MEX PWM: PWM124525

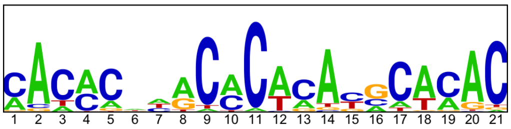


-----------------------------------------------------------------------------------------------

Location of top GRECO-BIT/MEX PWM in MAGIX peaks (up to the top 10,000)

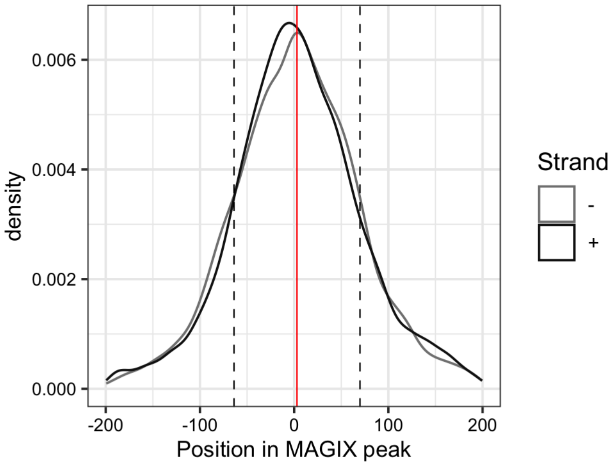

### ZNF208.DBD2

Top GRECO-BIT/MEX PWM: PWM003442

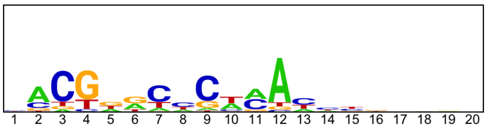


------------------------------------------------------------------------------------------------

Location of top GRECO-BIT/MEX PWM in MAGIX peaks (up to the top 10,000)

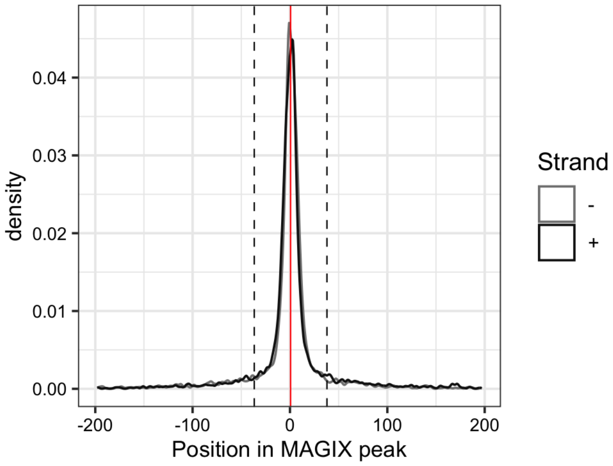

### ZNF208.DBD3

Top GRECO-BIT/MEX PWM: PWM003448

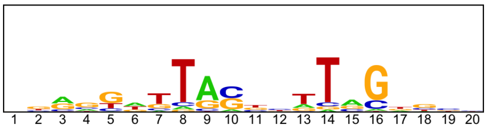


------------------------------------------------------------------------------------------------

Location of top GRECO-BIT/MEX PWM in MAGIX peaks (up to the top 10,000)

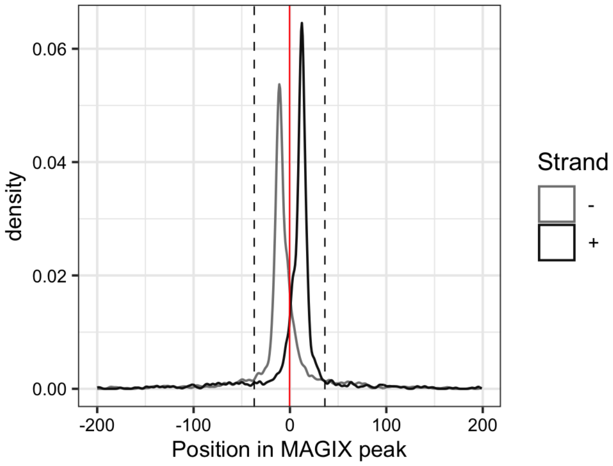

### ZNF215

Top GRECO-BIT/MEX PWM: PWM003473

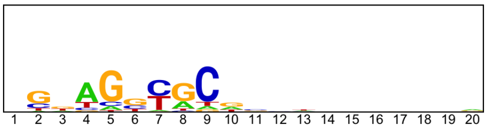


--------------------------------------------------------------------------------------------

Location of top GRECO-BIT/MEX PWM in MAGIX peaks (up to the top 10,000)

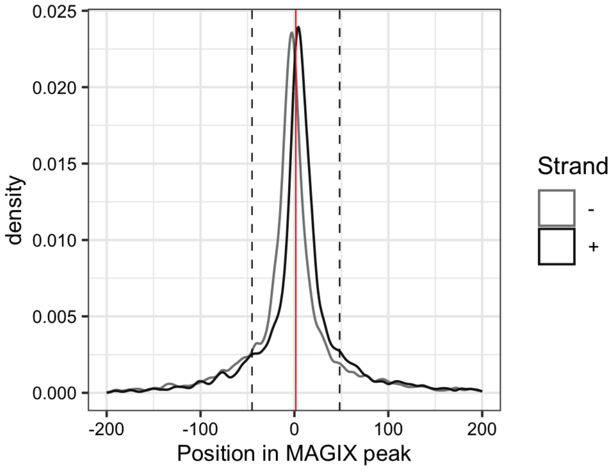


--------------------------------------------------------------------------------------------

AUROC of top GRECO-BIT/MEX PWM in sliding windows (500 peaks, 50 peak step) along the MAGIX peaks (up to the top 75,000) ranked by the MAGIX enrichment coefficient and fraction of overlapping ChIP peaks in the same window

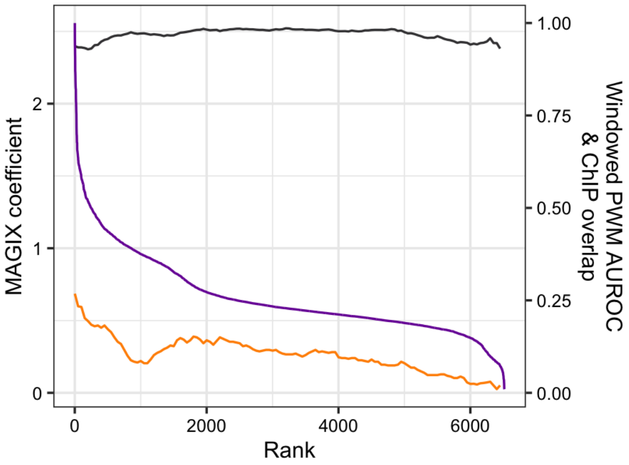

### ZNF226

Top GRECO-BIT/MEX PWM: PWM003505

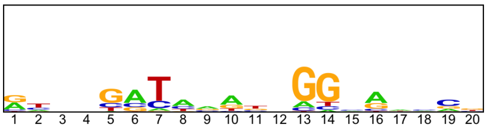


--------------------------------------------------------------------------------------------

Location of top GRECO-BIT/MEX PWM in MAGIX peaks (up to the top 10,000)

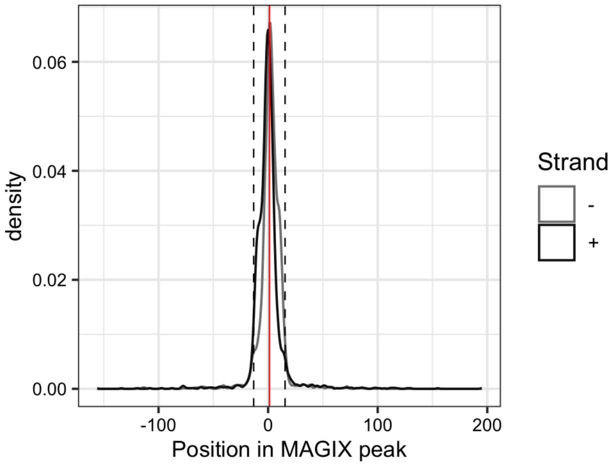


--------------------------------------------------------------------------------------------

AUROC of top GRECO-BIT/MEX PWM in sliding windows (500 peaks, 50 peak step) along the MAGIX peaks (up to the top 75,000) ranked by the MAGIX enrichment coefficient and fraction of overlapping ChIP peaks in the same window

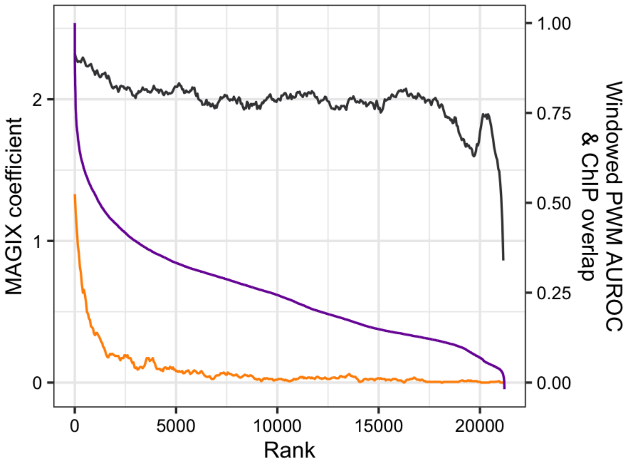

### ZNF229

Top GRECO-BIT/MEX PWM: PWM003525

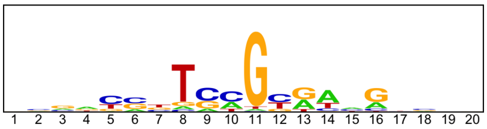


-----------------------------------------------------------------------------------------------

Location of top GRECO-BIT/MEX PWM in MAGIX peaks (up to the top 10,000)

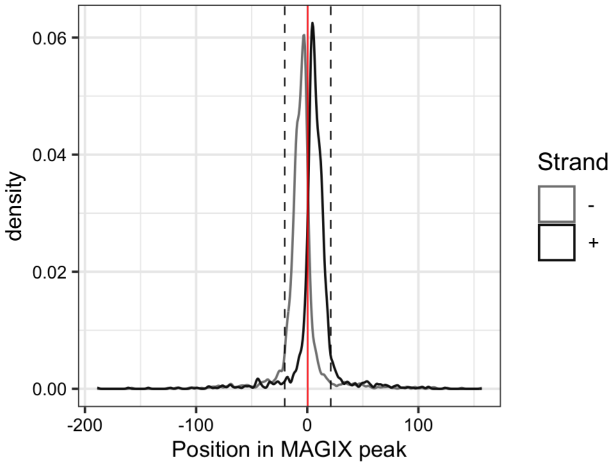

### ZNF233

Top GRECO-BIT/MEX PWM: PWM003570

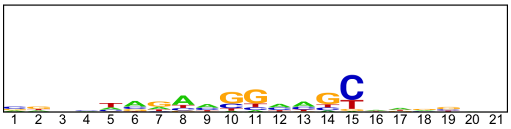


------------------------------------------------------------------------------------------------

Location of top GRECO-BIT/MEX PWM in MAGIX peaks (up to the top 10,000)

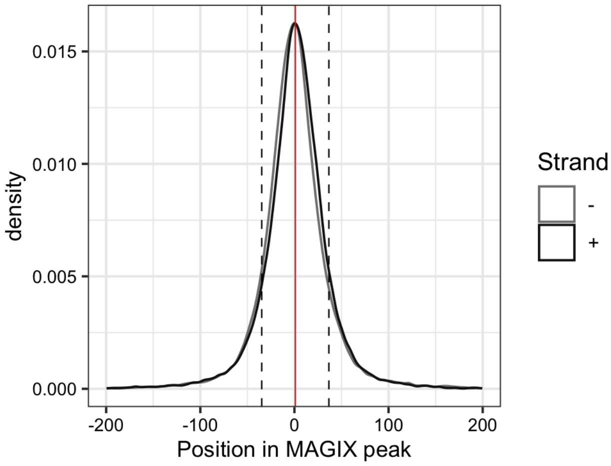


------------------------------------------------------------------------------------------------

AUROC of top GRECO-BIT/MEX PWM in sliding windows (500 peaks, 50 peak step) along the MAGIX peaks (up to the top 75,000) ranked by the MAGIX enrichment coefficient and fraction of overlapping ChIP peaks in the same window

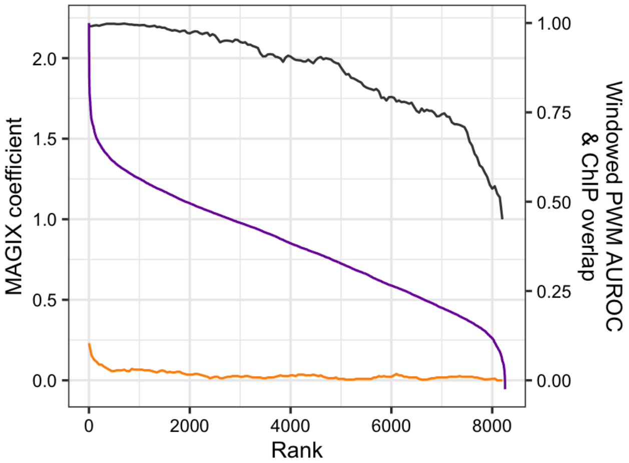

### ZNF234

Top GRECO-BIT/MEX PWM: PWM003611

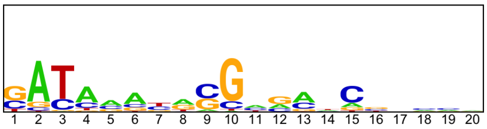

---

Location of top GRECO-BIT/MEX PWM in MAGIX peaks (up to the top 10,000)

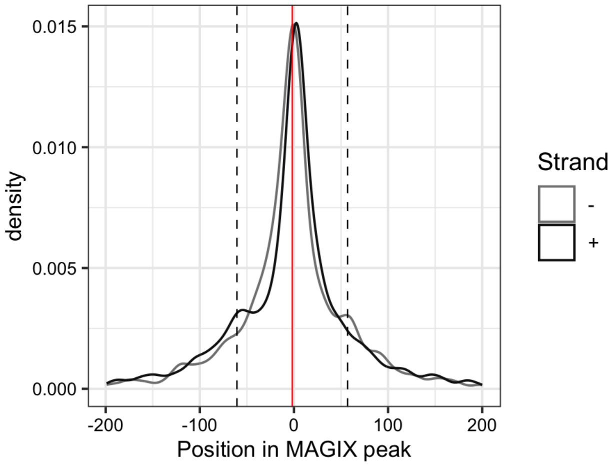


---

AUROC of top GRECO-BIT/MEX PWM in sliding windows (500 peaks, 50 peak step) along the MAGIX peaks (up to the top 75,000) ranked by the MAGIX enrichment coefficient and fraction of overlapping ChIP peaks in the same window

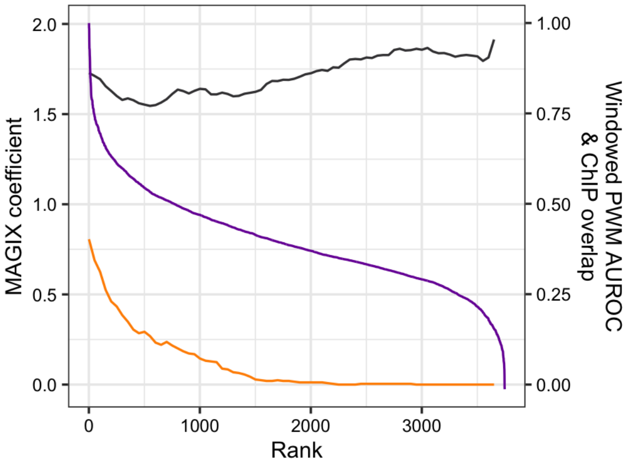

### ZNF250

Top GRECO-BIT/MEX PWM: PWM003634

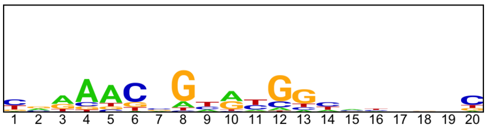


--------------------------------------------------------------------------------------------

Location of top GRECO-BIT/MEX PWM in MAGIX peaks (up to the top 10,000)

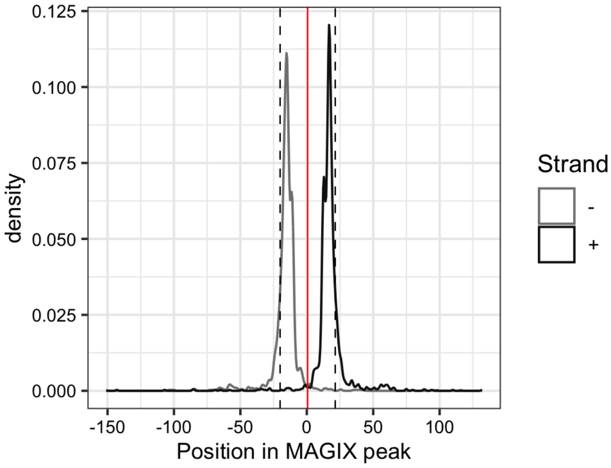


--------------------------------------------------------------------------------------------

AUROC of top GRECO-BIT/MEX PWM in sliding windows (500 peaks, 50 peak step) along the MAGIX peaks (up to the top 75,000) ranked by the MAGIX enrichment coefficient and fraction of overlapping ChIP peaks in the same window

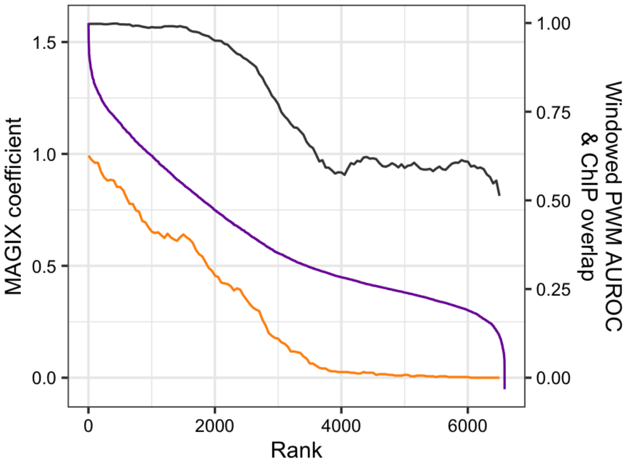

### ZNF251

Top GRECO-BIT/MEX PWM: PWM003657

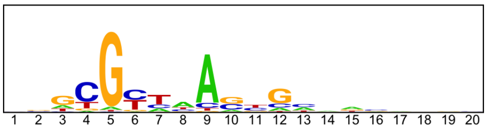


--------------------------------------------------------------------------------------------

Location of top GRECO-BIT/MEX PWM in MAGIX peaks (up to the top 10,000)

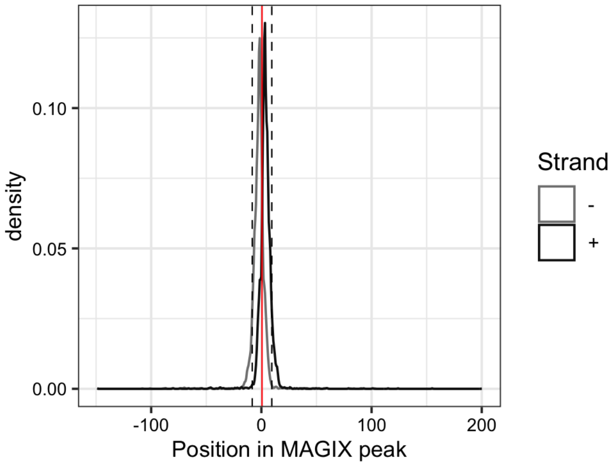


--------------------------------------------------------------------------------------------

AUROC of top GRECO-BIT/MEX PWM in sliding windows (500 peaks, 50 peak step) along the MAGIX peaks (up to the top 75,000) ranked by the MAGIX enrichment coefficient and fraction of overlapping ChIP peaks in the same window

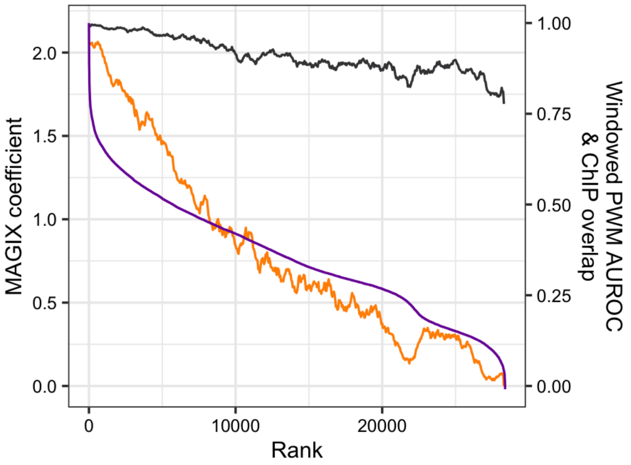

### ZNF260

Top GRECO-BIT/MEX PWM: PWM003683

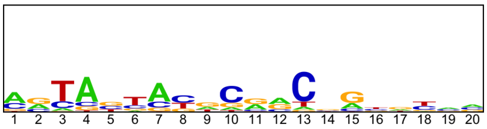


---

Location of top GRECO-BIT/MEX PWM in MAGIX peaks (up to the top 10,000)

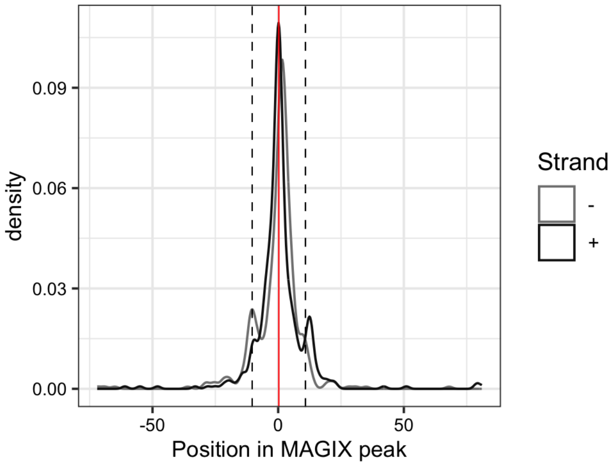


---

AUROC of top GRECO-BIT/MEX PWM in sliding windows (500 peaks, 50 peak step) along the MAGIX peaks (up to the top 75,000) ranked by the MAGIX enrichment coefficient and fraction of overlapping ChIP peaks in the same window

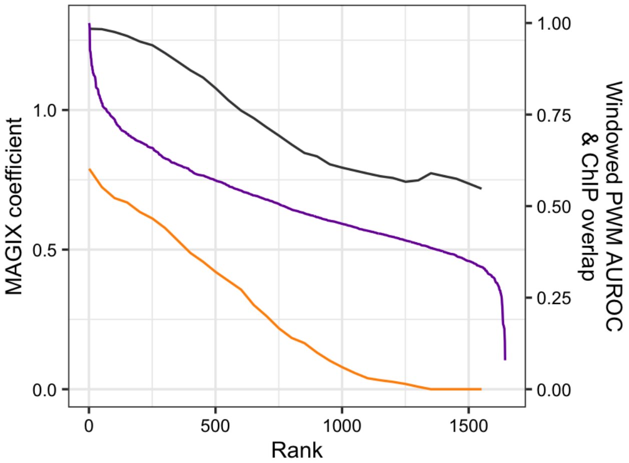

### ZNF264

Top GRECO-BIT/MEX PWM: PWM003714

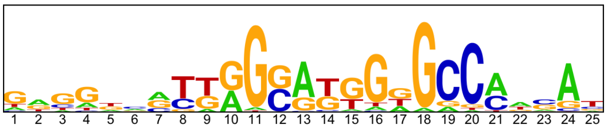

---

Location of top GRECO-BIT/MEX PWM in MAGIX peaks (up to the top 10,000)

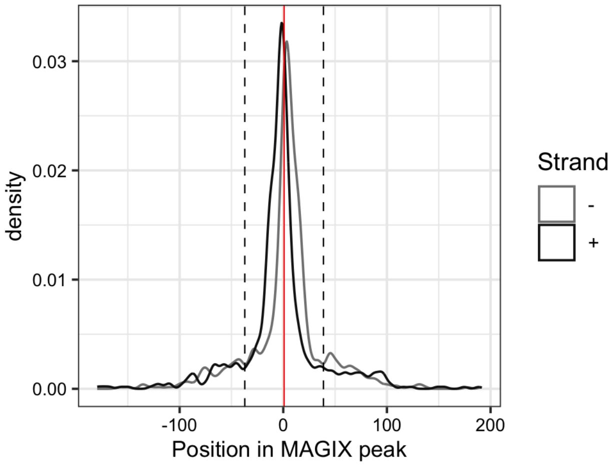


---

AUROC of top GRECO-BIT/MEX PWM in sliding windows (500 peaks, 50 peak step) along the MAGIX peaks (up to the top 75,000) ranked by the MAGIX enrichment coefficient and fraction of overlapping ChIP peaks in the same window

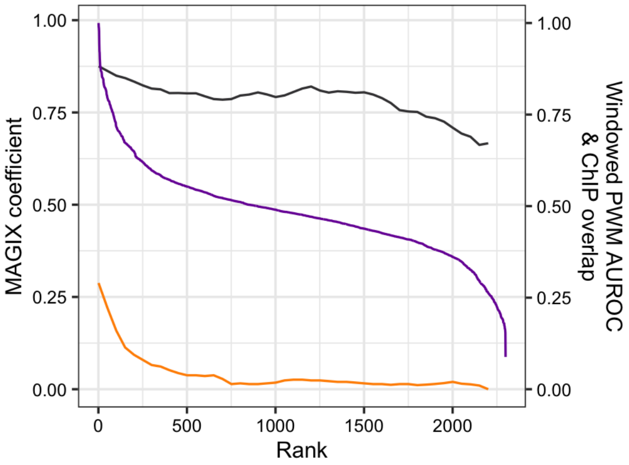

### ZNF275

Top GRECO-BIT/MEX PWM: PWM003734

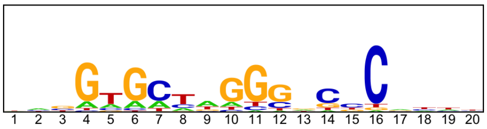


----------------------------------------------------------------------------------------------------

Location of top GRECO-BIT/MEX PWM in MAGIX peaks (up to the top 10,000)

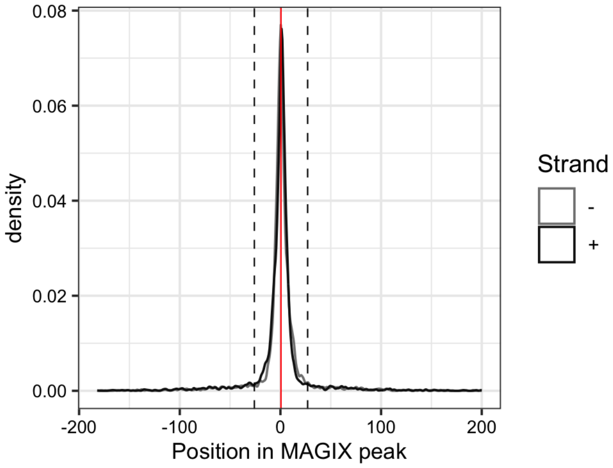

### ZNF286B

Top GRECO-BIT/MEX PWM: PWM003759

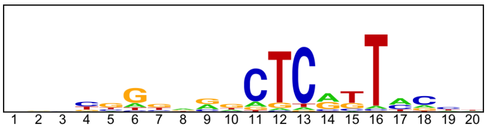


---

Location of top GRECO-BIT/MEX PWM in MAGIX peaks (up to the top 10,000)

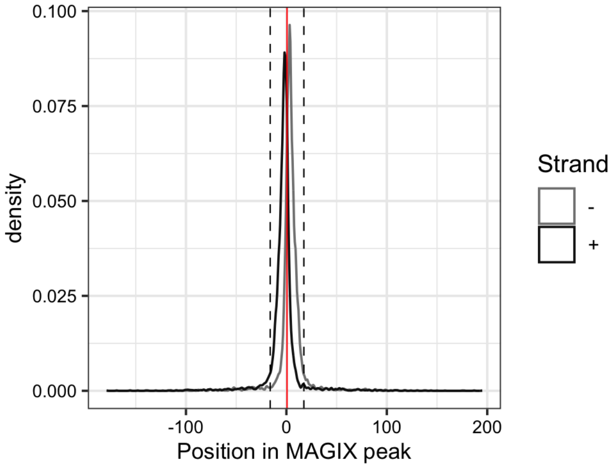


---

AUROC of top GRECO-BIT/MEX PWM in sliding windows (500 peaks, 50 peak step) along the MAGIX peaks (up to the top 75,000) ranked by the MAGIX enrichment coefficient and fraction of overlapping ChIP peaks in the same window

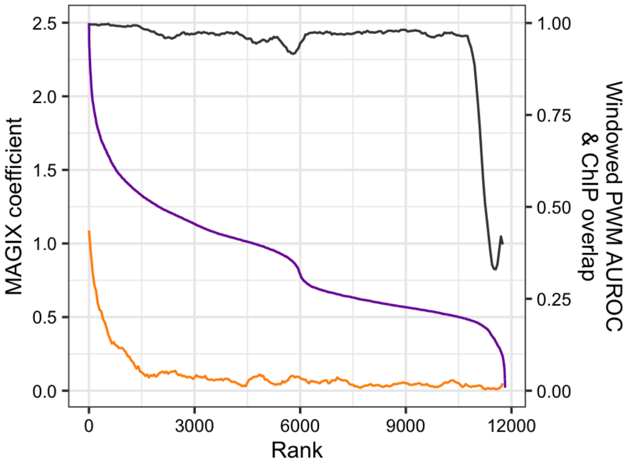

### ZNF292.DBD1

Top GRECO-BIT/MEX PWM: PWM003802

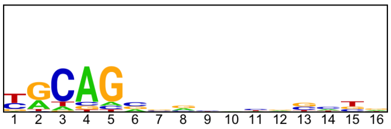


------------------------------------------------------------------------------------------------

Location of top GRECO-BIT/MEX PWM in MAGIX peaks (up to the top 10,000)

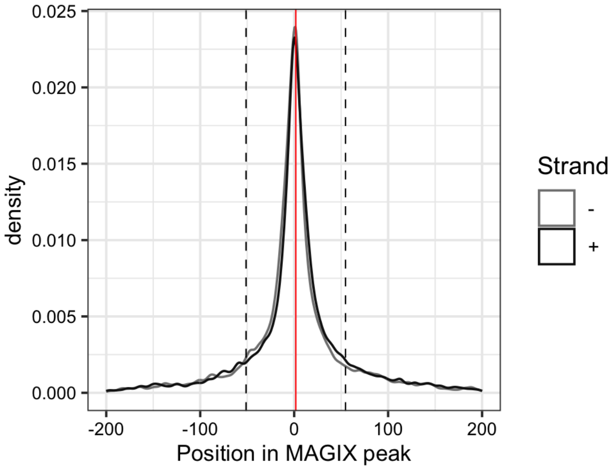

### ZNF292.DBD2

Top GRECO-BIT/MEX PWM: PWM003788

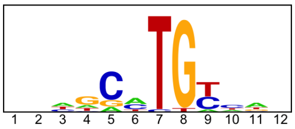


-------------------------------------------------------------------------------------------

Location of top GRECO-BIT/MEX PWM in MAGIX peaks (up to the top 10,000)

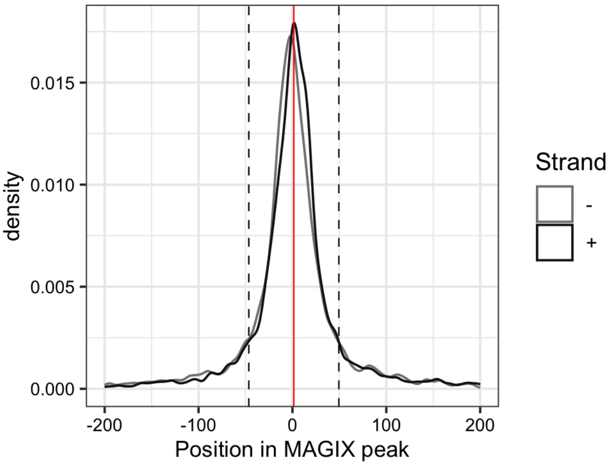

### ZNF322

Top GRECO-BIT/MEX PWM: PWM003869

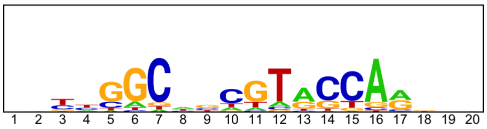


---

Location of top GRECO-BIT/MEX PWM in MAGIX peaks (up to the top 10,000)

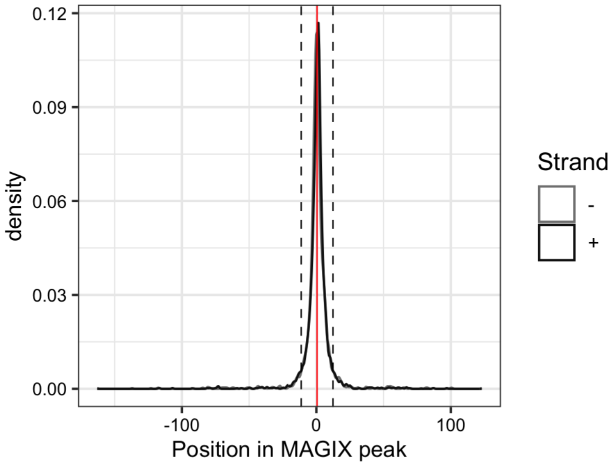


---

AUROC of top GRECO-BIT/MEX PWM in sliding windows (500 peaks, 50 peak step) along the MAGIX peaks (up to the top 75,000) ranked by the MAGIX enrichment coefficient and fraction of overlapping ChIP peaks in the same window

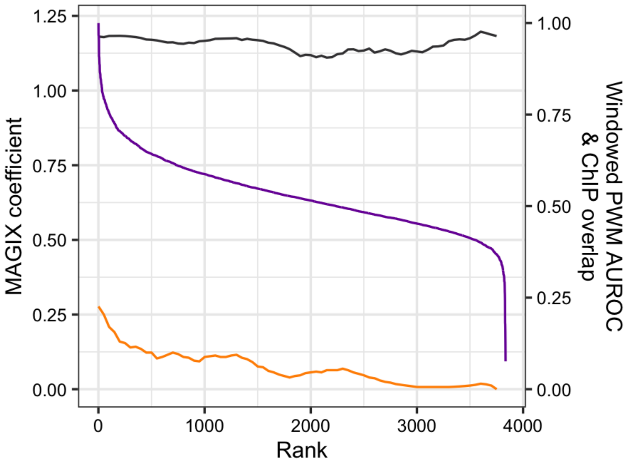

### ZNF335.DBD1

Top GRECO-BIT/MEX PWM: PWM003898

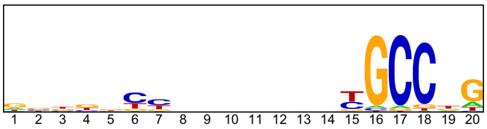


---

Location of top GRECO-BIT/MEX PWM in MAGIX peaks (up to the top 10,000)

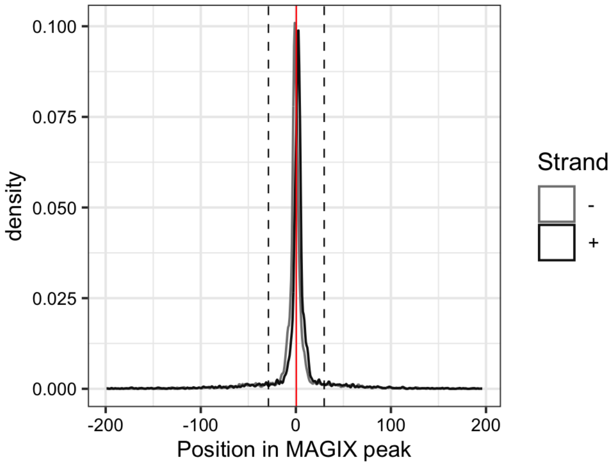

### ZNF335.DBD2

Top GRECO-BIT/MEX PWM: PWM003891

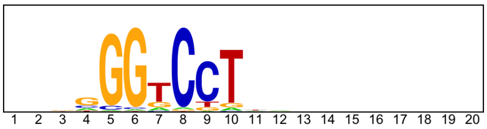


------------------------------------------------------------------------------------------------

Location of top GRECO-BIT/MEX PWM in MAGIX peaks (up to the top 10,000)

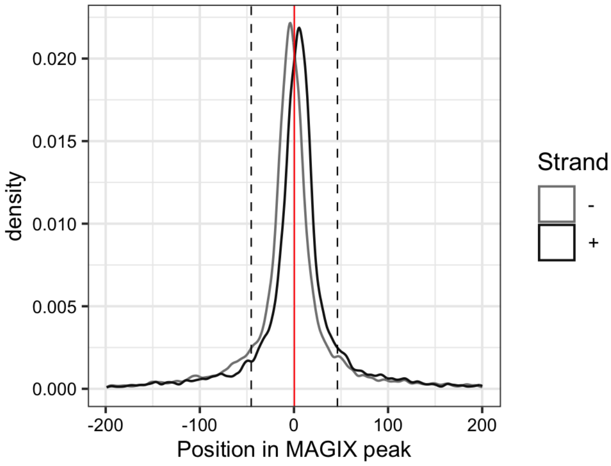

### ZNF347

Top GRECO-BIT/MEX PWM: PWM003926

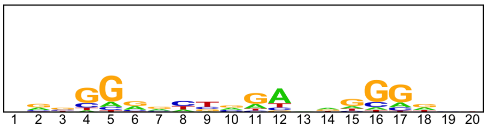


---

Location of top GRECO-BIT/MEX PWM in MAGIX peaks (up to the top 10,000)

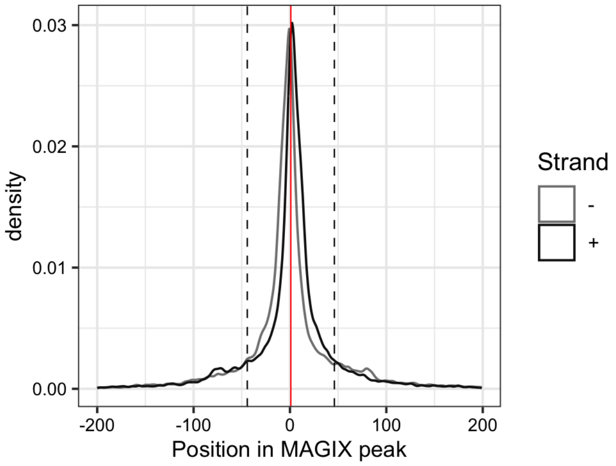


---

AUROC of top GRECO-BIT/MEX PWM in sliding windows (500 peaks, 50 peak step) along the MAGIX peaks (up to the top 75,000) ranked by the MAGIX enrichment coefficient and fraction of overlapping ChIP peaks in the same window

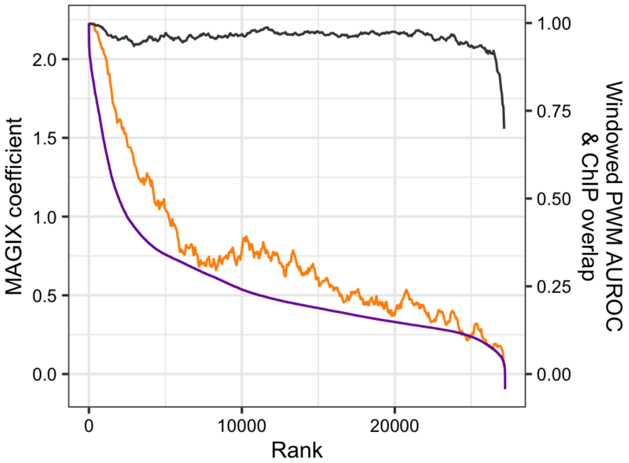

### ZNF35

Top GRECO-BIT/MEX PWM: PWM003953

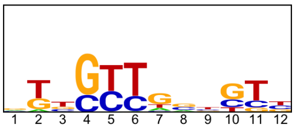


--------------------------------------------------------------------------------------------

Location of top GRECO-BIT/MEX PWM in MAGIX peaks (up to the top 10,000)

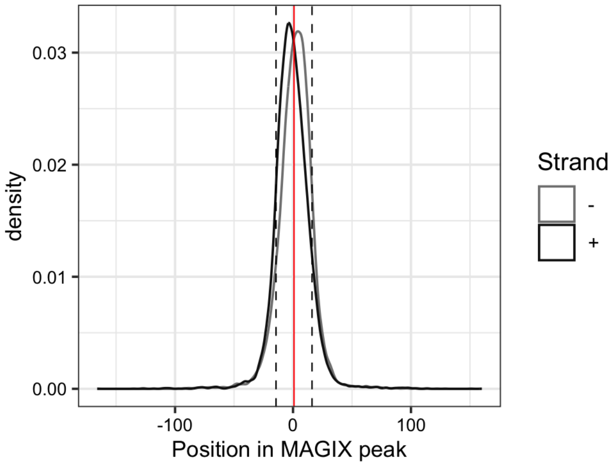


--------------------------------------------------------------------------------------------

AUROC of top GRECO-BIT/MEX PWM in sliding windows (500 peaks, 50 peak step) along the MAGIX peaks (up to the top 75,000) ranked by the MAGIX enrichment coefficient and fraction of overlapping ChIP peaks in the same window

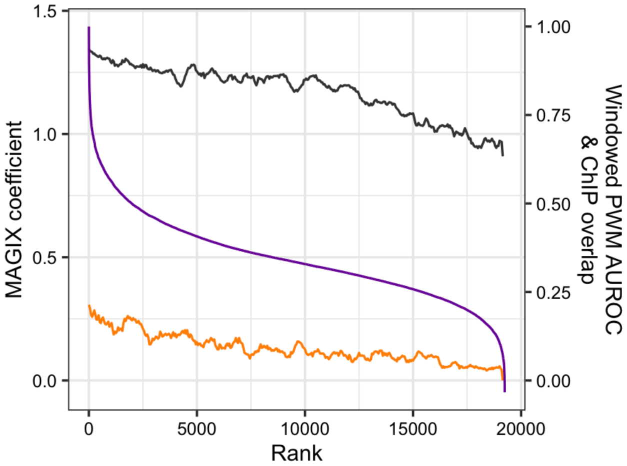

#### ZNF358

Top GRECO-BIT/MEX PWM: PWM003990

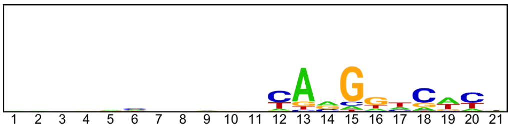

--------------------------------------------------------------------------------------------

Location of top GRECO-BIT/MEX PWM in MAGIX peaks (up to the top 10,000)

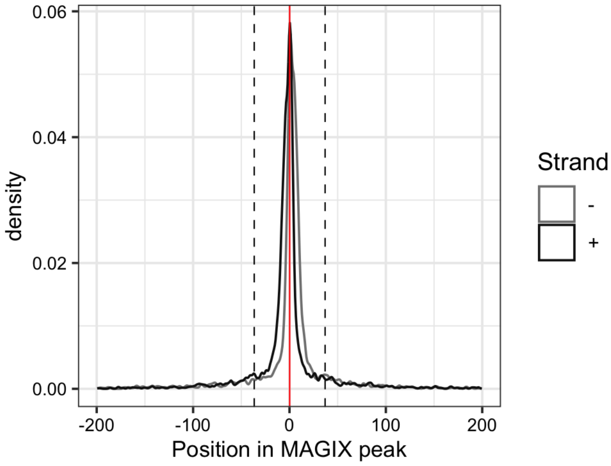

### ZNF362

Top GRECO-BIT/MEX PWM: PWM004037

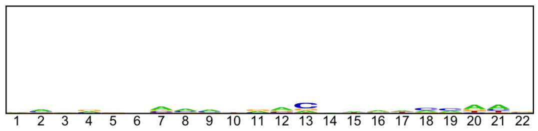

---

Location of top GRECO-BIT/MEX PWM in MAGIX peaks (up to the top 10,000)

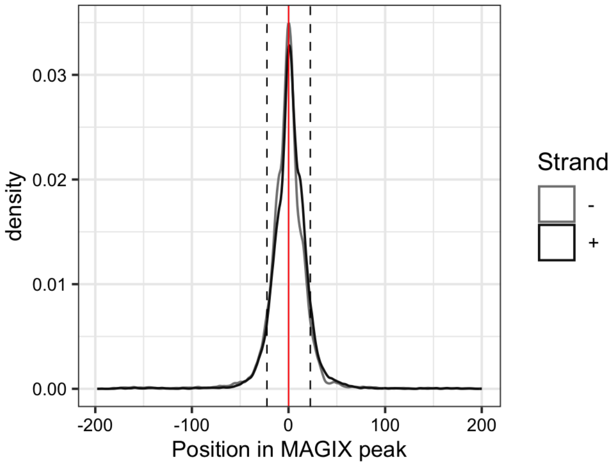


---

AUROC of top GRECO-BIT/MEX PWM in sliding windows (500 peaks, 50 peak step) along the MAGIX peaks (up to the top 75,000) ranked by the MAGIX enrichment coefficient and fraction of overlapping ChIP peaks in the same window

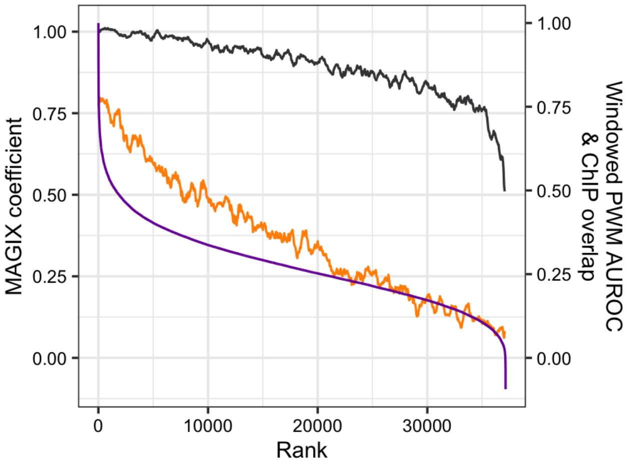

### ZNF367

Top GRECO-BIT/MEX PWM: PWM004062

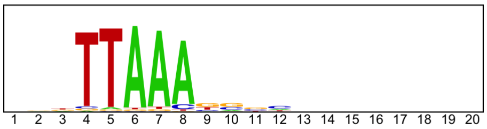


-----------------------------------------------------------------------------------------------

Location of top GRECO-BIT/MEX PWM in MAGIX peaks (up to the top 10,000)

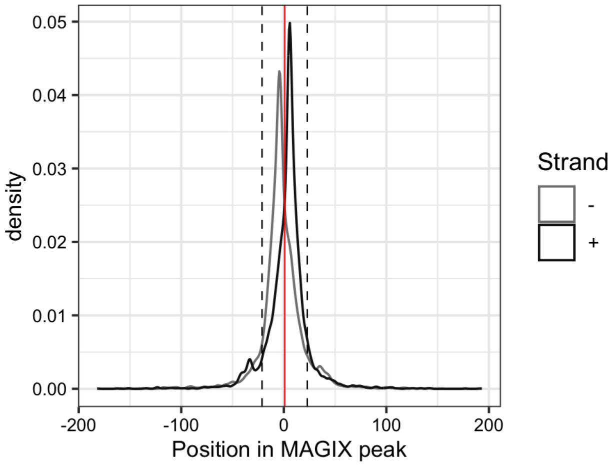


-----------------------------------------------------------------------------------------------

AUROC of top GRECO-BIT/MEX PWM in sliding windows (500 peaks, 50 peak step) along the MAGIX peaks (up to the top 75,000) ranked by the MAGIX enrichment coefficient and fraction of overlapping ChIP peaks in the same window

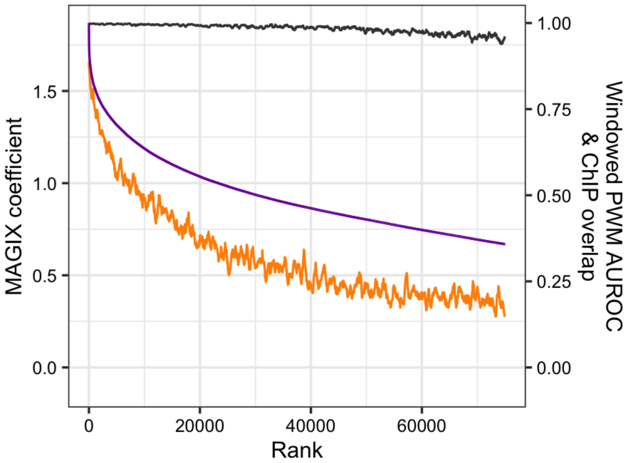

### ZNF384

Top GRECO-BIT/MEX PWM: PWM004120

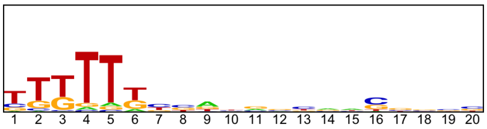


----------------------------------------------------------------------------------------------------

Location of top GRECO-BIT/MEX PWM in MAGIX peaks (up to the top 10,000)

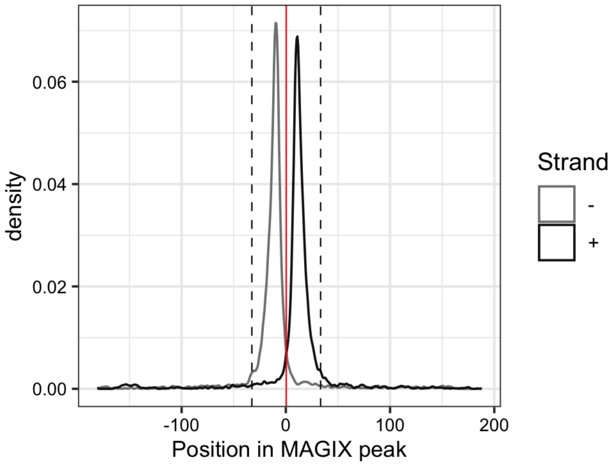


----------------------------------------------------------------------------------------------------

AUROC of top GRECO-BIT/MEX PWM in sliding windows (500 peaks, 50 peak step) along the MAGIX peaks (up to the top 75,000) ranked by the MAGIX enrichment coefficient and fraction of overlapping ChIP peaks in the same window

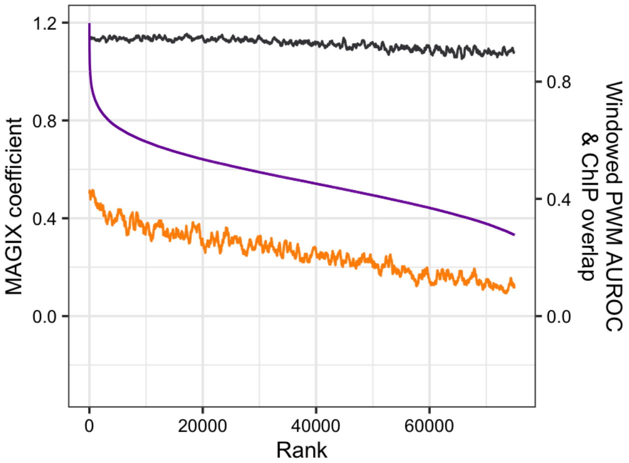

### ZNF395

Top GRECO-BIT/MEX PWM: PWM004151

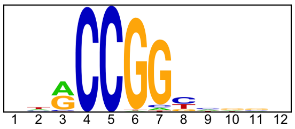


--------------------------------------------------------------------------------------------

Location of top GRECO-BIT/MEX PWM in MAGIX peaks (up to the top 10,000)

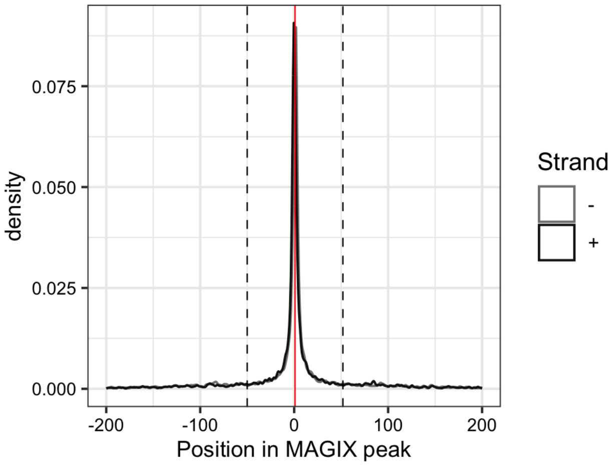


--------------------------------------------------------------------------------------------

AUROC of top GRECO-BIT/MEX PWM in sliding windows (500 peaks, 50 peak step) along the MAGIX peaks (up to the top 75,000) ranked by the MAGIX enrichment coefficient and fraction of overlapping ChIP peaks in the same window

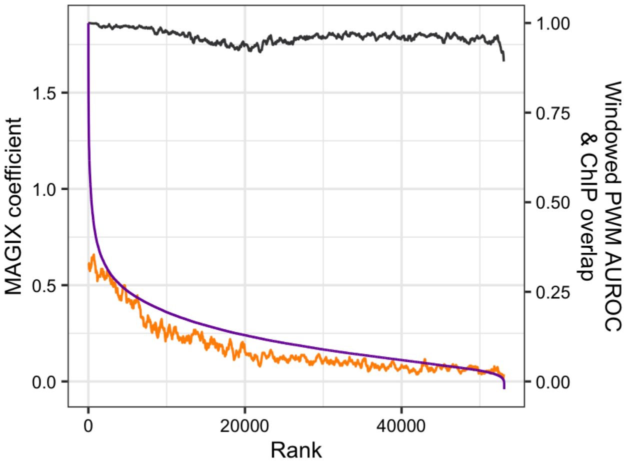

### ZNF407

Top GRECO-BIT/MEX PWM: PWM004189

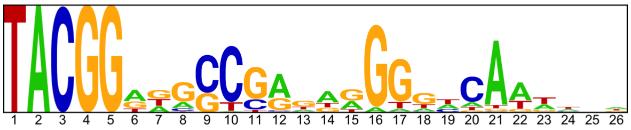


----------------------------------------------------------------------------------------------------

Location of top GRECO-BIT/MEX PWM in MAGIX peaks (up to the top 10,000)

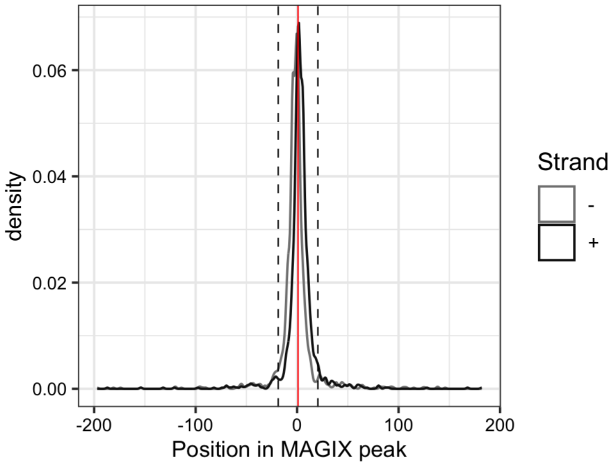


----------------------------------------------------------------------------------------------------

AUROC of top GRECO-BIT/MEX PWM in sliding windows (500 peaks, 50 peak step) along the MAGIX peaks (up to the top 75,000) ranked by the MAGIX enrichment coefficient and fraction of overlapping ChIP peaks in the same window

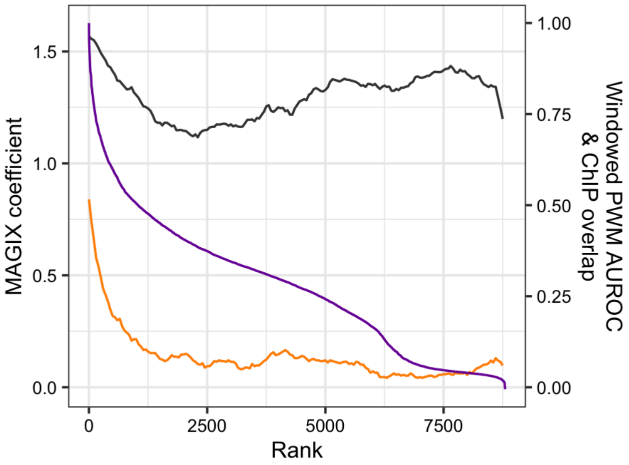

### ZNF43.DBD1

Top GRECO-BIT/MEX PWM: PWM004198

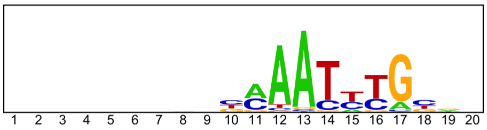

--------------------------------------------------------------------------------

Location of top GRECO-BIT/MEX PWM in MAGIX peaks (up to the top 10,000)

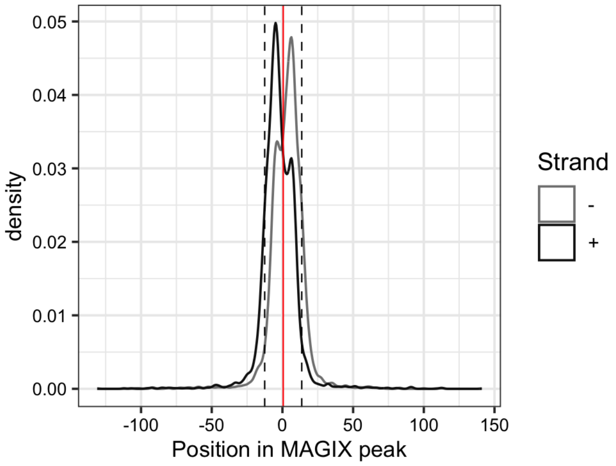

### ZNF43.DBD2

Top GRECO-BIT/MEX PWM: PWM004215

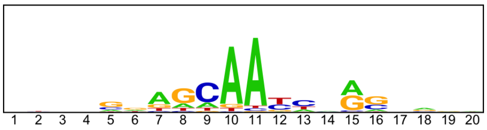


------------------------------------------------------------------------------------------------

Location of top GRECO-BIT/MEX PWM in MAGIX peaks (up to the top 10,000)

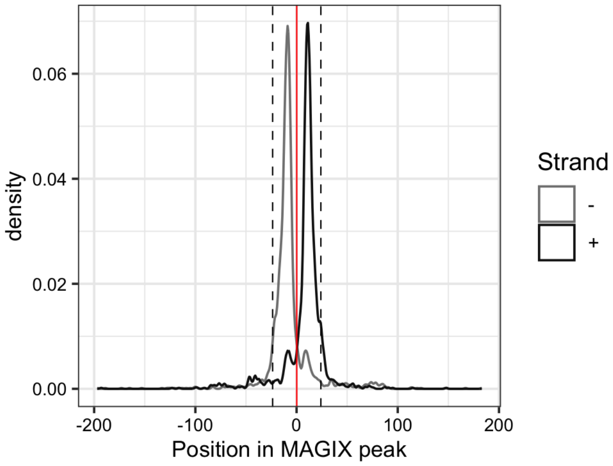

### ZNF43.FL

Top GRECO-BIT/MEX PWM: PWM004214

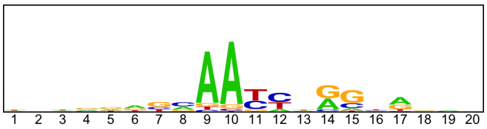


----------------------------------------------------------------------------------------------------

Location of top GRECO-BIT/MEX PWM in MAGIX peaks (up to the top 10,000)

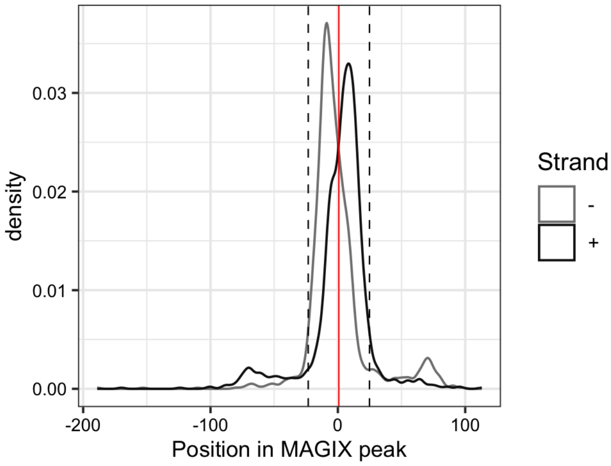


----------------------------------------------------------------------------------------------------

AUROC of top GRECO-BIT/MEX PWM in sliding windows (500 peaks, 50 peak step) along the MAGIX peaks (up to the top 75,000) ranked by the MAGIX enrichment coefficient and fraction of overlapping ChIP peaks in the same window

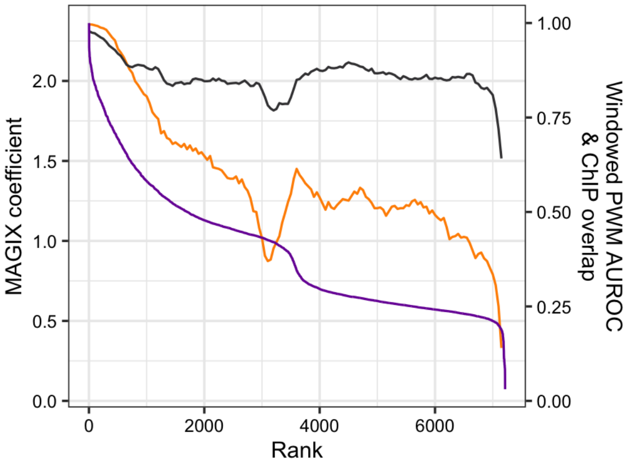

### ZNF436

Top GRECO-BIT/MEX PWM: PWM004243

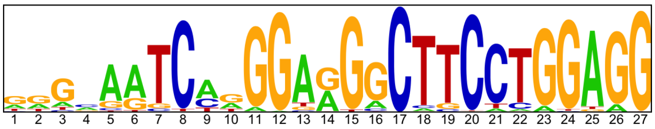


-----------------------------------------------------------------------------------------------

Location of top GRECO-BIT/MEX PWM in MAGIX peaks (up to the top 10,000)

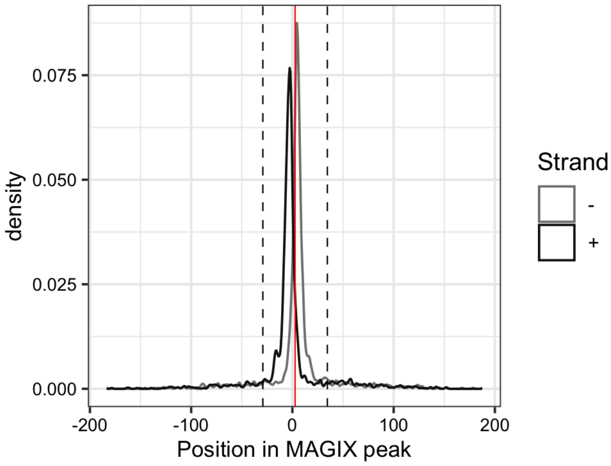


-----------------------------------------------------------------------------------------------

AUROC of top GRECO-BIT/MEX PWM in sliding windows (500 peaks, 50 peak step) along the MAGIX peaks (up to the top 75,000) ranked by the MAGIX enrichment coefficient and fraction of overlapping ChIP peaks in the same window

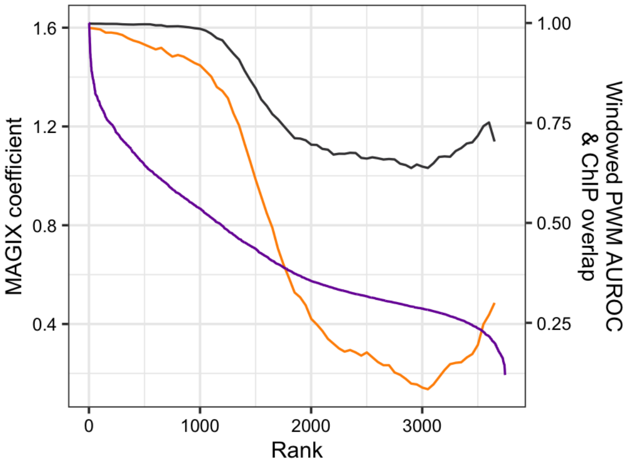

### ZNF470

Top GRECO-BIT/MEX PWM: PWM004282

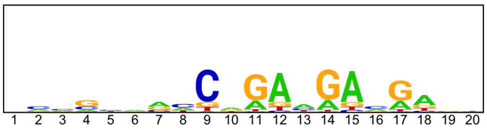

---

Location of top GRECO-BIT/MEX PWM in MAGIX peaks (up to the top 10,000)

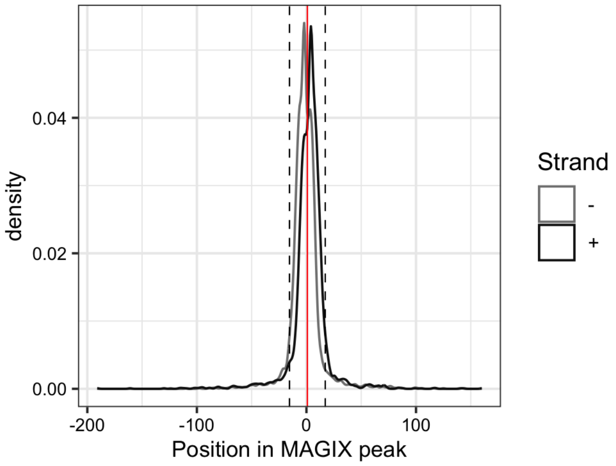


---

AUROC of top GRECO-BIT/MEX PWM in sliding windows (500 peaks, 50 peak step) along the MAGIX peaks (up to the top 75,000) ranked by the MAGIX enrichment coefficient and fraction of overlapping ChIP peaks in the same window

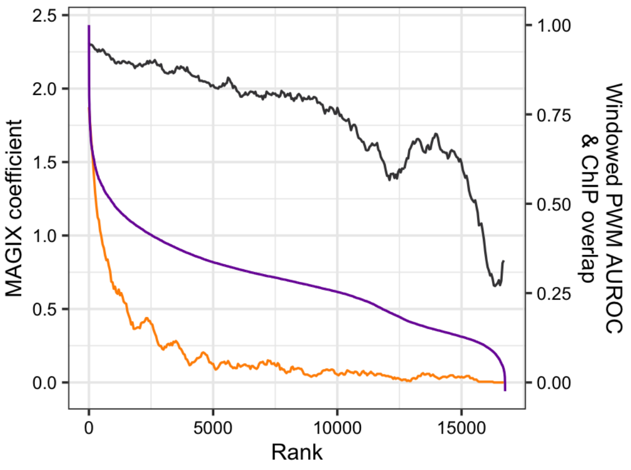

### ZNF471

Top GRECO-BIT/MEX PWM: PWM004337

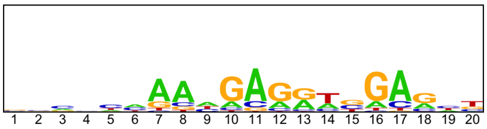


---

Location of top GRECO-BIT/MEX PWM in MAGIX peaks (up to the top 10,000)

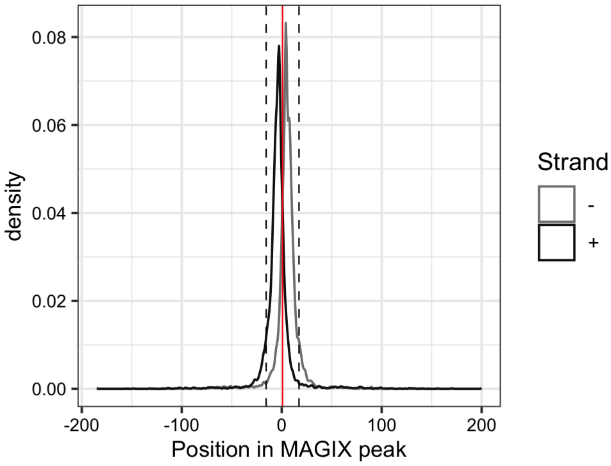


---

AUROC of top GRECO-BIT/MEX PWM in sliding windows (500 peaks, 50 peak step) along the MAGIX peaks (up to the top 75,000) ranked by the MAGIX enrichment coefficient and fraction of overlapping ChIP peaks in the same window

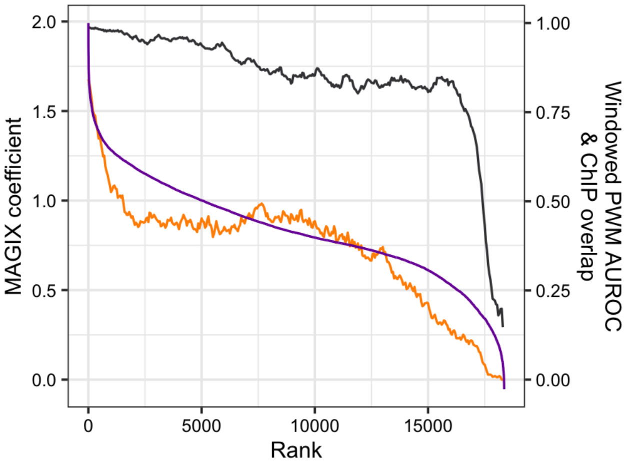

### ZNF48

Top GRECO-BIT/MEX PWM: PWM004344

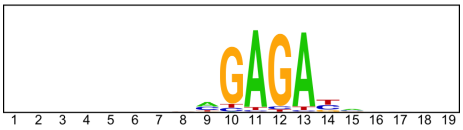


---

Location of top GRECO-BIT/MEX PWM in MAGIX peaks (up to the top 10,000)

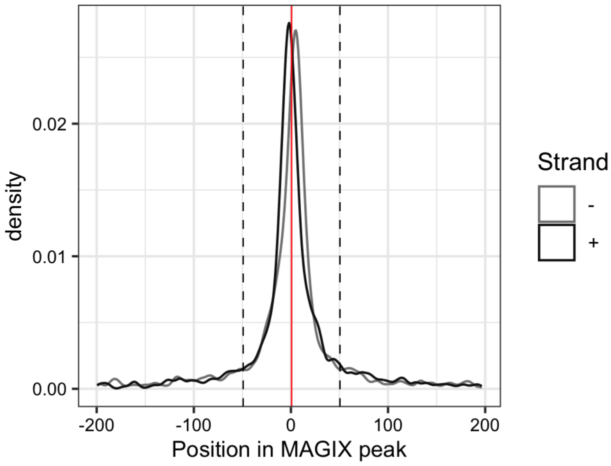


---

AUROC of top GRECO-BIT/MEX PWM in sliding windows (500 peaks, 50 peak step) along the MAGIX peaks (up to the top 75,000) ranked by the MAGIX enrichment coefficient and fraction of overlapping ChIP peaks in the same window

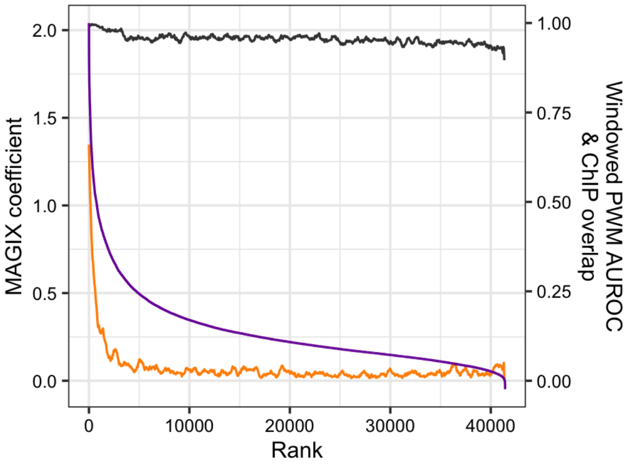

### ZNF493.DBD2

Top GRECO-BIT/MEX PWM: PWM004415

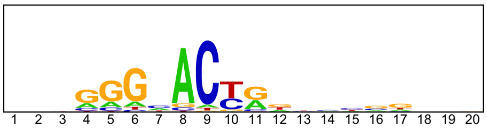

---

Location of top GRECO-BIT/MEX PWM in MAGIX peaks (up to the top 10,000)

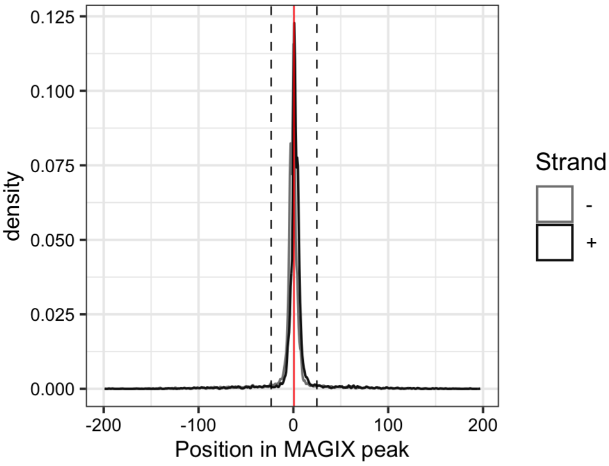

### ZNF493.FL

Top GRECO-BIT/MEX PWM: PWM004411

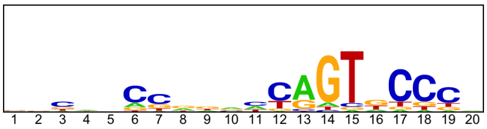


-----------------------------------------------------------------------------------------------

Location of top GRECO-BIT/MEX PWM in MAGIX peaks (up to the top 10,000)

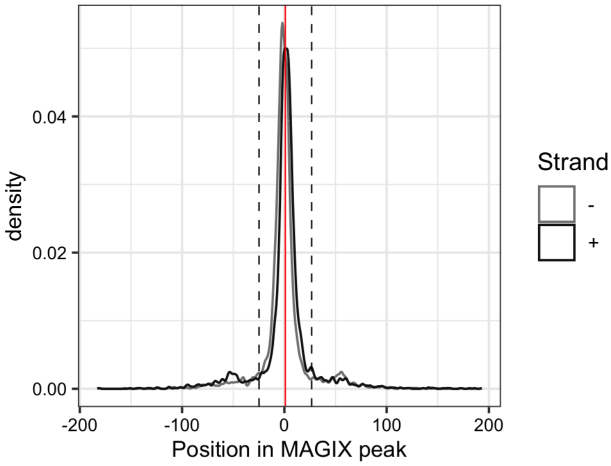


-----------------------------------------------------------------------------------------------

AUROC of top GRECO-BIT/MEX PWM in sliding windows (500 peaks, 50 peak step) along the MAGIX peaks (up to the top 75,000) ranked by the MAGIX enrichment coefficient and fraction of overlapping ChIP peaks in the same window

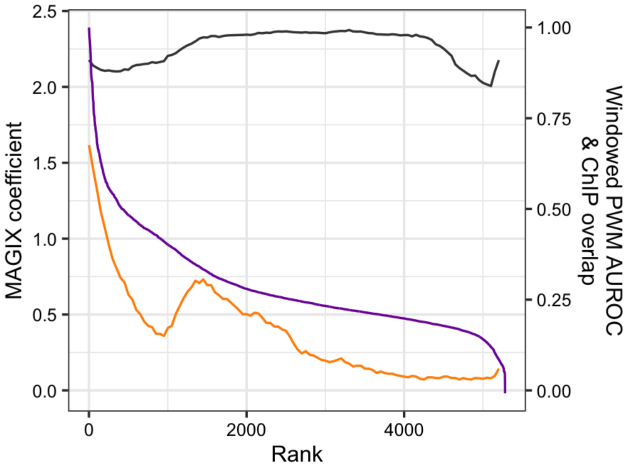

### ZNF497

Top GRECO-BIT/MEX PWM: PWM004430

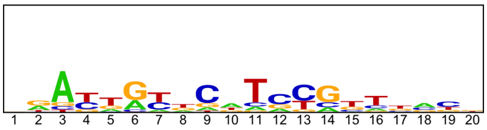


----------------------------------------------------------------------------------------------------

Location of top GRECO-BIT/MEX PWM in MAGIX peaks (up to the top 10,000)

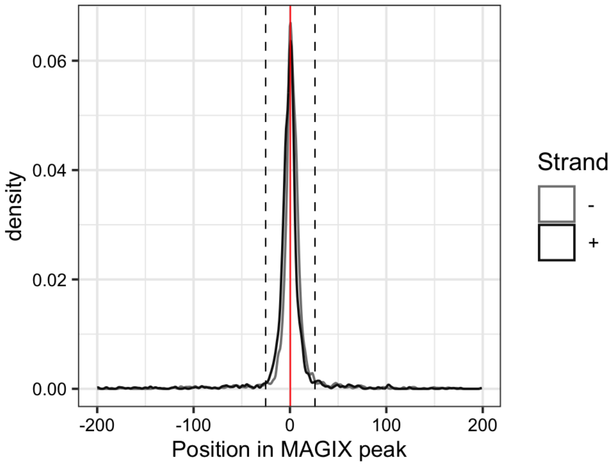

### ZNF500

Top GRECO-BIT/MEX PWM: PWM004463

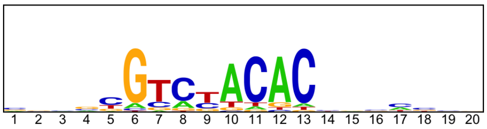


-----------------------------------------------------------------------------------------------

Location of top GRECO-BIT/MEX PWM in MAGIX peaks (up to the top 10,000)

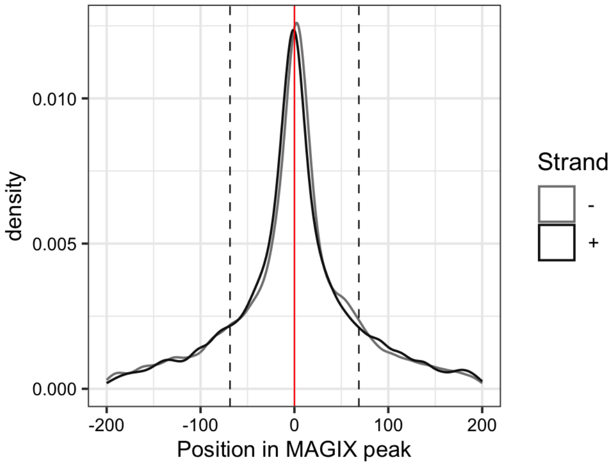


-----------------------------------------------------------------------------------------------

AUROC of top GRECO-BIT/MEX PWM in sliding windows (500 peaks, 50 peak step) along the MAGIX peaks (up to the top 75,000) ranked by the MAGIX enrichment coefficient and fraction of overlapping ChIP peaks in the same window

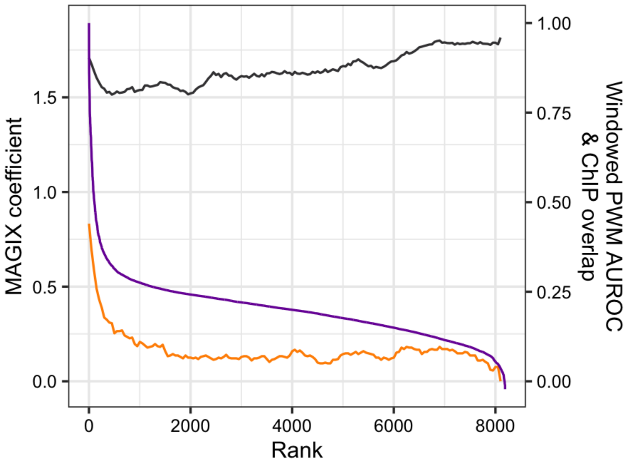

### ZNF510

Top GRECO-BIT/MEX PWM: PWM004519

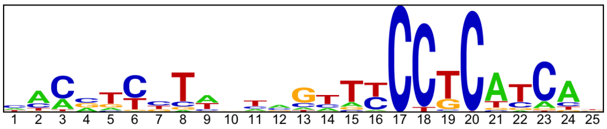

---

Location of top GRECO-BIT/MEX PWM in MAGIX peaks (up to the top 10,000)

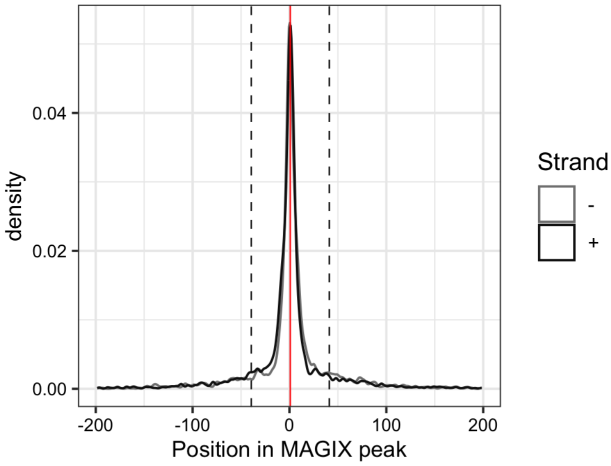


---

AUROC of top GRECO-BIT/MEX PWM in sliding windows (500 peaks, 50 peak step) along the MAGIX peaks (up to the top 75,000) ranked by the MAGIX enrichment coefficient and fraction of overlapping ChIP peaks in the same window

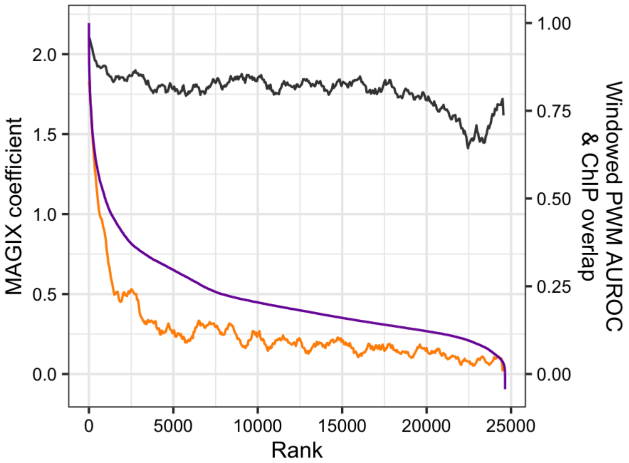

### ZNF518B

Top GRECO-BIT/MEX PWM: PWM004578

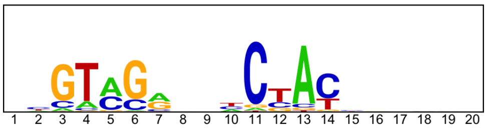


-------------------------------------------------------------------------------------------

Location of top GRECO-BIT/MEX PWM in MAGIX peaks (up to the top 10,000)

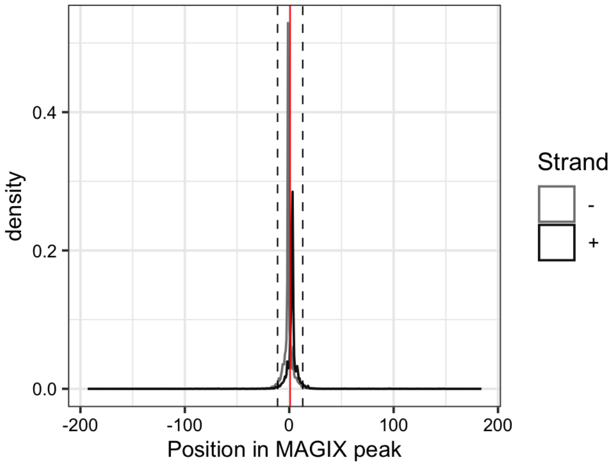


-------------------------------------------------------------------------------------------

AUROC of top GRECO-BIT/MEX PWM in sliding windows (500 peaks, 50 peak step) along the MAGIX peaks (up to the top 75,000) ranked by the MAGIX enrichment coefficient and fraction of overlapping ChIP peaks in the same window

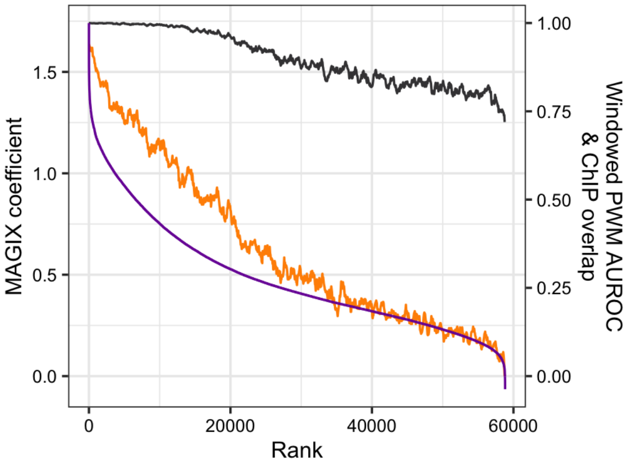

### ZNF532.DBD2

Top GRECO-BIT/MEX PWM: PWM004588

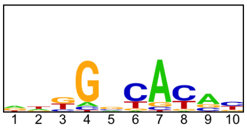


-----------------------------------------------------------------------------------------------

Location of top GRECO-BIT/MEX PWM in MAGIX peaks (up to the top 10,000)

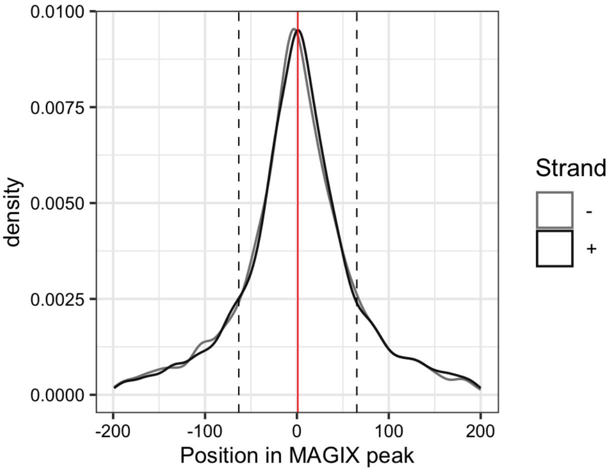

### ZNF536.DBD2

Top GRECO-BIT/MEX PWM: PWM004608

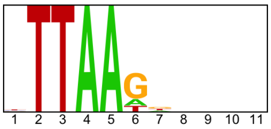


------------------------------------------------------------------------------------------------

Location of top GRECO-BIT/MEX PWM in MAGIX peaks (up to the top 10,000)

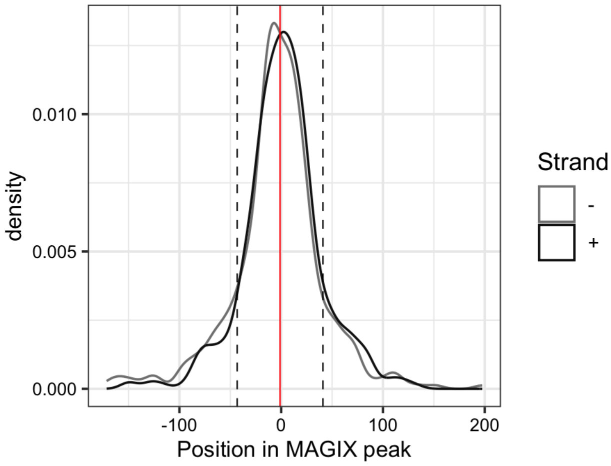

### ZNF551

Top GRECO-BIT/MEX PWM: PWM004658

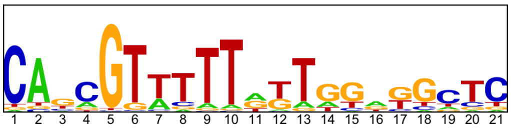


--------------------------------------------------------------------------------------------

Location of top GRECO-BIT/MEX PWM in MAGIX peaks (up to the top 10,000)

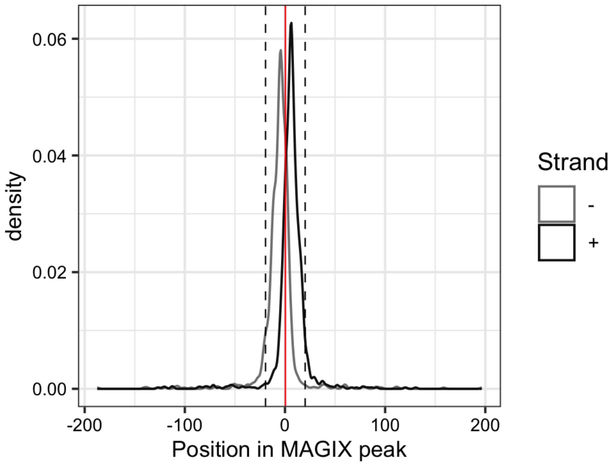


--------------------------------------------------------------------------------------------

AUROC of top GRECO-BIT/MEX PWM in sliding windows (500 peaks, 50 peak step) along the MAGIX peaks (up to the top 75,000) ranked by the MAGIX enrichment coefficient and fraction of overlapping ChIP peaks in the same window

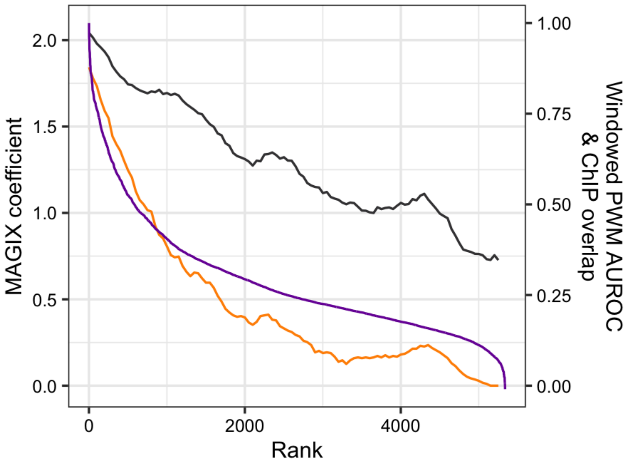

### ZNF568

Top GRECO-BIT/MEX PWM: PWM004669

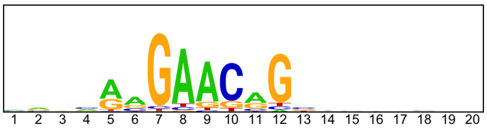


-----------------------------------------------------------------------------------------------

Location of top GRECO-BIT/MEX PWM in MAGIX peaks (up to the top 10,000)

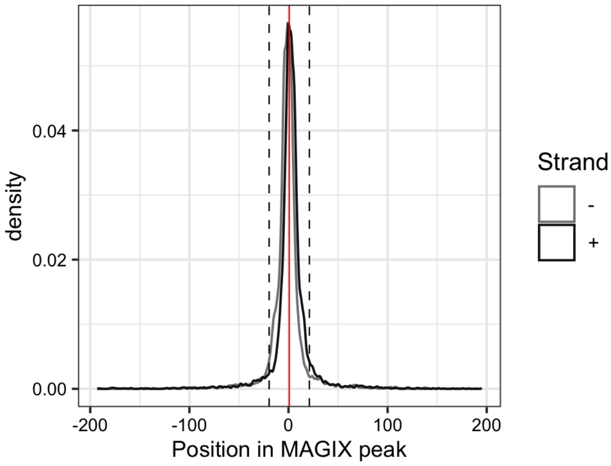

### ZNF569

Top GRECO-BIT/MEX PWM: PWM004689

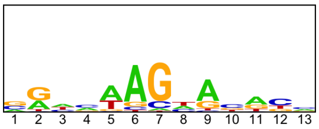


--------------------------------------------------------------------------------------------

Location of top GRECO-BIT/MEX PWM in MAGIX peaks (up to the top 10,000)

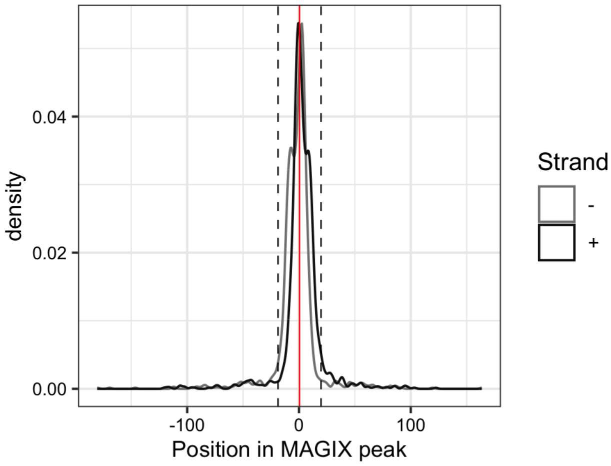

### ZNF57

Top GRECO-BIT/MEX PWM: PWM004716

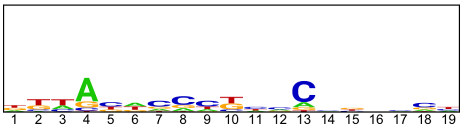


------------------------------------------------------------------------------------------

Location of top GRECO-BIT/MEX PWM in MAGIX peaks (up to the top 10,000)

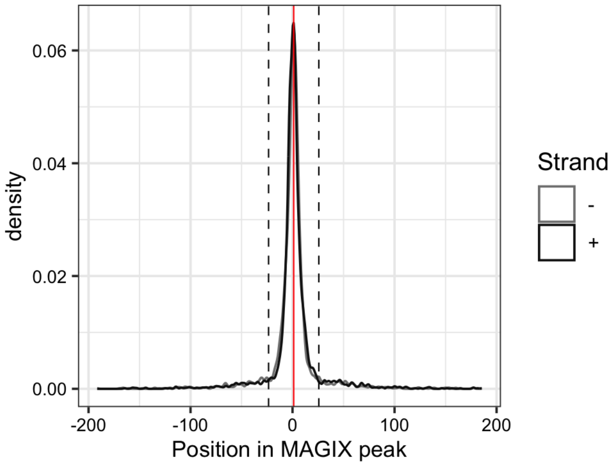


------------------------------------------------------------------------------------------

AUROC of top GRECO-BIT/MEX PWM in sliding windows (500 peaks, 50 peak step) along the MAGIX peaks (up to the top 75,000) ranked by the MAGIX enrichment coefficient and fraction of overlapping ChIP peaks in the same window

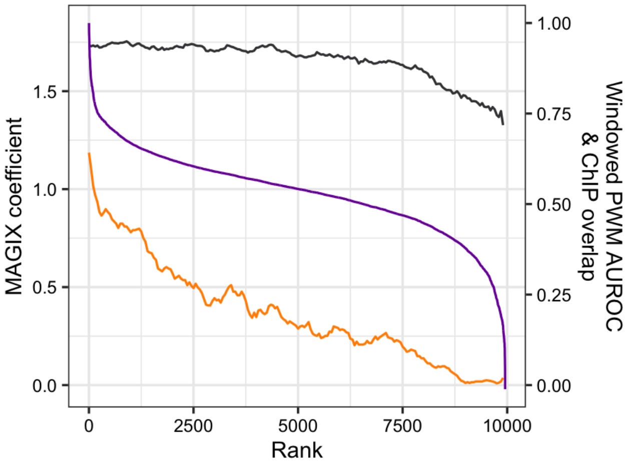

### ZNF575

Top GRECO-BIT/MEX PWM: PWM004745

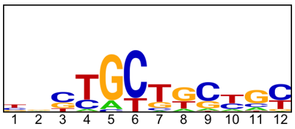


---

Location of top GRECO-BIT/MEX PWM in MAGIX peaks (up to the top 10,000)

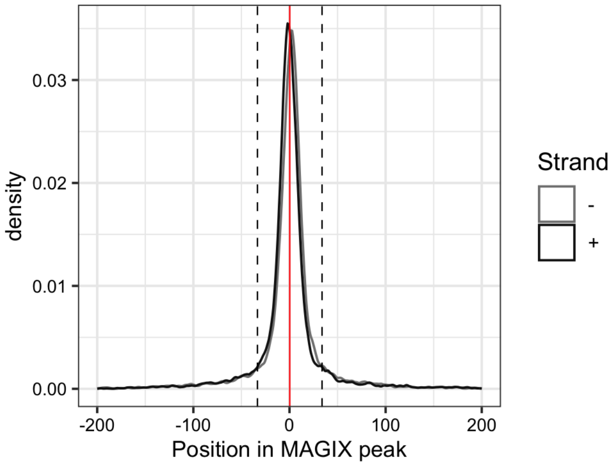


---

AUROC of top GRECO-BIT/MEX PWM in sliding windows (500 peaks, 50 peak step) along the MAGIX peaks (up to the top 75,000) ranked by the MAGIX enrichment coefficient and fraction of overlapping ChIP peaks in the same window

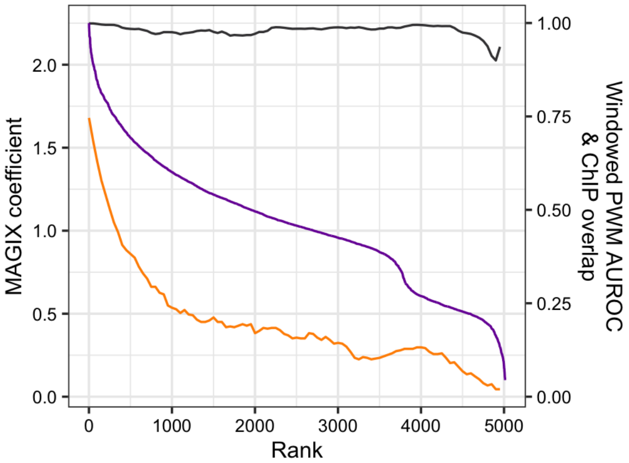

### ZNF587B

Top GRECO-BIT/MEX PWM: PWM004785

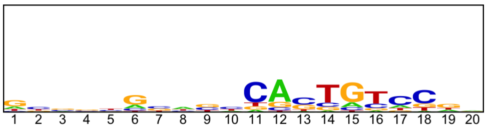


-----------------------------------------------------------------------------------------------

Location of top GRECO-BIT/MEX PWM in MAGIX peaks (up to the top 10,000)

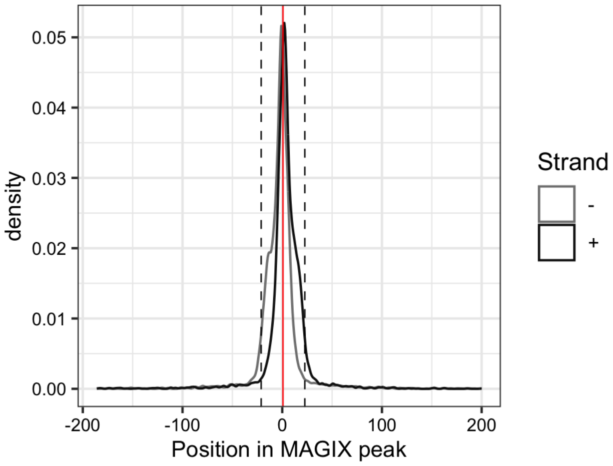

### ZNF596

Top GRECO-BIT/MEX PWM: PWM004806

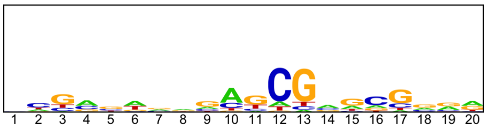


-----------------------------------------------------------------------------------------------

Location of top GRECO-BIT/MEX PWM in MAGIX peaks (up to the top 10,000)

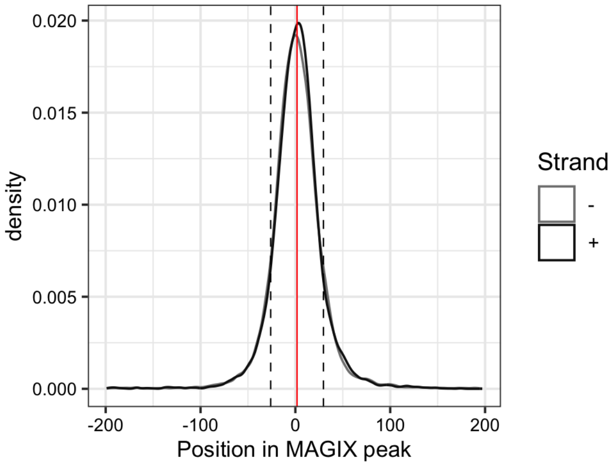


-----------------------------------------------------------------------------------------------

AUROC of top GRECO-BIT/MEX PWM in sliding windows (500 peaks, 50 peak step) along the MAGIX peaks (up to the top 75,000) ranked by the MAGIX enrichment coefficient and fraction of overlapping ChIP peaks in the same window

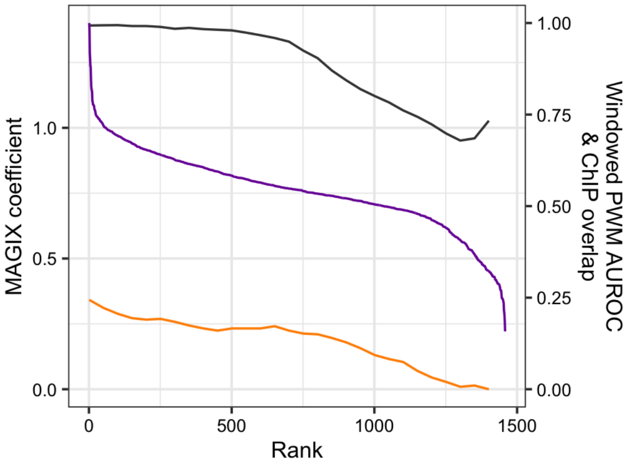

### ZNF606

Top GRECO-BIT/MEX PWM: PWM004837

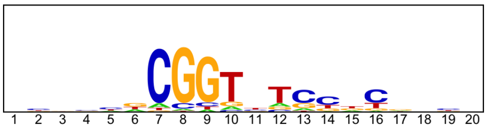


----------------------------------------------------------------------------------------------------

Location of top GRECO-BIT/MEX PWM in MAGIX peaks (up to the top 10,000)

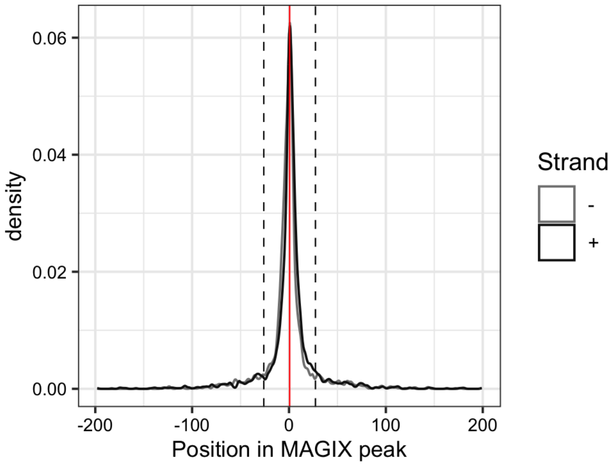


----------------------------------------------------------------------------------------------------

AUROC of top GRECO-BIT/MEX PWM in sliding windows (500 peaks, 50 peak step) along the MAGIX peaks (up to the top 75,000) ranked by the MAGIX enrichment coefficient and fraction of overlapping ChIP peaks in the same window

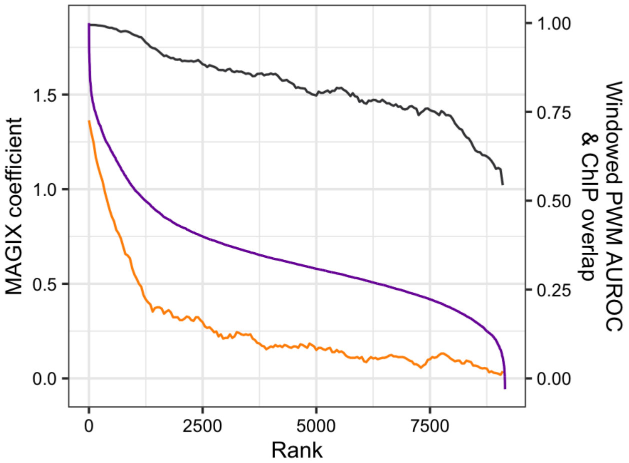

### ZNF618

Top GRECO-BIT/MEX PWM: PWM004873

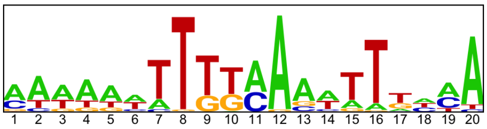


--------------------------------------------------------------------------------------------

Location of top GRECO-BIT/MEX PWM in MAGIX peaks (up to the top 10,000)

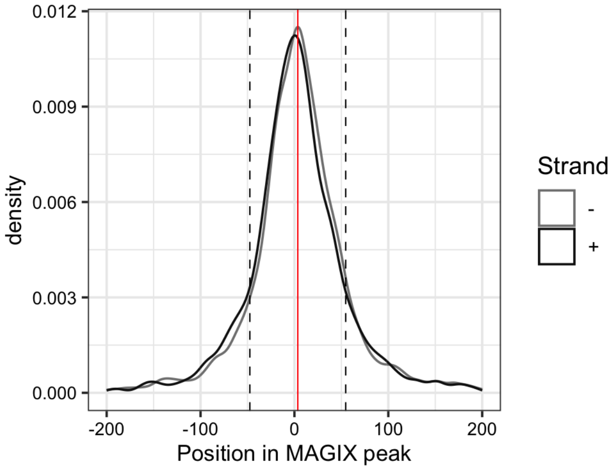

### ZNF623

Top GRECO-BIT/MEX PWM: PWM004915

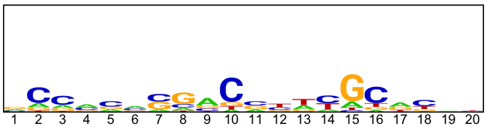


---

Location of top GRECO-BIT/MEX PWM in MAGIX peaks (up to the top 10,000)

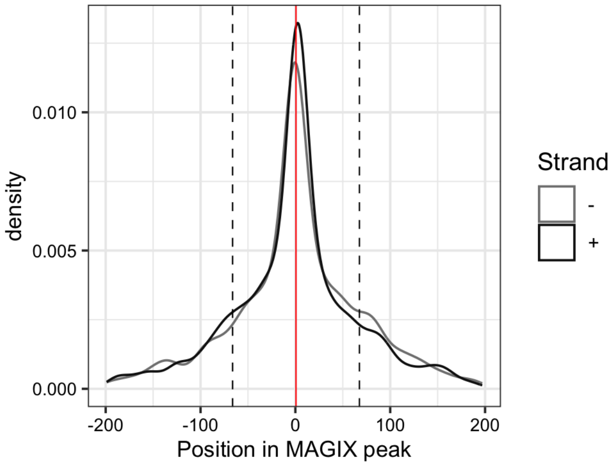


---

AUROC of top GRECO-BIT/MEX PWM in sliding windows (500 peaks, 50 peak step) along the MAGIX peaks (up to the top 75,000) ranked by the MAGIX enrichment coefficient and fraction of overlapping ChIP peaks in the same window

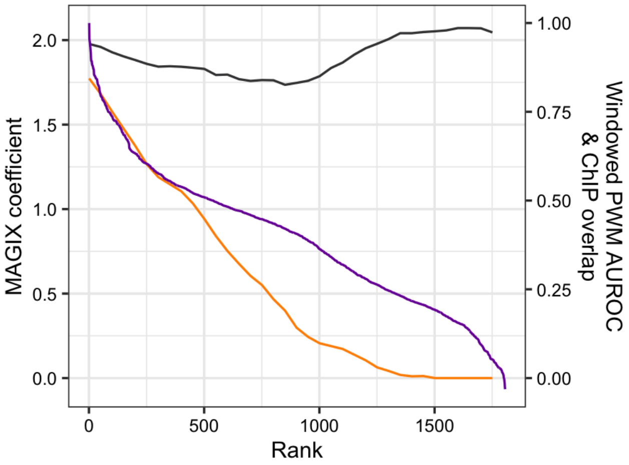

### ZNF646.DBD1

Top GRECO-BIT/MEX PWM: PWM163370

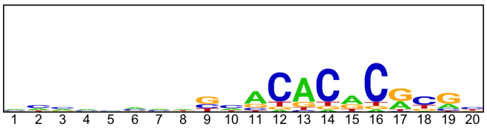

-----------------------------------------------------------------------------------------------

Location of top GRECO-BIT/MEX PWM in MAGIX peaks (up to the top 10,000)

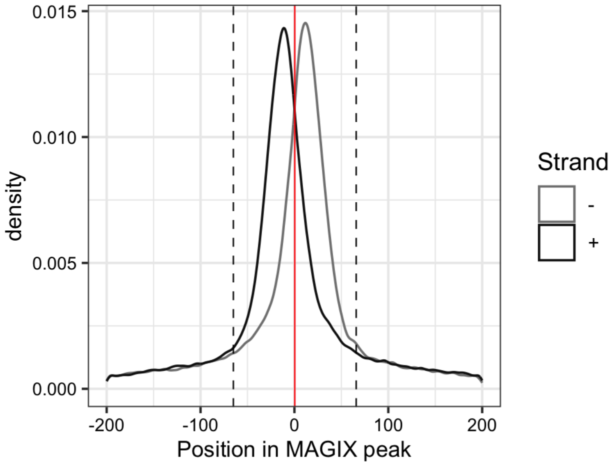

### ZNF646.DBD2

Top GRECO-BIT/MEX PWM: PWM163643

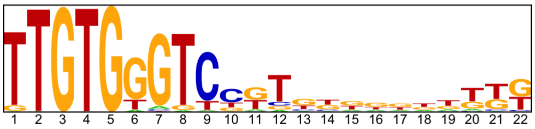


--------------------------------------------------------------------------------------------

Location of top GRECO-BIT/MEX PWM in MAGIX peaks (up to the top 10,000)

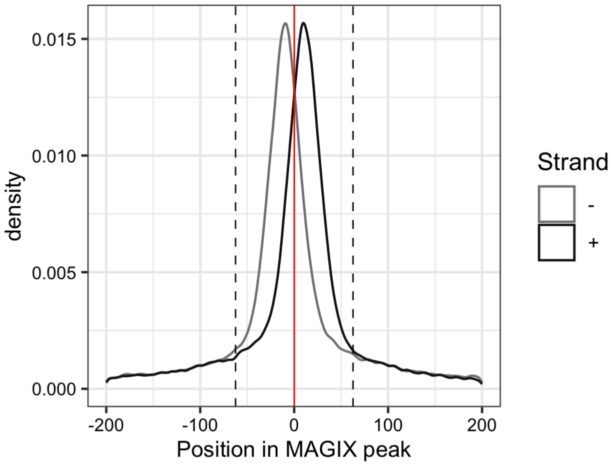

### ZNF646.DBD3

Top GRECO-BIT/MEX PWM: PWM004958

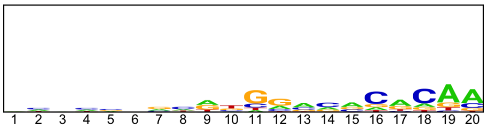


----------------------------------------------------------------------------------------------------

Location of top GRECO-BIT/MEX PWM in MAGIX peaks (up to the top 10,000)

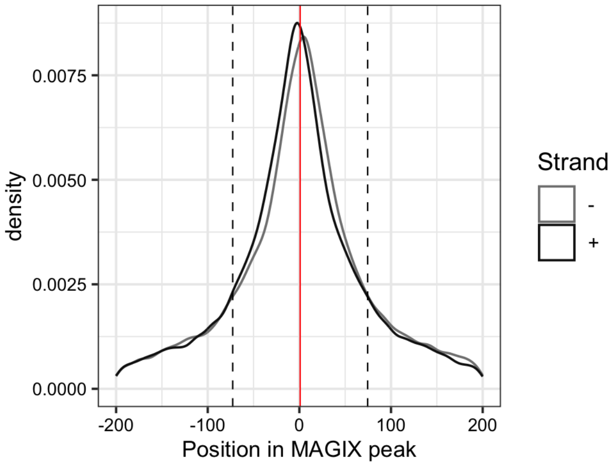

### ZNF648

Top GRECO-BIT/MEX PWM: PWM004990

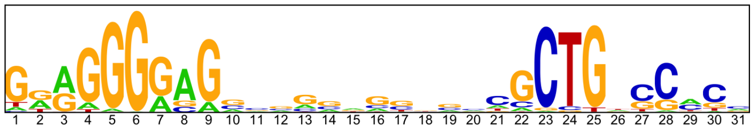

----------------------------------------------------------------------------------------------------

Location of top GRECO-BIT/MEX PWM in MAGIX peaks (up to the top 10,000)

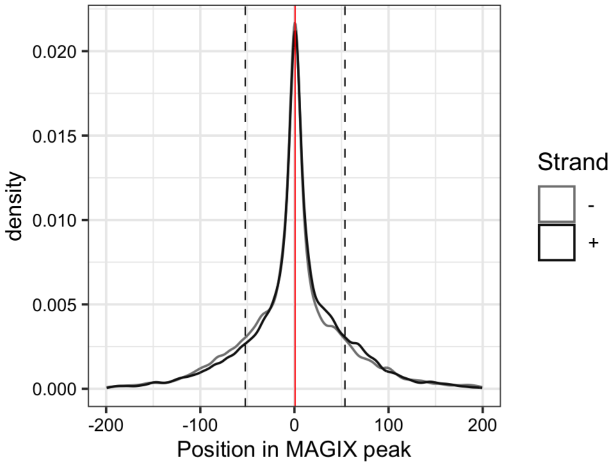


----------------------------------------------------------------------------------------------------

AUROC of top GRECO-BIT/MEX PWM in sliding windows (500 peaks, 50 peak step) along the MAGIX peaks (up to the top 75,000) ranked by the MAGIX enrichment coefficient and fraction of overlapping ChIP peaks in the same window

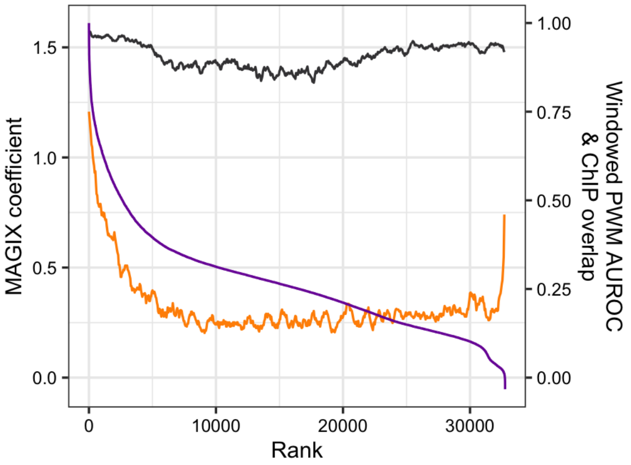

### ZNF66

Top GRECO-BIT/MEX PWM: PWM005025

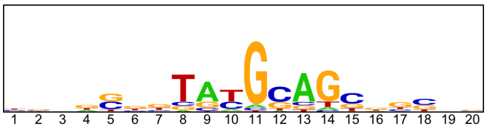


--------------------------------------------------------------------------------

Location of top GRECO-BIT/MEX PWM in MAGIX peaks (up to the top 10,000)

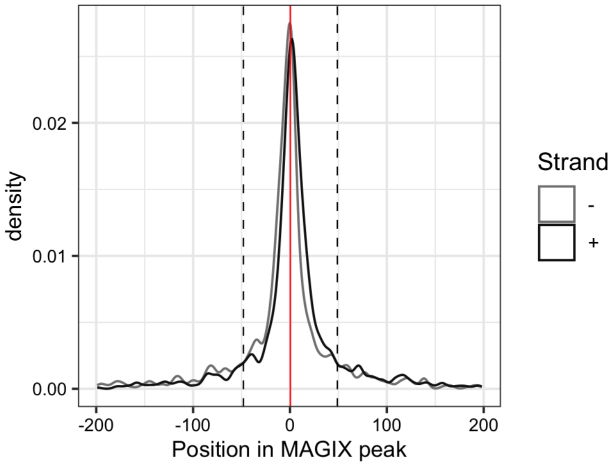


--------------------------------------------------------------------------------

AUROC of top GRECO-BIT/MEX PWM in sliding windows (500 peaks, 50 peak step) along the MAGIX peaks (up to the top 75,000) ranked by the MAGIX enrichment coefficient and fraction of overlapping ChIP peaks in the same window

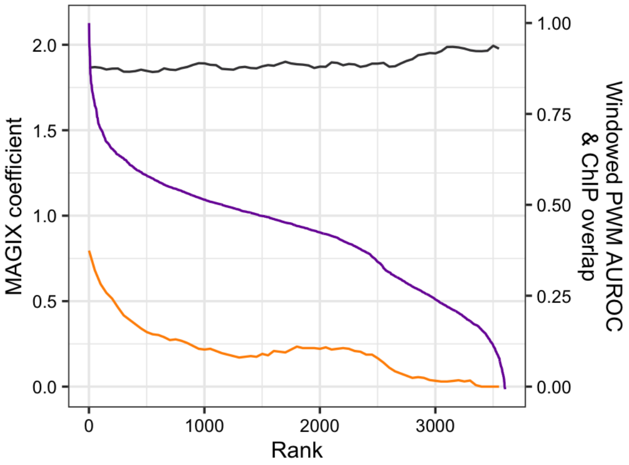

### ZNF665

Top GRECO-BIT/MEX PWM: PWM005065

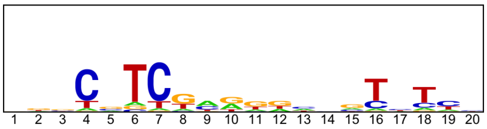


-----------------------------------------------------------------------------------------------

Location of top GRECO-BIT/MEX PWM in MAGIX peaks (up to the top 10,000)

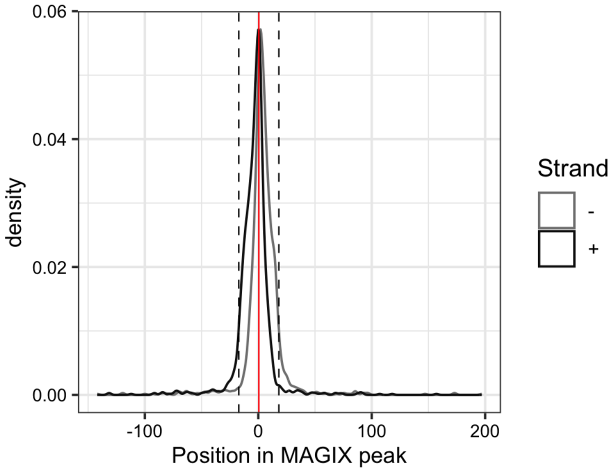


-----------------------------------------------------------------------------------------------

AUROC of top GRECO-BIT/MEX PWM in sliding windows (500 peaks, 50 peak step) along the MAGIX peaks (up to the top 75,000) ranked by the MAGIX enrichment coefficient and fraction of overlapping ChIP peaks in the same window

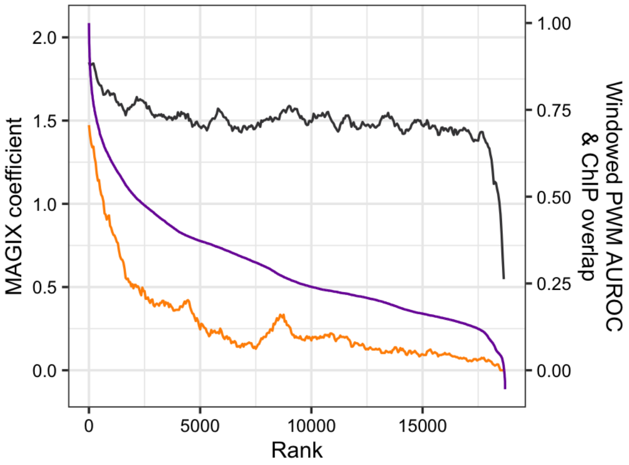

### ZNF668.DBD1

Top GRECO-BIT/MEX PWM: PWM005082

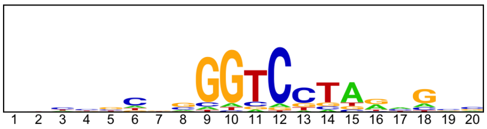


------------------------------------------------------------------------------------------------

Location of top GRECO-BIT/MEX PWM in MAGIX peaks (up to the top 10,000)

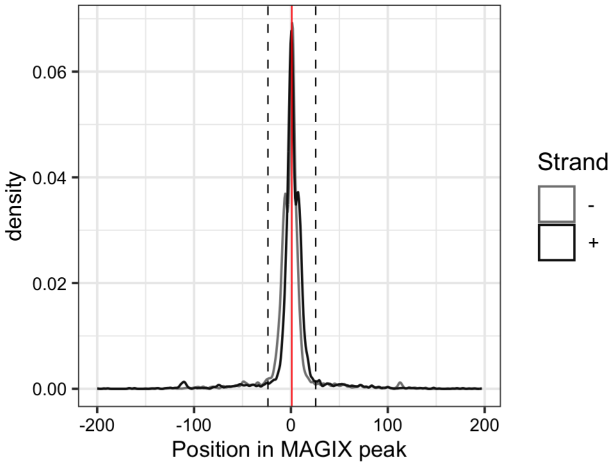

### ZNF668.FL

Top GRECO-BIT/MEX PWM: PWM005084

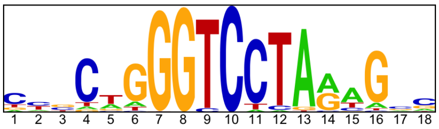


--------------------------------------------------------------------------------------------

Location of top GRECO-BIT/MEX PWM in MAGIX peaks (up to the top 10,000)

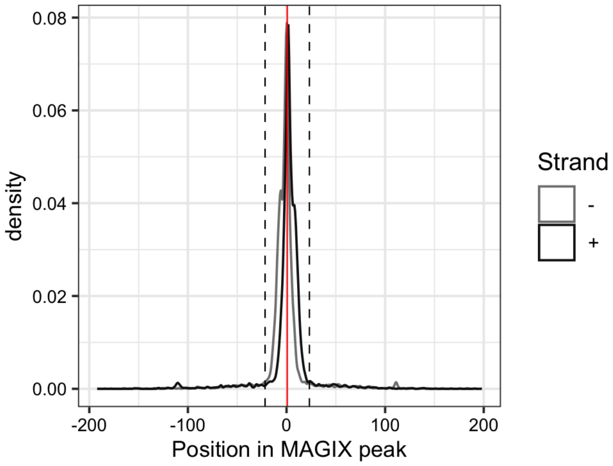

### ZNF672

Top GRECO-BIT/MEX PWM: PWM005120

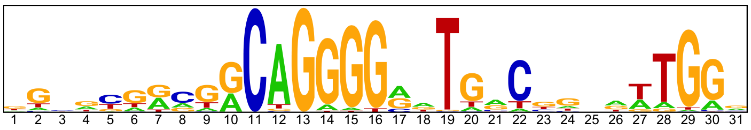


-------------------------------------------------------------------------------------------

Location of top GRECO-BIT/MEX PWM in MAGIX peaks (up to the top 10,000)

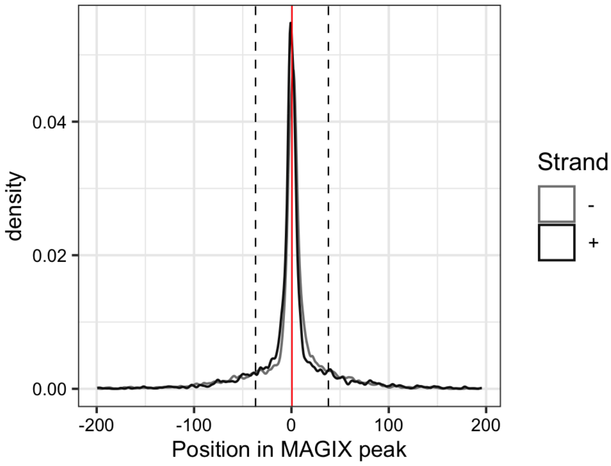


-------------------------------------------------------------------------------------------

AUROC of top GRECO-BIT/MEX PWM in sliding windows (500 peaks, 50 peak step) along the MAGIX peaks (up to the top 75,000) ranked by the MAGIX enrichment coefficient and fraction of overlapping ChIP peaks in the same window

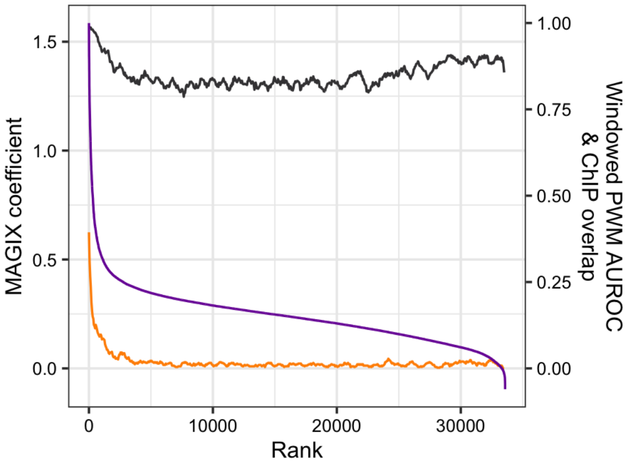

### ZNF676

Top GRECO-BIT/MEX PWM: PWM005154

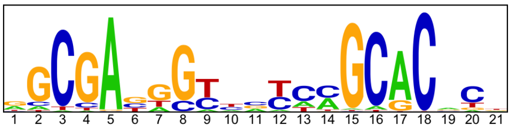

---

Location of top GRECO-BIT/MEX PWM in MAGIX peaks (up to the top 10,000)

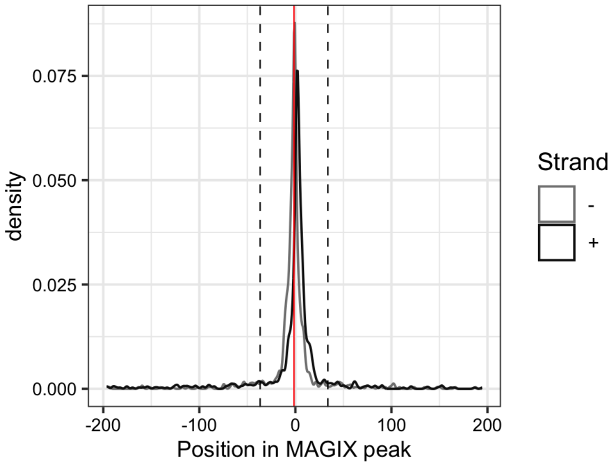


---

AUROC of top GRECO-BIT/MEX PWM in sliding windows (500 peaks, 50 peak step) along the MAGIX peaks (up to the top 75,000) ranked by the MAGIX enrichment coefficient and fraction of overlapping ChIP peaks in the same window

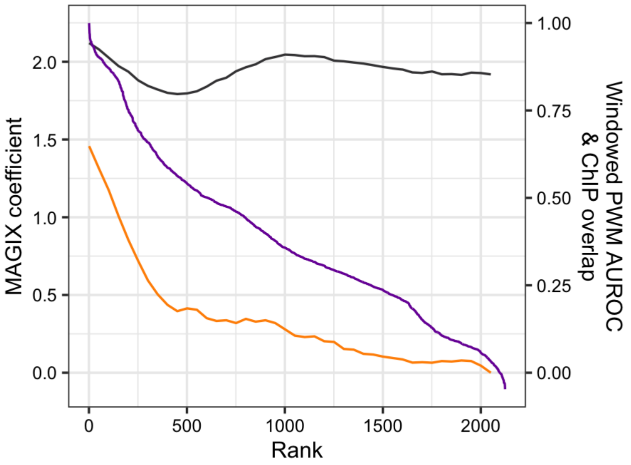

### ZNF678

Top GRECO-BIT/MEX PWM: PWM005176

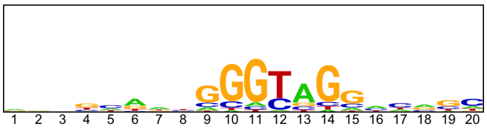


----------------------------------------------------------------------------------------------

Location of top GRECO-BIT/MEX PWM in MAGIX peaks (up to the top 10,000)

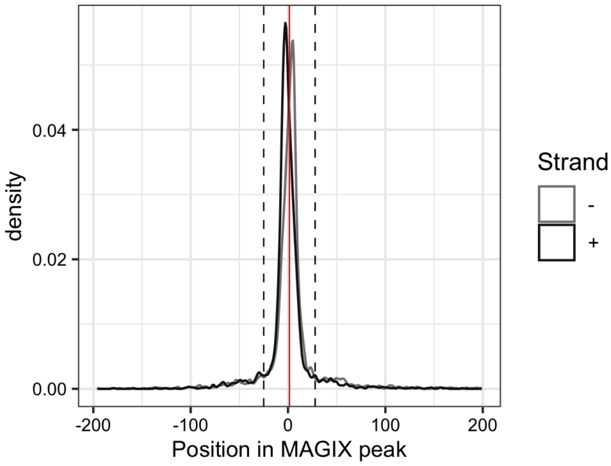


----------------------------------------------------------------------------------------------

AUROC of top GRECO-BIT/MEX PWM in sliding windows (500 peaks, 50 peak step) along the MAGIX peaks (up to the top 75,000) ranked by the MAGIX enrichment coefficient and fraction of overlapping ChIP peaks in the same window

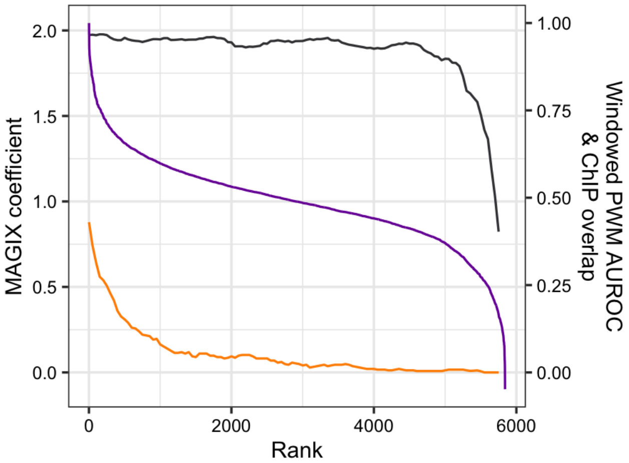

### ZNF683

Top GRECO-BIT/MEX PWM: PWM005199

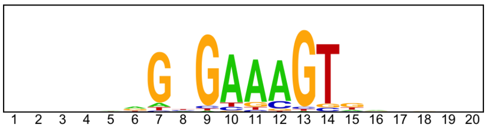


--------------------------------------------------------------------------------------------

Location of top GRECO-BIT/MEX PWM in MAGIX peaks (up to the top 10,000)

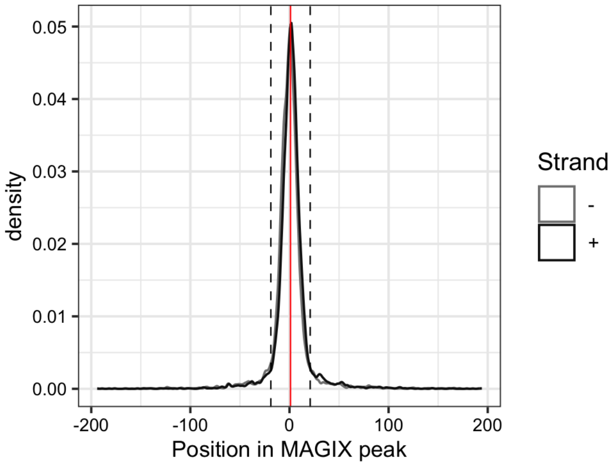


--------------------------------------------------------------------------------------------

AUROC of top GRECO-BIT/MEX PWM in sliding windows (500 peaks, 50 peak step) along the MAGIX peaks (up to the top 75,000) ranked by the MAGIX enrichment coefficient and fraction of overlapping ChIP peaks in the same window

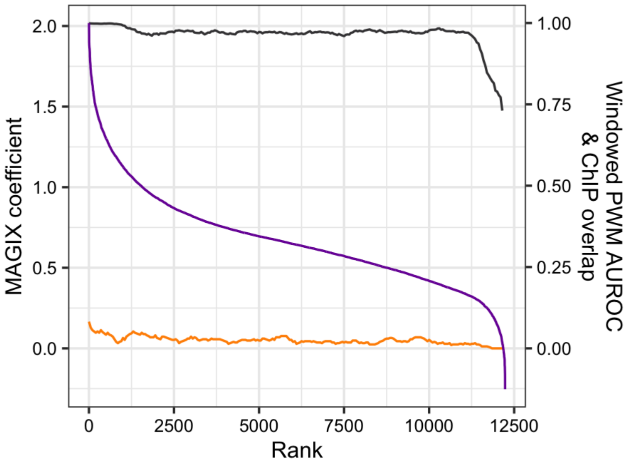

### ZNF689

Top GRECO-BIT/MEX PWM: PWM005247

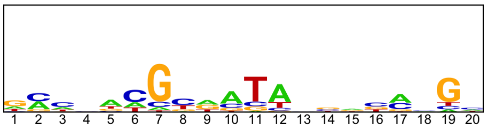


---

Location of top GRECO-BIT/MEX PWM in MAGIX peaks (up to the top 10,000)

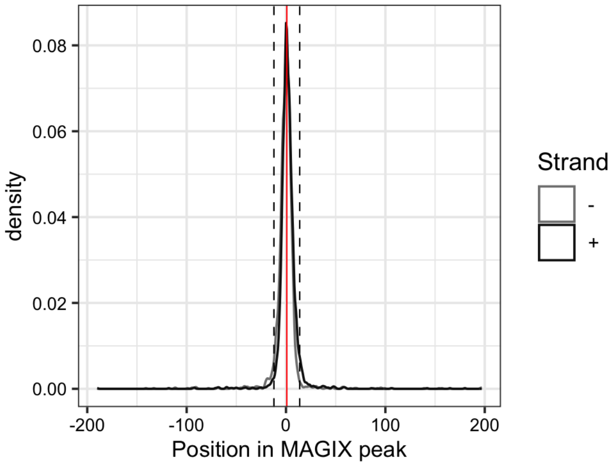


---

AUROC of top GRECO-BIT/MEX PWM in sliding windows (500 peaks, 50 peak step) along the MAGIX peaks (up to the top 75,000) ranked by the MAGIX enrichment coefficient and fraction of overlapping ChIP peaks in the same window

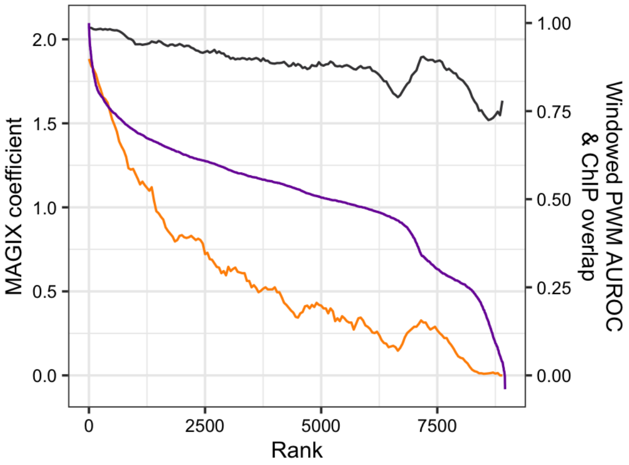

### ZNF696

Top GRECO-BIT/MEX PWM: PWM005274

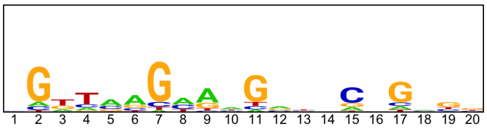


---

Location of top GRECO-BIT/MEX PWM in MAGIX peaks (up to the top 10,000)

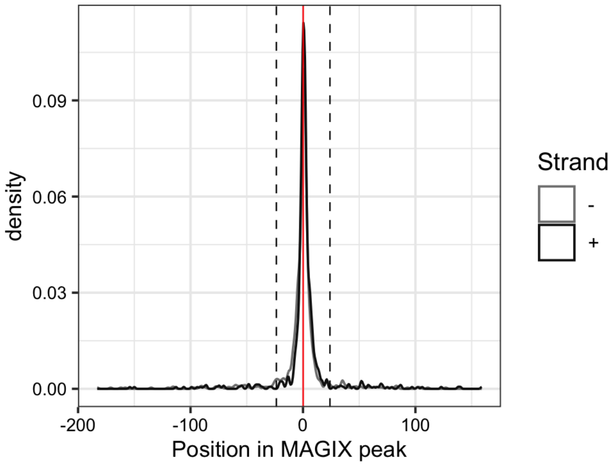


---

AUROC of top GRECO-BIT/MEX PWM in sliding windows (500 peaks, 50 peak step) along the MAGIX peaks (up to the top 75,000) ranked by the MAGIX enrichment coefficient and fraction of overlapping ChIP peaks in the same window

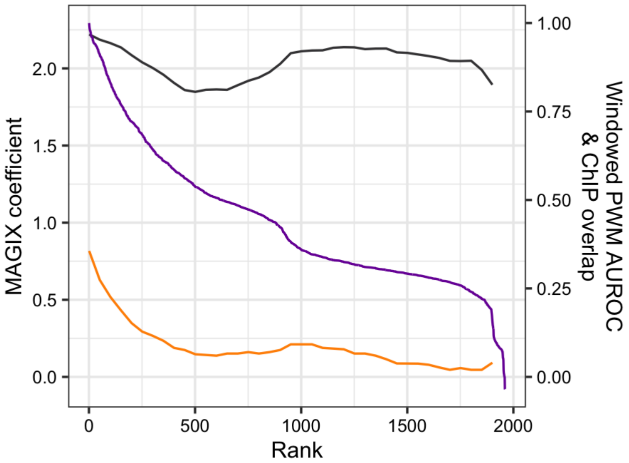

### ZNF699

Top GRECO-BIT/MEX PWM: PWM005313

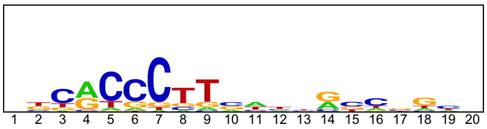


---

Location of top GRECO-BIT/MEX PWM in MAGIX peaks (up to the top 10,000)

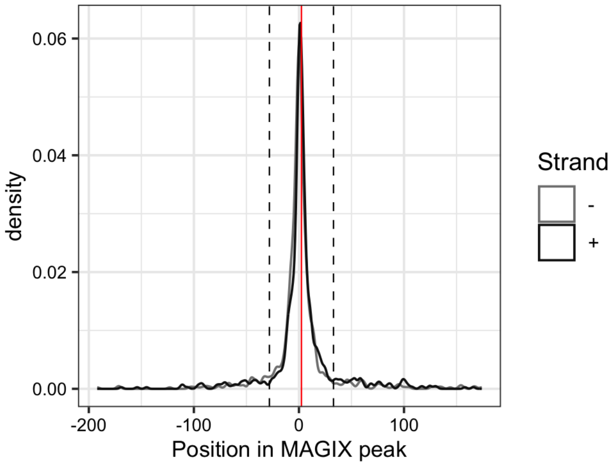


---

AUROC of top GRECO-BIT/MEX PWM in sliding windows (500 peaks, 50 peak step) along the MAGIX peaks (up to the top 75,000) ranked by the MAGIX enrichment coefficient and fraction of overlapping ChIP peaks in the same window

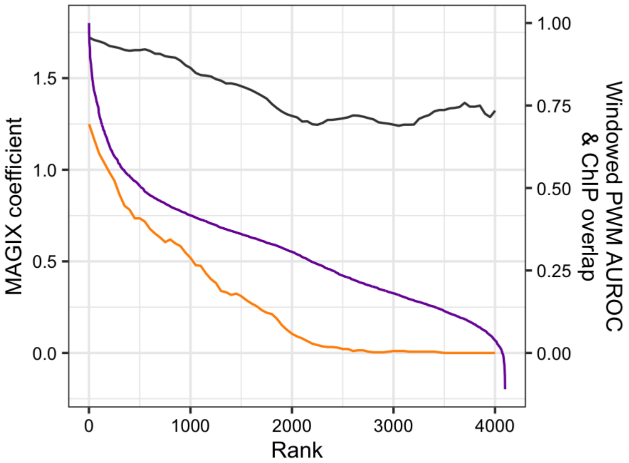

### ZNF70

Top GRECO-BIT/MEX PWM: PWM005343

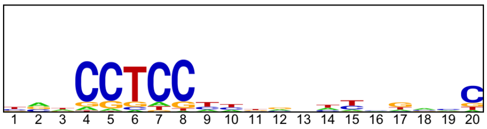


---

Location of top GRECO-BIT/MEX PWM in MAGIX peaks (up to the top 10,000)

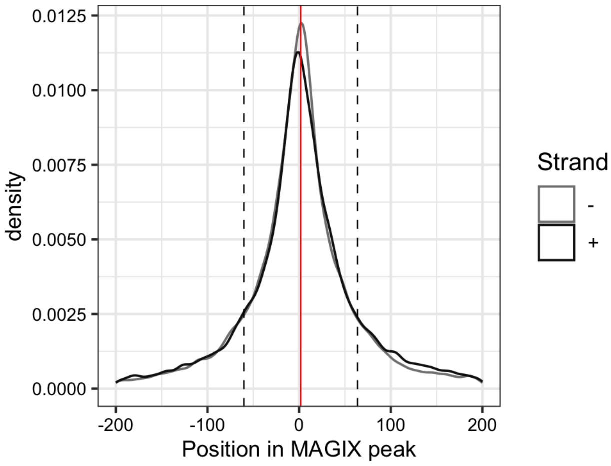


---

AUROC of top GRECO-BIT/MEX PWM in sliding windows (500 peaks, 50 peak step) along the MAGIX peaks (up to the top 75,000) ranked by the MAGIX enrichment coefficient and fraction of overlapping ChIP peaks in the same window

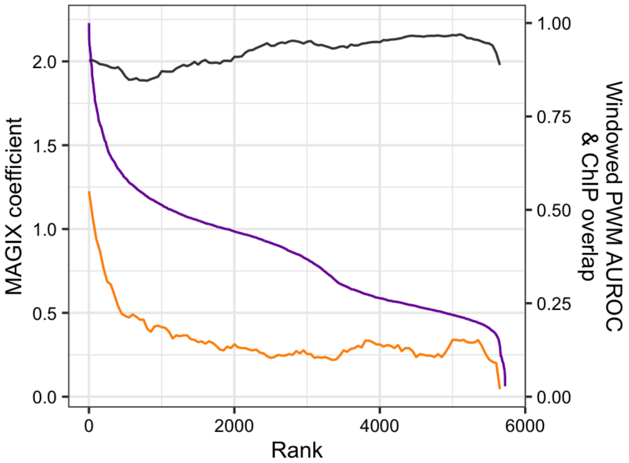

### ZNF700.DBD2

Top GRECO-BIT/MEX PWM: PWM005386

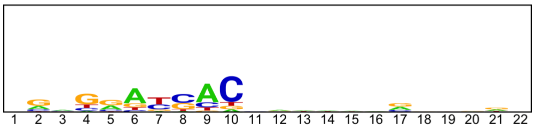


---

Location of top GRECO-BIT/MEX PWM in MAGIX peaks (up to the top 10,000)

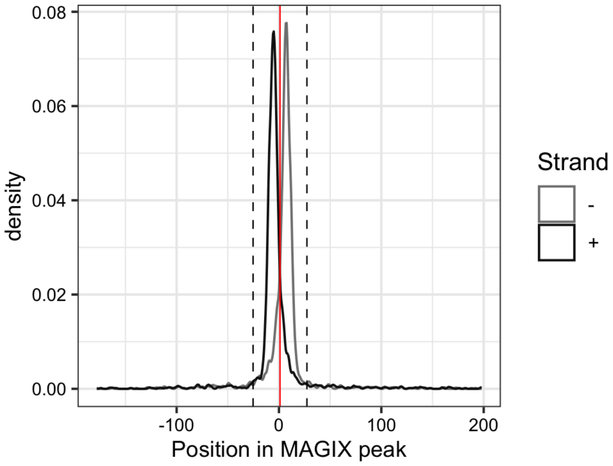

### ZNF721.DBD1

Top GRECO-BIT/MEX PWM: PWM005501

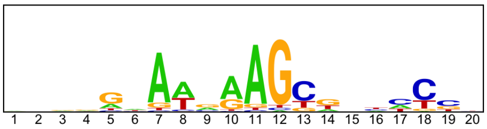


--------------------------------------------------------------------------------------------

Location of top GRECO-BIT/MEX PWM in MAGIX peaks (up to the top 10,000)

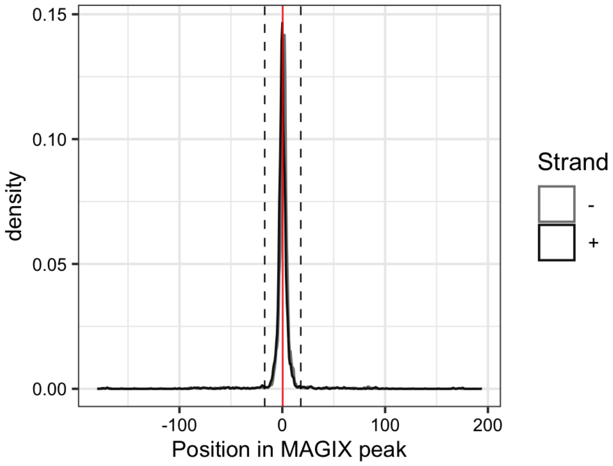

### ZNF721.DBD2

Top GRECO-BIT/MEX PWM: PWM005498

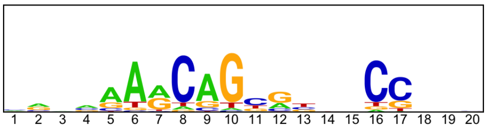


-----------------------------------------------------------------------------------------------

Location of top GRECO-BIT/MEX PWM in MAGIX peaks (up to the top 10,000)

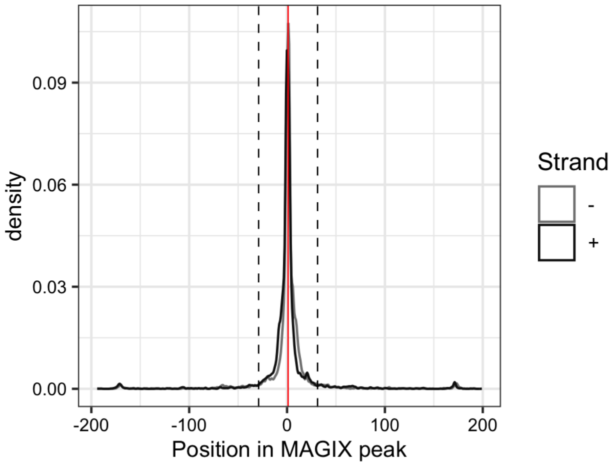

### ZNF721.FL

Top GRECO-BIT/MEX PWM: PWM005515

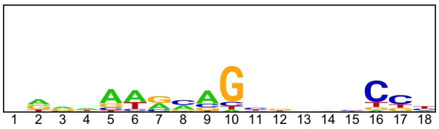


--------------------------------------------------------------------------------

Location of top GRECO-BIT/MEX PWM in MAGIX peaks (up to the top 10,000)

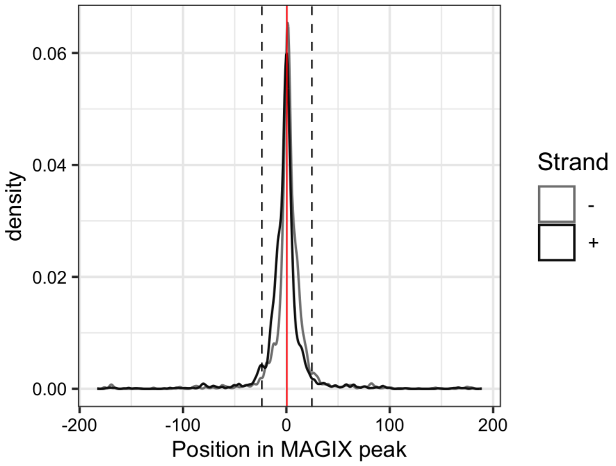


--------------------------------------------------------------------------------

AUROC of top GRECO-BIT/MEX PWM in sliding windows (500 peaks, 50 peak step) along the MAGIX peaks (up to the top 75,000) ranked by the MAGIX enrichment coefficient and fraction of overlapping ChIP peaks in the same window

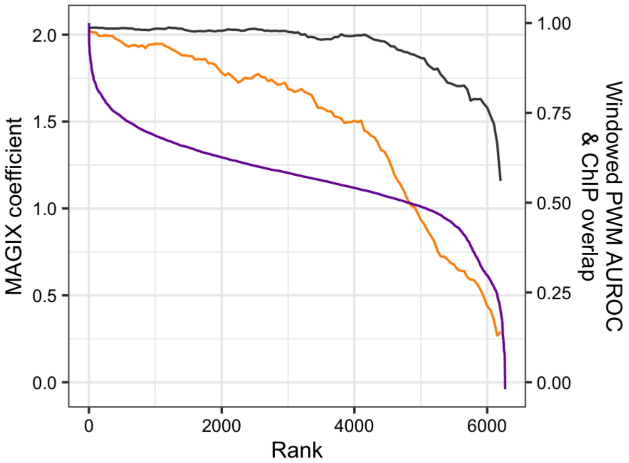

### ZNF724

Top GRECO-BIT/MEX PWM: PWM005525

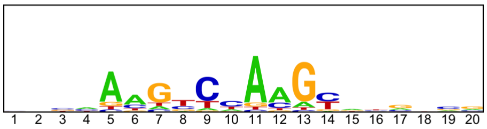


-----------------------------------------------------------------------------------------------

Location of top GRECO-BIT/MEX PWM in MAGIX peaks (up to the top 10,000)

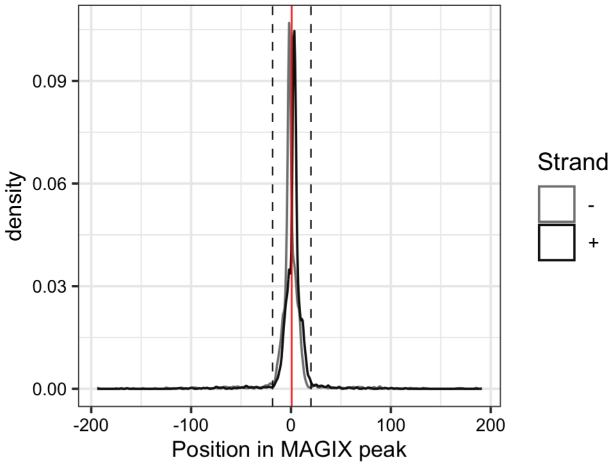


-----------------------------------------------------------------------------------------------

AUROC of top GRECO-BIT/MEX PWM in sliding windows (500 peaks, 50 peak step) along the MAGIX peaks (up to the top 75,000) ranked by the MAGIX enrichment coefficient and fraction of overlapping ChIP peaks in the same window

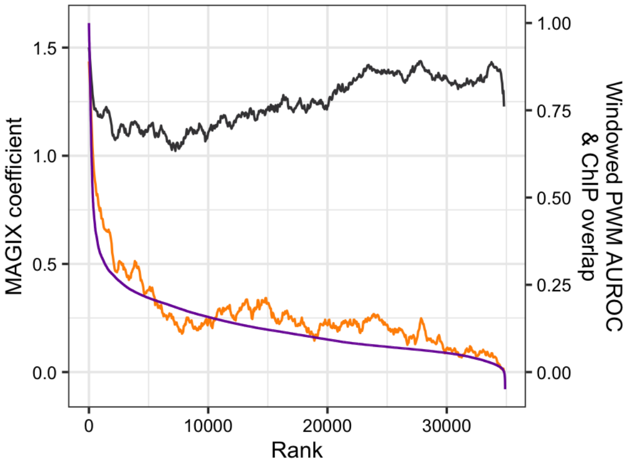

### ZNF726.DBD2

Top GRECO-BIT/MEX PWM: PWM005557

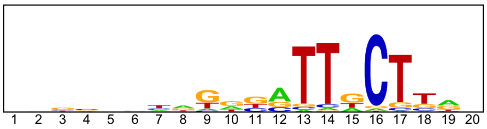


---

Location of top GRECO-BIT/MEX PWM in MAGIX peaks (up to the top 10,000)

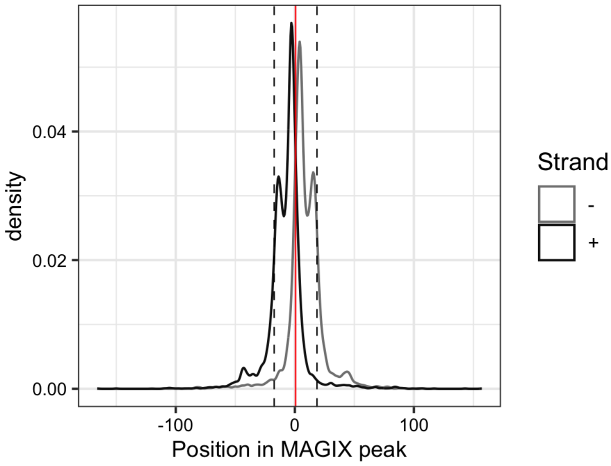

### ZNF726.FL

Top GRECO-BIT/MEX PWM: PWM005561

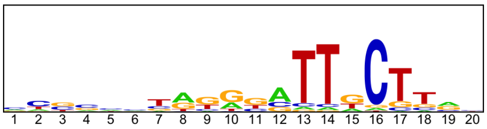


--------------------------------------------------------------------------------------------

Location of top GRECO-BIT/MEX PWM in MAGIX peaks (up to the top 10,000)

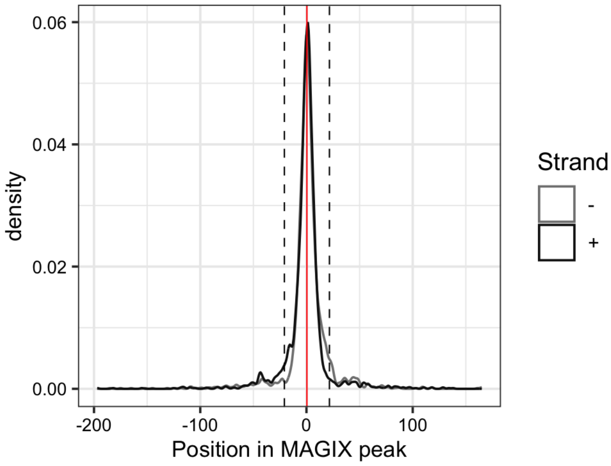


--------------------------------------------------------------------------------------------

AUROC of top GRECO-BIT/MEX PWM in sliding windows (500 peaks, 50 peak step) along the MAGIX peaks (up to the top 75,000) ranked by the MAGIX enrichment coefficient and fraction of overlapping ChIP peaks in the same window

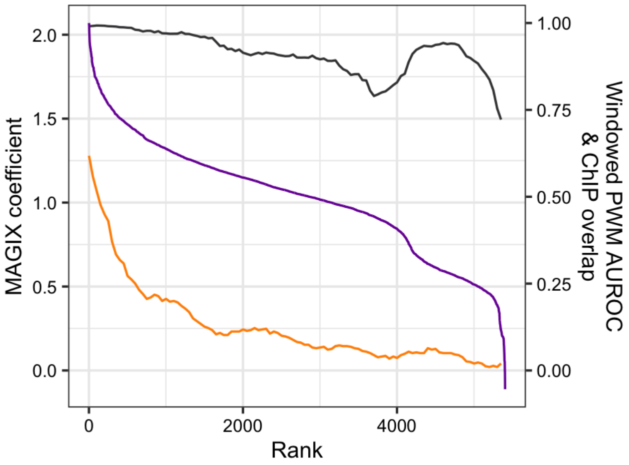

### ZNF728

Top GRECO-BIT/MEX PWM: PWM005595

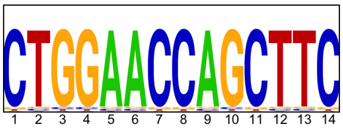


---

Location of top GRECO-BIT/MEX PWM in MAGIX peaks (up to the top 10,000)

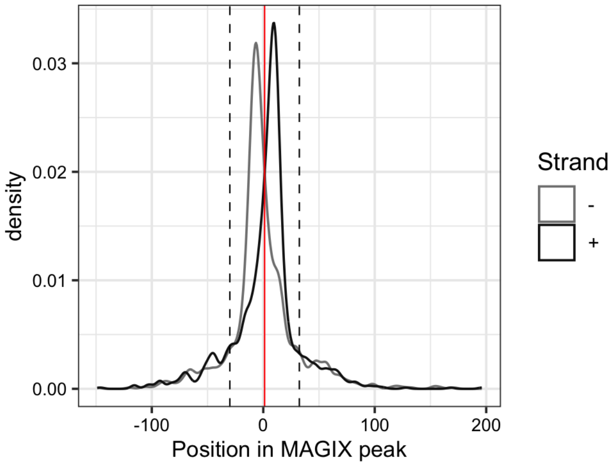


---

AUROC of top GRECO-BIT/MEX PWM in sliding windows (500 peaks, 50 peak step) along the MAGIX peaks (up to the top 75,000) ranked by the MAGIX enrichment coefficient and fraction of overlapping ChIP peaks in the same window

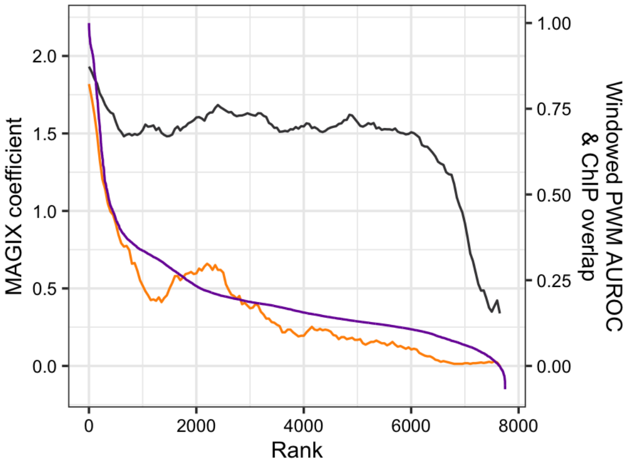

### ZNF729.DBD1

Top GRECO-BIT/MEX PWM: PWM180359

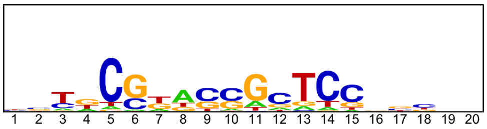


--------------------------------------------------------------------------------------------

Location of top GRECO-BIT/MEX PWM in MAGIX peaks (up to the top 10,000)

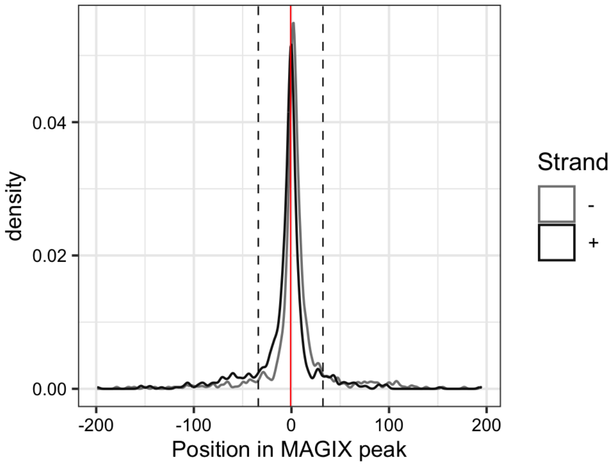

### ZNF729.DBD2

Top GRECO-BIT/MEX PWM: PWM180029

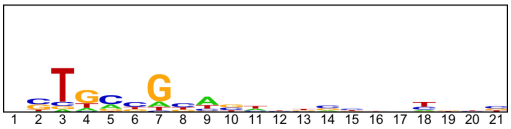


---

Location of top GRECO-BIT/MEX PWM in MAGIX peaks (up to the top 10,000)

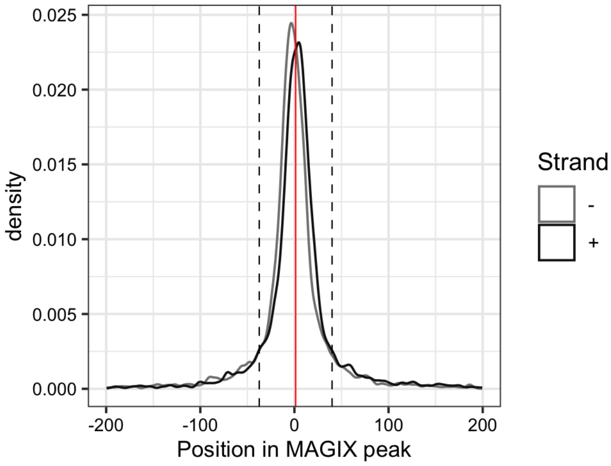

### ZNF729.DBD3

Top GRECO-BIT/MEX PWM: PWM005604

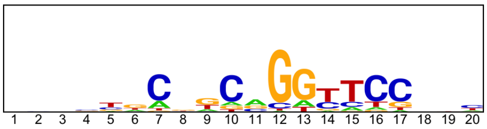


----------------------------------------------------------------------------------------------------

Location of top GRECO-BIT/MEX PWM in MAGIX peaks (up to the top 10,000)

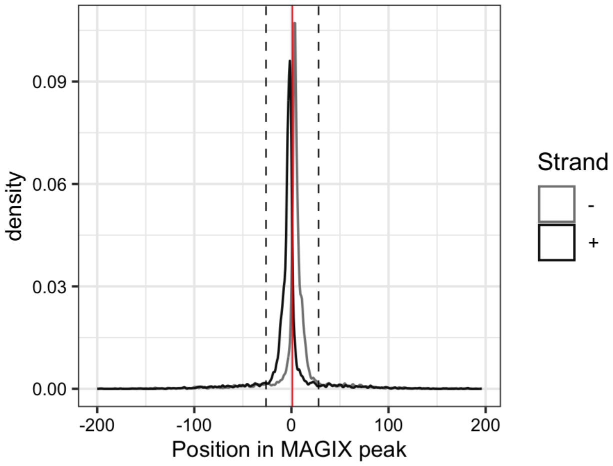

### ZNF732

Top GRECO-BIT/MEX PWM: PWM005631

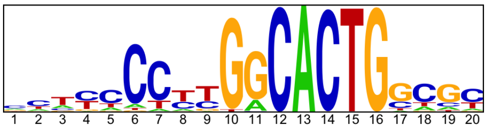

---

Location of top GRECO-BIT/MEX PWM in MAGIX peaks (up to the top 10,000)

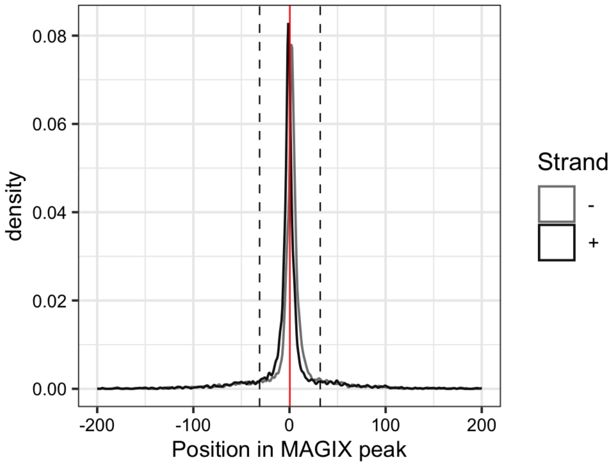


---

AUROC of top GRECO-BIT/MEX PWM in sliding windows (500 peaks, 50 peak step) along the MAGIX peaks (up to the top 75,000) ranked by the MAGIX enrichment coefficient and fraction of overlapping ChIP peaks in the same window

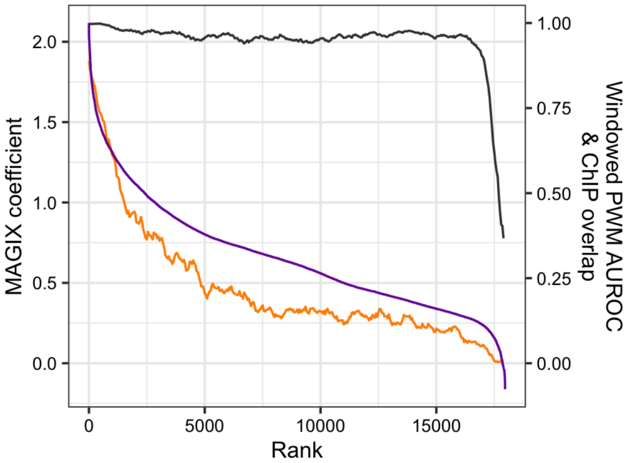

### ZNF746

Top GRECO-BIT/MEX PWM: PWM005659

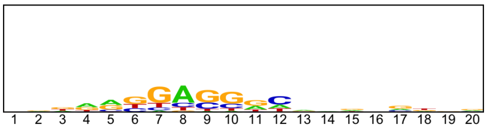


---

Location of top GRECO-BIT/MEX PWM in MAGIX peaks (up to the top 10,000)

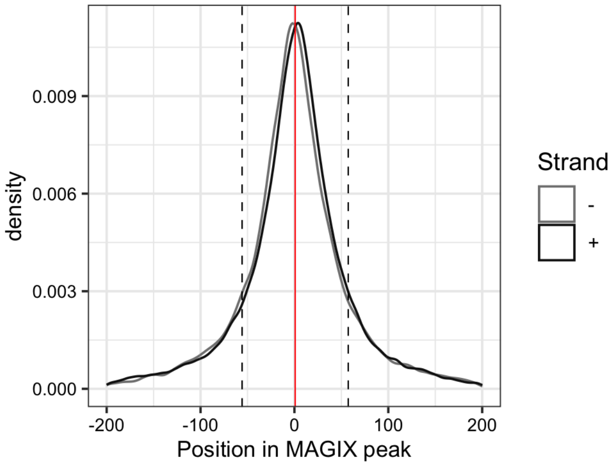


---

AUROC of top GRECO-BIT/MEX PWM in sliding windows (500 peaks, 50 peak step) along the MAGIX peaks (up to the top 75,000) ranked by the MAGIX enrichment coefficient and fraction of overlapping ChIP peaks in the same window

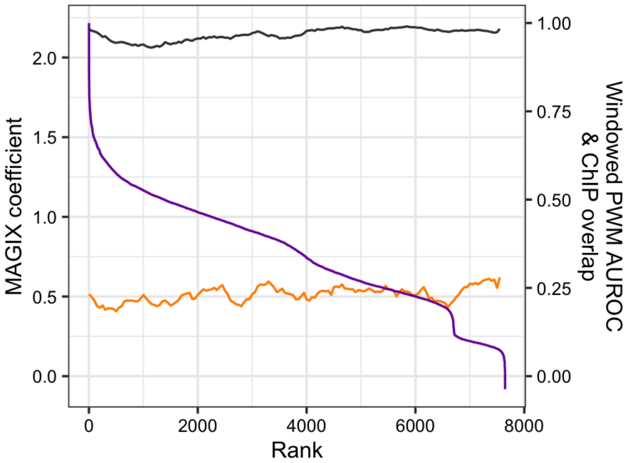

### ZNF770

Top GRECO-BIT/MEX PWM: PWM005690

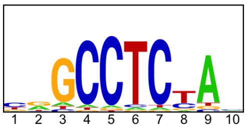


---

Location of top GRECO-BIT/MEX PWM in MAGIX peaks (up to the top 10,000)

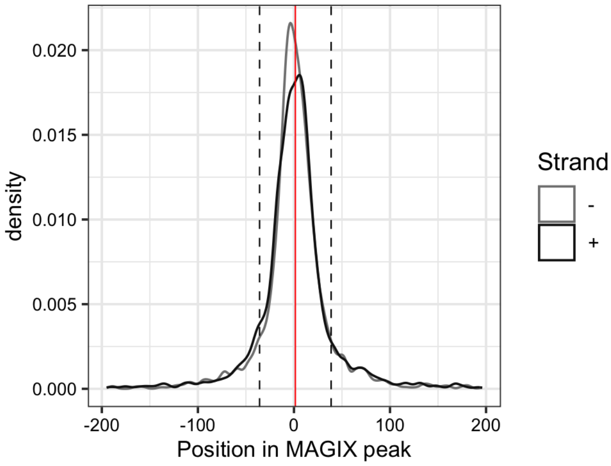


---

AUROC of top GRECO-BIT/MEX PWM in sliding windows (500 peaks, 50 peak step) along the MAGIX peaks (up to the top 75,000) ranked by the MAGIX enrichment coefficient and fraction of overlapping ChIP peaks in the same window

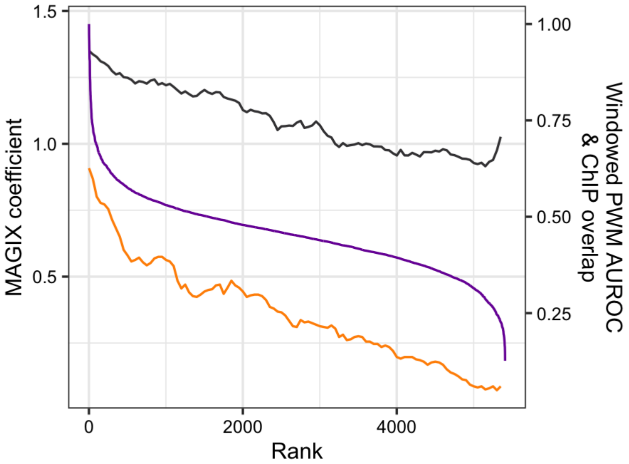

### ZNF772

Top GRECO-BIT/MEX PWM: PWM005715

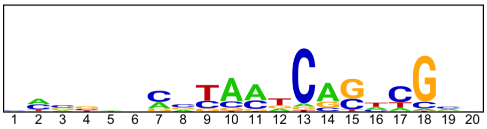


-----------------------------------------------------------------------------------------------

Location of top GRECO-BIT/MEX PWM in MAGIX peaks (up to the top 10,000)

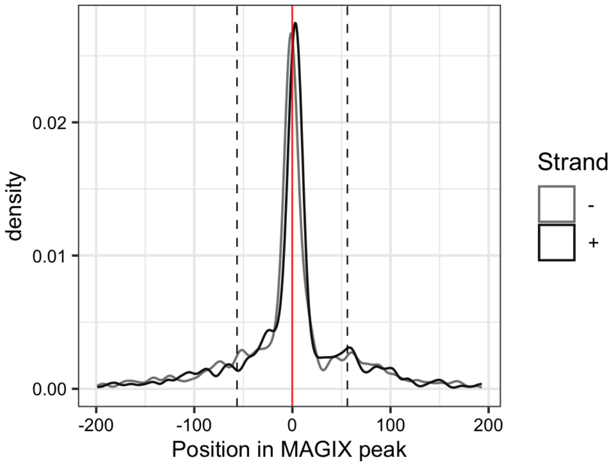


-----------------------------------------------------------------------------------------------

AUROC of top GRECO-BIT/MEX PWM in sliding windows (500 peaks, 50 peak step) along the MAGIX peaks (up to the top 75,000) ranked by the MAGIX enrichment coefficient and fraction of overlapping ChIP peaks in the same window

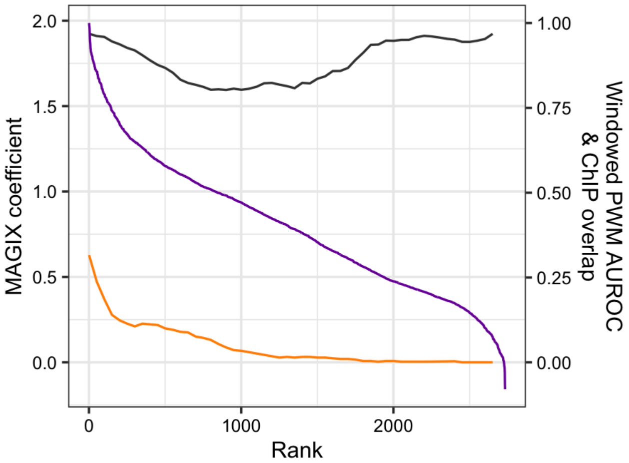

### ZNF773

Top GRECO-BIT/MEX PWM: PWM005749

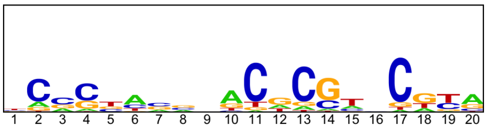


---

Location of top GRECO-BIT/MEX PWM in MAGIX peaks (up to the top 10,000)

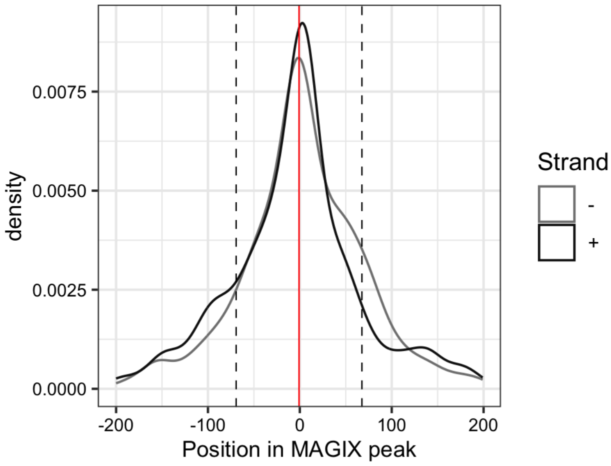


---

AUROC of top GRECO-BIT/MEX PWM in sliding windows (500 peaks, 50 peak step) along the MAGIX peaks (up to the top 75,000) ranked by the MAGIX enrichment coefficient and fraction of overlapping ChIP peaks in the same window

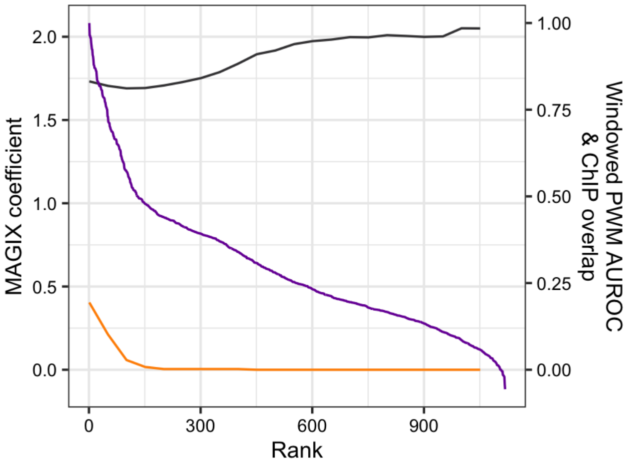

### ZNF775

Top GRECO-BIT/MEX PWM: PWM005771

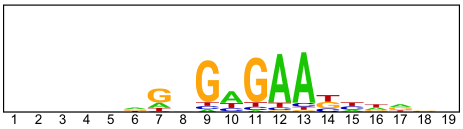


-----------------------------------------------------------------------------------------------

Location of top GRECO-BIT/MEX PWM in MAGIX peaks (up to the top 10,000)

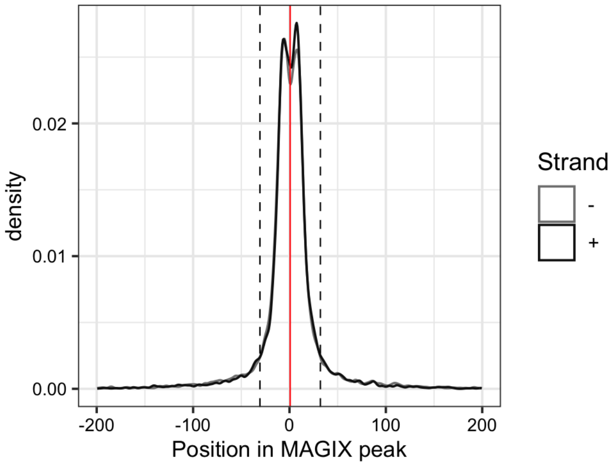


-----------------------------------------------------------------------------------------------

AUROC of top GRECO-BIT/MEX PWM in sliding windows (500 peaks, 50 peak step) along the MAGIX peaks (up to the top 75,000) ranked by the MAGIX enrichment coefficient and fraction of overlapping ChIP peaks in the same window

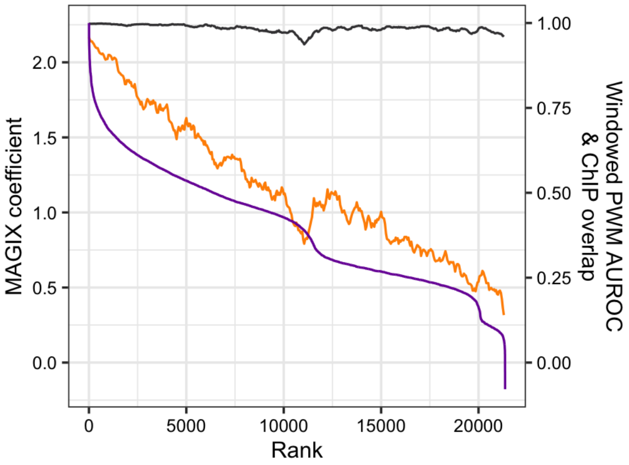

### ZNF780B.DBD1

Top GRECO-BIT/MEX PWM: PWM005795

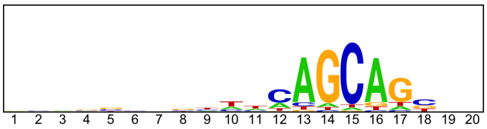


------------------------------------------------------------------------------------------------

Location of top GRECO-BIT/MEX PWM in MAGIX peaks (up to the top 10,000)

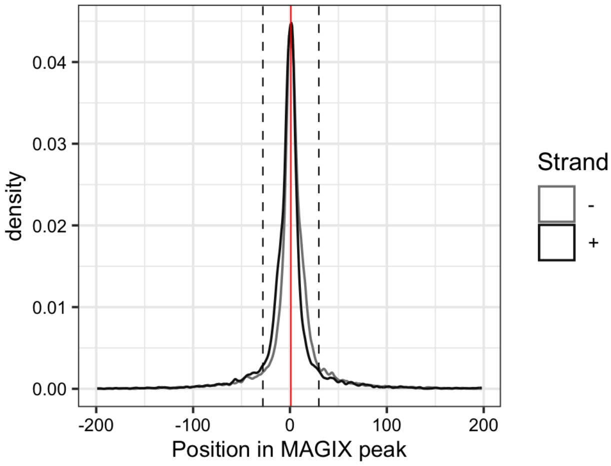

### ZNF780B.FL

Top GRECO-BIT/MEX PWM: PWM005796

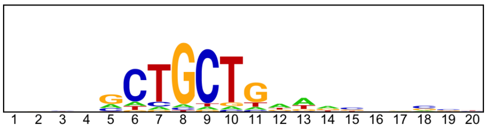


--------------------------------------------------------------------------------------------

Location of top GRECO-BIT/MEX PWM in MAGIX peaks (up to the top 10,000)

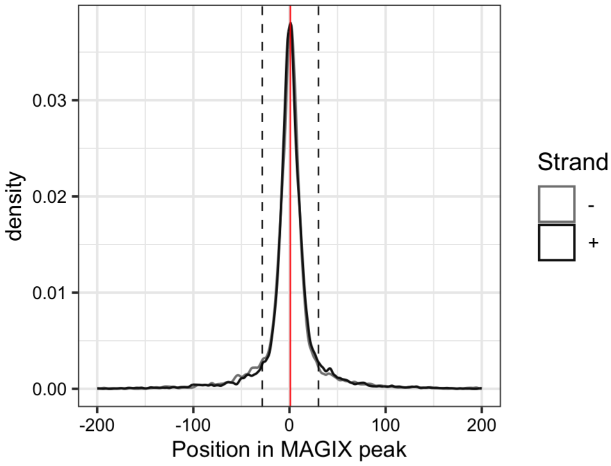


--------------------------------------------------------------------------------------------

AUROC of top GRECO-BIT/MEX PWM in sliding windows (500 peaks, 50 peak step) along the MAGIX peaks (up to the top 75,000) ranked by the MAGIX enrichment coefficient and fraction of overlapping ChIP peaks in the same window

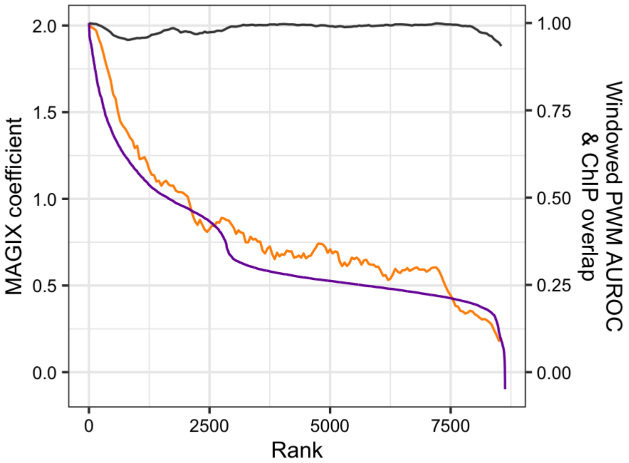

### ZNF800

Top GRECO-BIT/MEX PWM: PWM005848

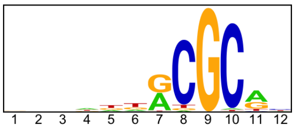

---

Location of top GRECO-BIT/MEX PWM in MAGIX peaks (up to the top 10,000)

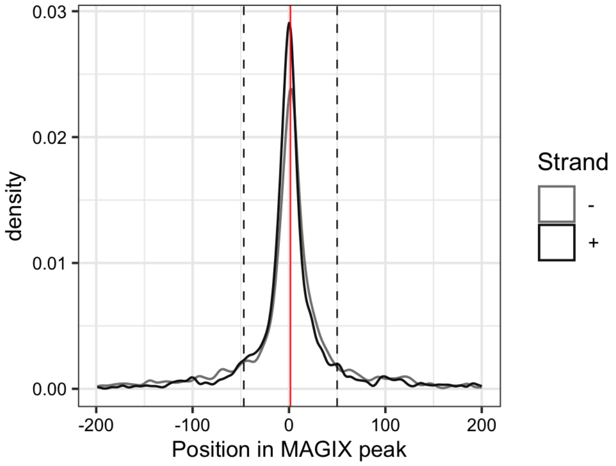


---

AUROC of top GRECO-BIT/MEX PWM in sliding windows (500 peaks, 50 peak step) along the MAGIX peaks (up to the top 75,000) ranked by the MAGIX enrichment coefficient and fraction of overlapping ChIP peaks in the same window

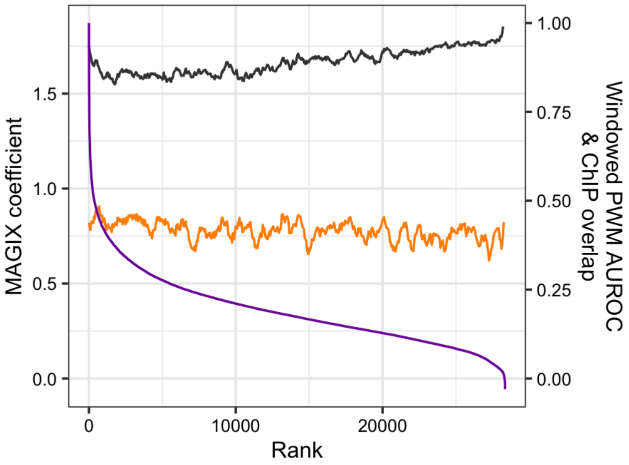

### ZNF813

Top GRECO-BIT/MEX PWM: PWM005866

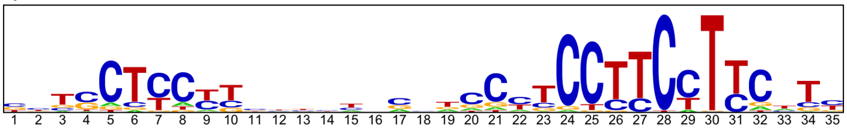

----------------------------------------------------------------------------------------------------

Location of top GRECO-BIT/MEX PWM in MAGIX peaks (up to the top 10,000)

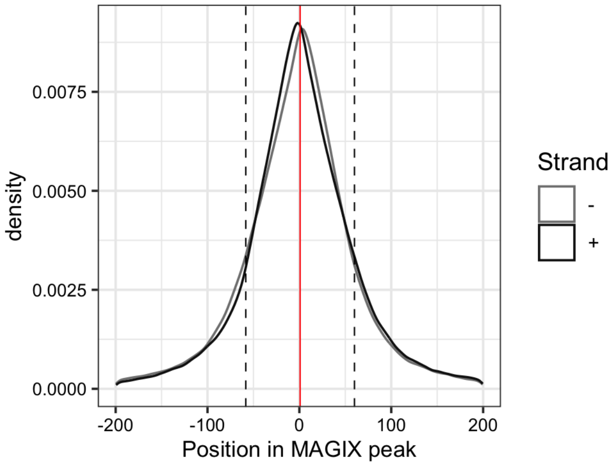

### ZNF814

Top GRECO-BIT/MEX PWM: PWM005898

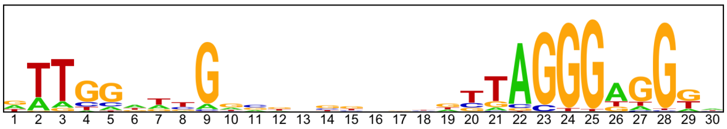


---

Location of top GRECO-BIT/MEX PWM in MAGIX peaks (up to the top 10,000)

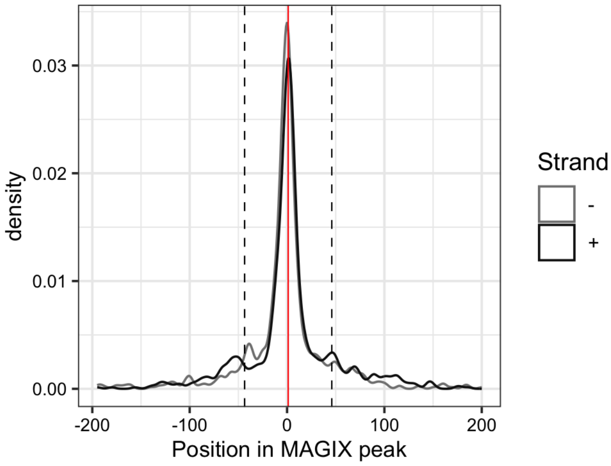


---

AUROC of top GRECO-BIT/MEX PWM in sliding windows (500 peaks, 50 peak step) along the MAGIX peaks (up to the top 75,000) ranked by the MAGIX enrichment coefficient and fraction of overlapping ChIP peaks in the same window

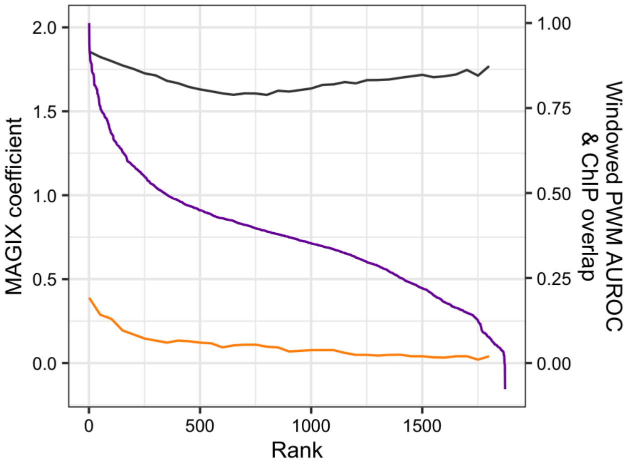

### ZNF827.DBD1

Top GRECO-BIT/MEX PWM: PWM005918

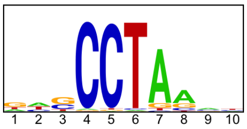

---

Location of top GRECO-BIT/MEX PWM in MAGIX peaks (up to the top 10,000)

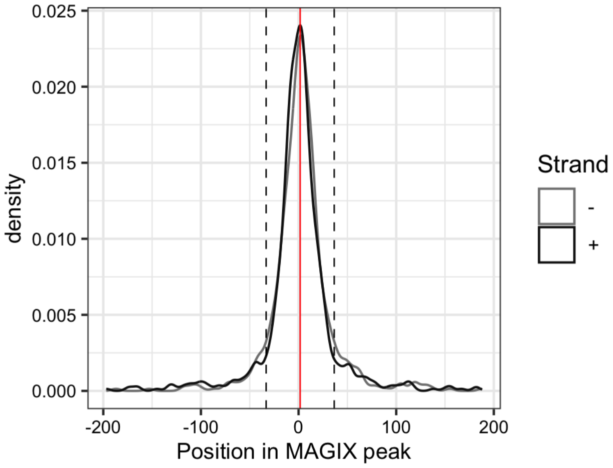

### ZNF83

Top GRECO-BIT/MEX PWM: PWM005938

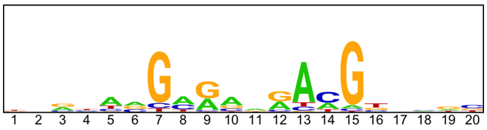


----------------------------------------------------------------------------------------------------

Location of top GRECO-BIT/MEX PWM in MAGIX peaks (up to the top 10,000)

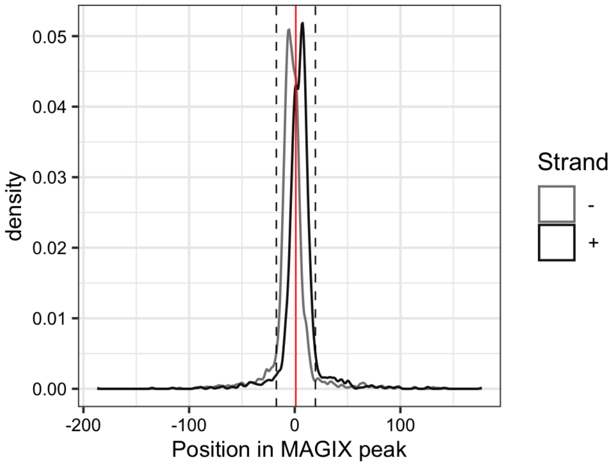

### ZNF831

Top GRECO-BIT/MEX PWM: PWM005969

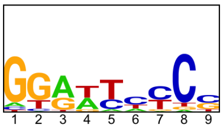


---------------------------------------------------------------------------------------------

Location of top GRECO-BIT/MEX PWM in MAGIX peaks (up to the top 10,000)

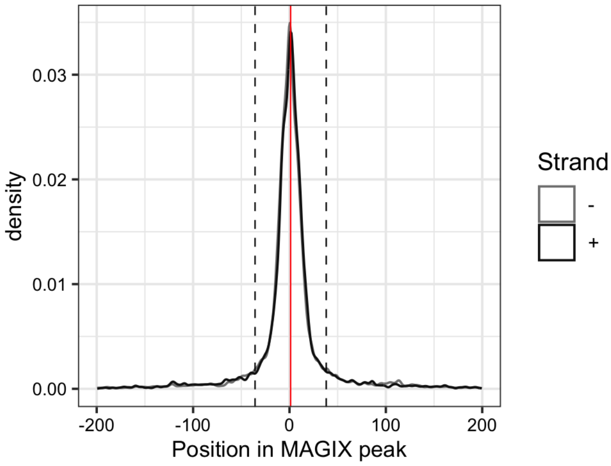


---------------------------------------------------------------------------------------------

AUROC of top GRECO-BIT/MEX PWM in sliding windows (500 peaks, 50 peak step) along the MAGIX peaks (up to the top 75,000) ranked by the MAGIX enrichment coefficient and fraction of overlapping ChIP peaks in the same window

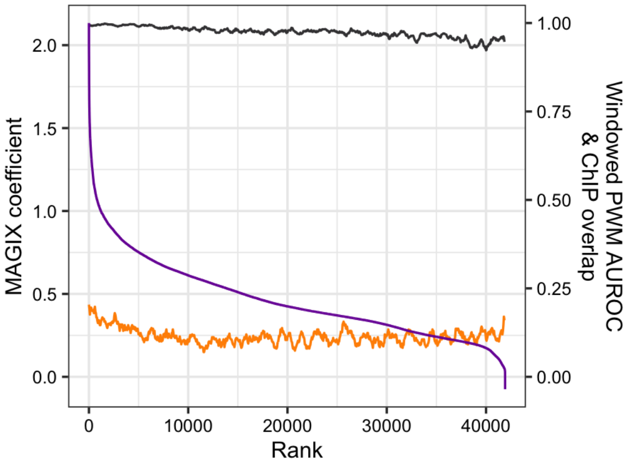

### ZNF836.DBD1

Top GRECO-BIT/MEX PWM: PWM190008

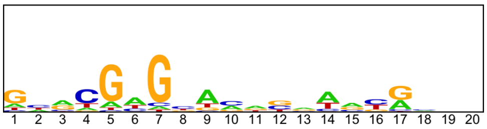


----------------------------------------------------------------------------------------------------

Location of top GRECO-BIT/MEX PWM in MAGIX peaks (up to the top 10,000)

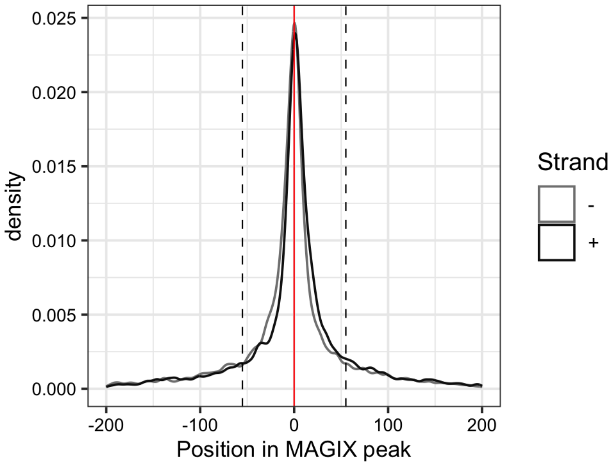

### ZNF836.DBD2

Top GRECO-BIT/MEX PWM: PWM006000

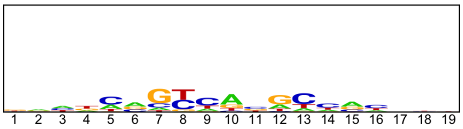


--------------------------------------------------------------------------------------------

Location of top GRECO-BIT/MEX PWM in MAGIX peaks (up to the top 10,000)

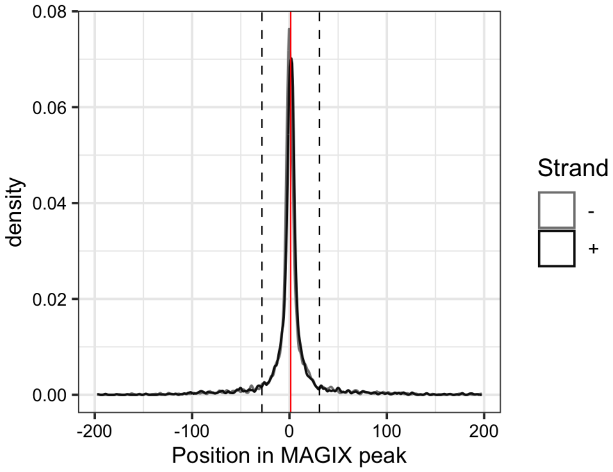

### ZNF836.FL

Top GRECO-BIT/MEX PWM: PWM006005

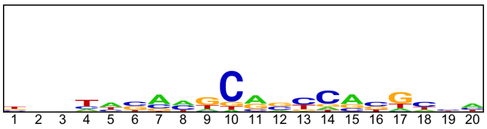


--------------------------------------------------------------------------------------------

Location of top GRECO-BIT/MEX PWM in MAGIX peaks (up to the top 10,000)

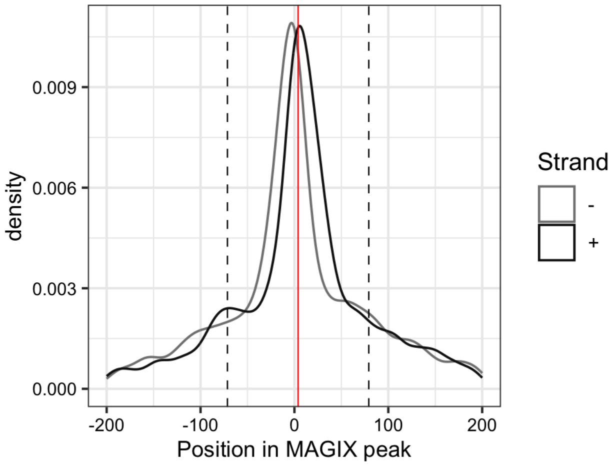


--------------------------------------------------------------------------------------------

AUROC of top GRECO-BIT/MEX PWM in sliding windows (500 peaks, 50 peak step) along the MAGIX peaks (up to the top 75,000) ranked by the MAGIX enrichment coefficient and fraction of overlapping ChIP peaks in the same window

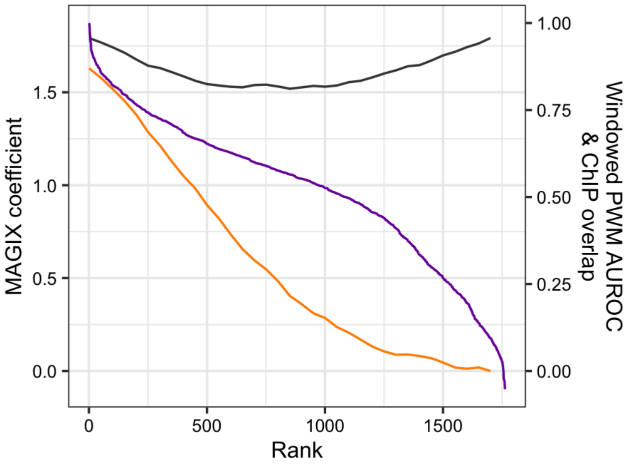

### ZNF841.DBD2

Top GRECO-BIT/MEX PWM: PWM006038

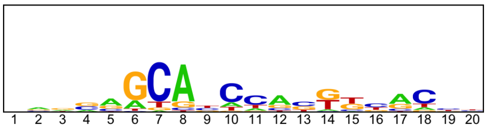


------------------------------------------------------------------------------------------------

Location of top GRECO-BIT/MEX PWM in MAGIX peaks (up to the top 10,000)

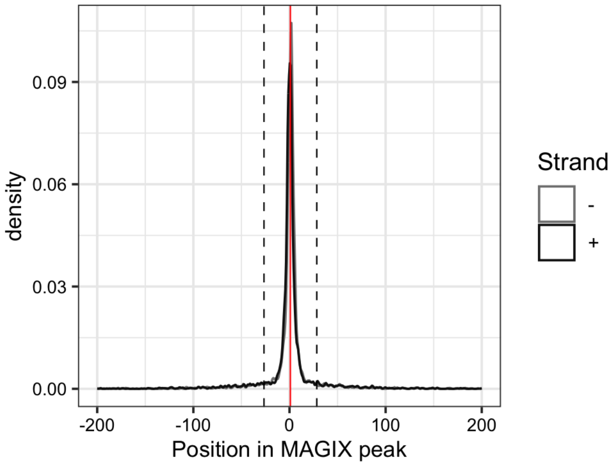

### ZNF841.FL

Top GRECO-BIT/MEX PWM: PWM006030

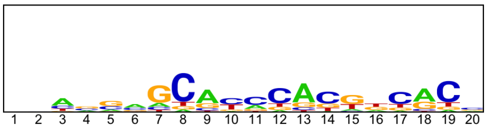


----------------------------------------------------------------------------------------------------

Location of top GRECO-BIT/MEX PWM in MAGIX peaks (up to the top 10,000)

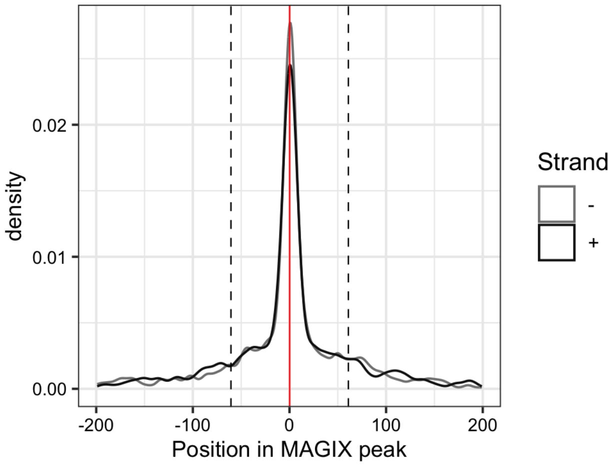


----------------------------------------------------------------------------------------------------

AUROC of top GRECO-BIT/MEX PWM in sliding windows (500 peaks, 50 peak step) along the MAGIX peaks (up to the top 75,000) ranked by the MAGIX enrichment coefficient and fraction of overlapping ChIP peaks in the same window

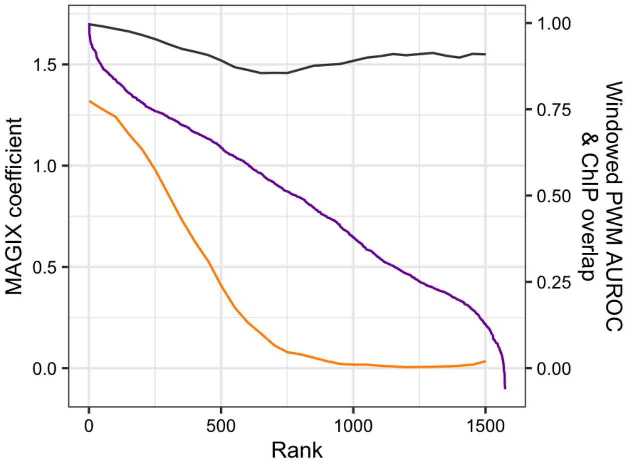

### ZNF845.DBD1

Top GRECO-BIT/MEX PWM: PWM192799

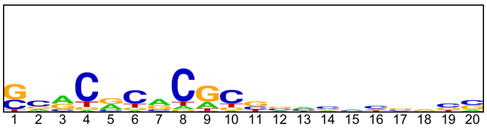


--------------------------------------------------------------------------------------------

Location of top GRECO-BIT/MEX PWM in MAGIX peaks (up to the top 10,000)

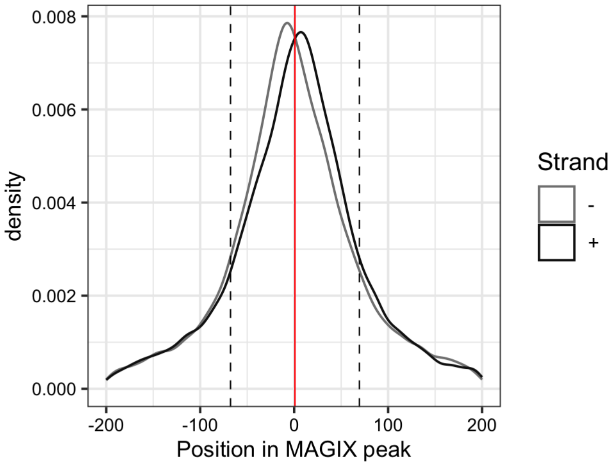

### ZNF845.DBD2

Top GRECO-BIT/MEX PWM: PWM006060

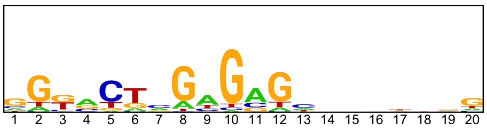

---

Location of top GRECO-BIT/MEX PWM in MAGIX peaks (up to the top 10,000)

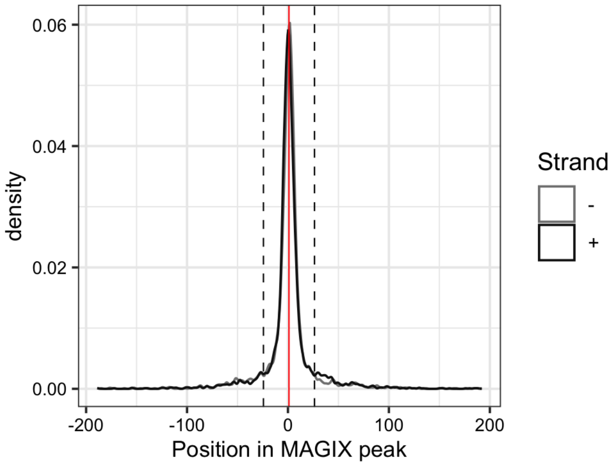

### ZNF850.DBD2

Top GRECO-BIT/MEX PWM: PWM006084

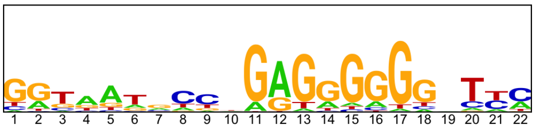

-------------------------------------------------------------------------------------------

Location of top GRECO-BIT/MEX PWM in MAGIX peaks (up to the top 10,000)

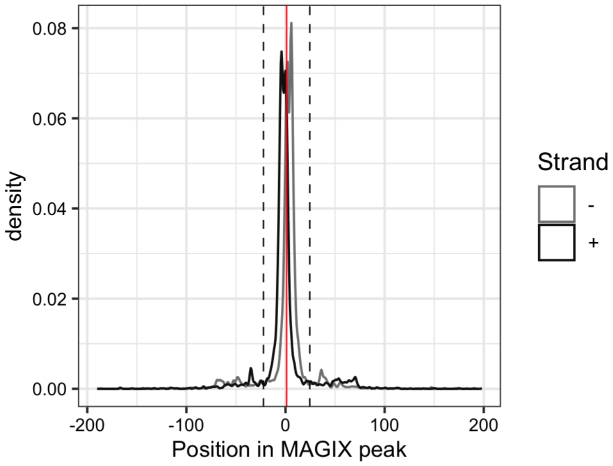

### ZNF853

Top GRECO-BIT/MEX PWM: PWM006104

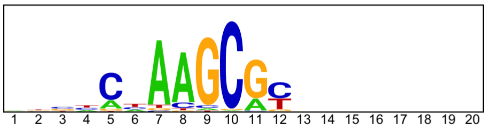


----------------------------------------------------------------------------------------------

Location of top GRECO-BIT/MEX PWM in MAGIX peaks (up to the top 10,000)

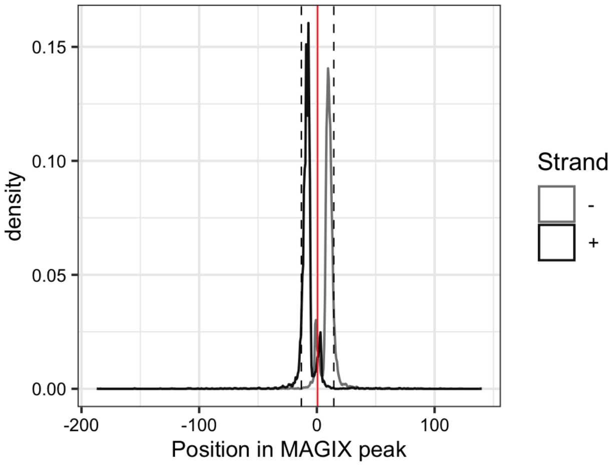

### ZNF865.DBD1

Top GRECO-BIT/MEX PWM: PWM006141

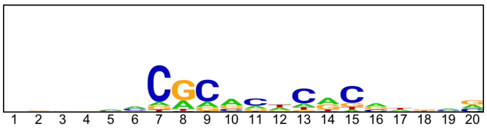


------------------------------------------------------------------------------------------------

Location of top GRECO-BIT/MEX PWM in MAGIX peaks (up to the top 10,000)

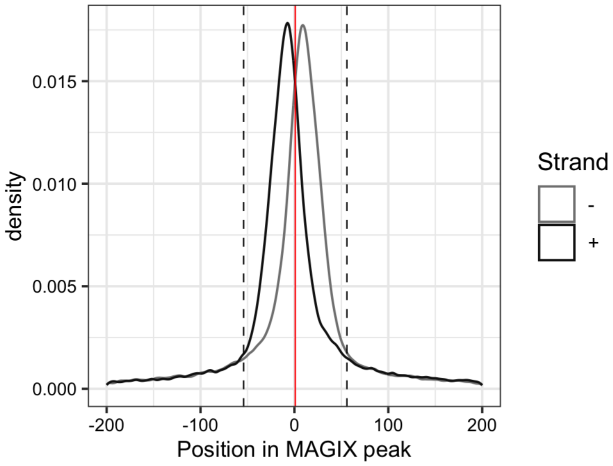

### ZNF865.DBD2

Top GRECO-BIT/MEX PWM: PWM006139

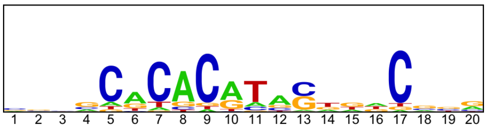


------------------------------------------------------------------------------------------------

Location of top GRECO-BIT/MEX PWM in MAGIX peaks (up to the top 10,000)

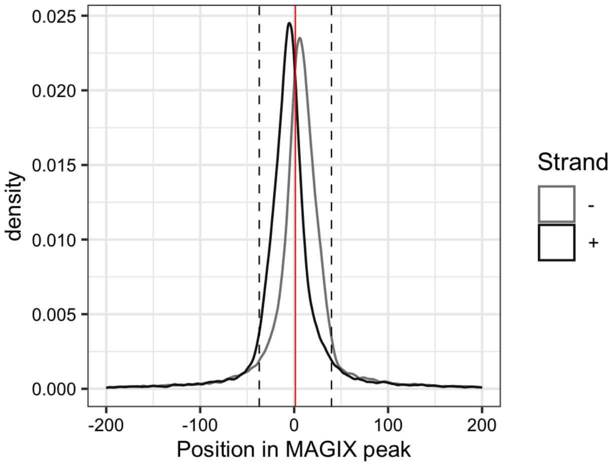

### ZSCAN2

Top GRECO-BIT/MEX PWM: PWM006244

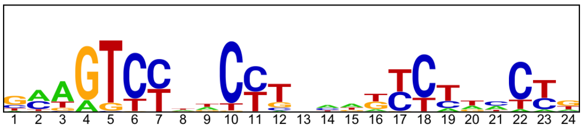


--------------------------------------------------------------------------------

Location of top GRECO-BIT/MEX PWM in MAGIX peaks (up to the top 10,000)

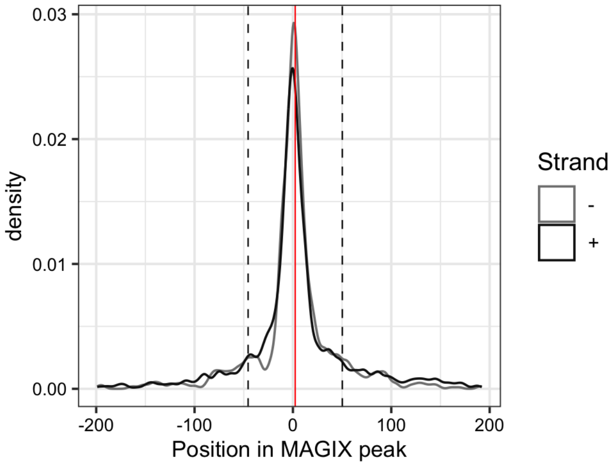


--------------------------------------------------------------------------------

AUROC of top GRECO-BIT/MEX PWM in sliding windows (500 peaks, 50 peak step) along the MAGIX peaks (up to the top 75,000) ranked by the MAGIX enrichment coefficient and fraction of overlapping ChIP peaks in the same window

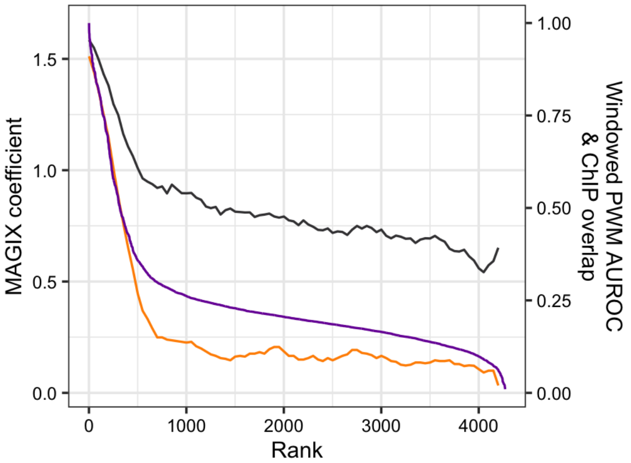

### ZSCAN25

Top GRECO-BIT/MEX PWM: PWM006291

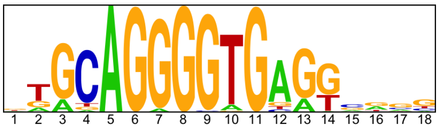


---

Location of top GRECO-BIT/MEX PWM in MAGIX peaks (up to the top 10,000)

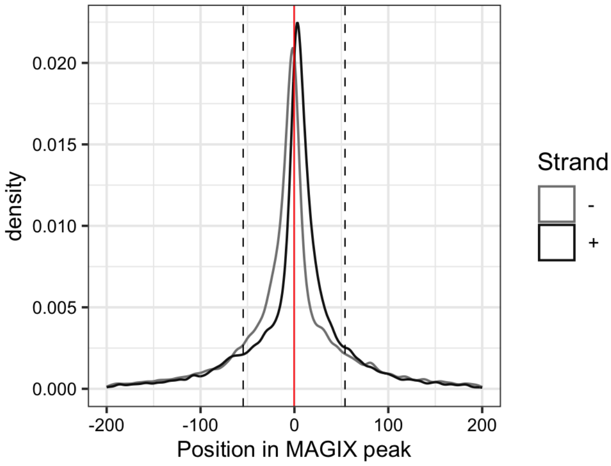


---

AUROC of top GRECO-BIT/MEX PWM in sliding windows (500 peaks, 50 peak step) along the MAGIX peaks (up to the top 75,000) ranked by the MAGIX enrichment coefficient and fraction of overlapping ChIP peaks in the same window

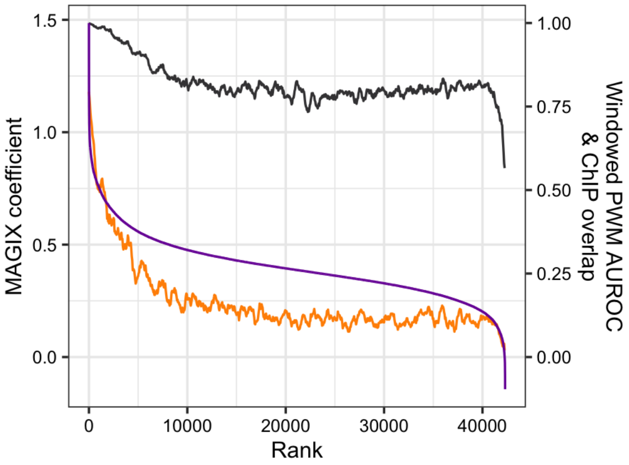

### ZSCAN4

Top GRECO-BIT/MEX PWM: PWM006306

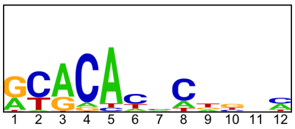


-----------------------------------------------------------------------------------------------

Location of top GRECO-BIT/MEX PWM in MAGIX peaks (up to the top 10,000)

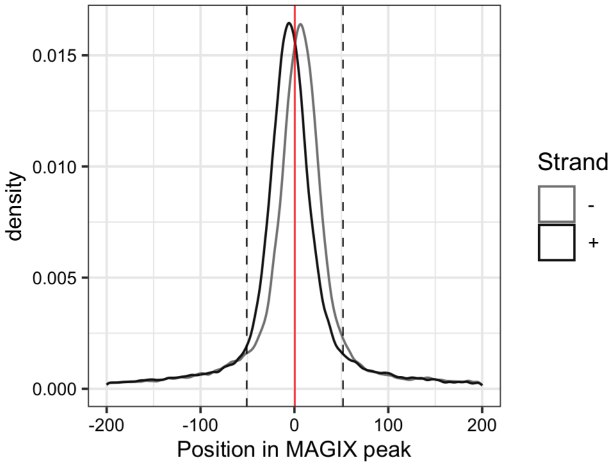
