## Supplementary Document S3 for "GHT-SELEX demonstrates unexpectedly high intrinsic sequence specificity and complex DNA binding of many human transcription factors"

### Enrichment of k-mers between HT-SELEX replicate experiments

This document displays plots of k-mers enriched in experiments for the same TF expressed in different or same expression system and describes what is the most likely explanation for differences seen in kmers. See **Table S6** for details of each pairwise comparison.

**C and G rich sequences**
There is a fairly common tendency for low level poly-C/poly-G sequences to be enriched in one sample but not the other. This outcome is however uncorrelated with the protein expression system, or with any other experimental variable. The initial HT-SELEX pool it is biased towards the same sequences, presumably due to bias in the commercial DNA synthesis. Examples of this are visible for example in comparison on page: 4,

**Enrichment of sequences that are complementary to Illumina flanking regions**
In this artifact the experiment enriches sequences that have partial matches to constant regions that are used in the Illumina sequencing. We do not know the mechanism behind this but it is likely derived from selection artifact of mis-hybridized ligands and occurs due to presence of partially single stranded DNA in the experiments. A common kmer like this is ACACGACGC. This is clearly visible for example in experiment pair on page: 26, where it enriches alongside genuine target site for DACH1.

**Enrichment of mono- and homodimeric binding sites for FLI1**
FLI1, a positive control, is known to bind both as mono- and homodimers, and selection pressure for them differs based on protein concentration as higher concentration of TF will drive enrichment of homodimer (See Jolma et al. 2013, PMID: 23332764); we observe what appears to be differential abundance of the two modes in different experiments. An example of an experiment pair with similar enrichment for both monomeric and dimeric FLI1 target sites is on page 33 with example 9-mers being ACCGGAAGT (monomer) and  CGGATATCC (dimer). This differential enrichment is visible in comparisons on pages: 33, 37, 38 and 39.

**Enrichment of aptameric partially single stranded DNA molecules in experiments for SOX-proteins**
SOX proteins can bind both doublestranded DNA and short aptamers formed from partially single stranded DNA and this is visible in several experiments for SRY, SOX15 and  SOX2 proteins. select commonly sequences that are likely to be stem-loop DNA aptamers formed from partially single stranded DNA (also previously described in Jolma et al. 2013), because the bases have symmetric dependencies on the opposite sides of the motif. An example of a commonly enriched aptameric 9-mer is AATGACATT whereas GAACAATGG is a commonly entriched 9-mer with a double stranded DNA target site. This differential enrichment is visible in experiment comparisons on pages: SOX15: 140, 143, 145 and 146; SOX2: 148, 150, 151, 153, 154 and 156; SRY: 178, 180 and 181

**CAMTA1**

Lysate vs Lysate, same construct

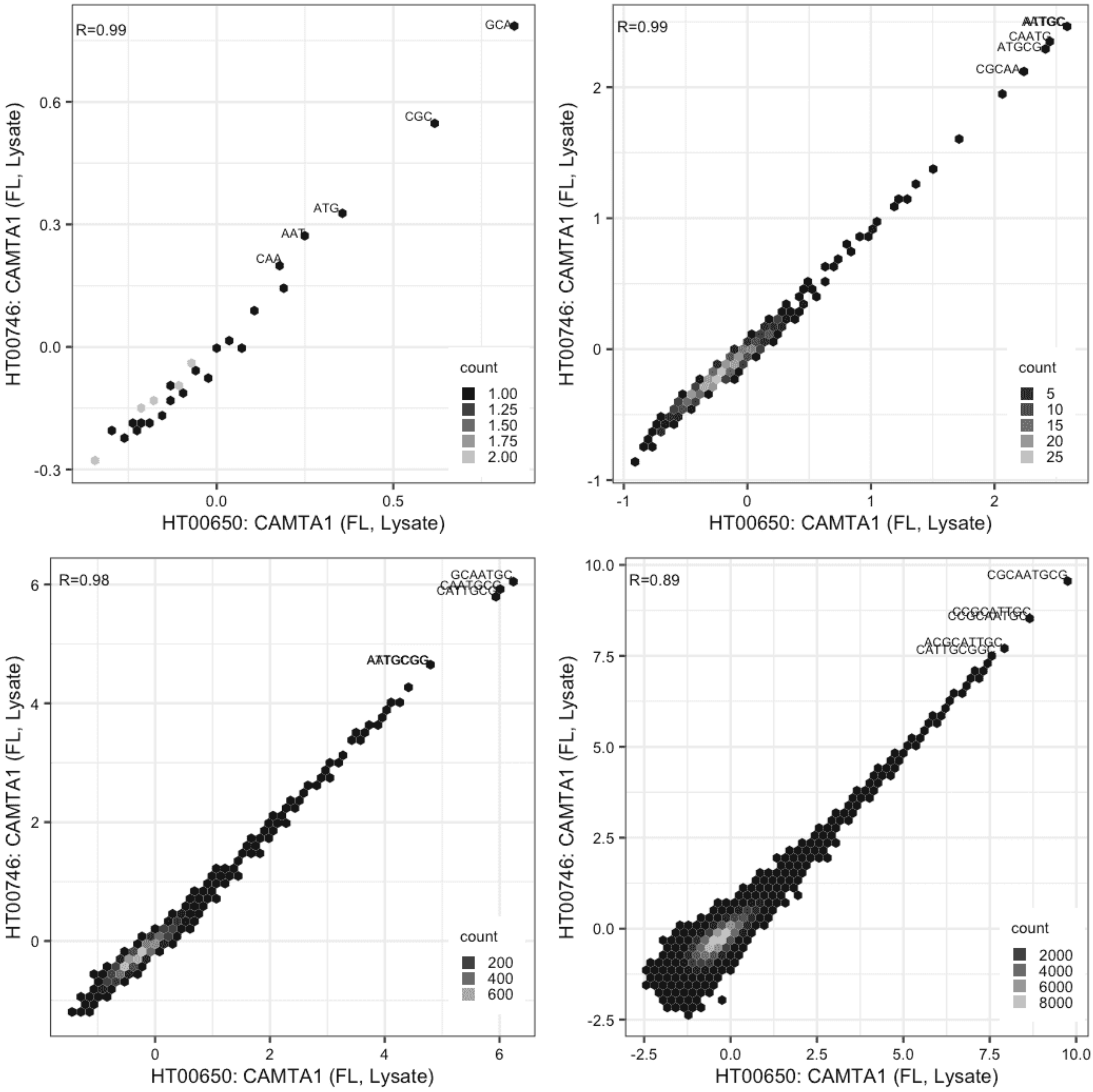

**CAMTA1**

Lysate vs GFPIVT, different constructs

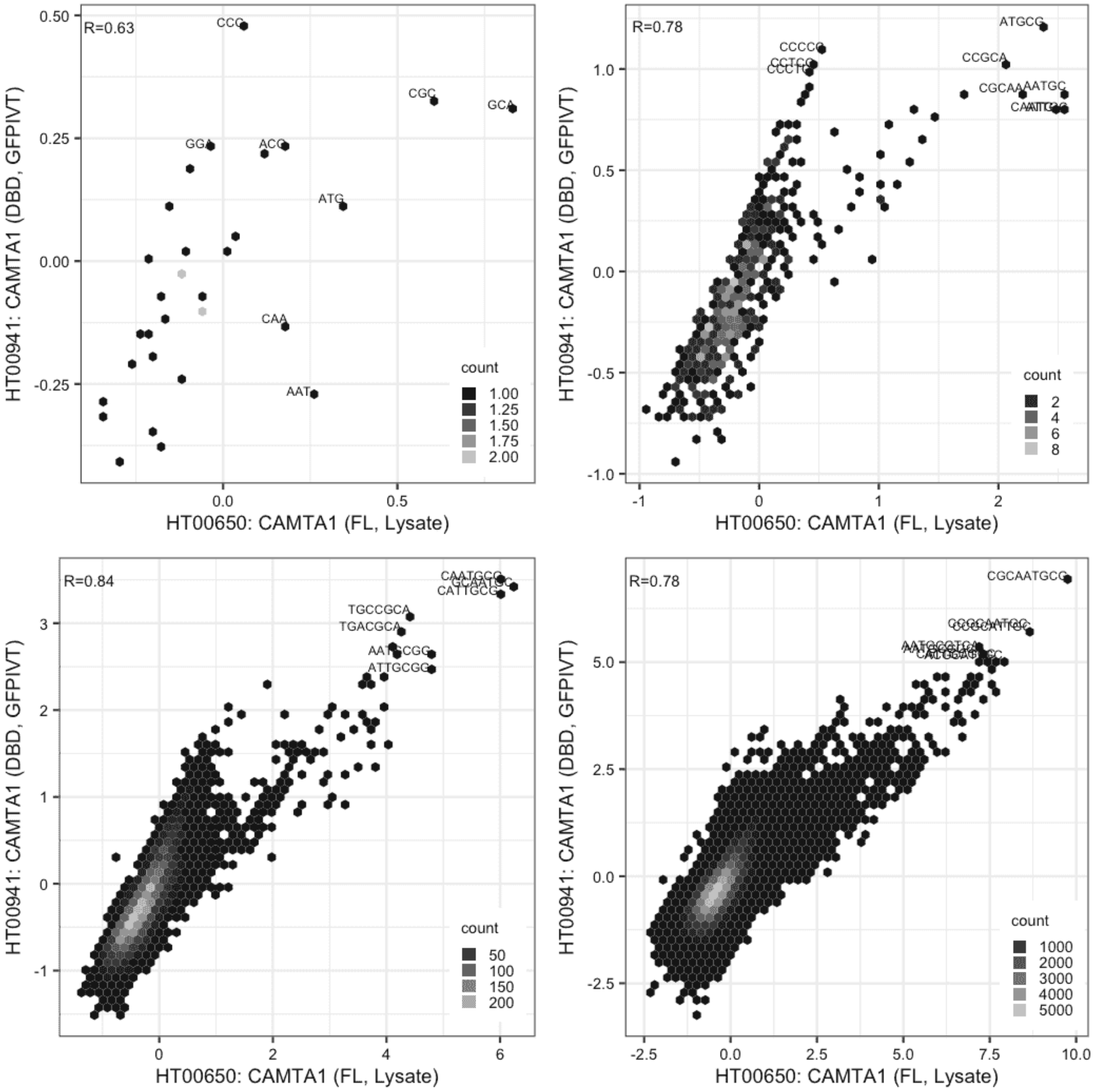

**CAMTA1**

Lysate vs GFPIVT, different constructs

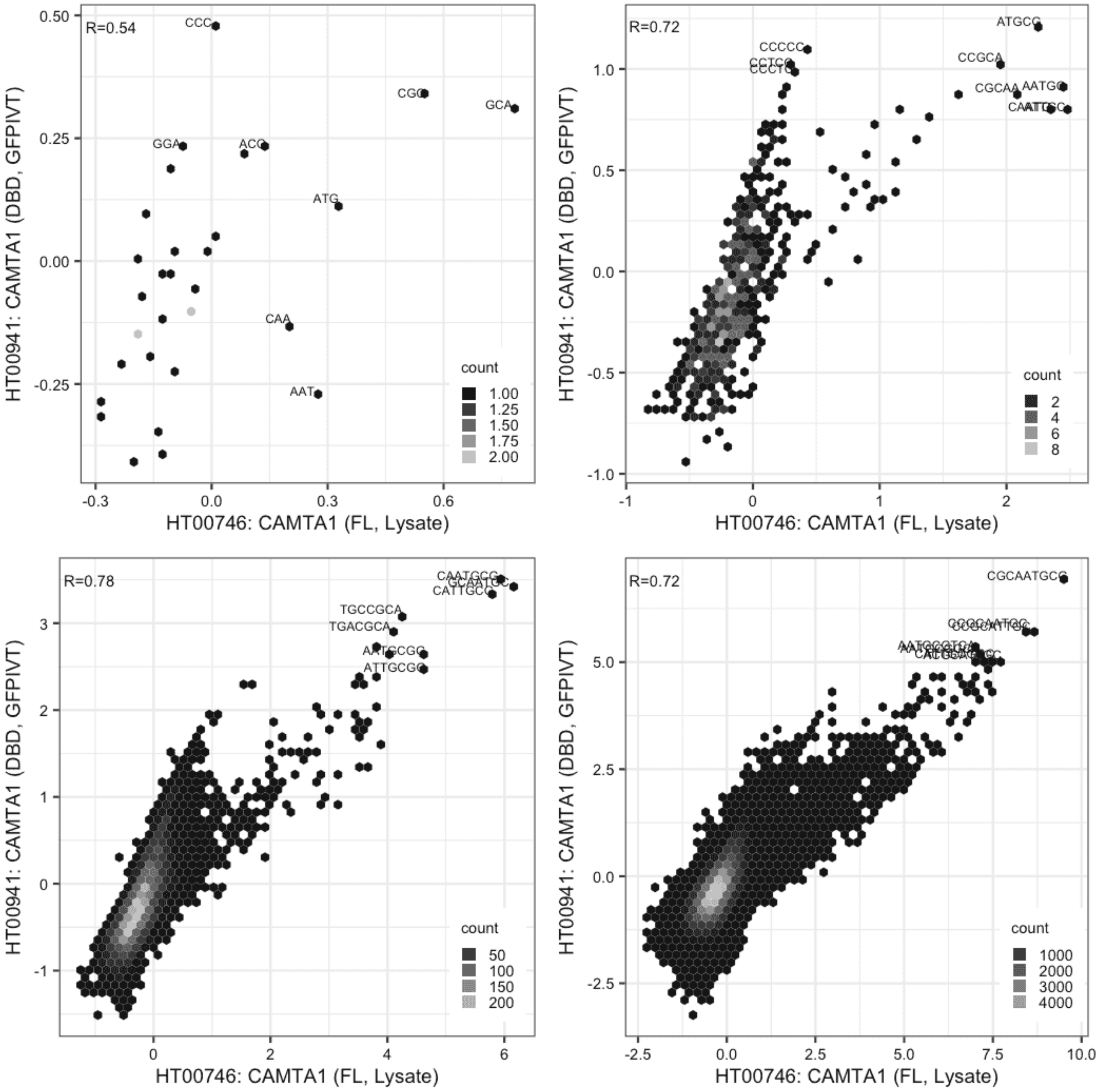

**CASZ1**

Lysate vs Lysate, same construct

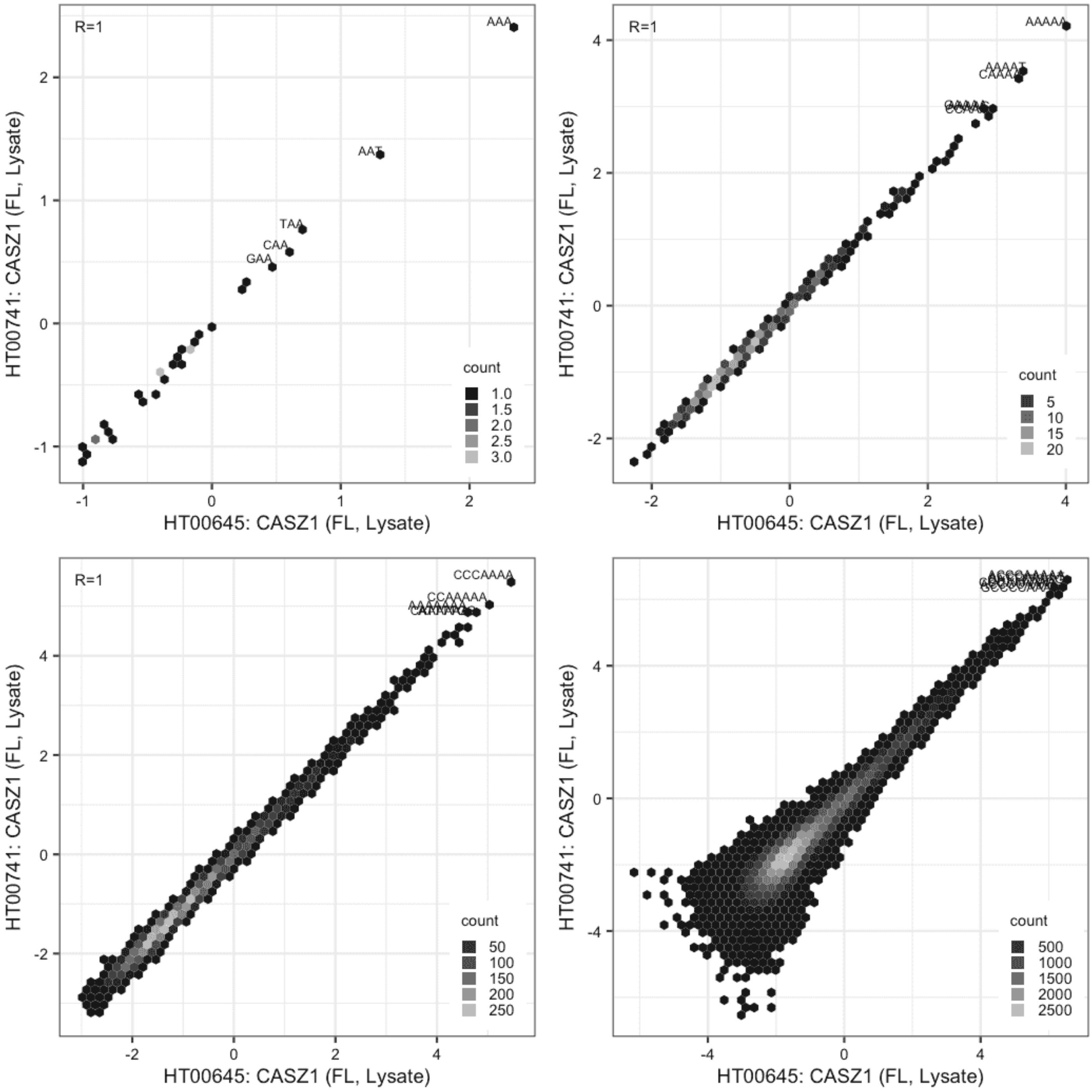

**CASZ1**

Lysate vs GFPIVT, same construct

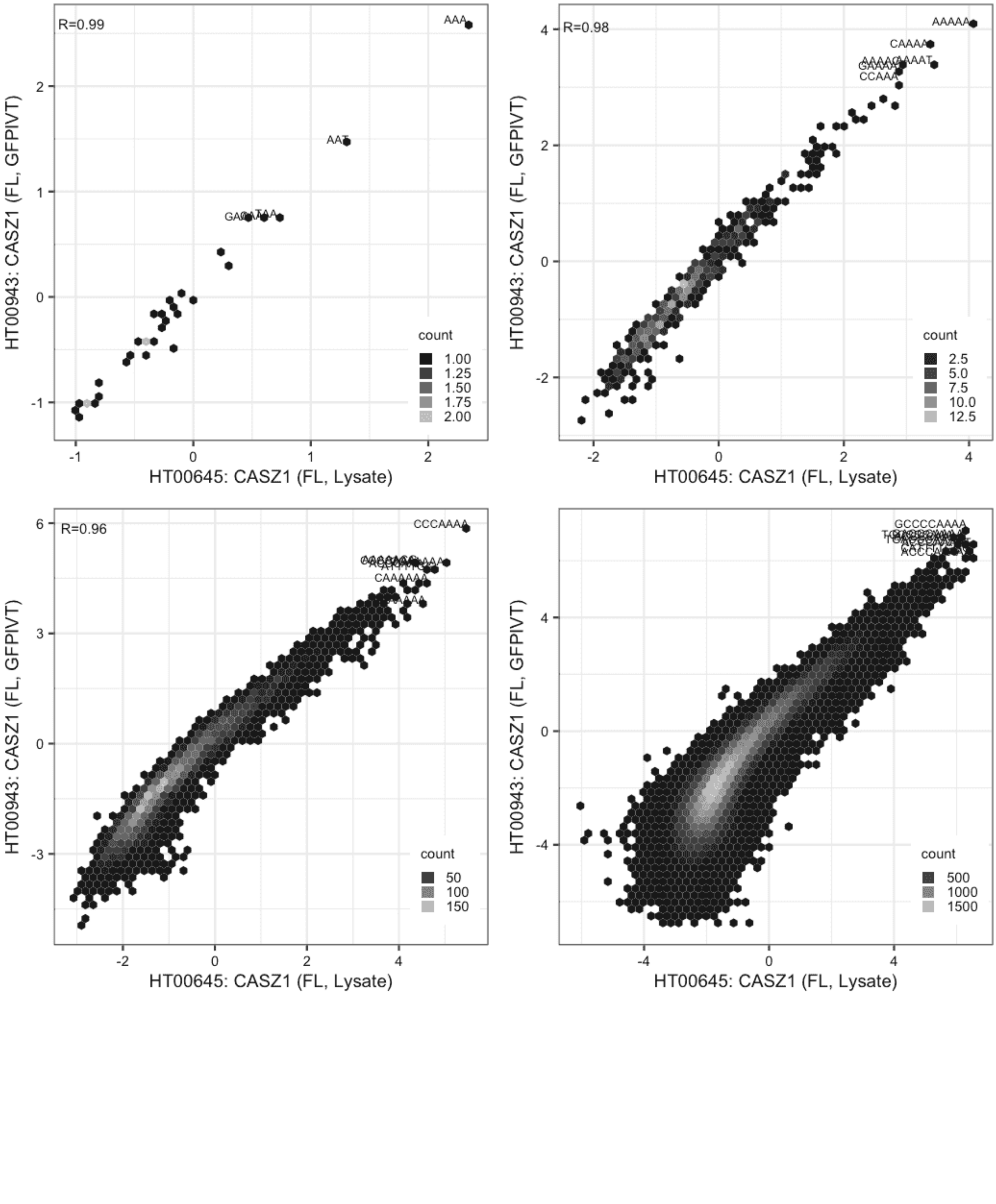

**CASZ1**

Lysate vs GFPIVT, same construct

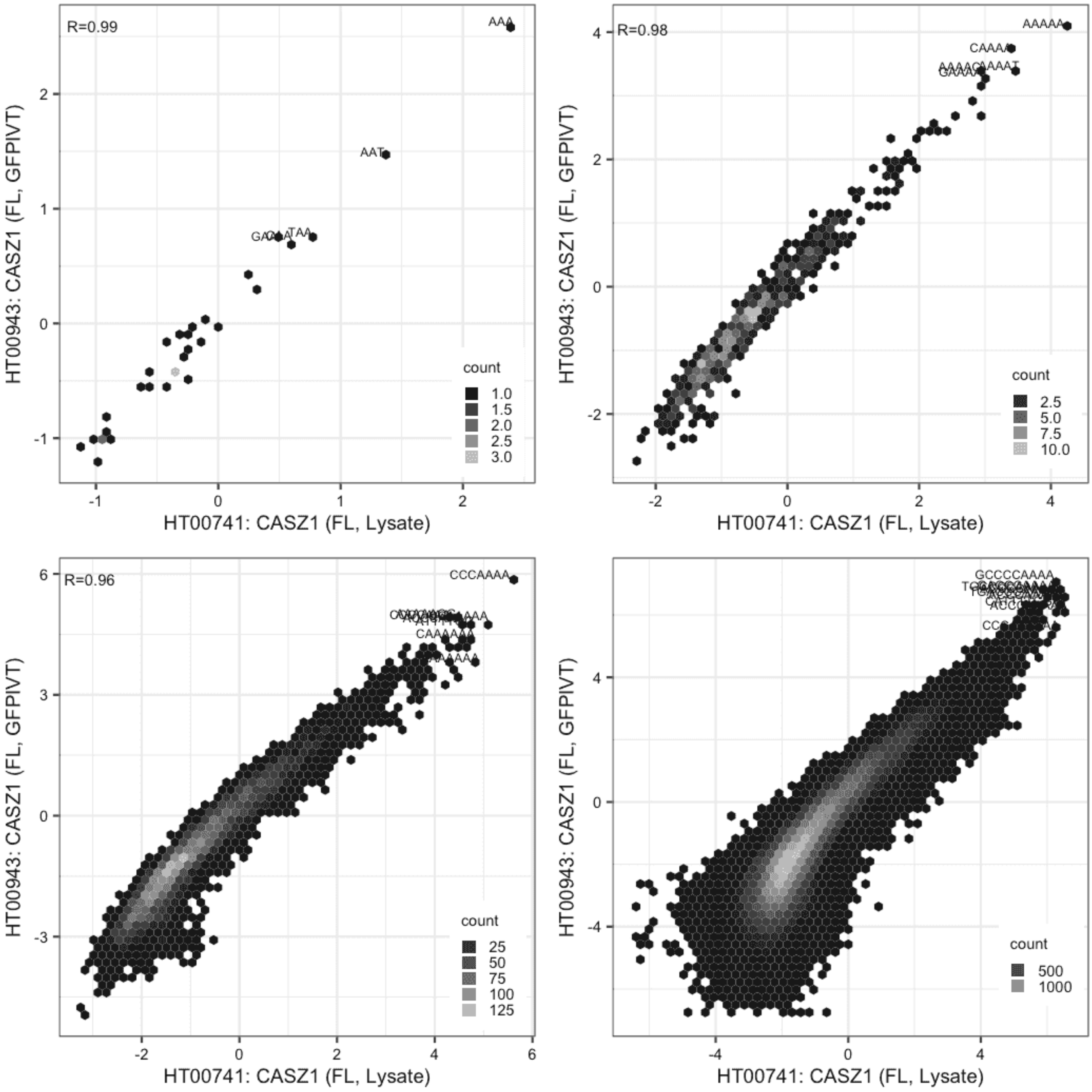

#### CREB3L3

IVT vs GFPIVT, different constructs

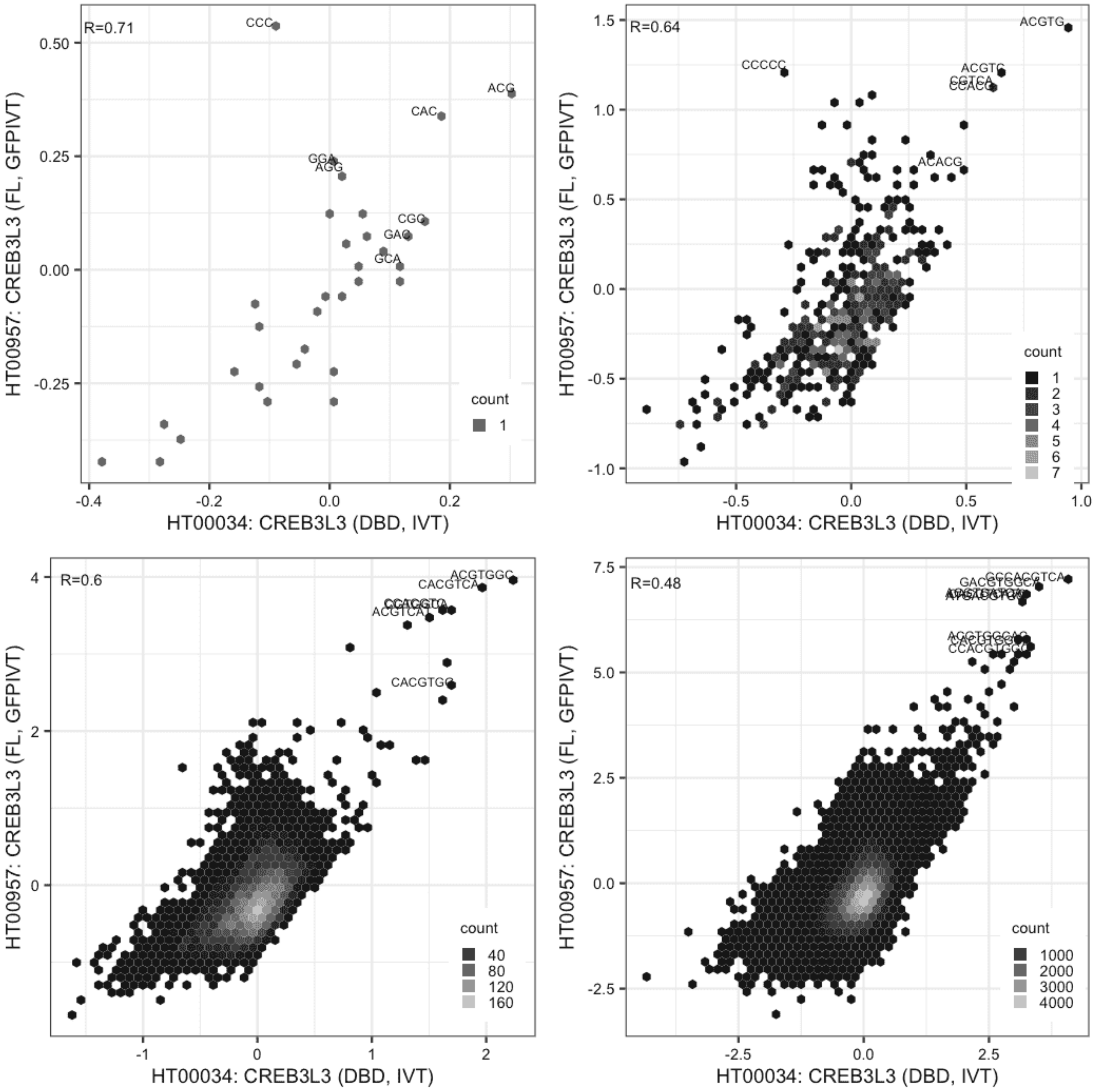

**CREB3L3**

Lysate vs IVT, different constructs

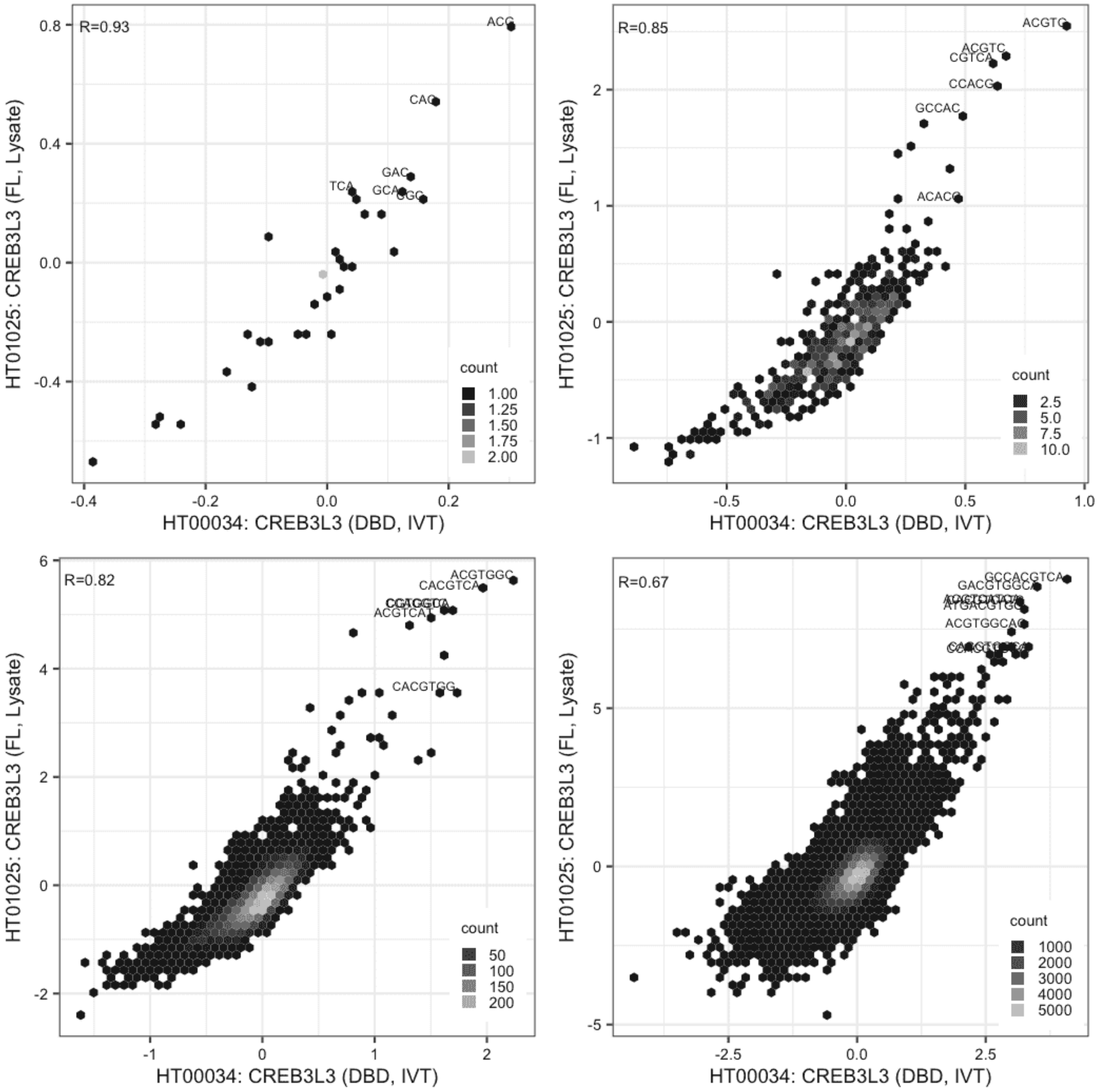

#### CREB3L3

Lysate vs IVT, different constructs

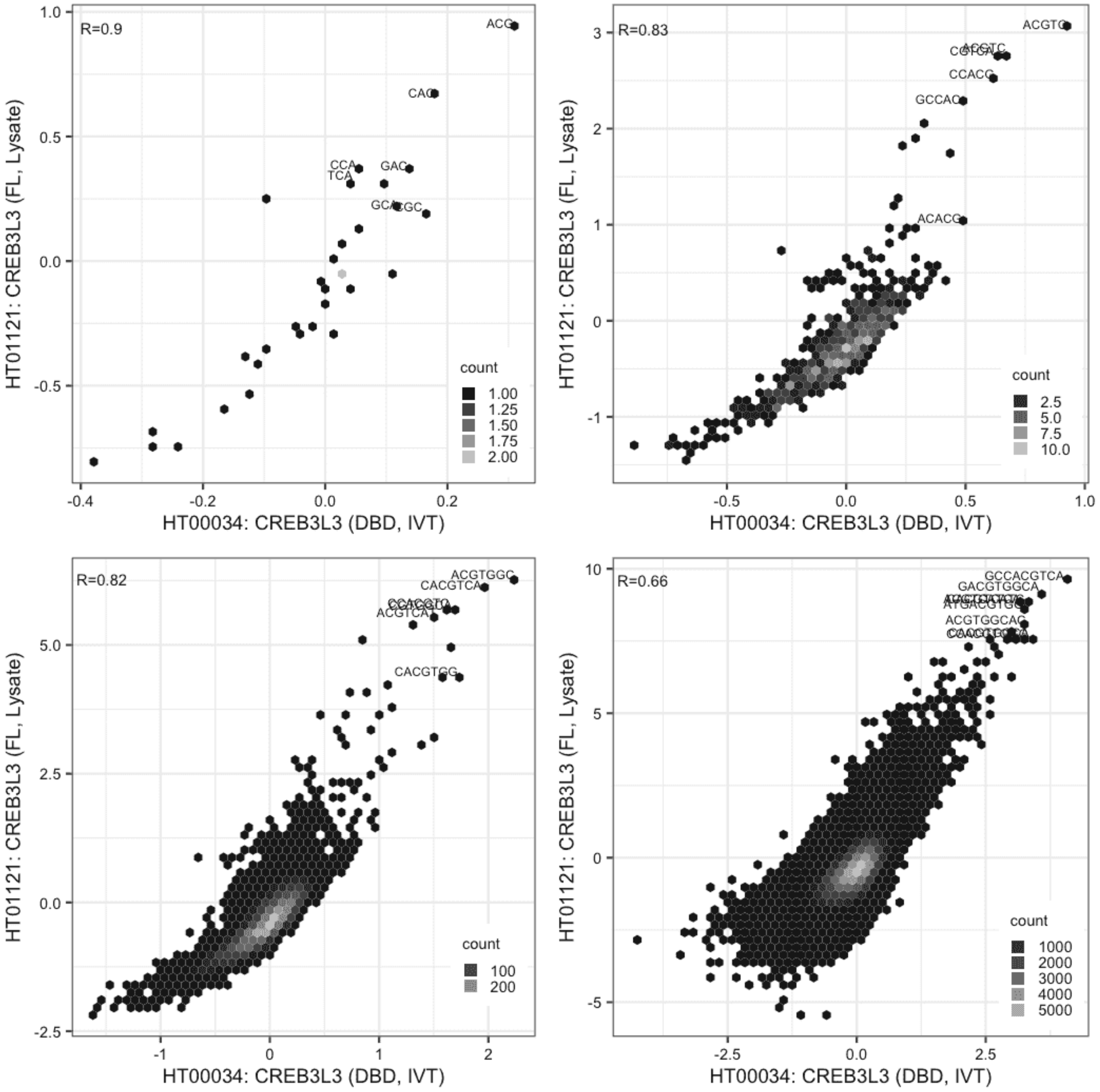

#### CREB3L3

Lysate vs GFPIVT, same construct

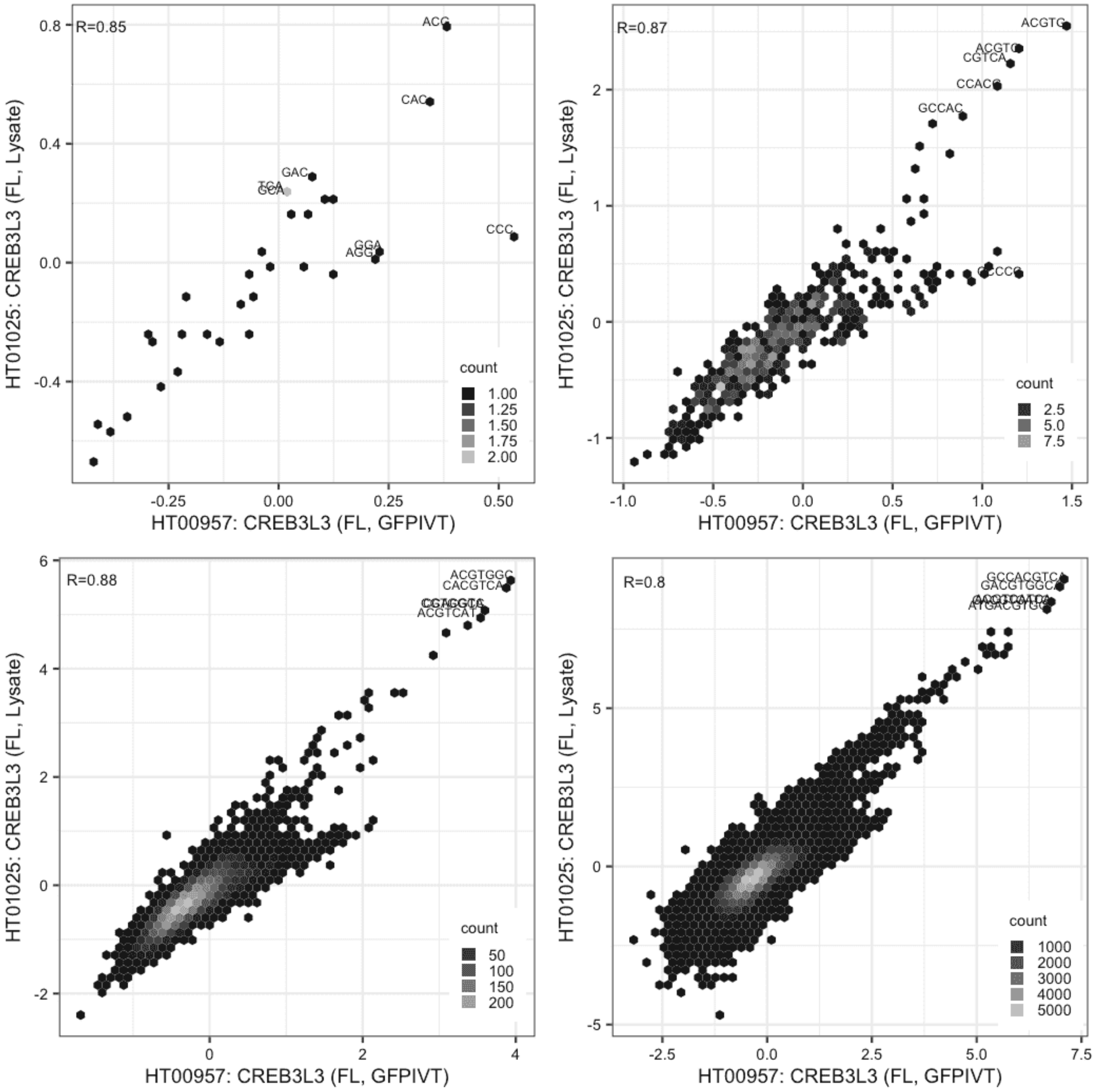

#### CREB3L3

Lysate vs GFPIVT, same construct

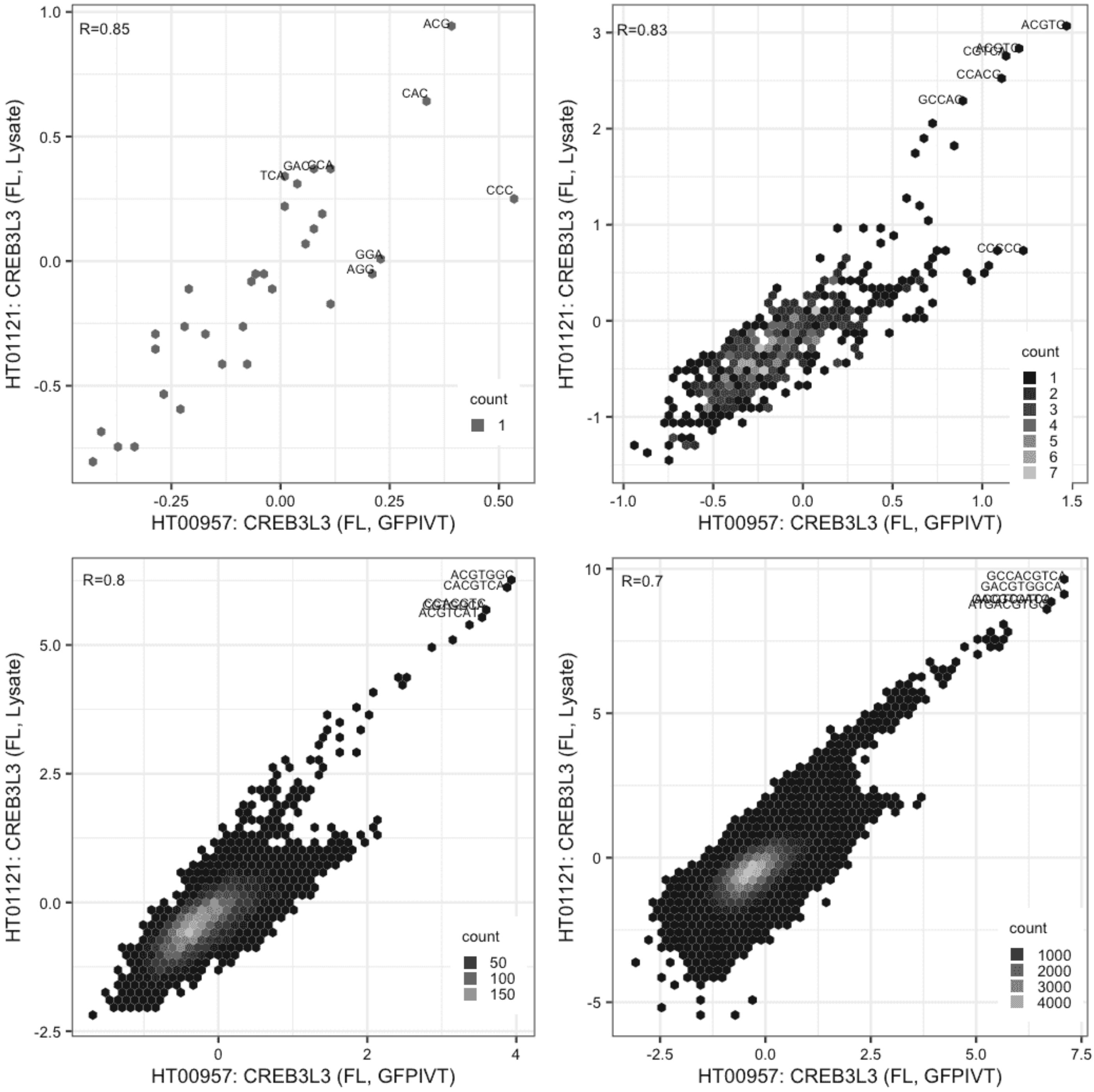

### CREB3L3

Lysate vs Lysate, same construct

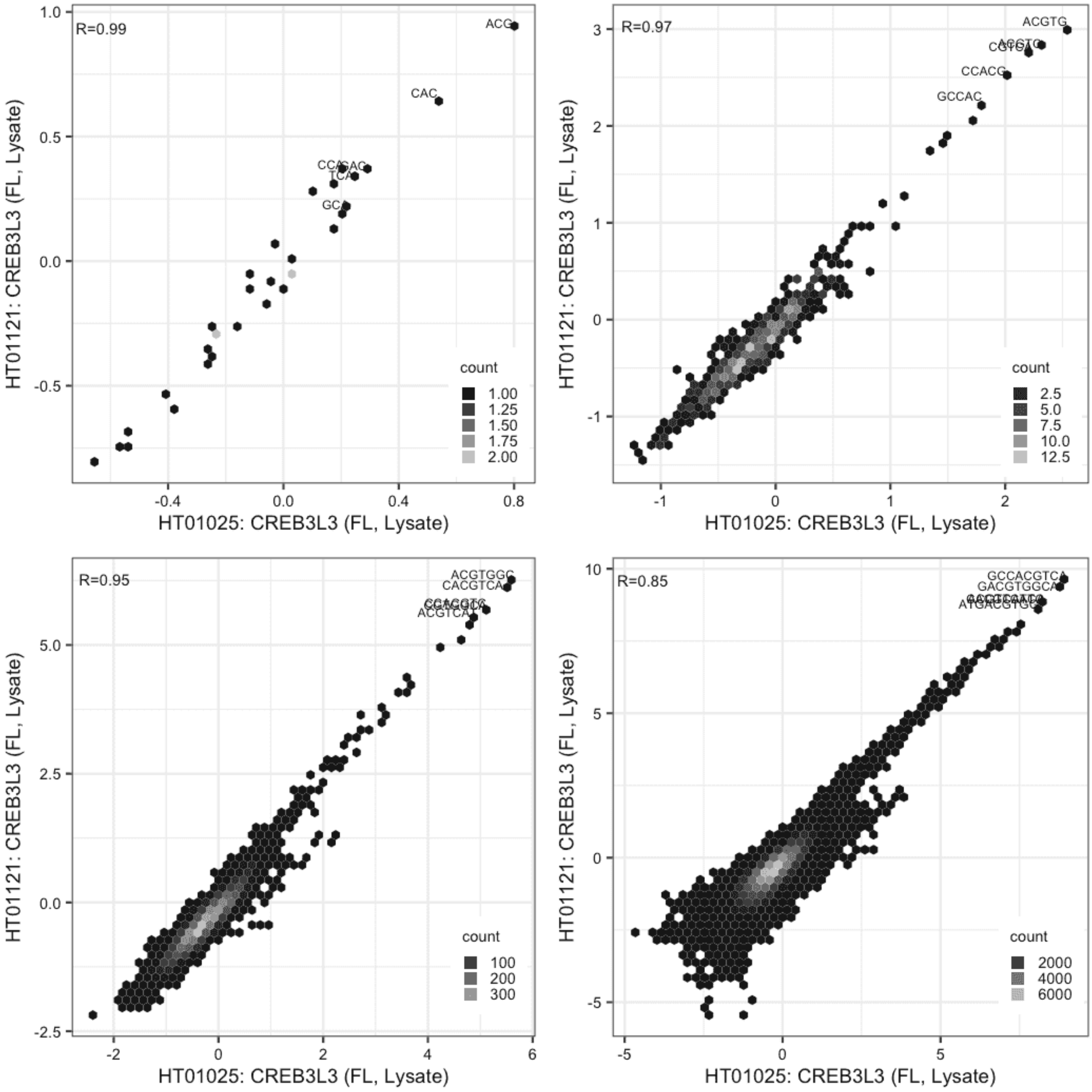

**CXXC4**

IVT vs IVT, different constructs

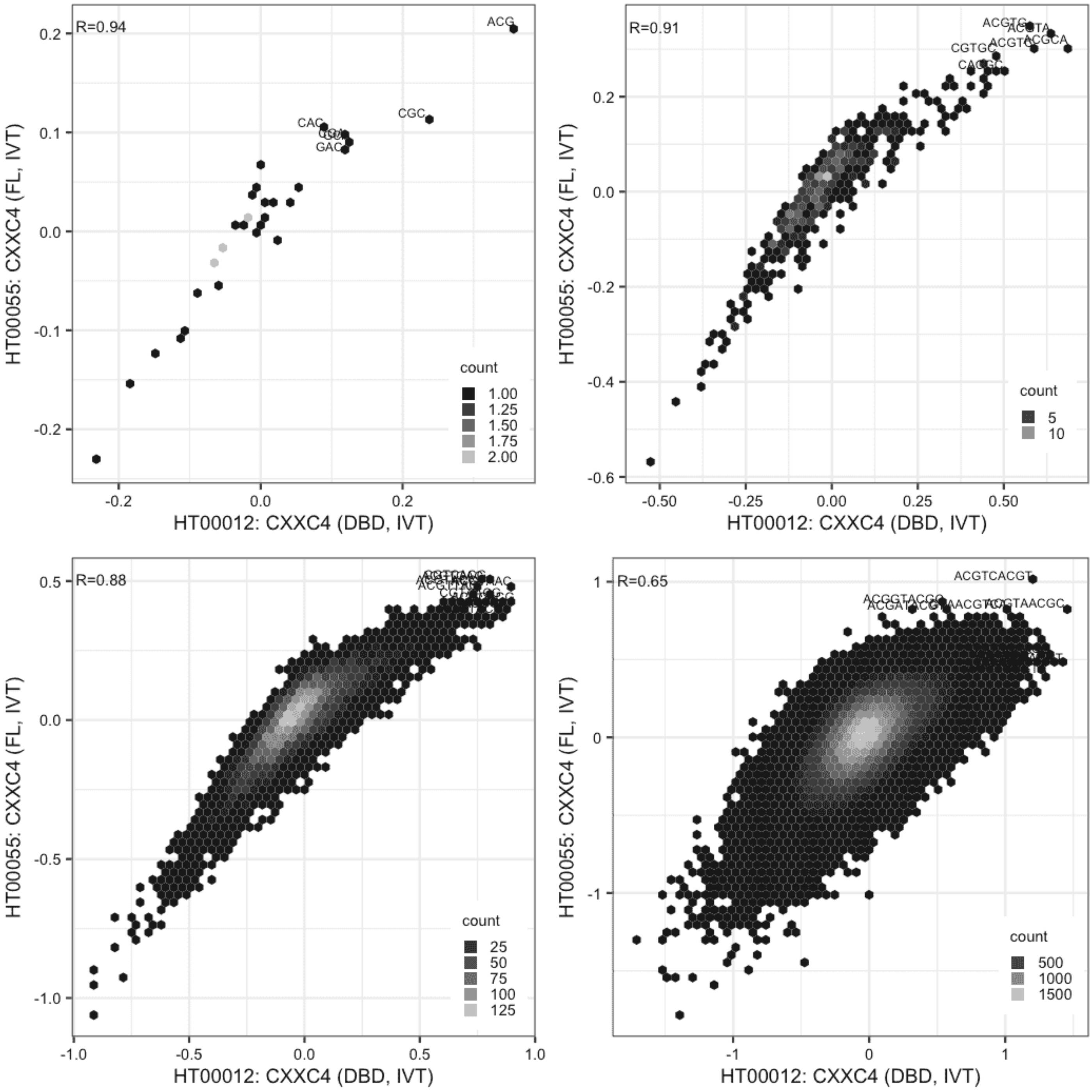

**CXXC4**

Lysate vs IVT, different constructs

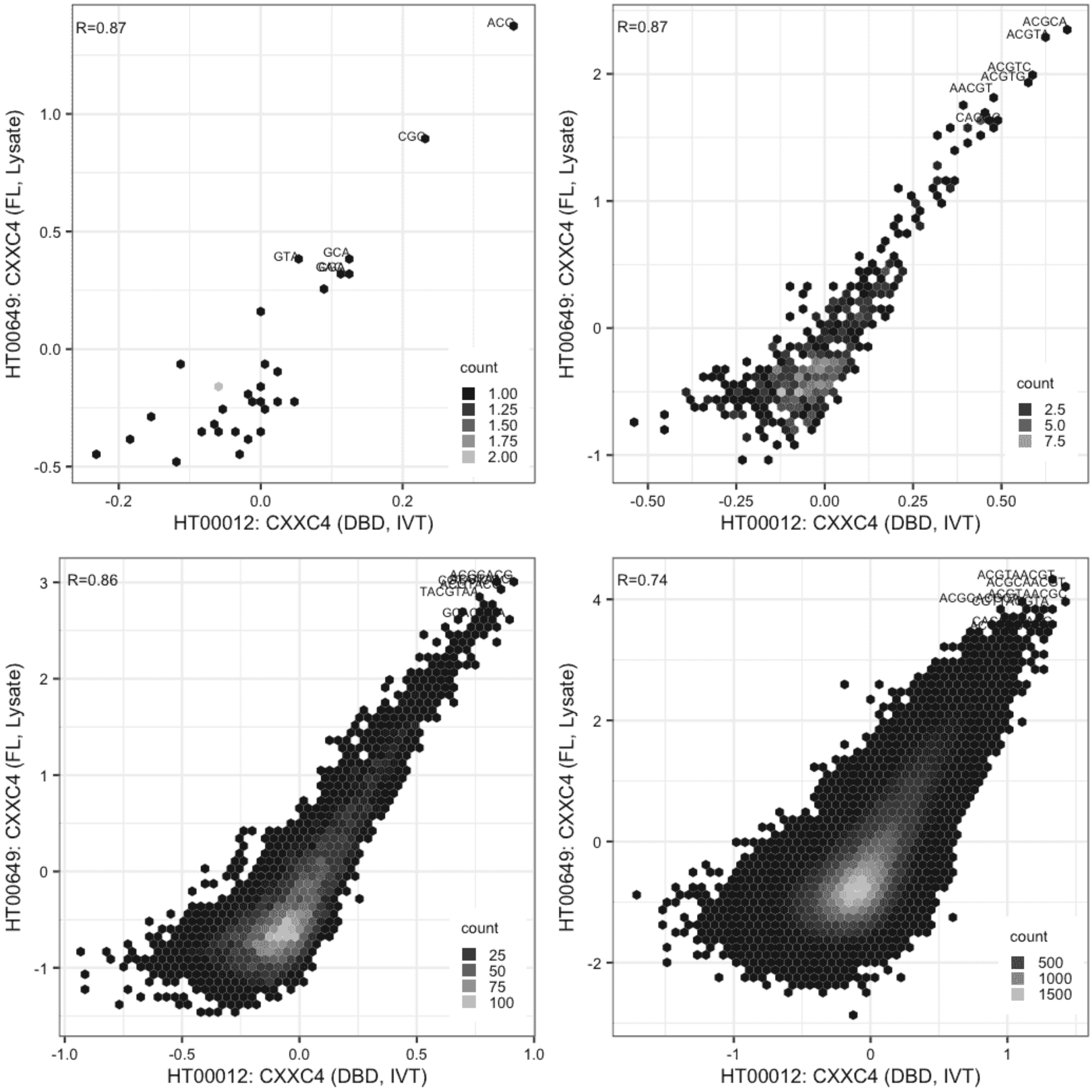

### CXXC4

Lysate vs IVT, different constructs

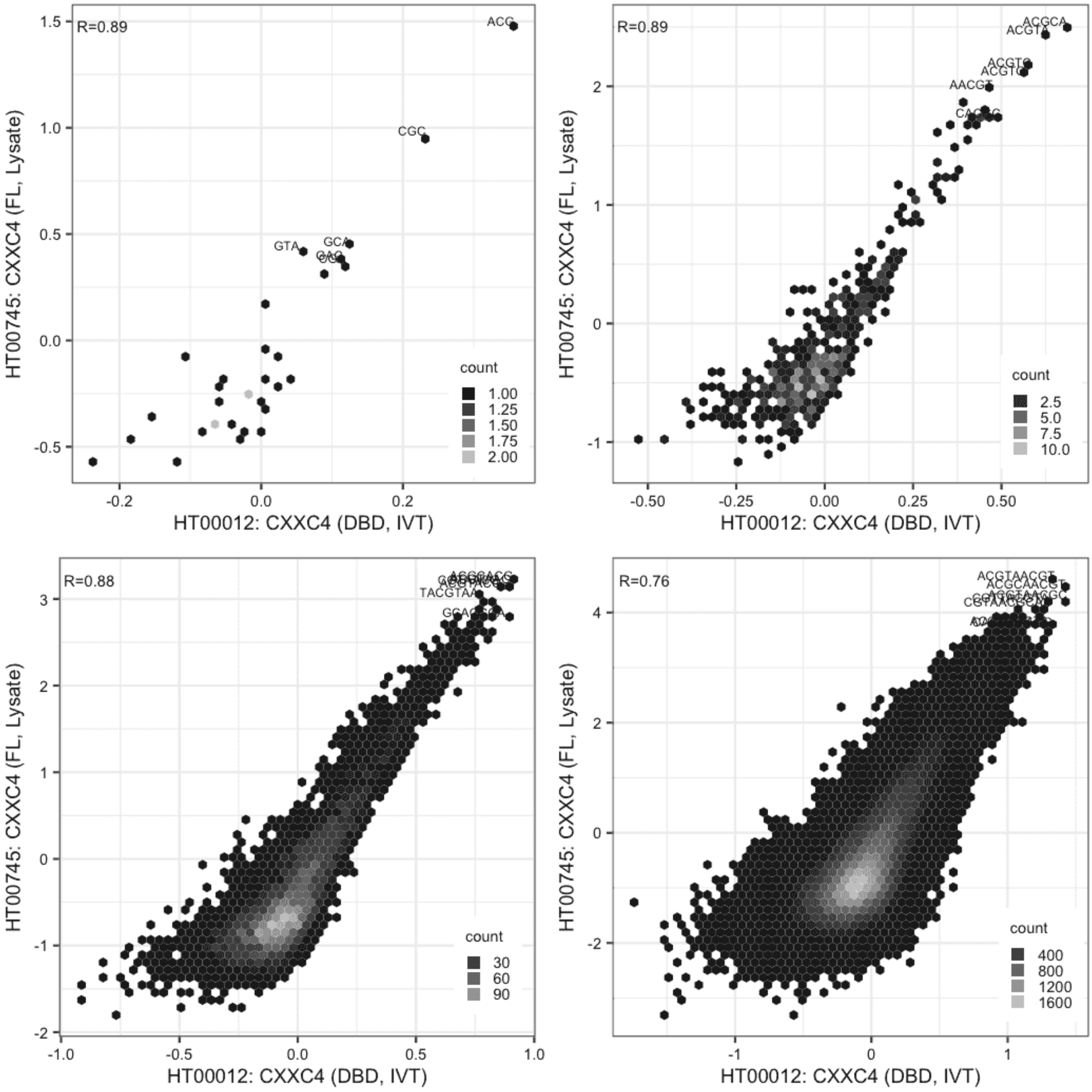

**CXXC4**

IVT vs GFPIVT, different constructs

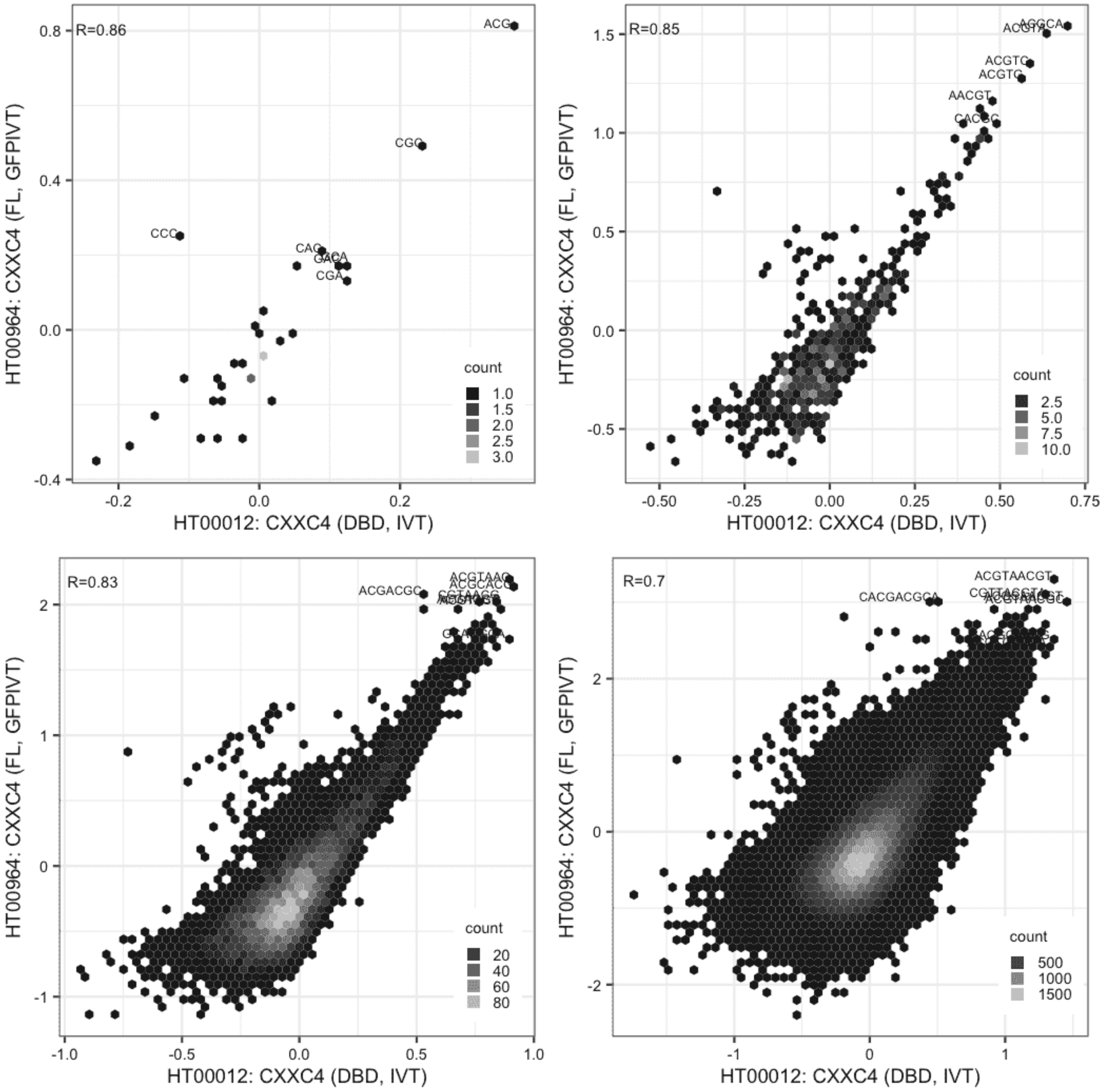

**CXXC4**

Lysate vs IVT, same construct

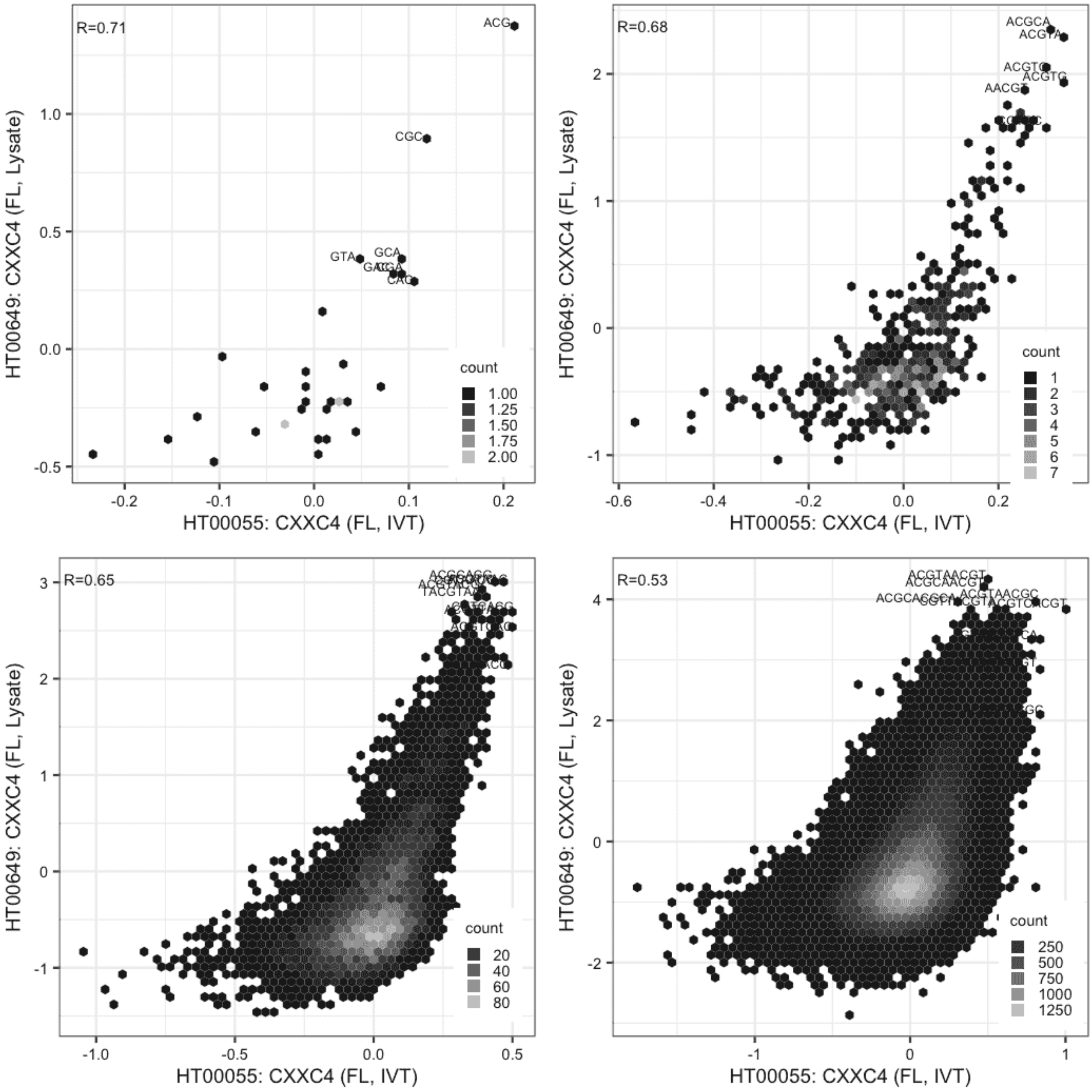

**CXXC4**

Lysate vs IVT, same construct

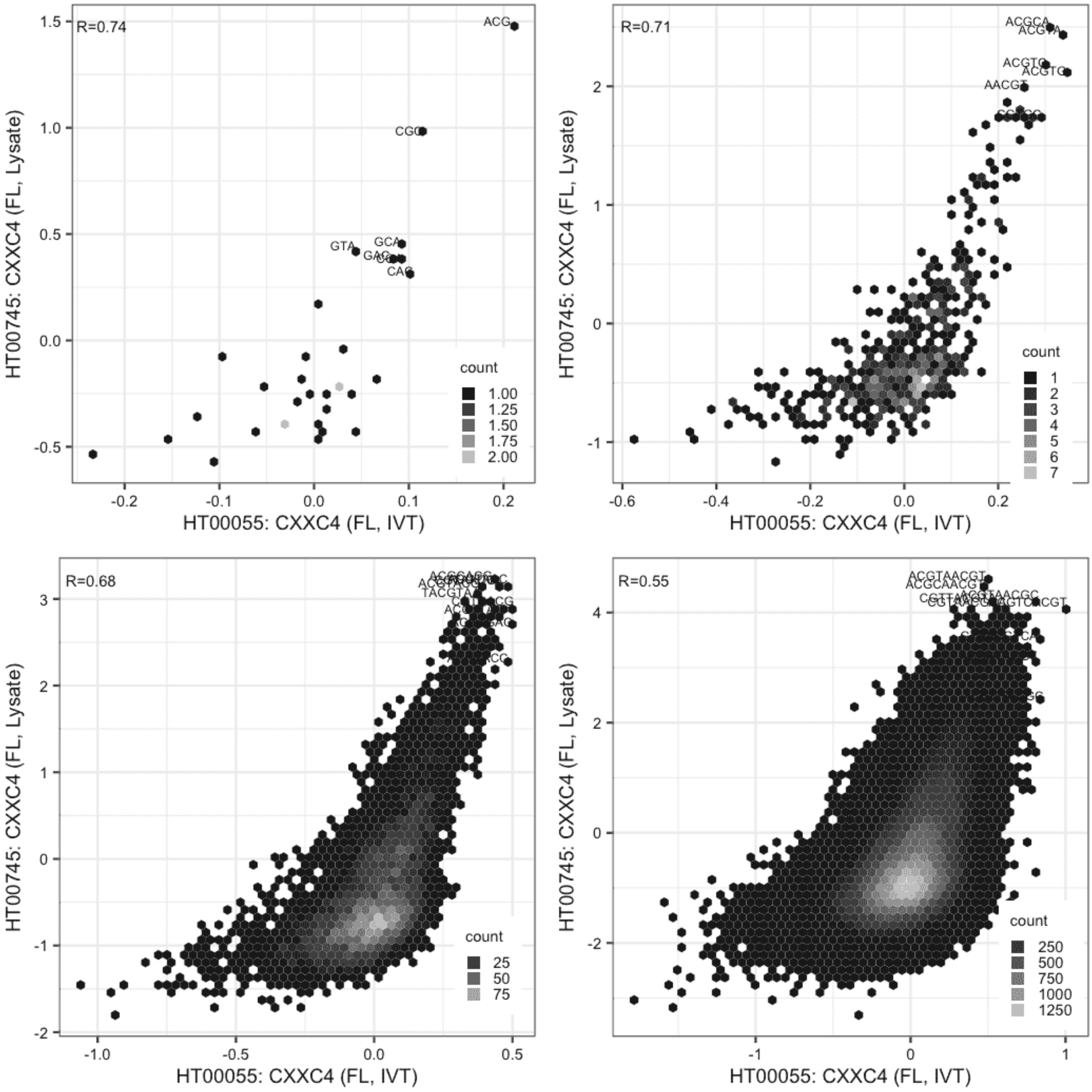

**CXXC4**

IVT vs GFPIVT, same construct

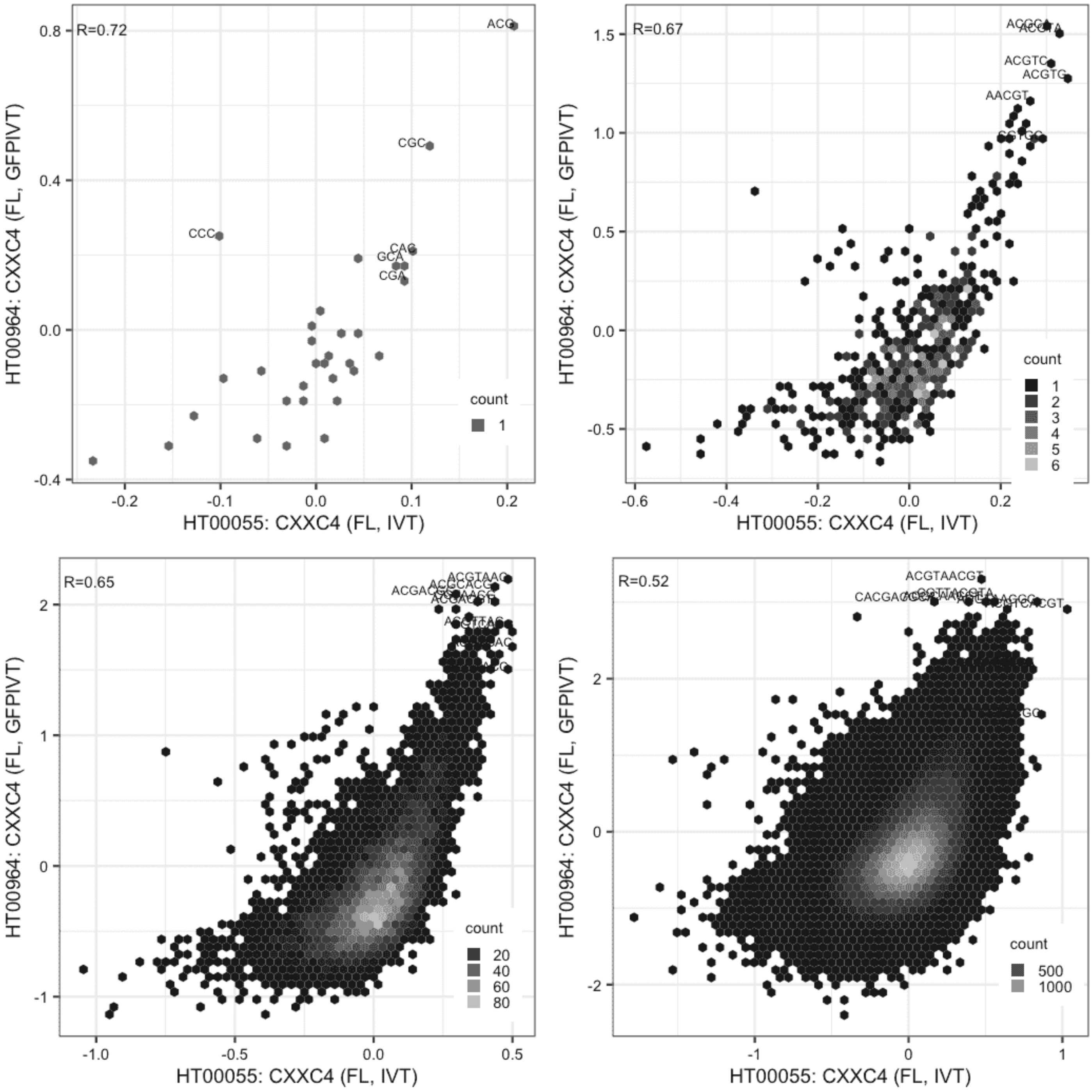

**CXXC4**

Lysate vs Lysate, same construct

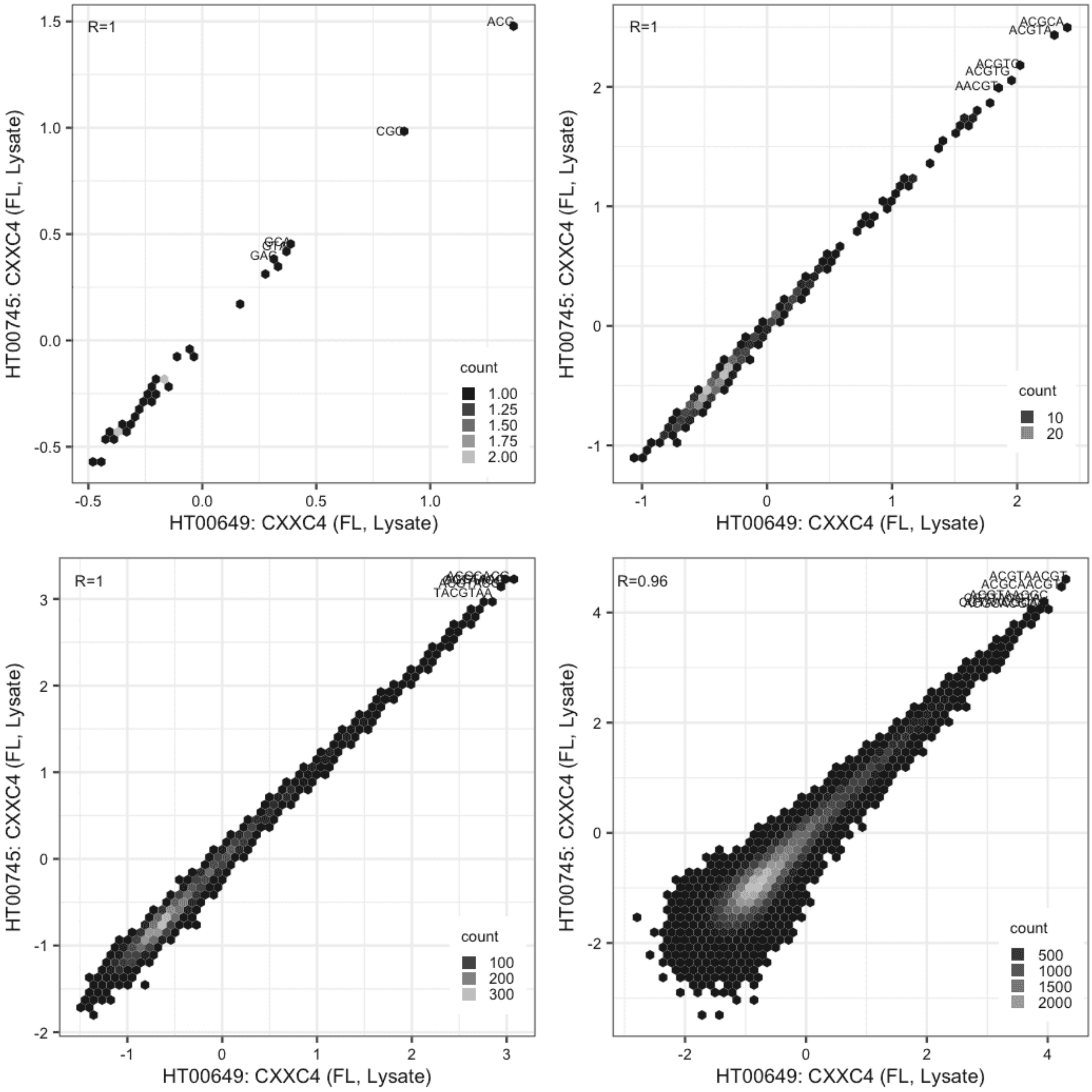

**CXXC4**

Lysate vs GFPIVT, same construct

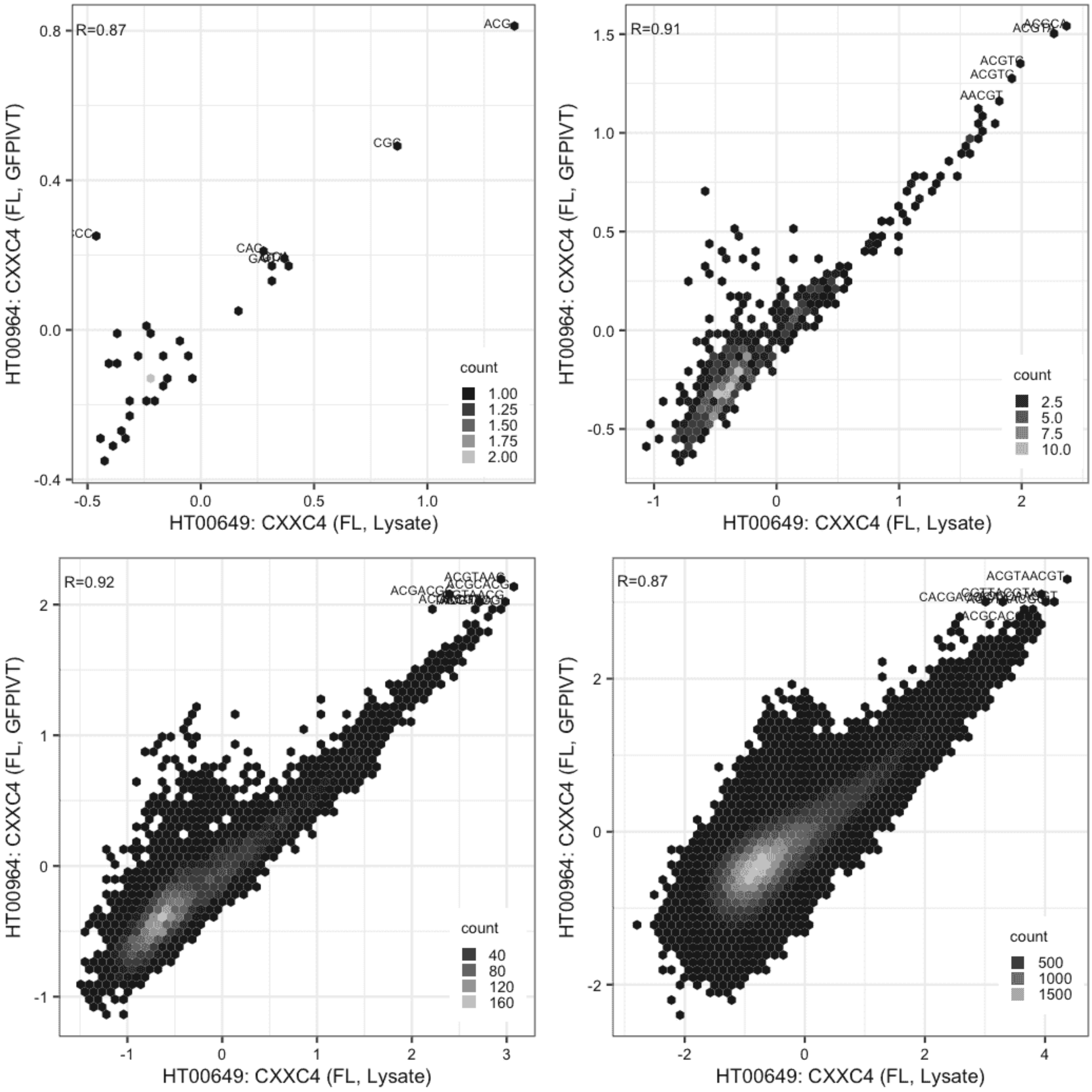

**CXXC4**

Lysate vs GFPIVT, same construct

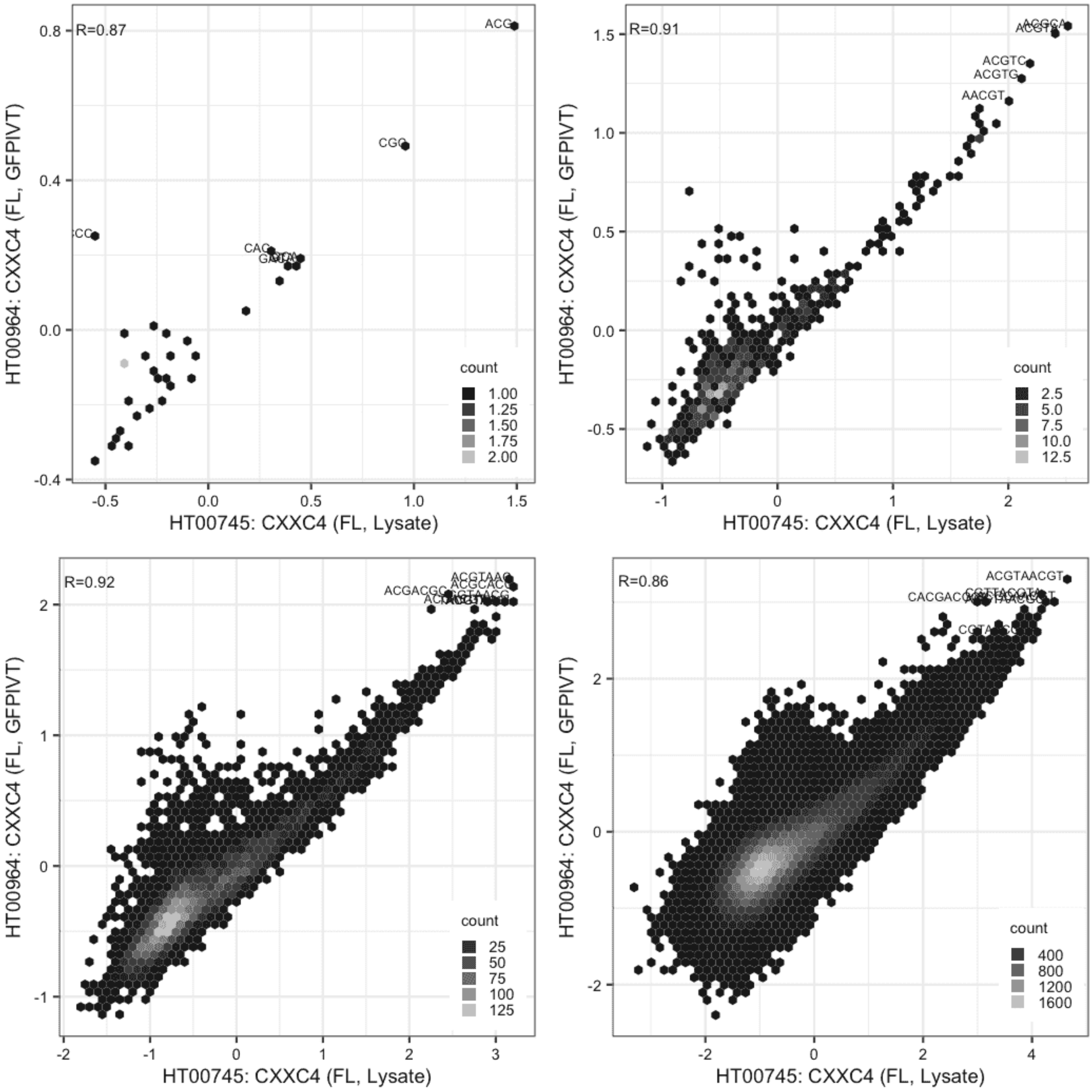

**DACH1**

Lysate vs GFPIVT, different constructs

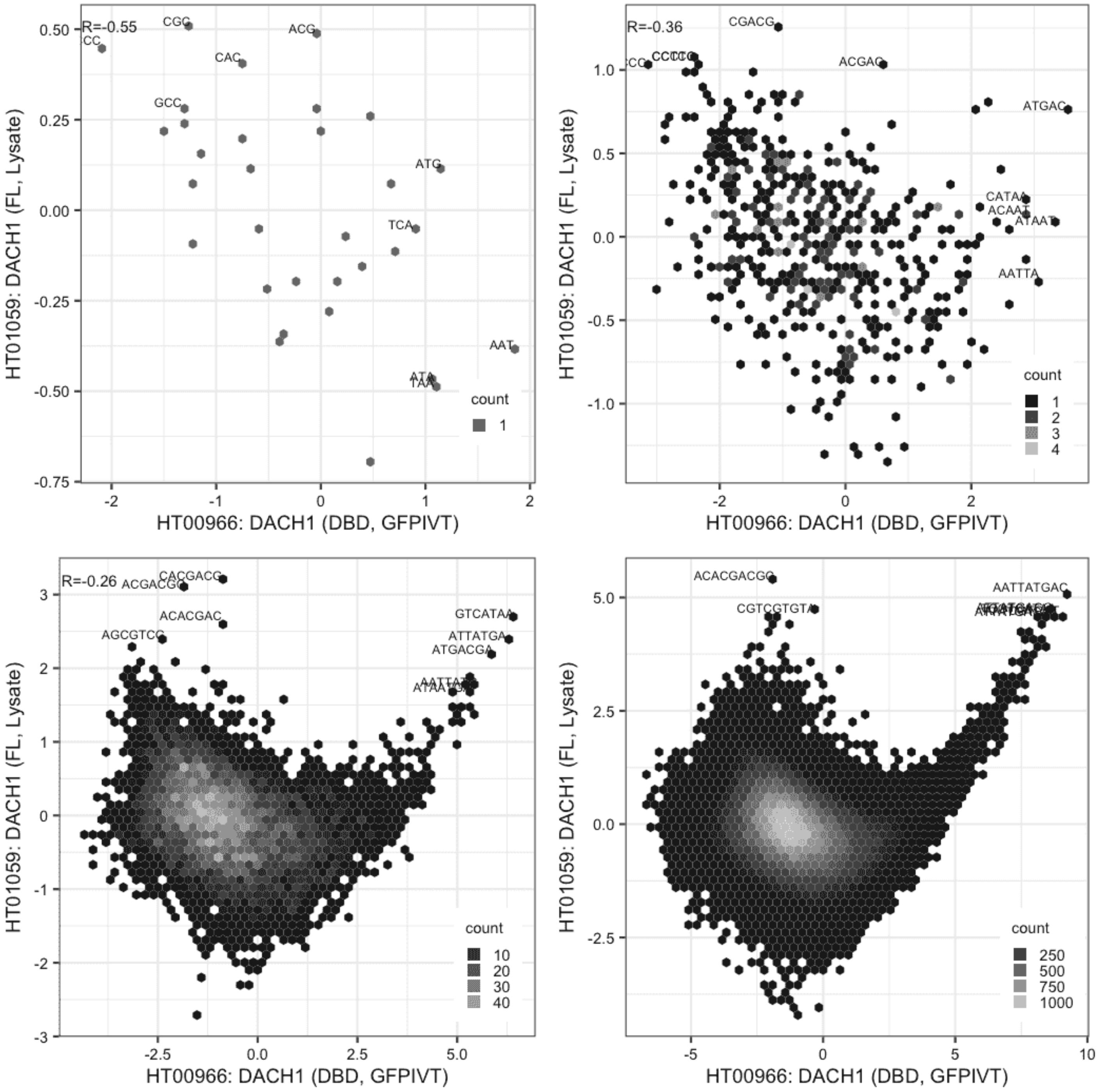

**DACH1**

Lysate vs GFPIVT, different constructs

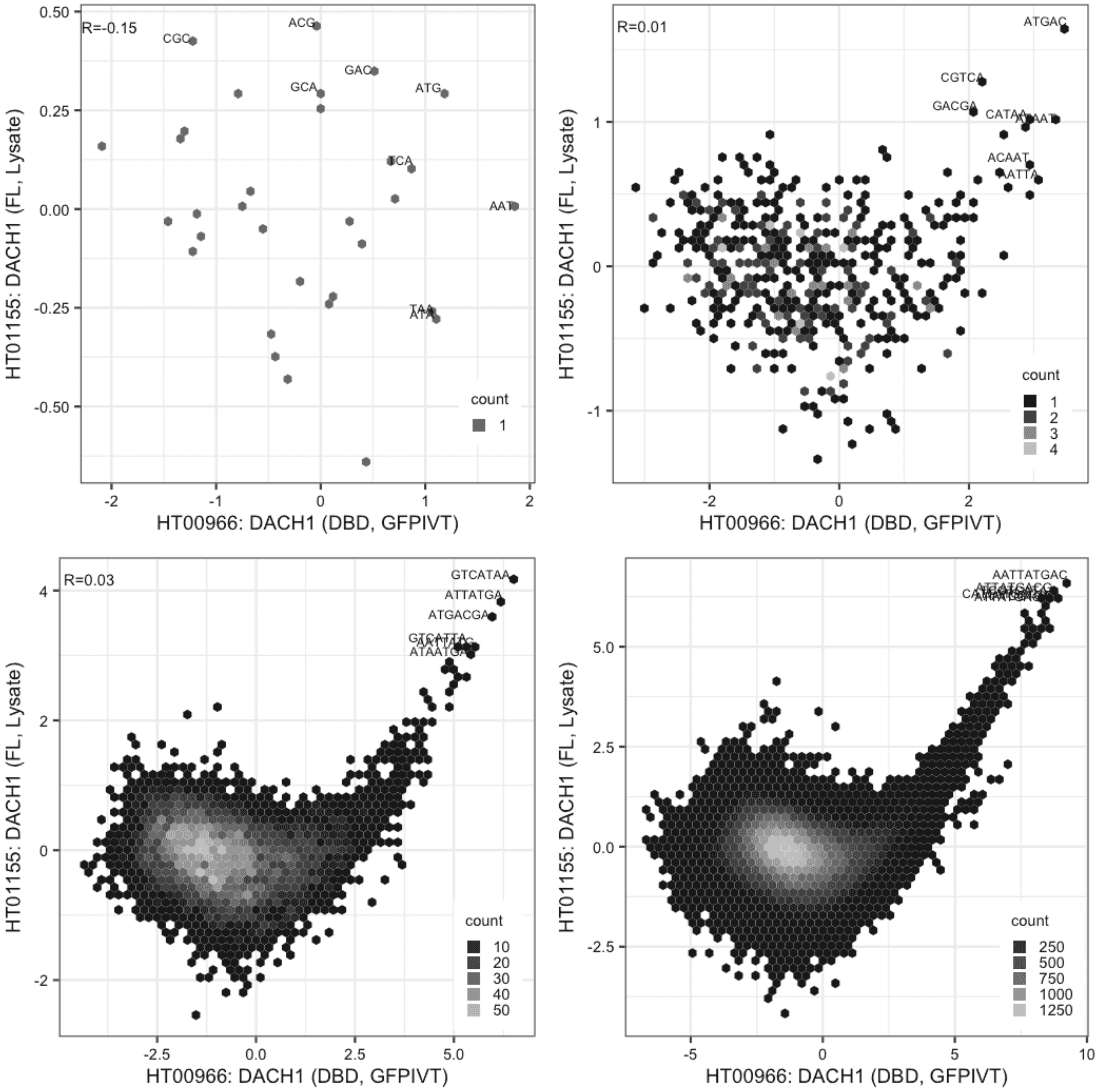

#### DACH1

Lysate vs Lysate, same construct

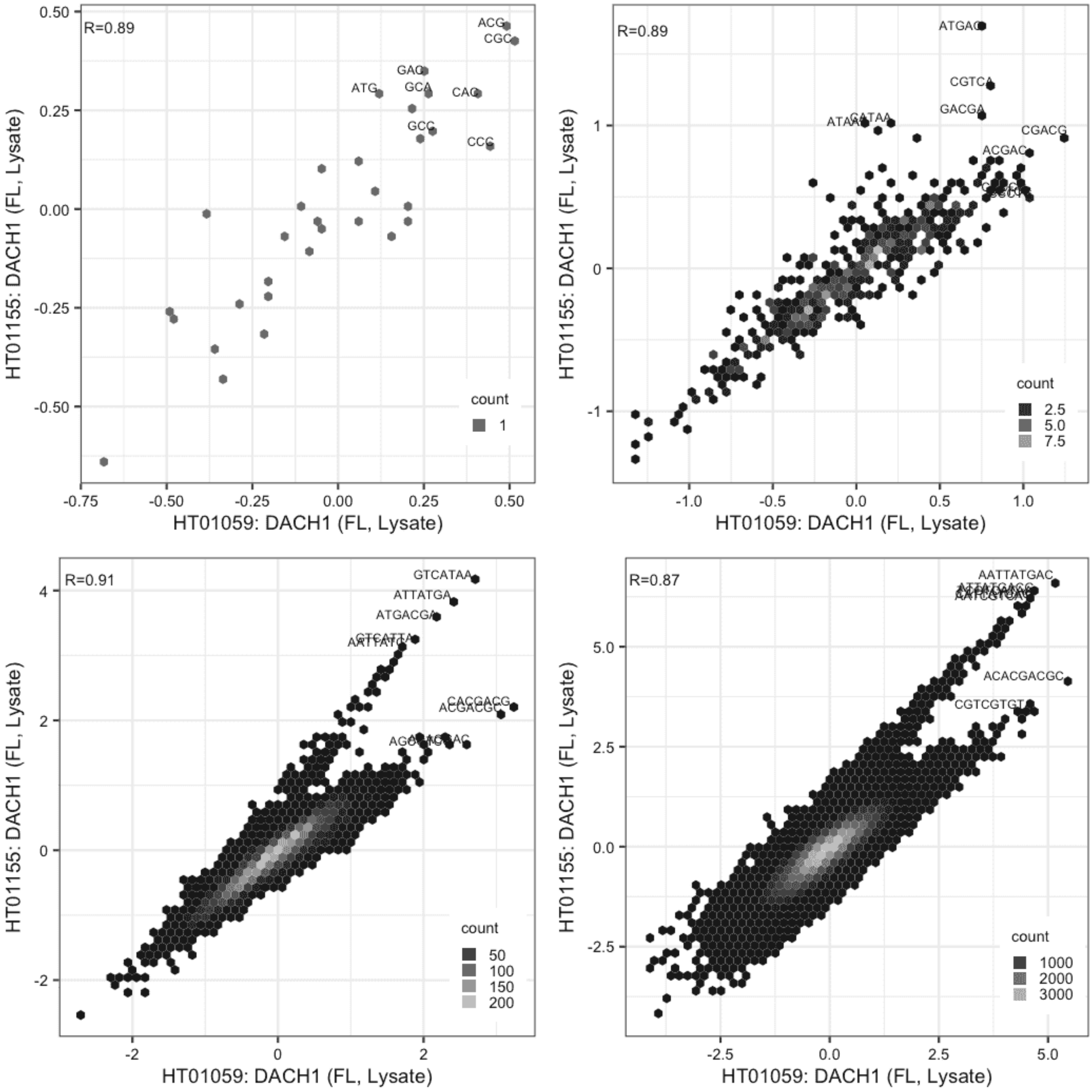

**DMTF1**

Lysate vs GFPIVT, same construct

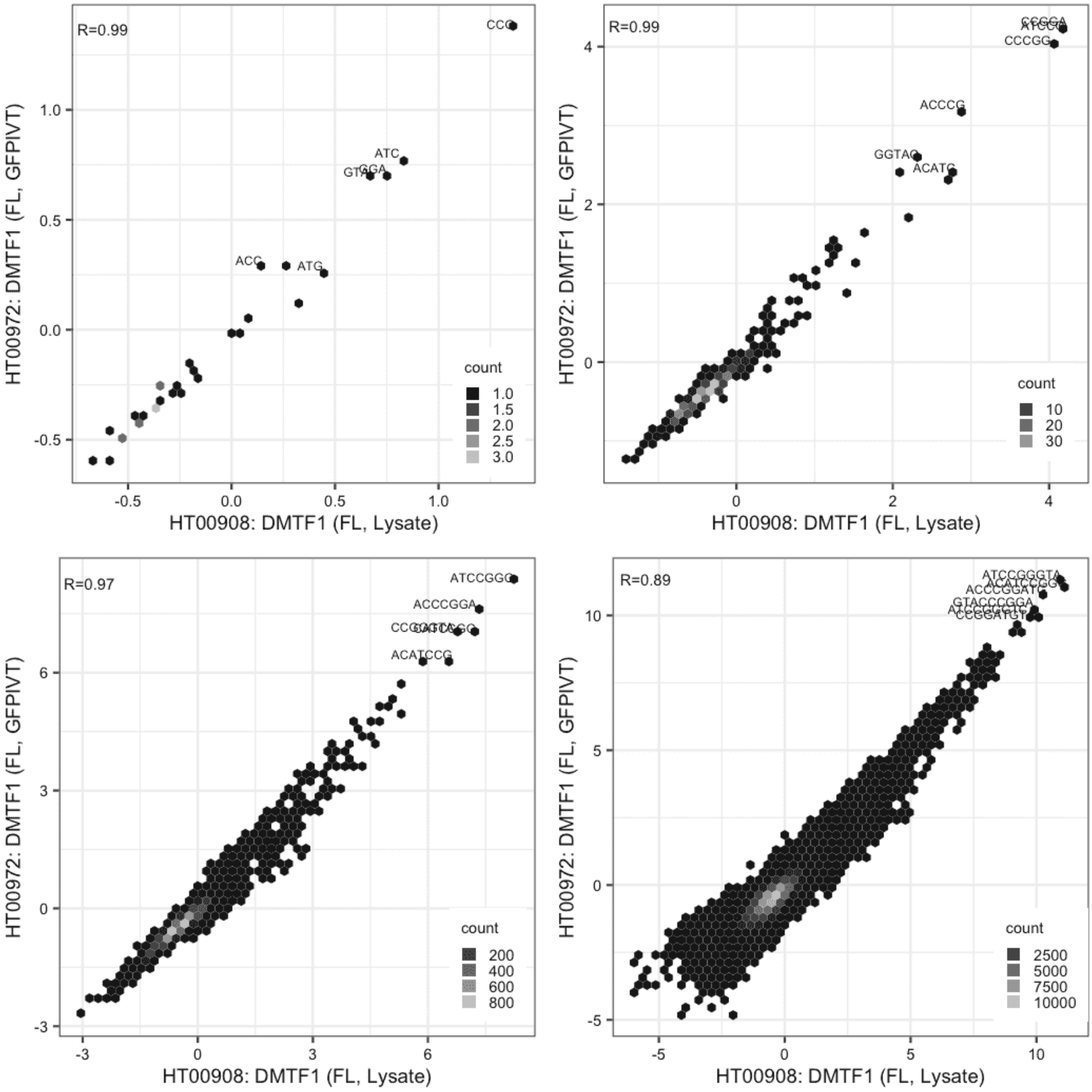

**DNTTIP1**

Lysate vs GFPIVT, same construct

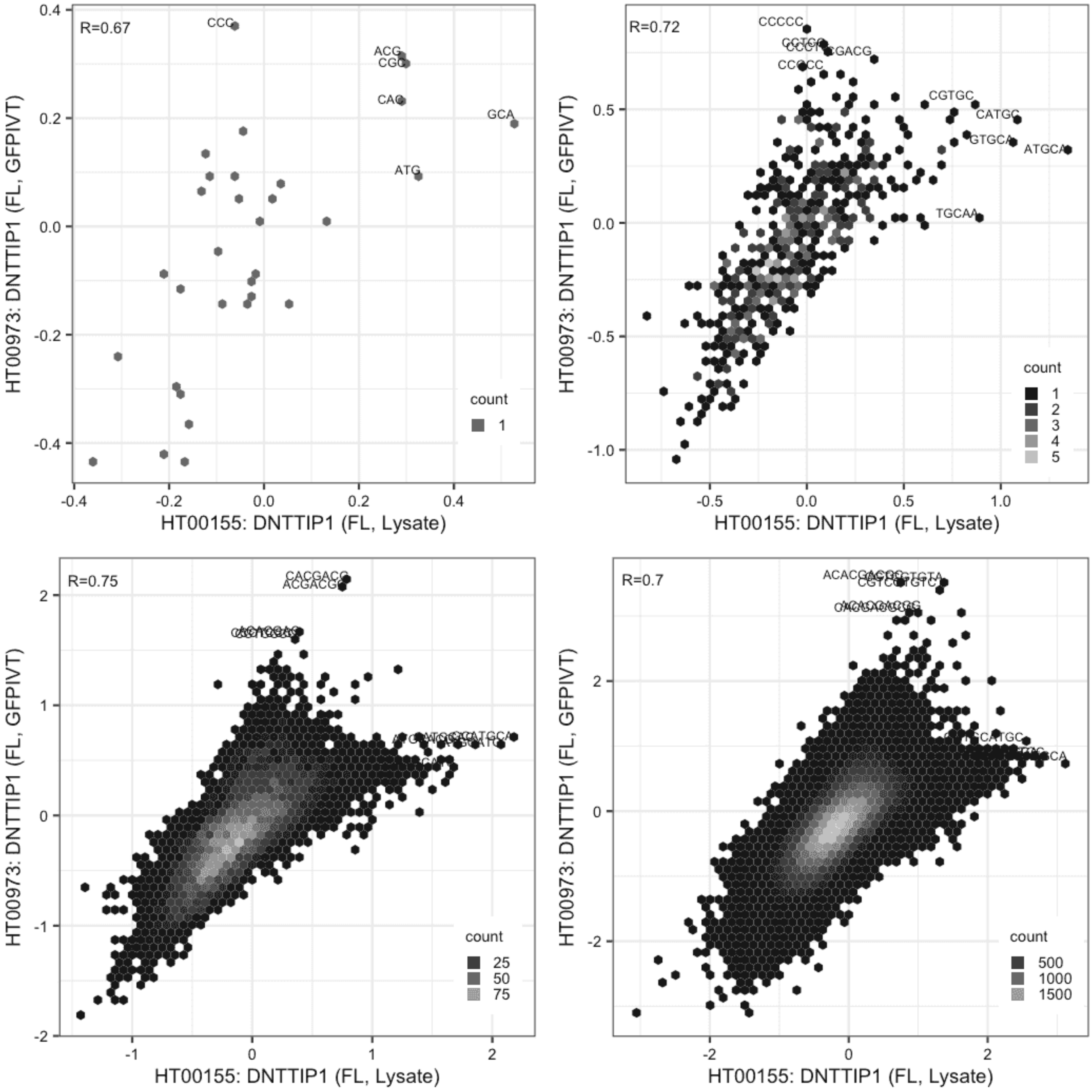

#### ELF3

Lysate vs GFPIVT, same construct

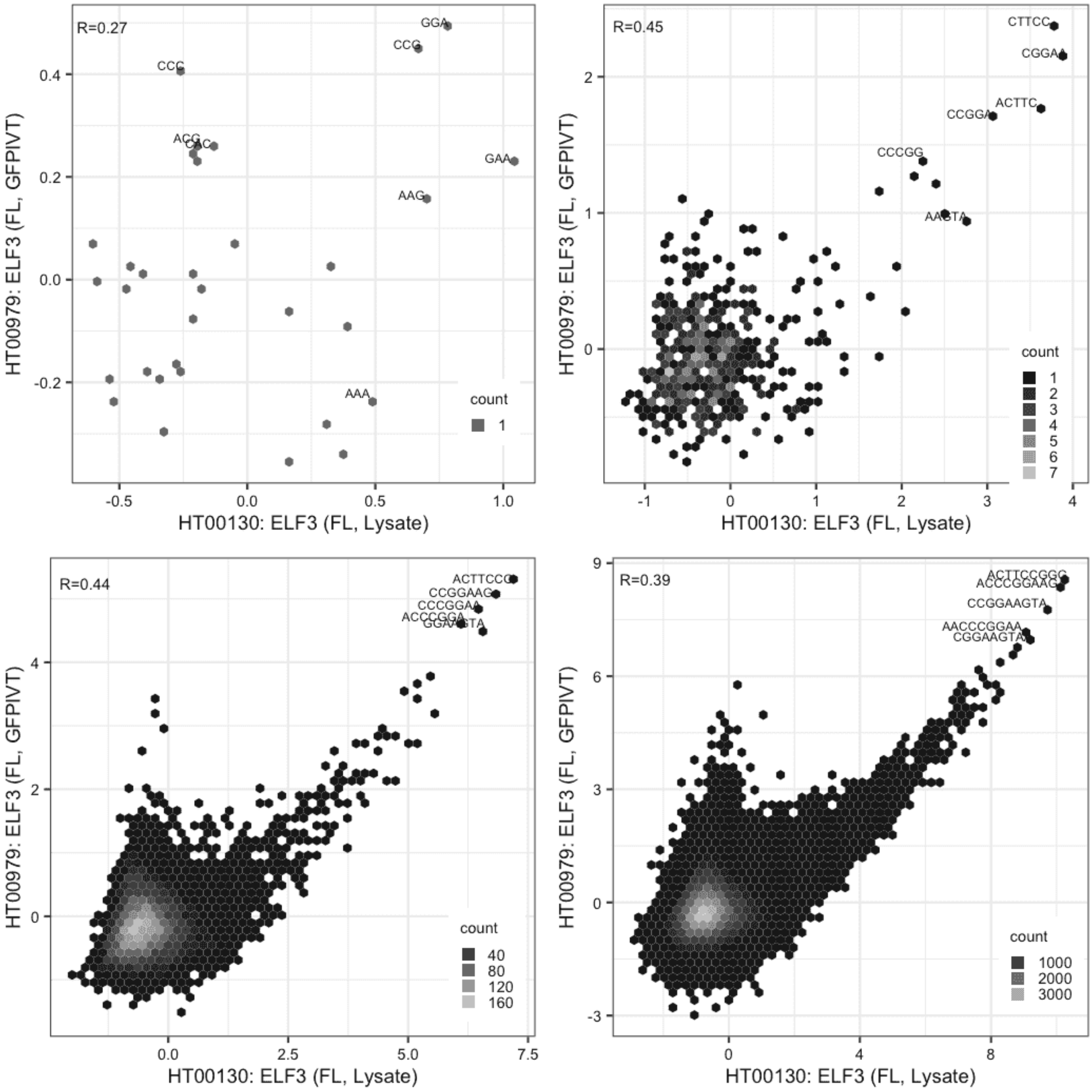

**FBXL19**

Lysate vs IVT, different constructs

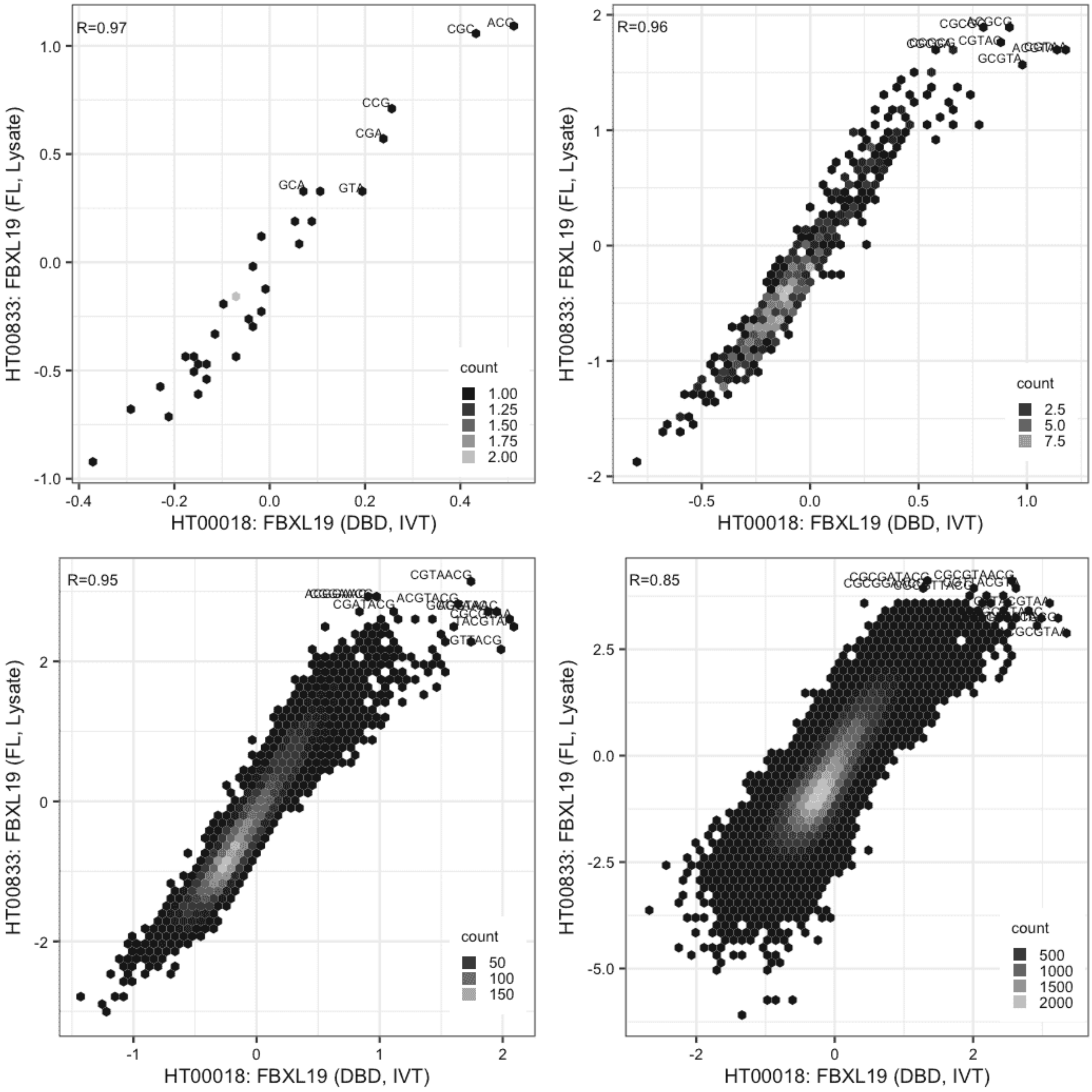

**FBXL19**

IVT vs GFPIVT, same construct

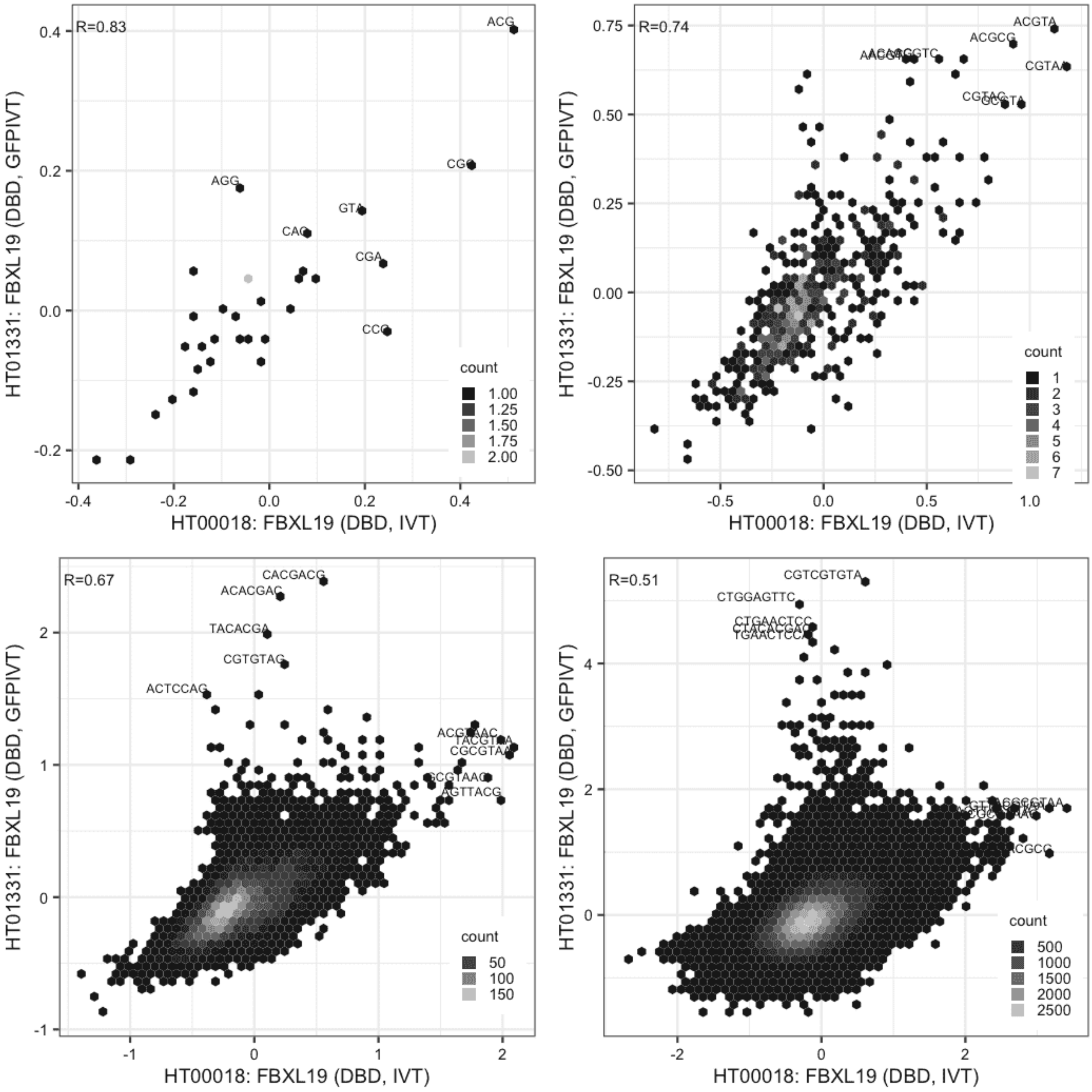

**FBXL19**

Lysate vs GFPIVT, different constructs

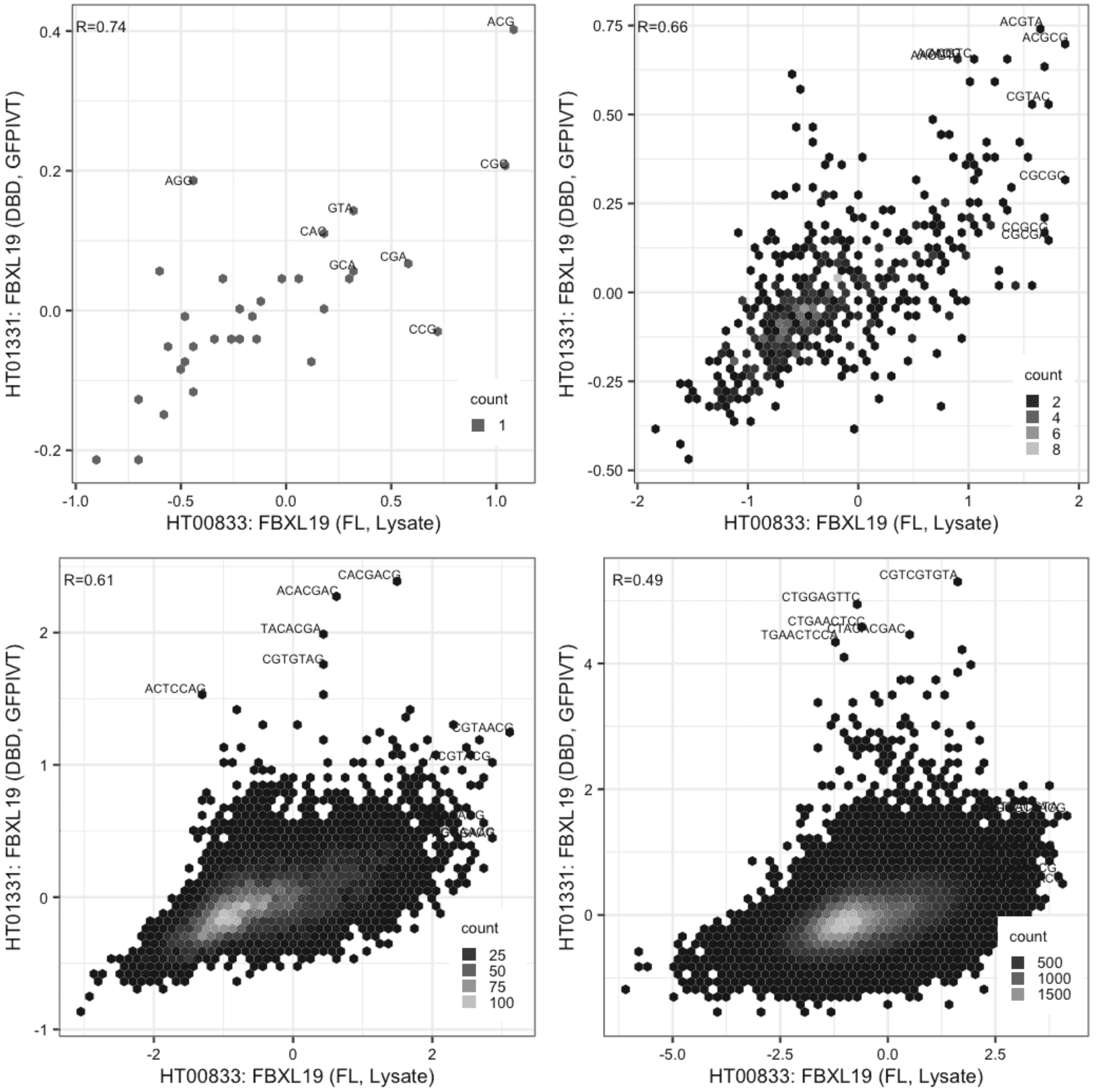

### FLI1

IVT vs IVT, same construct

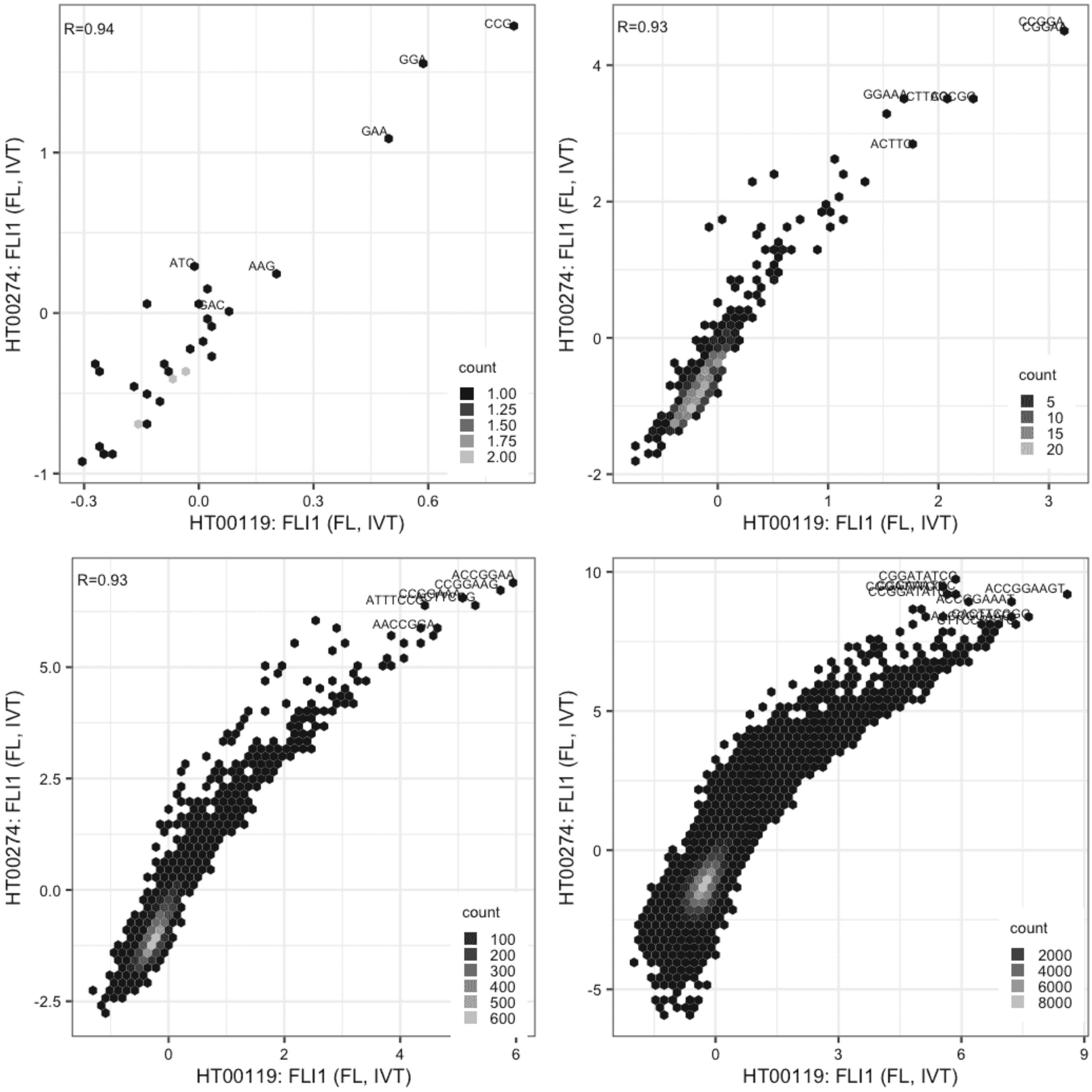

**FLI1**

Lysate vs IVT, same construct

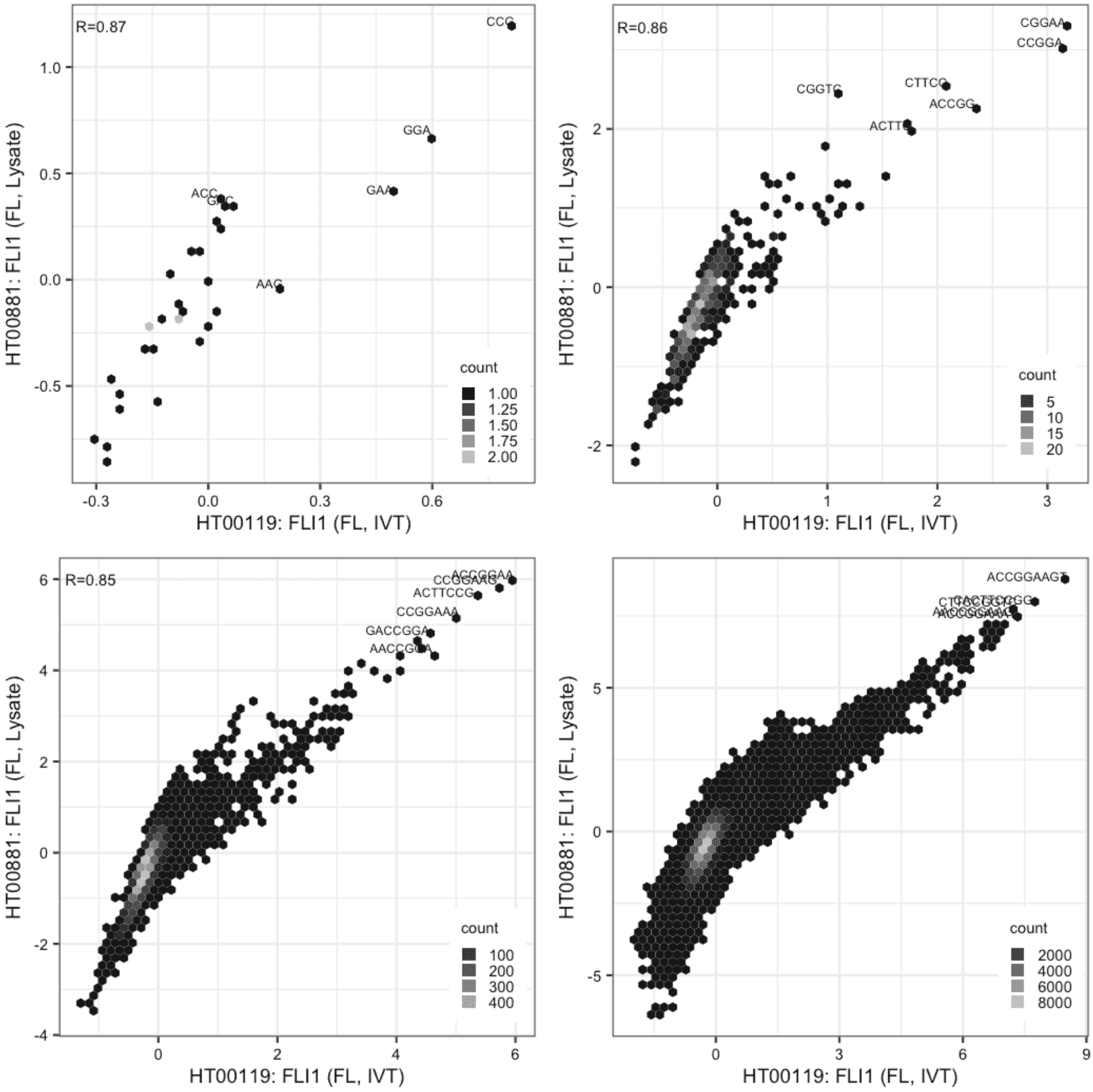

**FLI1**

Lysate vs IVT, same construct

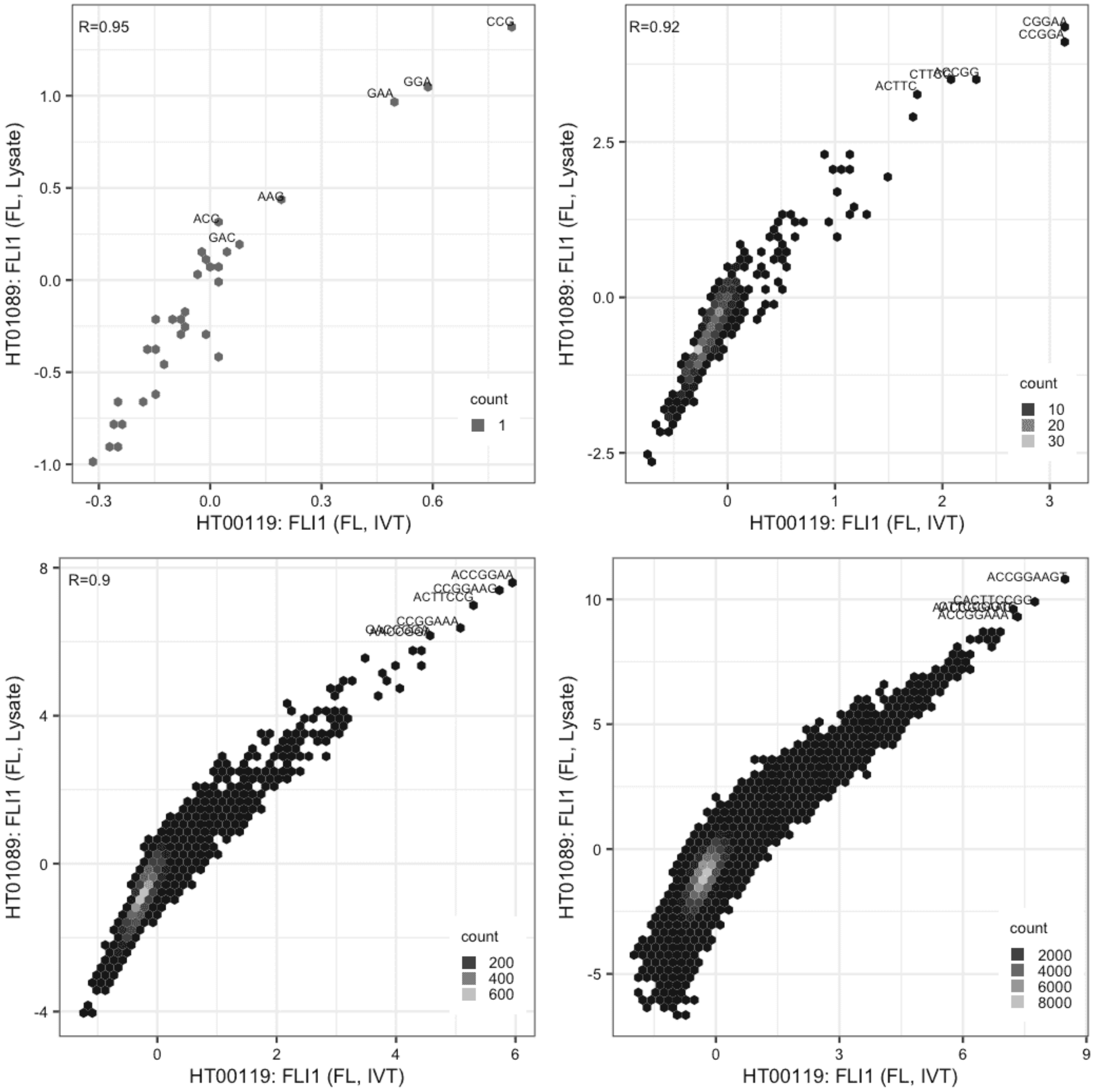

### FLI1

Lysate vs IVT, same construct

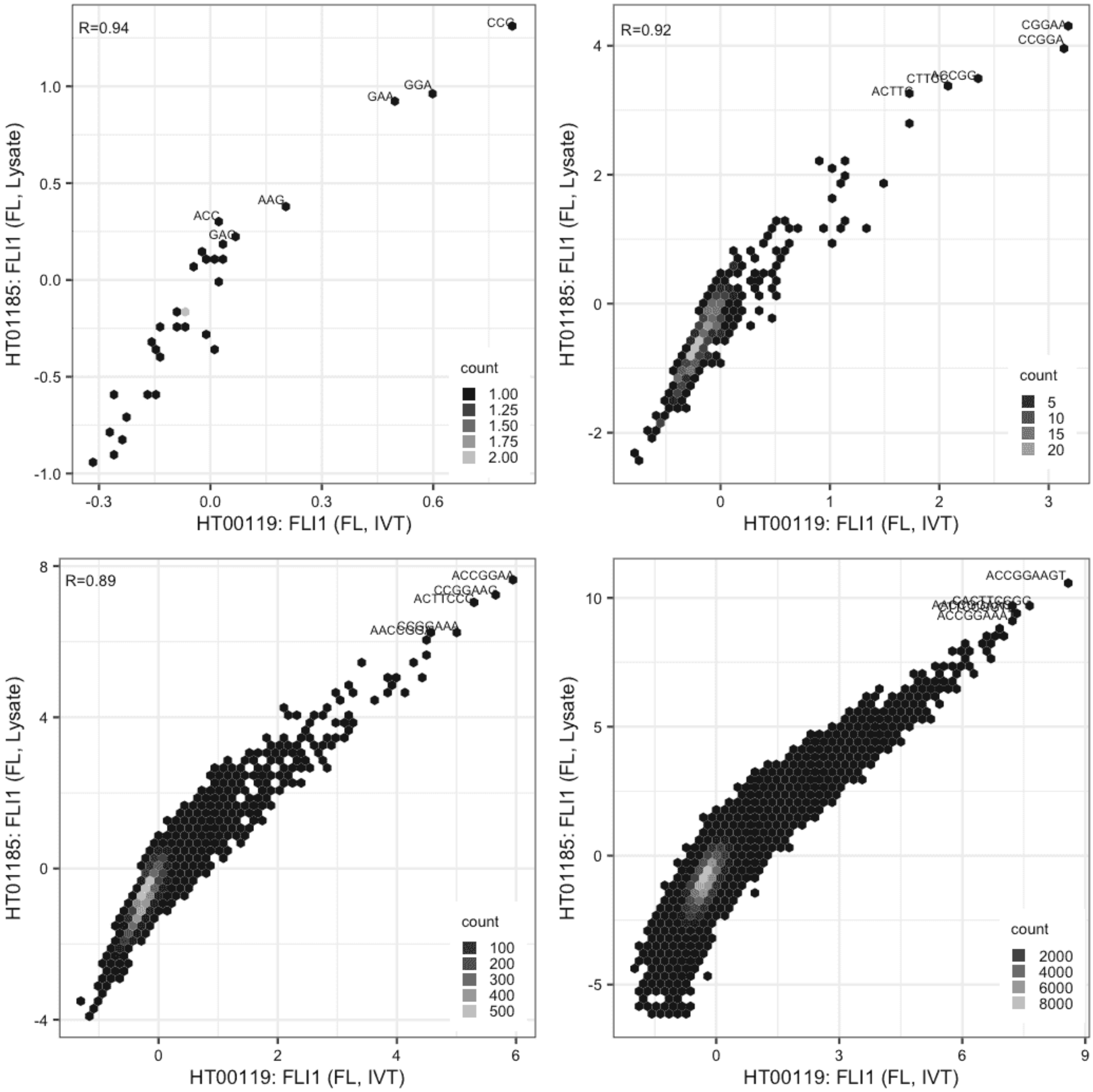

### FLI1

Lysate vs IVT, same construct

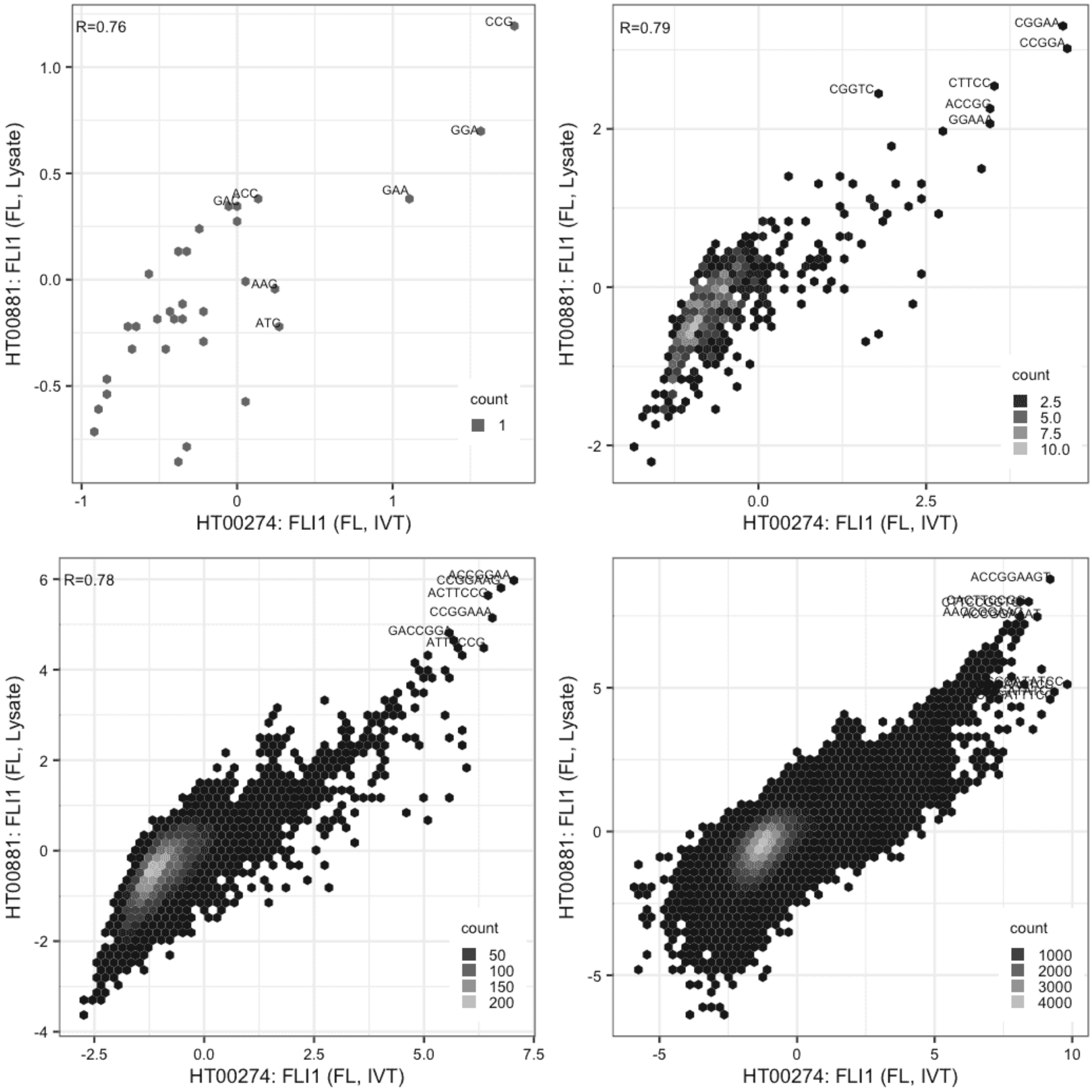

#### FLI1

Lysate vs IVT, same construct

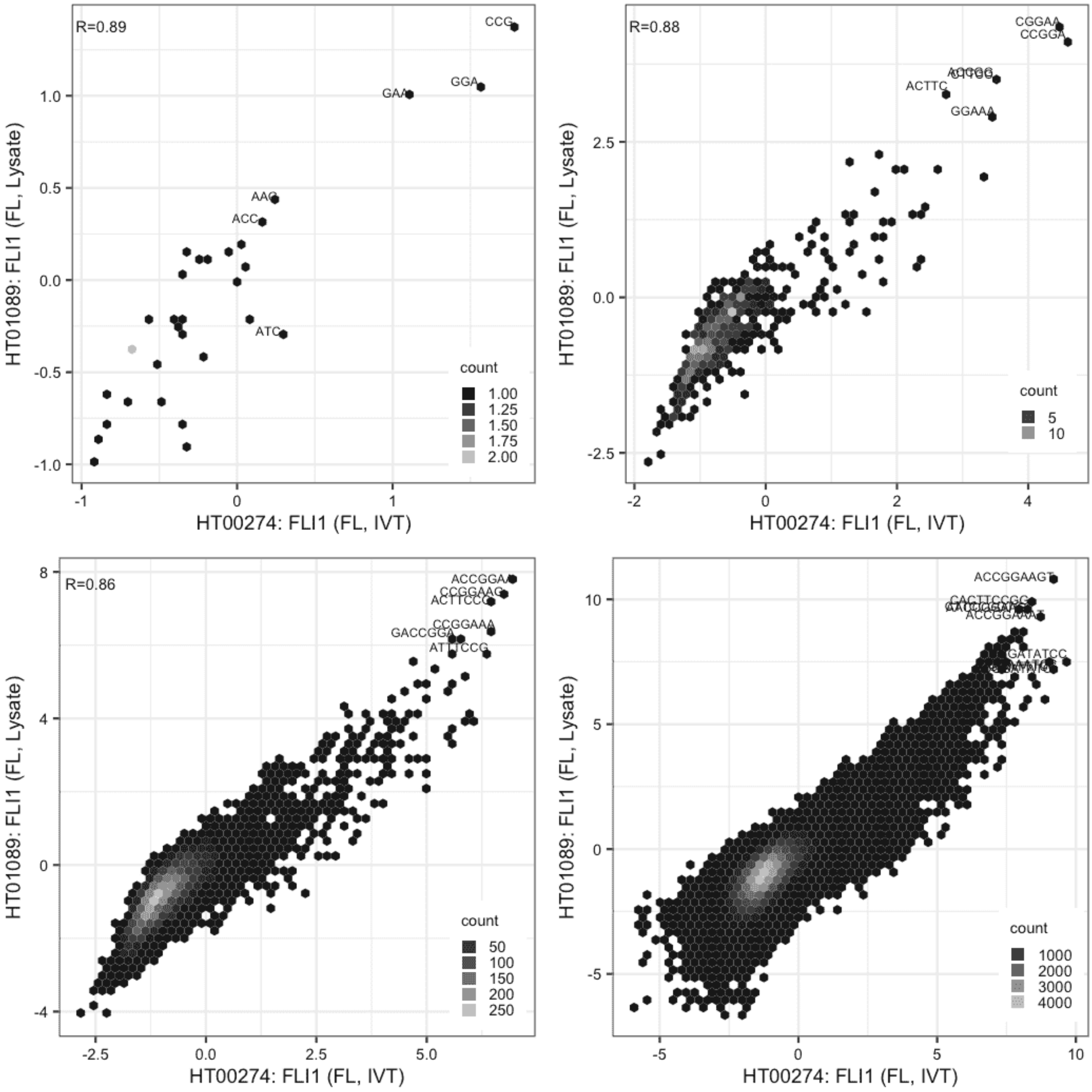

#### FLI1

Lysate vs IVT, same construct

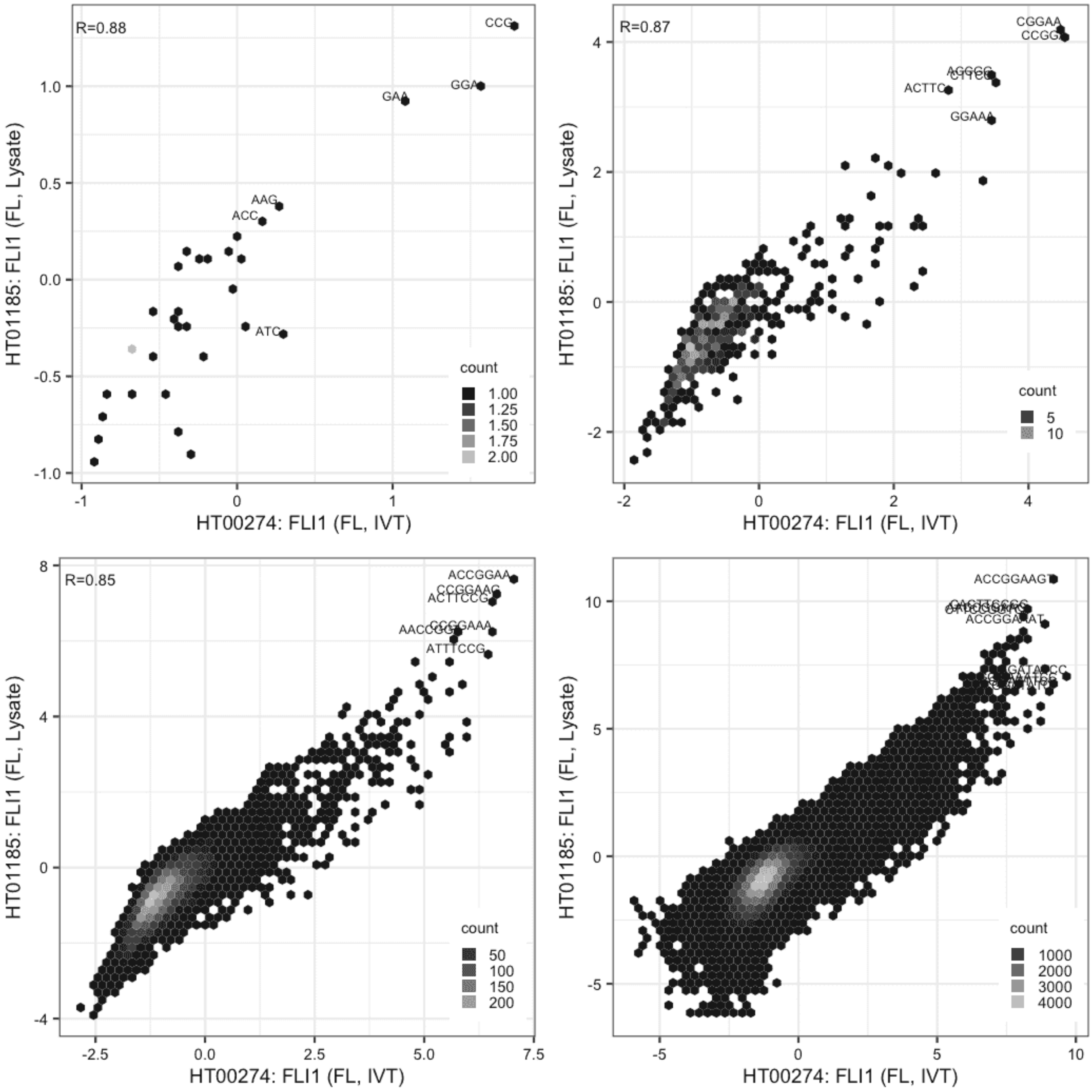

**FLI1**

Lysate vs Lysate, same construct

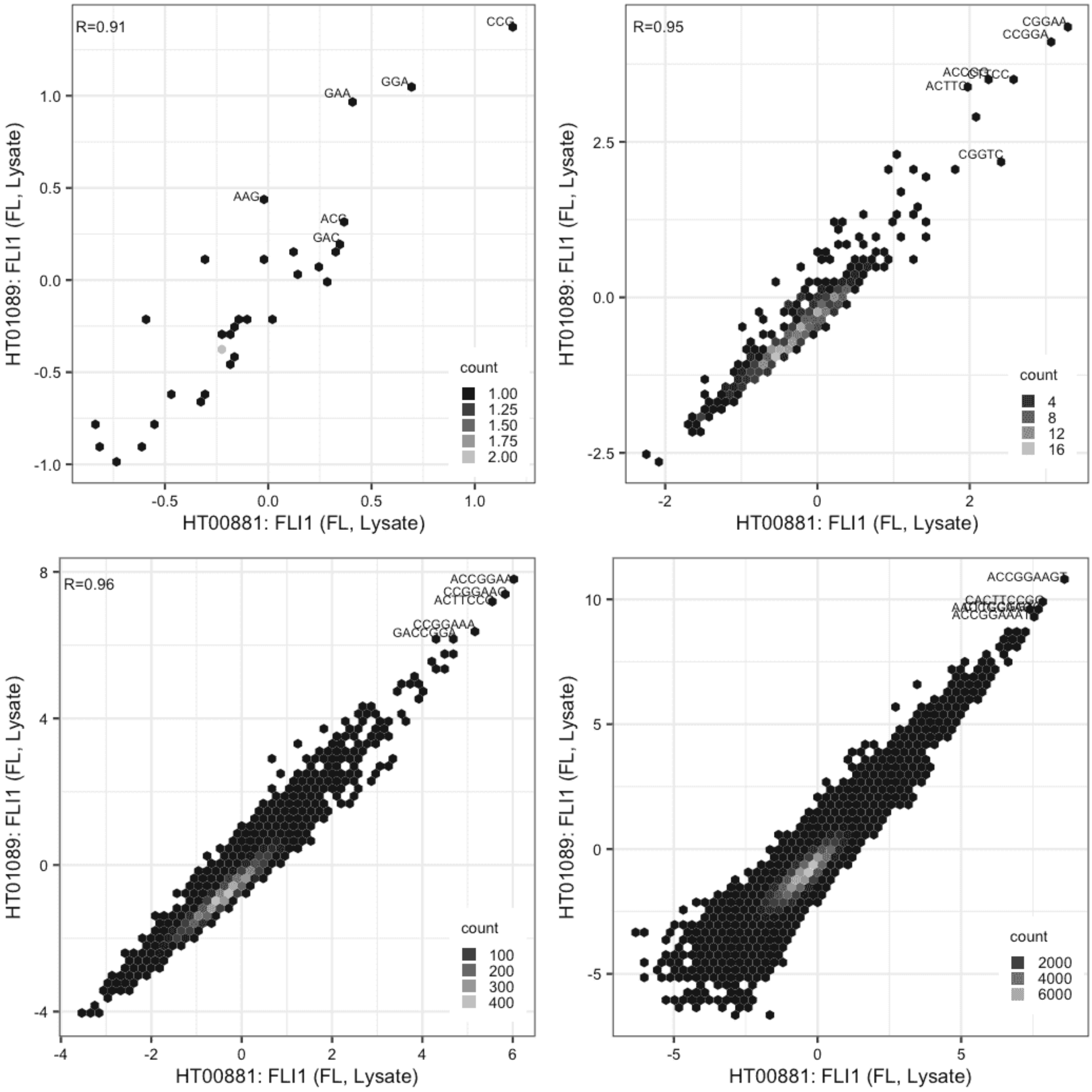

**FLI1**

Lysate vs Lysate, same construct

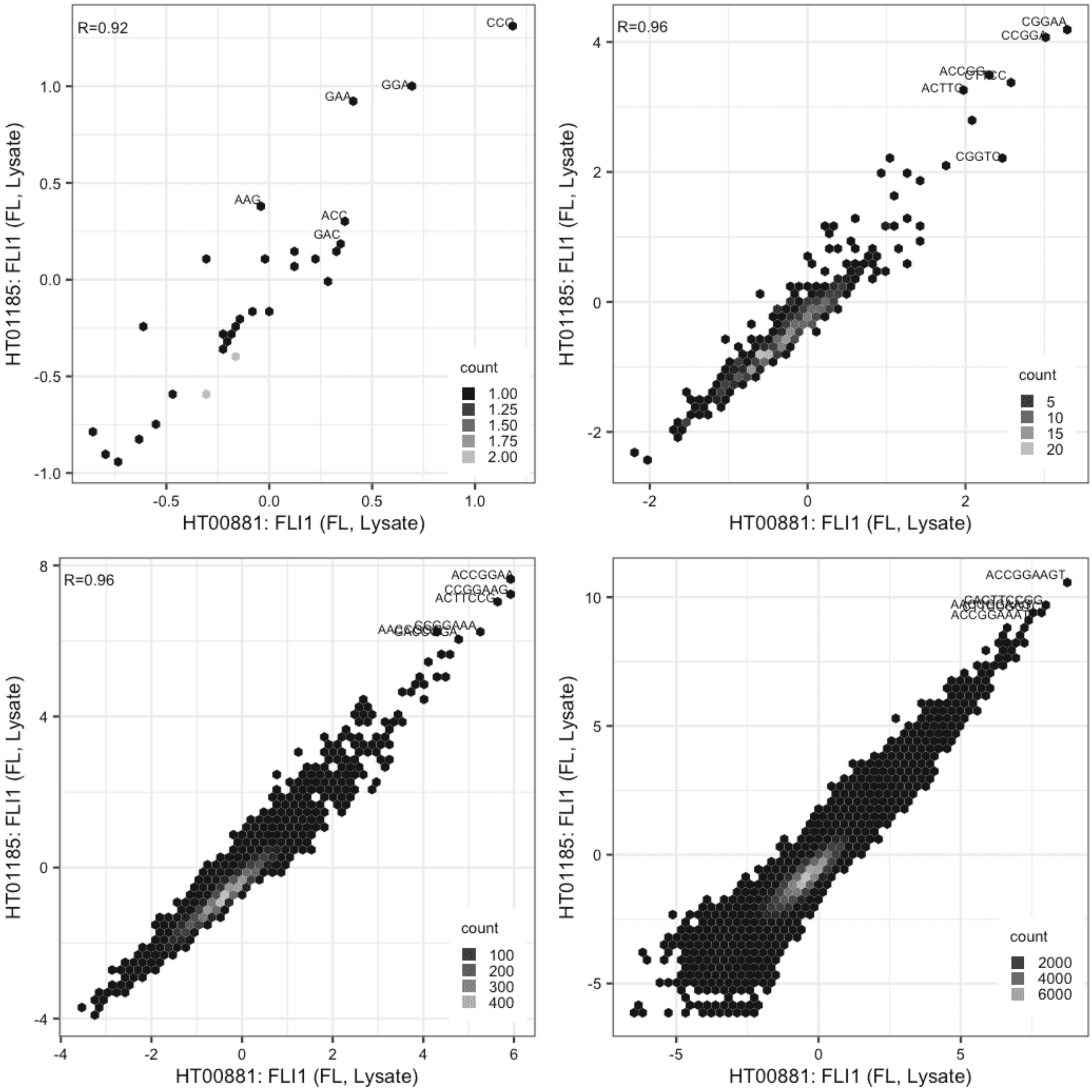

#### FLI1

Lysate vs Lysate, same construct

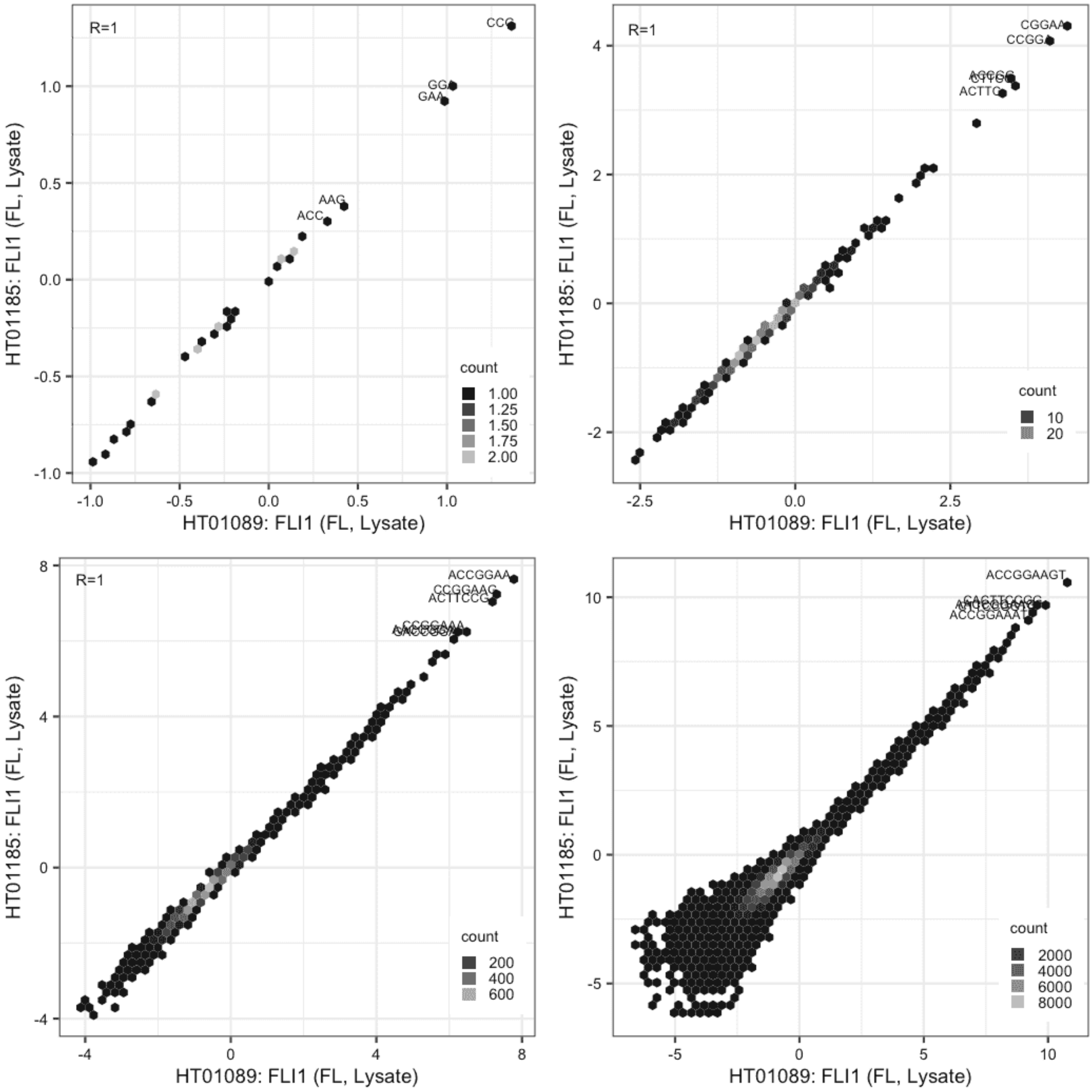

**FOSL2**

Lysate vs IVT, same construct

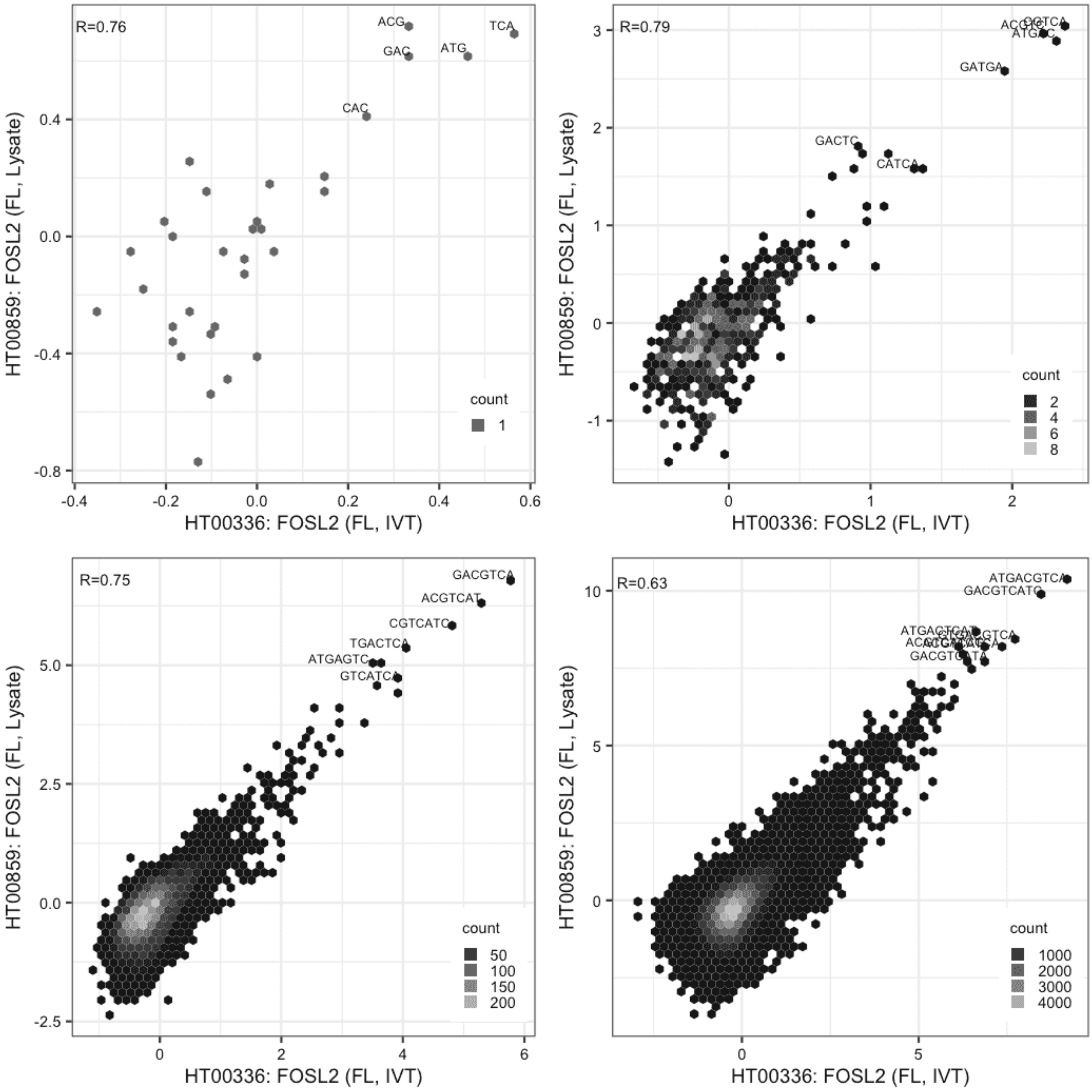

#### GABPA

Lysate vs IVT, same construct

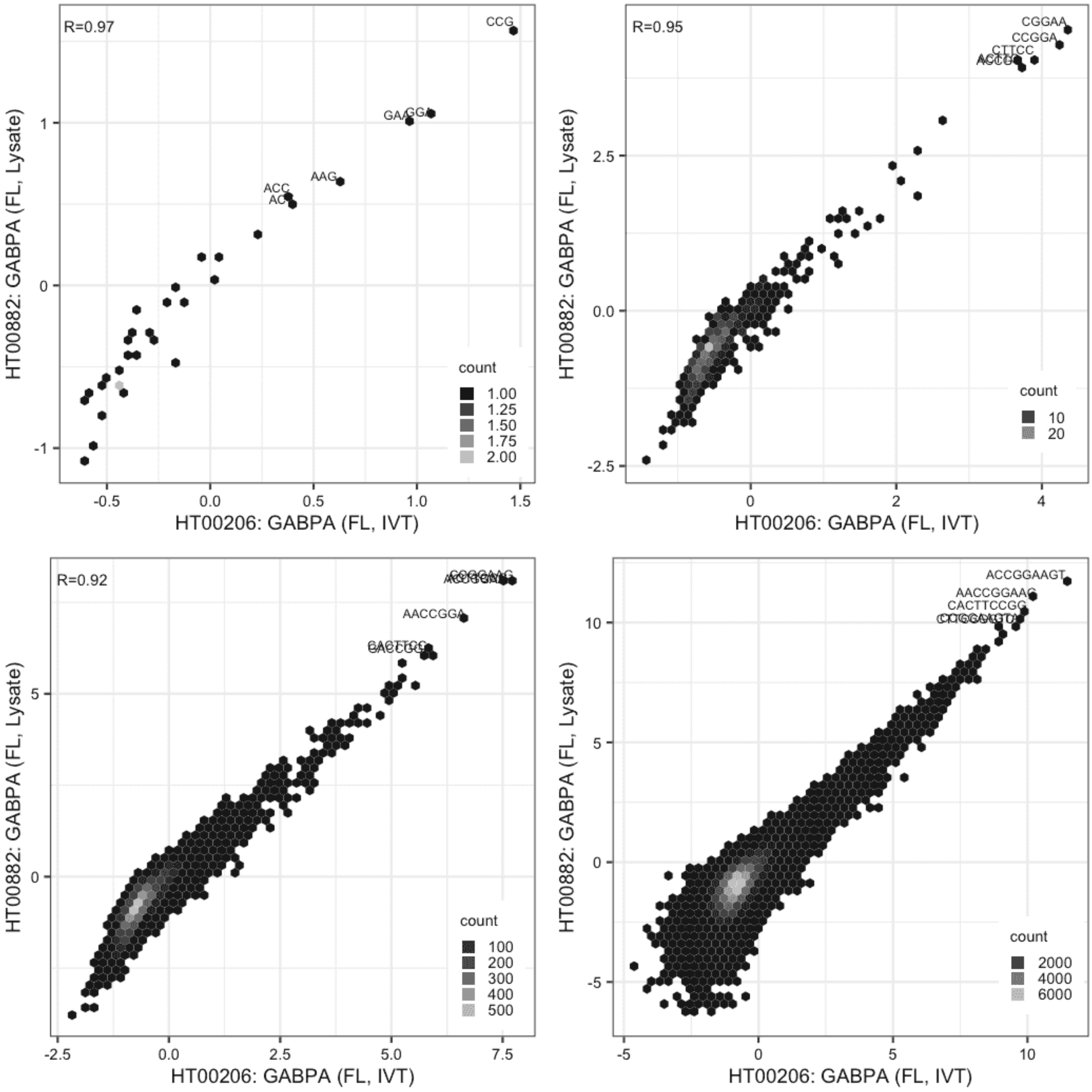

**GCM1**

Lysate vs IVT, same construct

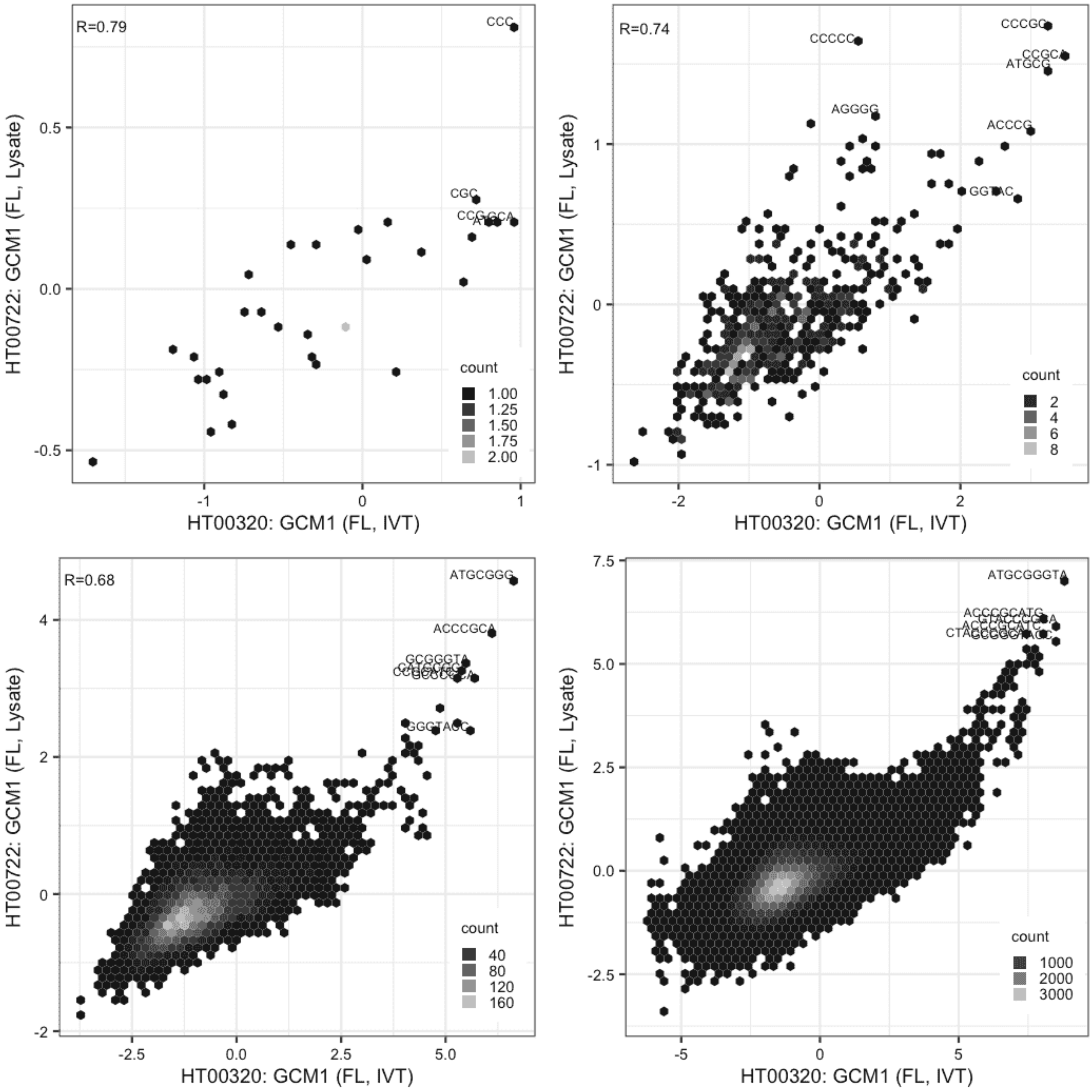

**GCM1**

Lysate vs IVT, same construct

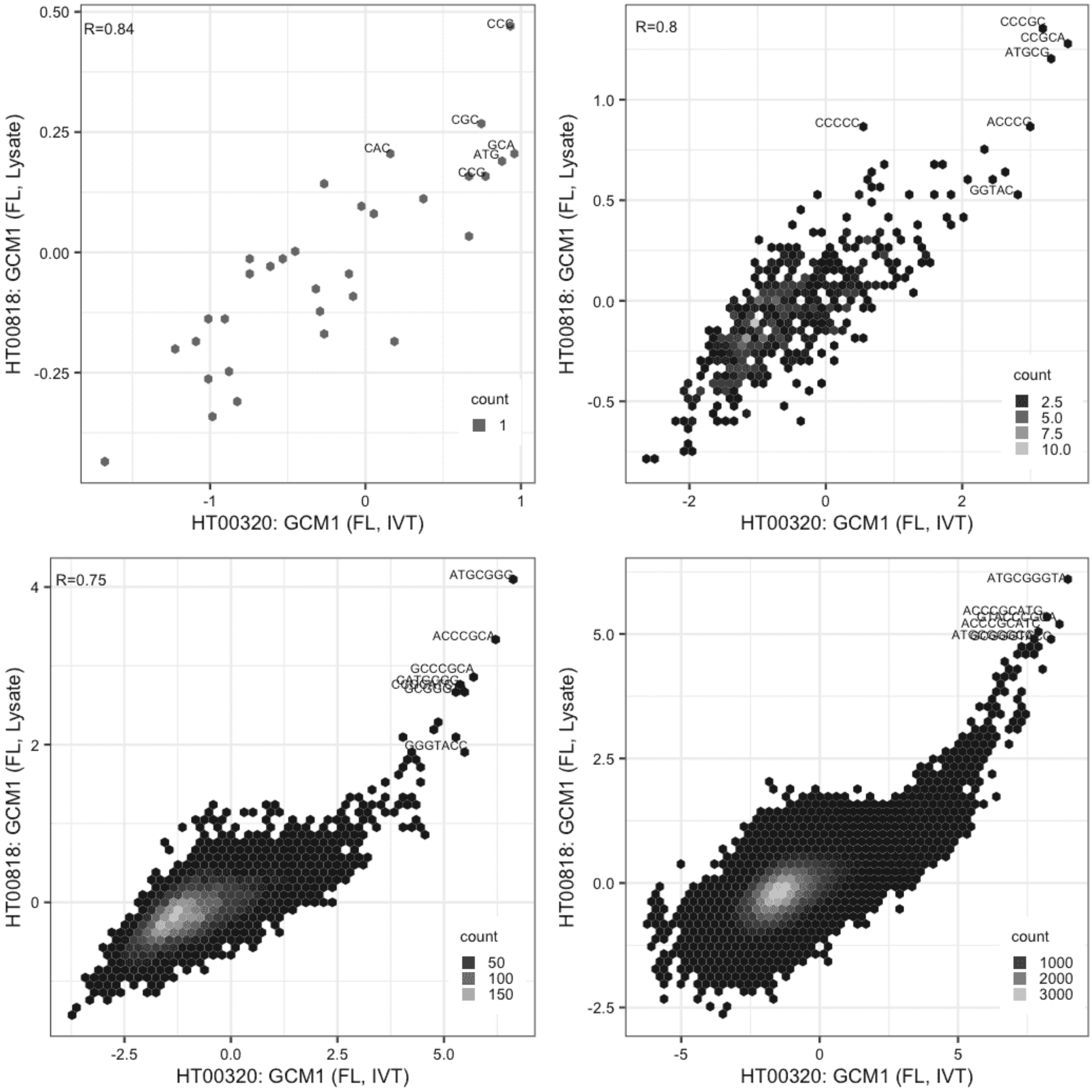

**GCM1**

Lysate vs Lysate, same construct

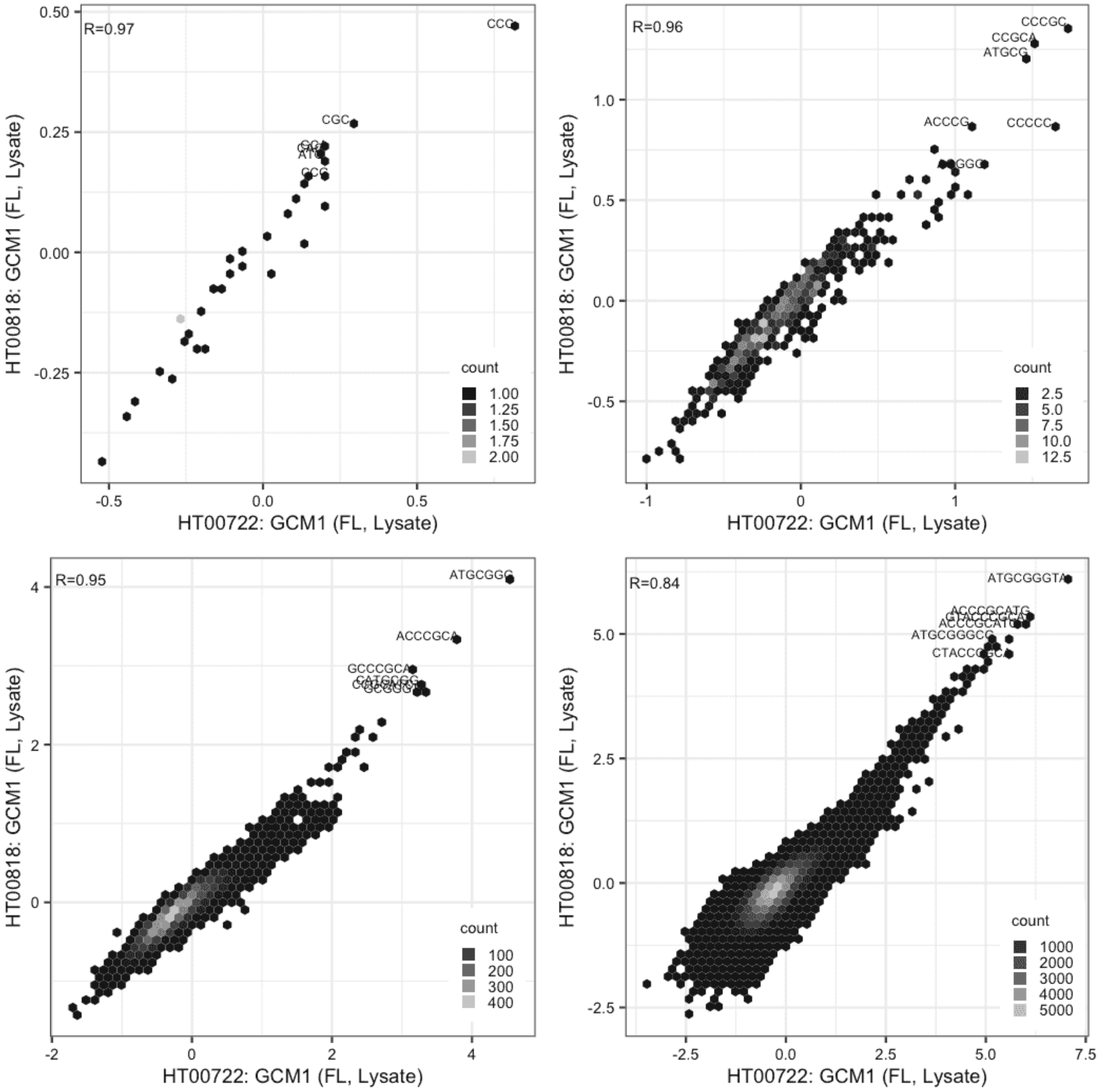

#### JRK

Lysate vs IVT, different constructs

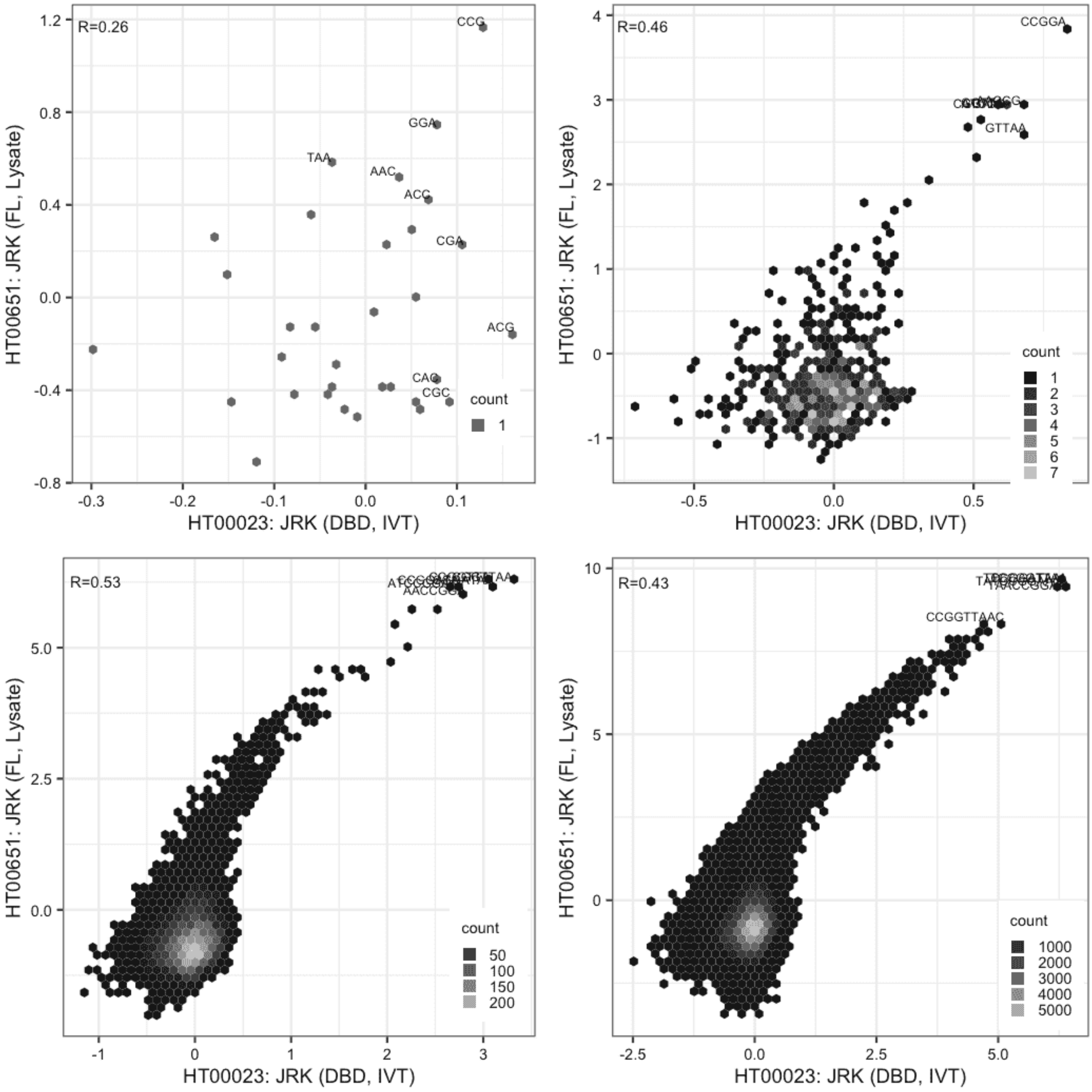

#### JRK

Lysate vs IVT, different constructs

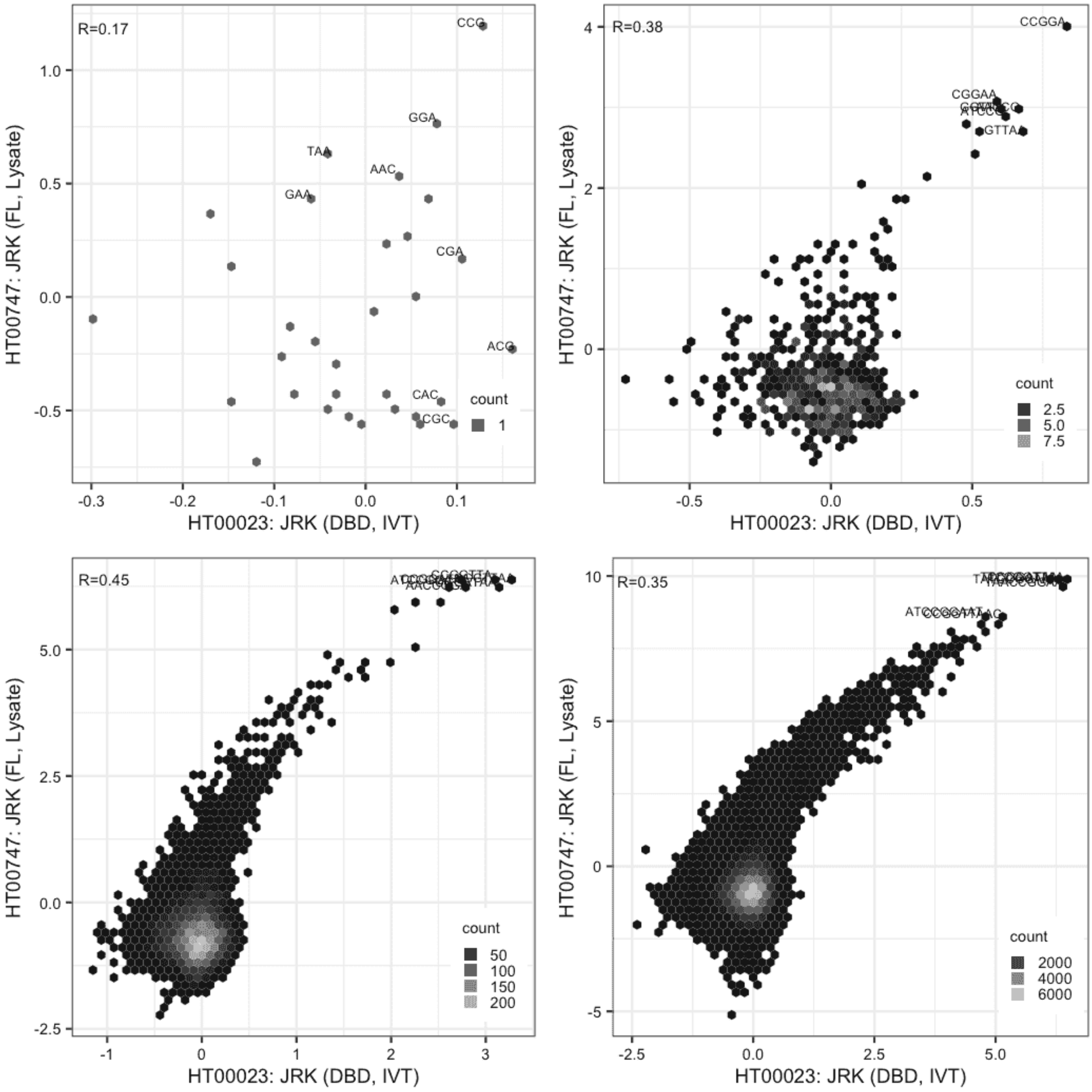

#### JRK

IVT vs GFPIVT, same construct

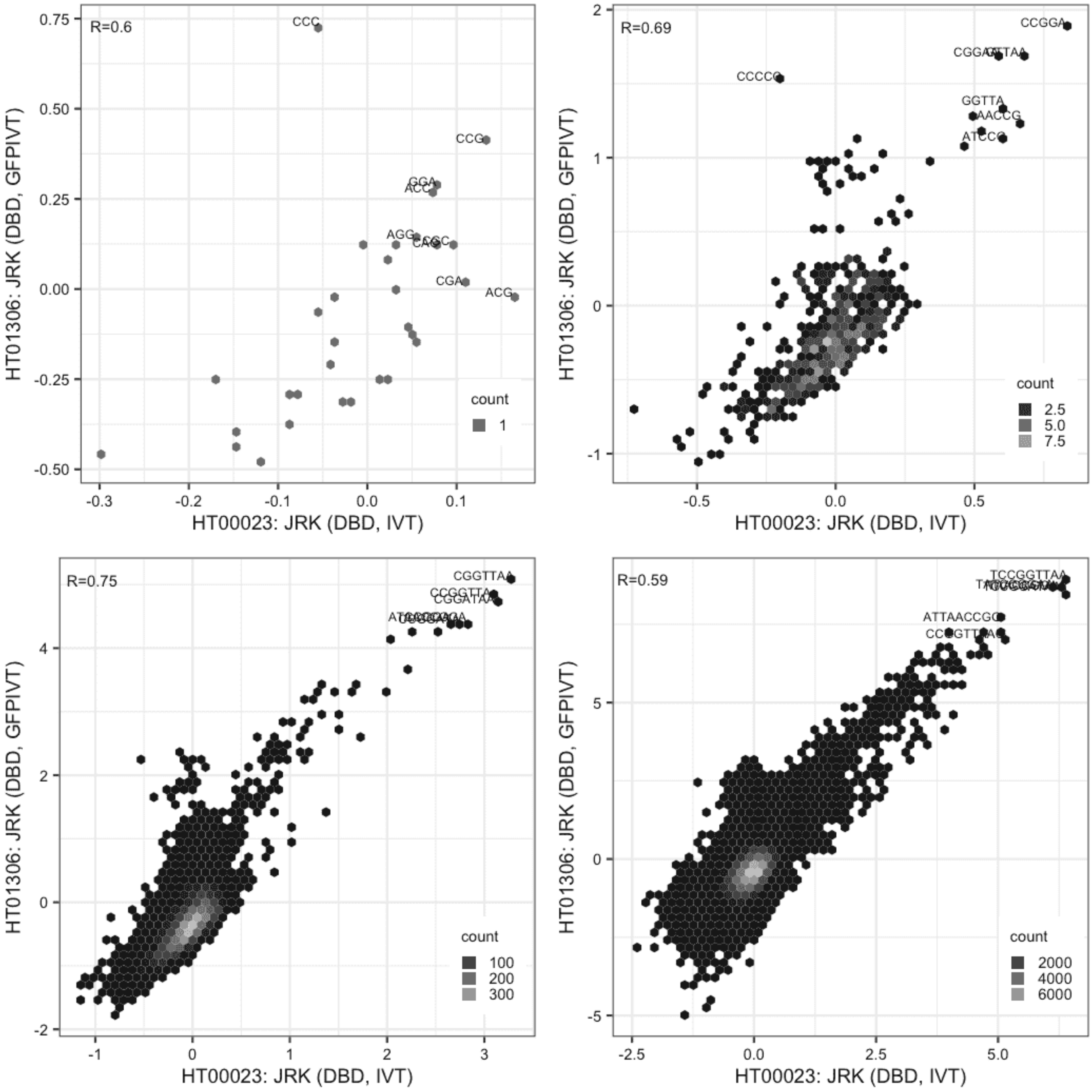

#### JRK

IVT vs GFPIVT, different constructs

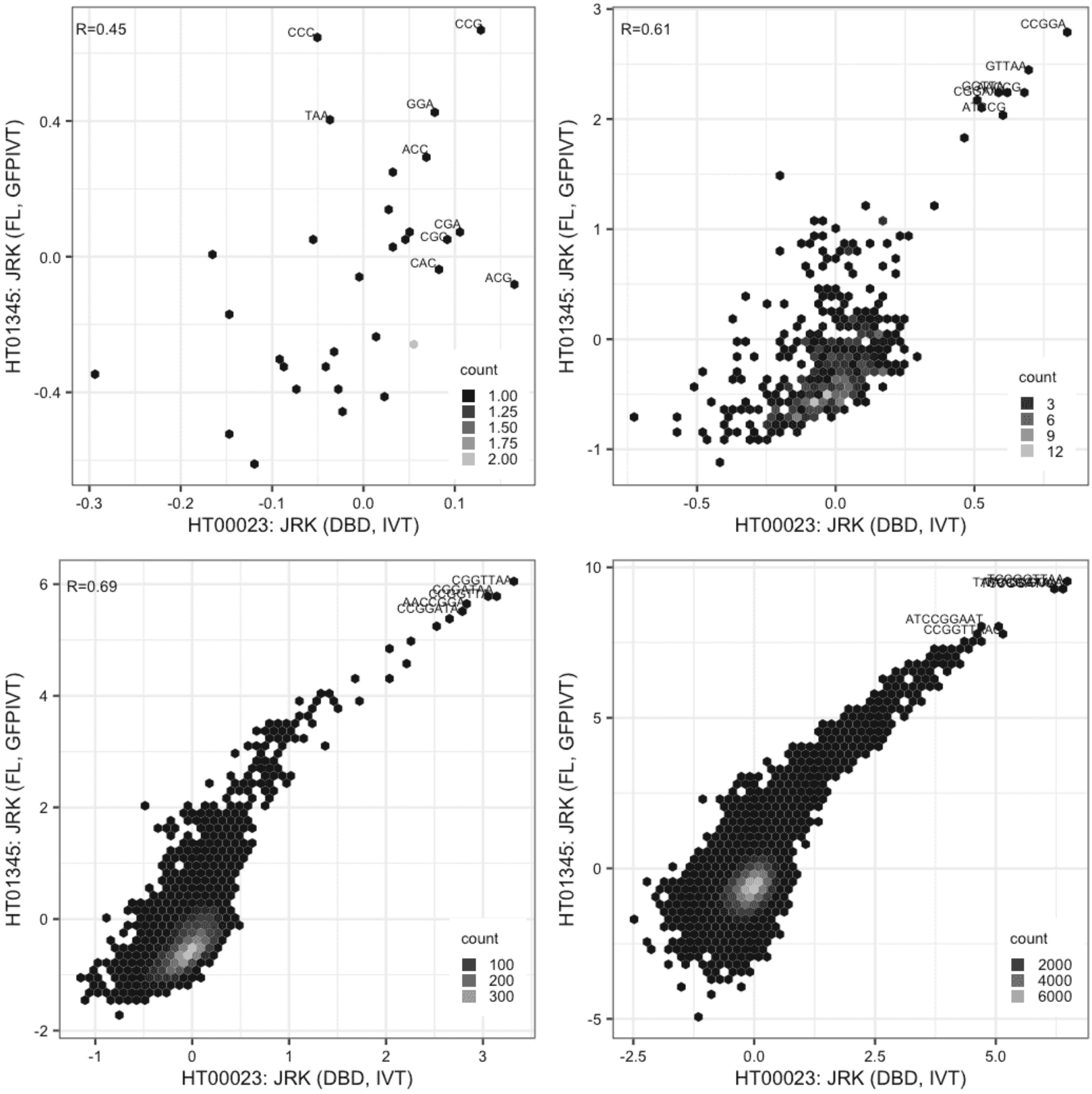

#### JRK

Lysate vs Lysate, same construct

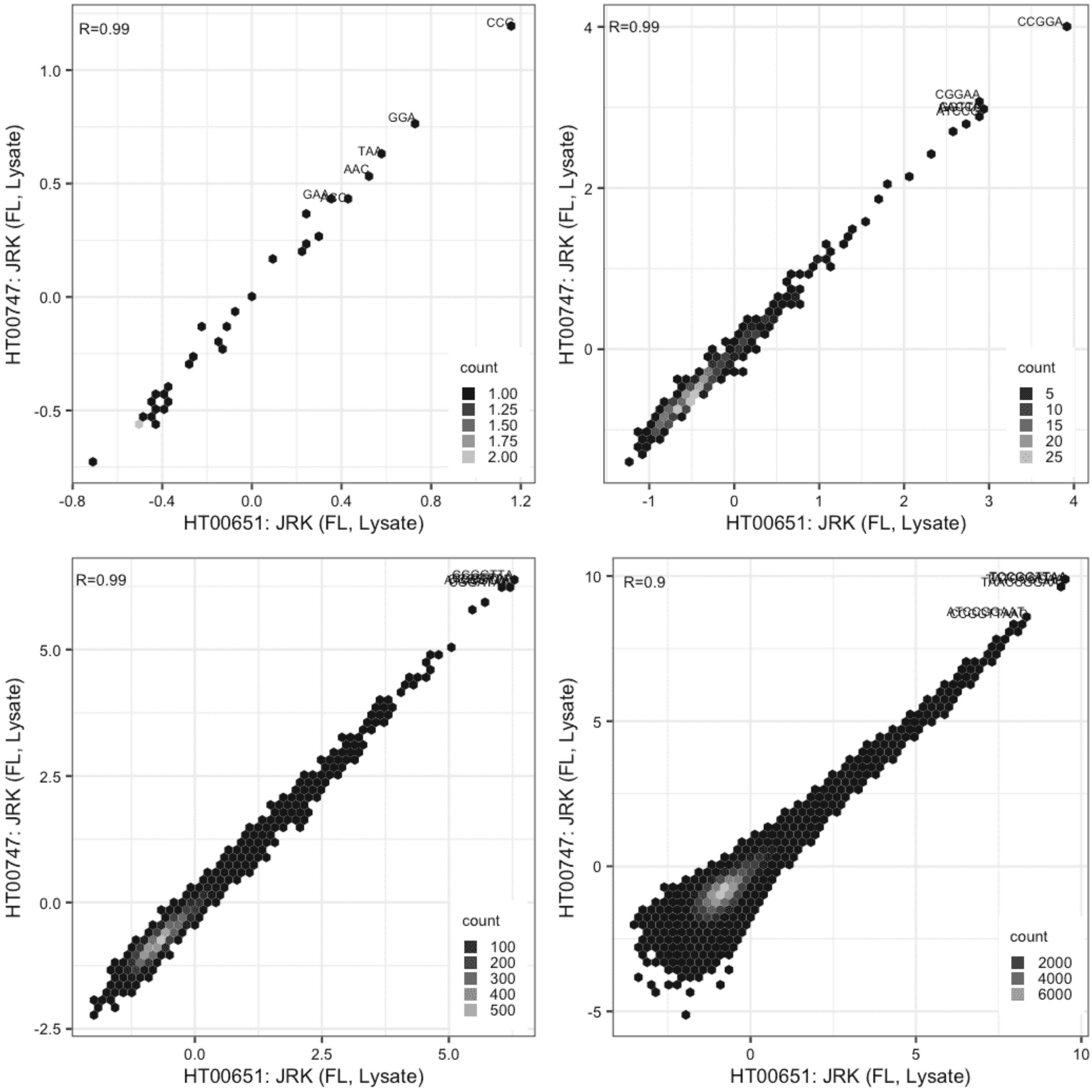

#### JRK

Lysate vs GFPIVT, different constructs

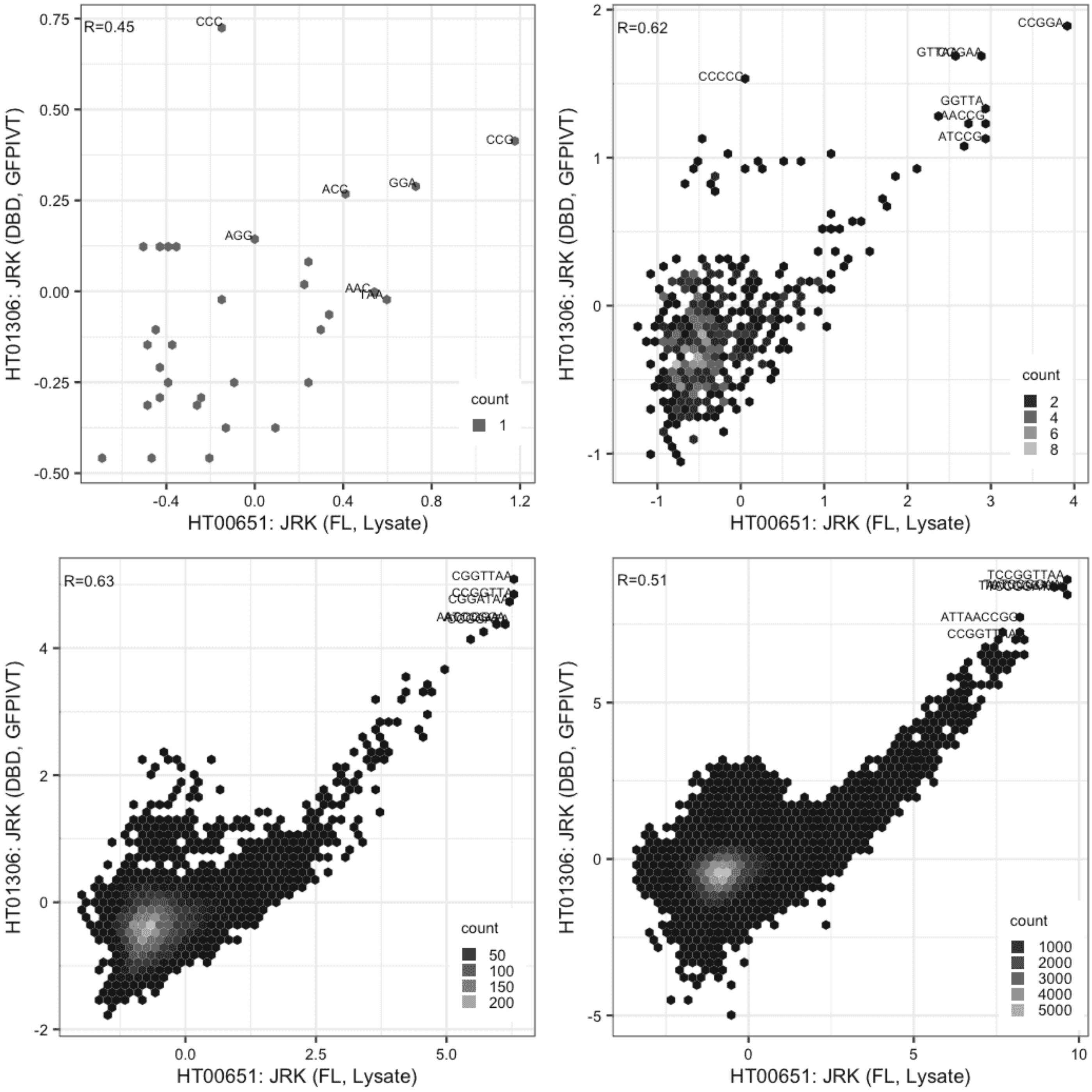

**JRK**

Lysate vs GFPIVT, same construct

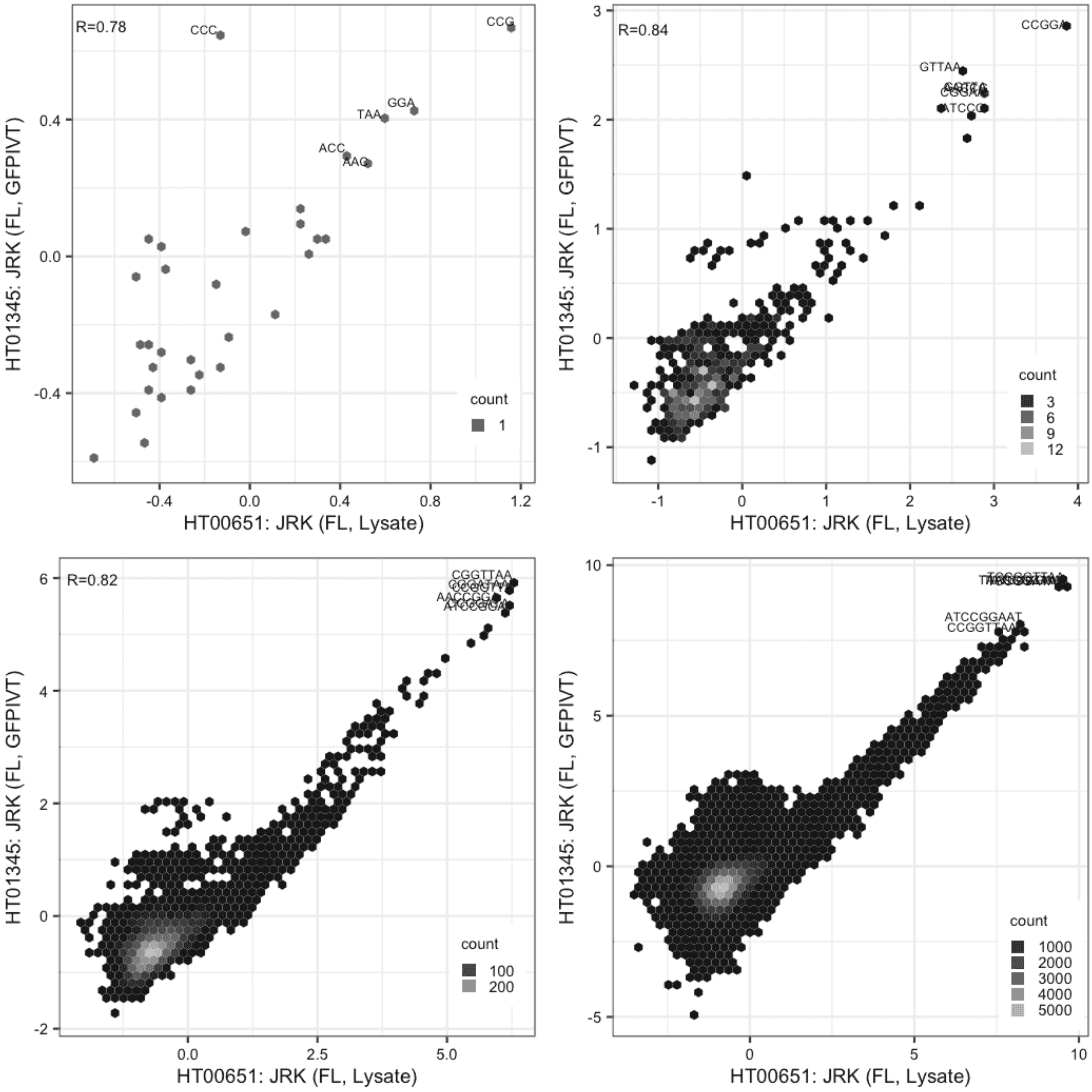

### JRK

Lysate vs GFPIVT, different constructs

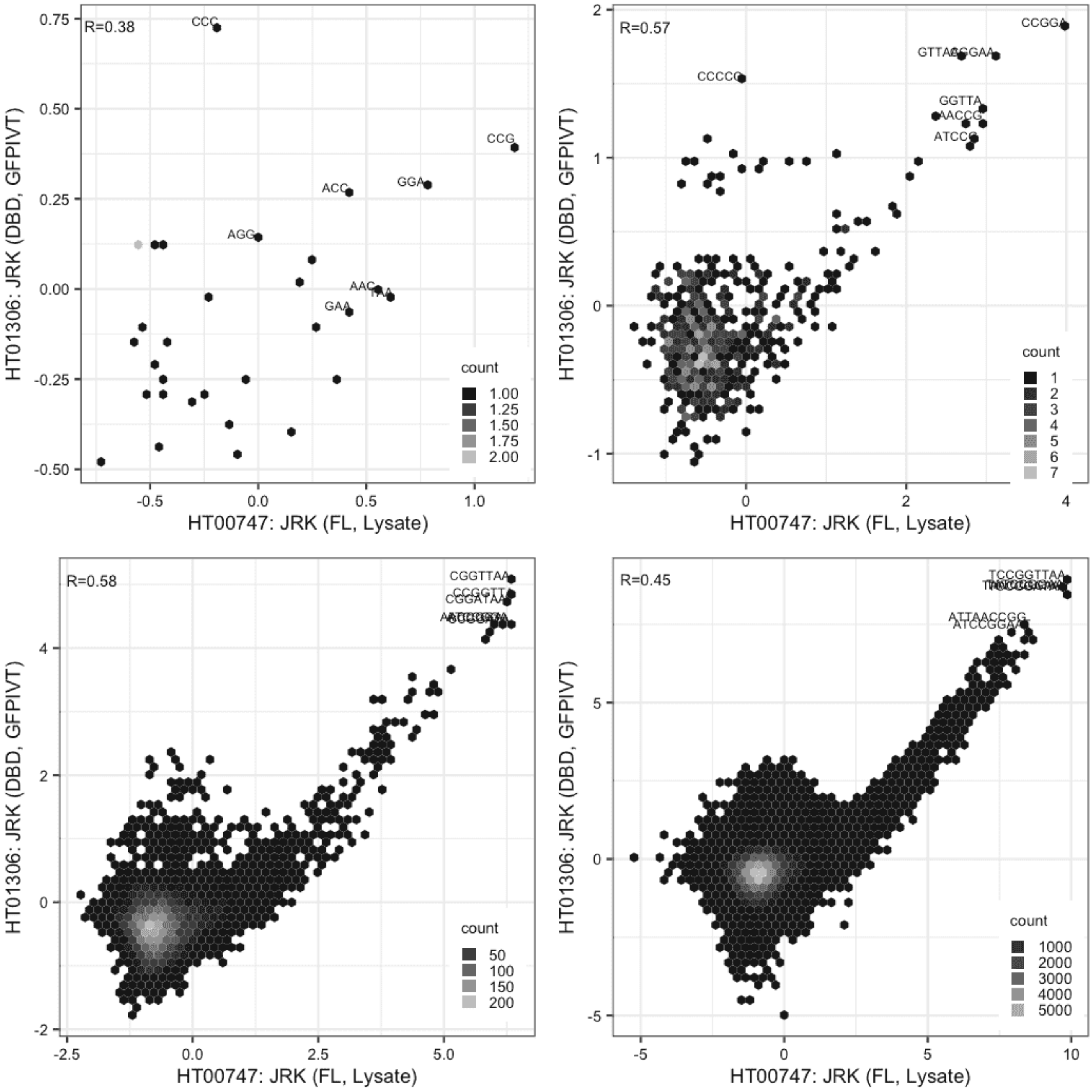

#### JRK

Lysate vs GFPIVT, same construct

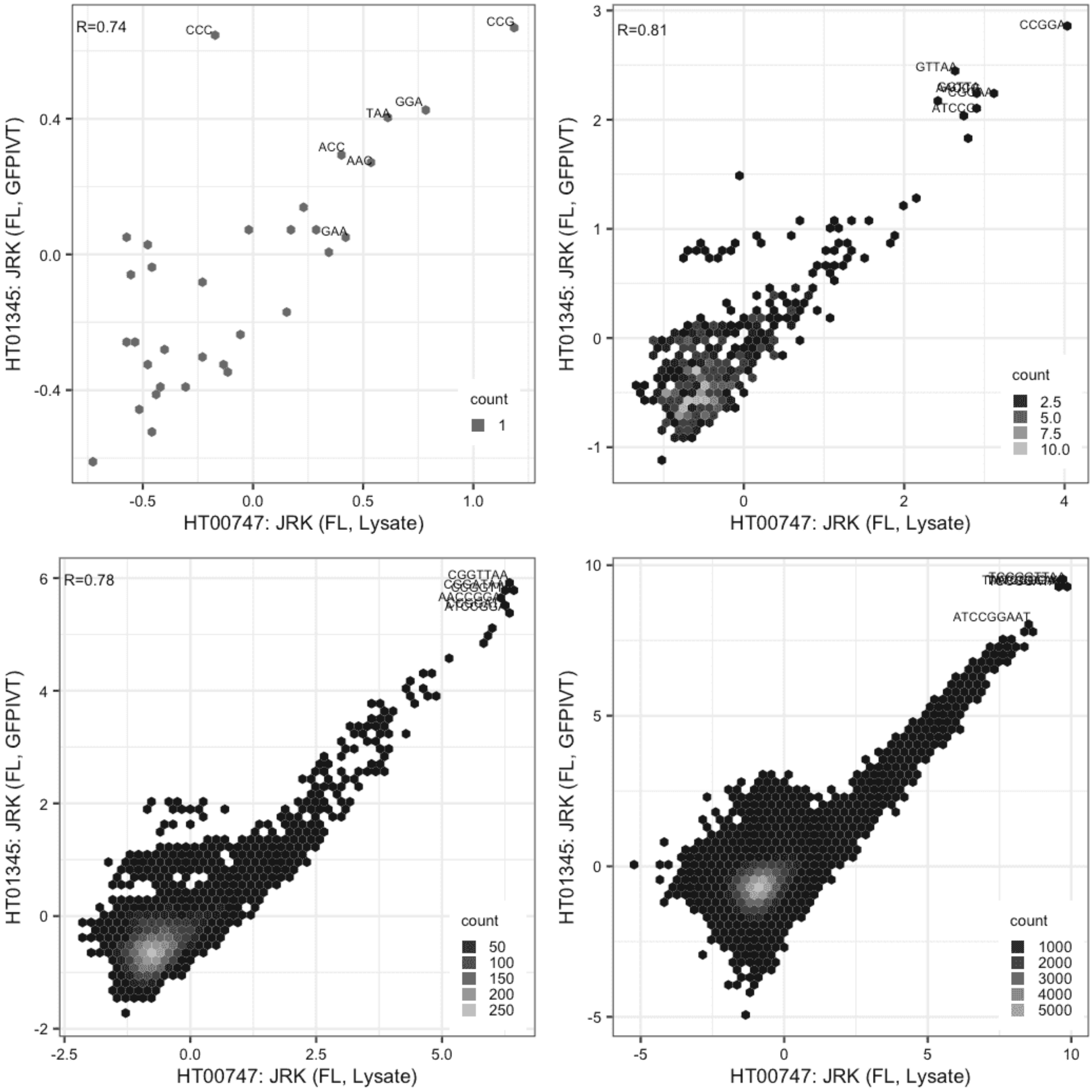

**JRK**

GFPIVT vs GFPIVT, different constructs

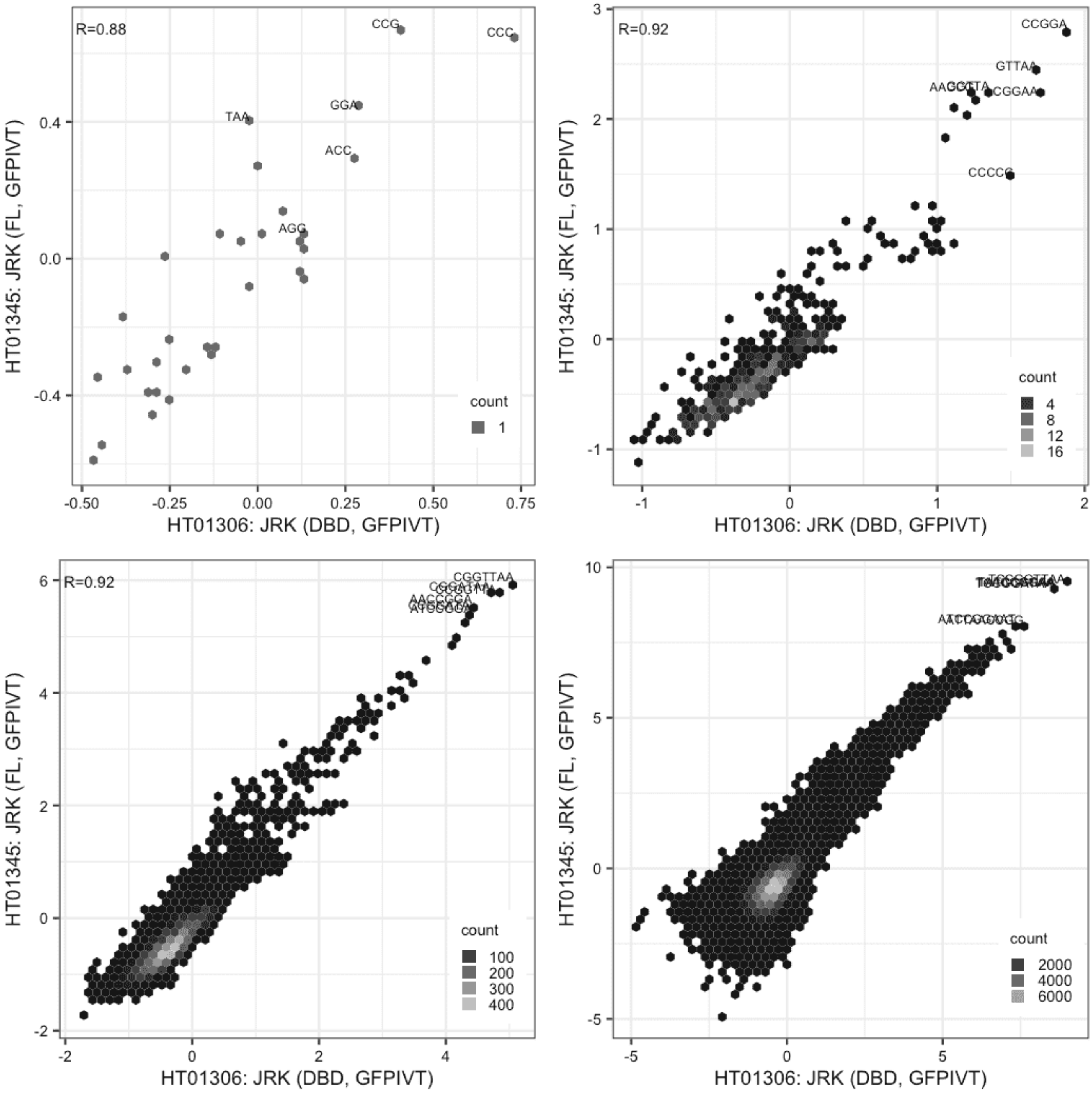

**KDM2A**

Lysate vs GFPIVT, different constructs

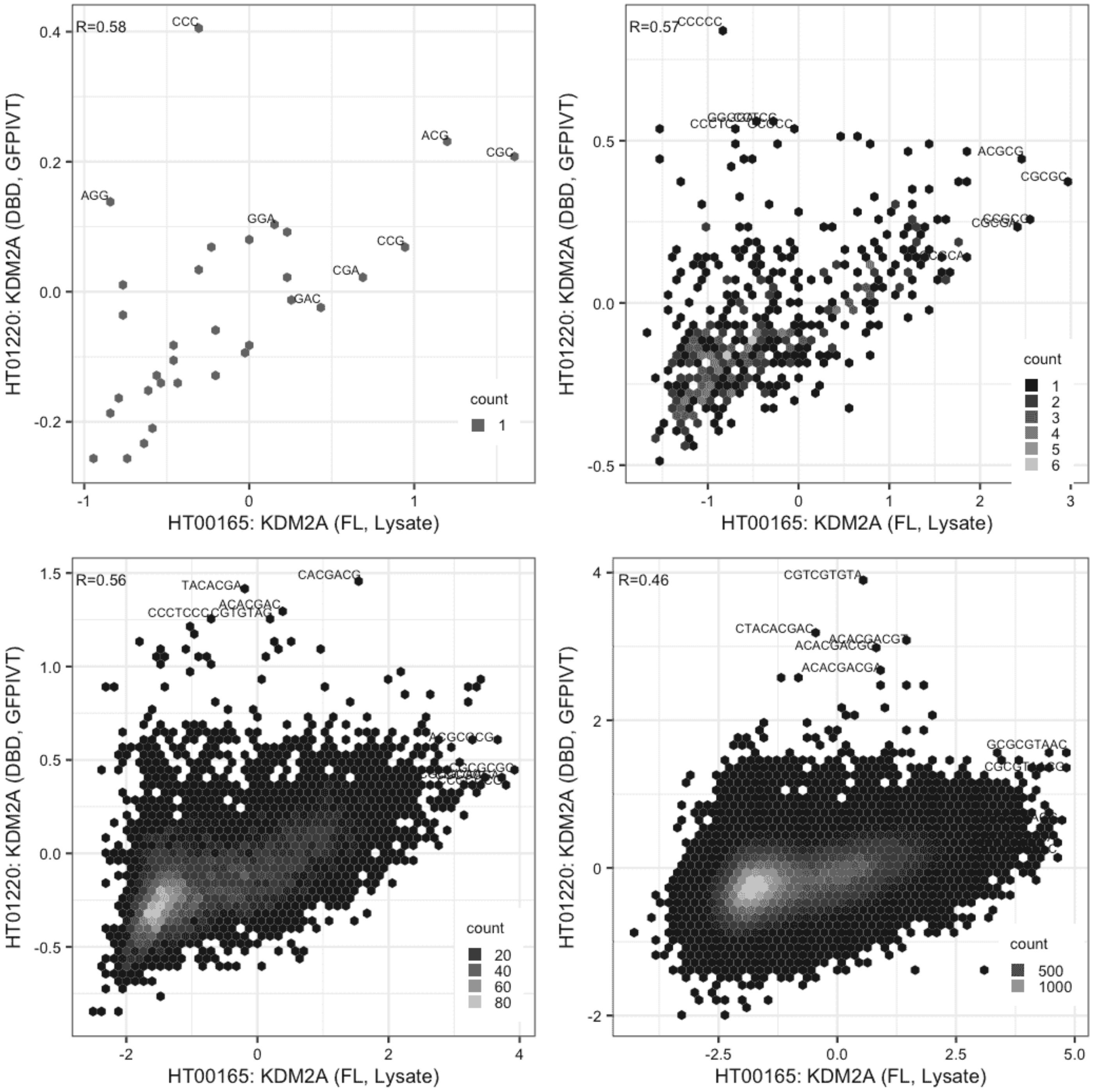

**LEF1**

Lysate vs IVT, same construct

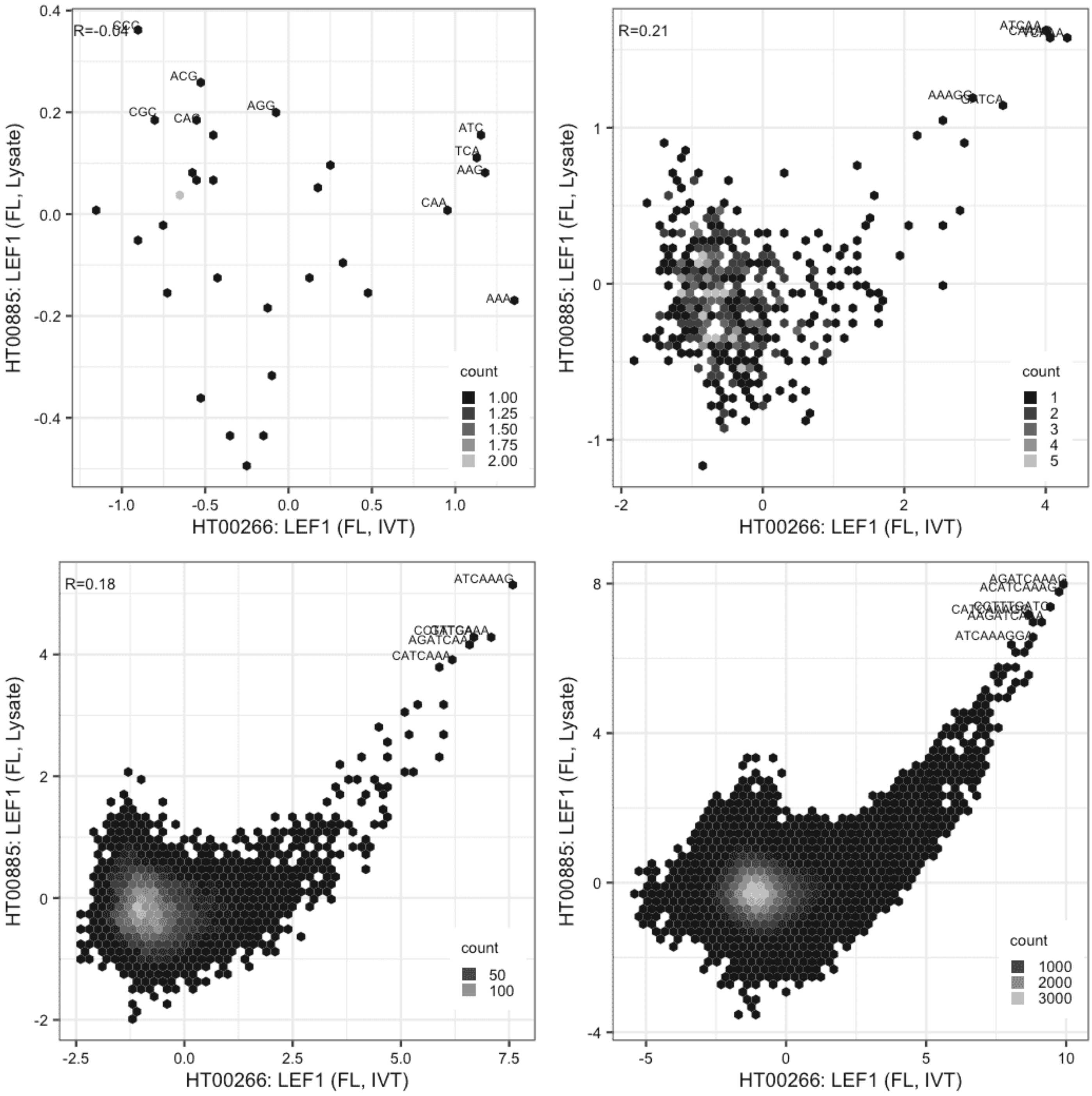

### LEUTX

Lysate vs Lysate, same construct

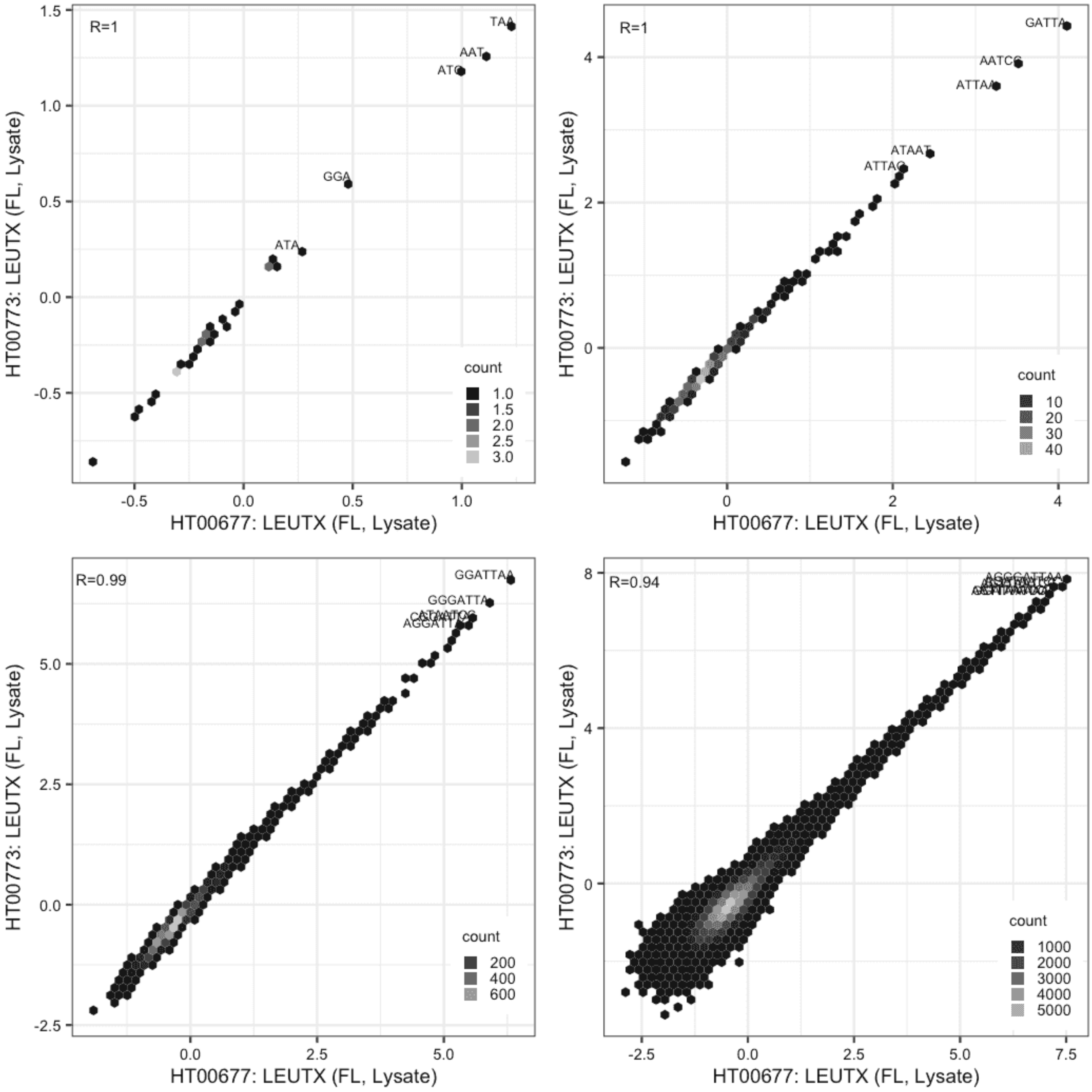

**LEUTX**

Lysate vs GFPIVT, same construct

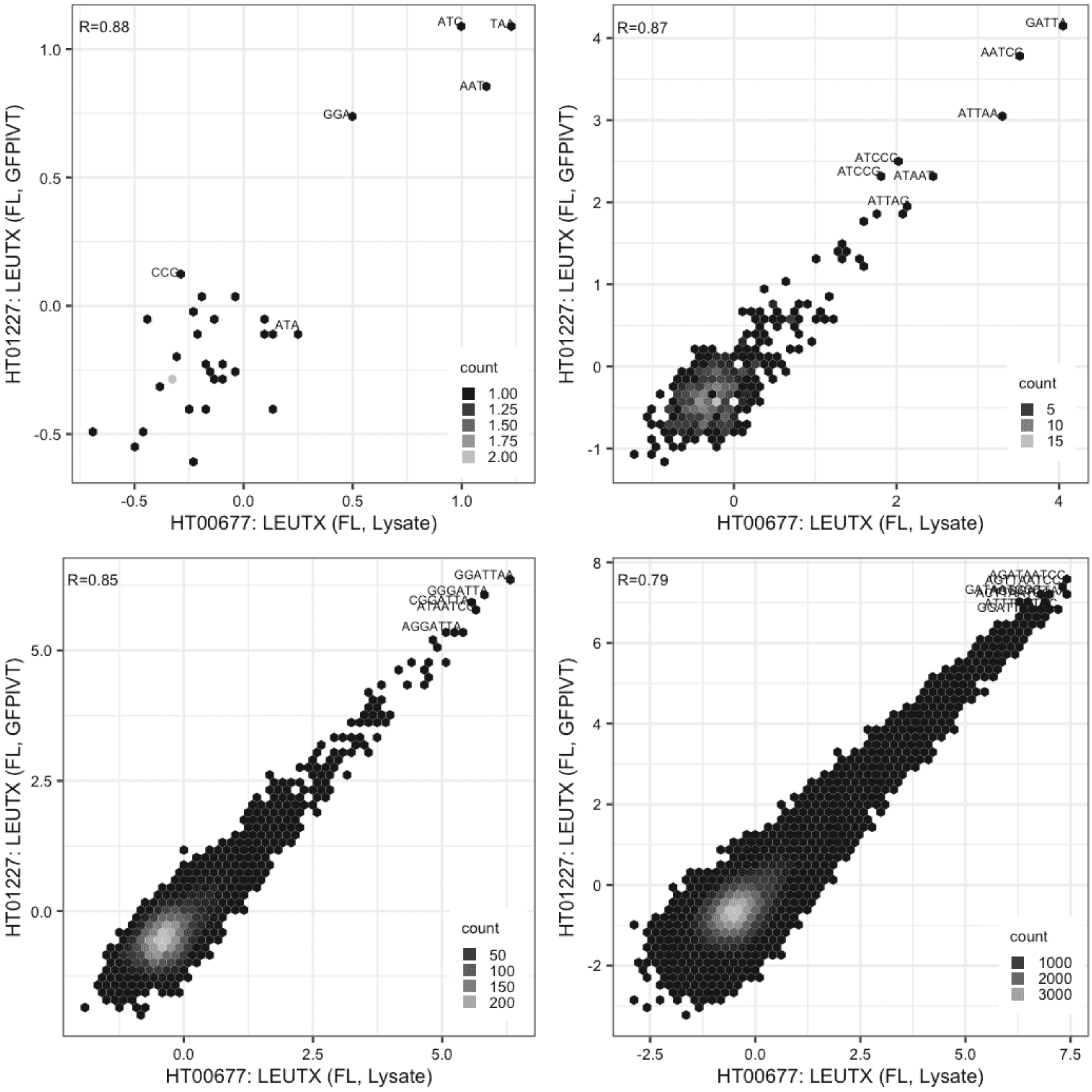

#### LEUTX

Lysate vs GFPIVT, same construct

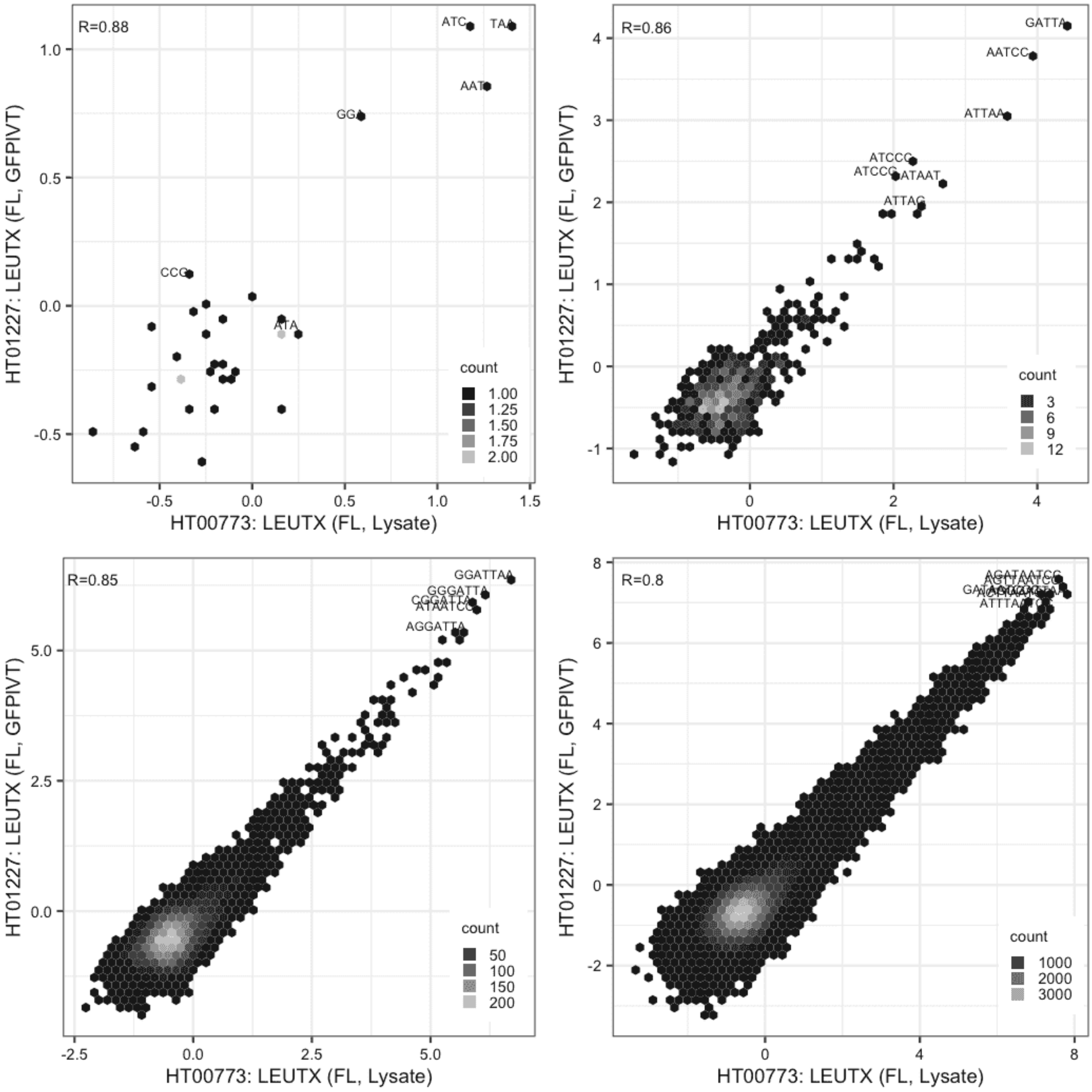

#### MAX

Lysate vs IVT, same construct

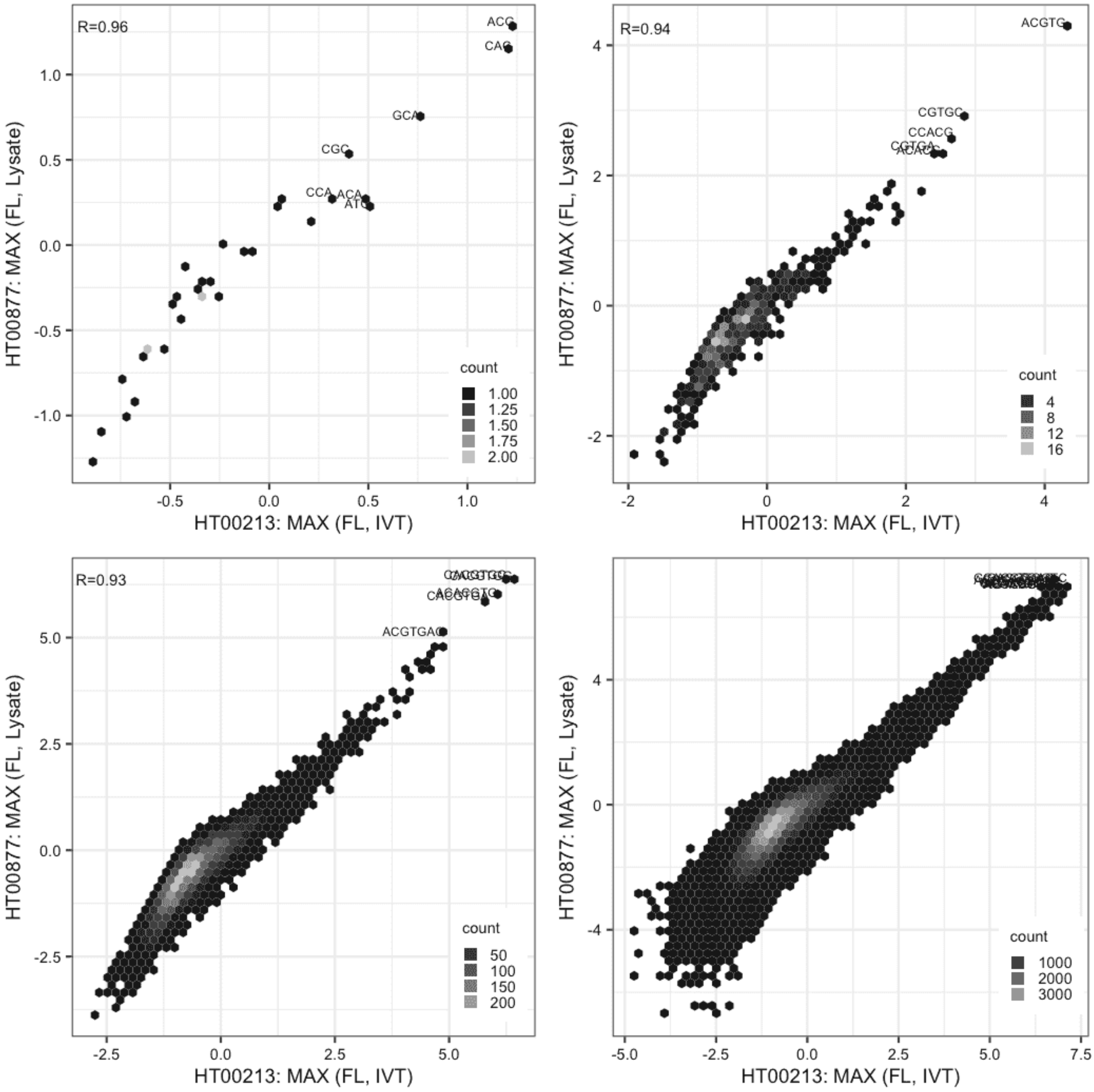

### MAX

Lysate vs IVT, same construct

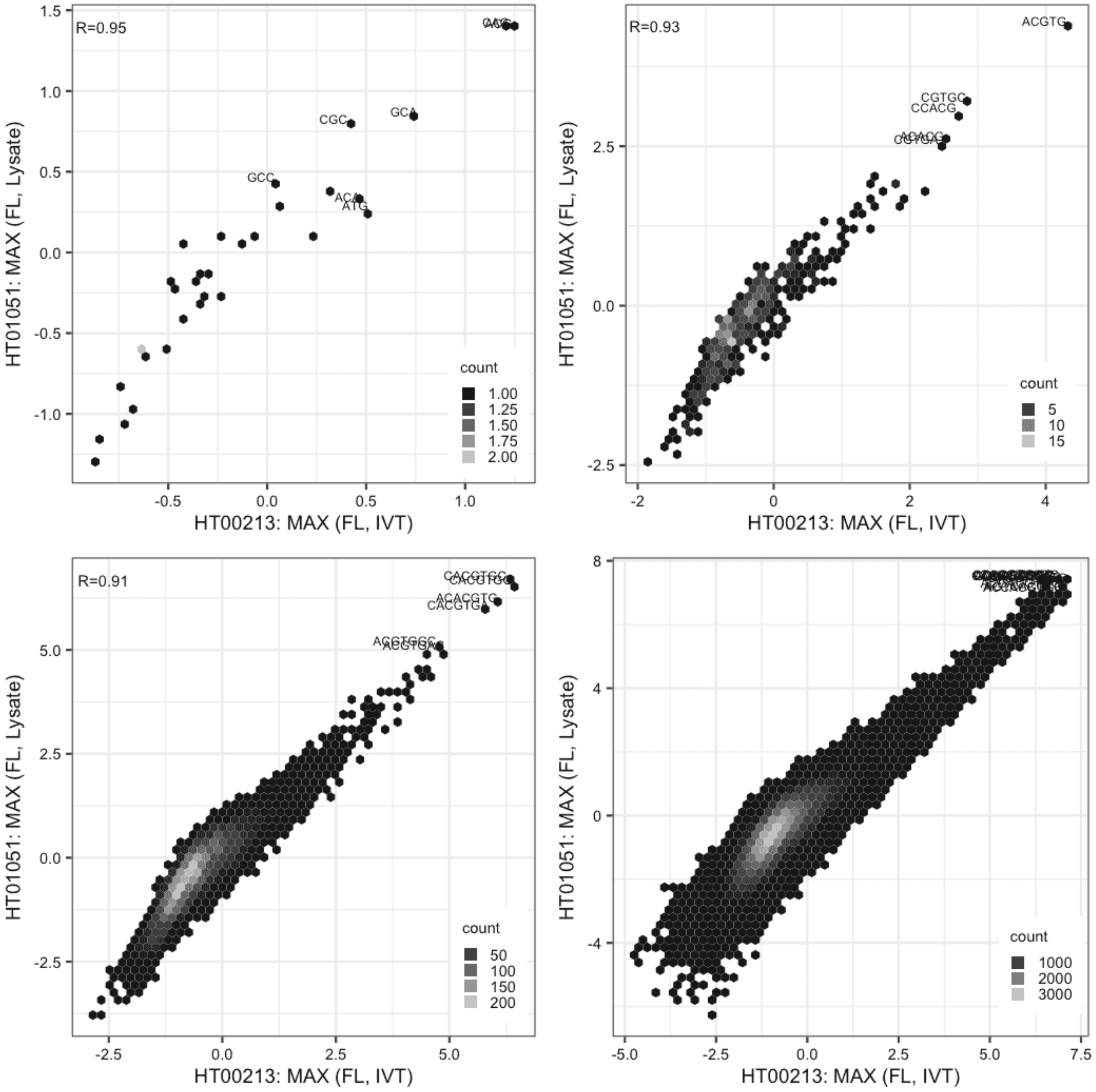

#### MAX

Lysate vs IVT, same construct

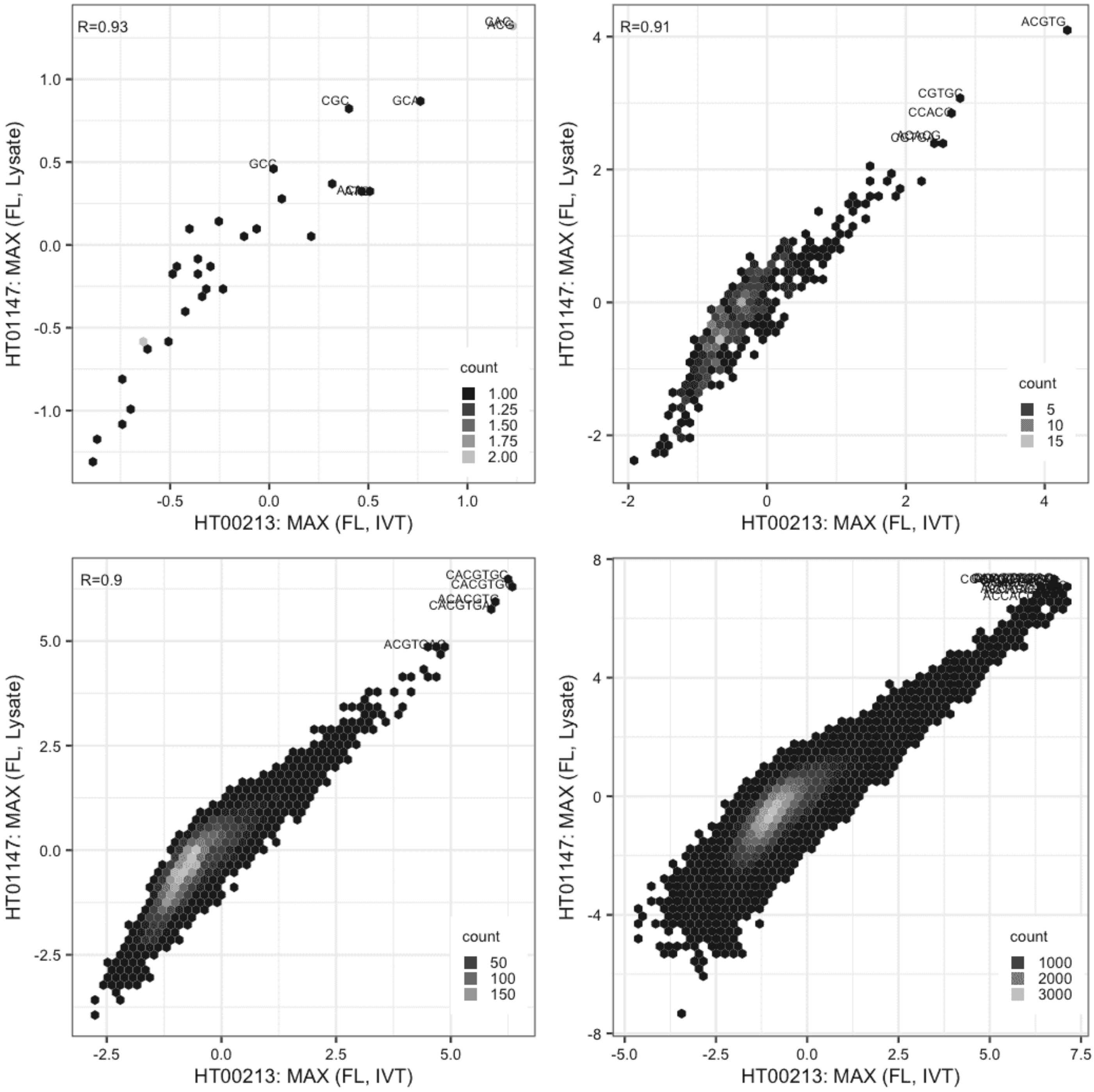

#### MAX

Lysate vs Lysate, same construct

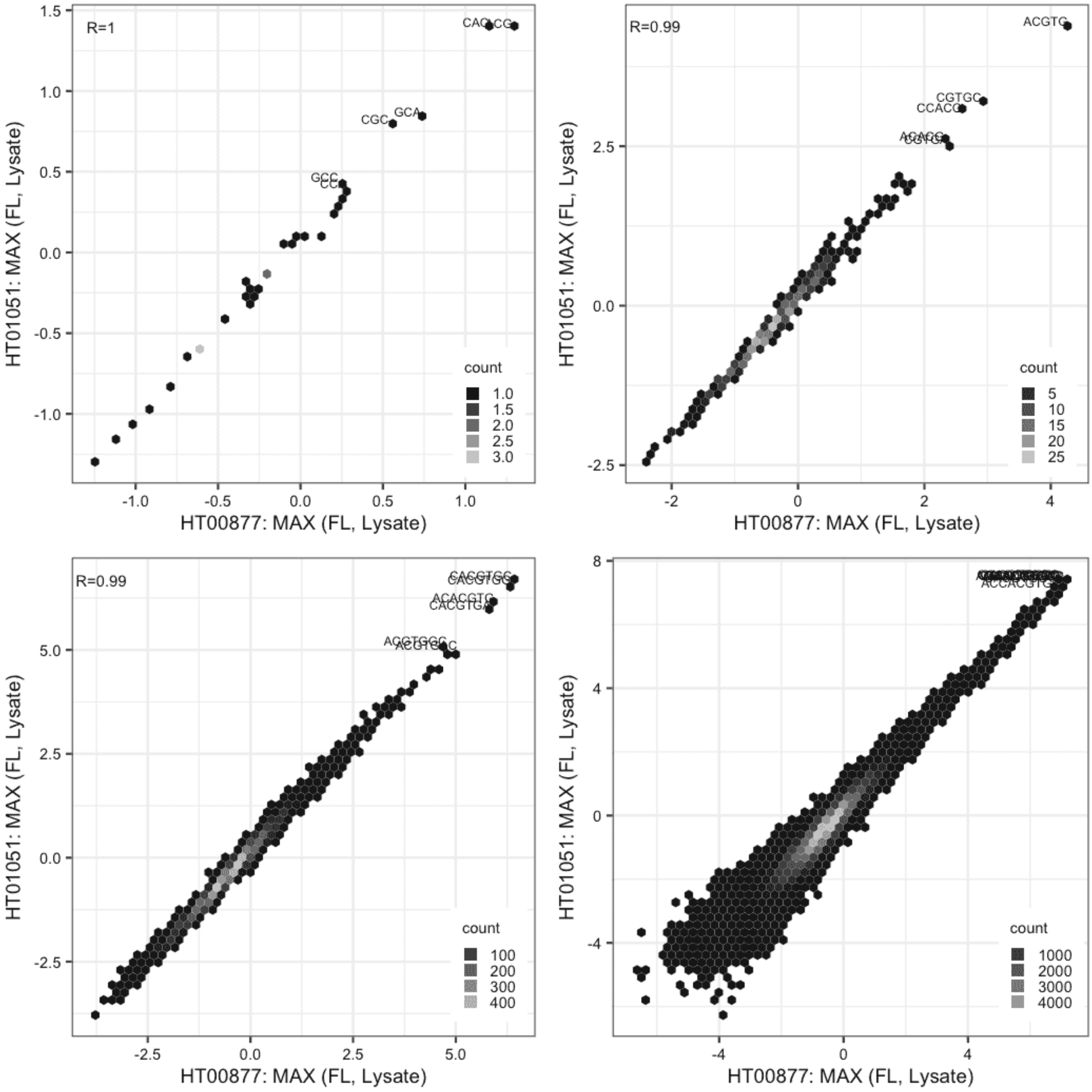

#### MAX

Lysate vs Lysate, same construct

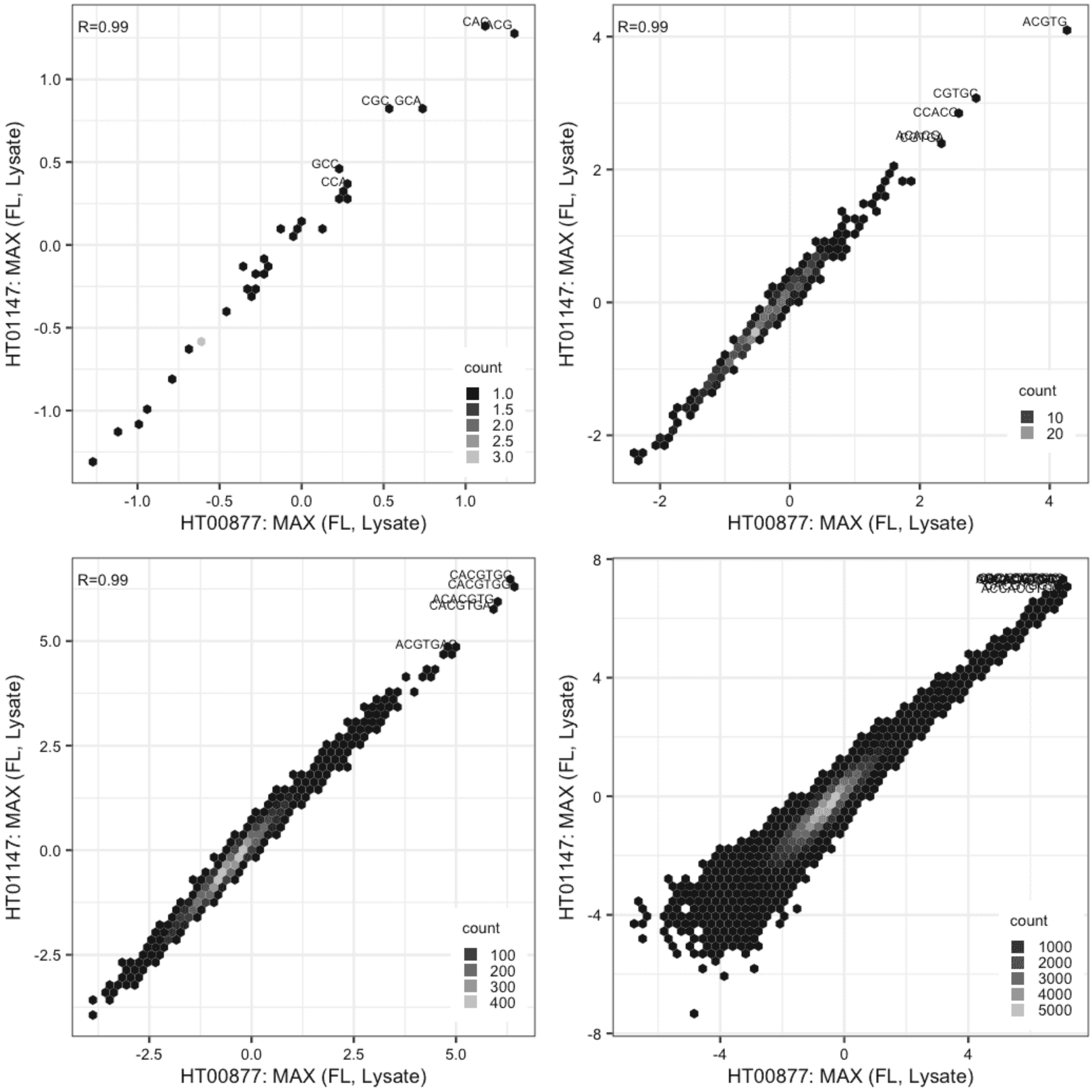

### MAX

Lysate vs Lysate, same construct

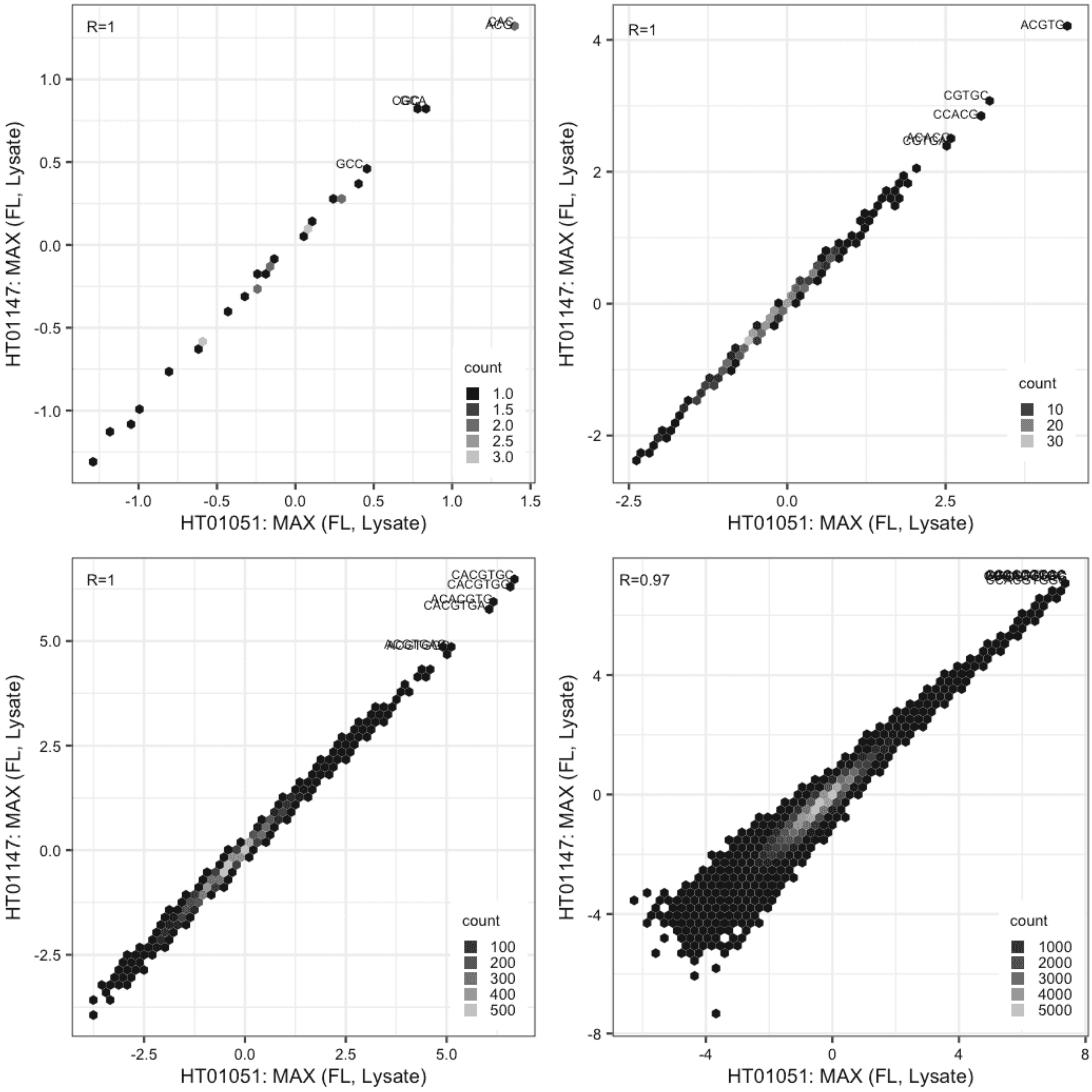

**MKX**

IVT vs IVT, different constructs

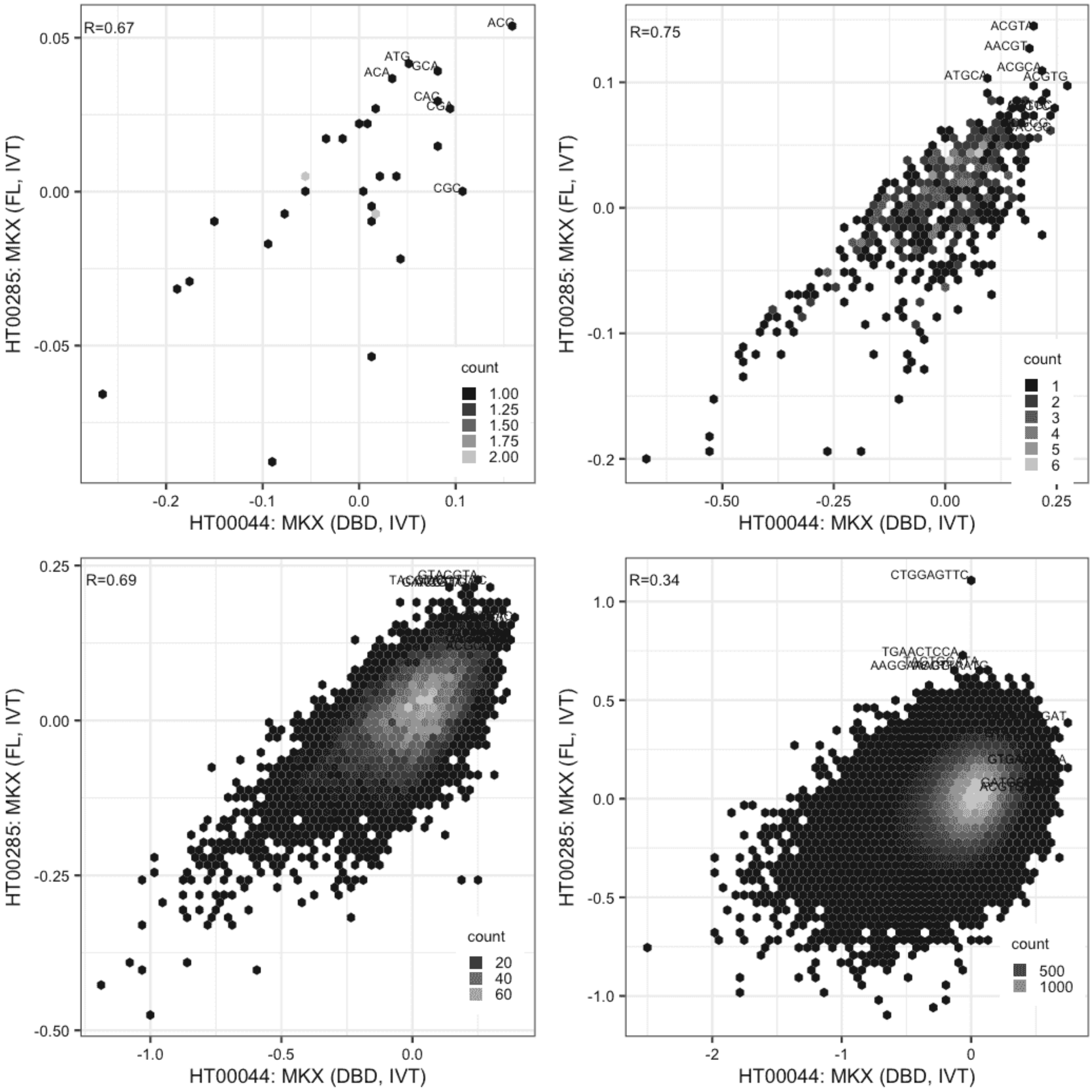

**MKX**

Lysate vs IVT, different constructs

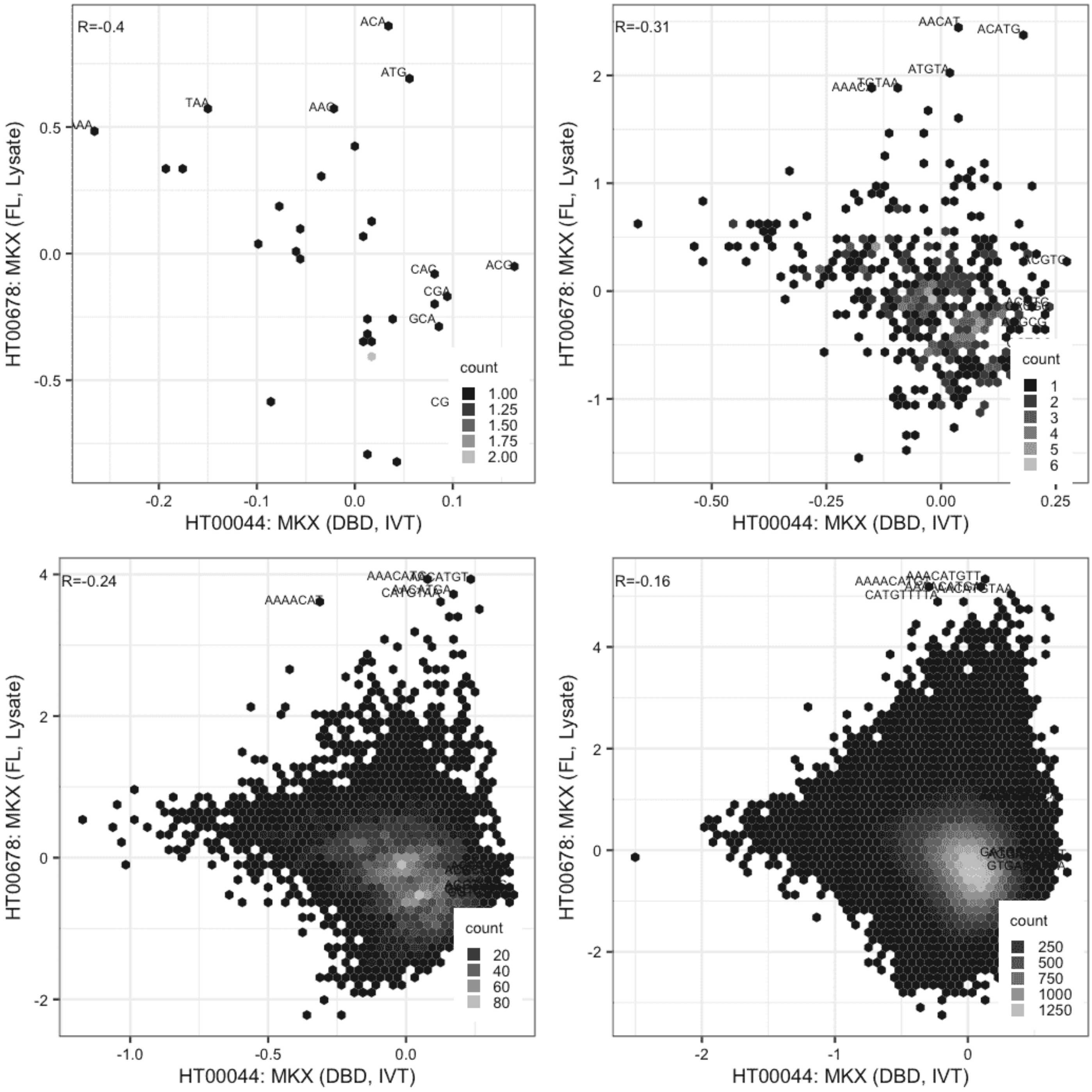

### MKX

Lysate vs IVT, different constructs

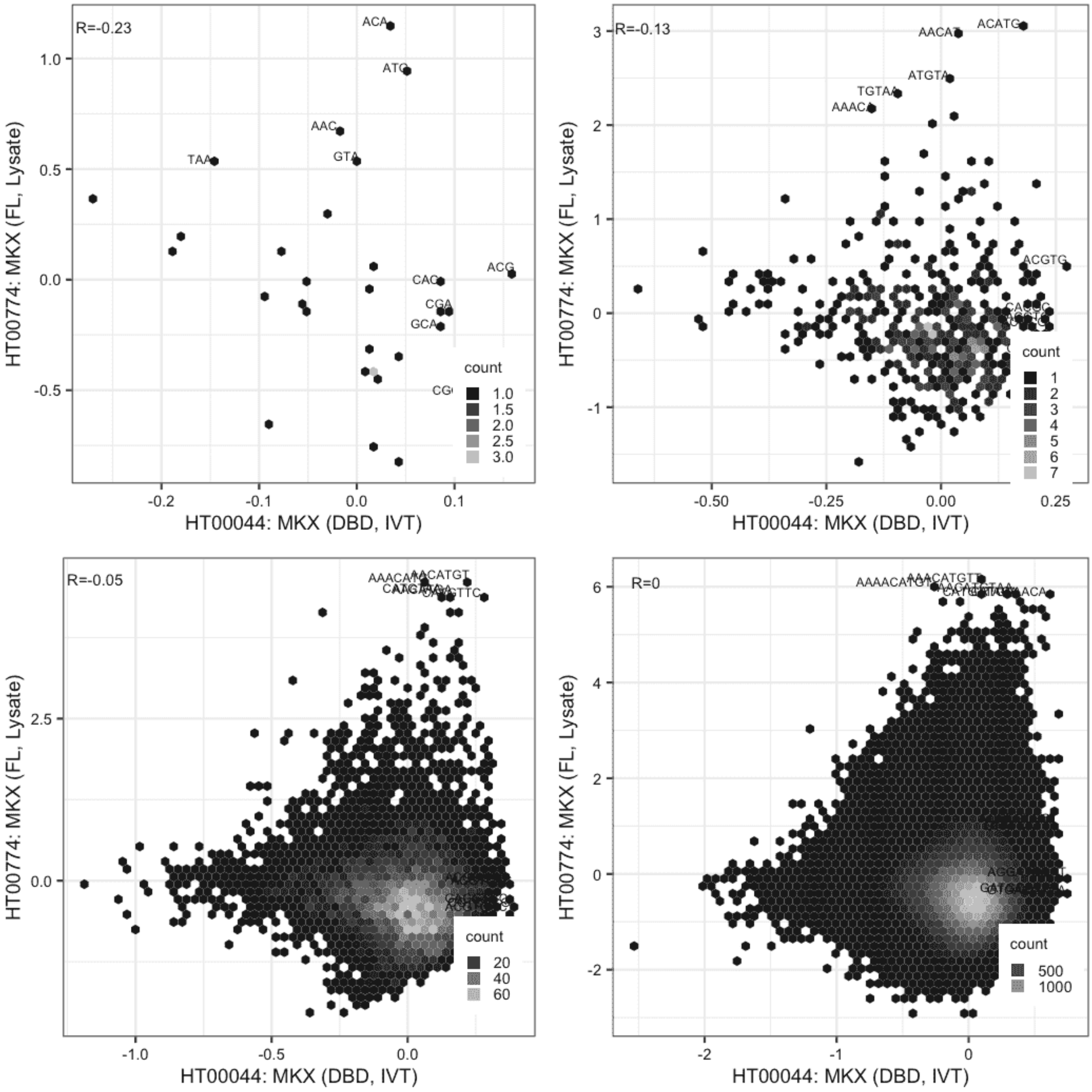

### MKX

IVT vs GFPIVT, different constructs

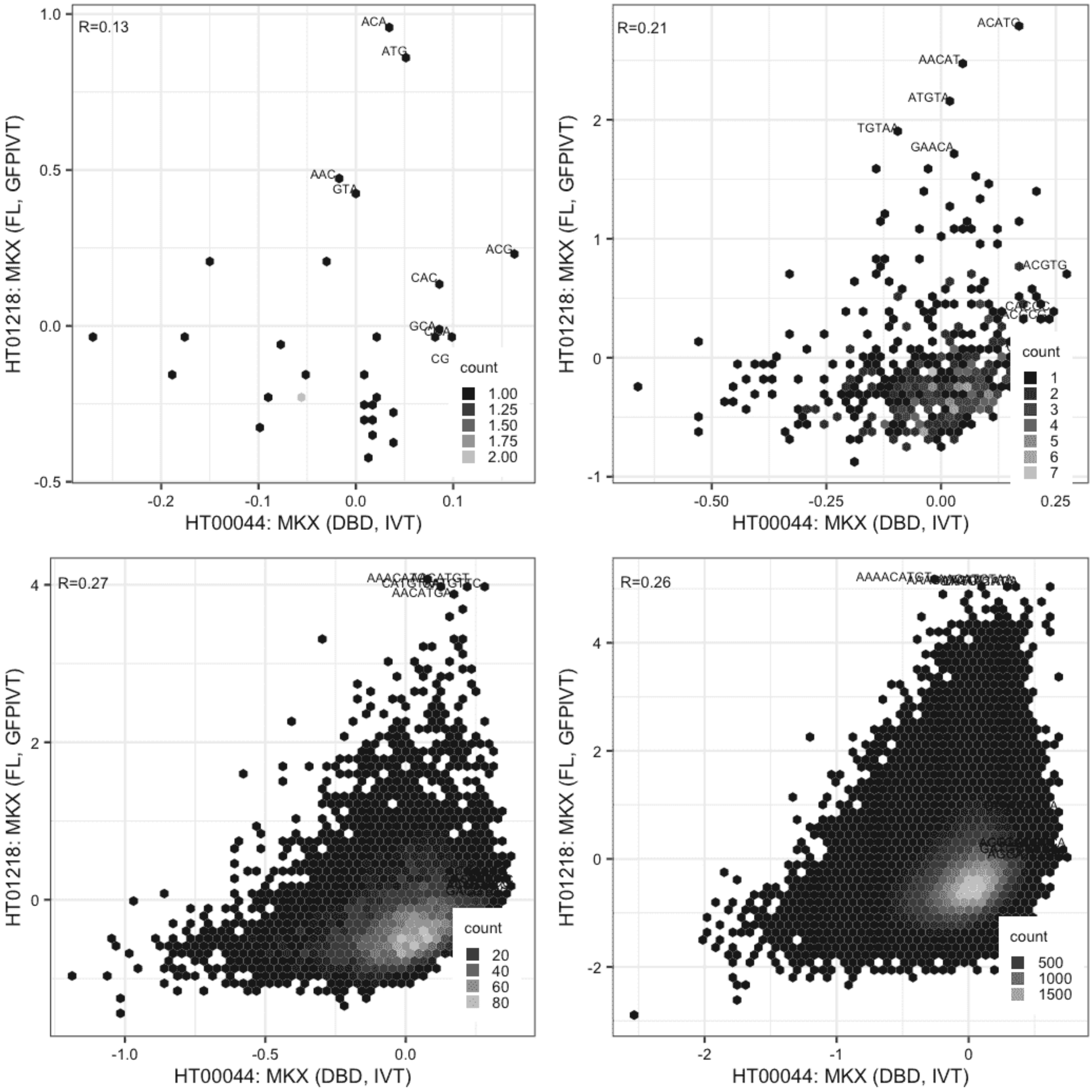

### MKX

Lysate vs IVT, same construct

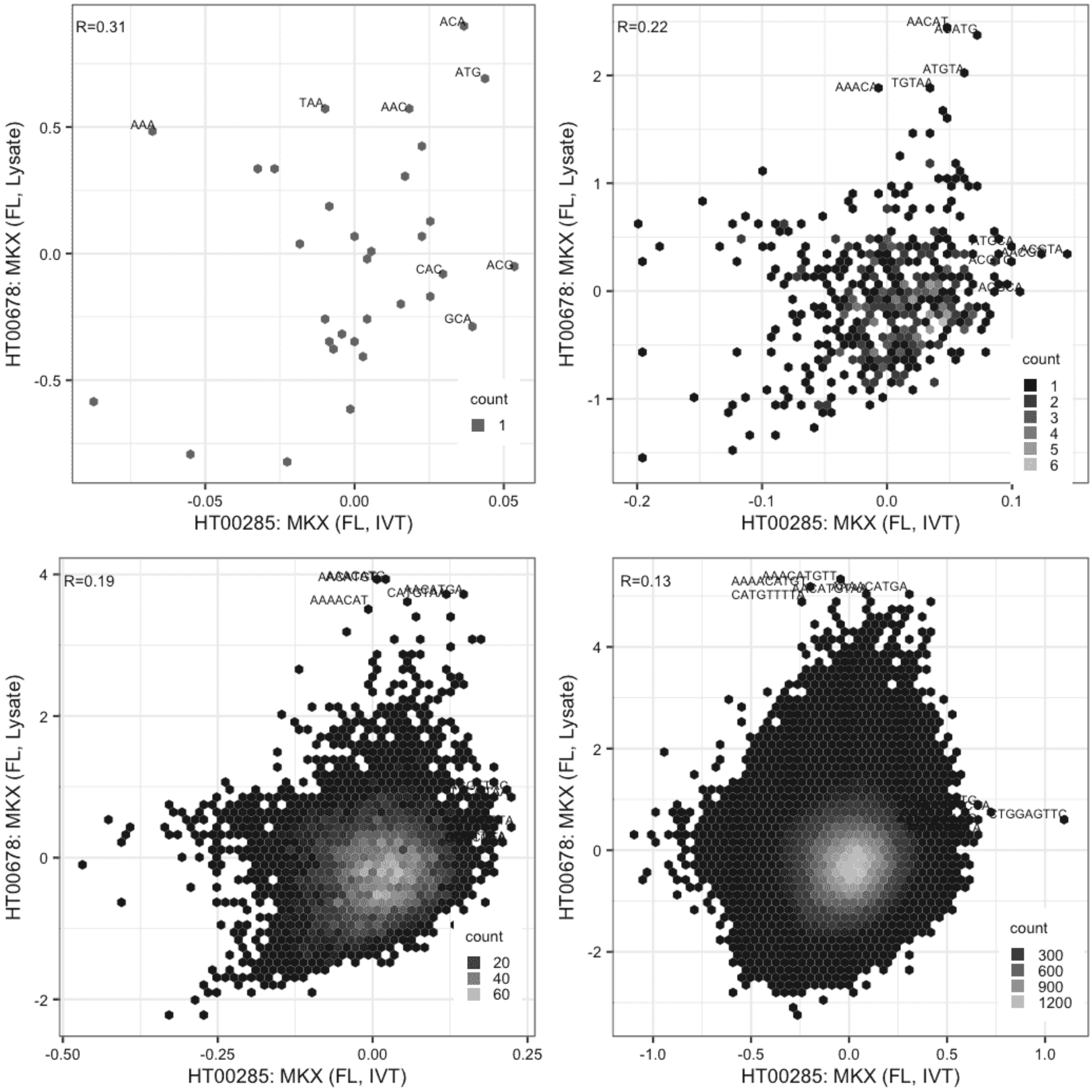

**MKX**

Lysate vs IVT, same construct

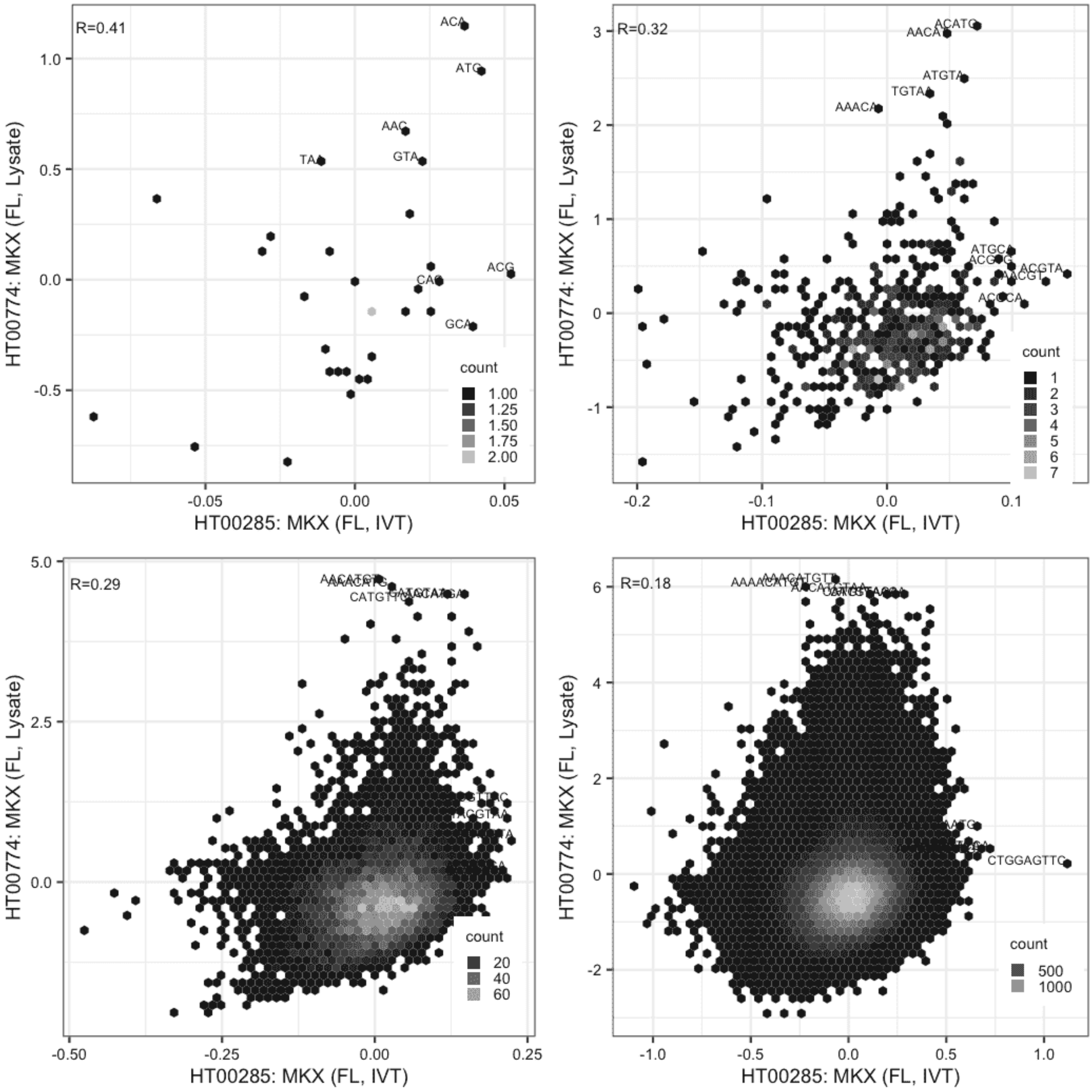

**MKX**

IVT vs GFPIVT, same construct

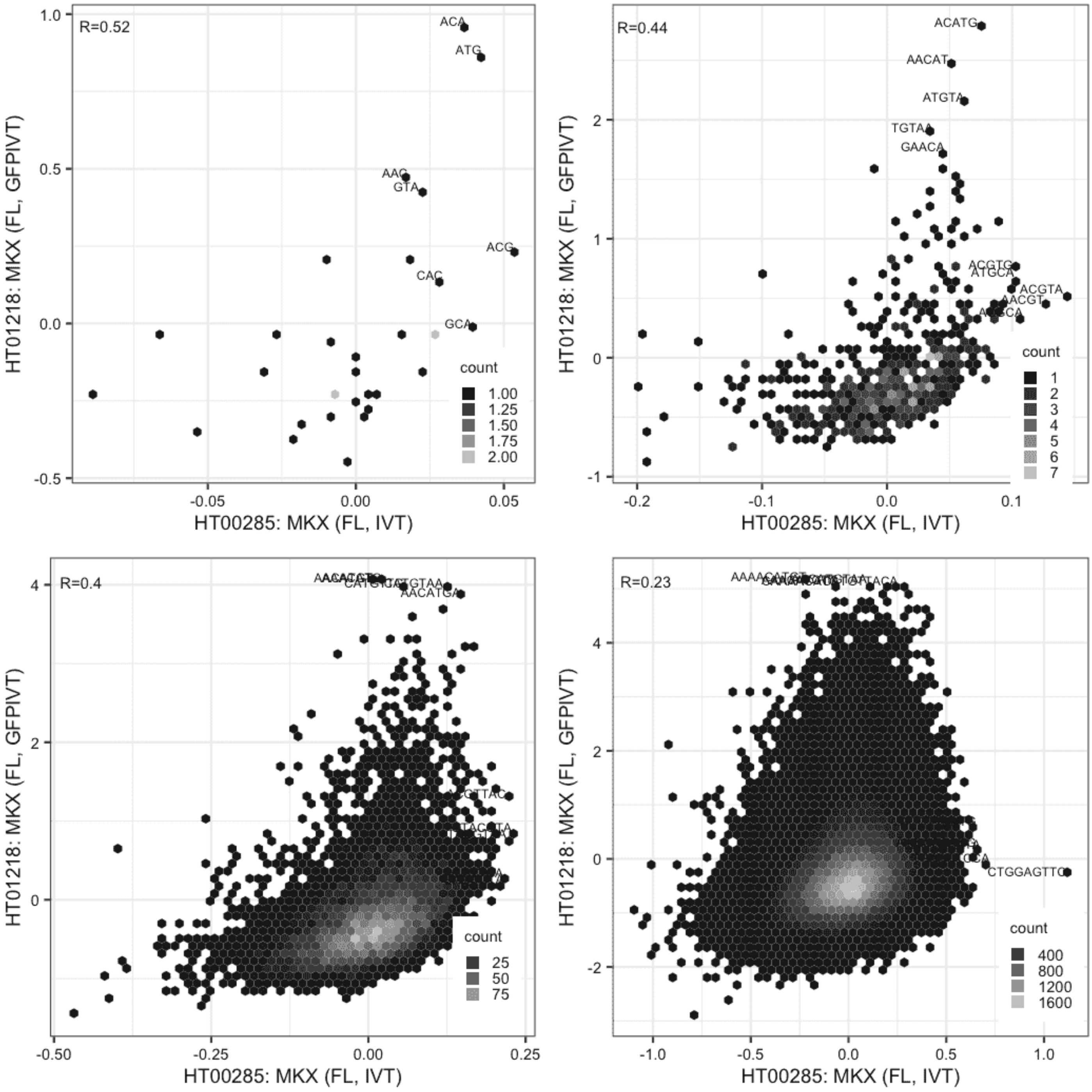

**MKX**

Lysate vs Lysate, same construct

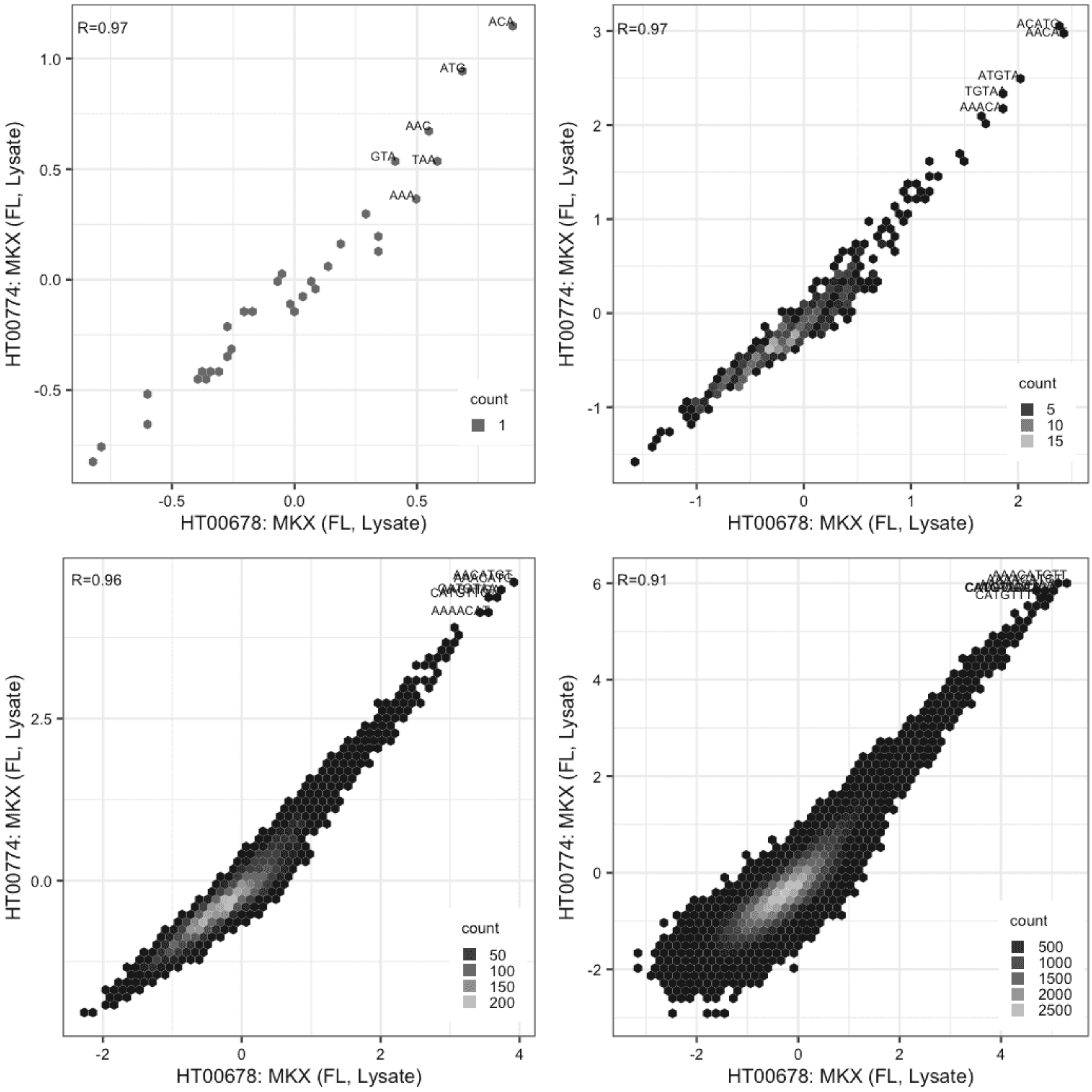

**MKX**

Lysate vs GFPIVT, same construct

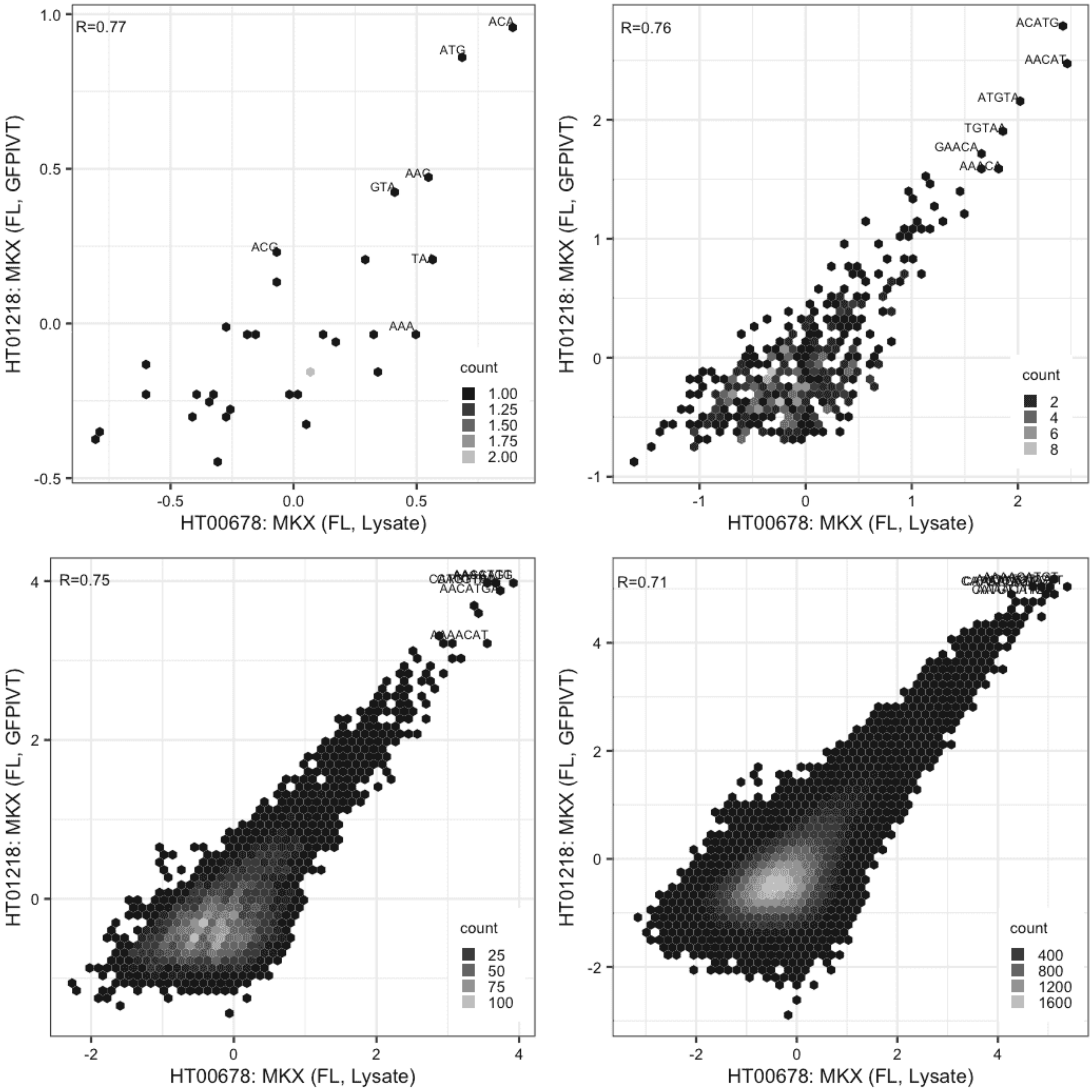

**MKX**

Lysate vs GFPIVT, same construct

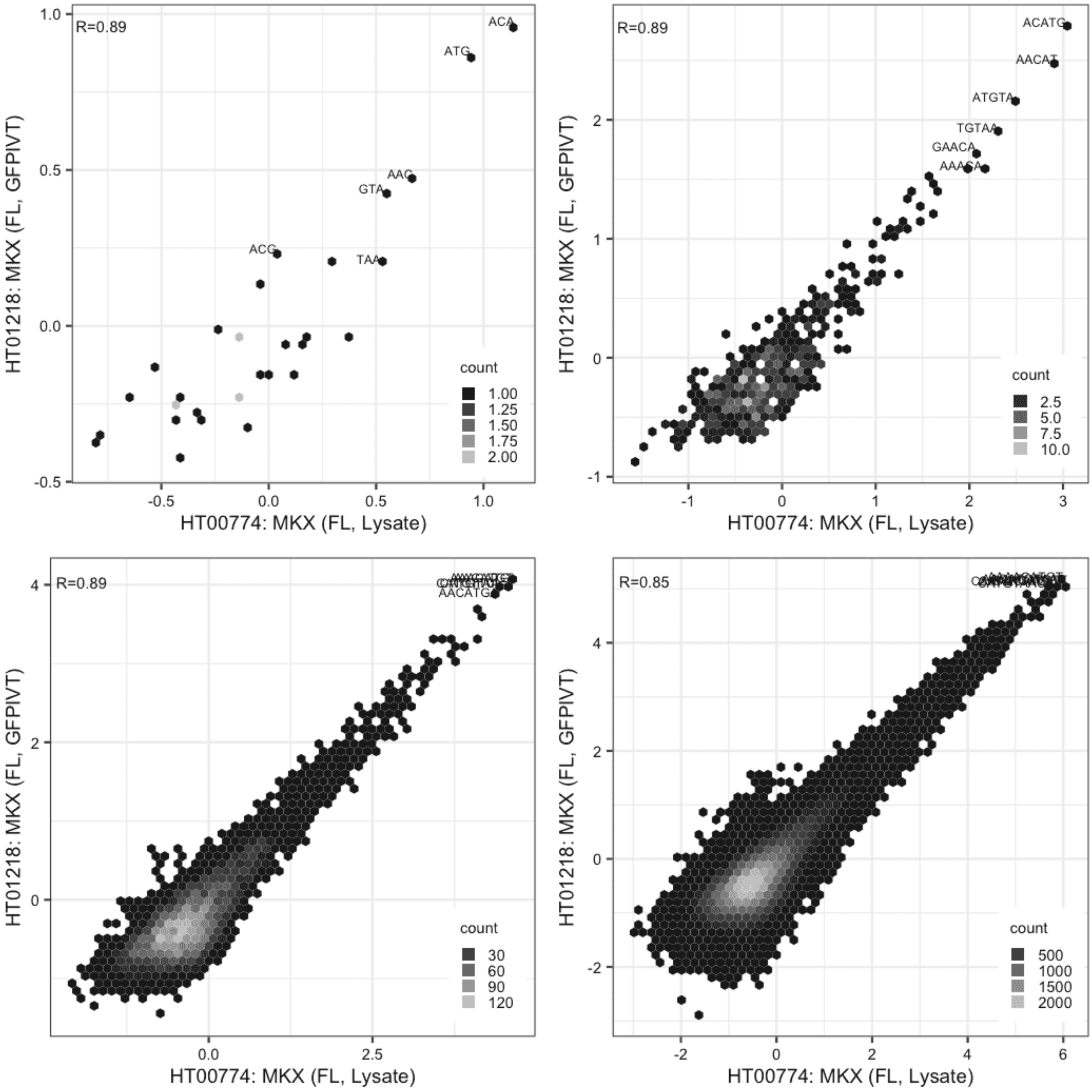

#### MSANTD1

Lysate vs IVT, different constructs

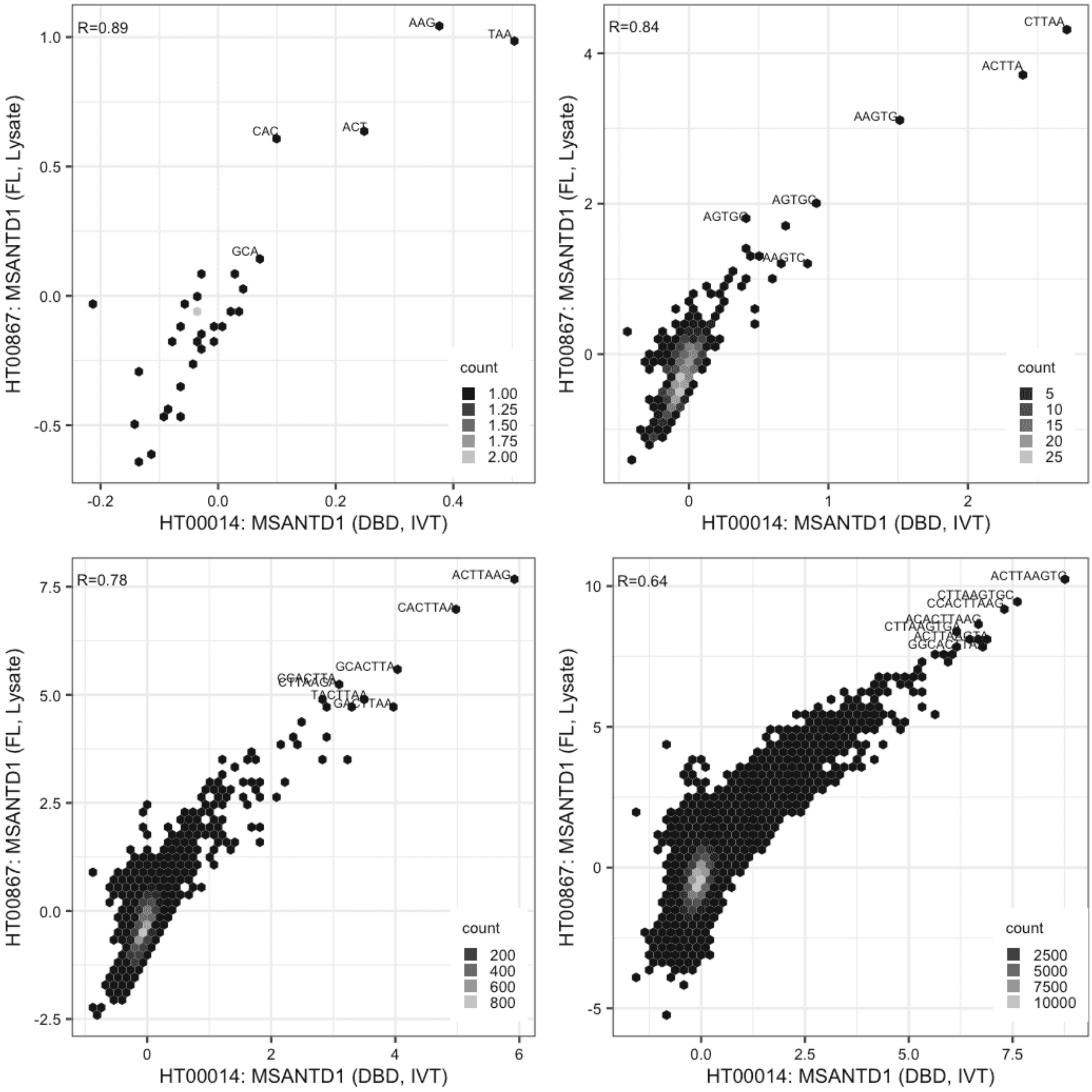

**MSANTD1**

IVT vs GFPIVT, different constructs

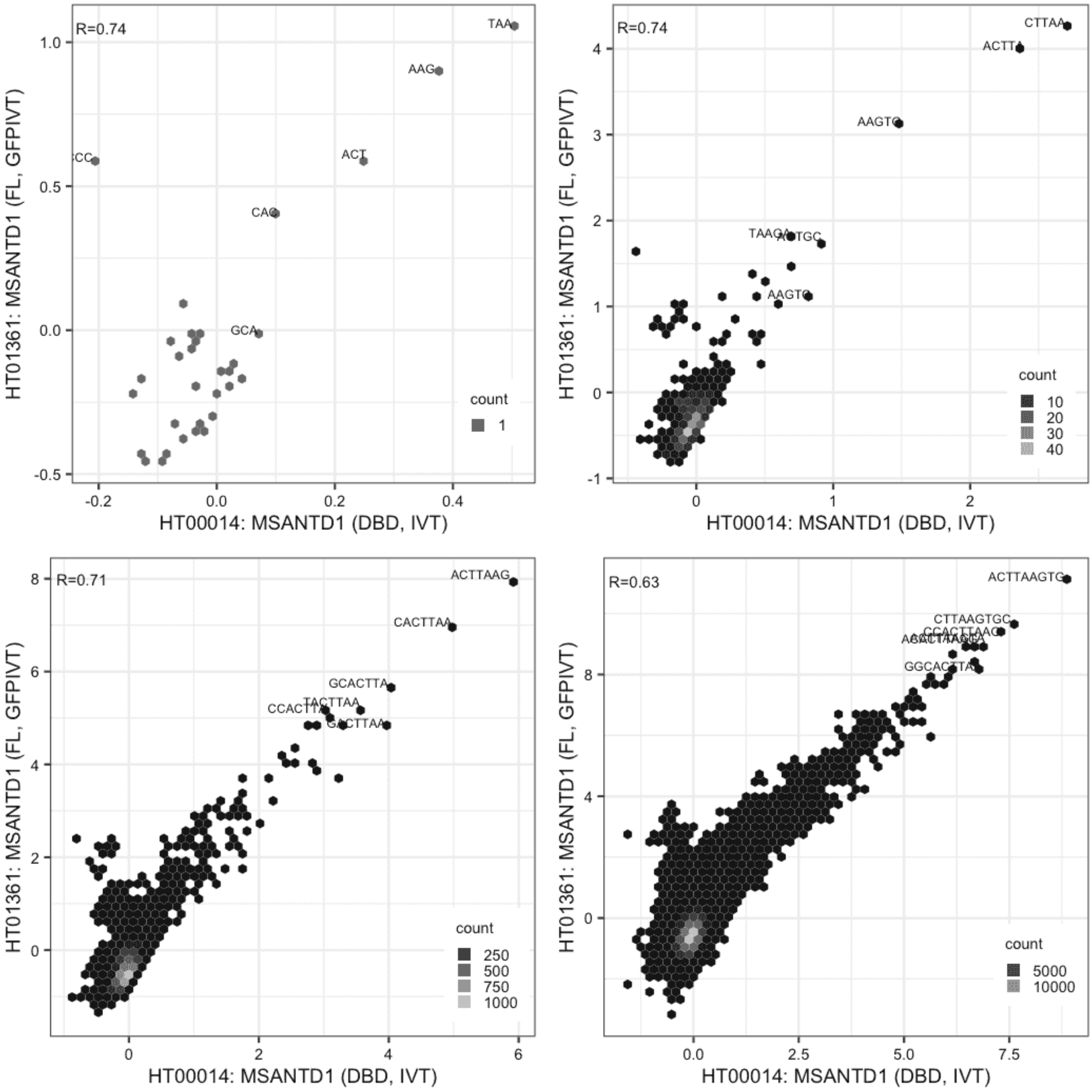

**MSANTD1**

Lysate vs GFPIVT, same construct

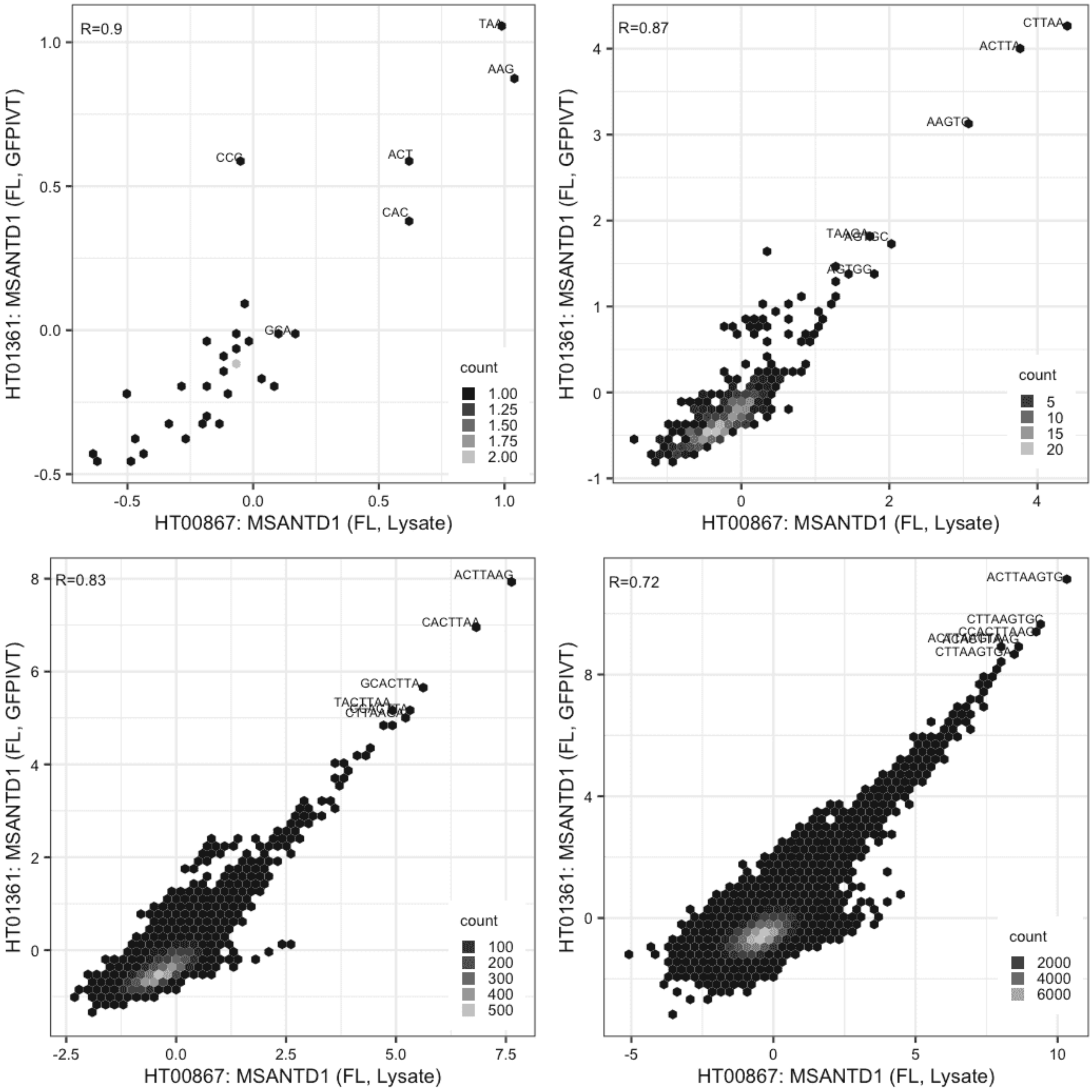

### MSANTD4

Lysate vs GFPIVT, same construct

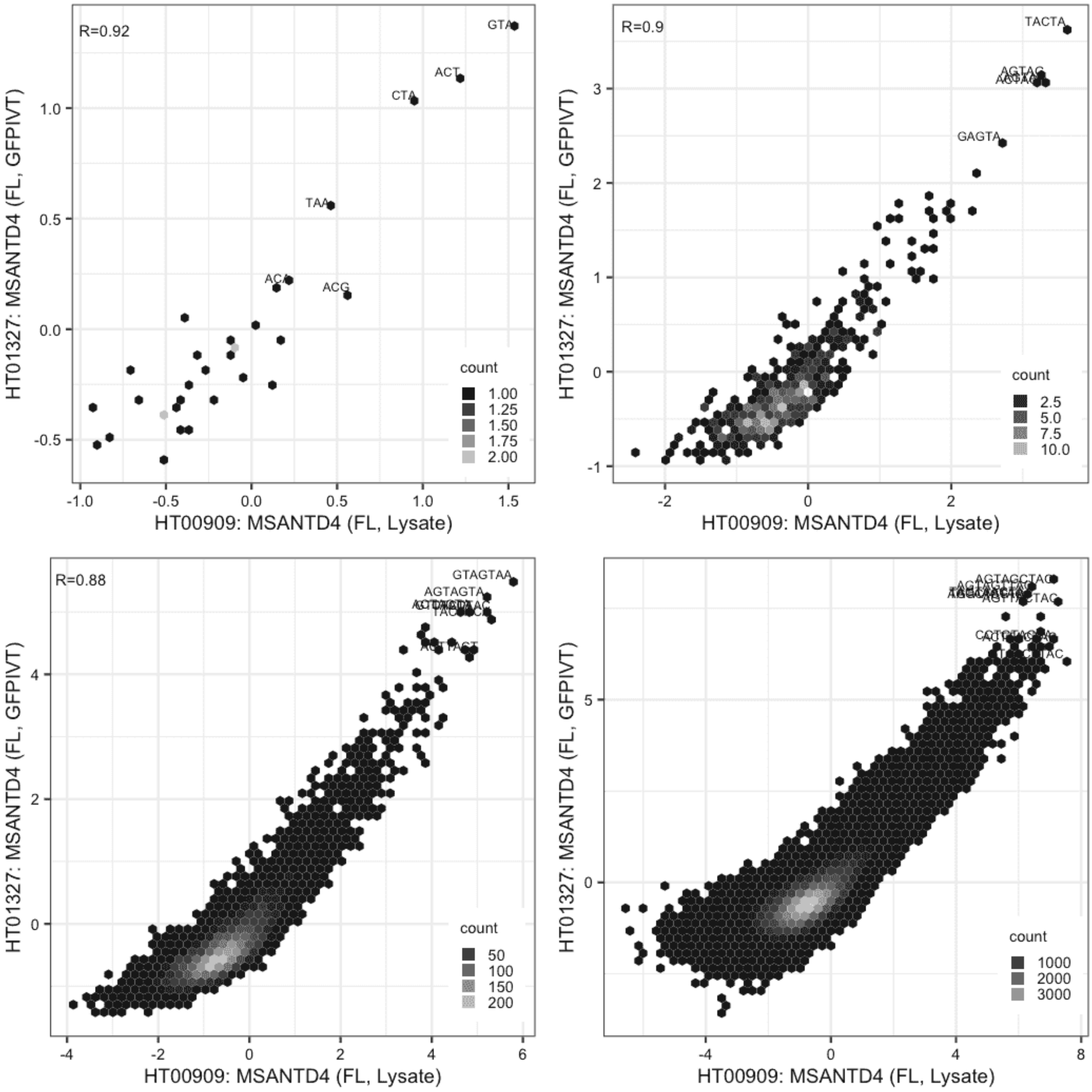

**MYF6**

Lysate vs IVT, same construct

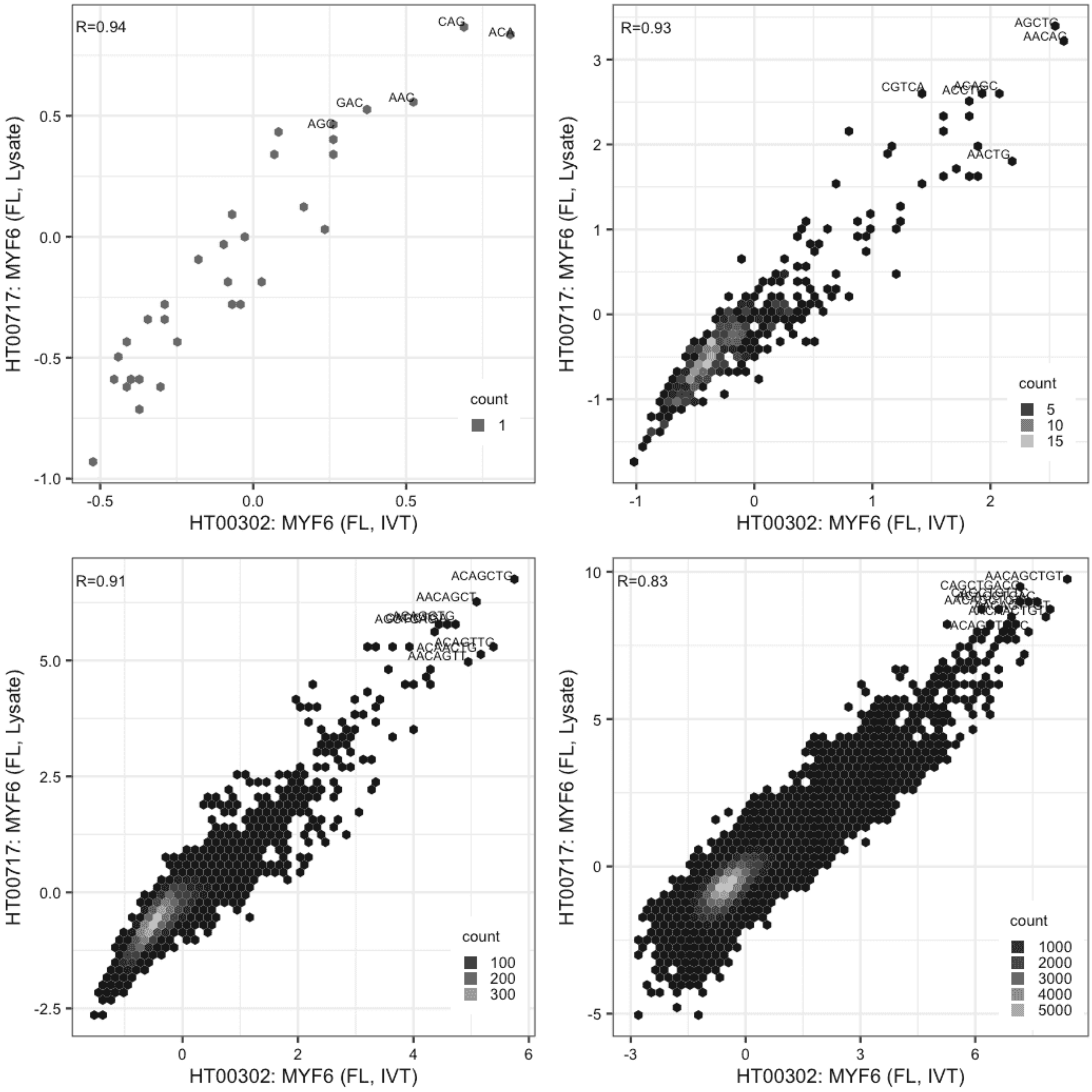

**MYF6**

Lysate vs IVT, same construct

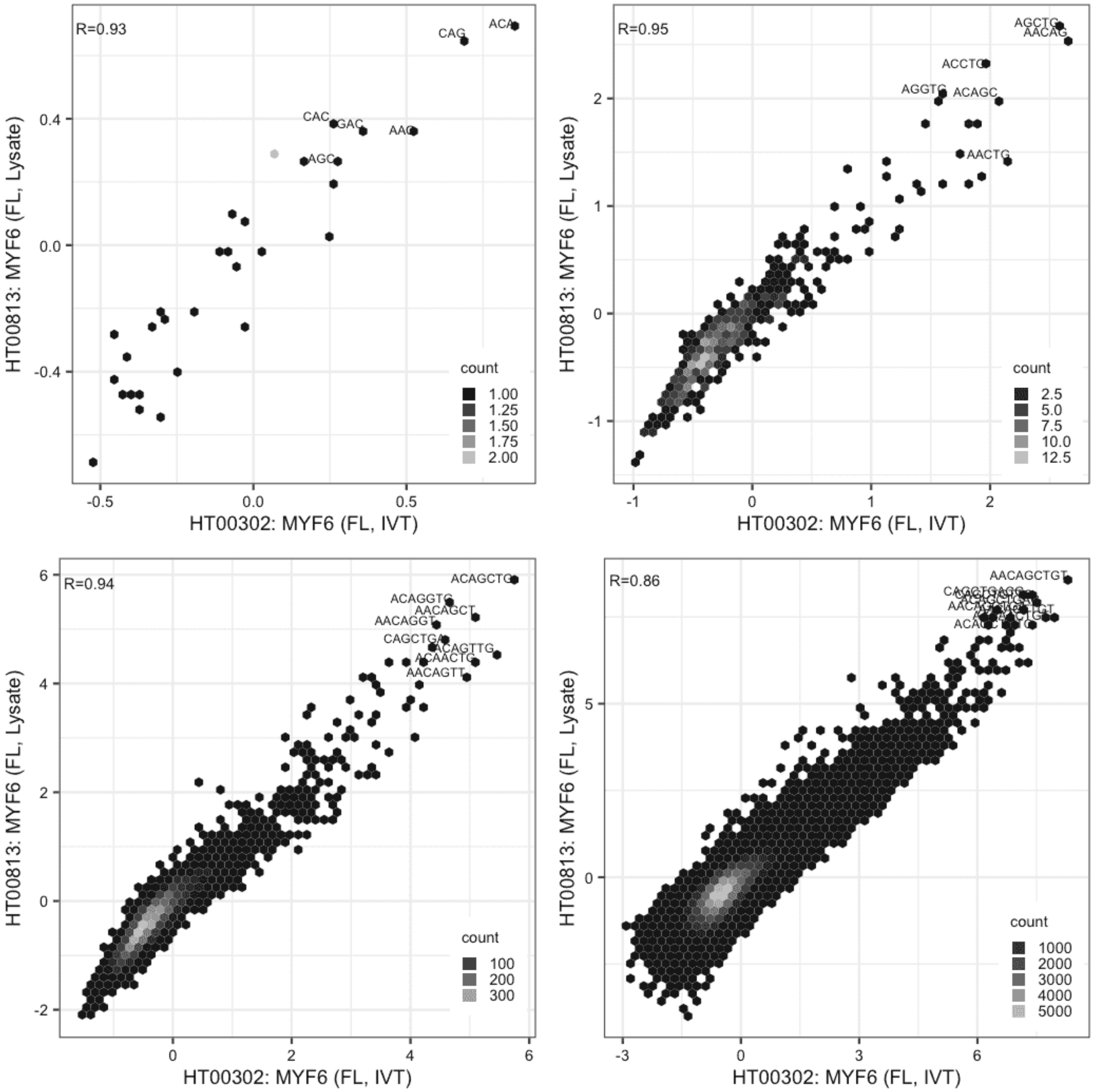

**MYF6**

Lysate vs Lysate, same construct

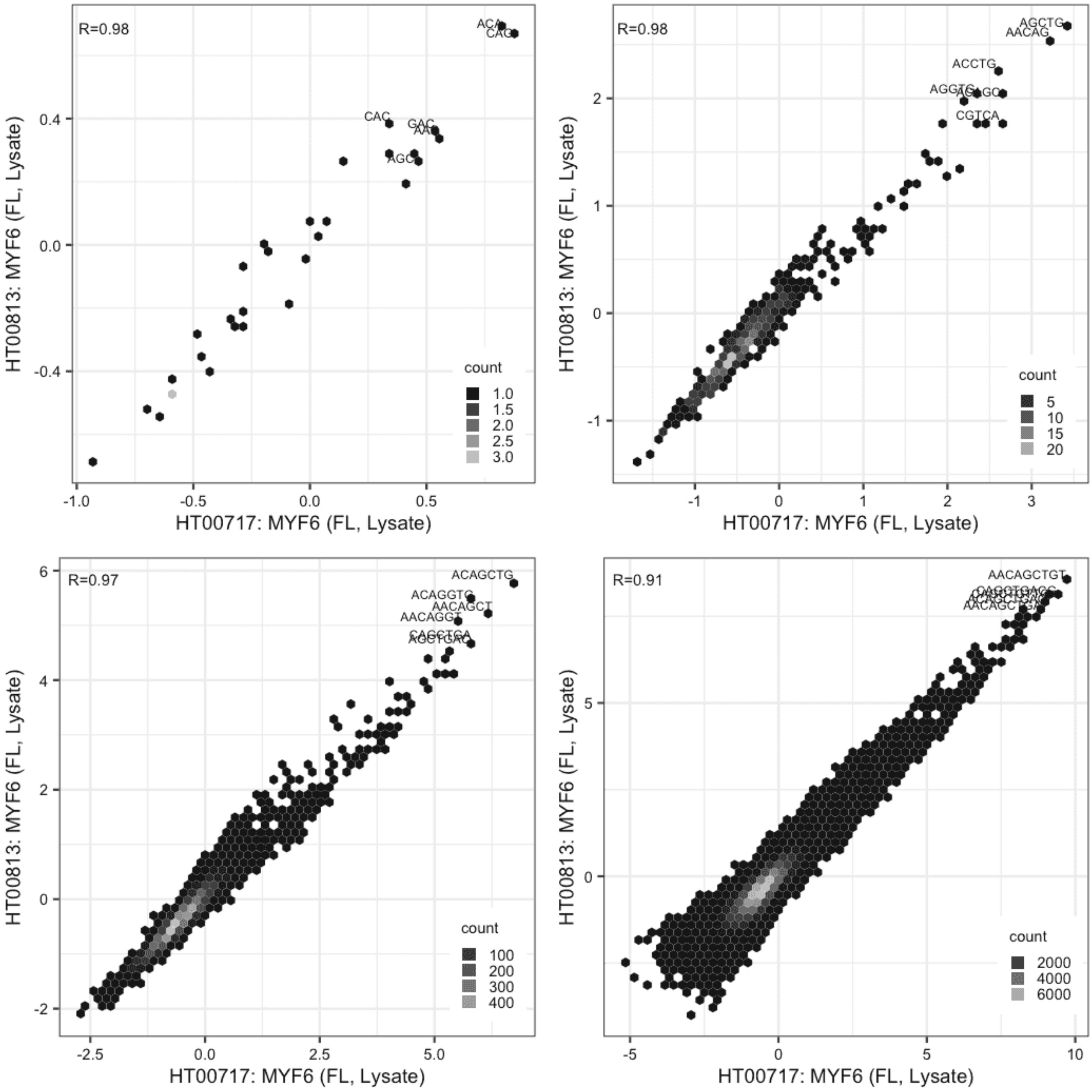

**MYPOP**

IVT vs IVT, different constructs

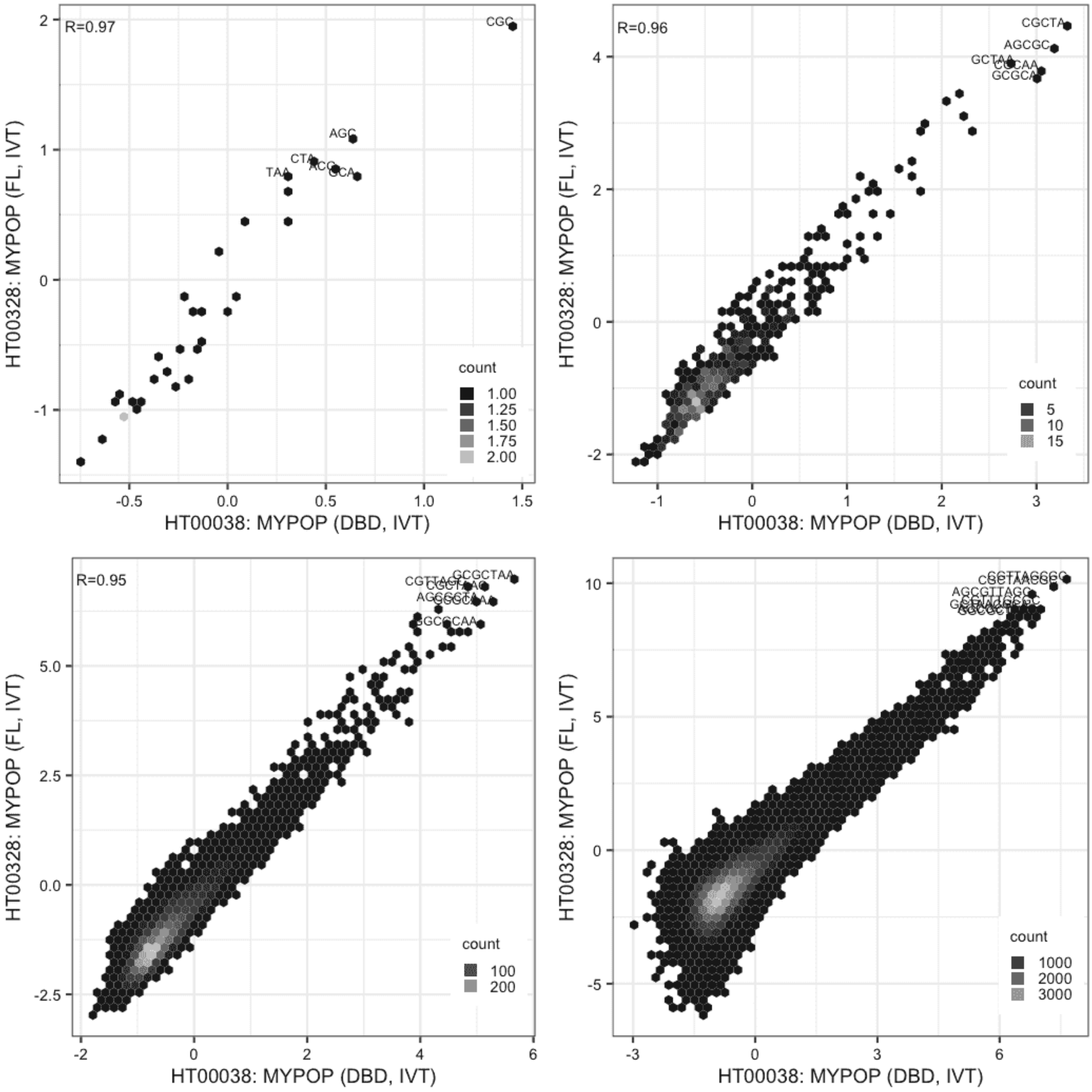

**MYPOP**

Lysate vs IVT, different constructs

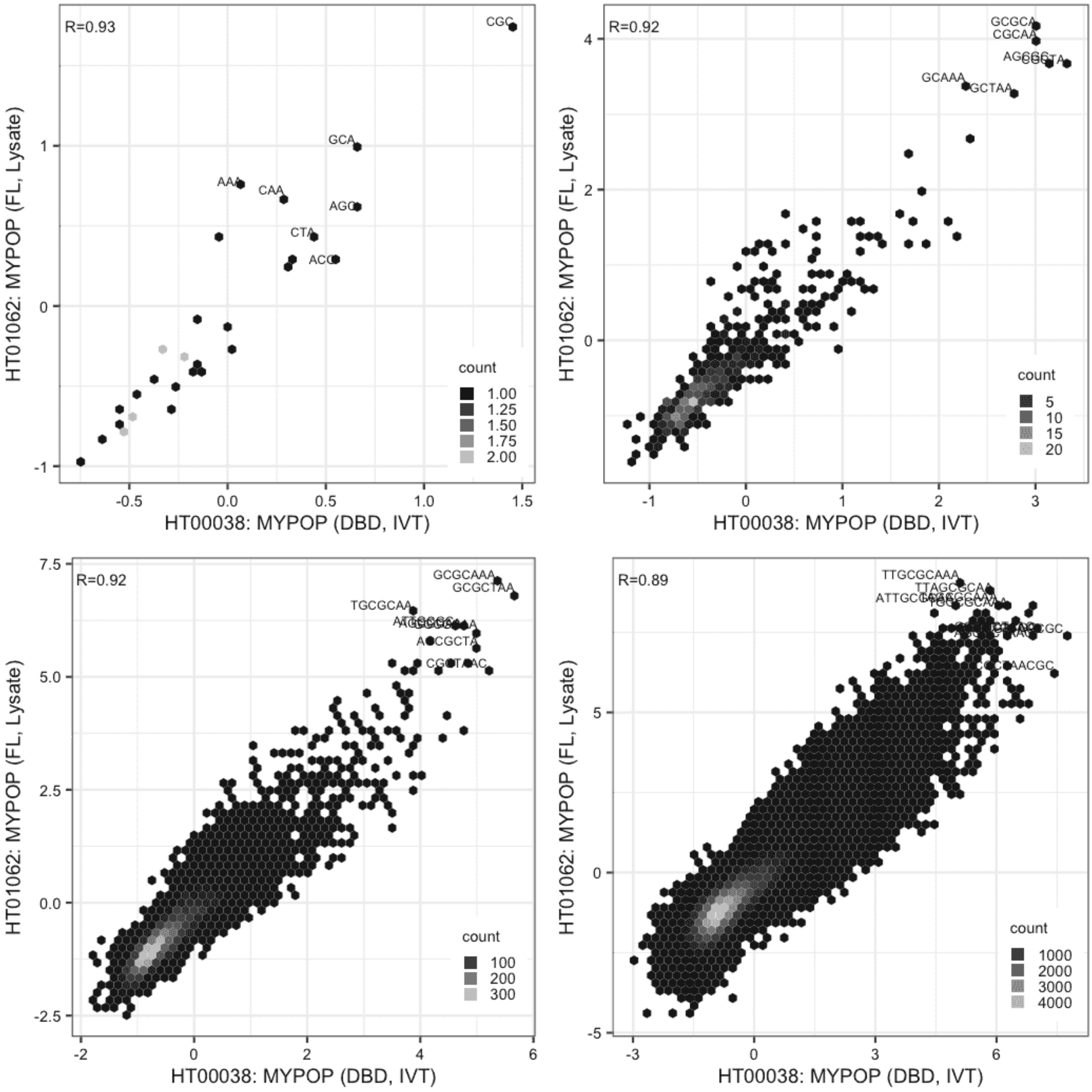

**MYPOP**

Lysate vs IVT, different constructs

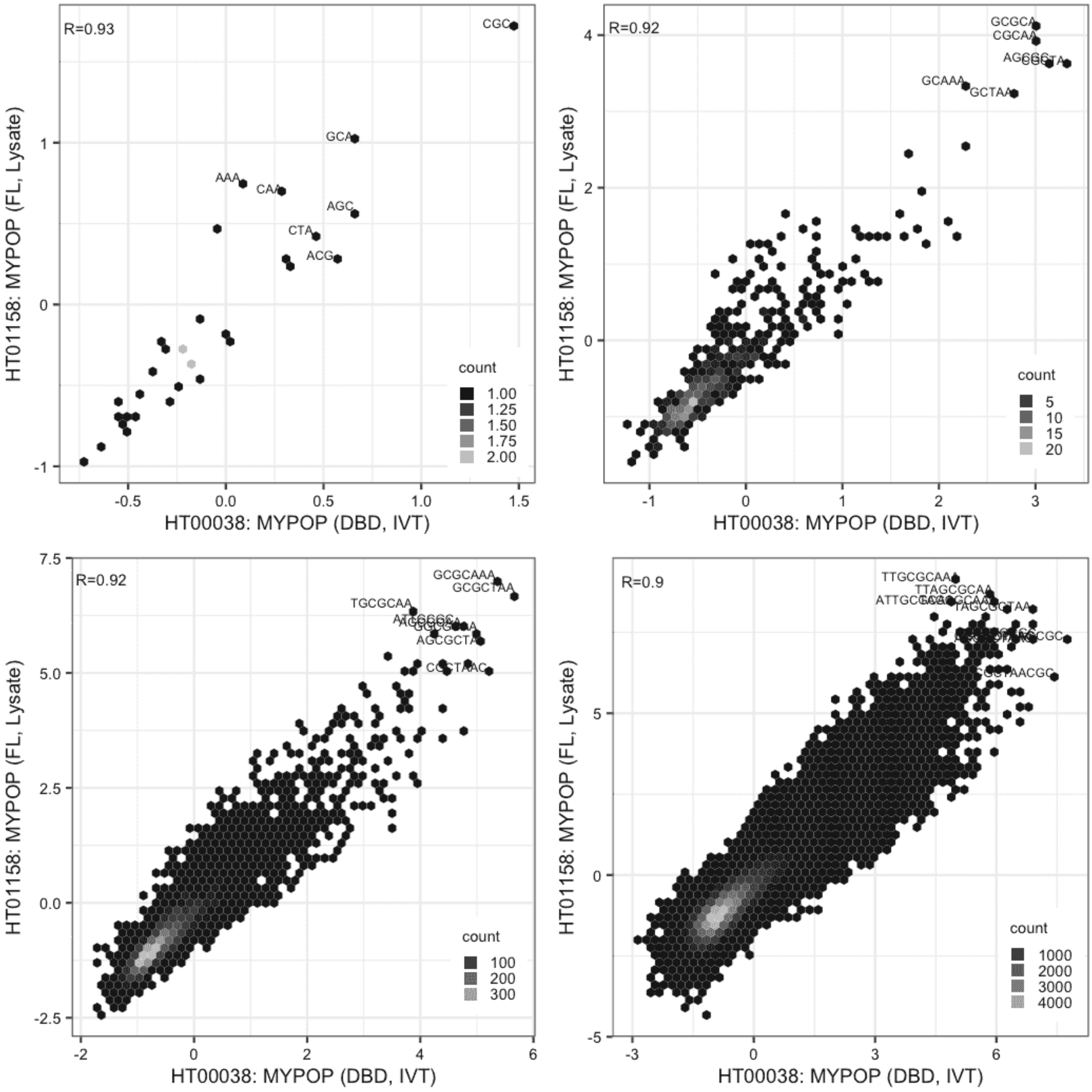

### MYPOP

IVT vs GFPIVT, different constructs

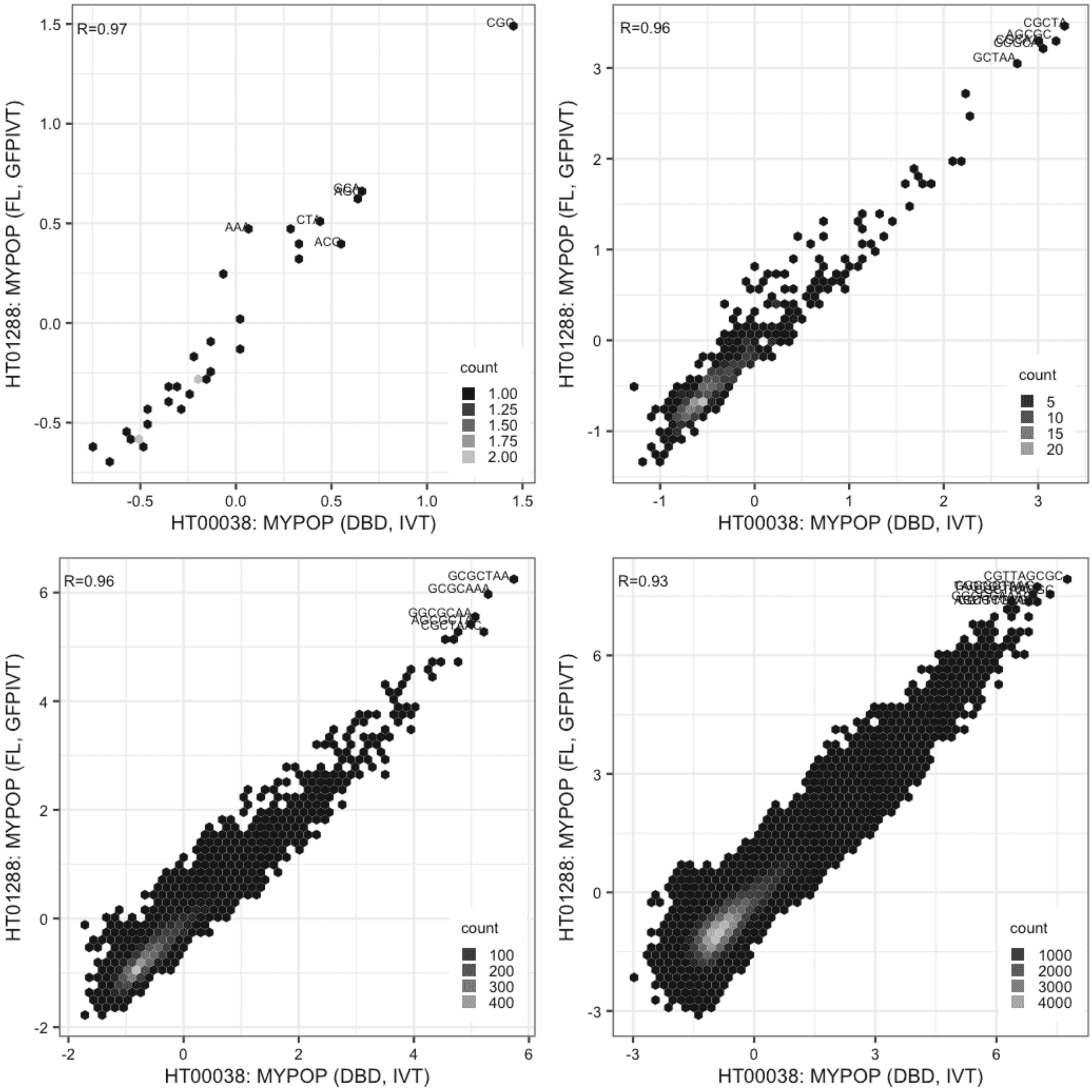

**MYPOP**

Lysate vs IVT, same construct

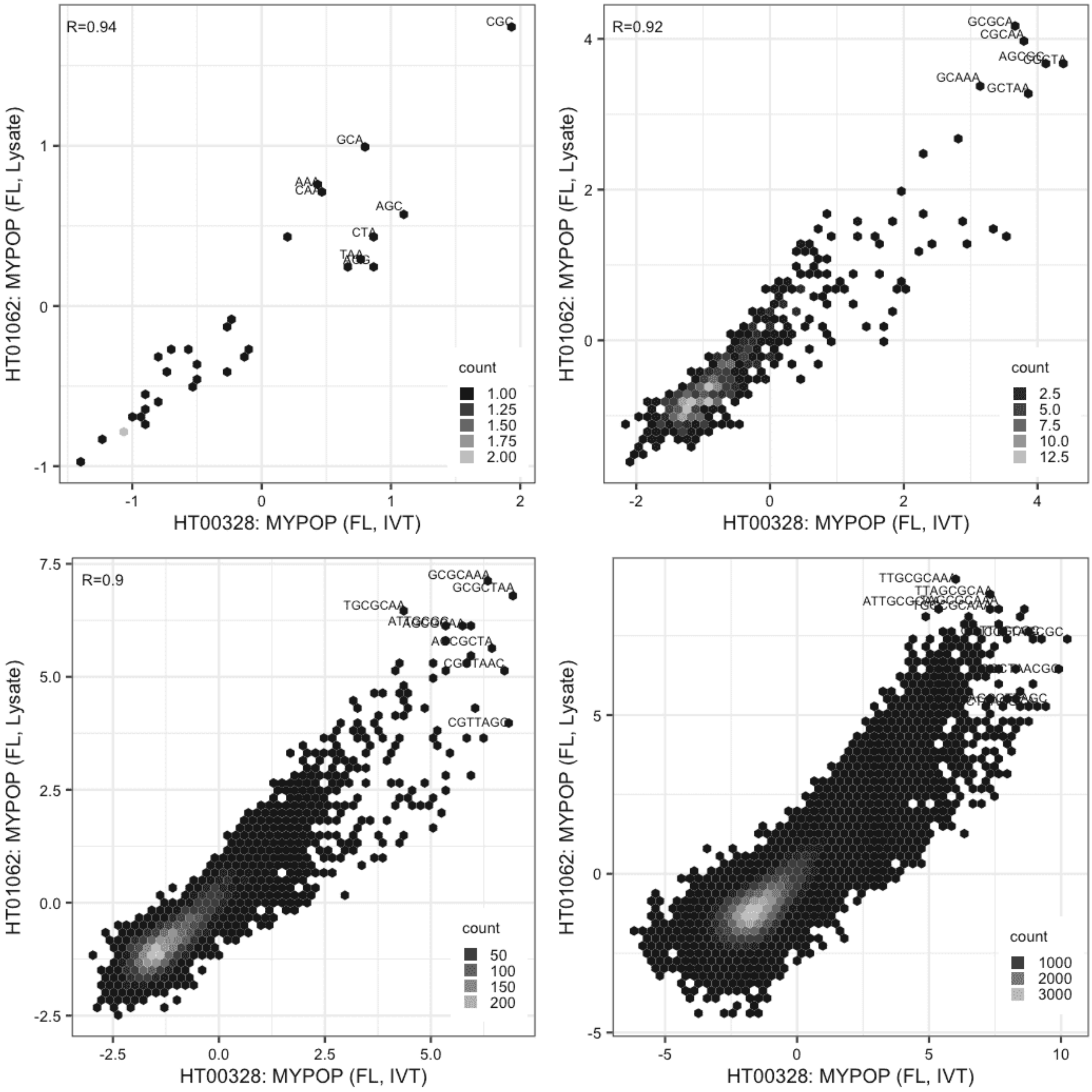

**MYPOP**

Lysate vs IVT, same construct

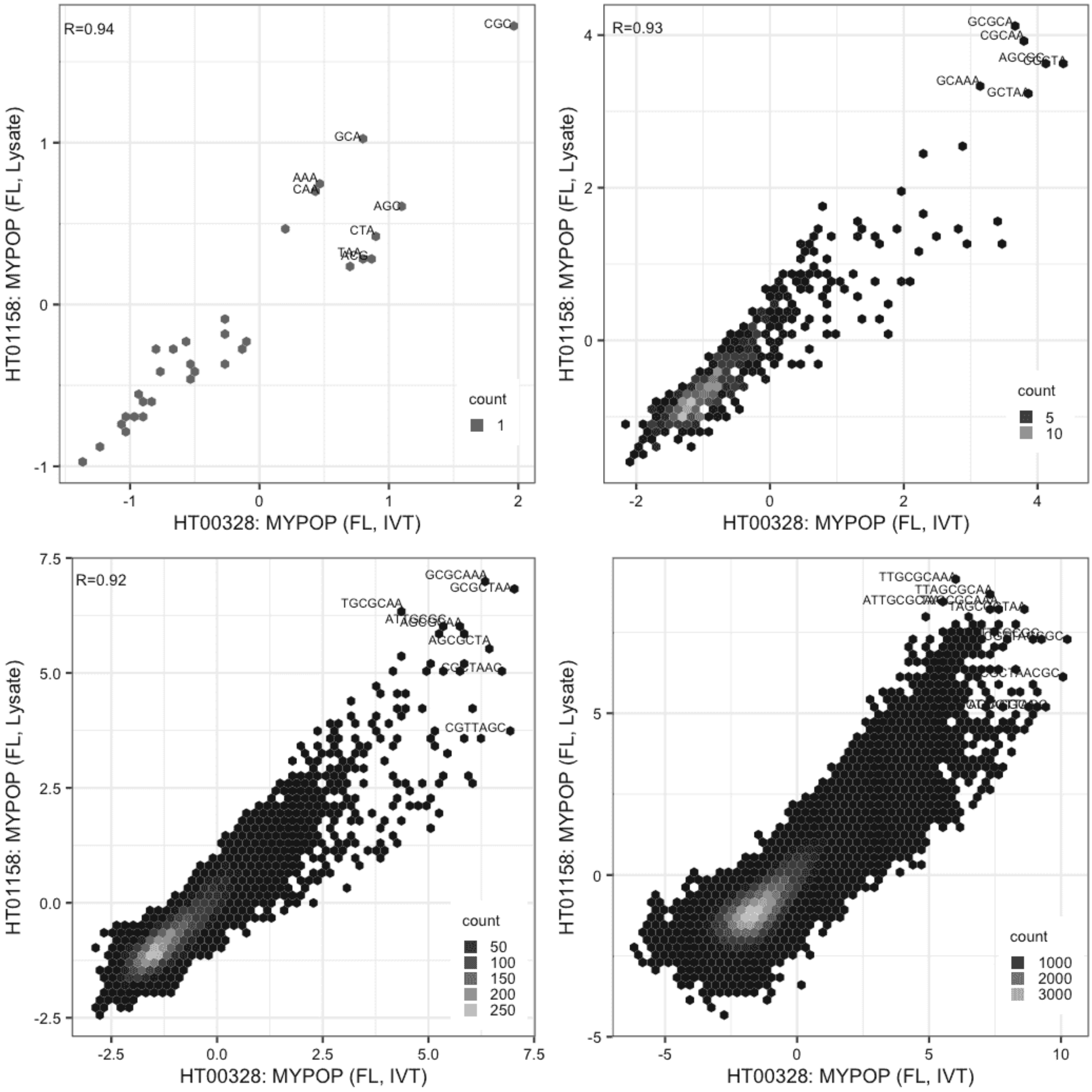

**MYPOP**

IVT vs GFPIVT, same construct

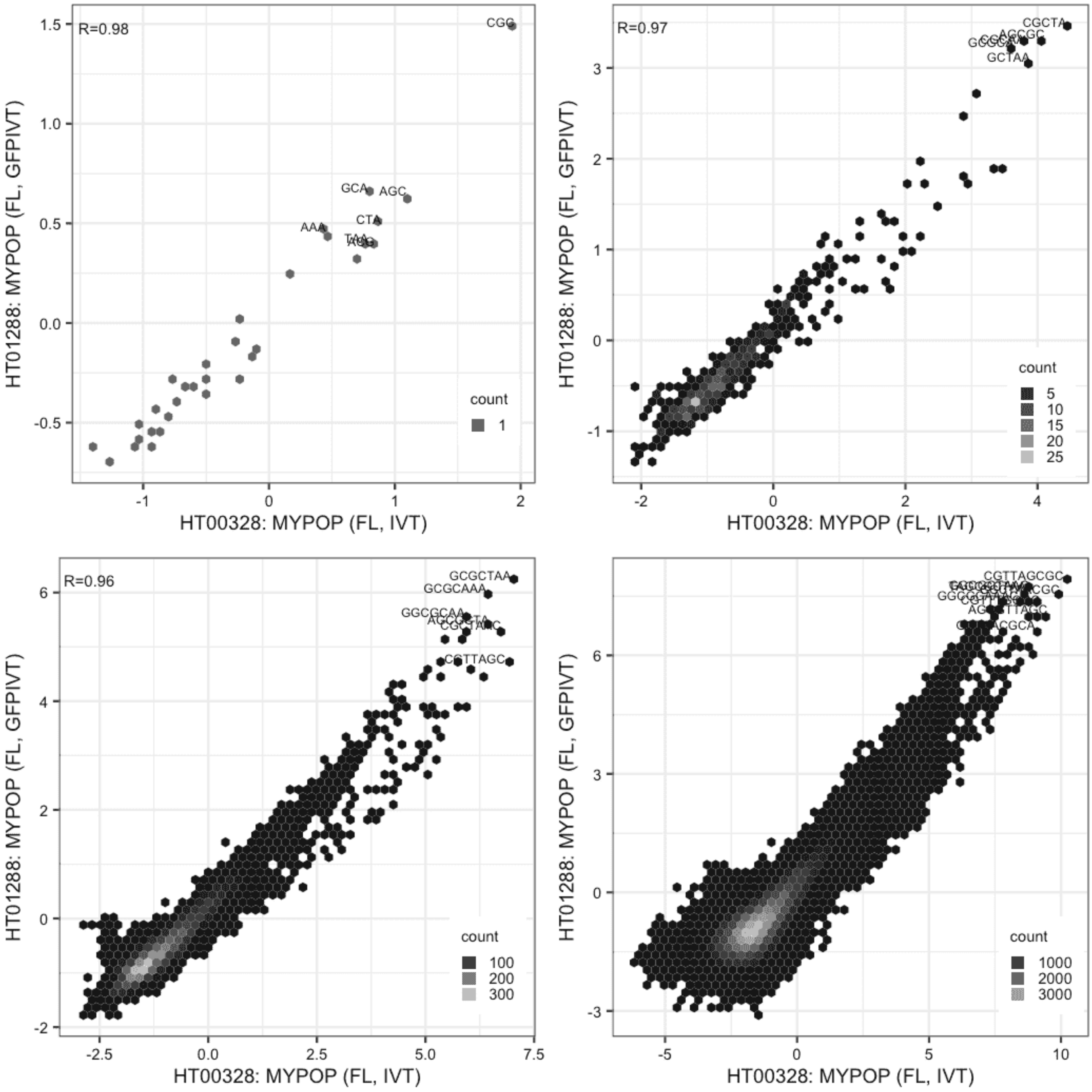

**MYPOP**

Lysate vs Lysate, same construct

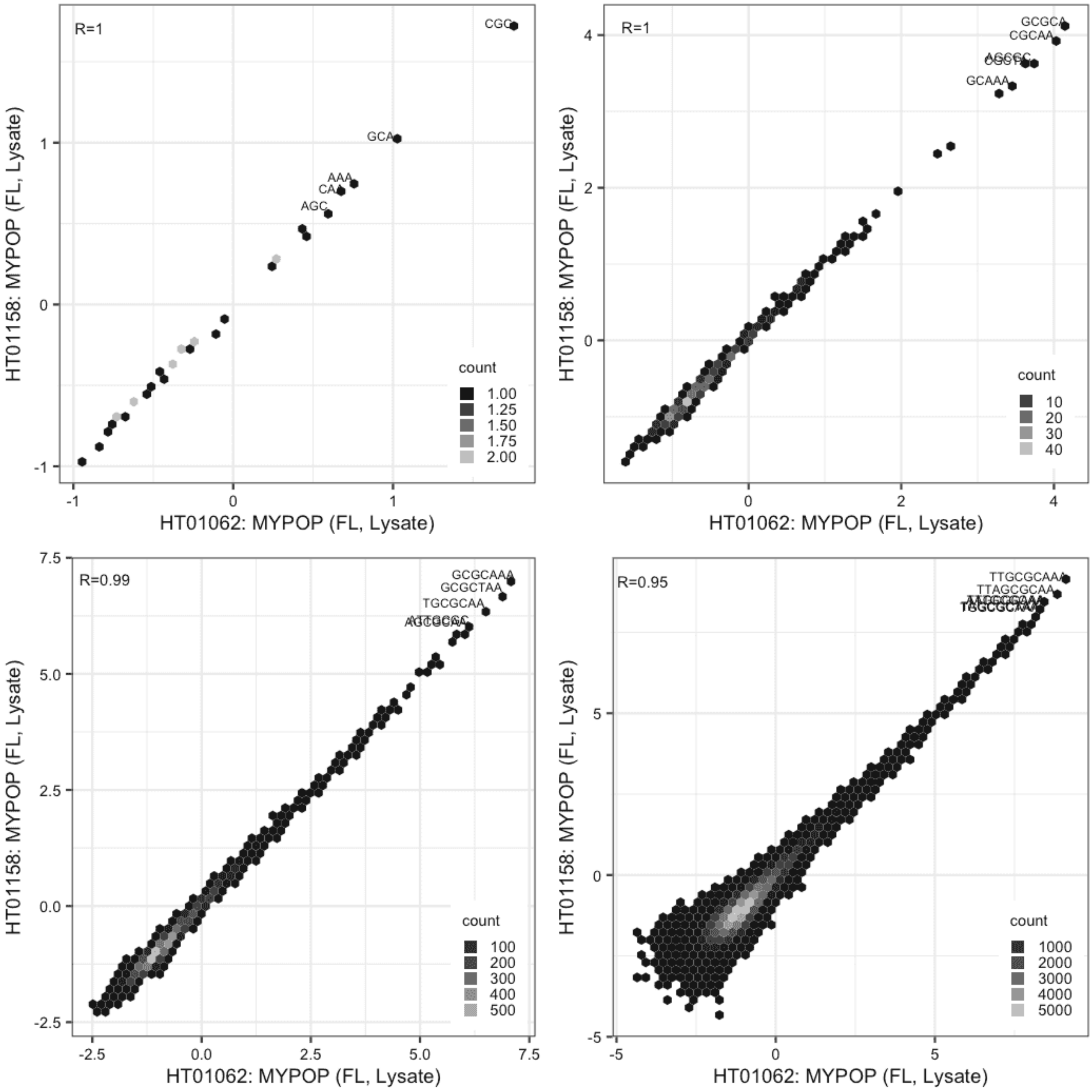

**MYPOP**

Lysate vs GFPIVT, same construct

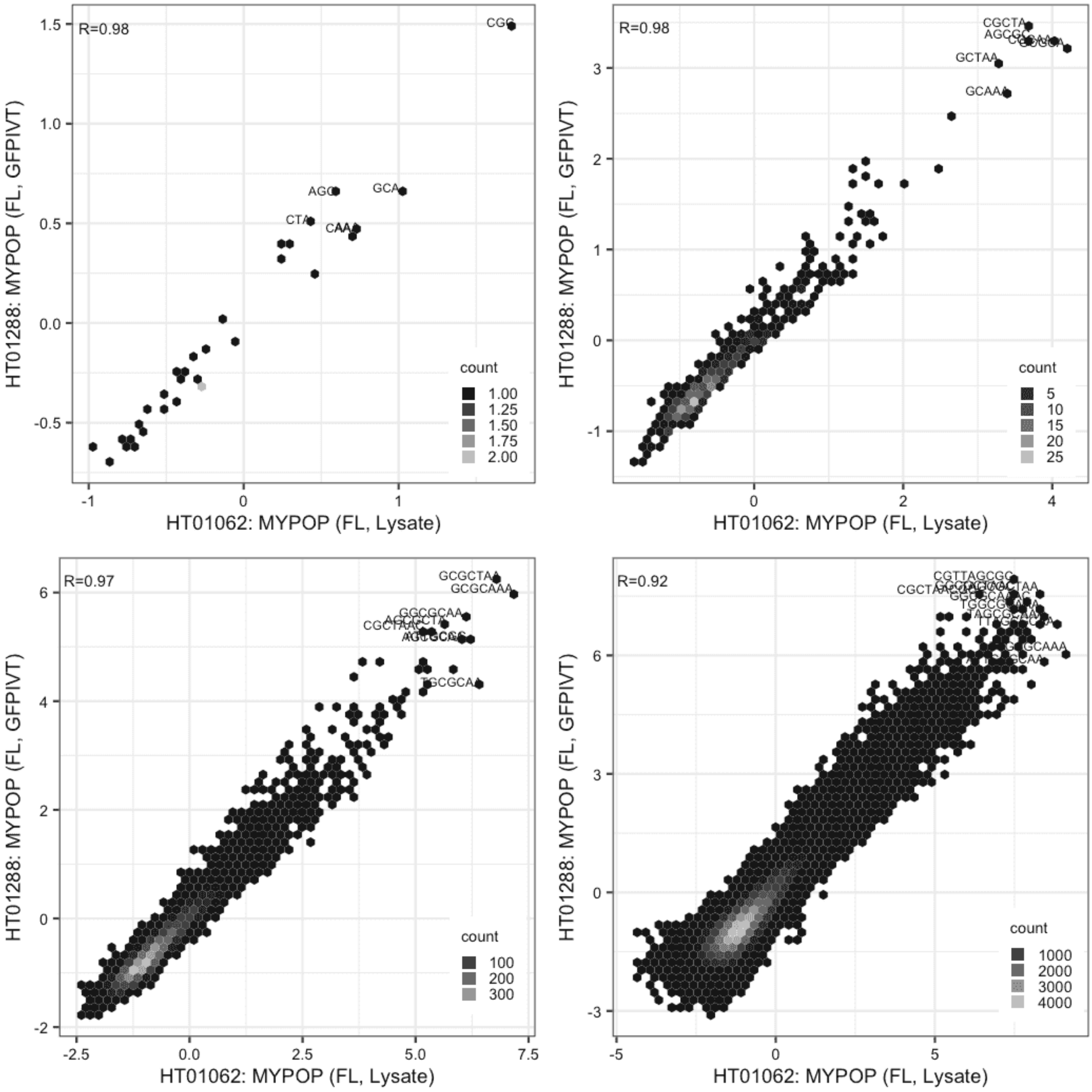

**MYPOP**

Lysate vs GFPIVT, same construct

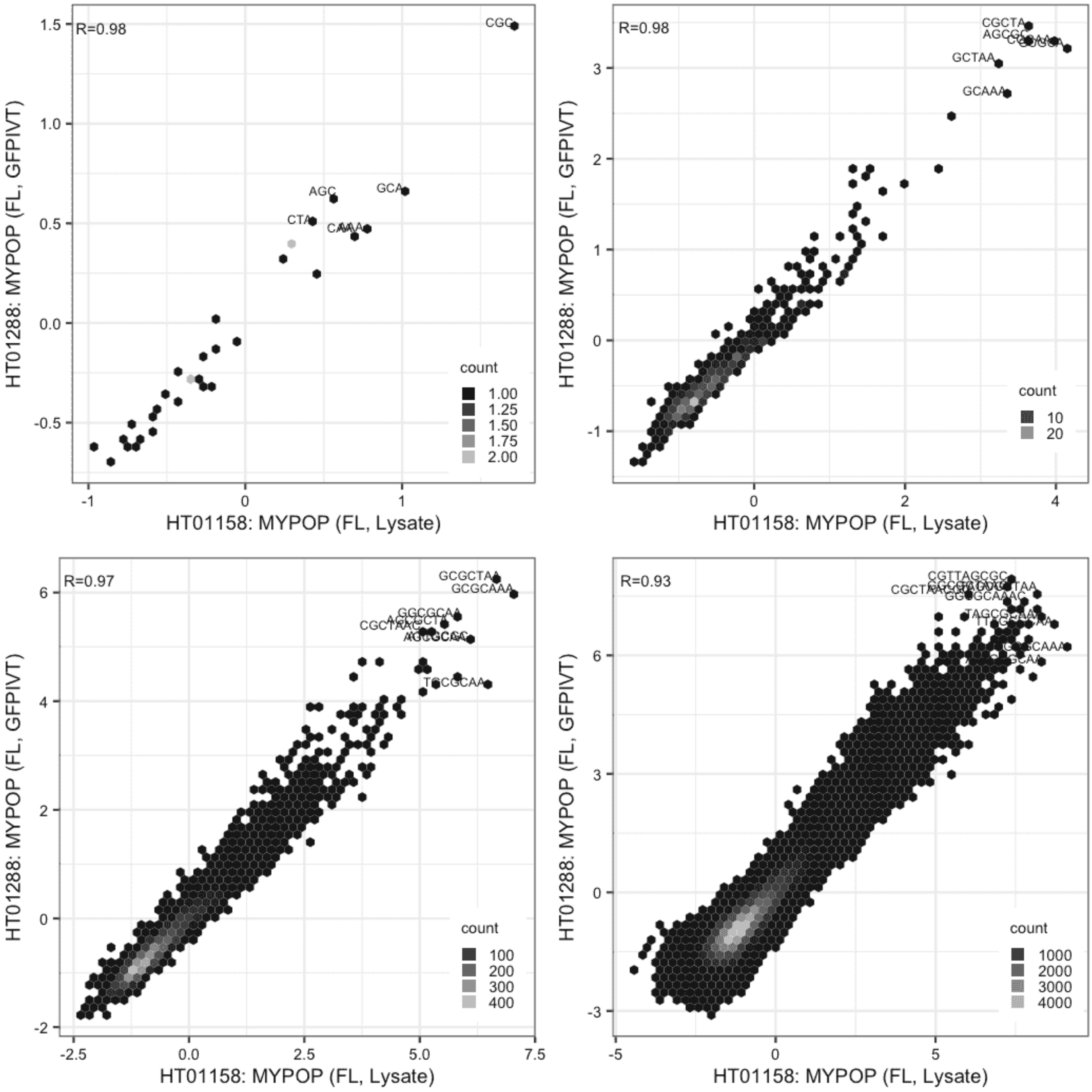

**NACC2**

Lysate vs IVT, different constructs

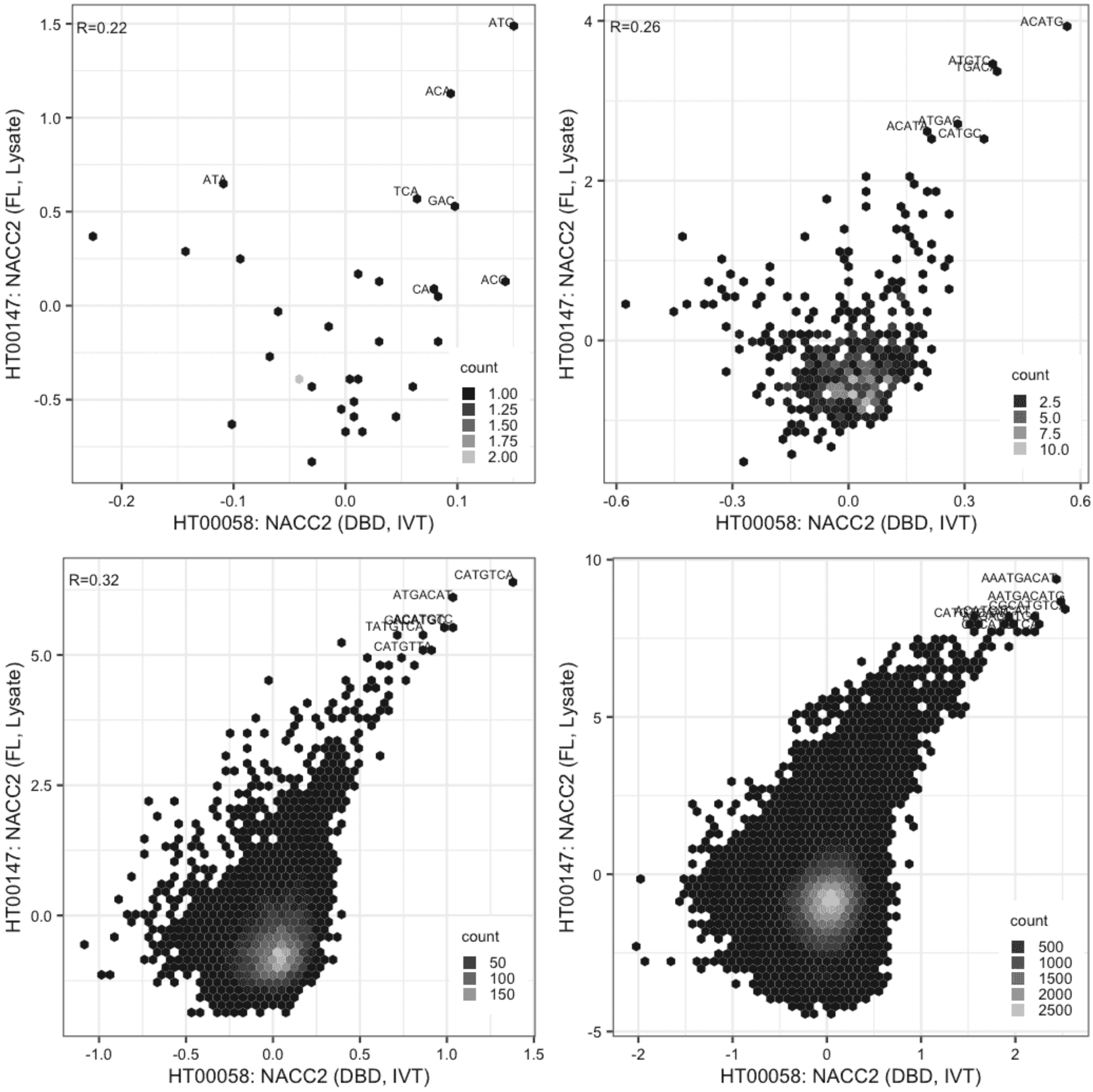

#### NACC2

IVT vs IVT, different constructs

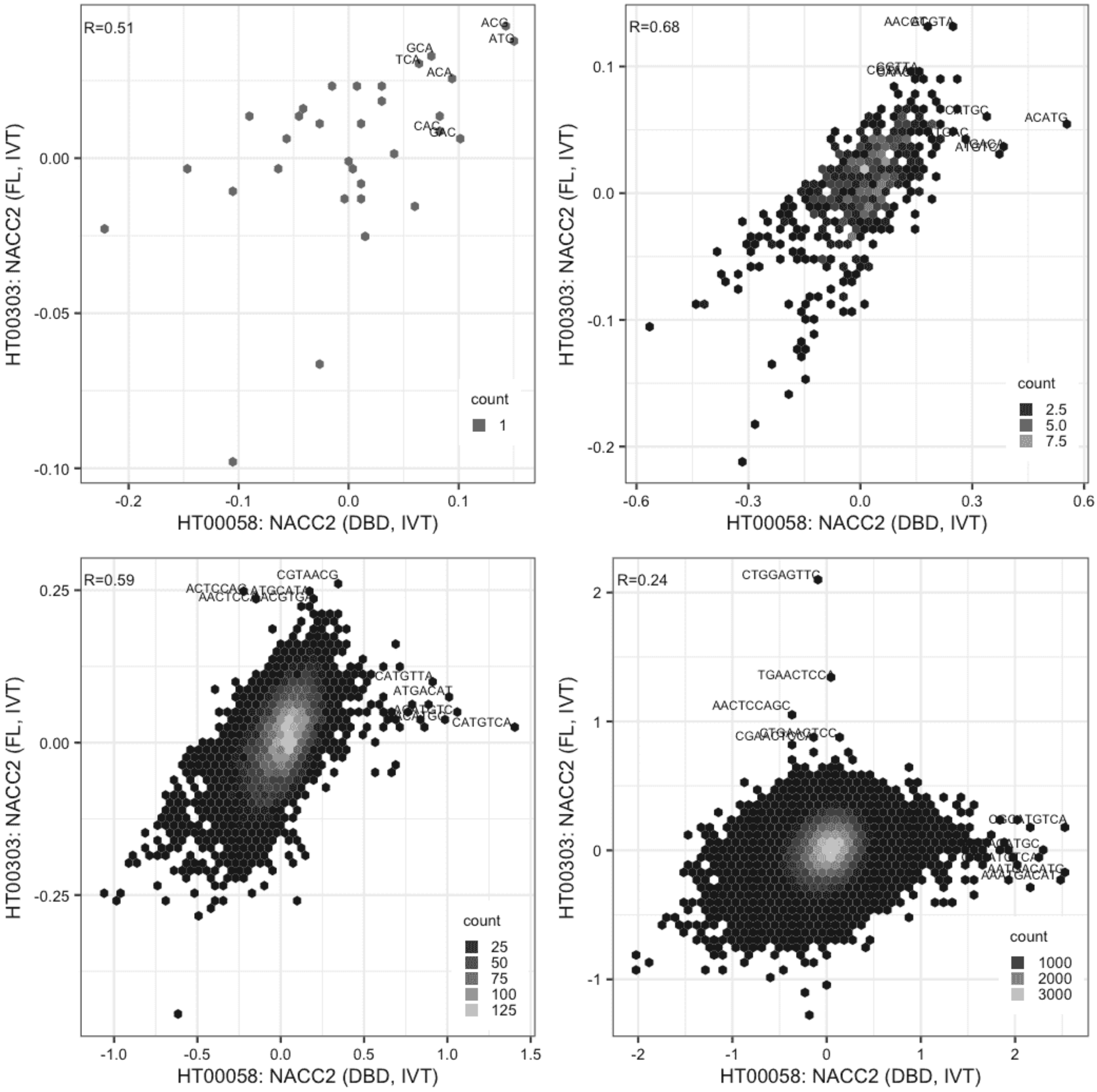

**NACC2**

IVT vs GFPIVT, same construct

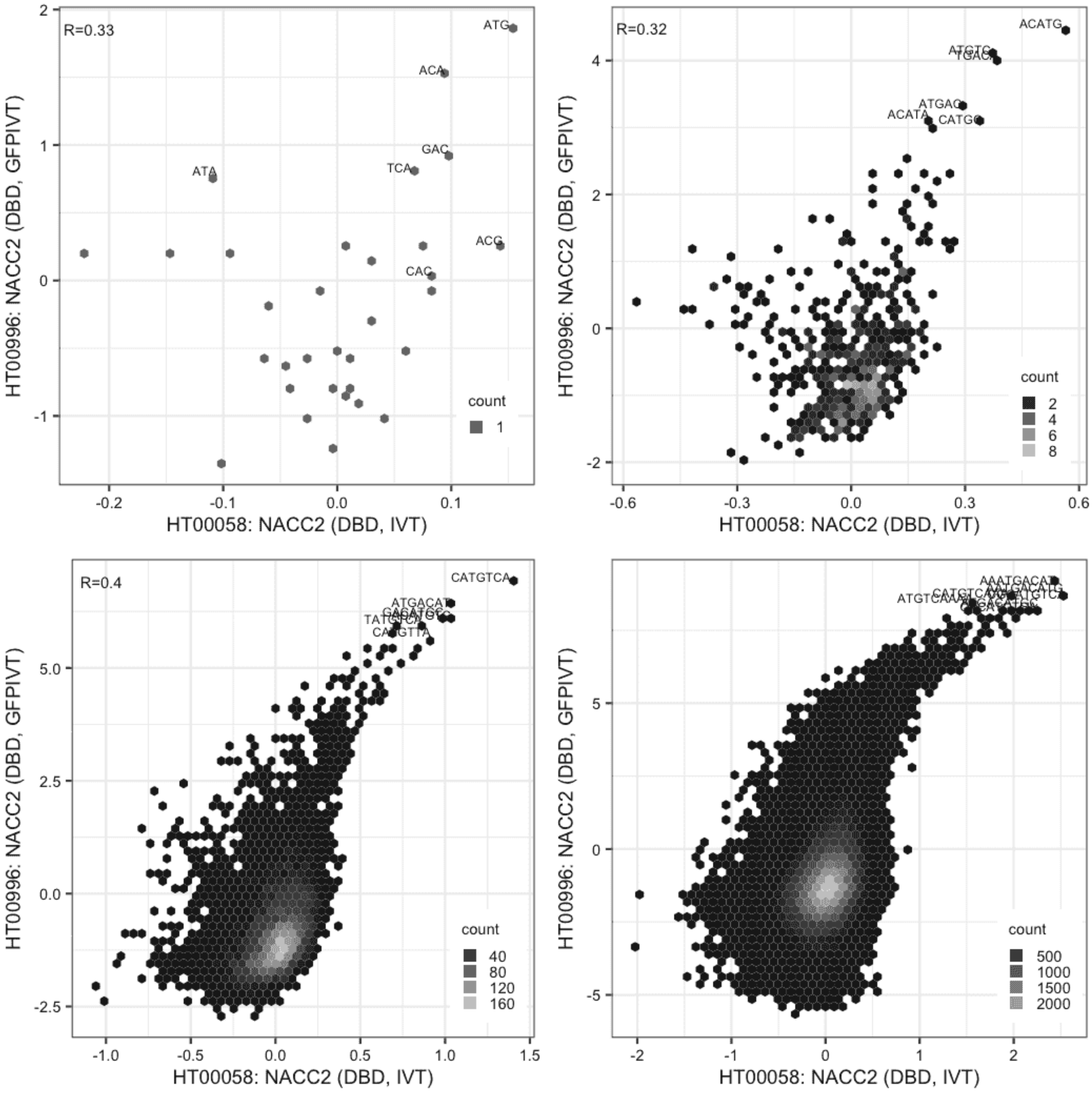

#### NACC2

Lysate vs IVT, same construct

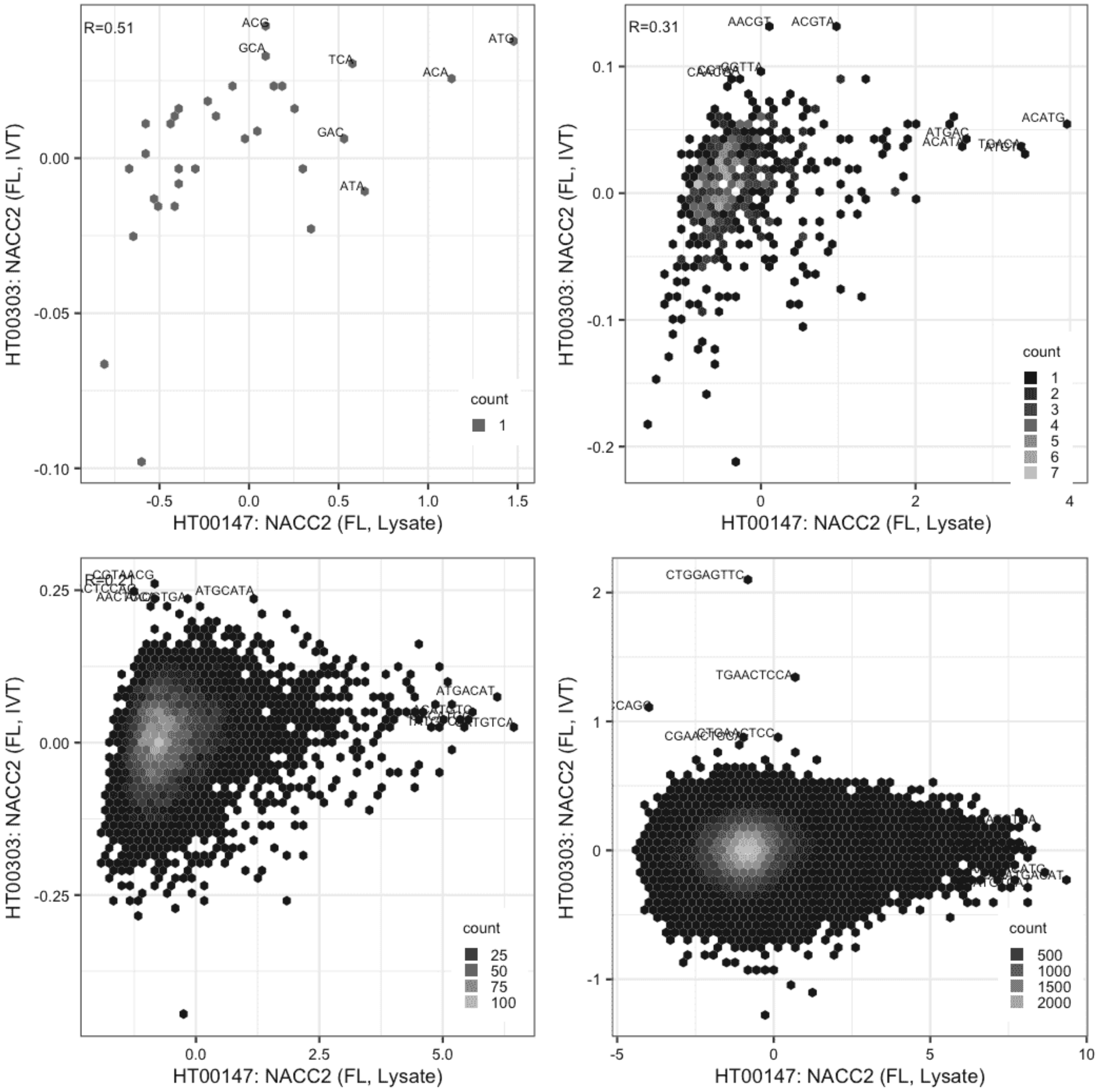

#### NACC2

Lysate vs GFPIVT, different constructs

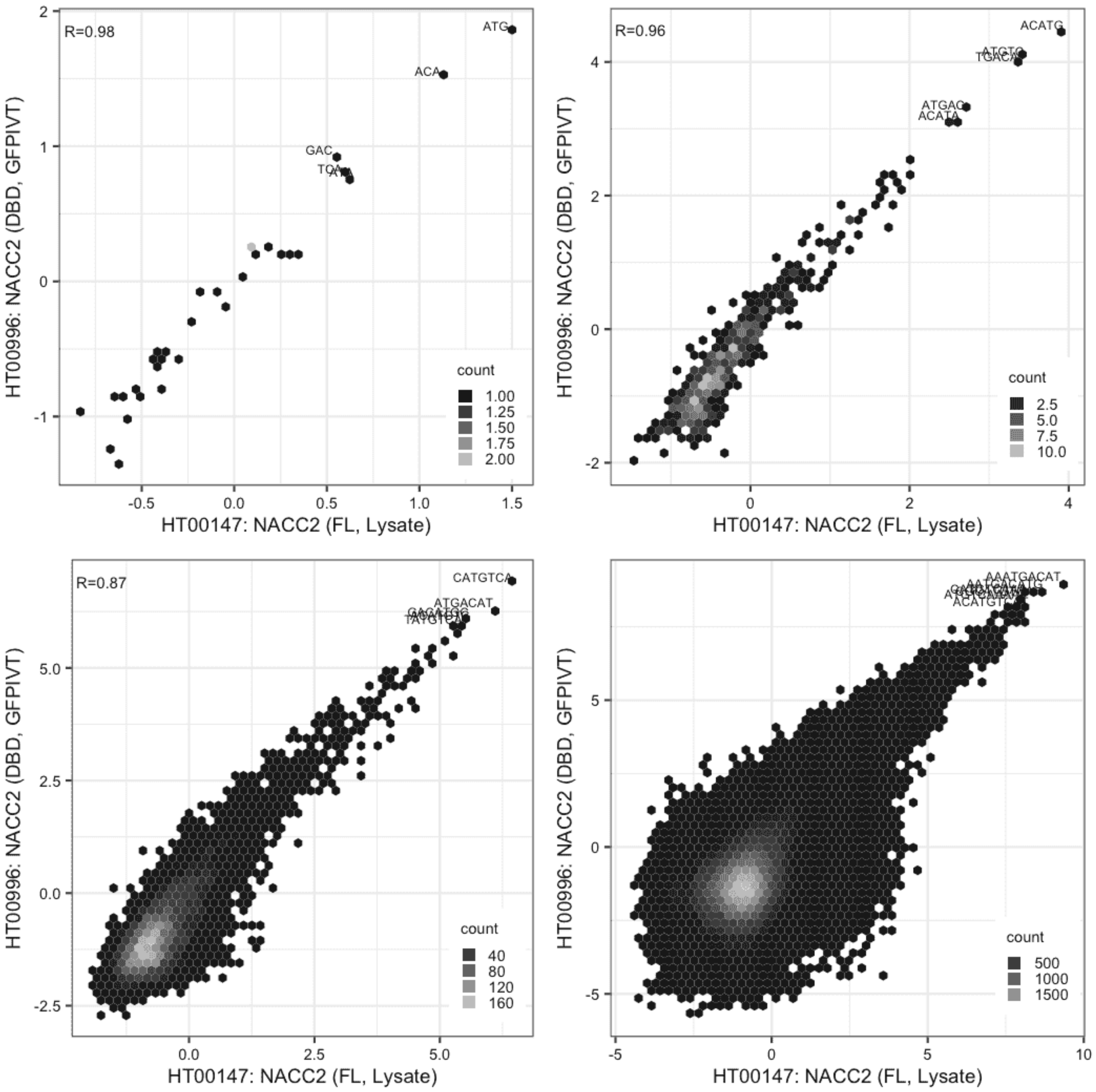

### NACC2

IVT vs GFPIVT, different constructs

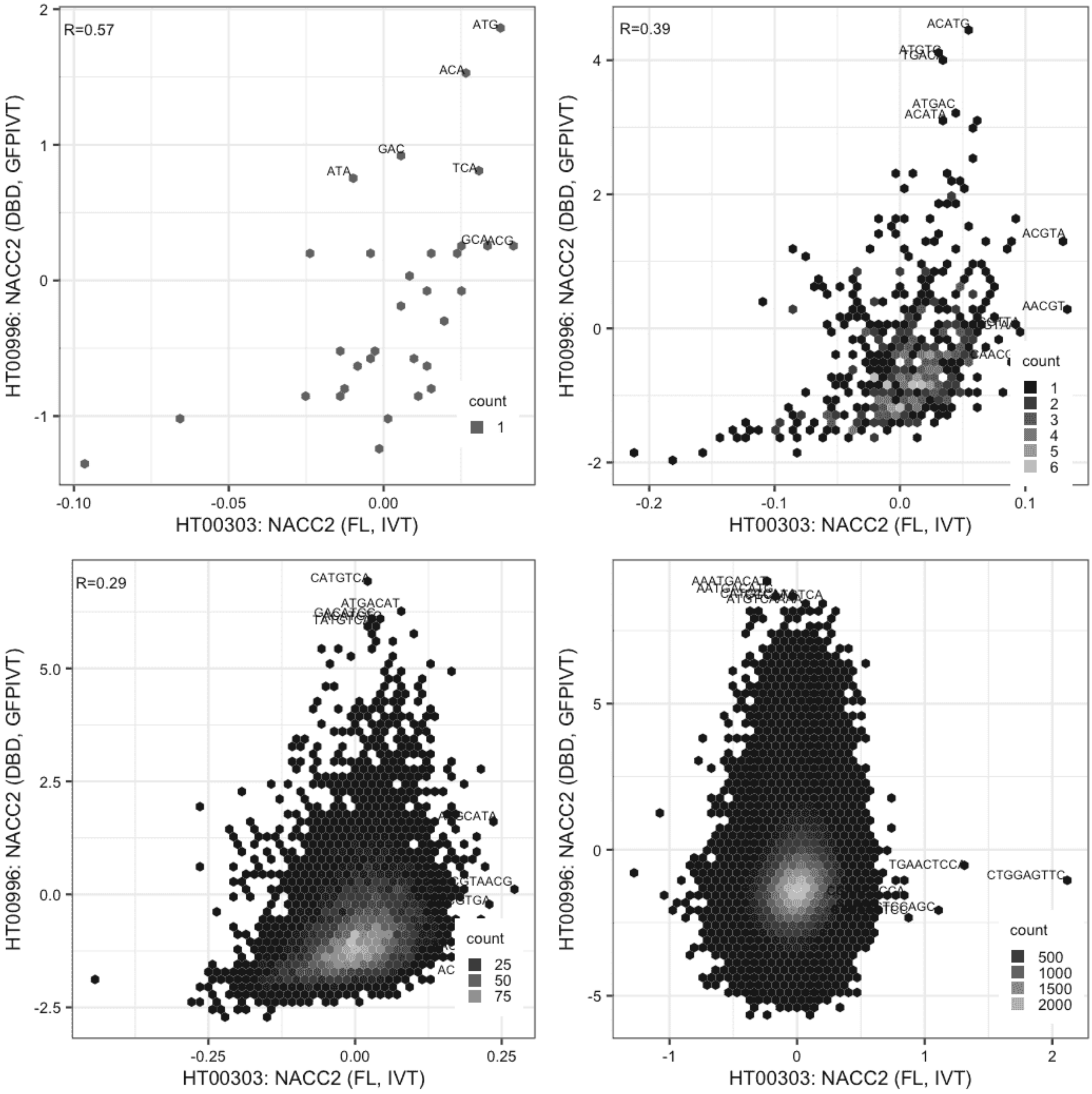

**NFKB1**

Lysate vs IVT, same construct

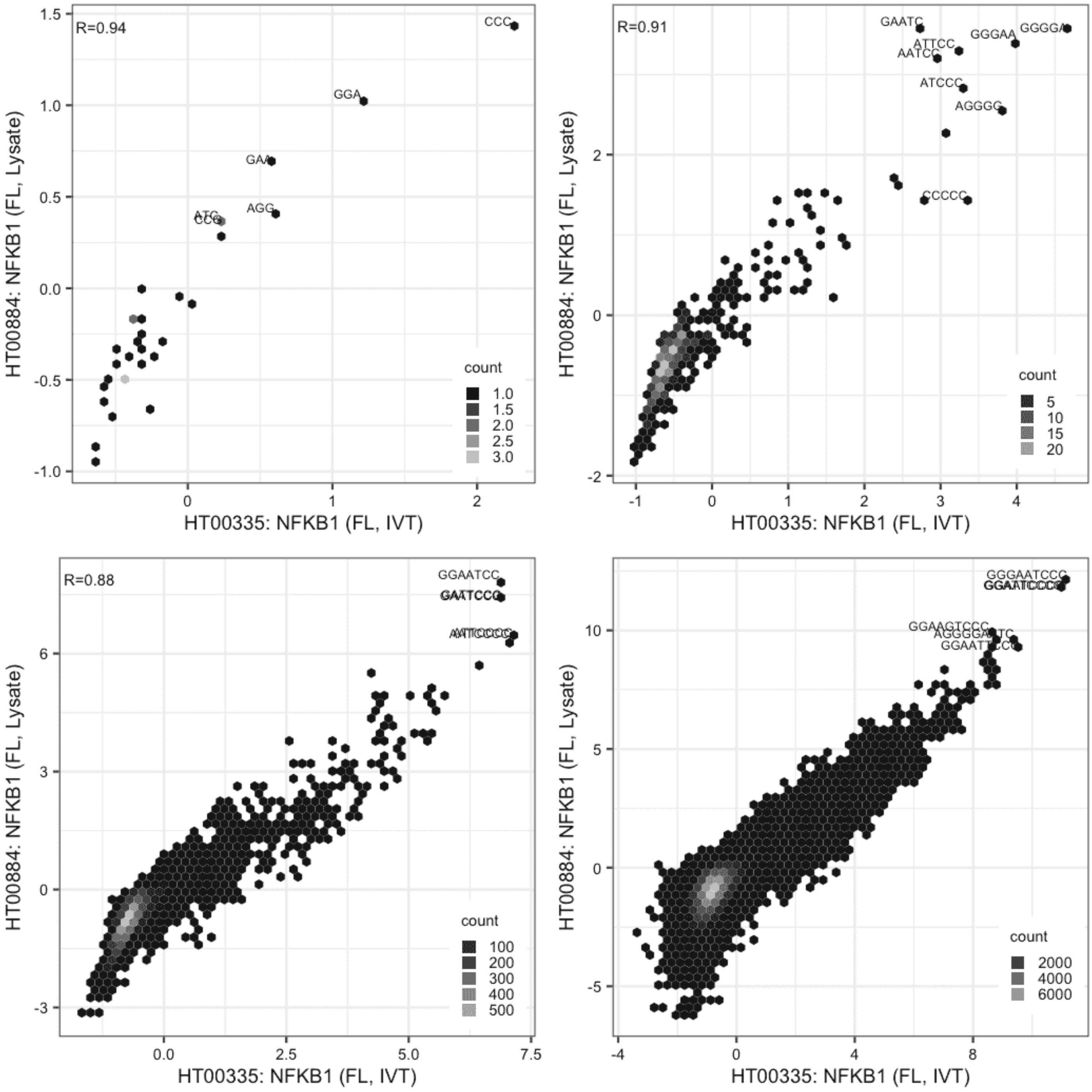

**NR1H4**

Lysate vs IVT, same construct

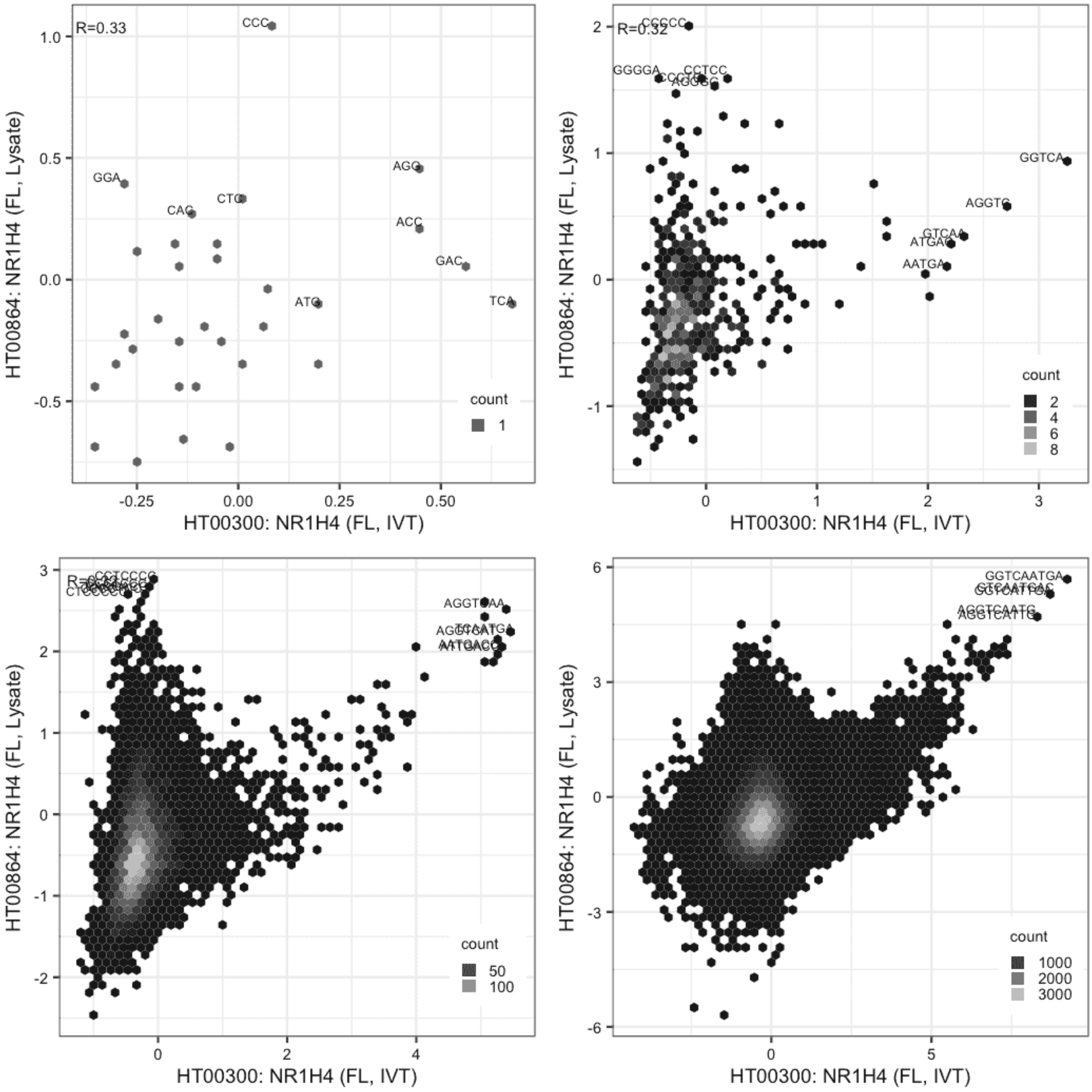

**PAX7**

Lysate vs IVT, same construct

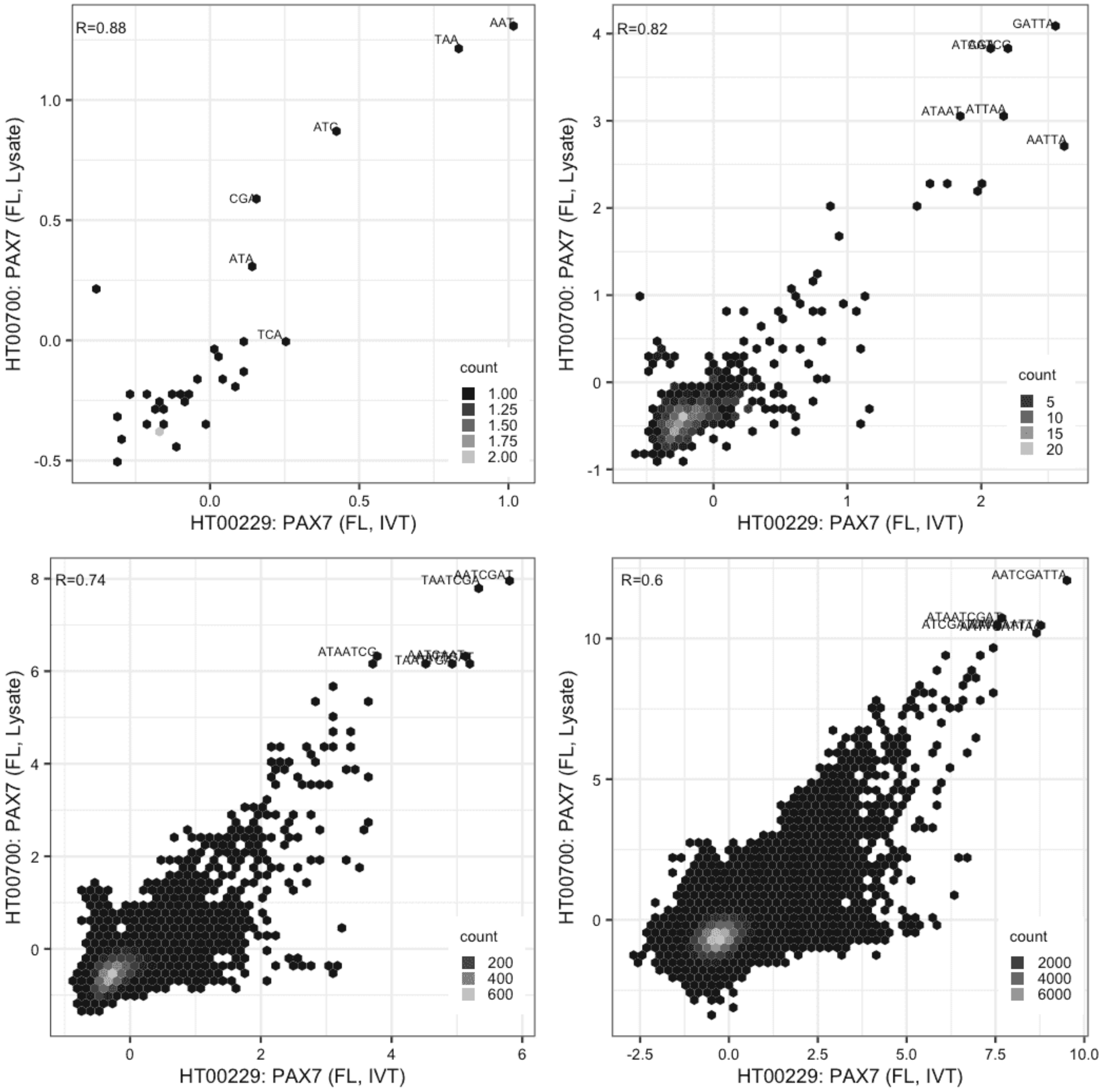

**PAX7**

Lysate vs IVT, same construct

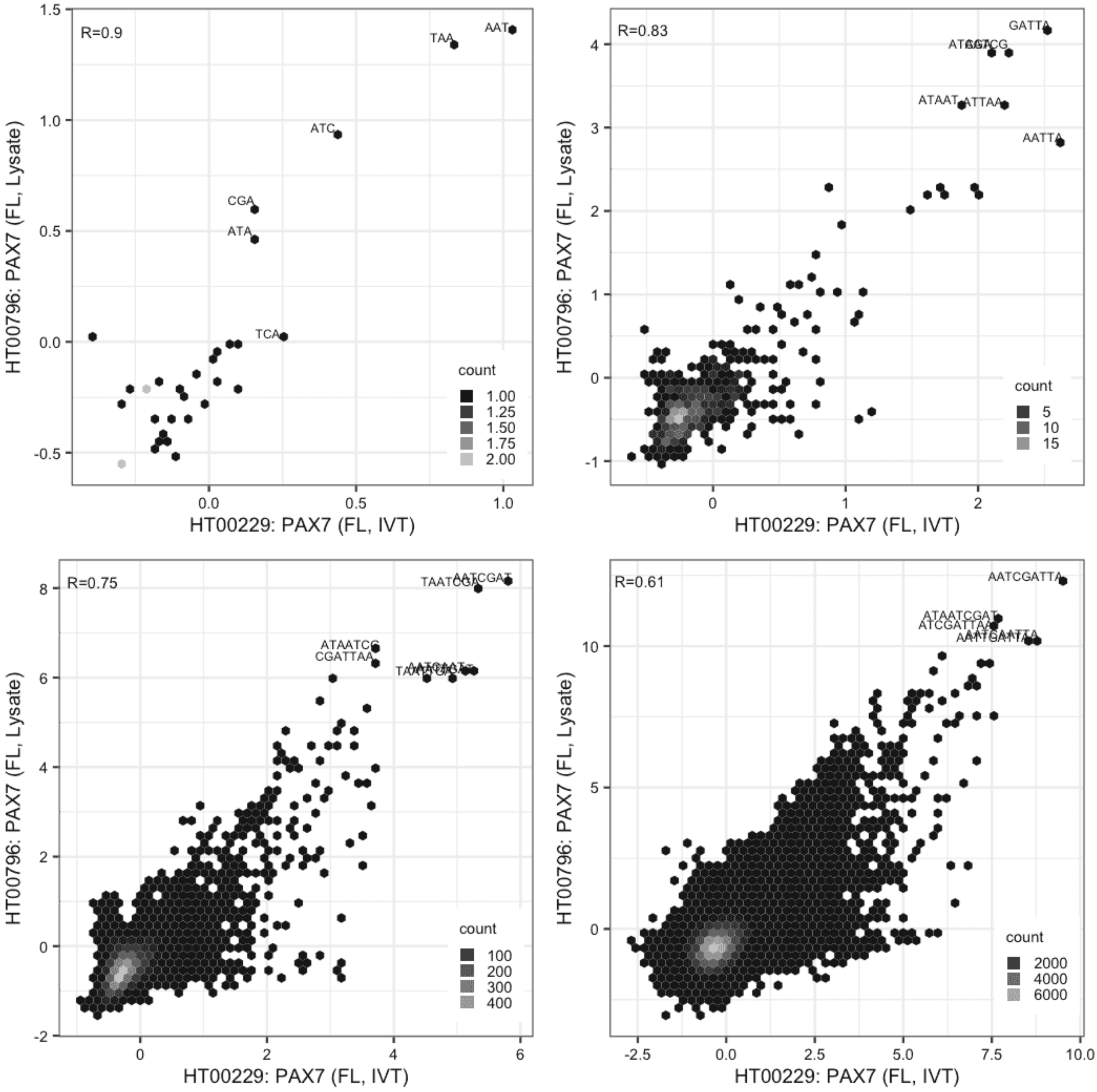

**PAX7**

Lysate vs Lysate, same construct

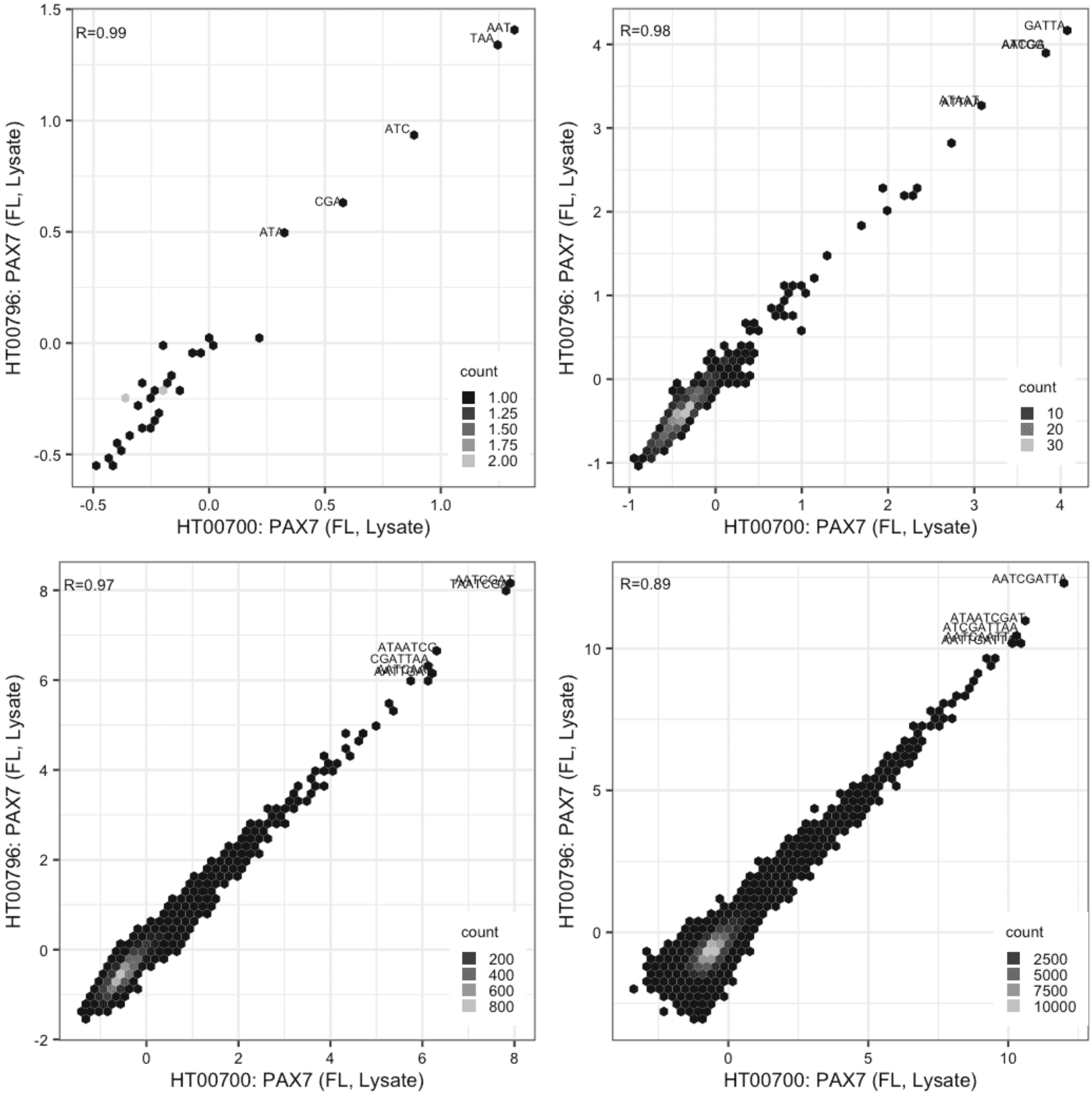

**POGK**

Lysate vs IVT, different constructs

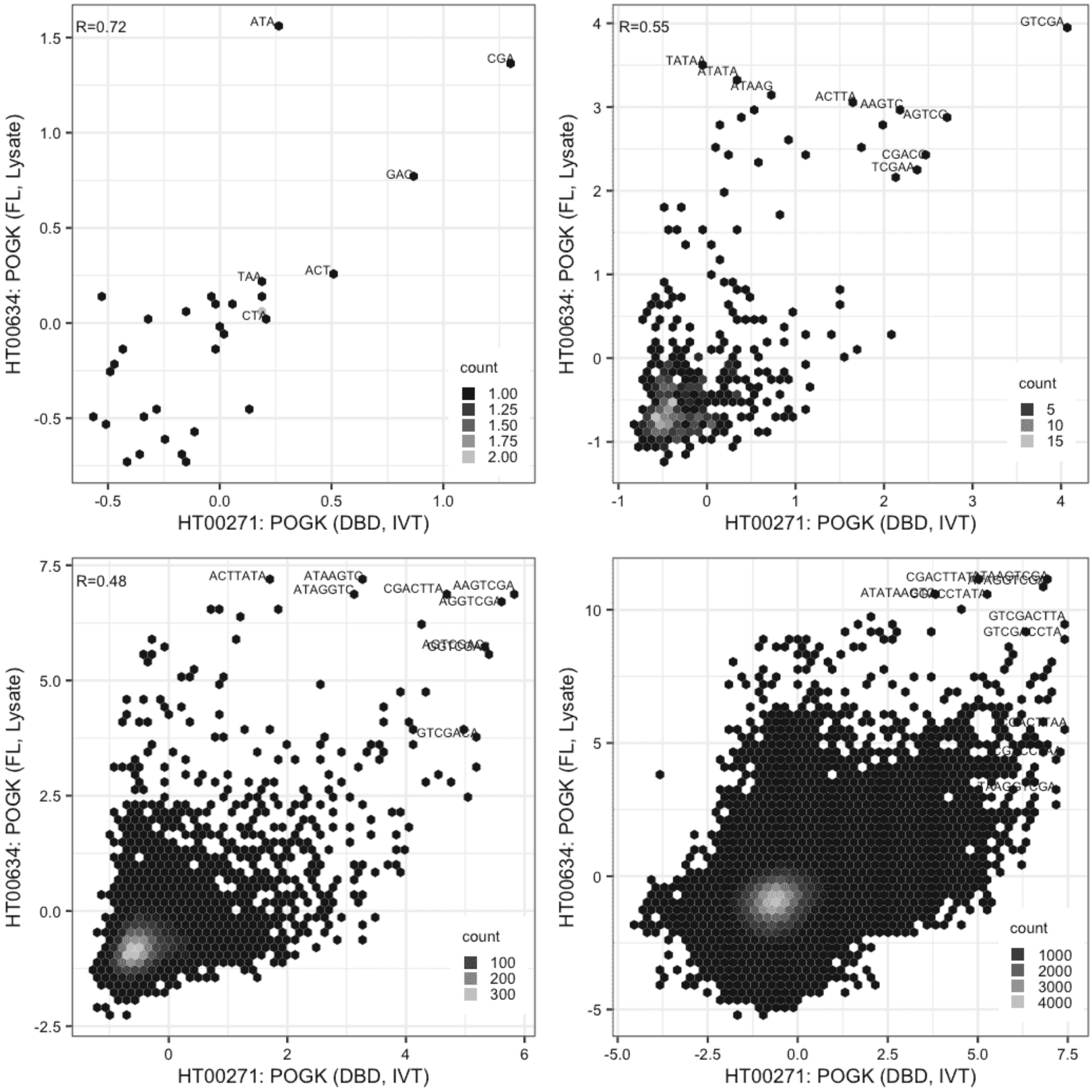

#### PRDM10

Lysate vs Lysate, same construct

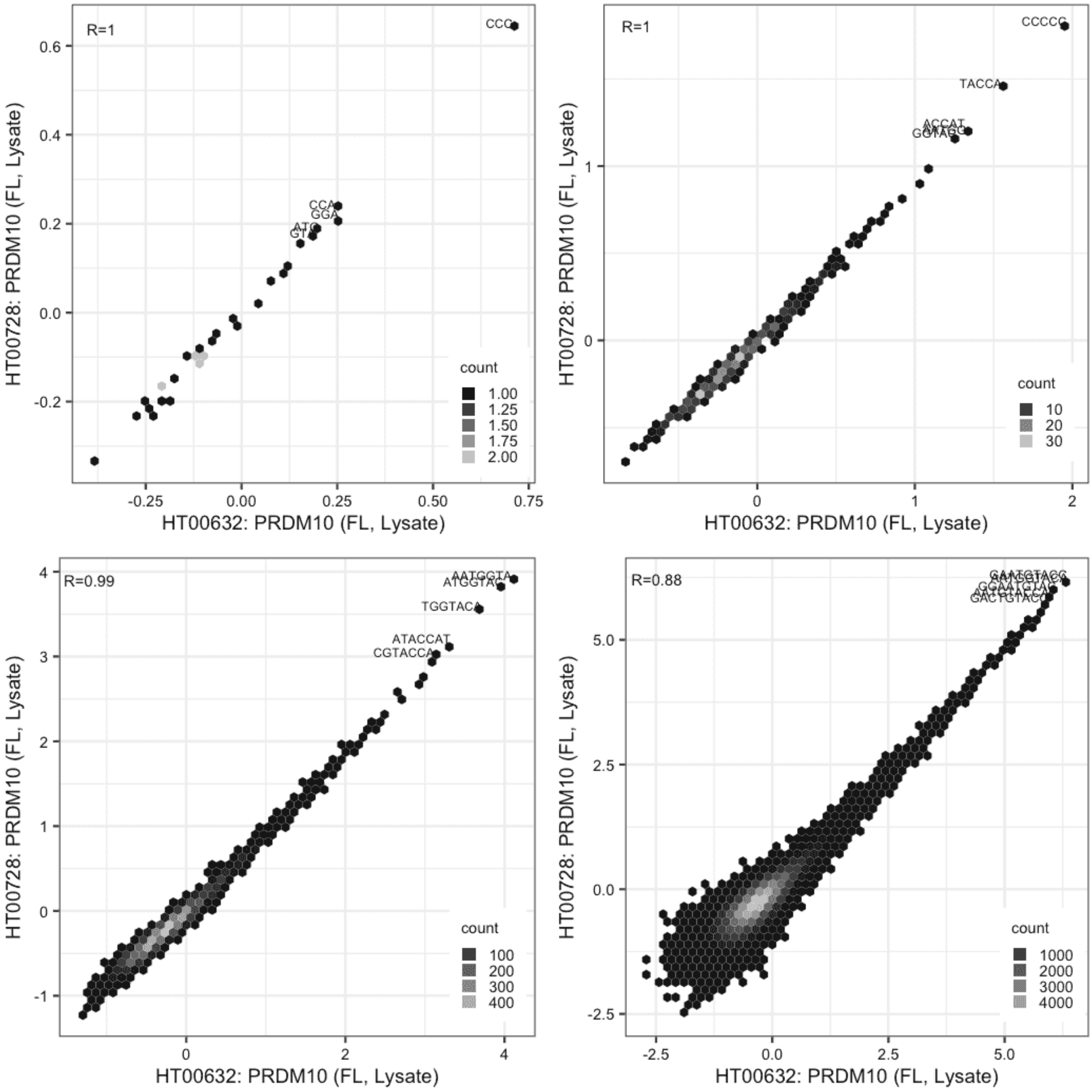

**PRDM10**

Lysate vs GFPIVT, different constructs

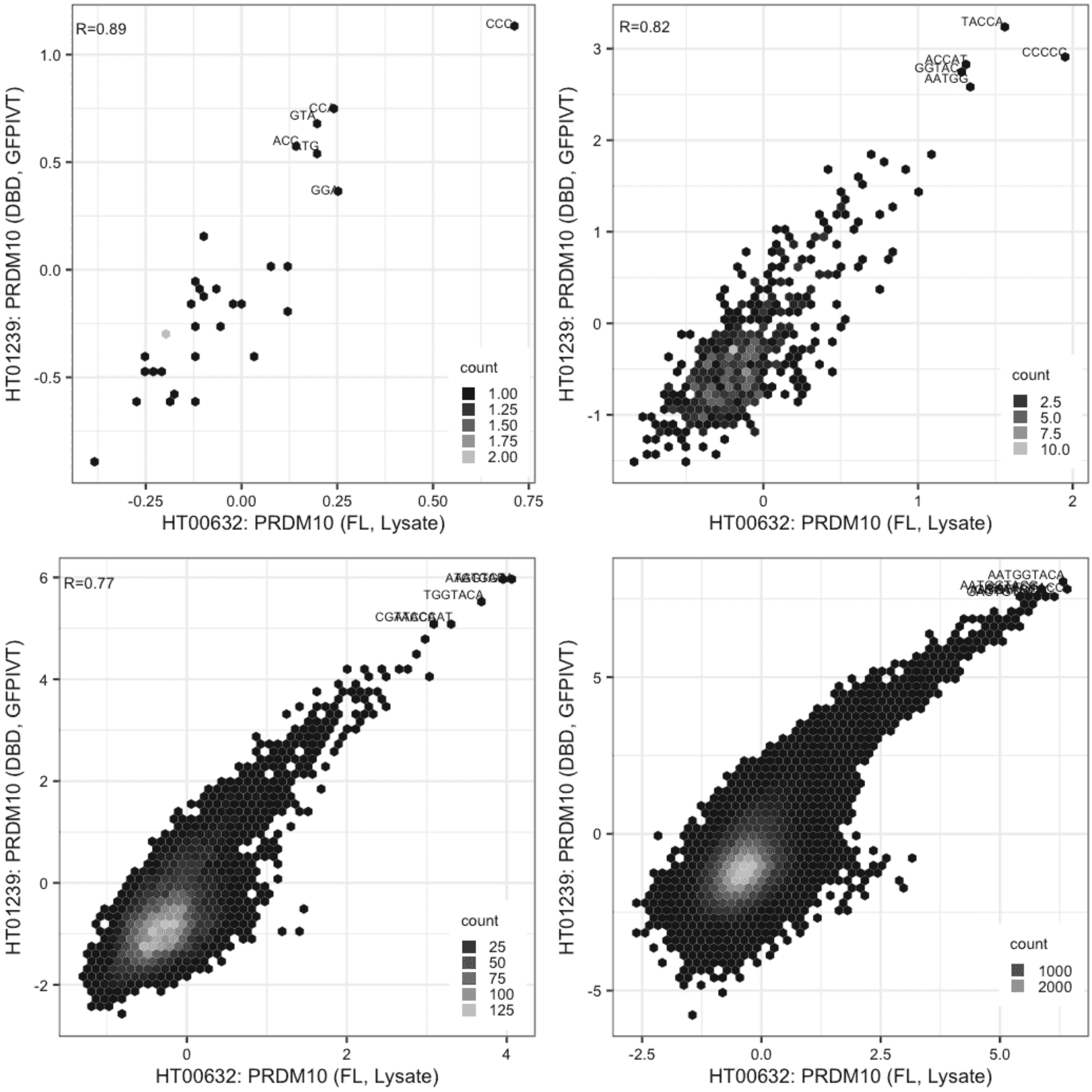

### PRDM10

Lysate vs GFPIVT, different constructs

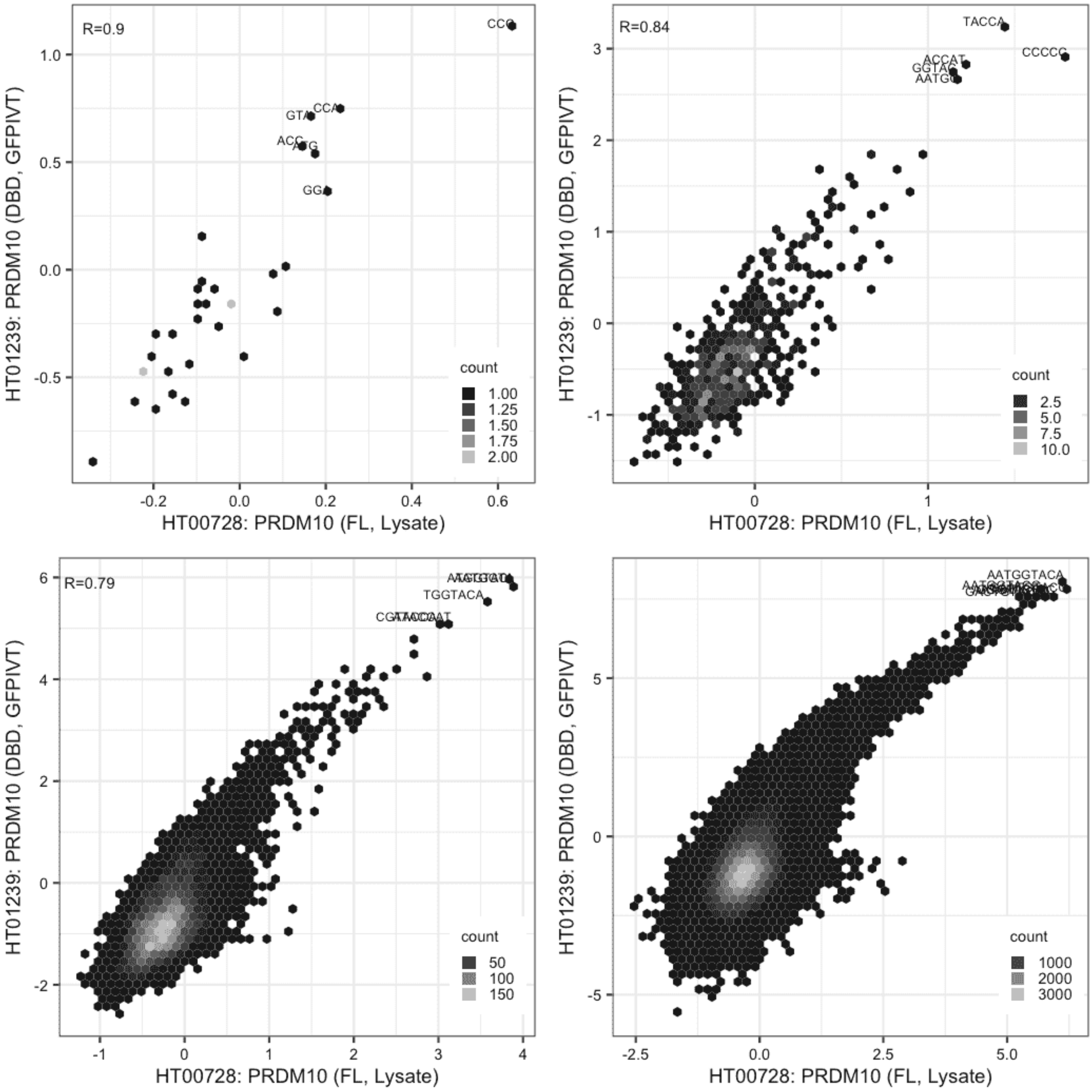

**PRDM13**

Lysate vs IVT, different constructs

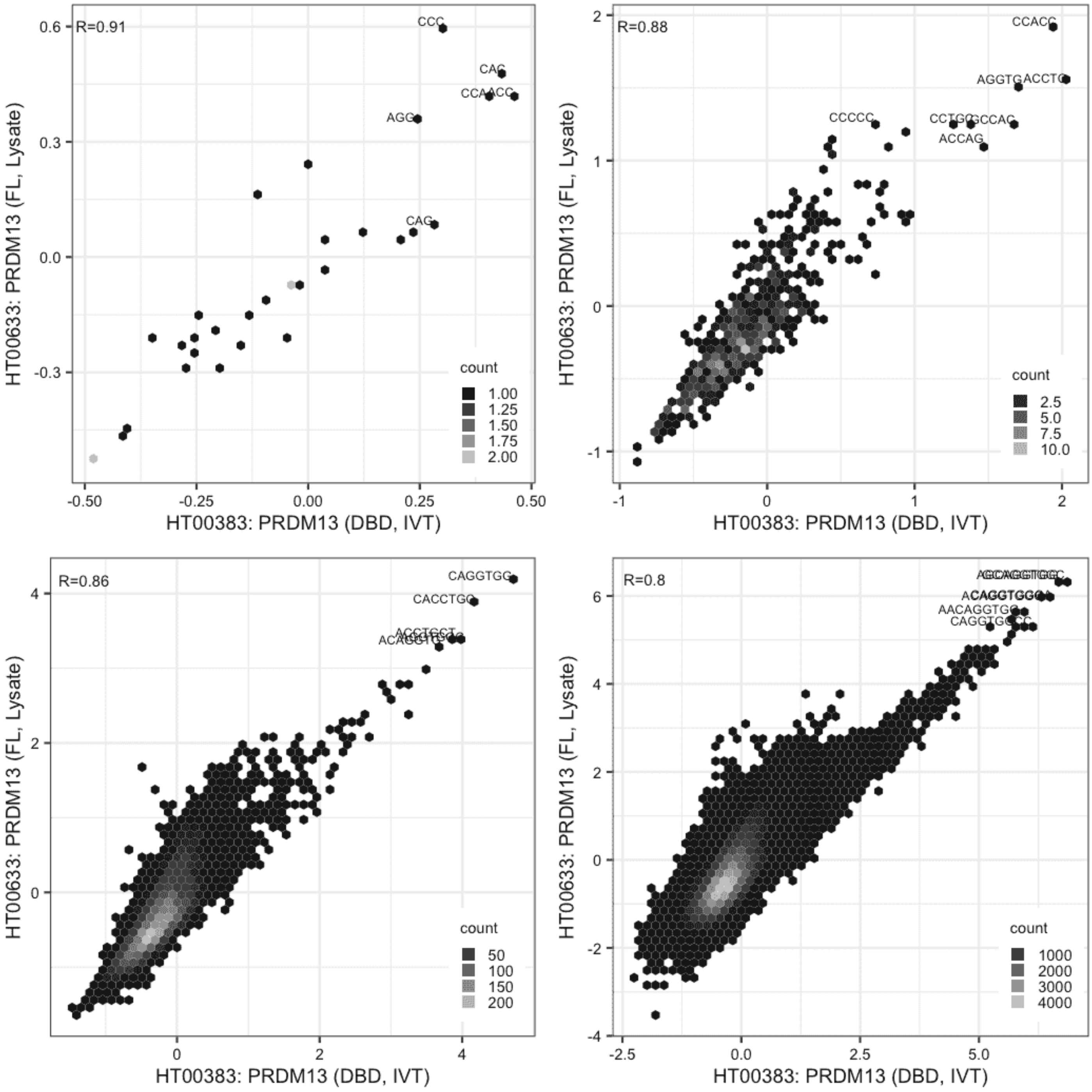

**PRDM13**

Lysate vs IVT, different constructs

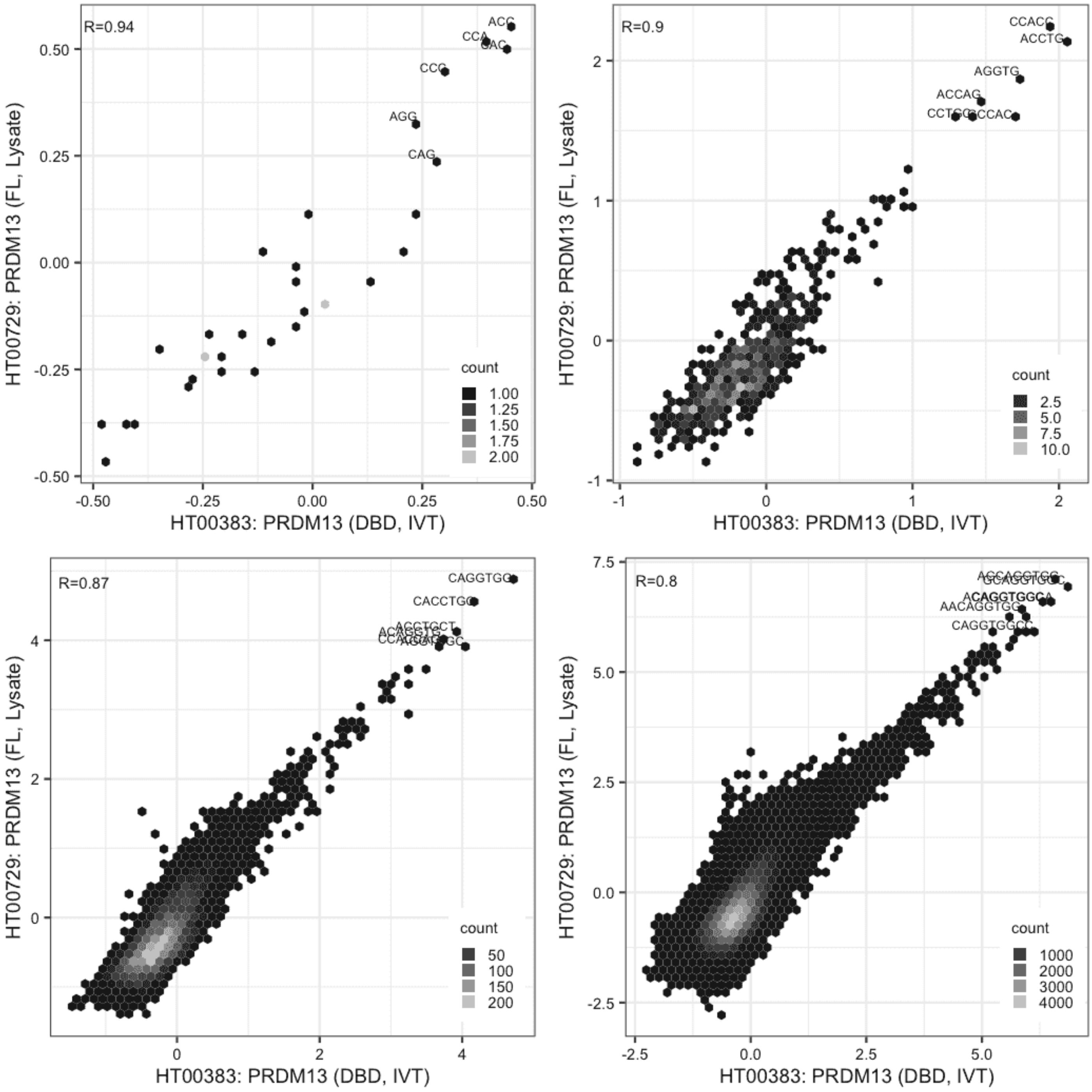

**PRDM13**

IVT vs GFPIVT, same construct

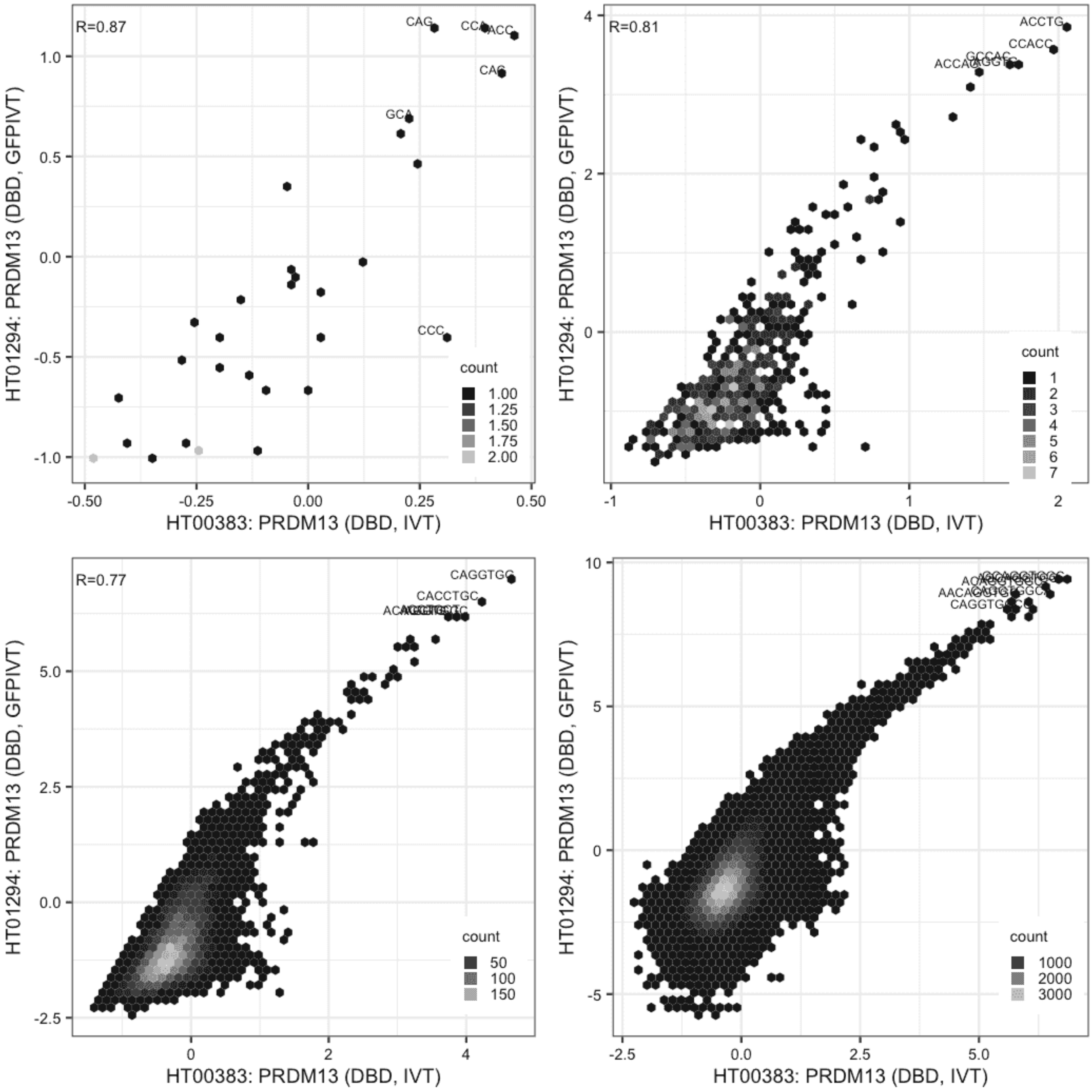

**PRDM13**

IVT vs GFPIVT, different constructs

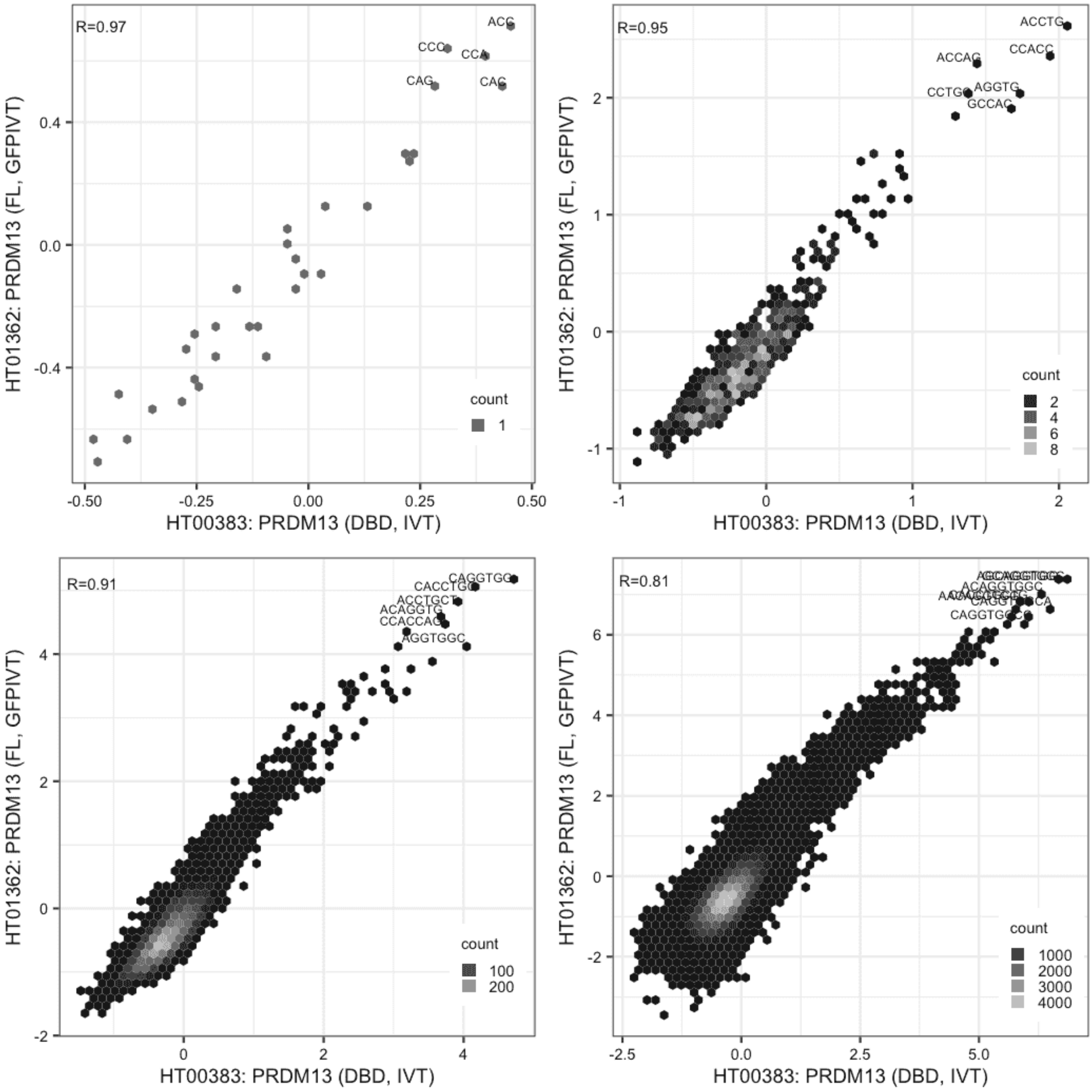

**PRDM13**

Lysate vs Lysate, same construct

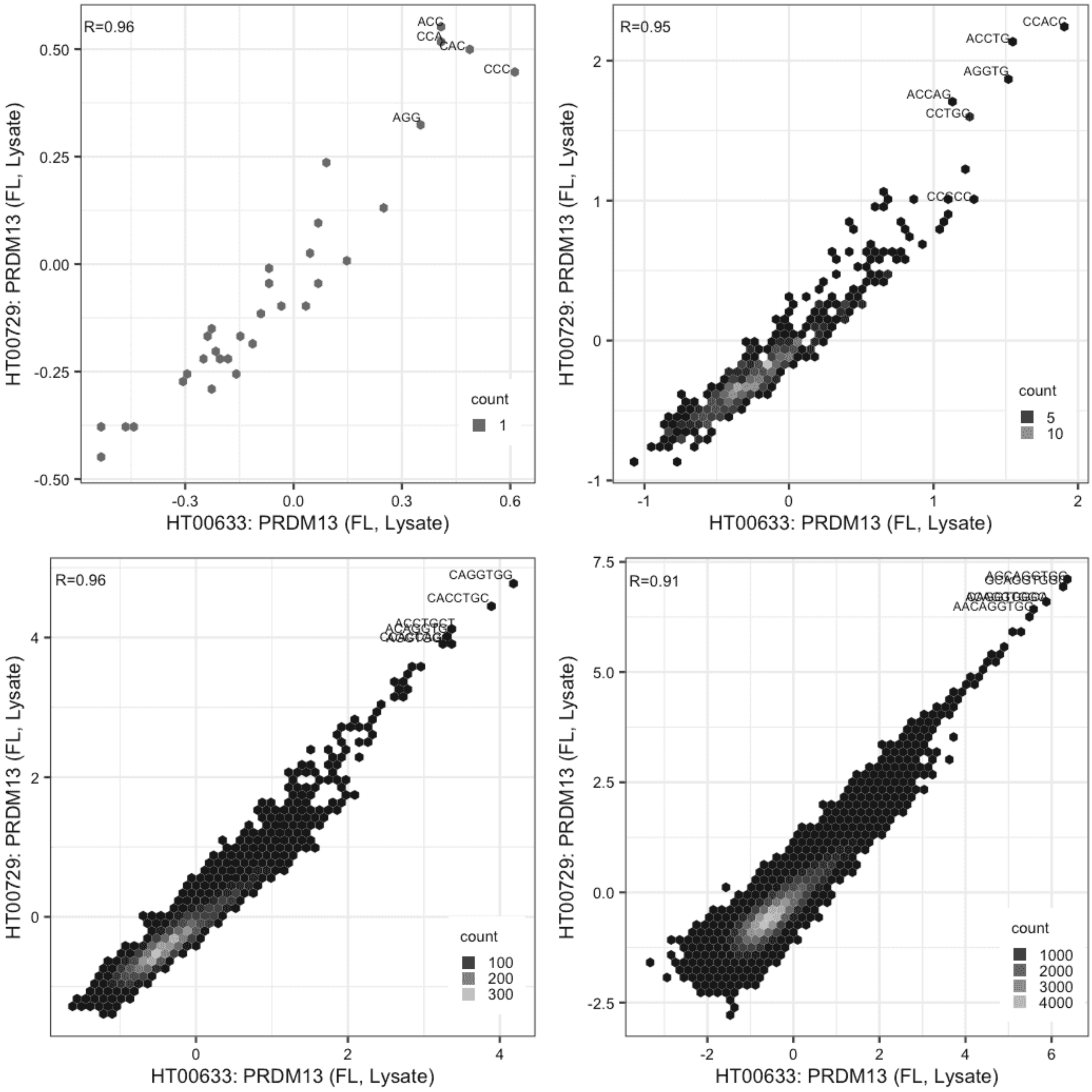

**PRDM13**

Lysate vs GFPIVT, different constructs

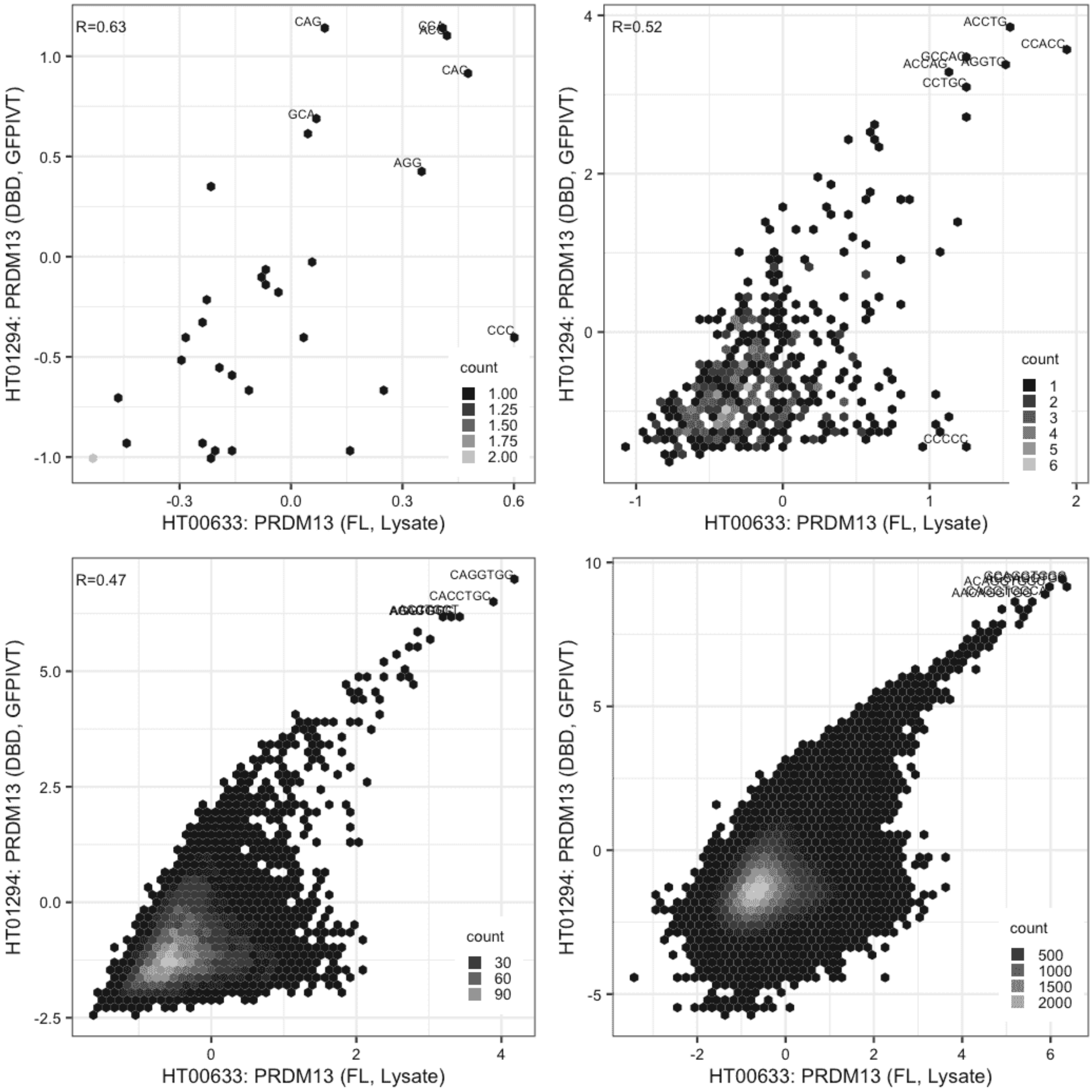

**PRDM13**

Lysate vs GFPIVT, same construct

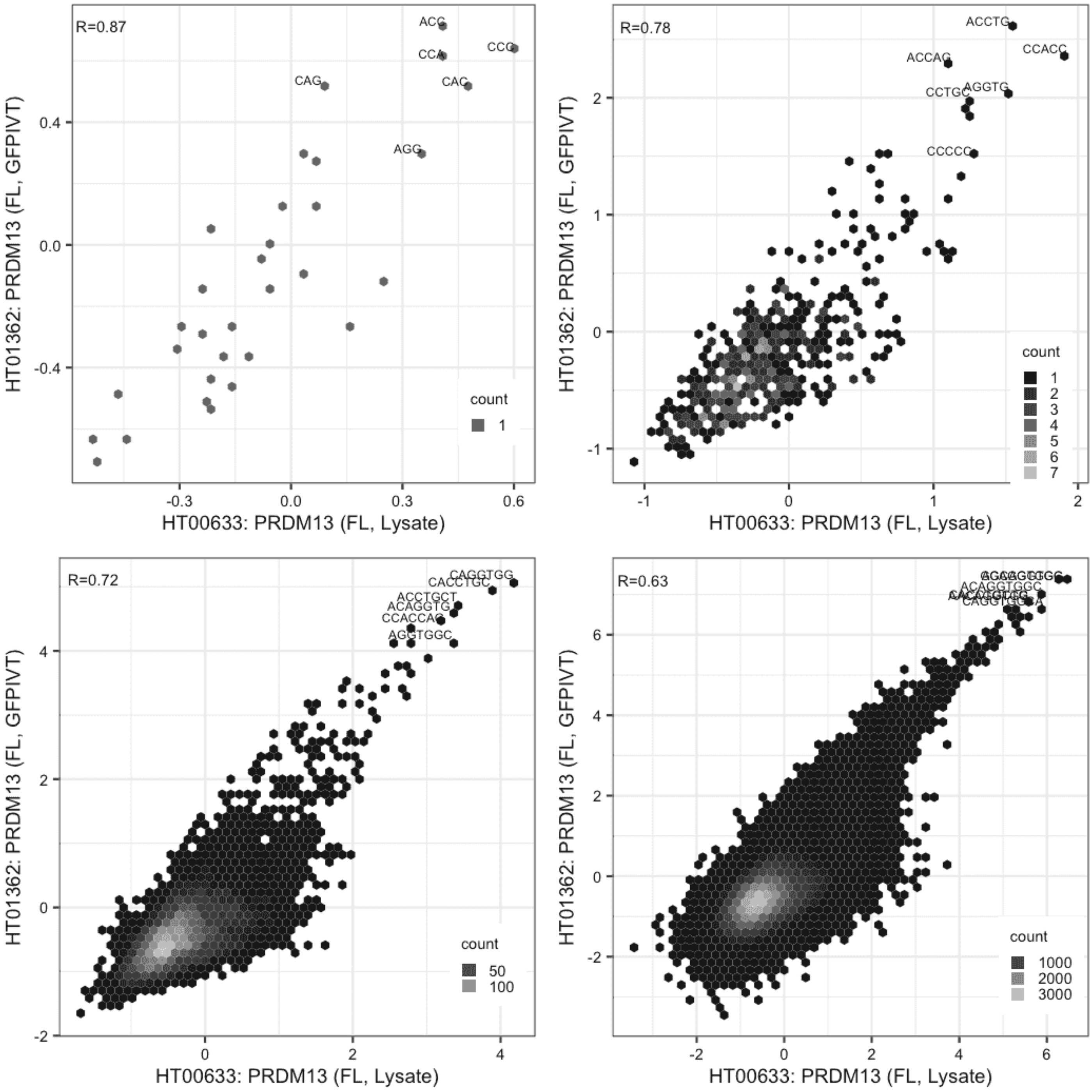

**PRDM13**

Lysate vs GFPIVT, different constructs

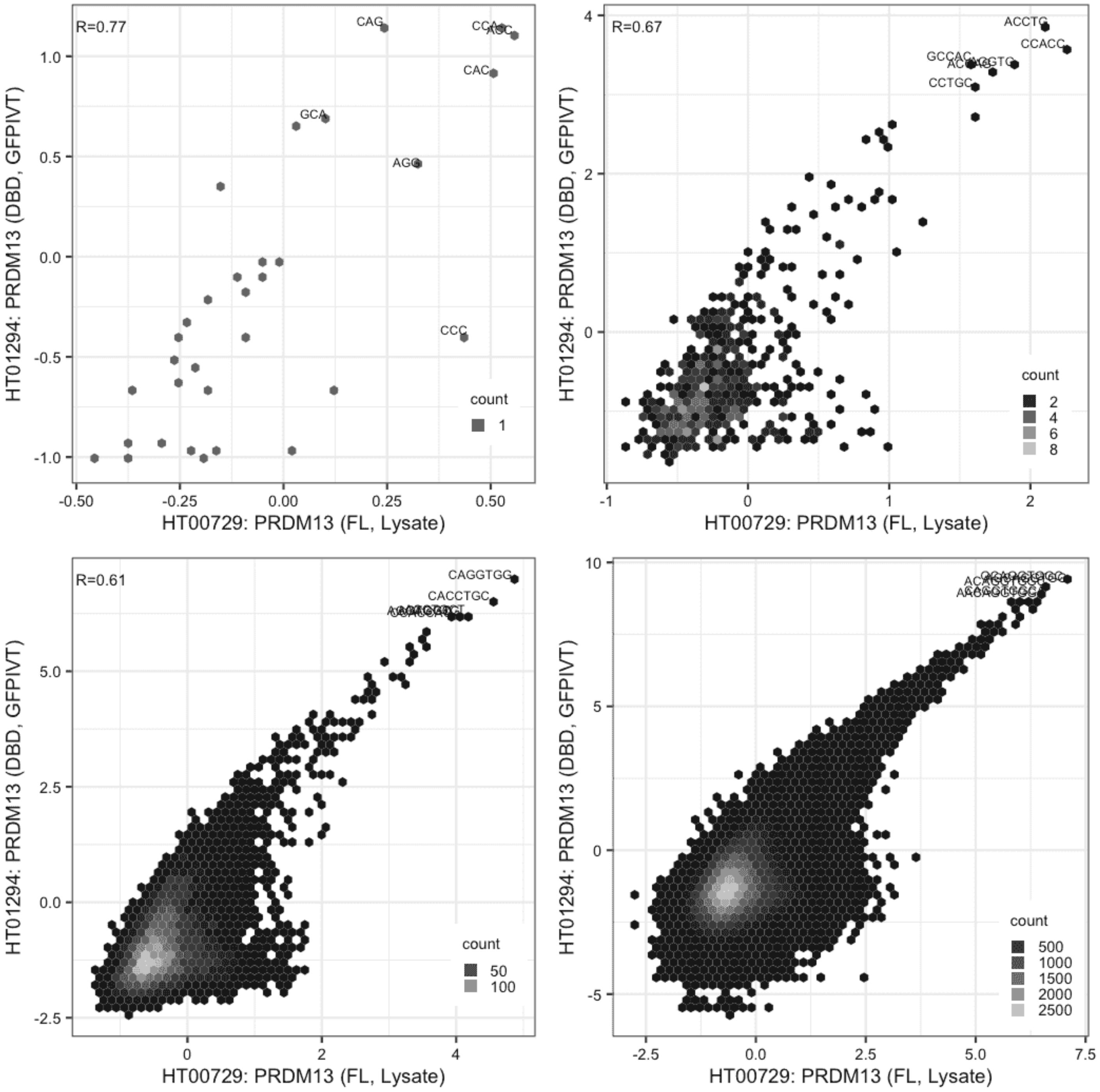

**PRDM13**

Lysate vs GFPIVT, same construct

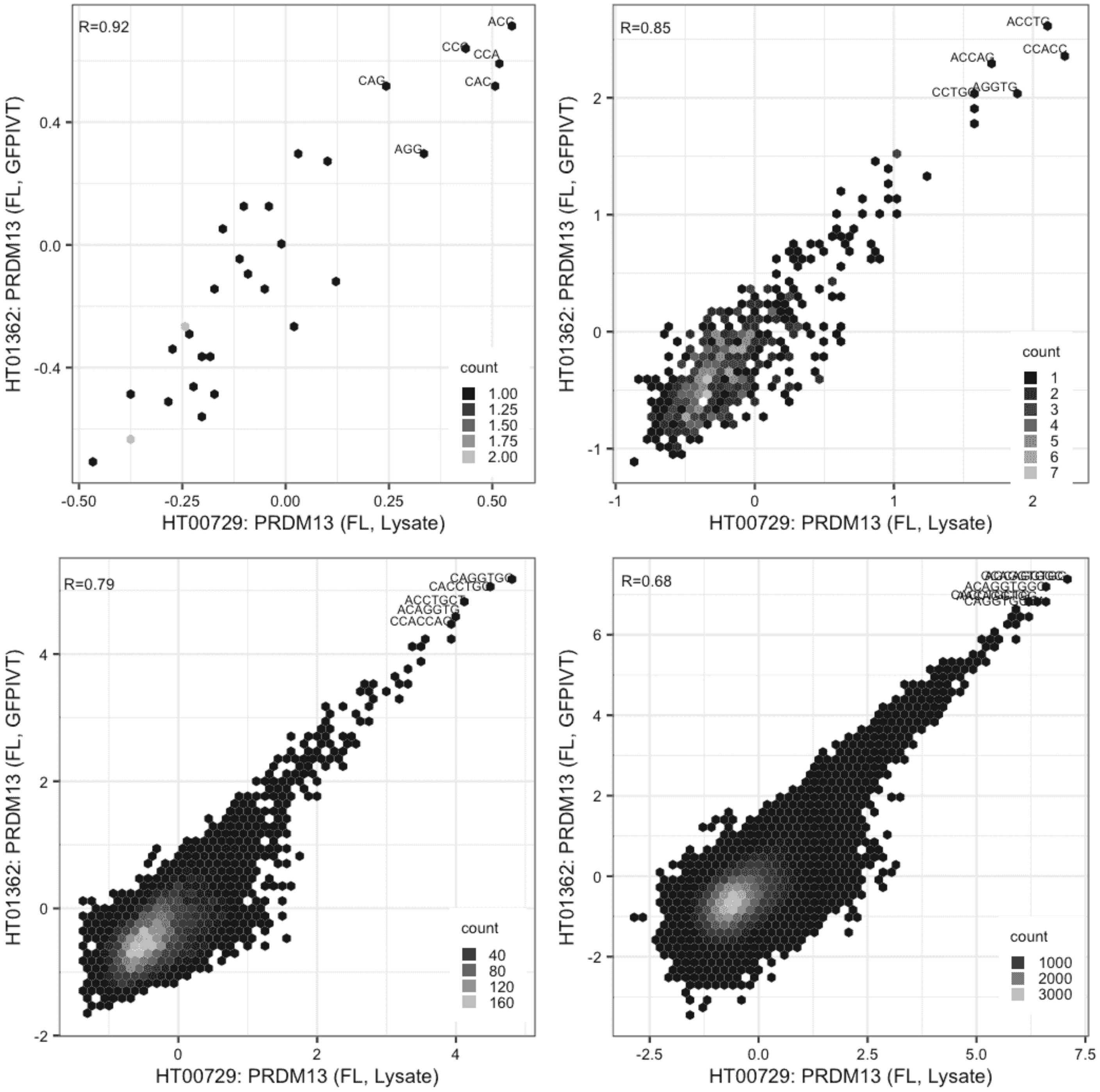

**PRDM13**

GFPIVT vs GFPIVT, different constructs

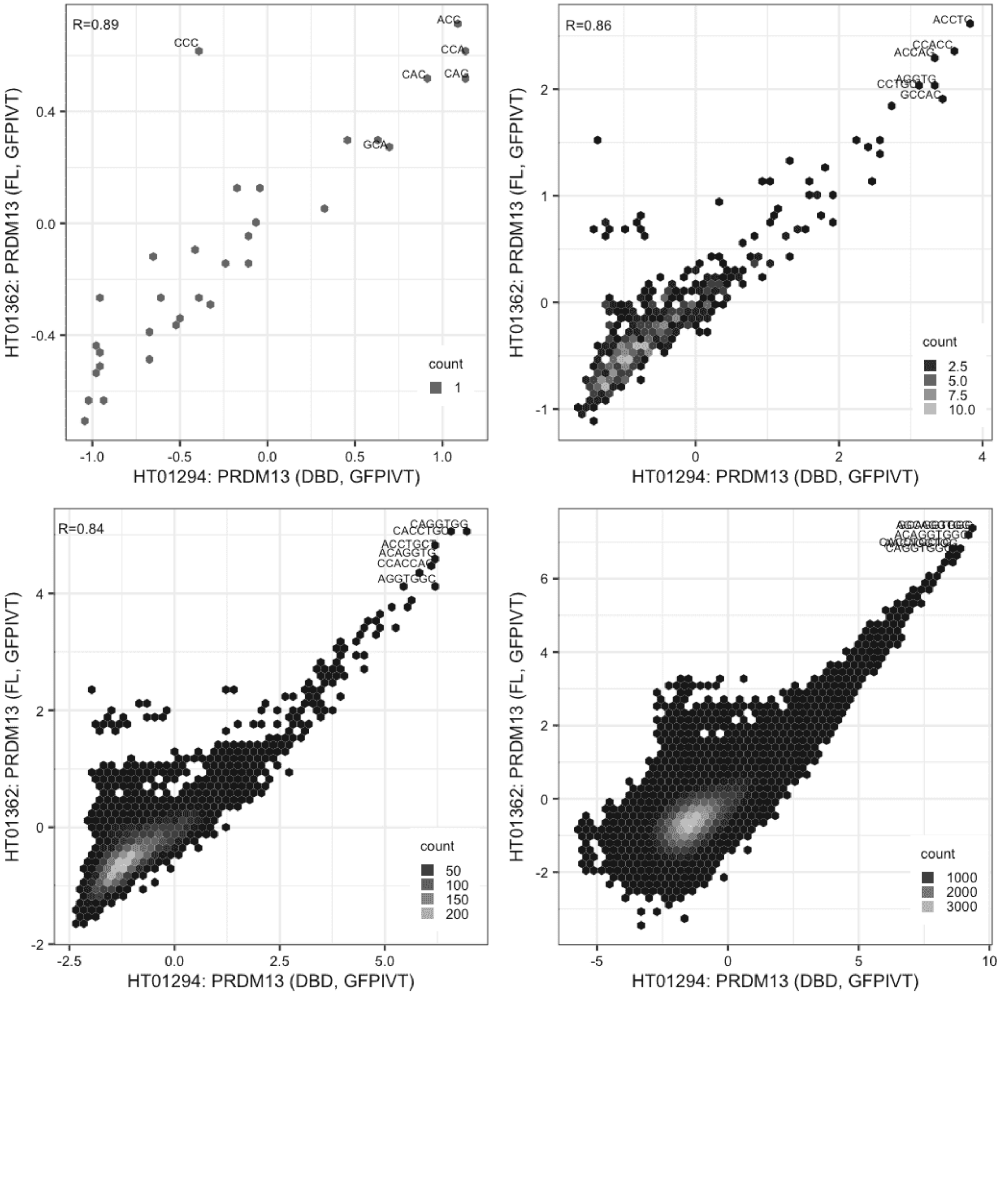

**PRDM5**

Lysate vs Lysate, same construct

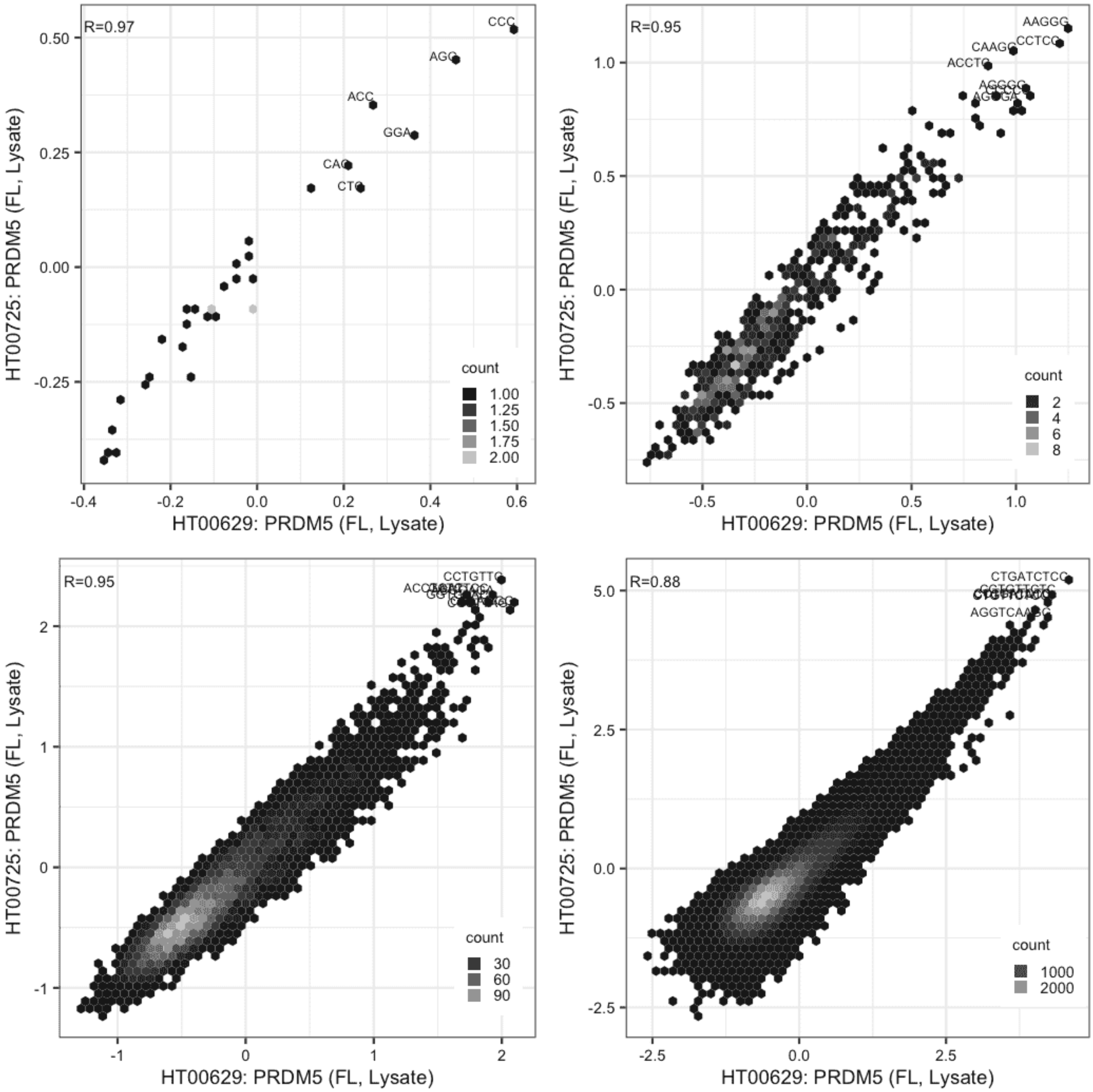

**PRDM5**

Lysate vs GFPIVT, same construct

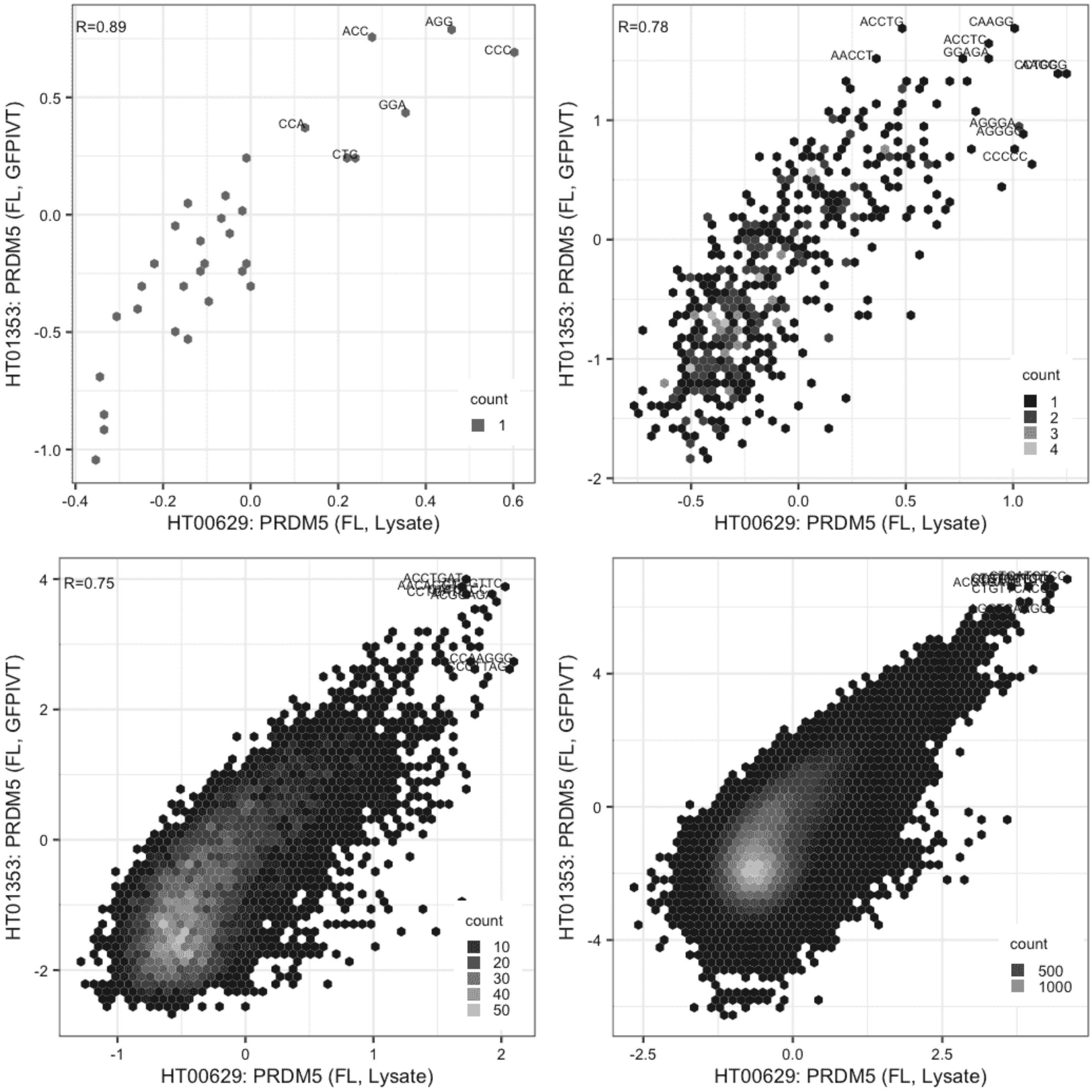

**PRDM5**

Lysate vs GFPIVT, different constructs

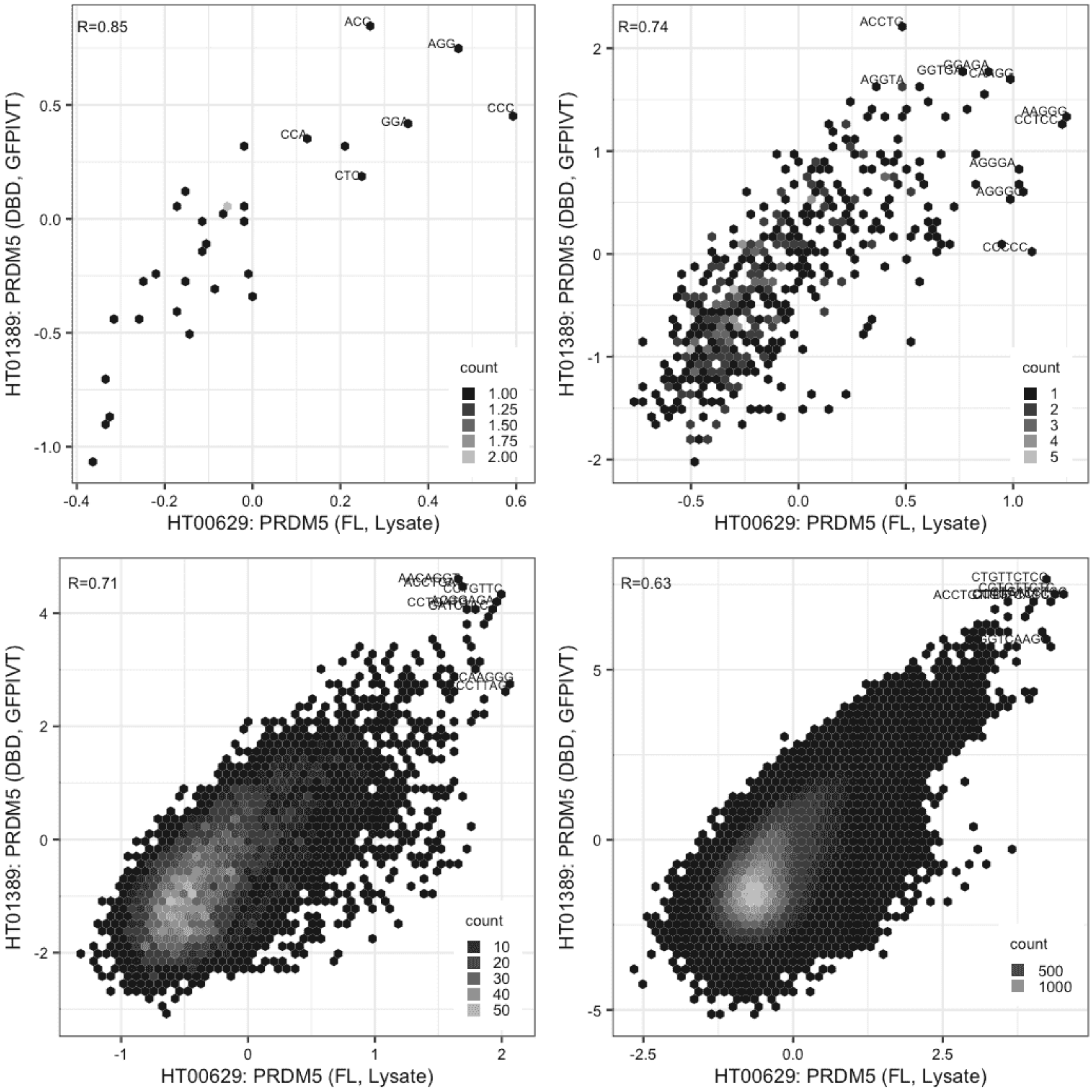

**PRDM5**

Lysate vs GFPIVT, same construct

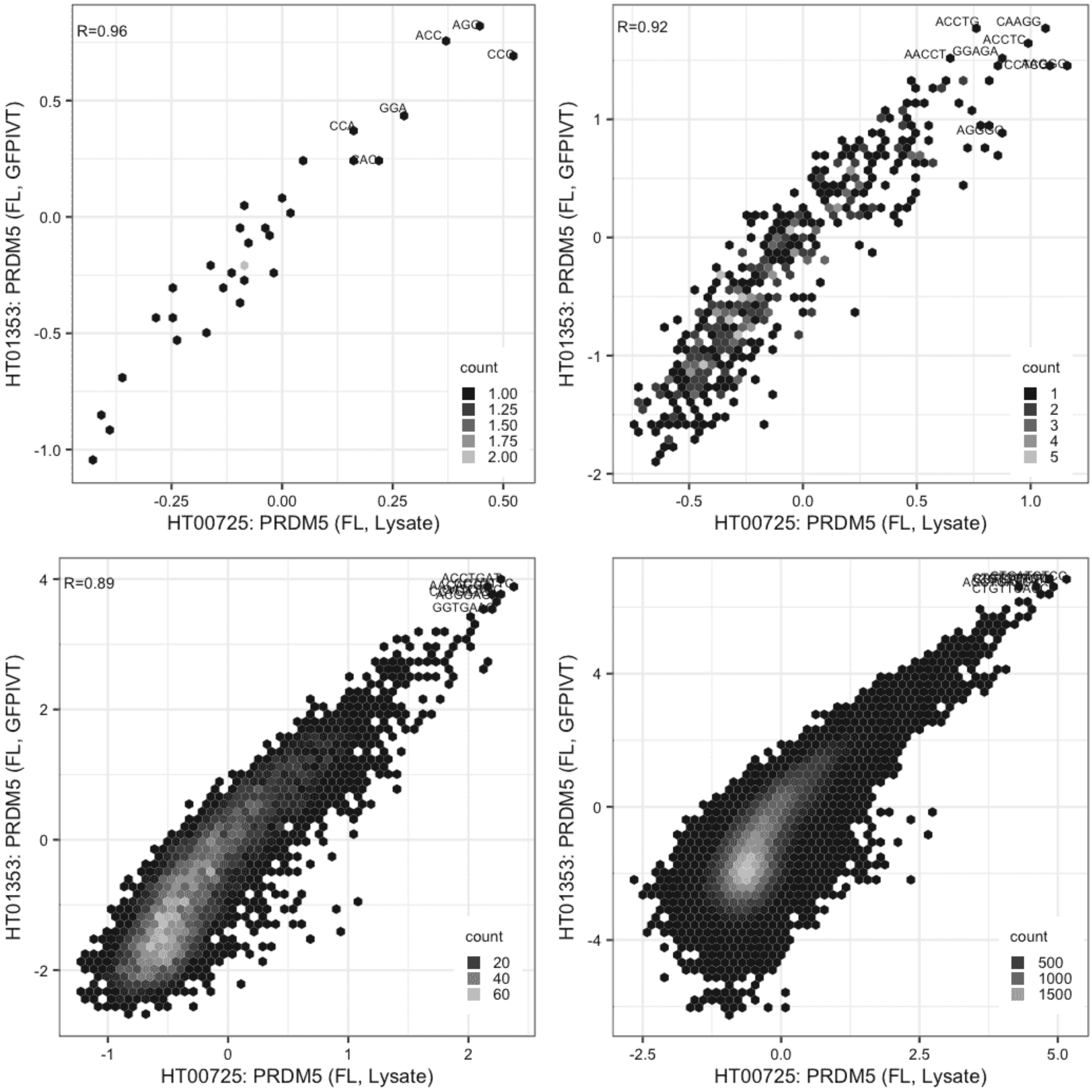

**PRDM5**

Lysate vs GFPIVT, different constructs

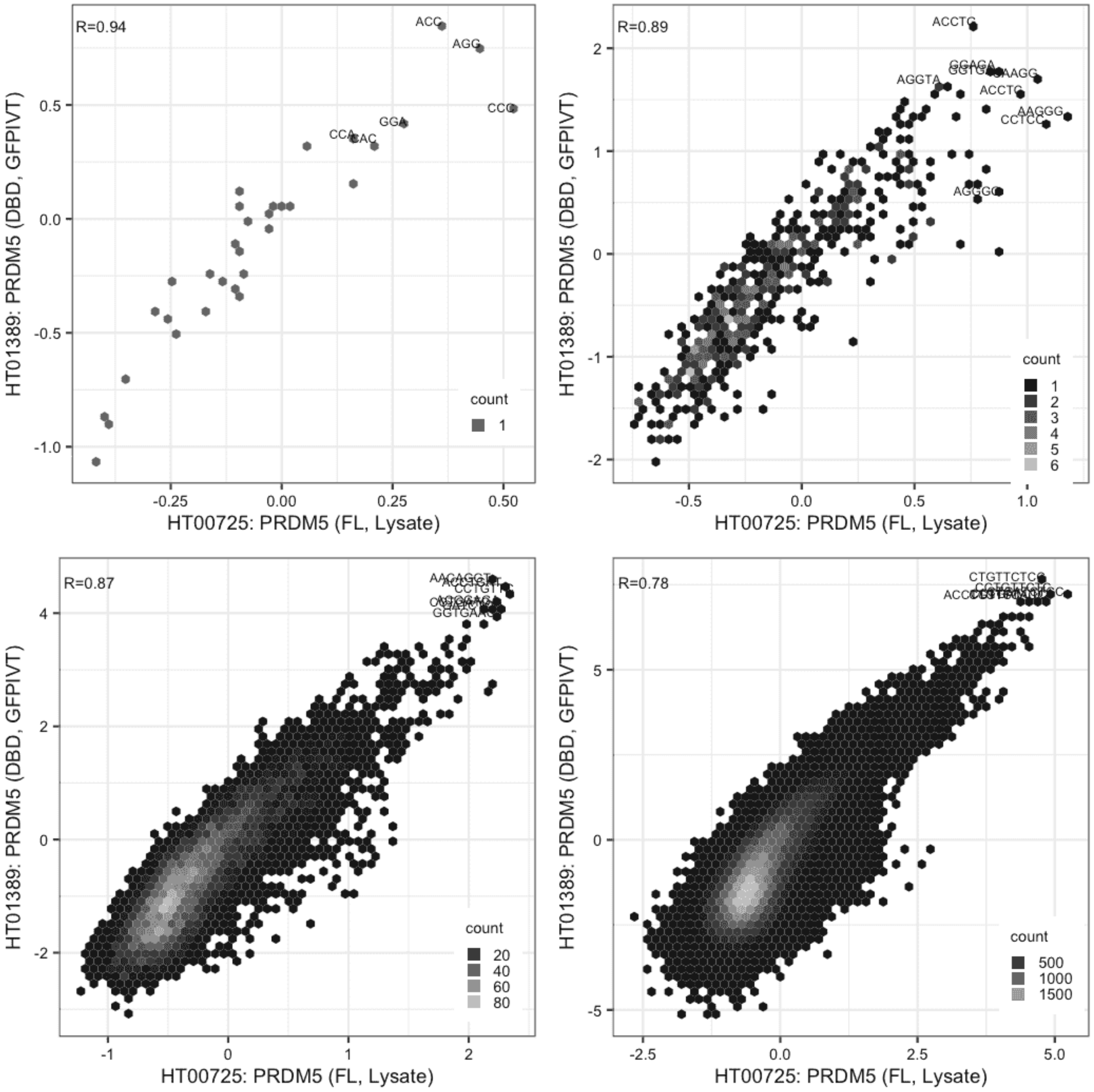

**PRDM5**

GFPIVT vs GFPIVT, different constructs

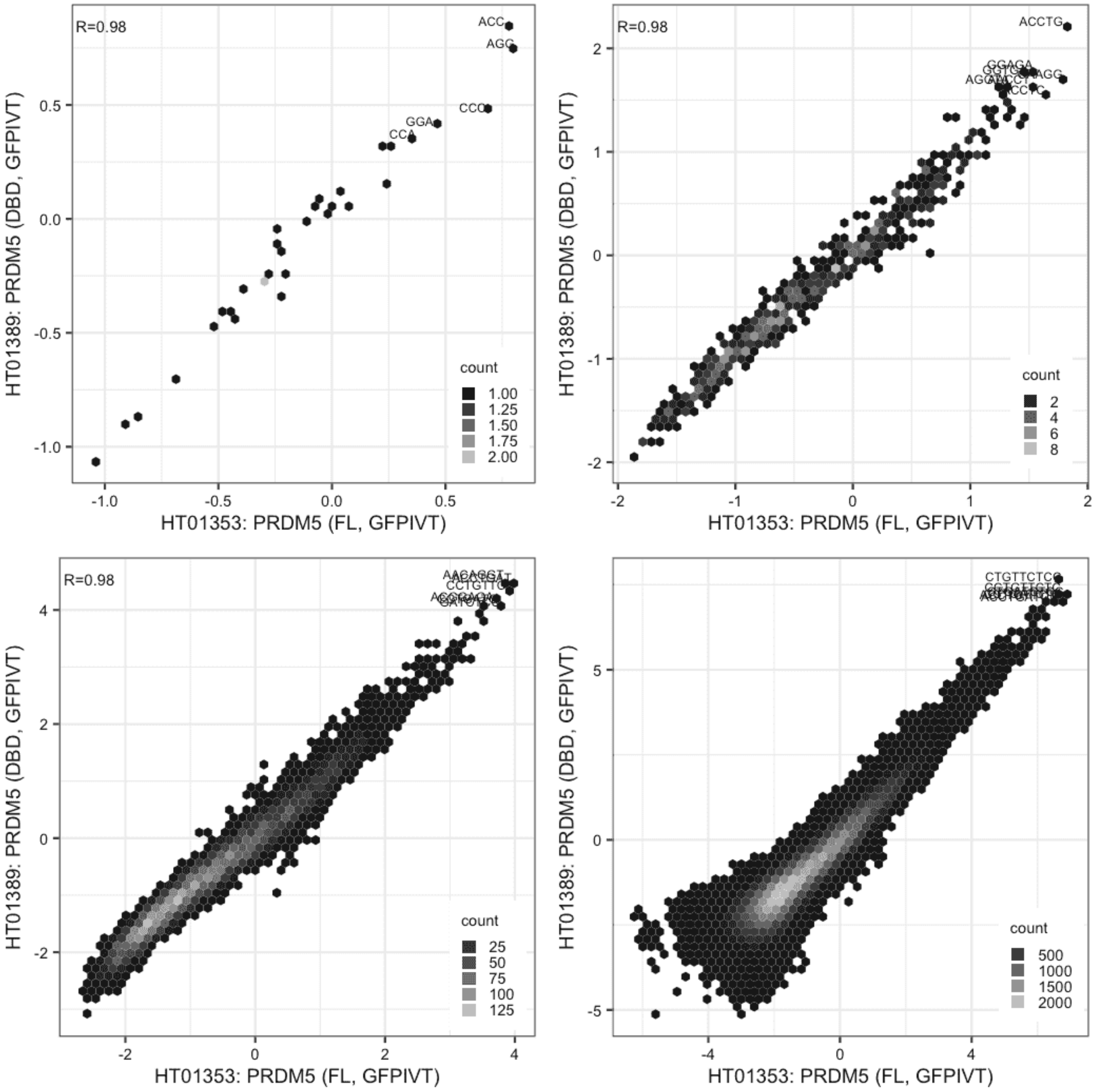

**RARA**

Lysate vs IVT, same construct

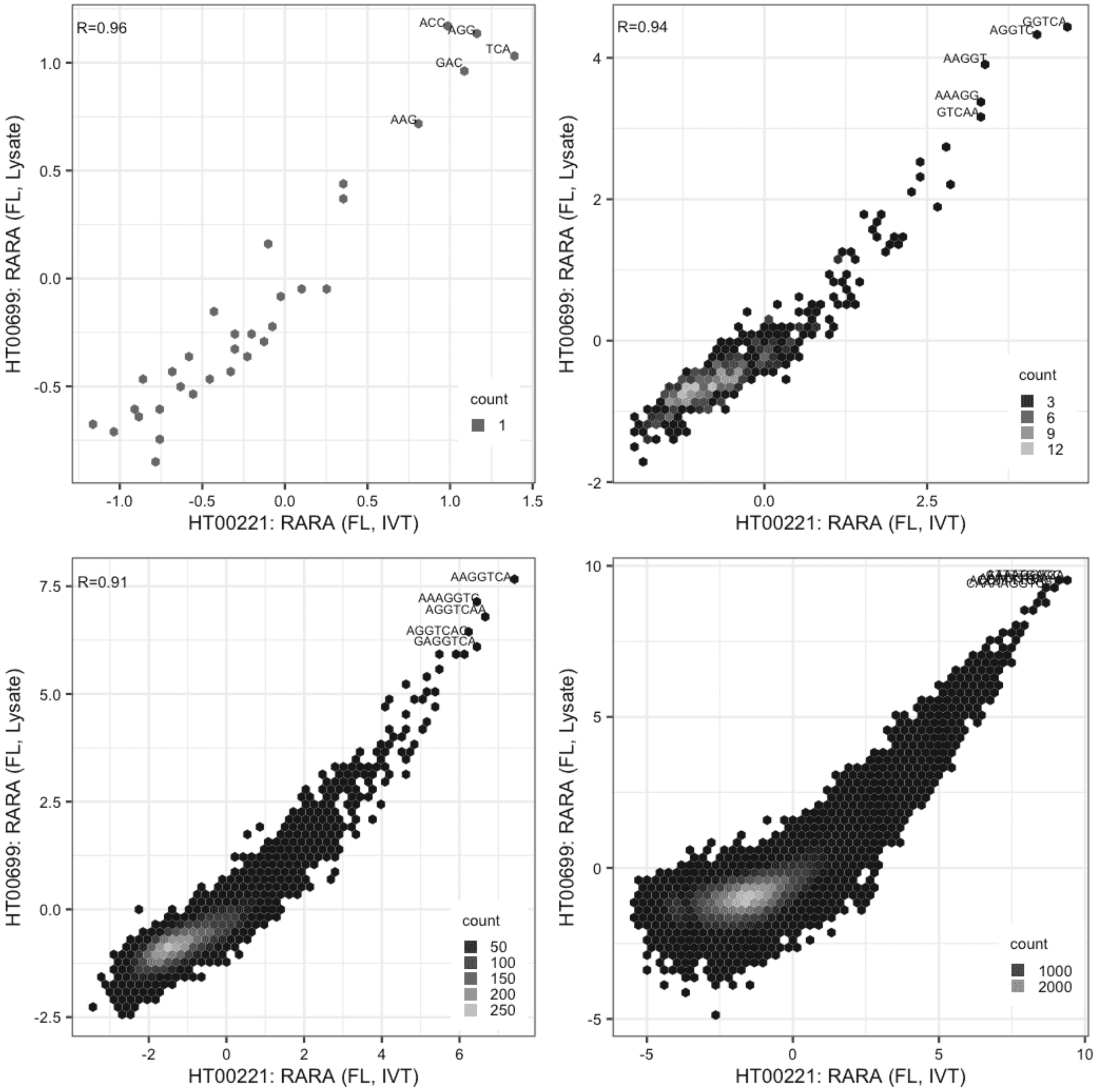

### RARA

Lysate vs IVT, same construct

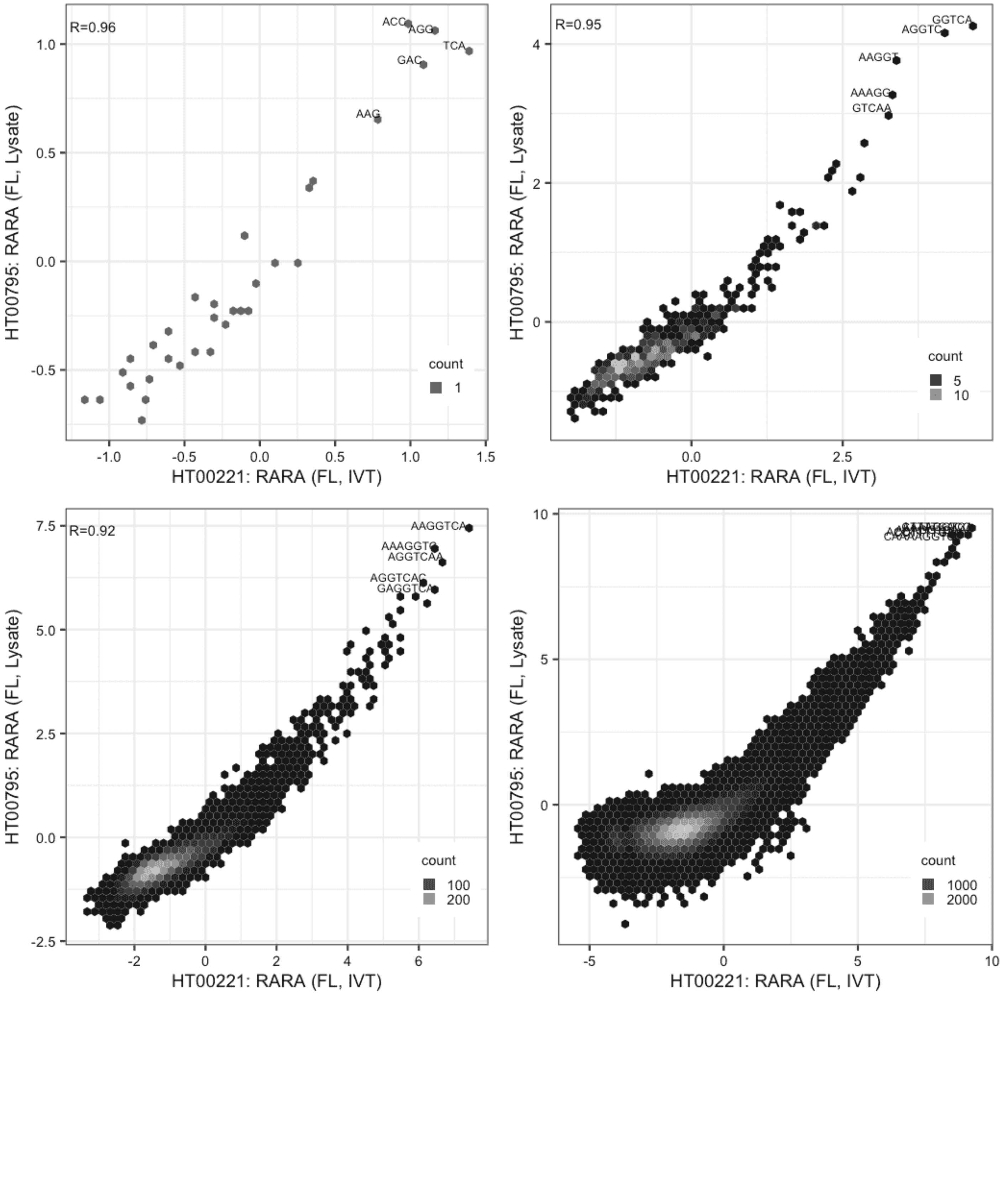

#### RARA

Lysate vs Lysate, same construct

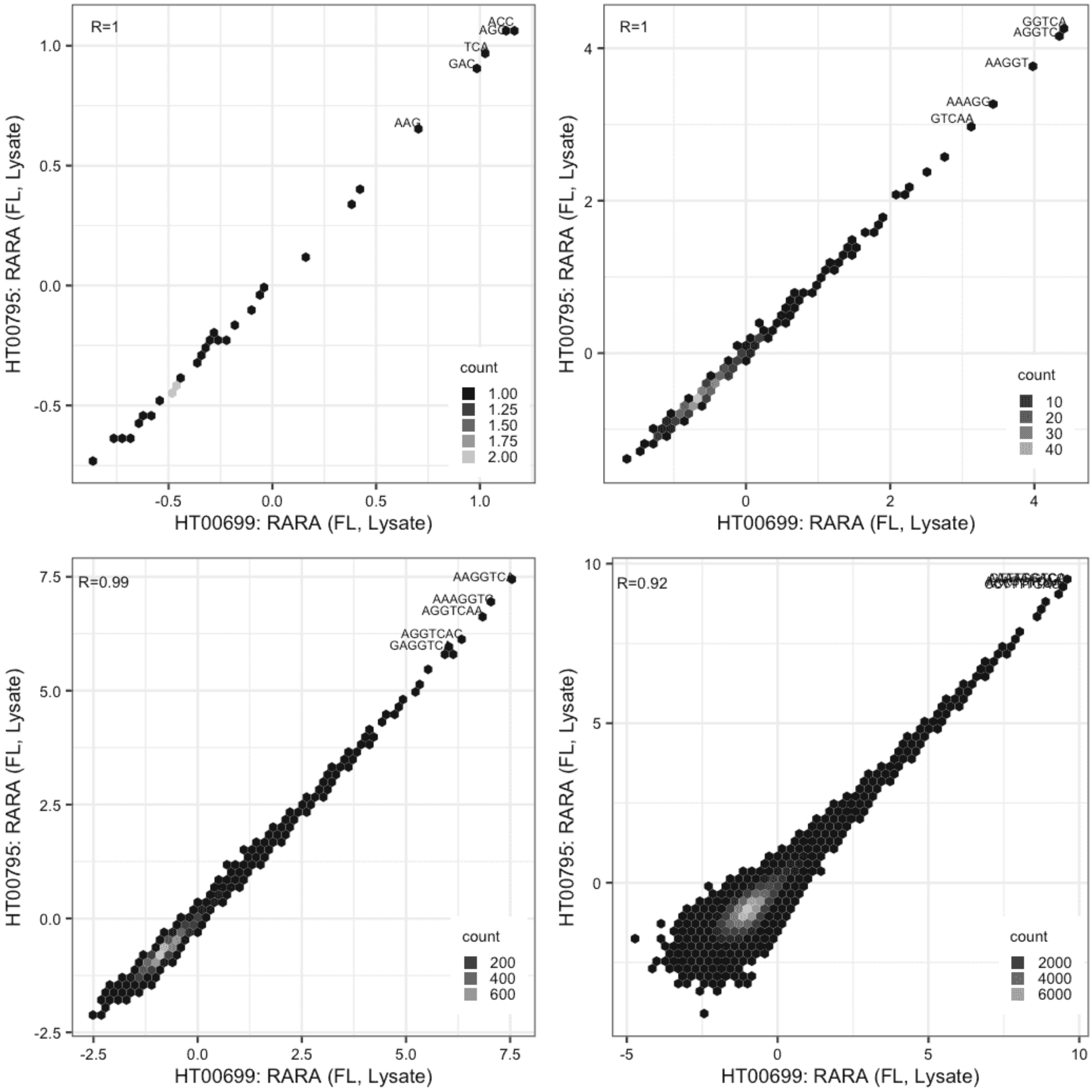

#### RORB

Lysate vs IVT, same construct

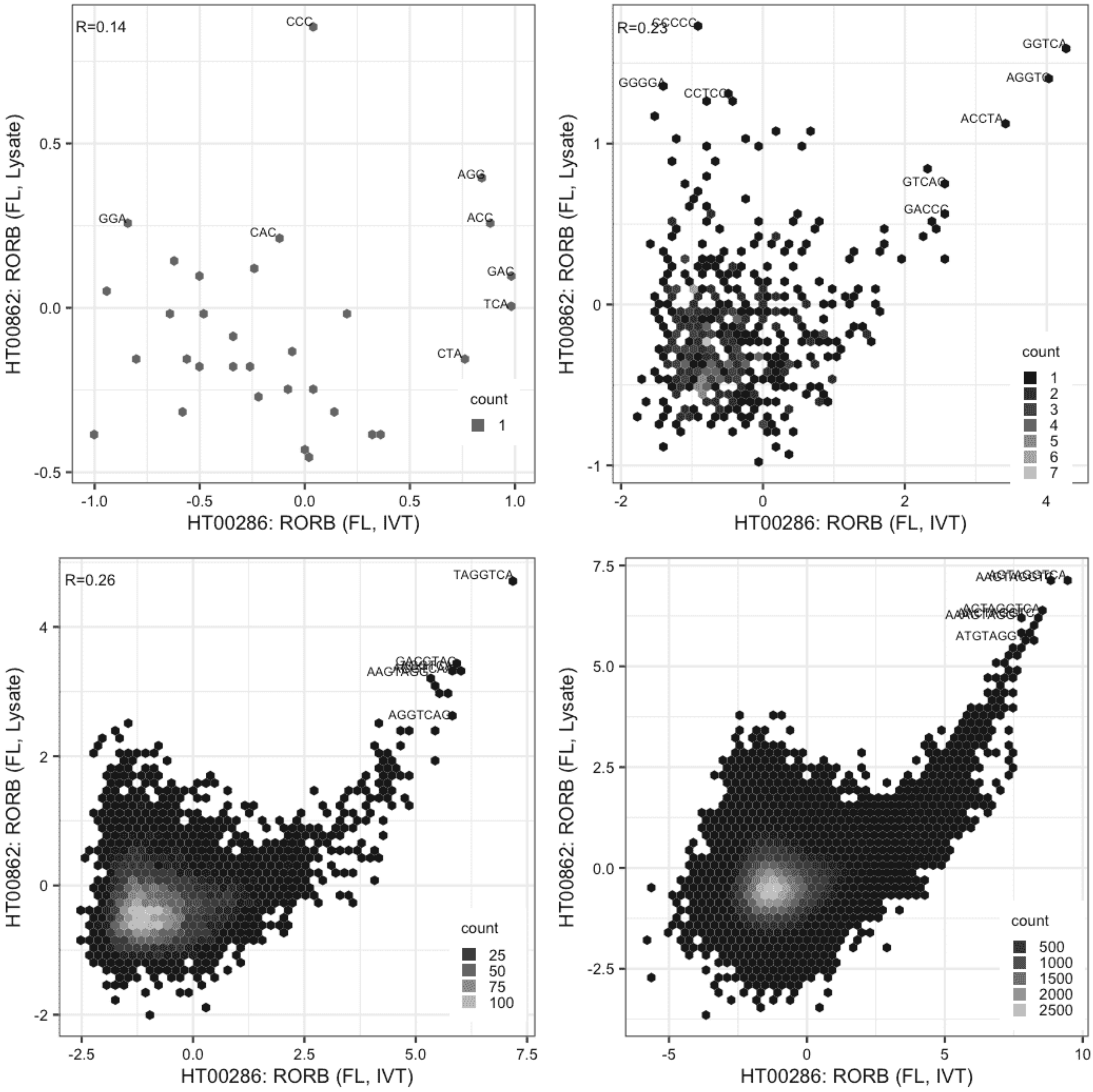

**RXRA**

IVT vs IVT, same construct

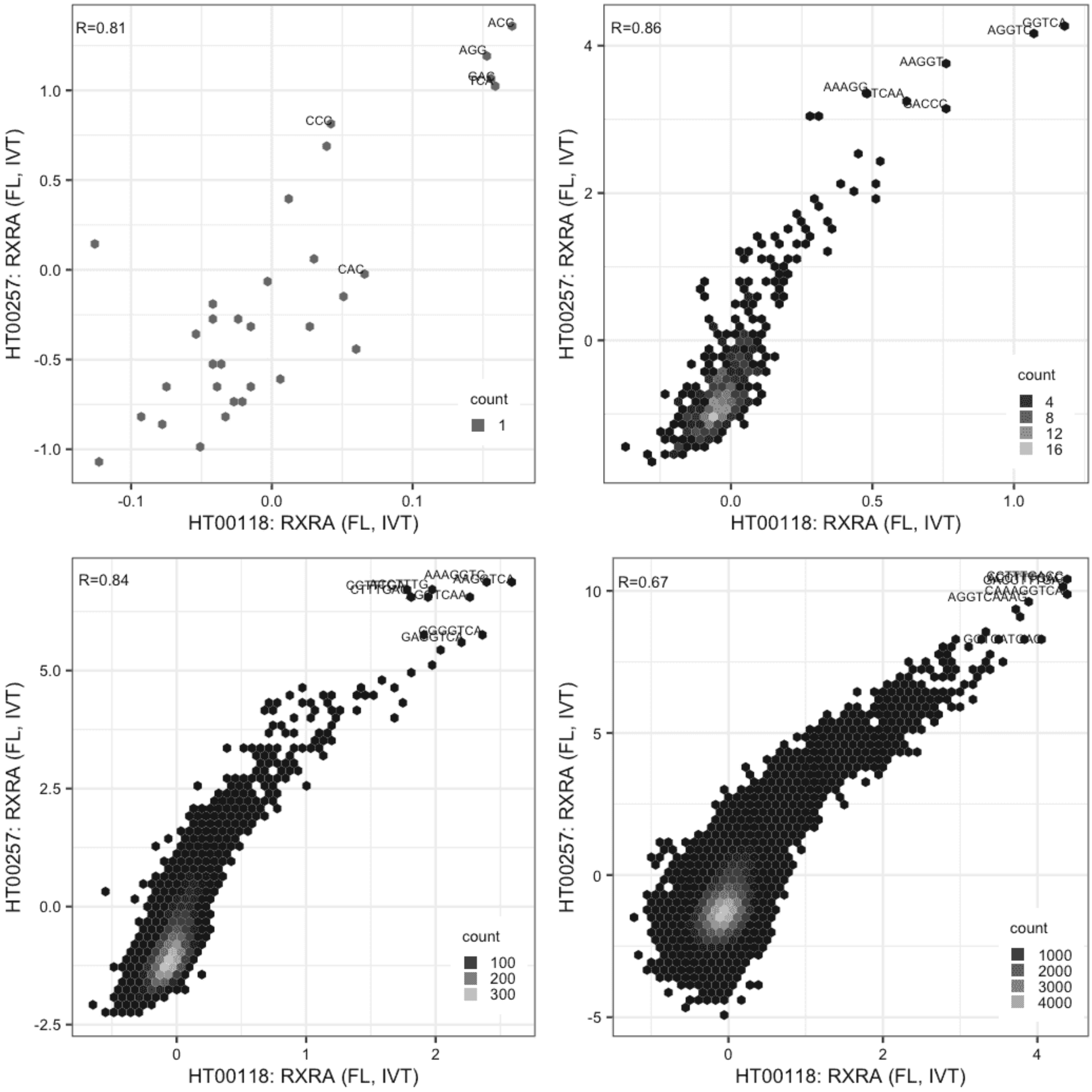

**RXRA**

Lysate vs IVT, same construct

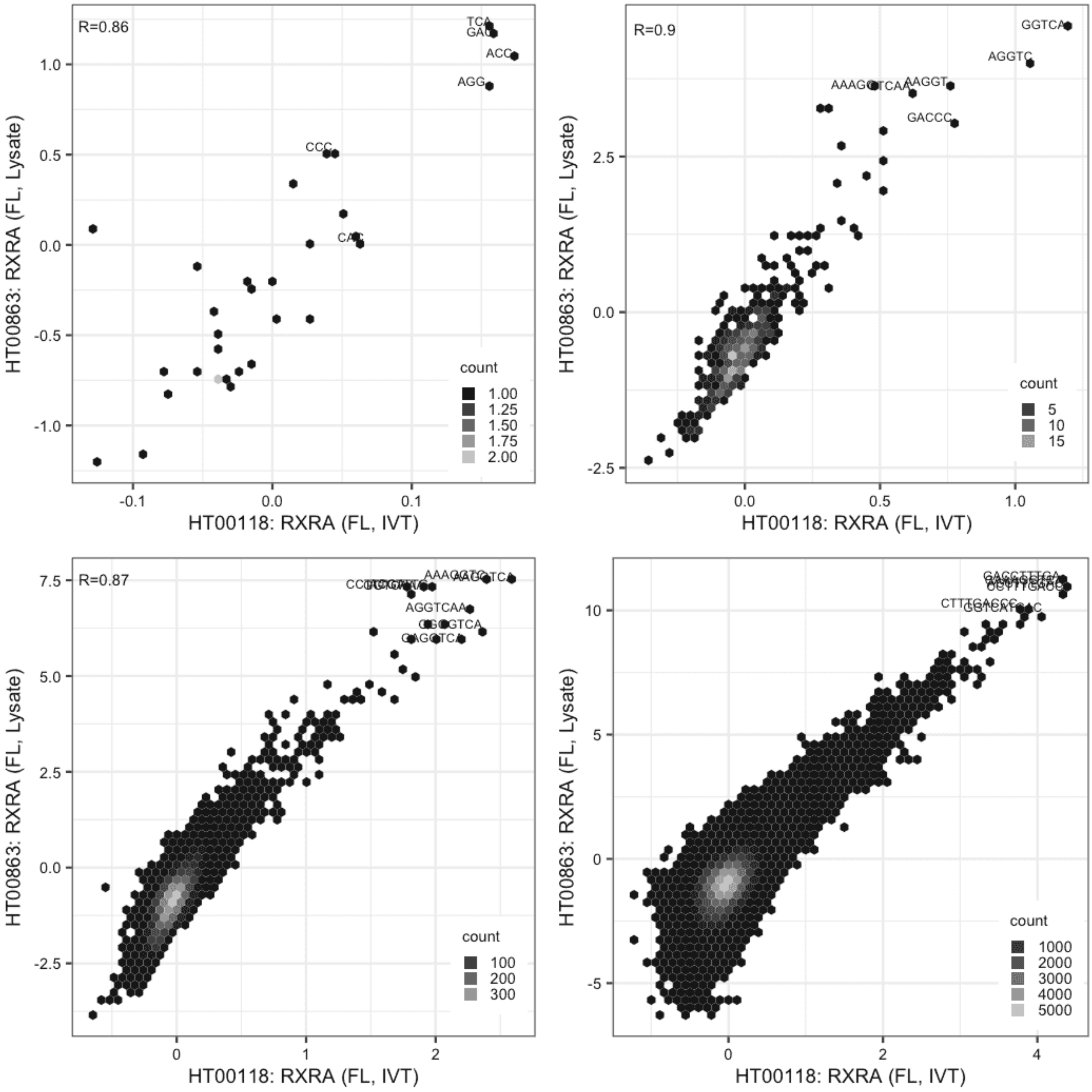

### RXRA

Lysate vs IVT, same construct

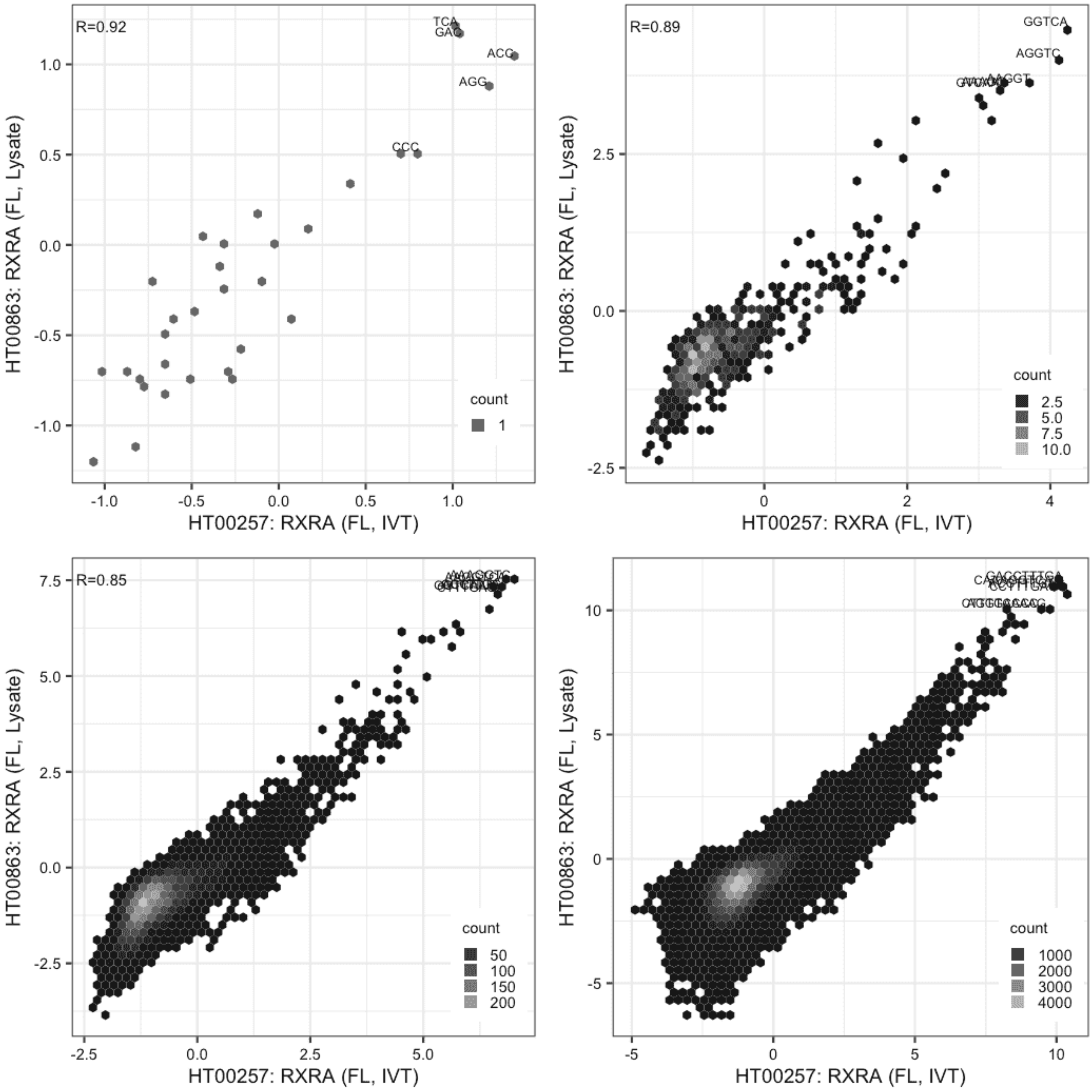

**SLC2A4RG**

Lysate vs IVT, different constructs

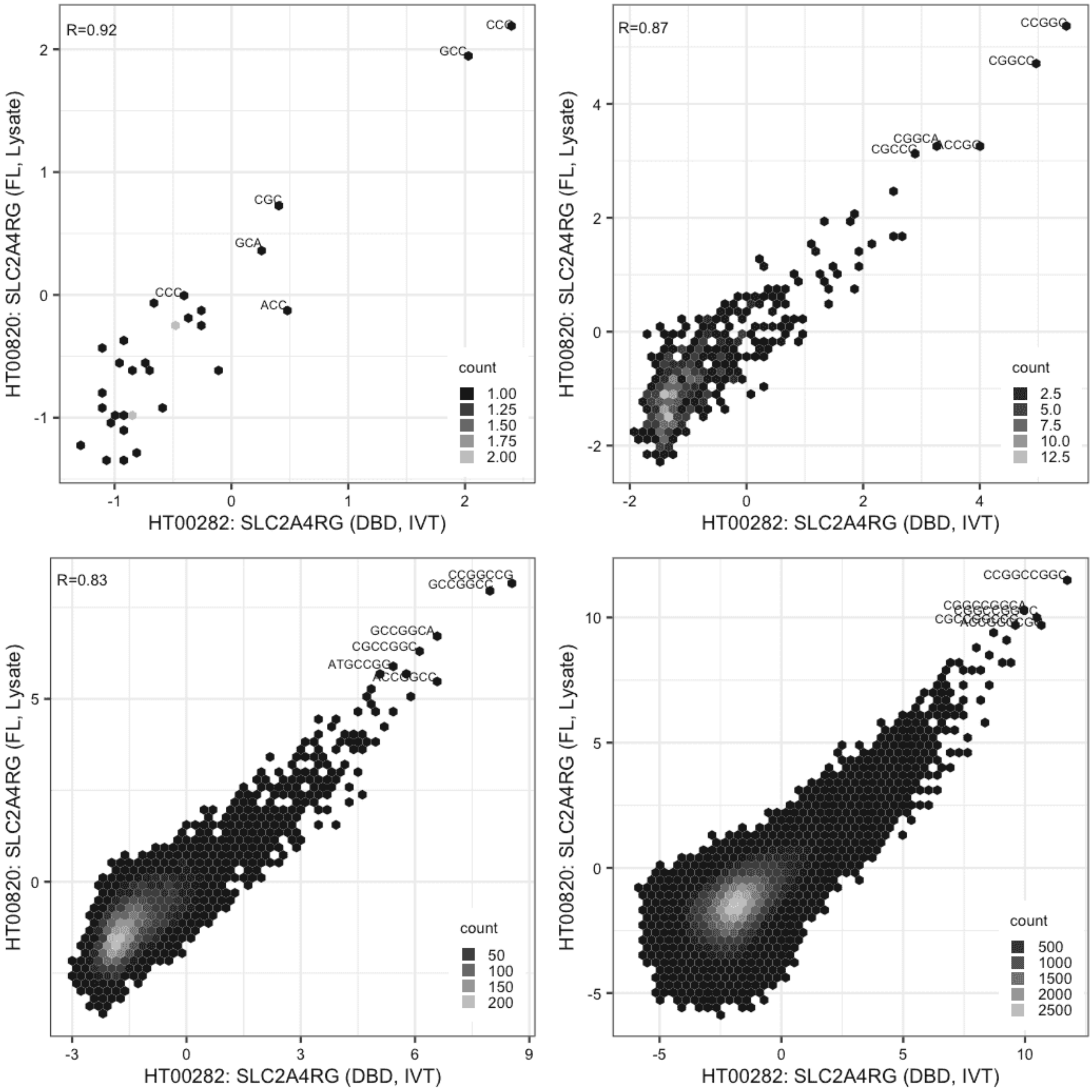

**SLC2A4RG**

IVT vs GFPIVT, different constructs

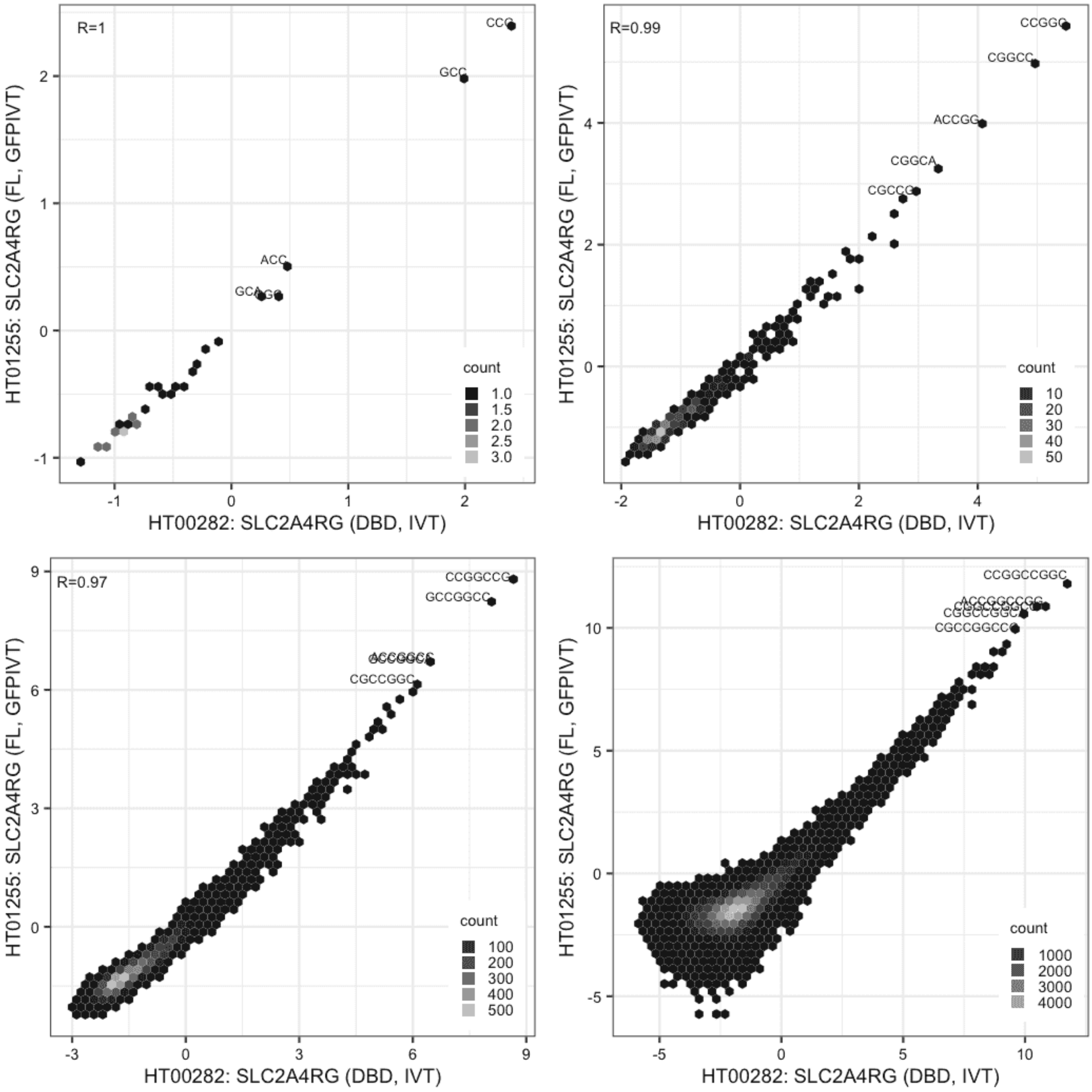

**SLC2A4RG**

Lysate vs GFPIVT, same construct

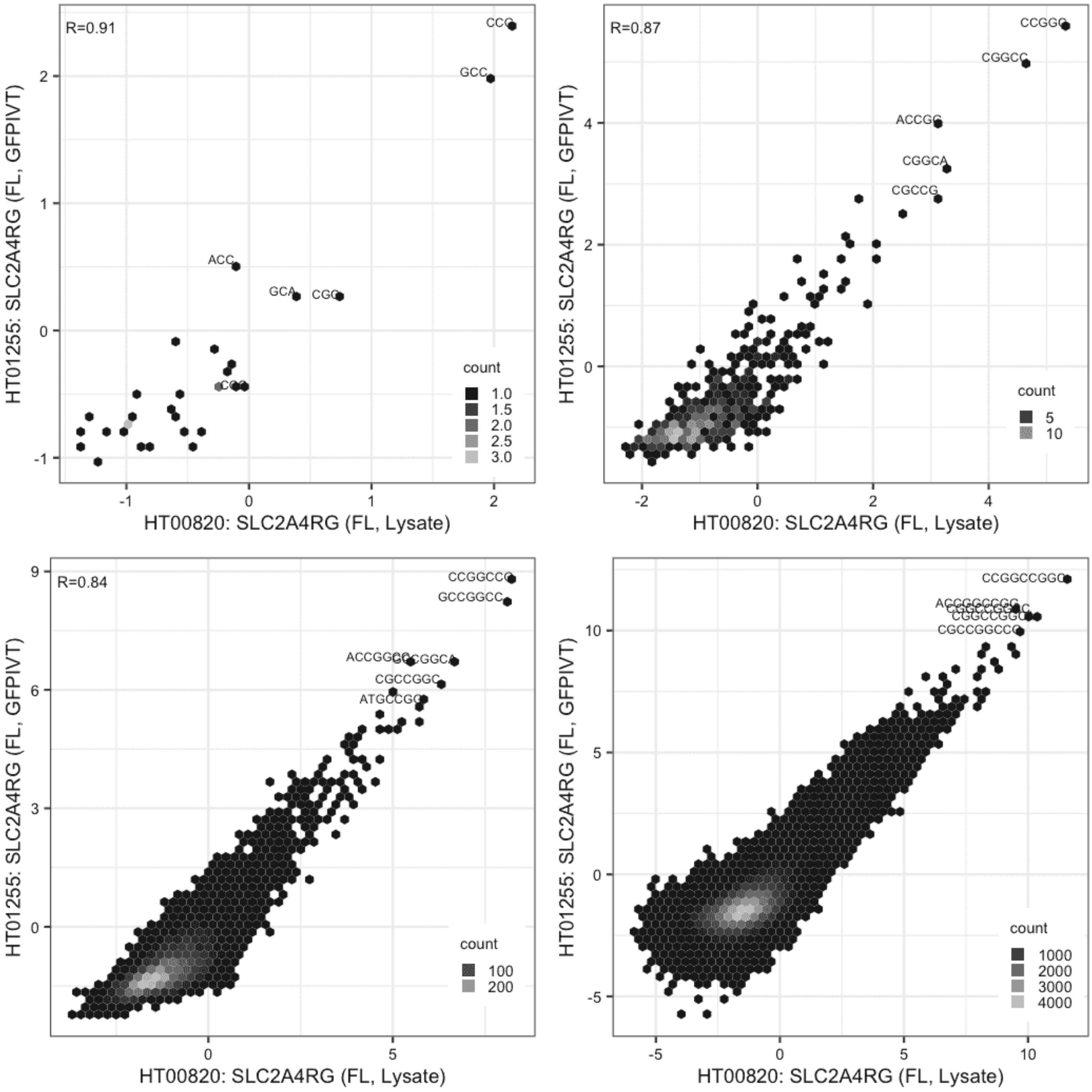

**SOX15**

Lysate vs IVT, same construct

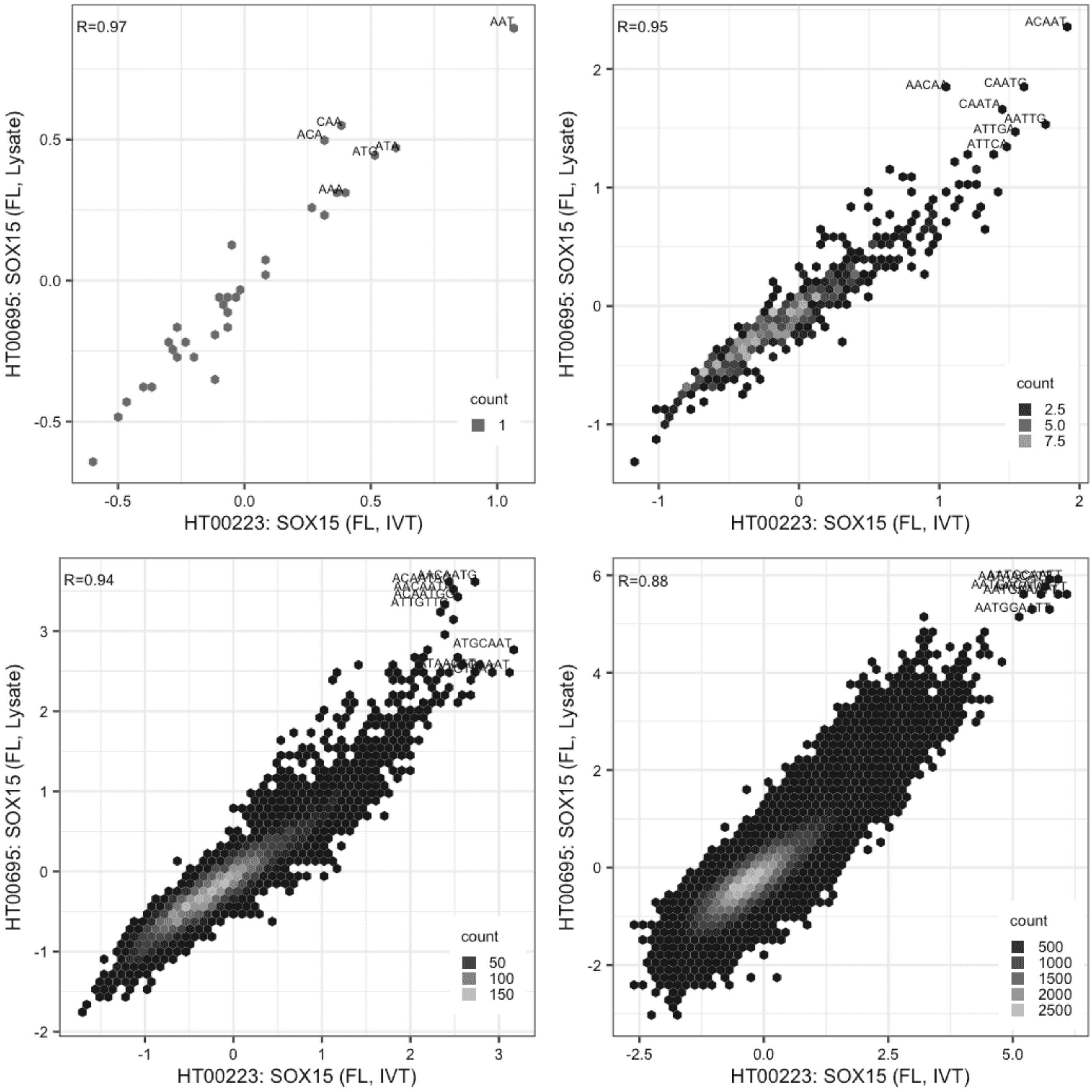

**SOX15**

Lysate vs IVT, same construct

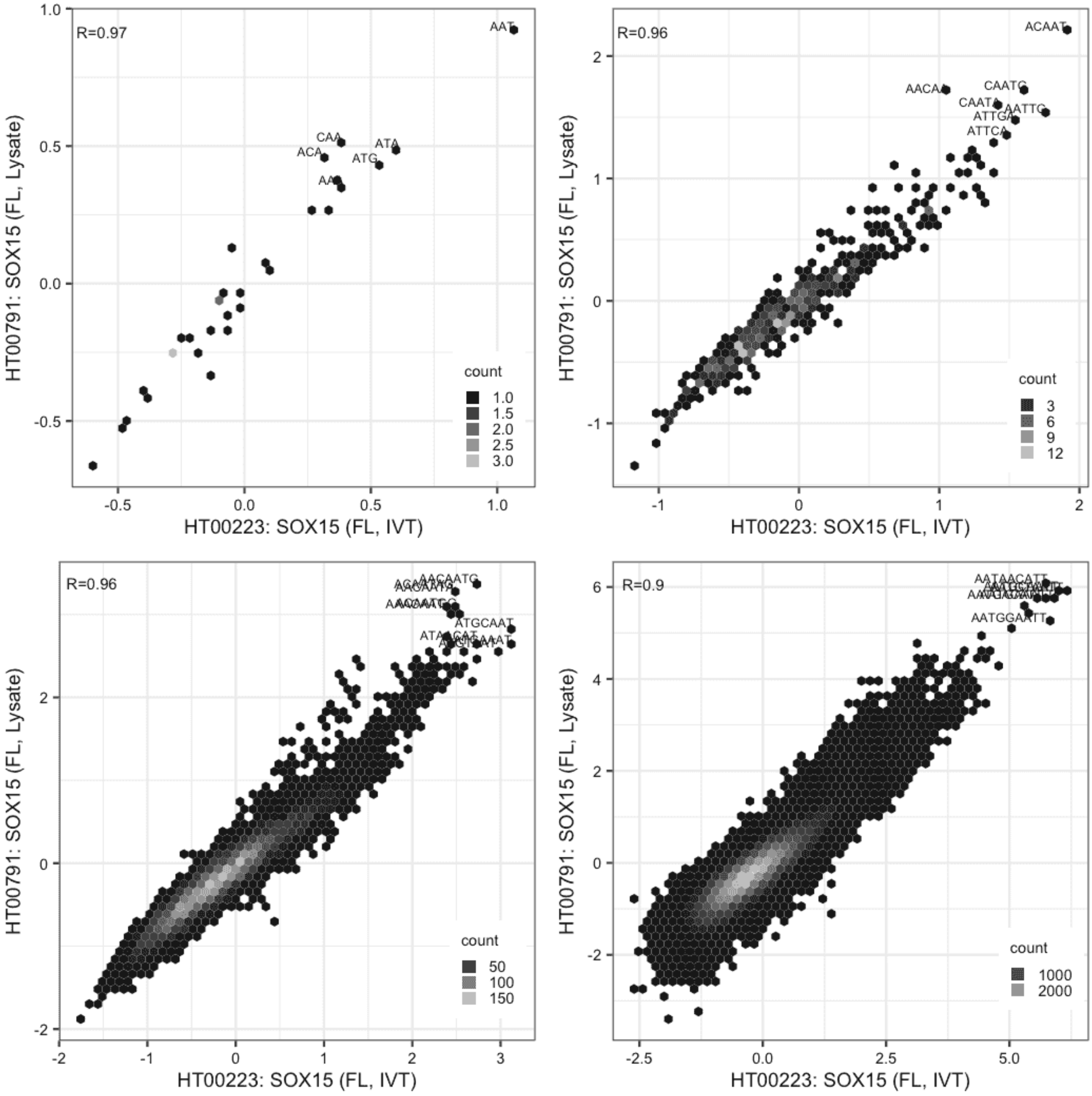

**SOX15**

Lysate vs IVT, same construct

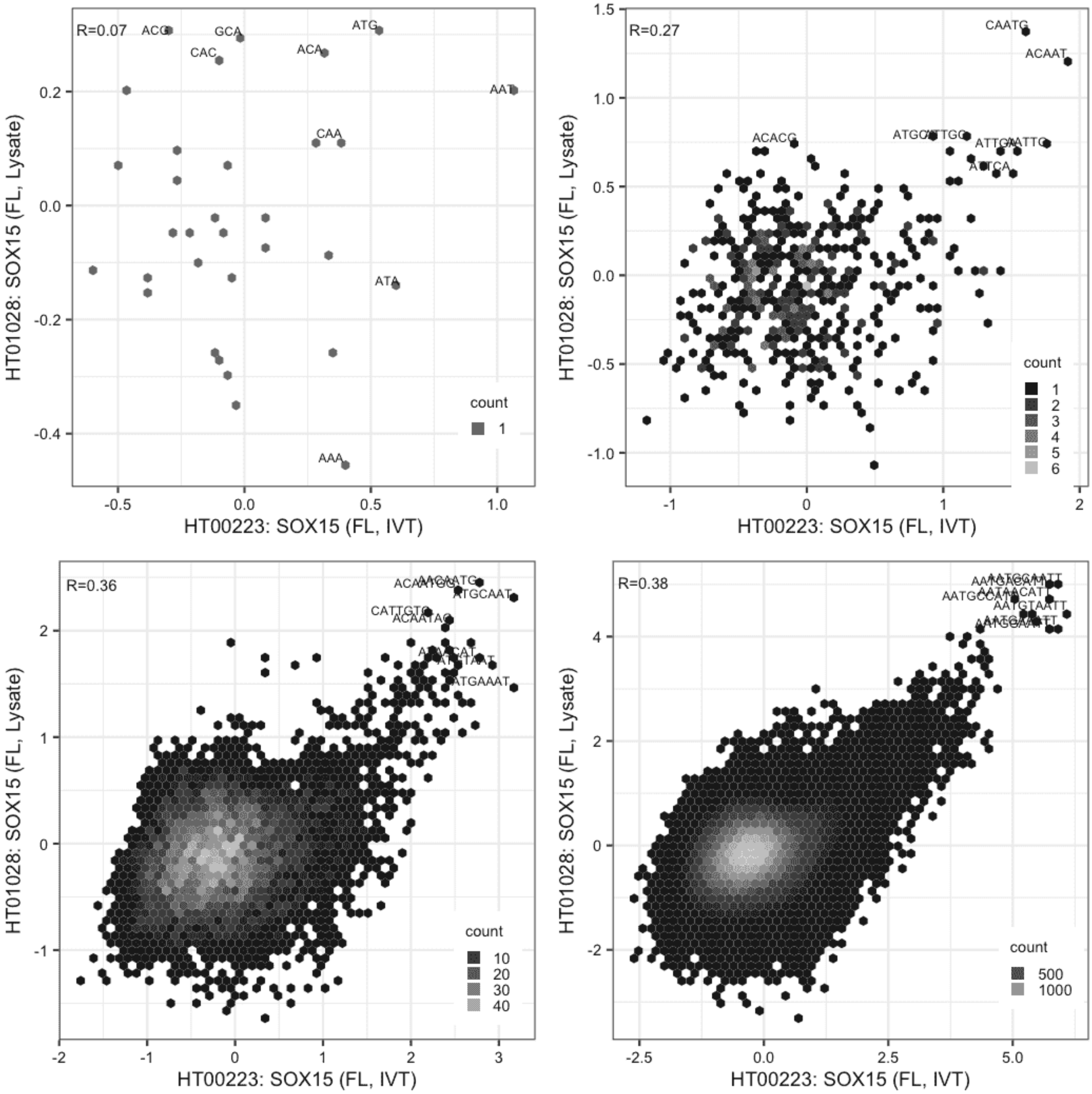

**SOX15**

Lysate vs IVT, same construct

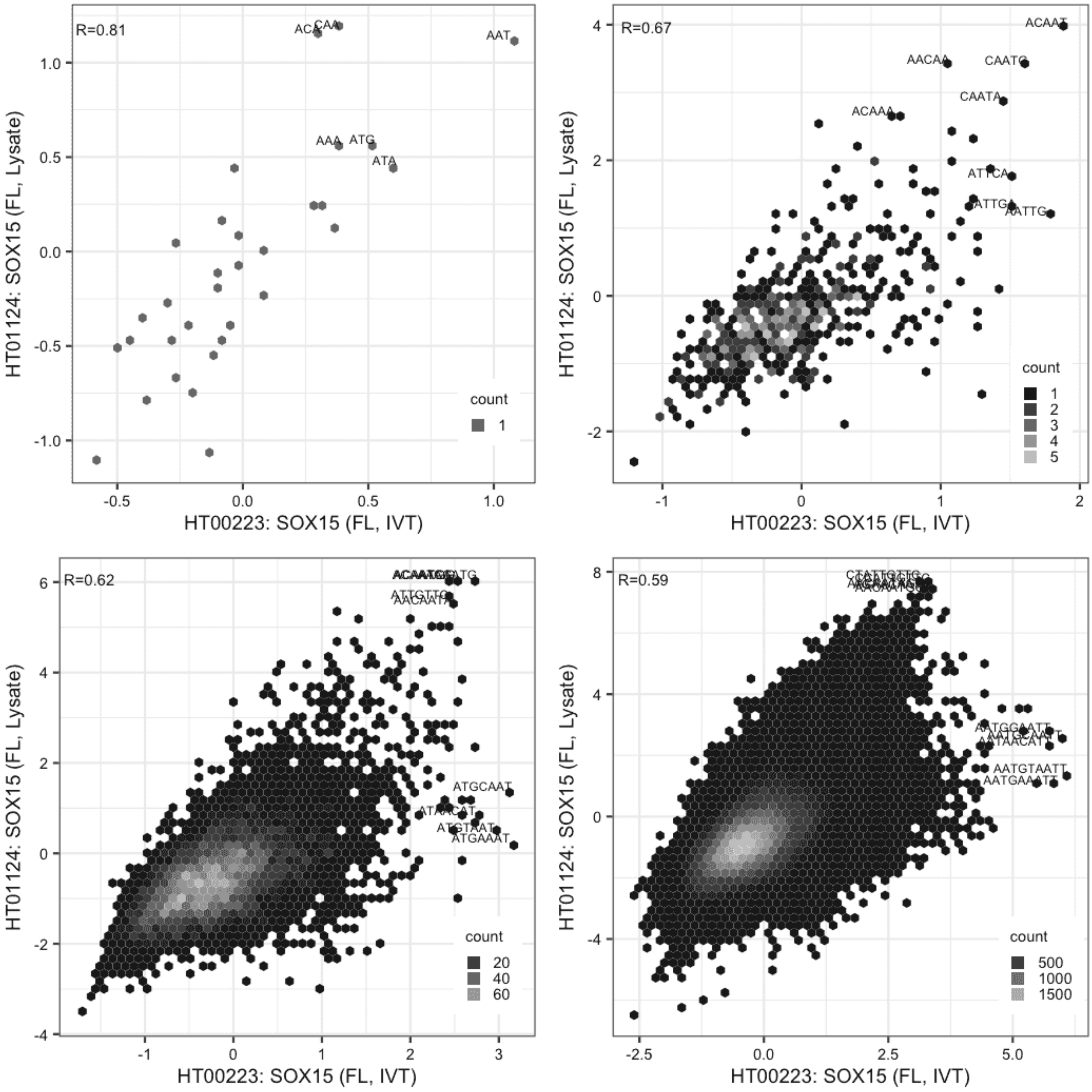

**SOX15**

Lysate vs Lysate, same construct

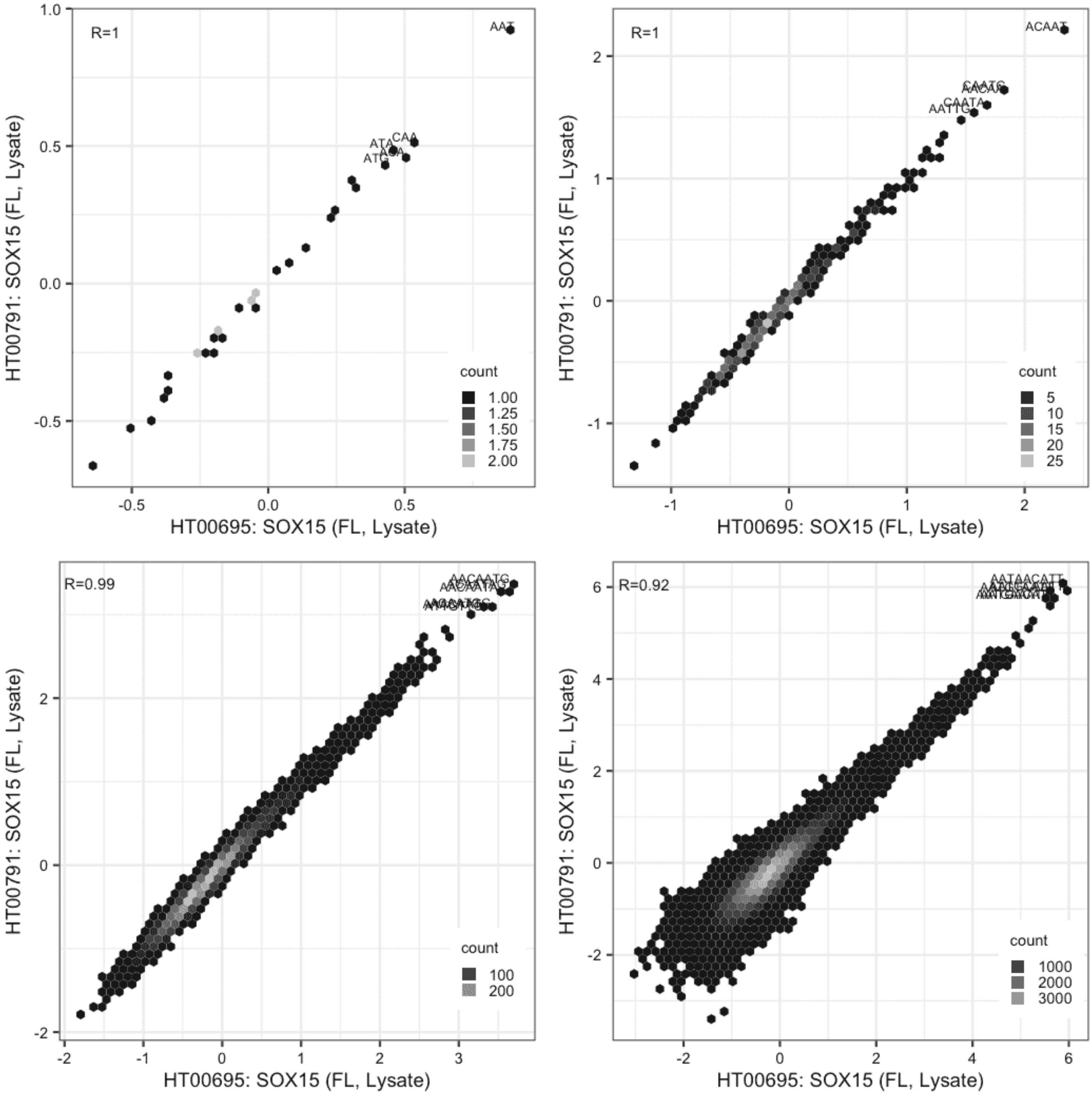

**SOX15**

Lysate vs Lysate, same construct

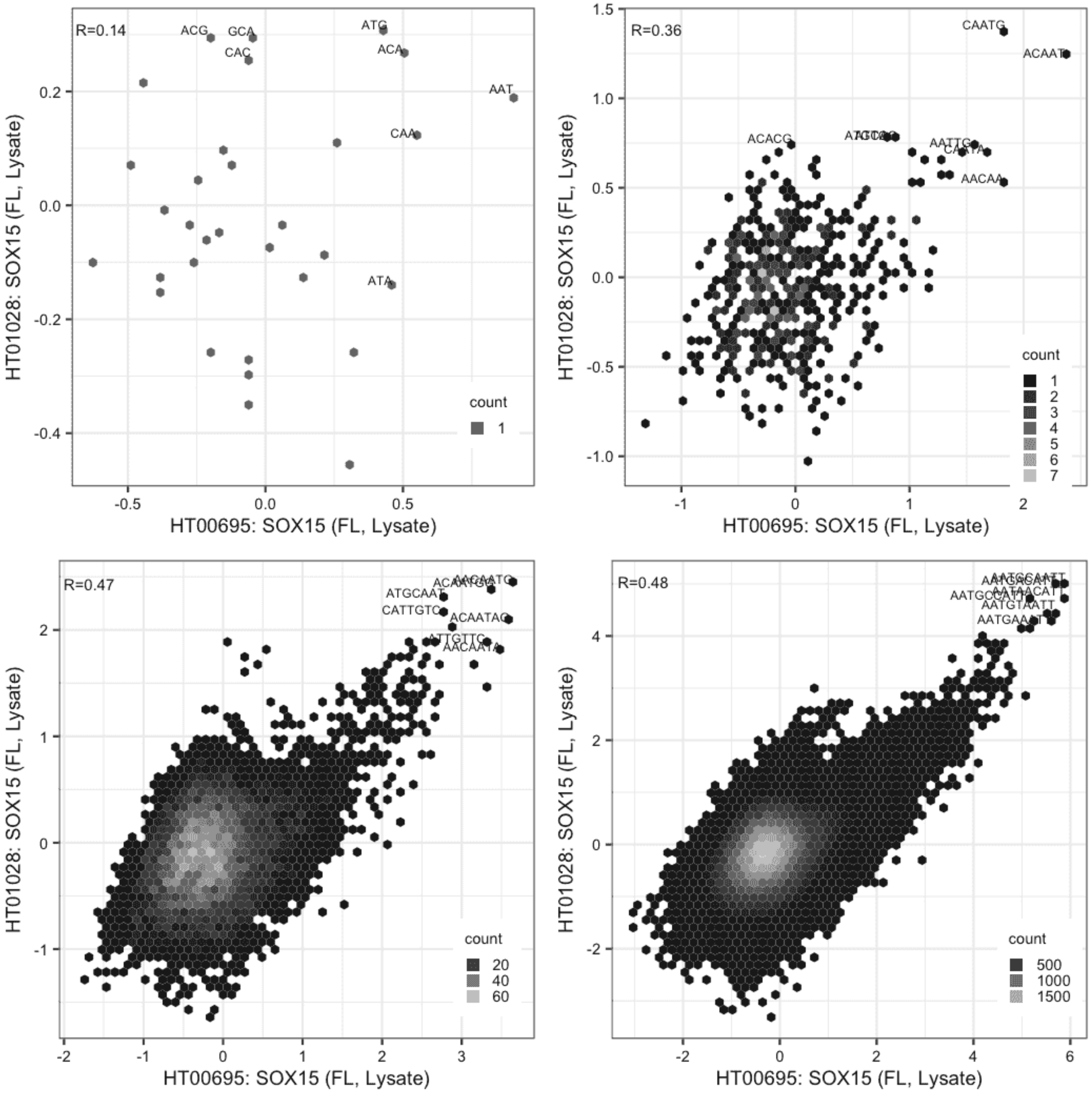

**SOX15**

Lysate vs Lysate, same construct

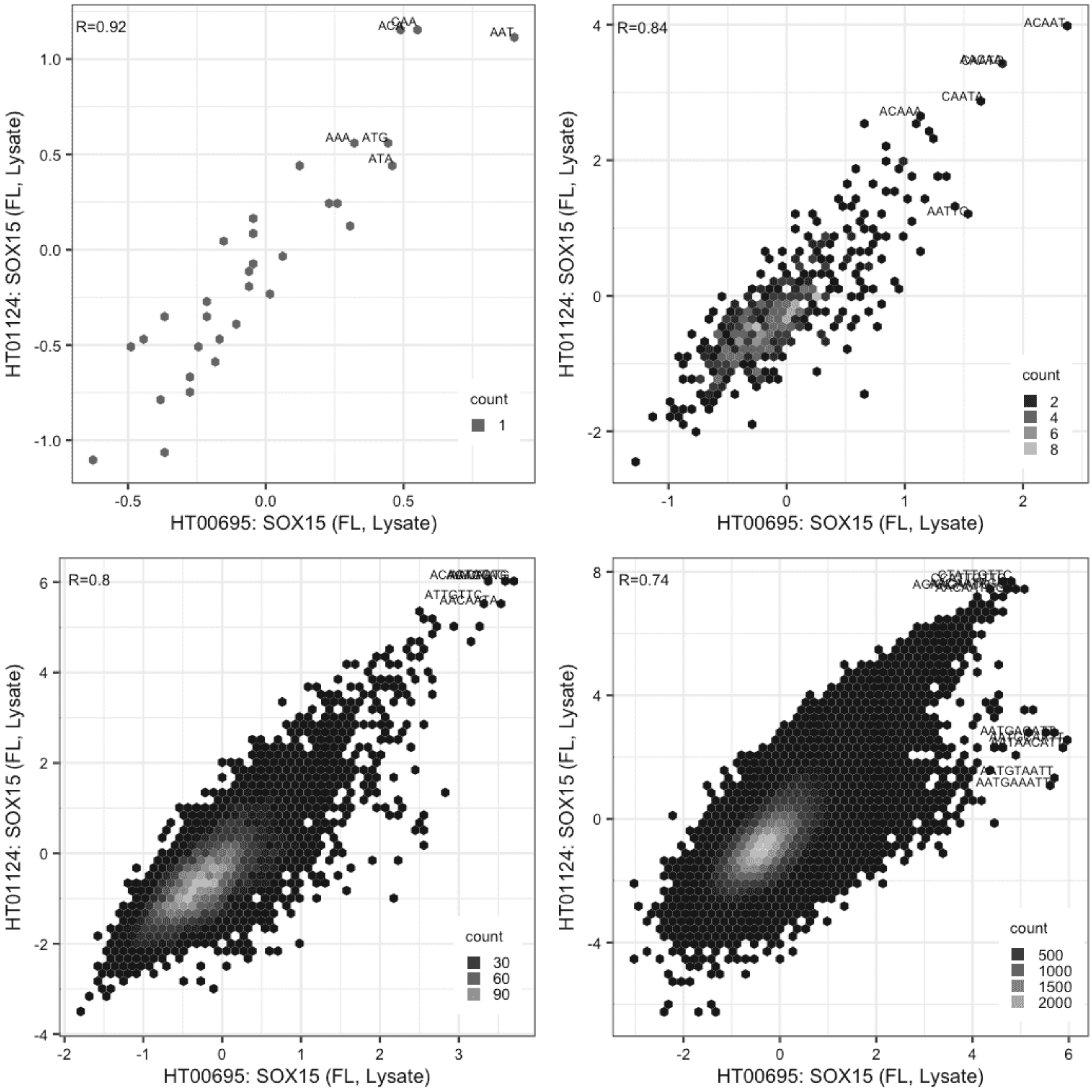

#### SOX15

Lysate vs Lysate, same construct

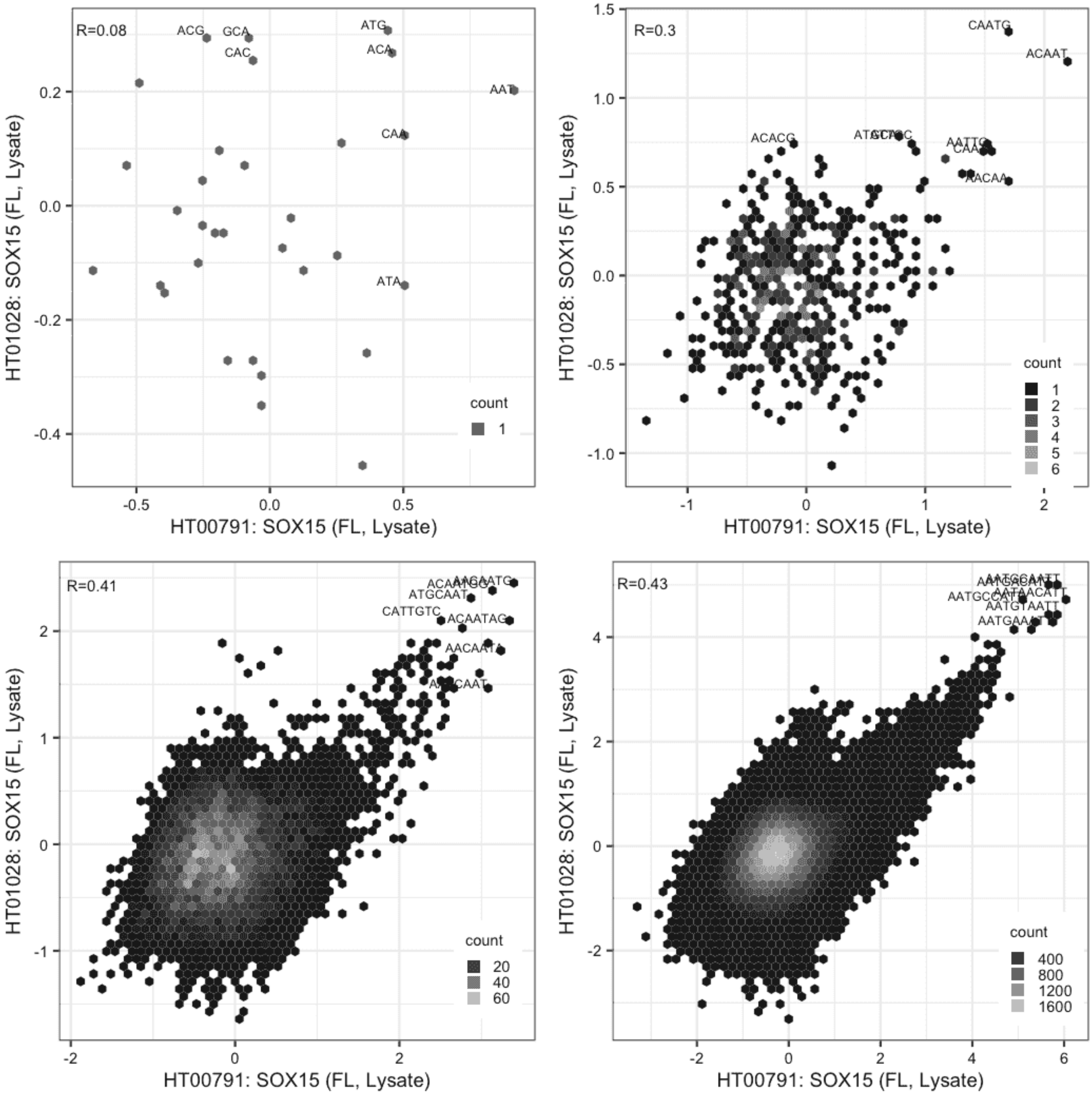

**SOX15**

Lysate vs Lysate, same construct

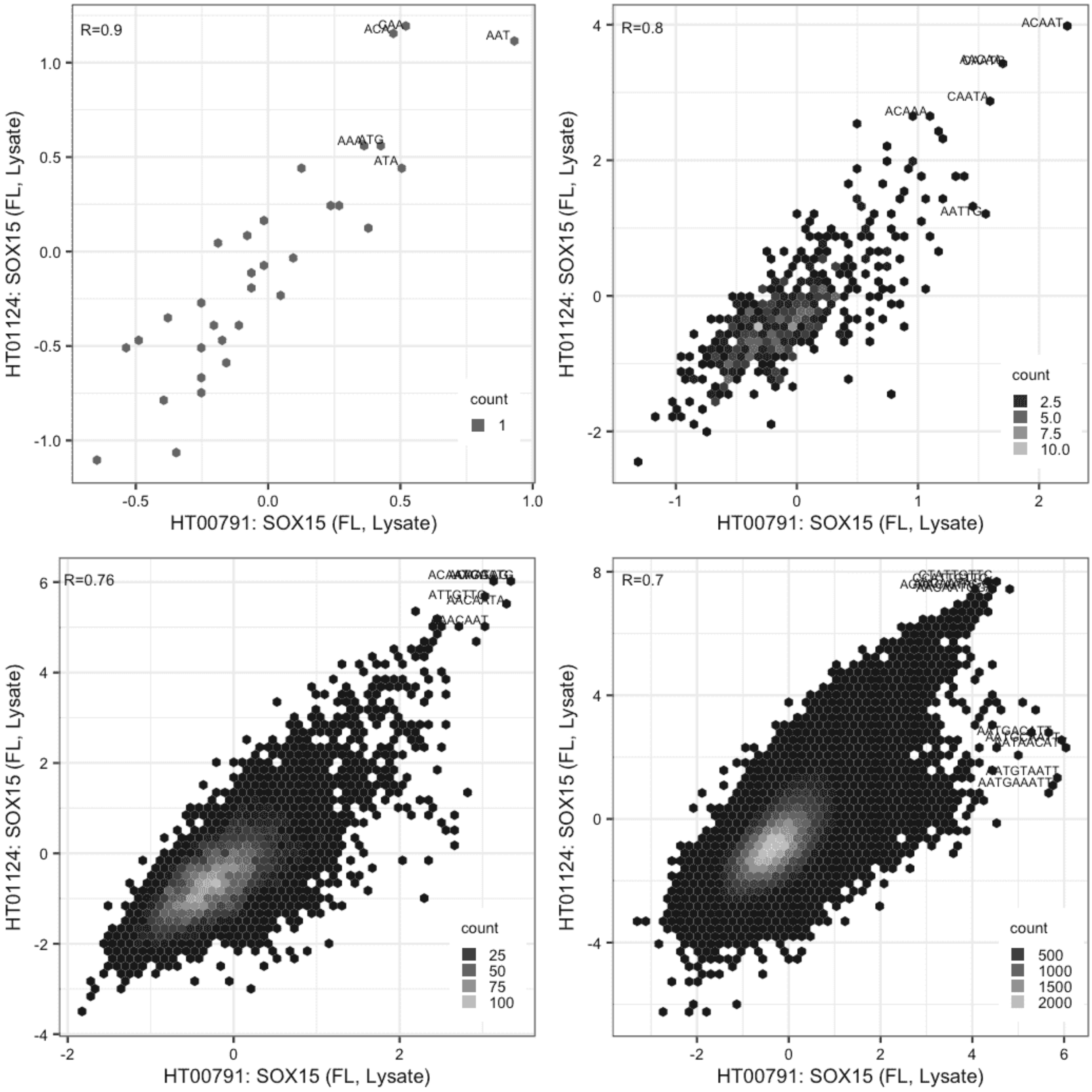

#### SOX15

Lysate vs Lysate, same construct

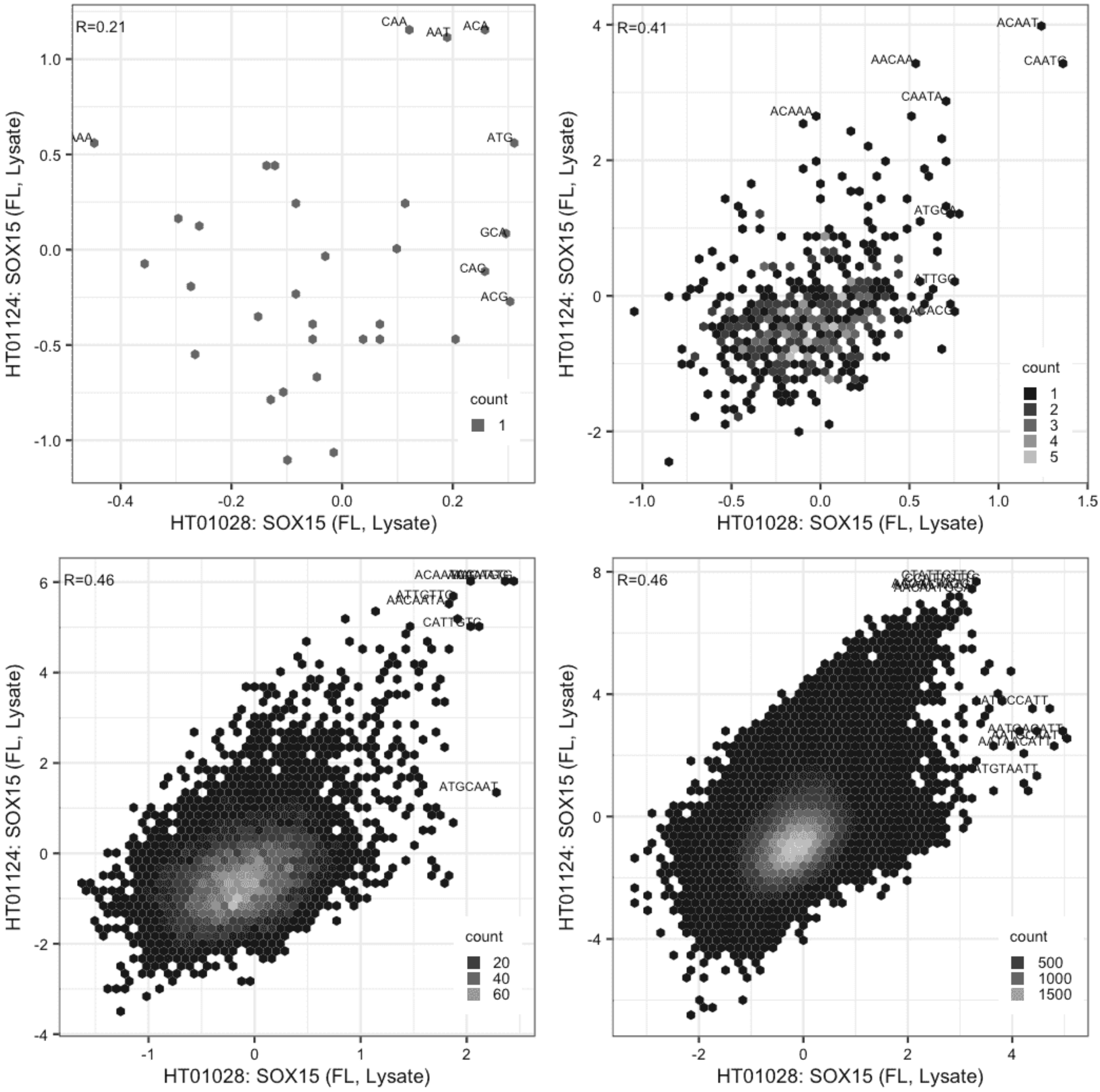

#### SOX2

IVT vs IVT, same construct

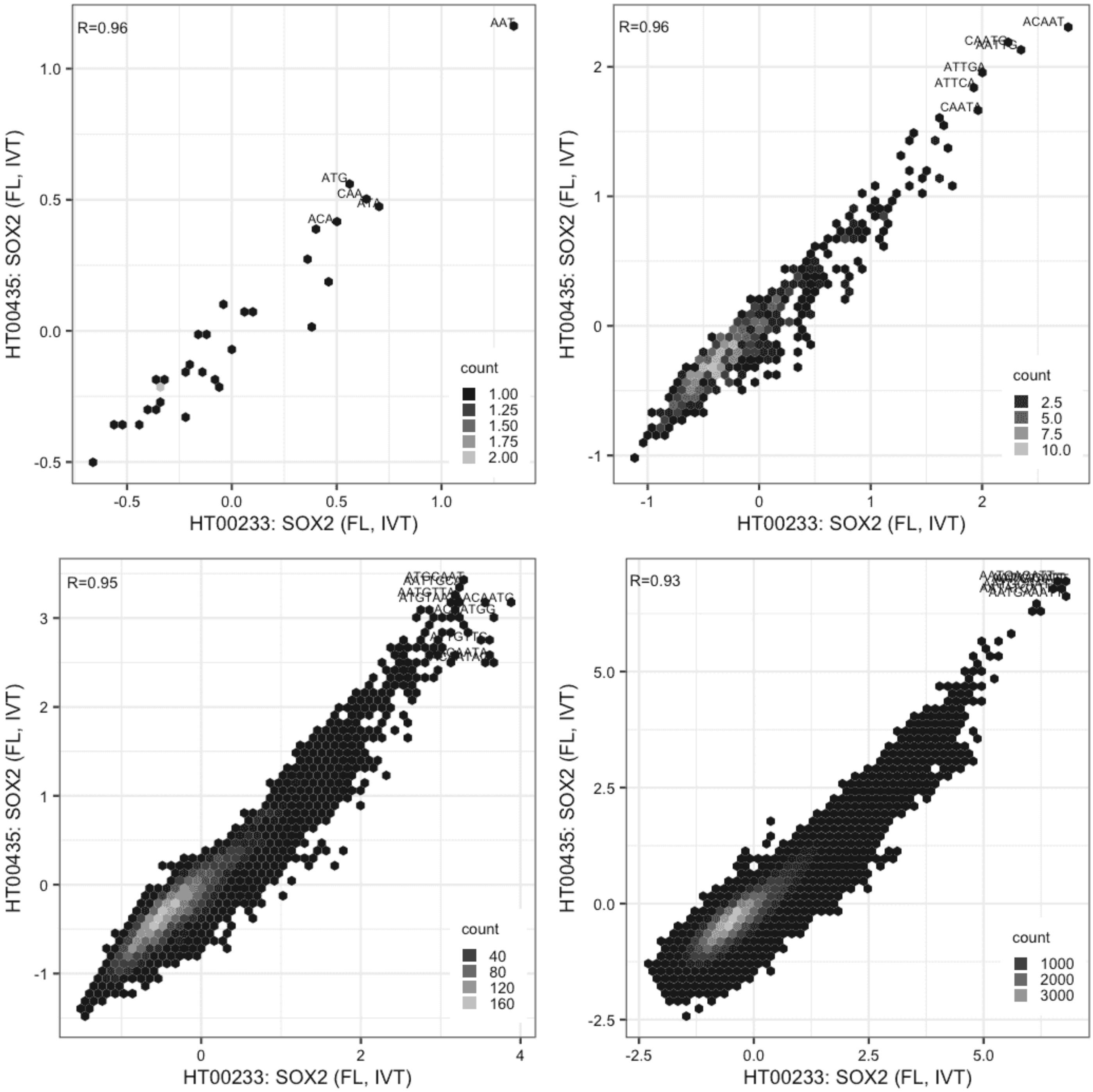

#### SOX2

Lysate vs IVT, same construct

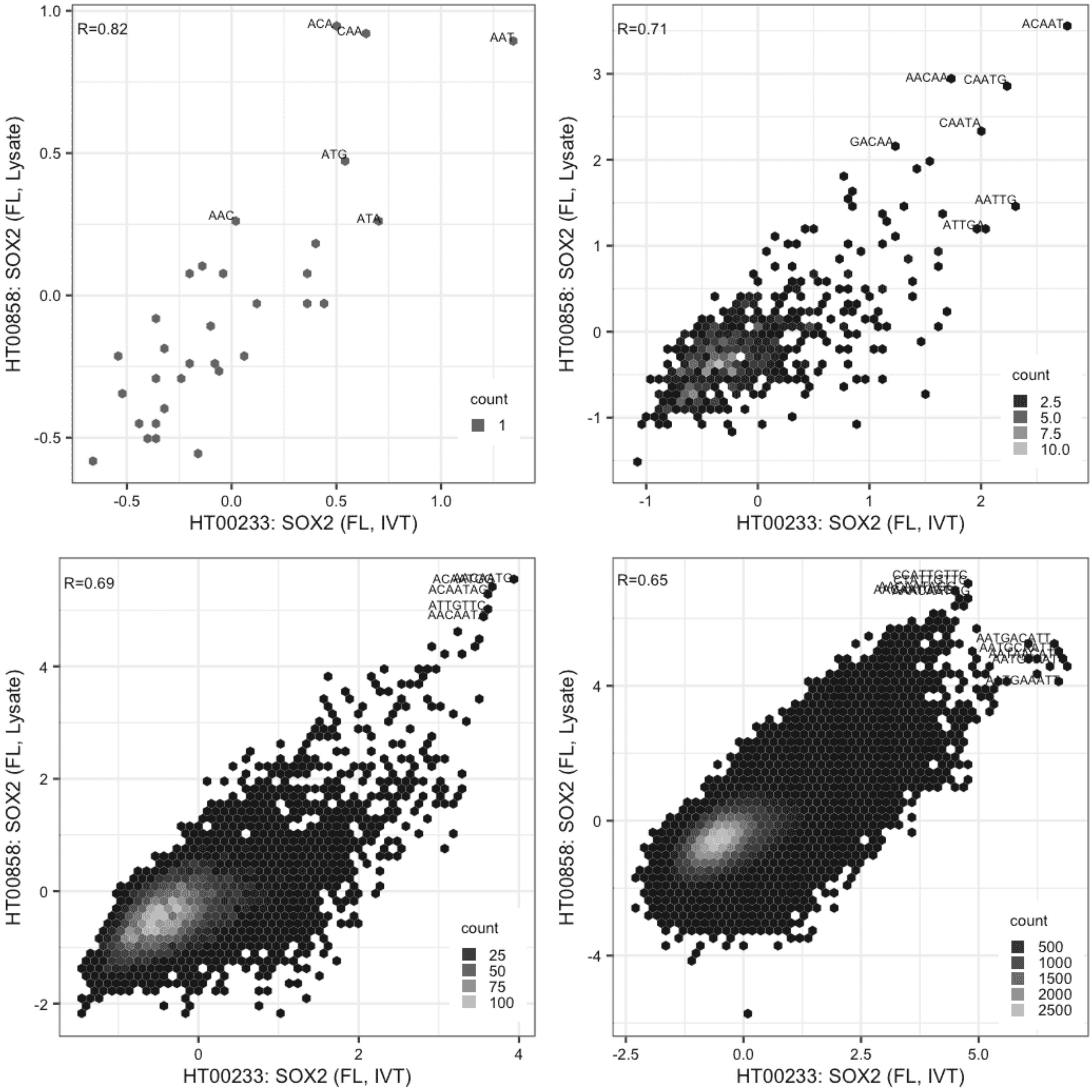

#### SOX2

Lysate vs IVT, same construct

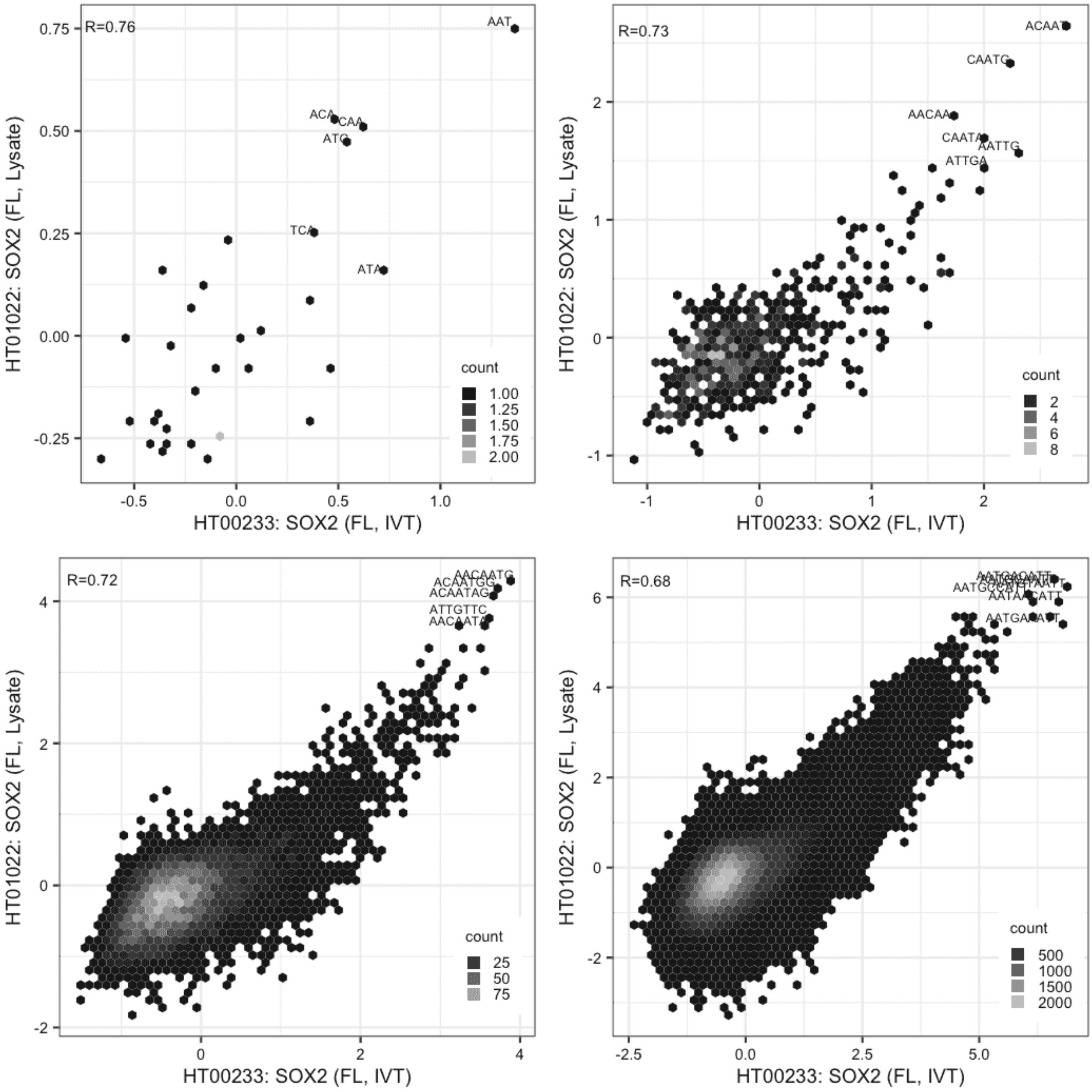

**SOX2**

Lysate vs IVT, same construct

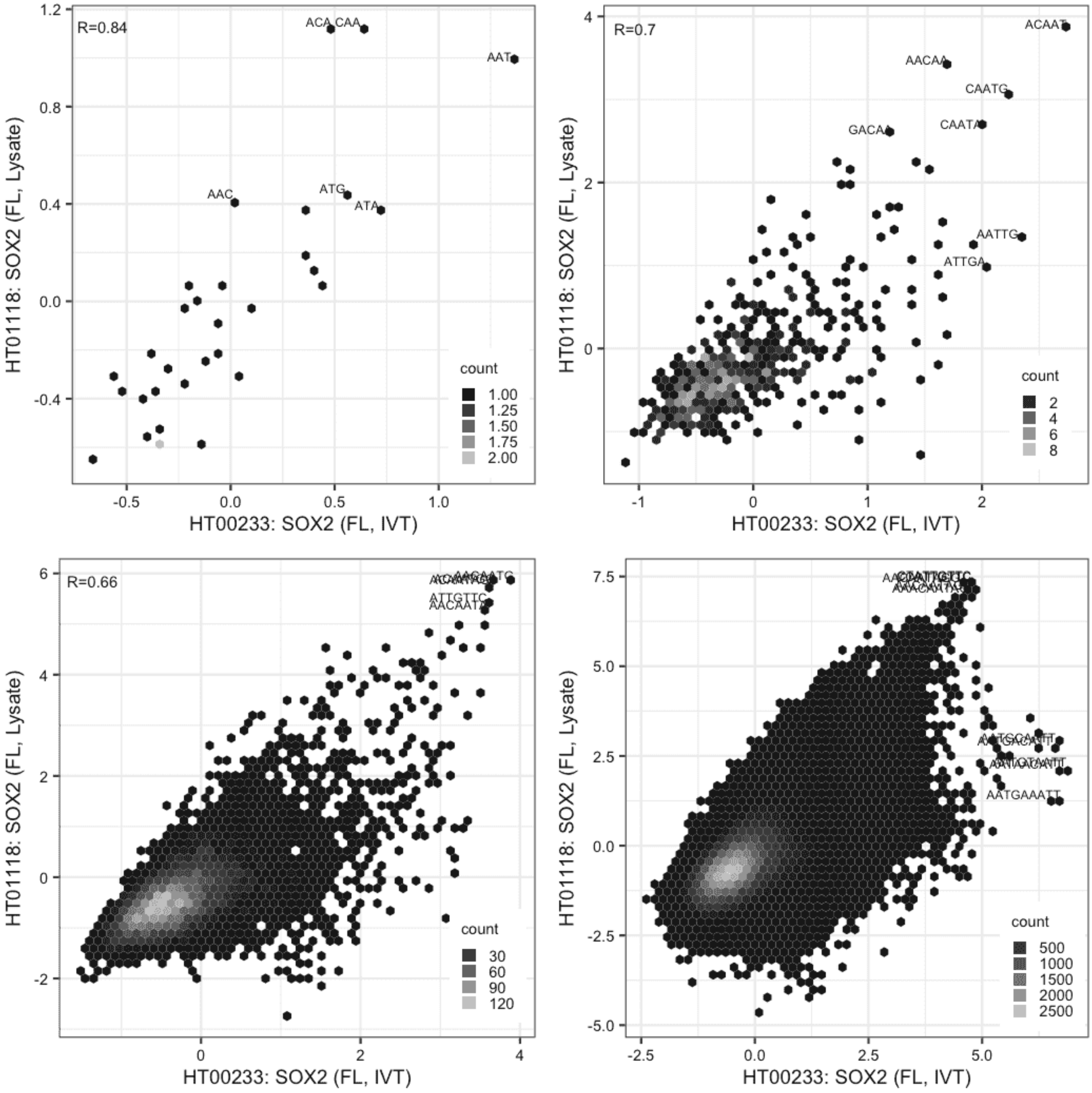

#### SOX2

Lysate vs IVT, same construct

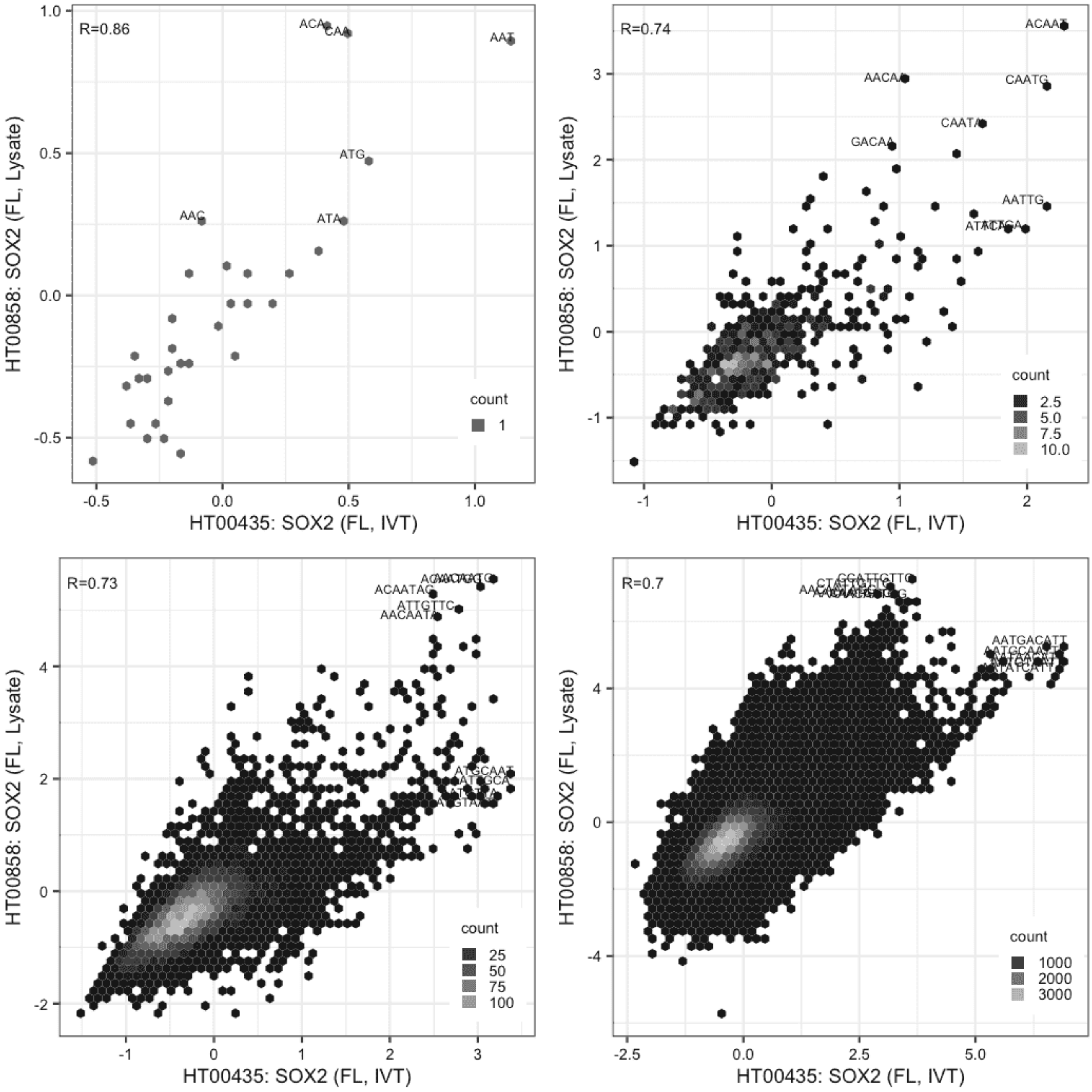

**SOX2**

Lysate vs IVT, same construct

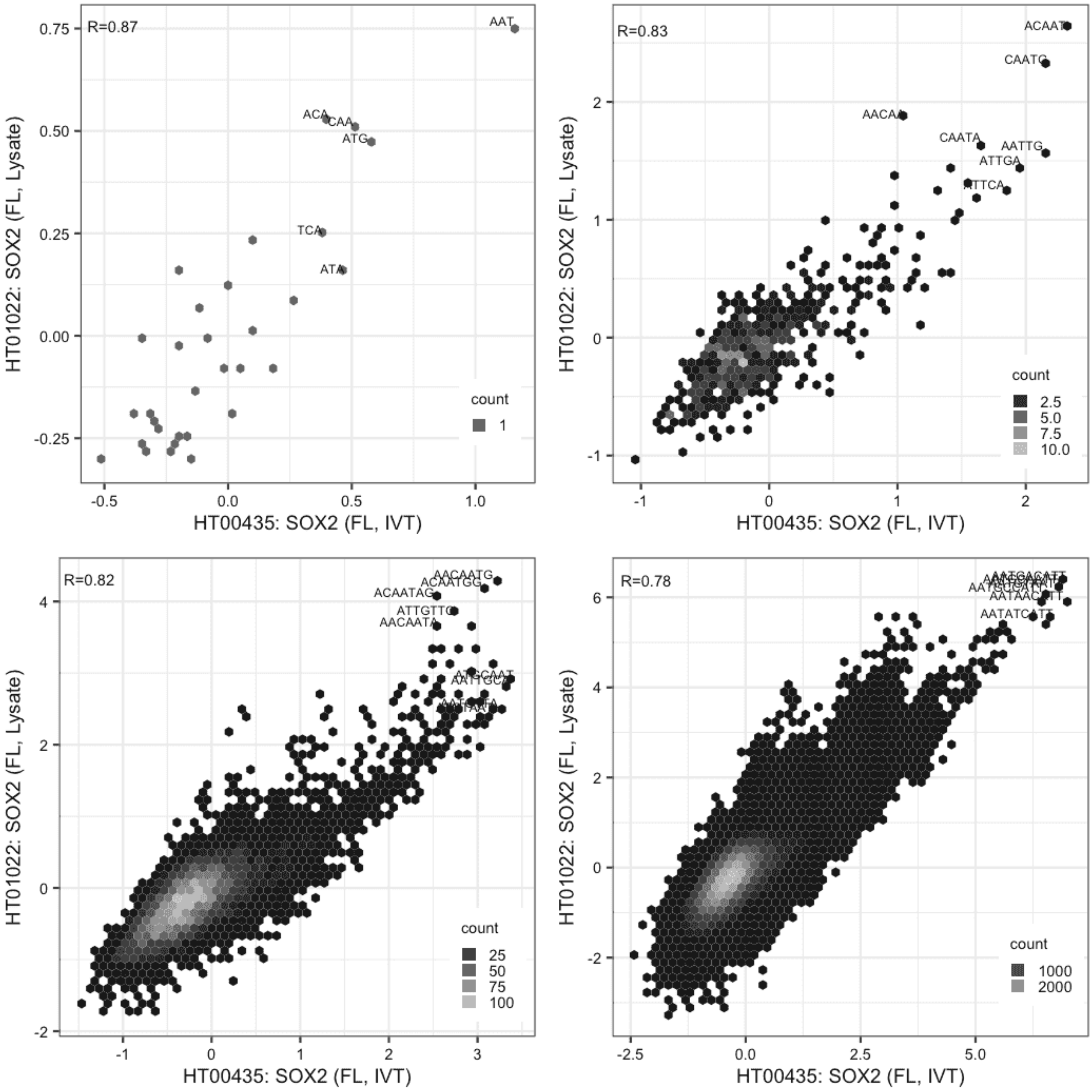

#### SOX2

Lysate vs IVT, same construct

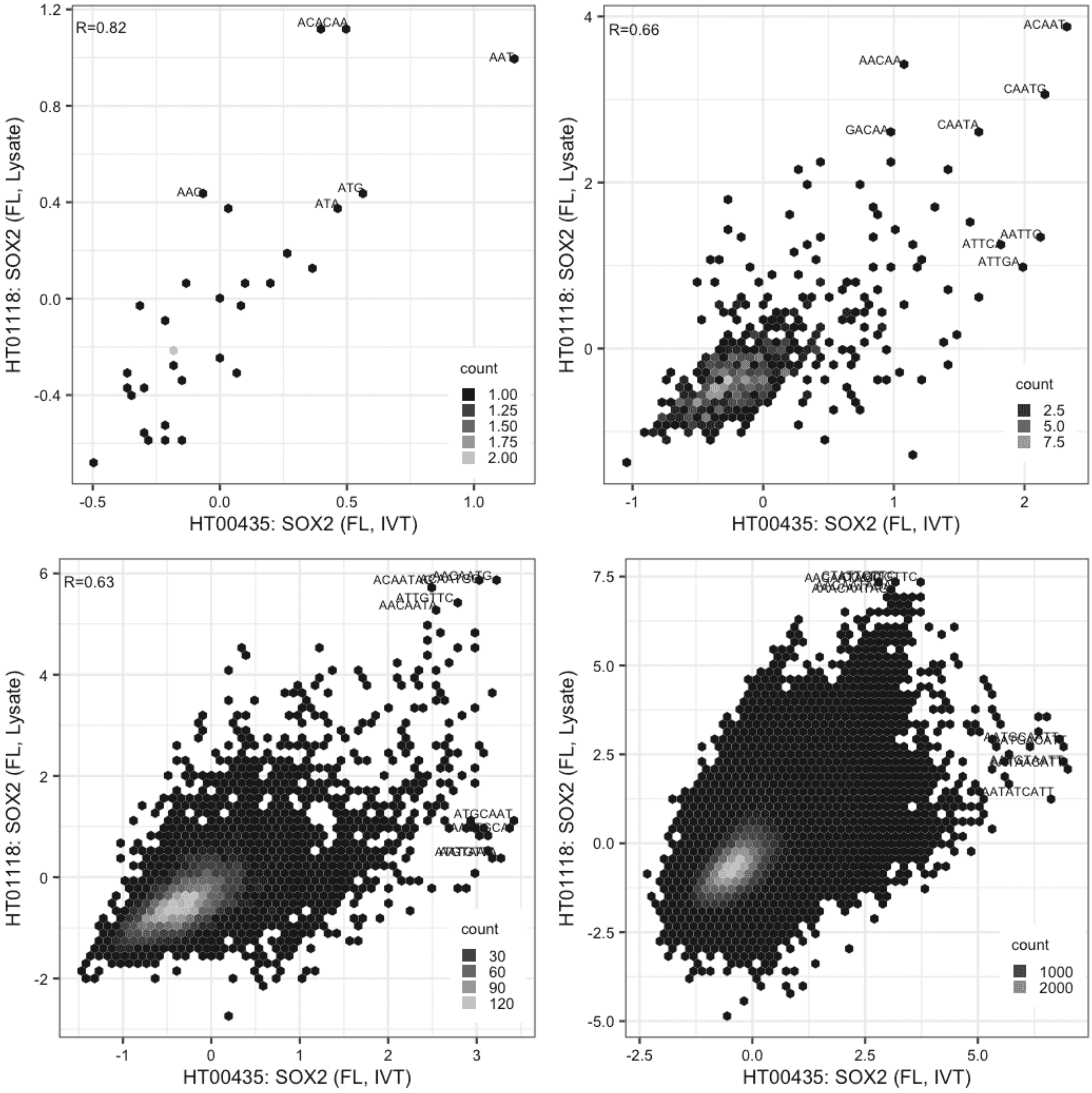

#### SOX2

Lysate vs Lysate, same construct

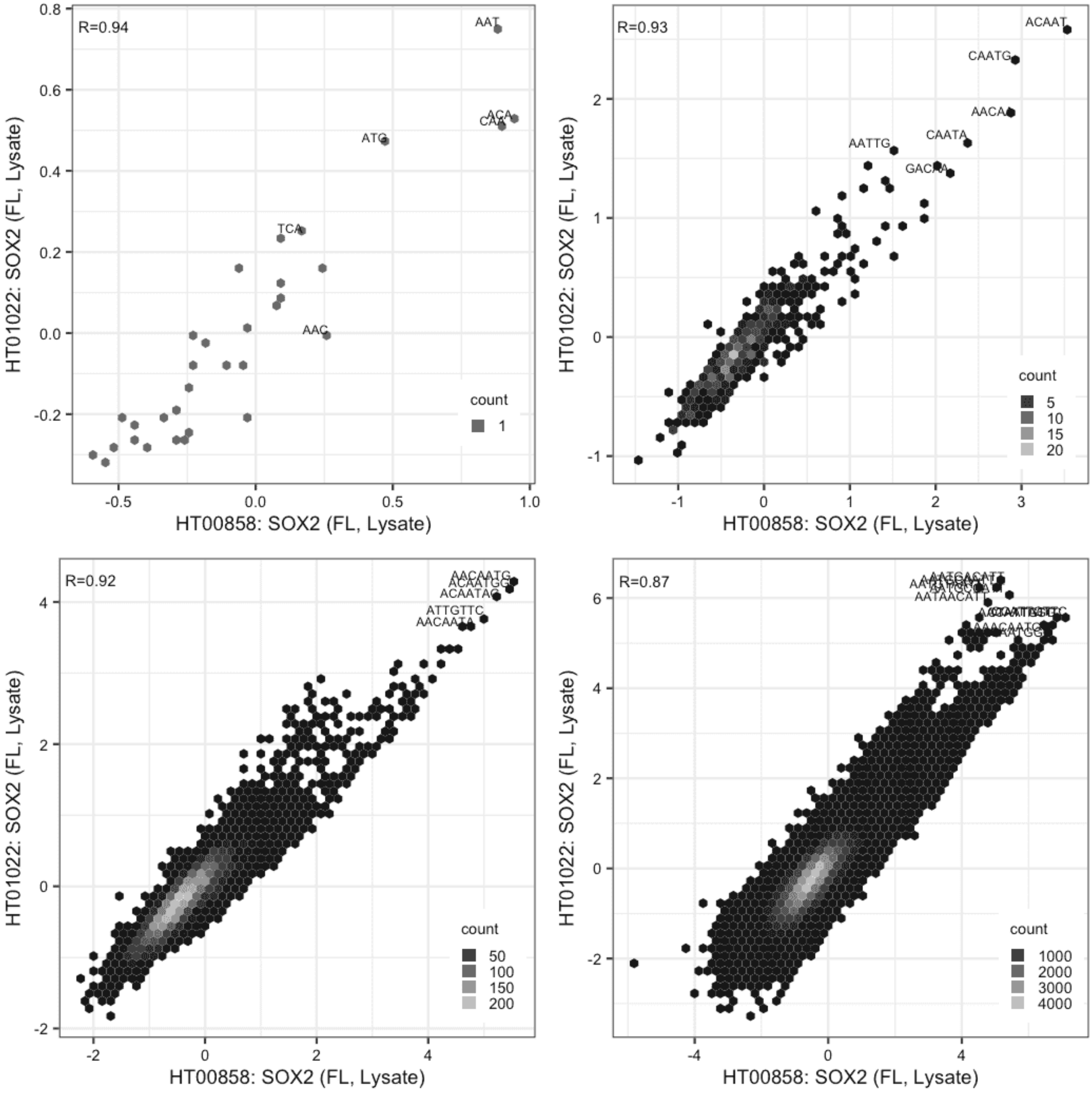

**SOX2**

Lysate vs Lysate, same construct

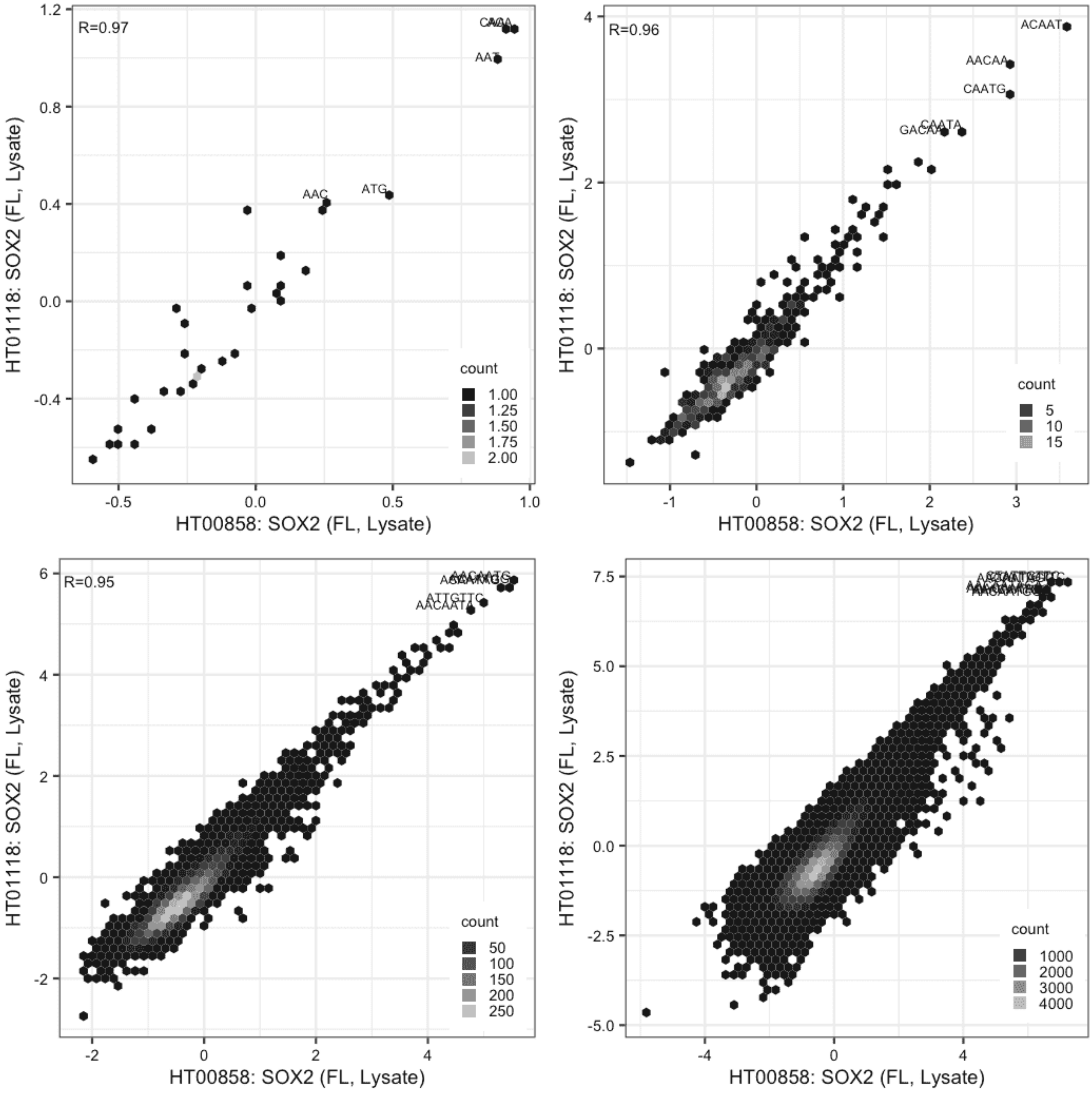

### SOX2

Lysate vs Lysate, same construct

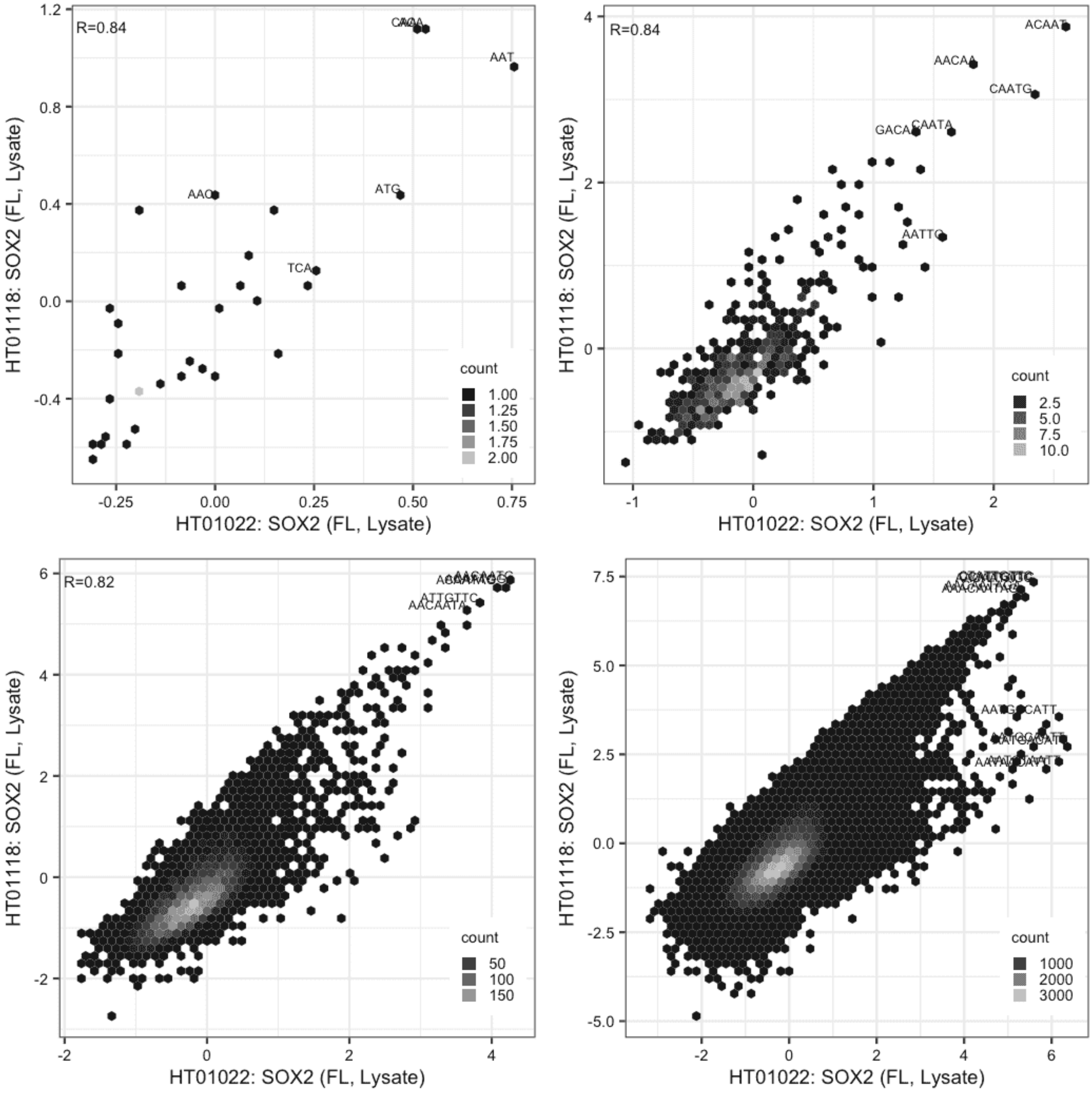

**SP140L**

IVT vs IVT, different constructs

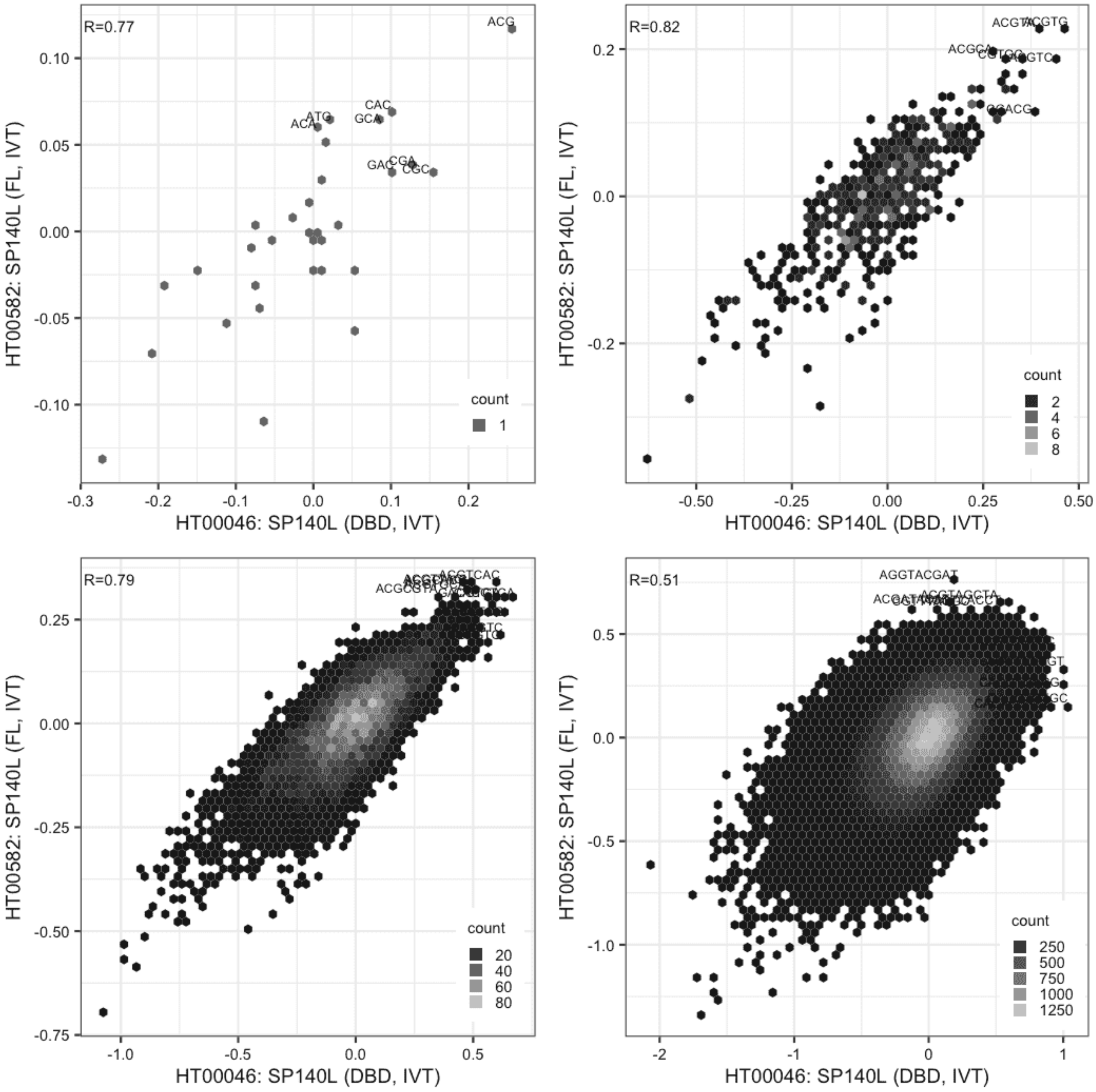

**SP140L**

Lysate vs IVT, different constructs

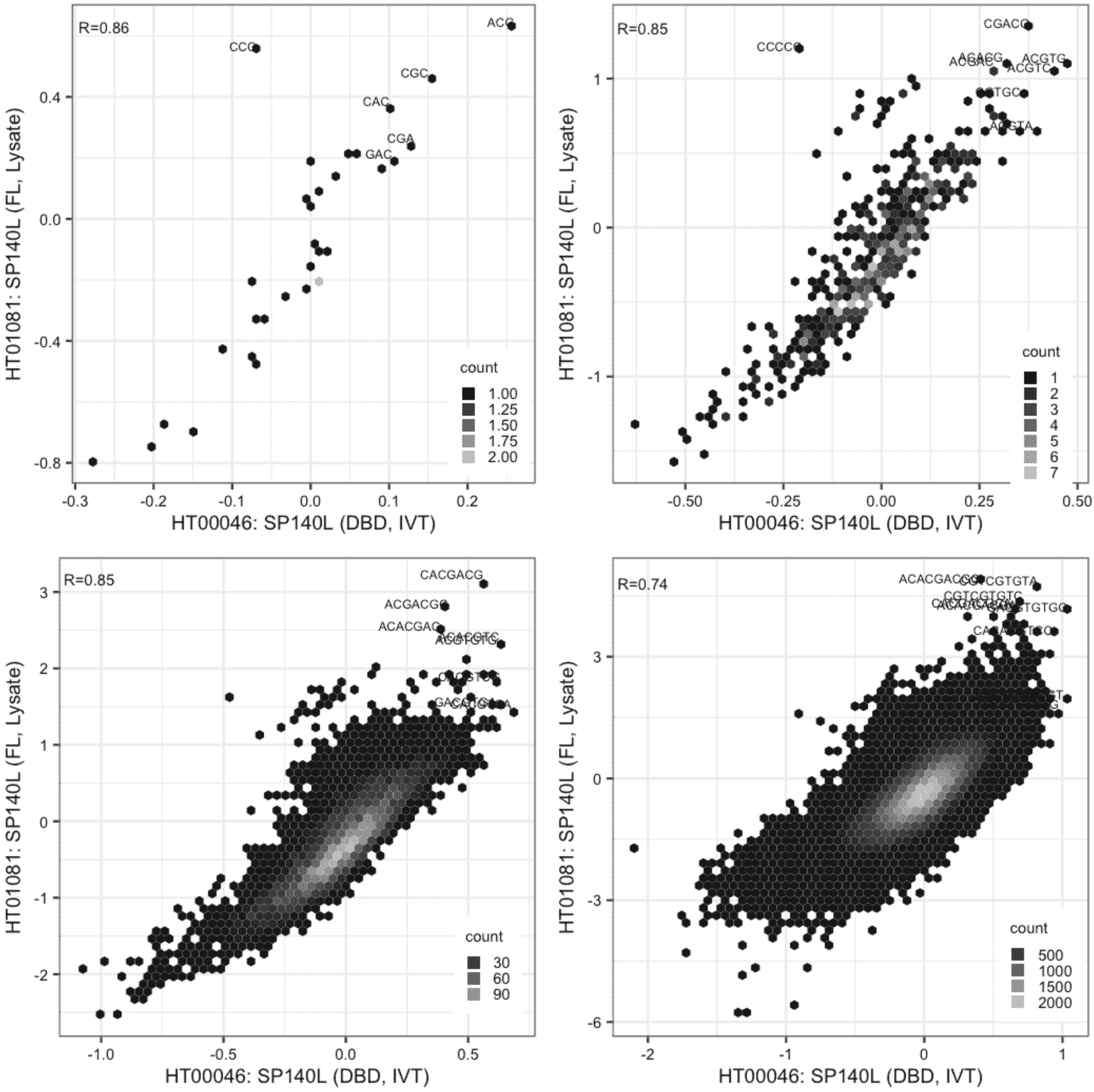

**SP140L**

Lysate vs IVT, different constructs

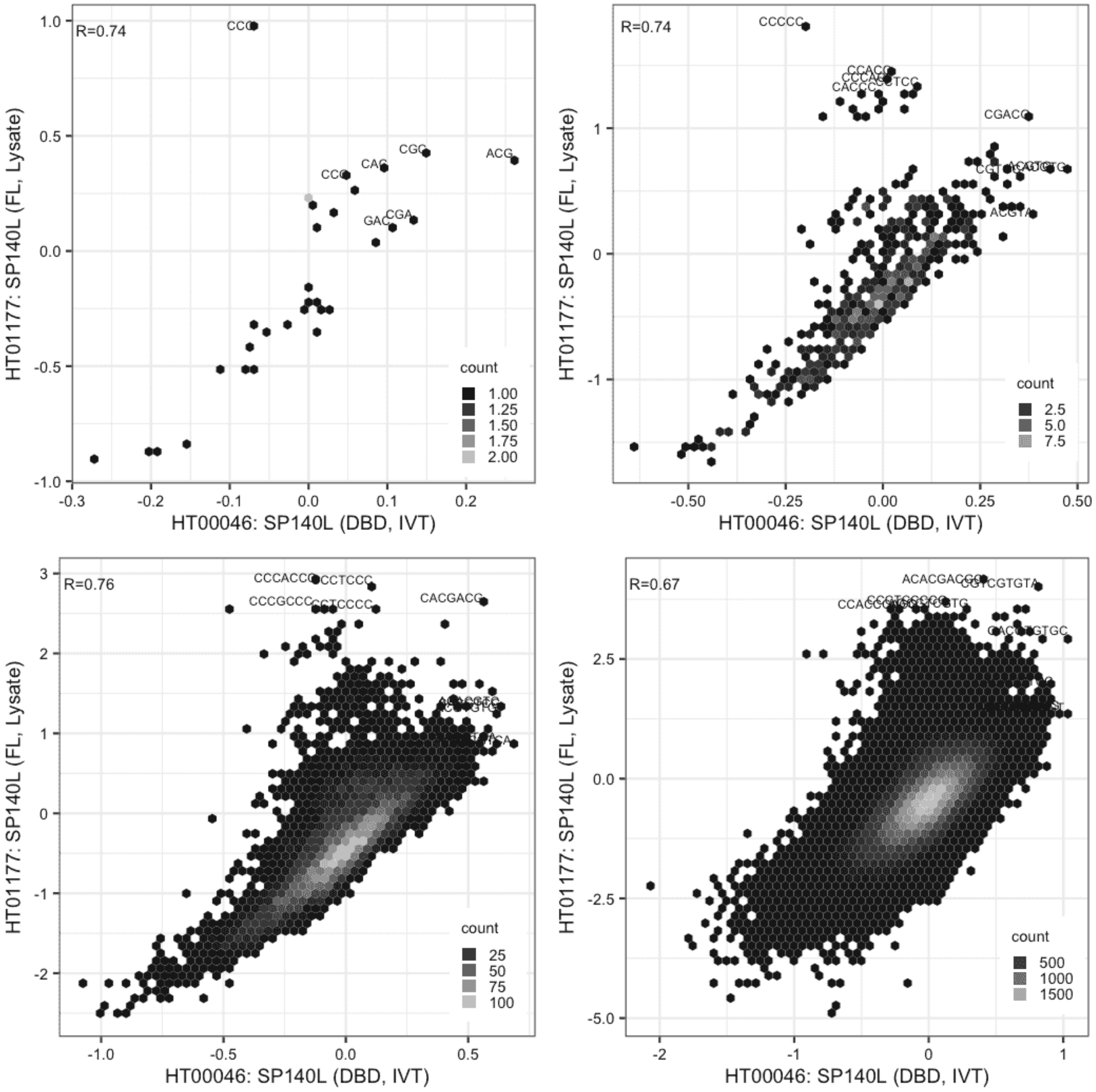

**SP140L**

IVT vs GFPIVT, different constructs

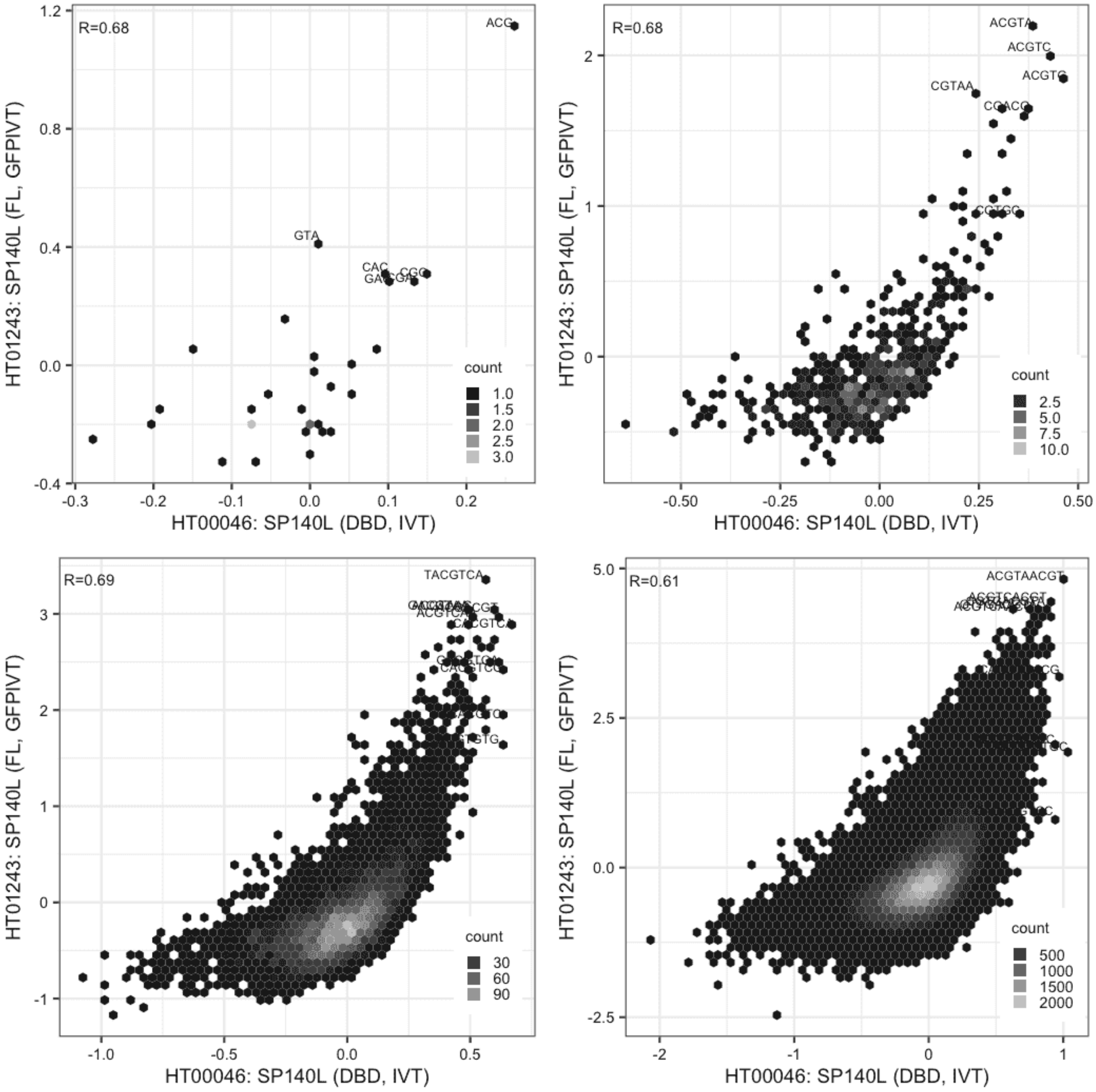

**SP140L**

IVT vs GFPIVT, same construct

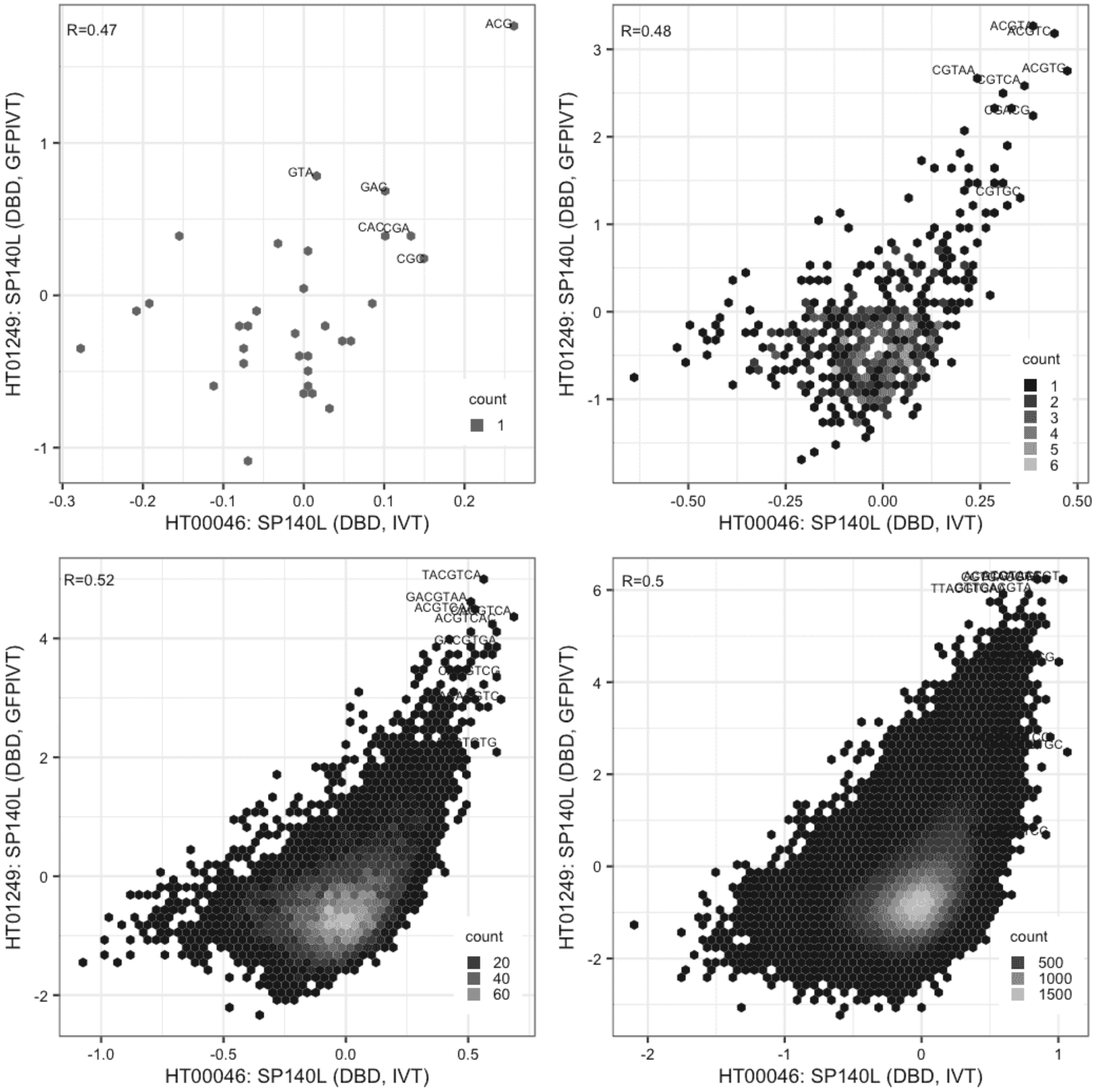

**SP140L**

Lysate vs IVT, same construct

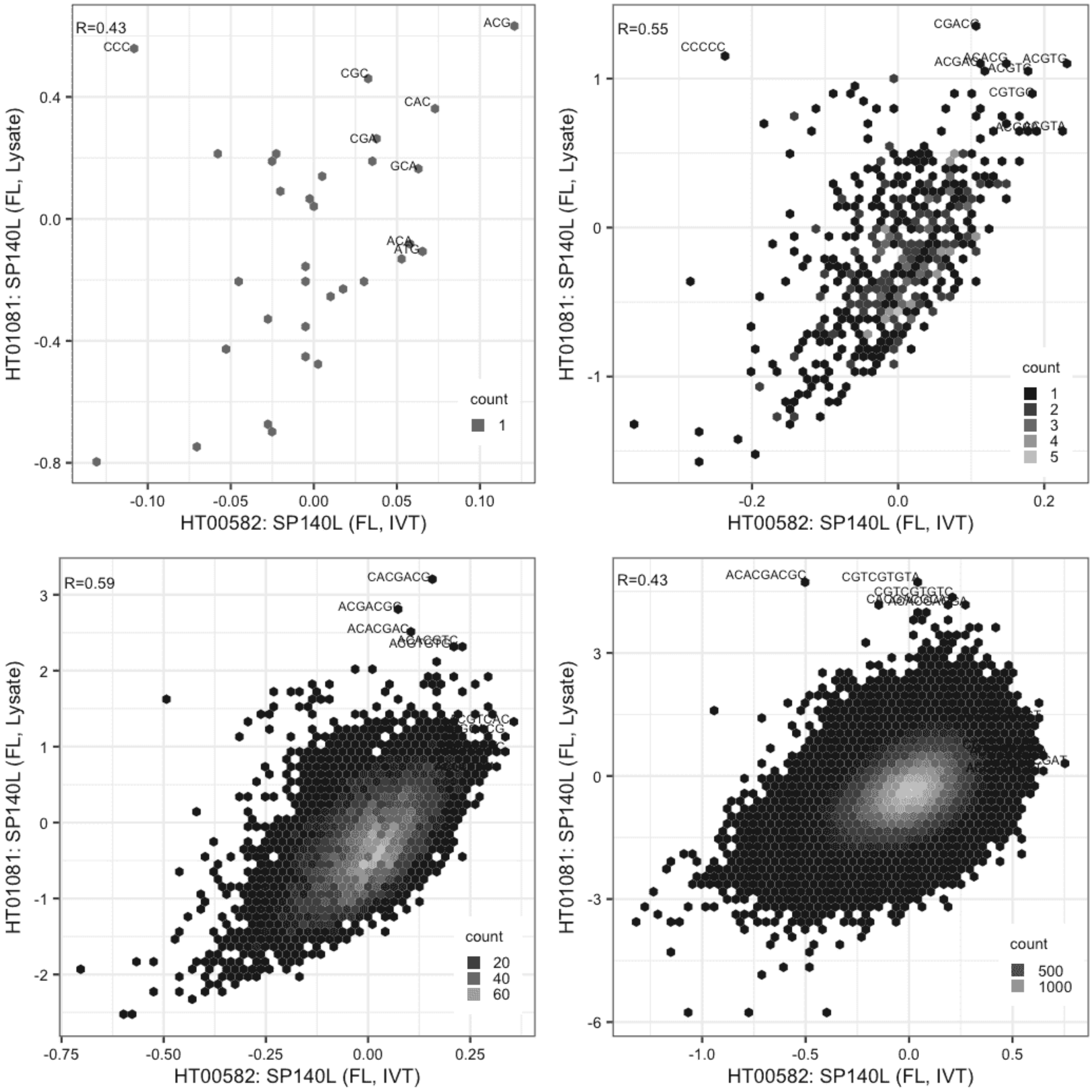

**SP140L**

Lysate vs IVT, same construct

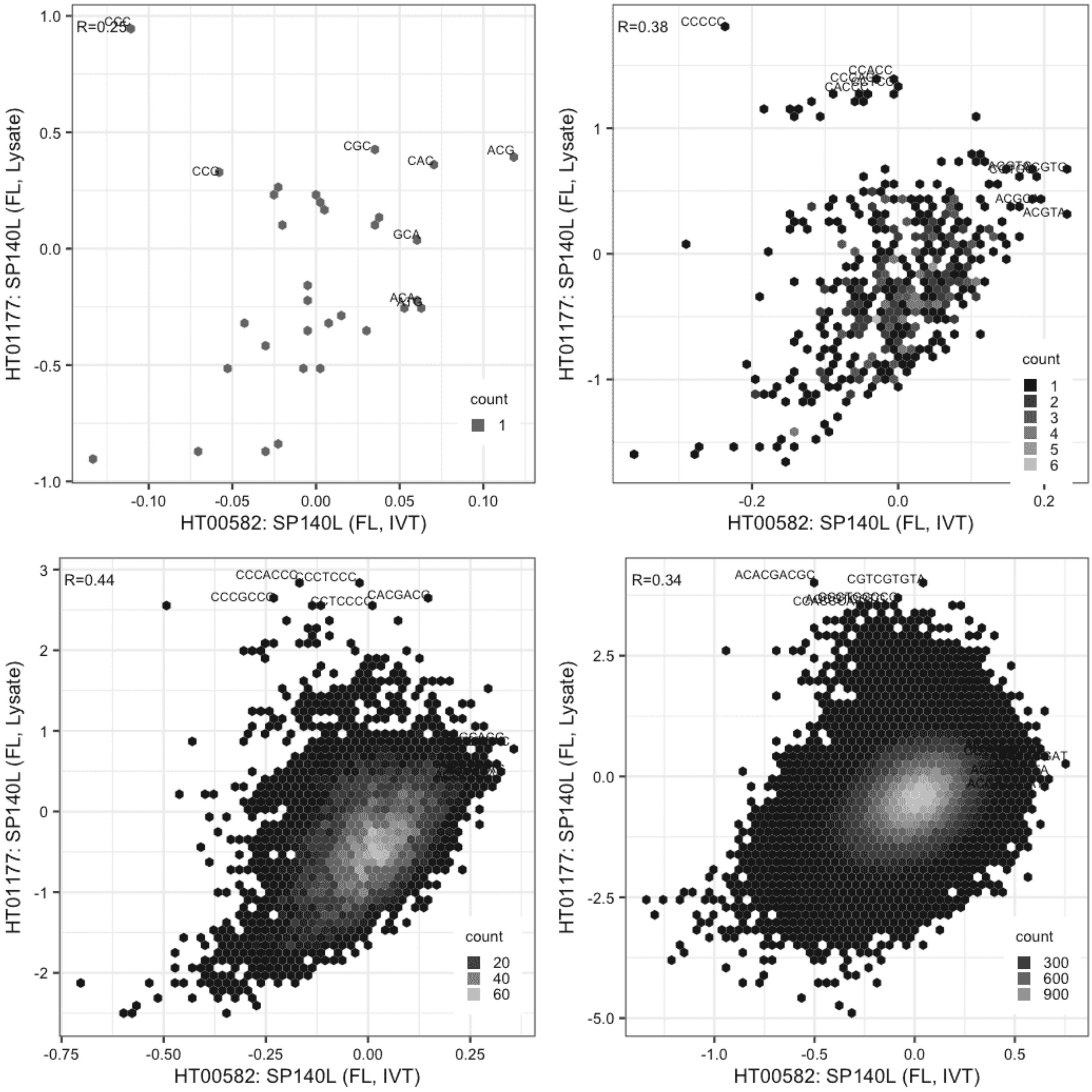

**SP140L**

IVT vs GFPIVT, same construct

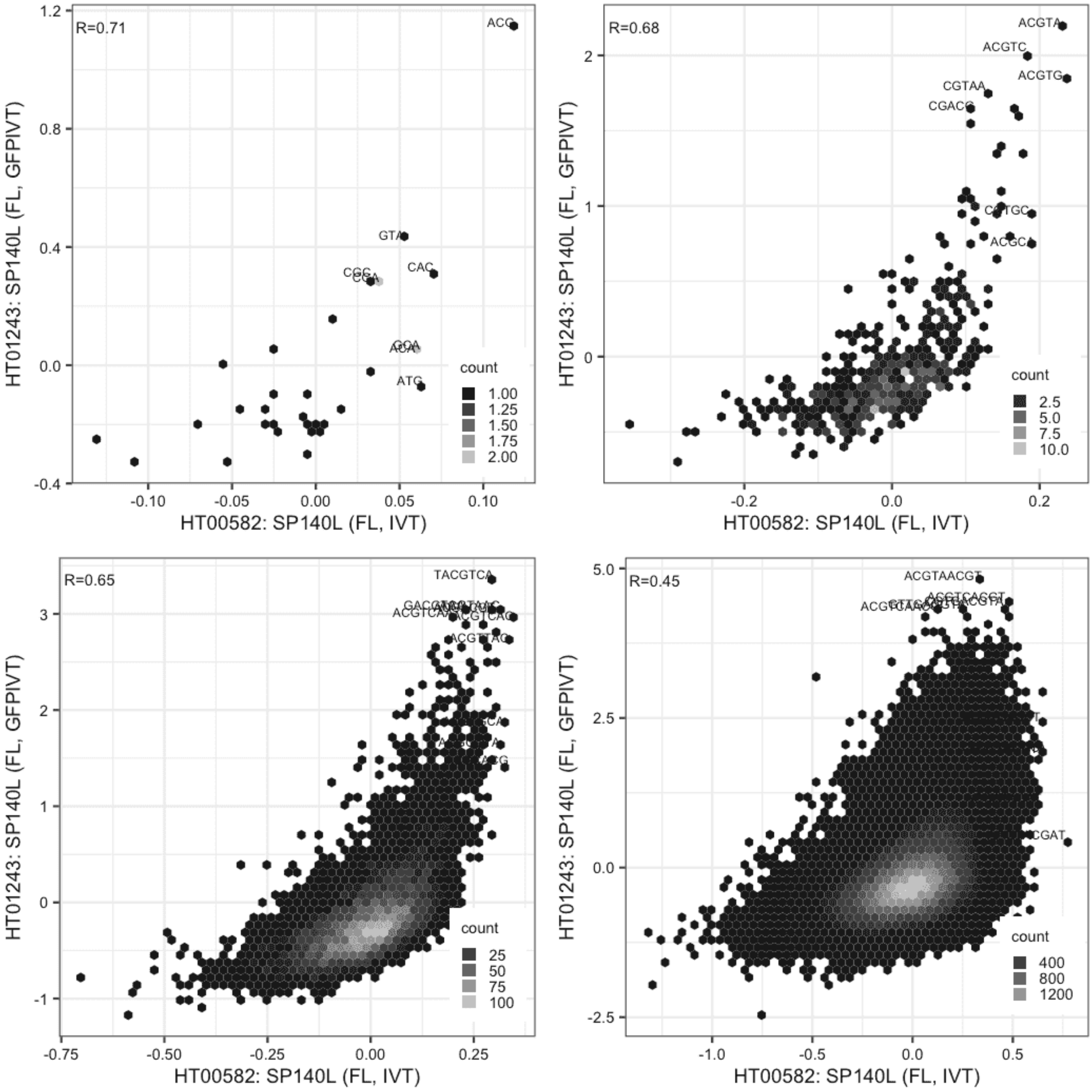

# SP140L

IVT vs GFPIVT, different constructs

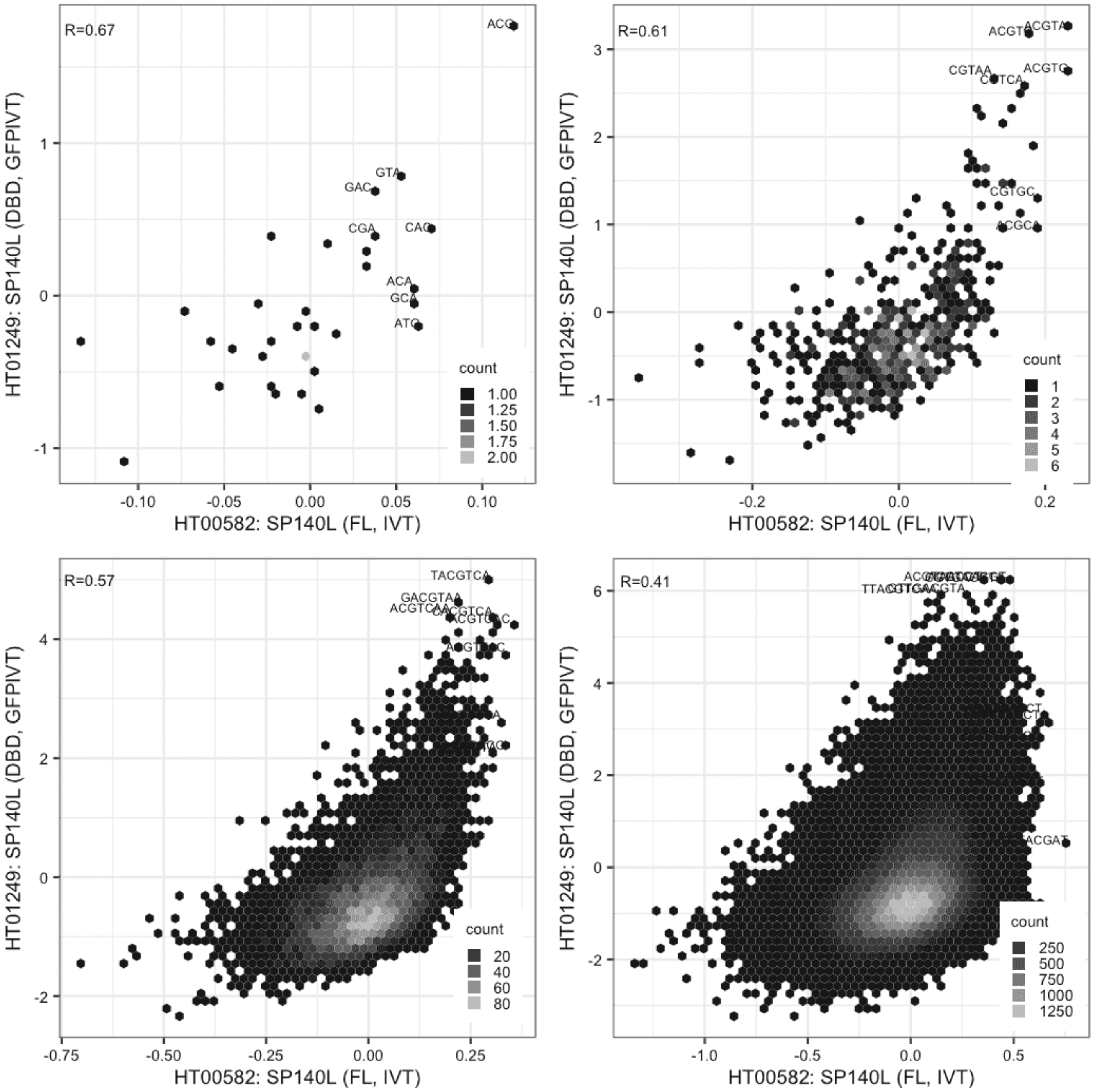

**SP140L**

Lysate vs Lysate, same construct

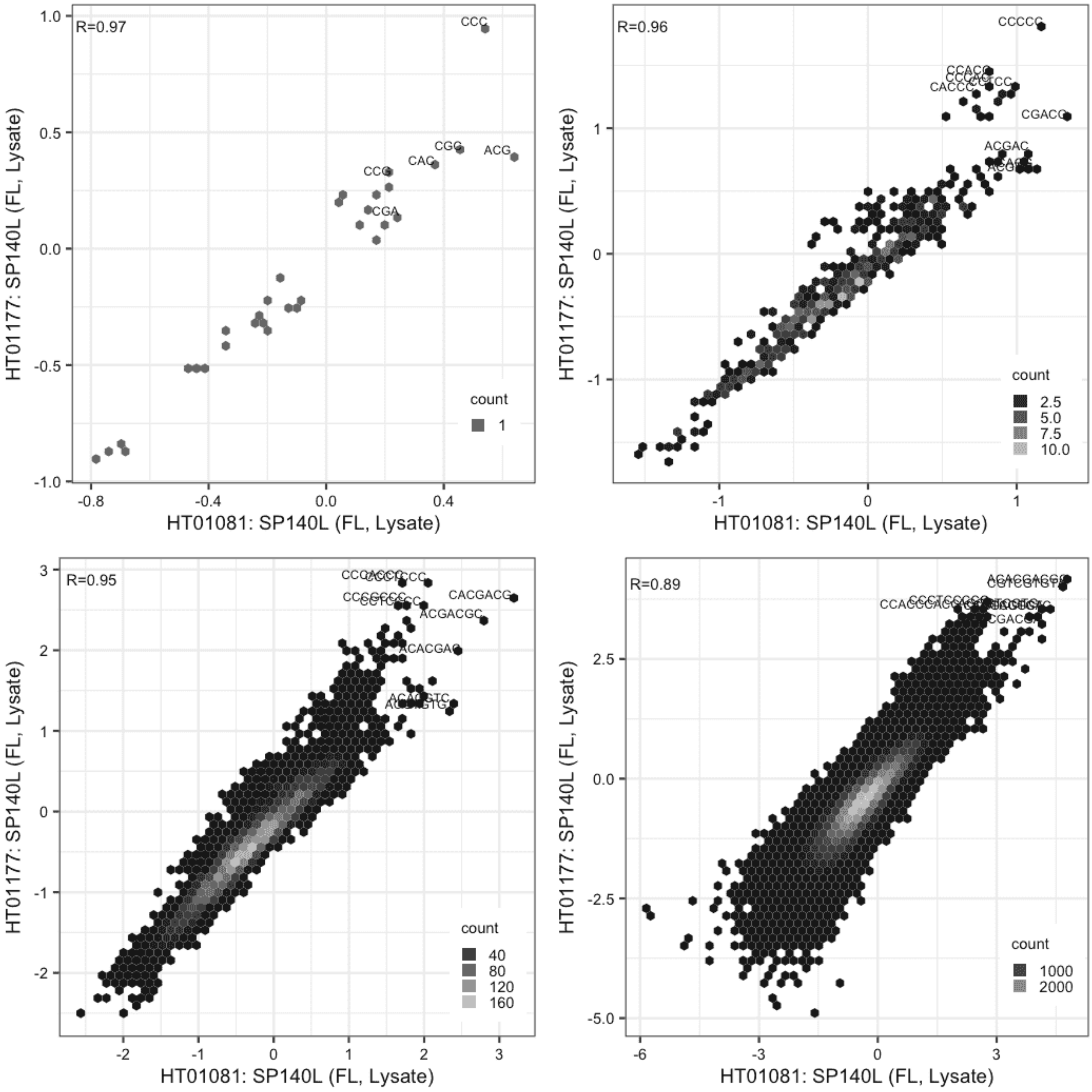

**SP140L**

Lysate vs GFPIVT, same construct

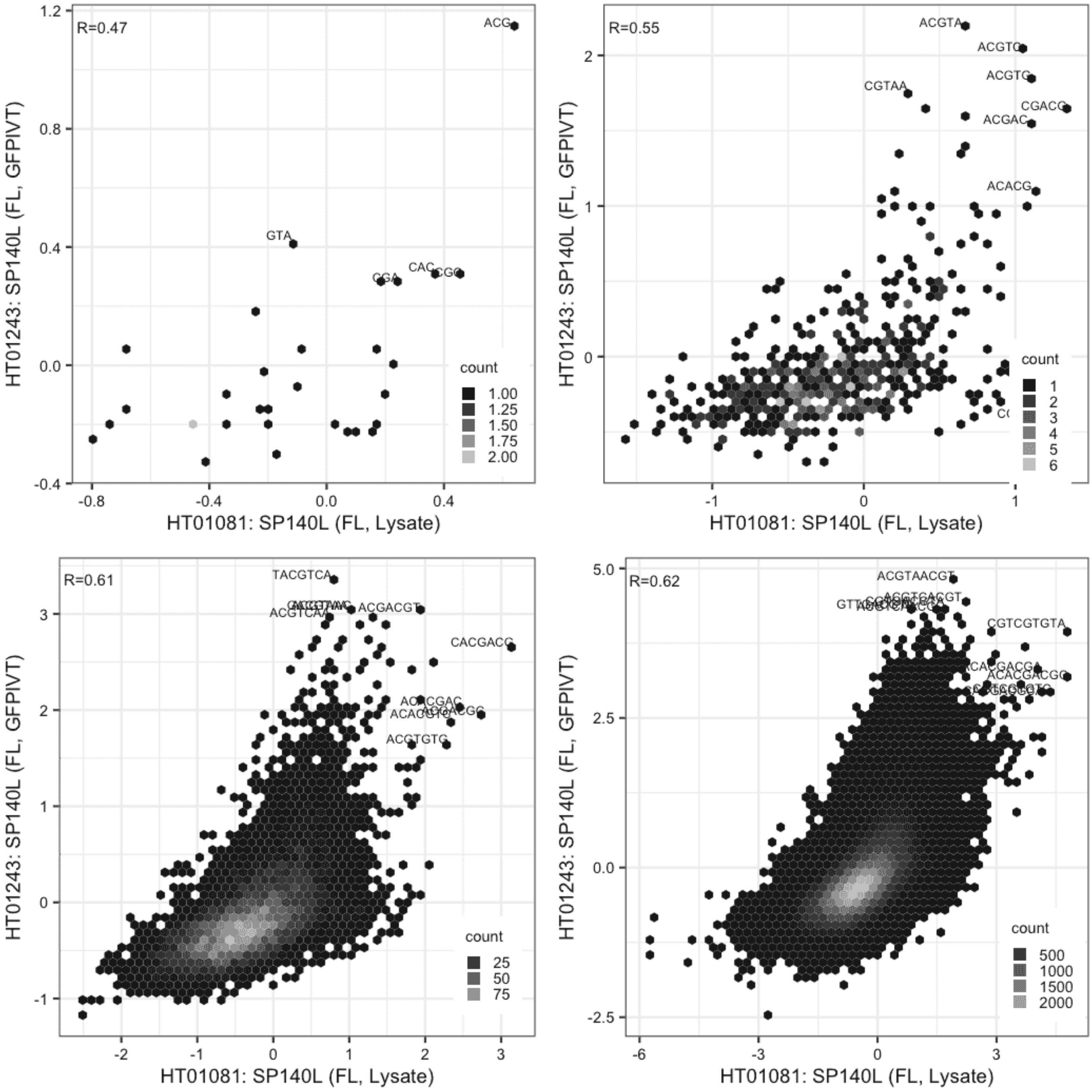

## SP140L

Lysate vs GFPIVT, different constructs

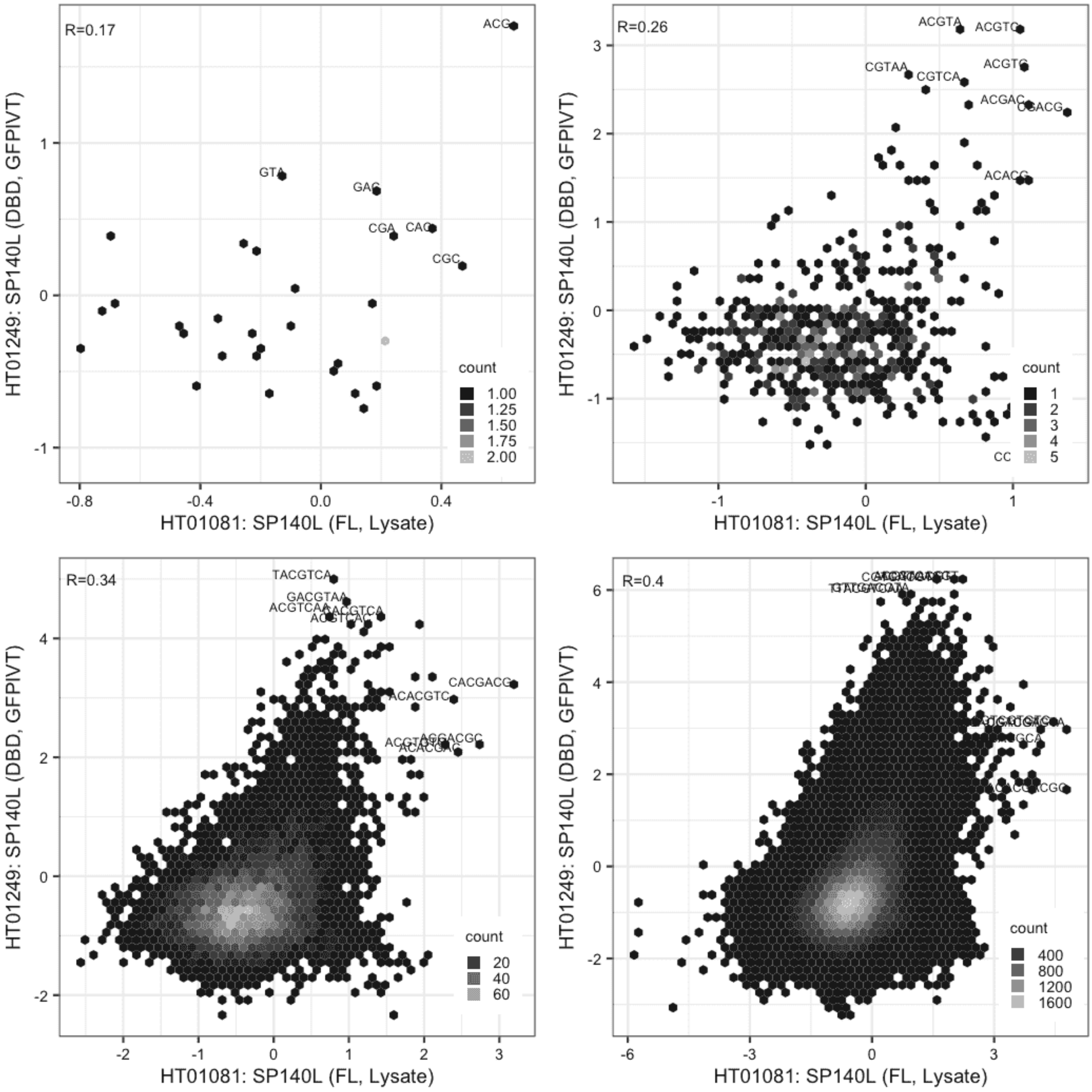

## SP140L

Lysate vs GFPIVT, same construct

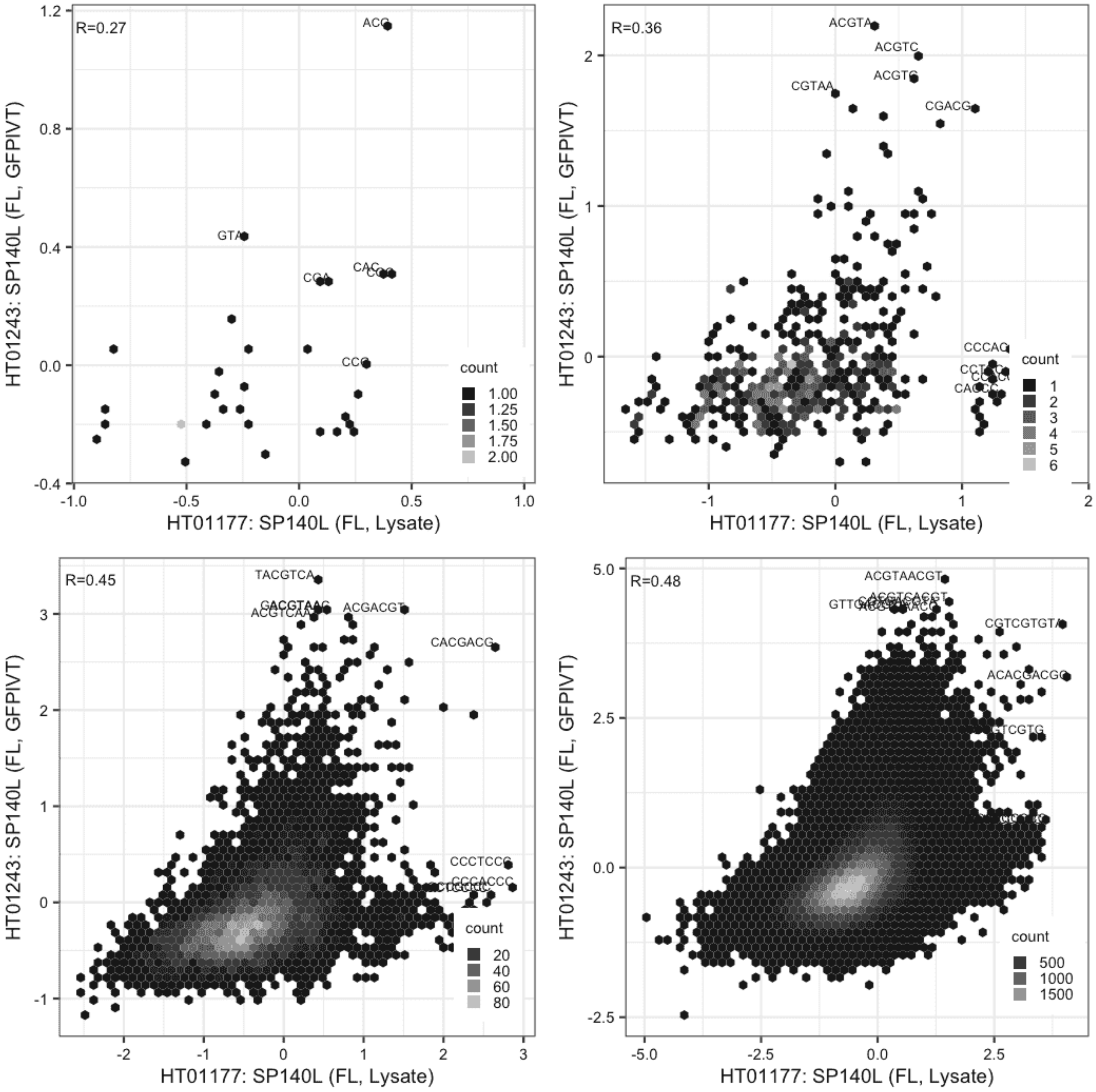

# SP140L

Lysate vs GFPIVT, different constructs

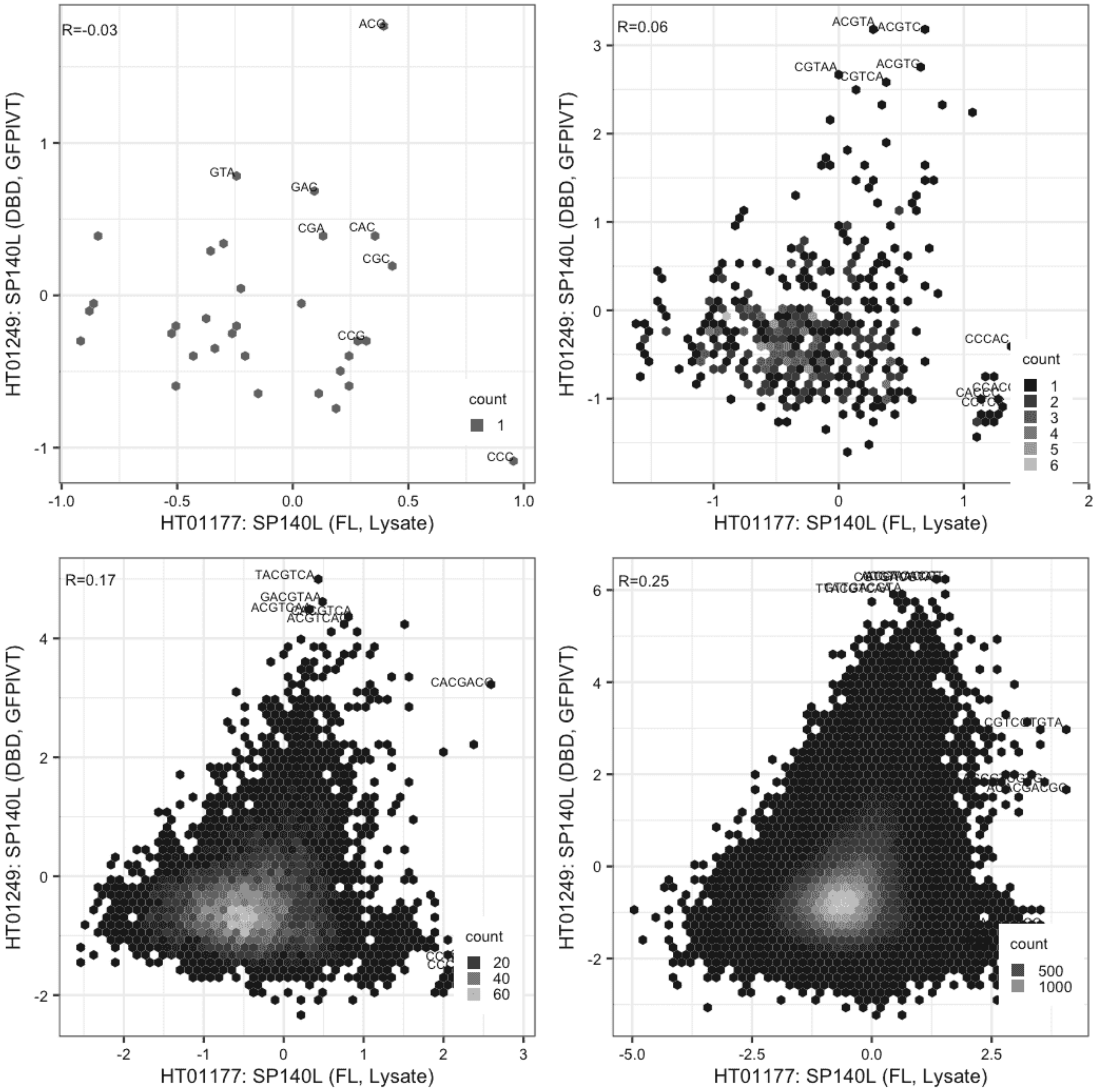

## SP140L

GFPIVT vs GFPIVT, different constructs

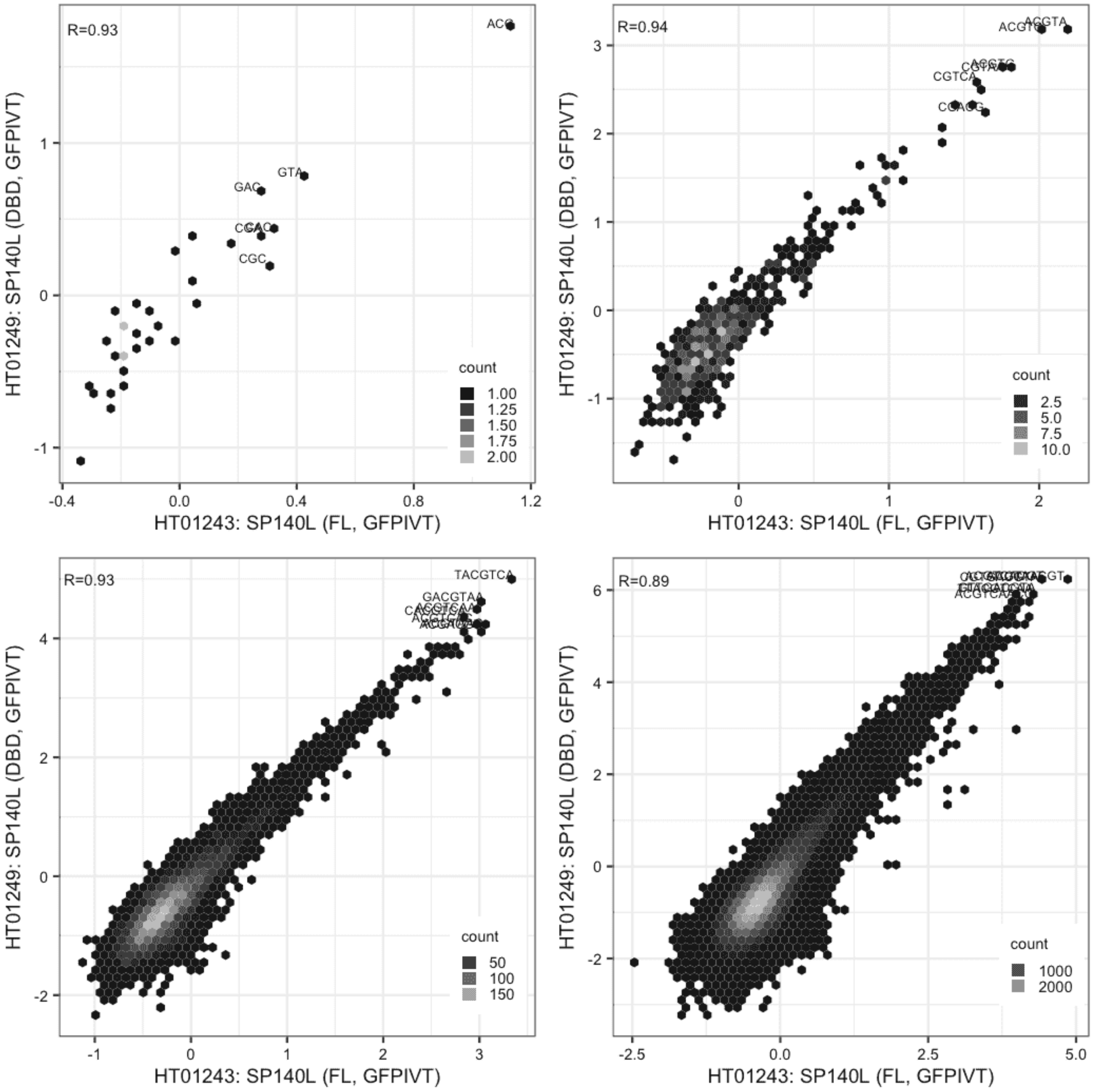

**SRY**

IVT vs IVT, same construct

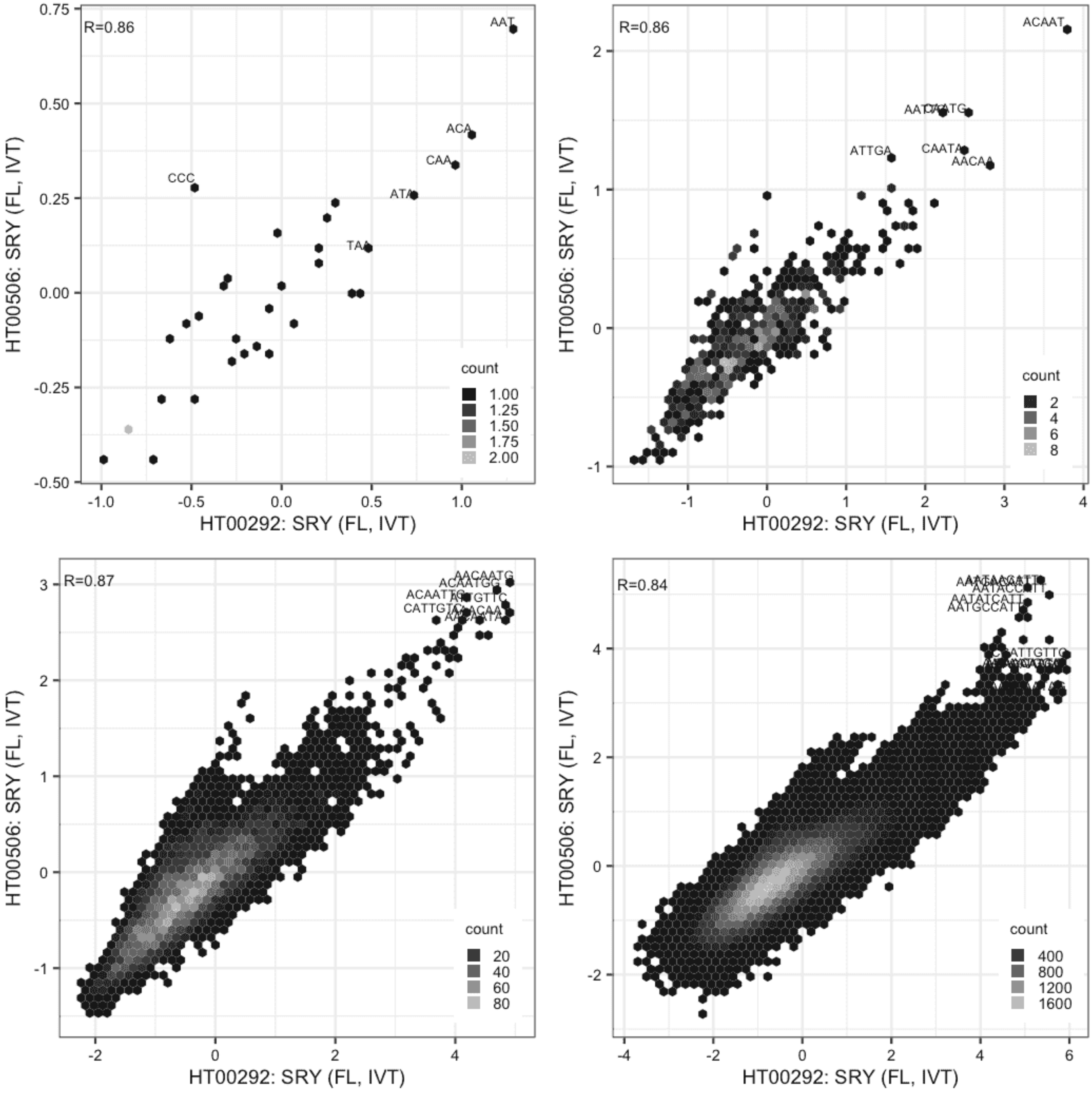

#### SRY

Lysate vs IVT, same construct

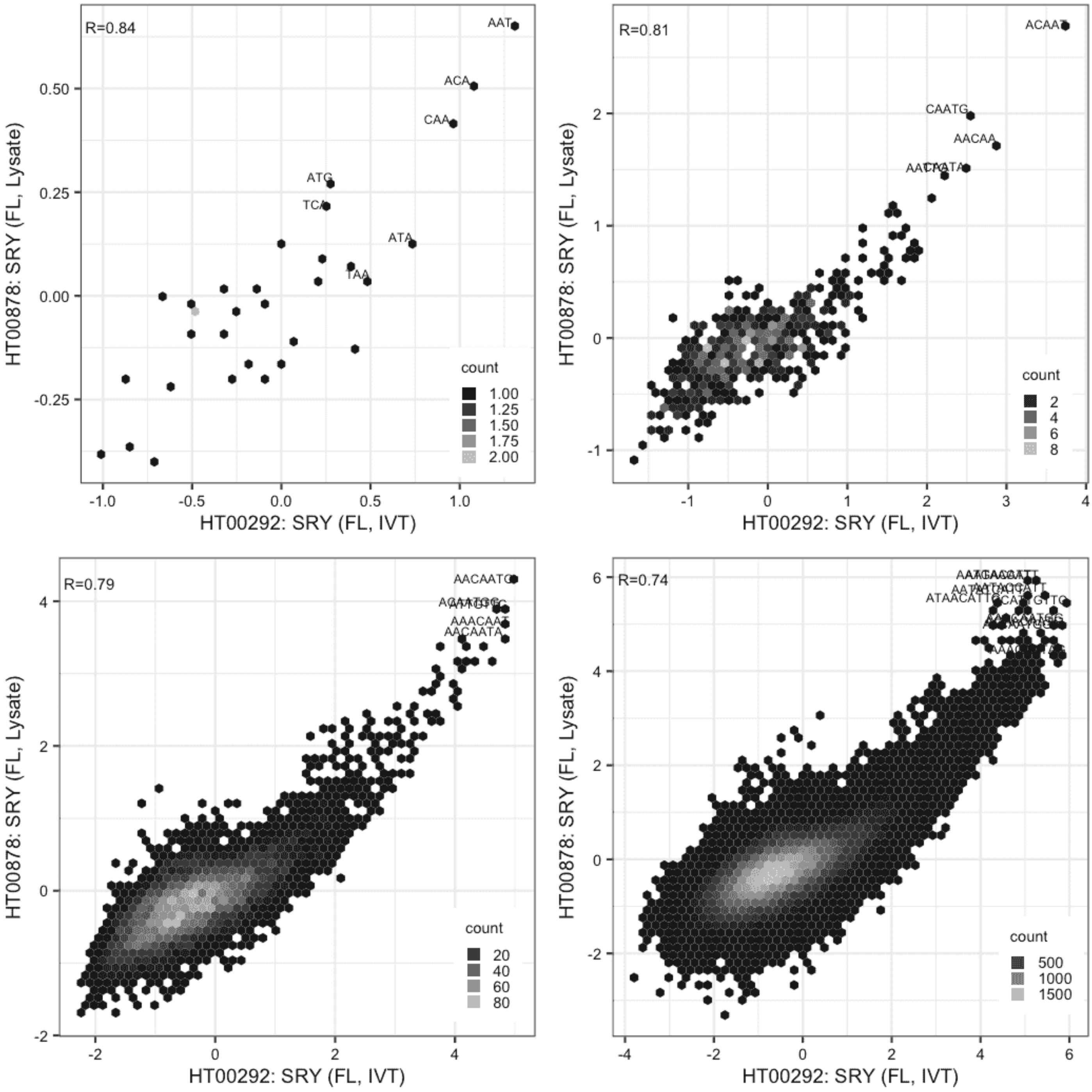

#### SRY

Lysate vs IVT, same construct

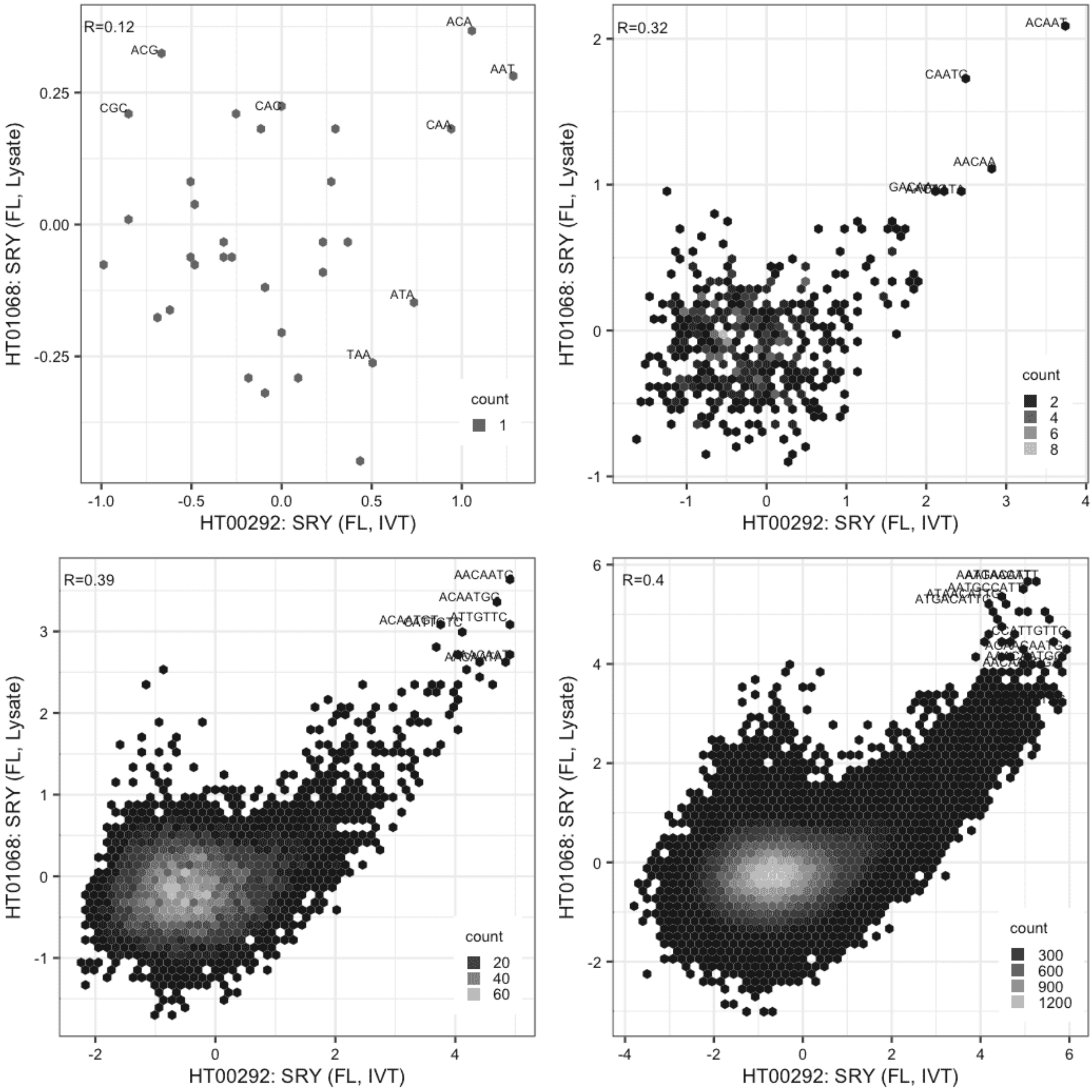

**SRY**

Lysate vs IVT, same construct

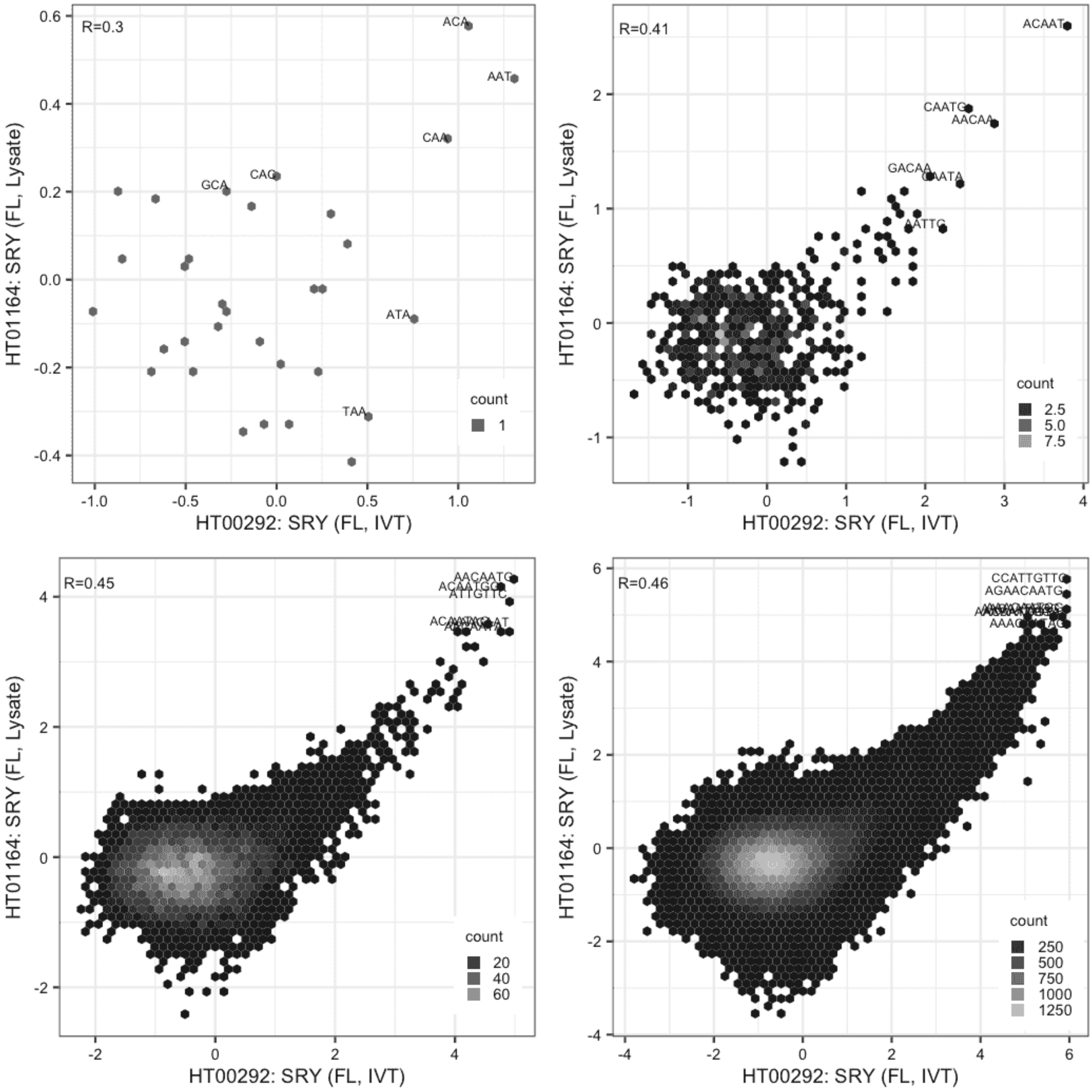

**SRY**

Lysate vs IVT, same construct

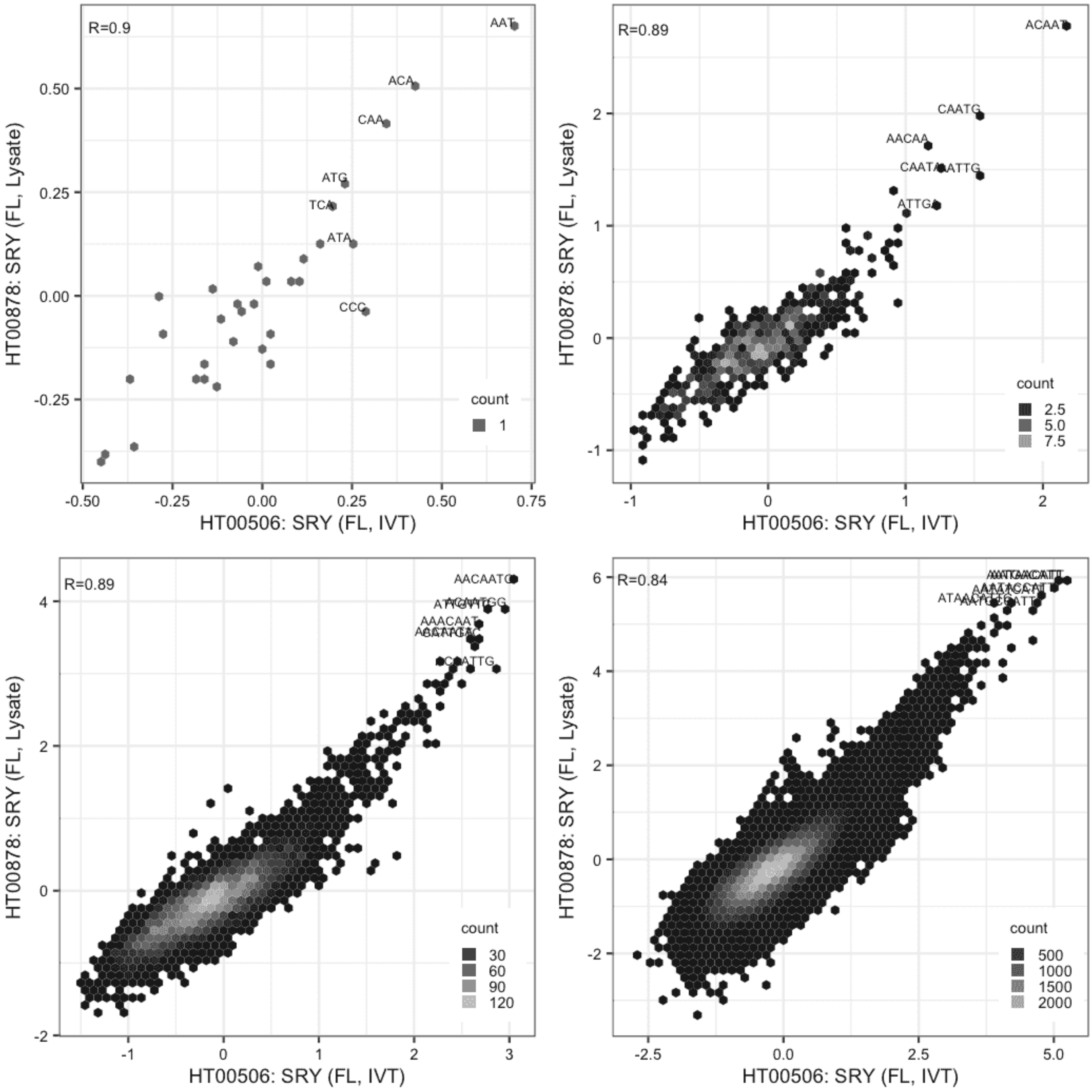

#### SRY

Lysate vs IVT, same construct

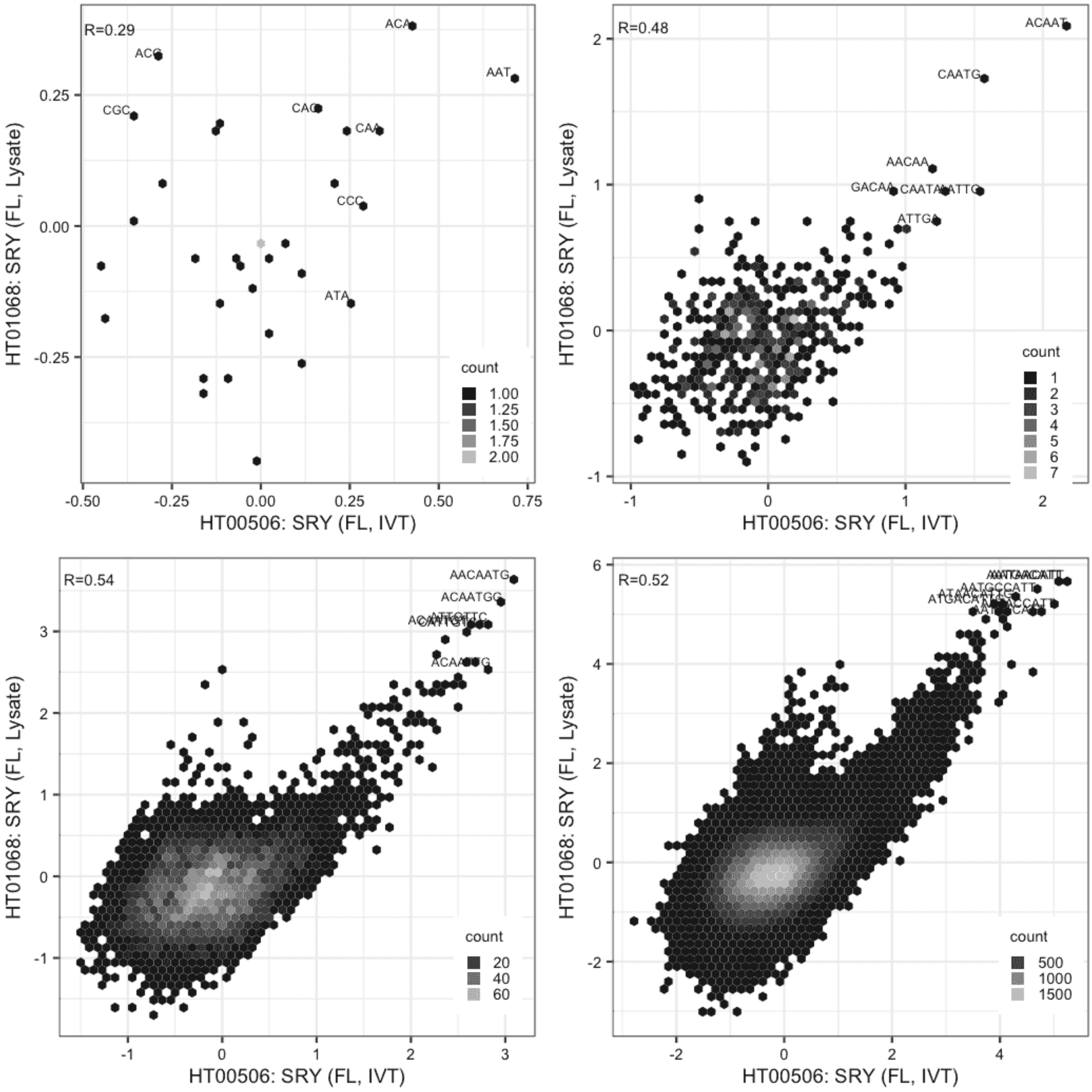

#### SRY

Lysate vs IVT, same construct

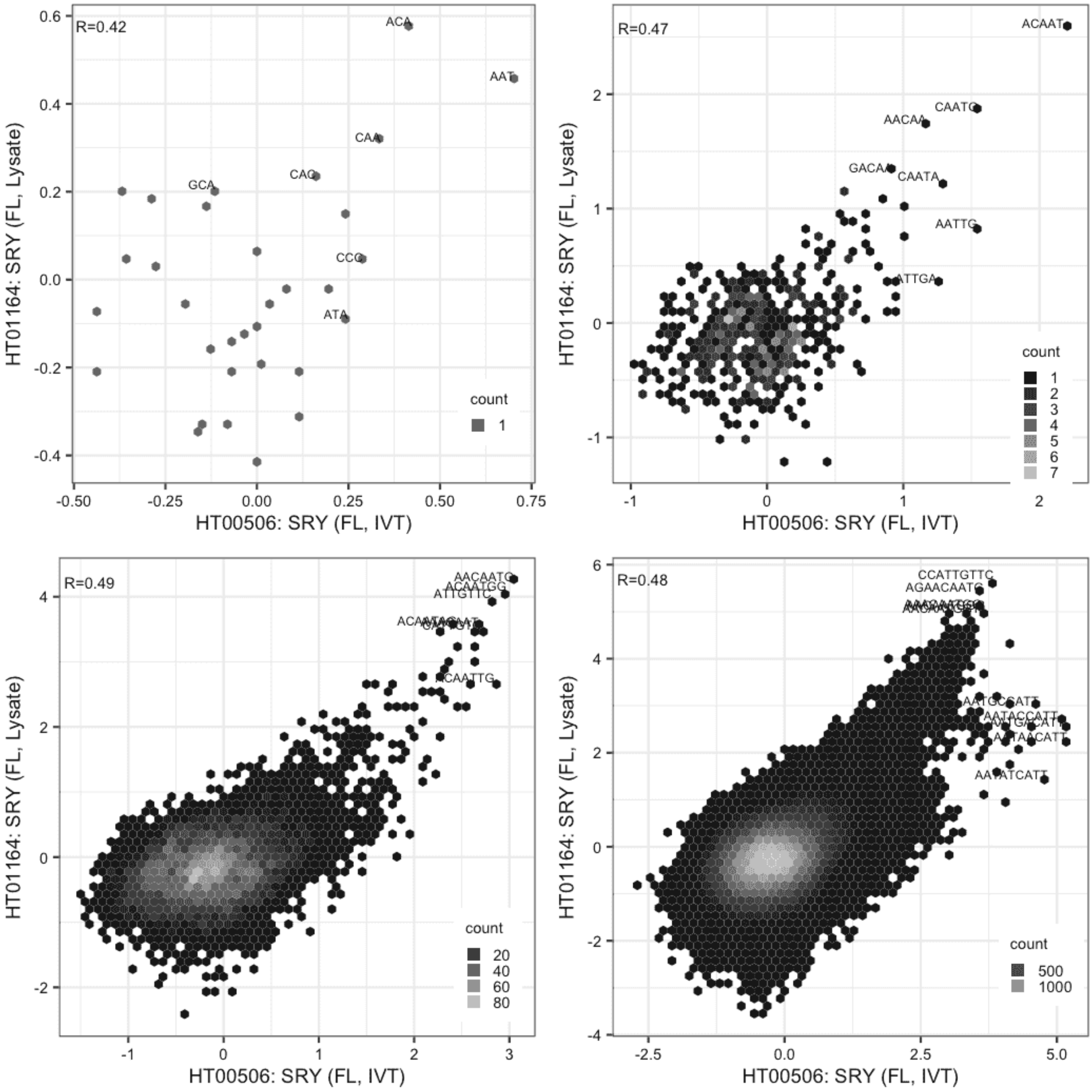

**SRY**

Lysate vs Lysate, same construct

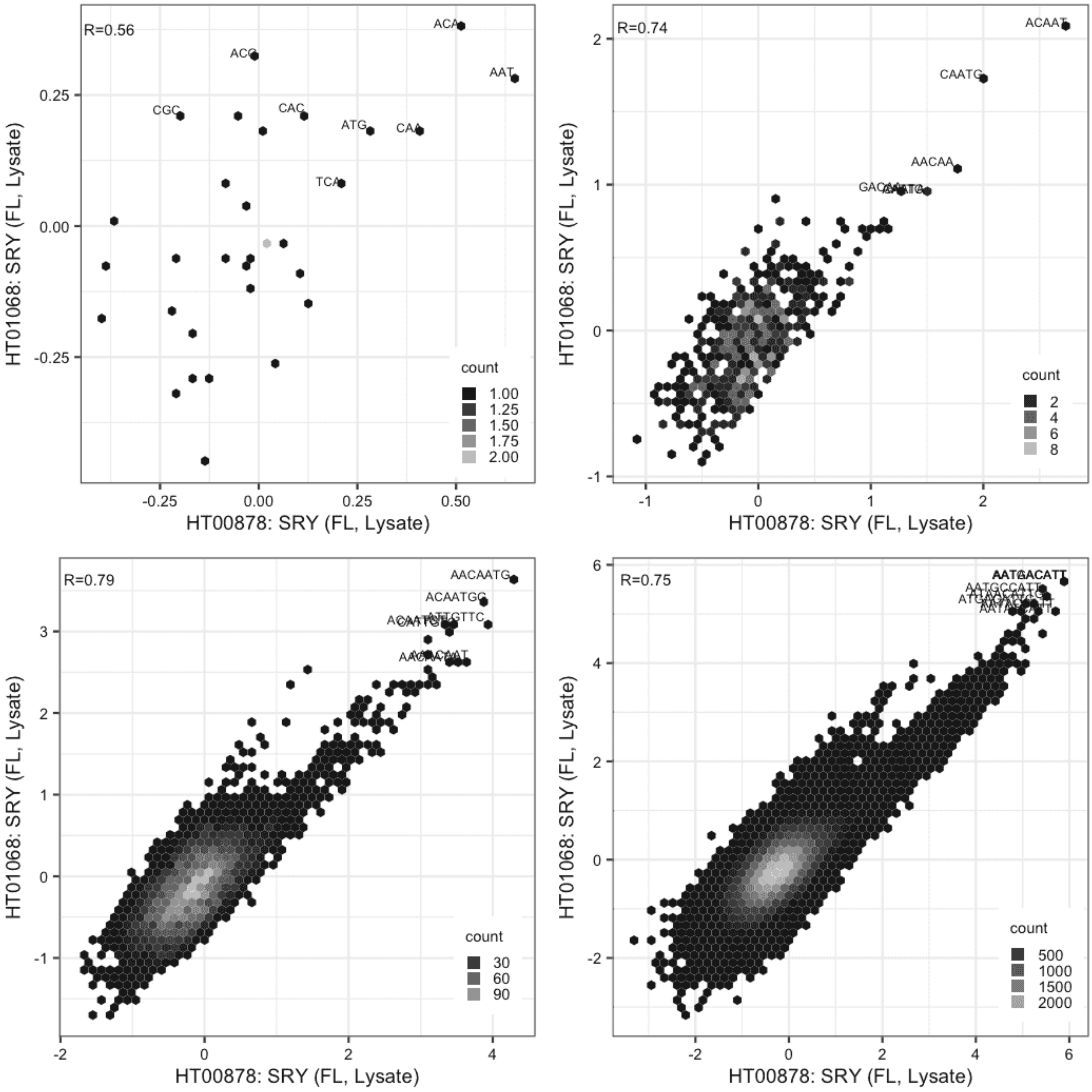

#### SRY

Lysate vs Lysate, same construct

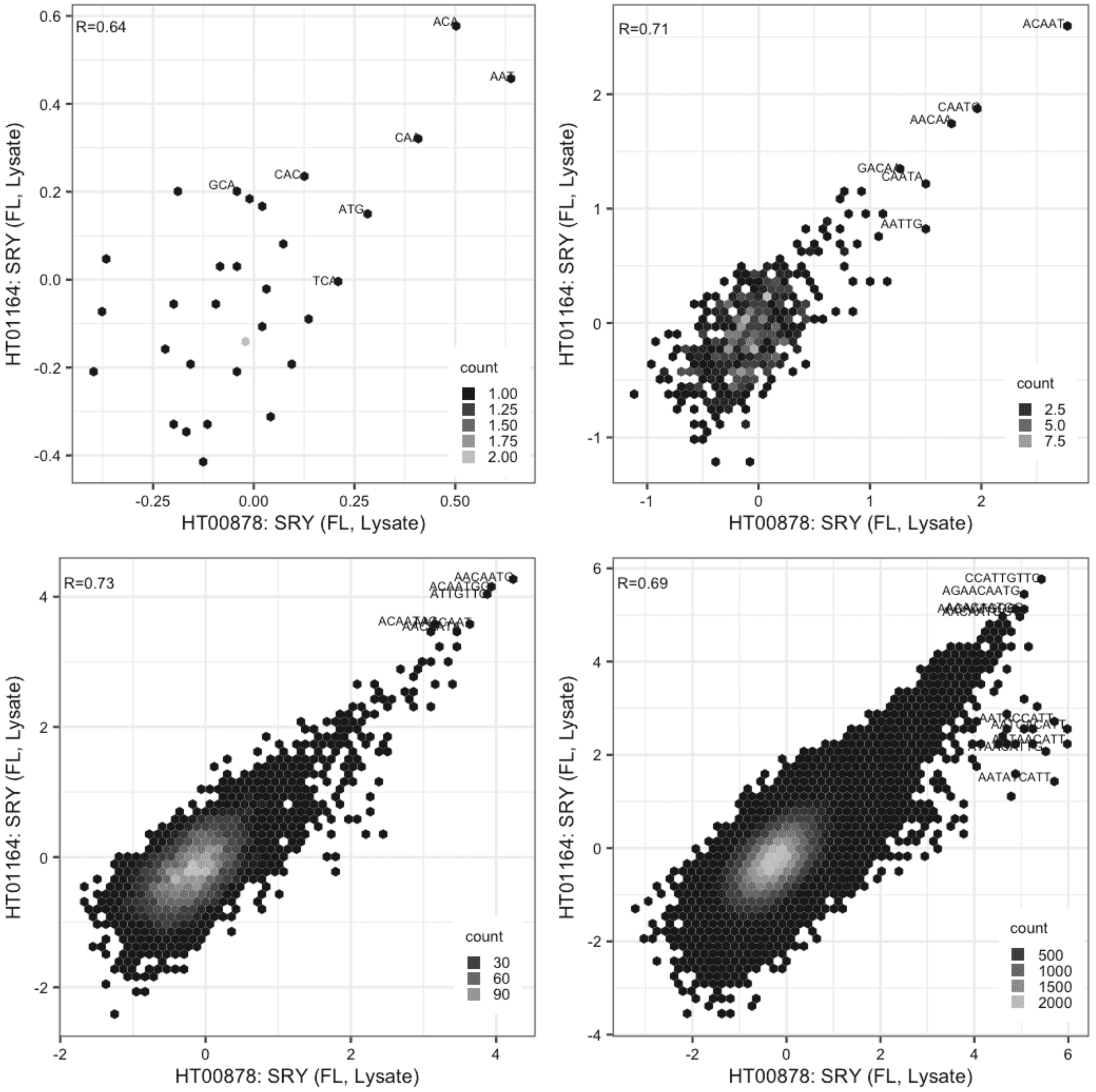

**SRY**

Lysate vs Lysate, same construct

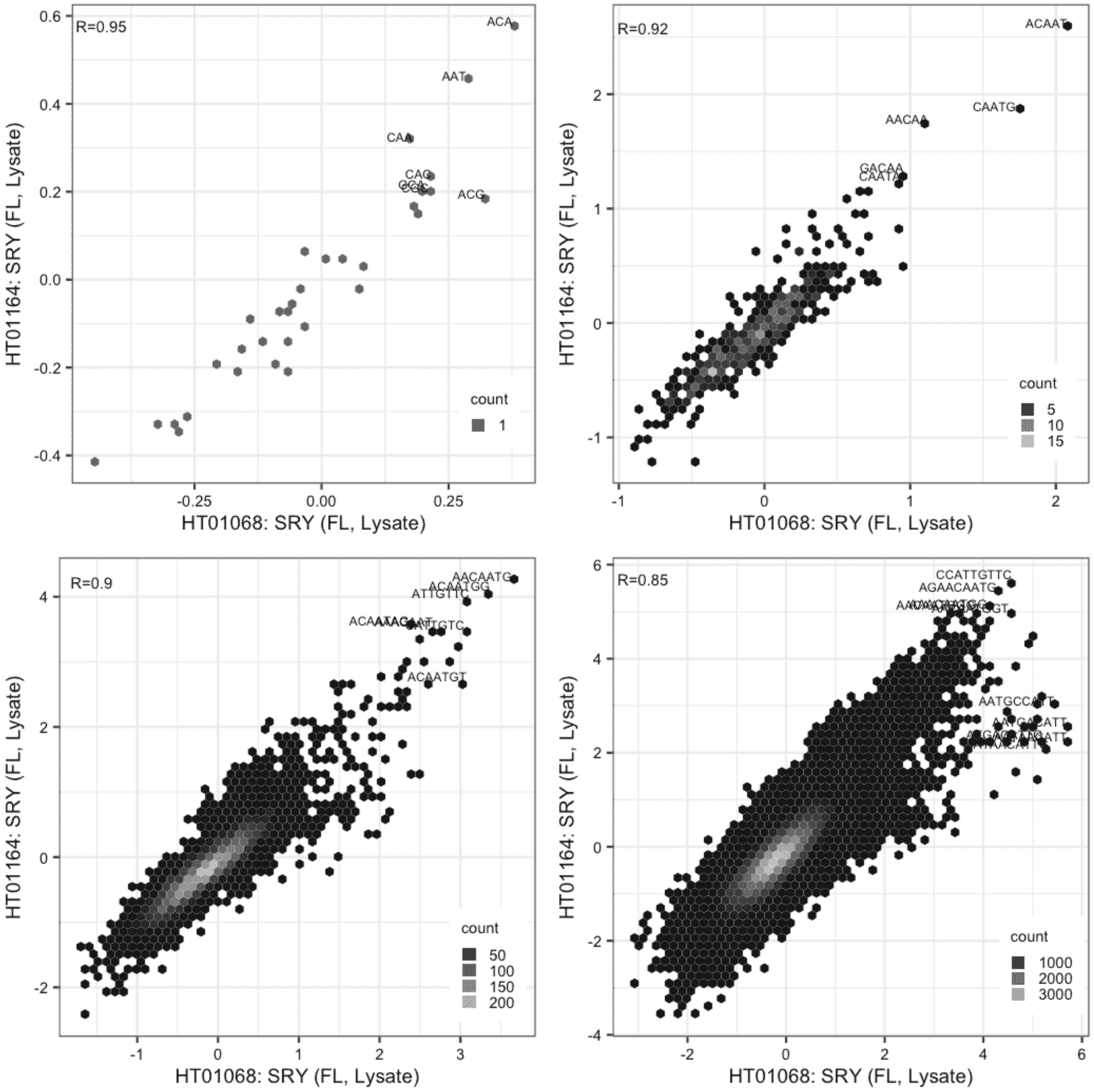

### TET3

IVT vs GFPIVT, same construct

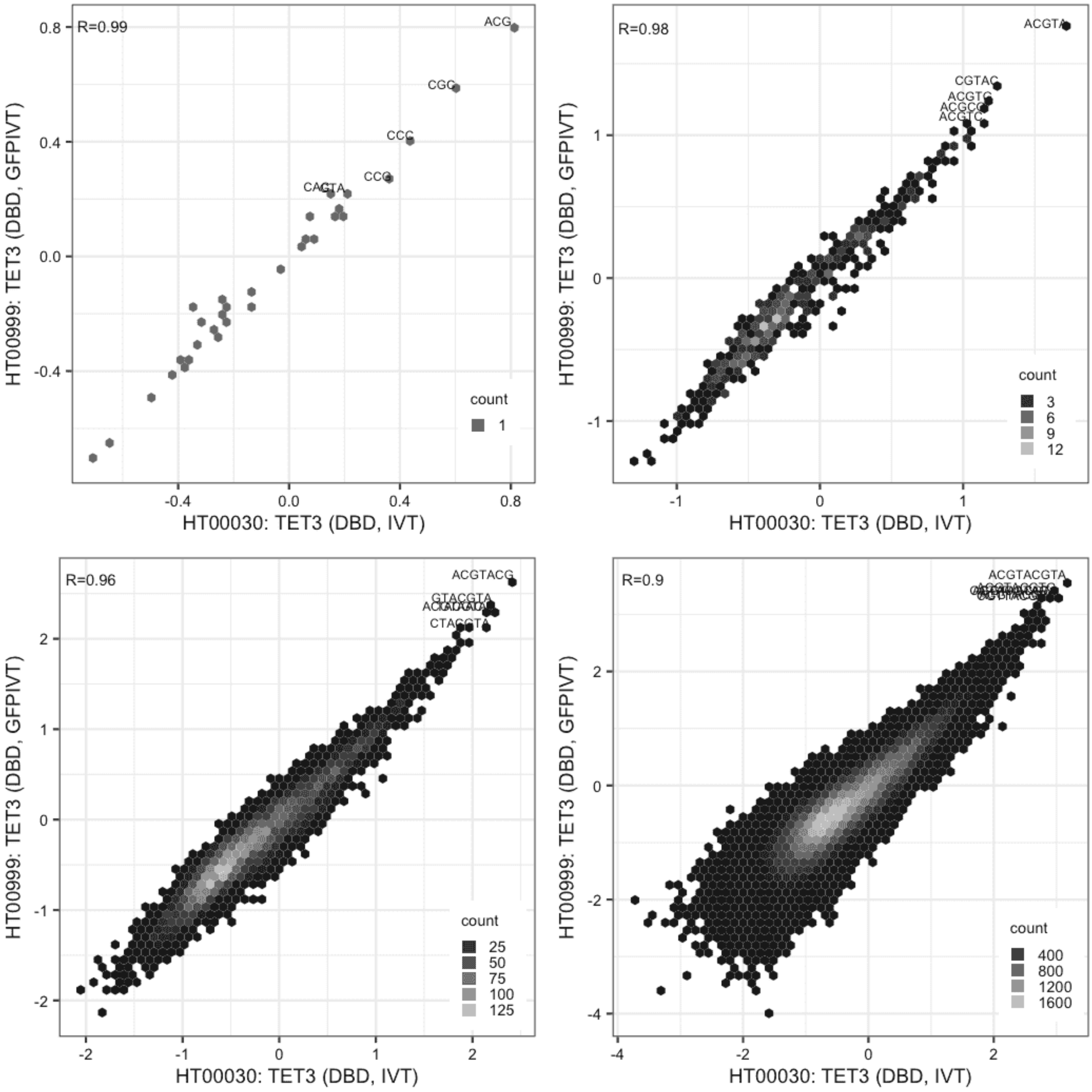

**TET3**

Lysate vs IVT, different constructs

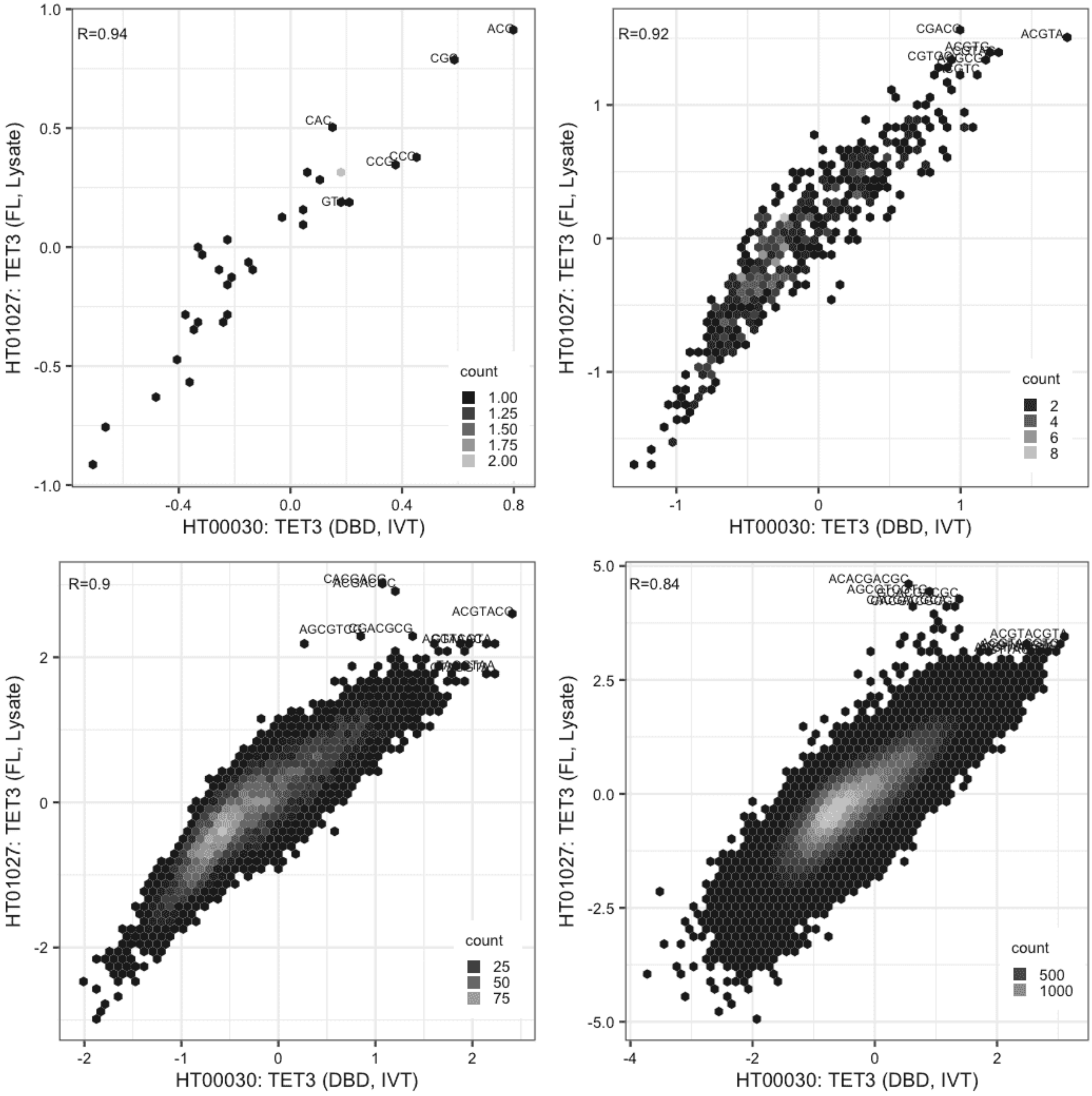

#### TET3

Lysate vs IVT, different constructs

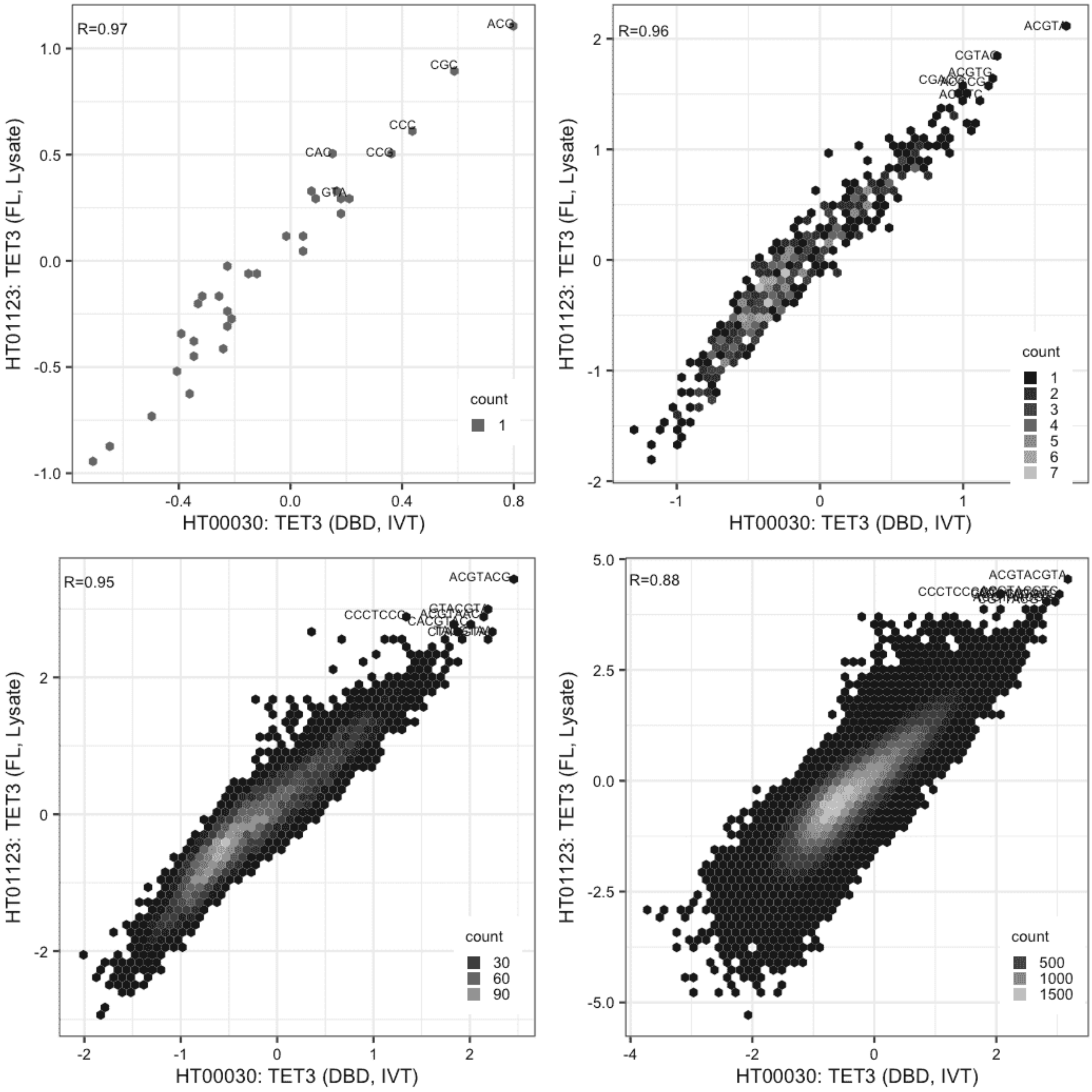

**TET3**

Lysate vs GFPIVT, different constructs

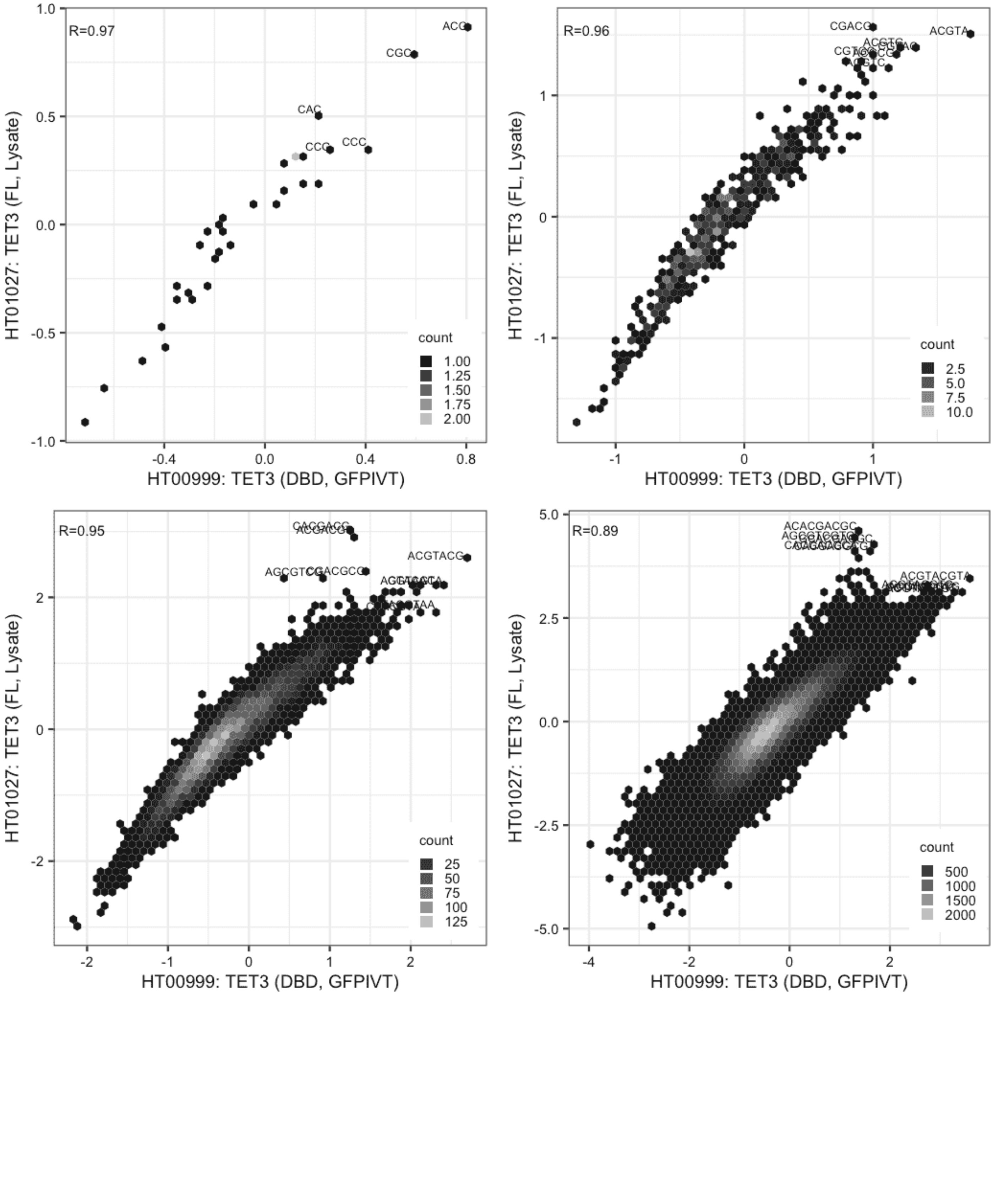

#### TET3

Lysate vs GFPIVT, different constructs

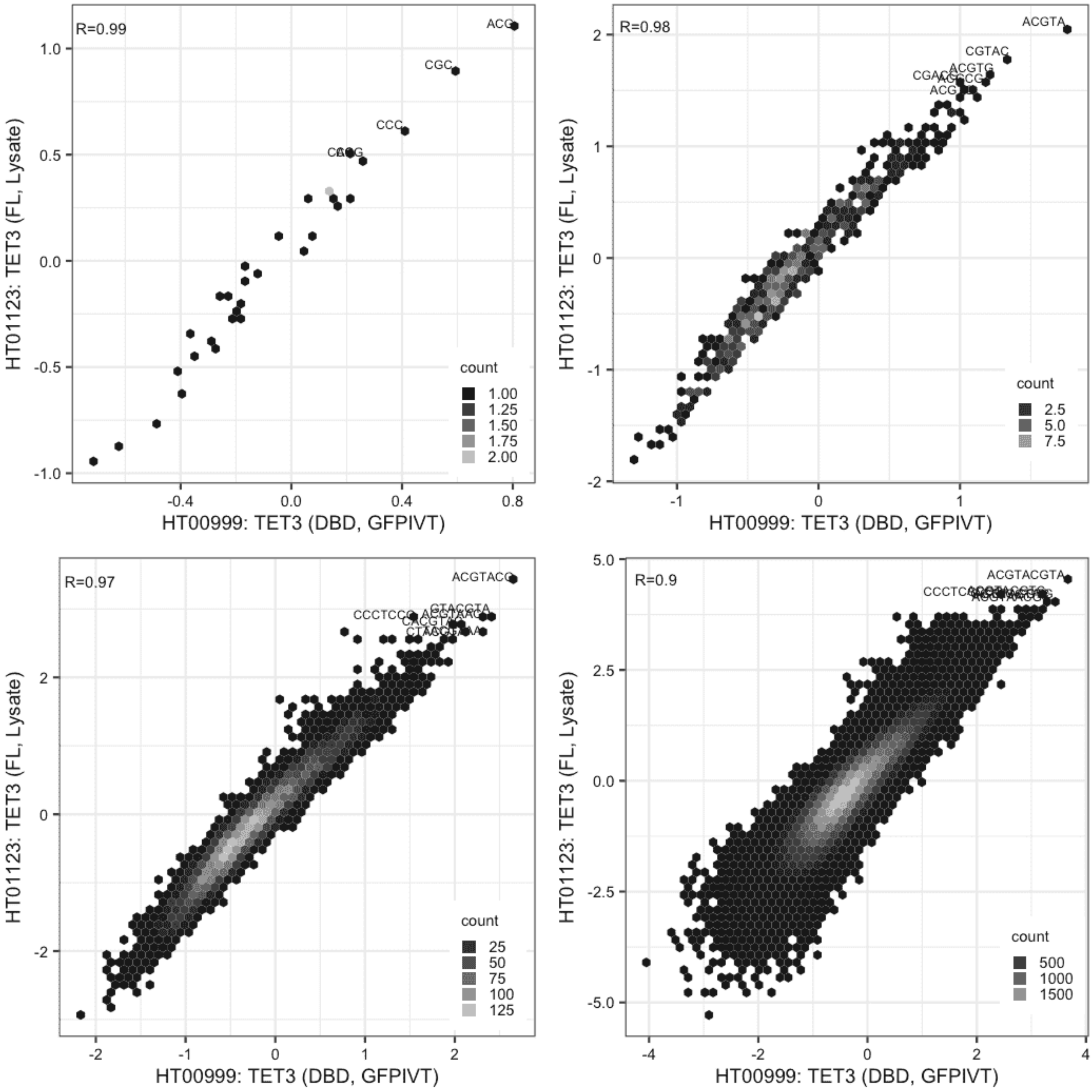

**TET3**

Lysate vs Lysate, same construct

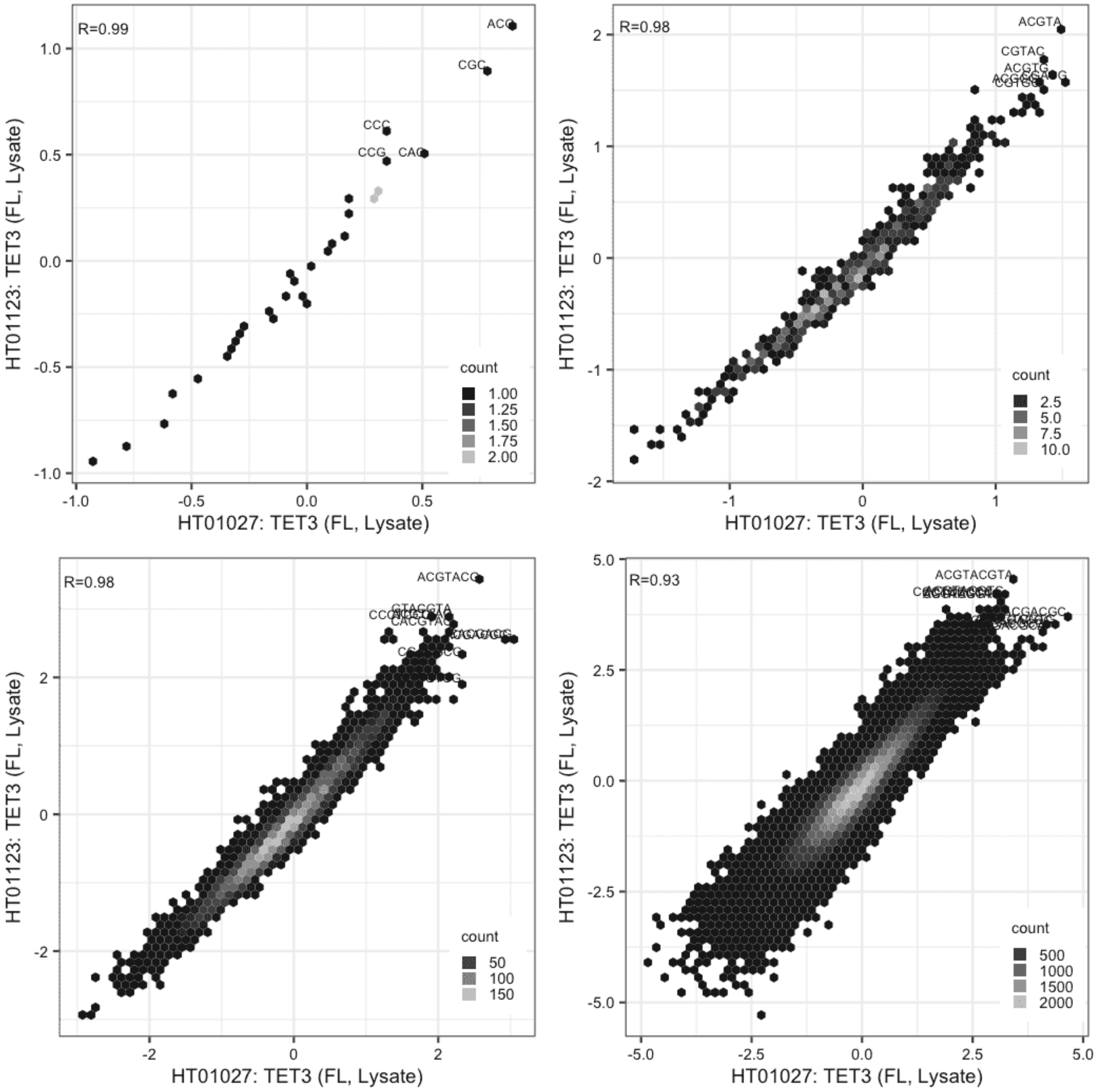

#### TIGD3

Lysate vs Lysate, same construct

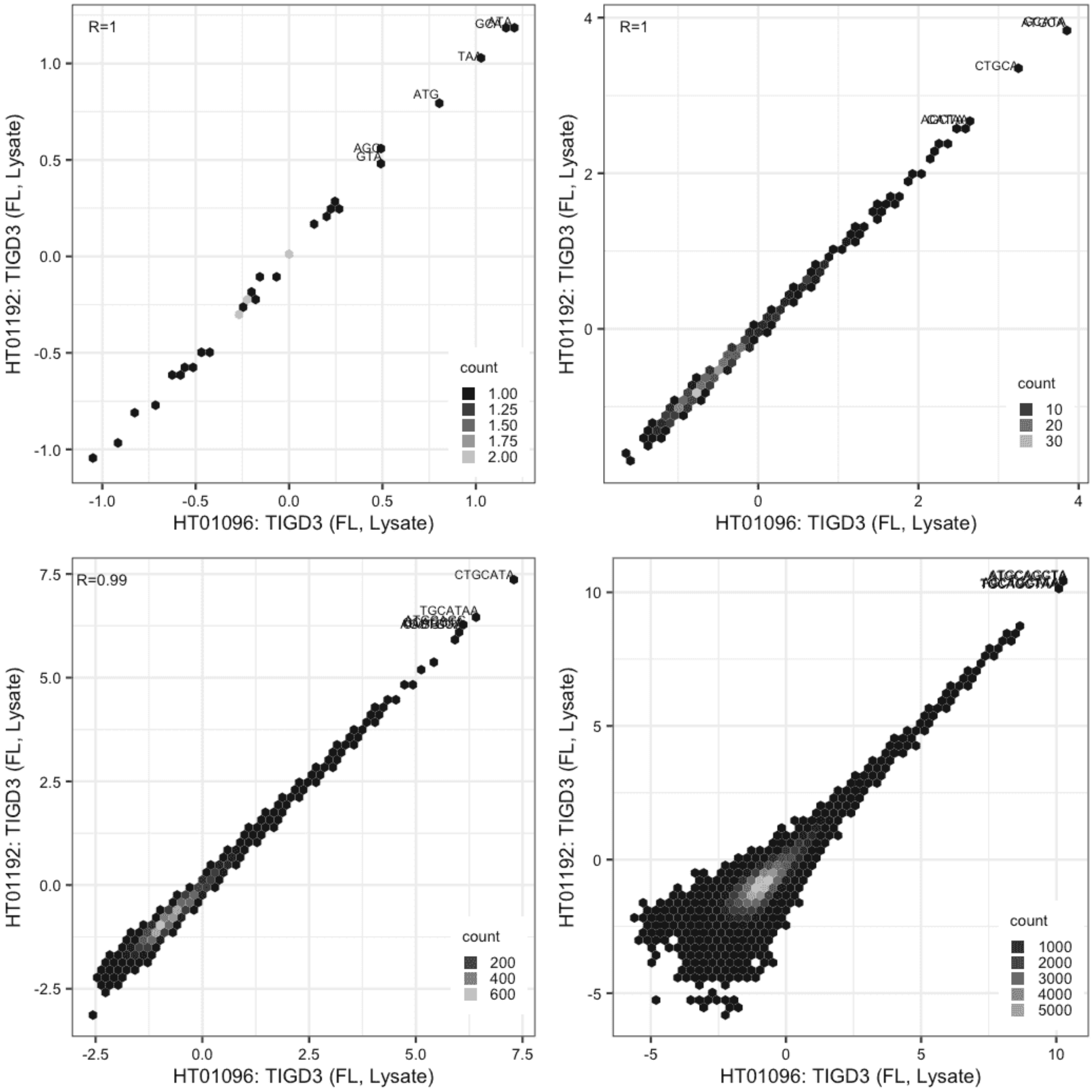

**TIGD3**

Lysate vs GFPIVT, same construct

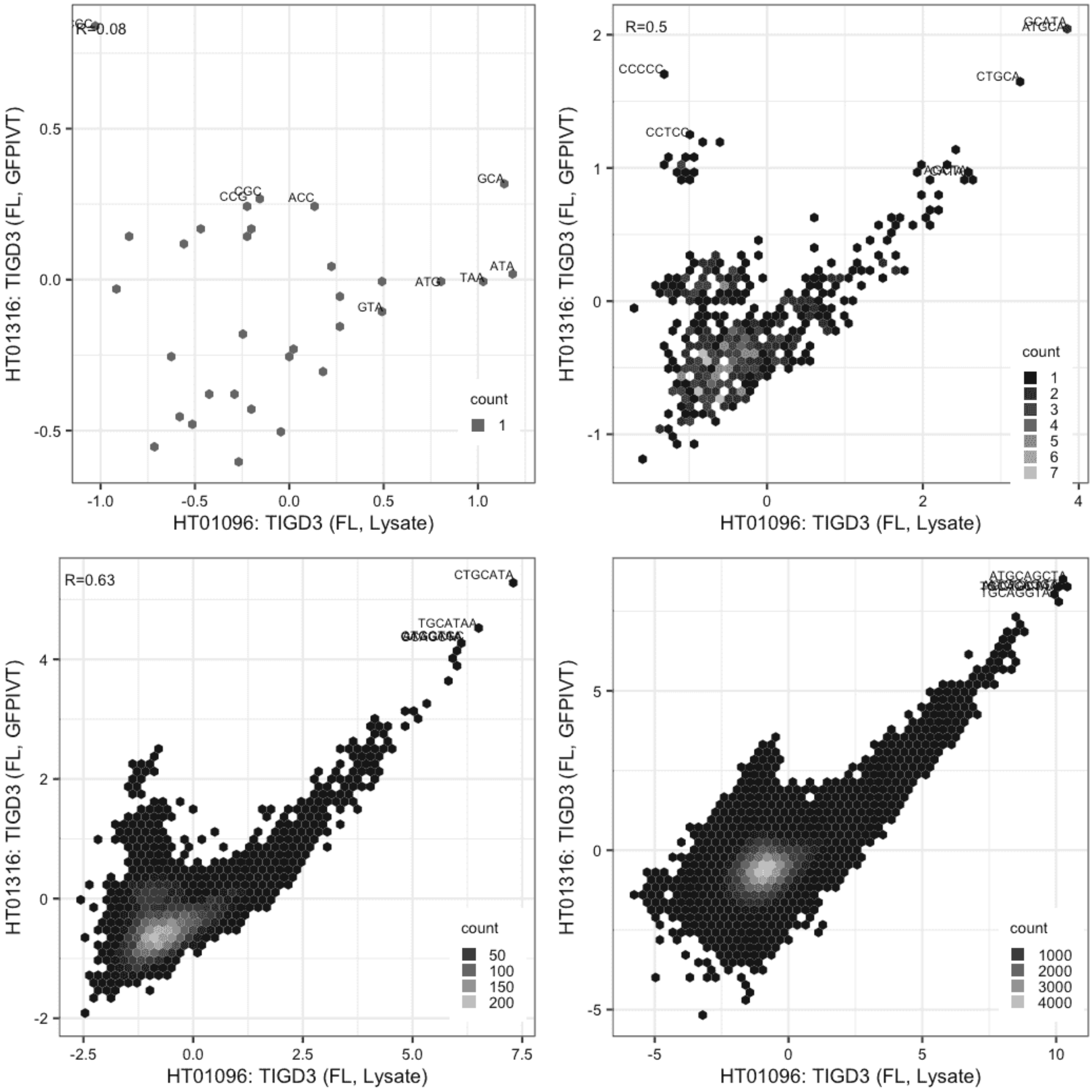

**TIGD3**

Lysate vs GFPIVT, same construct

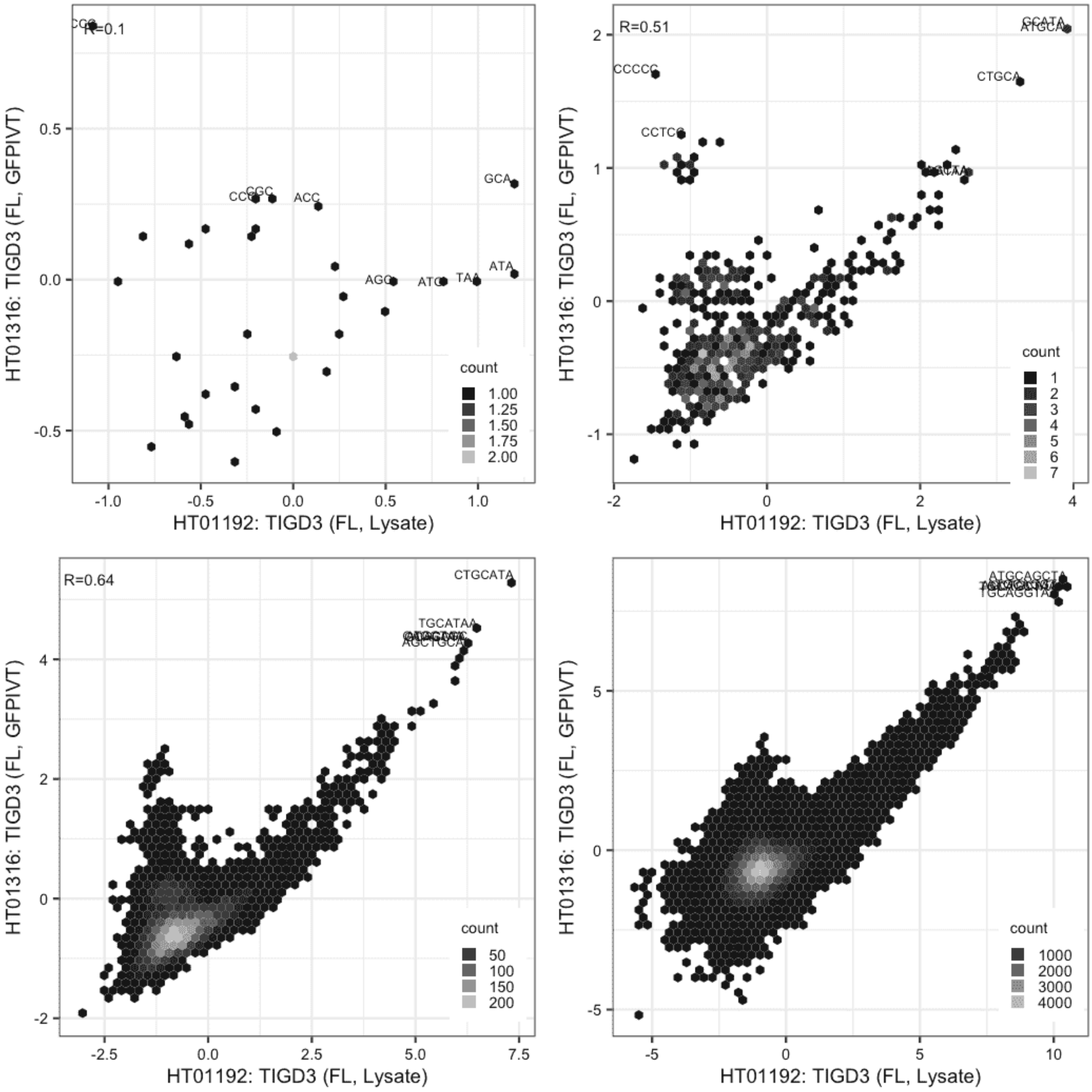

**TIGD4**

IVT vs GFPIVT, same construct

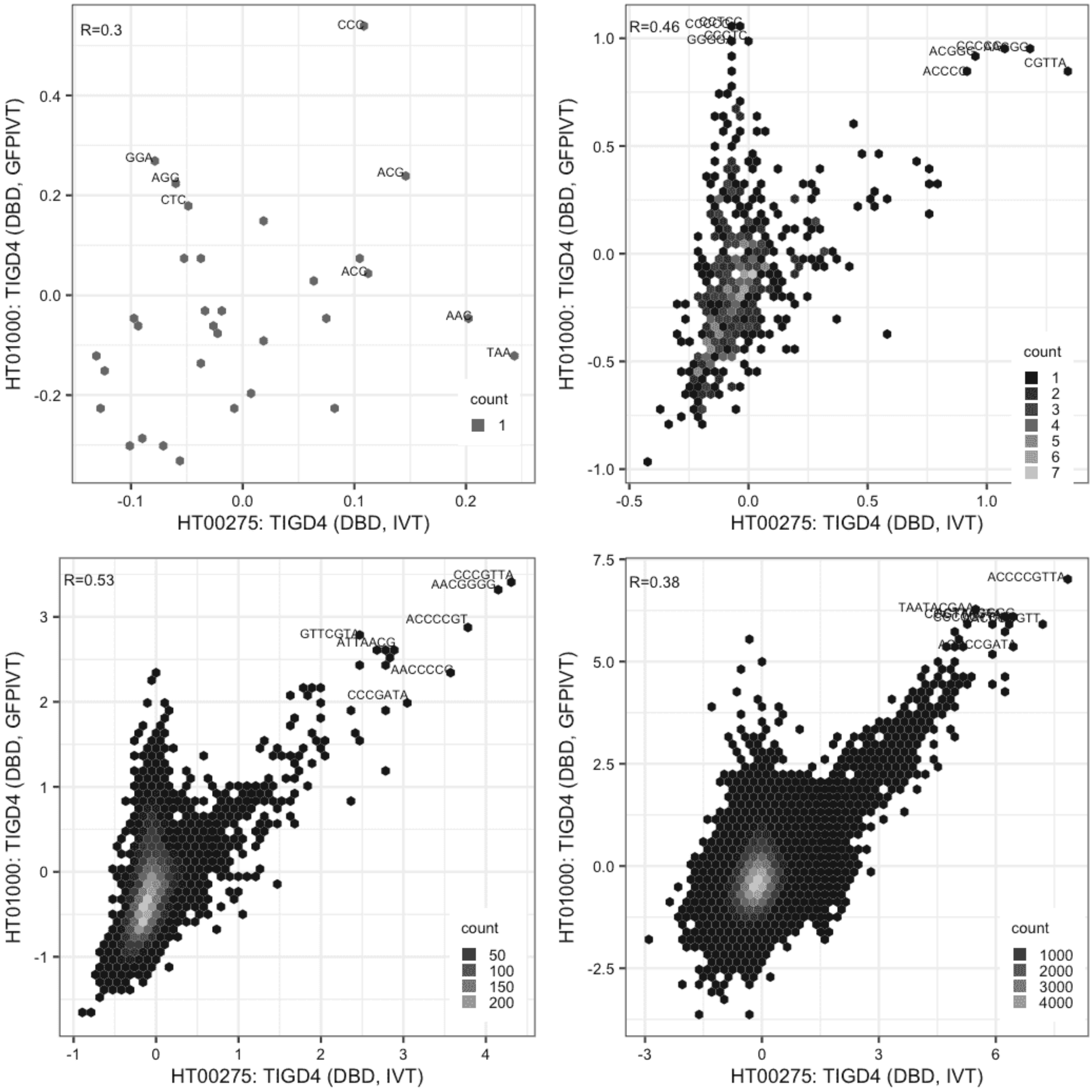

**TIGD4**

Lysate vs IVT, different constructs

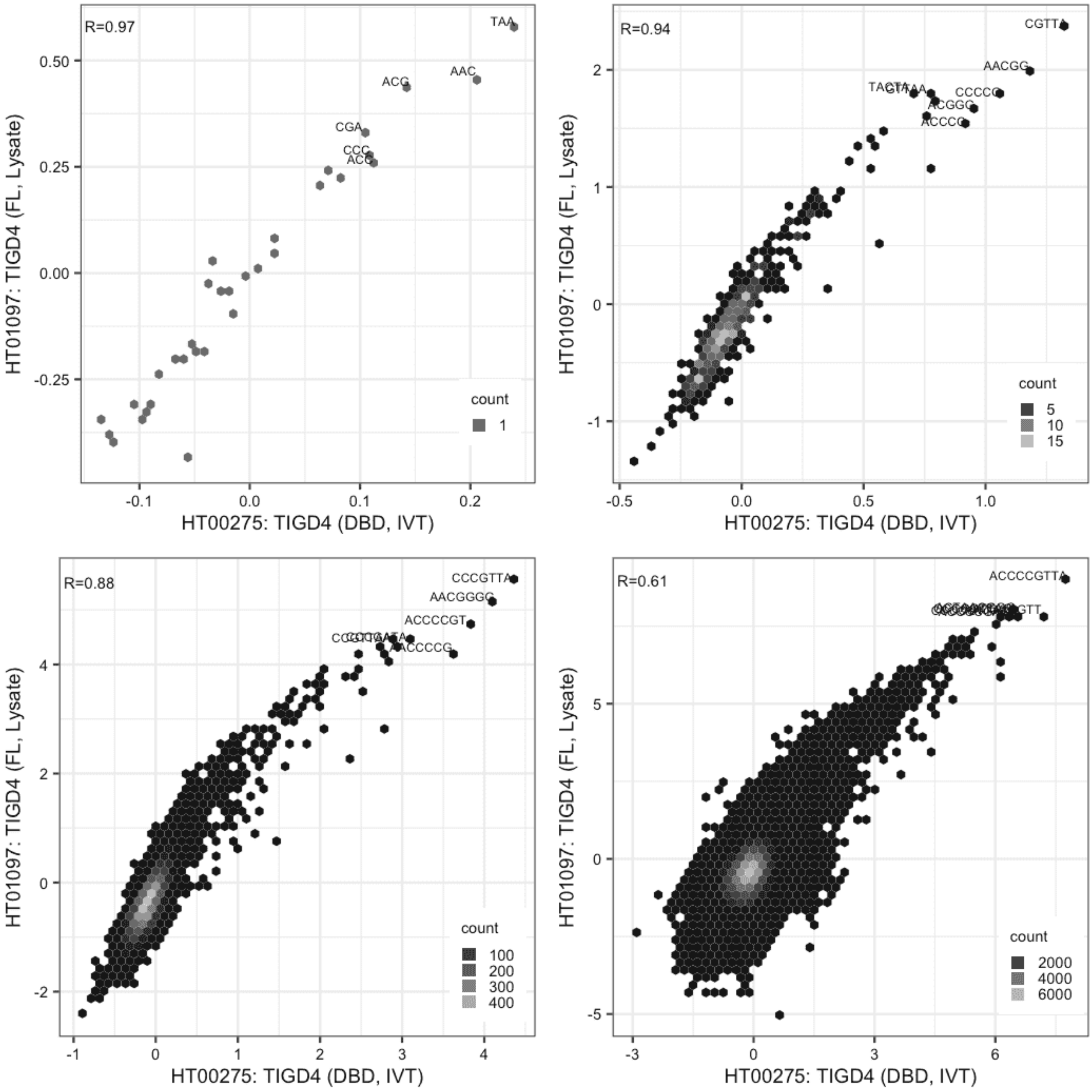

**TIGD4**

Lysate vs IVT, different constructs

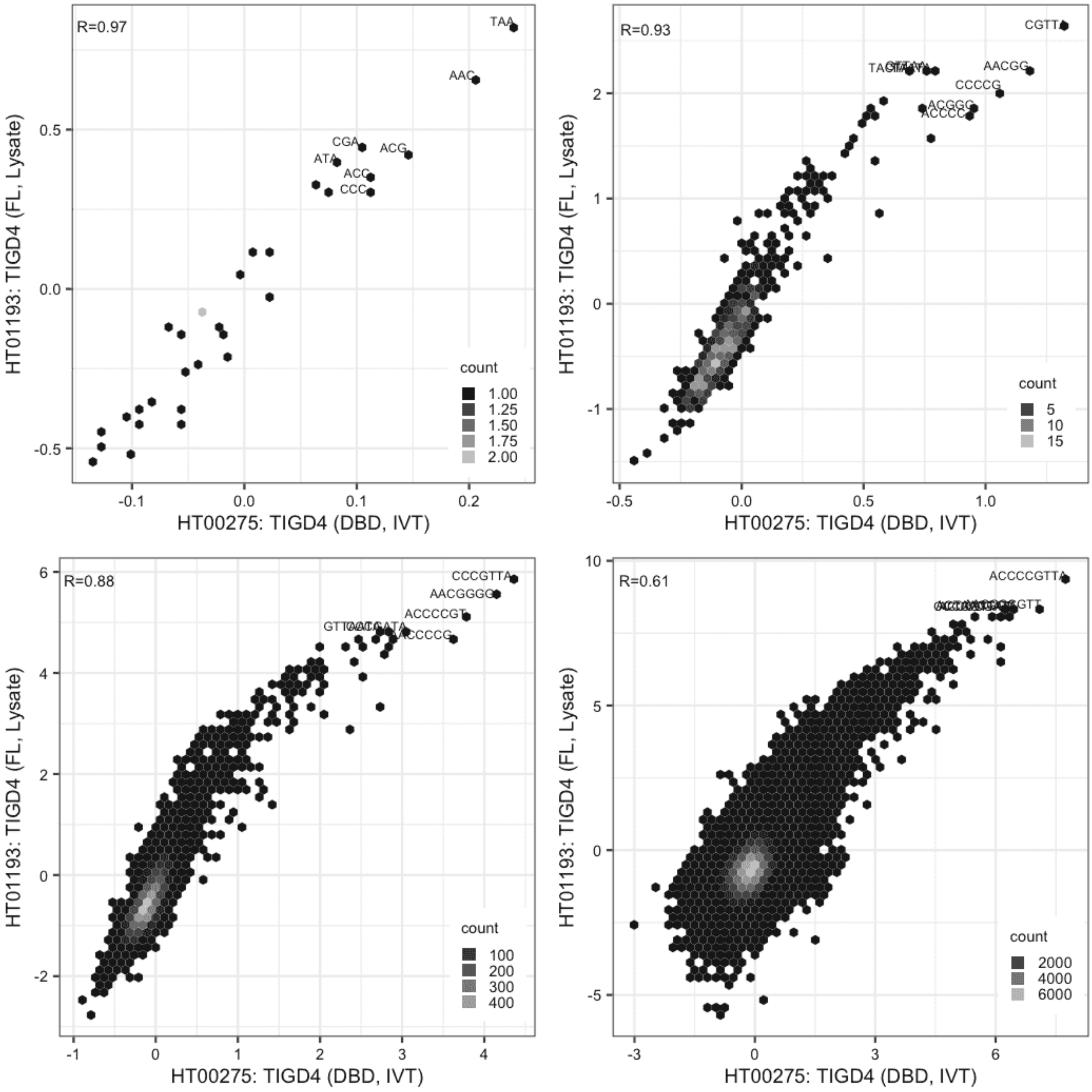

**TIGD4**

IVT vs GFPIVT, different constructs

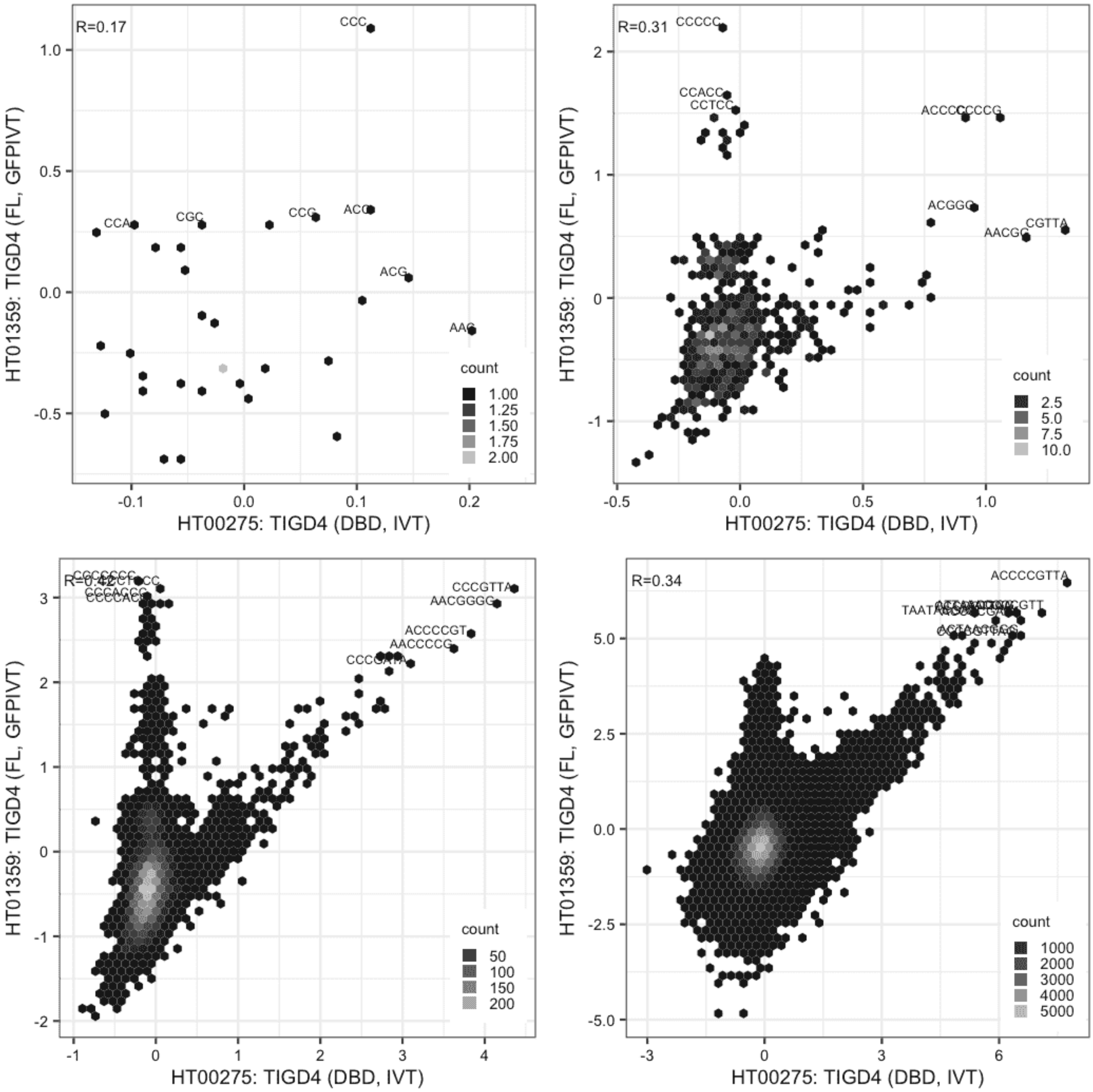

### TIGD4

Lysate vs GFPIVT, different constructs

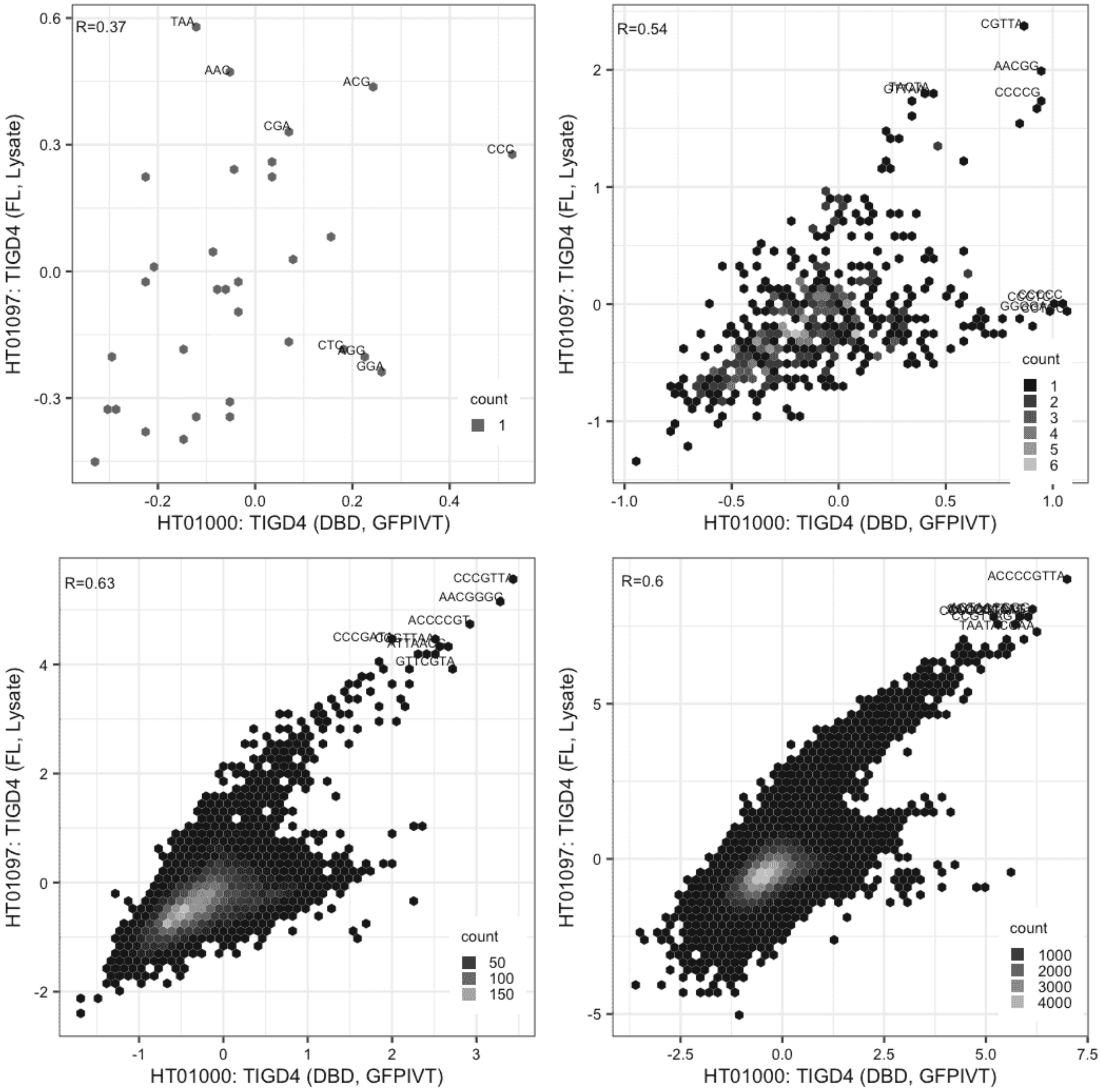

**TIGD4**

Lysate vs GFPIVT, different constructs

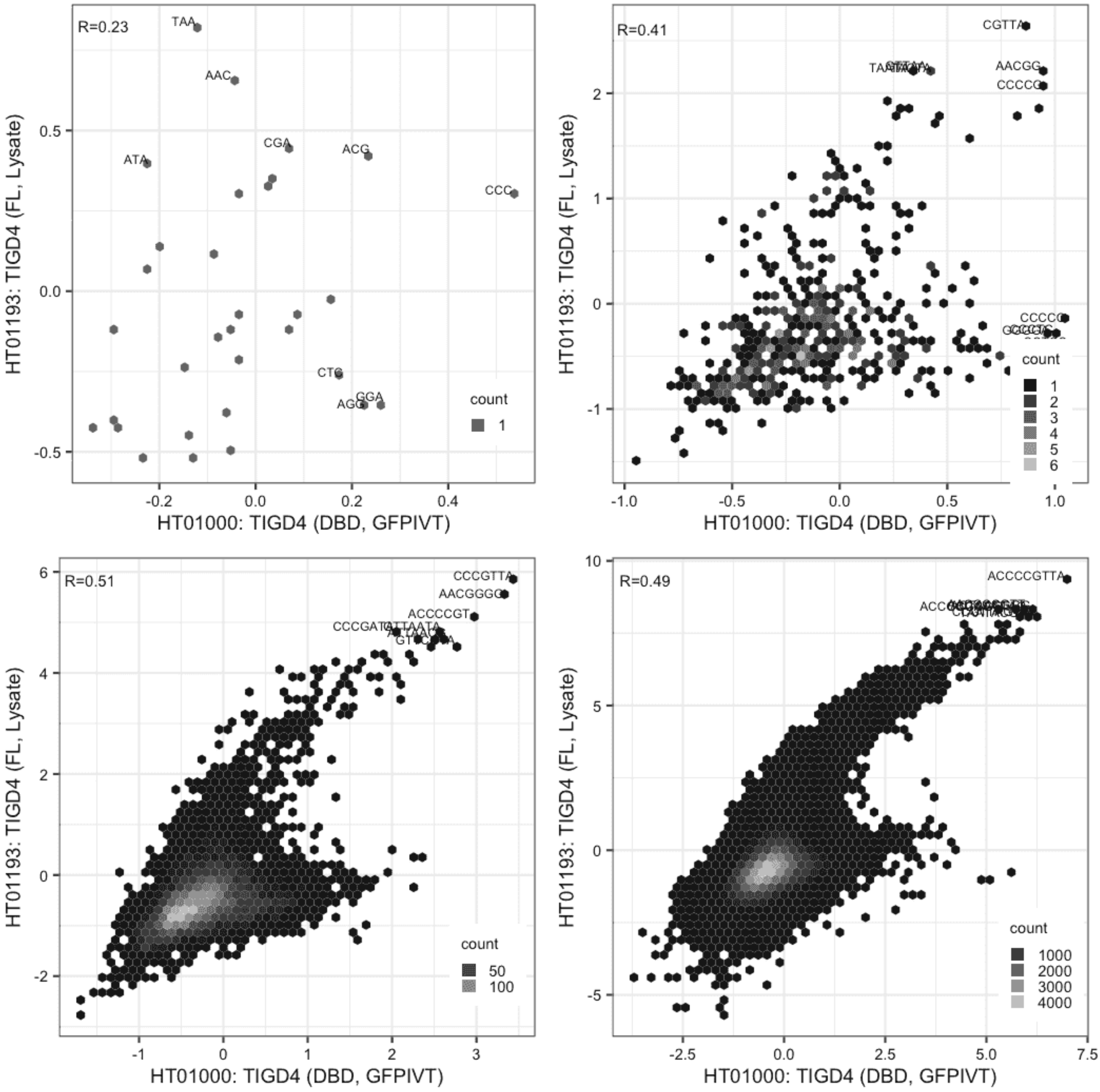

### TIGD4

GFPIVT vs GFPIVT, different constructs

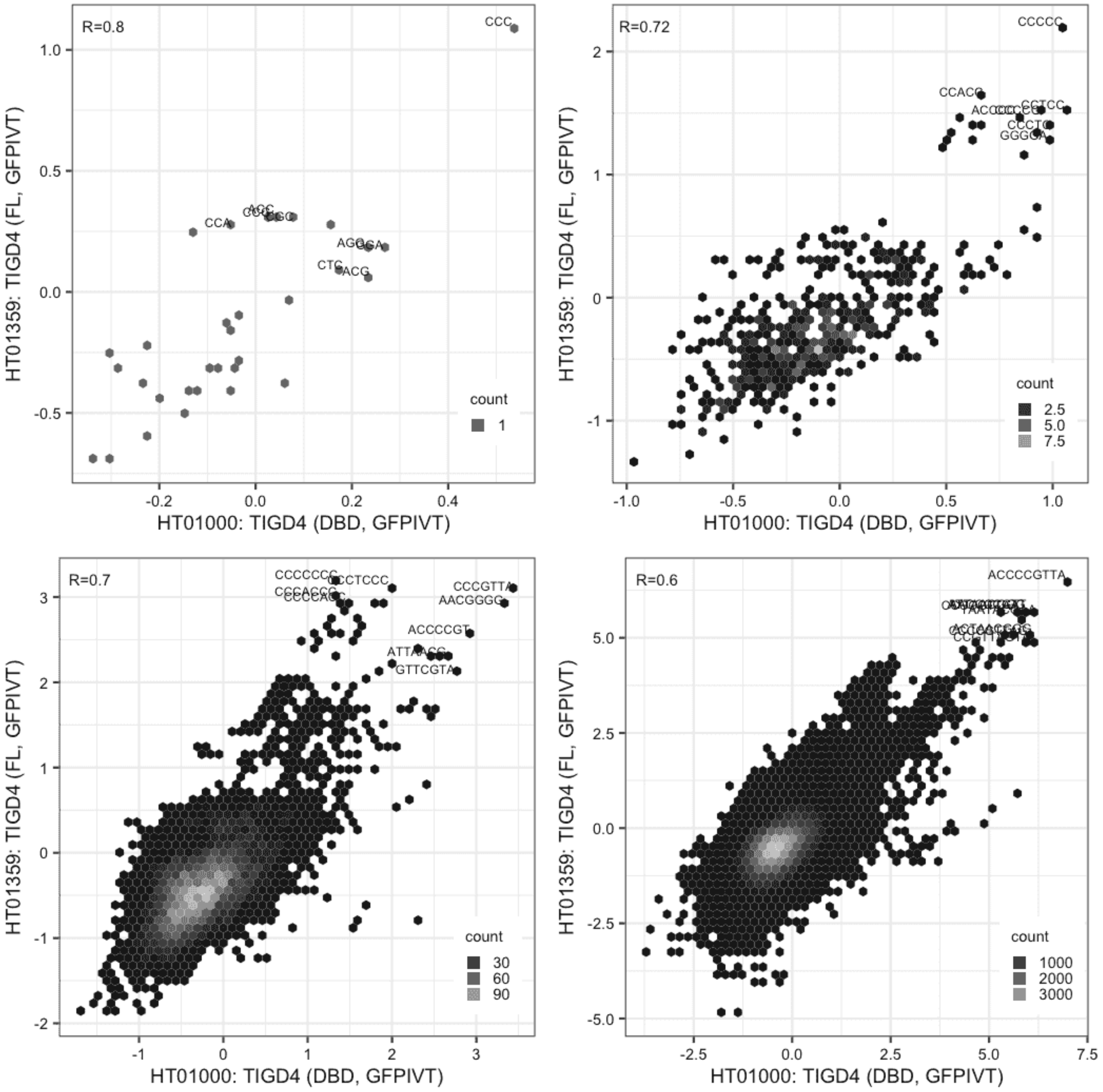

#### TIGD4

Lysate vs Lysate, same construct

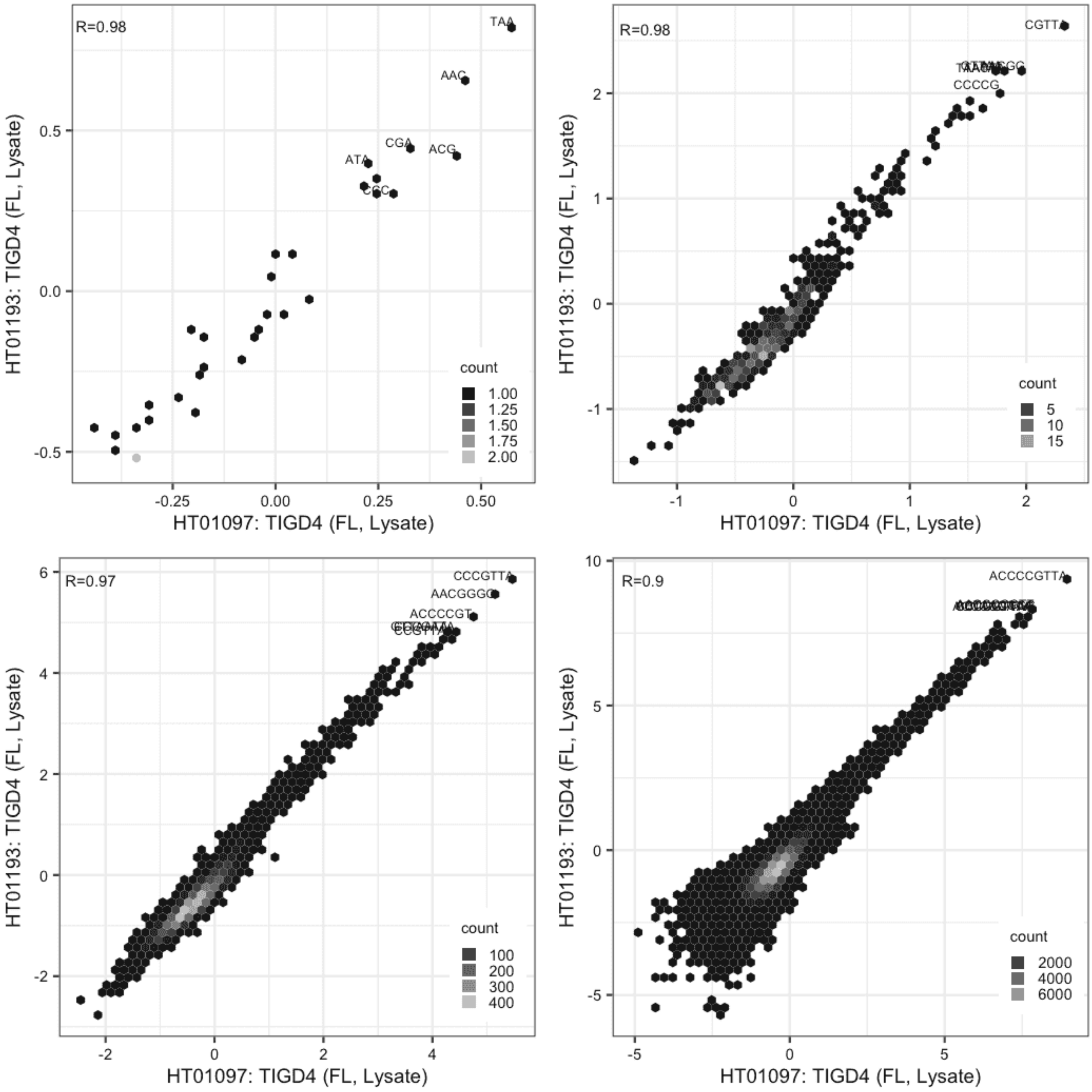

### TIGD4

Lysate vs GFPIVT, same construct

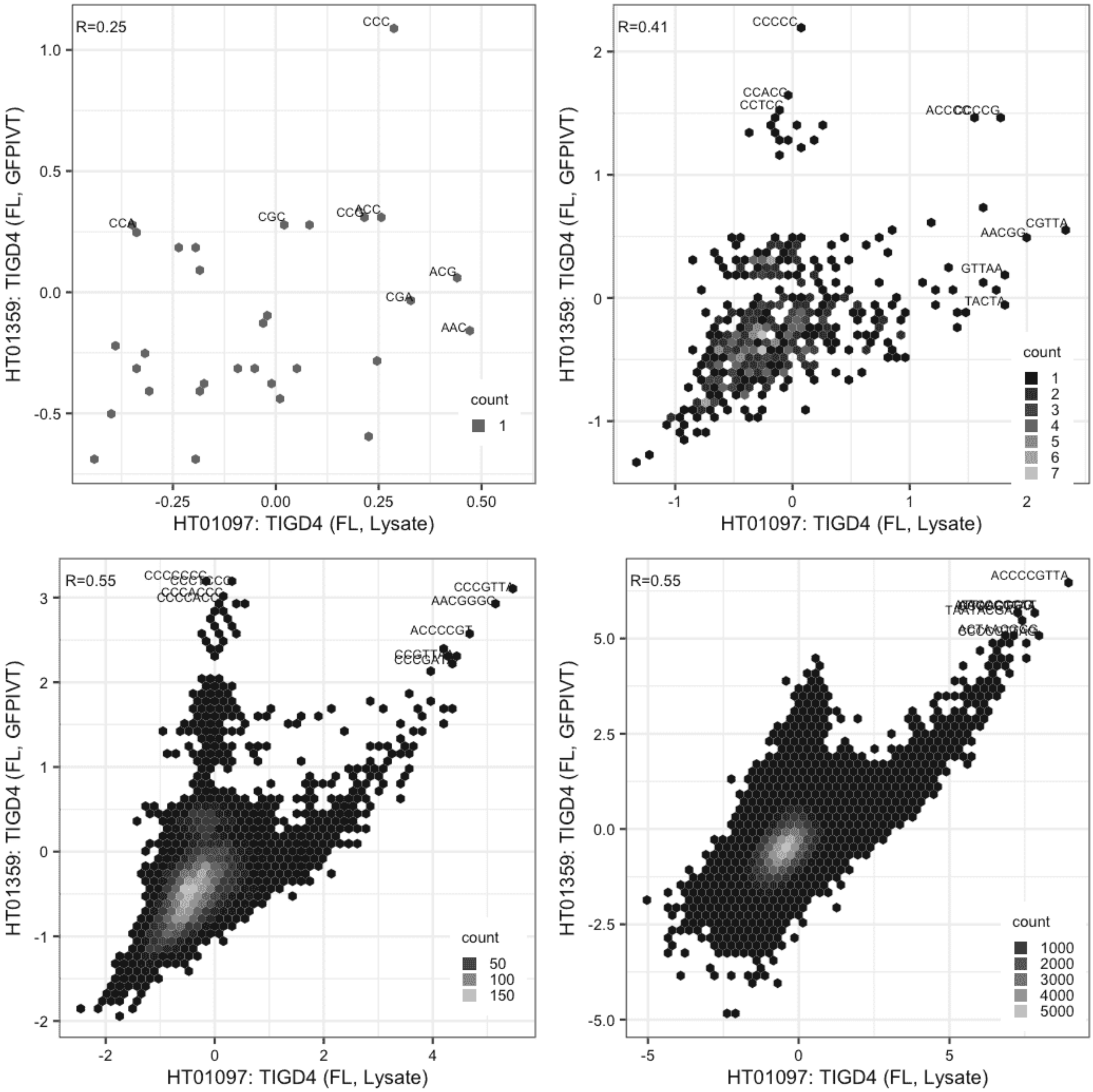

#### TIGD4

Lysate vs GFPIVT, same construct

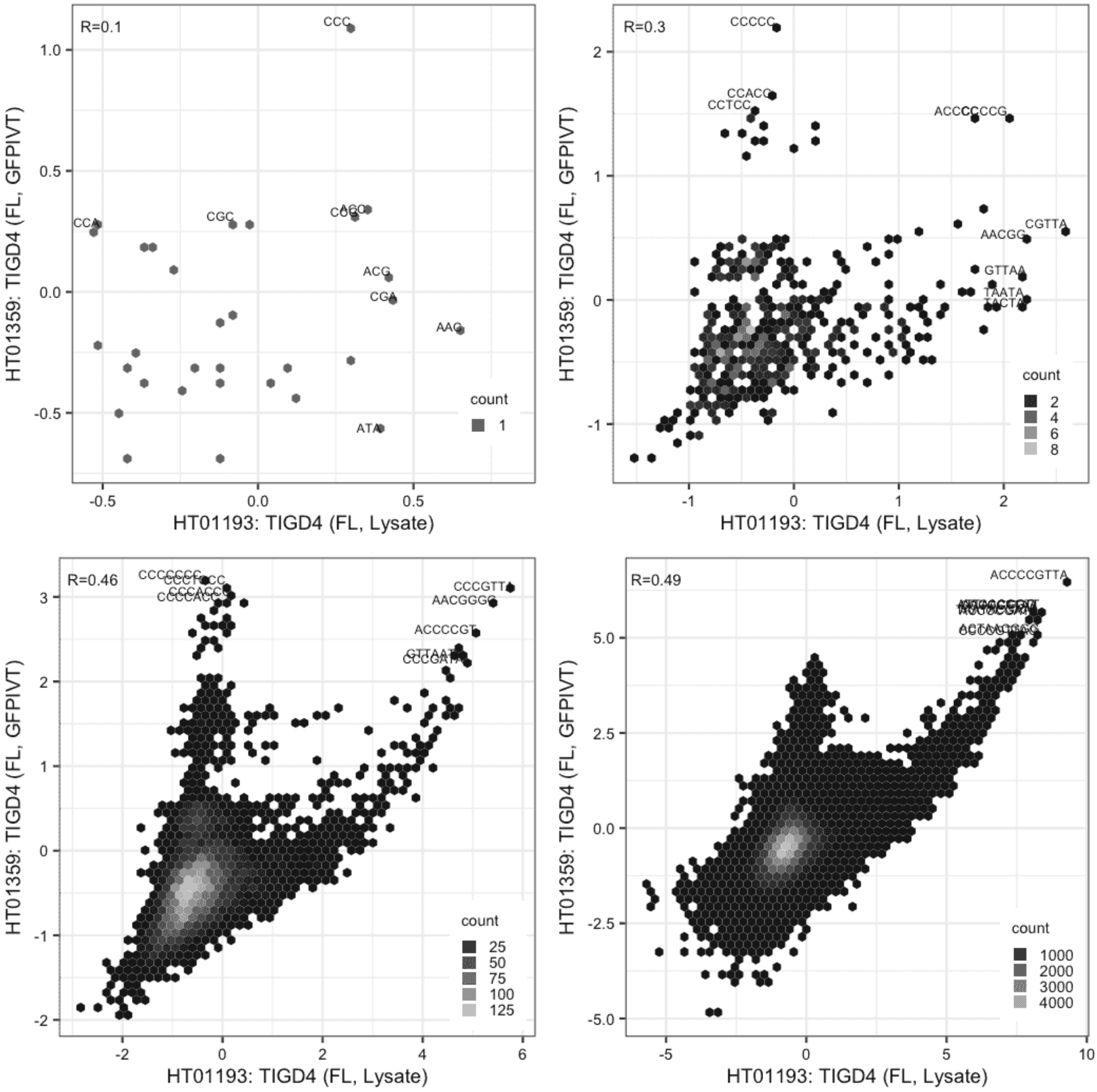

#### TIGD5

IVT vs GFPIVT, same construct

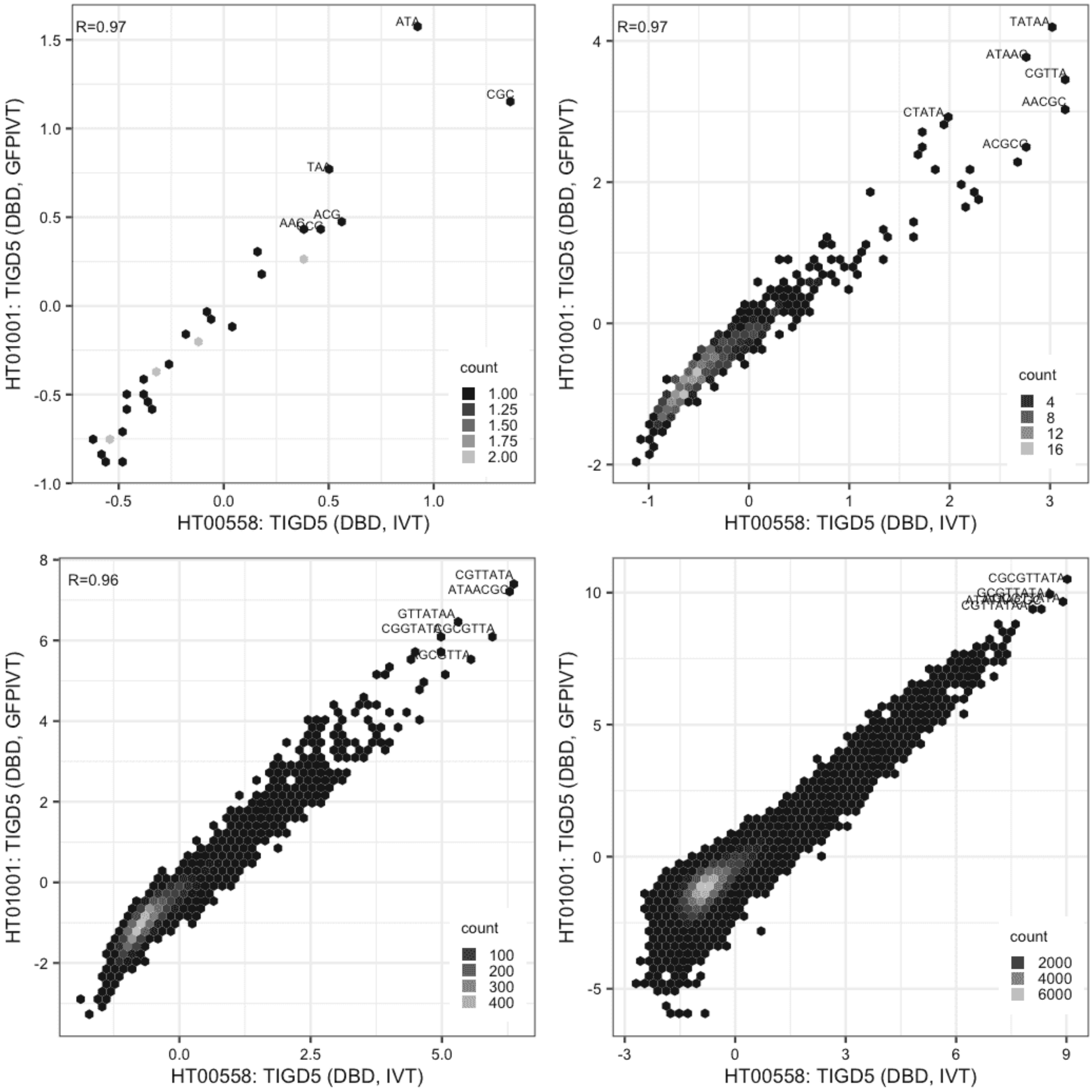

**TIGD5**

Lysate vs IVT, different constructs

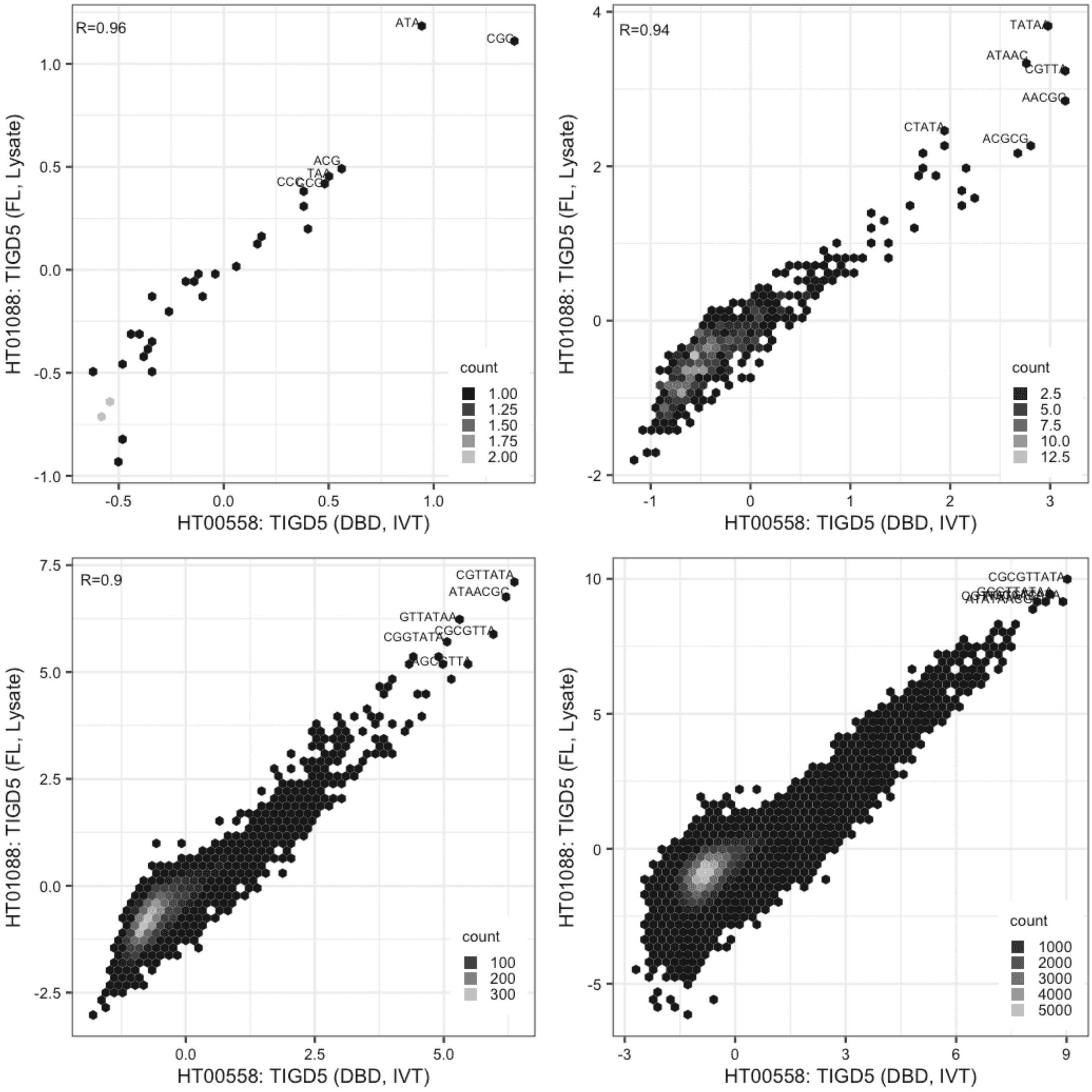

### TIGD5

Lysate vs IVT, different constructs

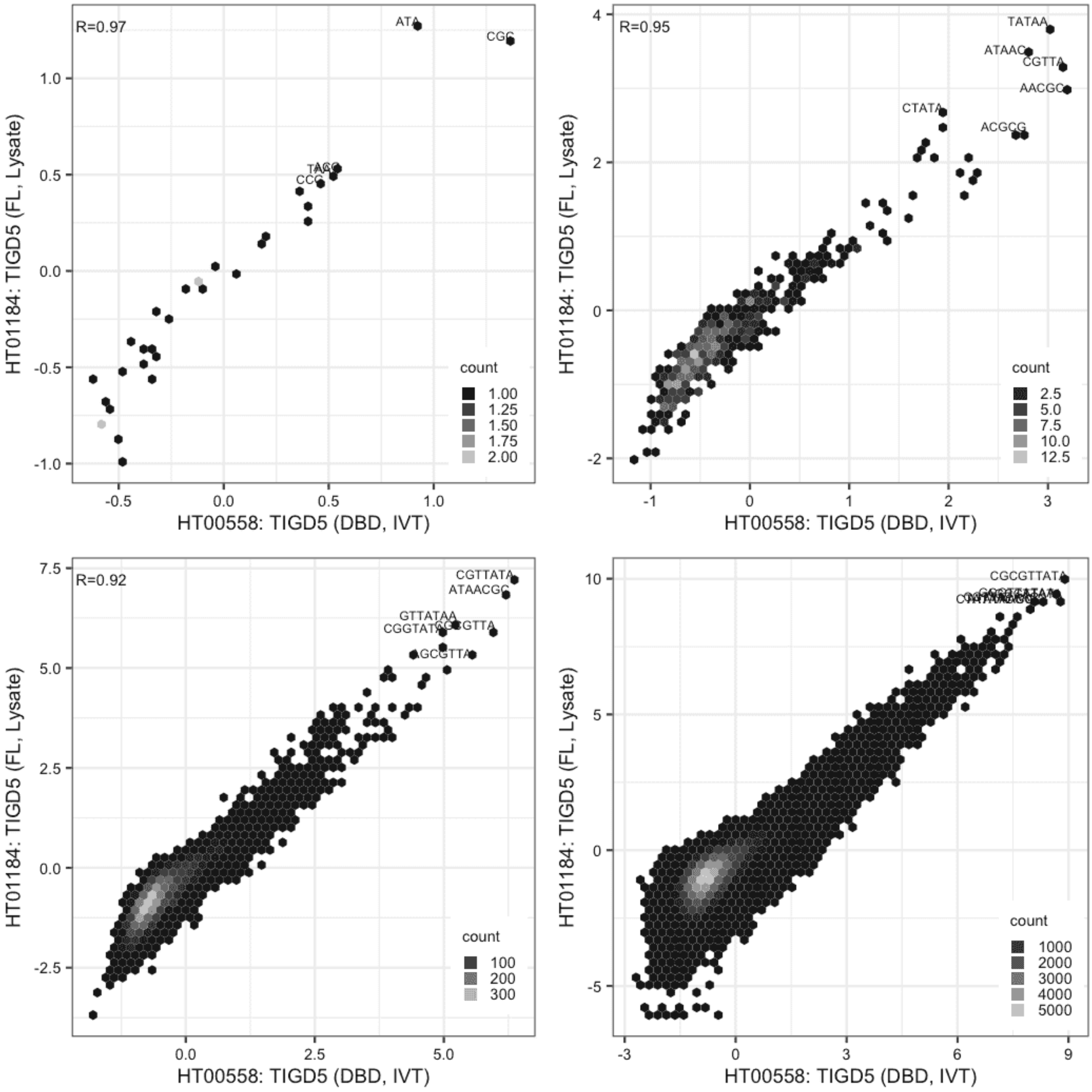

**TIGD5**

IVT vs GFPIVT, different constructs

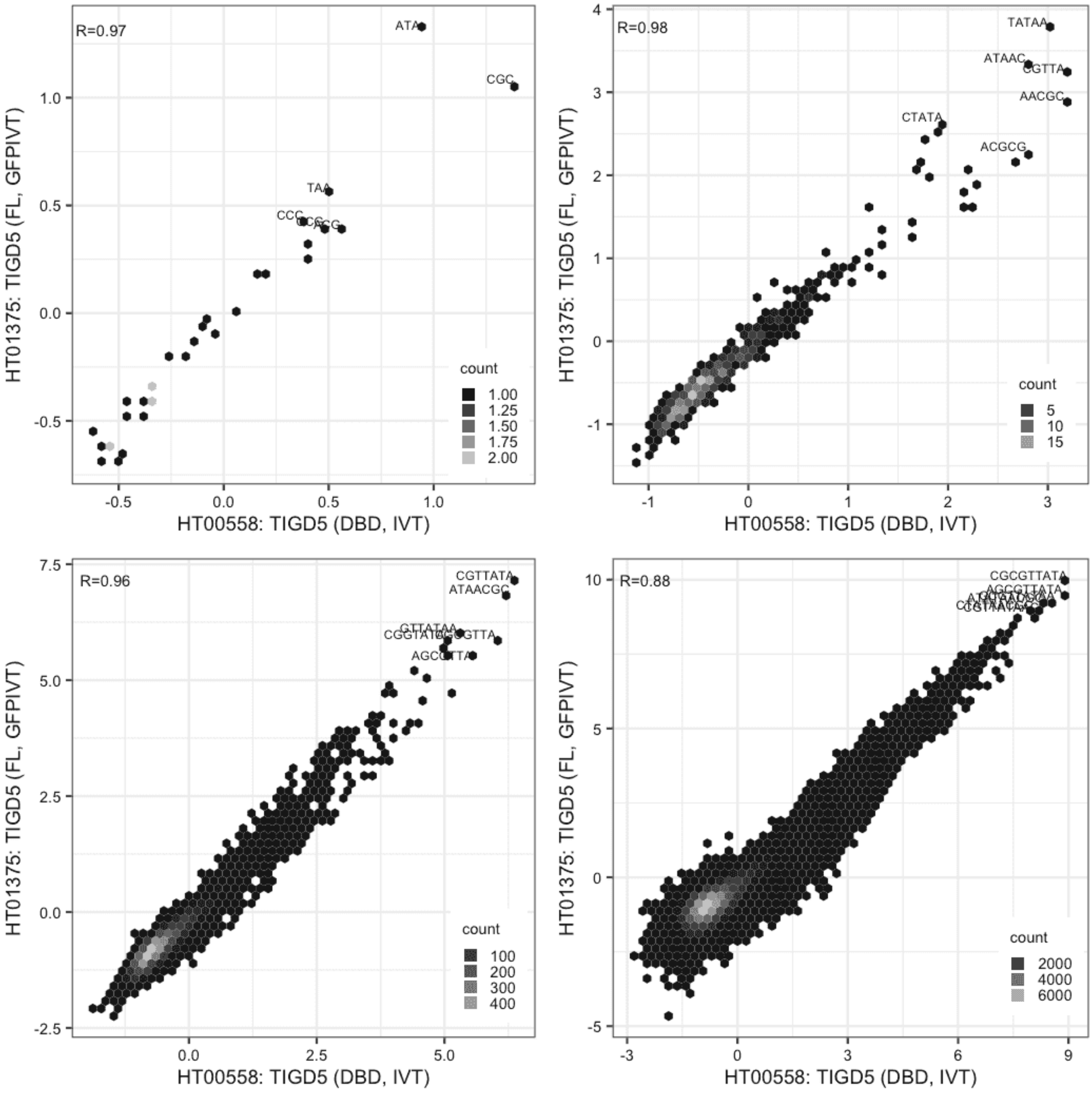

**TIGD5**

Lysate vs GFPIVT, different constructs

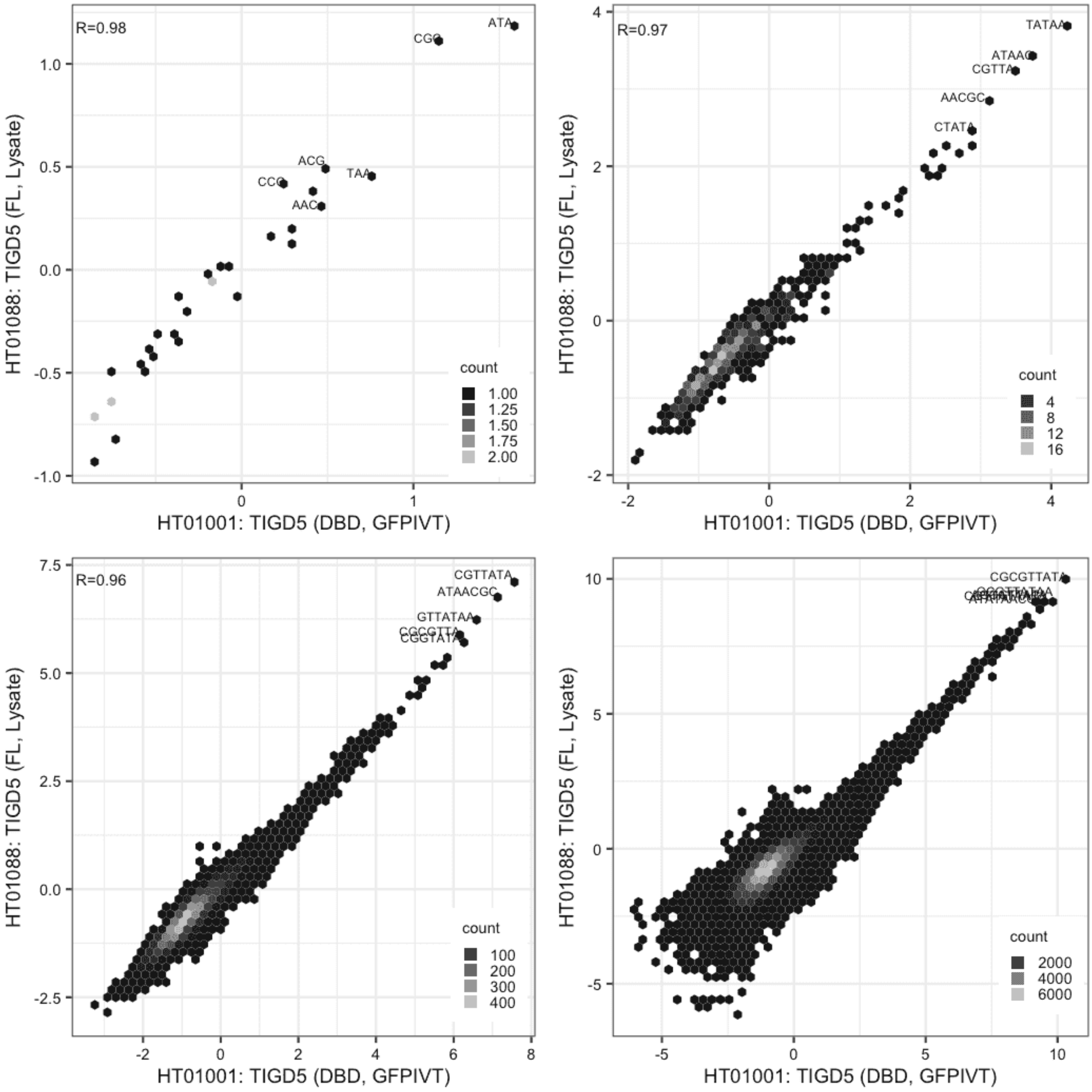

**TIGD5**

Lysate vs GFPIVT, different constructs

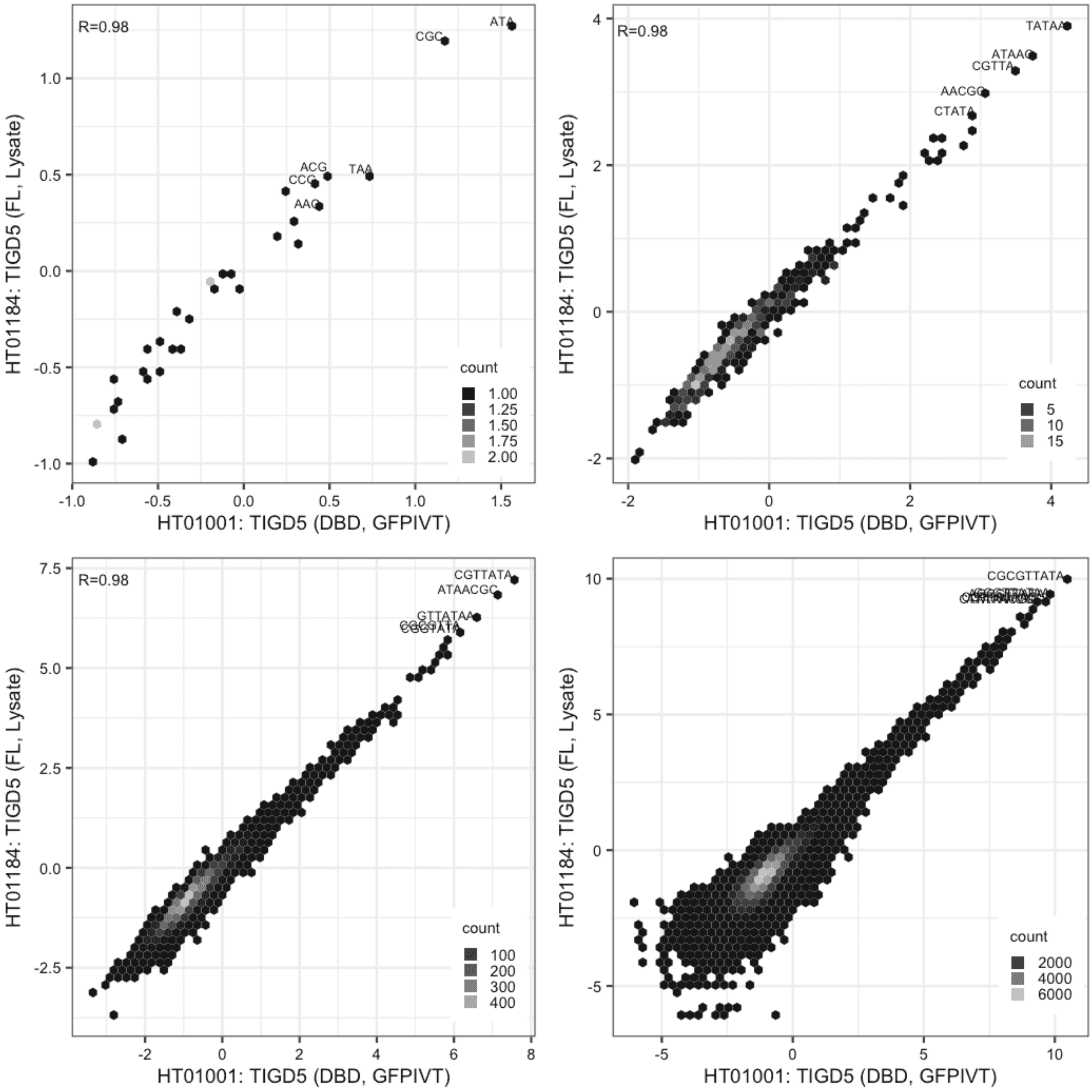

#### TIGD5

GFPIVT vs GFPIVT, different constructs

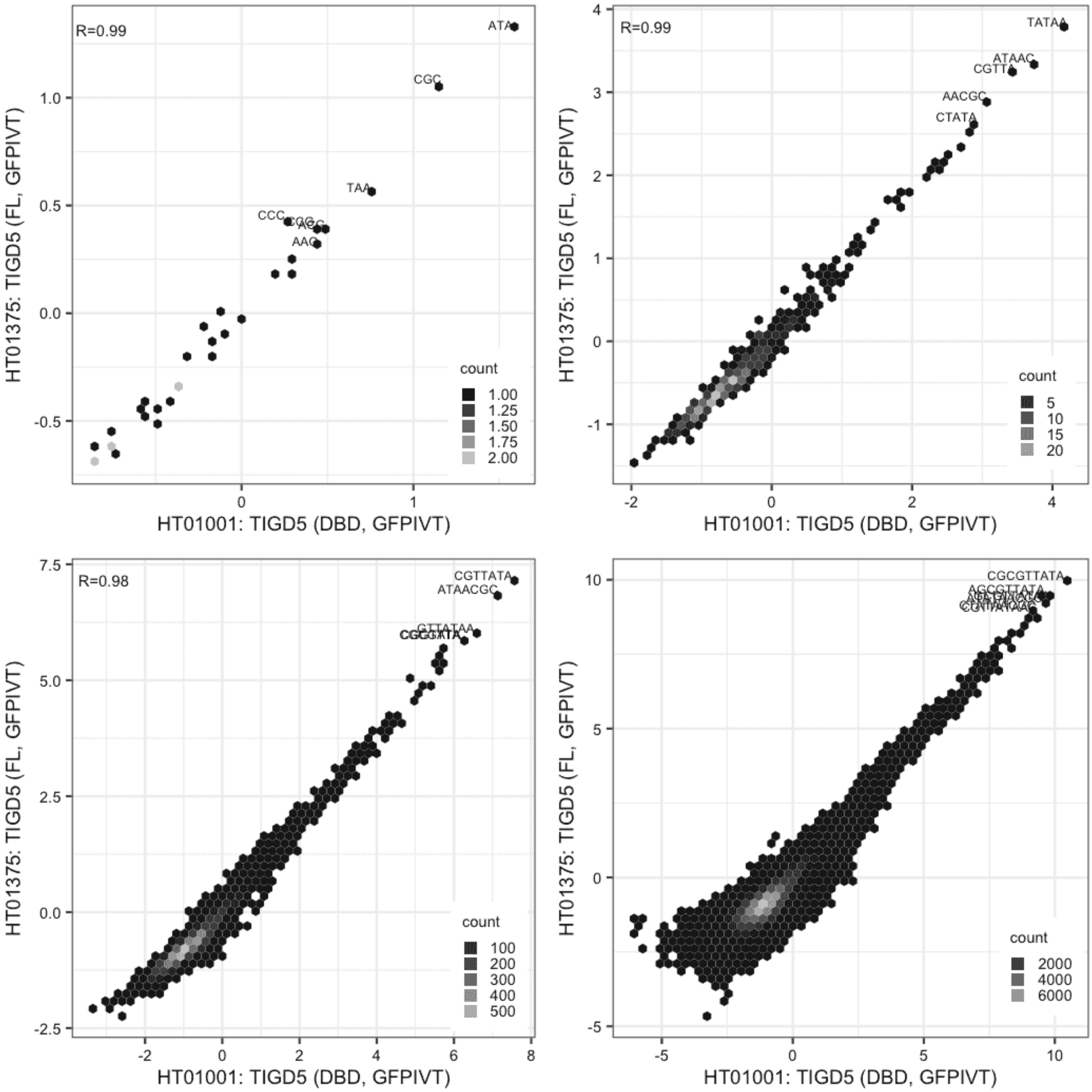

**TIGD5**

Lysate vs Lysate, same construct

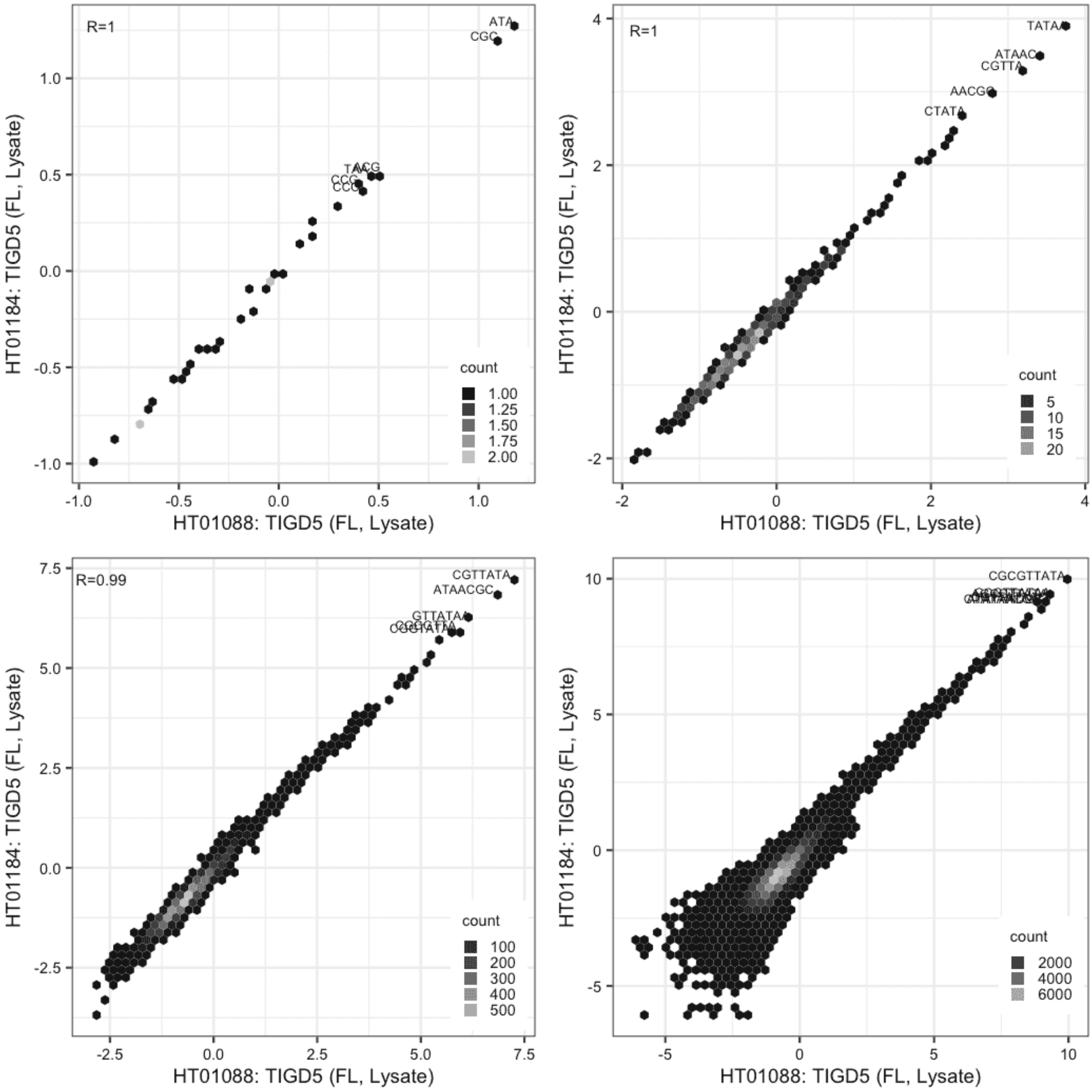

**TIGD5**

Lysate vs GFPIVT, same construct

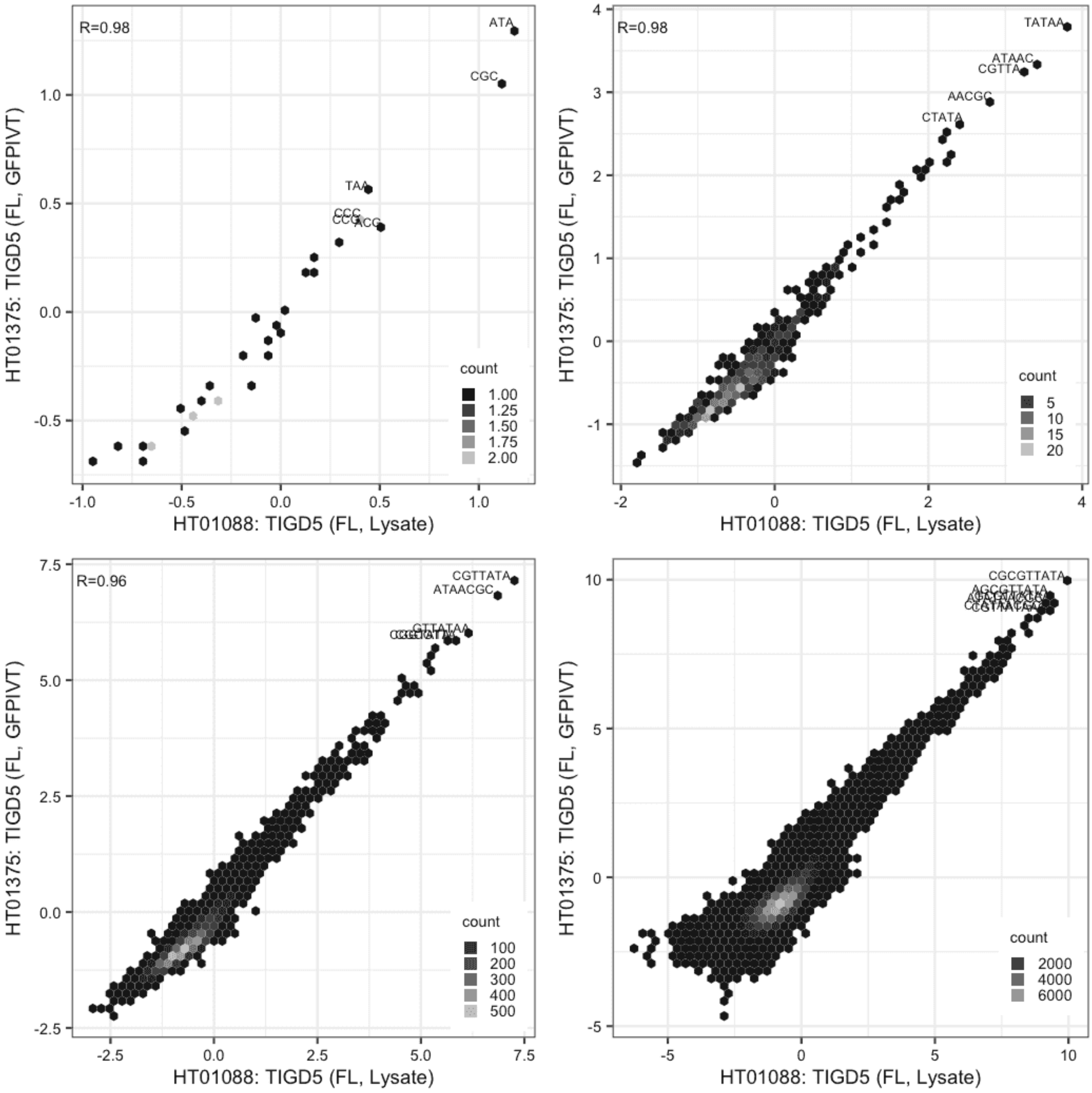

#### TIGD5

Lysate vs GFPIVT, same construct

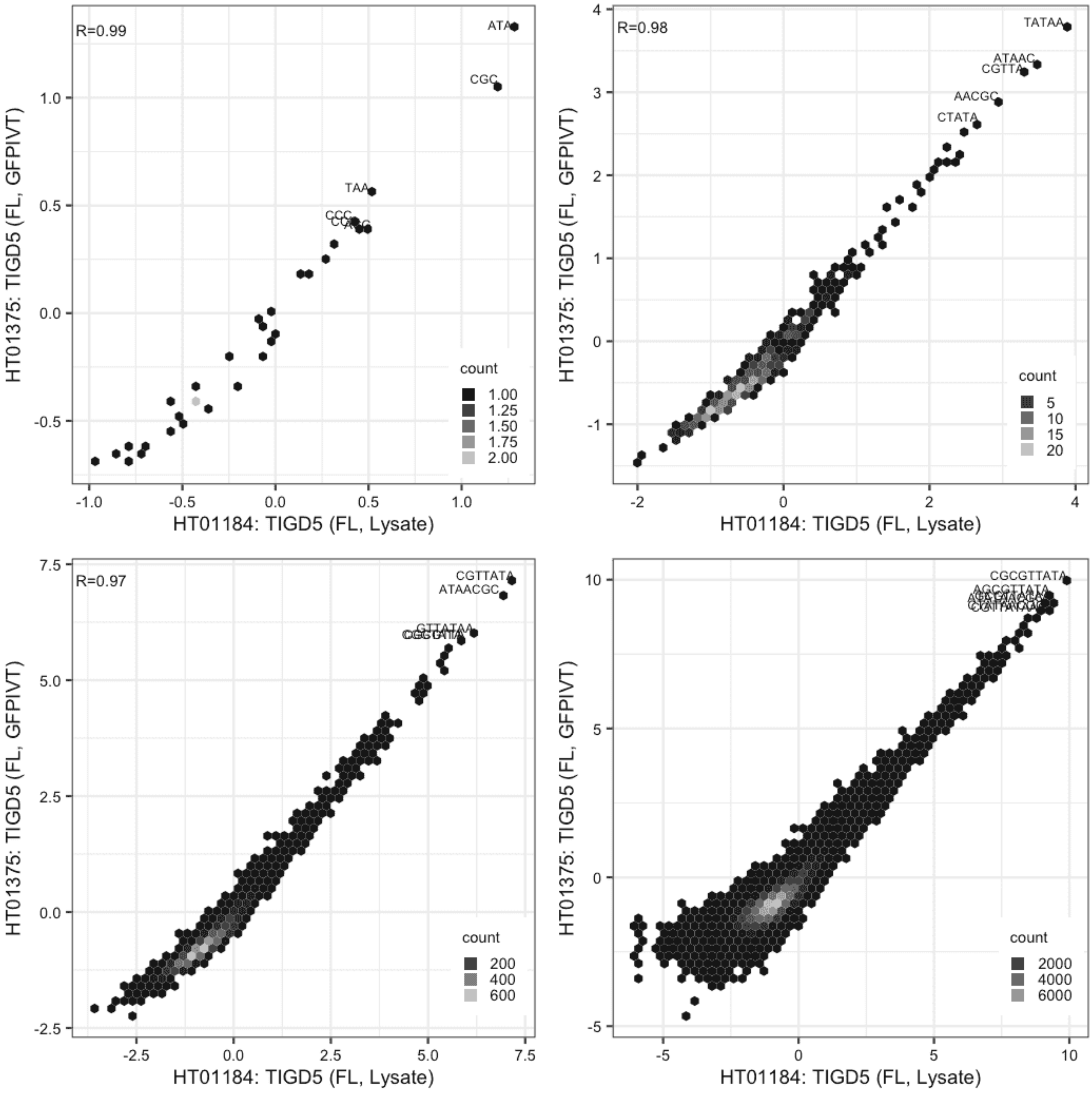

**TPRX1**

Lysate vs IVT, different constructs

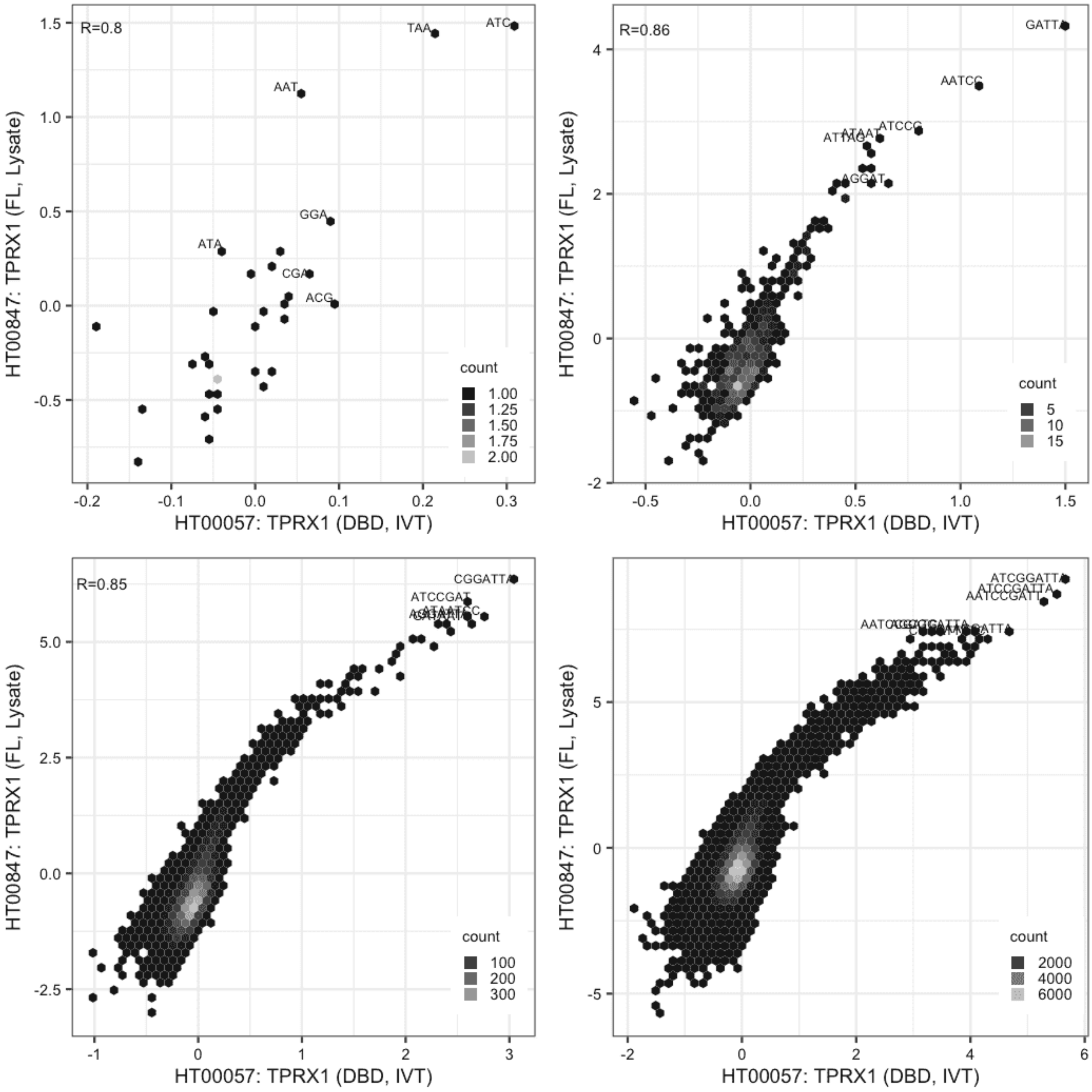

**TPRX1**

IVT vs GFPIVT, same construct

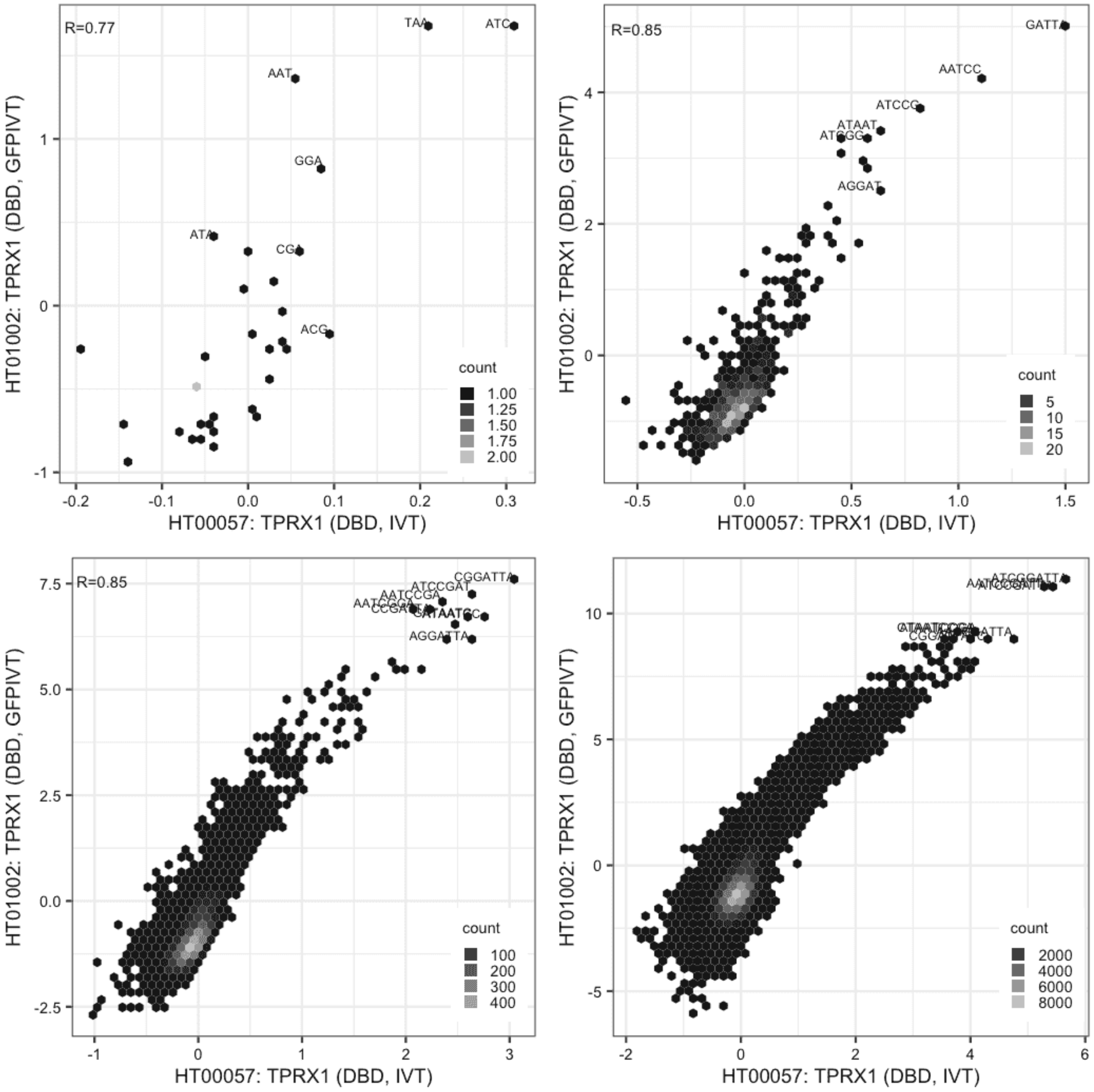

**TPRX1**

IVT vs GFPIVT, different constructs

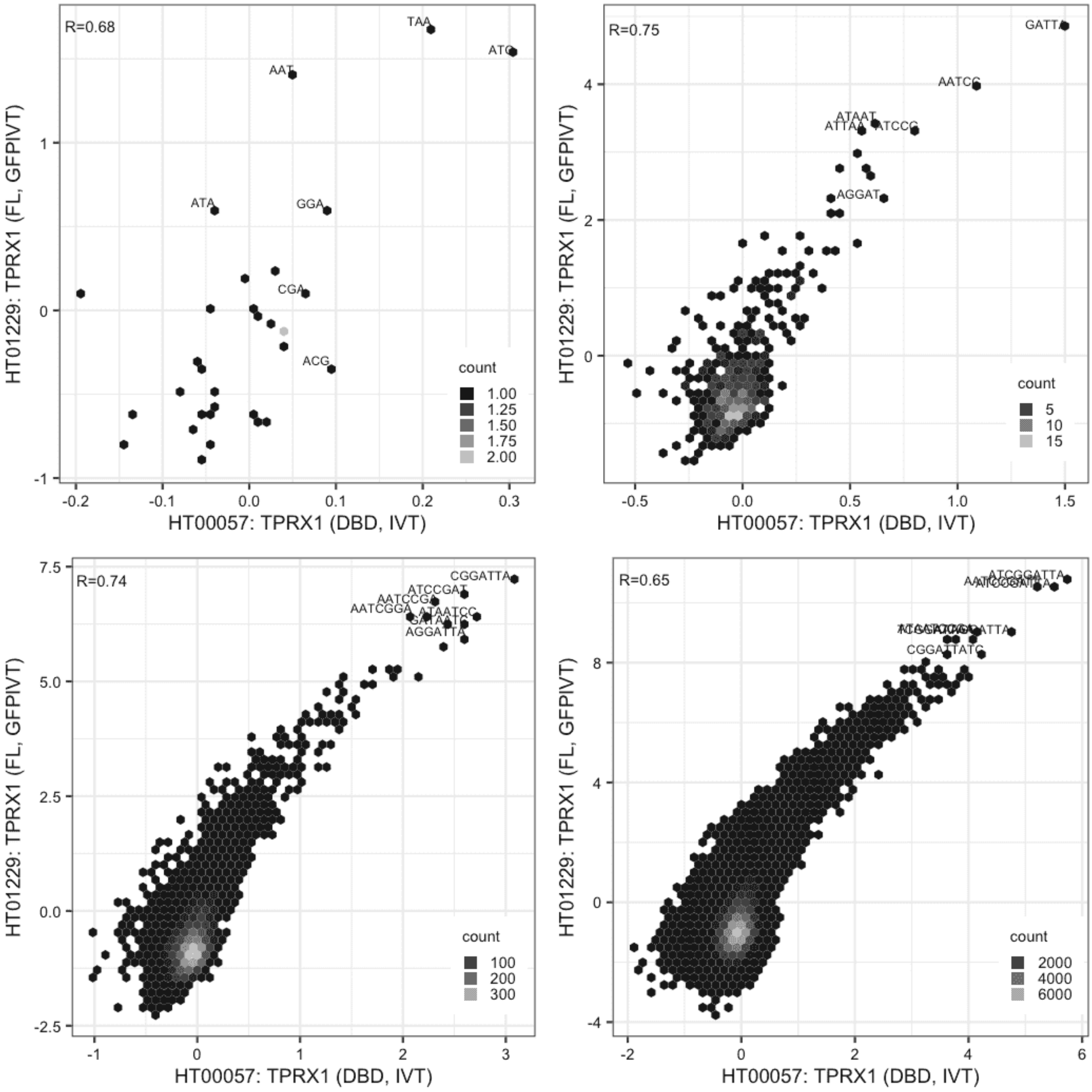

#### TPRX1

Lysate vs GFPIVT, different constructs

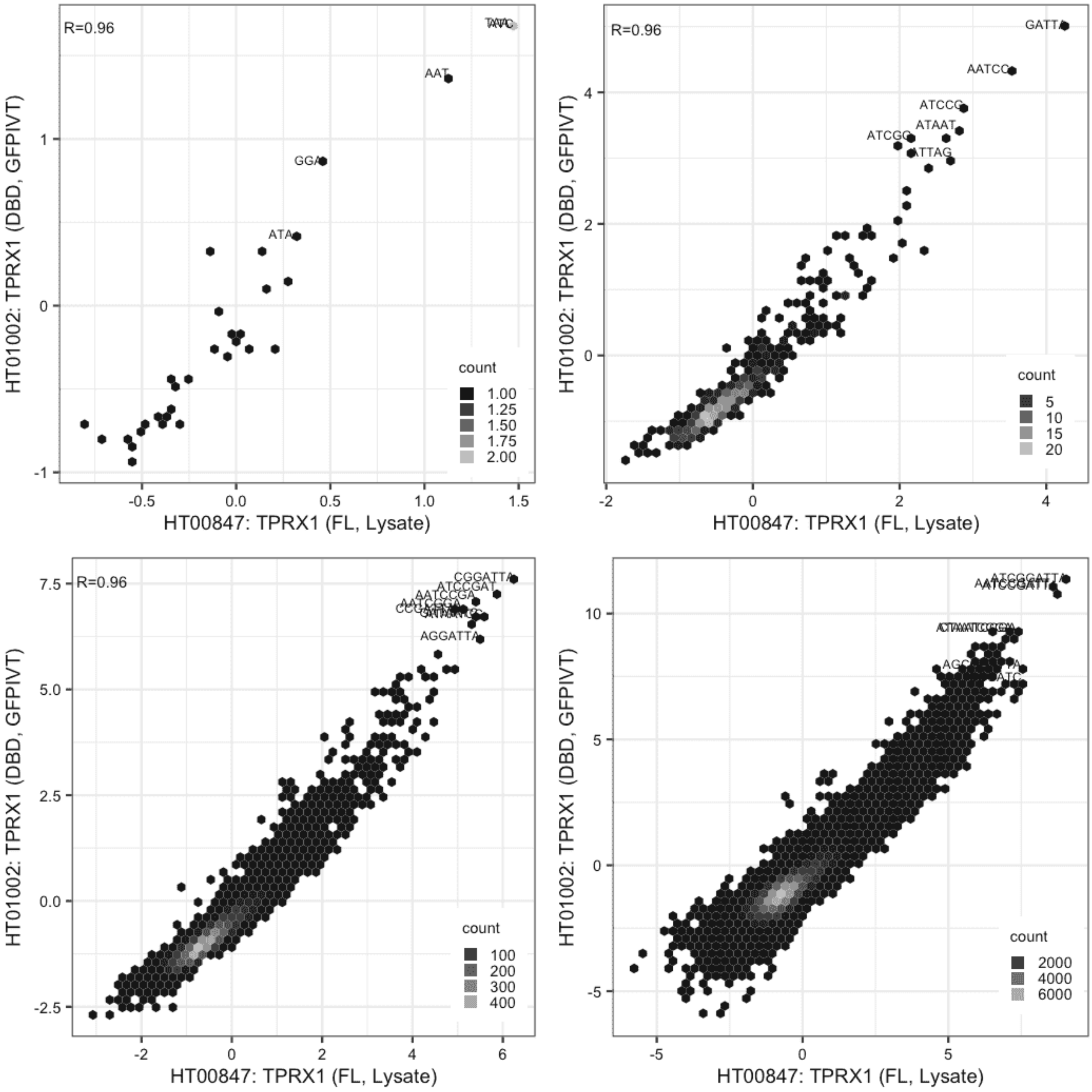

**TPRX1**

Lysate vs GFPIVT, same construct

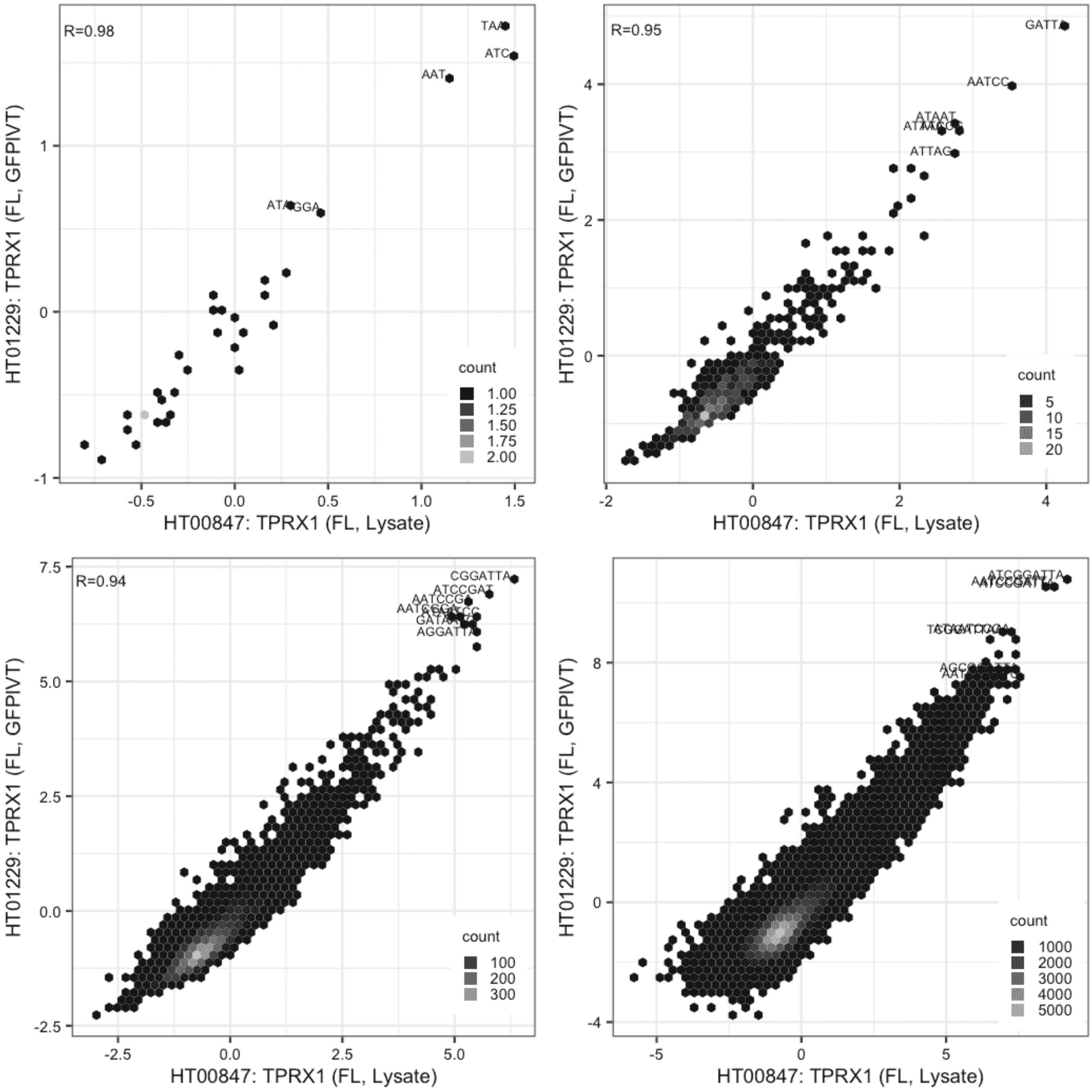

### TPRX1

GFPIVT vs GFPIVT, different constructs

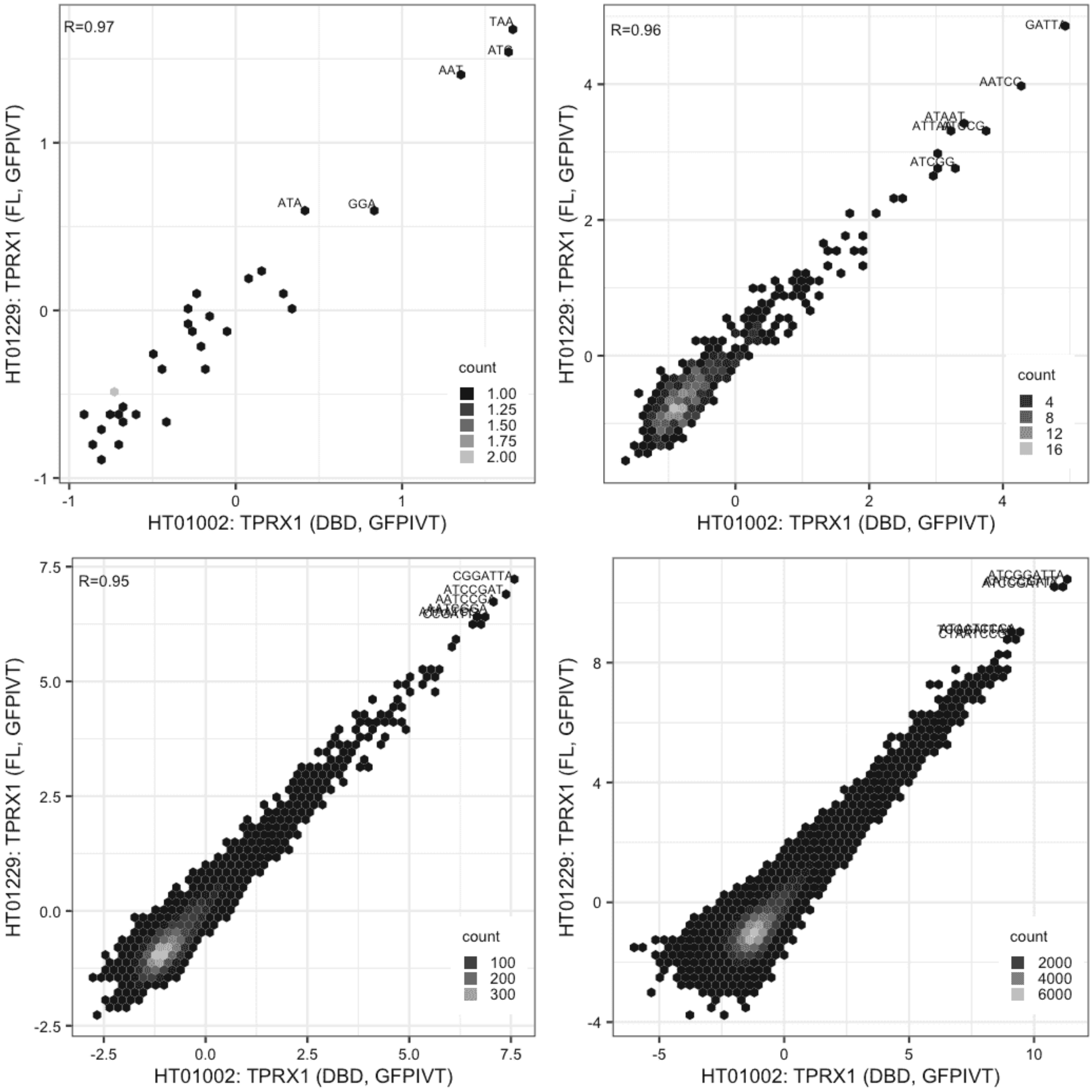

#### USF3

Lysate vs IVT, different constructs

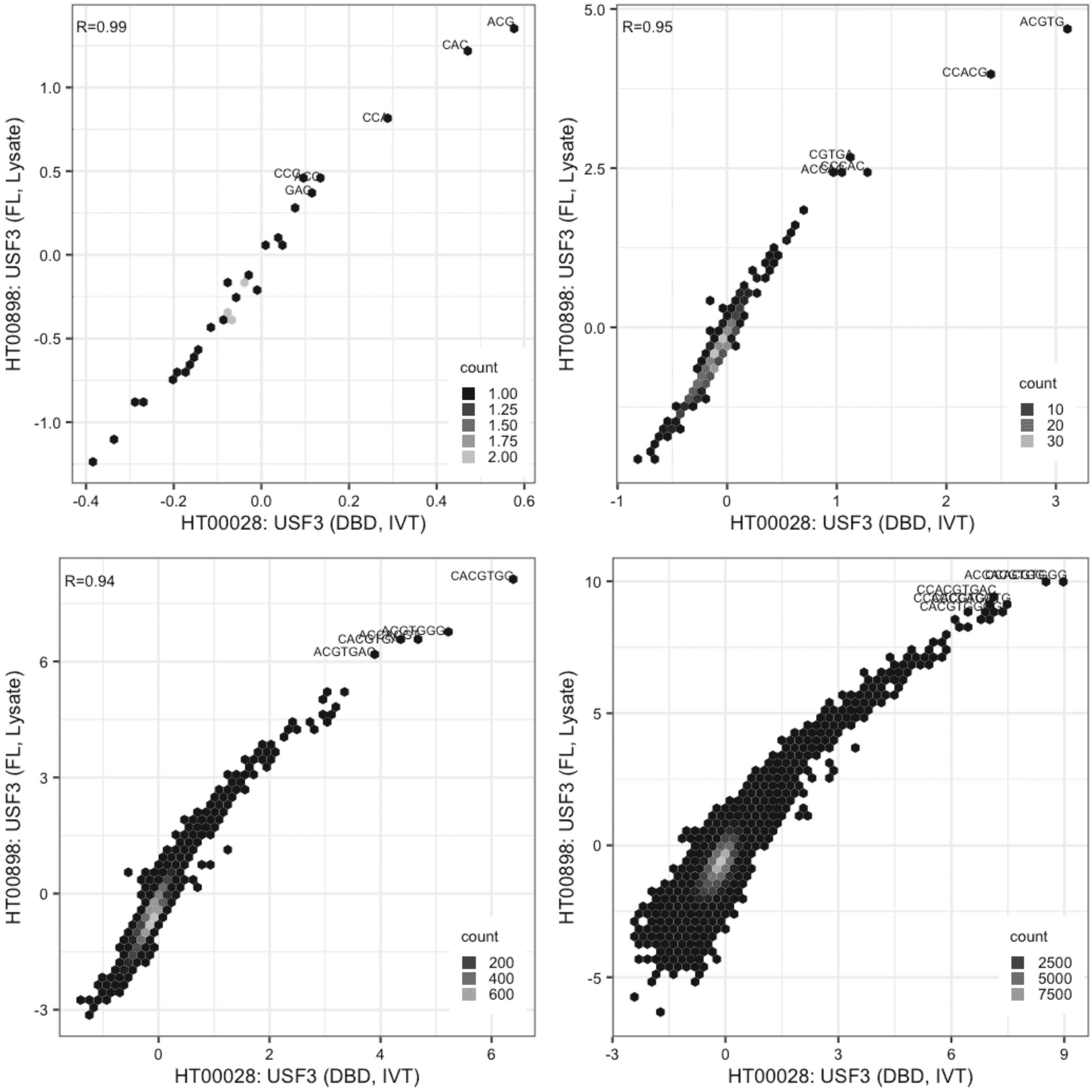

**USF3**

IVT vs GFPIVT, same construct

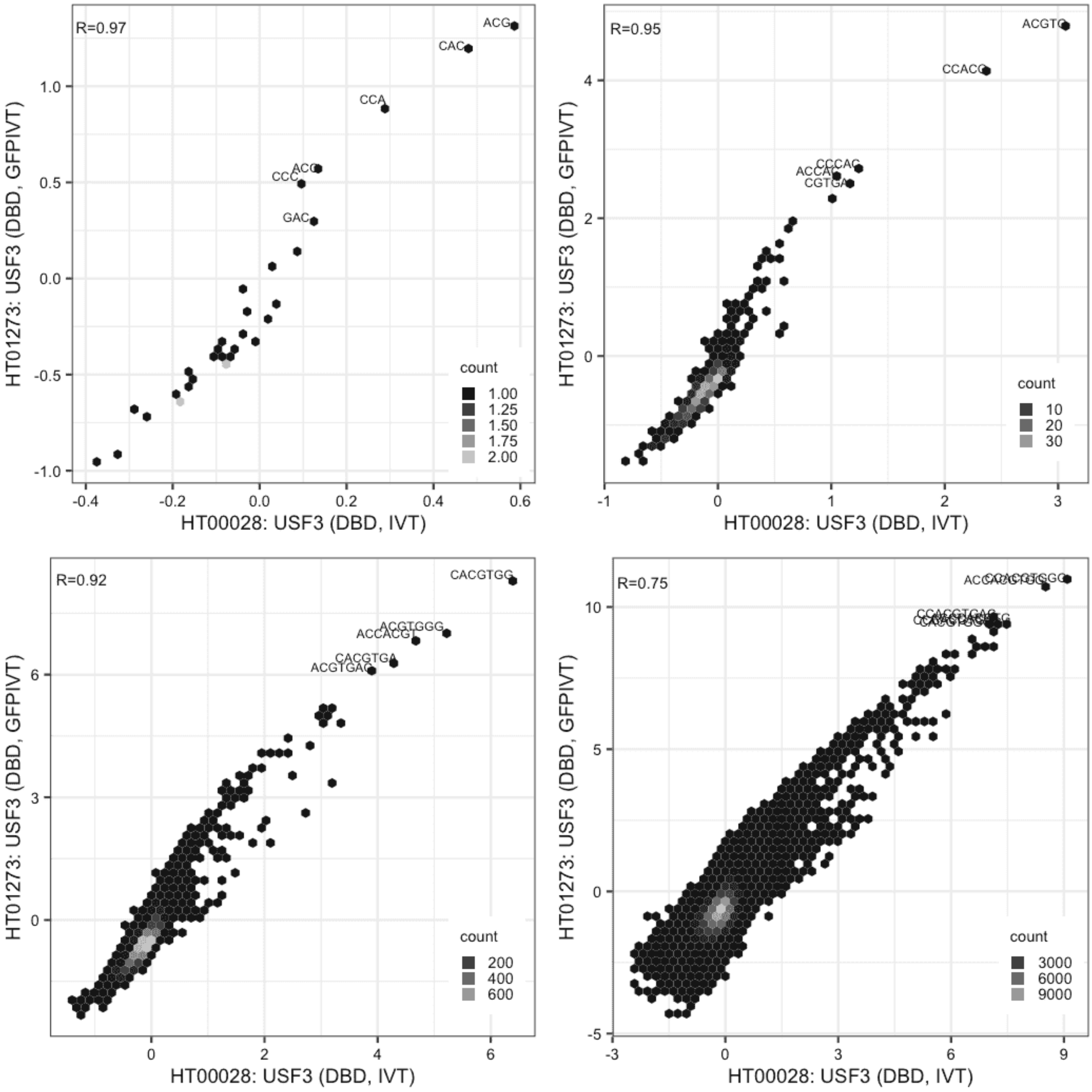

#### USF3

Lysate vs GFPIVT, different constructs

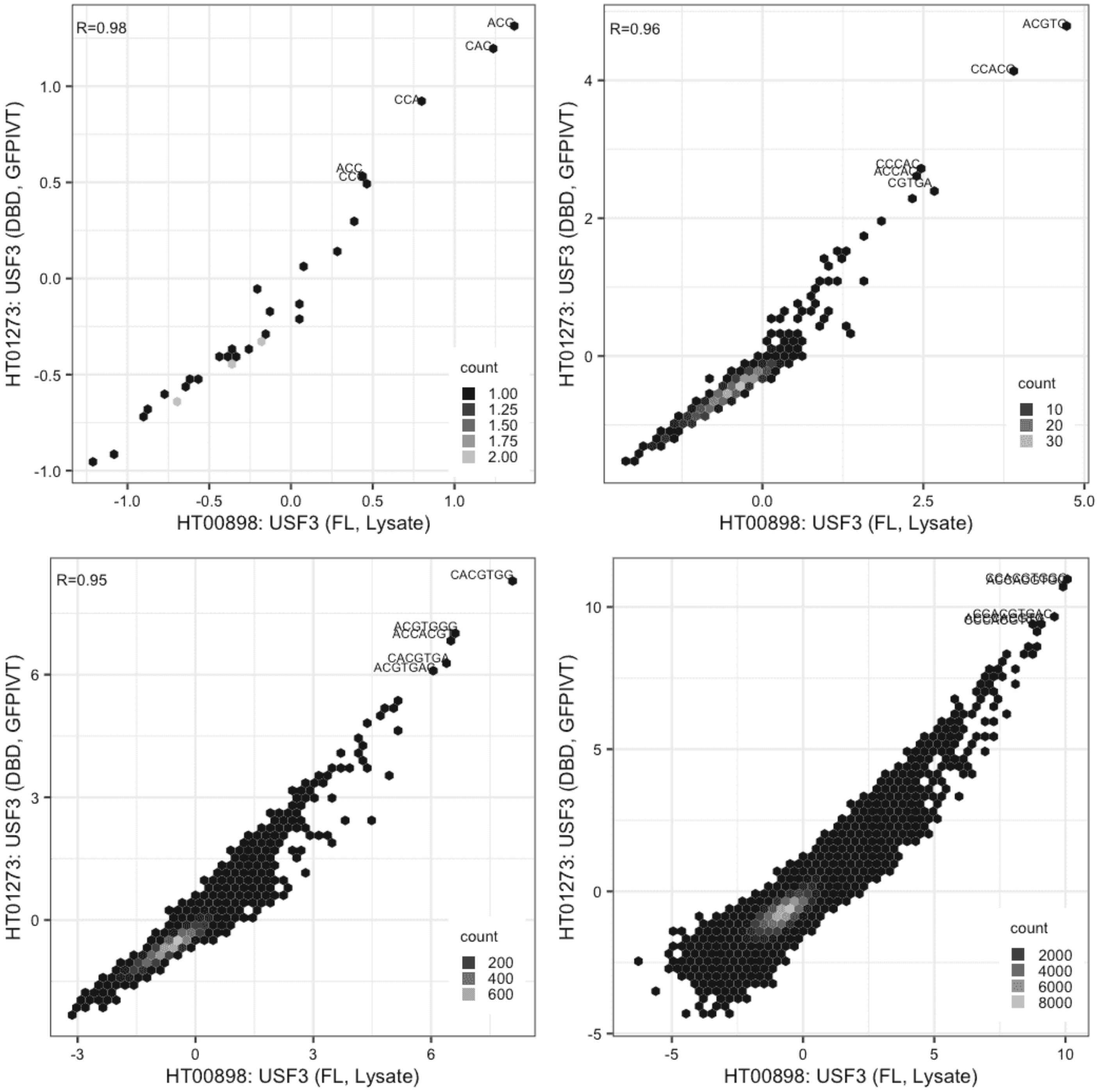

#### VDR

IVT vs IVT, same construct

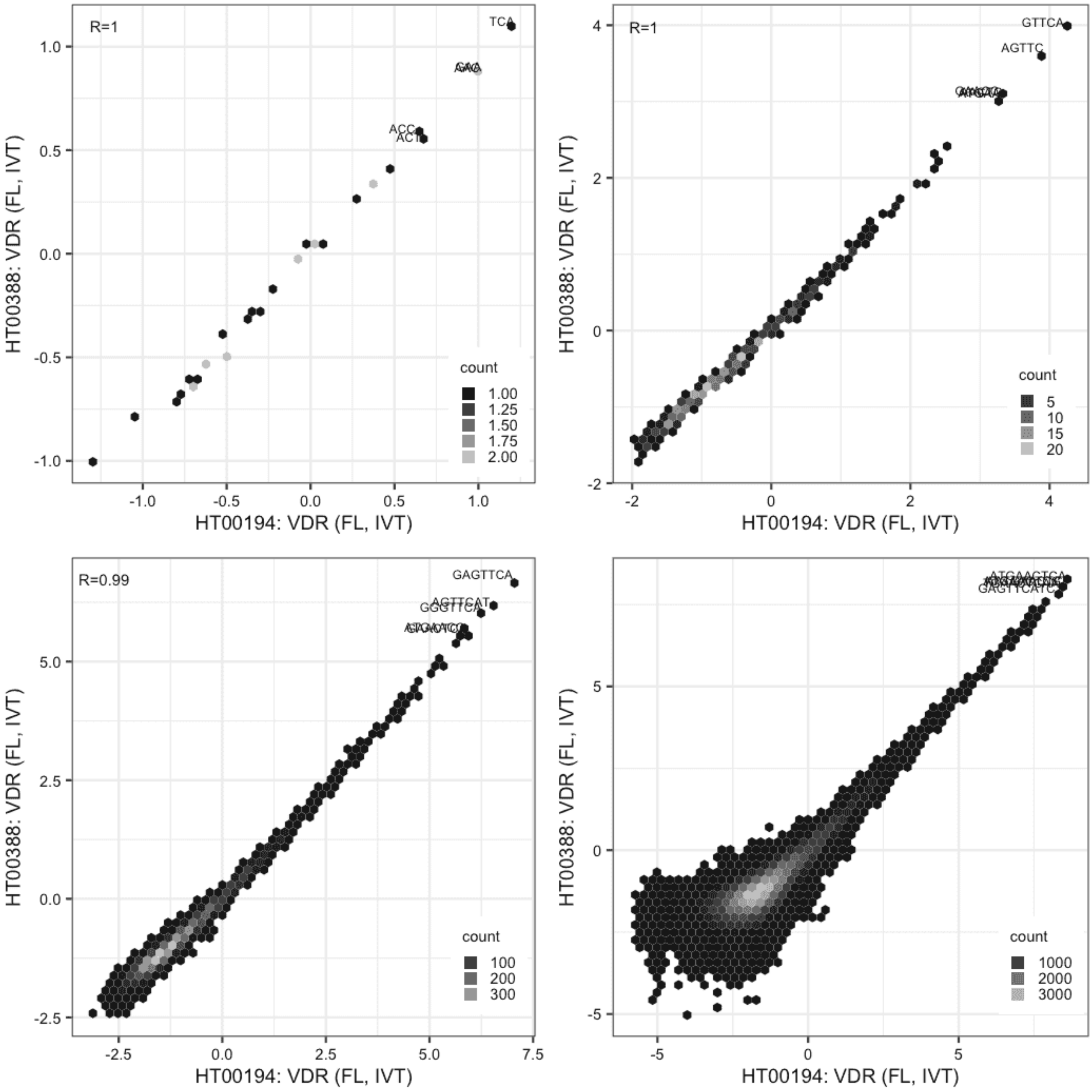

**VDR**

Lysate vs IVT, same construct

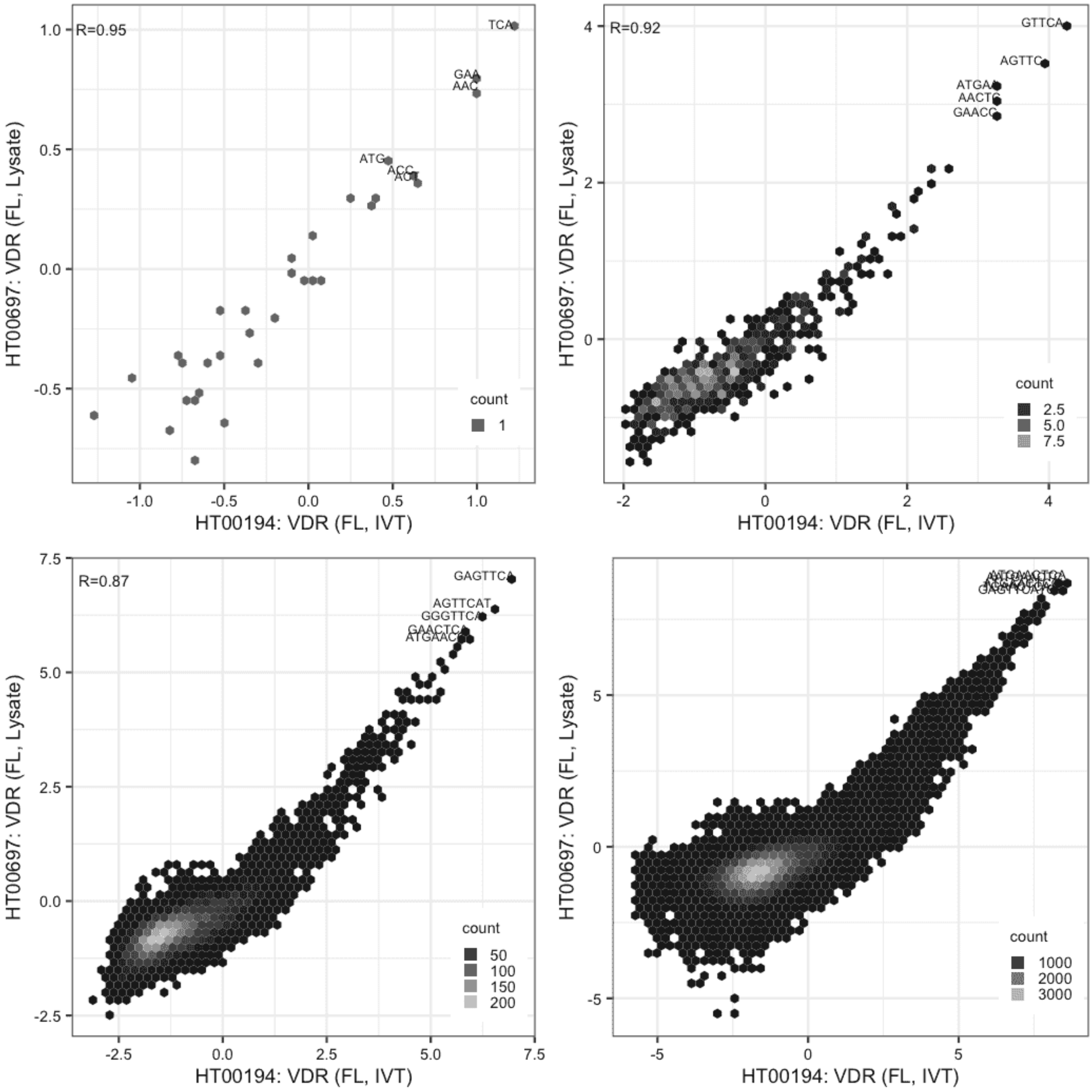

**VDR**

Lysate vs IVT, same construct

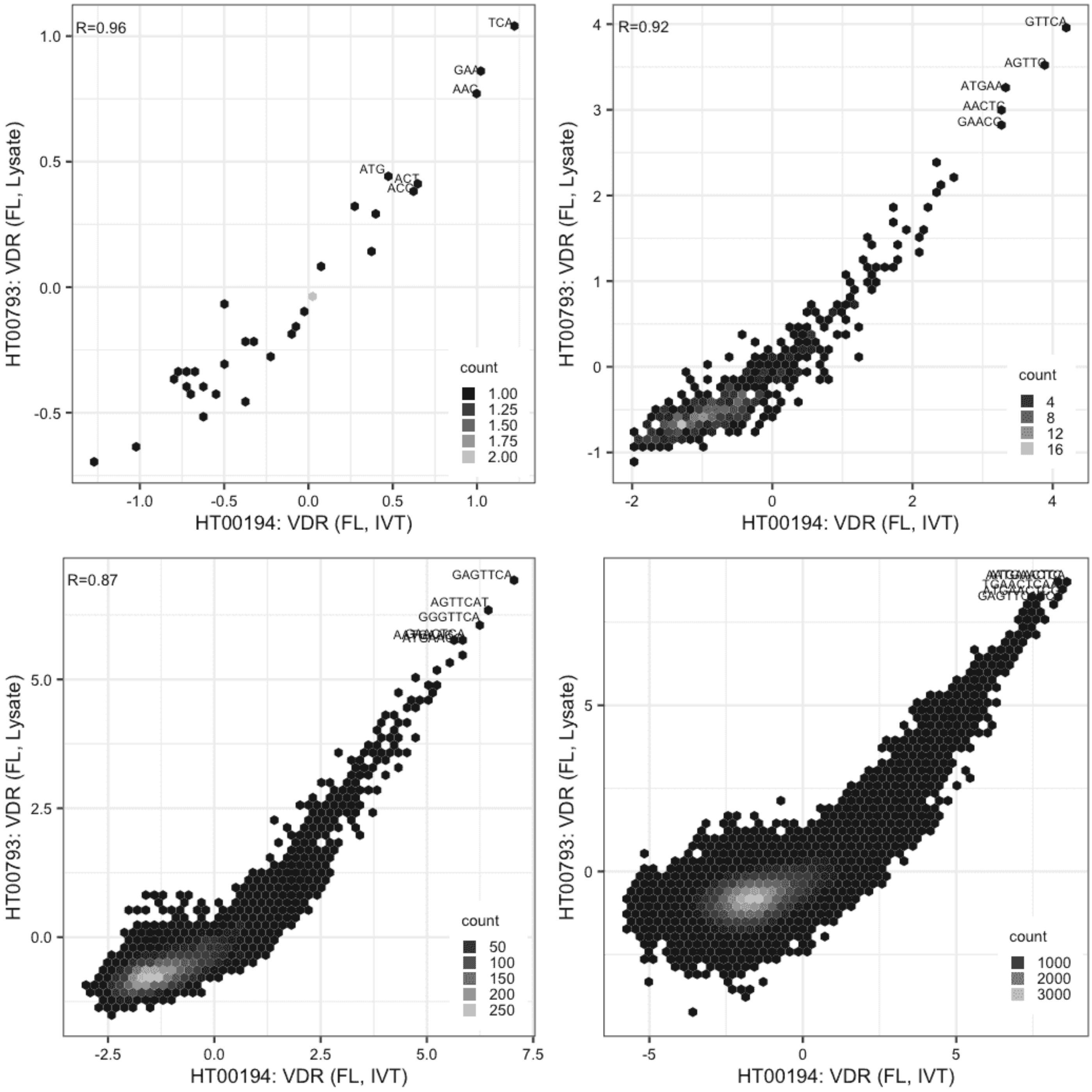

### VDR

Lysate vs IVT, same construct

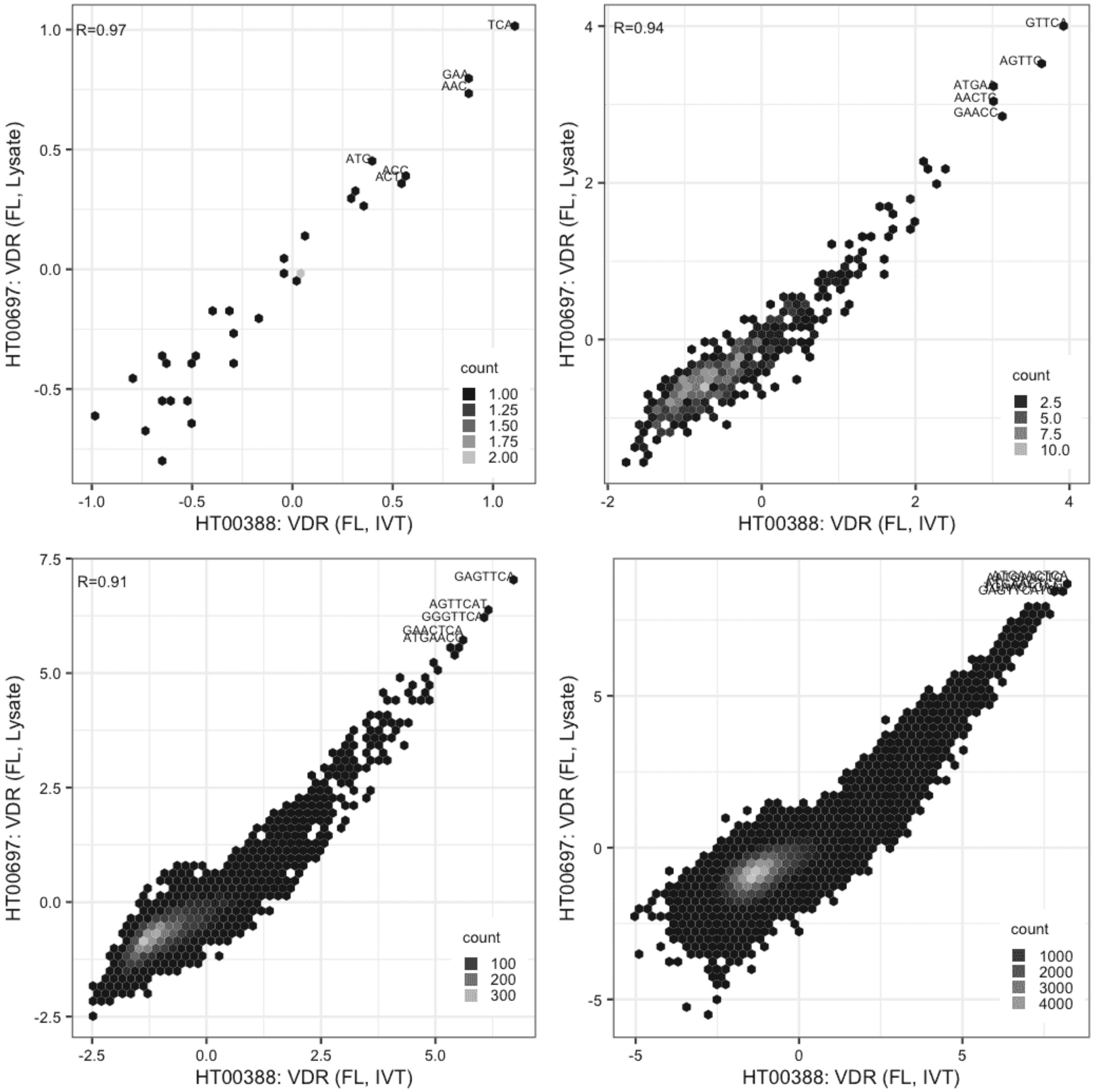

**VDR**

Lysate vs IVT, same construct

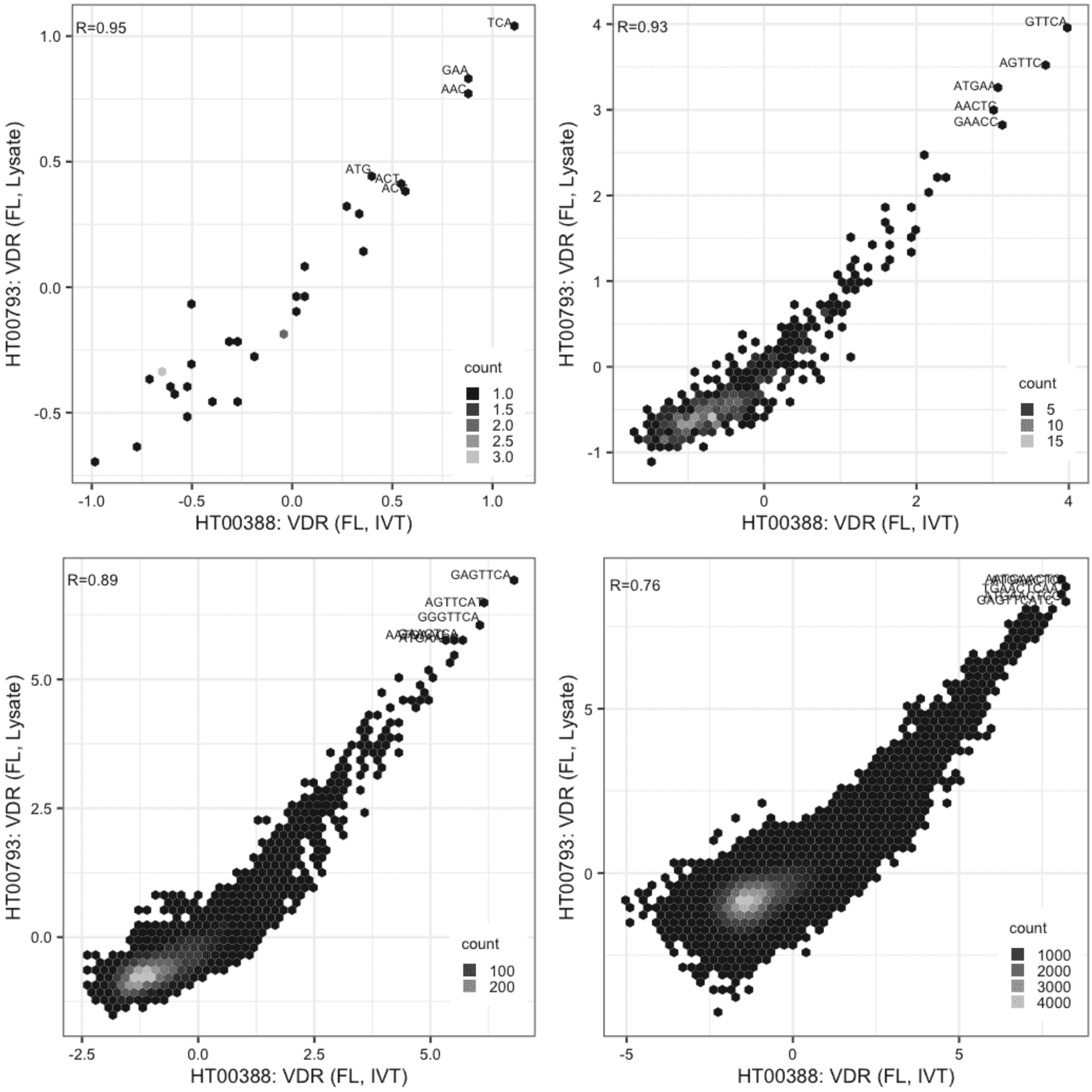

#### VDR

Lysate vs Lysate, same construct

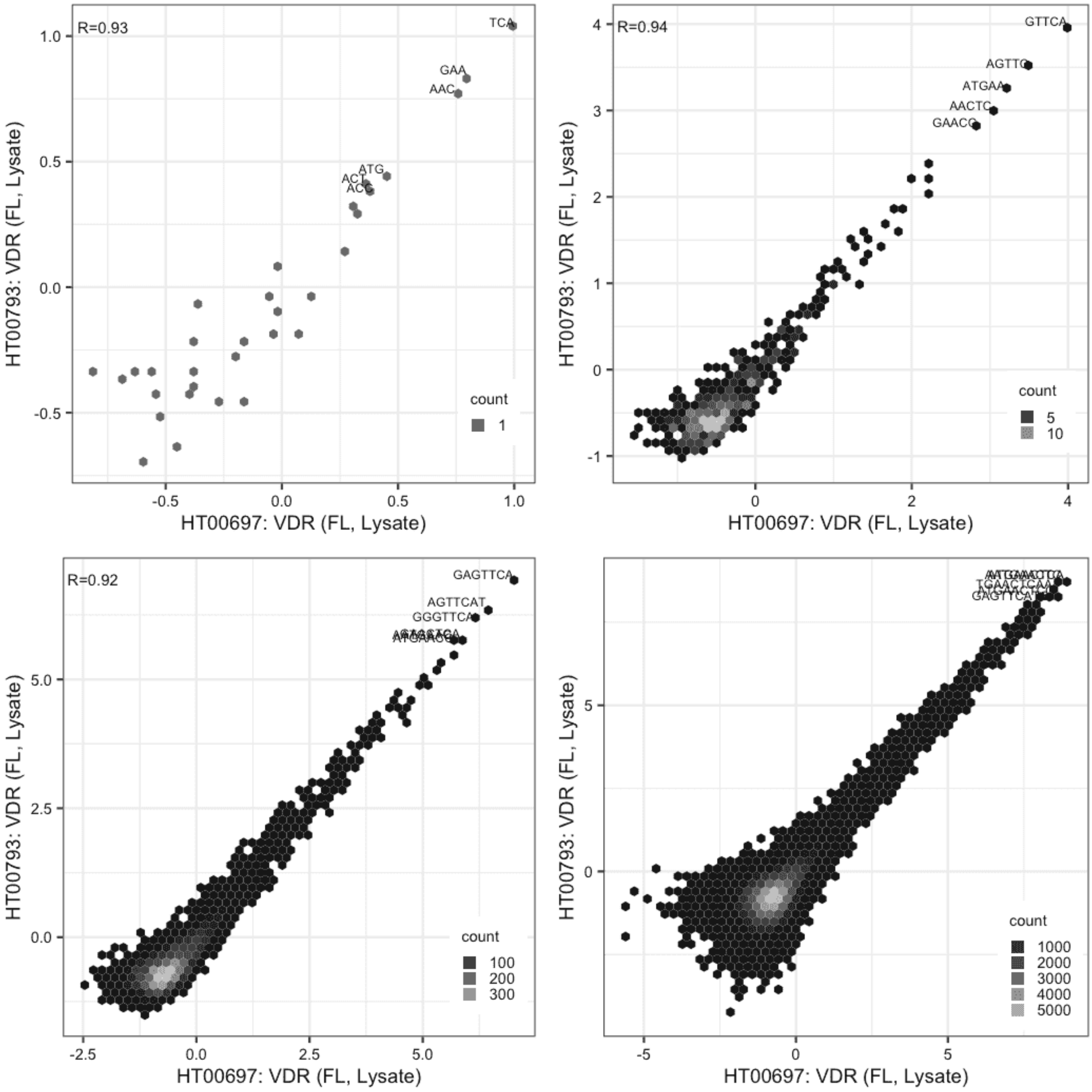

#### ZBED2

Lysate vs GFPIVT, same construct

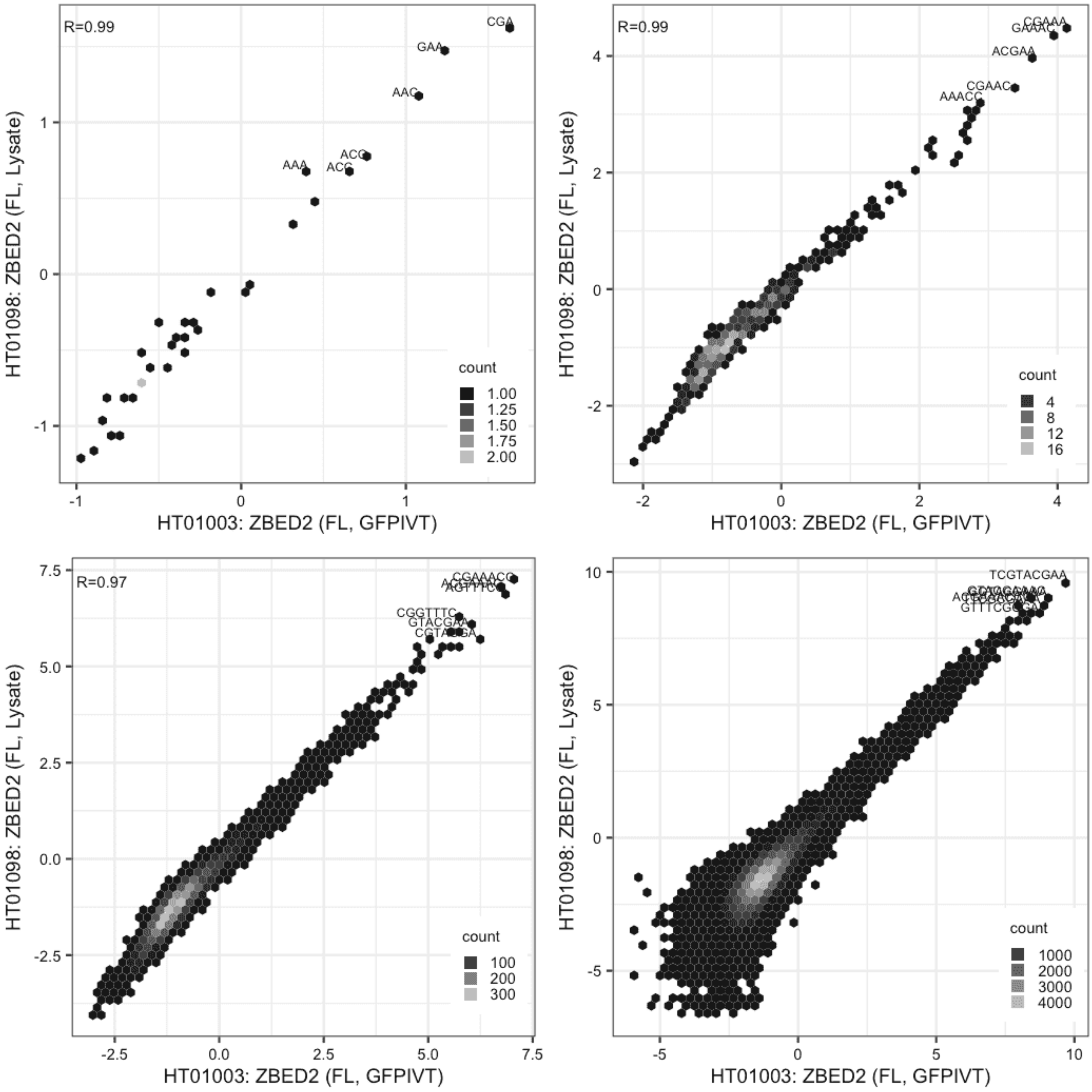

**ZBED2**

Lysate vs GFPIVT, same construct

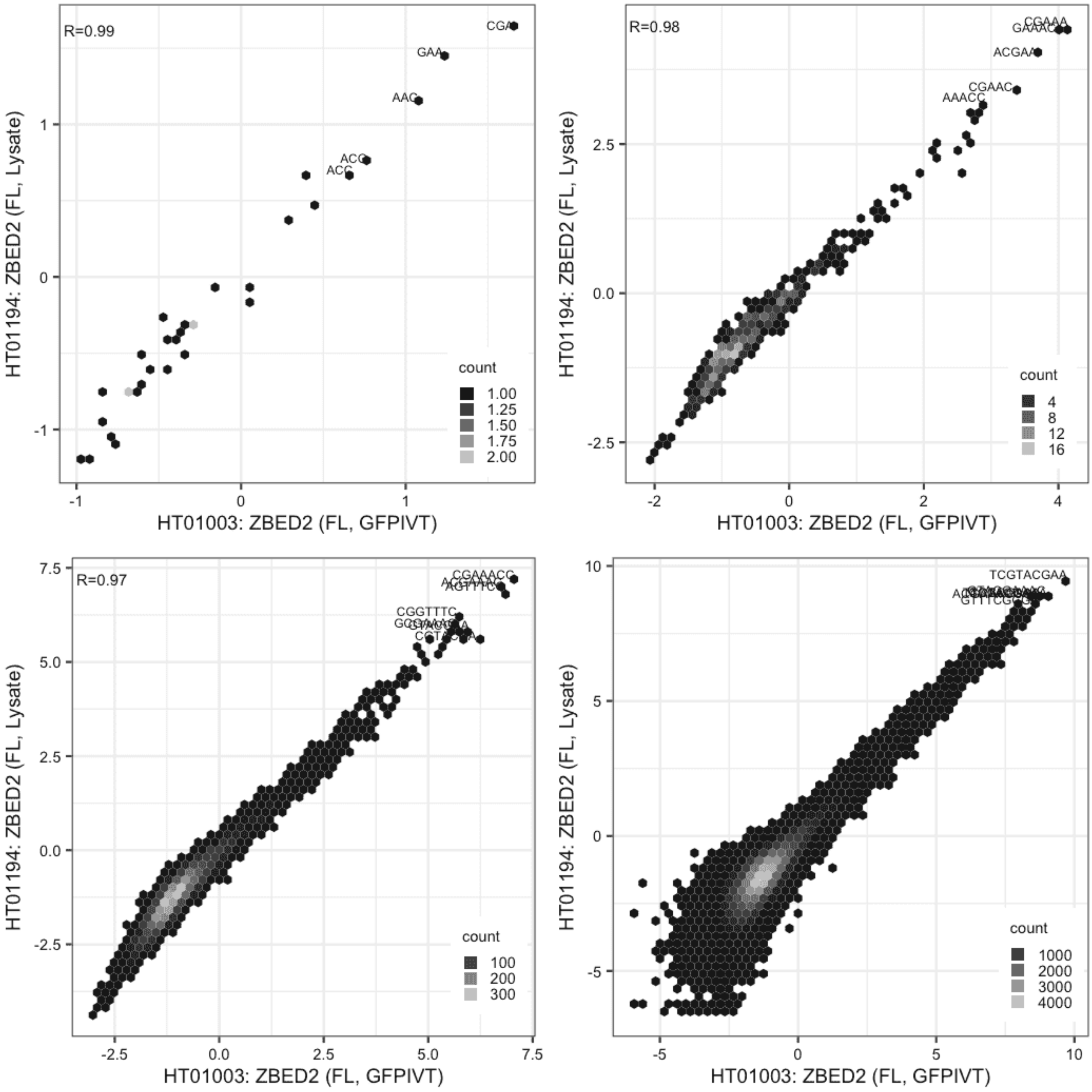

#### ZBED2

Lysate vs Lysate, same construct

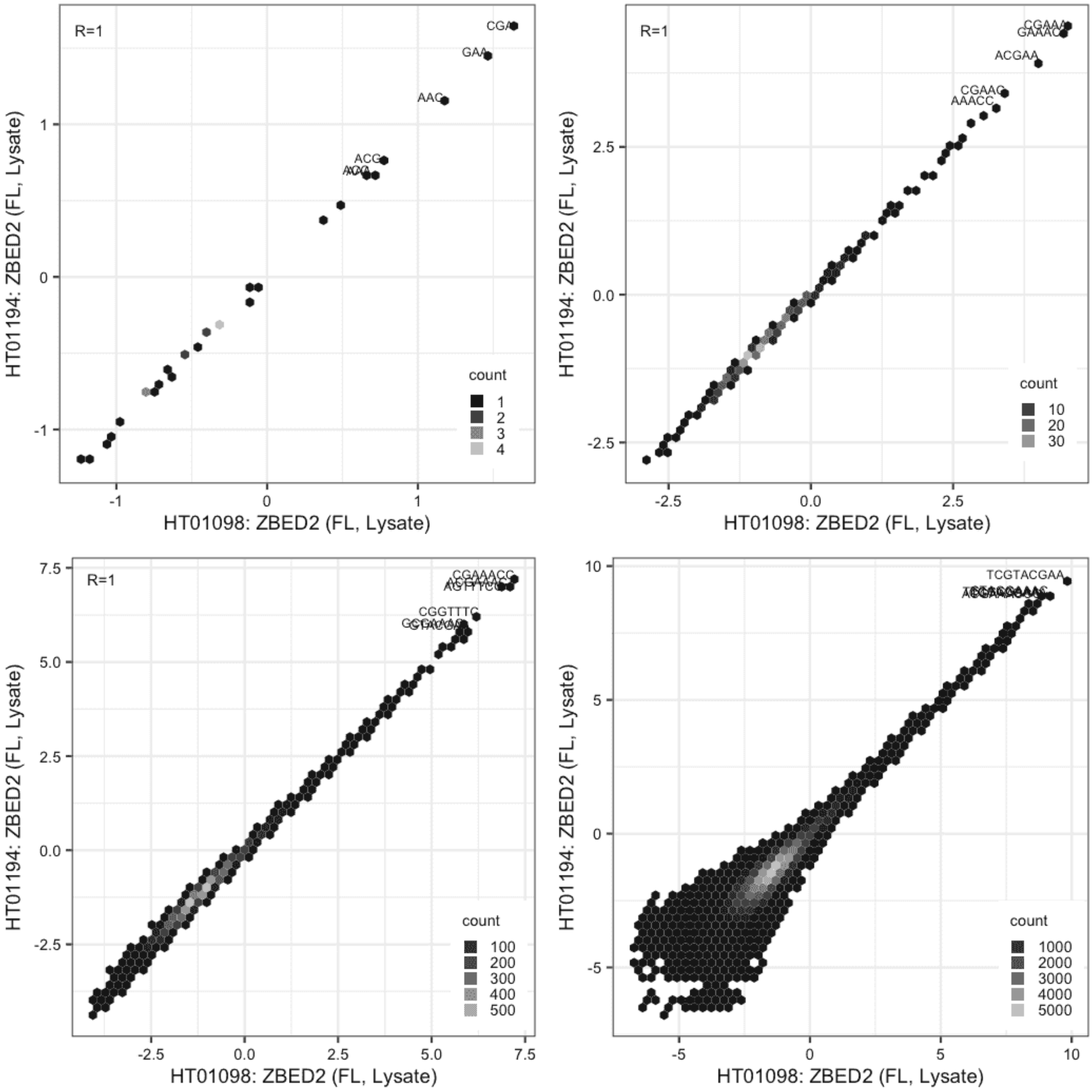

### ZBED5

IVT vs GFPIVT, same construct

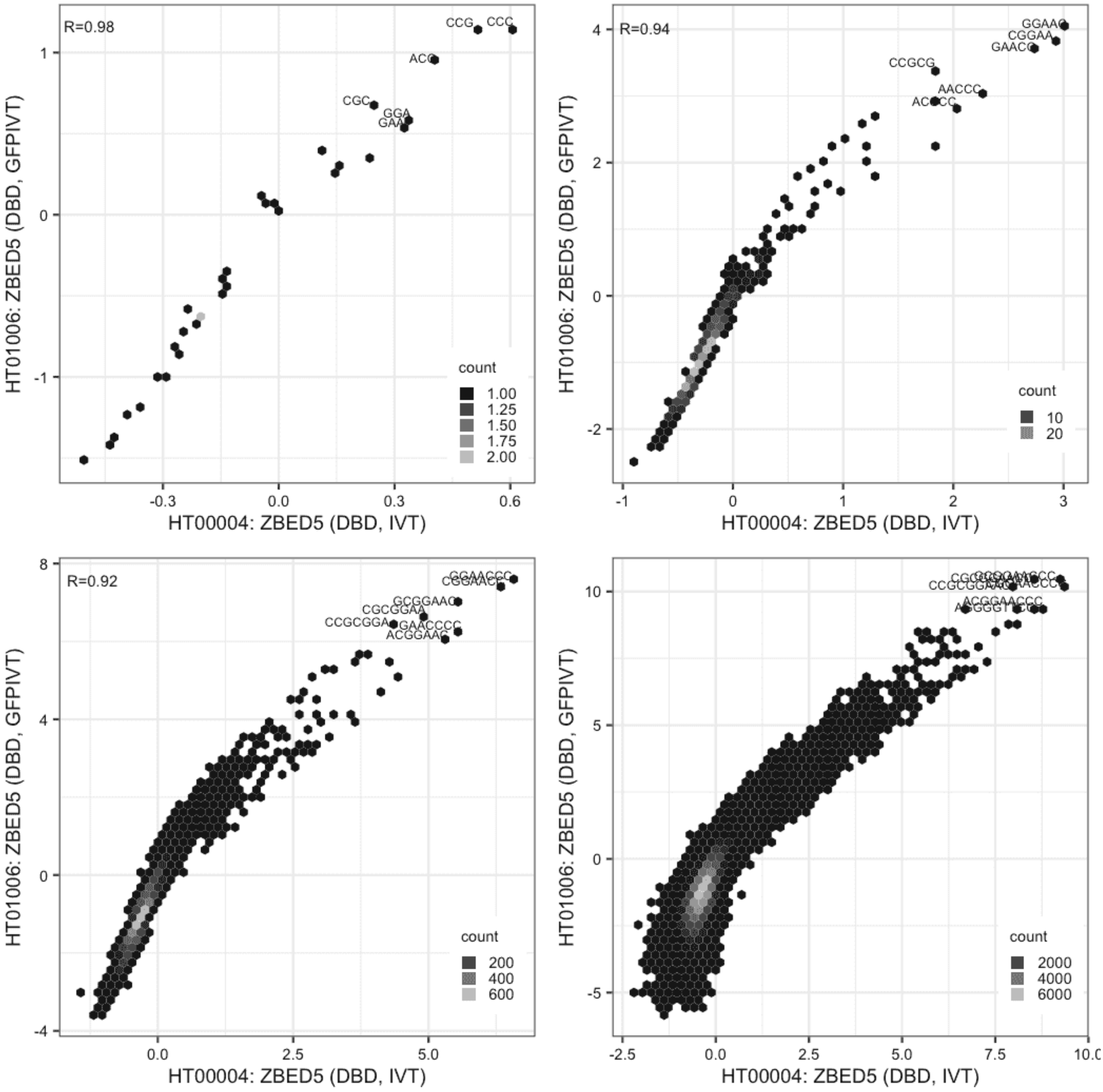

**ZBED5**

Lysate vs IVT, different constructs

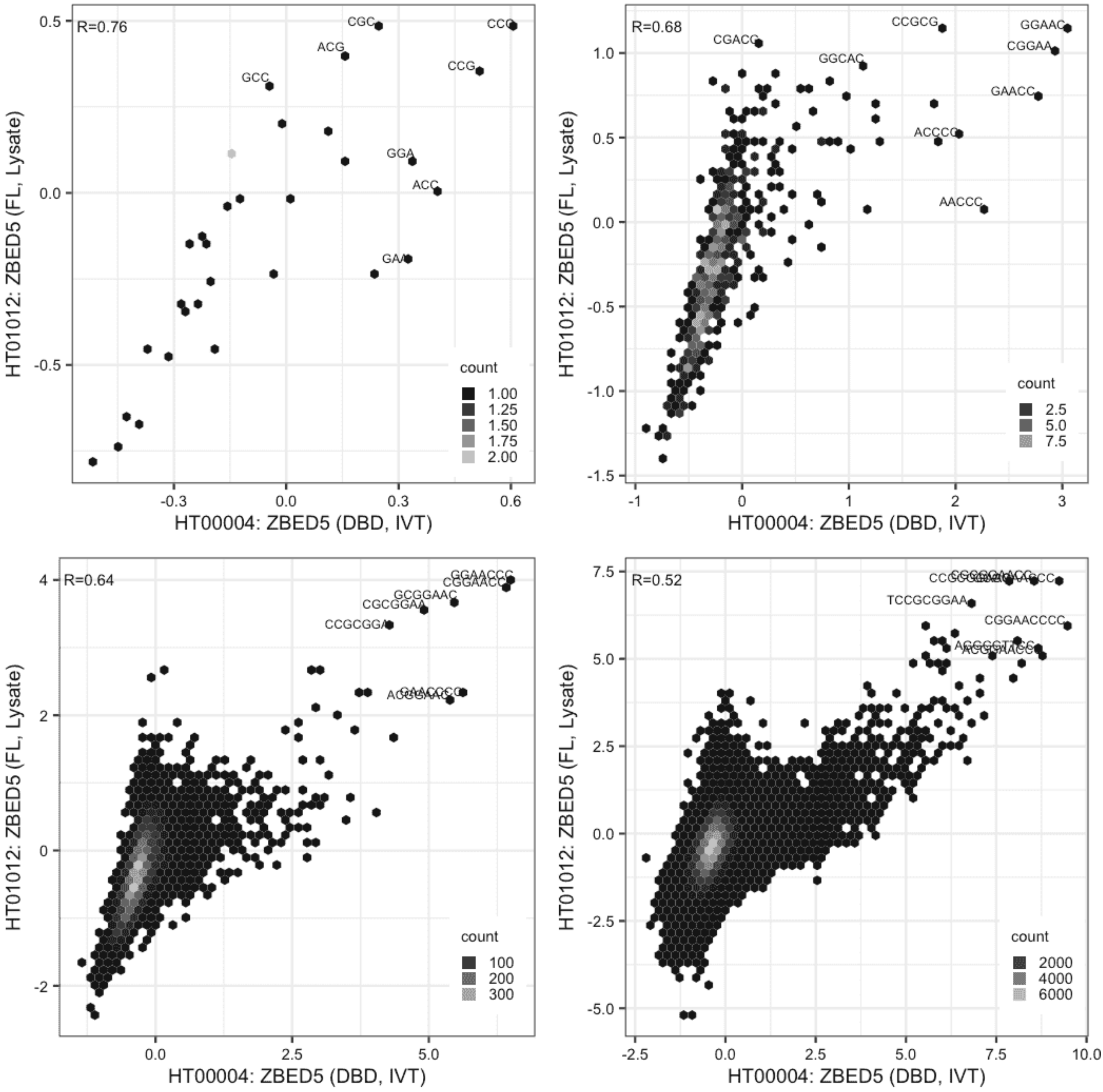

#### ZBED5

Lysate vs IVT, different constructs

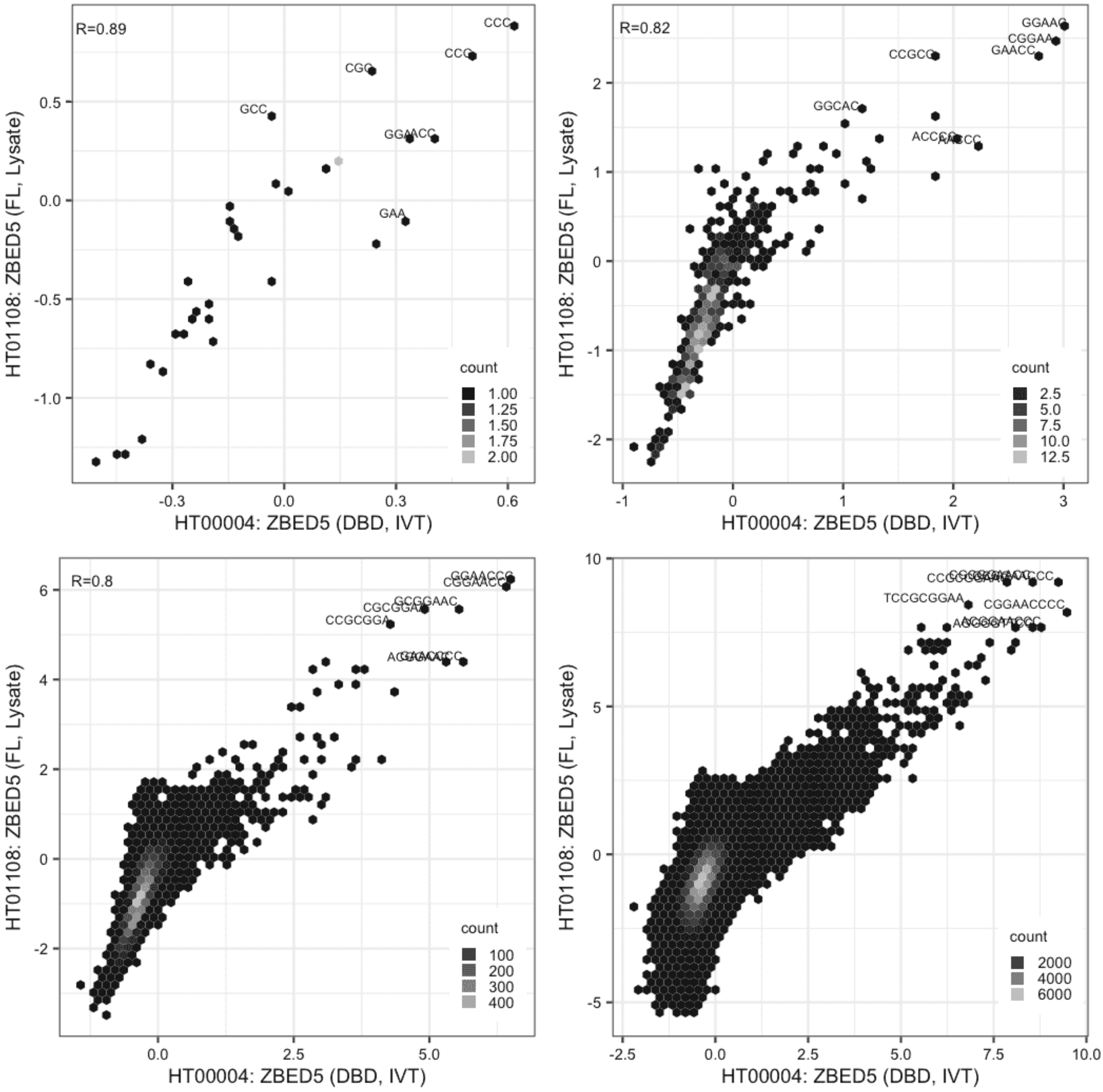

#### ZBED5

IVT vs GFPIVT, different constructs

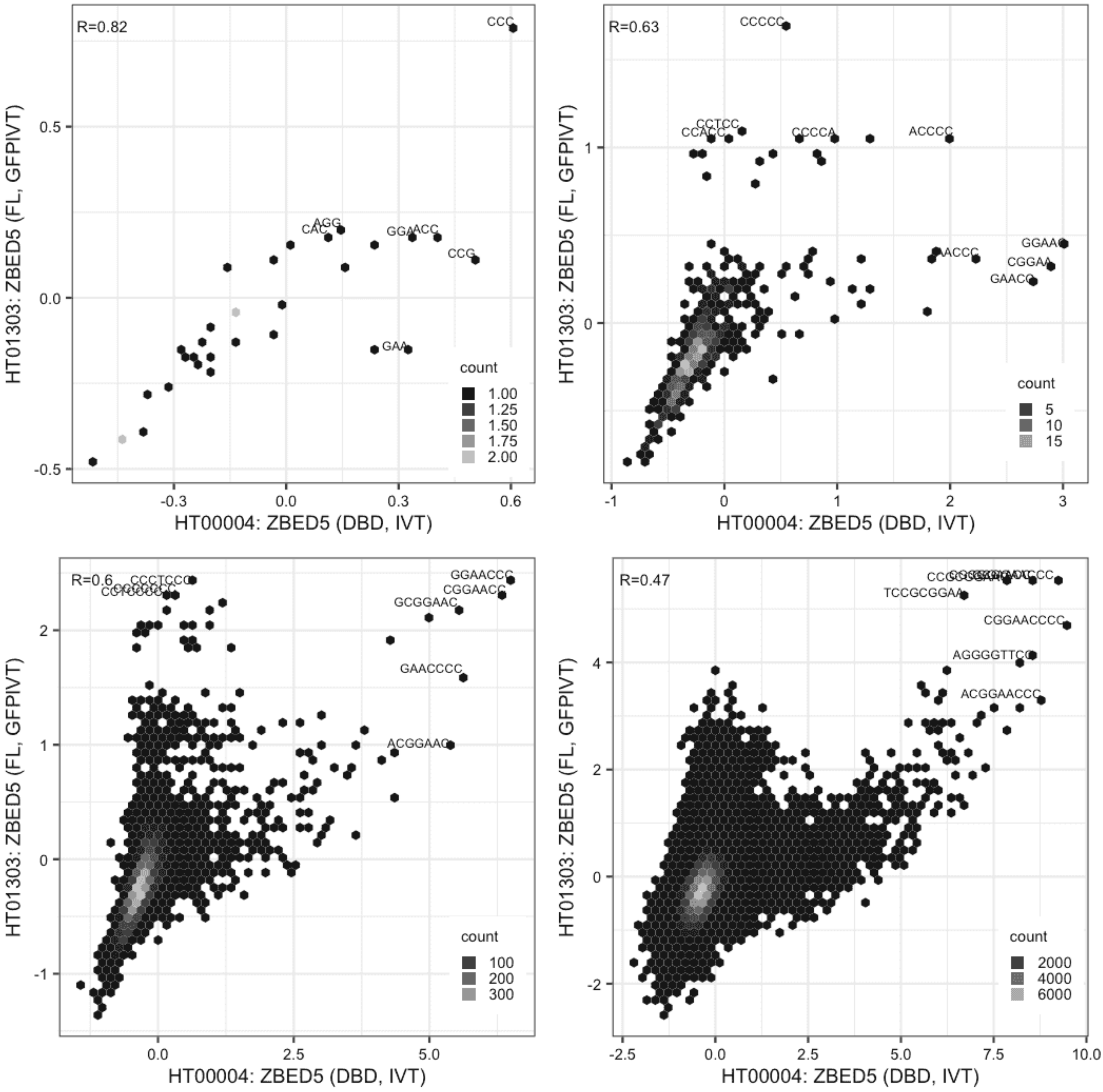

#### ZBED5

Lysate vs GFPIVT, different constructs

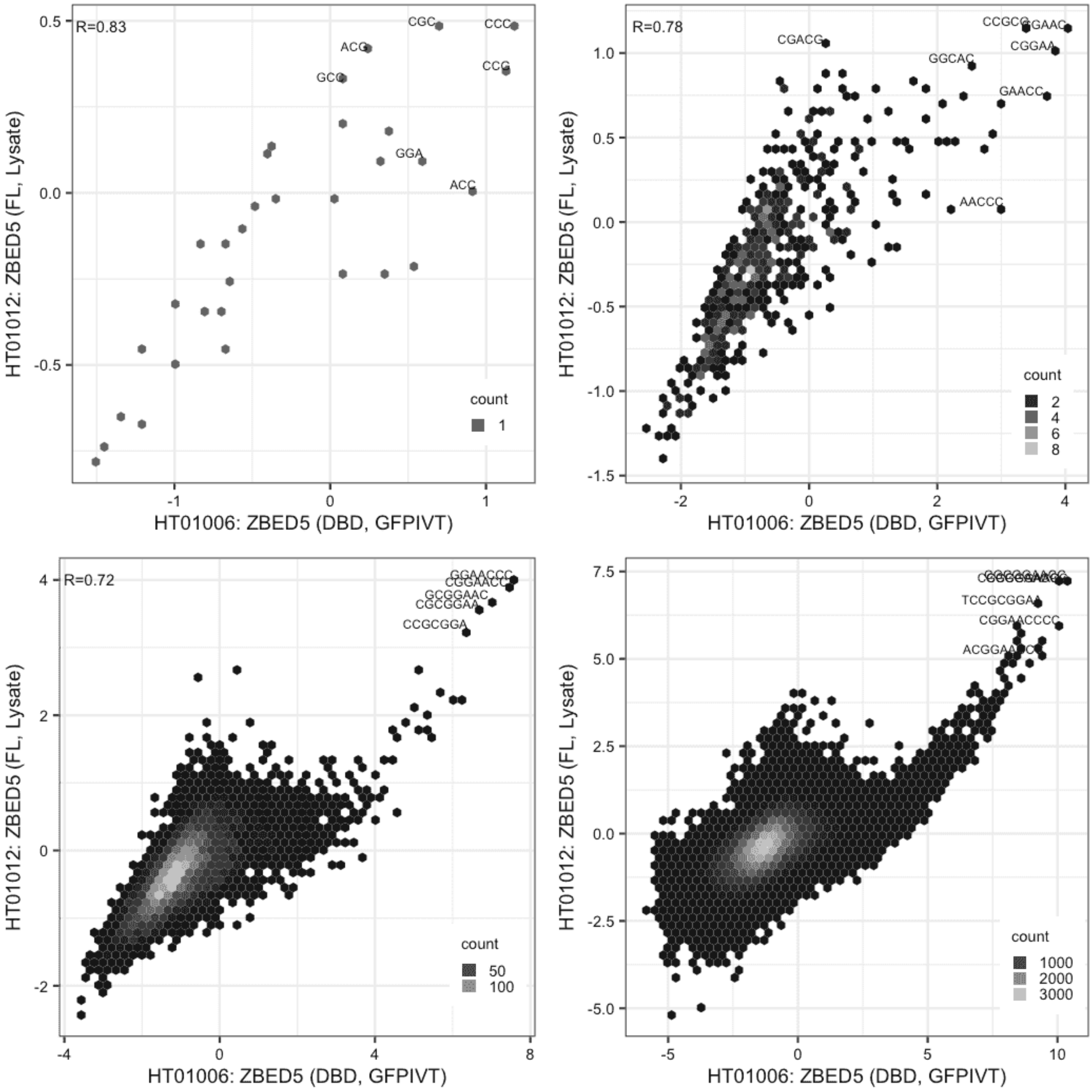

#### ZBED5

Lysate vs GFPIVT, different constructs

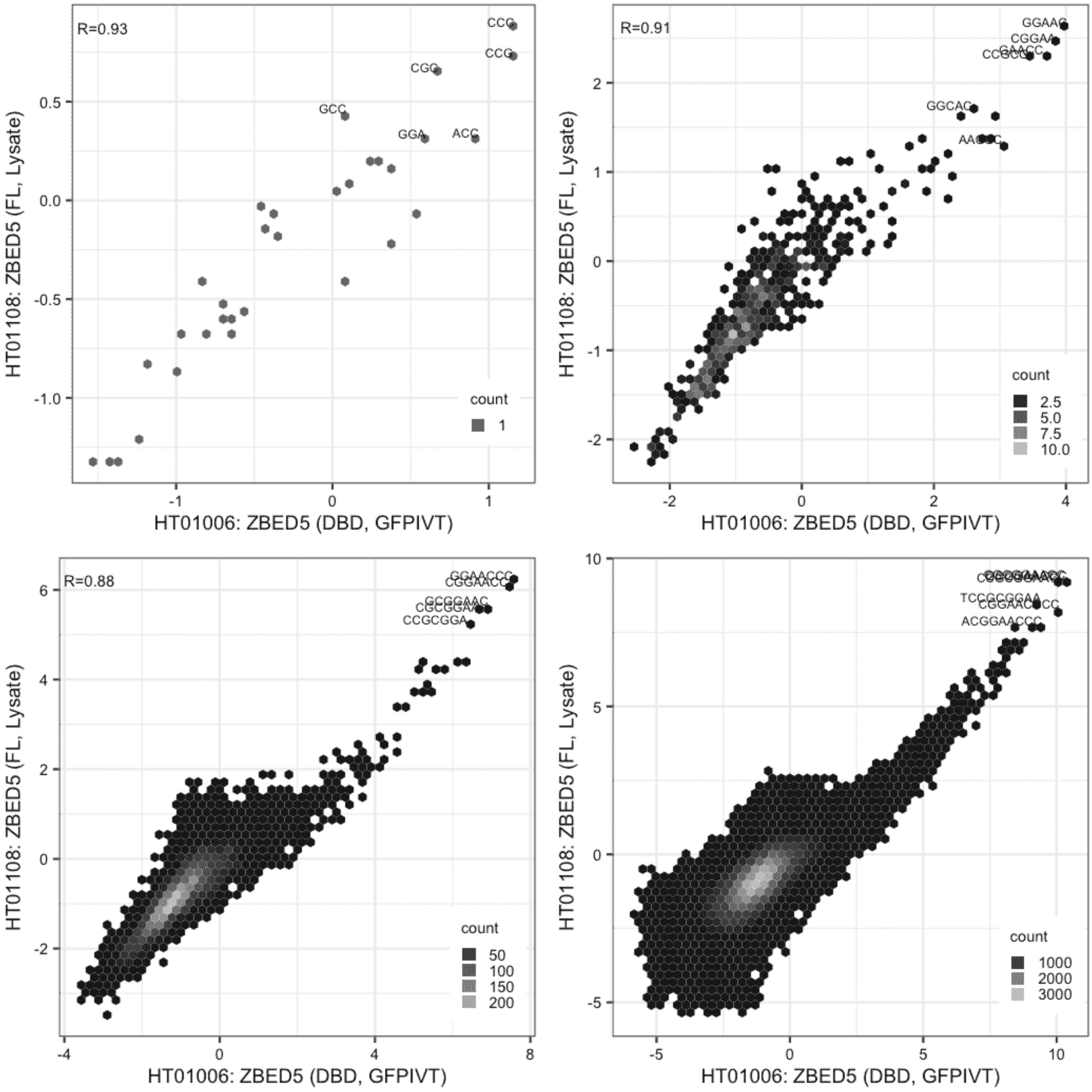

**ZBED5**

GFPIVT vs GFPIVT, different constructs

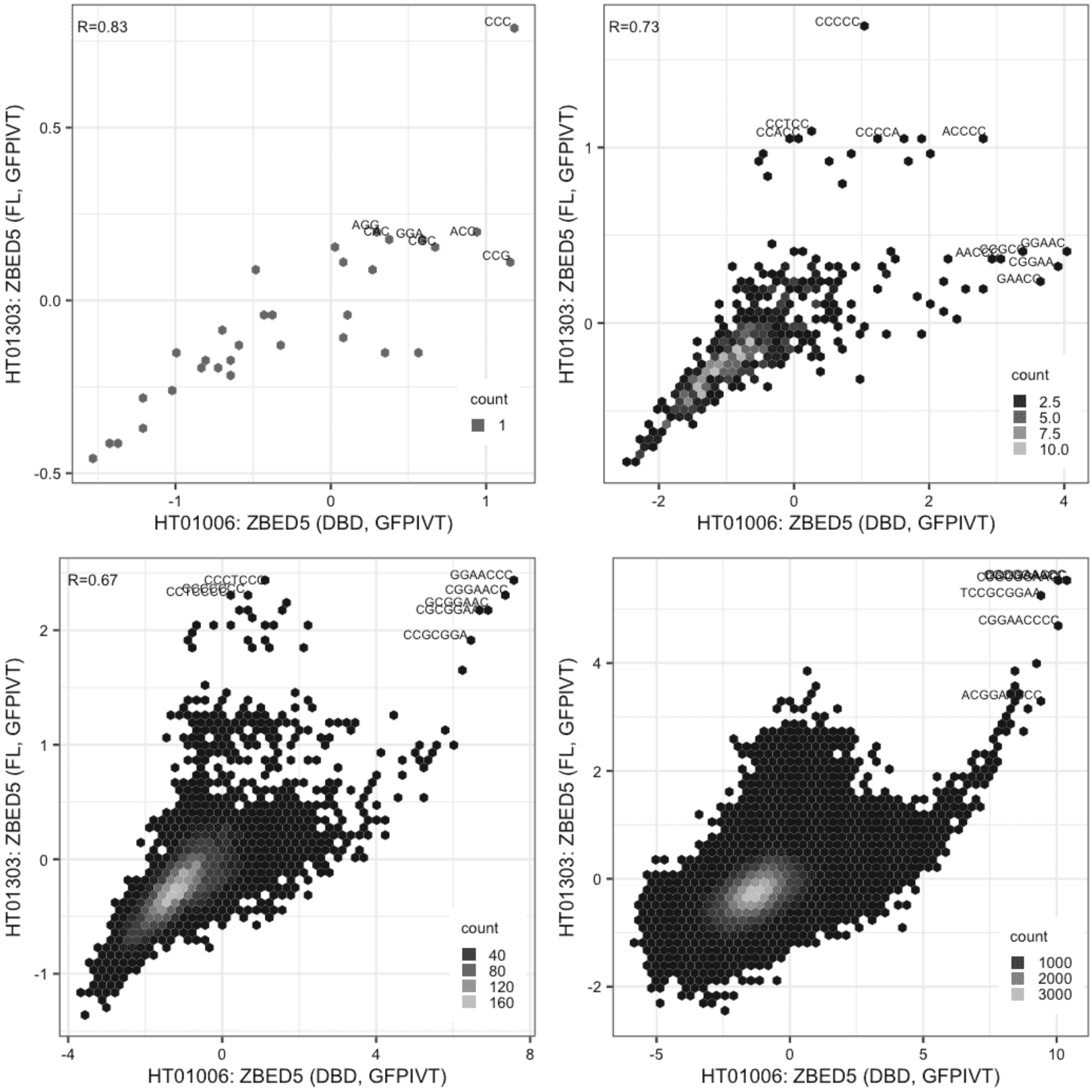

**ZBED5**

Lysate vs Lysate, same construct

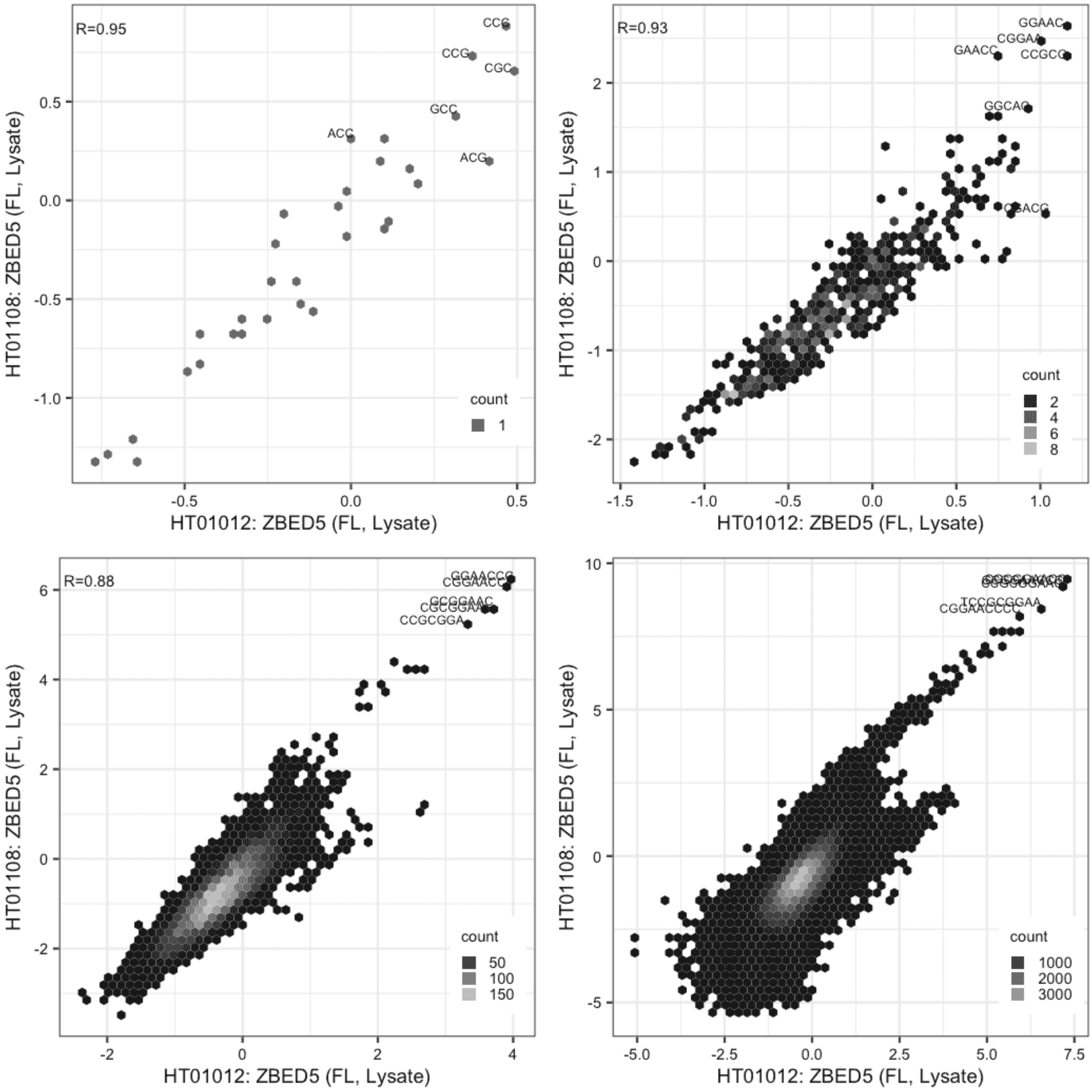

#### ZBED5

Lysate vs GFPIVT, same construct

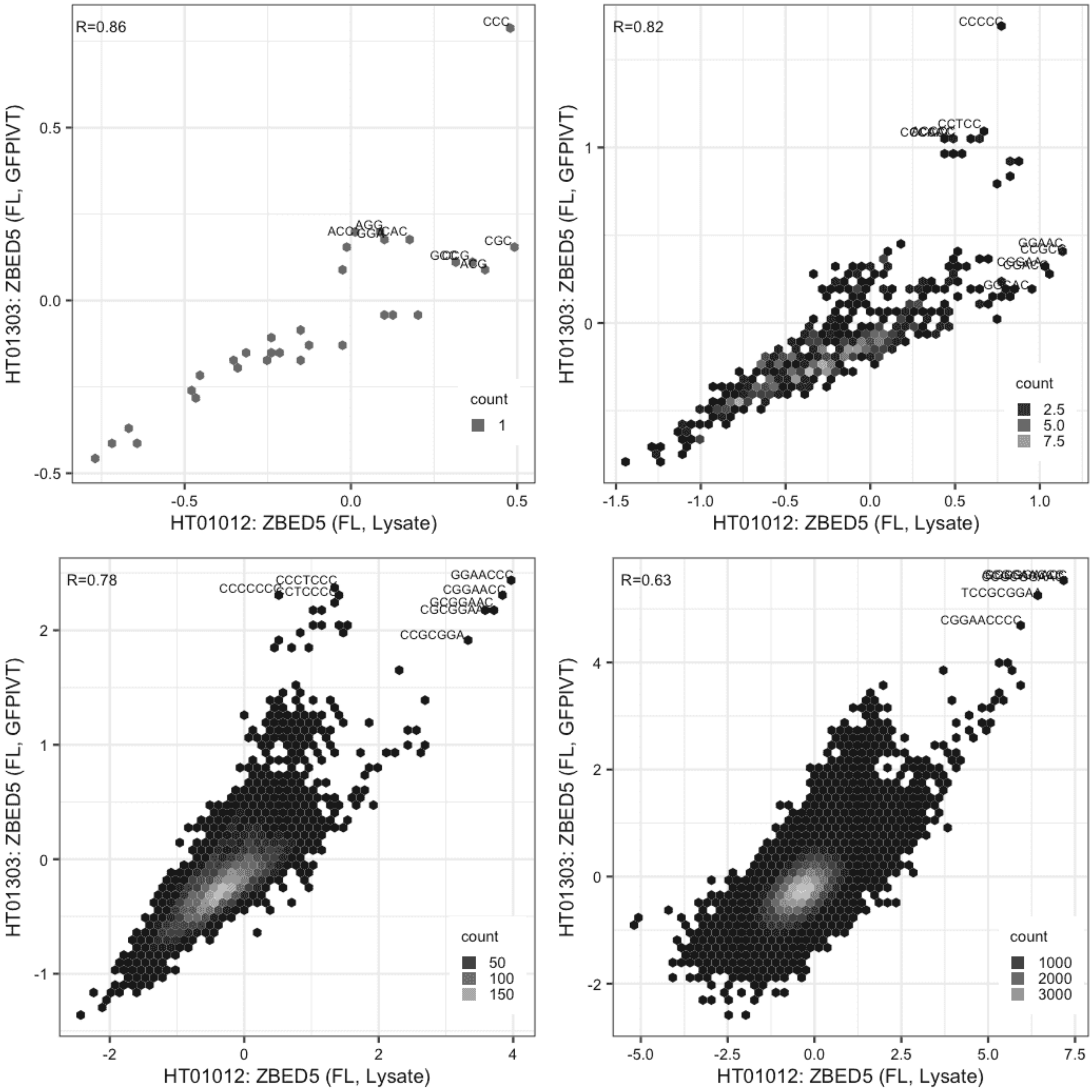

**ZBED5**

Lysate vs GFPIVT, same construct

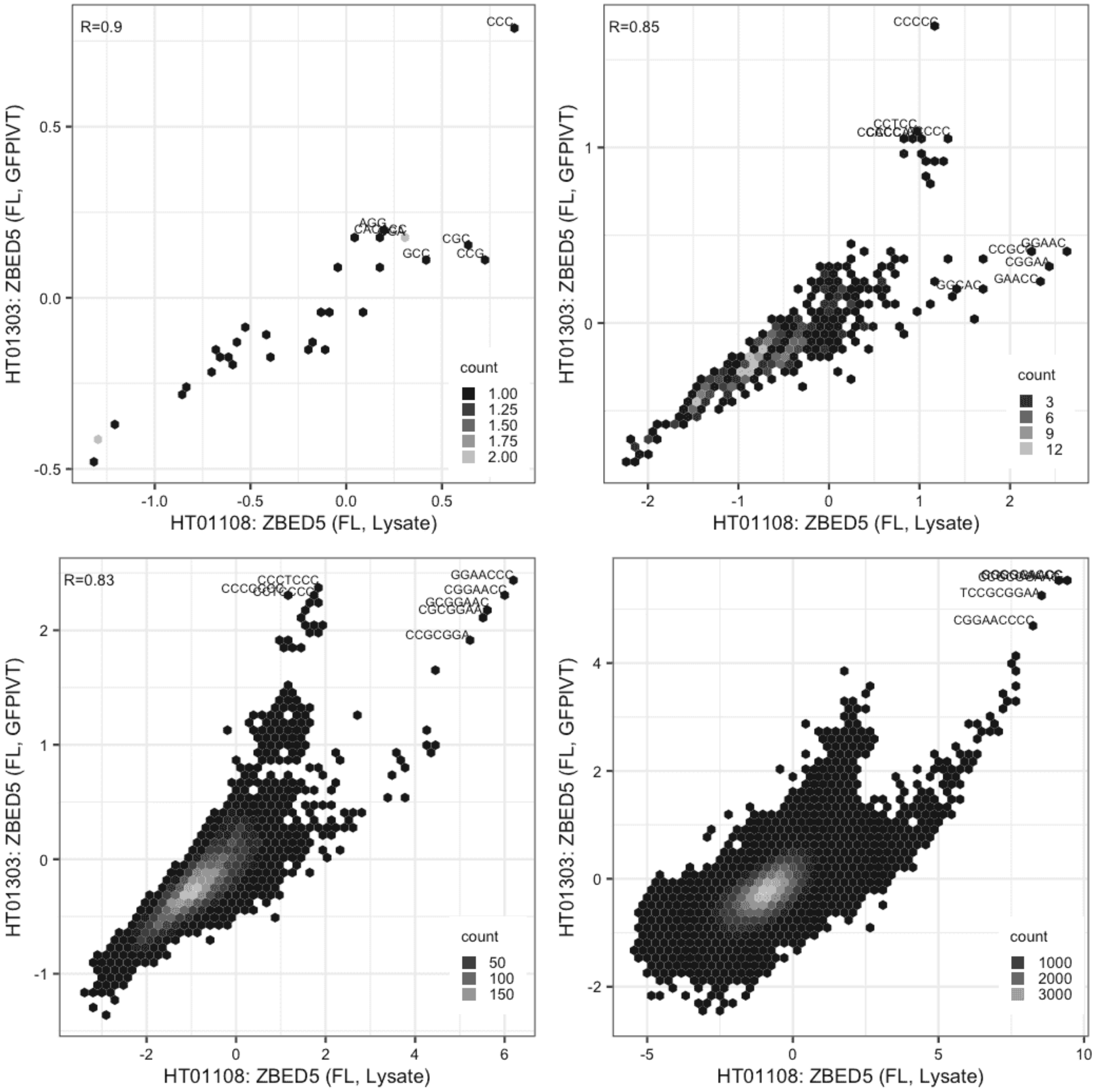

**ZBTB47**

Lysate vs Lysate, same construct

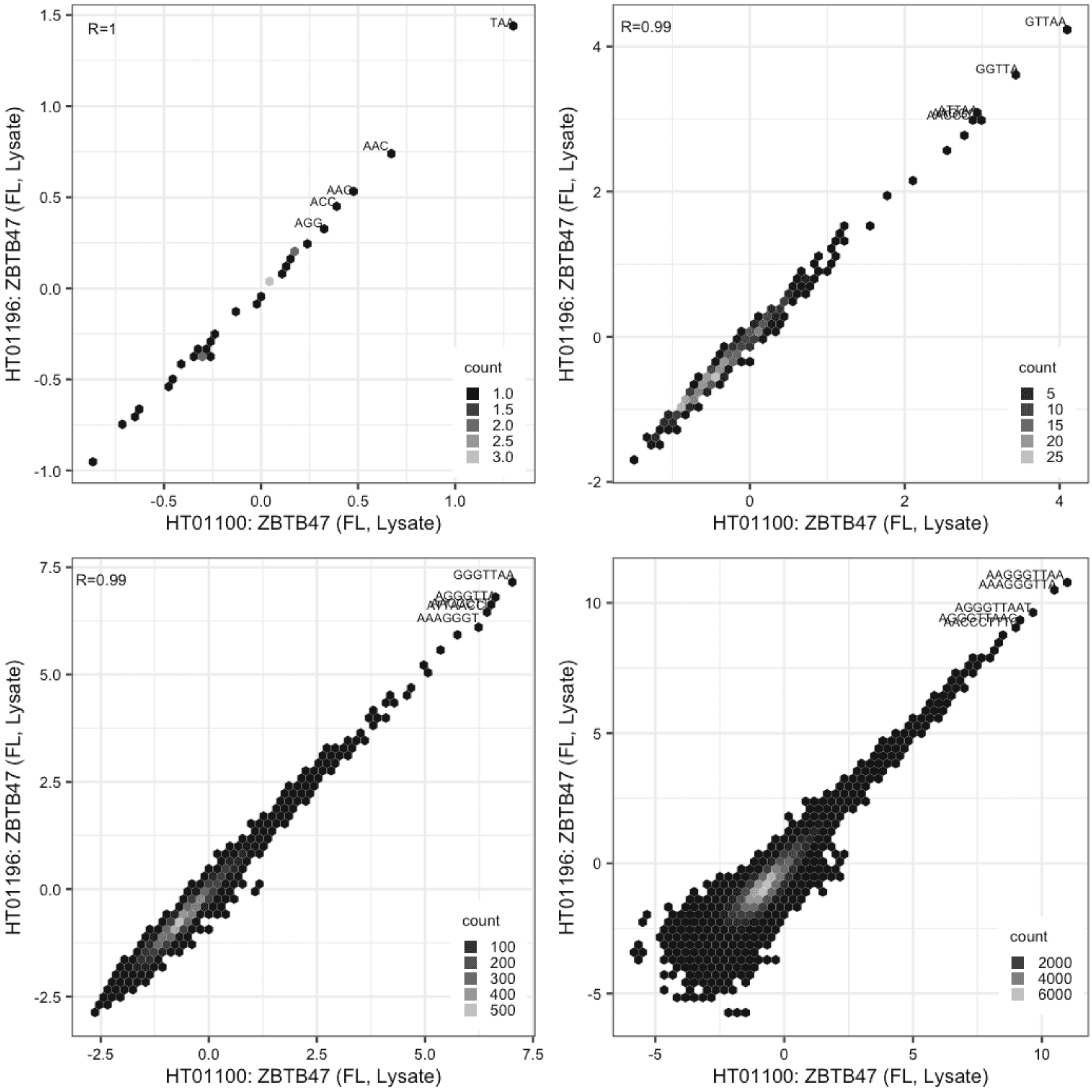

**ZBTB47**

Lysate vs GFPIVT, same construct

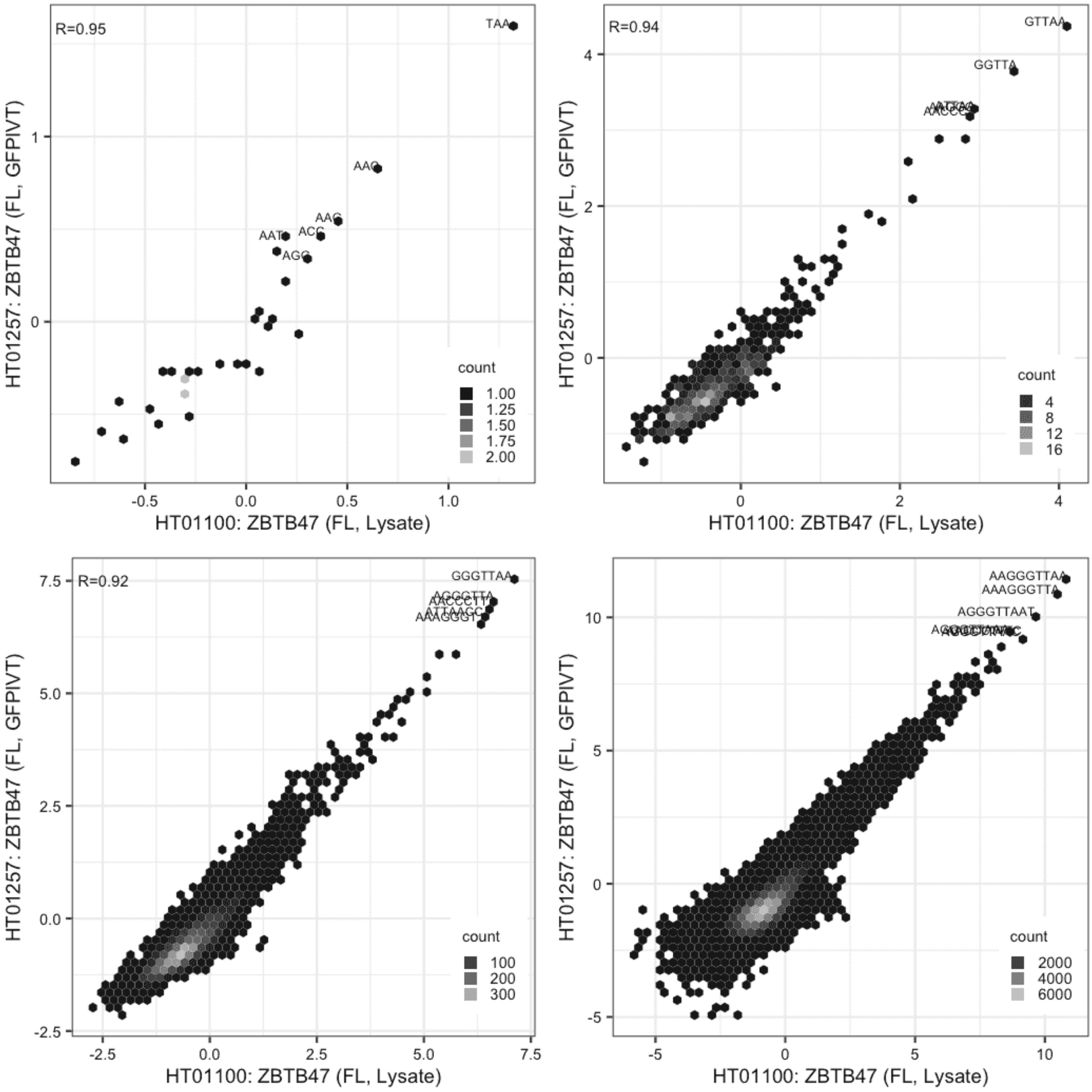

**ZBTB47**

Lysate vs GFPIVT, different constructs

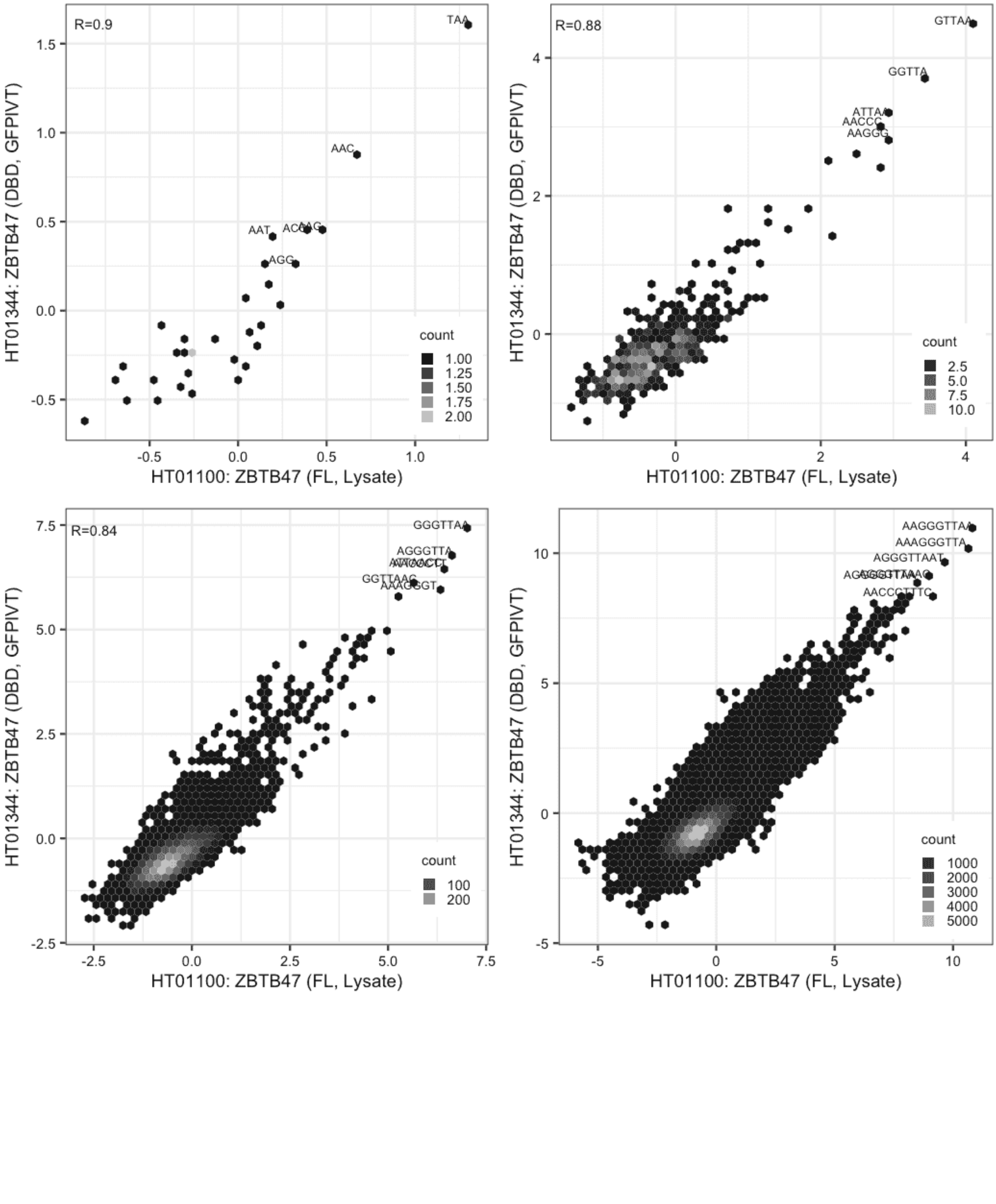

#### ZBTB47

Lysate vs GFPIVT, same construct

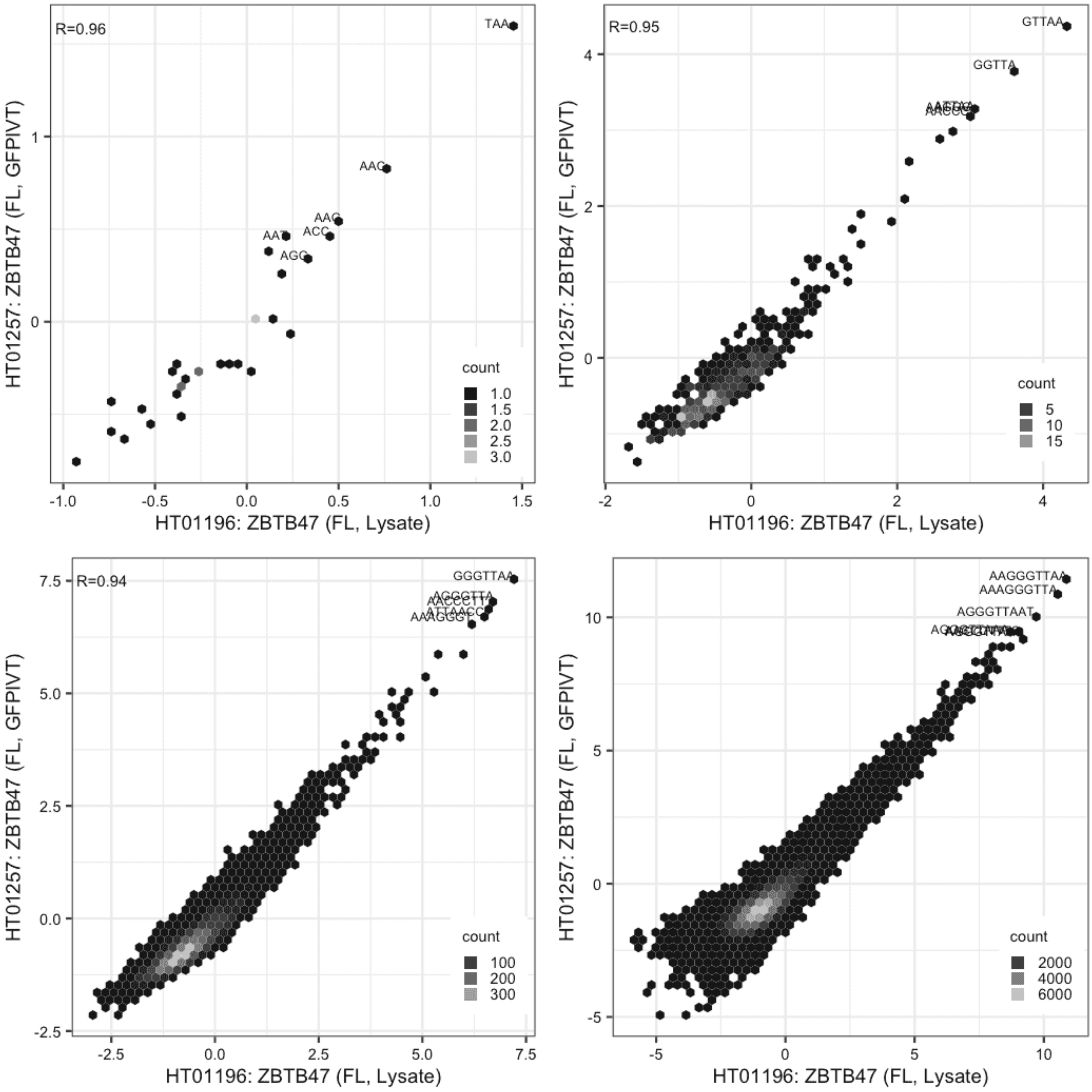

**ZBTB47**

Lysate vs GFPIVT, different constructs

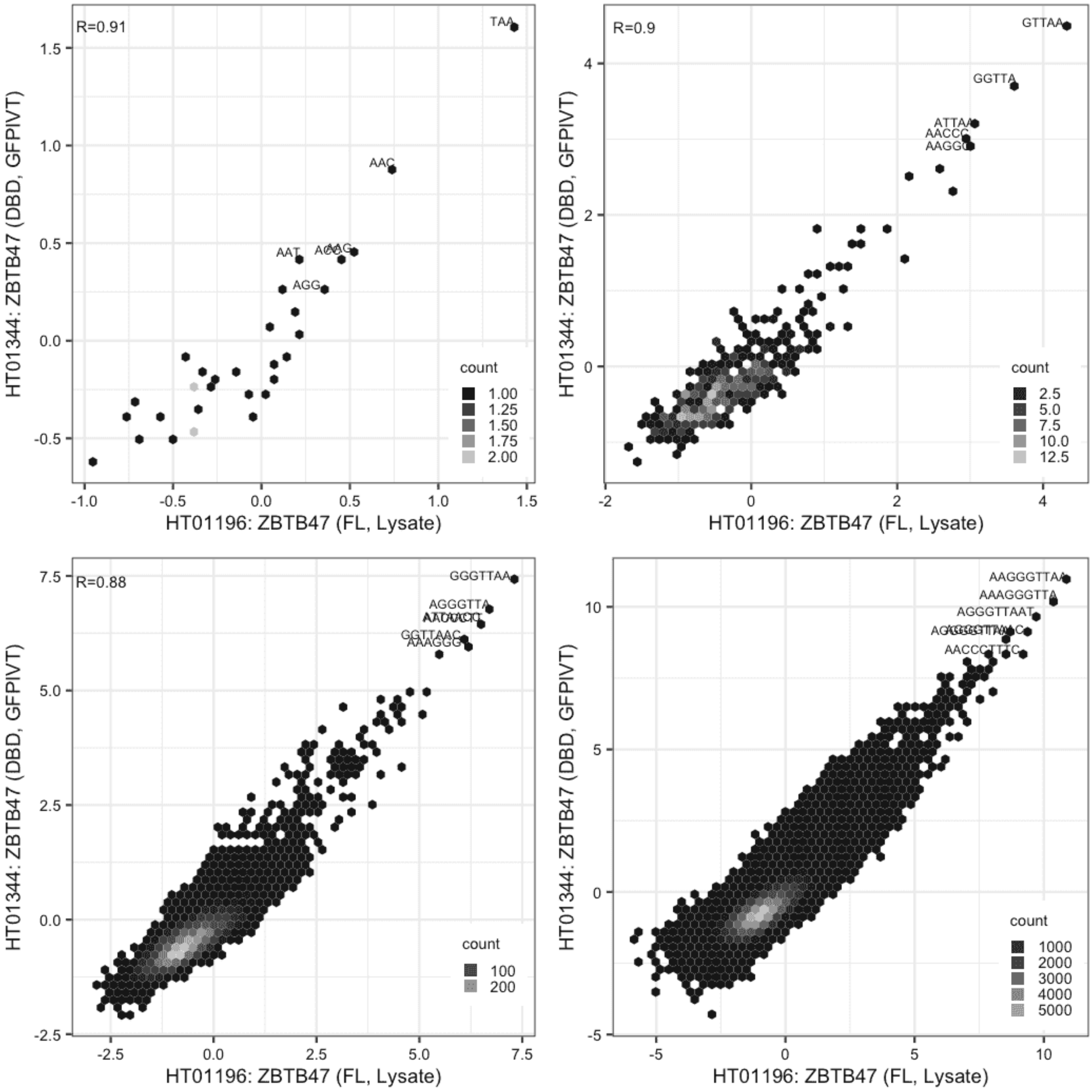

**ZBTB47**

GFPIVT vs GFPIVT, different constructs

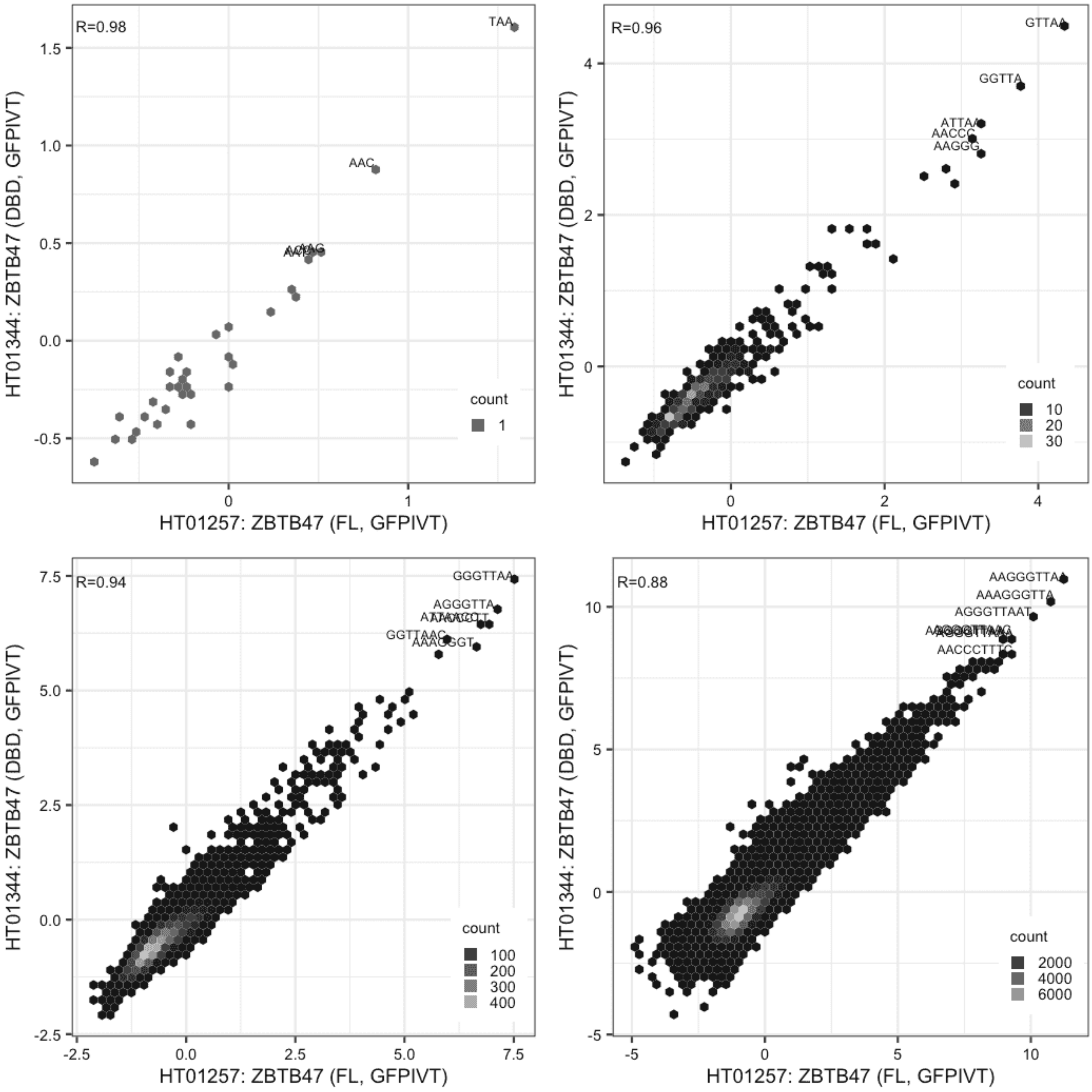

**ZBTB8A**

Lysate vs IVT, different constructs

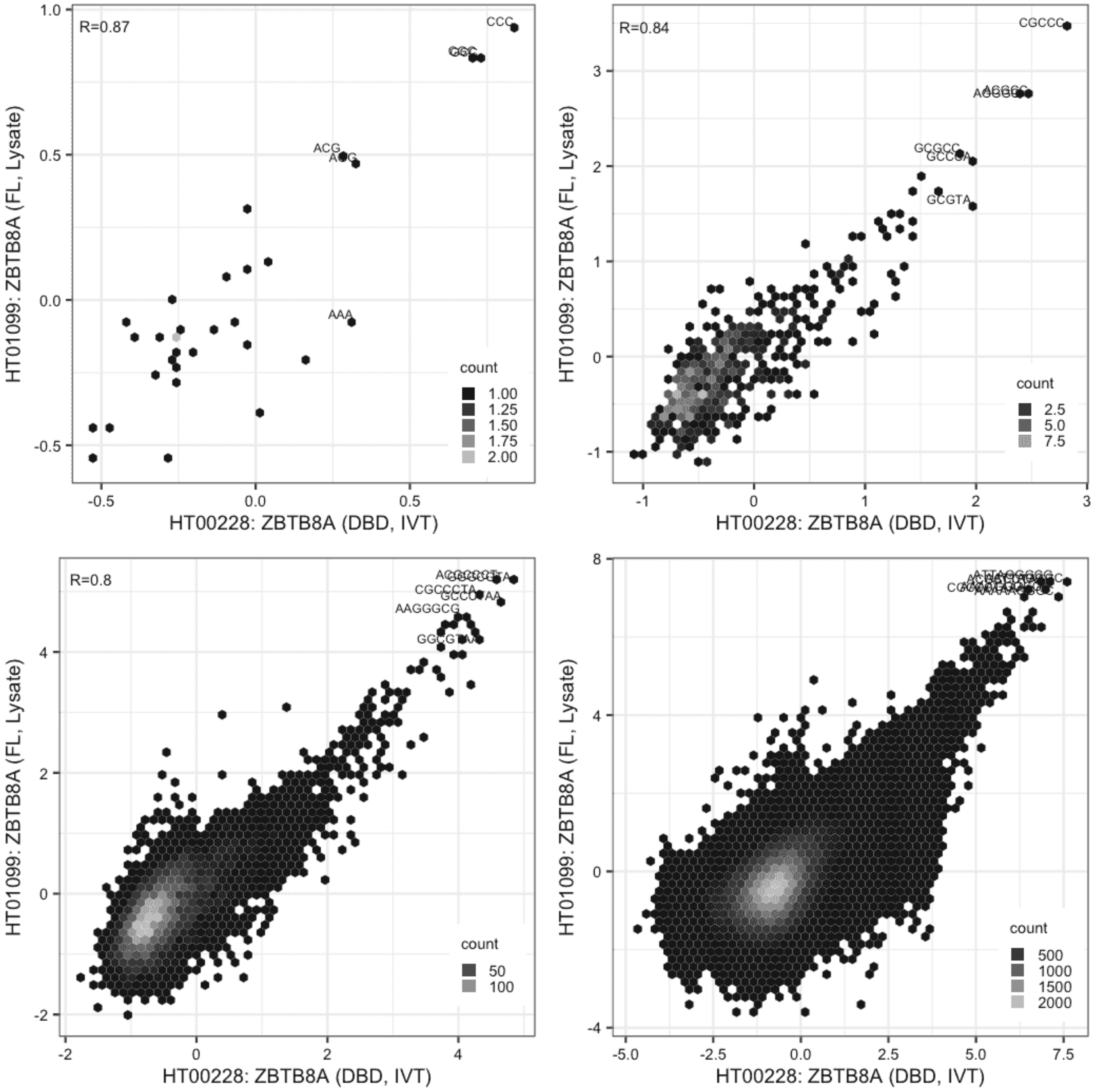

**ZBTB8A**

Lysate vs IVT, different constructs

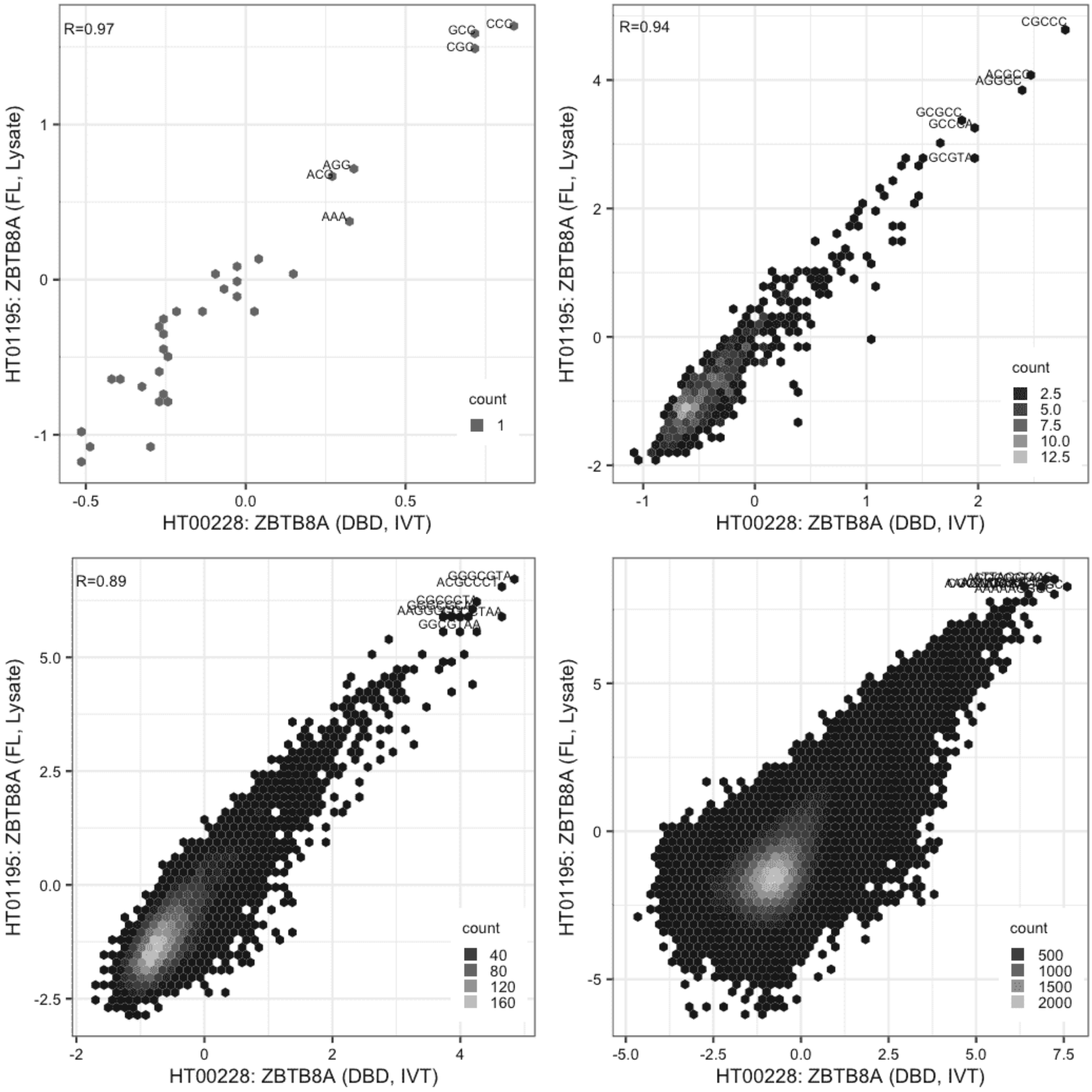

#### ZBTB8A

Lysate vs Lysate, same construct

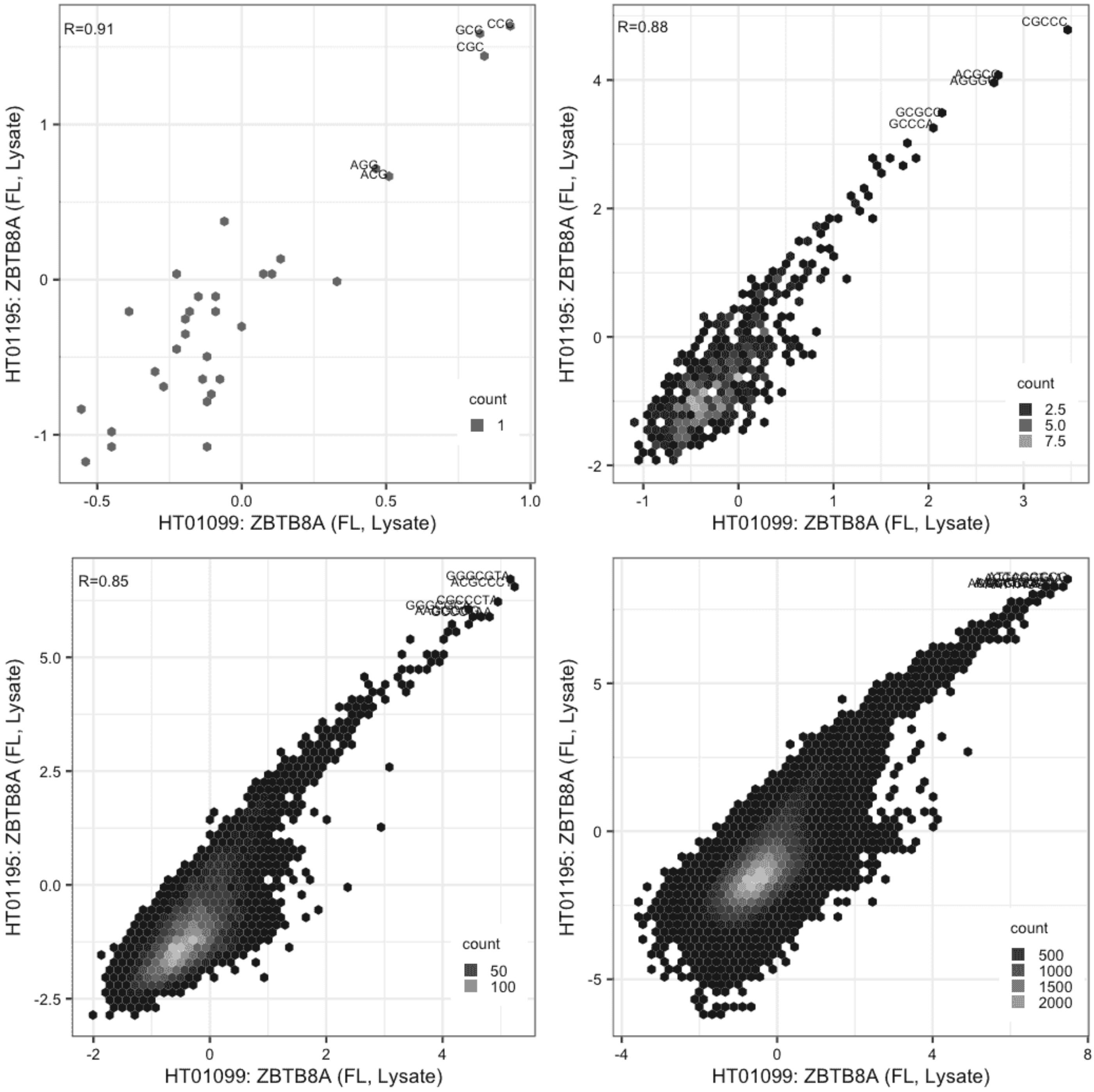

**ZBTB8B**

Lysate vs IVT, different constructs

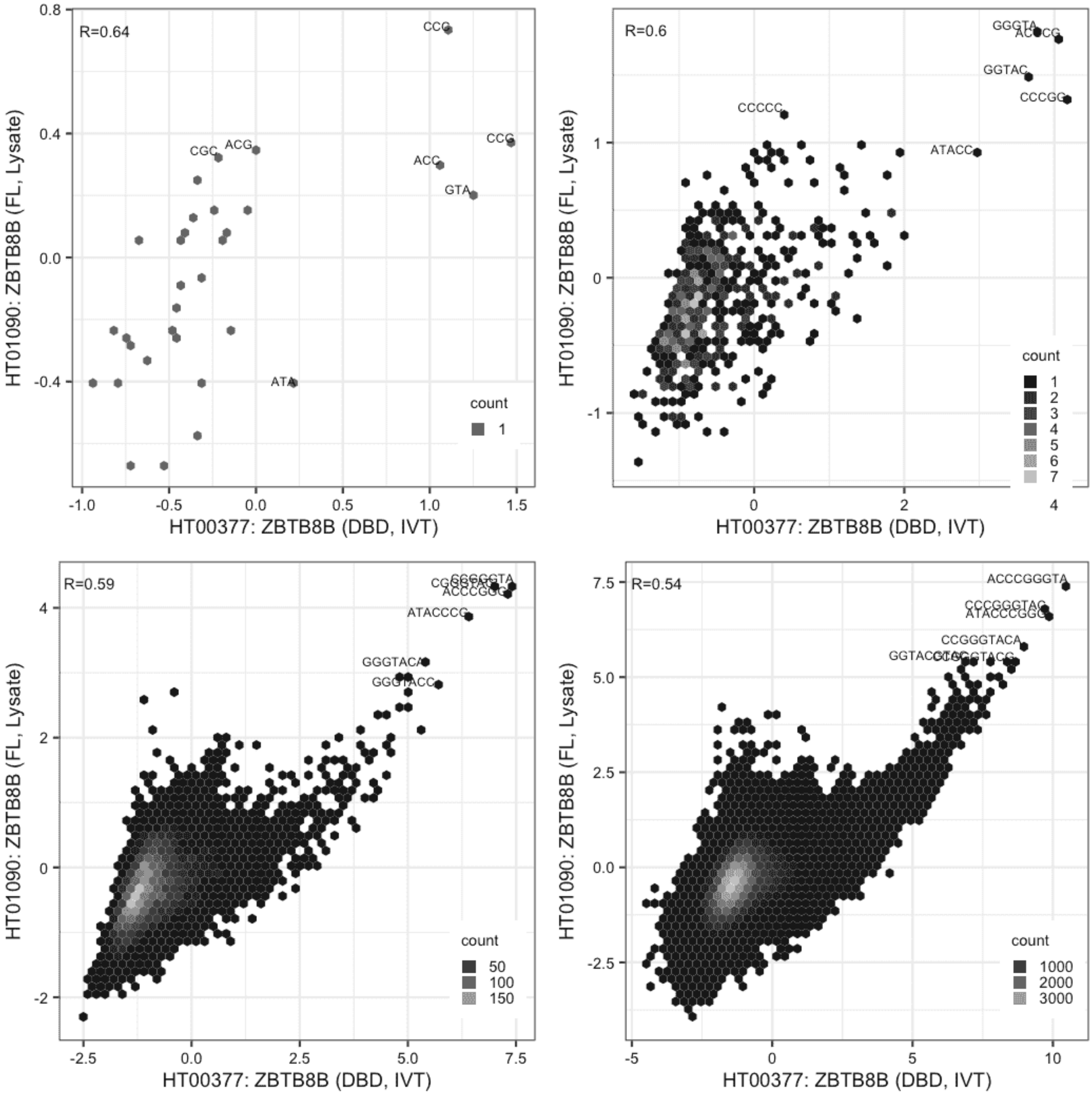

#### ZBTB8B

Lysate vs IVT, different constructs

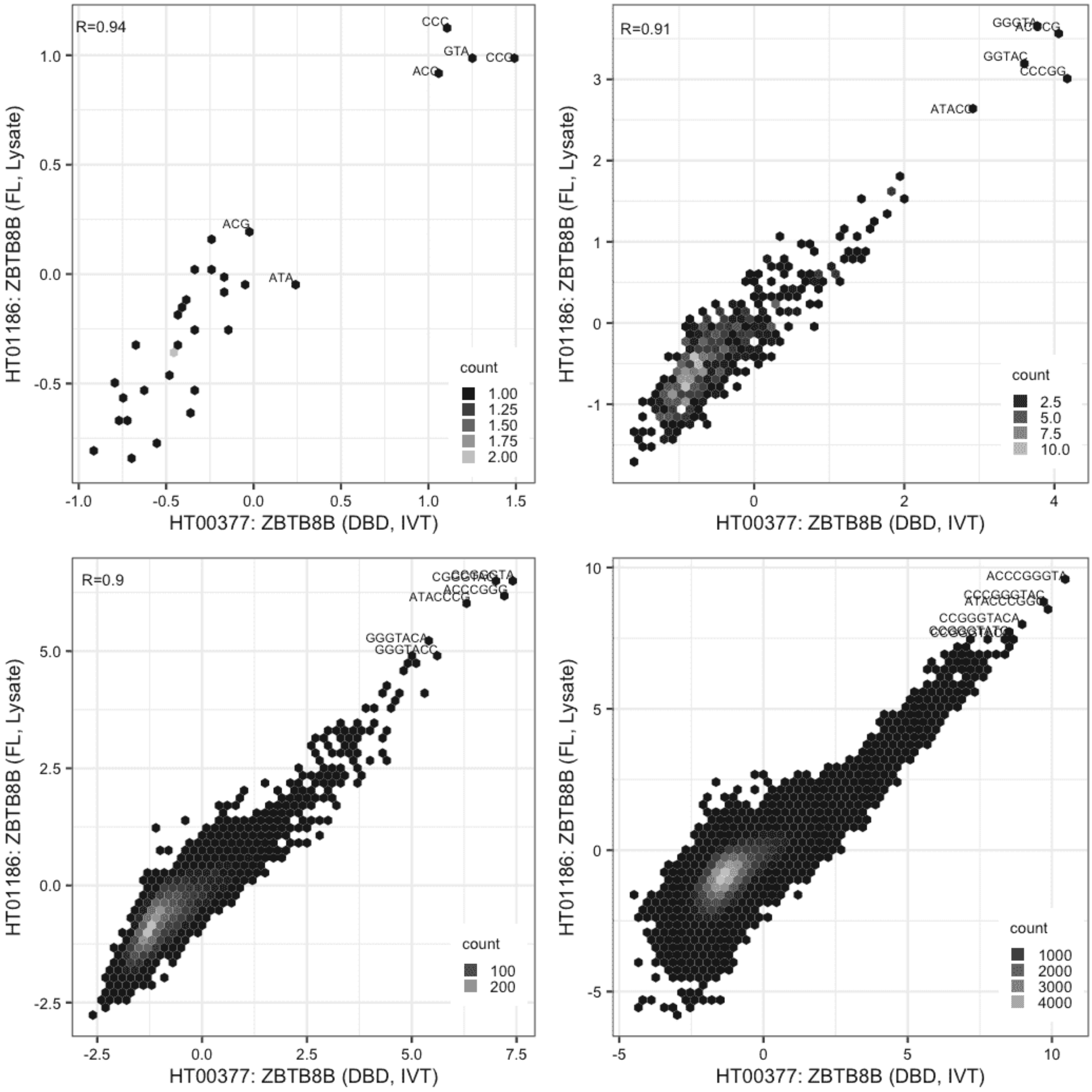

#### ZBTB8B

Lysate vs Lysate, same construct

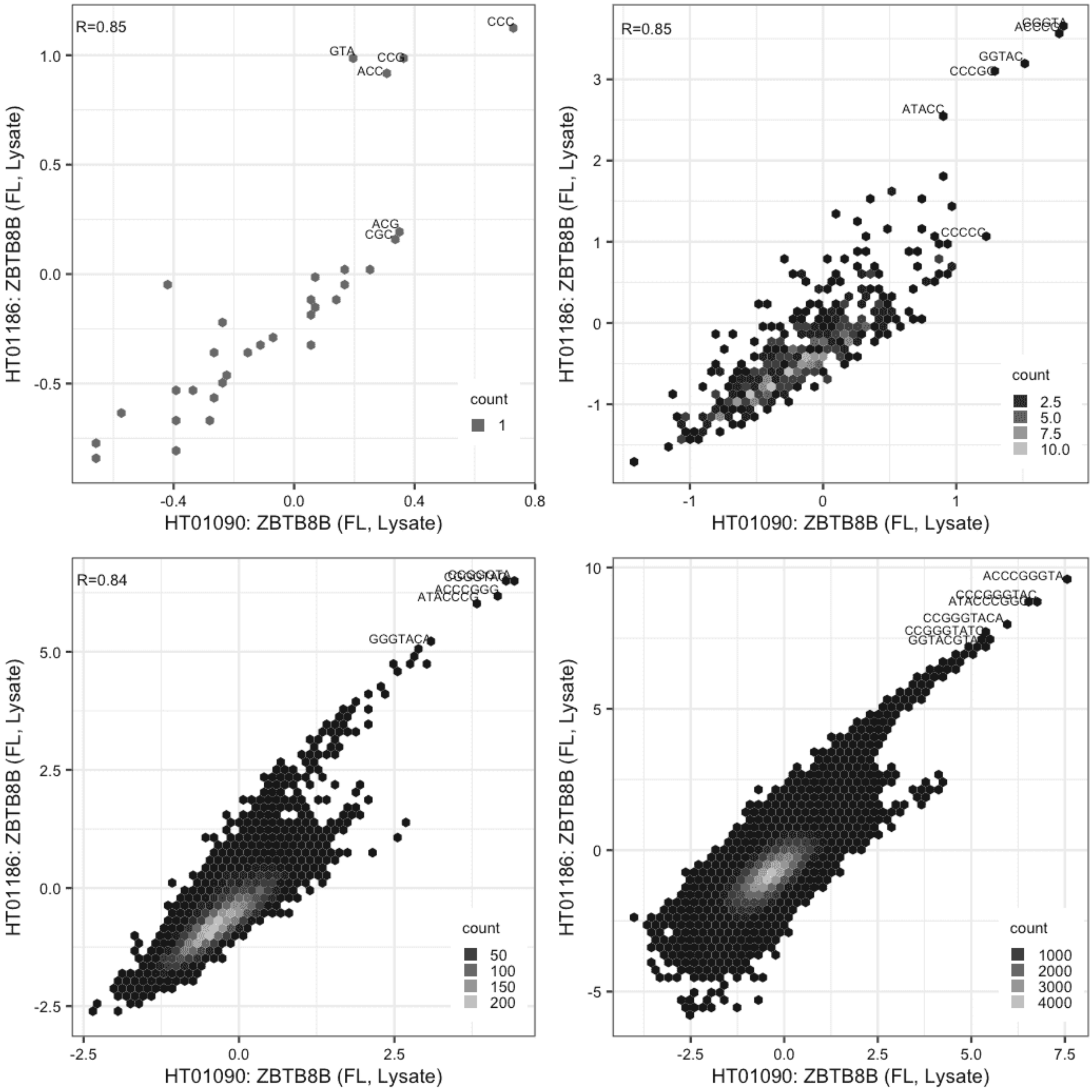

**ZFP3**

Lysate vs IVT, same construct

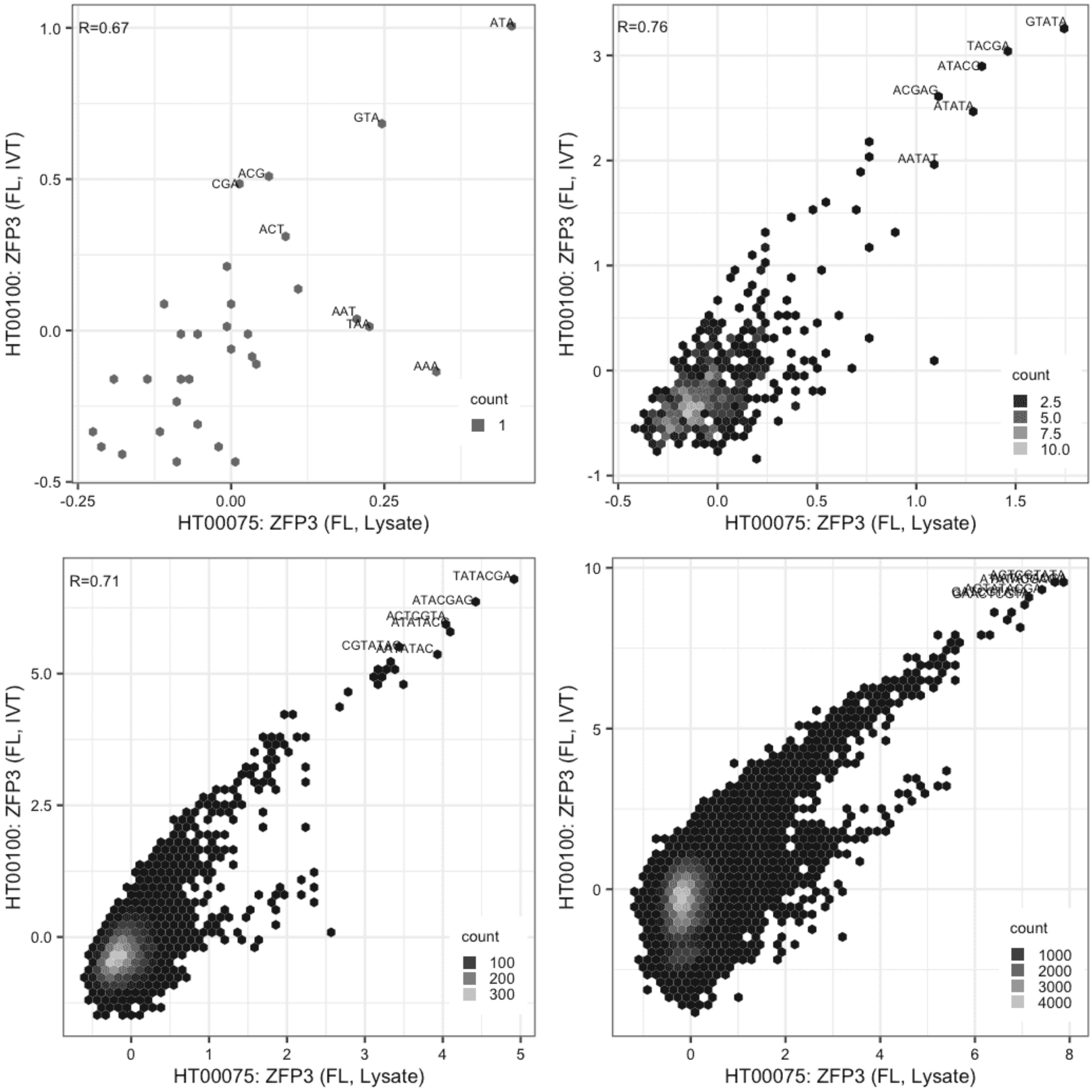

**ZGLP1**

Lysate vs IVT, different constructs

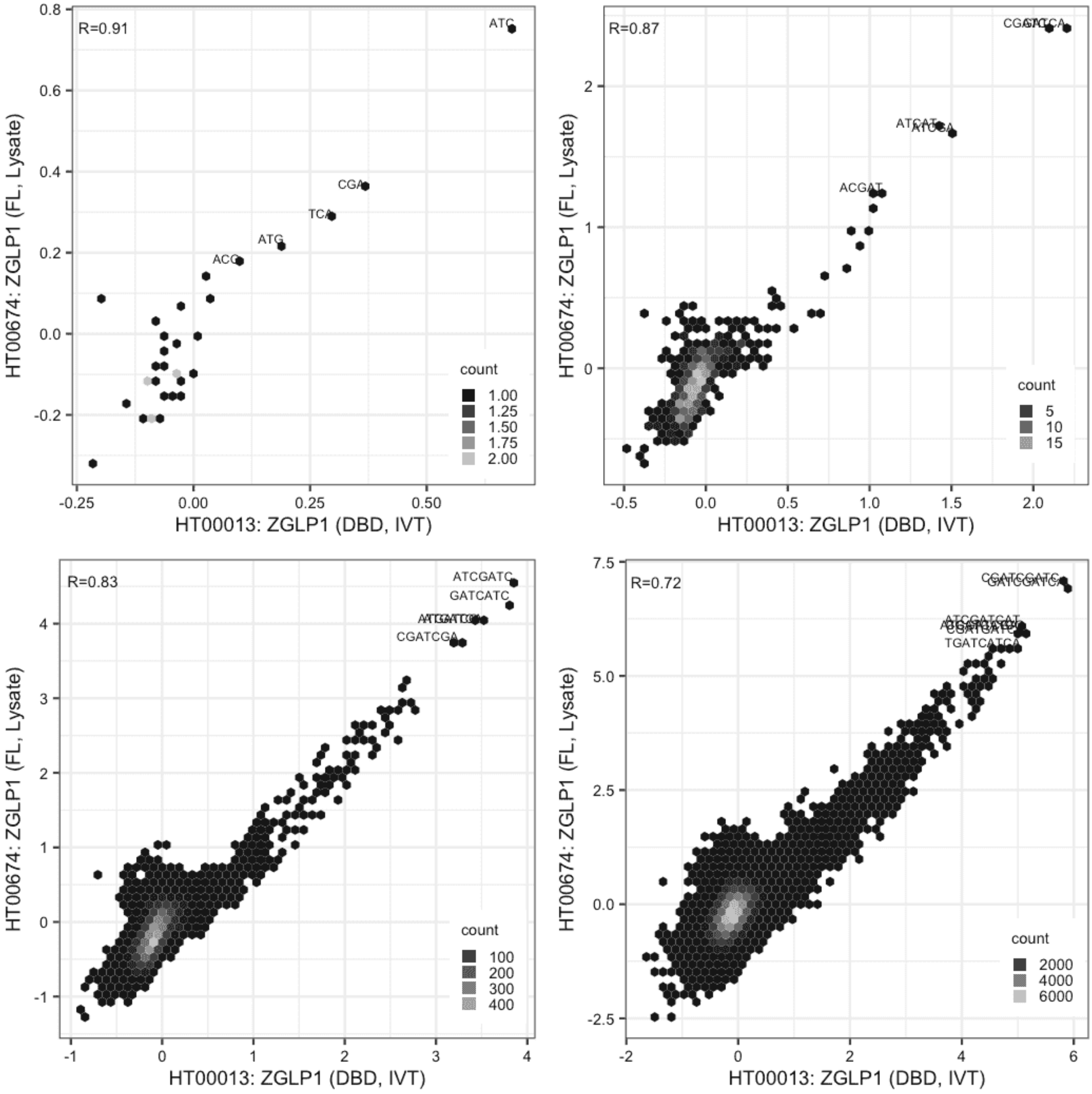

**ZGLP1**

Lysate vs IVT, different constructs

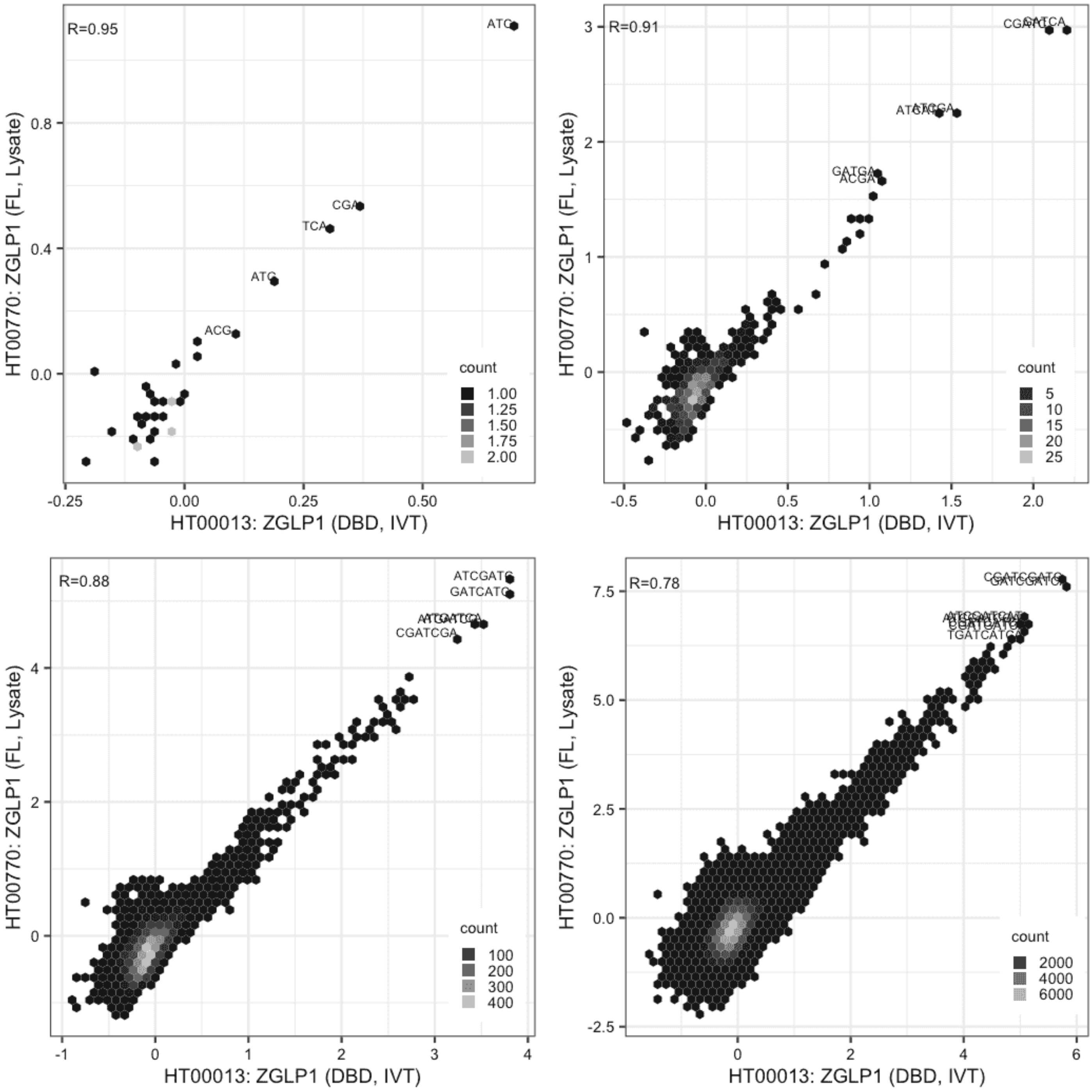

#### ZGLP1

Lysate vs Lysate, same construct

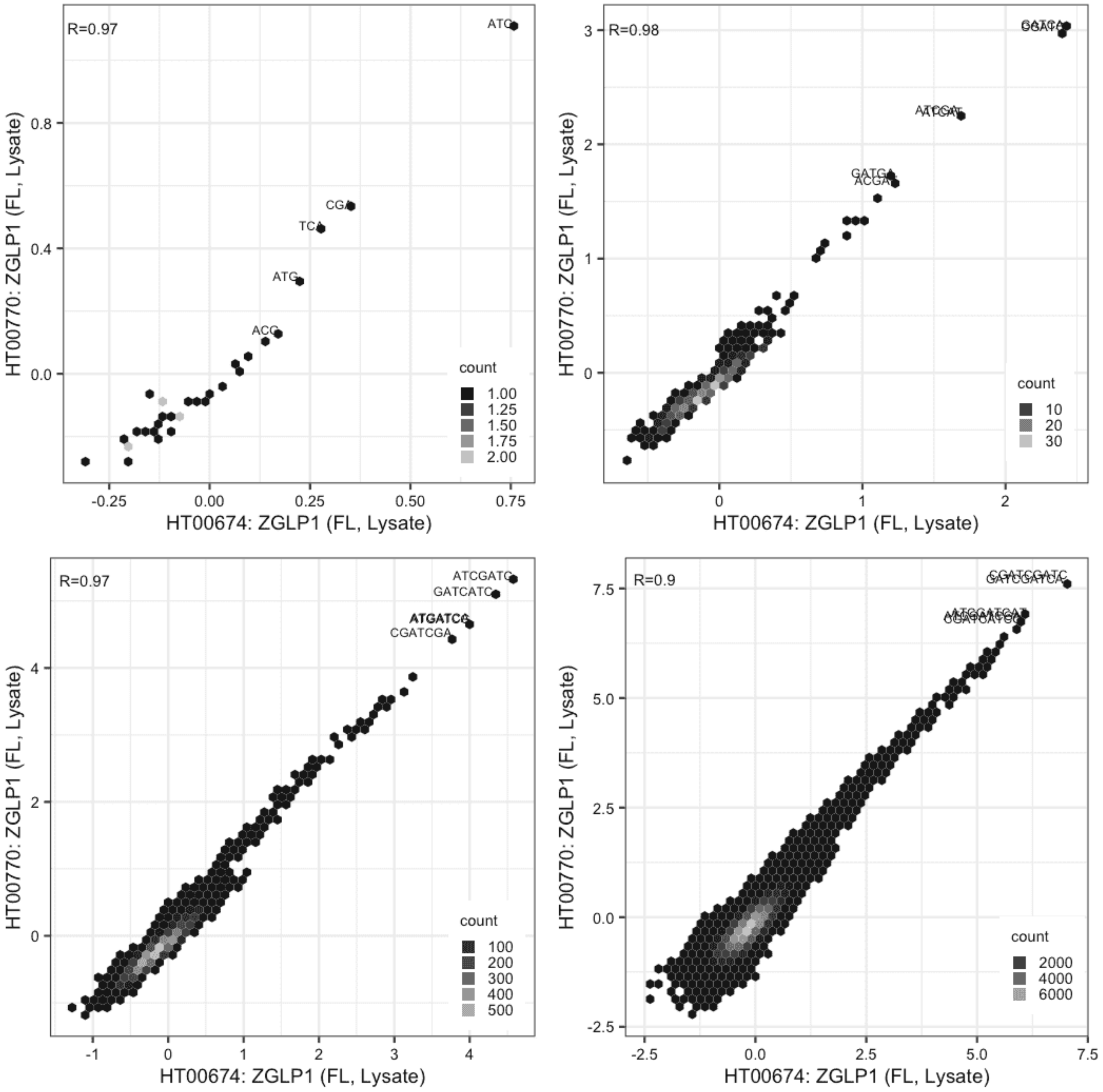

**ZNF131**

Lysate vs IVT, different constructs

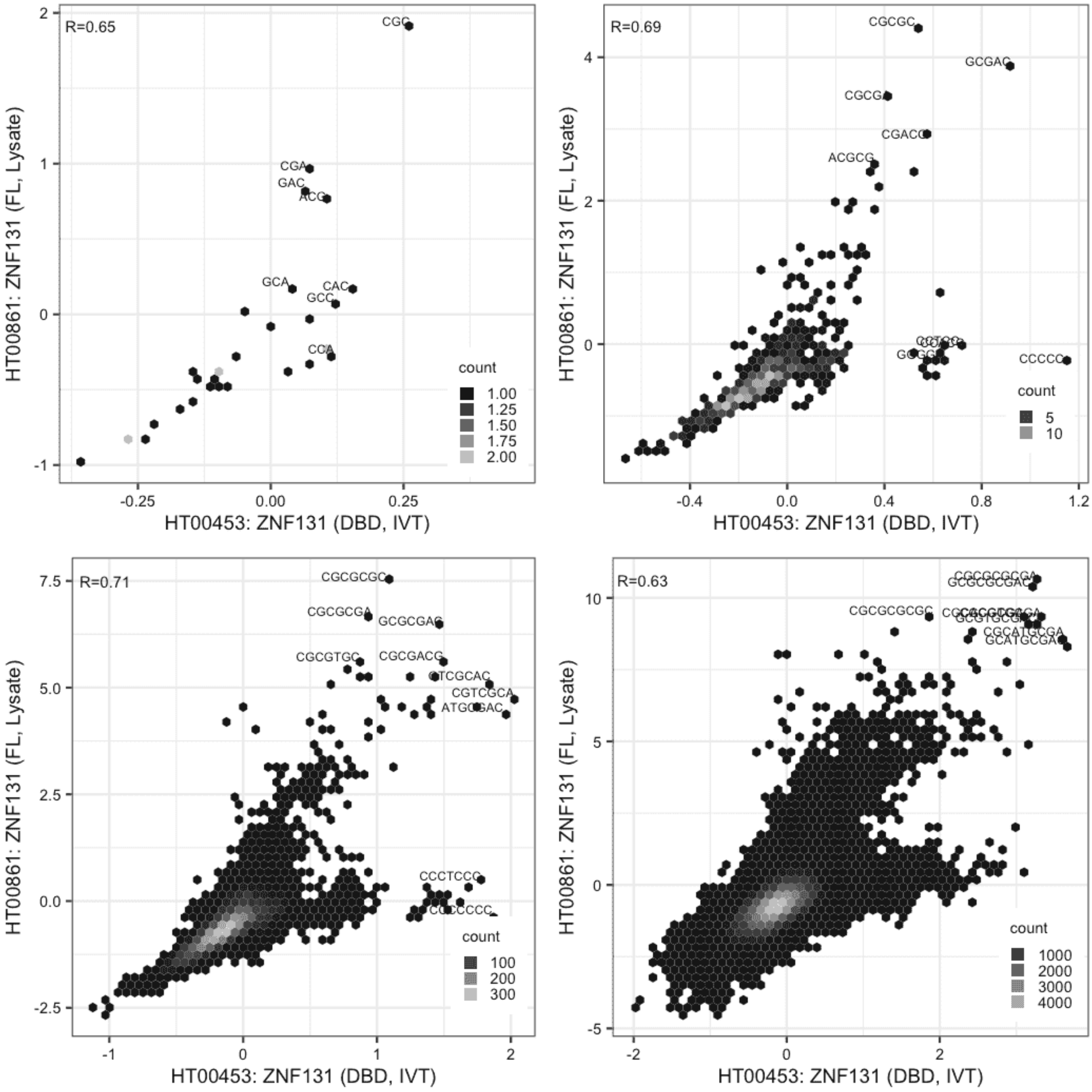

**ZNF131**

IVT vs GFPIVT, same construct

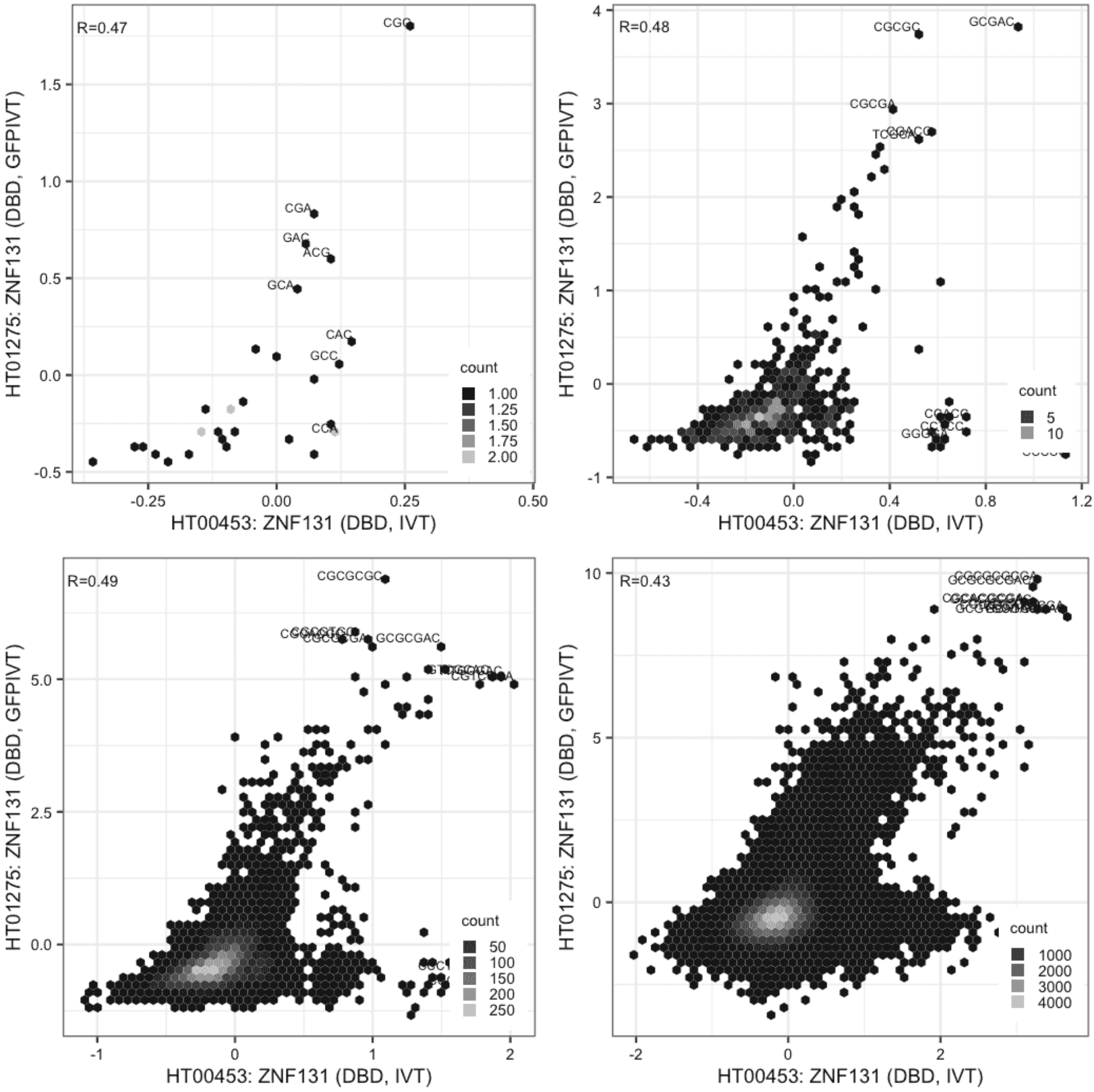

#### ZNF131

IVT vs GFPIVT, different constructs

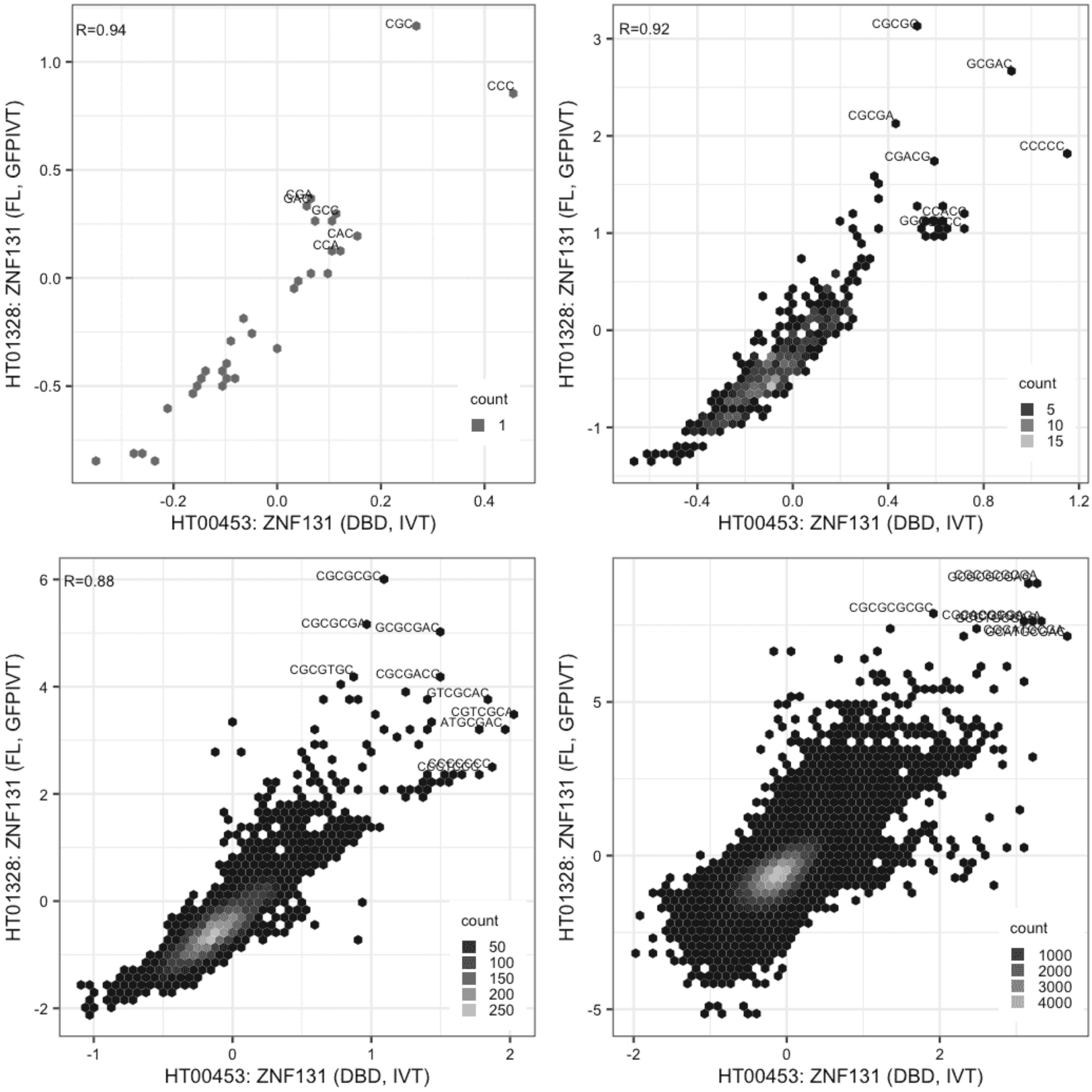

**ZNF131**

Lysate vs GFPIVT, different constructs

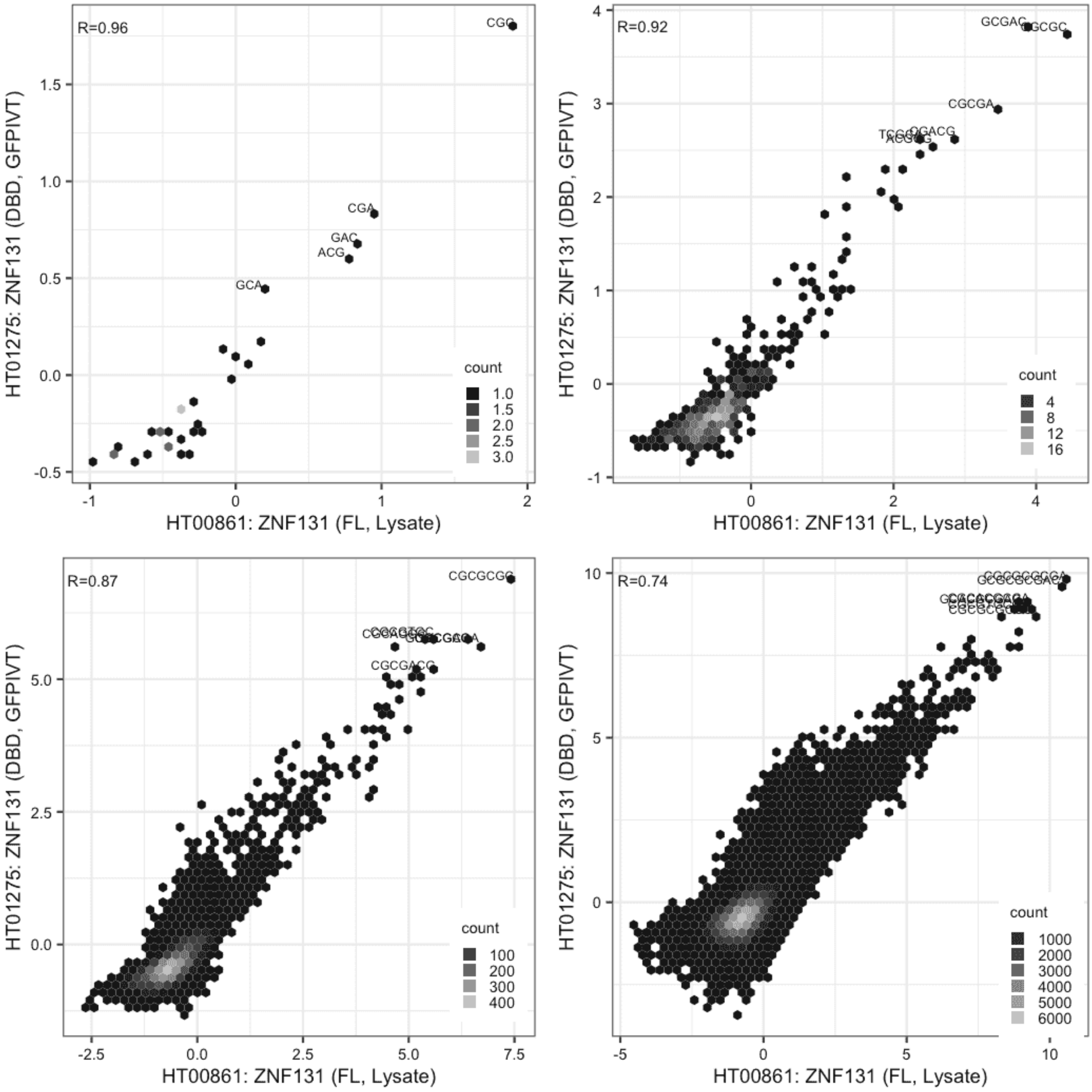

**ZNF131**

Lysate vs GFPIVT, same construct

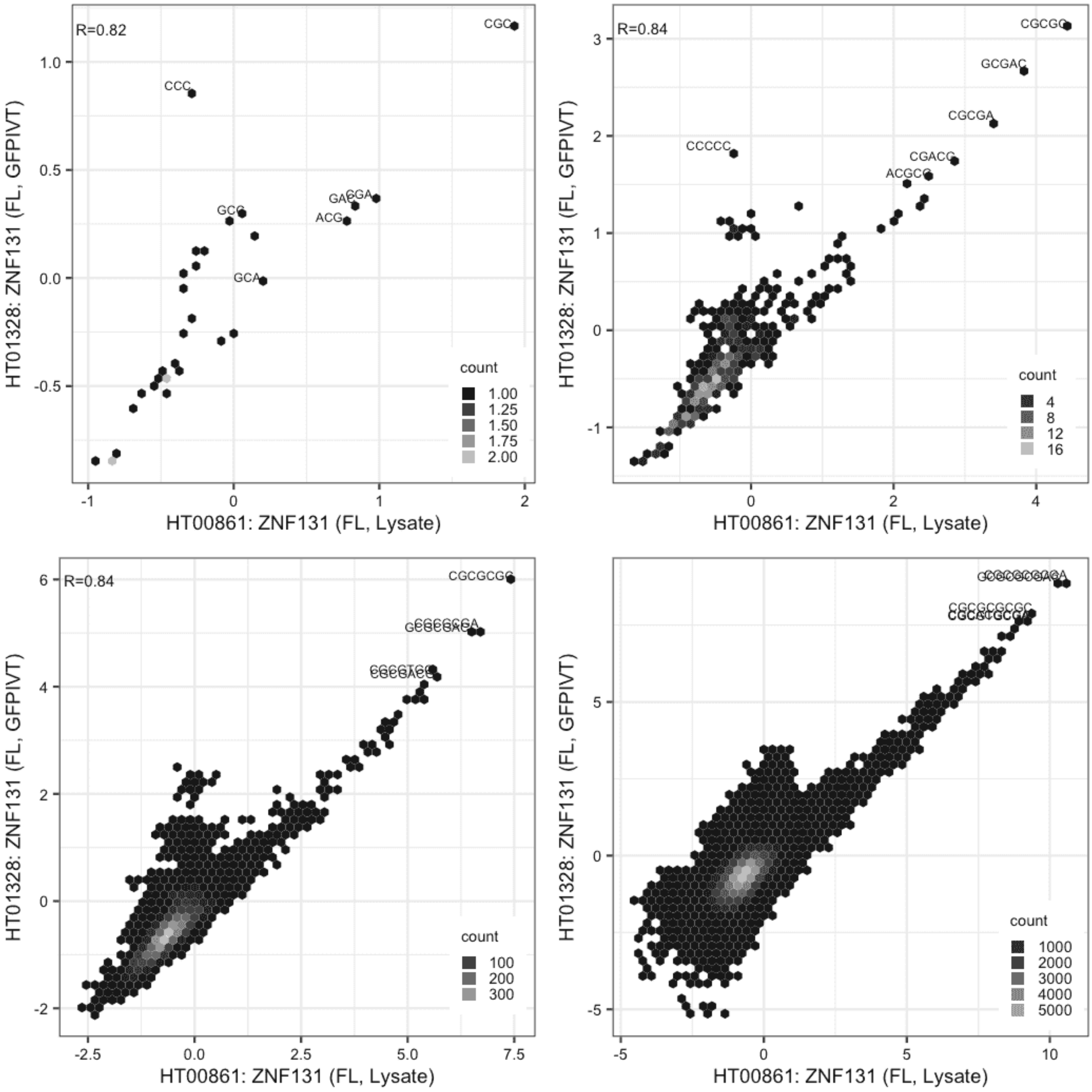

**ZNF131**

GFPIVT vs GFPIVT, different constructs

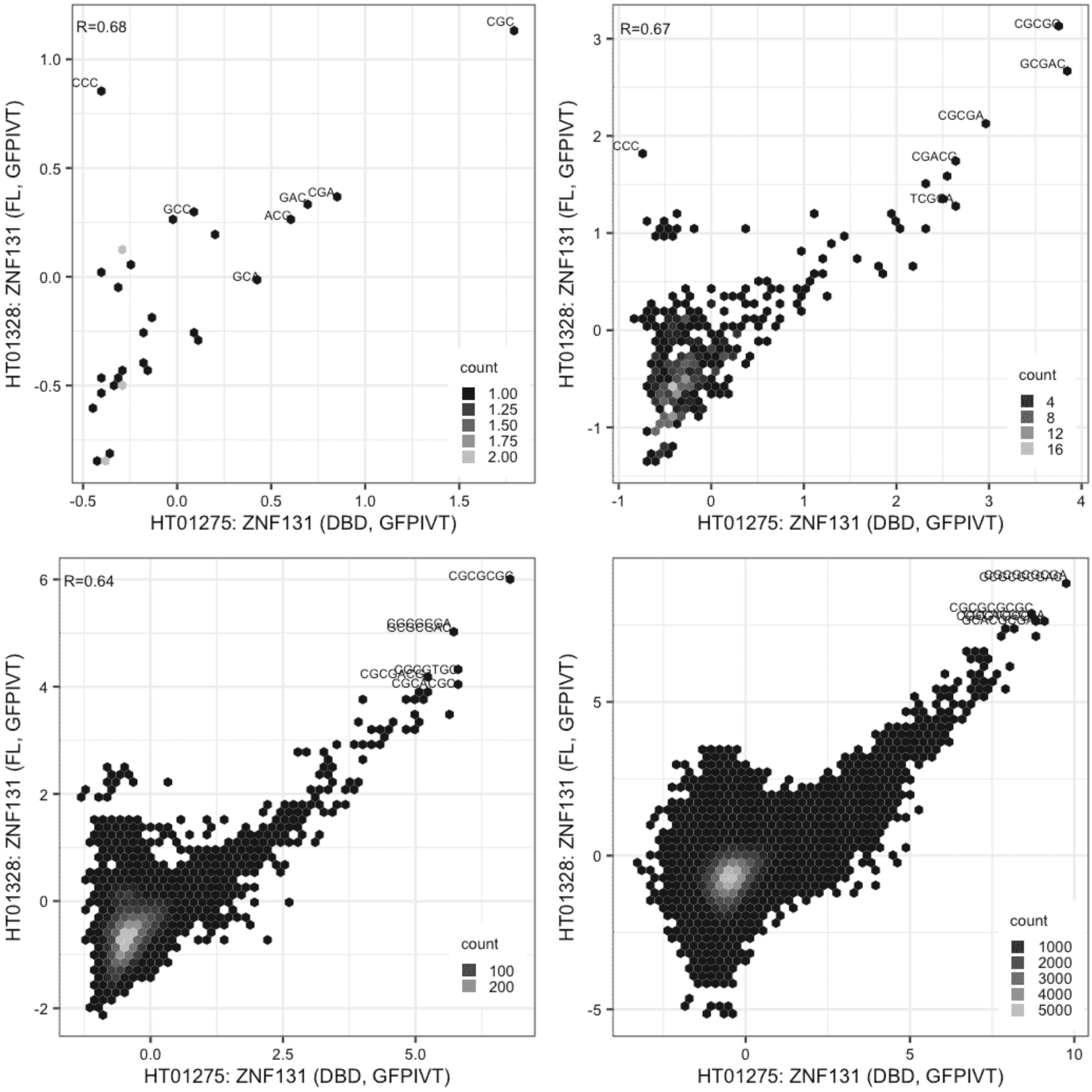

**ZNF362**

Lysate vs IVT, different constructs

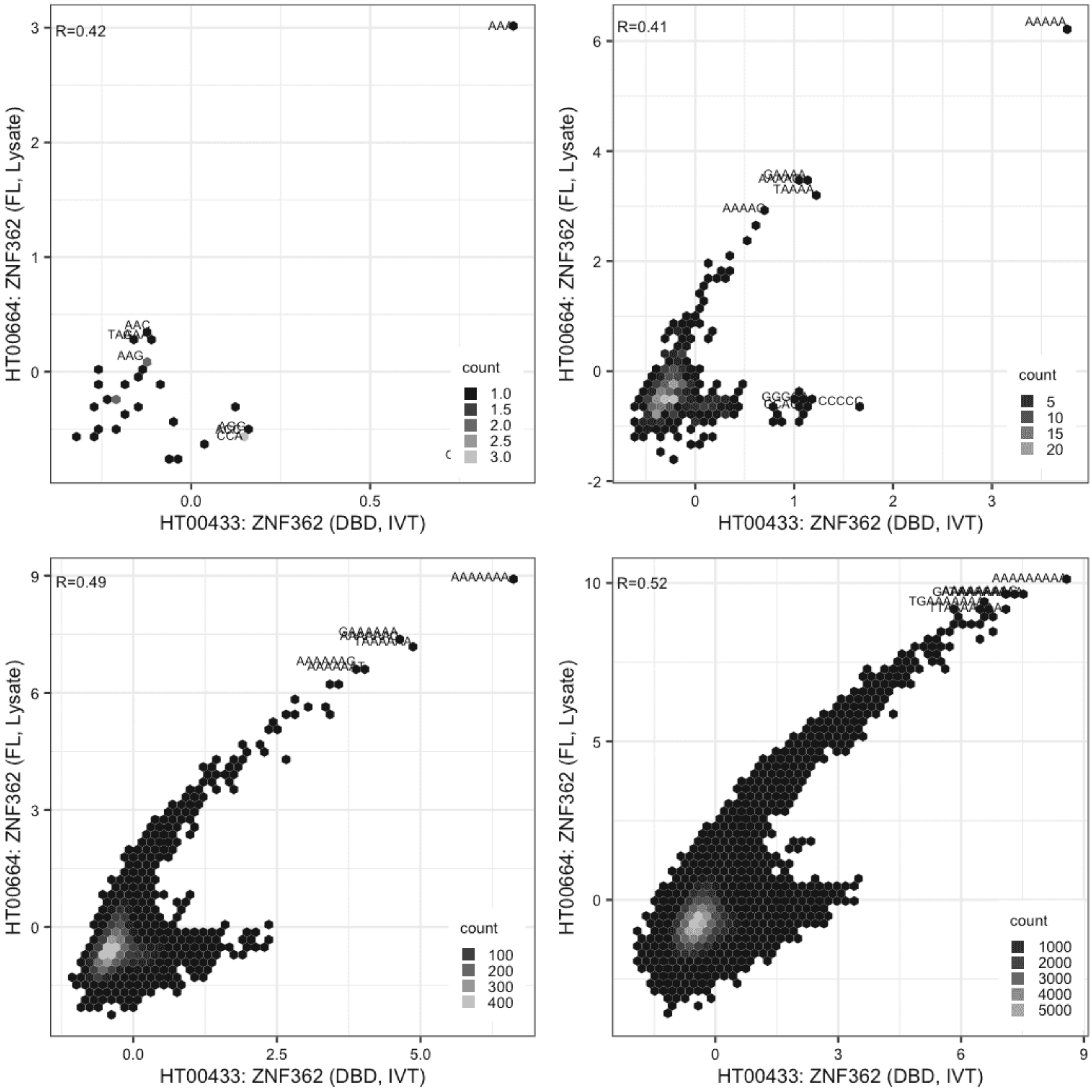

#### ZNF362

Lysate vs IVT, different constructs

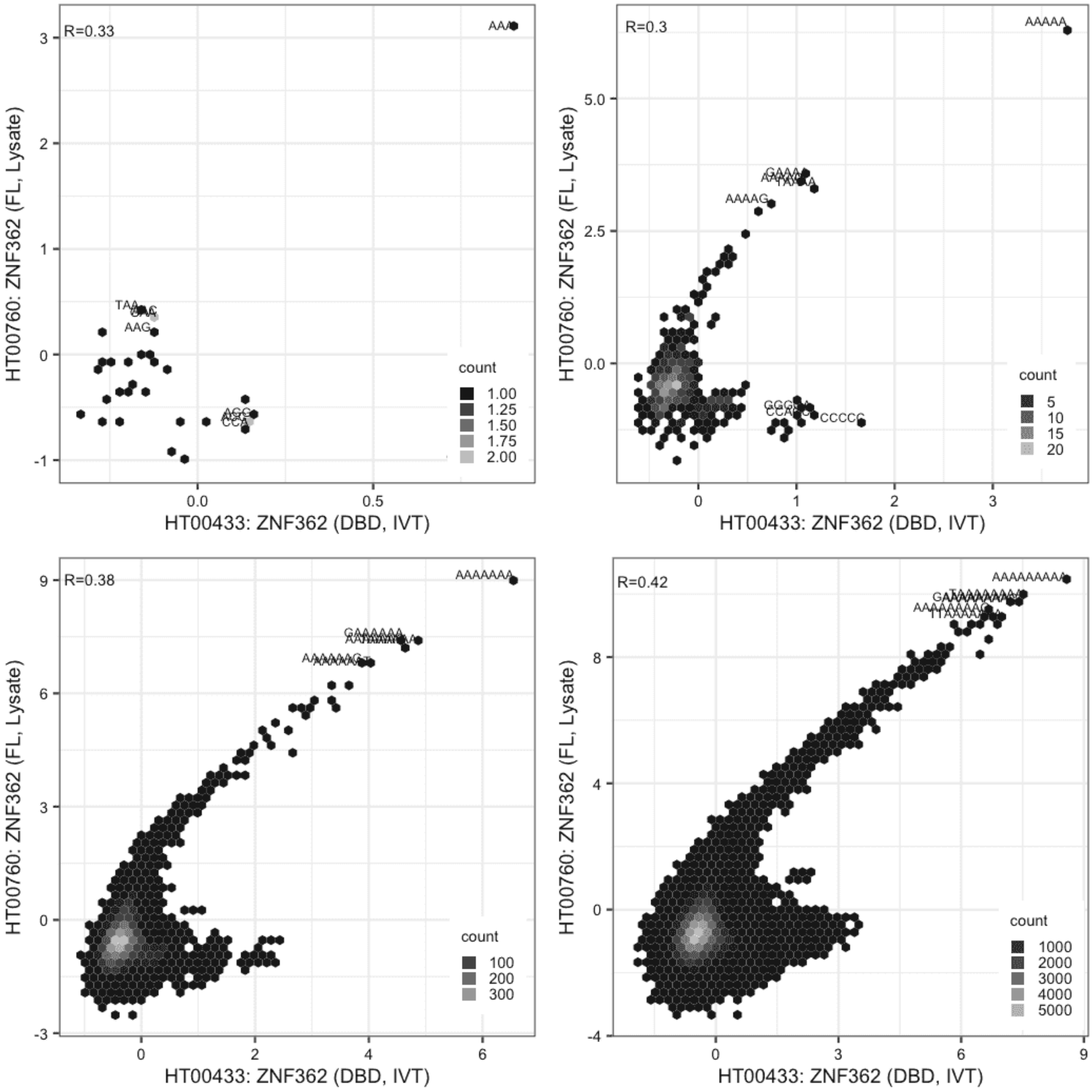

#### ZNF362

IVT vs GFPIVT, different constructs

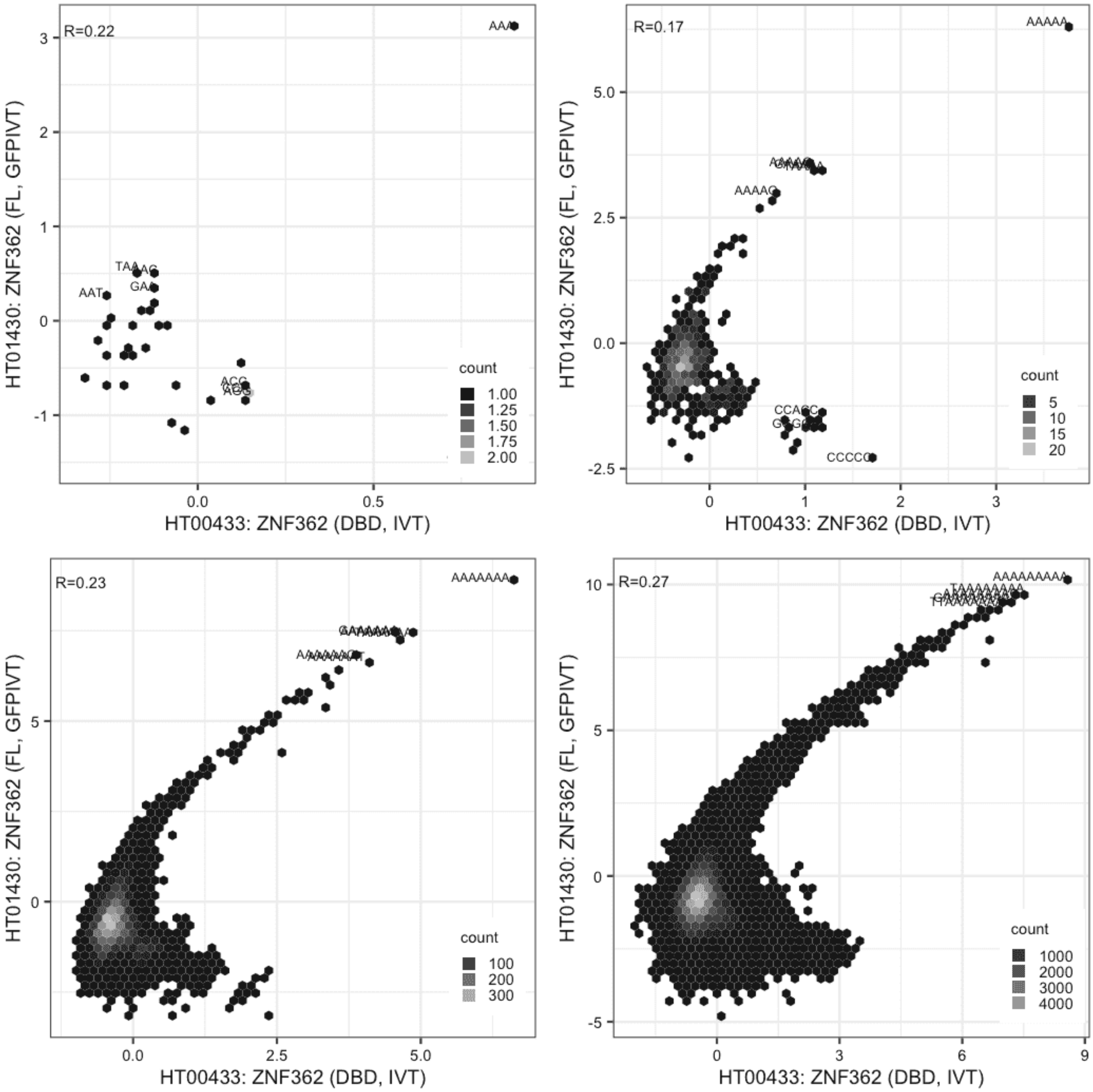

**ZNF362**

Lysate vs Lysate, same construct

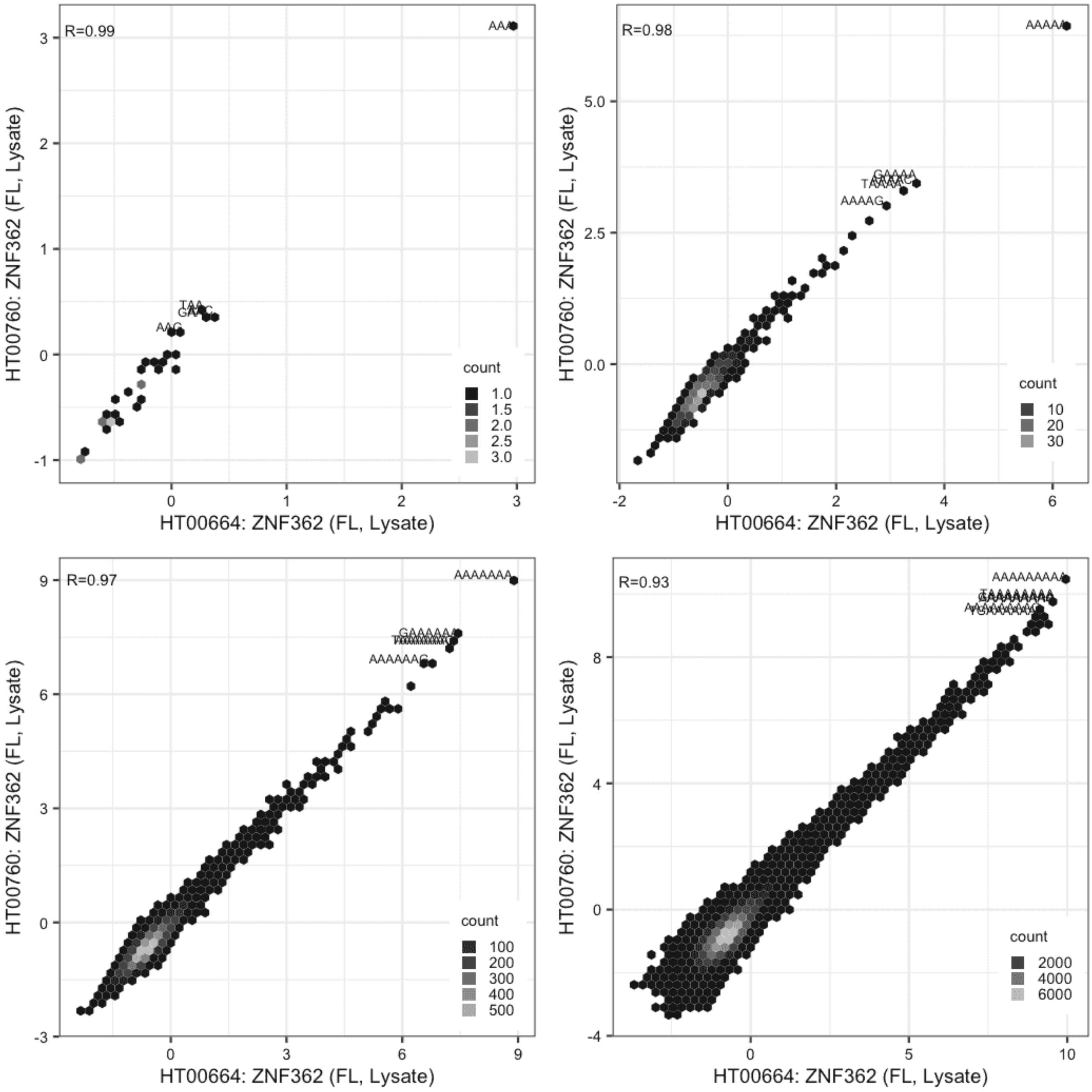

#### ZNF362

Lysate vs GFPIVT, same construct

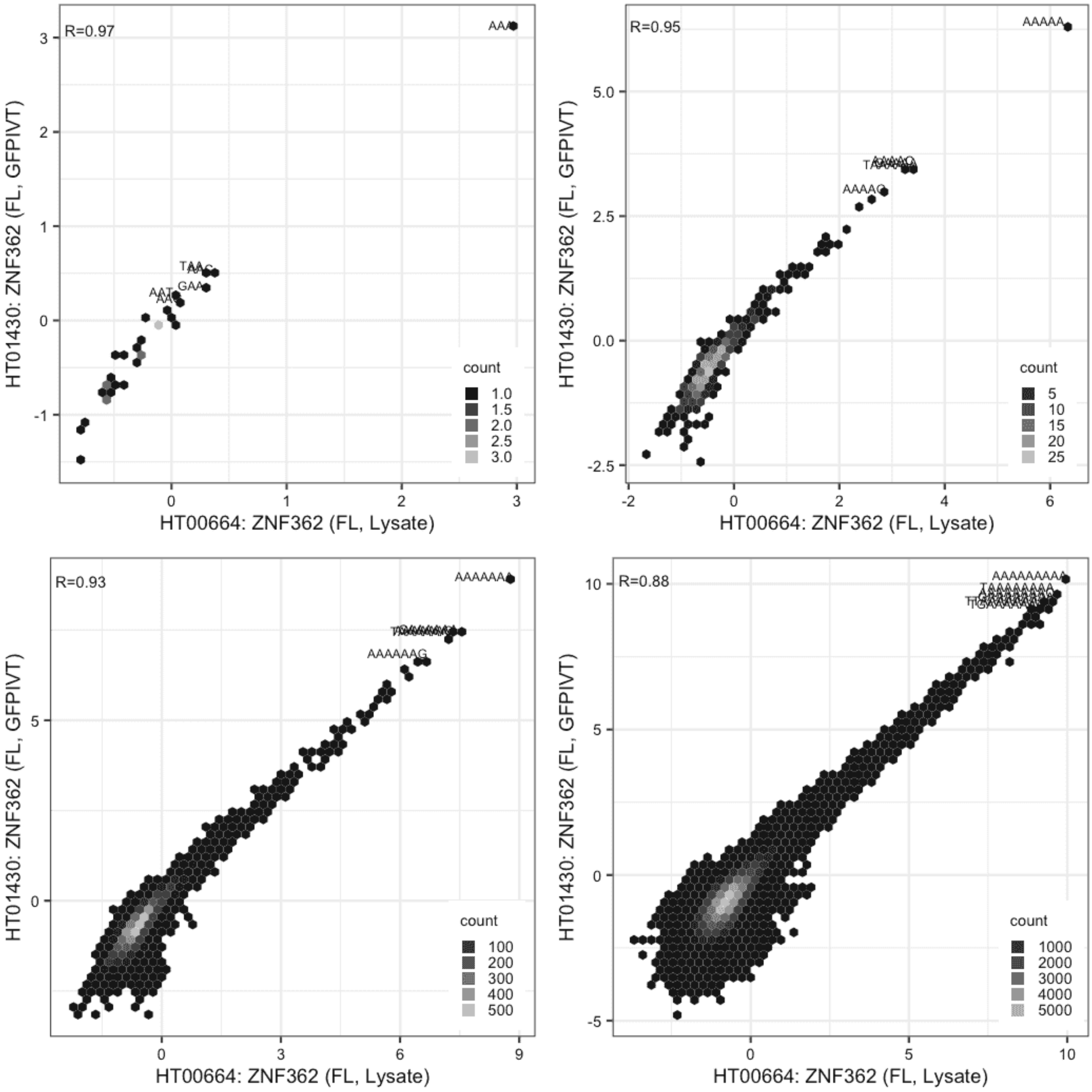

#### ZNF362

Lysate vs GFPIVT, same construct

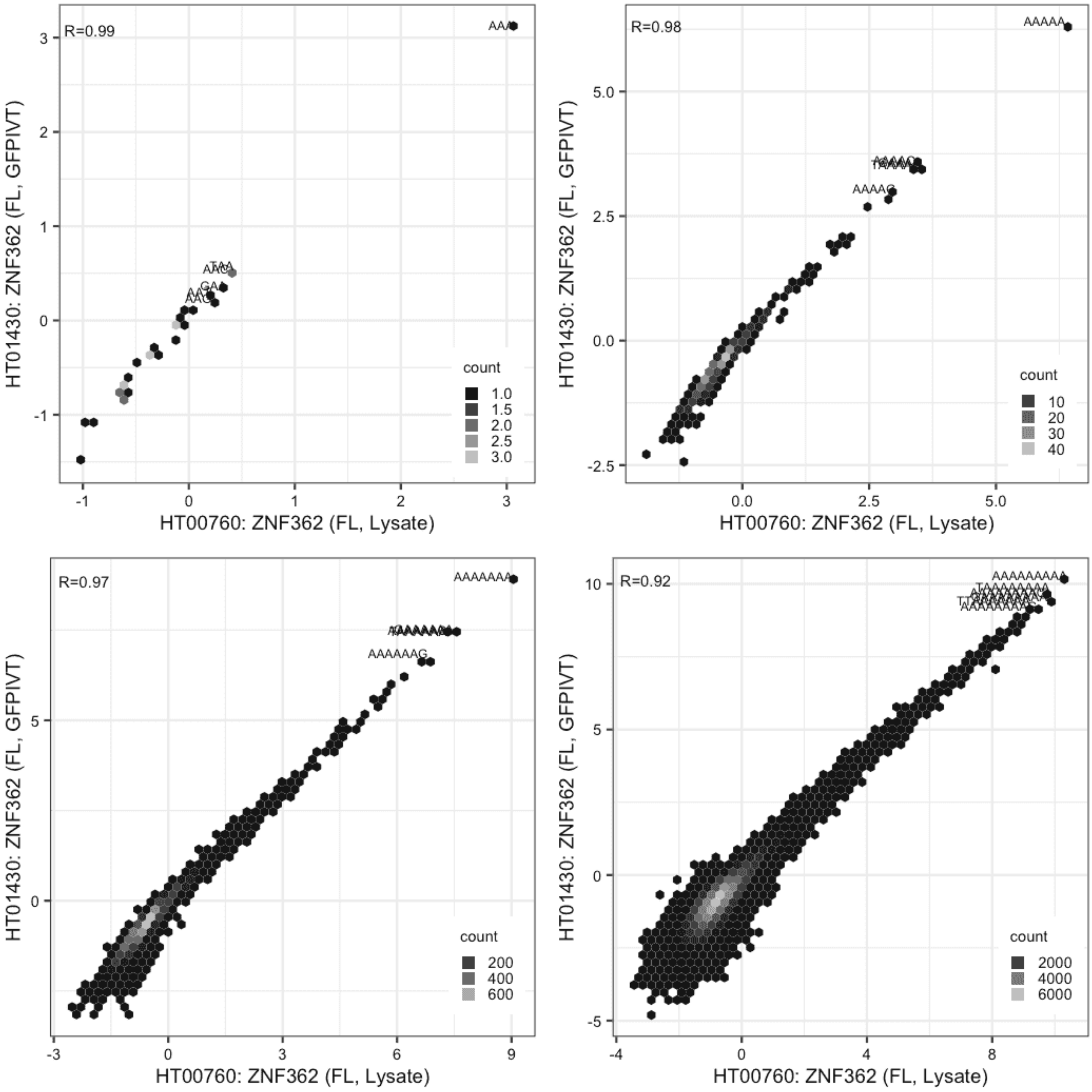

**ZNF367**

Lysate vs GFPIVT, same construct

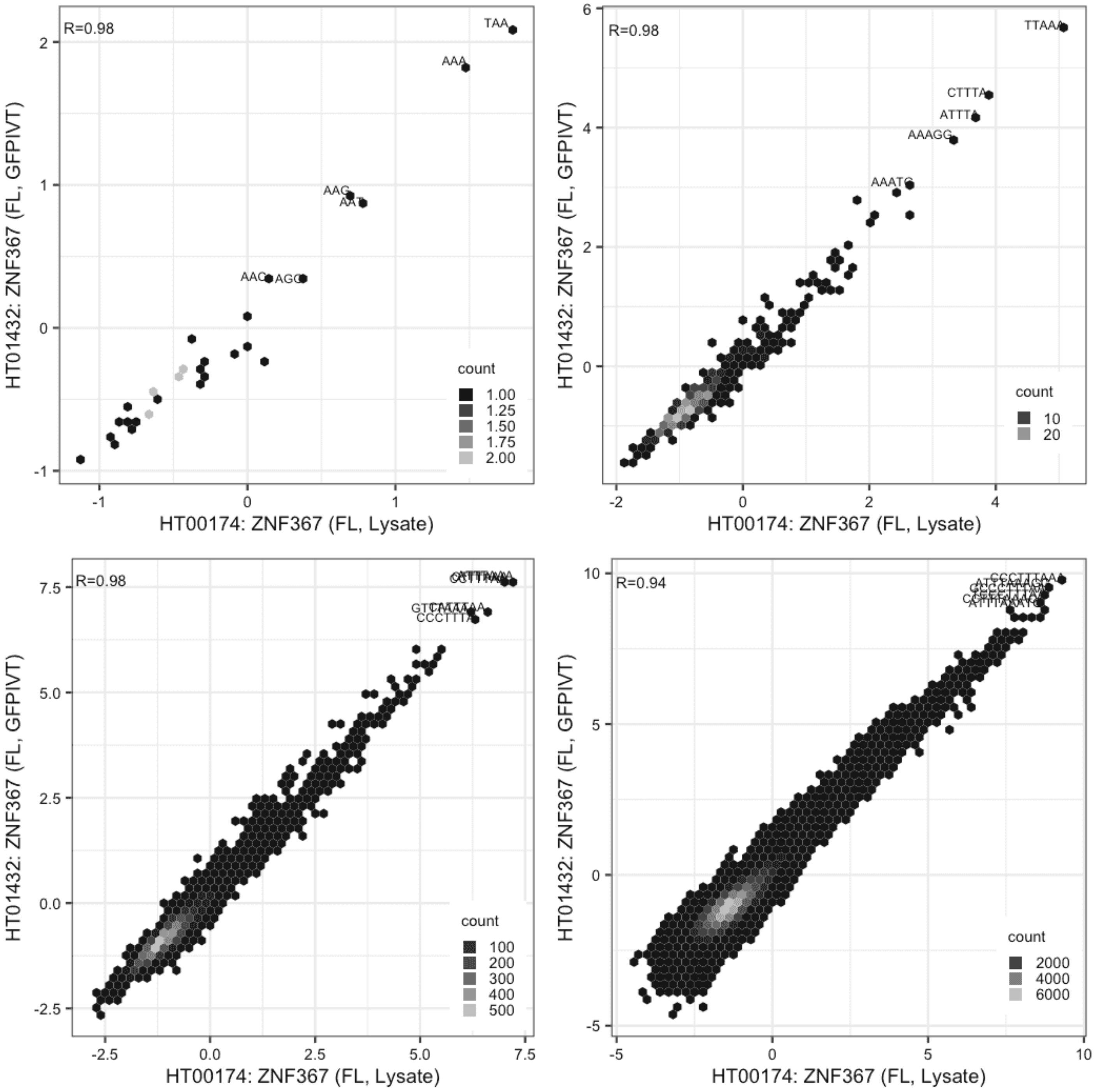

**ZNF395**

Lysate vs IVT, different constructs

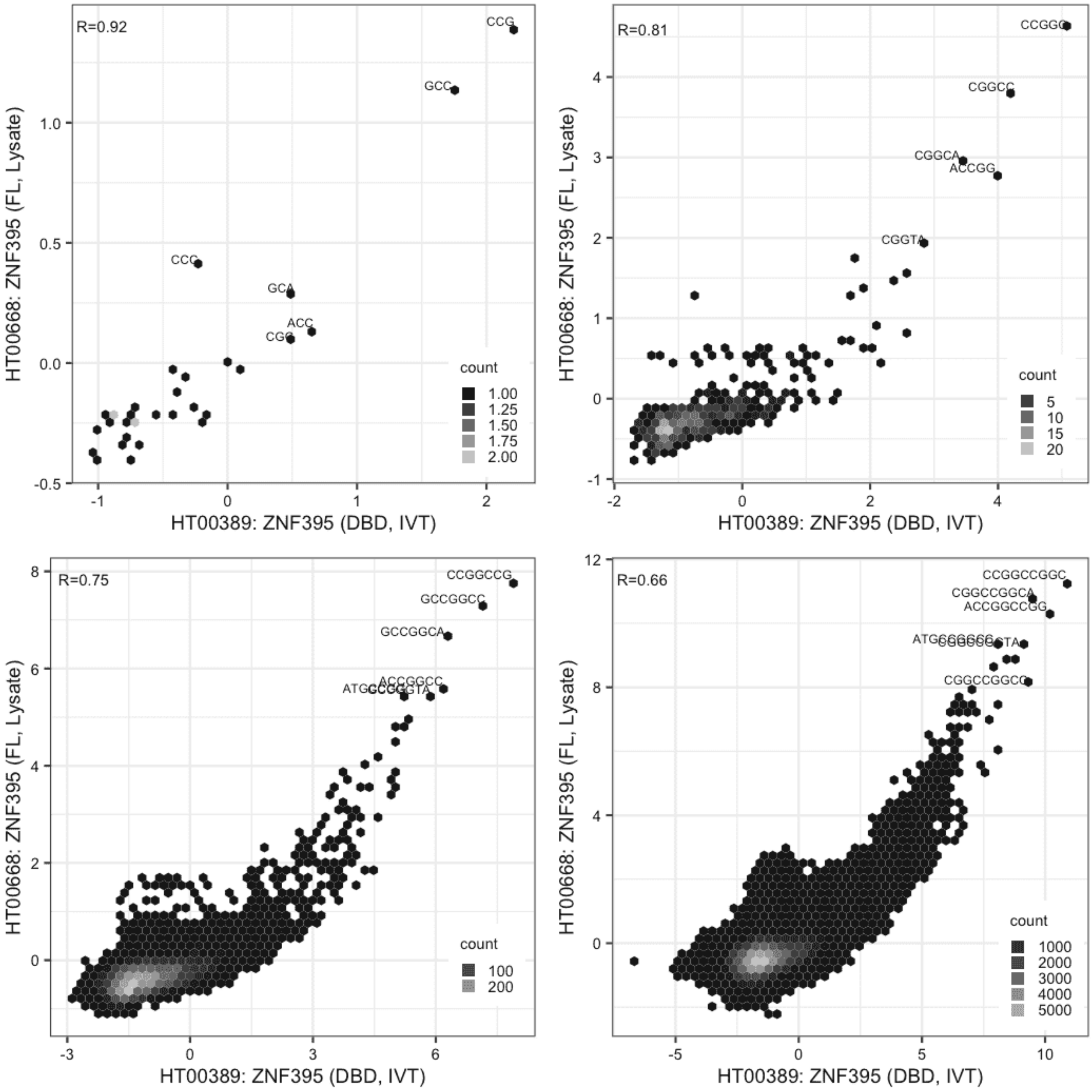

#### ZNF395

Lysate vs IVT, different constructs

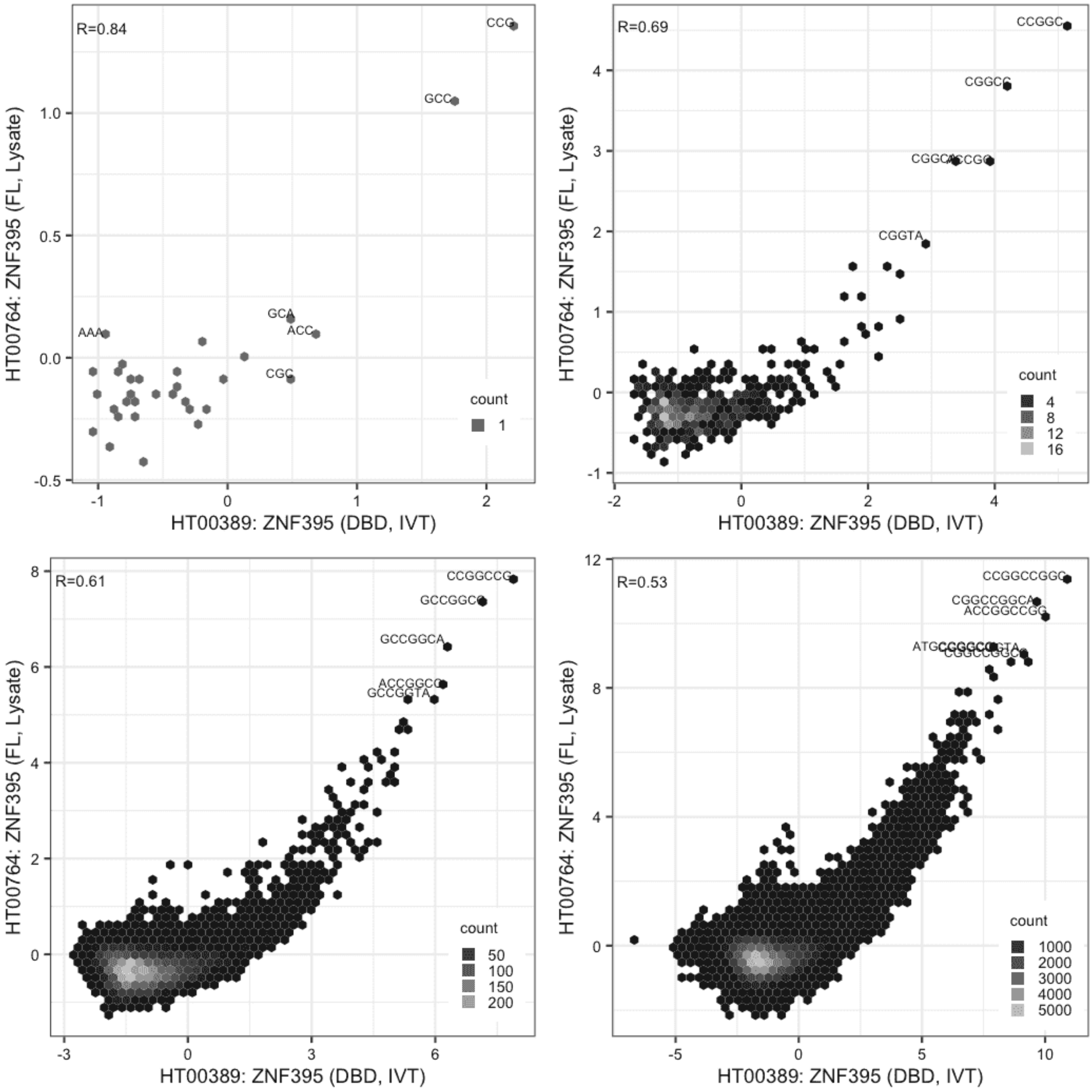

**ZNF395**

IVT vs GFPIVT, same construct

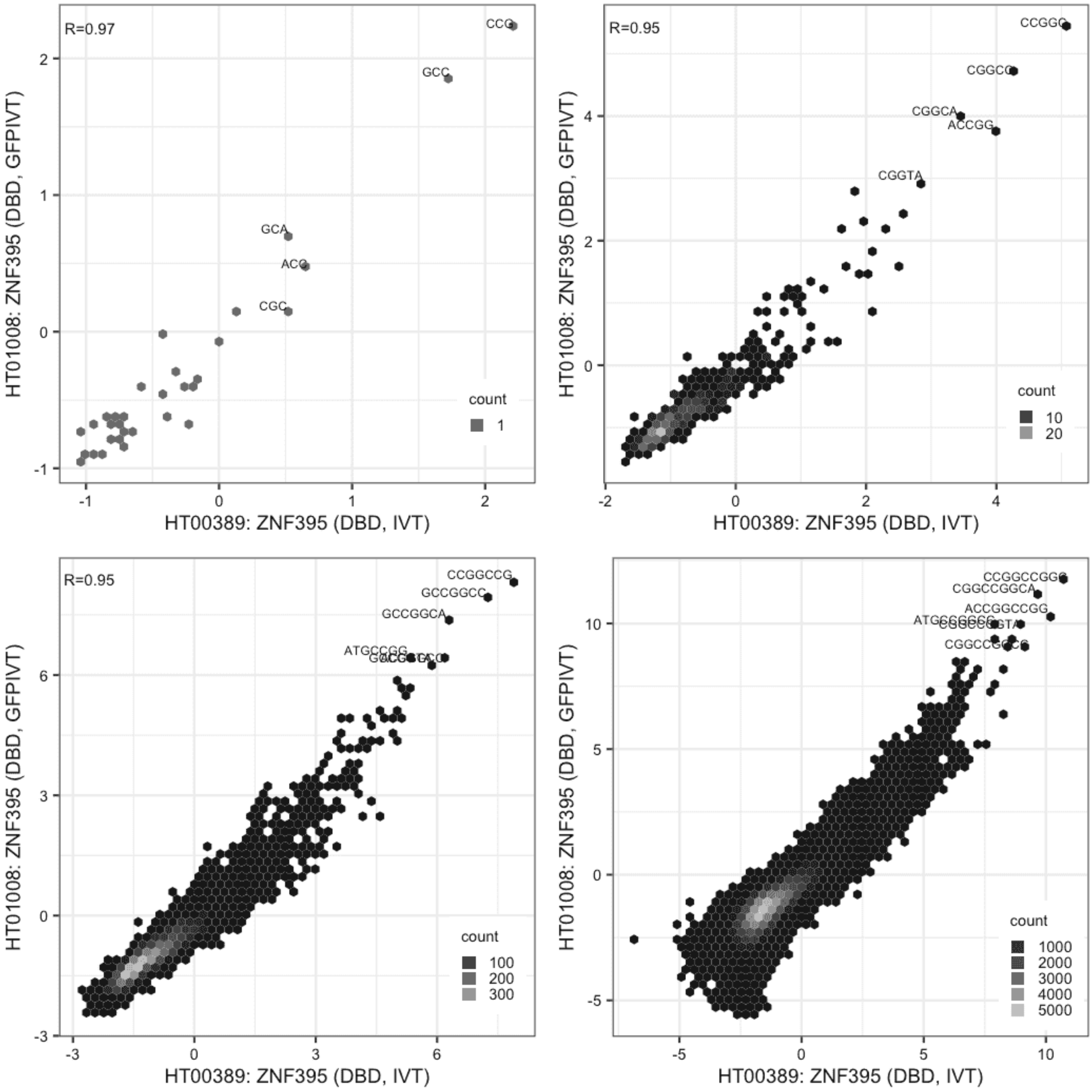

**ZNF395**

IVT vs GFPIVT, different constructs

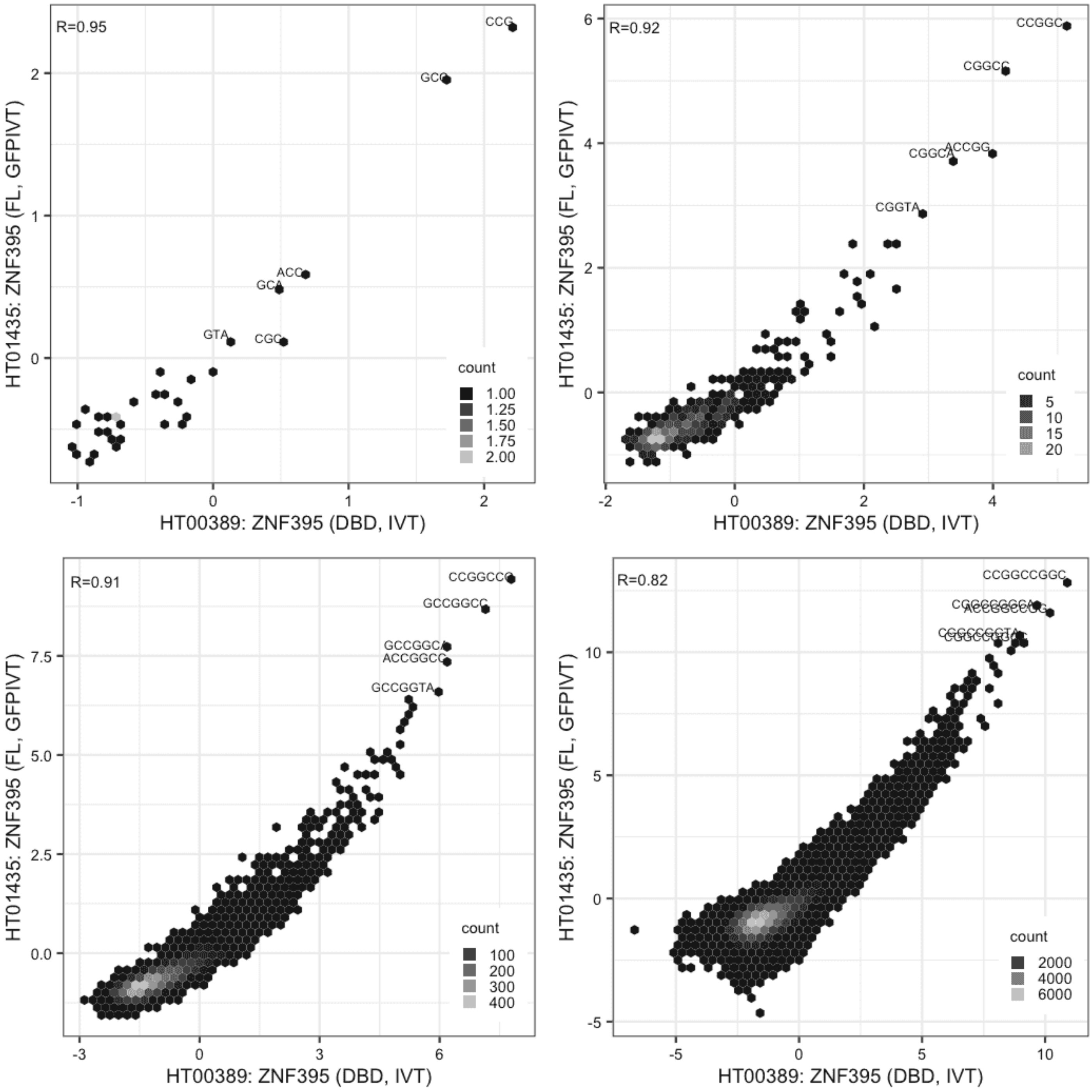

#### ZNF395

Lysate vs Lysate, same construct

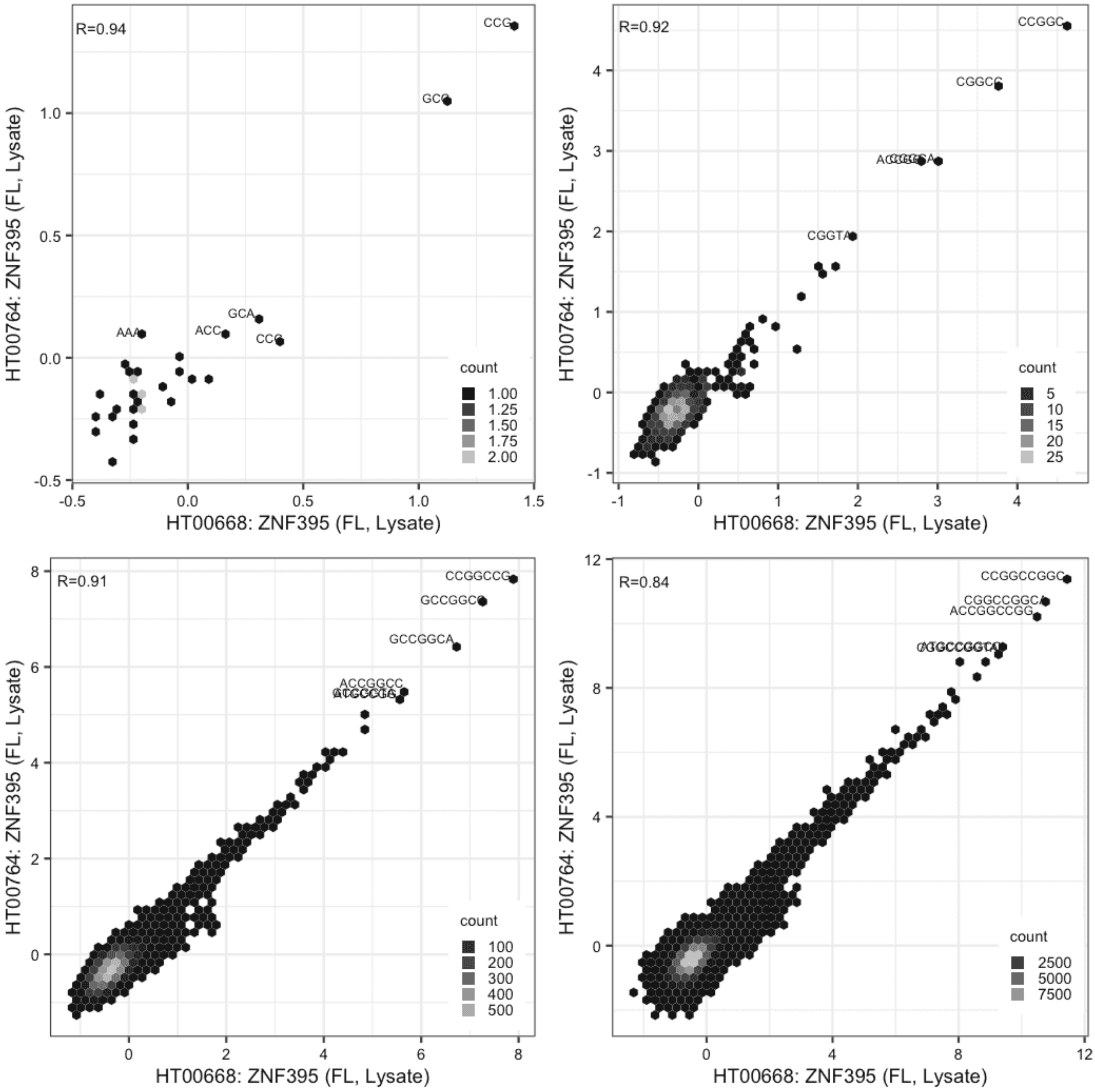

#### ZNF395

Lysate vs GFPIVT, different constructs

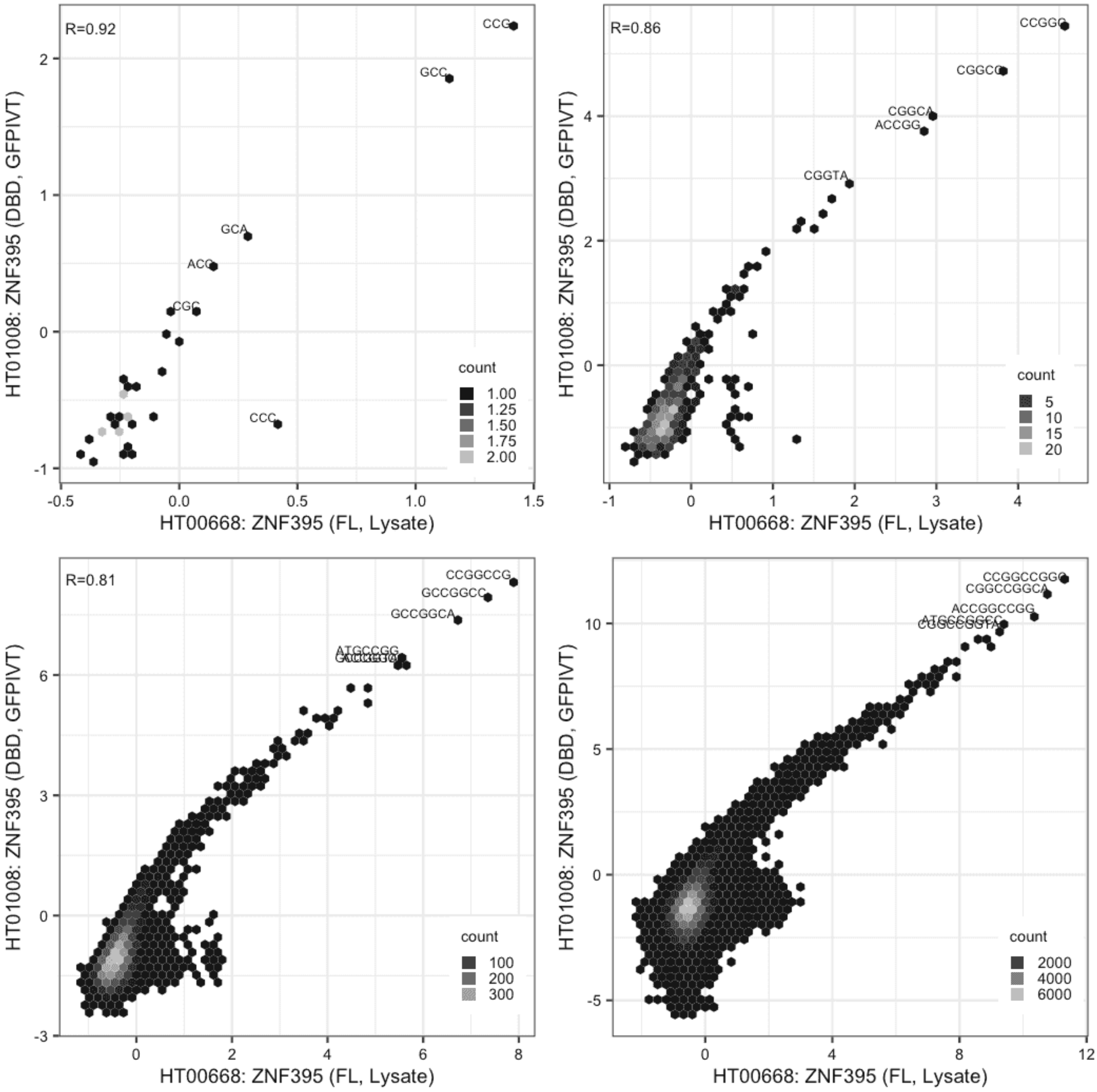

**ZNF395**

Lysate vs GFPIVT, same construct

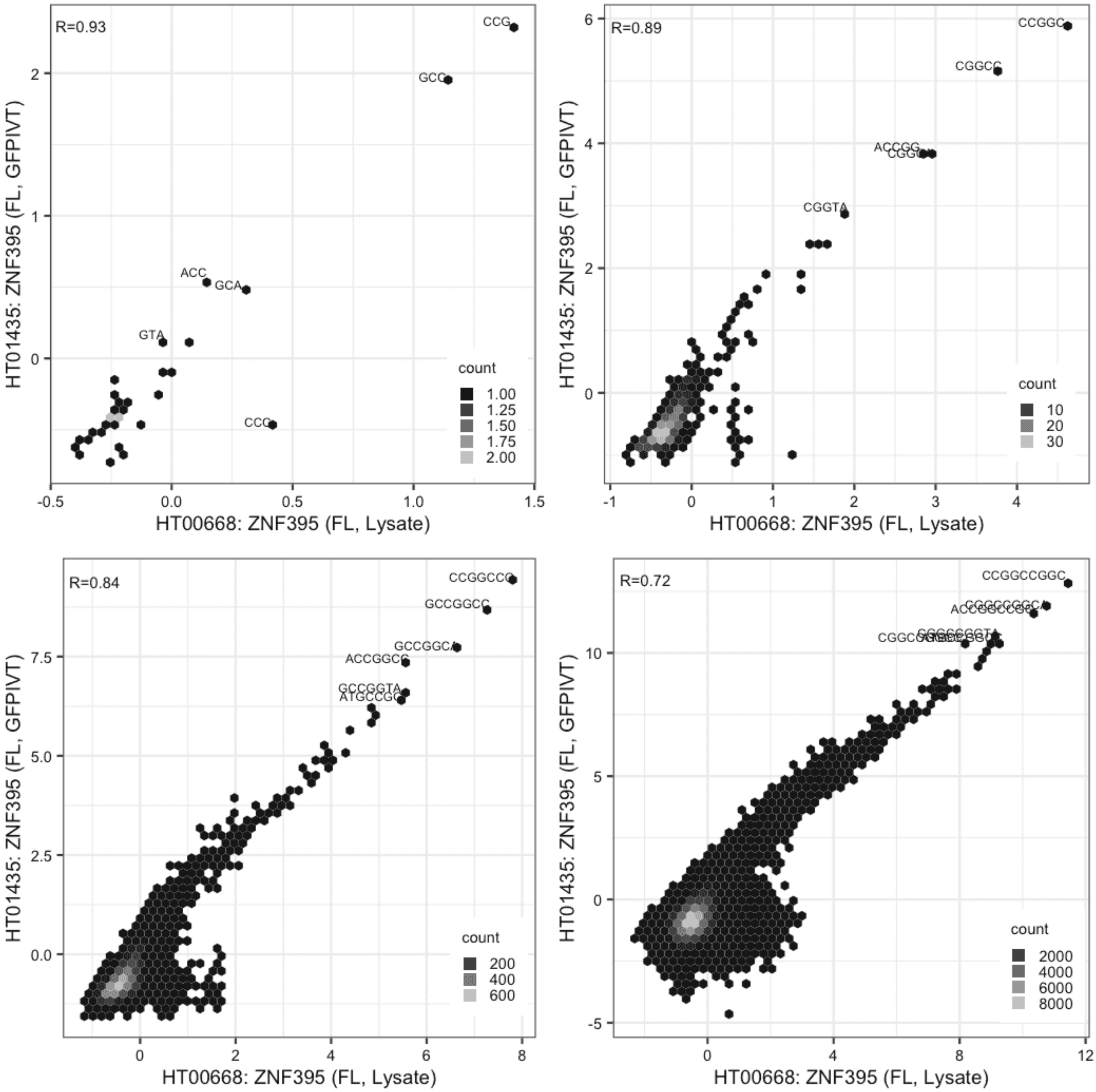

#### ZNF395

Lysate vs GFPIVT, different constructs

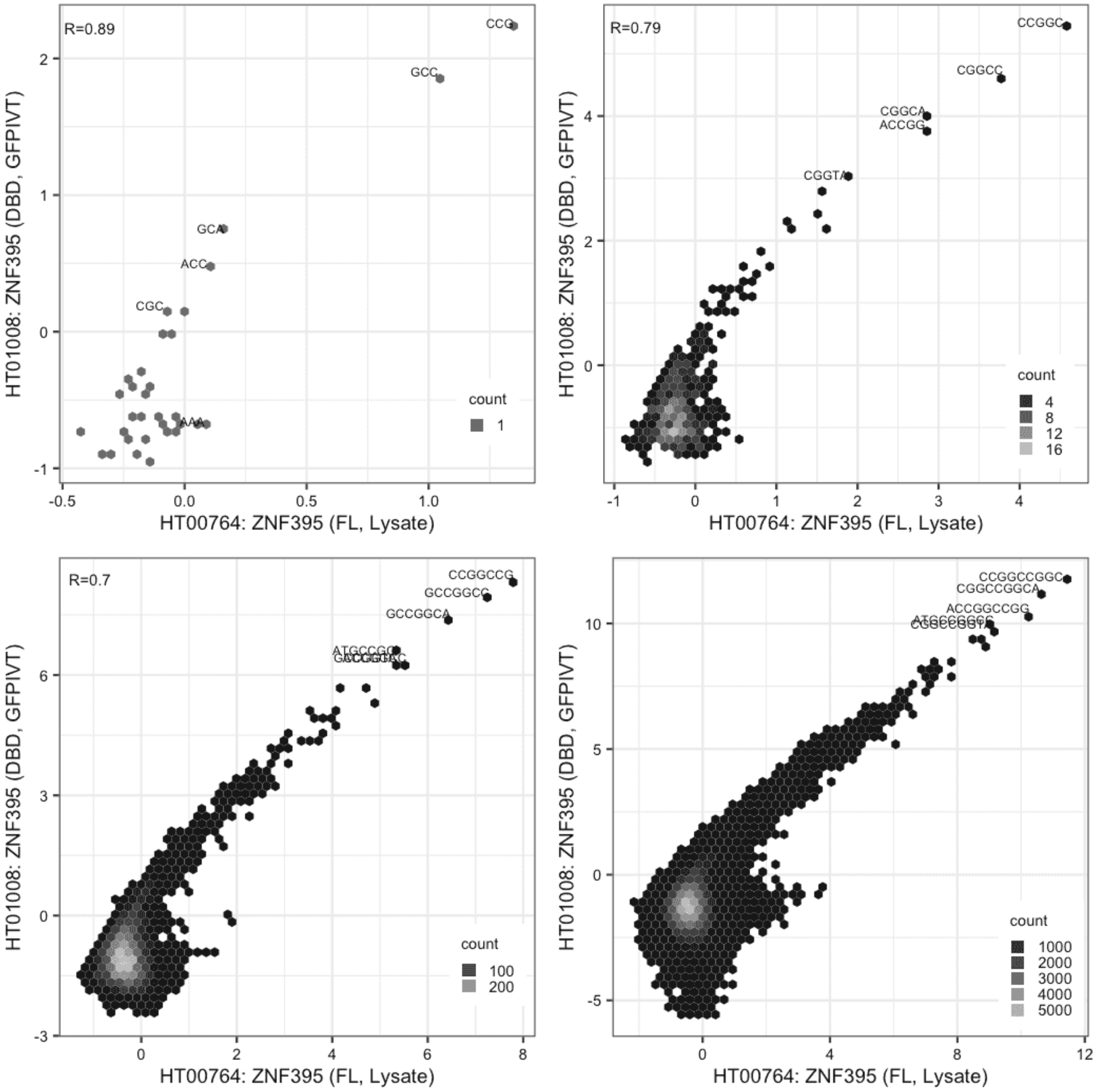

#### ZNF395

Lysate vs GFPIVT, same construct

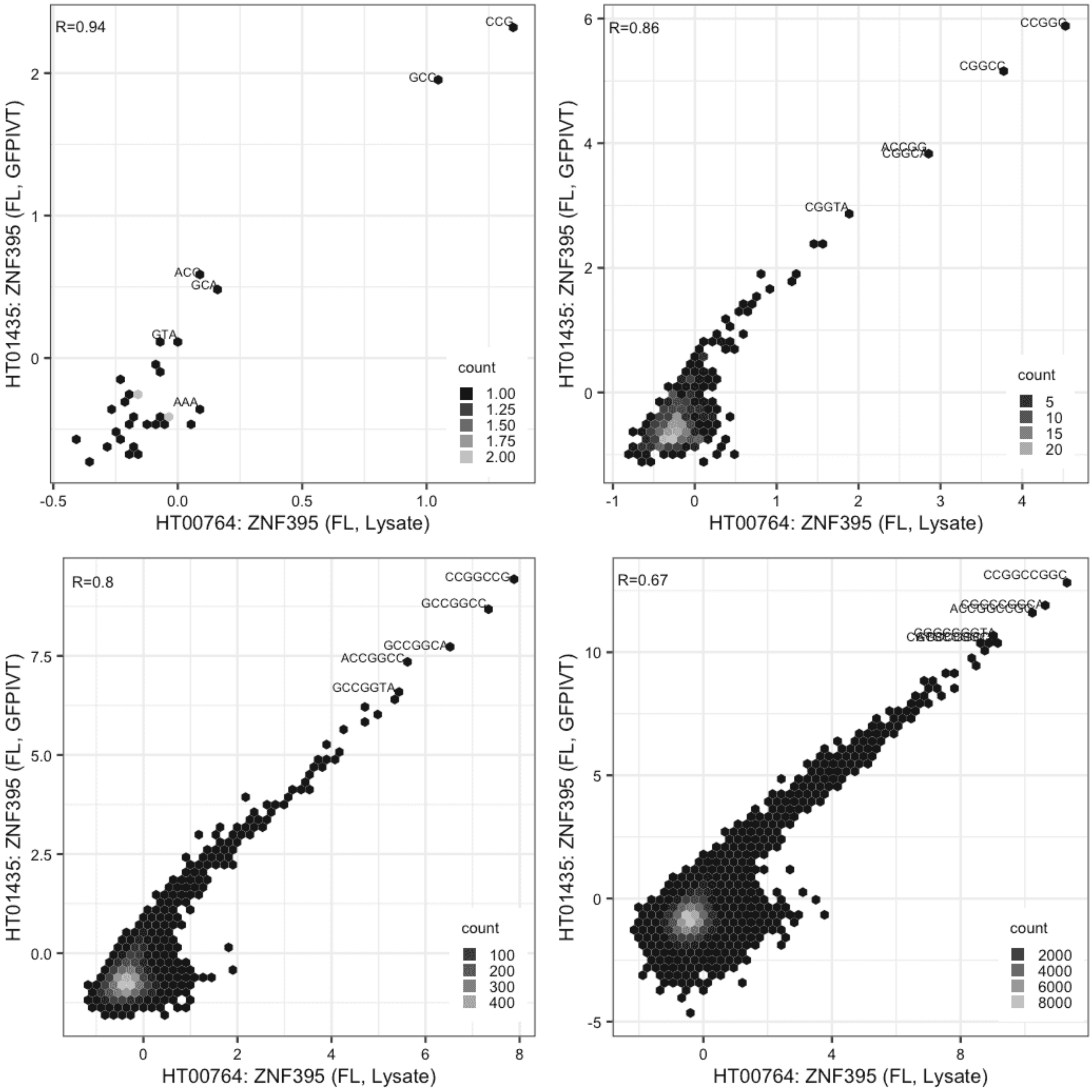

**ZNF395**

GFPIVT vs GFPIVT, different constructs

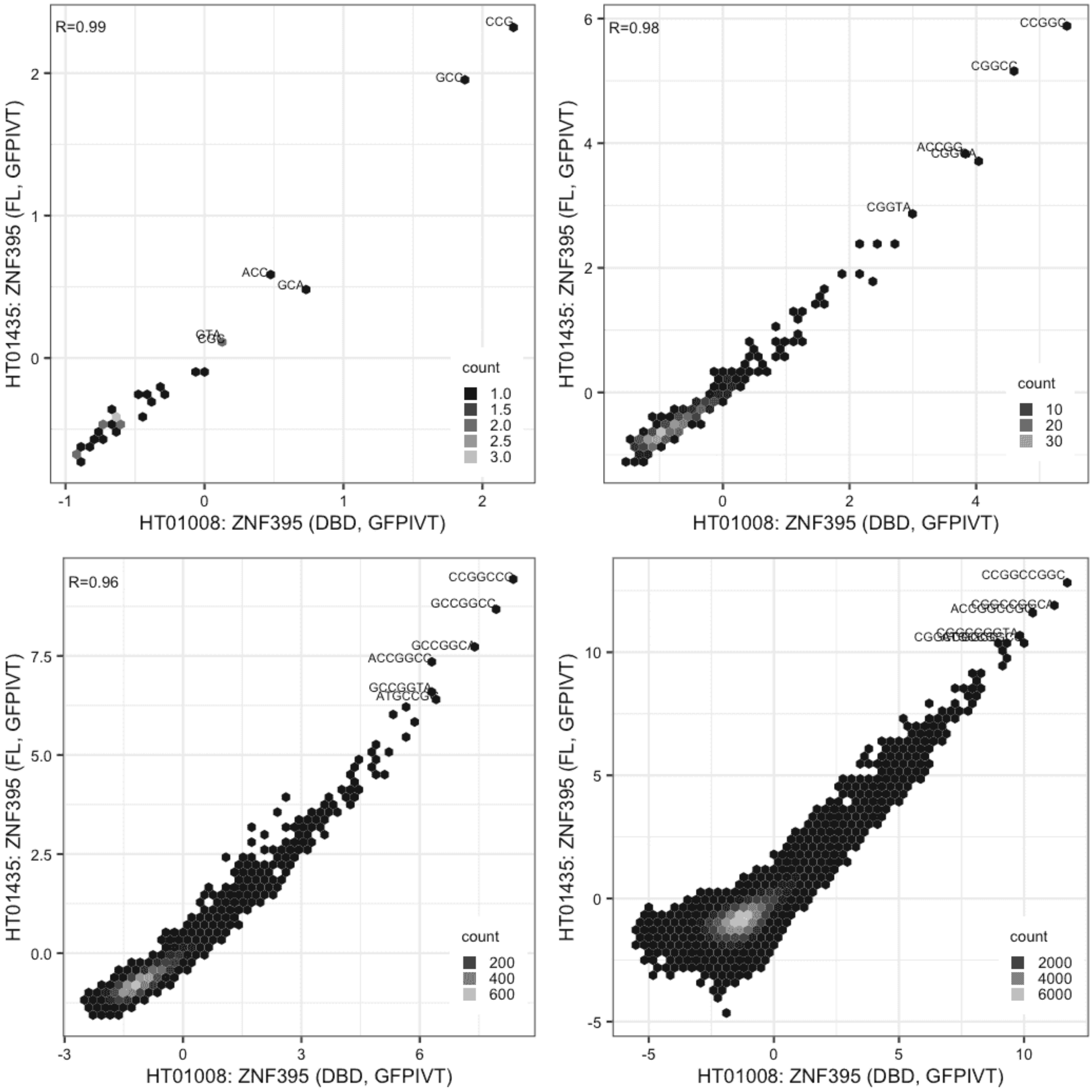

#### ZNF407

Lysate vs IVT, different constructs

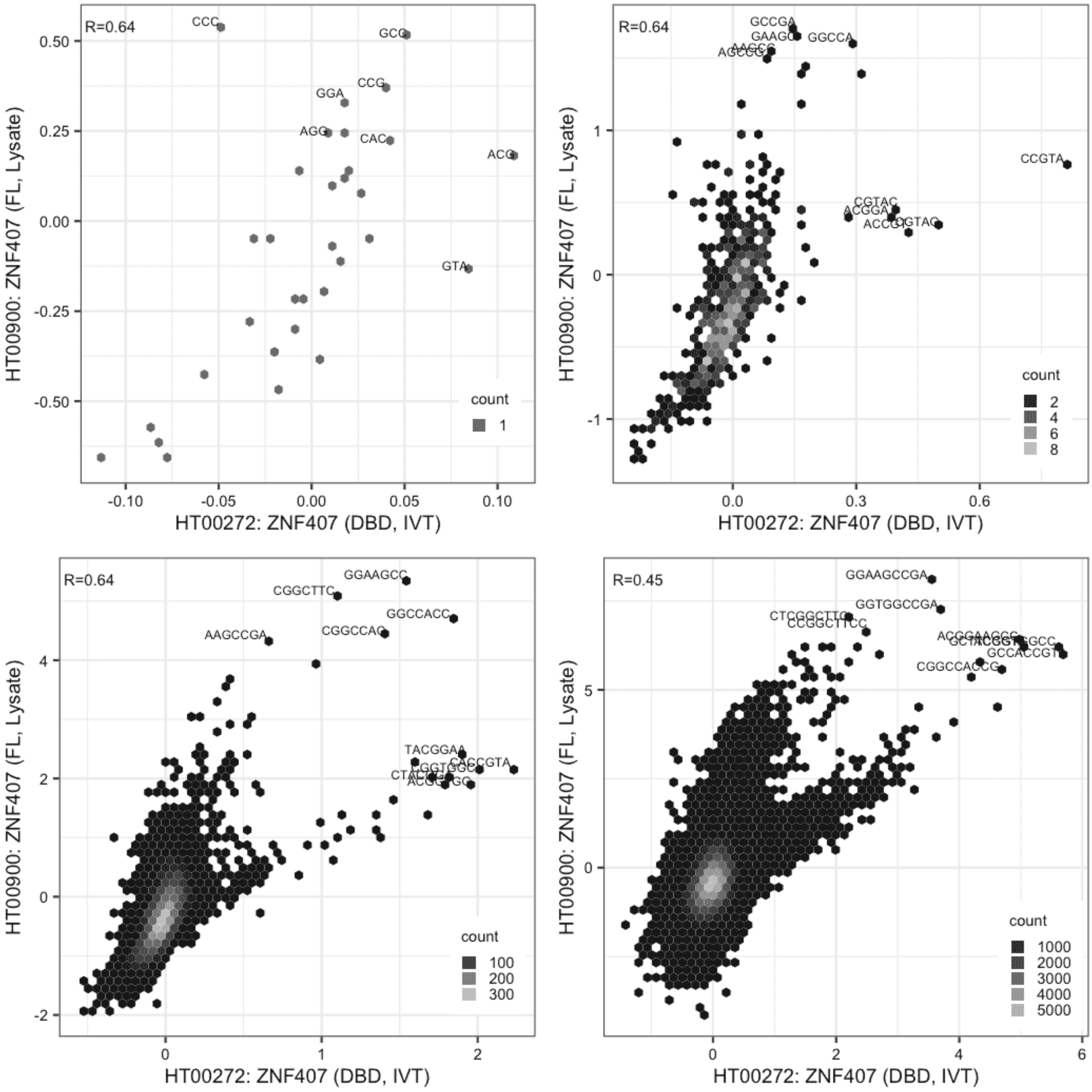

**ZNF407**

IVT vs GFPIVT, same construct

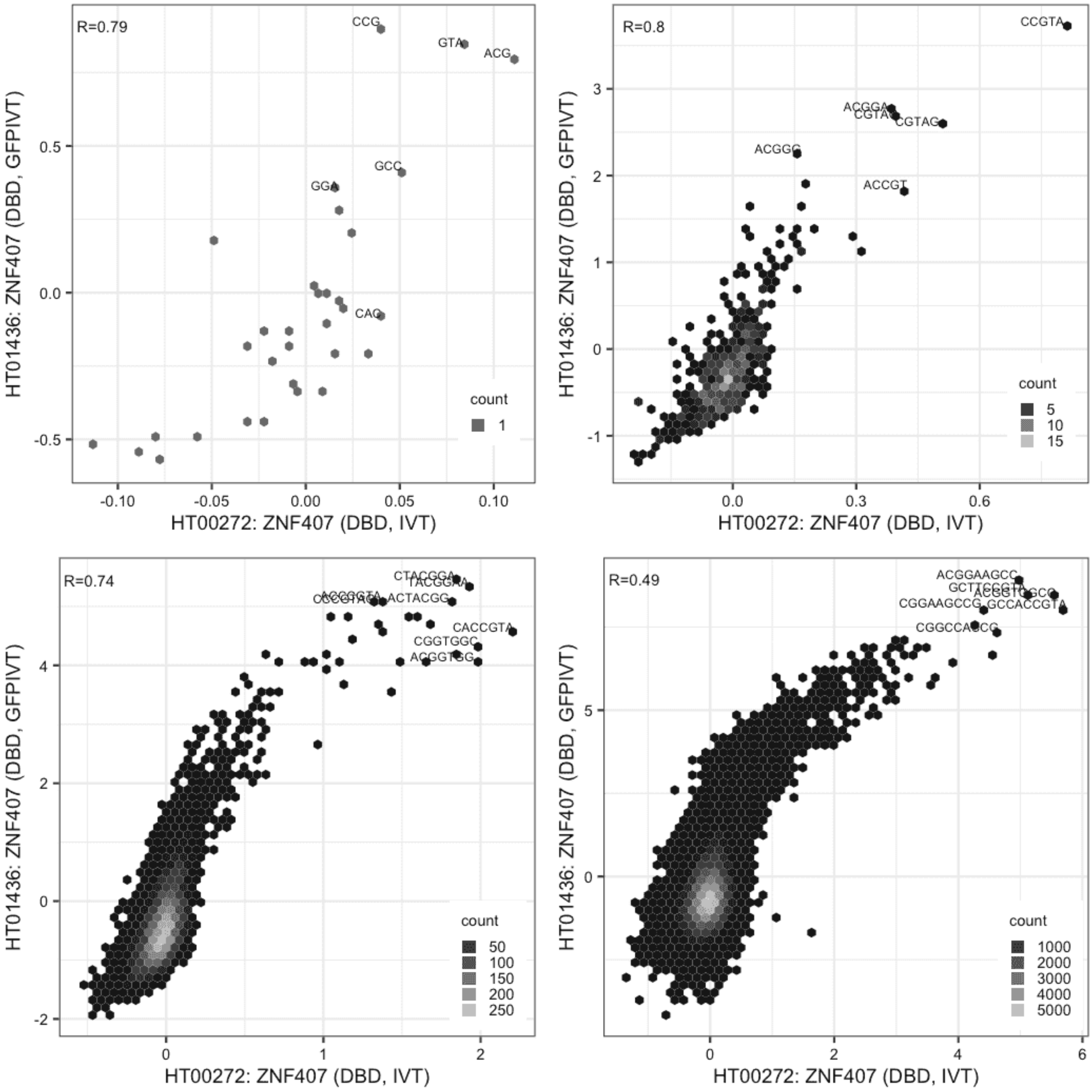

#### ZNF407

Lysate vs GFPIVT, different constructs

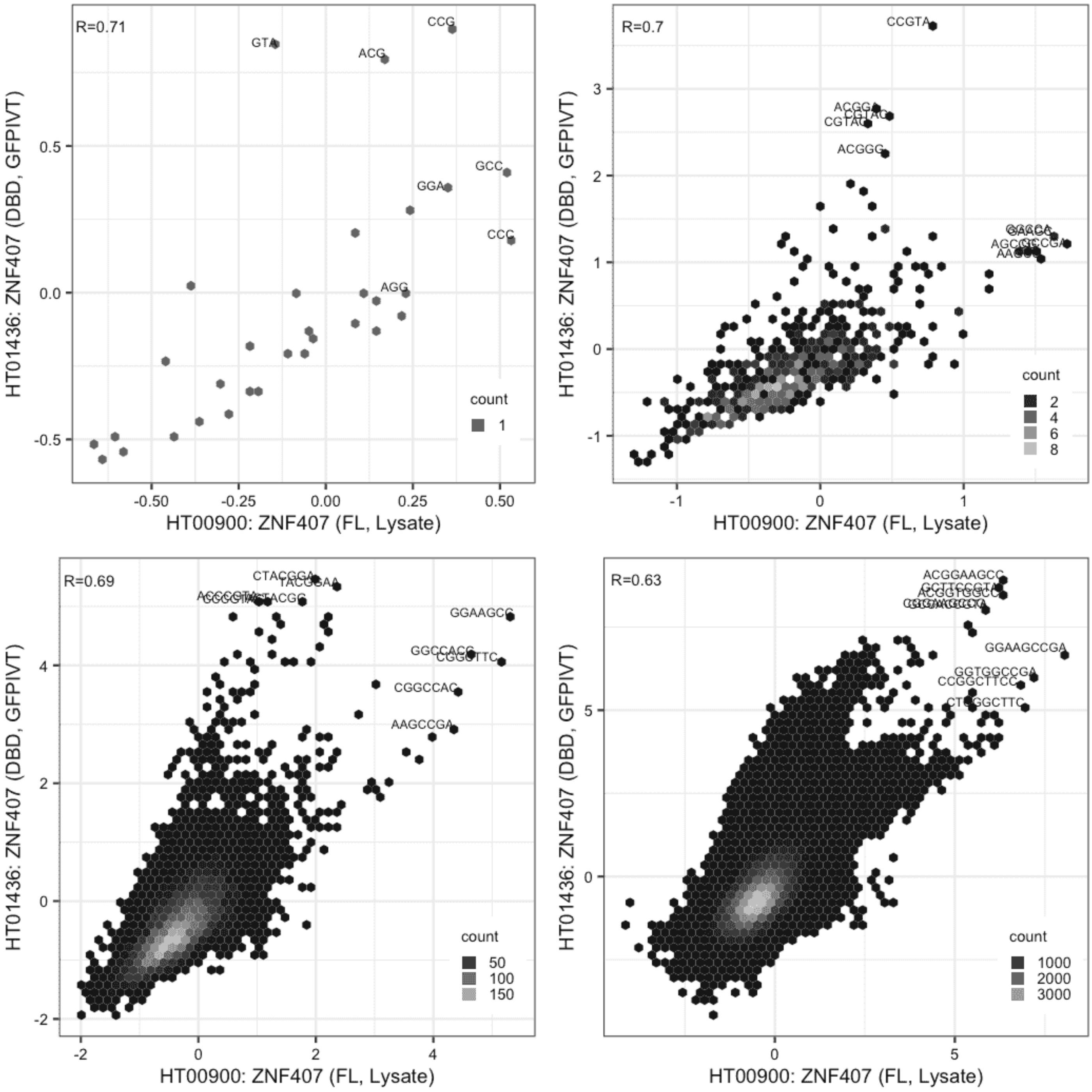

**ZNF48**

Lysate vs Lysate, same construct

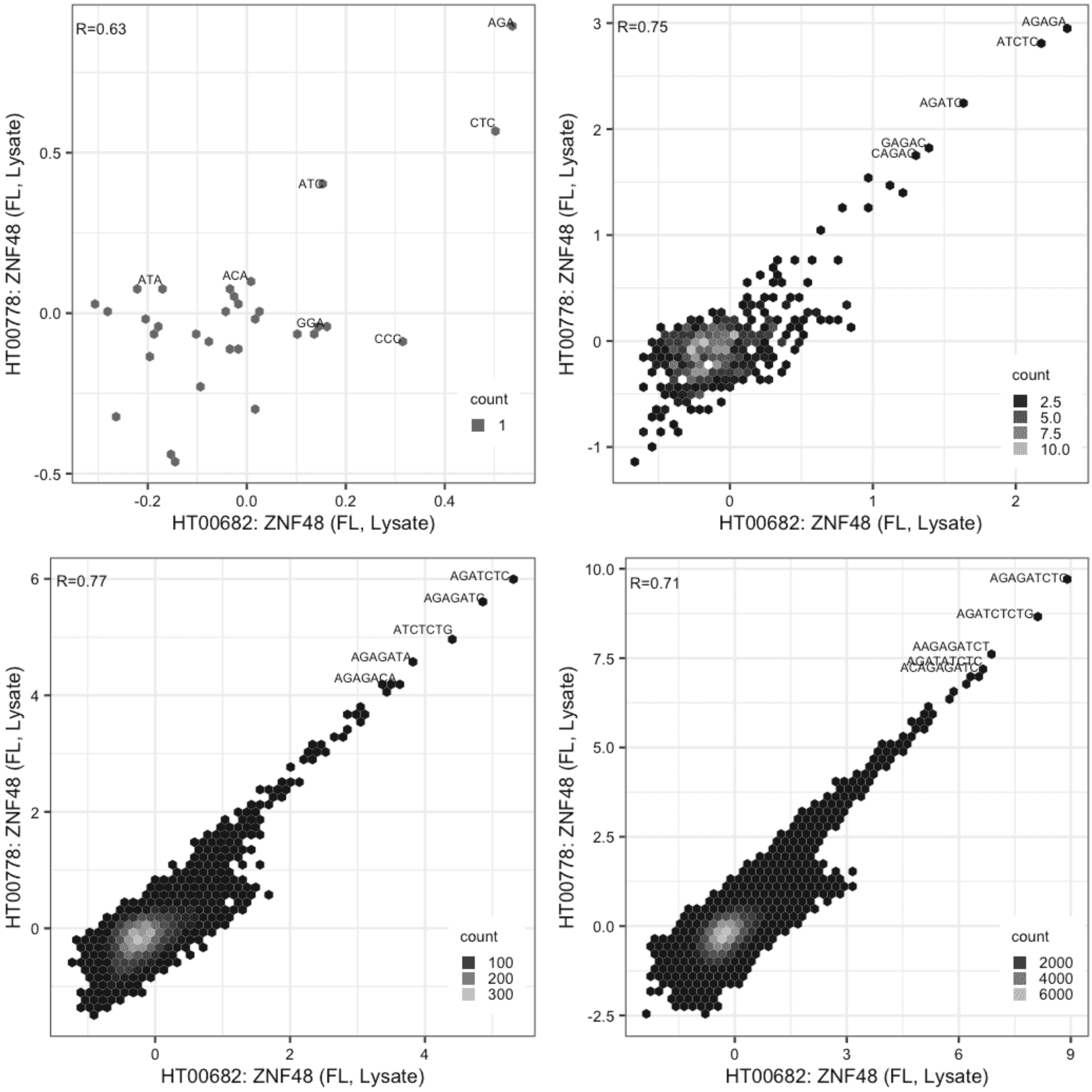

**ZNF48**

Lysate vs GFPIVT, same construct

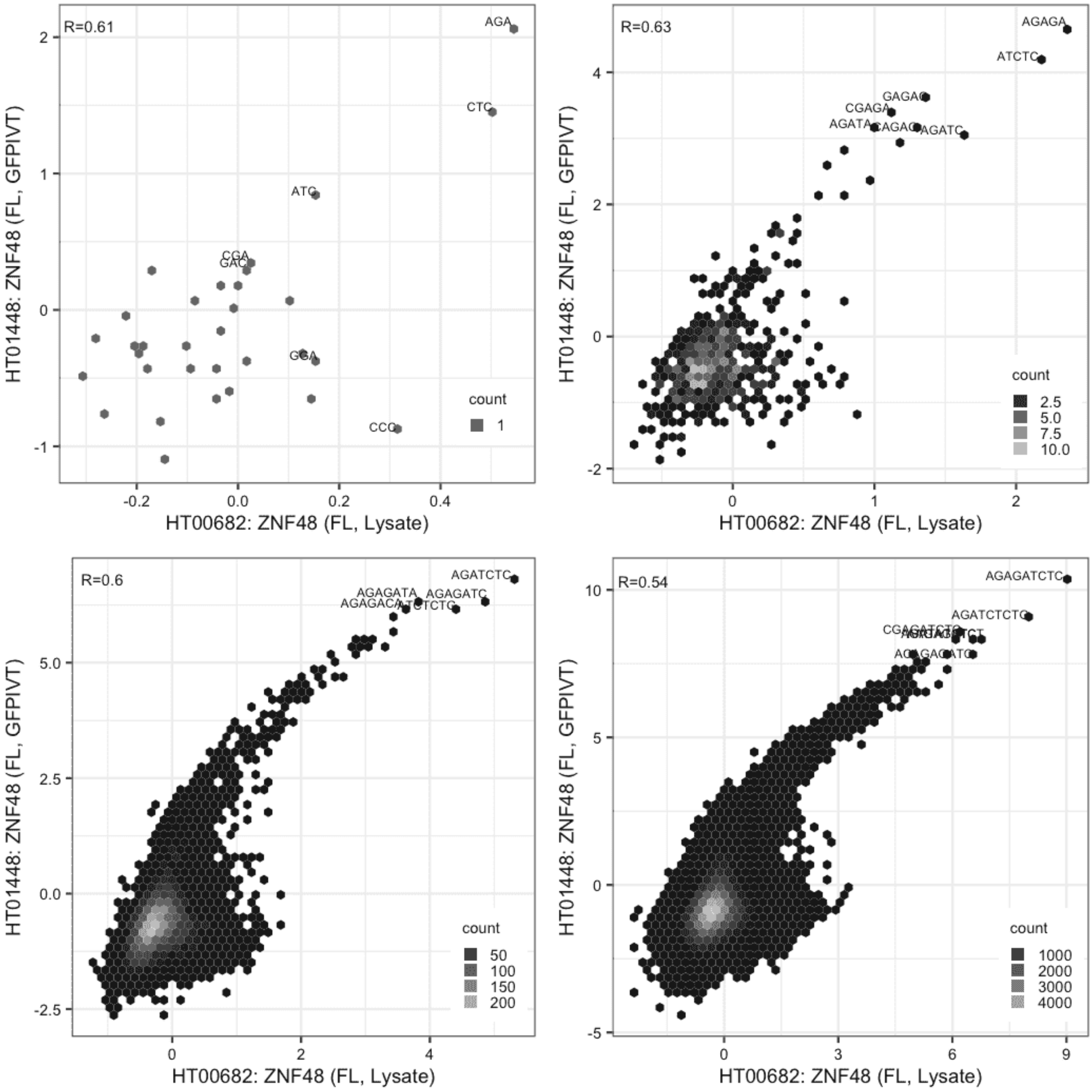

### ZNF48

Lysate vs GFPIVT, same construct

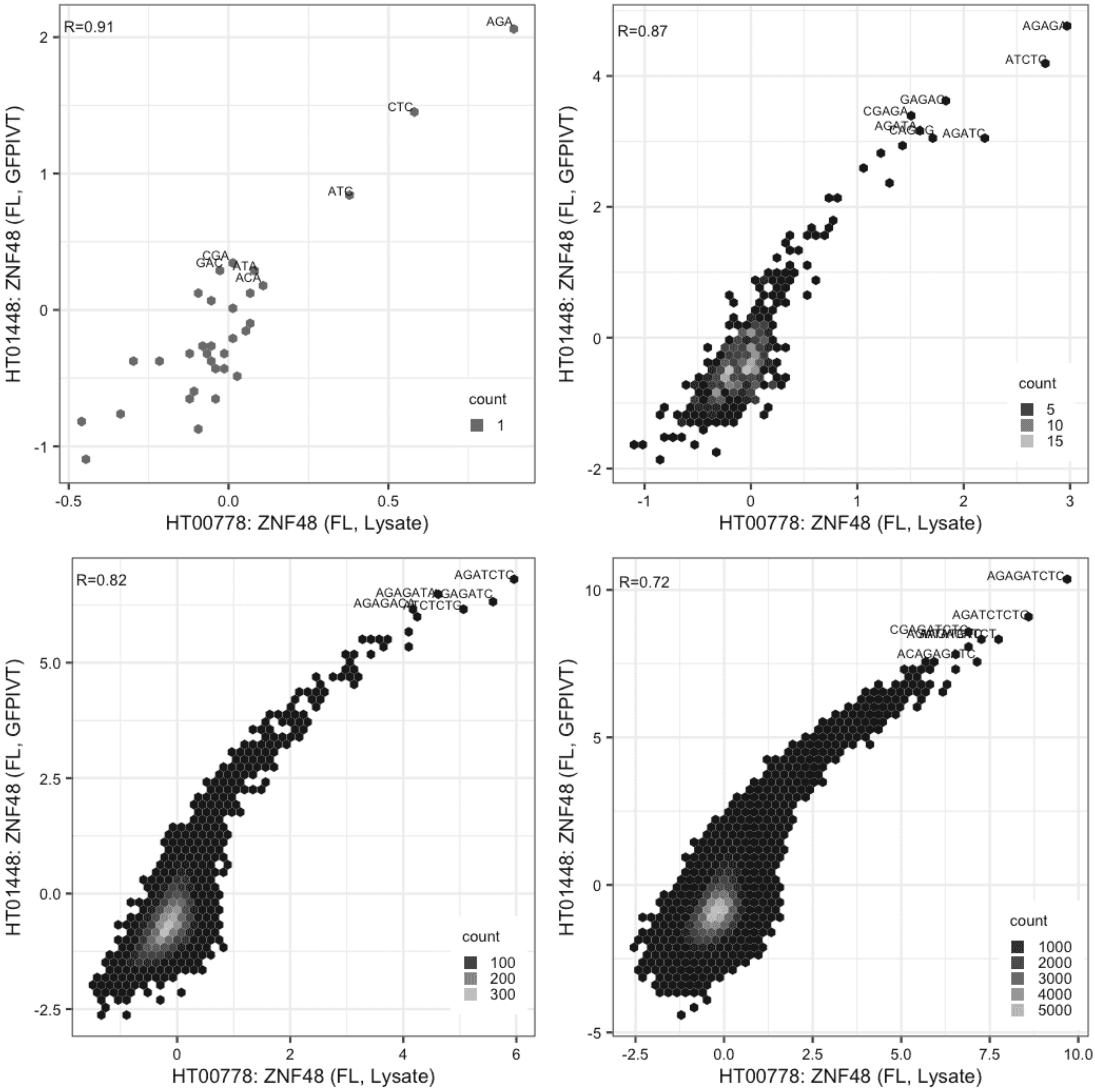

**ZNF518B**

Lysate vs Lysate, same construct

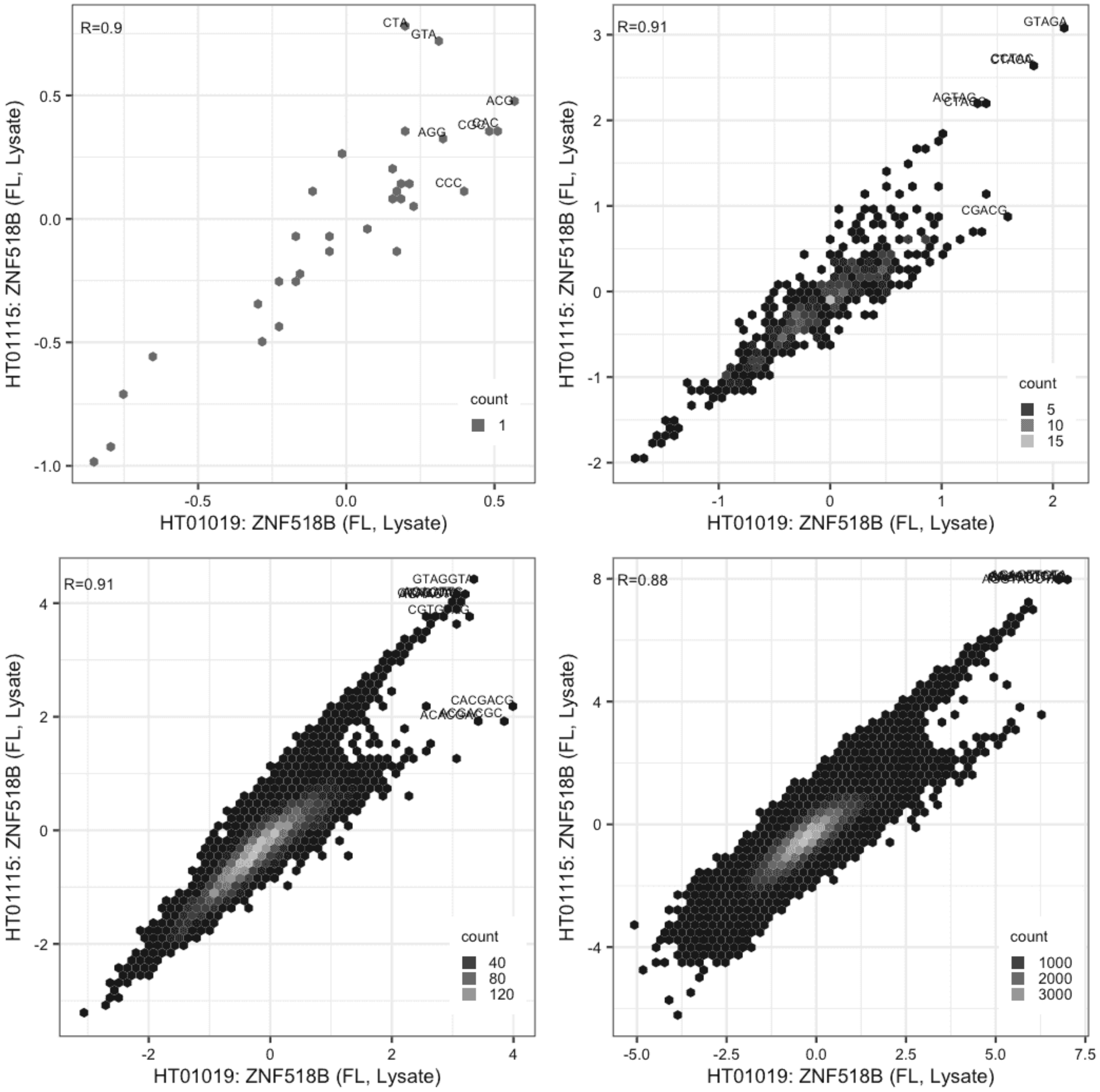

**ZNF518B**

Lysate vs GFPIVT, different constructs

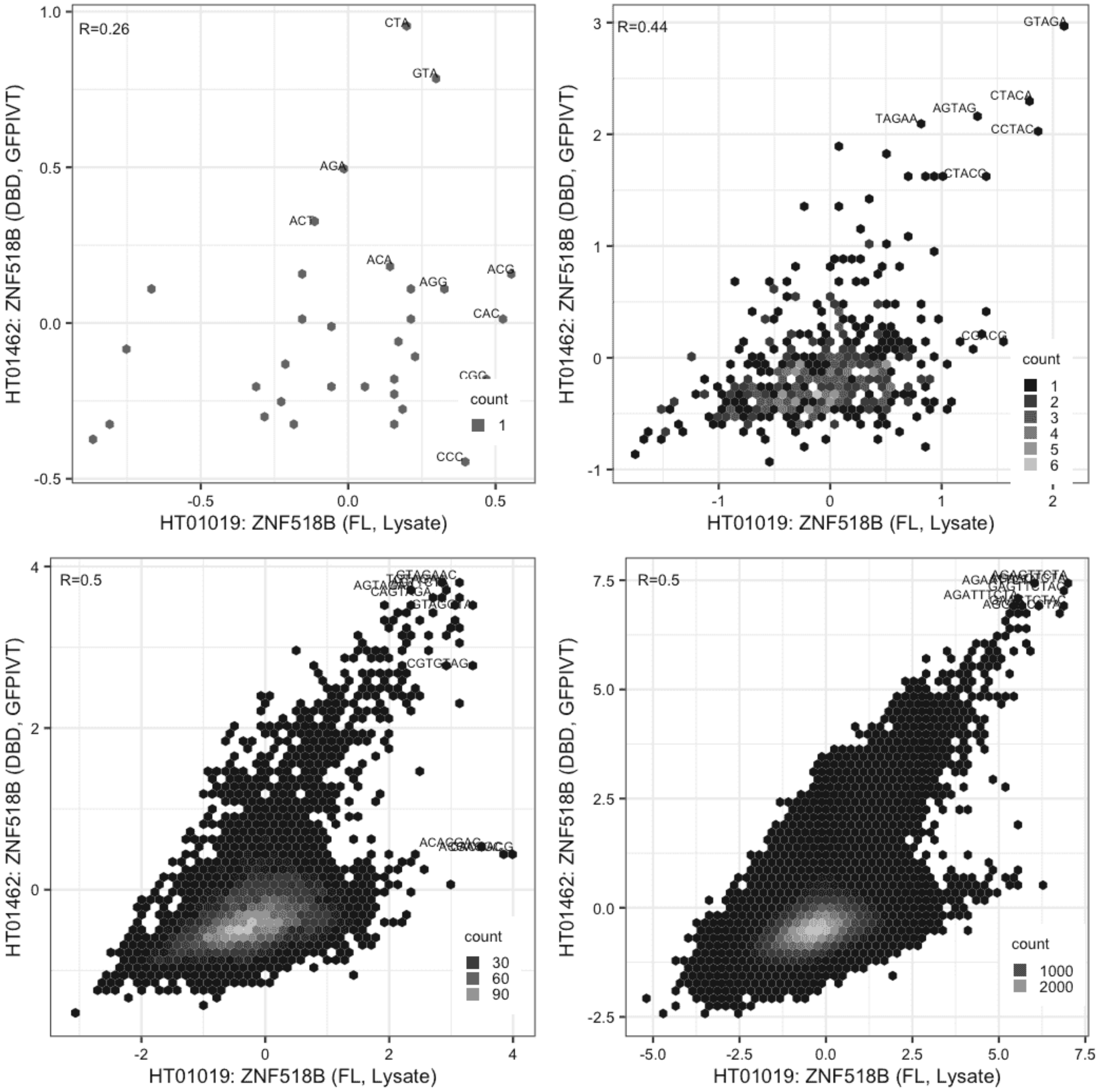

**ZNF518B**

Lysate vs GFPIVT, different constructs

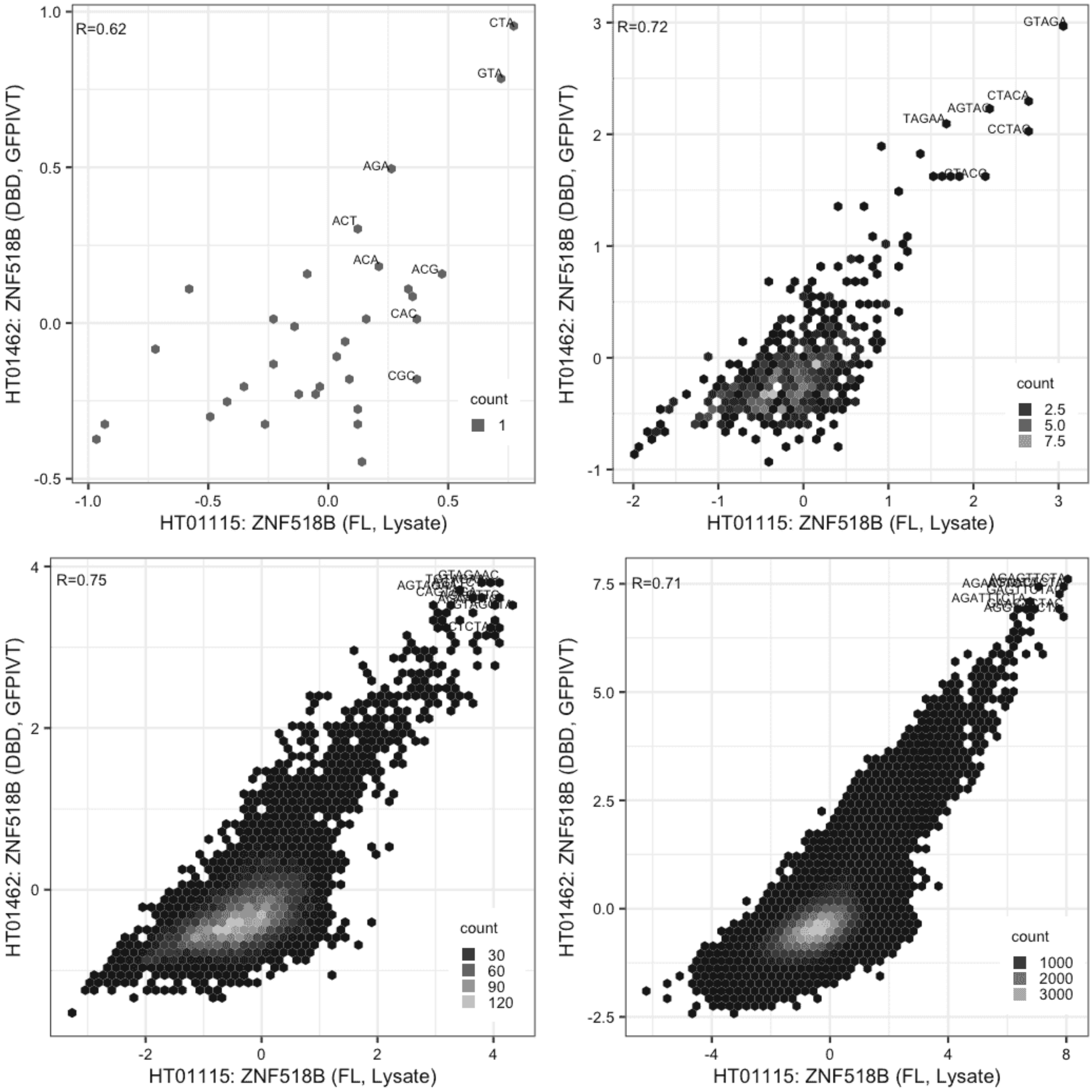

#### ZNF800

Lysate vs GFPIVT, different constructs

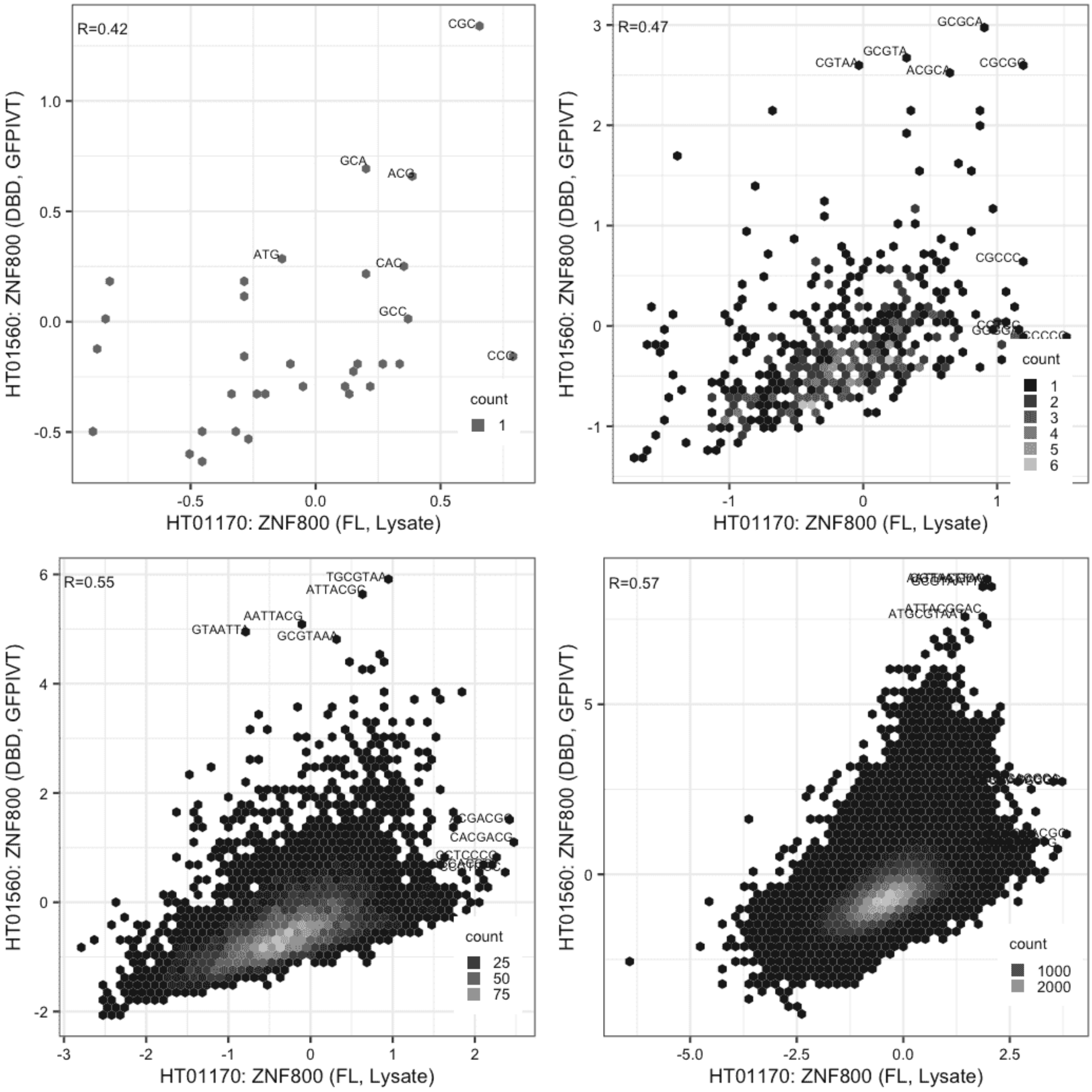

**ZNF800**

Lysate vs GFPIVT, same construct

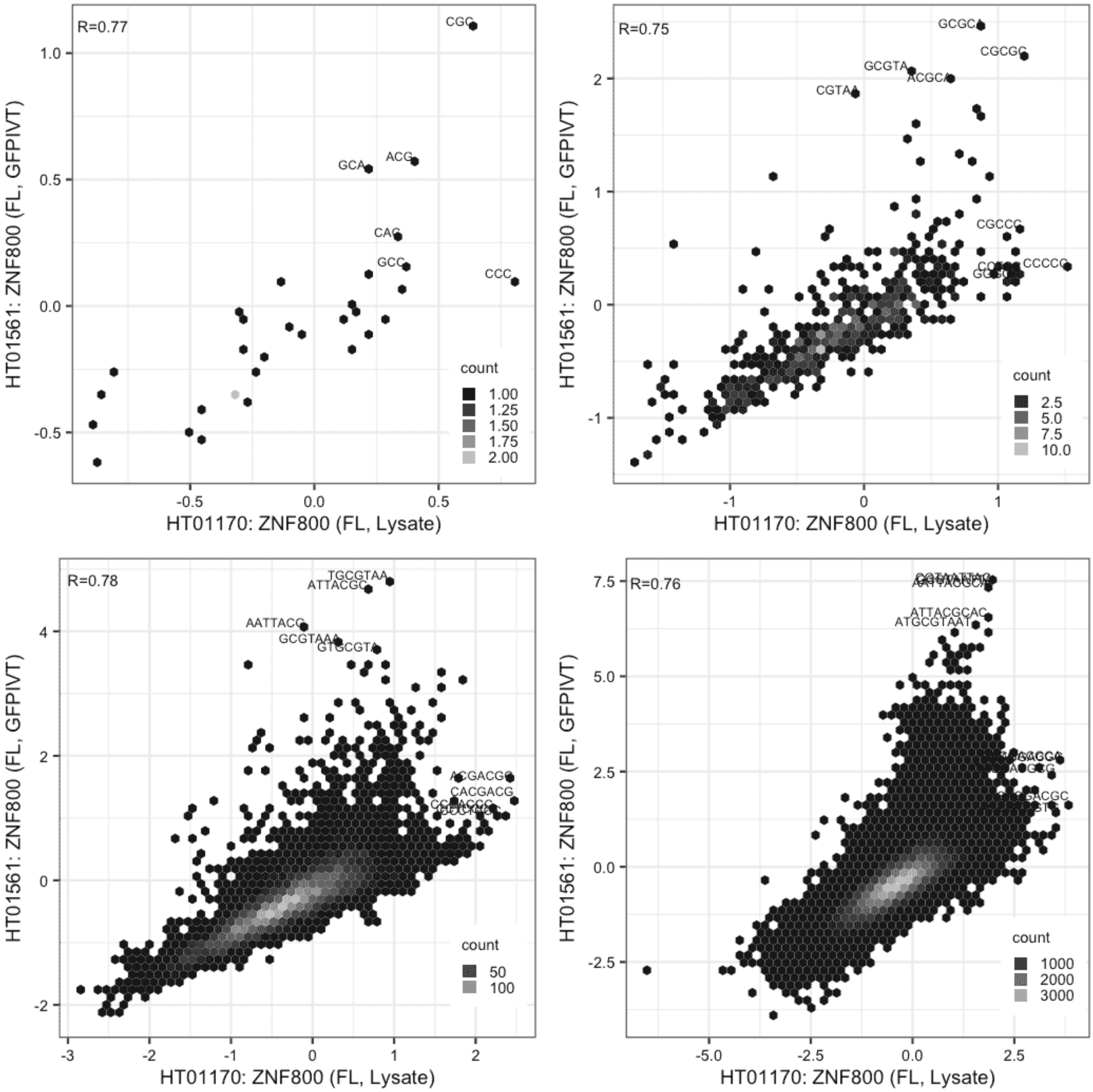

#### ZNF800

GFPIVT vs GFPIVT, different constructs

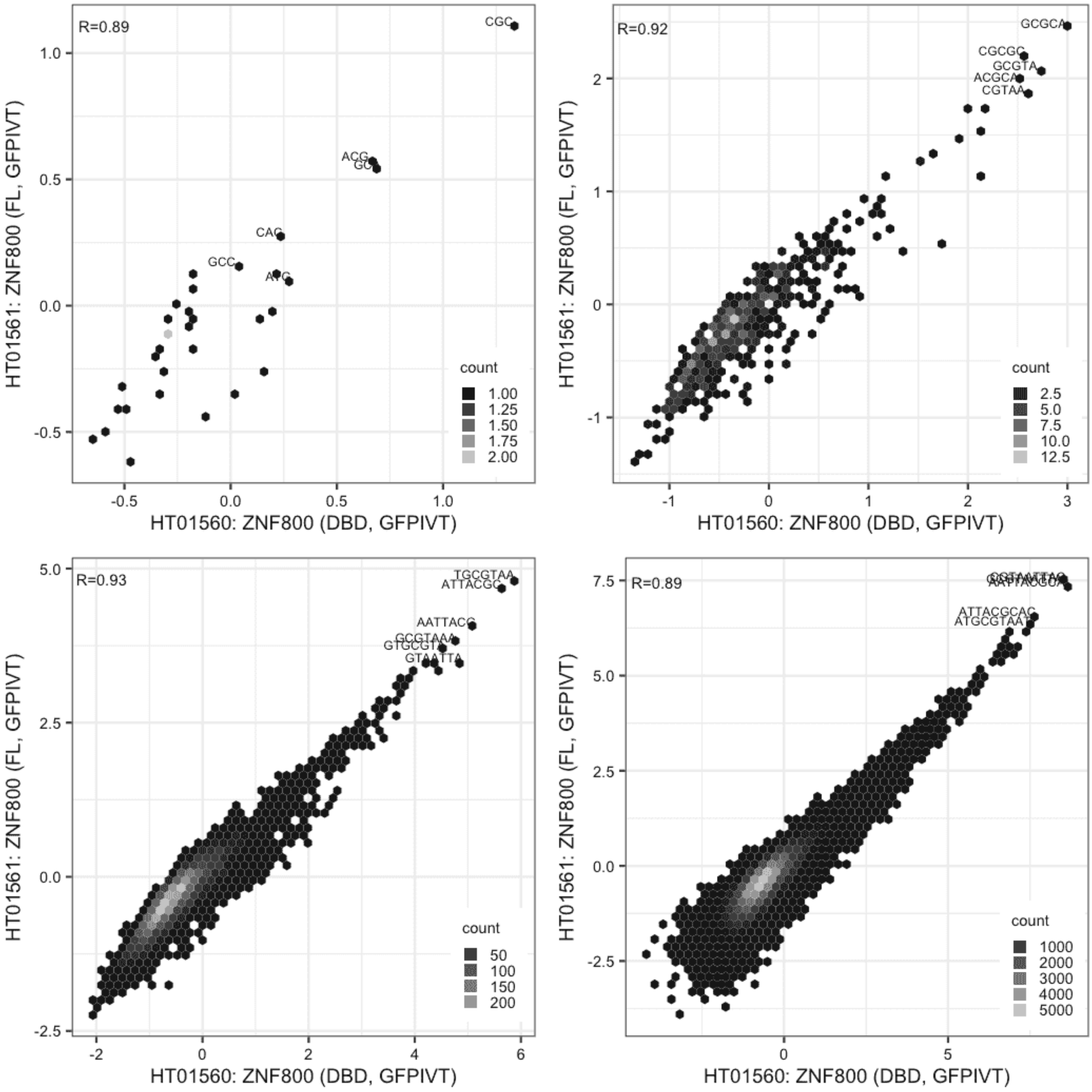

**ZSCAN25**

Lysate vs IVT, different constructs

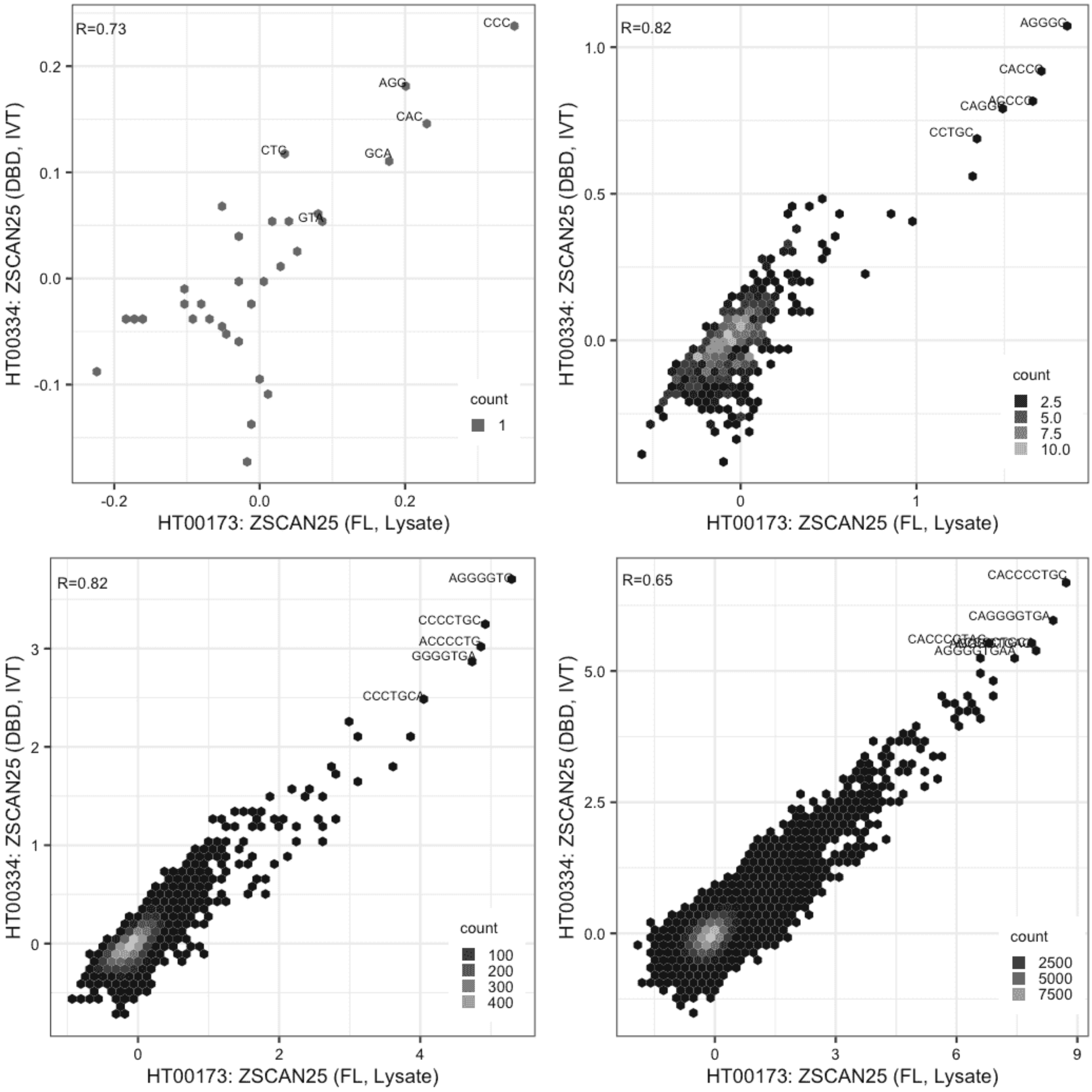
