## Supplementary Document S4 for "GHT-SELEX demonstrates unexpectedly high intrinsic sequence specificity and complex DNA binding of many human transcription factors"

#### Considerations of effects of partial protein products on modular C2H2 binding patterns

Protein production methods can generate partial proteins in some cases, and these can potentially impact modular binding results observed in GHT-SELEX. However, this should have very limited impact. For proteins with a single DNA-binding domain, a partial domain would likely be misfolded and/or not functional for DNA-binding, and thus inert in DNA-binding assays. For C2H2-zf proteins, truncations should lead to a bias towards binding sites that engage the C2H2-zf domains towards the N-terminus. Such a bias would lead to greater signal towards the left side in RCADEEM plots.
We assessed RCADEEM outputs for this type of an effect, and only three TFs, ZNF233, ZNF497 and ZNF853 displayed this type of modularity, However, none of these TFs, nor any of the modular binding TFs displayed in **Fig 6** showed relevant amounts of partial fragments in Western blots.

**Following pages**
**(***Second page, Panel A)* Summary of blot results. (*Second page, Panel B*) Modular C2H2-zfs shown in Fig6; (*Second page, Panel C*) C2H2-zfs ZNF233, ZNF497 and ZNF853, show sites in RCADEEM analysis that could have been affected by C-terminal truncations. Note, only ZNF853 show smaller than expected size fragments. (*Pages 2-6*) All western blot analyses: Wells have been annotated to display HNGC symbol of the TF, its expression construct along with gel and well references for all positions with the same TF; Whether the expression was deemed successful (black text) or unsuccessful (red text); If this protein expression yielded a successful experiment in GHT-SELEX (green bar) or not (red bar); If the TF is a C2H2 protein (black bar on top); If the C2H2 TF displayed modular binding specificity (blue bar) and the expected size of the fusion protein (asterisk over the blot).

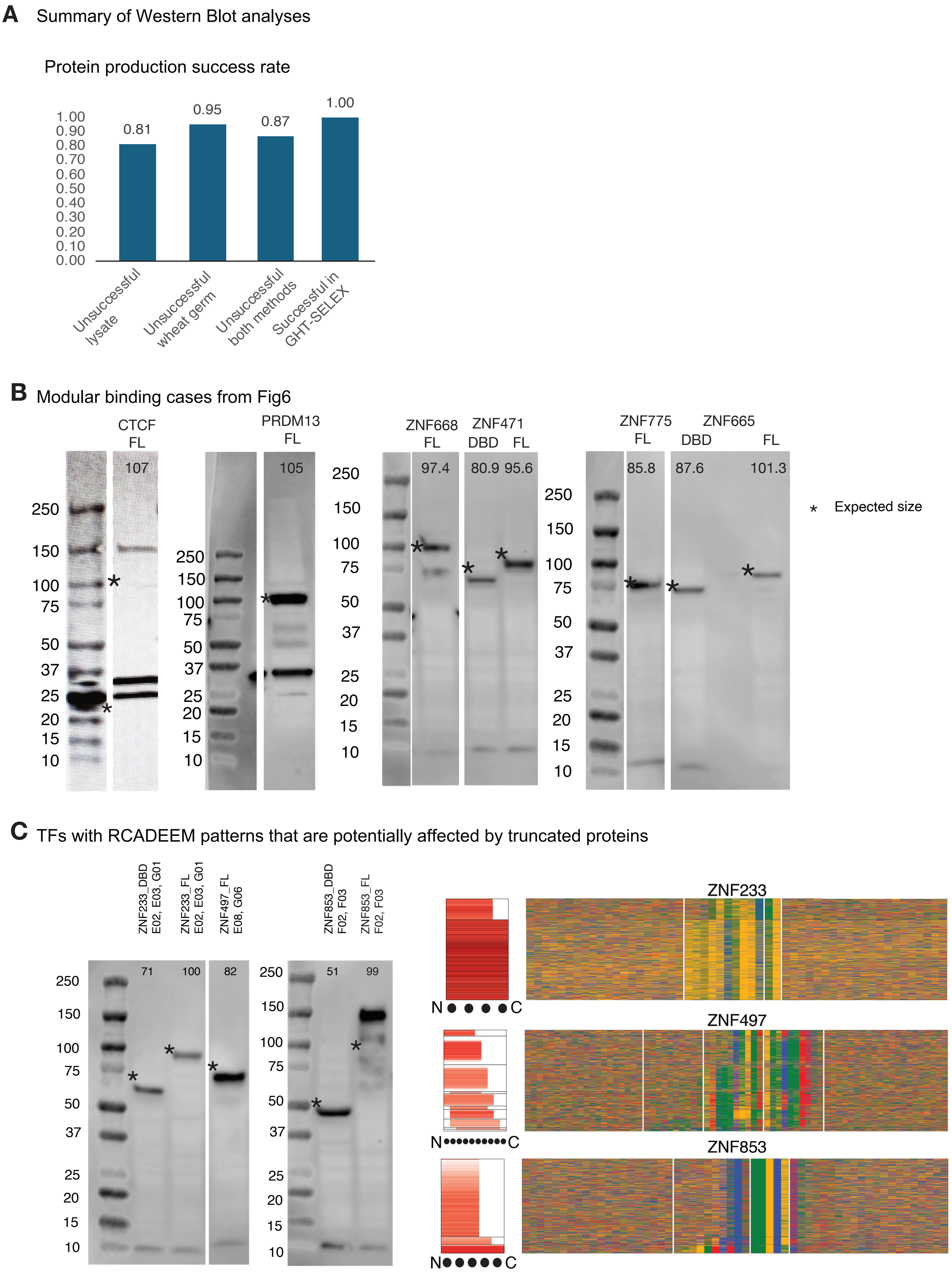
A Summary of Western Blot analyses
Protein production success rate
1.00
0.90
0.80
0.70
0.60
0.50
0.40
0.30
0.20
0.10
0.00
0.81
0.95
0.87
1.00
Unsuccessful lysate
Unsuccessful wheat germ
Unsuccessful both methods
Successful in GHT-SELEX
B Modular binding cases from Fig6
CTCF FL
107
PRDM13 FL
105
ZNF668 FL
97.4
ZNF471 DBD FL
80.9 95.6
ZNF775 FL
85.8
ZNF665 DBD FL
87.6
101.3
250
150
100
75
50
37
25
20
15
10
* Expected size
C TFs with RCADEEM patterns that are potentially affected by truncated proteins
ZNF233_DBD E02, E03, G01
ZNF233_FL E02, E03, G01
ZNF497_FL E08, G06
ZNF853_DBD F02, F03
ZNF853_FL F02, F03
71
100
82
51
99
ZNF233
ZNF497
ZNF853
N
C

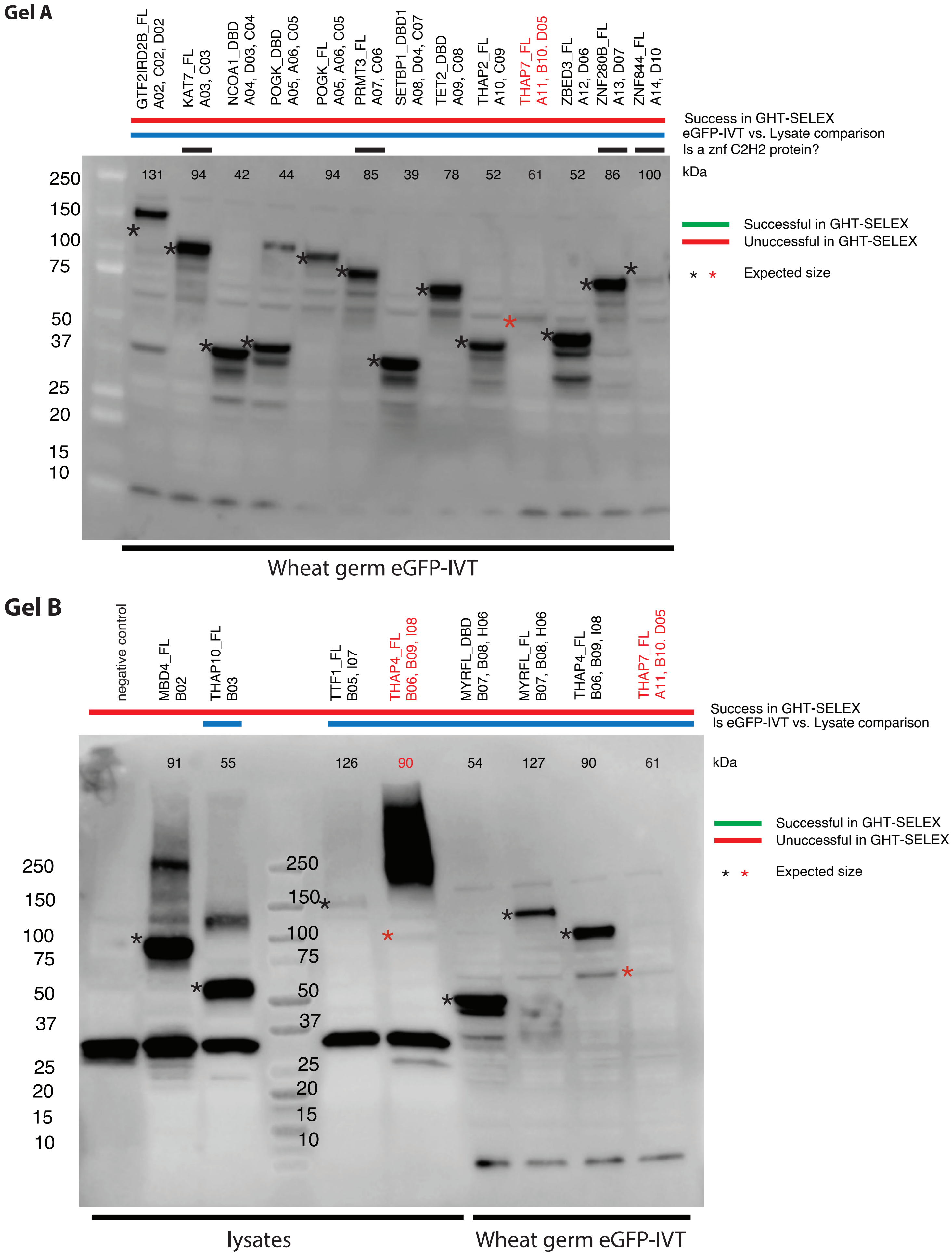
Gel A
GTF2IRD2B_FL A02, C02, D02
KAT7_FL A03, C03
NCOA1_DBD A04, D03, C04
POGK_DBD A05, A06, C05
POGK_FL A05, A06, C05
PRMT3_FL A07, C06
SETBP1_DBD1 A08, D04, C07
TET2_DBD A09, C08
THAP2_FL A10, C09
THAP7_FL A11, B10. D05
ZBED3_FL A12, D06
ZNF280B_FL A13, D07
ZNF844_FL A14, D10
Success in GHT-SELEX
eGFP-IVT vs. Lysate comparison
Is a znf C2H2 protein?
131 94 42 44 94 85 39 78 52 61 52 86 100
kDa
250
150
100
75
50
37
25
20
15
10
Successful in GHT-SELEX
Unuccessful in GHT-SELEX
Expected size
Wheat germ eGFP-IVT
Gel B
negative control
MBD4_FL B02
THAP10_FL B03
TTF1_FL B05, I07
THAP4_FL B06, B09, I08
MYRFL_DBD B07, B08, H06
MYRFL_FL B07, B08, H06
THAP4_FL B06, B09, I08
THAP7_FL A11, B10. D05
Success in GHT-SELEX
Is eGFP-IVT vs. Lysate comparison
91 55 126 90 54 127 90 61
kDa
250
150
100
75
50
37
25
20
15
10
Successful in GHT-SELEX
Unuccessful in GHT-SELEX
Expected size
lysates
Wheat germ eGFP-IVT

### Gel C

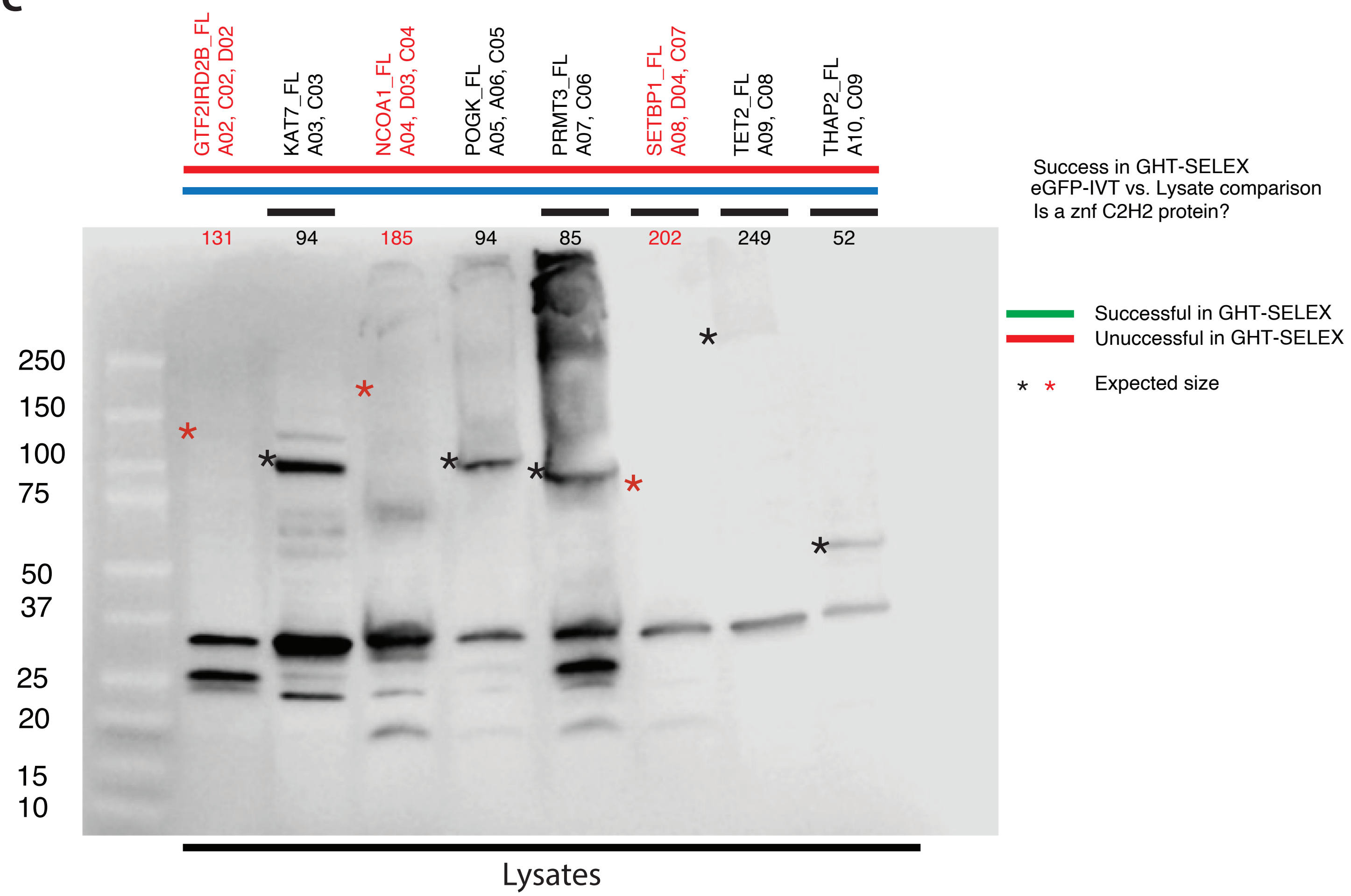
GTF2IRD2B_FL
A02, C02, D02
KAT7_FL
A03, C03
NCOA1_FL
A04, D03, C04
POGK_FL
A05, A06, C05
PRMT3_FL
A07, C06
SETBP1_FL
A08, D04, C07
TET2_FL
A09, C08
THAP2_FL
A10, C09
Success in GHT-SELEX
eGFP-IVT vs. Lysate comparison
Is a znf C2H2 protein?
131
94
185
94
85
202
249
52
Successful in GHT-SELEX
Unuccessful in GHT-SELEX
Expected size
250
150
100
75
50
37
25
20
15
10
Lysates

### Gel D

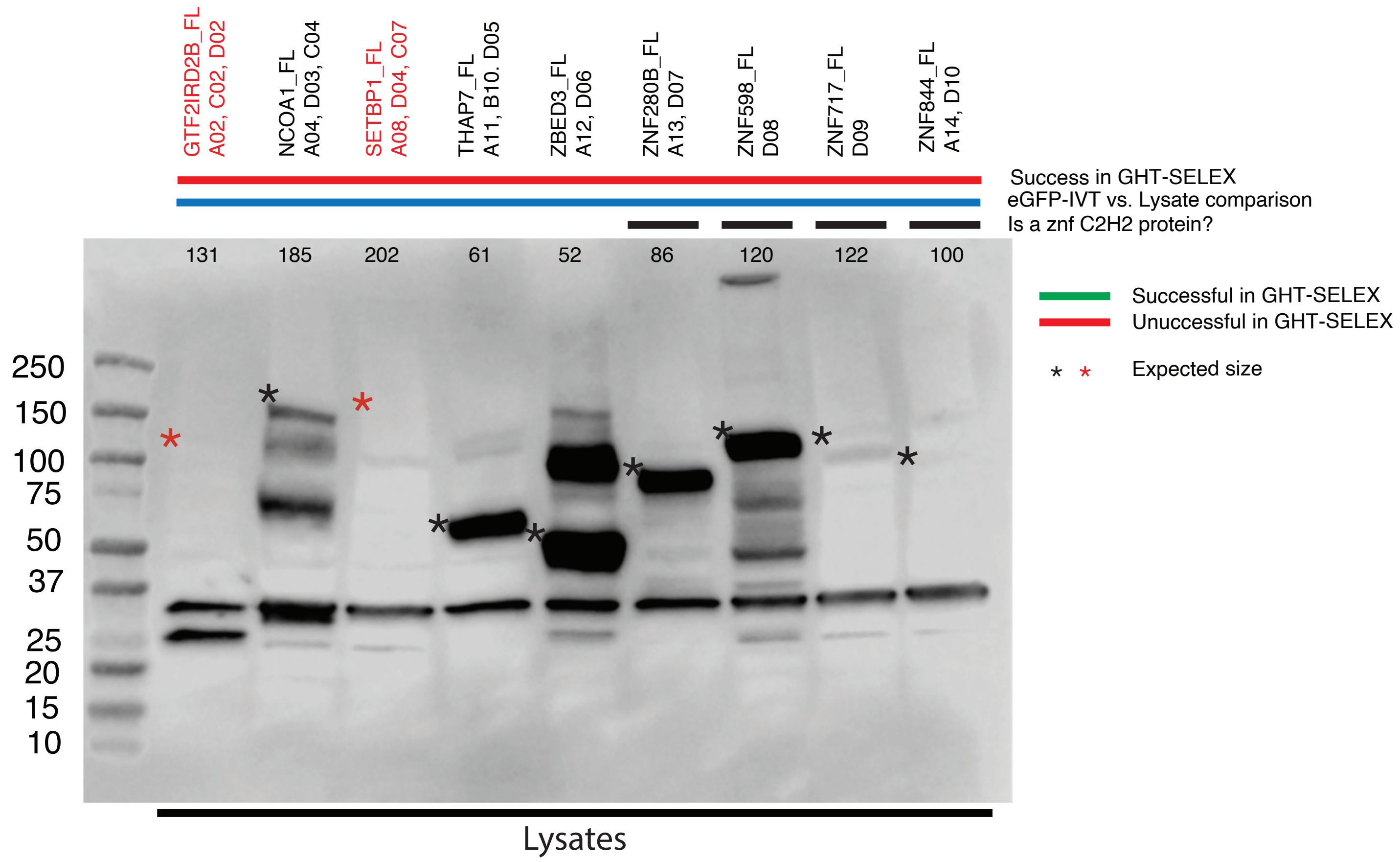
GTF2IRD2B_FL
A02, C02, D02
NCOA1_FL
A04, D03, C04
SETBP1_FL
A08, D04, C07
THAP7_FL
A11, B10. D05
ZBED3_FL
A12, D06
ZNF280B_FL
A13, D07
ZNF598_FL
D08
ZNF717_FL
D09
ZNF844_FL
A14, D10
Success in GHT-SELEX
eGFP-IVT vs. Lysate comparison
Is a znf C2H2 protein?
131
185
202
61
52
86
120
122
100
Successful in GHT-SELEX
Unuccessful in GHT-SELEX
Expected size
250
150
100
75
50
37
25
20
15
10
Lysates

#### Gel E

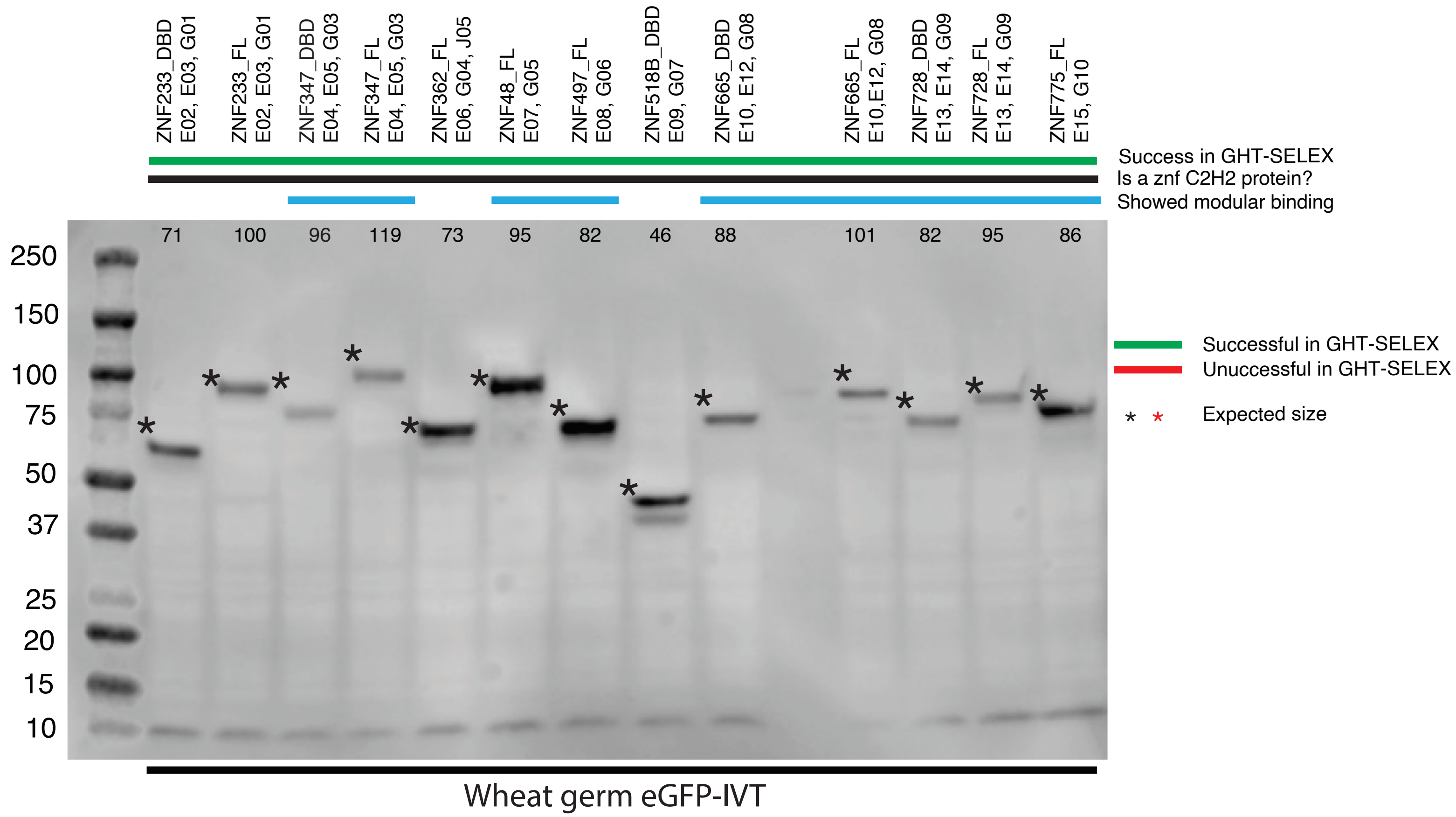
ZNF233_DBD E02, E03, G01
ZNF233_FL E02, E03, G01
ZNF347_DBD E04, E05, G03
ZNF347_FL E04, E05, G03
ZNF362_FL E06, G04, J05
ZNF48_FL E07, G05
ZNF497_FL E08, G06
ZNF518B_DBD E09, G07
ZNF665_DBD E10, E12, G08
ZNF665_FL E10,E12, G08
ZNF728_DBD E13, E14, G09
ZNF728_FL E13, E14, G09
ZNF775_FL E15, G10
Success in GHT-SELEX
Is a znf C2H2 protein?
Showed modular binding
71 100 96 119 73 95 82 46 88 101 82 95 86
250
150
100
75
50
37
25
20
15
10
Successful in GHT-SELEX
Unuccessful in GHT-SELEX
Expected size
Wheat germ eGFP-IVT

#### Gel F

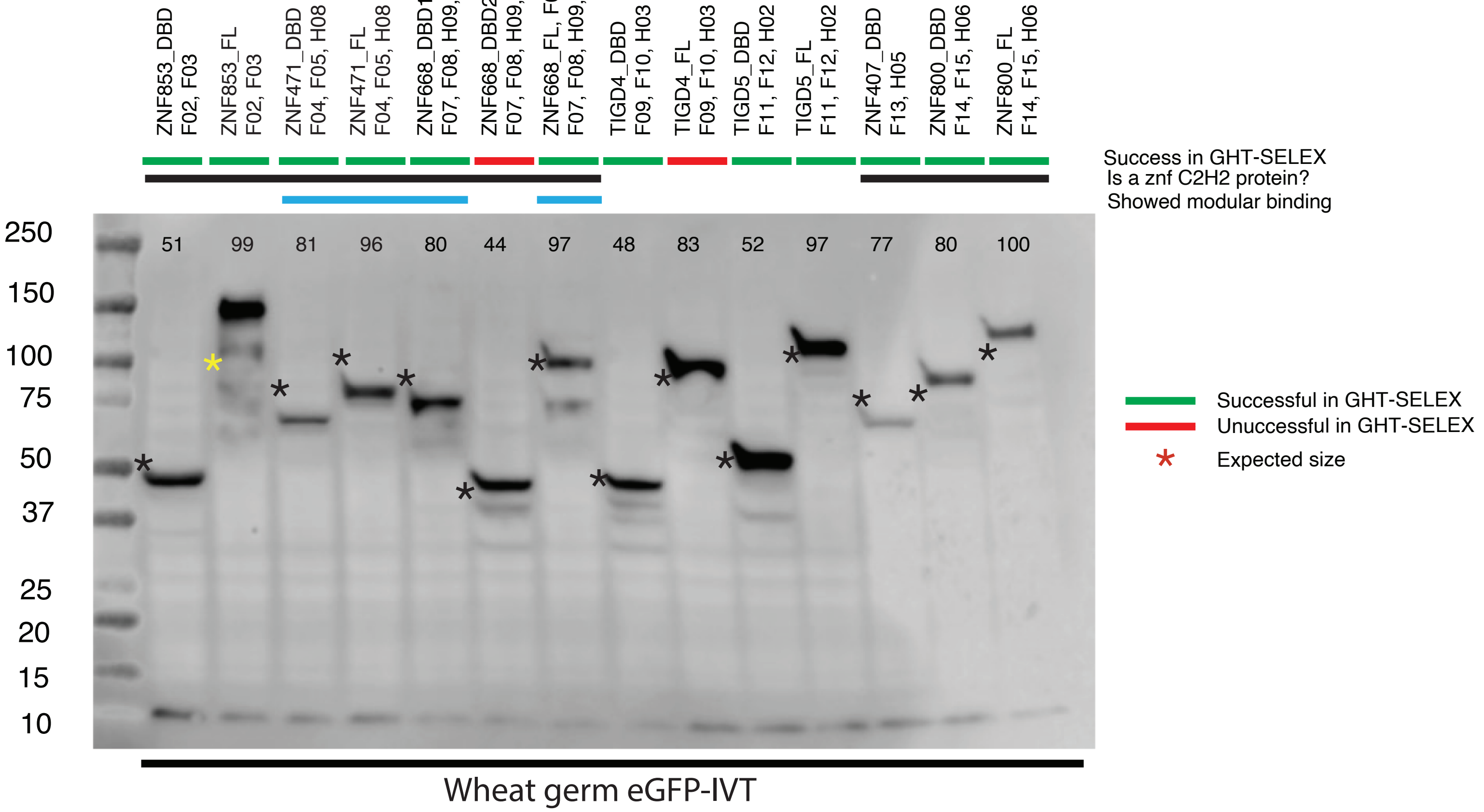
ZNF853_DBD F02, F03
ZNF853_FL F02, F03
ZNF471_DBD F04, F05, H08
ZNF471_FL F04, F05, H08
ZNF668_DBD1, F06, F07, F08, H09, I06
ZNF668_DBD2, F06, F07, F08, H09, I06
ZNF668_FL, F06, F07, F08, H09, I06
TIGD4_DBD F09, F10, H03
TIGD4_FL F09, F10, H03
TIGD5_DBD F11, F12, H02
TIGD5_FL F11, F12, H02
ZNF407_DBD F13, H05
ZNF800_DBD F14, F15, H06
ZNF800_FL F14, F15, H06
Success in GHT-SELEX
Is a znf C2H2 protein?
Showed modular binding
51 99 81 96 80 44 97 48 83 52 97 77 80 100
250
150
100
75
50
37
25
20
15
10
Successful in GHT-SELEX
Unuccessful in GHT-SELEX
Expected size
Wheat germ eGFP-IVT

#### Gel G

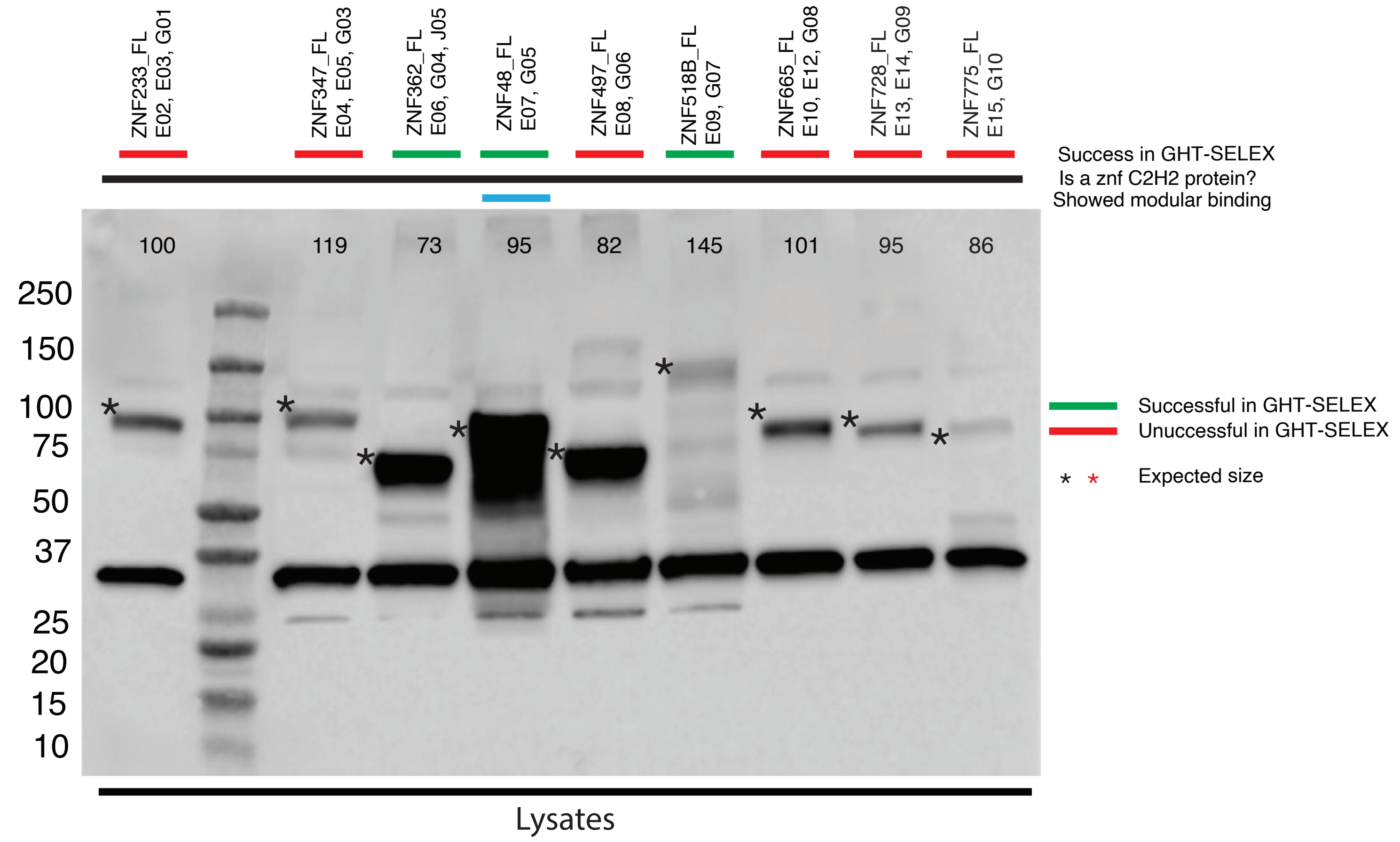
ZNF233_FL E02, E03, G01
ZNF347_FL E04, E05, G03
ZNF362_FL E06, G04, J05
ZNF48_FL E07, G05
ZNF497_FL E08, G06
ZNF518B_FL E09, G07
ZNF665_FL E10, E12, G08
ZNF728_FL E13, E14, G09
ZNF775_FL E15, G10
Success in GHT-SELEX
Is a znf C2H2 protein?
Showed modular binding
100
119
73
95
82
145
101
95
86
250
150
100
75
50
37
25
20
15
10
Successful in GHT-SELEX
Unuccessful in GHT-SELEX
* * Expected size
Lysates

#### Gel H

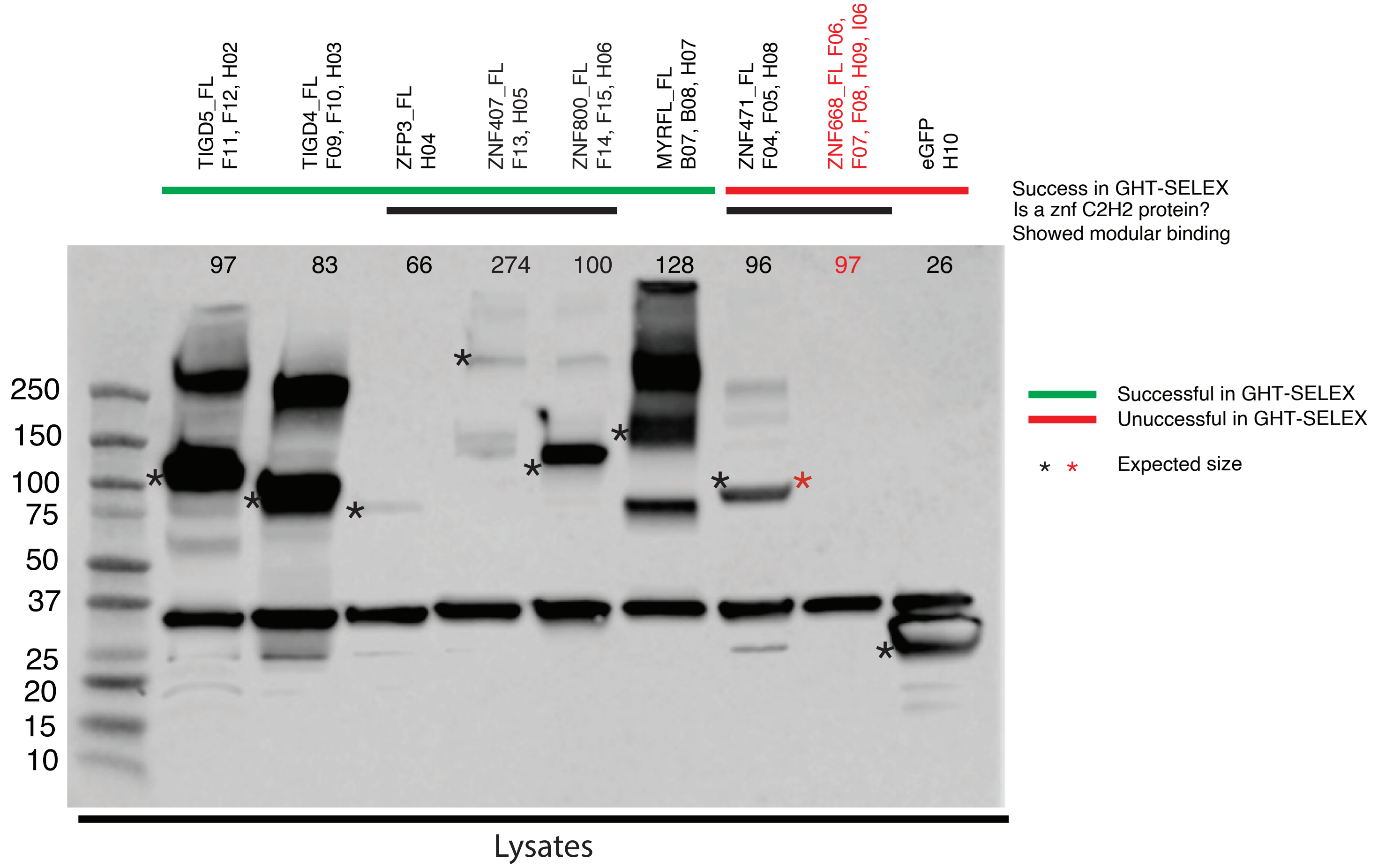
TIGD5_FL F11, F12, H02
TIGD4_FL F09, F10, H03
ZFP3_FL H04
ZNF407_FL F13, H05
ZNF800_FL F14, F15, H06
MYRFL_FL B07, B08, H07
ZNF471_FL F04, F05, H08
ZNF668_FL F06, F07, F08, H09, I06
eGFP H10
Success in GHT-SELEX
Is a znf C2H2 protein?
Showed modular binding
97
83
66
274
100
128
96
97
26
250
150
100
75
50
37
25
20
15
10
Successful in GHT-SELEX
Unuccessful in GHT-SELEX
* * Expected size
Lysates

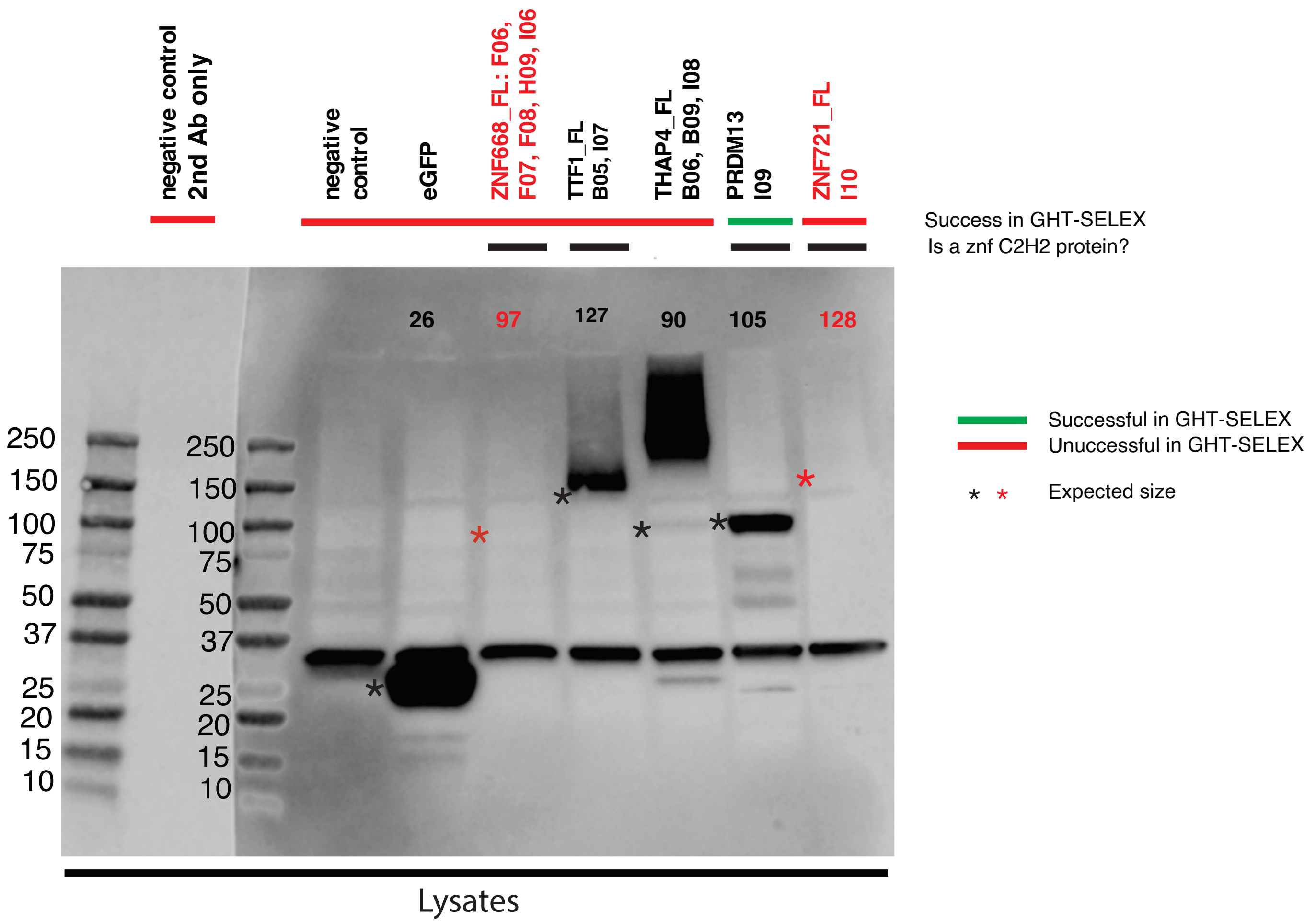
negative control
2nd Ab only
negative control
eGFP
ZNF668_FL: F06, F07, F08, H09, I06
TTF1_FL
B05, I07
THAP4_FL
B06, B09, I08
PRDM13
I09
ZNF721_FL
I10
Success in GHT-SELEX
Is a znf C2H2 protein?
26
97
127
90
105
128
250
150
100
75
50
37
25
20
15
10
250
150
100
75
50
37
25
20
15
10
Successful in GHT-SELEX
Unuccessful in GHT-SELEX
Expected size
Lysates

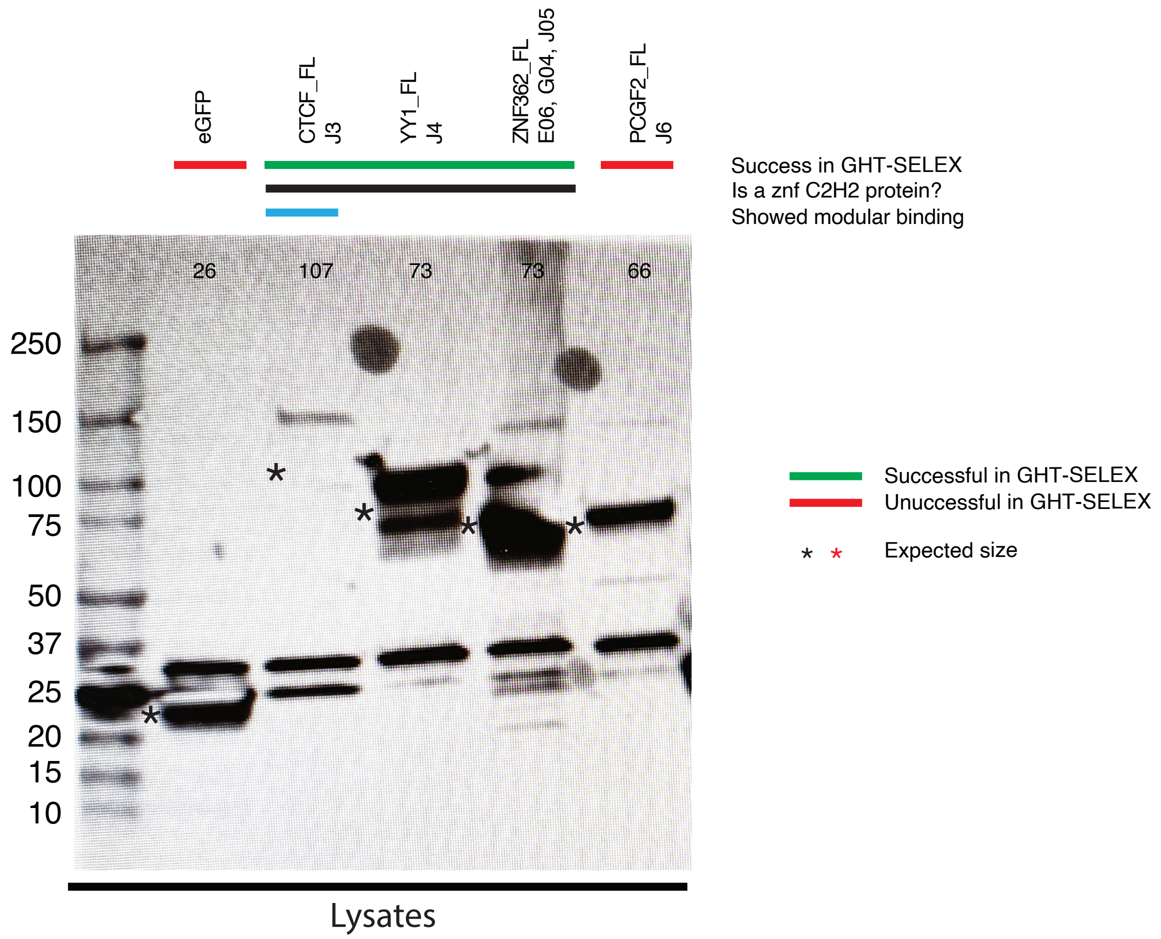
eGFP
CTCF_FL
J3
YY1_FL
J4
ZNF362_FL
E06, G04, J05
PCGF2_FL
J6
Success in GHT-SELEX
Is a znf C2H2 protein?
Showed modular binding
26
107
73
73
66
250
150
100
75
50
37
25
20
15
10
Successful in GHT-SELEX
Unuccessful in GHT-SELEX
Expected size
Lysates
