## Supplementary figures and images for "GHT-SELEX demonstrates unexpectedly high intrinsic sequence specificity and complex DNA binding of many human transcription factors"

### ZNF721_PWM005513_logo.png

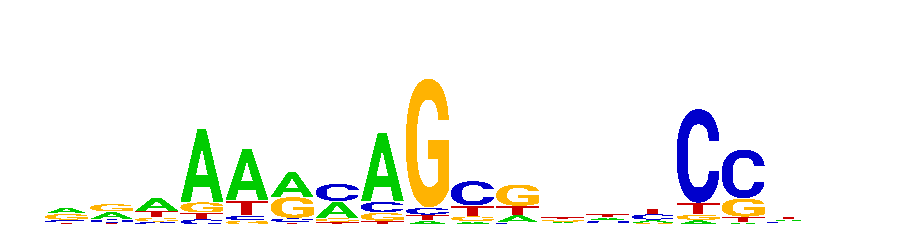
