## Supplementary figures and images for "GHT-SELEX demonstrates unexpectedly high intrinsic sequence specificity and complex DNA binding of many human transcription factors"

### AC092835_PWM006586_logo_REV.png

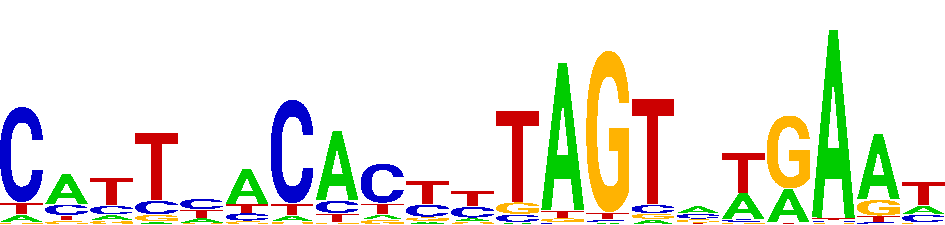

### CAMTA1_PWM000118_logo_REV.png

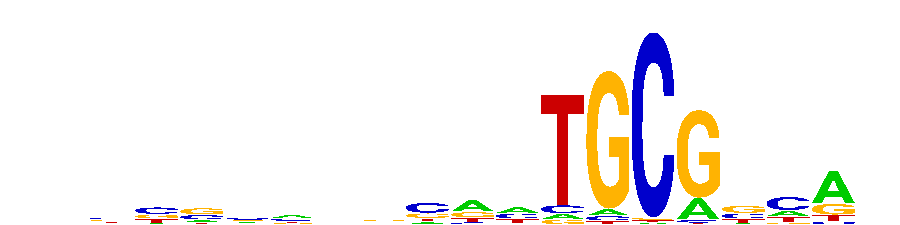

### CASZ1_PWM000182_logo_REV.png

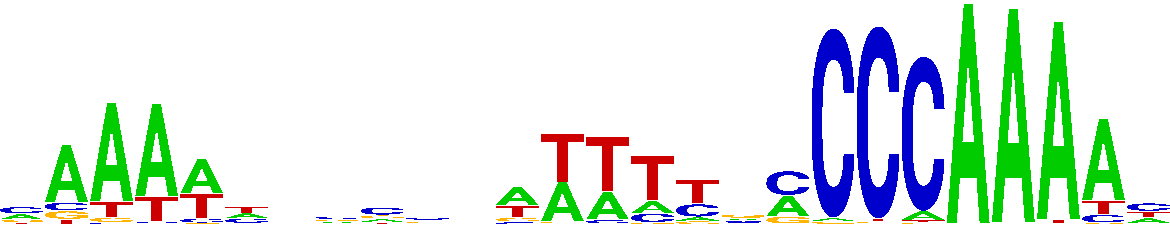

### CGGBP1_PWM000210_logo_REV.png

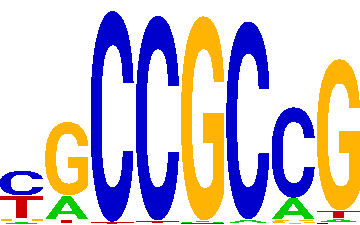

### DACH2_PWM021827_logo_REV.png

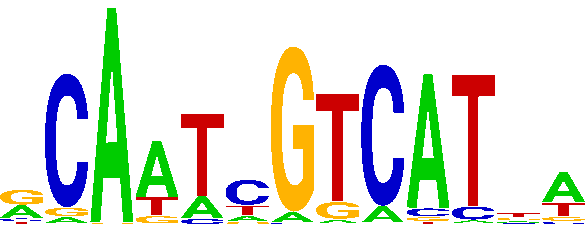

### ELF3_PWM000465_logo.png

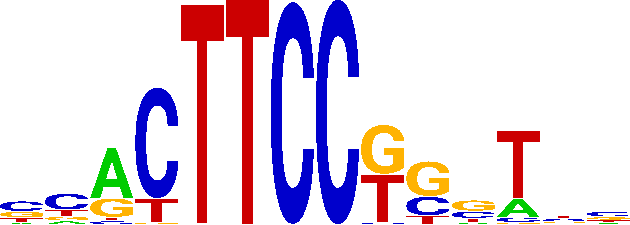

### GABPA_PWM000688_logo.png

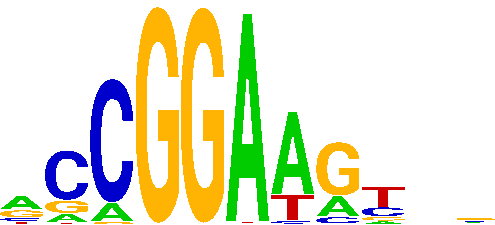

### GATAD2A_PWM000707_logo.png

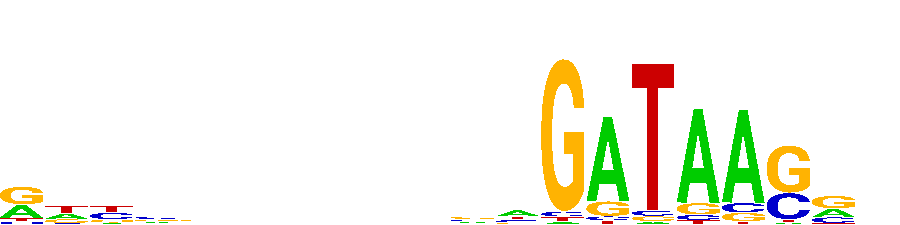

### GCM1_PWM000730_logo_REV.png

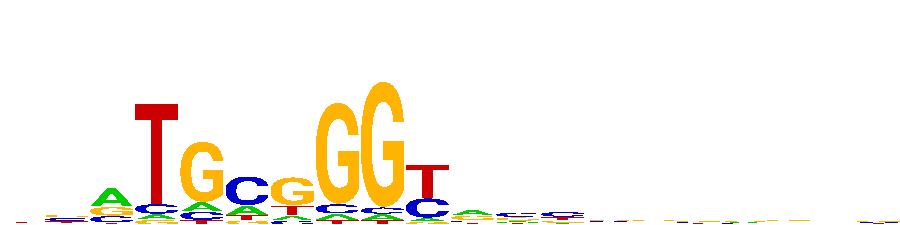

### HMGA1_PWM036826_logo_REV.png

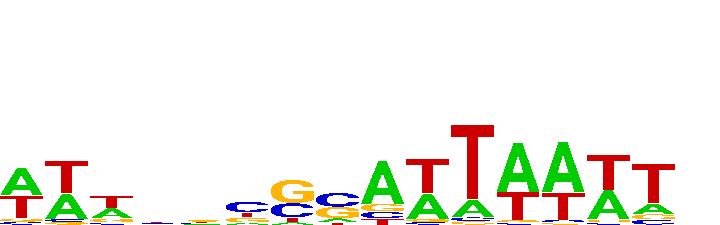

### LEF1_PWM000933_logo.png

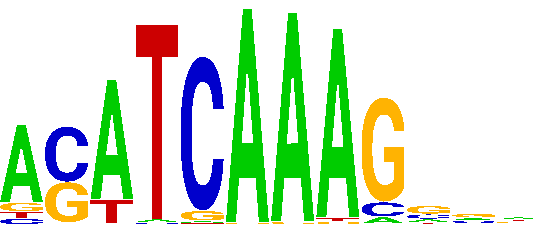

### LHX6_PWM000985_logo.png

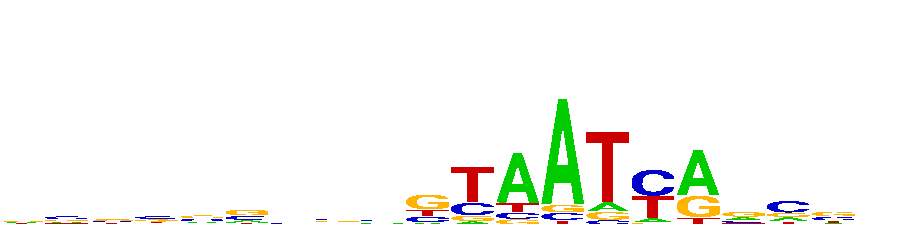

### MAX_PWM001025_logo.png

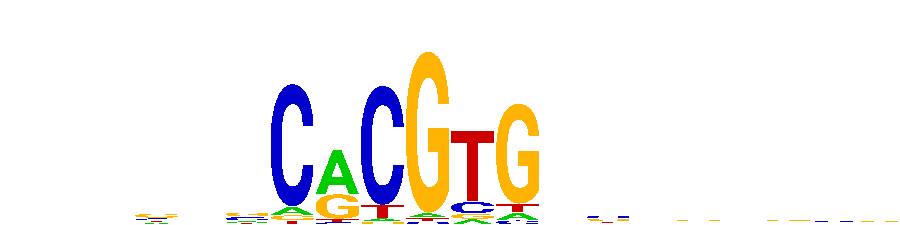

### MBD1_PWM001044_logo.png

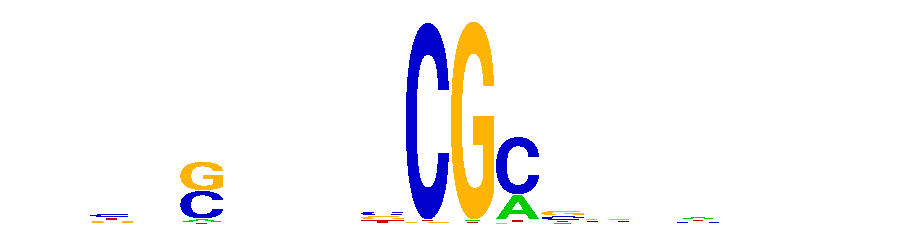

### MGA_PWM001108_logo.png

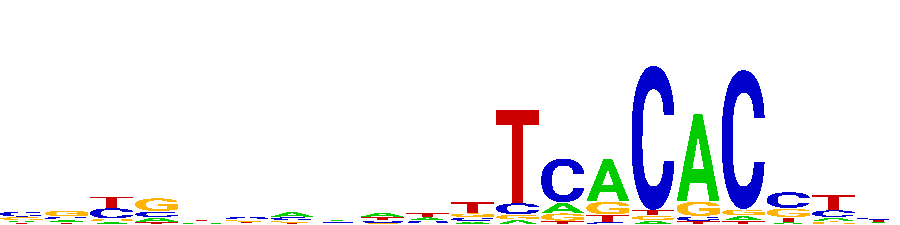

### MGA_PWM001108_logo_REV.png

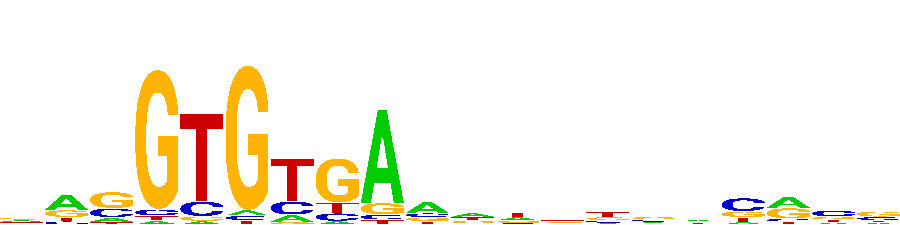

### MSANTD1_PWM001197_logo_REV.png

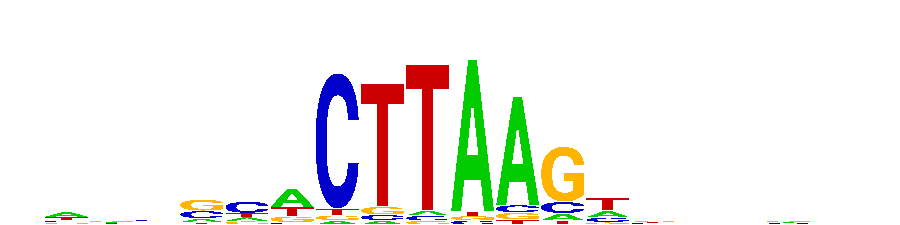

### MYPOP_PWM001308_logo.png

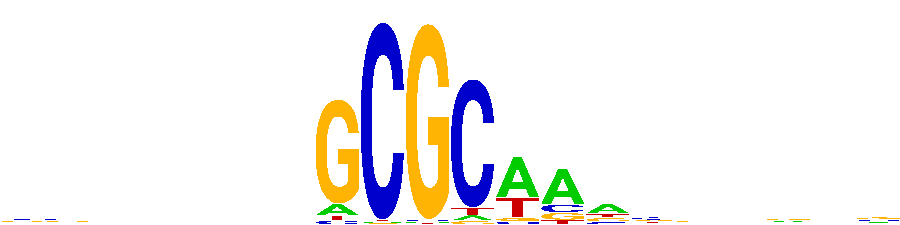

### MYSM1_PWM001347_logo.png

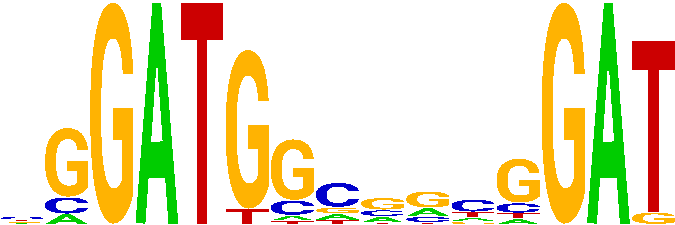

### NACC2_PWM001391_logo_REV.png

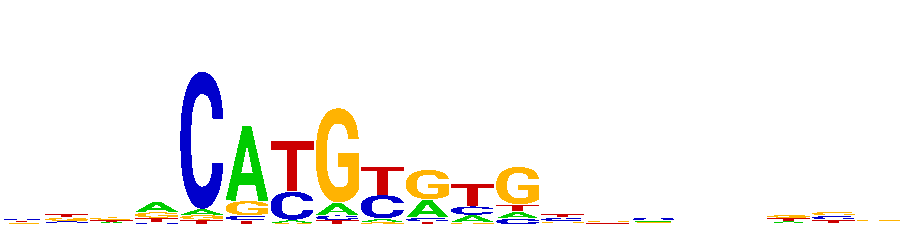

### NR1H4_PWM001456_logo_REV.png

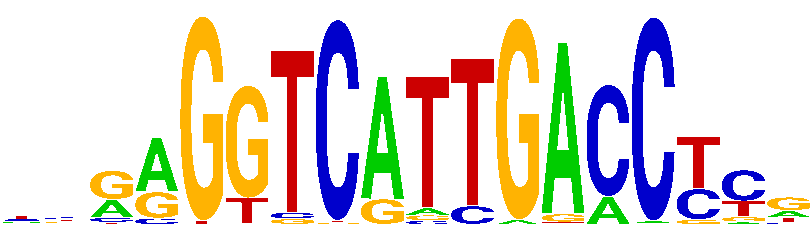

### PAX7_PWM001537_logo_REV.png

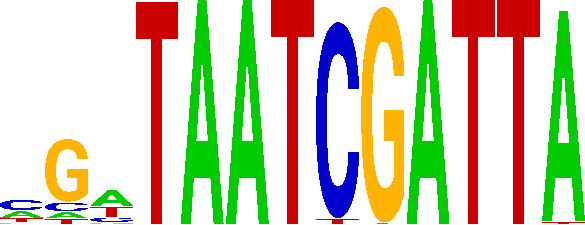

### POU5F1_PWM001584_logo_REV.png

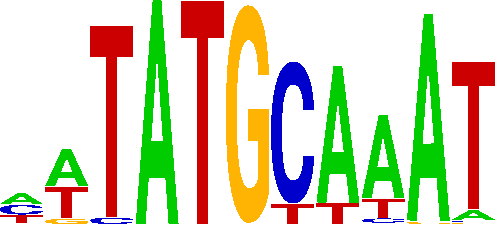

### PRDM5_PWM001672_logo.png

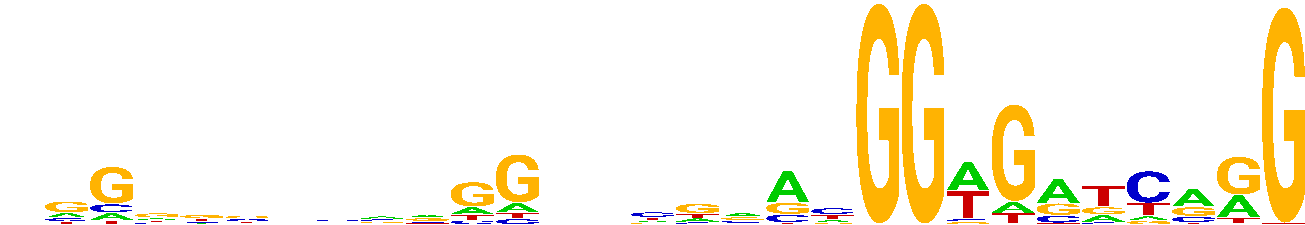

### PRDM5_PWM001672_logo_REV.png

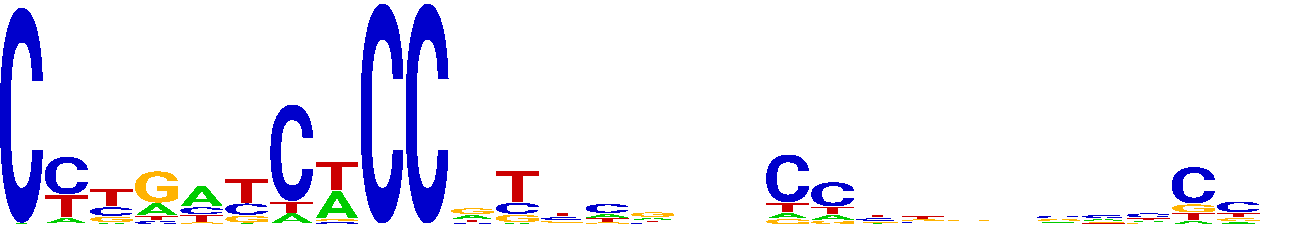

### RFX5_PWM001734_logo_REV.png

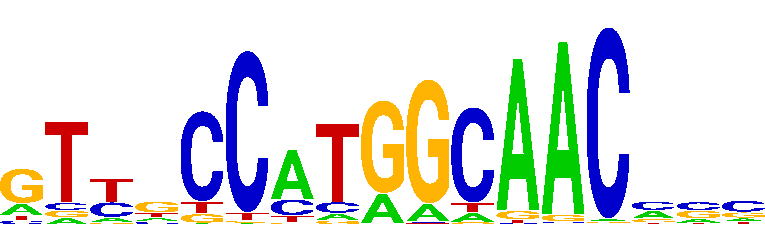

### RLF_PWM001787_logo.png

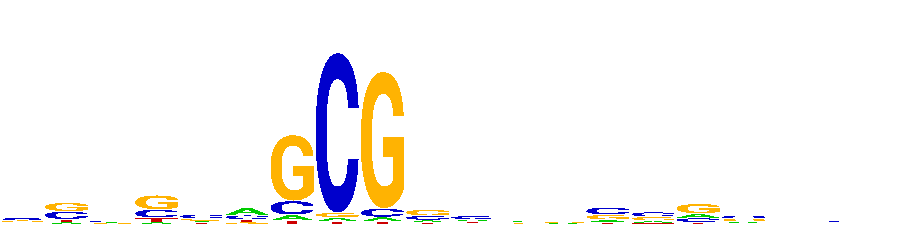

### RXRA_PWM001851_logo.png

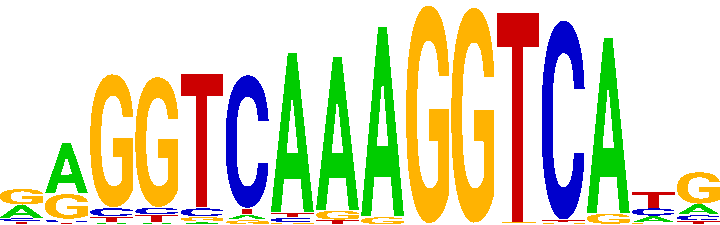

### SALL3_PWM001887_logo_REV.png

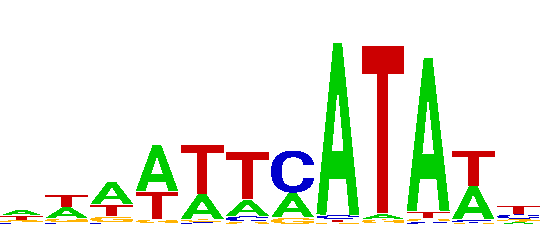

### SETBP1_PWM072635_logo.png

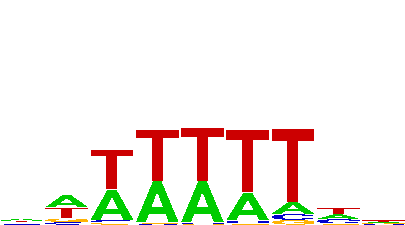

### SETBP1_PWM072635_logo_REV.png

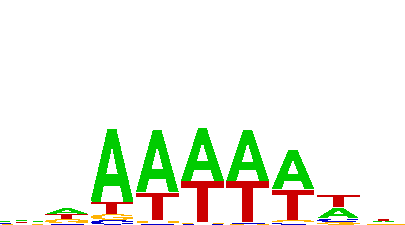

### SOX5_PWM077815_logo.png

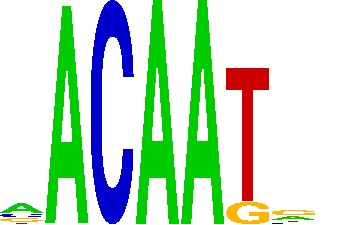

### SOX15_PWM001978_logo.png

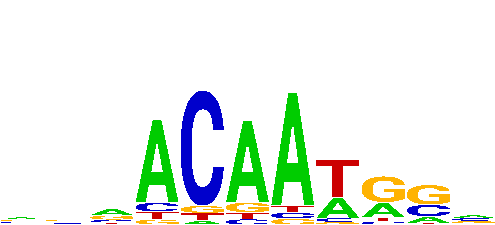

### SOX15_PWM001978_logo_REV.png

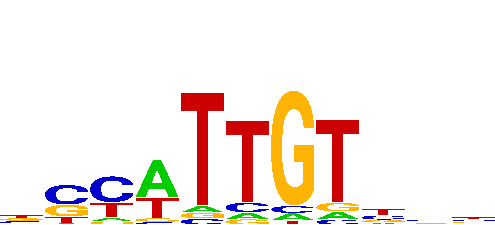

### SP140_PWM002077_logo_REV.png

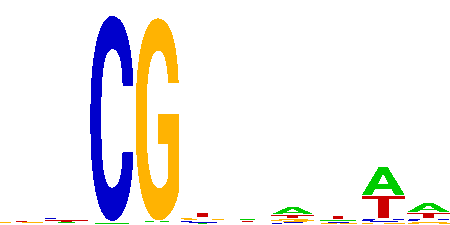

### SRY_PWM002141_logo_REV.png

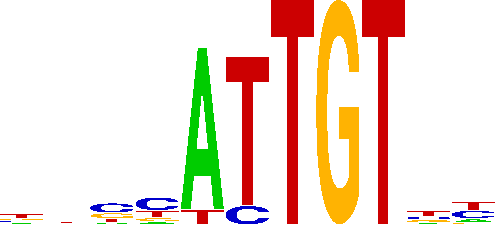

### TERF1_PWM002178_logo.png

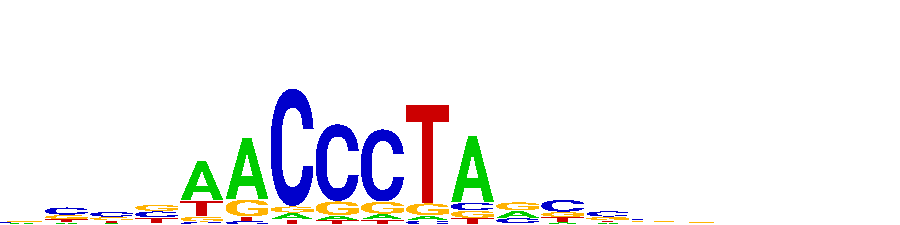

### TERF1_PWM002178_logo_REV.png

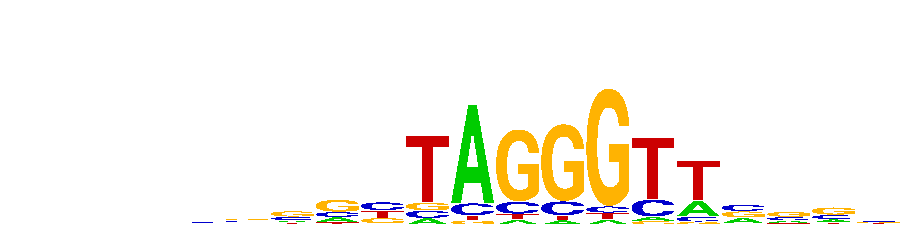

### TIGD4_PWM089799_logo.png

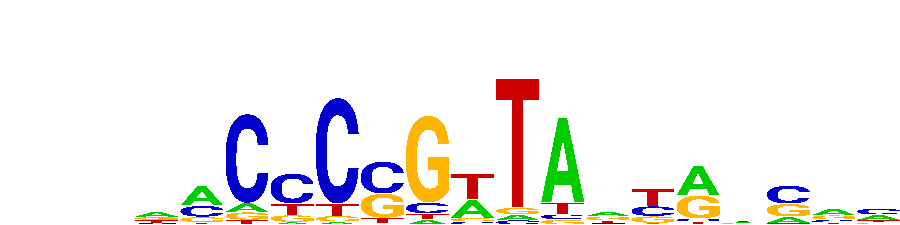

### TIGD7_PWM002338_logo_REV.png

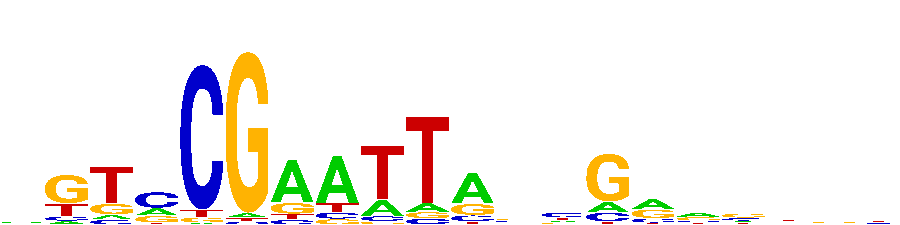

### TPRX1_PWM002375_logo.png

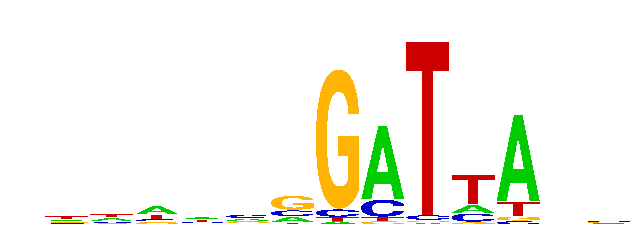

### USF3_PWM002464_logo.png

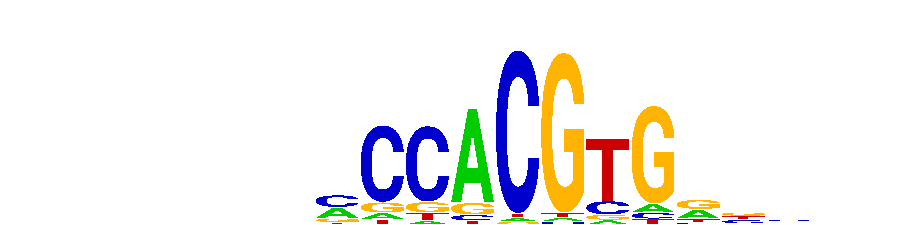

### USF3_PWM002464_logo_REV.png

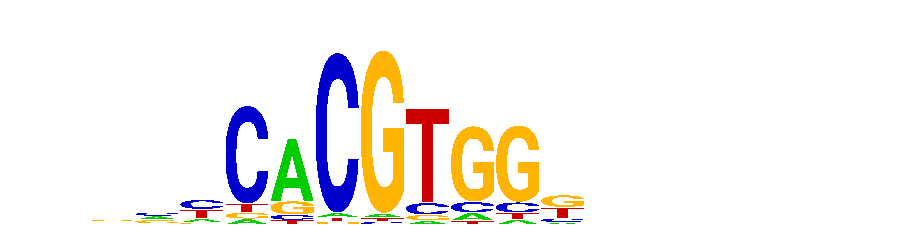

### ZBTB8B_PWM002864_logo_REV.png

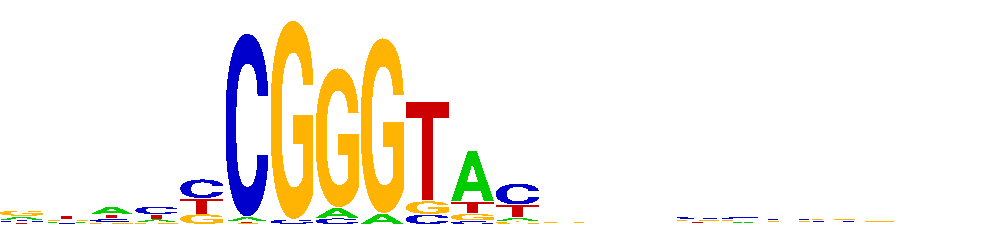

### ZBTB24_PWM002721_logo.png

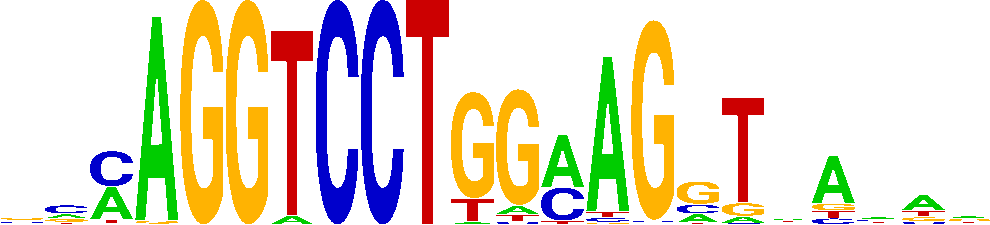

### ZBTB47_PWM002804_logo.png

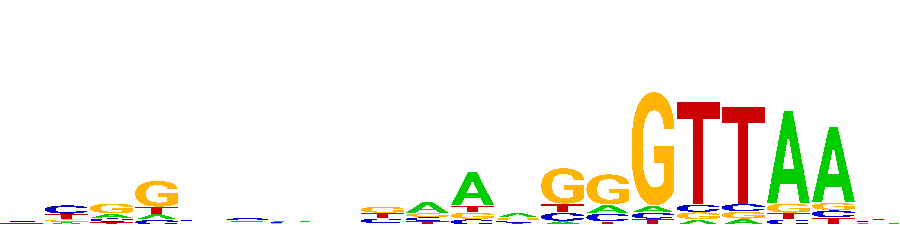

### ZFAT_PWM112021_logo.png

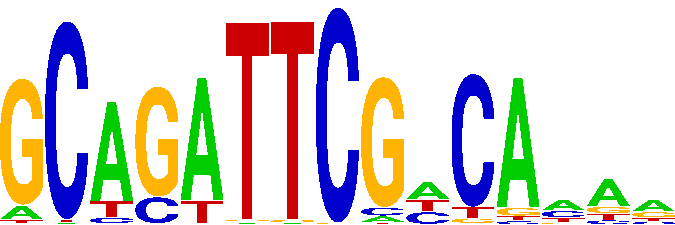

### ZFP3_PWM002952_logo.png

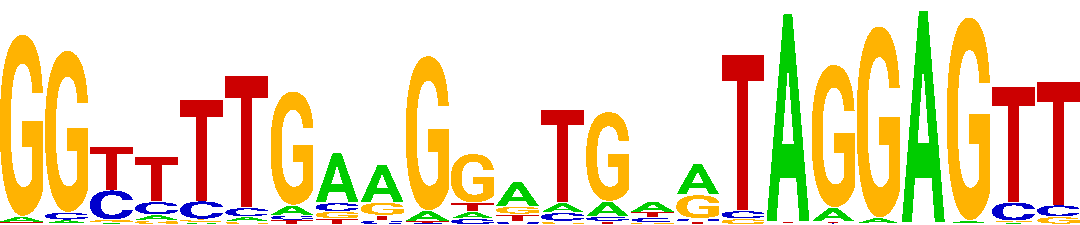

### ZGLP1_PWM003064_logo.png

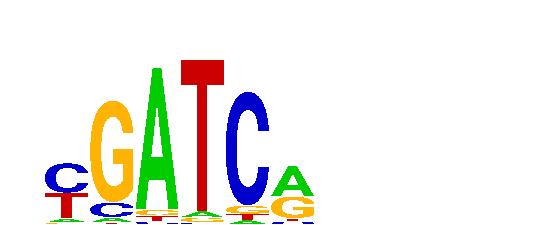

### ZKSCAN4_PWM003117_logo_REV.png

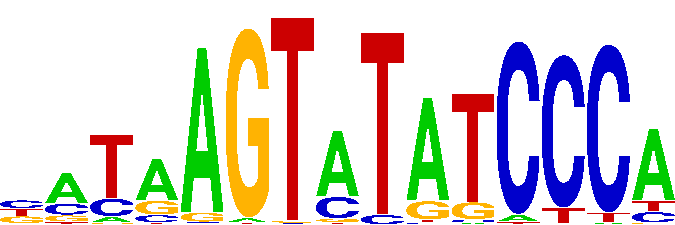

### ZNF14_PWM003249_logo_REV.png

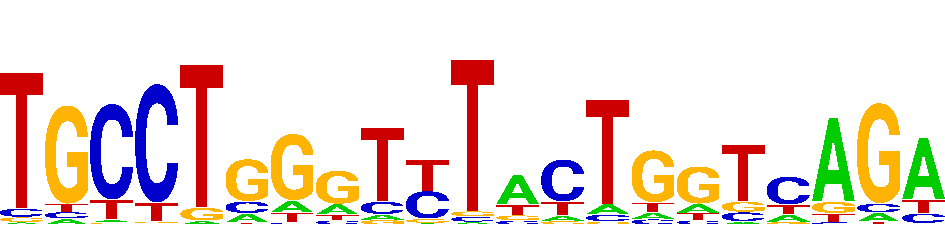

### ZNF16_PWM003365_logo.png

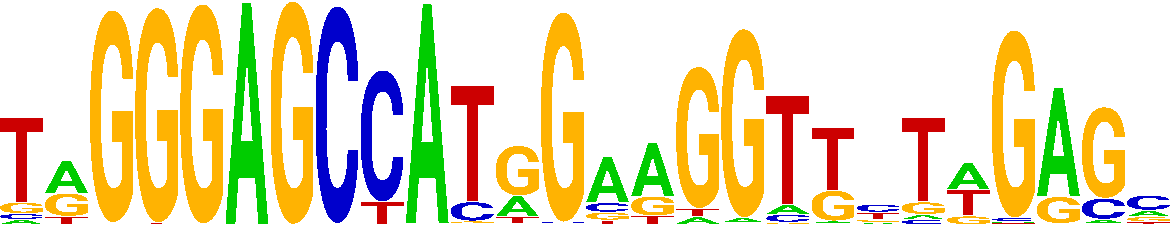

### ZNF18_PWM003405_logo_REV.png

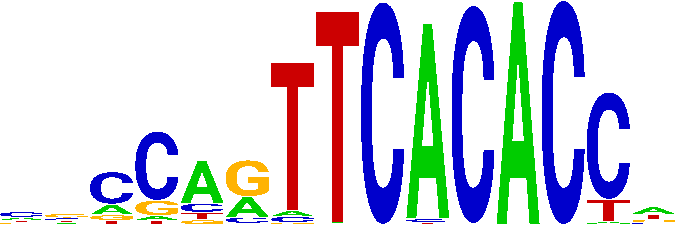

### ZNF20_PWM003431_logo.png

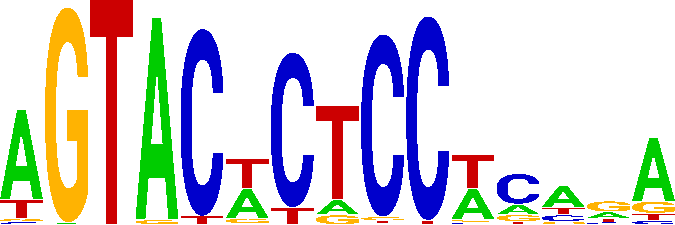

### ZNF35_PWM003959_logo_REV.png

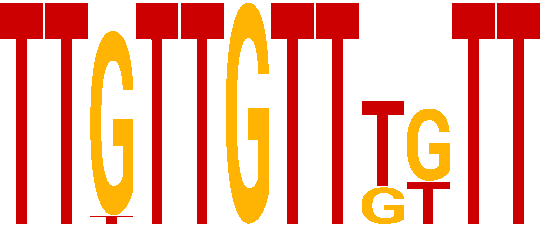

### ZNF43_PWM004214_logo_REV.png

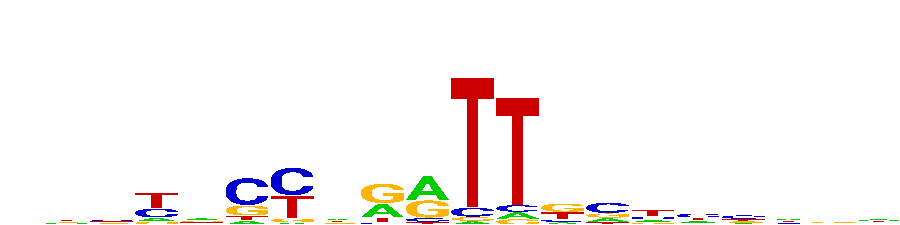

### ZNF57_PWM004724_logo.png

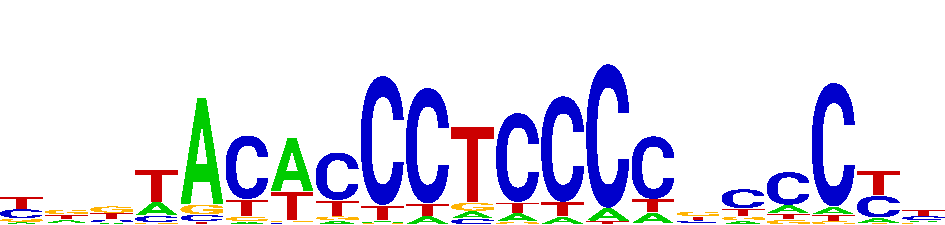

### ZNF70_PWM005355_logo_REV.png

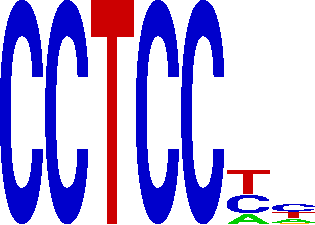

### ZNF92_PWM006209_logo.png

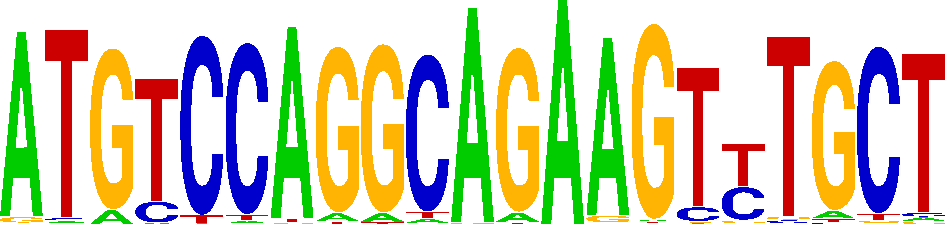

### ZNF160_PWM003374_logo_REV.png

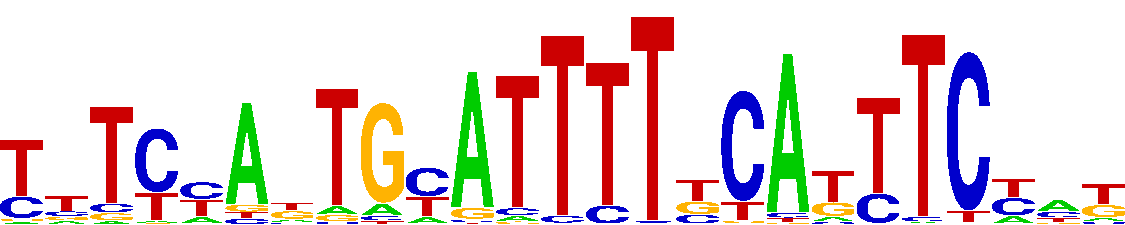

### ZNF226_PWM003524_logo_REV.png

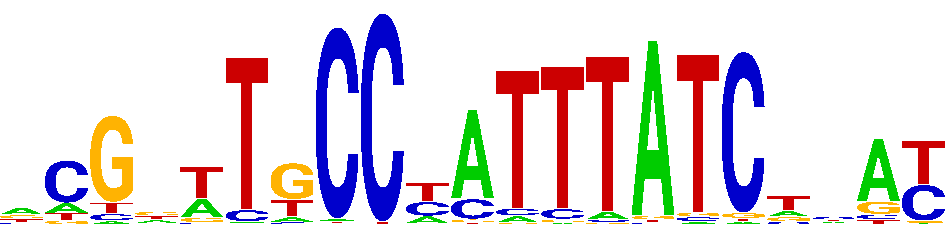

### ZNF234_PWM003628_logo.png

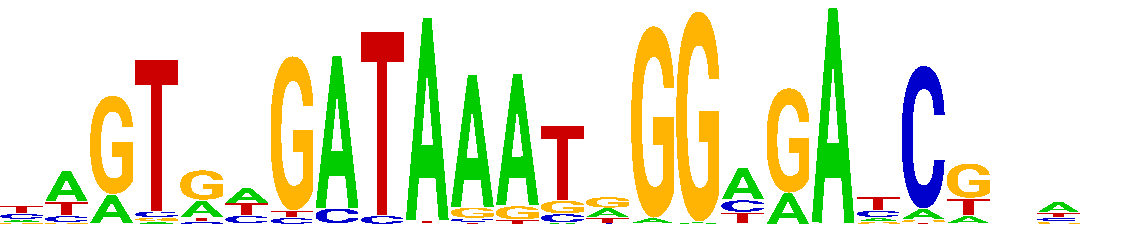

### ZNF251_PWM003658_logo.png

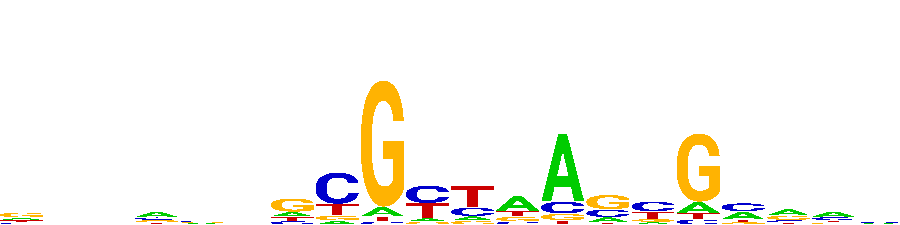

### ZNF251_PWM003658_logo_REV.png

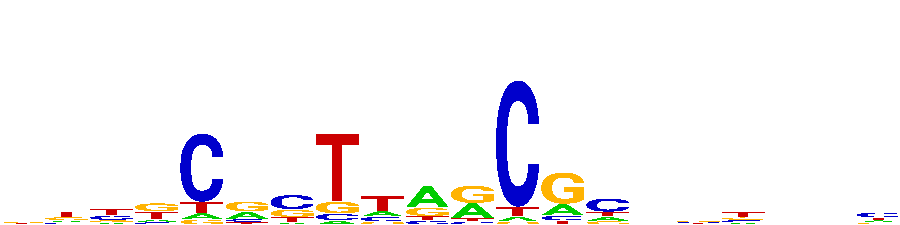

### ZNF264_PWM003714_logo.png

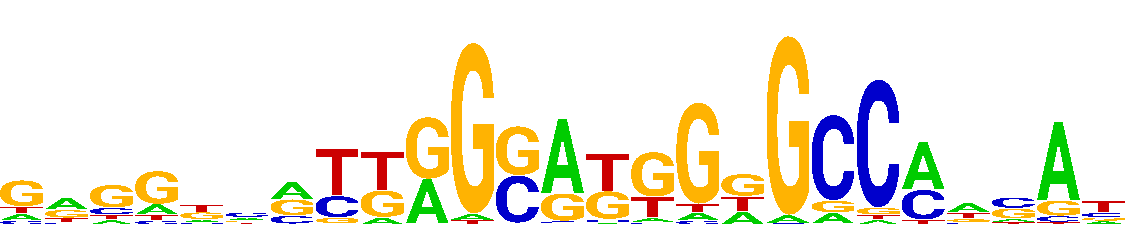

### ZNF362_PWM004025_logo_REV.png

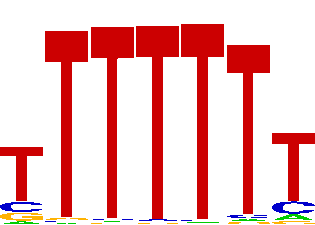

### ZNF367_PWM004066_logo.png

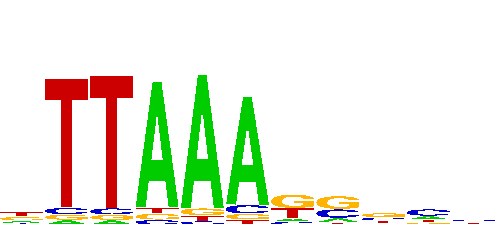

### ZNF454_PWM004269_logo.png

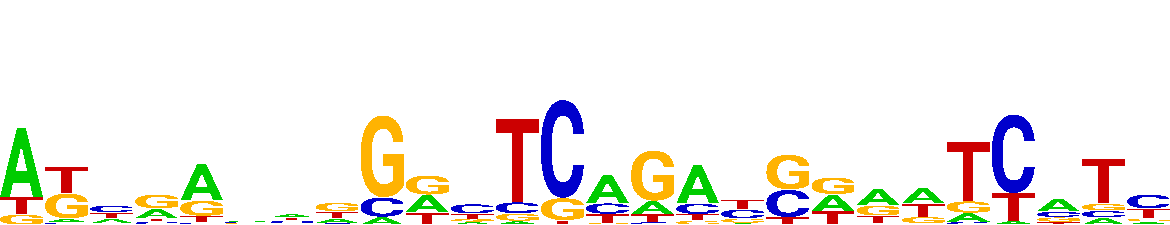

### ZNF471_PWM004322_logo_REV.png

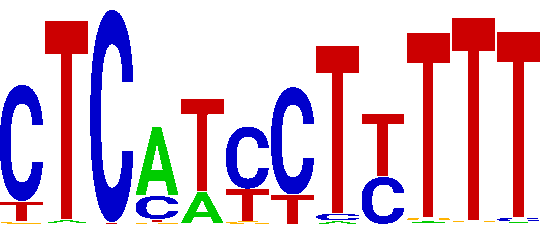

### ZNF490_PWM004386_logo.png

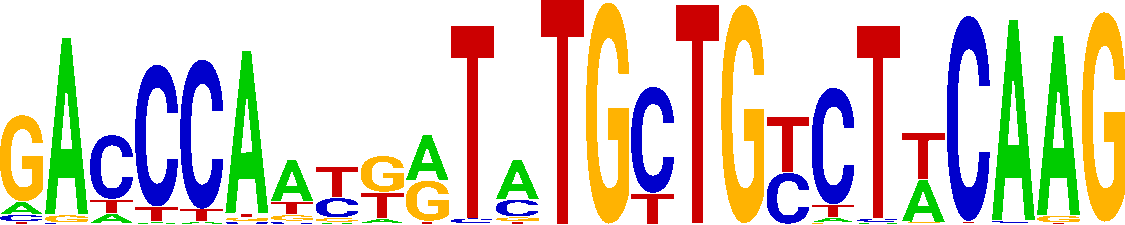

### ZNF490_PWM004386_logo_REV.png

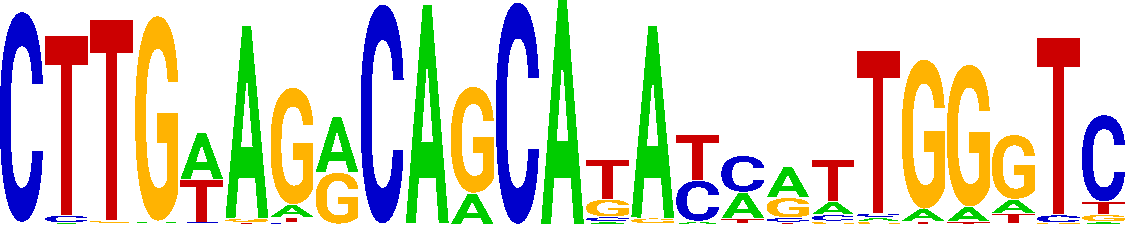

### ZNF493_PWM004421_logo.png

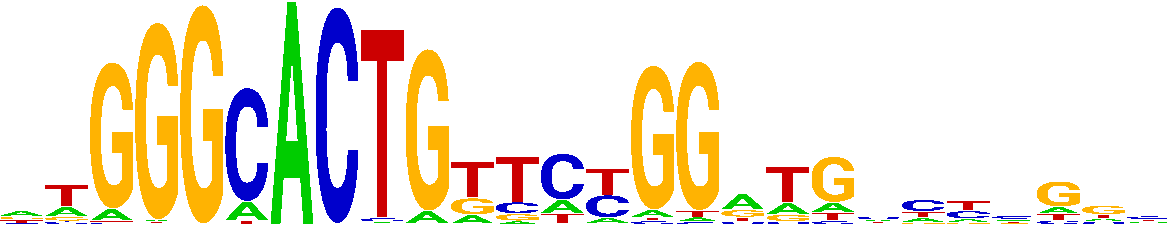

### ZNF500_PWM004464_logo_REV.png

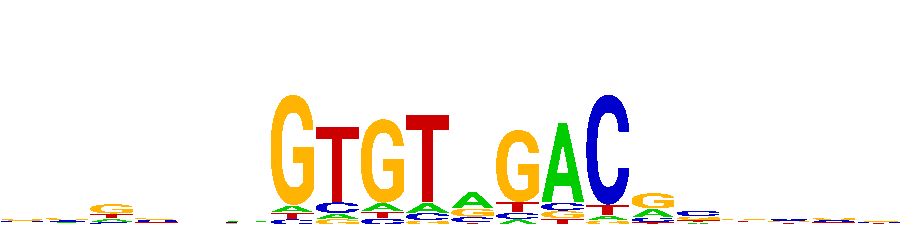

### ZNF507_PWM150320_logo.png

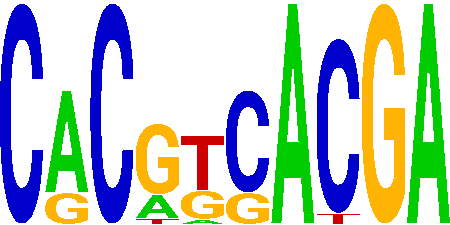

### ZNF507_PWM150320_logo_REV.png

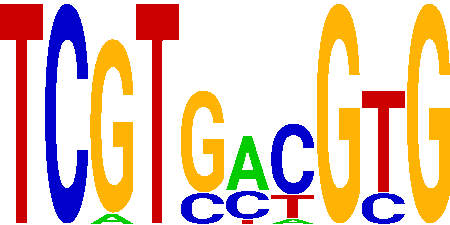

### ZNF516_PWM151871_logo.png

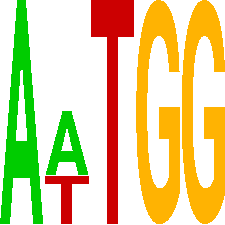

### ZNF518B_PWM004581_logo.png

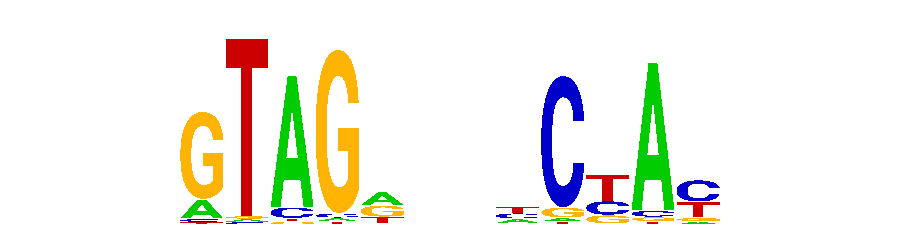

### ZNF518B_PWM004581_logo_REV.png

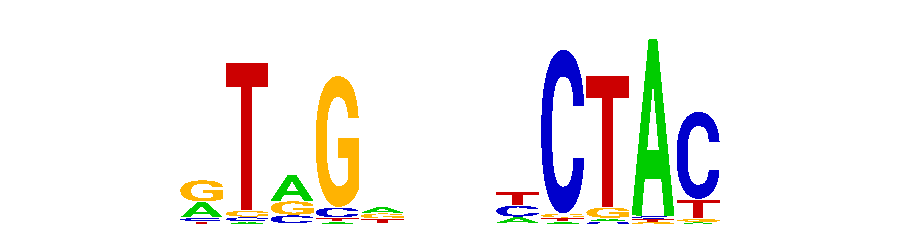

### ZNF536_PWM155337_logo_REV.png

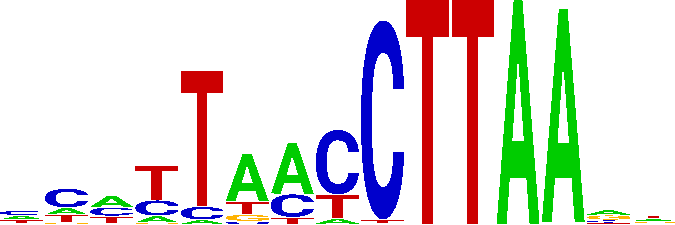

### ZNF569_PWM004692_logo_REV.png

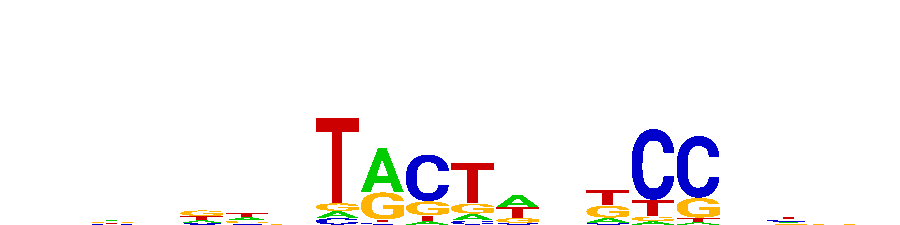

### ZNF587B_PWM160251_logo_REV.png

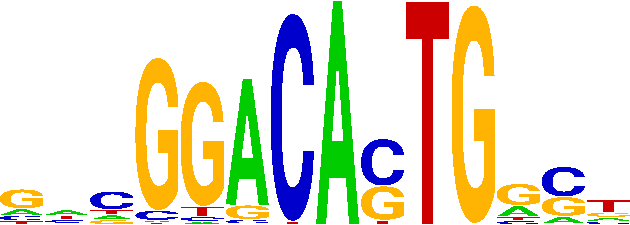

### ZNF623_PWM004913_logo_REV.png

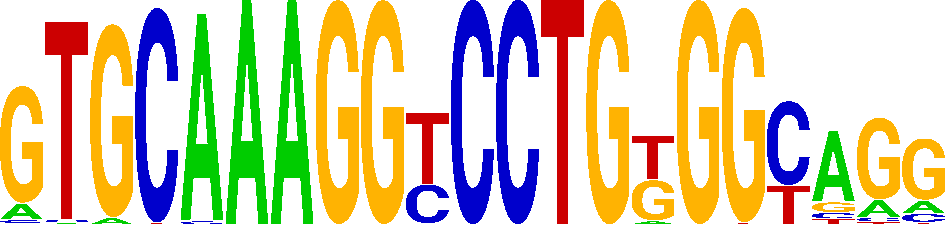

### ZNF668_PWM005084_logo.png

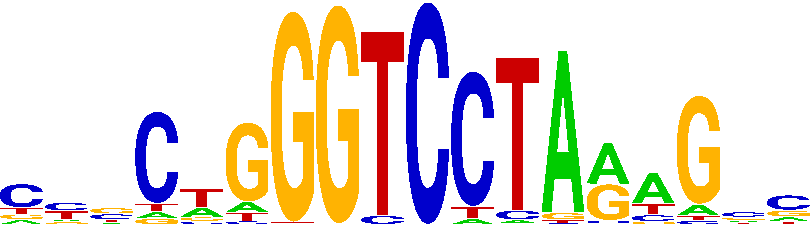

### ZNF708_PWM005442_logo_REV.png

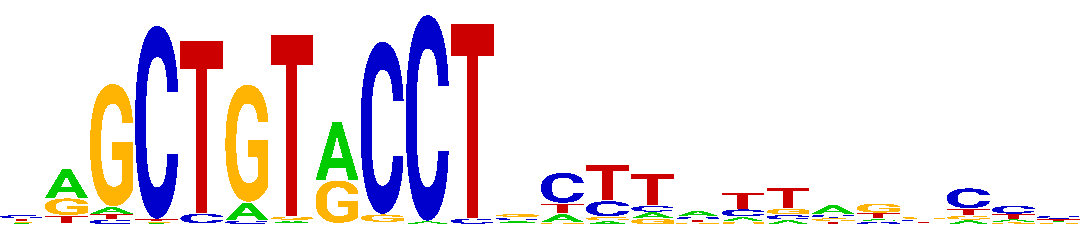

### ZNF721_PWM005513_logo_REV.png

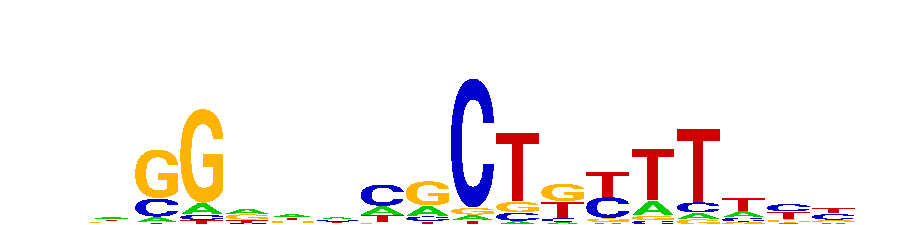

### ZNF728_PWM005588_logo.png

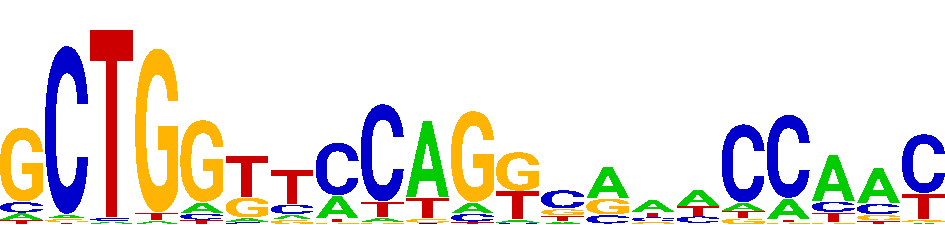

### ZNF728_PWM005588_logo_REV.png

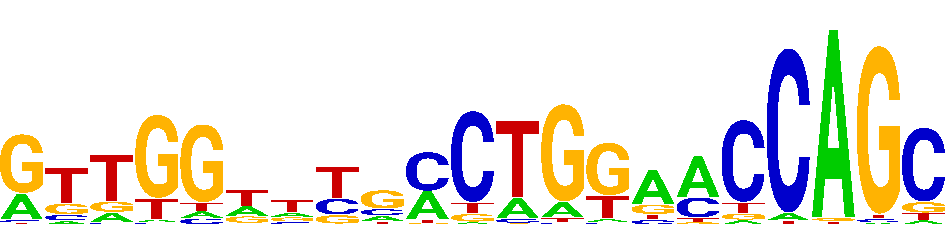

### ZNF729_PWM005614_logo_REV.png

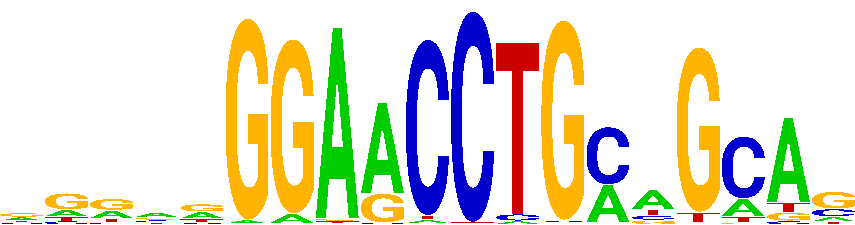

### ZNF732_PWM005632_logo.png

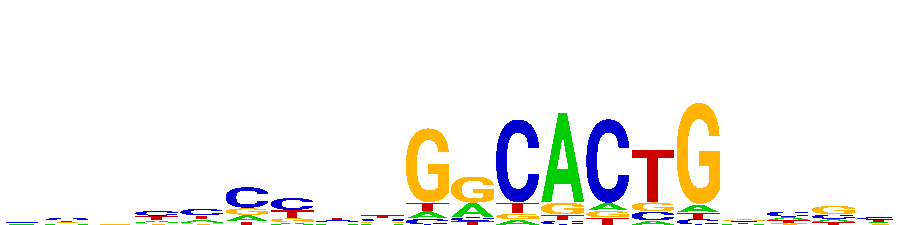

### ZNF770_PWM005686_logo_REV.png

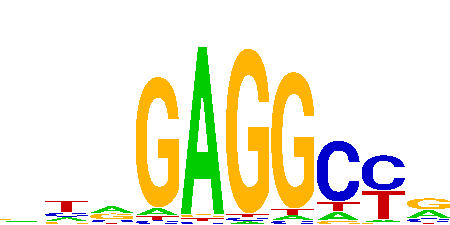

### ZNF841_PWM006037_logo.png

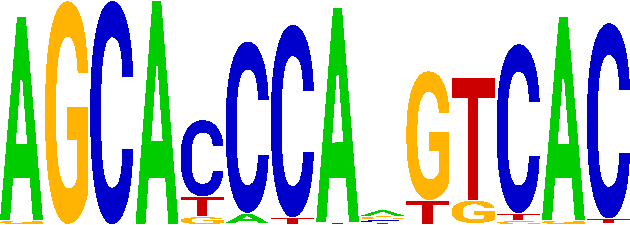

### ZNF845_PWM193356_logo.png

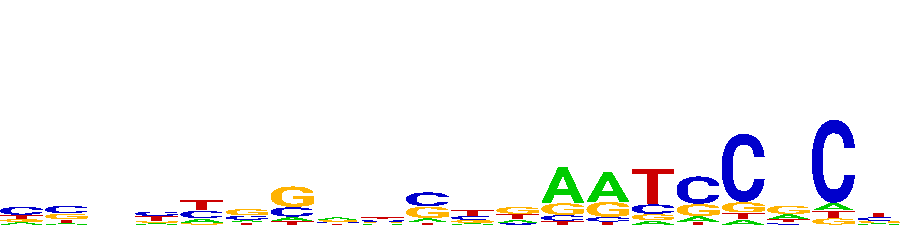

### ZNF850_PWM006089_logo.png

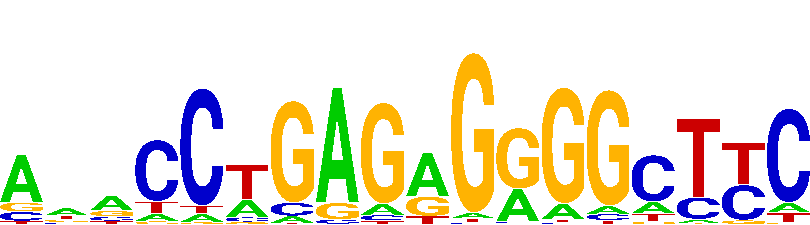

### ZNF850_PWM006089_logo_REV.png

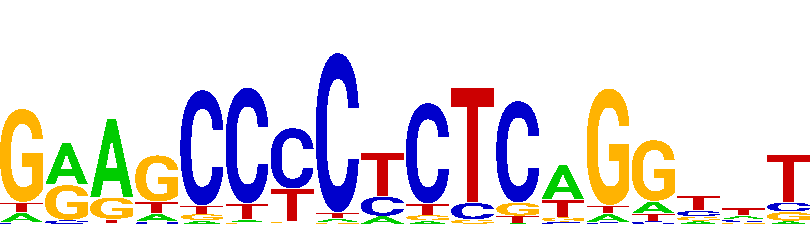

### ZNF853_PWM006118_logo_REV.png

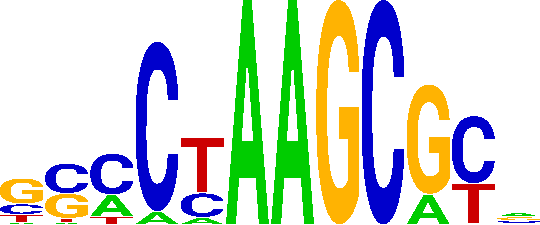

### ZNF878_PWM006160_logo_REV.png

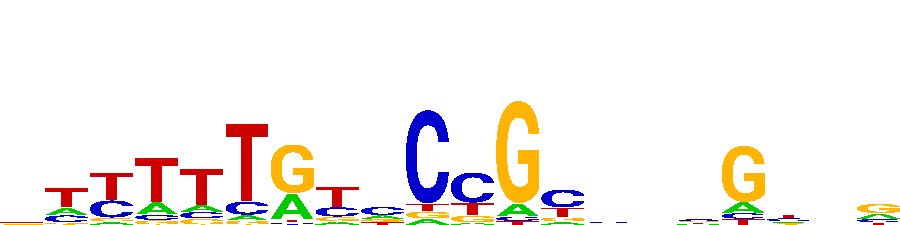

### ZNF888_PWM006179_logo.png

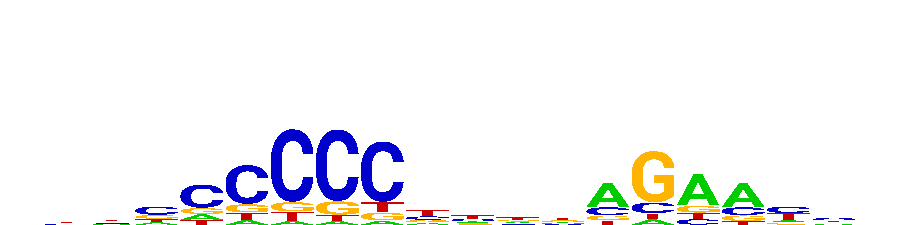

### ZSCAN12_PWM006222_logo.png

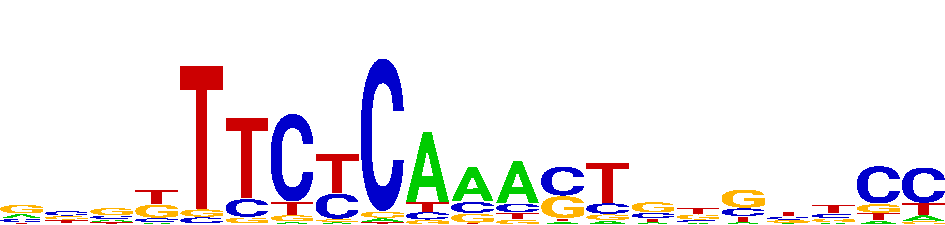

### ZSCAN25_PWM006286_logo.png

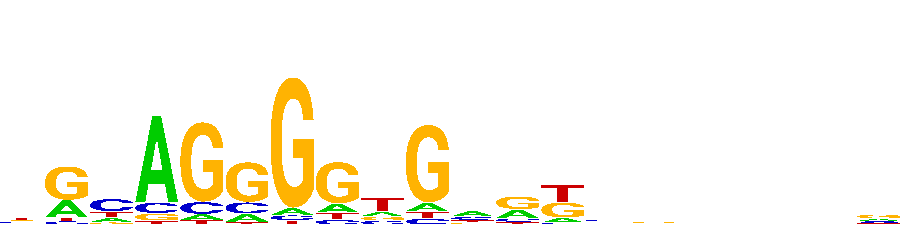

### ZXDB_PWM006360_logo_REV.png

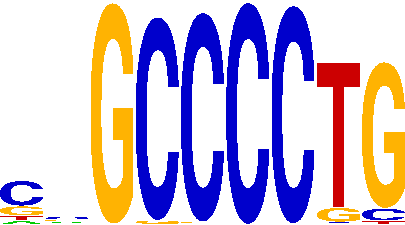
